# Hearing impairment due to *Mir183/96/182* mutations suggests both loss and gain of function effects

**DOI:** 10.1101/579003

**Authors:** Morag A. Lewis, Francesca Di Domenico, Neil J. Ingham, Haydn M. Prosser, Karen P. Steel

## Abstract

The microRNA miR-96 is important for hearing, as point mutations in humans and mice result in dominant progressive hearing loss. *Mir96* is expressed in sensory cells along with *Mir182* and *Mir183*, but the roles of these closely-linked microRNAs are as yet unknown. Here we analyse mice carrying null alleles of *Mir182*, and of *Mir183* and *Mir96* together to investigate their roles in hearing. We found that *Mir183*/*96* heterozygous mice had normal hearing and homozygotes were completely deaf with abnormal hair cell stereocilia bundles and reduced numbers of inner hair cell synapses at four weeks old. *Mir182* knockout mice developed normal hearing then exhibited progressive hearing loss. Our transcriptional analyses revealed significant changes in a range of other genes, but surprisingly there were fewer genes with altered expression in the organ of Corti of *Mir183/96* null mice compared with our previous findings in *Mir96^Dmdo^* mutants, which have a point mutation in the miR-96 seed region. This suggests the more severe phenotype of *Mir96^Dmdo^* mutants compared with *Mir183*/*96* mutants, including progressive hearing loss in *Mir96^Dmdo^* heterozygotes, is likely to be mediated by the gain of novel target genes in addition to the loss of its normal targets. We propose three mechanisms of action of mutant miRNAs; loss of targets that are normally completely repressed, loss of targets whose transcription is normally buffered by the miRNA, and gain of novel targets. Any of these mechanisms could lead to a partial loss of a robust cellular identity and consequent dysfunction.

## Introduction

The microRNAs miR-96, miR-182 and miR-183 are expressed together on a single transcript in sensory cells, including the retina and the hair cells of the inner ear [1, 2]. Point mutations in *Mir96* cause rapidly progressive hearing loss in the diminuendo mouse mutant (*Mir96^Dmdo^* [3]) and progressive hearing loss with later onset in human families [4, 5], and the diminuendo mutation has also been shown to delay maturation of the central auditory system [6]. In homozygous *Mir96^Dmdo^* mice, most of the cochlear hair cells die by 28 days after birth. However, this is not the cause of the hearing loss; even before the onset of normal hearing, homozygote hair cells fail to mature both morphologically and physiologically, remaining in their immature state, and heterozygote hair cells show a developmental delay. miR-96 is thus thought to be responsible for coordinating hair cell maturation [7, 8].

Overexpression of the three microRNAs also results in cochlear defects and hearing loss [9]. The complete loss of all mature miRNAs from the inner ear results in very early developmental defects including a severely truncated cochlear duct [10, 11]. miR-96, miR-182 and miR-183 have also been implicated in other diseases, including glaucoma [12], ischemic injury [13, 14] and spinal cord injury [15].

MicroRNAs regulate the expression of many other genes by targeting specific sequences in their mRNAs, leading to transcript destabilisation or translational inhibition. Transcriptome analyses of the *Mir96^Dmdo^* mouse organ of Corti showed that many genes were dysregulated in homozygotes, including several known to be important for hearing which appear to contribute to specific aspects of the diminuendo phenotype [3, 7, 8, 16]. However, the diminuendo mutation is a single base pair change in the seed region of the microRNA which is critical for correct targeting, and it is not clear to what extent the diminuendo mutant phenotype is the result of the loss of normal targets of miR-96, and how much is due to the gain of novel targets. We previously suggested that the progressive hearing loss was most likely due to the loss of normal target repression because all three point mutations in mouse and human *Mir96* lead to a similar phenotype which seems unlikely if the gain of novel targets is the main mechanism involved.

The regulatory network generated from *Mir96^Dmdo^* expression data [16] includes a number of genes known to be involved in deafness such as *Ptprq, Gfi1, Kcna10* and *Slc26a5* as well as new candidate genes. Manipulating this network may be a useful therapeutic approach to treating hearing loss due to hair cell dysfunction triggered by a broad range of factors, including both genetic variants and environmental insults. For example, *Trp53, Hif1a* and *Nfe2l2* are in the *Mir96^Dmdo^* network and are involved in cellular responses to stress [17]. In order to focus our translational efforts, it is important to understand better the molecular basis of the network. For this reason, we have analysed a second mutation of *Mir96* in this report, a double knockout of *Mir96* and *Mir183*, as well as a knockout of the closely linked *Mir182* gene, generated through a mouse miRNA knockout program [18]. We carried out Auditory Brainstem Response (ABR) and Distortion Product Otoacoustic Emission (DPOAE) tests to characterise their hearing, scanning electron and confocal microscopy to examine their hair cells and synapses, and mRNA sequencing (RNA-seq) to study their transcriptomes. While both new mouse mutants exhibit hearing loss, their electrophysiological and transcriptional phenotypes differ from the diminuendo mouse, with no sign of hearing loss in the heterozygotes, suggesting that the more severe phenotype of *Mir96^Dmdo^* mutants is likely to be mediated by the gain of novel target genes in addition to the loss of its normal targets.

## Results

### *Mir183*/*96* and *Mir182* knockout mice

Two mouse lines were used in this study; a knockout of *Mir182* (*Mir182^tm1Hmpr/Wtsi^*, referred to from here on as *Mir182^ko^*) and a double knockout of both *Mir183* and *Mir96* (*Mirc40^tm1Hmpr/WtsiOulu^*, referred to from here on as *Mir183/96^dko^*), which are only 116bp apart, making it technically challenging to generate two separate knockouts. The mice were generated and maintained on the C57BL/6N genetic background. C57BL/6 mice are known to have age-related hearing loss, partly due to the *Cdh23^ahl^* allele [19]. Higher frequencies are affected first, after four weeks of age, while the lower frequencies remain unaffected up to 6 months old [20]. We observed a similar pattern in wildtype mice from both the *Mir183*/*96^dko^* and *Mir182^ko^* lines, which exhibited mild progressive hearing loss at 24-42kHz from 8 weeks old but retained good hearing sensitivity at frequencies between 3-12kHz up to six months old (Fig. 1).

**Figure 1.**
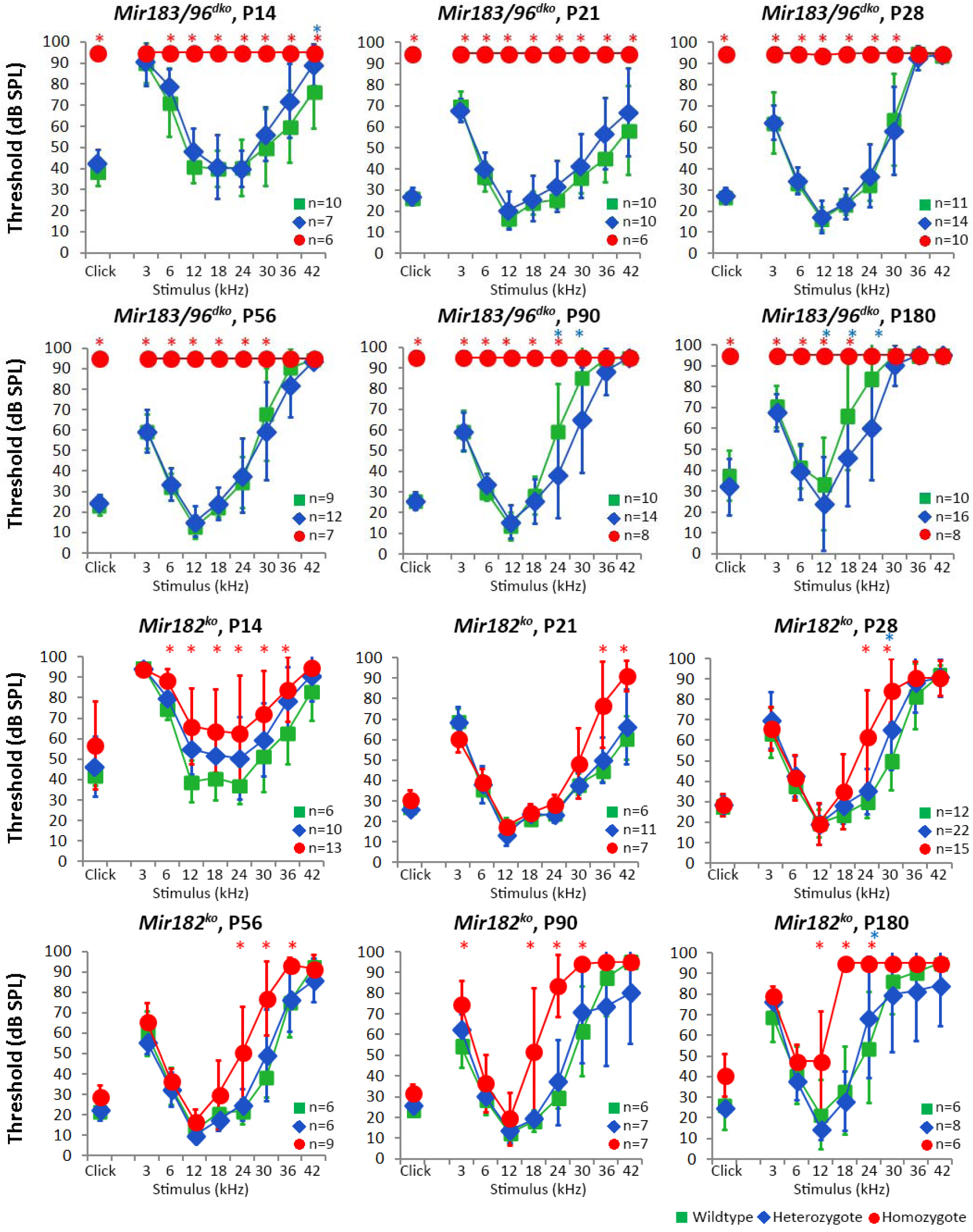
Mean ABR thresholds of *Mir183/96^dko^* and *Mir182^ko^* homozygous (red circles), heterozygous (blue diamonds) and wildtype (green squares) mice at P14, P21, P28, P56, P90 and P180. Homozygous *Mir183/96^dko^* mice show profound hearing loss at all ages tested, and points plotted at 95dB indicate no response at this level, the maximum SPL used. Homozygous *Mir182^ko^* mice display mildly raised thresholds at high frequencies, which slowly progresses to include the middle frequencies as the mice age. Heterozygotes from both lines show thresholds similar to wildtype mice. Number of mice of each genotype tested at each age is shown on the threshold plot. Error bars are standard deviation, and significant differences (P < 0.05, mixed linear model pairwise comparison) are marked by * in red (for a significant difference between wildtype and homozygote) or blue (for a significant difference between wildtype and heterozygote). See Supplementary Figures S2, S3 for individually plotted traces, and the supplementary data for p-values for all comparisons.

For the *Mir183*/*96^dko^* mice, 43 out of 242 mice (17.8%) from heterozygote by heterozygote matings were homozygous for the *Mir183*/*96* null allele, which is lower than the expected production (25%), suggesting that the absence of *Mir96* and/or *Mir183* has a small impact on viability (p=0.029, chi-squared test). For the *Mir182^ko^* mice, 42 homozygotes out of 152 pups in total (27.6%) were produced from heterozygote by heterozygote matings, which is consistent with the mutation having no impact on viability.

### Complete knockout of miRNA expression in the mutant organ of Corti

We carried out qPCR to test the expression levels of the three microRNAs in the organs of Corti of wildtypes, heterozygotes and homozygotes of each knockout. In *Mir183/96^dko^* homozygotes, there was no detectable expression of *Mir183* or *Mir96*. Likewise, in *Mir182^ko^* homozygotes there was no detectable expression of *Mir182* (Supplementary Fig. S1). The levels of expression in heterozygotes of both knockouts was variable, as was the expression of *Mir182* in *Mir183/96^dko^* homozygotes, and *Mir183* and *Mir96* in *Mir182^ko^* homozygotes. It is likely that this is because we lack a proper microRNA control for sensory tissue in the inner ear. We used *Mir99a*, which is expressed in almost all cell types in the cochlea, including hair and supporting cells [10], but because *Mir183*, *Mir182* and *Mir96* are also expressed in hair cells, it is possible that the mutant alleles affect the expression of *Mir99a* in hair cells, making it an unreliable calibrator between wildtype, heterozygote and homozygote. A better calibrator would be a microRNA expressed only in supporting cells and not in hair cells.

### Impaired auditory responses in homozygotes but normal thresholds in heterozygotes

*Mir183/96^dko^* homozygous mice were profoundly deaf, with most showing no response at the highest sound level tested (95dB SPL) at any of the ages tested (14 days to six months old). In contrast, *Mir182^ko^* homozygotes exhibited only a mild hearing loss starting at higher frequencies and progressing with age to lower frequencies (Fig. 1). The ABR thresholds of heterozygotes were normal at all ages tested (Fig. 1, see Supplementary Figures S2, S3 for individually plotted traces). ABR waveforms of *Mir183/96^dko^* heterozygotes and *Mir182^ko^* homozygotes were also similar to those of wildtype littermates at the equivalent sound pressure level above threshold (sensation level, SL) (Supplementary Fig. S4). We measured DPOAEs at 8 weeks old and found no difference in the amplitudes or thresholds between wildtype and heterozygous *Mir183/96^dko^* mice, while homozygotes had severely abnormal responses (Supplementary Fig. S6). *Mir182^ko^* homozygotes had raised DPOAE thresholds at high frequencies compared to wildtypes (Supplementary Fig. S6), which matched the difference in their ABR thresholds at 8 weeks (Fig. 1). *Mir182^ko^* mutant mice showed no sign of a vestibular defect (circling, head-bobbing or hyperactivity) up to six months old. However, *Mir183/96^dko^* homozygotes did show increasing incidence of hyperactivity with age (Supplementary Fig. S5).

### Heterozygous *Mir183/96^dko^* mice recover normally from noise exposure

As heterozygous *Mir183/96^dko^* mice showed no auditory deficit, in contrast to the hearing loss seen in diminuendo heterozygotes and in humans carrying one mutant *MIR96* allele, we asked if these heterozygous mice might be more sensitive to noise-induced damage. One day after noise exposure (2 hours of 8-16kHz octave-band noise at 96dB SPL) at 8 weeks old, both *Mir183/96^dko^* heterozygous and wildtype mice showed a marked increase in thresholds at 12kHz and above compared to unexposed control littermates (Fig. 2). Three days after noise exposure the 12kHz thresholds had recovered, but there was still a noticeable elevation at higher frequencies. By 7 days after exposure, all thresholds had recovered completely (Fig. 2). We measured the amplitude of wave 1 of the ABR waveform to look for a reduced neural response, which has been reported in CBA/CaJ mice after noise exposure [21] and is thought to be due to neuronal loss in the cochlea, but no difference was observed at 12 kHz (Fig. 2). At 24kHz, we observed a much greater effect, such that in most animals wave 1 was too poorly defined to measure the amplitude one day after exposure. However, both wildtype and heterozygote wave 1 amplitudes had recovered by 28 days after exposure, and did not look any different to mice which had not been exposed (Fig. 2; Supplementary Fig. S7).

**Figure 2.**
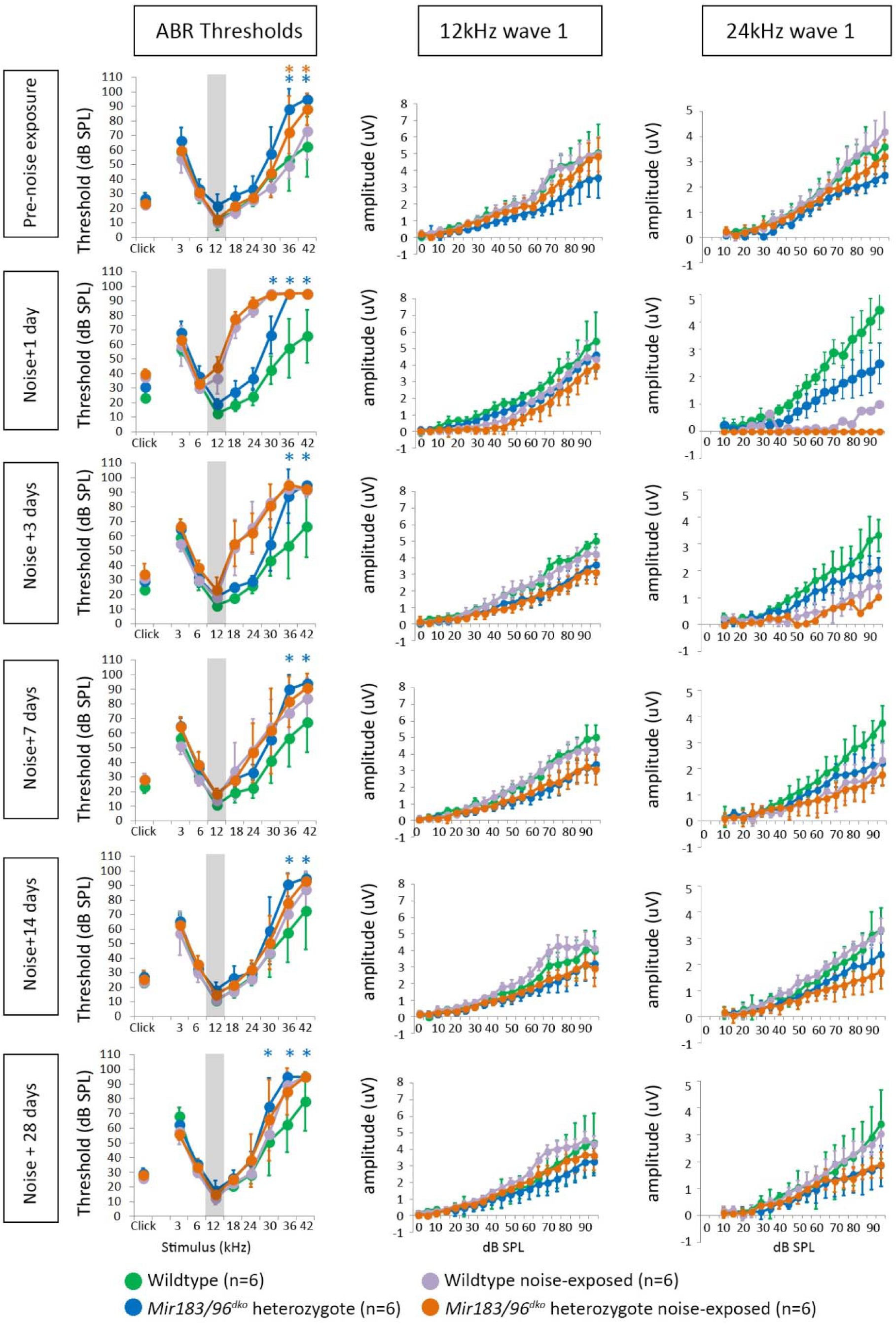
Mean ABR thresholds for wildtype and heterozygous *Mir183/96^dko^* mice subjected to noise exposure (95dB SPL, 8-16kHz, for two hours), before exposure and 1, 3, 7, 14 and 28 days after. An increase in thresholds at 12kHz and above is seen 1 day after exposure but thresholds have returned to normal by 28 days after. Significant differences (Bonferroni-corrected p < 0.05, mixed linear model pairwise comparison) are marked by * in blue (for a significant difference between wildtype unexposed mice and heterozygous unexposed mice) or orange (for a significant difference between wildtype noise-exposed mice and heterozygous noise-exposed mice). Please see the supplementary data for p-values for all comparisons. Wave 1 amplitudes at 12kHz and 24kHz are shown for each time point. No obvious effect is visible at 12kHz, but at 24kHz, one day after noise exposure, wave 1 was too poorly defined to measure the amplitude in all heterozygotes and all but one wildtype. By 28 days after exposure, both wildtype and heterozygote amplitudes have recovered to the normal range. 12 wildtype mice and 12 heterozygote mice were tested; 6 wildtypes were noise-exposed (violet) with 6 unexposed controls (green), and 6 heterozygotes were noise-exposed (orange) with 6 unexposed controls (blue). Error bars are standard deviation. The grey area on the threshold plots indicates the octave band of noise (8-16kHz).

Since there was a significant difference in the higher frequencies between the unexposed heterozygotes and wildtypes at 8 weeks old (Fig. 2, top left panel), but we did not see any difference in our original ABR tests (Fig. 1, P56), we compared the ABR thresholds from all mice tested at 8 weeks old (Supplementary Fig. S8). At high frequencies (30-42kHz) the thresholds were very variable in both heterozygotes and wildtypes. The differences in the means at 36kHz and 42kHz are statistically significant (P = 0.001, P=0.000; mixed linear model pairwise comparison), but given the variability we suggest that this is not biologically relevant.

### Hair cells and innervation

We examined the organ of Corti using scanning electron microscopy and found that hair cells in homozygous *Mir183/96^dko^* mice were severely affected at four weeks old (Fig. 3), with many hair bundles missing entirely. Where present, the stereocilia bundles of both outer and inner hair cells show splaying and fusion. The inner hair cells of *Mir183/96^dko^* heterozygotes are unaffected, but the outer hair cells’ upper surface appear slightly smaller and rounder in shape than the wildtype outer hair cells (Fig. 3). Their stereocilia bundles also appear smaller and more rounded than normal, with more pronounced tapering in height and overlap of shorter stereocilia rows with taller rows towards the two ends of each bundle.

**Figure 3.**
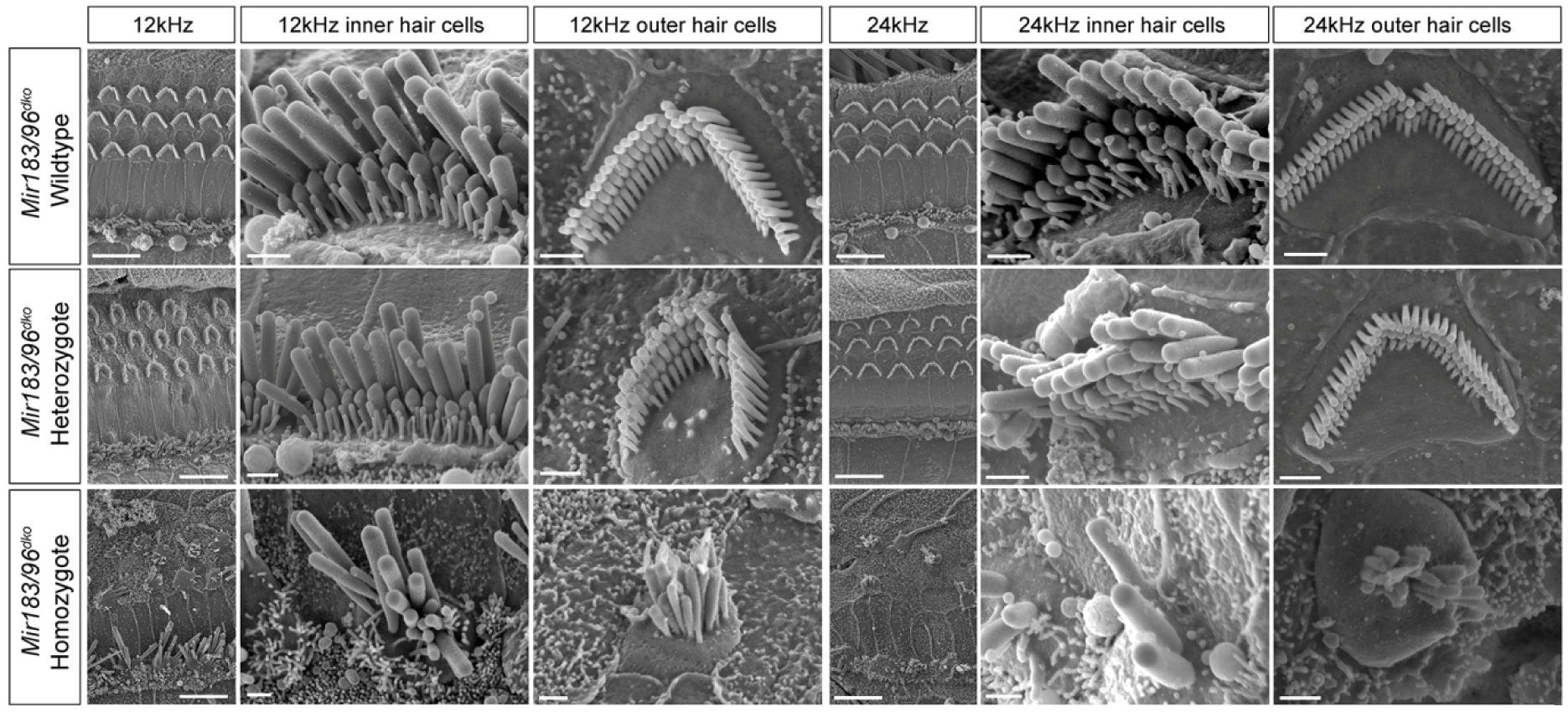
Scanning electron micrographs of *Mir183/96^dko^* mice at P28. Two best-frequency regions of the organ of Corti are shown; 12kHz (68% of the way along the organ of Corti from base to apex) and 24kHz (43% of the way along the organ of Corti from base to apex). For each region, the left-hand column shows a zoomed-out image with inner and outer hair cell rows (scale bars=10µm), and the other two columns show an inner and an outer hair cell close up (scale bars=1µm). The top row shows wildtype hair cells (n=6), the middle row shows heterozygote hair cells (n=6) and the bottom row shows homozygote hair cells (n=5).

Four week old *Mir182^ko^* heterozygotes and homozygotes showed no abnormalities of hair cells by scanning electron microscopy at either the 12kHz or 24kHz regions (Supplementary Fig. S9), corresponding to their normal ABR thresholds at that age. At eight weeks old, when hearing loss is evident at 24kHz and higher (Fig. 1), we also saw no systematic differences between wildtypes, heterozygotes and homozygotes (Fig. 4).

**Figure 4.**
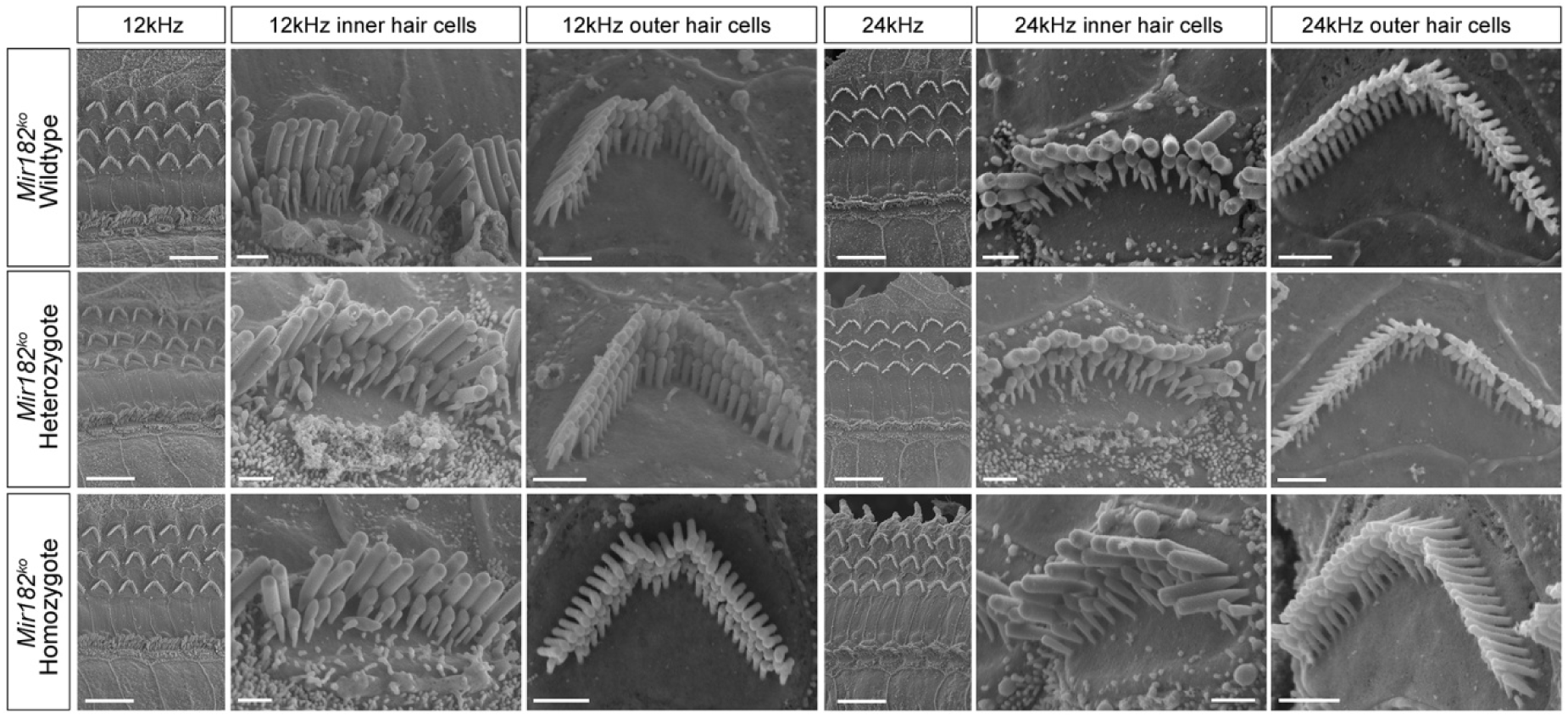
Scanning electron micrographs of *Mir182^ko^* mice at P56. Two best-frequency regions of the organ of Corti are shown; 12kHz (68% of the way along the organ of Corti from base to apex) and 24kHz (43% of the way along the organ of Corti from base to apex). For each region, the left-hand column shows a zoomed-out image with inner and outer hair cell rows (scale bars=10µm), and the other two columns show an inner and an outer hair cell close up (scale bars=1µm). The top row shows wildtype hair cells (n=1), the middle row shows heterozygote hair cells (n=2) and the bottom row shows homozygote hair cells (n=3).

The distribution of unmyelinated neurons appeared normal in both mutants using anti-neurofilament labelling (Supplementary Fig. S10). Synapses were examined using anti-Ribeye antibody to mark presynaptic ribbons and anti-GluR2 to mark postsynaptic densities, and a significant reduction was found in the number of colocalised pre- and postsynaptic markers in *Mir183/96^dko^* homozygotes (Fig. 5, p=0.016, one way ANOVA). No difference in synapse counts was observed in *Mir182^ko^* homozygotes.

**Figure 5.**
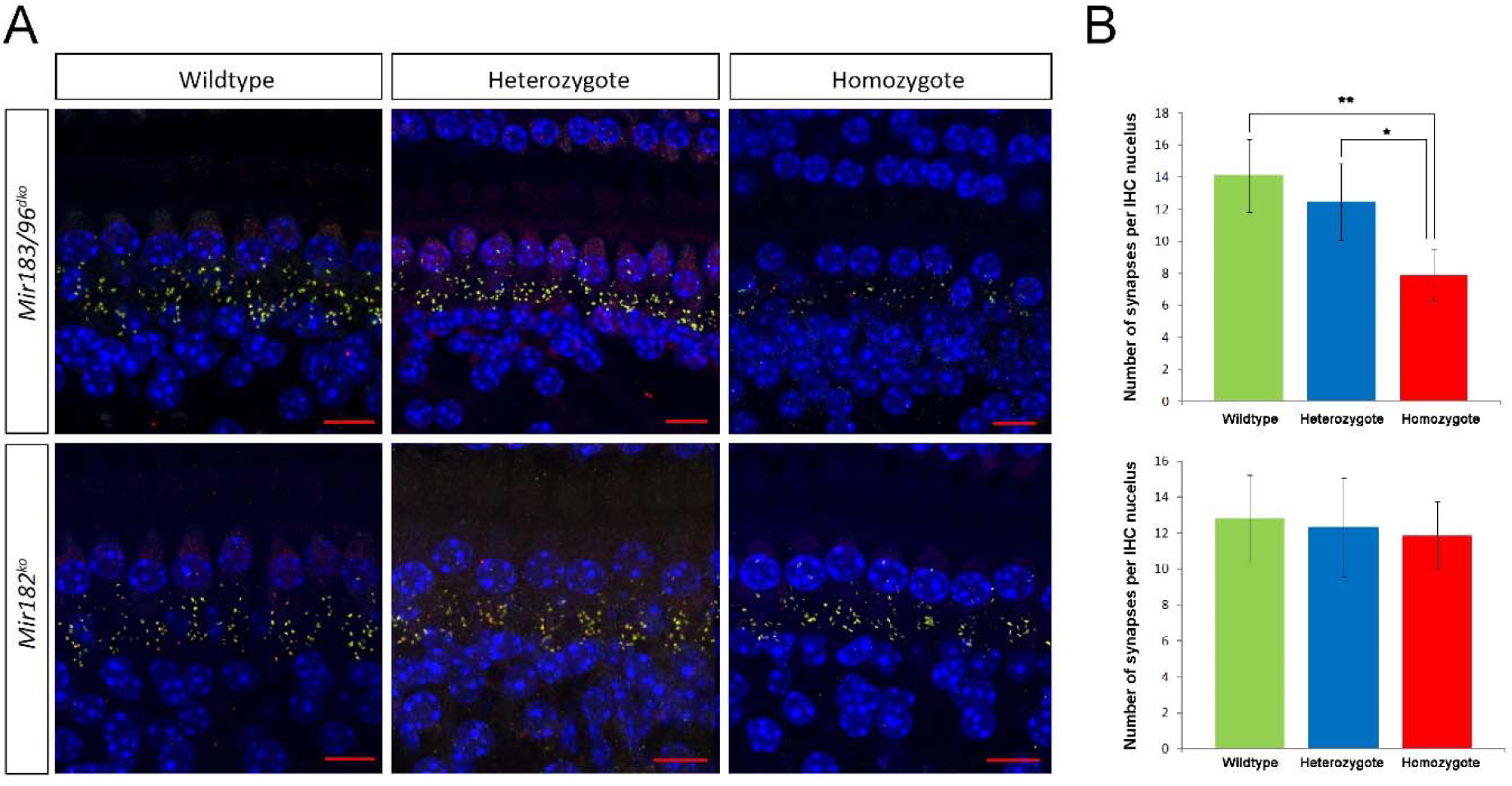
Colocalised pre- and postsynaptic densities in *Mir183/96^dko^* and *Mir182^ko^* mice. A. Synapses below inner hair cells (IHCs) in *Mir183/96^dko^* (top) and *Mir182^ko^* (bottom) wildtype (left), heterozygous (middle) and homozygous (right) knockout mice. Presynaptic ribbons are labelled with anti-Ribeye antibody (red) and postsynaptic densities with anti-GluR2 antibody (green); where they colocalise, the resulting colour is yellow. DAPI (blue) labelled the nuclei. Scale bar = 10µm. B. Mean counts of colocalised pre- and postsynaptic markers in wildtype (green), heterozygous (blue) and homozygous (red) *Mir183/96^dko^* (top) and *Mir182^ko^* (bottom) mice. There are significantly fewer colocalised synapses in *Mir183/96^dko^* homozygotes (n=3) compared to wildtypes (n=5, p = 0.016, one way ANOVA, Bonferroni-corrected p=0.02**) and also compared to heterozygotes (n=7, Bonferroni-corrected p=0.035*), but no significant differences in heterozygotes compared to wildtypes (Bonferroni-corrected p=1.0). No significant difference was seen between synapse counts in *Mir182^ko^* knockout heterozygotes (n=6), homozygotes (n=7) and wildtypes (n=4,p=0.818, one way ANOVA). Error bars show standard deviation.

### Transcriptome analysis reveals misregulation of gene expression in mutants

To investigate the impact of the *Mir182^ko^* and *Mir183/96^dko^* mutations on gene expression we carried out RNA-seq of isolated organ of Corti preparations from postnatal day (P)4 homozygotes and sex-matched littermate wildtype controls. This age was chosen to ensure all hair cells were still present and to facilitate comparison with our previous transcriptome data from *Mir96^Dmdo^* mice [3]. Thirty-four genes were identified as significantly misregulated (FDR < 0.05) in *Mir183/96^dko^* homozygotes, 22 upregulated and 12 downregulated compared with wildtype littermates. Many of the upregulated genes have sequences complementary to either the miR-96 seed region or the miR-183 seed region in their 3’UTRs (Table 1). Of this list of 34 genes, only *Hspa2*, *Ocm, Myo3a*, *Slc26a5*, *Slc52a3*, *St8sia3* and *Sema3e* were previously found to be misregulated in *Mir96^Dmdo^* mice at P4 and/or P0 [3, 16], and in each case the misregulation is in the same direction (Table 1). We tested 19 genes of the 34, selected because they were reported to show a large difference in expression levels between sensory and non-sensory cells in the organ of Corti (http://www.umgear.org; [22, 23]). All but three were confirmed correct (Table 1, Supplementary Fig. S11); those that were not confirmed by qPCR were either not misregulated or failed the significance test, and no genes were significantly misregulated in the opposite direction. Of the genes misregulated in *Mir183/96^dko^* homozygotes, five are known deafness genes; *Myo3a*, *Slc26a5* and *Tmc1* underlie deafness in mice and humans [24–29], while mutations in *Sema3e* cause deafness in people [30], and mice homozygous for a mutant allele of *Ocm* exhibit progressive hearing loss [31].

**Table 1.**
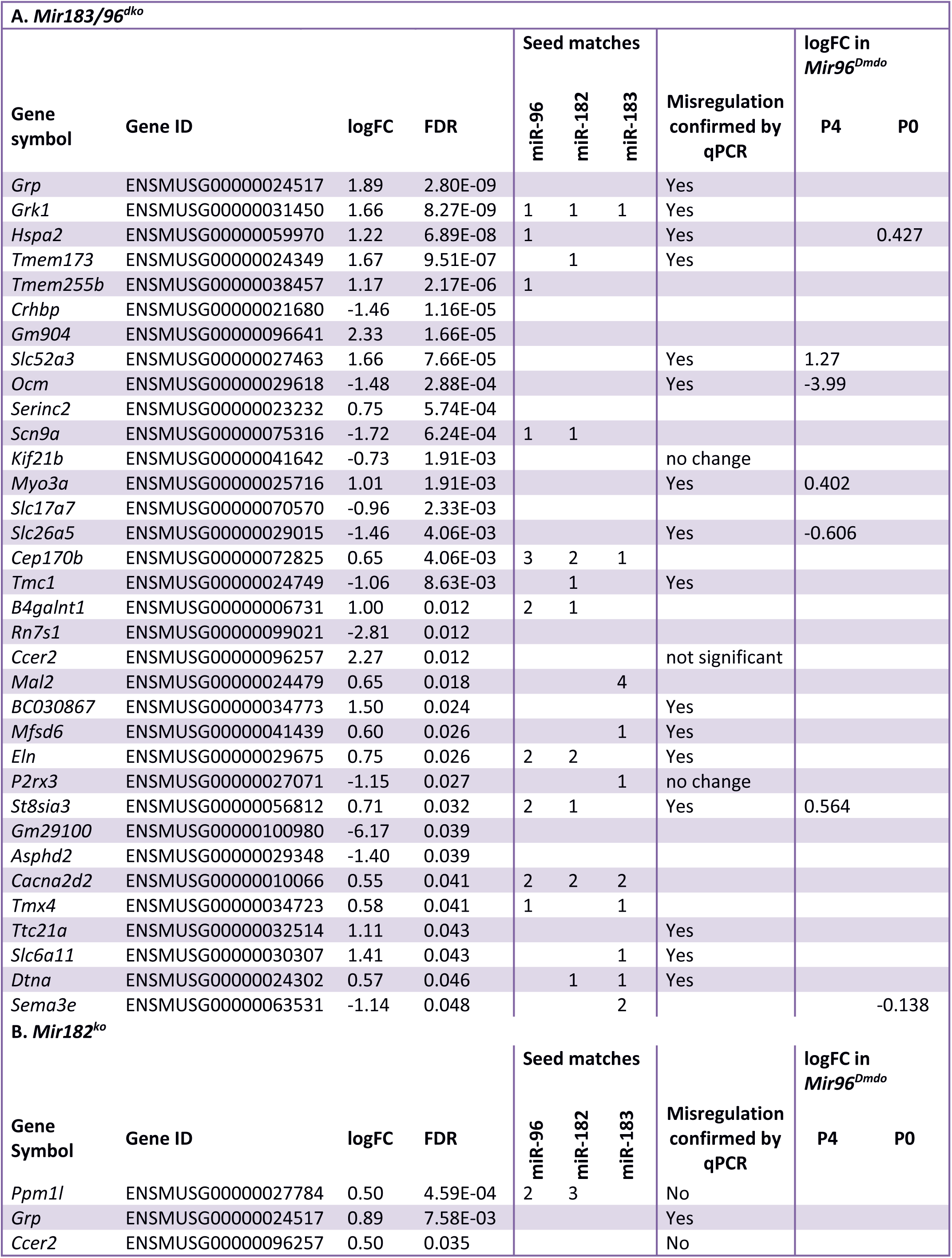
Genes with a false discovery rate (FDR) < 0.05 are shown for *Mir183/96^dko^* RNA-seq (top) and *Mir182^ko^* RNA-seq (bottom). The presence of sequence complementary to the three microRNA seed regions in the 3’UTR of each gene is shown, as is the significant fold change observed in *Mir96^Dmdo^* mice at either P4 or P0. Confirmation by qPCR indicates that the gene was shown by qPCR to be significantly misregulated in the same direction as found in the RNA-seq data.

It is possible that the difference between the transcriptomes of *Mir183/96^dko^* knockout mice and *Mir96^Dmdo^* mice is due to the different backgrounds. We looked for differences in predicted targets in the 3’UTRs of the C57BL/6NJ and C3H/HeJ genome sequences [32], which were the closest genome sequences to our mutant backgrounds available, and found that 1585 genes had the same number of seed matches, and 13 genes had seed matches in both strains, but not the same number of matches. 36 genes had no seed matches in the C57BL/6NJ sequence but had one or more match in the C3H/HeJ sequence, and 47 genes had no seed matches in the C3H/HeJ sequence, but one or more in the C57BL/6NJ sequence (Supplementary Table S1).

Three genes were found to be significantly upregulated (FDR < 0.05) in the *Mir182^ko^* homozygotes by RNA-seq, one of which, *Ppm1l*, has sequences complementary to the seed region of miR-182 (Table 1). The other two, *Grp* and *Ccer2*, were also upregulated in the *Mir183/96^dko^* homozygotes. No genes were significantly downregulated in *Mir182^ko^*. We tested the upregulated genes by qPCR, and also tested *Slc26a5* and *Ocm* (which were strongly downregulated in *Mir96^Dmdo^* homozygotes [3]) and found that only the upregulation of *Grp* was confirmed. *Ccer2* and *Ppm1l* were upregulated but not significantly, and *Slc26a5* and *Ocm* were downregulated but again, not significantly (Table 1, Supplementary Fig. S11).

To assess the impact of these miRNA knockouts on a genome-wide level, we used Sylamer [33] to measure the enrichment and depletion of all possible heptamers in the 3’UTRs of each total gene list, ranked from most upregulated to most downregulated irrespective of significance. In the *Mir183/96^dko^* gene list, the sequence complementary to the seed region of miR-96 was markedly enriched in the upregulated genes (red line, Fig. 6A), and the sequence complementary to the seed region of miR-183 has a small peak towards the centre of the graph (dark blue line, Fig. 6A). While the targets of miR-183 are not notably misregulated in this dataset, its signal is still distinct from all other miRNAs (blue line, Fig. 6A). There were no miRNA seed region heptamers enriched in the *Mir182^ko^* gene list (Fig. 6B), but the TATTTAT heptamer which is enriched in the *Mir182^ko^* downregulated genes (yellow line, Fig. 6B) resembles a portion of an AU-rich element. These are 50-150bp sequence elements found in 3’UTRs, and are typically involved in mRNA destabilising via a deadenylation-dependent mechanism (reviewed in [34, 35]). AU-rich elements, and TATTTAT in particular, are enriched in 3’UTR regions which have multiple RNA-binding protein binding sites, and it has been suggested that RNA-binding proteins compete with the RISC complex to bind to miRNA target sites within these regions [36]. It is possible that this TATTTAT signal, which is enriched in the genes downregulated in *Mir182^ko^* homozygote hair cells, is the result of a change in the binding of RNA-binding proteins in the absence of miR-182.

**Figure 6.**
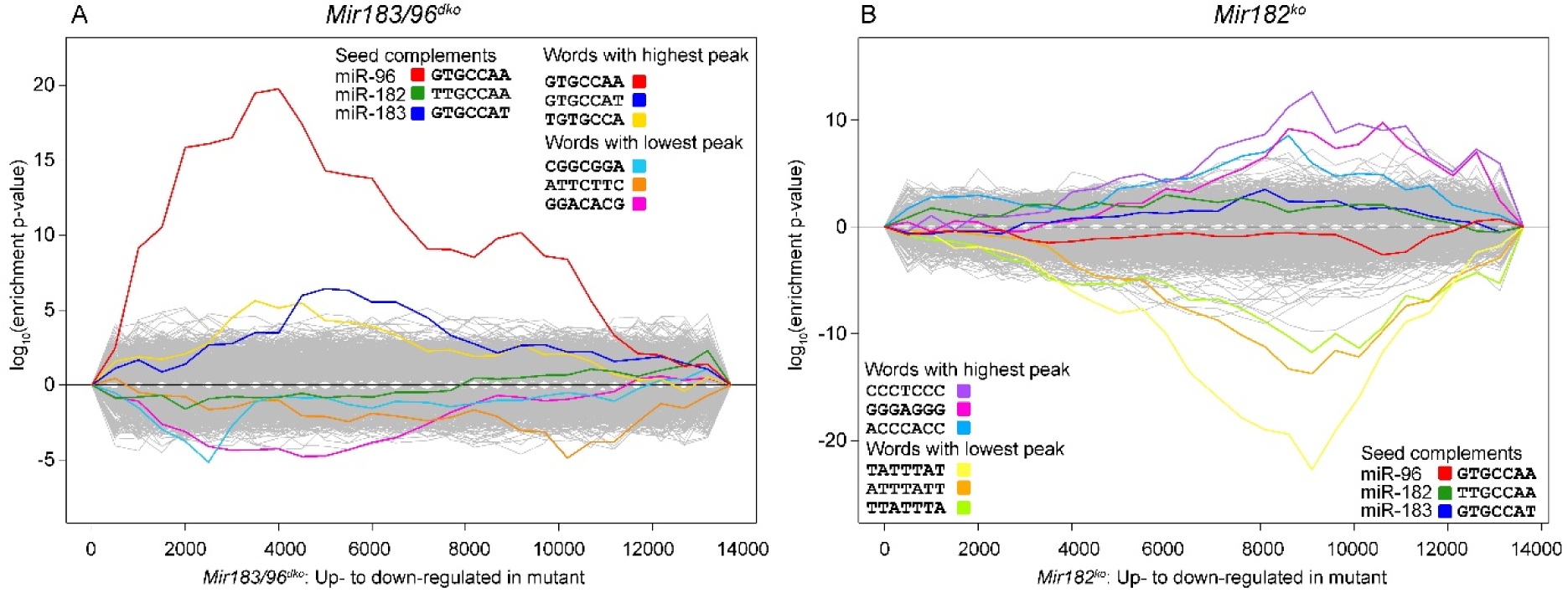
Sylamer analysis showing enrichment and depletion of heptamers in 3’UTRs in *Mir183/96^dko^* homozygous mice (A) and *Mir182^ko^* homozygous mice (B). The x-axis represents the sorted gene list from most up-regulated on the left to most down-regulated on the right. The y-axis shows the hypergeometric significance for enrichment or depletion of heptamers in 3’UTRs. UTRs are considered in bins, starting with the 500 most up-regulated genes on the left and increasing cumulatively until all genes have been considered. Lines indicate the enrichment of each heptamer; positive values indicate enrichment and negative values indicate depletion. The three words with the highest peaks and the three with the lowest peaks are highlighted in colour, as are the three heptamers complementary to the seed regions of miR-96, miR-182 and miR-183. The yellow line in the *Mir182^ko^* plot on the right, enriched in the downregulated genes of *Mir182^ko^* homozygotes, resembles a portion of an AU-rich element. This may reflect a change in protein binding to 3’UTRs in the absence of miR-182 (see Results for details).

### No evidence for differential splicing

Splicing patterns can be affected by a microRNA if it targets a splicing factor, as has been shown for miR-133a and nPTB [37], so we analysed the *Mir183/96^dko^* and *Mir182^ko^* RNA-seq data for evidence of alternative splicing events. Tools which test RNA-seq data for differential splicing of mRNAs are not as well established as those for measuring differential expression, and there is great variability in the results depending on which method is used [38]. We therefore used three different tools to test for differential splicing, which all take a different approach. Cuffdiff (from the Cufflinks package [39]) maps reads to individual transcripts in order to estimate the relative abundance of each isoform. This can introduce uncertainty in the case of reads which map to exonic regions shared by more than one transcript. JunctionSeq [40], is a count-based tool which tests read counts belonging to exonic regions specific to individual transcripts, avoiding the problem of shared regions but potentially missing isoforms which have no unique exons. Leafcutter, a very recently developed algorithm, tests for differential splicing by using reads which span exon-exon junctions, in effect focussing on the excised intron [41].

All three tools detected significant differential splicing in *Mir183/96^dko^* homozygotes compared to their wildtype littermates (Supplementary Table S2), but only three genes were identified by more than one tool; *Ppp3cb* and *Zfp618* (by Leafcutter and JunctionSeq), and *Il17re* (by Leafcutter and Cuffdiff). One of the advantages of Leafcutter and JunctionSeq is that they identify the sites at which they detect differential splicing occurring, which allowed us to test their results. We designed primers to cover the junctions and/or exons which were highly differentially spliced (15 genes; Supplementary Table S3) and sequenced PCR reactions from cDNA made from organ of Corti RNA from *Mir183/96^dko^* homozygotes and wildtypes. We found no evidence of the differential splicing detected by either tool, but evidence of a novel isoform of *Stard9* was detected by JunctionSeq. The novel isoform includes an exon (ENSMUSE00001437951) previously assigned to a transcript subject to nonsense-mediated decay (transcript ENSMUST00000140843), but the exons following ENSMUSE00001437951 in our results belong to the protein-coding transcript (ENSMUST00000180041). Exon ENSMUSE00001437951 is 72bp long, so does not introduce a frameshift, and its inclusion may result in a functional Stard9 protein with 24 extra amino acids. The novel splicing was present in both wildtype and homozygous *Mir183/96^dko^* RNA (Supplementary Fig. S12).

JunctionSeq did not predict any significant differential splicing in *Mir182^ko^* homozygotes. Leafcutter and Cuffdiff identified some genes with differential splicing (Supplementary Table S2), but there were no genes common to both predictors. Two genes, *Ube4a* and *Nrxn1*, were predicted to have alternative splicing present in the homozygote and not in the wildtype, but when we sequenced their mRNA from *Mir182^ko^* homozygotes and wildtypes, we found no differences in the splicing pattern.

### Immunohistochemistry confirms downregulation of *Ocm*

We carried out antibody stains on sections from the inner ear at P4, to check for the presence of Ocm (oncomodulin) and Slc26a5 (prestin) protein in the hair cells. Ocm staining is faint at P4, stronger in the basal turn of the cochlea, but prestin staining is clearly visible at that stage in wildtypes. We observed prestin staining in *Mir183/96^dko^* homozygotes, but no stain for Ocm, while both proteins were present in the wildtype littermate controls (Supplementary Fig. S13). Although immunohistochemistry is not a quantitative technique, this correlates with the qPCR results, which showed that *Ocm* RNA was nearly absent in *Mir183/96^dko^* homozygotes, while *Slc26a5* RNA levels were about 30% of wildtype levels (Supplementary Fig. S11). Ocm and Prestin staining was visible in *Mir182^ko^* homozygotes and their wildtype littermate controls (Supplementary Fig. S13).

### Network analysis suggests upstream regulators

We took three approaches to network analysis: Ingenuity Pathway Analysis, WGCNA, and network creation using existing publicly available regulatory data. First, we used Ingenuity Pathway Analysis to construct putative causal networks using the misregulated genes in each dataset with FDR<0.05, as follows. Upstream regulators which could explain the observed misregulation were identified through the Ingenuity Knowledge Base, a manually curated set of observations from the literature and from third-party databases. The activation or inhibition of each upstream regulator was calculated based on the observed misregulation of its target genes. Causal networks were then constructed to link the upstream regulators through a “root” regulator further upstream, again using the Ingenuity Knowledge Base (described in detail in [42]). The resulting networks were filtered based on their activation z-score, which is a measure of the match of observed and predicted misregulation patterns. The z-score is not just a measure of significance but also a prediction of the activation of the upstream root regulator [42]; scores below -2 are classed as significant by Ingenuity Pathway Analysis and indicate predicted downregulation or reduced activation of the root regulator, while scores above 2 are also considered significant and indicate predicted upregulation or increased activation of the root regulator. For the *Mir183/96^dko^* homozygotes, only four upstream regulators were given a z-score < -2, three more had z-scores < −1.7 and one had a z-score > 1.7. These eight upstream regulators, Bdnf, Cshl1, H2afx, Kcna3, Kit, Myocd, Prkaca and Pth, explain between them the misregulation of 14 genes (Supplementary Fig. S14), although they themselves are not misregulated (from the RNA-seq data). Most of the genes are predicted to be inhibited, so they cannot be directly linked to miR-96 (although Bdnf is a known target of miR-96 [43], which does not fit the predicted inhibition). Only Kcna3 is predicted to be activated (orange in Fig. 7), and it does not have any matches to the miR-96 or miR-183 seed regions in its 3’ UTR. Of these eight potential regulators, *Bdnf* and *Kit* are both known deafness genes [44, 45]. There were too few genes significantly misregulated in the *Mir182^ko^* RNA-seq data to obtain any causal networks with a significant activation z-score, but Ret and Adcyap1 were identified as immediate upstream regulators of Grp.

**Figure 7.**
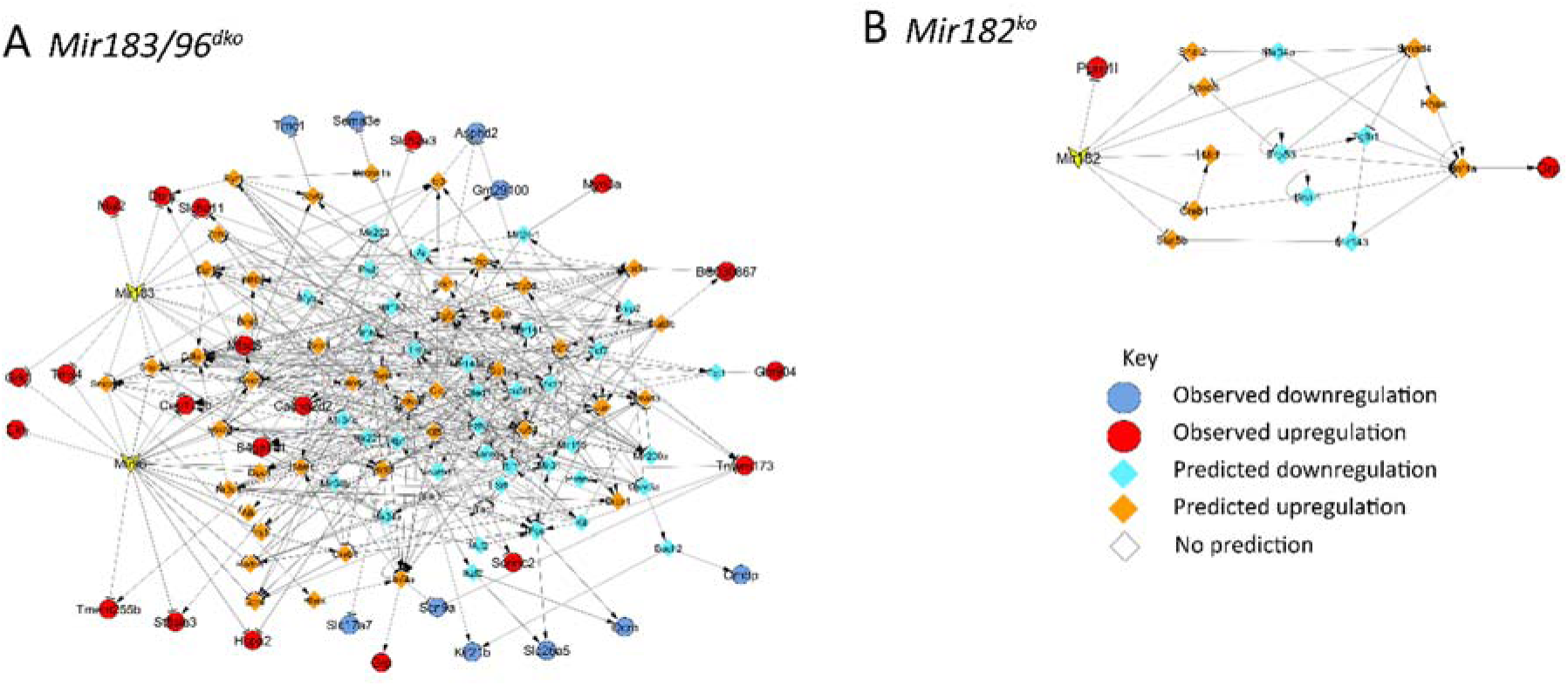
Networks drawn up from publicly available regulatory data, based on the *Mir183/96^dko^* (A), and *Mir182^ko^* (B) transcriptome data. Red and orange indicate upregulation and blue and turquoise downregulation; circles with black borders show known misregulation in the organ of Corti and diamonds without borders show predicted misregulation.

### Gene clustering using weighted gene correlation network analysis (WGCNA) suggests further regulators

Weighted gene correlation network analysis is a method of analysing transcriptome data to cluster genes into modules based on their expression across a number of individual samples without reference to the sample traits [46]. We used a Pearson correlation to cluster genes across all 24 samples (*Mir183/96^dko^* and *Mir182^ko^*) and obtained 29 consensus modules (including the reserved “grey” module, which consists of genes outside of all the other modules) (Supplementary Table S4).

These consensus modules are groups of genes with highly correlated expression profiles. For further analysis, we obtained each module’s eigengene, which represents all the genes within the module, and we calculated the correlation of each eigengene with the traits of the mice used for RNA-seq (Supplementary Fig. S15). We then clustered the eigengenes to identify meta-modules where eigengenes were highly correlated with each other (Supplementary Fig. S16). Of the consensus module eigengenes, three were highly correlated and clustered with the wildtype vs. *Mir183/96^dko^* homozygote trait (green, black, royal blue, Supplementary Fig. S16A). This means that the expression levels of the genes of the green, black and royal blue modules are correlated with each other and the genotypes of the *Mir183/96^dko^* mice. One module was highly correlated and clustered with the wildtype vs. *Mir182^ko^* homozygote trait (salmon, Supplementary Fig. S16B)

We chose nine modules for further exploration which had significant correlation with the wildtype vs. *Mir183/96^dko^* homozygote or wildtype vs. *Mir182^ko^* homozygote traits (Supplementary Fig. S15; black, green, royal blue, salmon, purple, grey60, light green and dark green modules). We also investigated the blue module, which contained the most differentially expressed genes (13, Supplementary Table S4)). We carried out enrichment analysis using PANTHER v14 [47], comparing the module genes to the GO biological process listing [48, 49] and the Reactome pathway database [50]. We found five of the modules had significant enrichment of GO and/or Reactome gene sets, defined as enrichment score ≥ 5 and corrected p-value (FDR) < 0.05. Genes in the black module were enriched for GO terms such as cytoplasmic translational initiation (GO:0002183) and positive regulation of mRNA catabolic process (GO:0061014), suggesting genes in this module are involved in mRNA processing. Genes in the light green module are also enriched in GO terms involved in translation (eg negative regulation of cytoplasmic translation (GO:2000766) and negative regulation of transcription by competitive promoter binding (GO:0010944)). Green module genes were enriched for GO terms involved in cell metabolism, such as mitochondrial respiratory chain complex I assembly (GO:0032981) and ATP synthesis coupled proton transport (GO:0015986), and the Reactome pathway enrichment shows a similar set of terms (eg Formation of ATP by chemiosmotic coupling (R-MMU-163210)). Blue module genes were enriched for just two GO terms (inner ear receptor cell fate commitment (GO:0060120) and auditory receptor cell fate commitment (GO:0009912)) and one Reactome pathway (Regulation of lipid metabolism by Peroxisome proliferator-activated receptor alpha (PPARalpha) (R-MMU-400206)). The purple module genes are enriched for the Reactome Rho GTPase pathways and for GO terms involved in development and transcription (eg neural tube closure (GO:0001843), regulation of neuron apoptotic process (GO:0043523) and negative regulation of transcription by RNA polymerase II (GO:0000122)) (see Supplementary Tables S5, S6 for full listings of enriched GO terms and Reactome pathways).

We also investigated potential transcription factors which could be responsible for the altered regulation of the genes within each module using oPOSSUM [51], which detects over-represented transcription factor binding sites in a set of genes. We obtained 29 transcription factor binding site profiles across the 9 modules with Z and Fisher scores above our chosen threshold of the mean + 1 standard deviation (Supplementary Fig. S17, Supplementary Table S4) [51]. Several of the profiles were shared between modules (Supplementary Fig. S18). Transcription factors implicated in the black, green, blue and light green modules include one deafness gene (*Foxi1* [52]) and several genes with miR-183/96/182 seed region matches, although none have yet been experimentally shown to be targets of any of the three microRNAs (Supplementary Table S7). Transcription factors implicated in the purple, royal blue and dark green modules also include genes with seed region matches, and Myc, which is also a deafness gene [53], is a potential regulator of the genes in the grey 60 module (Supplementary Table S7). None of the transcription factors were found to be misregulated in the RNA-seq data.

### Networks created using publicly available regulatory data

The IPA networks did not connect the misregulated genes to any of the three microRNAs, probably because each microRNA controls a regulatory cascade, and the IPA causal networks have a maximum depth of 3, which may not be enough to reach the level of direct targets. The WGCNA analysis identified modules of co-expressed genes and highlighted transcription factors which may be involved, but did not connect these directly to either the microRNAs or the misregulated genes.

In our previous study on genes misregulated in *Mir96^Dmdo^*, we used regulatory interactions described in the literature to create an internally consistent network of regulatory interactions connecting miR-96 to as many of the misregulated genes as possible [16]. For the current study, we automated the procedure using a perl script (available at https://github.com/moraglewis/PoPCoRN) and made use of publicly available regulatory data in addition to the manually curated regulatory data from the literature we compiled before.

The resulting *Mir183/96^dko^* network consists of 114 genes and 416 links. Thirty misregulated genes have been included, 13 of them predicted targets of either miR-183 or miR-96 or both. Two misregulated genes were left out completely due to having no known upstream regulators in our compiled regulatory interactions (*Ccer2*, and *Rn7s1*) (Fig. 7A). We carried out a network analysis using the Cytoscape Network Analyser tool to calculate the degree and betweenness centrality of each node (gene). A node’s degree is the number of edges connecting it to other nodes, and its betweenness centrality measures how important it is for connecting distant parts of the network. The node with the highest degree is *Trp53*, with 23 edges. Most nodes have 9 or fewer edges (Fig. 7A), but 34 have 10 or more edges, including seven which are direct targets of miR-96, miR-183 or both (listed in Table 2). None of these 34 nodes with 10 or more edges are known to be misregulated in *Mir183/96^dko^* homozygotes, but eight are known deafness genes (*Fos* [54], *Foxo3* [55], *Kit* [45], *Mir96*, *Nfkb1* [56], *Pkd1* [57], *Rest* [58] and *Tnf* [59]. Among these 34 nodes are *Trp53*, *Sp1*, *Tgfb1*, *Tnf* and *Fos*, which were also genes of interest in the manually-created network based on the *Mir96^Dmdo^* microarray data [16].

**Table 2.** Nodes with high edge count in the *Mir183/96^dko^* network.

| Name | Deafness<br>genes | microRNA<br>target | Betweenness<br>Centrality | Edge<br>count |
| --- | --- | --- | --- | --- |
| Trp53 |  |  | 0.09422711 | 23 |
| Foxo1 |  | Yes (both) | 0.08571448 | 21 |
| Sp1 |  |  | 0.06173317 | 18 |
| Stat5a |  |  | 0.03490919 | 17 |
| Jun |  |  | 0.04868463 | 16 |
| Nfkb1 | Yes |  | 0.03928189 | 16 |
| Hnf4a |  |  | 0.04198241 | 15 |
| Fos | Yes |  | 0.04456005 | 13 |
| Tnf | Yes |  | 0.05881021 | 12 |
| Tgfb1 |  |  | 0.02574521 | 12 |
| Mir155 |  |  | 0.01078878 | 12 |
| E2f2 |  |  | 0.04550675 | 11 |
| Pkd1 | Yes |  | 0.02115304 | 11 |
| Tcf7 |  |  | 0.02033637 | 11 |
| Kit | Yes |  | 0.0124918 | 11 |
| Ezh2 |  |  | 0.03284497 | 10 |
| Zfp36 |  |  | 0.02575902 | 10 |
| Mir200a |  |  | 0.02321144 | 10 |
| Elk1 |  |  | 0.01517501 | 10 |
| Cdkn1a |  | Yes (miR-96) | 0.03321781 | 23 |
| Mir96 | Yes |  | 0.04831501 | 22 |
| Rest | Yes |  | 0.03424156 | 17 |
| Mir183 |  |  | 0 | 16 |
| Egr1 |  | Yes (miR-183) | 0.0752374 | 15 |
| Stat5b |  |  | 0.00432024 | 14 |
| Nr3c1 |  | Yes (miR-96) | 0.04511961 | 13 |
| Bcl6 |  |  | 0.03888551 | 13 |
| Foxo3 | Yes | Yes (miR-96) | 0.02891802 | 13 |
| Mir141 |  |  | 0.04075138 | 12 |
| Smad4 |  | Yes (both) | 0.05817905 | 11 |
| Mir145a |  |  | 0.02869746 | 11 |
| Mir182 |  |  | 0.02760045 | 11 |
| Cebpa |  |  | 0.00923269 | 11 |
| Zeb1 | Yes | Yes (both) | 0.03369301 | 10 |

The *Mir182^ko^* network is very small, being based on three input genes: *Ppm1l*, *Ccer2* and *Grp*. *Ccer2* could not be included, due to the lack of any known regulators, and *Ppm1l* is a predicted direct target of miR-182. Multiple potential pathways link *Grp* to miR-182, but all of them work through *Hnf4a*, which upregulates *Grp* (Fig. 7B).

To validate the network analyses, we carried out qRTPCR on 11 genes from the *Mir183/96^dko^* network and two from the *Mir182^ko^* network (Fig. 7). Our choice was based on several factors, including their importance within the network, their edge count and close links to important genes (eg *Ocm* and *Slc26a5*). *Fos* and *Trp53* were also predicted to be involved by the IPA analysis (Supplementary Fig. S14). However, we found all the genes had very variable expression levels in homozygotes when compared to littermate wildtypes (Supplementary Fig. S11). It’s possible that this is the result of expression in different areas of the inner ear. We controlled for the quantity of sensory tissue using *Jag1*, but the inner ear is a complex organ with many cellular subtypes, and if a gene is expressed in multiple cell types, controlling for overall tissue quantity (using *Hprt*) and sensory tissue quantity (using *Jag1*) may not be sufficient to permit detection of small changes in mRNA levels in the sensory tissue. In addition, qRTPCR can only detect mRNA levels, and can’t directly measure protein levels or protein activity.

## Discussion

### *Mir183*/*96^dko^* mice have a less severe phenotype than *Mir96^Dmdo^* mice

Mice heterozygous for the *Mir96^Dmdo^* point mutation exhibit early-onset rapidly progressive hearing loss; even at P15 they have very raised thresholds [8]. We initially concluded that this was more likely to be due to haploinsufficiency than to the gain of novel targets, because humans heterozygous for different point mutations also had progressive hearing loss [4]. However, heterozygous *Mir183/96^dko^* mice have ABR thresholds and DPOAE responses resembling the wildtype, and there is no difference in thresholds or wave 1 amplitudes between heterozygotes and wildtypes after noise exposure. Two recent studies of different knockout alleles targeting the entire microRNA cluster (miR-183/96/182) found that heterozygotes also exhibited normal hearing [60, 61]. It is possible that the more severe phenotype seen in *Mir96^Dmdo^* heterozygotes is due to the acquisition of new targets by the mutant microRNA, or it could be an effect of the different background as the *Mir96^Dmdo^* allele was generated by ENU mutagenesis on a C3HeB/FeJ background in contrast to the C57BL/6N background of the *Mir183/96^dko^* allele or the reported knockouts of the entire cluster, which were created in 129S2 [62] or 129SV6 [60] ES cells which were then crossed onto the C57BL/6J background [60, 62]. Mice homozygous for the *Mir183/96^dko^* allele showed no auditory brainstem responses at all ages tested from 14 days old onwards, and in this they resemble the *Mir96^Dmdo^* homozygotes, where the compound action potentials recorded from the round window of the cochlea were undetectable at four weeks old [3].

The vestibular phenotype is also different; *Mir96^Dmdo^* homozygotes circle by three weeks old, whereas a milder phenotype of variable hyperactivity was observed only in some *Mir183/96^dko^* homozygotes, the prevalence increasing with age (Supplementary Fig. S5). Circling is a more severe manifestation of vestibular dysfunction than hyperactivity. This could again be due to the C3HeB/FeJ background, because *Mir96^Dmdo^* mice carry the *Pde6b^rd1^* mutation causing retinal degeneration [63] and are blind by adulthood. The *Mir183/96^dko^* allele was generated on a C57BL/6N background, which lacks the *Pde6b^rd1^* mutation but has the *Crb1^rd8^* mutation, which leads to variably penetrant retinal dysplasia [64]. Other ocular abnormalities have also been observed in C57BL/6N mice, such as lens abnormalities and vitreous crystalline deposits [64]. However, no ocular abnormalities have been reported for these mice in the IMPC pipeline (https://www.mousephenotype.org/data/genes/MGI:3619440), so it is likely that they have sufficient vision to partially compensate for the lack of vestibular sensory input, reducing the severity of the observed phenotype. It is notable that mice lacking the entire miR-183/96/182 cluster (on a mixed background of C57BL/6J and either 129S2 or 129SV6) exhibit both persistent circling behaviour and retinal defects [60, 62].

### Stereocilia bundles and innervation in *Mir183*/*96^dko^* mice and *Mir96^Dmdo^* mice

The hair cells of *Mir96^Dmdo^* homozygous mice are present at four days old but appear abnormal, and by 28 days old have degenerated almost completely [3]. In *Mir183/96^dko^* homozygotes, however, some hair cell stereocilia bundles are still visible at P28, although they are severely disorganised (Fig. 3). In *Mir183/96^dko^* heterozygotes, IHC stereocilia are mostly normal and the OHC stereocilia bundles appear to be slightly rounded in arrangement (Fig. 3), but this is not reflected in their ABR thresholds or DPOAE responses, which are normal (Fig. 1 and Supplementary Fig. S5). *Mir96^Dmdo^* heterozygotes, which are deaf by P28, have a more severe phenotype, with fused stereocilia and hair cell degeneration as well as misshapen stereocilia bundles in outer hair cells, and smaller stereocilia bundles in inner hair cells [3].

*Mir96^Dmdo^* homozygotes exhibit disorganised innervation [8], but we did not see that degree of disorganisation in neurofilament-labelled preparations of *Mir183/96^dko^* homozygotes (Supplementary Fig. S10). However, we found significantly fewer colocalised pre- and postsynaptic densities under inner hair cells of *Mir183/96^dko^* homozygotes indicating synaptic defects (Fig. 5). Heterozygote *Mir183/96^dko^* mice showed no differences in innervation or synapse counts compared to wildtypes (Supplementary Fig. S10, Fig. 5). Similar to the physiological phenotype, the structural phenotype of *Mir183/96^dko^* heterozygotes is much less severe than that of *Mir96^Dmdo^* heterozygotes.

### The *Mir183/96^dko^* transcriptome shows fewer genes are affected than in the *Mir96^Dmdo^* transcriptome

The *Mir183/96^dko^* transcriptome bears some resemblance to that of the *Mir96^Dmdo^* transcriptome from our previous studies [3, 16], but, as with the physiological and structural phenotypes, the effect of missing both miR-183 and miR-96 appears to be milder than the effect of a point mutation in the miR-96 seed region. Only 34 genes were identified as significantly misregulated by RNA-seq in the current study of *Mir183/96^dko^*, compared to 86 genes found to be significantly misregulated in the *Mir96^Dmdo^* P4 microarray [3], and only seven genes are similarly misregulated in both datasets: *Hspa2*, *Ocm, Myo3a*, *Slc26a5*, *Slc52a3*, *St8sia3* and *Sema3e* (Table 1). Some of the differences will be due to certain genes not being represented on the microarray assay when it was carried out, such as *Ccer2*, and some may be the result of the different genetic background. We aimed to minimise this by use of the sex-matched wildtype littermates as controls, but it isn’t possible to entirely eliminate the effect of genetic background; there are 83 genes which lack miR-96 seed region matches in one background compared to the other (Supplementary Table S1), and any of those may be contributing to the phenotype. Much of the difference, however, is likely to be due to the loss of miR-183 in the *Mir183/96^dko^* homozygotes and the presence of the mutant miR-96 in *Mir96^Dmdo^* homozygotes. We were unable to confirm any of the predicted differential mRNA splicing in *Mir183/96^dko^* homozygotes, suggesting that differential splicing is unlikely to play a large part in the mutant phenotypes.

### Network analyses of the *Mir183/96^dko^* transcriptome suggest multiple potential regulators

We took three approaches to find intermediate regulators, and obtained 8 upstream regulators from the Ingenuity Pathway Analysis causal network analysis, 29 transcription factor profiles from the WGCNA module analyses (Supplementary Table S7), and 34 highly connected nodes from our regulatory data-based network construction (Table 2). There are no genes suggested by all three approaches, although each one suggests multiple plausible candidates, such as *Fos*, *Kit*, *Foxi1*, *Myc*, *Zeb1* and *Foxo1*. However, when we tested a selection of intermediate genes, we found their expression was very variable between different homozygotes (Supplementary Fig. S11). This was true even of *Ikzf2*, which is known to directly regulate *Ocm* and *Slc26a5* [65], two critical genes for outer hair cell function which are strongly downregulated in both *Mir96^Dmdo^* [3, 16] and *Mir183/96^dko^*.

### Identifying candidate direct targets of miR-96 in hair cells

The only network approach which suggested direct targets of miR-96 in the inner ear was our regulatory data-based approach, which identifies candidate targets that have been experimentally confirmed. There were 21 direct targets of miR-96 in total, 10 of which were genes found to be upregulated in the RNA-seq data. We used the gEAR dataset comparison tool [66] and data from mouse hair cells compared to the rest of the cochlear duct at p0 [22] to identify which of the 21 direct targets were excluded from hair cells but present in the rest of the cochlear duct. We found eight which showed this pattern of expression: *Snai2*, *Zeb1*, *Irs1*, *Nr3c1*, *Foxo1*, *Alk*, *Eln* and *Rad51*. Three of these are known deafness genes: *Snai2*, which is involved in melanocyte development [67], *Zeb1*, which is required for the specification of mesenchymal identity and repression of epithelial identity [68], and *Irs1* [69]. The precise role of Irs1 in the function of the cochlea has not yet been elucidated, but since Zeb1 and Snai2 are known to be required for the development of non-sensory cells in the cochlear duct, it may be that miR-96 plays a role in establishing and/or maintaining the repression of non-hair cell genes in developing hair cells.

*Nr3c1* and *Foxo1* have the most downstream links of the 21 direct targets, predicted to regulate 9 and 12 genes respectively*. Foxo1* encodes a forkhead family transcription factor, and the *Nr3c1* gene encodes the glucocorticoid receptor GR, which is known to be expressed in the inner ear [70, 71]. No hearing or vestibular phenotypes have been reported on the Mouse Genome Informatics resource (http://www.informatics.jax.org) [72] for mice carrying mutations in either gene, but it is likely that the hearing of *Nr3c1* mutants has simply never been checked. The hearing of *Foxo1* knockout heterozygotes has been tested through the IMPC phenotyping pipeline and they were found to have normal hearing (http://www.mousephenotype.org, [73]) but since the knockout is homozygous lethal, there is no data for the effect of the absence of *Foxo1* on the inner ear.

Only one of these eight genes was found to be upregulated in *Mir183/96^dko^* homozygotes in the RNA-seq data. *Eln* has two matches to the miR-96 seed region in its 3’UTR, and is thus a potential direct target of miR-96. It is a connective tissue protein and a component of the extracellular matrix [74], and it plays a regulatory role in controlling vascular smooth muscle cells via a GPCR pathway [75], but its downstream targets, and its role in the inner ear, are not yet known.

### The gain of novel targets plays an important role in the phenotype resulting from a point mutation in *Mir96*

We originally hypothesised that the major effect of the *Mir96^Dmdo^* mutation on hearing was most likely mediated by a reduced level of downregulation of the normal target mRNAs [3], because two different point mutations in the human *MIR96* seed region also led to hearing loss but each of the three seed region point mutations are predicted to have different novel targets. However, the difference in the transcriptomes, along with the less severe phenotype including normal ABR thresholds of *Mir183/96^dko^* heterozygotes, suggest that the gain of novel target mRNAs is also important for the diminuendo phenotype. From the microarray carried out on *Mir96^Dmdo^* P4 organ of Corti, we found 19 genes which were significantly downregulated in the mutant and which bore matches to the mutant seed region in their 3’ UTR [3]. The list includes one known deafness gene, *Ptprq* [76], as well as *Chrna1*, which is expressed in the organ of Corti from early postnatal stages onwards [77]. The hair cells of mice homozygous for a null allele of *Ptprq* closely resemble those seen in the *Mir96^Dmdo^* homozygotes and heterozygotes at P4 [7], so it’s possible that the more severe phenotype seen in *Mir96^Dmdo^* homozygotes is due in part to the downregulation of *Ptprq* by the mutant miR-96.

### A model for mechanisms of action of mutant microRNAs

It has been suggested that miRNAs act in two ways; first, they repress targets to prevent translation, resulting in mutually exclusive expression of the miRNA and the target, and second, they buffer transcriptional noise, in which case the miRNA and its targets are co-expressed in the same cell [78]. Our approach should be able to detect both effects, but if a target is highly expressed in the non-sensory epithelial cells, the difference in expression between wildtype and homozygote due to derepression in the hair cells may not be detectable. Our current transcriptome data, therefore, is more likely to highlight targets which are being buffered rather than targets which are completely repressed by miR-96 [68]. This may explain why likely target genes like *Zeb1*, *Foxo1* and *Nr3c1* have not been found to be significantly misregulated in our transcriptome data. It may also explain the variability of the network genes we tested (Fig. S11E,F) - in the absence of the miRNA buffering, transcriptional noise has increased. This effect would be exacerbated by a mutated seed region, such as in the *Mir96^Dmdo^* mutant; not only would the normal buffering effect be gone, but multiple other genes would be misregulated within the hair cell, further disrupting normal cell function, as indeed we observed in the *Mir96^Dmdo^* transcriptome analyses [3, 16].

We suggest that the consistent misregulation of *Ocm*, *Slc26a5*, *Myo3a*, *Sema3e* and *Slc52a3* observed in both the *Mir96^Dmdo^* [3, 16] and *Mir183/96^dko^* homozygotes is the result of the lack of repression of genes that would usually not be expressed in hair cells at all, for example *Zeb1*, *Foxo1*, and *Nr3c1*. The links between the direct targets of miR-96 and the consistently misregulated downstream genes are yet to be discovered, but we suggest *Ikzf2* and *Fos* are likely to be involved (Fig. 8A, B).

**Figure 8.**
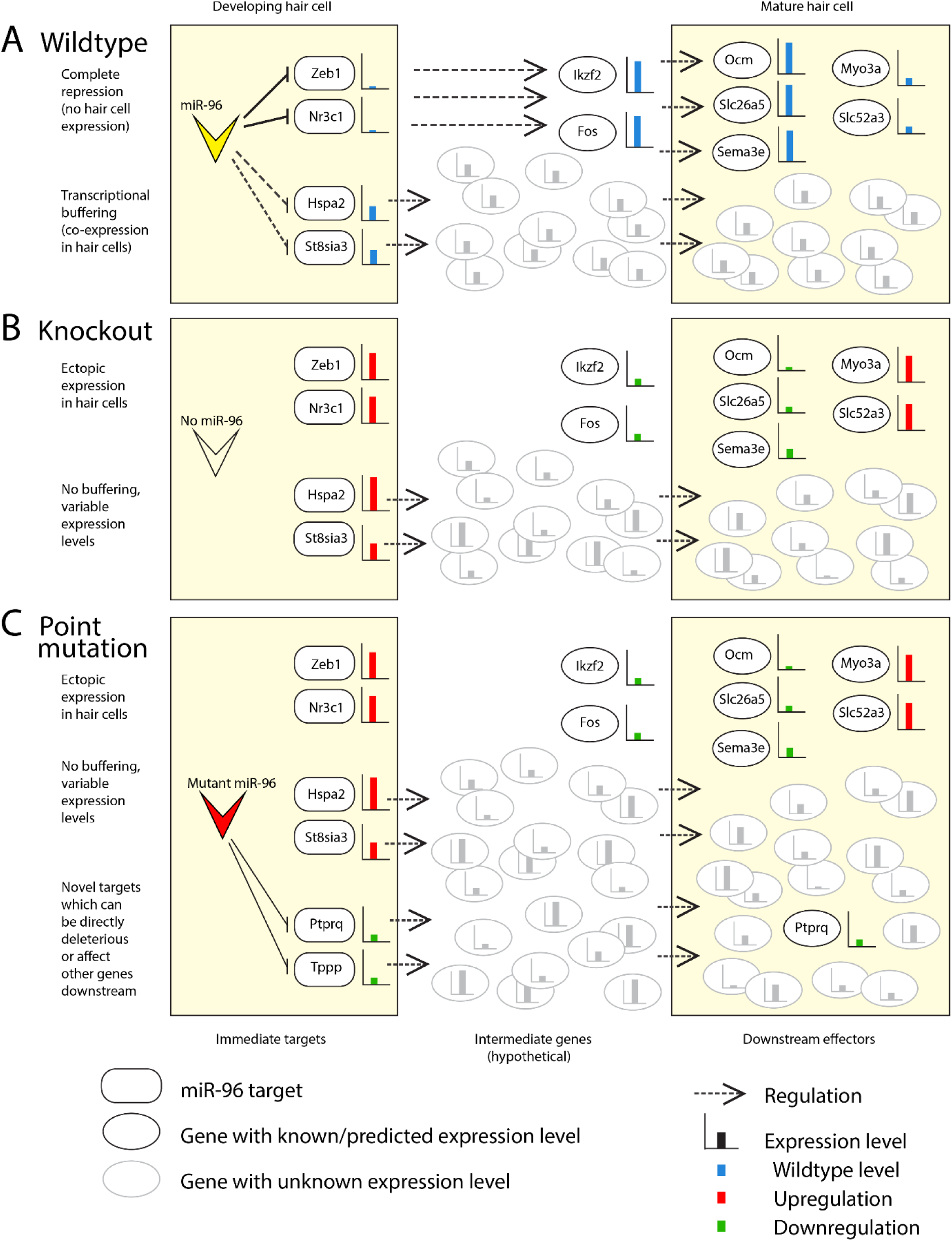
Diagram of the modes of action of mutant microRNAs in the outer hair cell. A) In a wildtype hair cell, miR-96 represses some genes completely (here represented by *Zeb1* and *Nr3c1*) and buffers the expression levels of other genes (*Hspa2*, *St8sia3*). The result is a mature, functional outer hair cell with wildtype expression levels of important genes (eg *Ocm*, *Slc26a5*). B) In the absence of miR-96, as seen in the *Mir183/96^dko^* homozygote, there is both ectopic expression of targets such as *Zeb1* and *Nr3c1*, and variable expression of the buffered targets such as *Hspa2* and *St8sia3*. This leads to very variable expression levels of many genes (shown in grey), and the downregulation of genes critical for outer hair cell function, such as *Ocm* and *Slc26a5*. C) If the microRNA bears a point mutation, as in the *Mir96^Dmdo^* homozygote, then a third effect comes into play, namely the downregulation of genes which bear a match to the mutant seed region in their 3’UTR. In the case of *Mir96^Dmdo^*, this includes *Ptprq*, which is an essential gene for stereocilia bundle development [76]. Many other genes (shown in grey) will also be misregulated through the gain of novel targets, further contributing to the functional failure of the outer hair cell. Expression levels are indicated by bar charts; blue bars show wildtype expression levels in (A), while in (B) and (C), red bars indicate upregulation and green bars indicate downregulation.

There are multiple genes which are known or potential miR-96 targets, are upregulated in miR-96 mutants, and are expressed in hair cells, such as *Hspa2*, *St8sia3* and *Grk1*. We suggest these are examples of genes whose expression is buffered by miR-96, not fully repressed but maintained at a consistent level. The loss of this buffering results in variable expression and transcriptional noise, not just of these genes (which will all be upregulated to varying degrees) but also of any genes which they regulate. This transcriptional noise is less likely to be consistent (at the gene level) between mice but will nonetheless contribute to the degraded functionality of the hair cell (Fig. 8A, B).

Finally, genes which are novel targets of a mutant microRNA have the potential to fulfil both roles. In the case of *Mir96^Dmdo^*, *Ptprq* is downregulated and it’s possible this is due to the mutant microRNA, since *Ptprq* bears a complementary match to the *Mir96^Dmdo^* seed region in its 3’ UTR. A reduction in *Ptprq* in the hair cells will contribute directly to the failure of the hair cells to mature properly. Other novel targets may also be directly affecting the hair cells, or may be adding to the overall transcriptional noise, or both (Fig. 8C).

### The role of *Mir183* in hearing remains unclear

The lack of a *Mir183*-specific mutation means that the effect of miR-183 alone is difficult to ascertain from our data. The relatively low mid-range peak for the miR-183 seed region in our Sylamer analysis of the RNA-seq data (Fig. 6A) implies that the lack of miR-183 has less of a global effect on the transcriptome than the lack of miR-96, and the literature-based regulatory network analysis reflects this; apart from its predicted targets, there are no downstream genes whose misregulation can be attributed to miR-183 alone (Fig. 7A). In that network, the only miR-183 targets which regulate downstream genes are also miR-96 targets.

### Mice lacking *Mir182* display mild hearing loss with no obvious changes in stereocilia bundles or hair cell innervation

The *Mir182^ko^* heterozygotes have audiograms which resemble the wildtype, while the homozygotes exhibit mild hearing loss at the higher frequencies (Fig. 1). No difference in hair cell stereocilia bundles, innervation or synapse counts was observed between the *Mir182^ko^* wildtype, heterozygous and homozygous mice at P28 (Supplementary Figures S9, S10, Fig. 5), but at that age *Mir182^ko^* homozygotes still have normal hearing. However, even at P56, when *Mir182^ko^* homozygotes exhibit high frequency hearing loss, the hair cells appear unaffected (Fig. 4). This is a completely different phenotype from either the *Mir96^Dmdo^* or the *Mir183/96^dko^* mice and may explain the paucity of significantly misregulated genes in the RNA-seq data from the P4 organ of Corti. This may be too early to see much effect of the absence of miR-182 on the transcriptome. The lack of significantly enriched heptamers corresponding to the miR-182 seed region in the Sylamer analysis for miR-182 also supports this hypothesis (Fig. 6B). The network drawn from the *Mir182^ko^* transcriptome may show the start of the misregulation cascade due to the microRNA knockout, and the transcription factors suggested by the WGCNA and IPA analyses may be involved, but further studies at a later age would be required for confirmation. None of the predicted differential mRNA splicing in *Mir182^ko^* homozygotes was confirmed.

### Relevance to human *MIR96* mutations

In this study we have shown that the phenotype of mice lacking miR-96 entirely is less severe than that of mice carrying a point mutation in the miR-96 seed region. Mice heterozygous for the *Mir183/96^dko^* allele display no auditory phenotype when compared to wildtypes, while *Mir96^Dmdo^* heterozygotes have rapid, severe progressive hearing loss. In our first study of the diminuendo point mutation, we concluded that the phenotype caused by the mutant miR-96 was most likely the result of loss of normal miR-96 targets, because all three mutations described (two in humans and one in the mouse) resulted in hearing loss [3, 4]. Our data from the current study, however, suggest that the gain of novel targets also plays an important role in the phenotype caused by mutations in miR-96. This has important implications for understanding the effect of mutant microRNAs in the human population. So far all three reported human mutations in miR-96 have been point mutations, two in the seed region and one in the stem region of the pre-miRNA [4, 5], and while all the individuals carrying those point mutations have exhibited some degree of progressive hearing loss, the phenotypes differ between people, as do the phenotypes of the mice carrying the *Mir96^Dmdo^* mutation and the *Mir183/96^dko^* allele. Determining the misregulated pathways common to all *Mir96* mutations will be important not only for furthering our understanding of the role of this microRNA in hair cell development but also for developing therapeutic interventions, not only for people with *MIR96* mutations but also more generally for hearing loss associated with hair cell defects.

## Materials and Methods

Ethics approval: Mouse studies were carried out in accordance with UK Home Office regulations and the UK Animals (Scientific Procedures) Act of 1986 (ASPA) under UK Home Office licences, and the study was approved by both the Wellcome Trust Sanger Institute and the King’s College London Ethical Review Committees. Mice were culled using methods approved under these licences to minimize any possibility of suffering.

### Mice

The miR-183/96 and miR-182 knockouts in C57BL/6N derived JM8.A3 ES cells were generated as previously described [18]. On chromosome 6 regions 30169424bp to 30169772bp (NCBIM38) for miR-182 and 30169424bp to 30169772bp (NCBIM38) for miR-183/96 were replaced with the PuroΔTK selection cassette. For both knockouts the PuroΔtk gene was subsequently deleted by transient transfection with Cre recombinase leading to recombination of the loxP sites that flanked the selection marker under 2-Fluoro-2-deoxy-1D-arabinofuranosyl-5-iodouracil (FIAU) selection. Heterozygous targeted ES cells were microinjected into C57BL/6N embryos for chimaera production. Mice were maintained on a C57BL/6N background. Both lines are available through EMMA (*Mir183/96^dko^* mice: EM:10856; *Mir182^ko^* mice: EM:12223).

### Auditory Brainstem Response

The hearing of *Mir183/96^dko^* and *Mir182^ko^* homozygote, heterozygote and wildtype littermates was tested using the Auditory Brainstem Response, following the protocol described in [79]. Animals were sedated using a ketamine/xylazine mix (10mg ketamine and 0.1mg xylazine in 0.1ml per 10g body weight) and recovered where possible using atipamezole (0.01mg atipamezole in 0.1ml per 10g body weight) to enable recurrent testing of the same cohort of mice. All injections were intraperitoneal. Responses were recorded from three subcutaneous needle electrodes placed one over the left bulla (reference), one over the right bulla (ground) and one on the vertex (active). We used a broadband click stimulus and 3kHz, 6kHz, 12kHz, 18kHz, 24kHz, 30kHz, 36kHz and 42kHz pure tone frequencies, at sound levels from 0-95dB, in 5dB steps. 256 stimulus presentations per frequency were carried out per frequency and sound level, and responses were averaged to produce the ABR waveform. The threshold for each stimulus, the lowest intensity at which a waveform could be distinguished visually, was identified using a stack of response waveforms. Mice were tested at 14 days old (P14), P21, P28 ±1 day, P56 ±2 days, P90 ±2 days and P180 ±3 days. Any hyperactivity was noted prior to anaesthesia. Wave 1 amplitudes were calculated using ABR Notebook software (courtesy of MC Liberman, Harvard Medical School/Massachusetts Eye and Ear).

### Distortion Product Otoacoustic Emission (DPOAE) measurements

We measured DPOAEs in mice aged 8 weeks old, anaesthetised with an intraperitoneal injection of 0.1ml / 10g of a solution of 20% urethane. Experiments were performed using Tucker Davis Technologies (TDT) BioSigRZ software driving a TDT RZ6 auditory processor and a pair of TDT MF1 magnetic loudspeakers. Signals were recorded via an Etymotic ER-10B+ low-noise DPOAE microphone. Stimulus tones (f1 & f2) were presented and microphone signals recorded via a closed-field acoustic system sealed into the auditory meatus of the mouse. Stimulus tones were presented at an f2:f1 ratio of 1.2. f2 tones were presented at frequencies to match ABR measurements (6, 12, 18, 24, 30 and 36 kHz). f1 was presented at levels from 0–85 dB in 5dB steps. f2 was presented at 10 dB below the level of f1. The magnitude of the 2f1-f2 DPOAE component was extracted from a fast Fourier transform of the recorded microphone signal and plotted as a function of f2 level. For each f2 level, the 20 spectral line magnitudes surrounding the 2f1-f2 frequency were averaged to form a mean noise floor estimate for each measurement. DPOAE threshold was defined as the lowest f2 stimulus level where the emission magnitude exceeded 2 standard deviations above the mean noise floor.

### Noise exposure

Wildtype and heterozygous *Mir183/96^dko^* mice (P55±1 day) were subjected to an 8-16kHz octave-band noise at 96dB SPL for two hours while awake and unrestrained in separate small cages within an exposure chamber designed to provide a uniform sound field (for chamber details, see [80]). Band pass noise was generated digitally using TDT RPvdsEx software, converted to an analogue signal using a TDT RZ6 auditory processor, and amplified using a Brüel and Kjær Type 2716C power amplifier. It was delivered to a compression driver (JBL 2446H, Northridge, CA) connected to a flat front biradial horn (JBL 2380A, Northridge, CA) secured to the roof of the sound box. ABRs were carried out the day before, and 1, 3, 7, 14 and 28 days after the noise exposure. Unexposed littermates were used as controls and went through the same set of ABR measurements.

### Genotyping

*Mir183/96^dko^* knockout mice were genotyped by PCR analysis using primers spanning the introduced deletion (Supplementary Table S3). The wildtype band is 841bp and the mutant band 645bp. *Mir182^ko^* mice were genotyped in a similar fashion with one of two primer sets (Supplementary Table S3). The wildtype band for the first set is 495bp and the mutant band 457bp. The wildtype band for the second set is 247bp, and the mutant band is 209bp.

### Scanning electron microscopy

The inner ears of wildtype, heterozygote and homozygote mice at P28 (*Mir183/96^dko^*, *Mir182^ko^*) and P56 (*Mir182^ko^*) were fixed in 2.5% glutaraldehyde in 0.1M sodium cacodylate buffer with 3mM CaCl_2_ at room temperature for two hours. Cochleae were finely dissected in PBS and processed according to the OTOTO method [81]. Samples were dehydrated using an ethanol series, critical point dried and mounted for examination. Low resolution images were taken to identify the 12kHz region of the cochlea (using the frequency-place map described by Müller et al [82]). Higher-resolution images were taken using a JEOL JSM 7800 Prime scanning electron microscope under a standard magnification (60x) to show the whole organ of Corti, and at higher magnifications to examine hair cell rows (2000x) and individual hair cells (15000-23000x). Whole images have been adjusted in Photoshop to normalise dynamic range across all panels, and rotated where necessary to present a uniform orientation.

### Wholemount immunostaining and confocal microscopy

The cochleae of P28 mice were fixed in 4% paraformaldehyde in PBS, washed in PBS, and decalcified in 10x ethylenediaminetetraacetic acid (EDTA) for 2 hours. After fine dissection, samples were blocked in 5% normal horse serum (NHS), 1% bovine serum albumin (BSA) and 0.3% Triton X-100 in PBS for 45 minutes at room temperature, then immunostained in 1% NHS and 0.3% Triton X-100 in PBS, as described in [83]. The primary antibodies used were anti-NFH (Abcam, cat. no: ab4680, diluted 1:800), anti-GluR2 (Millipore, cat. no: MAB397, diluted 1:200) and anti-Ribeye (Synaptic Systems, cat. no: 192 103, diluted 1:500), and the secondary antibodies were Alexa Fluor 488 goat anti-chicken (Invitrogen, cat. no: A11039, diluted 1:300), Alexa Fluor 546 goat anti-rabbit (Invitrogen, cat. no: A11035, diluted 1:300) and Alexa Fluor 488 goat anti-mouse (Invitrogen, cat. no: A21131, diluted 1:300). Samples were mounted in ProLong Gold antifade mounting medium with DAPI (Life Technologies, cat. no: P36931), or Vectashield Mounting Medium with DAPI (Vector Laboratories, cat. no: H-1200), and imaged with a Zeiss Imager 710 confocal microscope (plan-APOCHROMAT 63x Oil DIC objective) interfaced with ZEN 2010 software, or a Nikon A1R point-scanning confocal microscope (Plan Apo VC 60x/1.4NA oil objective) using NIS Elements v4.2 software (Nikon Instruments UK). Confocal z-stacks were obtained with a z-step size of 0.25µm (for synapses) or 0.4µm (for innervation). For synapse counting, a low magnification image was taken of approximately 20 inner hair cells in the 12kHz region, then a close-up image (2.5x optical zoom) was taken from both extremes of the low magnification image to ensure no synapses were counted twice. Synapses were counted from the maximum projection of the close up images using the FIJI plugin of ImageJ, and divided by the number of hair cell nuclei visible in the field of view (between 5 and 13, depending on which microscope was used) to obtain the number of synapses/hair cell. Where synapses were counted in both ears from the same mouse, the count was averaged before inclusion in the data. Whole images were processed in Adobe Photoshop, rotated where necessary, and adjusted so that all channels were equally visible.

### Dissection and RNA extraction

The organs of Corti of four-day-old (P4) mice were dissected during a fixed time window (between 6 and 7.5 hours after lights on) to avoid circadian variation, and stored at −20°C in RNAlater stabilisation reagent (Ambion). RNA was extracted from both organs of Corti using either QIAshredder columns (QIAgen, cat. no. 79654) and the RNeasy mini kit (QIAgen, cat. no. 74104), or the Lexogen SPLIT kit (Lexogen, cat. no. 008.48), following the manufacturer’s instructions. RNA concentration was measured using a Nanodrop spectrophotometer (ND-8000).

### RNA-seq

RNA from both ears of six wildtype and six sex-matched homozygote mutant littermates from each mutant line were used for RNA-seq. Samples were not pooled. Strand-specific libraries were prepared using the NuGEN Ovation Mouse RNA-Seq System 1-16 kit (NuGEN, cat. no. 0348) and sequenced on an Illumina HiSeq 2500 machine as paired-end 125bp reads. The resulting reads were quality checked using FastQC 0.11.4 [84] and trimmed with Trimmomatic 0.35 [85]; adapters were removed, trailing ends were clipped where the quality was low and a sliding window approach used to control for quality across the entire read. Finally, reads with 36 basepairs or fewer were discarded, because the shorter a read, the less likely it is to map uniquely to the genome. Reads were assembled to GRCm38 using Hisat2 version 2.0.2beta [86]. Bam files were soft-clipped beyond end-of-reference alignments and MAPQ scores set to 0 for unmapped reads using Picard 2.1.0 (Broad Institute, Cambridge, Mass. http://broadinstitute.github.io/picard) and checked for quality using QoRTS [87]. The QoRTS tool also generates count data (in the same format as HTSeq), and these were used with edgeR [88] to carry out a generalized linear model likelihood ratio test. Splicing analyses were performed using Cuffdiff (Cufflinks) [39], JunctionSeq [40] and Leafcutter [41].

### Transcriptome and network analysis

Sylamer 18-131 [33] was used to examine genes ranked in order of their misregulation from up- to downregulated for over- and under-represented heptamers in their 3’ UTRs. Ingenuity Pathway Analysis (IPA) was used to generate potential upstream regulators for the affected genes, using the causal network analysis module. The WGCNA R package [89] was used to carry out weighted gene correlation network analysis. Gene counts from the QoRTS tool were prepared for WGCNA using DESeq2 [90] and transformed with a variance stabilising transformation. Batch effects were controlled for using the limma R package [91]. Module enrichment analysis was carried out using PANTHER v14 [47], and oPOSSUM [51] was used to assess transcription factor binding site overrepresentation in each module. Motif detection in a set of genes can be affected by differing GC composition in the genes used as a “background” set, so background gene sets of 4,000-5,000 genes were selected for each module such that the GC content matched that of the genes of interest. oPOSSUM assigns two scores to each transcription factor profile, the Z-score (which assesses whether the rate of occurrence of a given motif in the genes of interest differs significantly from the expected rate calculated from the background genes) and the Fisher score (which compares the proportion of genes of interest which contain a given motif to the proportion of the background genes containing that motif in order to determine the probability of a non-random association between the motif and the genes of interest) [51]. We chose to use a threshold of the mean + 1 standard deviation for each score.

### RTPCR and qPCR

Organ of Corti RNA was normalised to the same concentration within each litter, then treated with DNAse 1 (Sigma, cat. no: AMPD1) before cDNA creation. cDNA was made using Superscript II Reverse Transcriptase (Invitrogen, cat. no: 11904-018) or M-MLV Reverse Transcriptase (Invitrogen, cat. no: 28025-013) or Precision Reverse Transcription Premix (PrimerDesign, cat. no: RT-premix2). MicroRNA cDNA was made using the miRCURY LNA RT Kit (QIAGEN, cat. no: 339340). Primers for sequencing cDNA for testing differential splicing were designed using Primer3 [92] (Supplementary Table S3). Sanger sequencing was carried out by Source Bioscience and analysed using Gap4 [93]. Quantitative RT-PCR was carried out on a CFX Connect qPCR machine (Bio-Rad), using probes from Applied Biosystems and QIAGEN (see Supplementary Table S3 for primer/probe details) and Sso-Advanced Master Mix (Bio-Rad, cat. no: 1725281) or the miRCURY LNA SYBR Green PCR Kit (QIAGEN, cat. no: 339345) for miRNA qPCR. Relative expression levels were calculated using the 2^-ΔΔct^ equation [94], with *Hprt* as an internal control for all protein-coding genes except *Ocm* and *Slc26a5*, which are specifically expressed in hair cells, for which *Jag1* was used as an internal control, because it is expressed in supporting cells of the organ of Corti [95, 96]. For all other genes and microRNAs, the quantity of sensory tissue present was checked using *Jag1*, and pairs were only used if their *Jag1* levels did not differ by more than ±20%. For the microRNA qPCR, the internal control was *Mir99a*, which is expressed in most cell types in the cochlea [10]. At least three technical replicates of each sample were carried out for each reaction, and at least four biological replicates were tested per probe (see legends of Supplementary Figures S1, S11 for numbers for each probe).

### Statistics

For qPCR data, the Wilcoxon rank sum test (Mann-Whitney U test) was chosen to determine significance, because it is a suitable test for small sample sizes and populations of unknown characteristics [97]. For the microRNA qPCR and synapse count data, we used a one-way ANOVA because three groups were being compared. Post-hoc test p-values were adjusted using the Bonferroni correction. For repeated ABR threshold analyses, the thresholds were not normally distributed, so the data were first transformed using the arcsine transformation then analysed using separate linear models for each frequency with a compound symmetric covariance structure and restricted Maximum Likelihood Estimation [98]. This allowed the inclusion of all available data, unlike the repeated measures ANOVA, which requires the discarding of a subject if any data points are missed (for example, if a mouse died before the final ABR measurement) [99]. For each stimulus the double interaction of genotype and age was measured, followed by Bonferroni correction for multiple testing. Wilcoxon rank sum tests were carried out using R, and the one way ANOVAs, arcsine transformation and mixed model linear pairwise comparison were done with SPSS v25.

### Immunohistochemistry

Samples from P4 pups were collected, fixed in 10% formalin, embedded in paraffin wax and cut into 8μm sections. Immunohistochemistry was carried out using a Ventana Discovery machine and reagents according to the manufacturer’s instructions (DABMap^TM^ Kit (cat.no 760-124), Hematoxylin (cat.no 760-2021), Bluing reagent (cat.no 760-2037), CC1 (cat.no 950-124), EZPrep (cat.no 950-100), LCS (cat.no 650-010), RiboWash (cat.no 760-105), Reaction Buffer (cat.no 95-300), and RiboCC (cat.no 760-107)). For each antibody, at least three wildtype/homozygote littermate pairs were tested, and from each animal, at least five mid-modiolar sections were used per antibody. Primary antibodies used were rabbit anti-Ocm (Abcam, cat. no: ab150947, diluted 1:50) and goat anti-Prestin (*Slc26a5*) (Santa Cruz, cat. no: sc-22692, diluted 1:50), and secondary antibodies were anti-goat (Jackson ImmunoResearch, cat.no 705-065-147, diluted 1:100), and anti-rabbit (Jackson ImmunoResearch, cat.no 711-065-152, diluted 1:100). Antibodies were diluted in staining solution (10% foetal calf serum, 0.1% Triton, 2% BSA and 0.5% sodium azide in PBS). A Zeiss Axioskop 2 microscope with a Plan Neofluar 63x 1.4NA objective was used to examine slides, and photos were taken using a Zeiss Axiocam camera and the associated Axiocam software. Images were processed in Adobe Photoshop; minimal adjustments were made, including rotation and resizing. Where image settings were altered, the adjustment was applied equally to wildtype and mutant photos and to the whole image.

### Prediction of potential causal regulatory networks

In our previous analysis [16], we used interactions from the literature to connect miR-96 to as many of the misregulated genes in *Mir96^Dmdo^* homozygotes as possible. In order to automate this procedure, a custom perl script (prediction of potential causal regulatory networks, PoPCoRN) was written to make use of publically available regulatory data from ArrayExpress and the associated Expression Atlas [100, 101], ORegAnno [102], miRTarBase [103], TransmiR [104] and TRRUST [105] (Table 3). All these data are based on experimental evidence, although we did make use of human regulatory interactions by converting human gene IDs to mouse gene IDs where there was a one-to-one orthologue (using Ensembl). Interactions from our previous network [16], which were obtained from the literature using Ingenuity IPA, were also added, as were regulations reported in [106], which is not available through ArrayExpress but is a particularly relevant study, since it reports the results of microarrays carried out on RNA from mouse inner ears. MicroRNA targets confirmed in the literature were included, as were experimentally validated targets listed in miRTarBase as having “strong experimental evidence”, either reporter assay or Western blot [103], and genes predicted or confirmed to be miR-96 targets by our previous studies [3, 16]. Finally, genes upregulated in *Mir183/96^dko^* homozygotes with heptamers complementary to either the miR-96 or the miR-183 seed region in their 3’UTR were included as targets of the relevant microRNA(s) (Table 1, top). Similarly, genes upregulated in *Mir182^ko^* homozygotes with heptamers complementary to the miR-182 seed region in their 3’UTR were included as targets of miR-182 (Table 1, bottom). This resulted in a list of 97062 unique links of the form <gene A> <interaction> <gene B>.

**Table 3.** Data used as a basis for building the regulatory networks.

| Database | Format example | Number of Unique Entries | Reference |
| --- | --- | --- | --- |
| ArrayExpress and the Gene Expression Atlas | If a null allele of gene A results in a decrease in expression of gene B, gene A upregulates gene B | 87424 | [100, 101] |
| ORegAnno | If there is a binding site in gene B for gene A, and the outcome of the interaction is NEGATIVE, gene A downregulates gene B | 1044 | [102] |
| miRTarBase | If microRNA X has target gene A, X downregulates A | 6199 | [103] |
| TransmiR | If gene A represses microRNA X, A downregulates X | 1596 | [104] |
| TRRUST | If the relationship between gene A and gene B is activation, gene A upregulates gene B | 8265 | [105] |
| n/a | Various | 519 | [16, 106] |
| n/a | If a null allele of microRNA X results in an increase in expression of gene A, and gene A has seed region matches for X, microRNA X downregulates gene A | 18 | Current study |

All potential links between the ultimate regulators (*Mir183* and *Mir96*, *Mir96* alone, or *Mir182*) and the misregulated genes were then identified, and the direction of misregulation of intermediate genes predicted. Starting with the known misregulated genes, each upstream regulator was given a score based on the direction of regulation of the known misregulated genes (Supplementary Fig. S19). This process iterated until the gene(s) at the top of the cascade (in this case, *Mir96*, *Mir183* or *Mir182*) were reached. Consistent links were kept, the shortest paths between microRNA and misregulated genes identified, and the final network was written out in the simple interaction format (sif) and viewed using Cytoscape [107] (Supplementary Fig. S18).

In order to test the PoPCoRN tool, we searched for studies which measured the misregulation of a set of genes in a system where a specific upstream regulator was either induced or silenced, and which also identified and confirmed the misregulation of an intermediate gene (where necessary, the data for these intermediate genes was removed from the input prior to network creation). We found ten suitable studies from which we were able to create networks and test the predicted misregulation of 14 intermediate genes. The tool made correct predictions for seven of the genes and incorrect predictions for three of them (Table 4). For the remaining four, it did not predict either up- or downregulation, which points to a deficiency of underlying data (Table 4). This will always be a problem for a tool based on existing data which does not attempt extrapolation.

**Table 4.** Data used for checking the PoPCoRN predictions.

| Upstream regulator | Perturbation | Tissue | Age | Intermediate gene | Paper confirmation | PoPCoRN prediction | Number of input genes | Reference |
| --- | --- | --- | --- | --- | --- | --- | --- | --- |
| *Tbx1* | Reduced function due to missense mutation | Inner ear | E16.5 | *Esrrb* | Downregulated | Downregulated | 39 | [111] |
| *Stk11* | Knockout | Chondrocytes | P30 | *Runx2* | Upregulated | Upregulated | 962 | [112] |
| *Stk11* | Knockout | Chondrocytes | P30 | *Nfatc1* | Upregulated | Downregulated | 962 | [112] |
| *Stk11* | Knockout | Chondrocytes | P30 | *Mir134* | Upregulated | Upregulated | 962 | [112] |
| *Yy1* | Knockout | Haematopoietic stem cells | - | *Kit* | Downregulated | Upregulated | 55 | [113] |
| *Sox2* | Induced knockout | Acinar cells of salivary gland | E16.5 | *Sox10* | Downregulated | Upregulated | 7 | [114] |
| *Trp53* | Knockout | Cultured osteoblasts | - | *Runx2* | Upregulated | Upregulated | 3 | [115] |
| *Trp53* | Knockout | Cultured osteoblasts | - | *Sp7* | Upregulated | Upregulated | 3 | [115] |
| *Yy1* | Inhibition by siRNA | HUVECs | - | *Rbpj* | Upregulated | Upregulated | 15 | [116] |
| *Brca1* | Inhibition by siRNA | HUVECs | - | *Cdkn1a* | Downregulated | Downregulated | 3 | [117] |
| *Hif1a* | Induction by hypoxia | HepG2/HeLa cells | - | *Bclaf1* | Upregulated | No prediction | 4 | [118] |
| *Gata4* | Inhibition by shRNA | Dental pulp stem cells | - | *Fbp1* | Upregulated | No prediction | 9 | [119] |
| *Arntl* | Knockout | Mandibular condyle | P28 | *Ihh* | Downregulated | No prediction | 7 | [120] |
| *Arntl* | Knockout | Mandibular condyle | P28 | *Ptch1* | Downregulated | No prediction | 7 | [120] |

Many similar approaches to network creation have been described previously (for example, [42, 108–110]), but they address the case where the upstream regulator or regulators remain unknown, for example when comparing samples from healthy and diseased tissues, and one of their main aims is to identify potential upstream regulators. Our implementation asks a different question, exploring the downstream regulatory cascade of a known regulator. In the case of miR-96, and potentially many other regulators involved in disease, the genes involved in mediating their effect are candidate therapeutic targets.

### Strain differences in seed region matches

The C57BL/6NJ genomic sequence is available from the Mouse Genomes Project [32] via the Ensembl browser [121], but the *Mir96^Dmdo^* mice were made and maintained on the C3HeB/FeJ background, which is not one of the sequenced strains. Instead, we made use of genomic sequence from the C3H/HeJ strain, which is closely related to C3HeB/FeJ and is one of the other strains sequenced as part of the Mouse Genomes Project. We searched the 3’UTRs of each strain for the complement of the miR-96 seed region (GTGCCAA). We removed genes without an MGI ID, since without a consistent ID the corresponding gene in the other strain could not be identified. 1733 genes were left with at least one match to the miR-96 seed region in their 3’ UTRs in one or both strains. 21 genes had no 3’UTR sequence available for C57BL/6NJ, and 31 genes had no 3’UTR sequence available for C3H/HeJ.

## Data and software availability

The datasets and computer code produced in this study are available in the following databases:

RNA-Seq data: ArrayExpress E-MTAB-5800 (https://www.ebi.ac.uk/arrayexpress/experiments/E-MTAB-5800).
PoPCoRN network construction script: GitHub (https://github.com/moraglewis/PoPCoRN)

## Disclosure of interest

The authors declare that they have no conflicts of interest.

## Funding

This project was supported by Action on Hearing Loss (Pauline Ashley Fellowship to MAL), the MRC (KPS, MC_qA137918, G0300212) and the Wellcome Trust (098051, KPS: 100699). The Zeiss Imager Confocal Microscope is maintained with a Wellcome Trust grant (WT089622MA). No role was taken by any funding body in designing the study, in collection, analysis or interpretation of data, or in the writing of this manuscript.

## Authors’ contributions

MAL and KPS conceived and designed the experiments. MAL, FDD and NJI performed the experiments. MAL, FDD, NJI and KPS analysed the data. MAL designed and wrote the PoPCoRN software. HMP contributed the *Mir183/96^dko^* and *Mir182^ko^* mice. MAL and KPS wrote the paper, and all authors reviewed it.

## Acknowledgements

The RNA-Seq was carried out by the National Institute for Health Research (NIHR) Biomedical Research Centre based at Guy’s and St Thomas’ NHS Foundation Trust and King’s College London. We thank Matthew Arno and Sucharitha Balu for doing the RNA library preps and sample submission. The confocal microscopy was carried out with help from the Nikon Imaging Centre at Kings College London, and the scanning electron microscopy was done at the King’s College London Centre for Ultrastructural Imaging. We are grateful to Allan Bradley for access to the *Mir183/96^dko^* and *Mir182^ko^* mice. We thank Holly Smith for her work on the immunohistochemistry. We are very grateful to Dr Lawrence Moon for help and advice with the statistics.

## Notes

https://www.ebi.ac.uk/arrayexpress/experiments/E-MTAB-5800

http://github.com/moraglewis/PoPCoRN

### Competing Interest Statement

The authors have declared no competing interest.

### Summary of Updates

Addition of Supplementary Data file containing all data underlying graphs presented as figures.

## References

1. Weston MD, Pierce ML, Rocha-Sanchez S, Beisel KW, Soukup GA: MicroRNA gene expression in the mouse inner ear. Brain Res 2006, 1111:95–104.

2. Xu S, Witmer PD, Lumayag S, Kovacs B, Valle D: MicroRNA (miRNA) transcriptome of mouse retina and identification of a sensory organ-specific miRNA cluster. J Biol Chem 2007, 282:25053–25066.

3. Lewis MA, Quint E, Glazier AM, Fuchs H, De Angelis MH, Langford C, van Dongen S, Abreu-Goodger C, Piipari M, Redshaw N, et al: An ENU-induced mutation of miR-96 associated with progressive hearing loss in mice. Nat Genet 2009, 41:614–618.

4. Mencia A, Modamio-Hoybjor S, Redshaw N, Morin M, Mayo-Merino F, Olavarrieta L, Aguirre LA, del Castillo I, Steel KP, Dalmay T, et al: Mutations in the seed region of human miR-96 are responsible for nonsyndromic progressive hearing loss. Nat Genet 2009, 41:609–613.

5. Solda G, Robusto M, Primignani P, Castorina P, Benzoni E, Cesarani A, Ambrosetti U, Asselta R, Duga S: A novel mutation within the MIR96 gene causes non-syndromic inherited hearing loss in an Italian family by altering pre-miRNA processing. Hum Mol Genet 2012, 21:577–585.

6. Schluter T, Berger C, Rosengauer E, Fieth P, Krohs C, Ushakov K, Steel KP, Avraham KB, Hartmann AK, Felmy F, Nothwang HG: miR-96 is required for normal development of the auditory hindbrain. Hum Mol Genet 2018, 27:860–874.

7. Chen J, Johnson SL, Lewis MA, Hilton JM, Huma A, Marcotti W, Steel KP: A reduction in Ptprq associated with specific features of the deafness phenotype of the miR-96 mutant mouse diminuendo. Eur J Neurosci 2014, 39:744–756.

8. Kuhn S, Johnson SL, Furness DN, Chen J, Ingham N, Hilton JM, Steffes G, Lewis MA, Zampini V, Hackney CM, et al: miR-96 regulates the progression of differentiation in mammalian cochlear inner and outer hair cells. Proc Natl Acad Sci U S A 2011, 108:2355–2360.

9. Weston MD, Tarang S, Pierce ML, Pyakurel U, Rocha-Sanchez SM, McGee J, Walsh EJ, Soukup GA: A mouse model of miR-96, miR-182 and miR-183 misexpression implicates miRNAs in cochlear cell fate and homeostasis. Sci Rep 2018, 8:3569.

10. Friedman LM, Dror AA, Mor E, Tenne T, Toren G, Satoh T, Biesemeier DJ, Shomron N, Fekete DM, Hornstein E, Avraham KB: MicroRNAs are essential for development and function of inner ear hair cells in vertebrates. Proc Natl Acad Sci U S A 2009, 106:7915–7920.

11. Soukup GA, Fritzsch B, Pierce ML, Weston MD, Jahan I, McManus MT, Harfe BD: Residual microRNA expression dictates the extent of inner ear development in conditional Dicer knockout mice. Dev Biol 2009, 328:328–341.

12. Liu Y, Bailey JC, Helwa I, Dismuke WM, Cai J, Drewry M, Brilliant MH, Budenz DL, Christen WG, Chasman DI, et al: A Common Variant in MIR182 Is Associated With Primary Open-Angle Glaucoma in the NEIGHBORHOOD Consortium. Invest Ophthalmol Vis Sci 2016, 57:4528–4535.

13. Cui H, Yang L: Analysis of microRNA expression detected by microarray of the cerebral cortex after hypoxic-ischemic brain injury. J Craniofac Surg 2013, 24:2147–2152.

14. Duan X, Gan J, Peng DY, Bao Q, Xiao L, Wei L, Wu J: Identification and functional analysis of microRNAs in rats following focal cerebral ischemia injury. Mol Med Rep 2019, 19:4175–4184.

15. Ling W, Xu X, Liu J: A causal relationship between the neurotherapeutic effects of miR182/7a and decreased expression of PRDM5. Biochem Biophys Res Commun 2017, 490:1–7.

16. Lewis MA, Buniello A, Hilton JM, Zhu F, Zhang WI, Evans S, van Dongen S, Enright AJ, Steel KP: Exploring regulatory networks of miR-96 in the developing inner ear. Sci Rep 2016, 6:23363.

17. Simmons SO, Fan CY, Ramabhadran R: Cellular stress response pathway system as a sentinel ensemble in toxicological screening. Toxicol Sci 2009, 111:202–225.

18. Prosser HM, Koike-Yusa H, Cooper JD, Law FC, Bradley A: A resource of vectors and ES cells for targeted deletion of microRNAs in mice. Nat Biotechnol 2011, 29:840–845.

19. Noben-Trauth K, Zheng QY, Johnson KR: Association of cadherin 23 with polygenic inheritance and genetic modification of sensorineural hearing loss. Nat Genet 2003, 35:21–23.

20. Li HS, Borg E: Age-related loss of auditory sensitivity in two mouse genotypes. Acta Otolaryngol 1991, 111:827–834.

21. Kujawa SG, Liberman MC: Adding insult to injury: cochlear nerve degeneration after “temporary” noise-induced hearing loss. J Neurosci 2009, 29:14077–14085.

22. Cai T, Jen HI, Kang H, Klisch TJ, Zoghbi HY, Groves AK: Characterization of the transcriptome of nascent hair cells and identification of direct targets of the Atoh1 transcription factor. J Neurosci 2015, 35:5870–5883.

23. Elkon R, Milon B, Morrison L, Shah M, Vijayakumar S, Racherla M, Leitch CC, Silipino L, Hadi S, Weiss-Gayet M, et al: RFX transcription factors are essential for hearing in mice. Nat Commun 2015, 6:8549.

24. Kurima K, Peters LM, Yang Y, Riazuddin S, Ahmed ZM, Naz S, Arnaud D, Drury S, Mo J, Makishima T, et al: Dominant and recessive deafness caused by mutations of a novel gene, TMC1, required for cochlear hair-cell function. Nat Genet 2002, 30:277–284.

25. Liberman MC, Gao J, He DZ, Wu X, Jia S, Zuo J: Prestin is required for electromotility of the outer hair cell and for the cochlear amplifier. Nature 2002, 419:300–304.

26. Liu XZ, Ouyang XM, Xia XJ, Zheng J, Pandya A, Li F, Du LL, Welch KO, Petit C, Smith RJ, et al: Prestin, a cochlear motor protein, is defective in non-syndromic hearing loss. Hum Mol Genet 2003, 12:1155–1162.

27. Vreugde S, Erven A, Kros CJ, Marcotti W, Fuchs H, Kurima K, Wilcox ER, Friedman TB, Griffith AJ, Balling R, et al: Beethoven, a mouse model for dominant, progressive hearing loss DFNA36. Nat Genet 2002, 30:257–258.

28. Walsh T, Walsh V, Vreugde S, Hertzano R, Shahin H, Haika S, Lee MK, Kanaan M, King MC, Avraham KB: From flies’ eyes to our ears: mutations in a human class III myosin cause progressive nonsyndromic hearing loss DFNB30. Proc Natl Acad Sci U S A 2002, 99:7518–7523.

29. Walsh VL, Raviv D, Dror AA, Shahin H, Walsh T, Kanaan MN, Avraham KB, King MC: A mouse model for human hearing loss DFNB30 due to loss of function of myosin IIIA. Mamm Genome 2011, 22:170–177.

30. Lalani SR, Safiullah AM, Molinari LM, Fernbach SD, Martin DM, Belmont JW: SEMA3E mutation in a patient with CHARGE syndrome. J Med Genet 2004, 41:e94.

31. Tong B, Hornak AJ, Maison SF, Ohlemiller KK, Liberman MC, Simmons DD: Oncomodulin, an EF-Hand Ca2+ Buffer, Is Critical for Maintaining Cochlear Function in Mice. J Neurosci 2016, 36:1631–1635.

32. Adams DJ, Doran AG, Lilue J, Keane TM: The Mouse Genomes Project: a repository of inbred laboratory mouse strain genomes. Mamm Genome 2015, 26:403–412.

33. van Dongen S, Abreu-Goodger C, Enright AJ: Detecting microRNA binding and siRNA off-target effects from expression data. Nat Methods 2008, 5:1023–1025.

34. Barreau C, Paillard L, Osborne HB: AU-rich elements and associated factors: are there unifying principles? Nucleic Acids Res 2005, 33:7138–7150.

35. Chen CY, Shyu AB: AU-rich elements: characterization and importance in mRNA degradation. Trends Biochem Sci 1995, 20:465–470.

36. Plass M, Rasmussen SH, Krogh A: Highly accessible AU-rich regions in 3’ untranslated regions are hotspots for binding of regulatory factors. PLoS Comput Biol 2017, 13:e1005460.

37. Boutz PL, Chawla G, Stoilov P, Black DL: MicroRNAs regulate the expression of the alternative splicing factor nPTB during muscle development. Genes Dev 2007, 21:71–84.

38. Liu R, Loraine AE, Dickerson JA: Comparisons of computational methods for differential alternative splicing detection using RNA-seq in plant systems. BMC Bioinformatics 2014, 15:364.

39. Trapnell C, Williams BA, Pertea G, Mortazavi A, Kwan G, van Baren MJ, Salzberg SL, Wold BJ, Pachter L: Transcript assembly and quantification by RNA-Seq reveals unannotated transcripts and isoform switching during cell differentiation. Nat Biotechnol 2010, 28:511–515.

40. Hartley SW, Mullikin JC: Detection and visualization of differential splicing in RNA-Seq data with JunctionSeq. Nucleic Acids Res 2016, 44:e127.

41. Li YI, Knowles DA, Pritchard JK: LeafCutter: Annotation-free quantification of RNA splicing. bioRxiv 2016.

42. Kramer A, Green J, Pollard J, Jr., Tugendreich S: Causal analysis approaches in Ingenuity Pathway Analysis. Bioinformatics 2014, 30:523–530.

43. Li H, Gong Y, Qian H, Chen T, Liu Z, Jiang Z, Wei S: Brain-derived neurotrophic factor is a novel target gene of the has-miR-183/96/182 cluster in retinal pigment epithelial cells following visible light exposure. Mol Med Rep 2015, 12:2793–2799.

44. Agerman K, Hjerling-Leffler J, Blanchard MP, Scarfone E, Canlon B, Nosrat C, Ernfors P: BDNF gene replacement reveals multiple mechanisms for establishing neurotrophin specificity during sensory nervous system development. Development 2003, 130:1479–1491.

45. Deol MS: The origin of the acoustic ganglion and effects of the gene dominant spotting (Wv) in the mouse. J Embryol Exp Morphol 1970, 23:773–784.

46. Zhang B, Horvath S: A general framework for weighted gene co-expression network analysis. Stat Appl Genet Mol Biol 2005, 4:Article17.

47. Mi H, Muruganujan A, Ebert D, Huang X, Thomas PD: PANTHER version 14: more genomes, a new PANTHER GO-slim and improvements in enrichment analysis tools. Nucleic Acids Res 2019, 47:D419–D426.

48. Ashburner M, Ball CA, Blake JA, Botstein D, Butler H, Cherry JM, Davis AP, Dolinski K, Dwight SS, Eppig JT, et al: Gene ontology: tool for the unification of biology. The Gene Ontology Consortium. Nat Genet 2000, 25:25–29.

49. Consortium TGO: The Gene Ontology Resource: 20 years and still GOing strong. Nucleic Acids Res 2019, 47:D330–D338.

50. Fabregat A, Jupe S, Matthews L, Sidiropoulos K, Gillespie M, Garapati P, Haw R, Jassal B, Korninger F, May B, et al: The Reactome Pathway Knowledgebase. Nucleic Acids Res 2018, 46:D649–D655.

51. Kwon AT, Arenillas DJ, Worsley Hunt R, Wasserman WW: oPOSSUM-3: advanced analysis of regulatory motif over-representation across genes or ChIP-Seq datasets. G3 (Bethesda) 2012, 2:987–1002.

52. Hulander M, Wurst W, Carlsson P, Enerback S: The winged helix transcription factor Fkh10 is required for normal development of the inner ear. Nat Genet 1998, 20:374–376.

53. Wei K, Chen J, Akrami K, Galbraith GC, Lopez IA, Chen F: Neural crest cell deficiency of c-myc causes skull and hearing defects. Genesis 2007, 45:382–390.

54. Paylor R, Johnson RS, Papaioannou V, Spiegelman BM, Wehner JM: Behavioral assessment of c-fos mutant mice. Brain Res 1994, 651:275-282.

55. Gilels F, Paquette ST, Zhang J, Rahman I, White PM: Mutation of Foxo3 causes adult onset auditory neuropathy and alters cochlear synapse architecture in mice. J Neurosci 2013, 33:18409–18424.

56. Lang H, Schulte BA, Zhou D, Smythe N, Spicer SS, Schmiedt RA: Nuclear factor kappaB deficiency is associated with auditory nerve degeneration and increased noise-induced hearing loss. J Neurosci 2006, 26:3541–3550.

57. Steigelman KA, Lelli A, Wu X, Gao J, Lin S, Piontek K, Wodarczyk C, Boletta A, Kim H, Qian F, et al: Polycystin-1 is required for stereocilia structure but not for mechanotransduction in inner ear hair cells. J Neurosci 2011, 31:12241–12250.

58. Nakano Y, Kelly MC, Rehman AU, Boger ET, Morell RJ, Kelley MW, Friedman TB, Banfi B: Defects in the Alternative Splicing-Dependent Regulation of REST Cause Deafness. Cell 2018, 174:536–548 e521.

59. Oishi N, Chen J, Zheng HW, Hill K, Schacht J, Sha SH: Tumor necrosis factor-alpha-mutant mice exhibit high frequency hearing loss. J Assoc Res Otolaryngol 2013, 14:801–811.

60. Fan J, Jia L, Li Y, Ebrahim S, May-Simera H, Wood A, Morell RJ, Liu P, Lei J, Kachar B, et al: Maturation arrest in early postnatal sensory receptors by deletion of the miR-183/96/182 cluster in mouse. Proc Natl Acad Sci U S A 2017, 114:E4271–E4280.

61. Geng R, Furness DN, Muraleedharan CK, Zhang J, Dabdoub A, Lin V, Xu S: The microRNA-183/96/182 Cluster is Essential for Stereociliary Bundle Formation and Function of Cochlear Sensory Hair Cells. Sci Rep 2018, 8:18022.

62. Lumayag S, Haldin CE, Corbett NJ, Wahlin KJ, Cowan C, Turturro S, Larsen PE, Kovacs B, Witmer PD, Valle D, et al: Inactivation of the microRNA-183/96/182 cluster results in syndromic retinal degeneration. Proc Natl Acad Sci U S A 2013, 110:E507–516.

63. Pittler SJ, Baehr W: Identification of a nonsense mutation in the rod photoreceptor cGMP phosphodiesterase beta-subunit gene of the rd mouse. Proc Natl Acad Sci U S A 1991, 88:8322–8326.

64. Moore BA, Roux MJ, Sebbag L, Cooper A, Edwards SG, Leonard BC, Imai DM, Griffey S, Bower L, Clary D, et al: A Population Study of Common Ocular Abnormalities in C57BL/6N rd8 Mice. Invest Ophthalmol Vis Sci 2018, 59:2252–2261.

65. Chessum L, Matern MS, Kelly MC, Johnson SL, Ogawa Y, Milon B, McMurray M, Driver EC, Parker A, Song Y, et al: Helios is a key transcriptional regulator of outer hair cell maturation. Nature 2018, 563:696–700.

66. The gEAR portal [http://gear.igs.umaryland.edu/#]

67. Sanchez-Martin M, Rodriguez-Garcia A, Perez-Losada J, Sagrera A, Read AP, Sanchez-Garcia I: SLUG (SNAI2) deletions in patients with Waardenburg disease. Hum Mol Genet 2002, 11:3231–3236.

68. Hertzano R, Elkon R, Kurima K, Morrisson A, Chan SL, Sallin M, Biedlingmaier A, Darling DS, Griffith AJ, Eisenman DJ, Strome SE: Cell type-specific transcriptome analysis reveals a major role for Zeb1 and miR-200b in mouse inner ear morphogenesis. PLoS Genet 2011, 7:e1002309.

69. DeMambro VE, Kawai M, Clemens TL, Fulzele K, Maynard JA, Marin de Evsikova C, Johnson KR, Canalis E, Beamer WG, Rosen CJ, Donahue LR: A novel spontaneous mutation of Irs1 in mice results in hyperinsulinemia, reduced growth, low bone mass and impaired adipogenesis. J Endocrinol 2010, 204:241–253.

70. Erichsen S, Stierna P, Bagger-Sjoback D, Curtis LM, Rarey KE, Schmid W, Hultcrantz M: Distribution of Na,K-ATPase is normal in the inner ear of a mouse with a null mutation of the glucocorticoid receptor. Hear Res 1998, 124:146–154.

71. ten Cate WJ, Curtis LM, Small GM, Rarey KE: Localization of glucocorticoid receptors and glucocorticoid receptor mRNAs in the rat cochlea. Laryngoscope 1993, 103:865–871.

72. Smith CL, Blake JA, Kadin JA, Richardson JE, Bult CJ: Mouse Genome Database (MGD)-2018: knowledgebase for the laboratory mouse. Nucleic Acids Res 2018, 46:D836–D842.

73. Dickinson ME, Flenniken AM, Ji X, Teboul L, Wong MD, White JK, Meehan TF, Weninger WJ, Westerberg H, Adissu H, et al: High-throughput discovery of novel developmental phenotypes. Nature 2016, 537:508–514.

74. Daamen WF, Quaglino D: Signaling pathways in elastic tissues. Cell Signal 2019, 63:109364.

75. Karnik SK, Brooke BS, Bayes-Genis A, Sorensen L, Wythe JD, Schwartz RS, Keating MT, Li DY: A critical role for elastin signaling in vascular morphogenesis and disease. Development 2003, 130:411–423.

76. Goodyear RJ, Legan PK, Wright MB, Marcotti W, Oganesian A, Coats SA, Booth CJ, Kros CJ, Seifert RA, Bowen-Pope DF, Richardson GP: A receptor-like inositol lipid phosphatase is required for the maturation of developing cochlear hair bundles. J Neurosci 2003, 23:9208–9219.

77. Roux I, Wu JS, McIntosh JM, Glowatzki E: Assessment of the expression and role of the alpha1-nAChR subunit in efferent cholinergic function during the development of the mammalian cochlea. J Neurophysiol 2016, 116:479–492.

78. Hornstein E, Shomron N: Canalization of development by microRNAs. Nat Genet 2006, 38 Suppl:S20-24.

79. Ingham NJ, Pearson S, Steel KP: Using the Auditory Brainstem Response (ABR) to Determine Sensitivity of Hearing in Mutant Mice. Curr Protoc Mouse Biol 2011, 1:279–287.

80. Holme RH, Steel KP: Progressive hearing loss and increased susceptibility to noise-induced hearing loss in mice carrying a Cdh23 but not a Myo7a mutation. J Assoc Res Otolaryngol 2004, 5:66–79.

81. Hunter-Duvar IM: A technique for preparation of cochlear specimens for assessment with the scanning electron microscope. *Acto Otoloaryng Suppl* 1978, 351:3–23.

82. Muller M, von Hunerbein K, Hoidis S, Smolders JW: A physiological place-frequency map of the cochlea in the CBA/J mouse. Hear Res 2005, 202:63–73.

83. Buniello A, Ingham NJ, Lewis MA, Huma AC, Martinez-Vega R, Varela-Nieto I, Vizcay-Barrena G, Fleck RA, Houston O, Bardhan T, et al: Wbp2 is required for normal glutamatergic synapses in the cochlea and is crucial for hearing. EMBO Mol Med 2016, 8:191–207.

84. FastQC: a quality control tool for high throughput sequence data. [http://www.bioinformatics.babraham.ac.uk/projects/fastqc/]

85. Bolger AM, Lohse M, Usadel B: Trimmomatic: a flexible trimmer for Illumina sequence data. Bioinformatics 2014, 30:2114–2120.

86. Kim D, Langmead B, Salzberg SL: HISAT: a fast spliced aligner with low memory requirements. Nat Methods 2015, 12:357–360.

87. Hartley SW, Mullikin JC: QoRTs: a comprehensive toolset for quality control and data processing of RNA-Seq experiments. BMC Bioinformatics 2015, 16:224.

88. Robinson MD, McCarthy DJ, Smyth GK: edgeR: a Bioconductor package for differential expression analysis of digital gene expression data. Bioinformatics 2010, 26:139–140.

89. Langfelder P, Horvath S: WGCNA: an R package for weighted correlation network analysis. BMC Bioinformatics 2008, 9:559.

90. Love MI, Huber W, Anders S: Moderated estimation of fold change and dispersion for RNA-seq data with DESeq2. Genome Biol 2014, 15:550.

91. Ritchie ME, Phipson B, Wu D, Hu Y, Law CW, Shi W, Smyth GK: limma powers differential expression analyses for RNA-sequencing and microarray studies. Nucleic Acids Res 2015, 43:e47.

92. Untergasser A, Cutcutache I, Koressaar T, Ye J, Faircloth BC, Remm M, Rozen SG: Primer3--new capabilities and interfaces. Nucleic Acids Res 2012, 40:e115.

93. Bonfield JK, Smith K, Staden R: A new DNA sequence assembly program. Nucleic Acids Res 1995, 23:4992–4999.

94. Livak KJ, Schmittgen TD: Analysis of relative gene expression data using real-time quantitative PCR and the 2(T)(-Delta Delta C) method. Methods 2001, 25:402–408.

95. Morrison A, Hodgetts C, Gossler A, Hrabe de Angelis M, Lewis J: Expression of Delta1 and Serrate1 (Jagged1) in the mouse inner ear. Mech Dev 1999, 84:169–172.

96. Zine A, Van De Water TR, de Ribaupierre F: Notch signaling regulates the pattern of auditory hair cell differentiation in mammals. Development 2000, 127:3373–3383.

97. Bridge PD, Sawilowsky SS: Increasing physicians’ awareness of the impact of statistics on research outcomes: comparative power of the t-test and and Wilcoxon Rank-Sum test in small samples applied research. J Clin Epidemiol 1999, 52:229–235.

98. Duricki DA, Soleman S, Moon LD: Analysis of longitudinal data from animals with missing values using SPSS. Nat Protoc 2016, 11:1112–1129.

99. Krueger C, Tian L: A comparison of the general linear mixed model and repeated measures ANOVA using a dataset with multiple missing data points. Biol Res Nurs 2004, 6:151–157.

100. Athar A, Fullgrabe A, George N, Iqbal H, Huerta L, Ali A, Snow C, Fonseca NA, Petryszak R, Papatheodorou I, et al: ArrayExpress update - from bulk to single-cell expression data. Nucleic Acids Res 2019, 47:D711–D715.

101. Petryszak R, Keays M, Tang YA, Fonseca NA, Barrera E, Burdett T, Fullgrabe A, Fuentes AM, Jupp S, Koskinen S, et al: Expression Atlas update--an integrated database of gene and protein expression in humans, animals and plants. Nucleic Acids Res 2016, 44:D746–752.

102. Lesurf R, Cotto KC, Wang G, Griffith M, Kasaian K, Jones SJ, Montgomery SB, Griffith OL: ORegAnno 3.0: a community-driven resource for curated regulatory annotation. Nucleic Acids Res 2016, 44:D126–132.

103. Chou CH, Shrestha S, Yang CD, Chang NW, Lin YL, Liao KW, Huang WC, Sun TH, Tu SJ, Lee WH, et al: miRTarBase update 2018: a resource for experimentally validated microRNA-target interactions. Nucleic Acids Res 2018, 46:D296–D302.

104. Tong Z, Cui Q, Wang J, Zhou Y: TransmiR v2.0: an updated transcription factor-microRNA regulation database. Nucleic Acids Res 2019, 47:D253–D258.

105. Han H, Cho JW, Lee S, Yun A, Kim H, Bae D, Yang S, Kim CY, Lee M, Kim E, et al: TRRUST v2: an expanded reference database of human and mouse transcriptional regulatory interactions. Nucleic Acids Res 2018, 46:D380–D386.

106. Hertzano R, Dror AA, Montcouquiol M, Ahmed ZM, Ellsworth B, Camper S, Friedman TB, Kelley MW, Avraham KB: Lhx3, a LIM domain transcription factor, is regulated by Pou4f3 in the auditory but not in the vestibular system. Eur J Neurosci 2007, 25:999–1005.

107. Shannon P, Markiel A, Ozier O, Baliga NS, Wang JT, Ramage D, Amin N, Schwikowski B, Ideker T: Cytoscape: a software environment for integrated models of biomolecular interaction networks. Genome Res 2003, 13:2498–2504.

108. Chindelevitch L, Ziemek D, Enayetallah A, Randhawa R, Sidders B, Brockel C, Huang ES: Causal reasoning on biological networks: interpreting transcriptional changes. Bioinformatics 2012, 28:1114–1121.

109. Fakhry CT, Choudhary P, Gutteridge A, Sidders B, Chen P, Ziemek D, Zarringhalam K: Interpreting transcriptional changes using causal graphs: new methods and their practical utility on public networks. BMC Bioinformatics 2016, 17:318.

110. Pollard J, Jr., Butte AJ, Hoberman S, Joshi M, Levy J, Pappo J: A computational model to define the molecular causes of type 2 diabetes mellitus. Diabetes Technol Ther 2005, 7:323–336.

111. Tian C, Johnson KR: TBX1 is required for normal stria vascularis and semicircular canal development. Dev Biol 2020, 457:91–103.

112. Liang S, Zhang JM, Lv ZT, Cheng P, Zhu WT, Chen AM: Identification of Skt11-regulated genes in chondrocytes by integrated bioinformatics analysis. Gene 2018, 677:340–348.

113. Lu Z, Hong CC, Kong G, Assumpcao A, Ong IM, Bresnick EH, Zhang J, Pan X: Polycomb Group Protein YY1 Is an Essential Regulator of Hematopoietic Stem Cell Quiescence. Cell Rep 2018, 22:1545–1559.

114. Emmerson E, May AJ, Nathan S, Cruz-Pacheco N, Lizama CO, Maliskova L, Zovein AC, Shen Y, Muench MO, Knox SM: SOX2 regulates acinar cell development in the salivary gland. Elife 2017, 6.

115. Artigas N, Gamez B, Cubillos-Rojas M, Sanchez-de Diego C, Valer JA, Pons G, Rosa JL, Ventura F: p53 inhibits SP7/Osterix activity in the transcriptional program of osteoblast differentiation. Cell Death Differ 2017, 24:2022–2031.

116. Zhang S, Kim JY, Xu S, Liu H, Yin M, Koroleva M, Guo J, Pei X, Jin ZG: Endothelial-specific YY1 governs sprouting angiogenesis through directly interacting with RBPJ. Proc Natl Acad Sci U S A 2020, 117:4792–4801.

117. Zeng ZM, Du HY, Xiong L, Zeng XL, Zhang P, Cai J, Huang L, Liu AW: BRCA1 protects cardiac microvascular endothelial cells against irradiation by regulating p21-mediated cell cycle arrest. Life Sci 2020, 244:117342.

118. Shao A, Lang Y, Wang M, Qin C, Kuang Y, Mei Y, Lin D, Zhang S, Tang J: Bclaf1 is a direct target of HIF-1 and critically regulates the stability of HIF-1alpha under hypoxia. Oncogene 2020.

119. Zhang Y, Fang M, Yang Z, Qin W, Guo S, Ma J, Chen W: GATA Binding Protein 4 Regulates Tooth Root Dentin Development via FBP1. Int J Biol Sci 2020, 16:181–193.

120. Yu S, Tang Q, Xie M, Zhou X, Long Y, Xie Y, Guo F, Chen L: Circadian BMAL1 regulates mandibular condyle development by hedgehog pathway. Cell Prolif 2020, 53:e12727.

121. Zerbino DR, Achuthan P, Akanni W, Amode MR, Barrell D, Bhai J, Billis K, Cummins C, Gall A, Giron CG, et al: Ensembl 2018. Nucleic Acids Res 2018, 46:D754–D761.

