## Supplementary Figures and Tables for "Hearing impairment due to *Mir183/96/182* mutations suggests both loss and gain of function effects"

**Supporting Information**

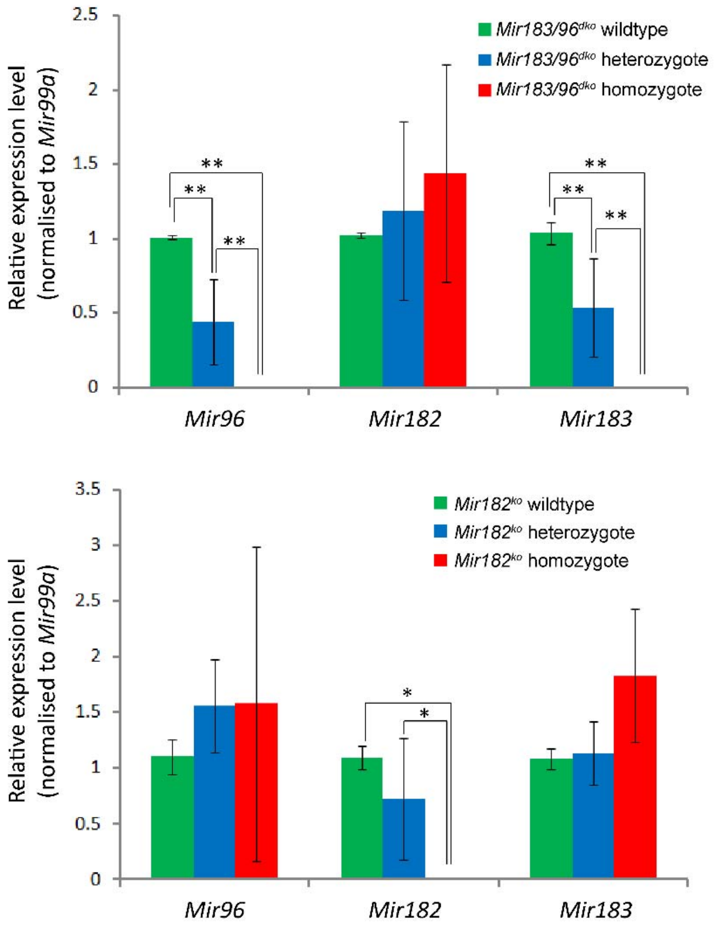

Supplementary Figure S1. Expression of *Mir96*, *Mir182* and *Mir183* in *Mir183/96*$^{dko}$ mutant mice (top) and *Mir182*$^{ko}$ mutant mice (bottom), relative to *Mir99a*, which is expressed in cochlear sensory epithelium. Homozygote (red; right bars) and heterozygote (blue; middle bars) expression levels have been normalised to expression in the wildtype (green; left bars). *Mir183/96*$^{dko}$: wildtype n=7, heterozygote n=5, homozygote n=6. One way ANOVA: Mir96 p<0.001 (wildtype vs. heterozygote Bonferroni-corrected p<0.001; wildtype vs. homozygote Bonferroni-corrected p<0.001; heterozygote vs. homozygote Bonferroni-corrected p=0.001) ; Mir182 p=0.37; Mir183 p<0.001 (wildtype vs. heterozygote Bonferroni-corrected p=0.001; wildtype vs. homozygote Bonferroni-corrected p<0.001; heterozygote vs. homozygote Bonferroni-corrected p<0.001). *Mir182*$^{ko}$: wildtype n=4, heterozygote n=4, homozygote n=4. One way ANOVA: Mir96 p=0.685; Mir182 p=0.003 (wildtype vs. heterozygote

Bonferroni-corrected p=0.397; wildtype vs. homozygote Bonferroni-corrected p=0.003; heterozygote vs. homozygote Bonferroni-corrected p=0.032); Mir183 p=0.04 (wildtype vs. heterozygote Bonferroni-corrected p=1.0; wildtype vs. homozygote Bonferroni-corrected p=0.068; heterozygote vs. homozygote Bonferroni-corrected p=0.094), Error bars are standard deviation (* = $P < 0.05$, ** = $P \leq 0.01$).

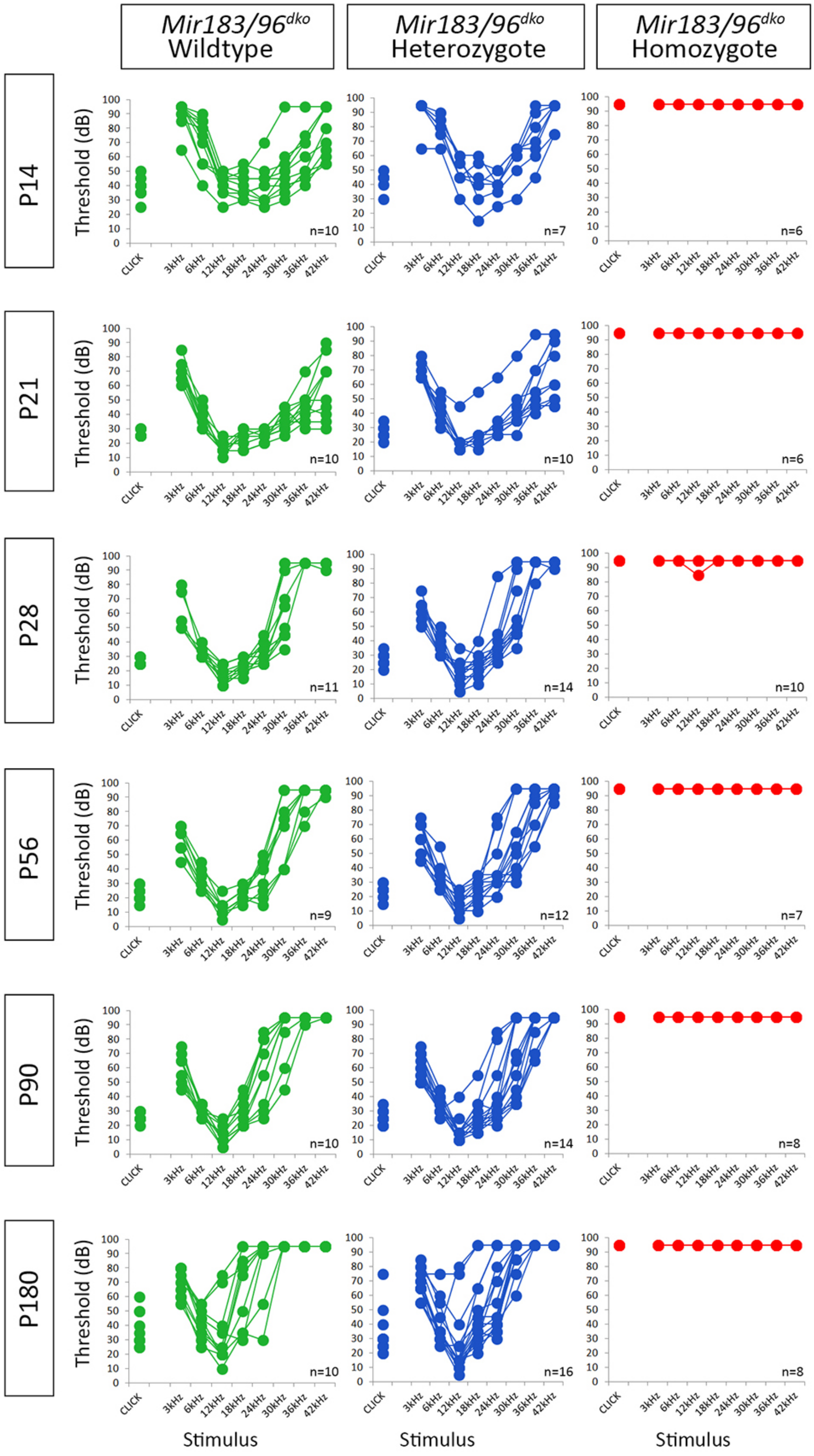
Mir183/96dko Wildtype
Mir183/96dko Heterozygote
Mir183/96dko Homozygote
P14
P21
P28
P56
P90
P180
Threshold (dB)
CLICK
3kHz
6kHz
12kHz
18kHz
24kHz
30kHz
36kHz
42kHz
n=10
n=7
n=6
n=10
n=10
n=6
n=11
n=14
n=10
n=9
n=12
n=7
n=10
n=14
n=8
n=10
n=16
n=8
Stimulus

Supplementary Figure S2. Individual ABR thresholds of wildtype, heterozygous and homozygous *Mir183/96*$^{dko}$ mice at all ages tested. Number of mice of each genotype tested at each age is shown on the threshold plot.

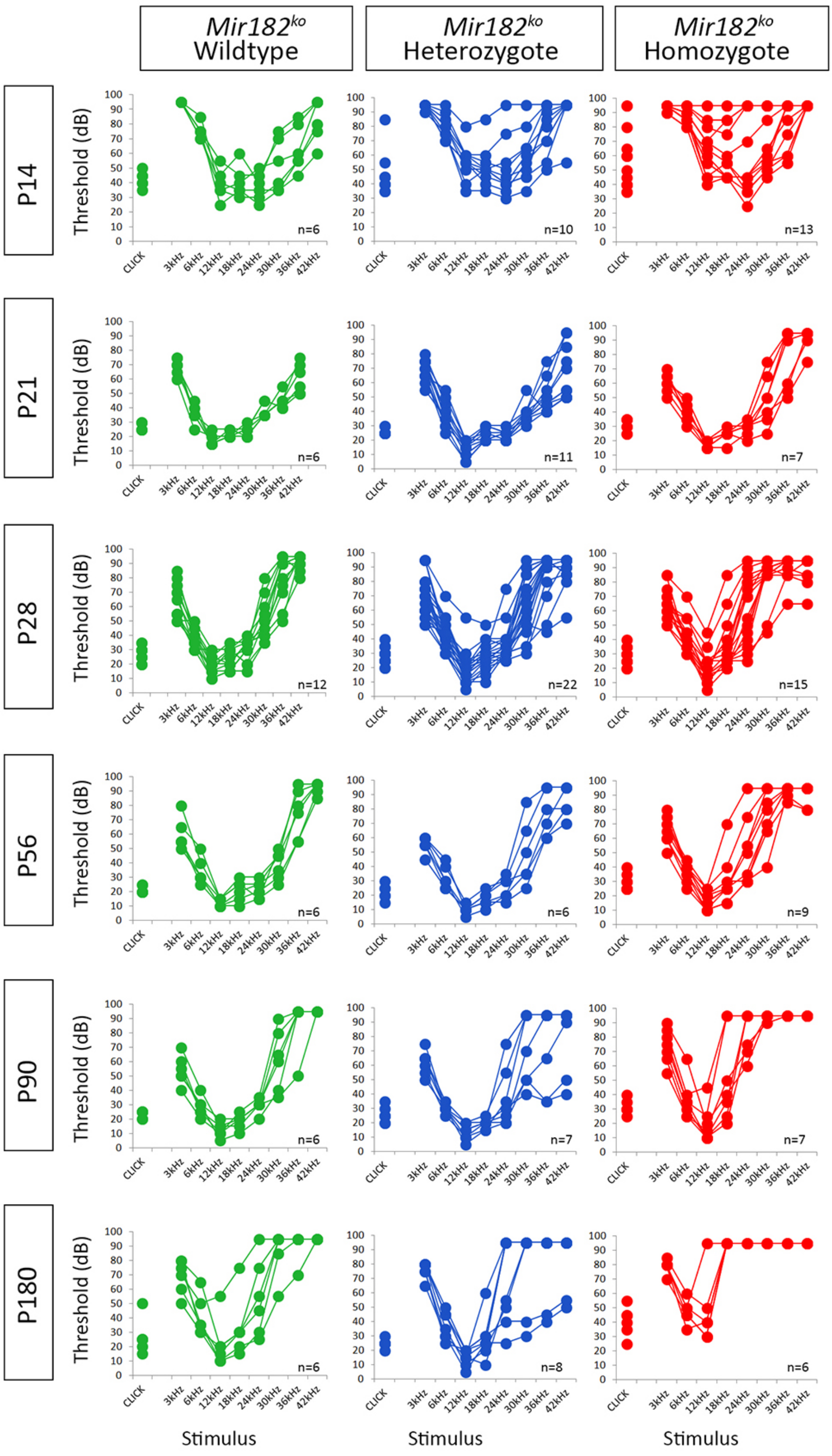

Mir182ko Wildtype
Mir182ko Heterozygote
Mir182ko Homozygote
P14
P21
P28
P56
P90
P180
Threshold (dB)
Stimulus
CLICK
3kHz
6kHz
12kHz
18kHz
24kHz
30kHz
36kHz
42kHz
n=6
n=10
n=13
n=6
n=11
n=7
n=12
n=22
n=15
n=6
n=6
n=9
n=6
n=7
n=7
n=6
n=8
n=6

Supplementary Figure S3. Individual ABR thresholds of wildtype, heterozygous and homozygous *Mir182*$^{ko}$ mice at all ages tested. Number of mice of each genotype tested at each age is shown on the threshold plot.

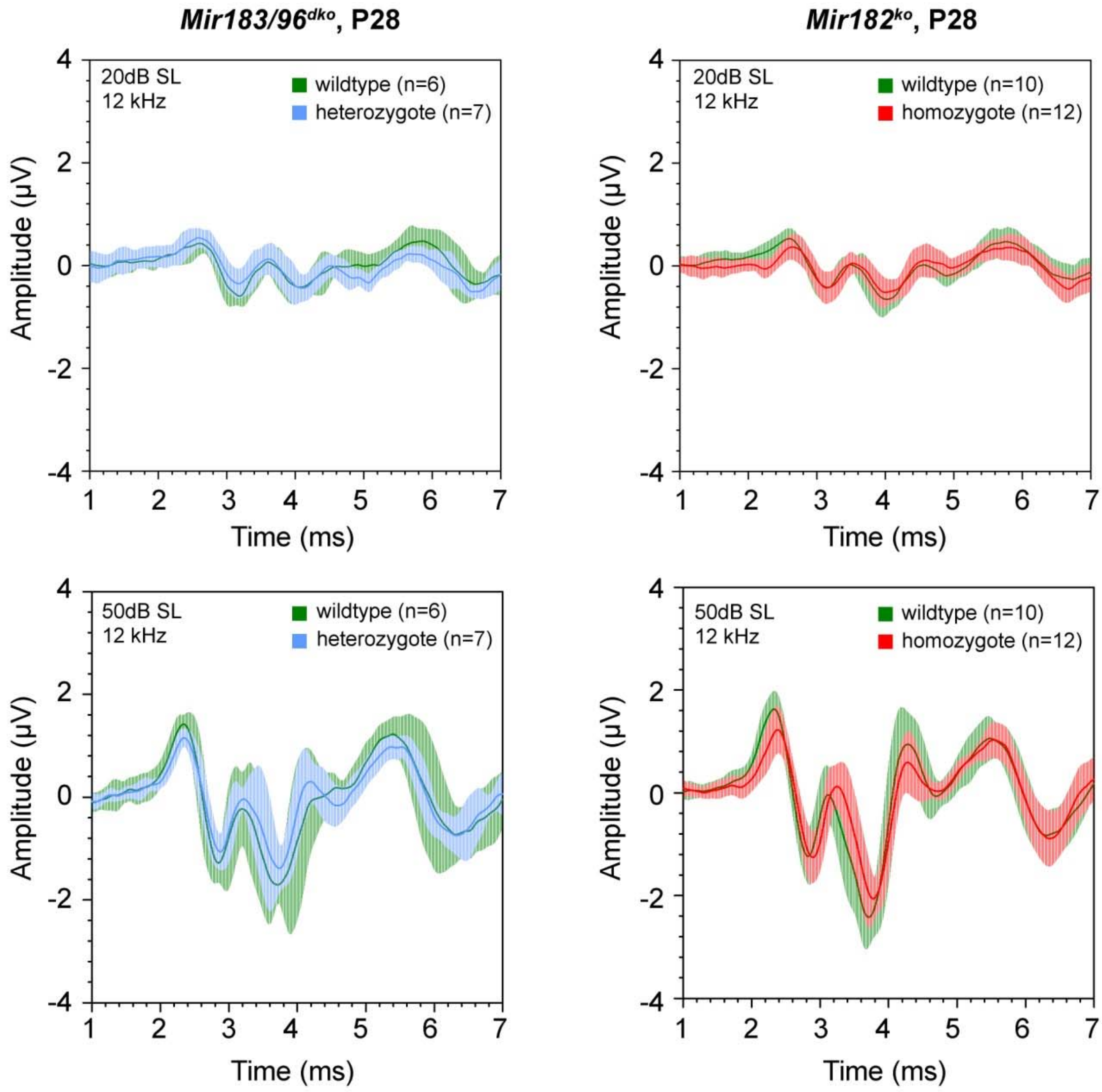

Supplementary Figure S4. Mean ABR waveforms at 12kHz, shown at 20dB (top) and 50dB (bottom) above threshold (sensation level, SL) ± standard deviation, at four weeks old. There is no obvious difference between *Mir183/96$^{dko}$* heterozygous (blue, n=7) and wildtype mice (green, n=6) (left), or between *Mir182$^{ko}$* homozygous (red, n=12) and wildtype mice (green, n=10) (right) at either sensation level.

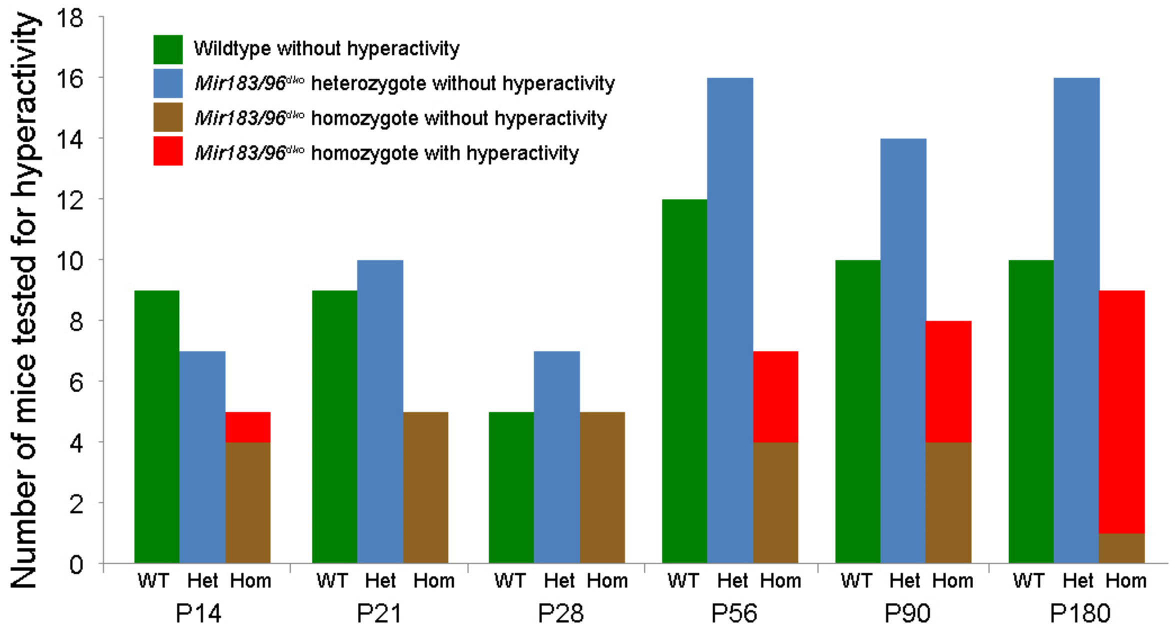

Supplementary Figure S5. Numbers of mice assessed for hyperactivity noted at different ages in wildtype (green, WT), heterozygote (blue, Het) and homozygote (red and brown, Hom) *Mir183/96*$^{dko}$ mice. Only homozygotes showed any vestibular phenotype, the incidence of which increased with age. Bright red indicates homozygous mice with hyperactive behaviour, and brown indicates homozygotes without hyperactivity.

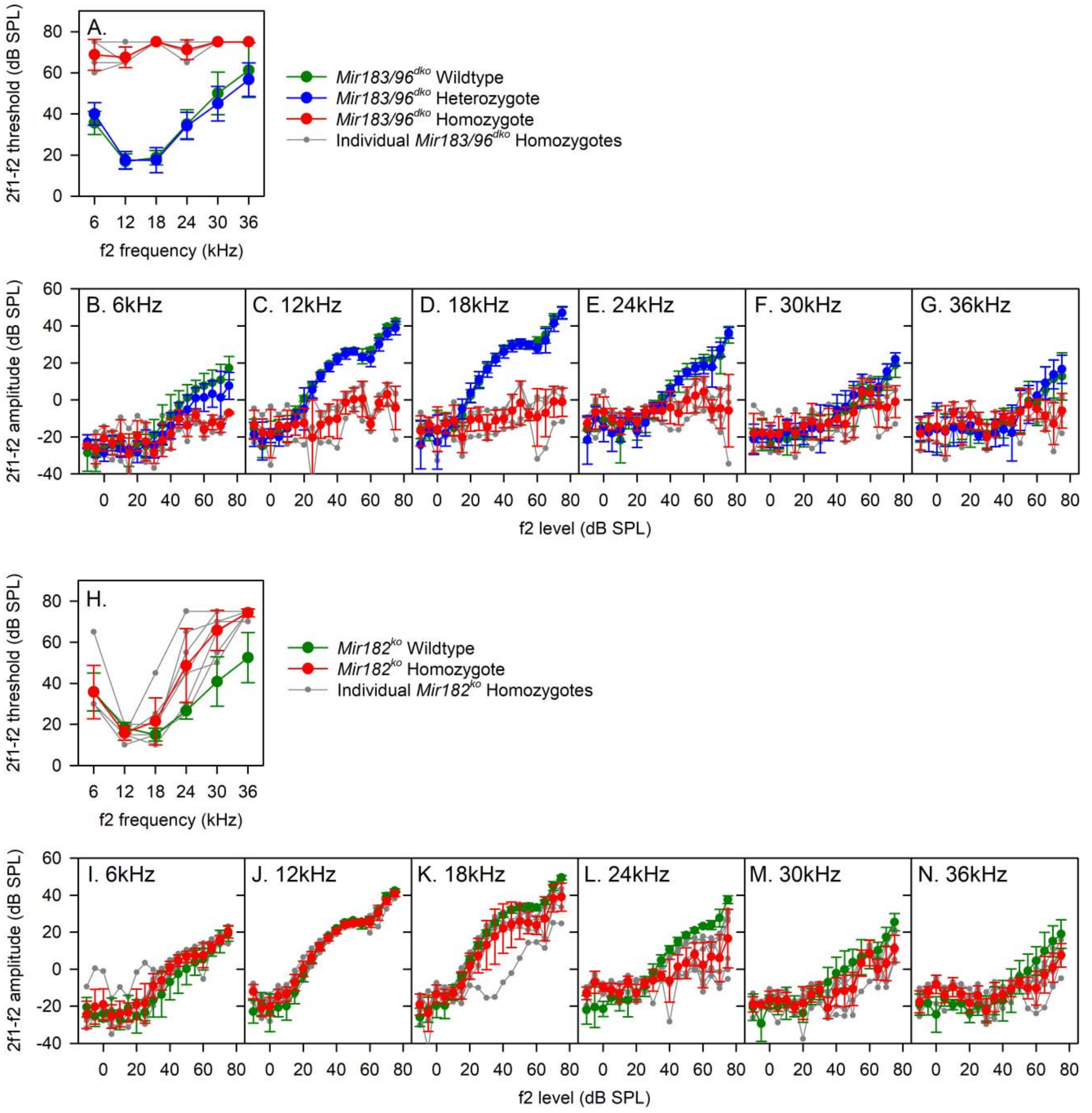

Supplementary Figure S6. Distortion Product Otoacoustic Emission (DPOAE) measurements from *Mir183/96*$^{dko}$ and *Mir182*$^{ko}$ mice. (A-G) DPOAEs recorded from *Mir183/96*$^{dko}$ mice; wildtype n=8, heterozygote n=6, homozygote n=4. (H-N) DPOAEs recorded from *Mir182*$^{ko}$ mice; wildtype n=6, homozygote n=7. Mean responses (±standard deviation) are indicated by green (wildtype), blue (heterozygote) and red (homozygote) lines & symbols. Responses from individual homozygote animals are indicated grey lines & symbols. (A, H) The threshold of the 2f1-f2 DPOAE (as defined in the methods) is plotted as a function of f2 frequency. (B-G, I-N). The amplitude of the 2f1-f2 DPOAE is plotted as a function of f2 level (dB SPL) for the range of f2 tones used; 6kHz (B & I), 12kHz (C & J), 18kHz (D & K), 24kHz (E & L), 30kHz (F & M) and 36kHz (G & N).

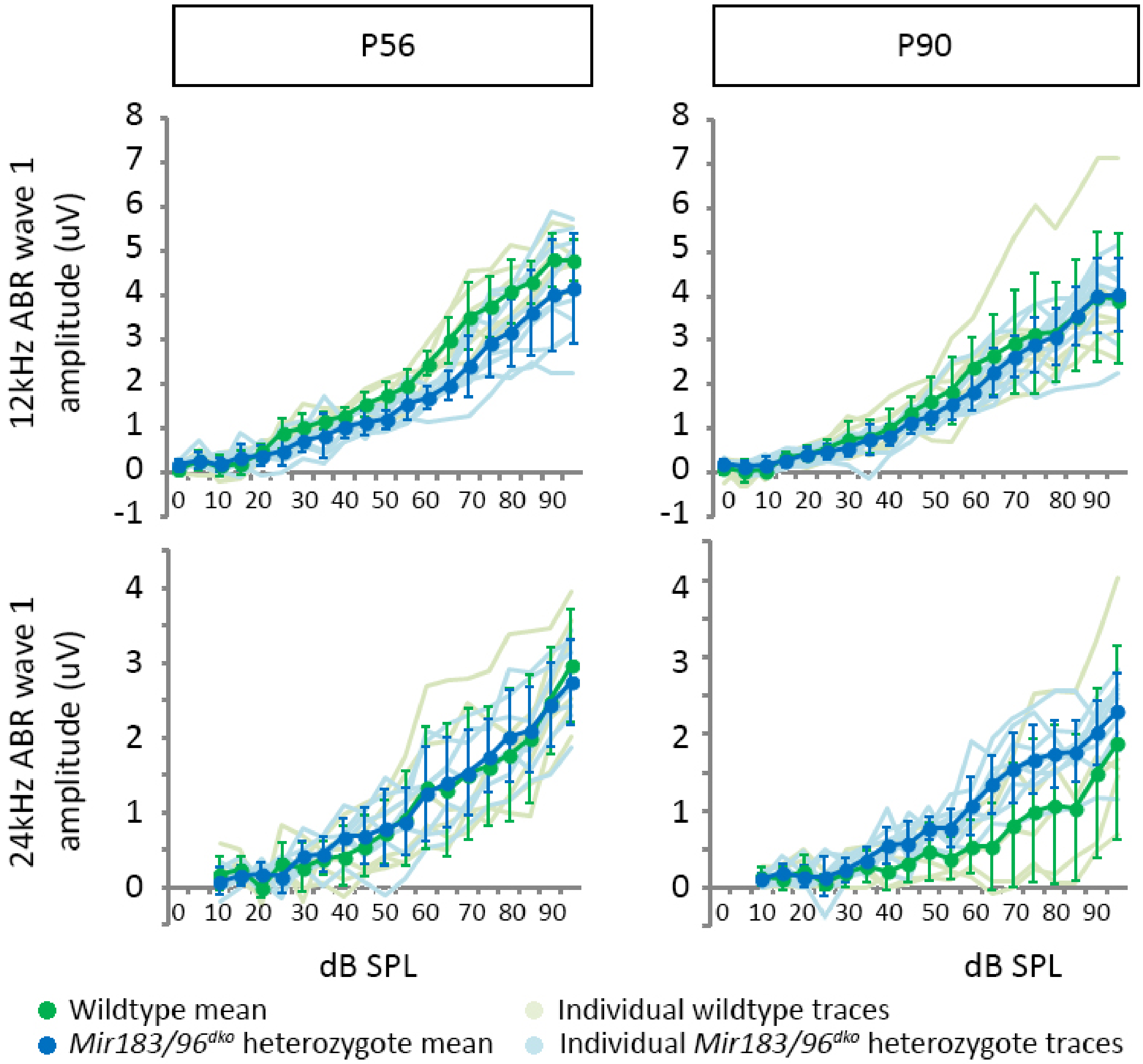

Supplementary Figure S7. ABR wave 1 amplitudes at 12kHz and 24kHz for *Mir183/96*$^{dko}$ wildtype (green) and heterozygous (blue) mice at P56 (n=6 wildtypes, n=9 heterozygotes) and P90 (n=7 wildtypes, n=12 heterozygotes). Individual wave 1 amplitudes are also plotted in pale green (wildtype) and pale blue (heterozygote). Heterozygous amplitudes appear similar to wildtype at both ages and both frequencies. Error bars are standard deviation.

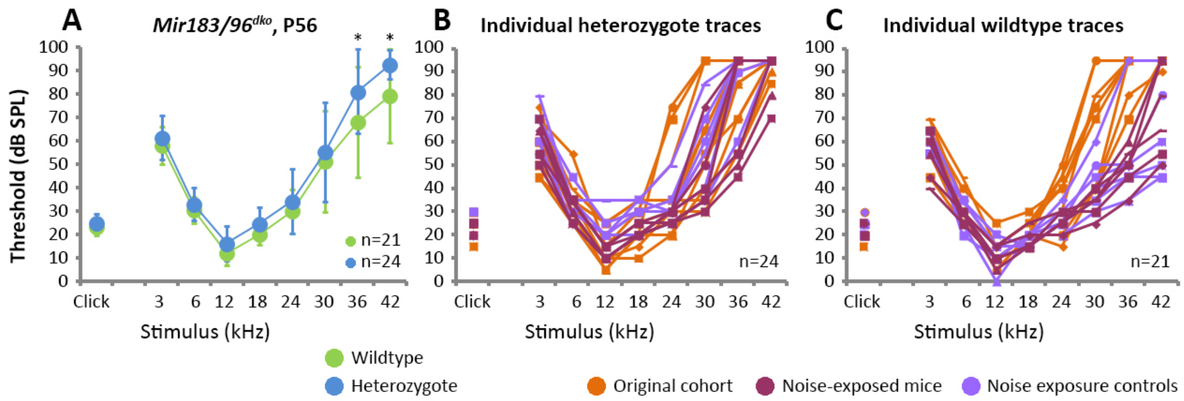

Supplementary Figure S8. All ABR thresholds from *Mir183/96*$^{dko}$ heterozygous and wildtype mice at 8 weeks old (53-58 days). (A)Means of all mice tested. Heterozygotes are shown in blue (n=24) and wildtypes in green (n=21). Error bars are standard deviation (* = Bonferroni-corrected p < 0.05, mixed linear model pairwise comparison; see supplementary data for all p-values). (B) Individual thresholds from heterozygous mice. (C) Individual thresholds from wildtype mice. In B and C, the original mice tested are coloured orange, the noise-exposed mice are coloured dark maroon and the noise exposure control mice are coloured lilac.

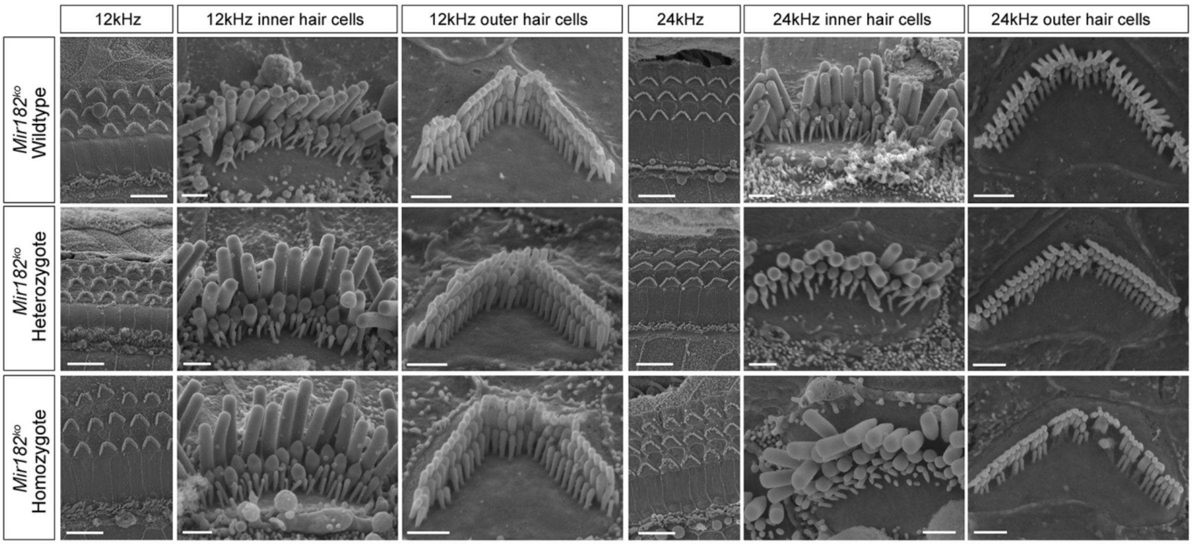

Supplementary Figure S9. Scanning electron micrographs of *Mir182*$^{ko}$ mice at P28. Two best-frequency regions of the organ of Corti are shown; 12kHz (68% of the way along the organ of Corti from base to apex) and 24kHz (43% of the way along the organ of Corti from base to apex). For each region, the left-hand column shows a zoomed-out image with inner and outer hair cell rows (scale bars=10µm), and the other two columns show an inner and an outer hair cell close up (scale bars=1µm). The top row shows wildtype hair cells (n=1), the middle row shows heterozygote hair cells (n=2) and the bottom row shows homozygote hair cells (n=1).

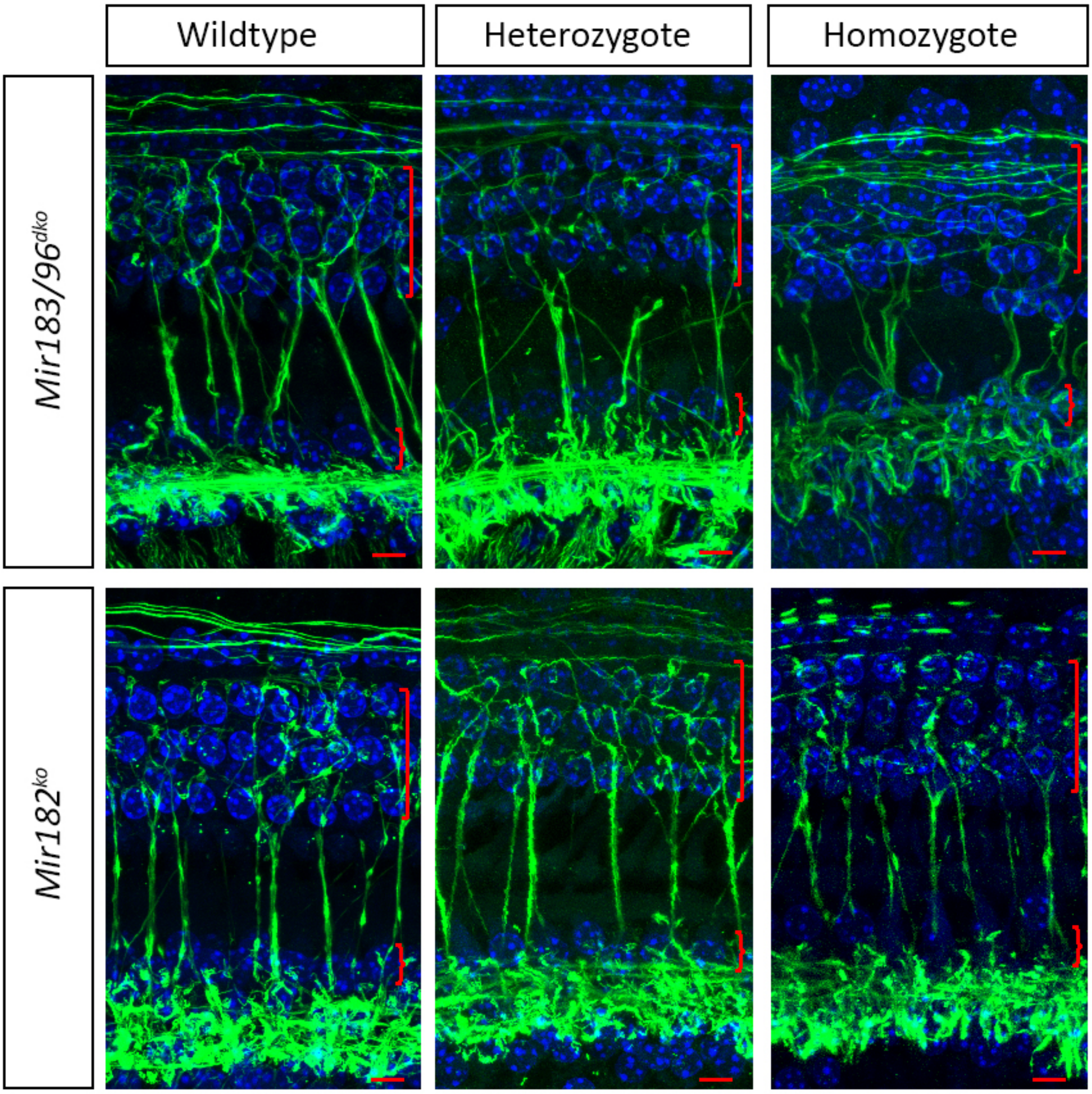

Supplementary Figure S10. Innervation of inner and outer hair cells of *Mir183/96*$^{dko}$ mice (wildtype n=9, heterozygote n=8, homozygote n=4), and *Mir182*$^{ko}$ mice (wildtype n=5, heterozygote n=6, homozygote n=10) at P28. Nerve fibres are stained with anti-neurofilament antibody (green) and nuclei are labelled with DAPI (blue). All panels show the 12kHz best-frequency region. Square brackets indicate the three rows of outer hair cell nuclei, and curly brackets the single row of inner hair cell nuclei. Scale bar = 5µm.

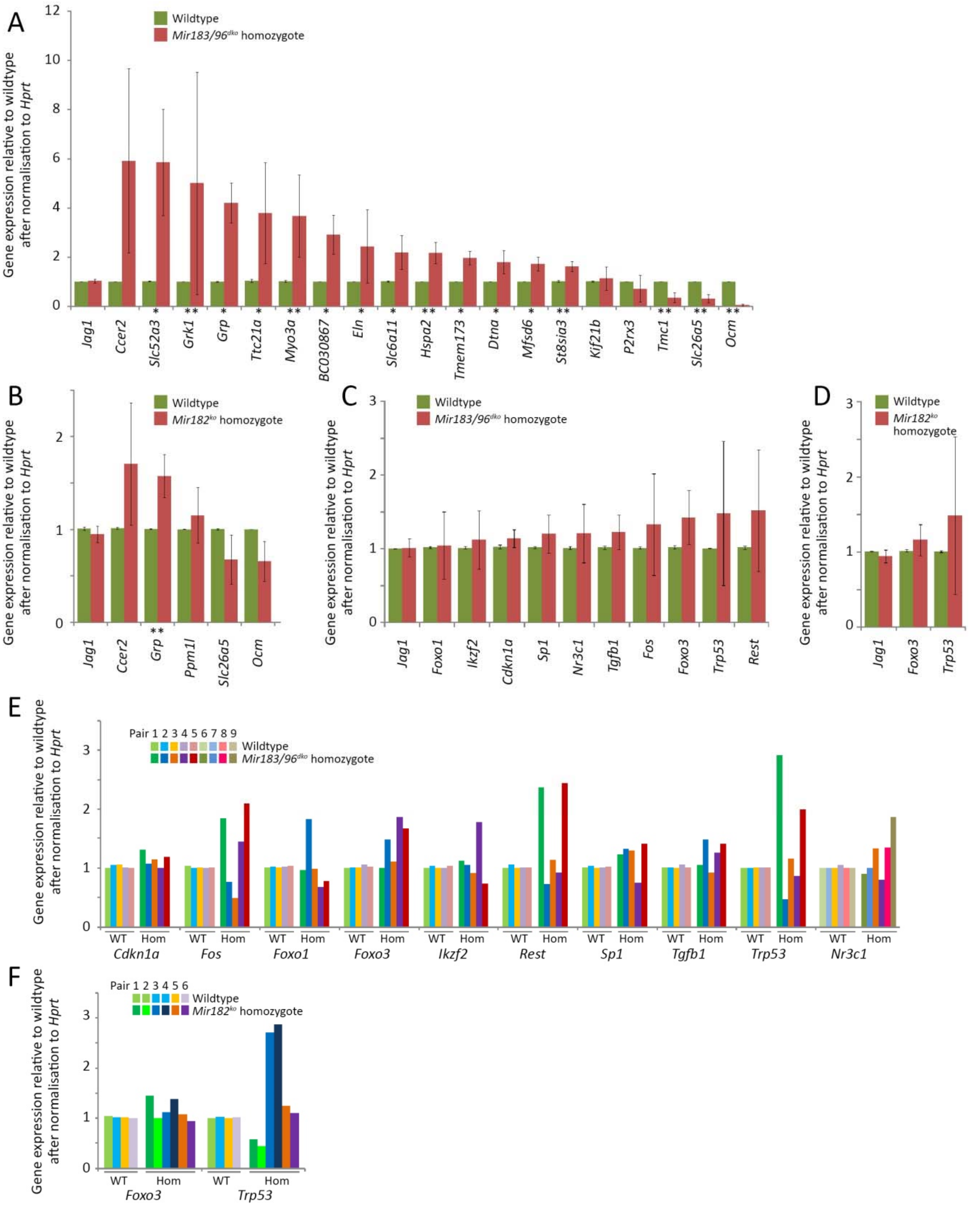

Supplementary Figure S11. Confirmation testing of RNA-seq results in *Mir183/96*$^{dko}$ (A) and *Mir182*$^{ko}$ (B) homozygotes and testing of network nodes in *Mir183/96*$^{dko}$ (C, E) and *Mir182*$^{ko}$ (D, F) homozygotes. qRTPCR was carried out on cDNA from P4 organs of Corti in wildtype (green; left bar in A-D) and homozygote (red; right bar in A-D) littermates to test gene expression changes. E and F show the expression levels of the network genes in individual wildtype-homozygote pairs, showing the high variability between mice. Error bars are standard deviation (* = P < 0.05, ** = P < 0.01). All p-values were calculated using the Wilcoxon rank sum test. (A) *Jag1* n=7 pairs, p=0.38; *Ccer2* n=6 pairs, p=0.065; *Slc52a3* n=6, pairs, p=0.0022; *Grk1* n=6 pairs, p=0.0022; *Grp* n=6 pairs, p= 0.0022; *Myo3a* n=6 pairs, p=0.0022; *Ttc21a* n=6 pairs, p=0.065; *BC030867* n=6 pairs, p=0.0022; *Slc6a11* n=6 pairs, p=0.0022; *Eln* n=6 pairs, p=0.0022; *Hspa2* n=6 pairs, p=0.0022; *Tmem173* n=6 pairs, p=0.0022; *Mfsd6* n=6 pairs, p=0.0022; *Dtna* n=6 pairs, p=0.0022; *St8sia3* n=6 pairs, p=0.015; *Kif21b* n=6 pairs, p=0.39; *P2rx3* n=6 pairs, p=0.065; *Tmc1* n=6 pairs, p=0.0022; *Slc26a5* n=6 wildtypes, 7 homozygotes, p=0.0012; *Ocm* n=6 wildtypes, 7 homozygotes, p=0.0012. (B) *Jag1* n=6 pairs, p=0.70; *Ccer2* n=6 pairs, p=0.065; *Grp* n=6 pairs, p=0.0022; *Ppm1l* n=6 pairs, p=0.39; *Slc26a5* n=6 pairs, p=0.065; *Ocm* n=6 pairs, p=0.065. (C) *Jag1* n=9 pairs, p=0.73; *Foxo1* n=5 pairs, p=0.15; *Ikzf2* n=5 pairs, p=0.69; *Cdkn1a* n=5 pairs, p=0.15; *Sp1* n=5 pairs, p=0.15; *Nr3c1* n=6 pairs, p=1; *Tgfb1* n=5 pairs, p=0.22; *Fos* n=5 pairs, p=0.69; *Foxo3* n=5 pairs, p=0.15; *Trp53* n=5 pairs, p=0.69; *Rest* n=5 pairs, p=0.69. (D) *Jag1* n=4 wildtypes, 6 homozygotes, p= 0.11; *Foxo3* n=4 wildtypes, 6 homozygotes p=0.48; *Trp53* n=4 wildtypes, 6 homozygotes p=0.48.

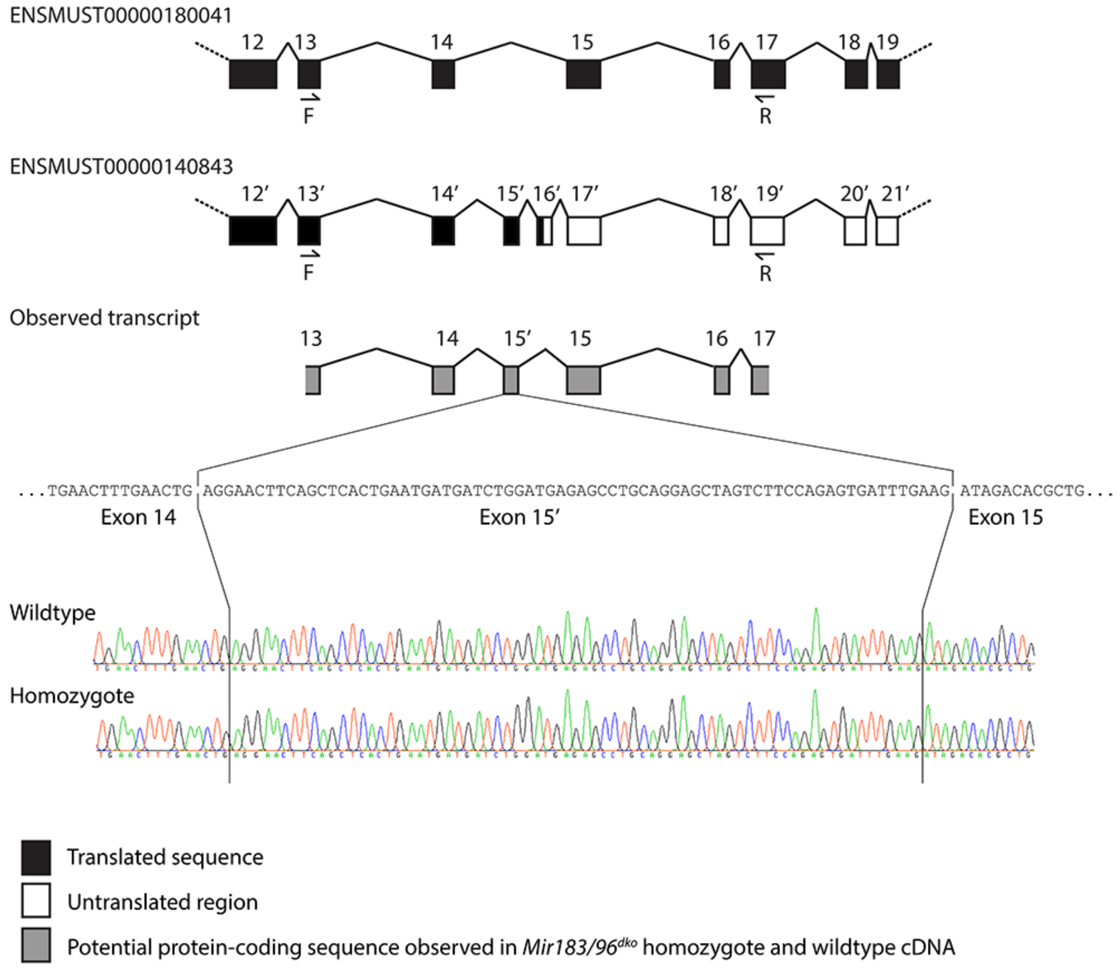

Supplementary Figure S12. Schematic of the novel splice pattern in *Stard9* predicted by JunctionSeq and observed in 4 wildtype and 4 homozygous *Mir183/96*$^{dko}$ mice. Exons 12-19 of the Ensembl protein-coding transcript ENSMUST00000180041 are shown at the top, and exons 12-21 of the nonsense-mediated decay transcript ENSMUST00000140843 underneath. We sequenced exons 13-17 from the protein-coding transcript (the positions of the primers used are marked with “F” and “R”) and found an exon between exons 14 and 15 corresponding to exon 15’ from the nonsense-mediated decay transcript (ENSMUSE00001437951). The sequence and traces are shown at the bottom. Both wildtype and homozygous sequences included exon 15’ and neither showed any sign of alternative splicing around it.

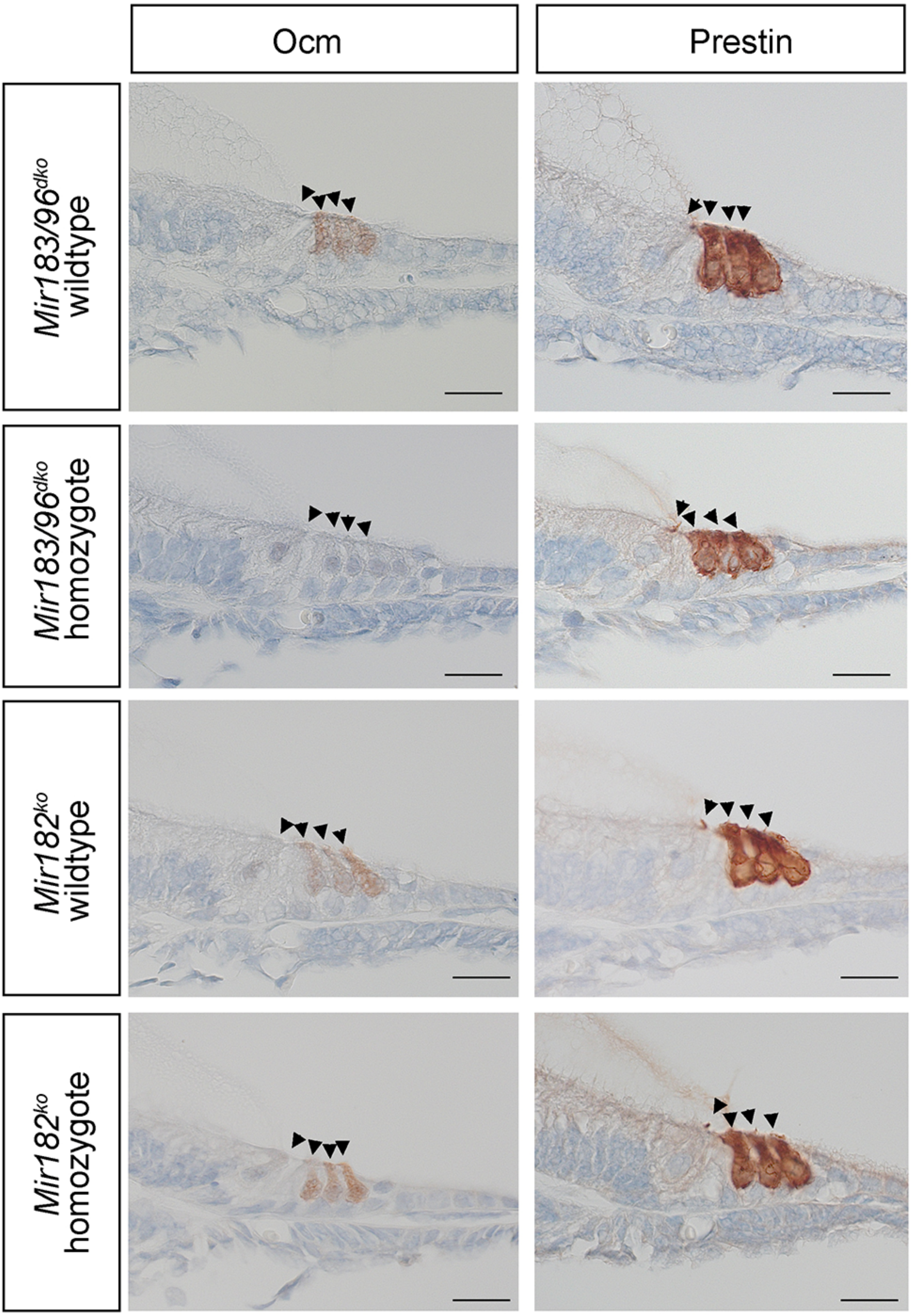

Ocm
Prestin
Mir183/96dko wildtype
Mir183/96dko homozygote
Mir182ko wildtype
Mir182ko homozygote

Supplementary Figure S13. Ocm (left) and Prestin (right) antibody stains in *Mir183/96*$^{dko}$ wildtypes and homozygotes , and *Mir182*$^{ko}$ wildtypes and homozygotes. No Ocm stain is visible in *Mir183/96*$^{dko}$ homozygotes. Hair cells are indicated by arrowheads. Scale bar = 10µm. 3 homozygotes and 3 wildtype littermates were tested with each antibody.

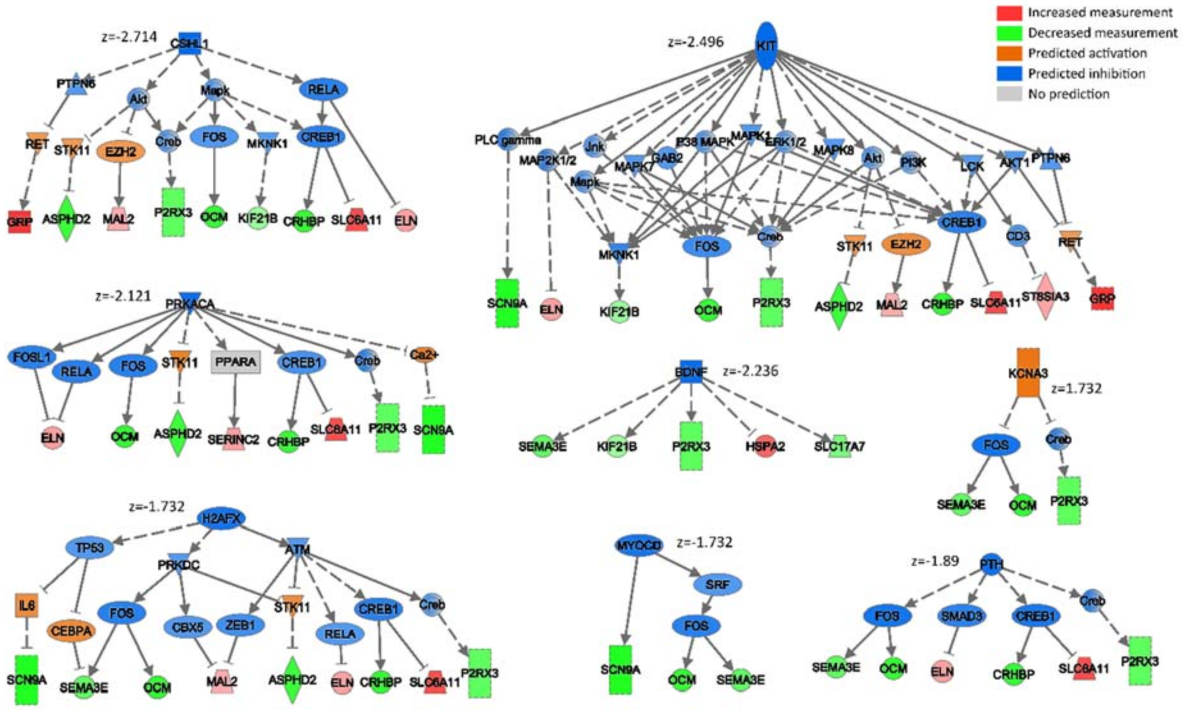

Supplementary Figure S14. Networks generated by Ingenuity Pathway Analysis from the *Mir183/96*$^{dko}$ RNA-seq data, showing predicted upstream regulators which may be responsible for some of the misregulation observed in the data. Misregulated genes are arranged on the lowest row, coloured according to observed misregulation (pink/red = upregulated, green = downregulated in mutants). The top row(s) contain predicted regulators (orange = predicted upregulation, blue = predicted downregulation). Predicted links inconsistent with the observed misregulation have been removed. The intensity of the colour indicates the level of observed or predicted misregulation. Dotted lines represent indirect regulation, and solid lines direct regulation. The z-score of each network, which is both a prediction of the direction of misregulation of the root regulator and a measure of the match of observed and predicted gene misregulation, is shown in the figure. A significant z-score is one with an absolute value greater than 2. A negative score indicates downregulation and a positive score upregulation of the root regulator. miR-96 is not one of the identified upstream regulators.

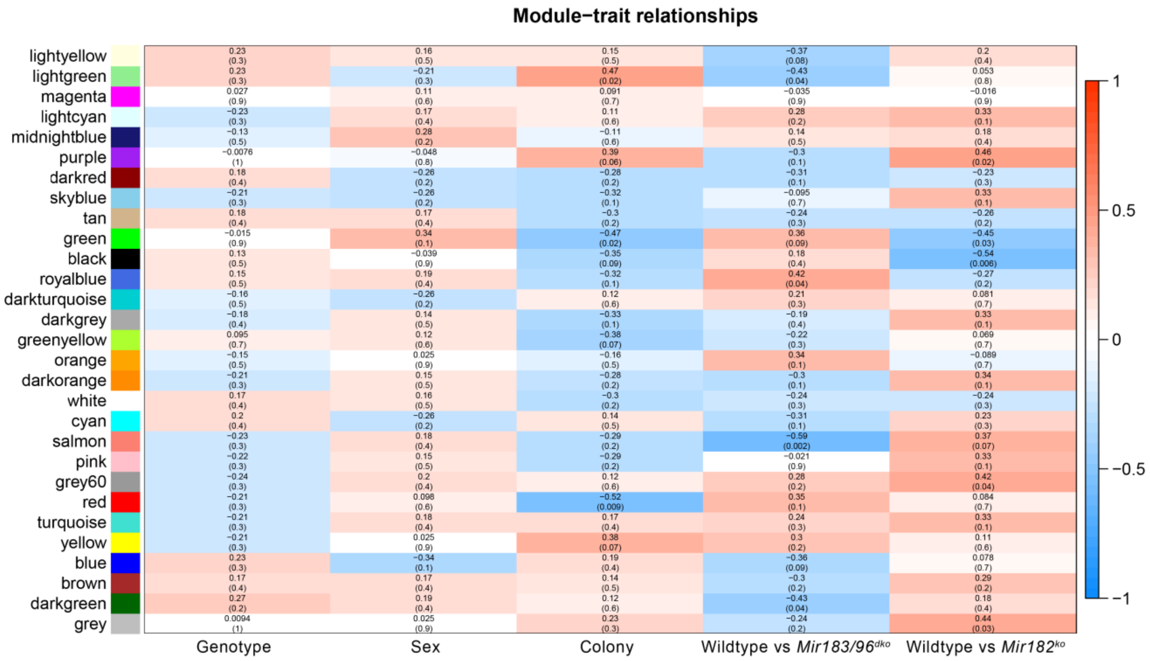

Supplementary Figure S15. Associations of the different module eigengenes (rows) with traits (columns). Cell colour indicates correlation level; each cell contains the correlation score and the p-value in brackets.

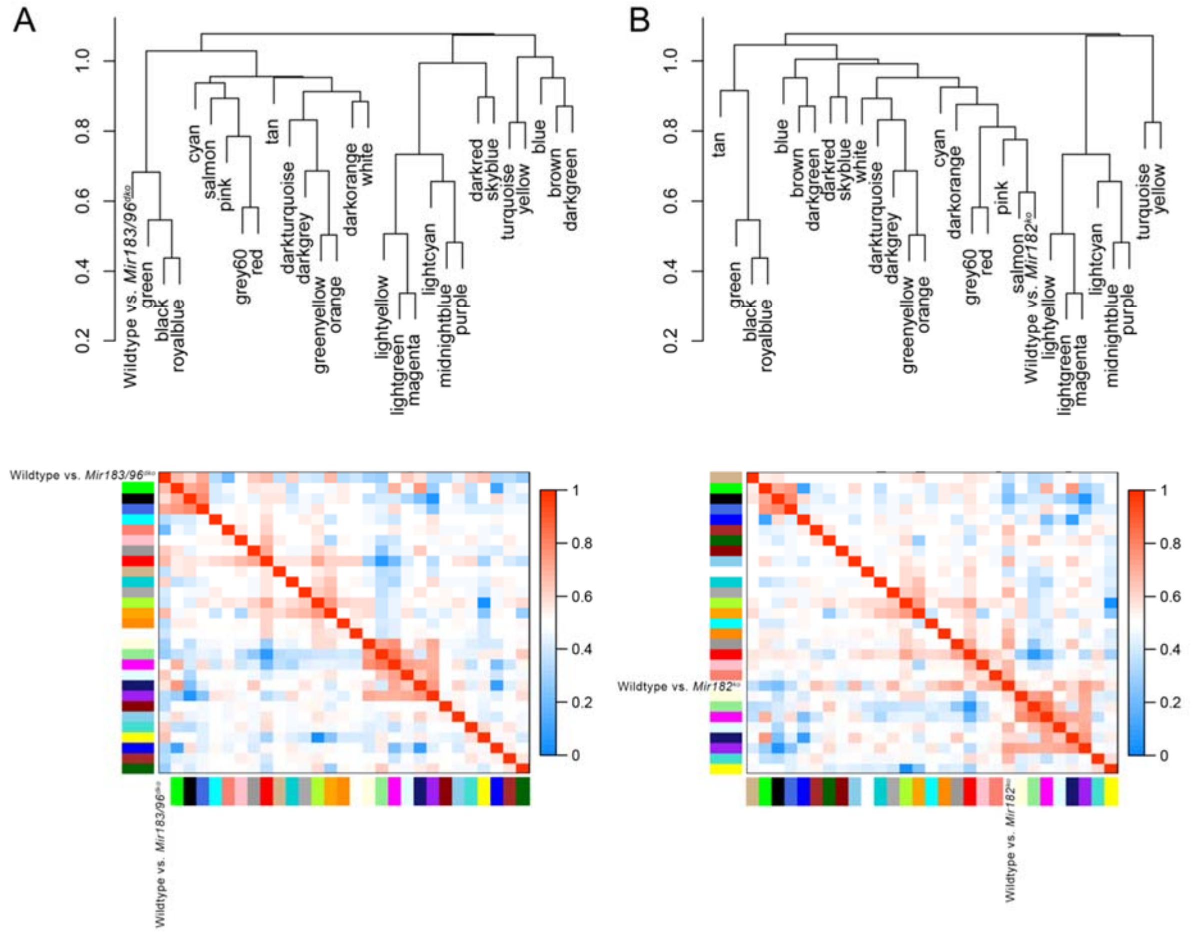

Supplementary Figure S16. Two visualisations of the relationships between the modules and the two genotype traits we examined (wildtype vs *Mir183/96*$^{dko}$ (A) and wildtype vs *Mir182*$^{ko}$ (B)). The top panel shows a hierarchical clustering dendrogram, and the bottom panel shows a heatmap of eigengene correlations. For the heatmaps, the row and column showing the correlation of the genotype trait with the module eigengenes corresponds to that trait's column in Supplementary Figure S15.

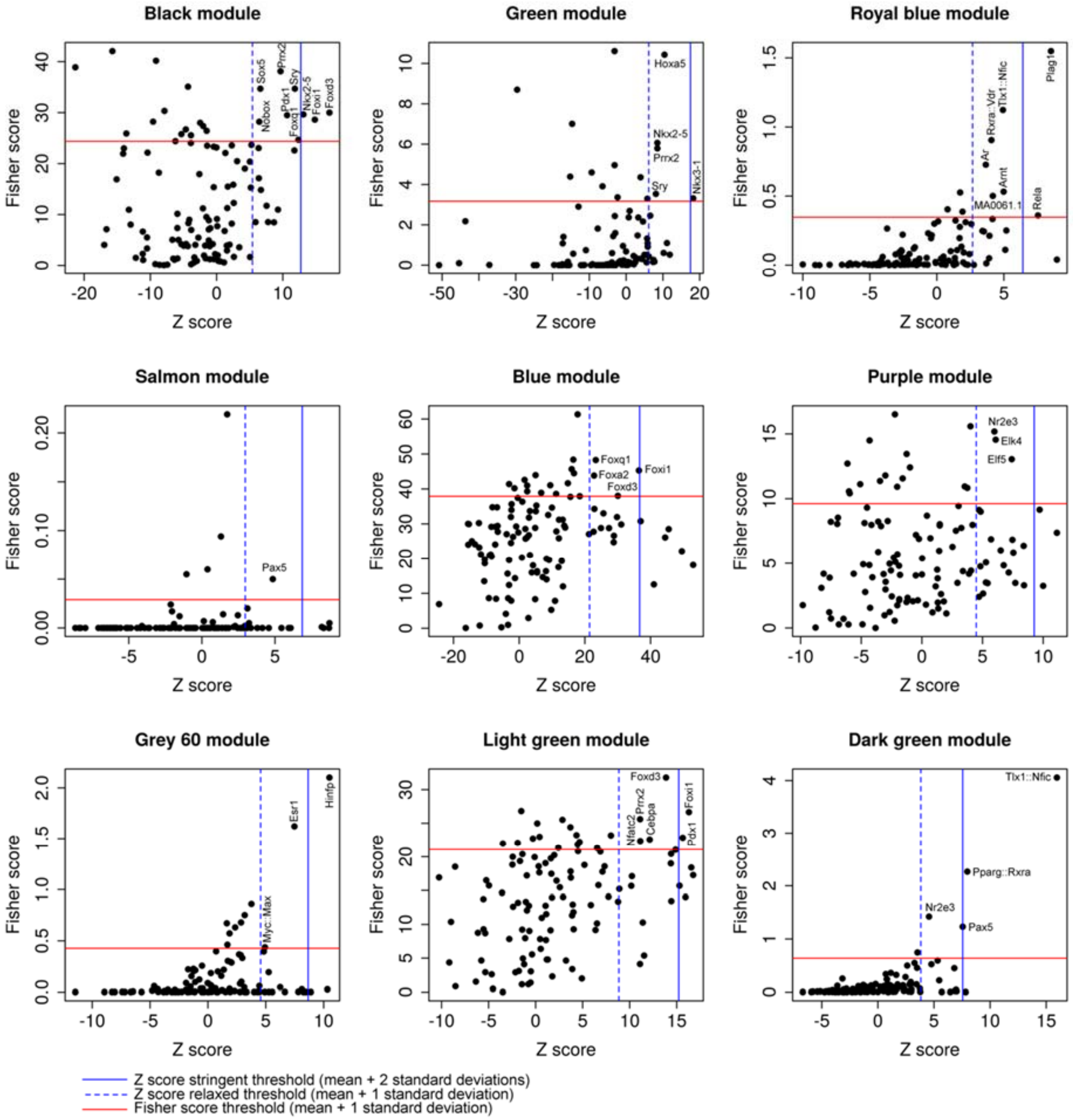

Supplementary Figure S17. Predicted transcription factors for each module, with Z score plotted along the x axis and Fisher score along the y axis. Chosen thresholds are shown in red (for Fisher score) and blue (Z score; the less stringent threshold is a broken line). Transcription factors scoring above both thresholds are labelled.

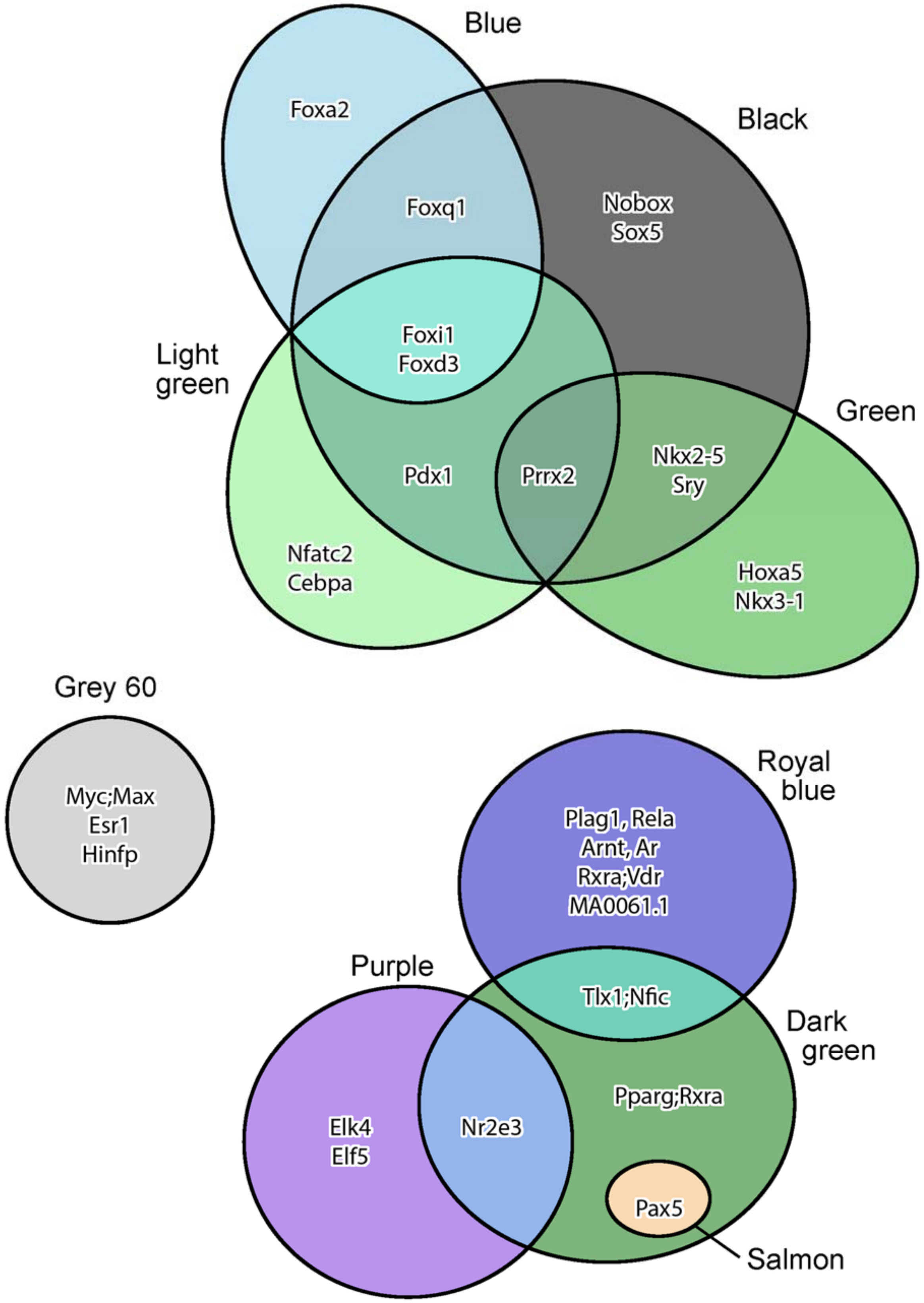

Supplementary Figure S18. Venn diagram showing which of the oPOSSUM transcription factor predictions are shared between modules.

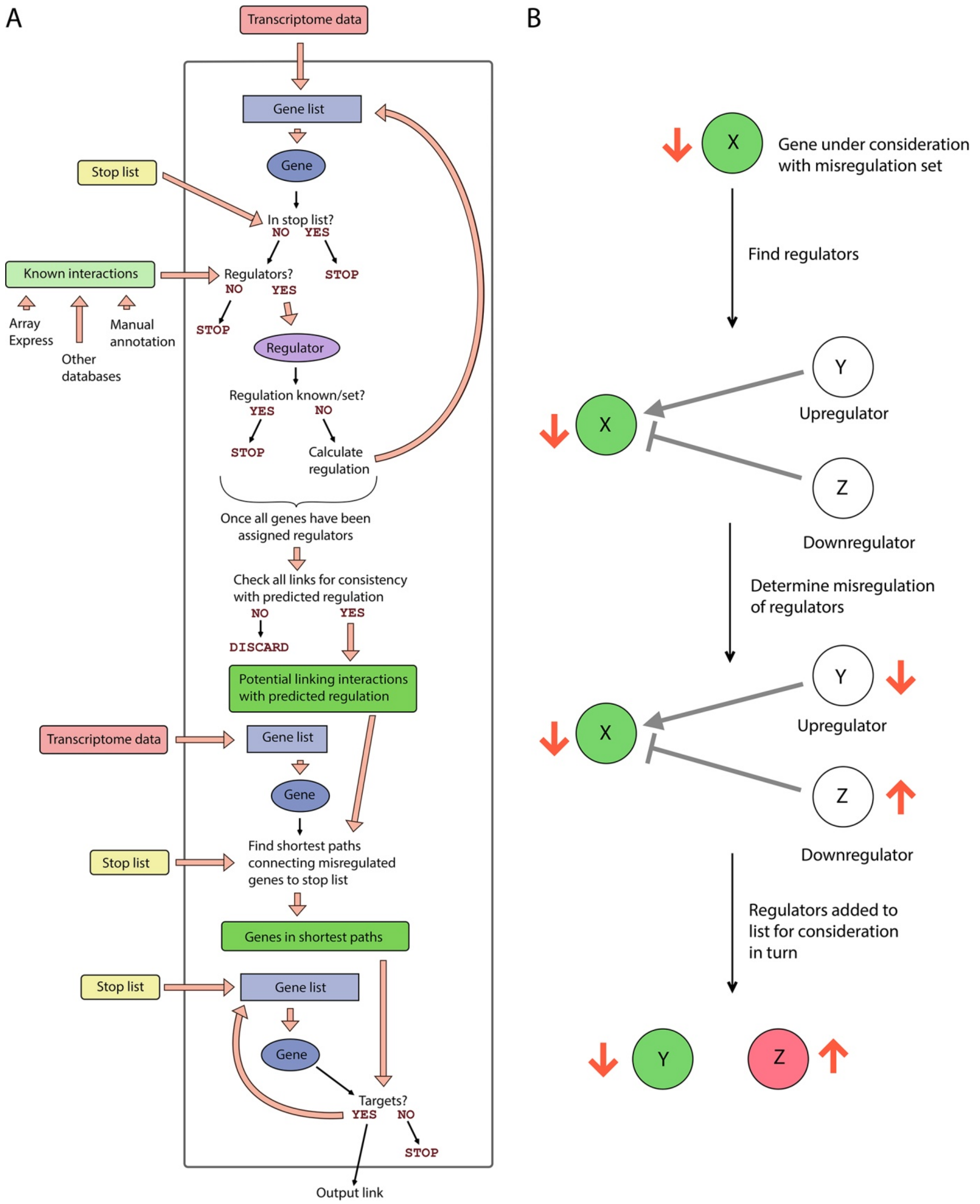

Supplementary Figure S19. The PoPCoRN script. (A) Schematic showing the steps of network construction carried out by the PoPCoRN script. The stop list is a list of genes at which the algorithm stops its upstream searching. In this case, the stop list consisted of the relevant microRNAs (*Mir182* for the *Mir182*$^{ko}$, and *Mir183* and *Mir96* for the *Mir183/96*$^{dko}$). (B) Diagram of the determination of direction of regulation of upstream regulators. Red arrows indicate direction of misregulation. Green indicates a downregulated gene, and pink an upregulated gene.

Supplementary Table S1. Matches for the complement of the miR-96 seed region (GTGCCAA) in 3'UTRs from C57BL/6 and C3H/HeJ sequence. Only genes with at least one match to the miR-96 seed region in their 3' UTRs in one or both strains are shown (1733 in total). The number of matches is colour-coded to aid viewing, from no matches (green) to four matches (red). Direct targets of miR-96 used for creating networks are indicated in the fourth column.

| | Number of miR-96 seed region matches in: | | |
|---|---|---|---|
| **Gene** | **C57BL/6N 3'UTR** | **C3H/HeJ 3'UTR** | **Predicted targets of miR-96** |
| 1110059E24Rik | 1 | 1 | |
| 1700011E24Rik | 1 | 1 | |
| 1700030J22Rik | 1 | 1 | |
| 1700031M16Rik | 1 | 1 | |
| 1810024B03Rik | 1 | 1 | |
| 2010315B03Rik | 1 | 1 | |
| 2210016F16Rik | 1 | 1 | |
| 2410016O06Rik | 1 | 1 | |
| 2610507B11Rik | 1 | 1 | |
| 2610528J11Rik | 1 | 1 | |
| 2700081O15Rik | 1 | 1 | |
| 2810021J22Rik | 1 | 1 | |
| 2810459M11Rik | 1 | 1 | |
| 4930533K18Rik | 1 | 1 | |
| 4931409K22Rik | 1 | 1 | |
| 4931428L18Rik | 1 | 1 | |
| 4933434E20Rik | 1 | 1 | |
| 5730480H06Rik | 1 | 1 | |
| 5730559C18Rik | 1 | 1 | |
| 6430573F11Rik | 1 | 1 | |
| 6820408C15Rik | 1 | 1 | |
| 9330182L06Rik | 1 | 1 | |
| 9430016H08Rik | 1 | 1 | |
| Aasdh | 1 | 1 | |
| Abat | 1 | 1 | |
| Abca1 | 1 | 1 | |
| Abca16 | 1 | 1 | |
| Abca2 | 1 | 1 | |
| Abcb11 | 1 | 1 | |
| Abhd18 | 1 | 1 | |
| Abi2 | 1 | 1 | |
| Acad9 | 1 | 1 | |
| Acod1 | 1 | 1 | |
| Acpp | 1 | 1 | |

| | Number of miR-96 seed region matches in: | | |
|---|---|---|---|
| **Gene** | **C57BL/6N 3'UTR** | **C3H/HeJ 3'UTR** | **Predicted targets of miR-96** |
| Acsl6 | 1 | 1 | |
| Actr1a | 1 | 1 | |
| Acvr2a | 1 | 1 | |
| Adam23 | 1 | 1 | |
| Adam28 | 1 | 1 | |
| Adamts4 | 1 | 1 | |
| Adarb2 | 1 | 1 | |
| Adat3 | 1 | 1 | |
| Adck5 | 1 | 1 | |
| Adcy10 | 1 | 1 | |
| Adgra2 | 1 | 1 | |
| Adgrb3 | 1 | 1 | |
| Adgre5 | 1 | 1 | |
| Adgrf5 | 1 | 1 | |
| Adgrg5 | 1 | 1 | |
| Adgrl1 | 1 | 1 | |
| Adgrv1 | 1 | 1 | |
| Adrbk1 | 1 | 1 | |
| Adsl | 1 | 1 | |
| AF529169 | 1 | 1 | |
| Afm | 1 | 1 | |
| Afmid | 1 | 1 | |
| Agbl5 | 1 | 1 | |
| Ahr | 1 | 1 | |
| AI182371 | 1 | 1 | |
| AI854703 | 1 | 1 | |
| Aifm2 | 1 | 1 | |
| Aifm3 | 1 | 1 | |
| Aipl1 | 1 | 1 | |
| Ajap1 | 1 | 1 | |
| Ak1 | 1 | 1 | |
| Akr1b10 | 1 | 1 | |
| Akr1c13 | 1 | 1 | |
| Akt3 | 1 | 1 | |
| Aldh1a2 | 1 | 1 | |
| Alg13 | 1 | 1 | |
| Alg14 | 1 | 1 | |
| Als2 | 1 | 1 | |
| Amacr | 1 | 1 | |
| Amer3 | 1 | 1 | |
| Angptl4 | 1 | 1 | |
| Angptl7 | 1 | 1 | |
| Ank1 | 1 | 1 | |

| | Number of miR-96 seed region matches in: | | |
|---|---|---|---|
| **Gene** | **C57BL/6N 3'UTR** | **C3H/HeJ 3'UTR** | **Predicted targets of miR-96** |
| Ankib1 | 1 | 1 | |
| Ankra2 | 1 | 1 | |
| Ankrd13c | 1 | 1 | |
| Ankrd27 | 1 | 1 | |
| Ankrd29 | 1 | 1 | |
| Ankrd45 | 1 | 1 | |
| Ano6 | 1 | 1 | |
| Anxa3 | 1 | 1 | |
| Ap1b1 | 1 | 1 | |
| Ap2b1 | 1 | 1 | |
| Ap3m1 | 1 | 1 | |
| Apaf1 | 1 | 1 | |
| Apba2 | 1 | 1 | |
| Aptx | 1 | 1 | |
| Ar | 1 | 1 | |
| Arhgap24 | 1 | 1 | |
| Arhgef12 | 1 | 1 | |
| Arhgef40 | 1 | 1 | |
| Arid3b | 1 | 1 | |
| Arid4b | 1 | 1 | |
| Arid5a | 1 | 1 | |
| Arl11 | 1 | 1 | |
| Arl5b | 1 | 1 | |
| Armcx6 | 1 | 1 | |
| Armt1 | 1 | 1 | |
| Arpc1b | 1 | 1 | |
| Arpc5l | 1 | 1 | |
| Arpp19 | 1 | 1 | |
| Arpp21 | 1 | 1 | |
| Arrb2 | 1 | 1 | |
| Arsj | 1 | 1 | |
| Ash1l | 1 | 1 | |
| Asph | 1 | 1 | |
| Atad3a | 1 | 1 | |
| Atf2 | 1 | 1 | |
| Atf3 | 1 | 1 | |
| Atf6 | 1 | 1 | |
| Atf7ip | 1 | 1 | |
| Atp11a | 1 | 1 | |
| Atp1b4 | 1 | 1 | |
| Atp2b4 | 1 | 1 | |
| Atp5g1 | 1 | 1 | |
| Atp6v0e2 | 1 | 1 | |

| | Number of miR-96 seed region matches in: | | |
|---|---|---|---|
| Gene | C57BL/6N 3'UTR | C3H/HeJ 3'UTR | Predicted targets of miR-96 |
| Atp6v1c1 | 1 | 1 | |
| Atp8b5 | 1 | 1 | |
| Avil | 1 | 1 | |
| B430305J03Rik | 1 | 1 | |
| B630019K06Rik | 1 | 1 | |
| Bace2 | 1 | 1 | |
| Bach2 | 1 | 1 | |
| Baiap2l2 | 1 | 1 | |
| Basp1 | 1 | 1 | |
| Baz1b | 1 | 1 | |
| Bbs9 | 1 | 1 | |
| Bbx | 1 | 1 | |
| Bcl2l11 | 1 | 1 | |
| Bcl7a | 1 | 1 | |
| Bcr | 1 | 1 | |
| Bdh1 | 1 | 1 | |
| Bhlhe40 | 1 | 1 | |
| Bicd2 | 1 | 1 | |
| Bmp2k | 1 | 1 | |
| Bnc2 | 1 | 1 | |
| Brd4 | 1 | 1 | |
| Brms1l | 1 | 1 | |
| Brwd3 | 1 | 1 | |
| Bscl2 | 1 | 1 | |
| Bsdc1 | 1 | 1 | |
| Btbd16 | 1 | 1 | |
| Btla | 1 | 1 | |
| C1d | 1 | 1 | |
| C1galt1 | 1 | 1 | |
| C1qtnf1 | 1 | 1 | |
| C2cd2 | 1 | 1 | |
| C3ar1 | 1 | 1 | |
| C530008M17Rik | 1 | 1 | |
| Cables1 | 1 | 1 | |
| Cabp1 | 1 | 1 | |
| Cacna2d2 | 1 | 1 | Yes |
| Cacng3 | 1 | 1 | |
| Cacng5 | 1 | 1 | |
| Calcrl | 1 | 1 | |
| Calm3 | 1 | 1 | |
| Caln1 | 1 | 1 | |
| Camk2n1 | 1 | 1 | |
| Camta1 | 1 | 1 | |

| | Number of miR-96 seed region matches in: | | |
|---|---|---|---|
| **Gene** | **C57BL/6N 3'UTR** | **C3H/HeJ 3'UTR** | **Predicted targets of miR-96** |
| Cand2 | 1 | 1 | |
| Capn13 | 1 | 1 | |
| Capn5 | 1 | 1 | |
| Capns1 | 1 | 1 | |
| Casp1 | 1 | 1 | |
| Casp2 | 1 | 1 | |
| Cav1 | 1 | 1 | |
| Cbarp | 1 | 1 | |
| Cbln2 | 1 | 1 | |
| Cbx6 | 1 | 1 | |
| Cc2d1b | 1 | 1 | |
| Ccdc114 | 1 | 1 | |
| Ccdc13 | 1 | 1 | |
| Ccdc190 | 1 | 1 | |
| Ccdc50 | 1 | 1 | |
| Ccdc60 | 1 | 1 | |
| Ccdc64 | 1 | 1 | |
| Ccdc87 | 1 | 1 | |
| Ccdc89 | 1 | 1 | |
| Ccl25 | 1 | 1 | |
| Ccl9 | 1 | 1 | |
| Ccng2 | 1 | 1 | |
| Ccnh | 1 | 1 | |
| Ccnt2 | 1 | 1 | |
| Cd160 | 1 | 1 | |
| Cd164 | 1 | 1 | |
| Cd200 | 1 | 1 | |
| Cd200r3 | 1 | 1 | |
| Cd248 | 1 | 1 | |
| Cd33 | 1 | 1 | |
| Cd47 | 1 | 1 | |
| Cdc27 | 1 | 1 | |
| Cdc37l1 | 1 | 1 | |
| Cdc42bpb | 1 | 1 | |
| Cdc42bpg | 1 | 1 | |
| Cdc42se1 | 1 | 1 | |
| Cdh20 | 1 | 1 | |
| Cdh7 | 1 | 1 | |
| Cdk5r2 | 1 | 1 | |
| Cdk7 | 1 | 1 | |
| Cdkal1 | 1 | 1 | |
| Cdyl2 | 1 | 1 | |
| Cecr6 | 1 | 1 | |

| | Number of miR-96 seed region matches in: | | |
|---|---|---|---|
| **Gene** | **C57BL/6N 3'UTR** | **C3H/HeJ 3'UTR** | **Predicted targets of miR-96** |
| Celf2 | 1 | 1 | |
| Celf5 | 1 | 1 | |
| Celf6 | 1 | 1 | |
| Celsr1 | 1 | 1 | |
| Celsr2 | 1 | 1 | Yes |
| Celsr3 | 1 | 1 | |
| Cenpb | 1 | 1 | |
| Cenpf | 1 | 1 | |
| Cep128 | 1 | 1 | |
| Cep97 | 1 | 1 | |
| Cers5 | 1 | 1 | |
| Chd5 | 1 | 1 | |
| Chic1 | 1 | 1 | |
| Chil5 | 1 | 1 | |
| Chmp2b | 1 | 1 | |
| Chn1 | 1 | 1 | |
| Chrd | 1 | 1 | |
| Chrna1 | 1 | 1 | |
| Chrna6 | 1 | 1 | |
| Chst1 | 1 | 1 | |
| Chst10 | 1 | 1 | |
| Clec11a | 1 | 1 | |
| Clec2e | 1 | 1 | |
| Clec9a | 1 | 1 | |
| Clvs2 | 1 | 1 | |
| Cnih3 | 1 | 1 | |
| Cnn3 | 1 | 1 | |
| Cnnm2 | 1 | 1 | |
| Cnnm3 | 1 | 1 | |
| Cnot6l | 1 | 1 | |
| Cnpy3 | 1 | 1 | |
| Cnr1 | 1 | 1 | |
| Coa4 | 1 | 1 | |
| Cobl | 1 | 1 | |
| Cog1 | 1 | 1 | |
| Cog5 | 1 | 1 | |
| Cog7 | 1 | 1 | |
| Cog8 | 1 | 1 | |
| Col13a1 | 1 | 1 | |
| Col23a1 | 1 | 1 | |
| Col25a1 | 1 | 1 | |
| Col27a1 | 1 | 1 | |
| Col5a1 | 1 | 1 | |

| | Number of miR-96 seed region matches in: | | |
|---|---|---|---|
| **Gene** | **C57BL/6N 3'UTR** | **C3H/HeJ 3'UTR** | **Predicted targets of miR-96** |
| Col9a1 | 1 | 1 | |
| Colec10 | 1 | 1 | |
| Copa | 1 | 1 | |
| Copg1 | 1 | 1 | |
| Coro2b | 1 | 1 | |
| Cox15 | 1 | 1 | |
| Cpd | 1 | 1 | |
| Cpeb4 | 1 | 1 | |
| Cpm | 1 | 1 | |
| Cpn2 | 1 | 1 | |
| Cpsf6 | 1 | 1 | |
| Cpsf7 | 1 | 1 | |
| Cramp1l | 1 | 1 | |
| Crb2 | 1 | 1 | |
| Creb3l2 | 1 | 1 | |
| Crebrf | 1 | 1 | |
| Csad | 1 | 1 | |
| Csf1 | 1 | 1 | |
| Csnk1d | 1 | 1 | |
| Csnk2a2 | 1 | 1 | |
| Cst6 | 1 | 1 | |
| Ctage5 | 1 | 1 | |
| Ctdspl | 1 | 1 | |
| Ctnnb1 | 1 | 1 | |
| Ctsb | 1 | 1 | |
| Cul4a | 1 | 1 | |
| Cxcr2 | 1 | 1 | |
| Cyb561d1 | 1 | 1 | |
| Cyb5r2 | 1 | 1 | |
| Cyb5rl | 1 | 1 | |
| Cyfip1 | 1 | 1 | |
| Cygb | 1 | 1 | |
| Cyp27b1 | 1 | 1 | |
| Cyp2j12 | 1 | 1 | |
| D10Wsu102e | 1 | 1 | |
| D430019H16Rik | 1 | 1 | |
| D5Ertd579e | 1 | 1 | |
| D730001G18Rik | 1 | 1 | |
| Dcaf10 | 1 | 1 | |
| Dcaf5 | 1 | 1 | |
| Dcdc2a | 1 | 1 | |
| Dchs1 | 1 | 1 | |
| Dclre1c | 1 | 1 | |

| | Number of miR-96 seed region matches in: | | |
|---|---|---|---|
| **Gene** | **C57BL/6N 3'UTR** | **C3H/HeJ 3'UTR** | **Predicted targets of miR-96** |
| Dcun1d3 | 1 | 1 | |
| Ddah1 | 1 | 1 | |
| Ddhd1 | 1 | 1 | |
| Ddr2 | 1 | 1 | |
| Ddx25 | 1 | 1 | |
| Decr1 | 1 | 1 | |
| Dennd2c | 1 | 1 | |
| Deptor | 1 | 1 | |
| Derl2 | 1 | 1 | |
| Desi1 | 1 | 1 | |
| Desi2 | 1 | 1 | |
| Dgkb | 1 | 1 | |
| Dgkz | 1 | 1 | |
| Dhcr24 | 1 | 1 | |
| Dirc2 | 1 | 1 | |
| Dlat | 1 | 1 | |
| Dlc1 | 1 | 1 | |
| Dlx2 | 1 | 1 | |
| Dnajb8 | 1 | 1 | |
| Dnajc1 | 1 | 1 | |
| Dnajc19 | 1 | 1 | |
| Dnajc3 | 1 | 1 | |
| Dnd1 | 1 | 1 | |
| Dnpep | 1 | 1 | |
| Dock1 | 1 | 1 | |
| Dock5 | 1 | 1 | |
| Dot1l | 1 | 1 | |
| Dph2 | 1 | 1 | |
| Dpp8 | 1 | 1 | |
| Dpy19l3 | 1 | 1 | |
| Dram1 | 1 | 1 | |
| Dsc1 | 1 | 1 | |
| Dsc2 | 1 | 1 | |
| Dsc3 | 1 | 1 | |
| Dsg2 | 1 | 1 | |
| Dsn1 | 1 | 1 | |
| Dtnb | 1 | 1 | |
| Dtx3l | 1 | 1 | |
| Duoxa1 | 1 | 1 | |
| Dusp12 | 1 | 1 | |
| Dusp13 | 1 | 1 | |
| Dynll1 | 1 | 1 | |
| E2f5 | 1 | 1 | |

| Gene | Number of miR-96 seed region matches in: C57BL/6N 3'UTR | C3H/HeJ 3'UTR | Predicted targets of miR-96 |
|---|---|---|---|
| Eaf1 | 1 | 1 | |
| Ebf3 | 1 | 1 | |
| Echdc1 | 1 | 1 | |
| Edem1 | 1 | 1 | |
| Efna1 | 1 | 1 | |
| Egfem1 | 1 | 1 | |
| Egfr | 1 | 1 | |
| Ehd1 | 1 | 1 | |
| Ehmt1 | 1 | 1 | |
| Eif4e2 | 1 | 1 | |
| Eif5 | 1 | 1 | |
| Eif5b | 1 | 1 | |
| Elac2 | 1 | 1 | |
| Elavl1 | 1 | 1 | |
| Elovl2 | 1 | 1 | |
| Emc7 | 1 | 1 | |
| En2 | 1 | 1 | |
| Epb41l3 | 1 | 1 | |
| Epha3 | 1 | 1 | |
| Epha5 | 1 | 1 | |
| Erc2 | 1 | 1 | |
| Ercc6l | 1 | 1 | |
| Ereg | 1 | 1 | |
| Ergic1 | 1 | 1 | |
| Ergic2 | 1 | 1 | |
| Erlin1 | 1 | 1 | |
| Erp29 | 1 | 1 | |
| Esp31 | 1 | 1 | |
| Esr1 | 1 | 1 | |
| Esyt2 | 1 | 1 | |
| Etf1 | 1 | 1 | |
| Etv3 | 1 | 1 | |
| Evx2 | 1 | 1 | |
| Exoc6b | 1 | 1 | |
| Eya3 | 1 | 1 | |
| Ezr | 1 | 1 | |
| F13a1 | 1 | 1 | |
| F8a | 1 | 1 | |
| Fadd | 1 | 1 | |
| Fads1 | 1 | 1 | |
| Faf1 | 1 | 1 | |
| Faf2 | 1 | 1 | |
| Fahd1 | 1 | 1 | |

| | Number of miR-96 seed region matches in: | | |
|---|---|---|---|
| **Gene** | **C57BL/6N 3'UTR** | **C3H/HeJ 3'UTR** | **Predicted targets of miR-96** |
| Fam107a | 1 | 1 | |
| Fam126b | 1 | 1 | |
| Fam129a | 1 | 1 | |
| Fam131a | 1 | 1 | |
| Fam132b | 1 | 1 | |
| Fam135a | 1 | 1 | |
| Fam160a2 | 1 | 1 | |
| Fam167a | 1 | 1 | |
| Fam175a | 1 | 1 | |
| Fam175b | 1 | 1 | |
| Fam178a | 1 | 1 | |
| Fam189a1 | 1 | 1 | |
| Fam195a | 1 | 1 | |
| Fam196a | 1 | 1 | |
| Fam199x | 1 | 1 | |
| Fam19a3 | 1 | 1 | |
| Fam234b | 1 | 1 | |
| Fam49b | 1 | 1 | |
| Fam53c | 1 | 1 | |
| Fam63a | 1 | 1 | |
| Fam65b | 1 | 1 | |
| Fam81a | 1 | 1 | |
| Farp1 | 1 | 1 | |
| Fbf1 | 1 | 1 | |
| Fbln5 | 1 | 1 | |
| Fbxl20 | 1 | 1 | |
| Fbxo22 | 1 | 1 | |
| Fbxo41 | 1 | 1 | |
| Fbxo7 | 1 | 1 | |
| Fbxw11 | 1 | 1 | |
| Fchsd1 | 1 | 1 | |
| Fem1b | 1 | 1 | |
| Fgf13 | 1 | 1 | |
| Fgf7 | 1 | 1 | |
| Fhdc1 | 1 | 1 | |
| Fhl1 | 1 | 1 | |
| Fign | 1 | 1 | |
| Filip1 | 1 | 1 | |
| Fkbp9 | 1 | 1 | |
| Flna | 1 | 1 | |
| Flot2 | 1 | 1 | |
| Fndc5 | 1 | 1 | |
| Fndc8 | 1 | 1 | |

| | Number of miR-96 seed region matches in: | | |
|---|---|---|---|
| **Gene** | **C57BL/6N 3'UTR** | **C3H/HeJ 3'UTR** | **Predicted targets of miR-96** |
| Fnip1 | 1 | 1 | |
| Foxf2 | 1 | 1 | |
| Foxk1 | 1 | 1 | |
| Foxl2 | 1 | 1 | |
| Foxn2 | 1 | 1 | |
| Foxn4 | 1 | 1 | |
| Foxo1 | 1 | 1 | Yes |
| Foxo4 | 1 | 1 | |
| Foxp2 | 1 | 1 | |
| Foxp3 | 1 | 1 | |
| Foxq1 | 1 | 1 | |
| Foxred2 | 1 | 1 | |
| Frem1 | 1 | 1 | |
| Frmd5 | 1 | 1 | |
| Frmpd4 | 1 | 1 | |
| Frs2 | 1 | 1 | |
| Fstl4 | 1 | 1 | |
| Fundc2 | 1 | 1 | |
| Furin | 1 | 1 | |
| Fut4 | 1 | 1 | |
| Fxyd5 | 1 | 1 | |
| Fyn | 1 | 1 | |
| G6pc2 | 1 | 1 | |
| Gabpb2 | 1 | 1 | |
| Gabra4 | 1 | 1 | |
| Gabrq | 1 | 1 | |
| Galk2 | 1 | 1 | |
| Galnt16 | 1 | 1 | |
| Galnt7 | 1 | 1 | |
| Galr1 | 1 | 1 | |
| Gap43 | 1 | 1 | |
| Gapdhs | 1 | 1 | |
| Gapvd1 | 1 | 1 | |
| Gas7 | 1 | 1 | |
| Gbas | 1 | 1 | |
| Gbf1 | 1 | 1 | |
| Gbp2b | 1 | 1 | |
| Gbp4 | 1 | 1 | |
| Gca | 1 | 1 | |
| Gcfc2 | 1 | 1 | |
| Gdnf | 1 | 1 | |
| Gfap | 1 | 1 | |
| Gfm2 | 1 | 1 | |

| | Number of miR-96 seed region matches in: | | |
|---|---|---|---|
| Gene | C57BL/6N 3'UTR | C3H/HeJ 3'UTR | Predicted targets of miR-96 |
| Gga2 | 1 | 1 | |
| Gga3 | 1 | 1 | |
| Ggta1 | 1 | 1 | |
| Gid4 | 1 | 1 | |
| Gif | 1 | 1 | |
| Gigyf2 | 1 | 1 | |
| Gja5 | 1 | 1 | |
| Gja8 | 1 | 1 | |
| Glis3 | 1 | 1 | |
| Glrb | 1 | 1 | |
| Glrx | 1 | 1 | |
| Gltscr1l | 1 | 1 | |
| Glul | 1 | 1 | |
| Gm11541 | 1 | 1 | |
| Gm20708 | 1 | 1 | |
| Gm21685 | 1 | 1 | |
| Gm21987 | 1 | 1 | |
| Gm27235 | 1 | 1 | |
| Gm2a | 1 | 1 | |
| Gm35549 | 1 | 1 | |
| Gm45062 | 1 | 1 | |
| Gm572 | 1 | 1 | |
| Gm6583 | 1 | 1 | |
| Gm6588 | 1 | 1 | |
| Gm973 | 1 | 1 | |
| Gm9903 | 1 | 1 | |
| Gm9956 | 1 | 1 | |
| Gmeb2 | 1 | 1 | |
| Gnai3 | 1 | 1 | |
| Gnao1 | 1 | 1 | |
| Gnas | 1 | 1 | |
| Gnb2 | 1 | 1 | |
| Gnb4 | 1 | 1 | |
| Gng4 | 1 | 1 | |
| Gng7 | 1 | 1 | |
| Gnl3l | 1 | 1 | |
| Gnptab | 1 | 1 | |
| Golm1 | 1 | 1 | |
| Gp1bb | 1 | 1 | |
| Gpc1 | 1 | 1 | Yes |
| Gpc3 | 1 | 1 | Yes |
| Gpihbp1 | 1 | 1 | |
| Gpm6a | 1 | 1 | |

| | Number of miR-96 seed region matches in: | | |
|---|---|---|---|
| **Gene** | **C57BL/6N 3'UTR** | **C3H/HeJ 3'UTR** | **Predicted targets of miR-96** |
| Gpr107 | 1 | 1 | |
| Gpr137 | 1 | 1 | |
| Gpr146 | 1 | 1 | |
| Gpr155 | 1 | 1 | |
| Gpr161 | 1 | 1 | |
| Gpr17 | 1 | 1 | |
| Gpr179 | 1 | 1 | |
| Gpr20 | 1 | 1 | |
| Gpr21 | 1 | 1 | |
| Gpr26 | 1 | 1 | |
| Gpr39 | 1 | 1 | |
| Gprc5b | 1 | 1 | |
| Gramd2 | 1 | 1 | |
| Gramd4 | 1 | 1 | |
| Grhl2 | 1 | 1 | |
| Gria1 | 1 | 1 | |
| Gria2 | 1 | 1 | |
| Grid1 | 1 | 1 | |
| Grid2 | 1 | 1 | |
| Grin2b | 1 | 1 | |
| Grm1 | 1 | 1 | |
| Gtf2a1 | 1 | 1 | |
| Gtf2h2 | 1 | 1 | |
| Gtf3c4 | 1 | 1 | |
| Gtpbp2 | 1 | 1 | |
| Gulp1 | 1 | 1 | |
| Gusb | 1 | 1 | |
| Gzf1 | 1 | 1 | |
| H13 | 1 | 1 | |
| H1foo | 1 | 1 | |
| H6pd | 1 | 1 | |
| Hacd2 | 1 | 1 | |
| Hap1 | 1 | 1 | |
| Hbegf | 1 | 1 | |
| Hbp1 | 1 | 1 | Yes |
| Hdac5 | 1 | 1 | |
| Hdac7 | 1 | 1 | |
| Hdgfrp3 | 1 | 1 | |
| Hdhd2 | 1 | 1 | |
| Heatr5a | 1 | 1 | |
| Helz | 1 | 1 | |
| Herpud1 | 1 | 1 | |
| Hfe2 | 1 | 1 | |

| | Number of miR-96 seed region matches in: | | |
|---|---|---|---|
| **Gene** | **C57BL/6N 3'UTR** | **C3H/HeJ 3'UTR** | **Predicted targets of miR-96** |
| Hhip | 1 | 1 | |
| Hif1an | 1 | 1 | |
| Hip1 | 1 | 1 | |
| Hipk1 | 1 | 1 | |
| Hmgcll1 | 1 | 1 | |
| Hmgcs1 | 1 | 1 | |
| Hmmr | 1 | 1 | |
| Homer1 | 1 | 1 | |
| Hook3 | 1 | 1 | |
| Hoxa5 | 1 | 1 | |
| Hpgds | 1 | 1 | |
| Hps1 | 1 | 1 | |
| Hras | 1 | 1 | |
| Hrh4 | 1 | 1 | |
| Hs6st1 | 1 | 1 | |
| Hsd17b12 | 1 | 1 | |
| Hsd3b7 | 1 | 1 | |
| Hsf5 | 1 | 1 | |
| Hspa13 | 1 | 1 | |
| Hspa2 | 1 | 1 | Yes |
| Htr1b | 1 | 1 | |
| Htr2c | 1 | 1 | |
| Hus1 | 1 | 1 | |
| Hyal1 | 1 | 1 | |
| Hyal2 | 1 | 1 | |
| Hyou1 | 1 | 1 | |
| Ids | 1 | 1 | |
| Idua | 1 | 1 | |
| Ifi35 | 1 | 1 | |
| Ifit1 | 1 | 1 | |
| Ifit1bl1 | 1 | 1 | |
| Ifit1bl2 | 1 | 1 | |
| Ifrd1 | 1 | 1 | |
| Ifrd2 | 1 | 1 | |
| Ift81 | 1 | 1 | |
| Igf2bp1 | 1 | 1 | |
| Igsf11 | 1 | 1 | |
| Ikzf4 | 1 | 1 | |
| Il10ra | 1 | 1 | |
| Il12a | 1 | 1 | |
| Il18rap | 1 | 1 | |
| Il1rap | 1 | 1 | |
| Il22ra1 | 1 | 1 | |

| | Number of miR-96 seed region matches in: | | |
|---|---|---|---|
| Gene | C57BL/6N 3'UTR | C3H/HeJ 3'UTR | Predicted targets of miR-96 |
| Ildr2 | 1 | 1 | |
| Imp4 | 1 | 1 | |
| Impact | 1 | 1 | |
| Impad1 | 1 | 1 | |
| Ina | 1 | 1 | |
| Inadl | 1 | 1 | |
| Inpp1 | 1 | 1 | |
| Inpp4b | 1 | 1 | |
| Inpp5a | 1 | 1 | |
| Ints6 | 1 | 1 | |
| Intu | 1 | 1 | |
| Invs | 1 | 1 | |
| Ipo11 | 1 | 1 | |
| Ipo9 | 1 | 1 | |
| Iqsec1 | 1 | 1 | |
| Iqsec3 | 1 | 1 | |
| Irf4 | 1 | 1 | |
| Irf6 | 1 | 1 | |
| Irs1 | 1 | 1 | Yes |
| Itga11 | 1 | 1 | |
| Itga3 | 1 | 1 | |
| Itga6 | 1 | 1 | |
| Itgb1 | 1 | 1 | |
| Itpr2 | 1 | 1 | |
| Itsn1 | 1 | 1 | |
| Jazf1 | 1 | 1 | |
| Jdp2 | 1 | 1 | |
| Jmjd1c | 1 | 1 | |
| Jmjd6 | 1 | 1 | |
| Kat5 | 1 | 1 | |
| Kcnc3 | 1 | 1 | |
| Kcnj6 | 1 | 1 | |
| Kcnk6 | 1 | 1 | |
| Kcnq5 | 1 | 1 | |
| Kctd12 | 1 | 1 | |
| Kdelr2 | 1 | 1 | |
| Kdelr3 | 1 | 1 | |
| Kidins220 | 1 | 1 | |
| Kif1b | 1 | 1 | |
| Kif24 | 1 | 1 | |
| Kif5a | 1 | 1 | |
| Kl | 1 | 1 | |
| Klf3 | 1 | 1 | |

| | Number of miR-96 seed region matches in: | | |
|---|---|---|---|
| **Gene** | **C57BL/6N 3'UTR** | **C3H/HeJ 3'UTR** | **Predicted targets of miR-96** |
| Klhl32 | 1 | 1 | |
| Klhl41 | 1 | 1 | |
| Klhl7 | 1 | 1 | |
| Klhl8 | 1 | 1 | |
| Klrg2 | 1 | 1 | |
| Kmt2a | 1 | 1 | |
| Knstrn | 1 | 1 | |
| Kpnb1 | 1 | 1 | |
| Kras | 1 | 1 | |
| L2hgdh | 1 | 1 | |
| L3hypdh | 1 | 1 | |
| Lactb2 | 1 | 1 | |
| Lamc1 | 1 | 1 | |
| Lamp2 | 1 | 1 | |
| Lcor | 1 | 1 | |
| Lcorl | 1 | 1 | |
| Lcp1 | 1 | 1 | |
| Ldb1 | 1 | 1 | |
| Ldb2 | 1 | 1 | |
| Ldlrad4 | 1 | 1 | |
| Lekr1 | 1 | 1 | |
| Letm1 | 1 | 1 | |
| Lhx4 | 1 | 1 | |
| Lhx6 | 1 | 1 | |
| Lhx9 | 1 | 1 | |
| Lingo2 | 1 | 1 | |
| Lingo4 | 1 | 1 | |
| Lmo3 | 1 | 1 | |
| Lmtk3 | 1 | 1 | |
| Lnx2 | 1 | 1 | |
| Loxl3 | 1 | 1 | |
| Lpp | 1 | 1 | |
| Lrat | 1 | 1 | |
| Lrfn5 | 1 | 1 | |
| Lrig1 | 1 | 1 | |
| Lrp11 | 1 | 1 | |
| Lrp2 | 1 | 1 | |
| Lrp4 | 1 | 1 | |
| Lrrc15 | 1 | 1 | |
| Lrrc17 | 1 | 1 | |
| Lrrc28 | 1 | 1 | |
| Lrrc30 | 1 | 1 | |
| Lrrc74a | 1 | 1 | |

| | Number of miR-96 seed region matches in: | | |
|---|---|---|---|
| **Gene** | **C57BL/6N 3'UTR** | **C3H/HeJ 3'UTR** | **Predicted targets of miR-96** |
| Lsg1 | 1 | 1 | |
| Lsm11 | 1 | 1 | |
| Luzp1 | 1 | 1 | |
| Lyplal1 | 1 | 1 | |
| Lyrm9 | 1 | 1 | |
| Lzts1 | 1 | 1 | |
| Mab21l2 | 1 | 1 | |
| Mafk | 1 | 1 | |
| Magi1 | 1 | 1 | |
| Magi3 | 1 | 1 | |
| Mamld1 | 1 | 1 | |
| Maoa | 1 | 1 | |
| Map2k1 | 1 | 1 | |
| Map2k3 | 1 | 1 | |
| Map3k2 | 1 | 1 | |
| Map3k7 | 1 | 1 | |
| Map4k2 | 1 | 1 | |
| Mapk10 | 1 | 1 | |
| Mapk11 | 1 | 1 | |
| Mapk8ip2 | 1 | 1 | |
| Mapk9 | 1 | 1 | |
| Mapre2 | 1 | 1 | |
| Mars2 | 1 | 1 | |
| Mastl | 1 | 1 | |
| Mbnl1 | 1 | 1 | |
| Mcc | 1 | 1 | |
| Mcf2 | 1 | 1 | |
| Mcmbp | 1 | 1 | |
| Mcmdc2 | 1 | 1 | |
| Mcu | 1 | 1 | |
| Mecom | 1 | 1 | |
| Mecp2 | 1 | 1 | |
| Med17 | 1 | 1 | |
| Mef2a | 1 | 1 | |
| Mefv | 1 | 1 | |
| Megf9 | 1 | 1 | |
| Meox2 | 1 | 1 | |
| Metap2 | 1 | 1 | |
| Mettl1 | 1 | 1 | |
| Mettl21e | 1 | 1 | |
| Mettl25 | 1 | 1 | |
| Mfap3l | 1 | 1 | |
| Mfhas1 | 1 | 1 | |

| | Number of miR-96 seed region matches in: | | |
|---|---|---|---|
| Gene | C57BL/6N 3'UTR | C3H/HeJ 3'UTR | Predicted targets of miR-96 |
| Mfng | 1 | 1 | |
| Mfrp | 1 | 1 | |
| Mfsd4a | 1 | 1 | |
| Mfsd5 | 1 | 1 | |
| Mgat5 | 1 | 1 | |
| Mib1 | 1 | 1 | |
| Mical3 | 1 | 1 | |
| Mid2 | 1 | 1 | |
| Mief2 | 1 | 1 | |
| Mier3 | 1 | 1 | |
| Mitf | 1 | 1 | Yes |
| Mkrn1 | 1 | 1 | |
| Mlec | 1 | 1 | |
| Mmp17 | 1 | 1 | |
| Mmp2 | 1 | 1 | |
| Mmrn2 | 1 | 1 | |
| Mms19 | 1 | 1 | |
| Mms22l | 1 | 1 | |
| Mob1b | 1 | 1 | |
| Mobp | 1 | 1 | |
| Mon2 | 1 | 1 | |
| Mpp1 | 1 | 1 | |
| Mpzl3 | 1 | 1 | |
| Mras | 1 | 1 | |
| Mrgpra6 | 1 | 1 | |
| Mrgprf | 1 | 1 | |
| Mroh5 | 1 | 1 | |
| Mrpl10 | 1 | 1 | |
| Mrpl55 | 1 | 1 | |
| Mrps11 | 1 | 1 | |
| Mrps17 | 1 | 1 | |
| Ms4a2 | 1 | 1 | |
| Msh5 | 1 | 1 | |
| Msl3 | 1 | 1 | |
| Msx2 | 1 | 1 | |
| Mta3 | 1 | 1 | |
| Mtf1 | 1 | 1 | |
| Mtif3 | 1 | 1 | |
| Mtmr10 | 1 | 1 | |
| Mtmr12 | 1 | 1 | |
| Mtmr3 | 1 | 1 | |
| Mtr | 1 | 1 | |
| Mtss1 | 1 | 1 | |

| | Number of miR-96 seed region matches in: | | |
|---|---|---|---|
| **Gene** | **C57BL/6N 3'UTR** | **C3H/HeJ 3'UTR** | **Predicted targets of miR-96** |
| Mtx3 | 1 | 1 | |
| Mvk | 1 | 1 | |
| Mxd1 | 1 | 1 | |
| Mxra7 | 1 | 1 | |
| Myh9 | 1 | 1 | |
| Myo16 | 1 | 1 | |
| Myo18b | 1 | 1 | |
| Myo19 | 1 | 1 | |
| Myo5a | 1 | 1 | |
| Myo5b | 1 | 1 | |
| Myrip | 1 | 1 | |
| N4bp1 | 1 | 1 | |
| Naa25 | 1 | 1 | |
| Naa50 | 1 | 1 | |
| Nabp1 | 1 | 1 | |
| Nalcn | 1 | 1 | |
| Nanos1 | 1 | 1 | |
| Napepld | 1 | 1 | |
| Nat8f2 | 1 | 1 | |
| Nat8f4 | 1 | 1 | |
| Nat8l | 1 | 1 | |
| Ncapd2 | 1 | 1 | |
| Nckap1l | 1 | 1 | |
| Ndst1 | 1 | 1 | |
| Nedd4l | 1 | 1 | |
| Nek6 | 1 | 1 | |
| Nek9 | 1 | 1 | |
| Nepro | 1 | 1 | |
| Neu4 | 1 | 1 | |
| Neurod4 | 1 | 1 | |
| Nfat5 | 1 | 1 | |
| Nfkb1 | 1 | 1 | |
| Nhlh2 | 1 | 1 | |
| Nhsl2 | 1 | 1 | |
| Nim1k | 1 | 1 | |
| Nit1 | 1 | 1 | |
| Nkain2 | 1 | 1 | |
| Nkap | 1 | 1 | |
| Nnt | 1 | 1 | |
| Nol9 | 1 | 1 | |
| Npnt | 1 | 1 | |
| Nptn | 1 | 1 | |
| Nptx1 | 1 | 1 | |

| | Number of miR-96 seed region matches in: | | |
|---|---|---|---|
| **Gene** | **C57BL/6N 3'UTR** | **C3H/HeJ 3'UTR** | **Predicted targets of miR-96** |
| Nptx2 | 1 | 1 | |
| Nr2e3 | 1 | 1 | |
| Nrarp | 1 | 1 | |
| Nrcam | 1 | 1 | |
| Nrros | 1 | 1 | |
| Nrsn2 | 1 | 1 | |
| Nrxn1 | 1 | 1 | |
| Nsg1 | 1 | 1 | |
| Nsmce4a | 1 | 1 | |
| Nts | 1 | 1 | |
| Nudcd3 | 1 | 1 | |
| Nudt7 | 1 | 1 | |
| Nudt8 | 1 | 1 | |
| Nup50 | 1 | 1 | |
| Nus1 | 1 | 1 | |
| Nwd1 | 1 | 1 | |
| Nwd2 | 1 | 1 | |
| Nyap2 | 1 | 1 | |
| Nynrin | 1 | 1 | |
| Ociad1 | 1 | 1 | |
| Odf2 | 1 | 1 | Yes |
| Ogdh | 1 | 1 | |
| Ogfr | 1 | 1 | |
| Ogg1 | 1 | 1 | |
| Ogt | 1 | 1 | |
| Olfr613 | 1 | 1 | |
| Olfr70 | 1 | 1 | |
| Olfr78 | 1 | 1 | |
| Olig3 | 1 | 1 | |
| Onecut3 | 1 | 1 | |
| Oog4 | 1 | 1 | |
| Opcml | 1 | 1 | |
| Opn1sw | 1 | 1 | |
| Oprm1 | 1 | 1 | |
| Osbpl2 | 1 | 1 | |
| Ostm1 | 1 | 1 | |
| Otog | 1 | 1 | |
| Otop3 | 1 | 1 | |
| Otud7a | 1 | 1 | |
| Ovol1 | 1 | 1 | |
| Oxsr1 | 1 | 1 | |
| P2rx4 | 1 | 1 | |
| P2ry14 | 1 | 1 | |

| | Number of miR-96 seed region matches in: | | |
|---|---|---|---|
| **Gene** | **C57BL/6N 3'UTR** | **C3H/HeJ 3'UTR** | **Predicted targets of miR-96** |
| P2ry2 | 1 | 1 | |
| P4ha1 | 1 | 1 | |
| Pabpc4 | 1 | 1 | |
| Pabpc6 | 1 | 1 | |
| Pacsin1 | 1 | 1 | |
| Pak1 | 1 | 1 | |
| Palb2 | 1 | 1 | |
| Panx2 | 1 | 1 | |
| Pappa | 1 | 1 | |
| Pappa2 | 1 | 1 | |
| Papss1 | 1 | 1 | |
| Paqr8 | 1 | 1 | |
| Pard3b | 1 | 1 | |
| Parm1 | 1 | 1 | |
| Parva | 1 | 1 | |
| Patz1 | 1 | 1 | |
| Pax7 | 1 | 1 | |
| Paxip1 | 1 | 1 | |
| Pbx1 | 1 | 1 | |
| Pcca | 1 | 1 | |
| Pcdh11x | 1 | 1 | |
| Pcdh15 | 1 | 1 | |
| Pcdh17 | 1 | 1 | |
| Pcolce2 | 1 | 1 | |
| Pde10a | 1 | 1 | |
| Pde11a | 1 | 1 | |
| Pde1a | 1 | 1 | |
| Pde1c | 1 | 1 | |
| Pde3a | 1 | 1 | |
| Pde7a | 1 | 1 | |
| Pdgfd | 1 | 1 | |
| Pdik1l | 1 | 1 | |
| Pdk2 | 1 | 1 | |
| Pdp2 | 1 | 1 | |
| Pdxk | 1 | 1 | |
| Pea15a | 1 | 1 | |
| Pear1 | 1 | 1 | |
| Peg3 | 1 | 1 | |
| Per2 | 1 | 1 | |
| Pet100 | 1 | 1 | |
| Pex11b | 1 | 1 | |
| Pggt1b | 1 | 1 | |
| Pgs1 | 1 | 1 | |

| | Number of miR-96 seed region matches in: | | |
|---|---|---|---|
| **Gene** | **C57BL/6N 3'UTR** | **C3H/HeJ 3'UTR** | **Predicted targets of miR-96** |
| Phactr4 | 1 | 1 | |
| Phc1 | 1 | 1 | |
| Phc3 | 1 | 1 | |
| Phf10 | 1 | 1 | |
| Phf7 | 1 | 1 | |
| Phip | 1 | 1 | |
| Phkb | 1 | 1 | |
| Pi4k2b | 1 | 1 | |
| Pick1 | 1 | 1 | |
| Piga | 1 | 1 | |
| Pik3c2a | 1 | 1 | |
| Pip4k2b | 1 | 1 | |
| Pkn2 | 1 | 1 | |
| Pkp4 | 1 | 1 | |
| Pla1a | 1 | 1 | |
| Plagl1 | 1 | 1 | |
| Plaur | 1 | 1 | |
| Plcb1 | 1 | 1 | |
| Plcb4 | 1 | 1 | |
| Plch1 | 1 | 1 | |
| Plch2 | 1 | 1 | |
| Plcxd3 | 1 | 1 | |
| Plekha6 | 1 | 1 | |
| Plekhf1 | 1 | 1 | |
| Plekhm1 | 1 | 1 | |
| Plod2 | 1 | 1 | |
| Plscr3 | 1 | 1 | |
| Plxdc1 | 1 | 1 | |
| Pmpca | 1 | 1 | |
| Pnkp | 1 | 1 | |
| Pnpla2 | 1 | 1 | |
| Pnpla8 | 1 | 1 | Yes |
| Podxl | 1 | 1 | |
| Pogk | 1 | 1 | |
| Poldip3 | 1 | 1 | |
| Polg | 1 | 1 | |
| Polr3f | 1 | 1 | |
| Polrmt | 1 | 1 | |
| Porcn | 1 | 1 | |
| Pou2f3 | 1 | 1 | |
| Pou3f2 | 1 | 1 | |
| Pou6f2 | 1 | 1 | |
| Ppbp | 1 | 1 | |

| | Number of miR-96 seed region matches in: | | |
|---|---|---|---|
| **Gene** | **C57BL/6N 3'UTR** | **C3H/HeJ 3'UTR** | **Predicted targets of miR-96** |
| Ppfibp2 | 1 | 1 | |
| Ppip5k1 | 1 | 1 | |
| Ppm1k | 1 | 1 | |
| Ppm1l | 1 | 1 | |
| Ppme1 | 1 | 1 | |
| Ppp1r12c | 1 | 1 | |
| Ppp1r7 | 1 | 1 | |
| Ppp1r9a | 1 | 1 | |
| Ppp1r9b | 1 | 1 | |
| Ppp2ca | 1 | 1 | |
| Ppp3r1 | 1 | 1 | |
| Ppp6c | 1 | 1 | |
| Prdm11 | 1 | 1 | |
| Prdm16 | 1 | 1 | |
| Prkag3 | 1 | 1 | |
| Prkar1a | 1 | 1 | |
| Prkca | 1 | 1 | |
| Prkce | 1 | 1 | |
| Prkrir | 1 | 1 | |
| Prlr | 1 | 1 | |
| Prom2 | 1 | 1 | |
| Prosc | 1 | 1 | |
| Prrg1 | 1 | 1 | |
| Prrg3 | 1 | 1 | |
| Prrt1 | 1 | 1 | |
| Prrt3 | 1 | 1 | |
| Prrt4 | 1 | 1 | |
| Prss33 | 1 | 1 | |
| Prtg | 1 | 1 | |
| Psd2 | 1 | 1 | |
| Pskh1 | 1 | 1 | |
| Psmb11 | 1 | 1 | |
| Psmd13 | 1 | 1 | |
| Ptger3 | 1 | 1 | |
| Ptges | 1 | 1 | |
| Ptgfr | 1 | 1 | |
| Ptgfrn | 1 | 1 | |
| Ptpn20 | 1 | 1 | |
| Ptpn9 | 1 | 1 | |
| Ptpra | 1 | 1 | |
| Pum2 | 1 | 1 | |
| Pus3 | 1 | 1 | |
| Pwp2 | 1 | 1 | |

| | Number of miR-96 seed region matches in: | | |
|---|---|---|---|
| Gene | C57BL/6N 3'UTR | C3H/HeJ 3'UTR | Predicted targets of miR-96 |
| Pycard | 1 | 1 | |
| Qk | 1 | 1 | |
| Qpctl | 1 | 1 | |
| R3hdm2 | 1 | 1 | |
| Rab11fip4 | 1 | 1 | |
| Rab23 | 1 | 1 | |
| Rab26 | 1 | 1 | |
| Rab27a | 1 | 1 | |
| Rab43 | 1 | 1 | |
| Rab8b | 1 | 1 | |
| Rad51b | 1 | 1 | |
| Ralgps1 | 1 | 1 | |
| Ranbp6 | 1 | 1 | |
| Rap1gap2 | 1 | 1 | |
| Rap2b | 1 | 1 | |
| Rapgef4 | 1 | 1 | |
| Rapgef5 | 1 | 1 | |
| Raph1 | 1 | 1 | |
| Rasa2 | 1 | 1 | |
| Rassf10 | 1 | 1 | |
| Rassf8 | 1 | 1 | |
| Rbbp6 | 1 | 1 | |
| Rbm15b | 1 | 1 | |
| Rbm20 | 1 | 1 | |
| Rbm25 | 1 | 1 | |
| Rbm26 | 1 | 1 | |
| Rbm38 | 1 | 1 | |
| Rbm48 | 1 | 1 | |
| Rbm4b | 1 | 1 | |
| Rcbtb2 | 1 | 1 | |
| Rdh10 | 1 | 1 | |
| Rdh19 | 1 | 1 | |
| Recql5 | 1 | 1 | |
| Reep2 | 1 | 1 | |
| Rev1 | 1 | 1 | Yes |
| Rev3l | 1 | 1 | |
| Rexo1 | 1 | 1 | |
| Rgs2 | 1 | 1 | |
| Rgs8 | 1 | 1 | |
| Rgsl1 | 1 | 1 | |
| Rhbdf1 | 1 | 1 | |
| Rhob | 1 | 1 | |
| Rhpn2 | 1 | 1 | |

| | Number of miR-96 seed region matches in: | | |
|---|---|---|---|
| Gene | C57BL/6N 3'UTR | C3H/HeJ 3'UTR | Predicted targets of miR-96 |
| Ric8b | 1 | 1 | |
| Rictor | 1 | 1 | |
| Rimklb | 1 | 1 | |
| Rims4 | 1 | 1 | |
| Rita1 | 1 | 1 | |
| Rnf103 | 1 | 1 | |
| Rnf112 | 1 | 1 | |
| Rnf114 | 1 | 1 | |
| Rnf139 | 1 | 1 | |
| Rnf152 | 1 | 1 | |
| Rnf169 | 1 | 1 | |
| Rnf183 | 1 | 1 | |
| Rnf207 | 1 | 1 | |
| Rnf217 | 1 | 1 | |
| Rnf225 | 1 | 1 | |
| Rnft2 | 1 | 1 | |
| Rp9 | 1 | 1 | |
| Rpa1 | 1 | 1 | |
| Rpe | 1 | 1 | |
| Rpl15 | 1 | 1 | |
| Rps6ka6 | 1 | 1 | |
| Rps6kb2 | 1 | 1 | |
| Rrnad1 | 1 | 1 | |
| Rrp7a | 1 | 1 | |
| Rtl1 | 1 | 1 | |
| Rufy1 | 1 | 1 | |
| Rufy2 | 1 | 1 | |
| Rundc3b | 1 | 1 | |
| Runx1 | 1 | 1 | |
| Rwdd2b | 1 | 1 | |
| Rxfp3 | 1 | 1 | |
| Rxra | 1 | 1 | |
| Ryk | 1 | 1 | |
| Samd10 | 1 | 1 | |
| Samd12 | 1 | 1 | |
| Sap18 | 1 | 1 | |
| Scaf11 | 1 | 1 | |
| Scamp3 | 1 | 1 | |
| Scarb1 | 1 | 1 | Yes |
| Scd1 | 1 | 1 | |
| Scyl3 | 1 | 1 | |
| Sdad1 | 1 | 1 | |
| Sdc2 | 1 | 1 | Yes |

| | Number of miR-96 seed region matches in: | | |
|---|---|---|---|
| Gene | C57BL/6N 3'UTR | C3H/HeJ 3'UTR | Predicted targets of miR-96 |
| Sec14l1 | 1 | 1 | |
| Sec62 | 1 | 1 | |
| Secisbp2l | 1 | 1 | |
| Sema5a | 1 | 1 | |
| Sema6a | 1 | 1 | |
| Sema6b | 1 | 1 | |
| Sept11 | 1 | 1 | |
| Serinc3 | 1 | 1 | |
| Serinc5 | 1 | 1 | |
| Serpina12 | 1 | 1 | |
| Serpinb2 | 1 | 1 | |
| Serpinb6b | 1 | 1 | |
| Serpinb8 | 1 | 1 | |
| Sesn1 | 1 | 1 | |
| Setd1b | 1 | 1 | |
| Sez6 | 1 | 1 | |
| Sfrp1 | 1 | 1 | |
| Sfxn5 | 1 | 1 | |
| Sgk3 | 1 | 1 | |
| Sh2b3 | 1 | 1 | |
| Sh3bp5 | 1 | 1 | |
| Sh3kbp1 | 1 | 1 | |
| Sh3pxd2a | 1 | 1 | |
| Shisa3 | 1 | 1 | |
| Shox2 | 1 | 1 | |
| Shq1 | 1 | 1 | |
| Shroom2 | 1 | 1 | |
| Sidt2 | 1 | 1 | |
| Siglech | 1 | 1 | |
| Sik2 | 1 | 1 | |
| Sike1 | 1 | 1 | |
| Sim1 | 1 | 1 | |
| Sirt7 | 1 | 1 | |
| Six1 | 1 | 1 | |
| Skap1 | 1 | 1 | |
| Slain2 | 1 | 1 | |
| Slc10a3 | 1 | 1 | |
| Slc12a5 | 1 | 1 | |
| Slc16a13 | 1 | 1 | |
| Slc16a9 | 1 | 1 | |
| Slc18a3 | 1 | 1 | |
| Slc1a2 | 1 | 1 | |
| Slc22a15 | 1 | 1 | |

| | Number of miR-96 seed region matches in: | | |
|---|---|---|---|
| Gene | C57BL/6N 3'UTR | C3H/HeJ 3'UTR | Predicted targets of miR-96 |
| Slc22a23 | 1 | 1 | |
| Slc25a1 | 1 | 1 | |
| Slc25a12 | 1 | 1 | |
| Slc25a25 | 1 | 1 | |
| Slc25a42 | 1 | 1 | |
| Slc25a44 | 1 | 1 | |
| Slc25a46 | 1 | 1 | |
| Slc2a13 | 1 | 1 | |
| Slc30a10 | 1 | 1 | |
| Slc30a3 | 1 | 1 | |
| Slc31a1 | 1 | 1 | |
| Slc35a1 | 1 | 1 | |
| Slc35e1 | 1 | 1 | |
| Slc35e4 | 1 | 1 | |
| Slc37a2 | 1 | 1 | |
| Slc38a4 | 1 | 1 | |
| Slc41a1 | 1 | 1 | |
| Slc43a2 | 1 | 1 | |
| Slc44a2 | 1 | 1 | |
| Slc44a5 | 1 | 1 | |
| Slc50a1 | 1 | 1 | |
| Slc5a3 | 1 | 1 | |
| Slc5a5 | 1 | 1 | |
| Slc5a7 | 1 | 1 | |
| Slc6a19 | 1 | 1 | |
| Slc6a2 | 1 | 1 | |
| Slc6a9 | 1 | 1 | |
| Slc7a8 | 1 | 1 | |
| Slc9a8 | 1 | 1 | |
| Slco1c1 | 1 | 1 | |
| Slco2b1 | 1 | 1 | |
| Slco3a1 | 1 | 1 | |
| Slfn5 | 1 | 1 | |
| Slitrk2 | 1 | 1 | |
| Slurp1 | 1 | 1 | |
| Smek1 | 1 | 1 | |
| Smg1 | 1 | 1 | |
| Smim1 | 1 | 1 | |
| Smpd4 | 1 | 1 | |
| Snx16 | 1 | 1 | |
| Snx21 | 1 | 1 | |
| Snx27 | 1 | 1 | |
| Soat1 | 1 | 1 | |

| | Number of miR-96 seed region matches in: | | |
|---|---|---|---|
| Gene | C57BL/6N 3'UTR | C3H/HeJ 3'UTR | Predicted targets of miR-96 |
| Soga1 | 1 | 1 | |
| Sorbs2 | 1 | 1 | |
| Sort1 | 1 | 1 | |
| Sost | 1 | 1 | |
| Sowahc | 1 | 1 | |
| Sox12 | 1 | 1 | |
| Sox8 | 1 | 1 | |
| Sp7 | 1 | 1 | |
| Spast | 1 | 1 | Yes |
| Spata13 | 1 | 1 | |
| Specc1 | 1 | 1 | |
| Speg | 1 | 1 | |
| Spen | 1 | 1 | |
| Sphkap | 1 | 1 | |
| Spin4 | 1 | 1 | |
| Spta1 | 1 | 1 | |
| Srpk1 | 1 | 1 | |
| Srrm3 | 1 | 1 | |
| Srrm4 | 1 | 1 | |
| Srxn1 | 1 | 1 | |
| Ssbp3 | 1 | 1 | |
| Ssc4d | 1 | 1 | |
| Sssca1 | 1 | 1 | |
| St3gal3 | 1 | 1 | |
| St8sia6 | 1 | 1 | |
| Stam | 1 | 1 | |
| Stard7 | 1 | 1 | |
| Stat5b | 1 | 1 | |
| Stat6 | 1 | 1 | |
| Stk10 | 1 | 1 | |
| Stk11 | 1 | 1 | |
| Stk16 | 1 | 1 | |
| Stk17b | 1 | 1 | |
| Stk25 | 1 | 1 | |
| Stmn2 | 1 | 1 | |
| Stmnd1 | 1 | 1 | |
| Stoml1 | 1 | 1 | |
| Strbp | 1 | 1 | |
| Strn4 | 1 | 1 | |
| Stx8 | 1 | 1 | |
| Stxbp4 | 1 | 1 | |
| Sugp2 | 1 | 1 | |
| Sult4a1 | 1 | 1 | |

| | Number of miR-96 seed region matches in: | | |
|---|---|---|---|
| **Gene** | **C57BL/6N 3'UTR** | **C3H/HeJ 3'UTR** | **Predicted targets of miR-96** |
| Sumf2 | 1 | 1 | |
| Surf6 | 1 | 1 | |
| Suv420h1 | 1 | 1 | |
| Sv2c | 1 | 1 | |
| Sybu | 1 | 1 | |
| Syn3 | 1 | 1 | |
| Syngr2 | 1 | 1 | |
| Syngr3 | 1 | 1 | |
| Synpo2 | 1 | 1 | |
| Sypl | 1 | 1 | |
| Tab1 | 1 | 1 | |
| Tab2 | 1 | 1 | |
| Tac1 | 1 | 1 | |
| Tacc1 | 1 | 1 | |
| Taok2 | 1 | 1 | |
| Tax1bp3 | 1 | 1 | |
| Tbc1d1 | 1 | 1 | |
| Tbc1d2 | 1 | 1 | |
| Tbc1d22b | 1 | 1 | |
| Tbc1d5 | 1 | 1 | |
| Tbl1x | 1 | 1 | |
| Tbx1 | 1 | 1 | |
| Tbx15 | 1 | 1 | |
| Tbx18 | 1 | 1 | |
| Tbx20 | 1 | 1 | |
| Tc2n | 1 | 1 | |
| Tceanc | 1 | 1 | |
| Tcf24 | 1 | 1 | |
| Tcf4 | 1 | 1 | |
| Tcf7l2 | 1 | 1 | |
| Tctn1 | 1 | 1 | |
| Tecrl | 1 | 1 | |
| Telo2 | 1 | 1 | |
| Tenm3 | 1 | 1 | |
| Terb1 | 1 | 1 | |
| Tes | 1 | 1 | |
| Tet1 | 1 | 1 | |
| Tfeb | 1 | 1 | |
| Tfpi | 1 | 1 | |
| Tg | 1 | 1 | |
| Tgfbi | 1 | 1 | |
| Tgfbr1 | 1 | 1 | |
| Thada | 1 | 1 | |

| | Number of miR-96 seed region matches in: | | |
|---|---|---|---|
| Gene | C57BL/6N 3'UTR | C3H/HeJ 3'UTR | Predicted targets of miR-96 |
| Thoc1 | 1 | 1 | |
| Thsd7a | 1 | 1 | |
| Tiam1 | 1 | 1 | |
| Timp3 | 1 | 1 | |
| Tinag | 1 | 1 | |
| Tiprl | 1 | 1 | |
| Tktl1 | 1 | 1 | |
| Tmbim6 | 1 | 1 | |
| Tmc3 | 1 | 1 | |
| Tmcc3 | 1 | 1 | |
| Tmco4 | 1 | 1 | |
| Tmed1 | 1 | 1 | |
| Tmem104 | 1 | 1 | |
| Tmem106b | 1 | 1 | |
| Tmem116 | 1 | 1 | |
| Tmem119 | 1 | 1 | |
| Tmem128 | 1 | 1 | |
| Tmem132e | 1 | 1 | |
| Tmem145 | 1 | 1 | |
| Tmem169 | 1 | 1 | |
| Tmem170b | 1 | 1 | |
| Tmem18 | 1 | 1 | |
| Tmem198 | 1 | 1 | |
| Tmem201 | 1 | 1 | |
| Tmem234 | 1 | 1 | |
| Tmem246 | 1 | 1 | |
| Tmem26 | 1 | 1 | |
| Tmem43 | 1 | 1 | |
| Tmem5 | 1 | 1 | |
| Tmem57 | 1 | 1 | |
| Tmem63c | 1 | 1 | |
| Tmem67 | 1 | 1 | |
| Tmem68 | 1 | 1 | |
| Tmppe | 1 | 1 | |
| Tmx1 | 1 | 1 | |
| Tmx4 | 1 | 1 | Yes |
| Tnfaip2 | 1 | 1 | |
| Tnfaip8l1 | 1 | 1 | |
| Tnfrsf11a | 1 | 1 | |
| Tnfrsf19 | 1 | 1 | |
| Tnfrsf1a | 1 | 1 | |
| Tnfsf13 | 1 | 1 | |
| Tnfsfm13 | 1 | 1 | |

| | Number of miR-96 seed region matches in: | | |
|---|---|---|---|
| Gene | C57BL/6N 3'UTR | C3H/HeJ 3'UTR | Predicted targets of miR-96 |
| Tnik | 1 | 1 | |
| Tnip3 | 1 | 1 | |
| Tnrc6b | 1 | 1 | |
| Tns3 | 1 | 1 | |
| Tor1b | 1 | 1 | |
| Tpcn1 | 1 | 1 | |
| Tpm1 | 1 | 1 | |
| Tpm2 | 1 | 1 | |
| Traf7 | 1 | 1 | |
| Trappc12 | 1 | 1 | |
| Treh | 1 | 1 | |
| Trib3 | 1 | 1 | |
| Trim27 | 1 | 1 | |
| Trim31 | 1 | 1 | |
| Trim44 | 1 | 1 | |
| Trim46 | 1 | 1 | |
| Trim55 | 1 | 1 | |
| Trim71 | 1 | 1 | |
| Trmt12 | 1 | 1 | |
| Trp53bp1 | 1 | 1 | |
| Trp73 | 1 | 1 | |
| Trpm5 | 1 | 1 | |
| Tsc22d3 | 1 | 1 | |
| Tsku | 1 | 1 | |
| Tsn | 1 | 1 | |
| Tspan1 | 1 | 1 | |
| Tspan14 | 1 | 1 | |
| Tspan5 | 1 | 1 | |
| Ttc9c | 1 | 1 | |
| Tti2 | 1 | 1 | |
| Ttl | 1 | 1 | |
| Ttll12 | 1 | 1 | |
| Txlna | 1 | 1 | |
| Ube2c | 1 | 1 | |
| Ube2f | 1 | 1 | |
| Ube2g1 | 1 | 1 | |
| Ubfd1 | 1 | 1 | |
| Ubox5 | 1 | 1 | |
| Ufd1l | 1 | 1 | |
| Ufm1 | 1 | 1 | |
| Uggt1 | 1 | 1 | |
| Ugt8a | 1 | 1 | |
| Uimc1 | 1 | 1 | |

| Gene | Number of miR-96 seed region matches in: C57BL/6N 3'UTR | C3H/HeJ 3'UTR | Predicted targets of miR-96 |
|---|---|---|---|
| Umad1 | 1 | 1 | |
| Umps | 1 | 1 | |
| Unc13c | 1 | 1 | |
| Unc45b | 1 | 1 | |
| Usf1 | 1 | 1 | |
| Usp29 | 1 | 1 | |
| Usp45 | 1 | 1 | |
| Uspl1 | 1 | 1 | |
| Utp14b | 1 | 1 | |
| Vamp3 | 1 | 1 | |
| Vamp4 | 1 | 1 | |
| Vamp8 | 1 | 1 | |
| Vangl1 | 1 | 1 | |
| Vangl2 | 1 | 1 | |
| Vapb | 1 | 1 | |
| Vat1 | 1 | 1 | |
| Vat1l | 1 | 1 | |
| Veph1 | 1 | 1 | |
| Vhl | 1 | 1 | |
| Vldlr | 1 | 1 | |
| Vmn1r65 | 1 | 1 | |
| Vps37c | 1 | 1 | |
| Vps37d | 1 | 1 | |
| Vstm2a | 1 | 1 | |
| Vsx1 | 1 | 1 | |
| Vwa5b1 | 1 | 1 | |
| Wbscr22 | 1 | 1 | |
| Wdr12 | 1 | 1 | |
| Wdr13 | 1 | 1 | |
| Wdr26 | 1 | 1 | |
| Wdr36 | 1 | 1 | |
| Wdr4 | 1 | 1 | |
| Wdr82 | 1 | 1 | |
| Wfdc5 | 1 | 1 | |
| Whamm | 1 | 1 | |
| Whsc1 | 1 | 1 | |
| Wipi2 | 1 | 1 | |
| Wnk1 | 1 | 1 | |
| Wnt5b | 1 | 1 | |
| Wwox | 1 | 1 | |
| Xiap | 1 | 1 | |
| Xkr4 | 1 | 1 | |
| Xkr5 | 1 | 1 | |

| | Number of miR-96 seed region matches in: | | |
|---|---|---|---|
| Gene | C57BL/6N 3'UTR | C3H/HeJ 3'UTR | Predicted targets of miR-96 |
| Xpo1 | 1 | 1 | |
| Xrcc3 | 1 | 1 | |
| Xylt1 | 1 | 1 | |
| Yipf4 | 1 | 1 | |
| Yipf6 | 1 | 1 | |
| Zak | 1 | 1 | |
| Zbtb22 | 1 | 1 | |
| Zbtb4 | 1 | 1 | |
| Zbtb7c | 1 | 1 | |
| Zc2hc1c | 1 | 1 | |
| Zc3h12b | 1 | 1 | |
| Zc3h7a | 1 | 1 | |
| Zcchc11 | 1 | 1 | |
| Zcchc24 | 1 | 1 | |
| Zcchc3 | 1 | 1 | |
| Zdbf2 | 1 | 1 | |
| Zdhhc17 | 1 | 1 | |
| Zdhhc18 | 1 | 1 | |
| Zdhhc20 | 1 | 1 | |
| Zdhhc24 | 1 | 1 | |
| Zdhhc3 | 1 | 1 | |
| Zfand2b | 1 | 1 | |
| Zfhx4 | 1 | 1 | |
| Zfp101 | 1 | 1 | |
| Zfp120 | 1 | 1 | |
| Zfp142 | 1 | 1 | |
| Zfp174 | 1 | 1 | |
| Zfp185 | 1 | 1 | |
| Zfp236 | 1 | 1 | |
| Zfp24 | 1 | 1 | |
| Zfp286 | 1 | 1 | |
| Zfp322a | 1 | 1 | |
| Zfp329 | 1 | 1 | |
| Zfp335 | 1 | 1 | |
| Zfp354c | 1 | 1 | |
| Zfp36l1 | 1 | 1 | |
| Zfp385b | 1 | 1 | |
| Zfp39 | 1 | 1 | |
| Zfp408 | 1 | 1 | |
| Zfp41 | 1 | 1 | |
| Zfp410 | 1 | 1 | |
| Zfp458 | 1 | 1 | |
| Zfp512b | 1 | 1 | |

| | Number of miR-96 seed region matches in: | | |
|---|---|---|---|
| Gene | C57BL/6N 3'UTR | C3H/HeJ 3'UTR | Predicted targets of miR-96 |
| Zfp568 | 1 | 1 | |
| Zfp592 | 1 | 1 | |
| Zfp593 | 1 | 1 | |
| Zfp597 | 1 | 1 | |
| Zfp606 | 1 | 1 | |
| Zfp607b | 1 | 1 | |
| Zfp609 | 1 | 1 | |
| Zfp64 | 1 | 1 | |
| Zfp704 | 1 | 1 | |
| Zfp72 | 1 | 1 | |
| Zfp780b | 1 | 1 | |
| Zfp87 | 1 | 1 | |
| Zhx1 | 1 | 1 | |
| Zhx2 | 1 | 1 | |
| Zim1 | 1 | 1 | |
| Zkscan1 | 1 | 1 | |
| Zkscan2 | 1 | 1 | |
| Zmym1 | 1 | 1 | |
| Zranb2 | 1 | 1 | |
| Zrsr1 | 1 | 1 | |
| Zyg11b | 1 | 1 | |
| 6430548M08Rik | 2 | 2 | |
| Aak1 | 2 | 2 | |
| Abcd1 | 2 | 2 | |
| Adamts10 | 2 | 2 | |
| Akap1 | 2 | 2 | |
| Anapc5 | 2 | 2 | |
| Ankfy1 | 2 | 2 | |
| Ankrd33b | 2 | 2 | |
| Aqp5 | 2 | 2 | Yes |
| Arf2 | 2 | 2 | Yes |
| Asb6 | 2 | 2 | |
| Atg16l1 | 2 | 2 | |
| Atg9a | 2 | 2 | |
| Atp13a4 | 2 | 2 | |
| Atxn1 | 2 | 2 | |
| Atxn3 | 2 | 2 | |
| Azin1 | 2 | 2 | |
| B4galnt1 | 2 | 2 | Yes |
| Bcl7c | 2 | 2 | |
| Bicd1 | 2 | 2 | |
| Brinp2 | 2 | 2 | |
| Brpf3 | 2 | 2 | |

| | Number of miR-96 seed region matches in: | | |
|---|---|---|---|
| **Gene** | **C57BL/6N 3'UTR** | **C3H/HeJ 3'UTR** | **Predicted targets of miR-96** |
| Cacna1c | 2 | 2 | |
| Cacnb1 | 2 | 2 | |
| Cacnb4 | 2 | 2 | |
| Camk1d | 2 | 2 | |
| Cbx5 | 2 | 2 | |
| Cd22 | 2 | 2 | |
| Cep170 | 2 | 2 | |
| Clasp1 | 2 | 2 | |
| Clmn | 2 | 2 | |
| Cnot9 | 2 | 2 | |
| Cpeb1 | 2 | 2 | |
| Creb1 | 2 | 2 | |
| D1Ertd622e | 2 | 2 | |
| Dab2ip | 2 | 2 | |
| Disp2 | 2 | 2 | |
| Drg1 | 2 | 2 | |
| Dtx4 | 2 | 2 | |
| Dus2 | 2 | 2 | |
| Eif4ebp2 | 2 | 2 | |
| Enpp1 | 2 | 2 | |
| Epha4 | 2 | 2 | |
| Ephb2 | 2 | 2 | |
| Ext1 | 2 | 2 | |
| Fbxl18 | 2 | 2 | |
| Fibcd1 | 2 | 2 | |
| Fosl2 | 2 | 2 | |
| Foxk2 | 2 | 2 | |
| Fut9 | 2 | 2 | |
| Gabra2 | 2 | 2 | |
| Gad2 | 2 | 2 | Yes |
| Galnt2 | 2 | 2 | |
| Glb1l | 2 | 2 | |
| Gm45140 | 2 | 2 | |
| Gna11 | 2 | 2 | |
| Gpm6b | 2 | 2 | |
| Gpr101 | 2 | 2 | |
| Gtf2h1 | 2 | 2 | |
| Gtf3c1 | 2 | 2 | |
| Gxylt1 | 2 | 2 | |
| Hmgxb4 | 2 | 2 | |
| Hr | 2 | 2 | |
| Jph2 | 2 | 2 | |
| Kif26b | 2 | 2 | |

| Gene | Number of miR-96 seed region matches in: C57BL/6N 3'UTR | C3H/HeJ 3'UTR | Predicted targets of miR-96 |
|---|---|---|---|
| Lair1 | 2 | 2 | |
| Ldb3 | 2 | 2 | |
| Lgr6 | 2 | 2 | |
| Lrrc3 | 2 | 2 | |
| Lrrtm2 | 2 | 2 | |
| Lrtm1 | 2 | 2 | |
| Man1a2 | 2 | 2 | |
| Mfsd12 | 2 | 2 | |
| Mmd2 | 2 | 2 | |
| Mpv17l | 2 | 2 | |
| Msantd4 | 2 | 2 | |
| Msn | 2 | 2 | |
| Mtor | 2 | 2 | |
| Mvb12b | 2 | 2 | |
| Mysm1 | 2 | 2 | |
| Natd1 | 2 | 2 | |
| Nhlrc3 | 2 | 2 | |
| Nova1 | 2 | 2 | |
| Nudt3 | 2 | 2 | |
| Osbpl10 | 2 | 2 | |
| Pax5 | 2 | 2 | |
| Pcif1 | 2 | 2 | |
| Pgap1 | 2 | 2 | |
| Phrf1 | 2 | 2 | |
| Pikfyve | 2 | 2 | |
| Pmepa1 | 2 | 2 | |
| Pofut1 | 2 | 2 | |
| Ppm1f | 2 | 2 | |
| Ppp1r12b | 2 | 2 | |
| Prpf19 | 2 | 2 | |
| Prrx1 | 2 | 2 | |
| Pura | 2 | 2 | |
| Pxdn | 2 | 2 | |
| Rab35 | 2 | 2 | |
| Rassf2 | 2 | 2 | |
| Rbm39 | 2 | 2 | |
| Rimkla | 2 | 2 | |
| Rptor | 2 | 2 | |
| Rsf1 | 2 | 2 | |
| Slc1a1 | 2 | 2 | Yes |
| Slc38a9 | 2 | 2 | |
| Slc7a2 | 2 | 2 | |
| Slco4c1 | 2 | 2 | |

| | Number of miR-96 seed region matches in: | | |
|---|---|---|---|
| Gene | C57BL/6N 3'UTR | C3H/HeJ 3'UTR | Predicted targets of miR-96 |
| Smad7 | 2 | 2 | |
| Socs6 | 2 | 2 | |
| Socs7 | 2 | 2 | |
| Sox5 | 2 | 2 | |
| Sox6 | 2 | 2 | |
| Sppl2a | 2 | 2 | |
| Spsb1 | 2 | 2 | |
| St3gal6 | 2 | 2 | |
| St6galnac3 | 2 | 2 | |
| St6galnac5 | 2 | 2 | |
| St7 | 2 | 2 | |
| St8sia3 | 2 | 2 | Yes |
| Sypl2 | 2 | 2 | |
| Tbc1d24 | 2 | 2 | |
| Tbr1 | 2 | 2 | |
| Tfcp2l1 | 2 | 2 | |
| Tm9sf4 | 2 | 2 | |
| Tmem150c | 2 | 2 | |
| Tmem163 | 2 | 2 | |
| Tmem186 | 2 | 2 | |
| Tnr | 2 | 2 | |
| Ttc38 | 2 | 2 | |
| Ube2l3 | 2 | 2 | |
| Uck2 | 2 | 2 | |
| Vtcn1 | 2 | 2 | |
| Vwa1 | 2 | 2 | |
| Zbtb1 | 2 | 2 | |
| Zbtb37 | 2 | 2 | |
| Zcchc16 | 2 | 2 | |
| Zfp334 | 2 | 2 | |
| Zfp626 | 2 | 2 | |
| Zfp961 | 2 | 2 | |
| C77370 | 3 | 3 | |
| Cdc73 | 3 | 3 | |
| Gng12 | 3 | 3 | |
| Hacd4 | 3 | 3 | |
| Jam2 | 3 | 3 | |
| Nlgn2 | 3 | 3 | |
| Plppr4 | 3 | 3 | |
| Snx30 | 3 | 3 | |
| Sun2 | 3 | 3 | |
| Tmem178b | 3 | 3 | |
| Ttyh3 | 3 | 3 | |

| | Number of miR-96 seed region matches in: | | |
|---|---|---|---|
| **Gene** | **C57BL/6N 3'UTR** | **C3H/HeJ 3'UTR** | **Predicted targets of miR-96** |
| Rab3c | 4 | 4 | |
| Sh3pxd2b | 4 | 4 | |
| Syt9 | 4 | 4 | |
| 4930402H24Rik | 0 | 1 | |
| 4930427A07Rik | 0 | 1 | |
| 6030445D17Rik | 0 | 1 | |
| Adamts2 | 0 | 1 | |
| Ahrr | 0 | 1 | |
| Ceacam3 | 0 | 1 | |
| Cyth3 | 0 | 1 | |
| Diablo | 0 | 1 | |
| Dlg1 | 0 | 1 | |
| Dnmt3a | 0 | 1 | |
| Dtwd2 | 0 | 1 | |
| Eif4g3 | 0 | 1 | |
| Ercc6l2 | 0 | 1 | |
| Gnpda1 | 0 | 1 | |
| Homer2 | 0 | 1 | |
| Iqck | 0 | 1 | |
| Khdrbs1 | 0 | 1 | |
| Masp1 | 0 | 1 | |
| Med1 | 0 | 1 | |
| Mtrr | 0 | 1 | |
| Ncan | 0 | 1 | |
| Pdcd10 | 0 | 1 | |
| Pik3ca | 0 | 1 | |
| Rln3 | 0 | 1 | |
| Rnasek | 0 | 1 | |
| Rnf121 | 0 | 1 | |
| Slc36a4 | 0 | 1 | |
| Slc39a10 | 0 | 1 | |
| Sstr3 | 0 | 1 | |
| Tead3 | 0 | 1 | |
| Timm17a | 0 | 1 | |
| Tmem52b | 0 | 1 | |
| Tmem71 | 0 | 1 | |
| Zfp449 | 0 | 1 | |
| Afap1l1 | 0 | 2 | |
| Hn1l | 0 | 2 | |
| 1810009A15Rik | 1 | 0 | |
| 1810032O08Rik | 1 | 0 | |
| 9030624J02Rik | 1 | 0 | |
| Ap3s1 | 1 | 0 | |

| | Number of miR-96 seed region matches in: | | |
|---|---|---|---|
| Gene | C57BL/6N 3'UTR | C3H/HeJ 3'UTR | Predicted targets of miR-96 |
| Arid1b | 1 | 0 | |
| Bhmt2 | 1 | 0 | |
| Caskin1 | 1 | 0 | |
| Casp9 | 1 | 0 | |
| Cds2 | 1 | 0 | |
| Ciao1 | 1 | 0 | |
| Ciapin1 | 1 | 0 | |
| Crim1 | 1 | 0 | |
| Ctdsp2 | 1 | 0 | |
| Ddx5 | 1 | 0 | |
| Dgkh | 1 | 0 | |
| Dgkk | 1 | 0 | |
| Gan | 1 | 0 | |
| Ginm1 | 1 | 0 | |
| Gm28042 | 1 | 0 | |
| Grem2 | 1 | 0 | |
| Grk6 | 1 | 0 | |
| Grm5 | 1 | 0 | |
| Mdm4 | 1 | 0 | |
| Miip | 1 | 0 | |
| Nos1ap | 1 | 0 | |
| Pithd1 | 1 | 0 | |
| Pm20d1 | 1 | 0 | |
| Pnma3 | 1 | 0 | |
| Ppp6r1 | 1 | 0 | |
| Rapgefl1 | 1 | 0 | |
| Rbfox1 | 1 | 0 | |
| Rnf222 | 1 | 0 | |
| Rrm2b | 1 | 0 | |
| Sh3bgrl2 | 1 | 0 | |
| Shc2 | 1 | 0 | |
| Slc22a5 | 1 | 0 | |
| Smo | 1 | 0 | |
| Srsf10 | 1 | 0 | |
| Tmem97 | 1 | 0 | |
| Tpi1 | 1 | 0 | |
| Tsfm | 1 | 0 | |
| Zbtb14 | 1 | 0 | |
| Zfp710 | 1 | 0 | |
| Aste1 | 1 | 2 | |
| Insig2 | 1 | 2 | Yes |
| Lrrc58 | 1 | 2 | |
| Phf20l1 | 1 | 2 | |

| Gene | Number of miR-96 seed region matches in: C57BL/6N 3'UTR | C3H/HeJ 3'UTR | Predicted targets of miR-96 |
|---|---|---|---|
| Tirap | 1 | 2 | |
| Zbtb41 | 1 | 2 | |
| Morf4l2 | 2 | 0 | |
| Pole3 | 2 | 0 | |
| Slc4a8 | 2 | 0 | |
| Spin1 | 2 | 0 | |
| Cep170b | 2 | 1 | Yes |
| Msh3 | 2 | 1 | |
| Sass6 | 2 | 1 | |
| Sptlc2 | 2 | 1 | |
| Zfp775 | 2 | 1 | |
| Tmem245 | 3 | 1 | |
| Adcy6 | 3 | 2 | Yes |
| 1190007I07Rik | 1 | Sequence unavailable | |
| BC024978 | 1 | Sequence unavailable | |
| Brwd1 | 1 | Sequence unavailable | |
| Camsap1 | 1 | Sequence unavailable | |
| Cd244 | 1 | Sequence unavailable | |
| Cfhr1 | 1 | Sequence unavailable | |
| Dusp16 | 1 | Sequence unavailable | |
| Fgd2 | 1 | Sequence unavailable | |
| Ints4 | 1 | Sequence unavailable | |
| Kctd14 | 1 | Sequence unavailable | |
| Krtap13 | 1 | Sequence unavailable | |
| Lix1 | 1 | Sequence unavailable | |
| Ptdss2 | 1 | Sequence unavailable | |
| Sgpl1 | 1 | Sequence unavailable | |
| Shc1 | 1 | Sequence unavailable | |
| Slc22a21 | 1 | Sequence unavailable | |
| Smg5 | 1 | Sequence unavailable | |
| Stk19 | 1 | Sequence unavailable | |
| Stk35 | 1 | Sequence unavailable | |
| Tdgf1 | 1 | Sequence unavailable | |
| Thoc2 | 1 | Sequence unavailable | |
| Zeb1 | 1 | Sequence unavailable | |
| Zfand5 | 1 | Sequence unavailable | |
| Zfp874a | 1 | Sequence unavailable | |
| Zkscan3 | 1 | Sequence unavailable | |
| Slc12a6 | 2 | Sequence unavailable | |
| Slc39a1 | 2 | Sequence unavailable | |
| Zfp111 | 2 | Sequence unavailable | |
| Zfp850 | 2 | Sequence unavailable | |
| Zscan20 | 2 | Sequence unavailable | |

| | Number of miR-96 seed region matches in: | | |
|---|---|---|---|
| **Gene** | **C57BL/6N 3'UTR** | **C3H/HeJ 3'UTR** | **Predicted targets of miR-96** |
| Lrch2 | 3 | Sequence unavailable | |
| Btg1 | Sequence unavailable | 1 | |
| Cgn | Sequence unavailable | 1 | |
| Cnot3 | Sequence unavailable | 1 | |
| Crb1 | Sequence unavailable | 1 | |
| Gm10065 | Sequence unavailable | 1 | |
| Hc | Sequence unavailable | 1 | |
| Itgb1bp1 | Sequence unavailable | 1 | |
| Kcnv1 | Sequence unavailable | 1 | |
| Lrp1 | Sequence unavailable | 1 | |
| Nacc1 | Sequence unavailable | 1 | |
| Ntn4 | Sequence unavailable | 1 | |
| Pclo | Sequence unavailable | 1 | |
| Pde8b | Sequence unavailable | 1 | |
| Phf20 | Sequence unavailable | 1 | |
| Pou2f2 | Sequence unavailable | 1 | |
| Rdh16 | Sequence unavailable | 1 | |
| Scn1a | Sequence unavailable | 1 | |
| Sh2d1a | Sequence unavailable | 1 | |
| Sp100 | Sequence unavailable | 1 | |
| Usf3 | Sequence unavailable | 1 | |
| Polq | Sequence unavailable | 2 | |

Supplementary Table S2. Genes with significant differential splicing predicted by Cuffdiff, Leafcutter and JunctionSeq in *Mir183/96*$^{dko}$ homozygotes (A) and *Mir182*$^{ko}$ homozygotes (B). * indicates which predictions were tested by sequencing. None of the differential splicing predictions were confirmed.

**(A) *Mir183/96*$^{dko}$**

| **Cuffdiff** | | **Leafcutter** | | **JunctionSeq** | |
|---|---|---|---|---|---|
| **Gene** | **Significance (adjusted p value)** | **Gene** | **Significance (FDR)** | **Gene** | **Significance (adjusted p value)** |
| Rars | 0.00972143 | Rnf157 | 0.000261808 | Wdpcp* | 0.000364 |
| Ncor1 | 0.00972143 | Ppp3cb | 0.000437474 | Dlgap3* | 0.000566 |
| Srr | 0.00972143 | Slc22a15 | 0.001567787 | Tm2d1 | 0.00114 |
| Npepps | 0.00972143 | Spag9 | 0.001780359 | Cdc73 | 0.00158 |
| Slc38a10 | 0.00972143 | Slc8a1 | 0.001780359 | Stard9* | 0.00158 |
| Naa35 | 0.00972143 | Pik3c2a | 0.002821861 | Nav2* | 0.00158 |
| Wdr37 | 0.00972143 | Tnc* | 0.003622023 | Insc | 0.00296 |
| Adgrb1 | 0.00972143 | Abi2 | 0.004435971 | Ddx11* | 0.00455 |
| Tmbim6 | 0.00972143 | Nsfl1c | 0.00585876 | Dnah8* | 0.00487 |
| Scn8a | 0.00972143 | Cap1 | 0.007645588 | Zfp618* | 0.0091 |
| Acvrl1 | 0.00972143 | Plscr3 | 0.007783801 | Igf2* | 0.0124 |
| Rapgef3 | 0.00972143 | Camk2g | 0.009450284 | Slc44a5* | 0.0151 |
| Cfap126 | 0.00972143 | Nfasc | 0.009450284 | Adamtsl4* | 0.024 |
| Tnk2 | 0.00972143 | Zfp280d | 0.010903811 | Ppp3cb* | 0.029 |
| A930003A15Rik,Gm27883 | 0.00972143 | Tmod1 | 0.018799915 | Nrxn2 | 0.029 |
| Atat1 | 0.00972143 | Skp1a | 0.026059743 | Rims3 | 0.0413 |
| Fam98a | 0.00972143 | Adgrl3 | 0.026059743 | Greb1l* | 0.0435 |
| Cabyr | 0.00972143 | Raver1 | 0.026059743 | Fam96a* | 0.0435 |
| Snhg1 | 0.00972143 | Zfp37 | 0.027564421 | Zfp280c | 0.0443 |
| Snx15 | 0.00972143 | Xpot | 0.030579786 | Ranbp17* | 0.0482 |
| Slc3a2 | 0.00972143 | Rnf180 | 0.030579786 | Onecut2 | 0.0482 |
| Fam107b | 0.00972143 | Sgms1 | 0.030579786 | Stx16 | 0.0482 |
| Scn2a1 | 0.00972143 | Fcrlb | 0.030579786 | Gpnmb* | 0.0482 |
| Fmn1 | 0.00972143 | Selenom | 0.030677894 | | |
| Smox | 0.00972143 | Ppip5k2 | 0.030677894 | | |
| Zeb2 | 0.00972143 | Rbms3 | 0.030898362 | | |
| P2rx3 | 0.00972143 | Sgcd | 0.031113858 | | |
| Ubr1 | 0.00972143 | Efemp1 | 0.031665004 | | |
| Tcfl5 | 0.00972143 | Slc4a1ap | 0.033113361 | | |

| | | | |
|---|---|---|---|
| Rhoc | 0.00972143 | Mgat5 | 0.035600128 |
| Mcoln3 | 0.00972143 | Clec16a | 0.036013137 |
| Cryz | 0.00972143 | Cobll1 | 0.036013137 |
| Sec24b | 0.00972143 | Kazn | 0.036013137 |
| Sec61b | 0.00972143 | Ubn2 | 0.036013137 |
| Megf6 | 0.00972143 | Snrk | 0.036013137 |
| Mrpl20 | 0.00972143 | Nckap5l | 0.037180232 |
| Tnfrsf4 | 0.00972143 | Mrpl24 | 0.037180232 |
| Ptbp3 | 0.00972143 | Ppfia2 | 0.037990292 |
| Brca2 | 0.00972143 | Nufip2 | 0.037990292 |
| G3bp2 | 0.00972143 | Copa | 0.037990292 |
| Slc15a4 | 0.00972143 | Fbxl20 | 0.040667577 |
| 5930412G12Rik | 0.00972143 | Efr3a | 0.040667577 |
| Auts2 | 0.00972143 | Pfdn5 | 0.040667577 |
| Il17re | 0.00972143 | Ablim1 | 0.040667577 |
| Sec13 | 0.00972143 | Gab2 | 0.040667577 |
| Clpb | 0.00972143 | Hpf1 | 0.040667577 |
| Anpep | 0.00972143 | Zfp651 | 0.040667577 |
| Spats2l | 0.00972143 | Rpl26 | 0.043842403 |
| Apc2 | 0.00972143 | 2810428I15Rik | 0.043842403 |
| Smarcc1 | 0.00972143 | Rnps1 | 0.044056022 |
| Birc3 | 0.00972143 | Tial1 | 0.044056022 |
| Cntn5 | 0.00972143 | Tbata | 0.044623796 |
| Gramd1b | 0.00972143 | Tlk2 | 0.044623796 |
| Agtr2 | 0.00972143 | Cryl1 | 0.044623796 |
| Gm27875,Jpx | 0.00972143 | Mdga1 | 0.044623796 |
| Ctps2 | 0.00972143 | Lcor | 0.044623796 |
| Wrn | 0.0191018 | Slc25a44 | 0.044623796 |
| | | Phf13 | 0.044623796 |
| | | Prkcsh | 0.044623796 |
| | | Cdv3 | 0.044623796 |
| | | Decr2 | 0.044950165 |
| | | Mbd5 | 0.044950165 |
| | | Carm1 | 0.046056306 |
| | | Calml4 | 0.046056306 |
| | | Cd200 | 0.046119016 |
| | | Pdzd2 | 0.04741293 |
| | | Zfpm2 | 0.04741293 |
| | | Arfgef2 | 0.04741293 |
| | | Zfp618 | 0.04741293 |
| | | Deaf1 | 0.04741293 |

| | | |
|---|---|---|
| | Appl1 | 0.047696496 |
| | Pdgfc | 0.047696496 |
| | Elovl1 | 0.047696496 |
| | Prickle2 | 0.047696496 |
| | Il17re | 0.047696496 |
| | Rpl10 | 0.04782345 |
| | Dtnb | 0.049562528 |
| | Nol4l | 0.049716919 |

***(B) Mir182$^{ko}$***

| **Cuffdiff** | | **Leafcutter** | |
|---|---|---|---|
| **Gene** | **Significance (adjusted p value)** | **Gene** | **Significance (FDR)** |
| Tada2a | 0.037425 | Ube4a* | 0.003625717 |
| Tnfaip2 | 0.037425 | Prdm4 | 0.013483712 |
| Psmd6 | 0.037425 | Irf2bp2 | 0.013483712 |
| Snhg7 | 0.037425 | Atrx | 0.013483712 |
| Ptpn4 | 0.037425 | Tmem184b | 0.016348513 |
| Ptpn3 | 0.037425 | Rad17 | 0.017626834 |
| Nfyc | 0.037425 | Pbrm1 | 0.017626834 |
| Rpl6 | 0.037425 | Nrxn1* | 0.017626834 |
| Auts2 | 0.037425 | Fubp3 | 0.017626834 |
| Casc1 | 0.037425 | Adgrl3 | 0.017626834 |
| Nr3c2 | 0.037425 | Ddx59 | 0.017790912 |
| Mre11a | 0.037425 | Arhgef1 | 0.017790912 |
| Usp2 | 0.037425 | Kctd17 | 0.029537826 |
| Ryk | 0.037425 | Kdm1a | 0.029865303 |
| | | Ptprt | 0.036976191 |
| | | Adamtsl1 | 0.036976191 |
| | | Cbx1 | 0.037836631 |
| | | Ttc3 | 0.037836631 |
| | | Rsrc1 | 0.037836631 |
| | | Pan3 | 0.037836631 |
| | | Sec23ip | 0.037836631 |
| | | Col4a5 | 0.037836631 |
| | | Slc39a8 | 0.038458003 |
| | | Ncor2 | 0.040186555 |
| | | Trit1 | 0.040463733 |
| | | Tfpi | 0.044934868 |

| | | |
|---|---|---|
| | Zfp719 | 0.045815348 |

Supplementary Table S3. (A) Genotyping primers. (B) Primers used for testing differential splicing. (C) Taqman primer/probe sets used for qRTPCR. (D) MicroRNA primer sets used for qRTPCR.

**(A) Primers for genotyping**

| Allele | Primer F (5'-3') | Primer R (5'-3') |
|---|---|---|
| Mir183/96$^{dko}$ | tattgggatgtgatgggaaactctg | tagcagaaggctagaccccaaagac |
| Mir182$^{ko}$ (1) | gggtacagtgcctttgagagcagt | gggaaacattaagggtcacttccag |
| Mir182$^{ko}$ (2) | gcttgaggaggttttacactgg | ttcctggtgatcggcagg |

**(B) Primers for sequencing**

| Gene | Transcript | Primer F (5'-3') | Primer R (5'-3') |
|---|---|---|---|
| Wdpcp | ENSMUST00000020568 | TCTTAGACAGAGGCTCACACC | CATGCGCCCTTCTTCTTGC |
| Dlgap3 | ENSMUST00000106094 | ACACCGAGAACAGGAGTCG | CTGGAAGGTGGGCACAGG |
| Stard9 | ENSMUST00000180041 | ACATCATCAACAAGCCACGG | TCTCTCTCAATCCACTGGCC |
| Ddx11 | ENSMUST00000163605 | TGCCCCTATTACGGAAGCC | CATGTACTGGAGCAACTGGG |
| Dnah8 | ENSMUST00000170651 | CAGGAGGGAGGAAGATGACG | GGTCAAAGTATCCACAGCCG |
| Zfp618 | ENSMUST00000030043 | CTGGAAGGAAAAGCGCGG | GATCCCGCATTCATAGGACC |
| Igf2 | ENSMUST00000000033 | GAGTTCAGAGAGGCCAAACG | TGTTCTGTTCTTCTCCTTGGG |
| Slc44a5 | ENSMUST00000089948 | TTAGCTACCTTCCCAGTGCG | AGATCCAAAGGCCAGAGACC |
| Adamtsl4 | ENSMUST00000117782 | TAACGCCATGCTCCTCCC | TGGGCTTCTGGATGTCTTGG |
| Ppp3cb | ENSMUST00000159027 | CAGTTTAATTGCTCTCCACATCC | CAGCACGCTTTCACTCTCC |
| Greb1l | ENSMUST00000048977 | TCAAACAGCCCACCAATTCC | CAAAGAGCAGAGGATTGTCGG |
| Fam69a | ENSMUST00000034945 | CGGATCATGGAAGAGAAAGCG | CACTGTTCCACGATTTCCCG |
| Ranbp17 | ENSMUST00000102815 | TTGACAATGTACTCCAGGCC | GGTTCTGTTCCACTCCTTCC |
| Gpnmb | ENSMUST00000031840 | TCTGCCCACATTATCAACACC | TCGGAGATGATCGTACAGGC |

| | | | |
|---|---|---|---|
| Tnc | ENSMUST00000107377 | GTTACCGCCTCAACTACAGC | TGAGGTTCTGGACAGTCTGG |
| Ube4a | ENSMUST00000117506 | TAGCTGTGAGGTGTCGTCG | AGTCATCTGGGCTTGCTGG |
| Nrxn1 | ENSMUST00000160844 | GCACACCTGATGATGGGC | GCTGATTTCCCTGTGTGAAGC |

**(C) qPCR primer/probes**

| Gene | Primer/probe catalogue number |
|---|---|
| Hprt | Mm01318747_g1 |
| Ppm1l | Mm00618786_m1 |
| Jag1 | Mm01270190_m1 |
| Ccer2 | Mm01179046_g1 |
| Grp | Mm00612977_m1 |
| Grk1 | Mm01220714_m1 |
| Tmc1 | 4331348 (manual design using Applied Biosystems software) |
| Hspa2 | Mm00434069_s1 |
| Myo3a | 4331348 (manual design using Applied Biosystems software) |
| St8sia3 | 4331348 (manual design using Applied Biosystems software) |
| Tmem173 | Mm01158119_g1 |
| Slc52a3 | Mm00510191_g1 |
| Kif21b | Mm01285309_g1 |
| Mfsd6 | Mm00505561_m1 |
| BC030867 | Mm00463529_m1 |
| Eln | Mm00514696_g1 |
| P2rx3 | Mm01278228_g1 |
| Ttc21a | Mm01351694_g1 |
| Slc6a11 | Mm01190466_m1 |

| | |
|---|---|
| Dtna | Mm01135282_m1 |
| Ikzf2 | Mm00496108_m1 |
| Rest | Mm00803267_m1 |
| Cebpa | Mm01265914_s1 |
| Cdkn1a | Mm00432448_m1 |
| Fos | Mm01302932_g1 |
| Nr3c1 | Mm01260500_m1 |
| Foxo3 | Mm01185722_m1 |
| Foxo1 | Mm00490672_m1 |
| Tgfb1 | Mm00441729_g1 |
| Sp1 | Mm03053855_g1 |
| Trp53 | Mm01337166_mH |
| Ocm | Mm00712881_g1 |
| Slc26a5 | 4331348 (manual design using Applied Biosystems software) |

**(D) microRNA qPCR primer sets**

| Gene | Primer set catalogue number |
|---|---|
| hsa-miR-96-5p | YP00204417 |
| mmu-miR-182-5p | YP00205089 |
| hsa-miR-183-5p | YP00206030 |
| hsa-miR-99a-5p | YP00204521 |

Supplementary Table S4. Modules detected by WGCNA, their correlation with the two genotype traits we examined (wildtype vs *Mir183/96*$^{dko}$ and wildtype vs *Mir182*$^{ko}$) and the number of genes and differentially expressed genes in each module. The first nine modules were chosen for further analysis.

| **Module** | **black** | **green** | **royal blue** | **salmon** | **blue** | **purple** | **grey60** | **light green** | **dark green** | **pink** |
|---|---|---|---|---|---|---|---|---|---|---|
| Correlation for wildtype vs. *Mir183/96*$^{dko}$ homozygote trait | 0.176 | 0.358 | 0.418 | -0.594 | -0.357 | -0.304 | 0.275 | -0.428 | -0.430 | -0.021 |
| p-value for wildtype vs. *Mir183/96*$^{dko}$ homozygote trait | 0.410 | 0.086 | 0.042 | 0.002 | 0.087 | 0.148 | 0.193 | 0.037 | 0.036 | 0.924 |
| Correlation for wildtype vs. *Mir182*$^{ko}$ homozygote trait | -0.543 | -0.454 | -0.270 | 0.375 | 0.078 | 0.457 | 0.416 | 0.053 | 0.181 | 0.328 |
| p-value for wildtype vs. *Mir182*$^{ko}$ homozygote trait | 0.006 | 0.026 | 0.203 | 0.071 | 0.717 | 0.025 | 0.043 | 0.807 | 0.398 | 0.117 |
| Number of genes | 858 | 1624 | 261 | 583 | 3307 | 695 | 314 | 305 | 240 | 851 |
| Number of differentially expressed genes (*Mir183/96*$^{dko}$) | 0 | 3 | 0 | 0 | 13 | 1 | 1 | 2 | 0 | 0 |
| Number of differentially expressed genes (*Mir182*$^{ko}$) | 0 | 0 | 0 | 0 | 2 | 0 | 0 | 0 | 0 | 0 |

| **Module** | **brown** | **cyan** | **dark grey** | **grey** | **dark red** | **dark turquoise** | **green yellow** | **dark orange** | **light cyan** | **light yellow** |
|---|---|---|---|---|---|---|---|---|---|---|
| Correlation for wildtype vs. *Mir183/96*$^{dko}$ homozygote trait | -0.297 | -0.313 | -0.194 | -0.245 | -0.312 | 0.210 | -0.218 | -0.305 | 0.279 | -0.368 |
| p-value for wildtype vs. *Mir183/96*$^{dko}$ homozygote trait | 0.159 | 0.136 | 0.364 | 0.250 | 0.138 | 0.325 | 0.306 | 0.147 | 0.186 | 0.077 |
| Correlation for wildtype vs. *Mir182*$^{ko}$ homozygote trait | 0.295 | 0.233 | 0.330 | 0.438 | -0.232 | 0.081 | 0.069 | 0.341 | 0.325 | 0.195 |
| p-value for wildtype vs. *Mir182*$^{ko}$ homozygote trait | 0.162 | 0.273 | 0.115 | 0.032 | 0.276 | 0.707 | 0.750 | 0.103 | 0.121 | 0.360 |
| Number of genes | 2257 | 575 | 131 | 5972 | 258 | 152 | 666 | 96 | 354 | 298 |
| Number of differentially expressed genes (*Mir183/96*$^{dko}$) | 4 | 0 | 0 | 1 | 0 | 0 | 0 | 0 | 0 | 0 |
| Number of differentially expressed genes (*Mir182*$^{ko}$) | 1 | 0 | 0 | 0 | 0 | 0 | 0 | 0 | 0 | 0 |

| **Module** | **magenta** | **red** | **orange** | **tan** | **sky blue** | **midnight blue** | **turquoise** | **white** | **yellow** |
|---|---|---|---|---|---|---|---|---|---|
| Correlation for wildtype vs. *Mir183/96*$^{dko}$ homozygote trait | -0.035 | 0.347 | 0.343 | -0.242 | -0.095 | 0.138 | 0.239 | -0.238 | 0.301 |
| p-value for wildtype vs. *Mir183/96*$^{dko}$ homozygote trait | 0.872 | 0.096 | 0.100 | 0.254 | 0.659 | 0.520 | 0.261 | 0.263 | 0.153 |
| Correlation for wildtype vs. *Mir182*$^{ko}$ homozygote trait | -0.016 | 0.084 | -0.089 | -0.259 | 0.326 | 0.177 | 0.334 | -0.237 | 0.114 |

| | | | | | | | | | |
|---|---|---|---|---|---|---|---|---|---|
| p-value for wildtype vs. *Mir182$^{ko}$* homozygote trait | 0.940 | 0.697 | 0.678 | 0.222 | 0.120 | 0.409 | 0.111 | 0.265 | 0.595 |
| Number of genes | 758 | 1019 | 104 | 622 | 86 | 483 | 8793 | 90 | 2225 |
| Number of differentially expressed genes (*Mir183/96$^{dko}$*) | 2 | 5 | 0 | 0 | 0 | 0 | 1 | 0 | 1 |
| Number of differentially expressed genes (*Mir182$^{ko}$*) | 0 | 0 | 0 | 0 | 0 | 0 | 0 | 0 | 0 |

Supplementary Table S5. REACTOME enrichment scores for the nine WGCNA modules tested. If no entries are shown for a module, no pathway terms were returned at all.

| **Black module** | **Mus musculus genes (22296)** | **Module genes (711)** | **Module genes (expected)** | **Module genes fold Enrichment** | **Module genes FDR** |
|---|---|---|---|---|---|
| mRNA Splicing - Major Pathway (R-MMU-72163) | 170 | 27 | 5.42 | 4.98 | 3.79E-08 |
| mRNA Splicing (R-MMU-72172) | 177 | 28 | 5.64 | 4.96 | 2.47E-08 |
| Processing of Capped Intron-Containing Pre-mRNA (R-MMU-72203) | 231 | 32 | 7.37 | 4.34 | 3.41E-08 |
| RNA Polymerase II Pre-transcription Events (R-MMU-674695) | 78 | 10 | 2.49 | 4.02 | 3.65E-02 |
| Metabolism of RNA (R-MMU-8953854) | 500 | 50 | 15.94 | 3.14 | 1.74E-08 |
| Cilium Assembly (R-MMU-5617833) | 184 | 16 | 5.87 | 2.73 | 4.13E-02 |
| Transcriptional Regulation by TP53 (R-MMU-3700989) | 276 | 23 | 8.8 | 2.61 | 7.71E-03 |
| RHO GTPase Effectors (R-MMU-195258) | 253 | 21 | 8.07 | 2.6 | 1.45E-02 |
| Antigen processing: Ubiquitination & Proteasome degradation (R-MMU-983168) | 315 | 23 | 10.05 | 2.29 | 3.57E-02 |
| Class I MHC mediated antigen processing & presentation (R-MMU-983169) | 370 | 26 | 11.8 | 2.2 | 3.33E-02 |
| Gene expression (Transcription) (R-MMU-74160) | 961 | 66 | 30.65 | 2.15 | 4.94E-06 |
| RNA Polymerase II Transcription (R-MMU-73857) | 851 | 56 | 27.14 | 2.06 | 1.70E-04 |

| | | | | | |
|---|---|---|---|---|---|
| Generic Transcription Pathway (R-MMU-212436) | 733 | 42 | 23.37 | 1.8 | 3.73E-02 |
| Post-translational protein modification (R-MMU-597592) | 1325 | 69 | 42.25 | 1.63 | 1.32E-02 |
| Metabolism of proteins (R-MMU-392499) | 1720 | 88 | 54.85 | 1.6 | 2.88E-03 |
| Unclassified (UNCLASSIFIED) | 12915 | 355 | 411.85 | 0.86 | 3.45E-03 |
| GPCR downstream signalling (R-MMU-388396) | 1270 | 12 | 40.5 | 0.3 | 3.48E-05 |
| Signaling by GPCR (R-MMU-372790) | 1294 | 12 | 41.26 | 0.29 | 2.94E-05 |
| G alpha (s) signalling events (R-MMU-418555) | 720 | 2 | 22.96 | 0.09 | 1.64E-05 |
| Olfactory Signaling Pathway (R-MMU-381753) | 585 | 0 | 18.66 | < 0.01 | 5.23E-06 |

| **Green module** | **Mus musculus genes (22296)** | **Module genes (711)** | **Module genes (expected)** | **Module genes fold Enrichment** | **Module genes FDR** |
|---|---|---|---|---|---|
| Resolution of AP sites via the single-nucleotide replacement pathway (R-MMU-110381) | 4 | 3 | 0.23 | 12.91 | 3.71E-02 |
| SLBP Dependent Processing of Replication-Dependent Histone Pre-mRNAs (R-MMU-77588) | 9 | 5 | 0.52 | 9.56 | 6.39E-03 |
| Receptor Mediated Mitophagy (R-MMU-8934903) | 11 | 6 | 0.64 | 9.39 | 2.32E-03 |
| Formation of ATP by chemiosmotic coupling (R-MMU-163210) | 19 | 10 | 1.1 | 9.06 | 4.80E-05 |
| Cristae formation (R-MMU-8949613) | 19 | 10 | 1.1 | 9.06 | 4.72E-05 |
| SLBP independent Processing of Histone Pre-mRNAs (R-MMU-111367) | 8 | 4 | 0.46 | 8.61 | 2.52E-02 |
| Complex I biogenesis (R-MMU-6799198) | 55 | 26 | 3.19 | 8.14 | 8.70E-12 |

| | | | | | |
|---|---|---|---|---|---|
| Respiratory electron transport, ATP synthesis by chemiosmotic coupling, and heat production by uncoupling proteins. (R-MMU-163200) | 85 | 40 | 4.94 | 8.1 | 1.01E-17 |
| Respiratory electron transport (R-MMU-611105) | 61 | 28 | 3.54 | 7.9 | 1.98E-12 |
| Mitochondrial biogenesis (R-MMU-1592230) | 27 | 12 | 1.57 | 7.65 | 2.40E-05 |
| Pink/Parkin Mediated Mitophagy (R-MMU-5205685) | 17 | 7 | 0.99 | 7.09 | 2.58E-03 |
| Mitochondrial translation elongation (R-MMU-5389840) | 83 | 33 | 4.82 | 6.85 | 4.69E-13 |
| Mitochondrial translation termination (R-MMU-5419276) | 85 | 33 | 4.94 | 6.68 | 6.33E-13 |
| Mitochondrial translation (R-MMU-5368287) | 86 | 33 | 5 | 6.61 | 6.80E-13 |
| The citric acid (TCA) cycle and respiratory electron transport (R-MMU-1428517) | 136 | 52 | 7.9 | 6.58 | 5.63E-20 |
| Mitophagy (R-MMU-5205647) | 24 | 9 | 1.39 | 6.46 | 6.89E-04 |
| Glycogen synthesis (R-MMU-3322077) | 15 | 5 | 0.87 | 5.74 | 3.02E-02 |
| Endosomal Sorting Complex Required For Transport (ESCRT) (R-MMU-917729) | 30 | 10 | 1.74 | 5.74 | 6.19E-04 |
| mTORC1-mediated signalling (R-MMU-166208) | 21 | 7 | 1.22 | 5.74 | 6.22E-03 |
| Regulation of RUNX2 expression and activity (R-MMU-8939902) | 50 | 16 | 2.9 | 5.51 | 1.58E-05 |
| Translation (R-MMU-72766) | 163 | 52 | 9.47 | 5.49 | 1.67E-17 |
| Citric acid cycle (TCA cycle) (R-MMU-71403) | 22 | 7 | 1.28 | 5.48 | 7.50E-03 |

| | | | | | |
|---|---|---|---|---|---|
| Autodegradation of Cdh1 by Cdh1:APC/C (R-MMU-174084) | 63 | 20 | 3.66 | 5.47 | 1.16E-06 |
| mRNA decay by 3' to 5' exoribonuclease (R-MMU-429958) | 16 | 5 | 0.93 | 5.38 | 3.69E-02 |
| AUF1 (hnRNP D0) binds and destabilizes mRNA (R-MMU-450408) | 55 | 17 | 3.19 | 5.32 | 1.16E-05 |
| Nonsense Mediated Decay (NMD) enhanced by the Exon Junction Complex (EJC) (R-MMU-975957) | 94 | 29 | 5.46 | 5.31 | 1.57E-09 |
| Nonsense-Mediated Decay (NMD) (R-MMU-927802) | 94 | 29 | 5.46 | 5.31 | 1.49E-09 |
| Translesion Synthesis by POLH (R-MMU-110320) | 20 | 6 | 1.16 | 5.17 | 2.03E-02 |
| Ubiquitin-dependent degradation of Cyclin D1 (R-MMU-69229) | 50 | 15 | 2.9 | 5.17 | 4.76E-05 |
| Ubiquitin-dependent degradation of Cyclin D (R-MMU-75815) | 50 | 15 | 2.9 | 5.17 | 4.68E-05 |
| APC/C:Cdc20 mediated degradation of Securin (R-MMU-174154) | 67 | 20 | 3.89 | 5.14 | 2.51E-06 |
| FBXL7 down-regulates AURKA during mitotic entry and in early mitosis (R-MMU-8854050) | 54 | 16 | 3.14 | 5.1 | 3.11E-05 |
| Autodegradation of the E3 ubiquitin ligase COP1 (R-MMU-349425) | 51 | 15 | 2.96 | 5.06 | 5.16E-05 |
| p53-Independent G1/S DNA damage checkpoint (R-MMU-69613) | 51 | 15 | 2.96 | 5.06 | 5.09E-05 |
| p53-Independent DNA Damage Response (R-MMU-69610) | 51 | 15 | 2.96 | 5.06 | 5.01E-05 |
| Ubiquitin Mediated Degradation of Phosphorylated Cdc25A (R-MMU-69601) | 51 | 15 | 2.96 | 5.06 | 4.94E-05 |
| APC/C:Cdc20 mediated degradation of Cyclin B (R-MMU-174048) | 24 | 7 | 1.39 | 5.02 | 1.07E-02 |

| | | | | | |
|---|---|---|---|---|---|
| CDK-mediated phosphorylation and removal of Cdc6 (R-MMU-69017) | 72 | 21 | 4.18 | 5.02 | 1.68E-06 |
| Detoxification of Reactive Oxygen Species (R-MMU-3299685) | 35 | 10 | 2.03 | 4.92 | 1.62E-03 |
| Degradation of GLI1 by the proteasome (R-MMU-5610780) | 56 | 16 | 3.25 | 4.92 | 4.13E-05 |
| Hedgehog ligand biogenesis (R-MMU-5358346) | 63 | 18 | 3.66 | 4.92 | 1.30E-05 |
| NoRC negatively regulates rRNA expression (R-MMU-427413) | 21 | 6 | 1.22 | 4.92 | 2.39E-02 |
| Degradation of DVL (R-MMU-4641258) | 56 | 16 | 3.25 | 4.92 | 4.06E-05 |
| mRNA Capping (R-MMU-72086) | 28 | 8 | 1.63 | 4.92 | 6.10E-03 |
| Regulation of RUNX3 expression and activity (R-MMU-8941858) | 53 | 15 | 3.08 | 4.87 | 6.88E-05 |
| Regulation of ornithine decarboxylase (ODC) (R-MMU-350562) | 50 | 14 | 2.9 | 4.82 | 1.35E-04 |
| Cdc20:Phospho-APC/C mediated degradation of Cyclin A (R-MMU-174184) | 72 | 20 | 4.18 | 4.78 | 5.82E-06 |
| APC/C:Cdh1 mediated degradation of Cdc20 and other APC/C:Cdh1 targeted proteins in late mitosis/early G1 (R-MMU-174178) | 72 | 20 | 4.18 | 4.78 | 5.62E-06 |
| Degradation of AXIN (R-MMU-4641257) | 54 | 15 | 3.14 | 4.78 | 8.04E-05 |
| SCF-beta-TrCP mediated degradation of Emi1 (R-MMU-174113) | 54 | 15 | 3.14 | 4.78 | 7.94E-05 |
| Oxygen-dependent proline hydroxylation of Hypoxia-inducible Factor Alpha (R-MMU-1234176) | 65 | 18 | 3.78 | 4.77 | 1.78E-05 |

| | | | | | |
|---|---|---|---|---|---|
| GLI3 is processed to GLI3R by the proteasome (R-MMU-5610785) | 58 | 16 | 3.37 | 4.75 | 5.13E-05 |
| NIK-->noncanonical NF-kB signaling (R-MMU-5676590) | 58 | 16 | 3.37 | 4.75 | 5.06E-05 |
| Energy dependent regulation of mTOR by LKB1-AMPK (R-MMU-380972) | 29 | 8 | 1.68 | 4.75 | 7.19E-03 |
| CDT1 association with the CDC6:ORC:origin complex (R-MMU-68827) | 58 | 16 | 3.37 | 4.75 | 4.98E-05 |
| Dectin-1 mediated noncanonical NF-kB signaling (R-MMU-5607761) | 58 | 16 | 3.37 | 4.75 | 4.90E-05 |
| APC:Cdc20 mediated degradation of cell cycle proteins prior to satisfation of the cell cycle checkpoint (R-MMU-179419) | 73 | 20 | 4.24 | 4.72 | 6.57E-06 |
| Stabilization of p53 (R-MMU-69541) | 55 | 15 | 3.19 | 4.7 | 9.03E-05 |
| SCF(Skp2)-mediated degradation of p27/p21 (R-MMU-187577) | 59 | 16 | 3.43 | 4.67 | 5.51E-05 |
| mRNA Splicing - Minor Pathway (R-MMU-72165) | 48 | 13 | 2.79 | 4.66 | 3.14E-04 |
| Cross-presentation of soluble exogenous antigens (endosomes) (R-MMU-1236978) | 48 | 13 | 2.79 | 4.66 | 3.11E-04 |
| APC-Cdc20 mediated degradation of Nek2A (R-MMU-179409) | 26 | 7 | 1.51 | 4.64 | 1.52E-02 |
| APC/C:Cdc20 mediated degradation of mitotic proteins (R-MMU-176409) | 75 | 20 | 4.36 | 4.59 | 8.92E-06 |
| Activation of APC/C and APC/C:Cdc20 mediated degradation of mitotic proteins (R-MMU-176814) | 76 | 20 | 4.41 | 4.53 | 9.77E-06 |

| | | | | | |
|---|---|---|---|---|---|
| Transcriptional regulation by RUNX2 (R-MMU-8878166) | 61 | 16 | 3.54 | 4.52 | 7.43E-05 |
| RNA Polymerase I Promoter Opening (R-MMU-73728) | 23 | 6 | 1.34 | 4.49 | 3.36E-02 |
| Cellular response to hypoxia (R-MMU-2262749) | 70 | 18 | 4.07 | 4.43 | 3.57E-05 |
| Regulation of Hypoxia-inducible Factor (HIF) by oxygen (R-MMU-1234174) | 70 | 18 | 4.07 | 4.43 | 3.50E-05 |
| Switching of origins to a post-replicative state (R-MMU-69052) | 90 | 23 | 5.23 | 4.4 | 2.71E-06 |
| p53-Dependent G1/S DNA damage checkpoint (R-MMU-69580) | 63 | 16 | 3.66 | 4.37 | 9.46E-05 |
| p53-Dependent G1 DNA Damage Response (R-MMU-69563) | 63 | 16 | 3.66 | 4.37 | 9.35E-05 |
| Metabolism of polyamines (R-MMU-351202) | 83 | 21 | 4.82 | 4.36 | 8.83E-06 |
| Negative epigenetic regulation of rRNA expression (R-MMU-5250941) | 24 | 6 | 1.39 | 4.3 | 3.87E-02 |
| Cap-dependent Translation Initiation (R-MMU-72737) | 68 | 17 | 3.95 | 4.3 | 6.73E-05 |
| Eukaryotic Translation Initiation (R-MMU-72613) | 68 | 17 | 3.95 | 4.3 | 6.64E-05 |
| Regulation of mRNA stability by proteins that bind AU-rich elements (R-MMU-450531) | 81 | 20 | 4.7 | 4.25 | 1.85E-05 |
| G1/S DNA Damage Checkpoints (R-MMU-69615) | 65 | 16 | 3.78 | 4.24 | 1.28E-04 |
| Activation of NF-kappaB in B cells (R-MMU-1169091) | 65 | 16 | 3.78 | 4.24 | 1.27E-04 |
| Asymmetric localization of PCP proteins (R-MMU-4608870) | 61 | 15 | 3.54 | 4.23 | 2.13E-04 |

| | | | | | |
|---|---|---|---|---|---|
| TP53 Regulates Metabolic Genes (R-MMU-5628897) | 49 | 12 | 2.85 | 4.22 | 1.28E-03 |
| UCH proteinases (R-MMU-5689603) | 86 | 21 | 5 | 4.2 | 1.28E-05 |
| Regulation of APC/C activators between G1/S and early anaphase (R-MMU-176408) | 82 | 20 | 4.76 | 4.2 | 2.09E-05 |
| Orc1 removal from chromatin (R-MMU-68949) | 70 | 17 | 4.07 | 4.18 | 8.28E-05 |
| Macroautophagy (R-MMU-1632852) | 66 | 16 | 3.83 | 4.17 | 1.46E-04 |
| E3 ubiquitin ligases ubiquitinate target proteins (R-MMU-8866654) | 33 | 8 | 1.92 | 4.17 | 1.31E-02 |
| Formation of the ternary complex, and subsequently, the 43S complex (R-MMU-72695) | 58 | 14 | 3.37 | 4.16 | 4.38E-04 |
| Glycogen metabolism (R-MMU-8982491) | 25 | 6 | 1.45 | 4.13 | 4.50E-02 |
| Processing of Capped Intronless Pre-mRNA (R-MMU-75067) | 25 | 6 | 1.45 | 4.13 | 4.48E-02 |
| FGFR2 alternative splicing (R-MMU-6803529) | 25 | 6 | 1.45 | 4.13 | 4.46E-02 |
| RNA Polymerase II Transcription Initiation And Promoter Clearance (R-MMU-76042) | 46 | 11 | 2.67 | 4.12 | 2.66E-03 |
| RNA Polymerase II Transcription Initiation (R-MMU-75953) | 46 | 11 | 2.67 | 4.12 | 2.64E-03 |
| RNA Polymerase II Transcription Pre-Initiation And Promoter Opening (R-MMU-73779) | 46 | 11 | 2.67 | 4.12 | 2.63E-03 |
| RNA Polymerase II Promoter Escape (R-MMU-73776) | 46 | 11 | 2.67 | 4.12 | 2.61E-03 |
| Assembly of the pre-replicative complex (R-MMU-68867) | 67 | 16 | 3.89 | 4.11 | 1.65E-04 |

| | | | | | |
|---|---|---|---|---|---|
| RUNX1 regulates transcription of genes involved in differentiation of HSCs (R-MMU-8939236) | 67 | 16 | 3.89 | 4.11 | 1.63E-04 |
| Ribosomal scanning and start codon recognition (R-MMU-72702) | 63 | 15 | 3.66 | 4.1 | 2.82E-04 |
| Regulation of PTEN stability and activity (R-MMU-8948751) | 68 | 16 | 3.95 | 4.05 | 1.88E-04 |
| Pyruvate metabolism and Citric Acid (TCA) cycle (R-MMU-71406) | 51 | 12 | 2.96 | 4.05 | 1.69E-03 |
| Regulation of RAS by GAPs (R-MMU-5658442) | 68 | 16 | 3.95 | 4.05 | 1.86E-04 |
| Downstream signaling events of B Cell Receptor (BCR) (R-MMU-1168372) | 77 | 18 | 4.47 | 4.02 | 7.45E-05 |
| Synthesis of active ubiquitin: roles of E1 and E2 enzymes (R-MMU-8866652) | 30 | 7 | 1.74 | 4.02 | 2.84E-02 |
| MAPK6/MAPK4 signaling (R-MMU-5687128) | 73 | 17 | 4.24 | 4.01 | 1.28E-04 |
| Cyclin E associated events during G1/S transition (R-MMU-69202) | 69 | 16 | 4.01 | 3.99 | 2.15E-04 |
| Regulation of mitotic cell cycle (R-MMU-453276) | 88 | 20 | 5.11 | 3.91 | 4.42E-05 |
| APC/C-mediated degradation of cell cycle proteins (R-MMU-174143) | 88 | 20 | 5.11 | 3.91 | 4.35E-05 |
| CLEC7A (Dectin-1) signaling (R-MMU-5607764) | 88 | 20 | 5.11 | 3.91 | 4.27E-05 |
| RNA Polymerase II Transcription Termination (R-MMU-73856) | 62 | 14 | 3.6 | 3.89 | 7.81E-04 |
| Cleavage of Growing Transcript in the Termination Region (R-MMU-109688) | 62 | 14 | 3.6 | 3.89 | 7.74E-04 |

| | | | | | |
|---|---|---|---|---|---|
| Formation of the Early Elongation Complex (R-MMU-113418) | 31 | 7 | 1.8 | 3.89 | 3.27E-02 |
| Cyclin A:Cdk2-associated events at S phase entry (R-MMU-69656) | 71 | 16 | 4.12 | 3.88 | 2.73E-04 |
| The role of GTSE1 in G2/M progression after G2 checkpoint (R-MMU-8852276) | 72 | 16 | 4.18 | 3.83 | 3.12E-04 |
| Recognition of DNA damage by PCNA-containing replication complex (R-MMU-110314) | 32 | 7 | 1.86 | 3.77 | 3.69E-02 |
| Synthesis of DNA (R-MMU-69239) | 119 | 26 | 6.91 | 3.76 | 5.14E-06 |
| DNA Replication Pre-Initiation (R-MMU-69002) | 83 | 18 | 4.82 | 3.73 | 1.52E-04 |
| M/G1 Transition (R-MMU-68874) | 83 | 18 | 4.82 | 3.73 | 1.50E-04 |
| DNA Damage Recognition in GG-NER (R-MMU-5696394) | 37 | 8 | 2.15 | 3.72 | 2.30E-02 |
| Degradation of beta-catenin by the destruction complex (R-MMU-195253) | 79 | 17 | 4.59 | 3.7 | 2.59E-04 |
| Deadenylation-dependent mRNA decay (R-MMU-429914) | 42 | 9 | 2.44 | 3.69 | 1.42E-02 |
| Formation of TC-NER Pre-Incision Complex (R-MMU-6781823) | 52 | 11 | 3.02 | 3.64 | 5.87E-03 |
| Interleukin-1 signaling (R-MMU-9020702) | 95 | 20 | 5.52 | 3.62 | 8.30E-05 |
| mTOR signalling (R-MMU-165159) | 38 | 8 | 2.21 | 3.62 | 2.61E-02 |
| RNA Polymerase II Pre-transcription Events (R-MMU-674695) | 78 | 16 | 4.53 | 3.53 | 6.70E-04 |
| Formation of Incision Complex in GG-NER (R-MMU-5696395) | 39 | 8 | 2.27 | 3.53 | 2.97E-02 |
| DNA Replication (R-MMU-69306) | 127 | 26 | 7.38 | 3.52 | 1.13E-05 |

| | | | | | |
|---|---|---|---|---|---|
| Metabolism of RNA (R-MMU-8953854) | 500 | 102 | 29.04 | 3.51 | 1.74E-21 |
| Protein ubiquitination (R-MMU-8852135) | 54 | 11 | 3.14 | 3.51 | 7.26E-03 |
| Gap-filling DNA repair synthesis and ligation in TC-NER (R-MMU-6782210) | 64 | 13 | 3.72 | 3.5 | 2.95E-03 |
| PTEN Regulation (R-MMU-6807070) | 114 | 23 | 6.62 | 3.47 | 4.26E-05 |
| Translesion synthesis by Y family DNA polymerases bypasses lesions on DNA template (R-MMU-110313) | 40 | 8 | 2.32 | 3.44 | 3.34E-02 |
| DNA Damage Bypass (R-MMU-73893) | 50 | 10 | 2.9 | 3.44 | 1.27E-02 |
| mRNA 3'-end processing (R-MMU-72187) | 55 | 11 | 3.19 | 3.44 | 8.01E-03 |
| Senescence-Associated Secretory Phenotype (SASP) (R-MMU-2559582) | 61 | 12 | 3.54 | 3.39 | 5.89E-03 |
| Transcription-Coupled Nucleotide Excision Repair (TC-NER) (R-MMU-6781827) | 77 | 15 | 4.47 | 3.35 | 1.69E-03 |
| G1/S Transition (R-MMU-69206) | 104 | 20 | 6.04 | 3.31 | 2.15E-04 |
| Nucleotide Excision Repair (R-MMU-5696398) | 104 | 20 | 6.04 | 3.31 | 2.13E-04 |
| Interleukin-1 family signaling (R-MMU-446652) | 111 | 21 | 6.45 | 3.26 | 1.71E-04 |
| Downstream TCR signaling (R-MMU-202424) | 90 | 17 | 5.23 | 3.25 | 9.25E-04 |
| mRNA Splicing (R-MMU-72172) | 177 | 33 | 10.28 | 3.21 | 2.97E-06 |
| Transcriptional regulation by RUNX3 (R-MMU-8878159) | 81 | 15 | 4.7 | 3.19 | 2.59E-03 |
| C-type lectin receptors (CLRs) (R-MMU-5621481) | 108 | 20 | 6.27 | 3.19 | 3.14E-04 |

| | | | | | |
|---|---|---|---|---|---|
| Dual incision in TC-NER (R-MMU-6782135) | 65 | 12 | 3.78 | 3.18 | 8.61E-03 |
| PCP/CE pathway (R-MMU-4086400) | 87 | 16 | 5.05 | 3.17 | 1.82E-03 |
| mRNA Splicing - Major Pathway (R-MMU-72163) | 170 | 31 | 9.87 | 3.14 | 9.09E-06 |
| Global Genome Nucleotide Excision Repair (GG-NER) (R-MMU-5696399) | 77 | 14 | 4.47 | 3.13 | 4.50E-03 |
| Formation of RNA Pol II elongation complex (R-MMU-112382) | 55 | 10 | 3.19 | 3.13 | 2.26E-02 |
| RNA Polymerase II Transcription Elongation (R-MMU-75955) | 55 | 10 | 3.19 | 3.13 | 2.25E-02 |
| S Phase (R-MMU-69242) | 147 | 26 | 8.54 | 3.05 | 6.41E-05 |
| Peroxisomal protein import (R-MMU-9033241) | 68 | 12 | 3.95 | 3.04 | 1.17E-02 |
| Hedgehog 'on' state (R-MMU-5632684) | 103 | 18 | 5.98 | 3.01 | 1.34E-03 |
| Processing of Capped Intron-Containing Pre-mRNA (R-MMU-72203) | 231 | 40 | 13.42 | 2.98 | 8.96E-07 |
| Cellular responses to external stimuli (R-MMU-8953897) | 416 | 72 | 24.16 | 2.98 | 1.97E-12 |
| Major pathway of rRNA processing in the nucleolus and cytosol (R-MMU-6791226) | 76 | 13 | 4.41 | 2.95 | 9.97E-03 |
| rRNA processing in the nucleus and cytosol (R-MMU-8868773) | 76 | 13 | 4.41 | 2.95 | 9.92E-03 |
| rRNA processing (R-MMU-72312) | 76 | 13 | 4.41 | 2.95 | 9.86E-03 |
| Antigen processing-Cross presentation (R-MMU-1236975) | 88 | 15 | 5.11 | 2.93 | 5.11E-03 |
| Mitotic Anaphase (R-MMU-68882) | 188 | 32 | 10.92 | 2.93 | 1.78E-05 |
| Mitotic Metaphase and Anaphase (R-MMU-2555396) | 189 | 32 | 10.98 | 2.92 | 1.84E-05 |

| | | | | | |
|---|---|---|---|---|---|
| Separation of Sister Chromatids (R-MMU-2467813) | 185 | 31 | 10.75 | 2.89 | 3.08E-05 |
| TNFR2 non-canonical NF-kB pathway (R-MMU-5668541) | 98 | 16 | 5.69 | 2.81 | 5.10E-03 |
| RNA polymerase II transcribes snRNA genes (R-MMU-6807505) | 74 | 12 | 4.3 | 2.79 | 2.11E-02 |
| RAB geranylgeranylation (R-MMU-8873719) | 62 | 10 | 3.6 | 2.78 | 4.31E-02 |
| Signaling by FGFR2 (R-MMU-5654738) | 75 | 12 | 4.36 | 2.75 | 2.29E-02 |
| TCR signaling (R-MMU-202403) | 107 | 17 | 6.21 | 2.74 | 4.65E-03 |
| ABC-family proteins mediated transport (R-MMU-382556) | 101 | 16 | 5.87 | 2.73 | 6.45E-03 |
| Cellular responses to stress (R-MMU-2262752) | 354 | 56 | 20.56 | 2.72 | 3.01E-08 |
| Hedgehog 'off' state (R-MMU-5610787) | 109 | 17 | 6.33 | 2.69 | 5.47E-03 |
| Beta-catenin independent WNT signaling (R-MMU-3858494) | 123 | 19 | 7.14 | 2.66 | 3.17E-03 |
| COPI-mediated anterograde transport (R-MMU-6807878) | 98 | 15 | 5.69 | 2.64 | 1.15E-02 |
| G2/M Checkpoints (R-MMU-69481) | 139 | 21 | 8.07 | 2.6 | 3.30E-03 |
| Gene Silencing by RNA (R-MMU-211000) | 73 | 11 | 4.24 | 2.59 | 4.46E-02 |
| Mitotic G1-G1/S phases (R-MMU-453279) | 133 | 20 | 7.72 | 2.59 | 4.68E-03 |
| Transcriptional regulation by RUNX1 (R-MMU-8878171) | 167 | 25 | 9.7 | 2.58 | 9.32E-04 |
| FCERI mediated NF-kB activation (R-MMU-2871837) | 114 | 17 | 6.62 | 2.57 | 7.78E-03 |
| COPII-mediated vesicle transport (R-MMU-204005) | 74 | 11 | 4.3 | 2.56 | 4.85E-02 |

| | | | | | |
|---|---|---|---|---|---|
| ER to Golgi Anterograde Transport (R-MMU-199977) | 155 | 23 | 9 | 2.55 | 1.90E-03 |
| Signaling by FGFR (R-MMU-190236) | 82 | 12 | 4.76 | 2.52 | 3.96E-02 |
| G2/M Transition (R-MMU-69275) | 177 | 25 | 10.28 | 2.43 | 1.66E-03 |
| Cell Cycle Checkpoints (R-MMU-69620) | 262 | 37 | 15.22 | 2.43 | 8.34E-05 |
| Metabolism of amino acids and derivatives (R-MMU-71291) | 249 | 35 | 14.46 | 2.42 | 1.48E-04 |
| Signaling by Hedgehog (R-MMU-5358351) | 143 | 20 | 8.31 | 2.41 | 7.10E-03 |
| Mitotic G2-G2/M phases (R-MMU-453274) | 179 | 25 | 10.4 | 2.4 | 1.85E-03 |
| Rab regulation of trafficking (R-MMU-9007101) | 115 | 16 | 6.68 | 2.4 | 2.31E-02 |
| Ub-specific processing proteases (R-MMU-5689880) | 169 | 23 | 9.82 | 2.34 | 6.00E-03 |
| Transport to the Golgi and subsequent modification (R-MMU-948021) | 185 | 25 | 10.75 | 2.33 | 3.47E-03 |
| Neddylation (R-MMU-8951664) | 246 | 33 | 14.29 | 2.31 | 4.35E-04 |
| Mitotic Spindle Checkpoint (R-MMU-69618) | 112 | 15 | 6.51 | 2.31 | 3.70E-02 |
| Signaling by the B Cell Receptor (BCR) (R-MMU-983705) | 145 | 19 | 8.42 | 2.26 | 2.23E-02 |
| TCF dependent signaling in response to WNT (R-MMU-201681) | 177 | 23 | 10.28 | 2.24 | 7.59E-03 |
| M Phase (R-MMU-68886) | 341 | 44 | 19.81 | 2.22 | 8.31E-05 |
| Transcriptional Regulation by TP53 (R-MMU-3700989) | 276 | 35 | 16.03 | 2.18 | 9.31E-04 |
| Asparagine N-linked glycosylation (R-MMU-446203) | 287 | 36 | 16.67 | 2.16 | 7.79E-04 |

| | | | | | |
|---|---|---|---|---|---|
| Organelle biogenesis and maintenance (R-MMU-1852241) | 211 | 26 | 12.26 | 2.12 | 7.47E-03 |
| Fc epsilon receptor (FCERI) signaling (R-MMU-2454202) | 158 | 19 | 9.18 | 2.07 | 3.49E-02 |
| PIP3 activates AKT signaling (R-MMU-1257604) | 228 | 27 | 13.24 | 2.04 | 1.07E-02 |
| Deubiquitination (R-MMU-5688426) | 237 | 28 | 13.77 | 2.03 | 8.53E-03 |
| Metabolism of proteins (R-MMU-392499) | 1720 | 203 | 99.9 | 2.03 | 1.67E-17 |
| Antigen processing: Ubiquitination & Proteasome degradation (R-MMU-983168) | 315 | 37 | 18.3 | 2.02 | 2.37E-03 |
| Signaling by WNT (R-MMU-195721) | 257 | 30 | 14.93 | 2.01 | 8.39E-03 |
| DNA Repair (R-MMU-73894) | 271 | 31 | 15.74 | 1.97 | 7.57E-03 |
| Membrane Trafficking (R-MMU-199991) | 570 | 65 | 33.11 | 1.96 | 4.17E-05 |
| Cell Cycle, Mitotic (R-MMU-69278) | 472 | 53 | 27.41 | 1.93 | 3.43E-04 |
| Intracellular signaling by second messengers (R-MMU-9006925) | 253 | 28 | 14.69 | 1.91 | 1.87E-02 |
| RNA Polymerase II Transcription (R-MMU-73857) | 851 | 91 | 49.43 | 1.84 | 8.45E-06 |
| Class I MHC mediated antigen processing & presentation (R-MMU-983169) | 370 | 39 | 21.49 | 1.81 | 8.21E-03 |
| Neutrophil degranulation (R-MMU-6798695) | 553 | 58 | 32.12 | 1.81 | 6.28E-04 |
| Vesicle-mediated transport (R-MMU-5653656) | 646 | 67 | 37.52 | 1.79 | 2.66E-04 |
| Metabolism (R-MMU-1430728) | 1818 | 185 | 105.59 | 1.75 | 9.81E-11 |
| Gene expression (Transcription) (R-MMU-74160) | 961 | 96 | 55.82 | 1.72 | 3.60E-05 |
| Cell Cycle (R-MMU-1640170) | 541 | 54 | 31.42 | 1.72 | 3.59E-03 |

| | | | | | |
|---|---|---|---|---|---|
| Generic Transcription Pathway (R-MMU-212436) | 733 | 72 | 42.57 | 1.69 | 6.94E-04 |
| Post-translational protein modification (R-MMU-597592) | 1325 | 125 | 76.96 | 1.62 | 1.83E-05 |
| Innate Immune System (R-MMU-168249) | 1039 | 90 | 60.35 | 1.49 | 3.91E-03 |
| Unclassified (UNCLASSIFIED) | 12915 | 622 | 750.13 | 0.83 | 3.38E-10 |
| Signal Transduction (R-MMU-162582) | 2644 | 109 | 153.57 | 0.71 | 1.55E-03 |
| Class A/1 (Rhodopsin-like receptors) (R-MMU-373076) | 309 | 7 | 17.95 | 0.39 | 4.48E-02 |
| GPCR ligand binding (R-MMU-500792) | 415 | 8 | 24.1 | 0.33 | 3.58E-03 |
| Developmental Biology (R-MMU-1266738) | 466 | 7 | 27.07 | 0.26 | 2.14E-04 |
| Signaling by GPCR (R-MMU-372790) | 1294 | 18 | 75.16 | 0.24 | 8.26E-13 |
| GPCR downstream signalling (R-MMU-388396) | 1270 | 17 | 73.76 | 0.23 | 7.11E-13 |
| G alpha (s) signalling events (R-MMU-418555) | 720 | 1 | 41.82 | 0.02 | 1.98E-14 |
| Formation of the cornified envelope (R-MMU-6809371) | 105 | 0 | 6.1 | < 0.01 | 3.42E-02 |
| Keratinization (R-MMU-6805567) | 140 | 0 | 8.13 | < 0.01 | 5.99E-03 |
| Olfactory Signaling Pathway (R-MMU-381753) | 585 | 0 | 33.98 | < 0.01 | 7.37E-13 |

| **Blue module** | **Mus musculus genes (22296)** | **Module genes (711)** | **Module genes (expected)** | **Module genes fold Enrichment** | **Module genes FDR** |
|---|---|---|---|---|---|
| Regulation of lipid metabolism by Peroxisome proliferator-activated receptor alpha (PPARalpha) (R-MMU-400206) | 20 | 10 | 1.9 | 5.28 | 1.71E-02 |

| | | | | | |
|---|---|---|---|---|---|
| RUNX1 interacts with co-factors whose precise effect on RUNX1 targets is not known (R-MMU-8939243) | 37 | 14 | 3.51 | 3.99 | 1.23E-02 |
| PKMTs methylate histone lysines (R-MMU-3214841) | 48 | 15 | 4.55 | 3.3 | 2.65E-02 |
| Protein-protein interactions at synapses (R-MMU-6794362) | 67 | 18 | 6.35 | 2.83 | 3.03E-02 |
| Chromatin modifying enzymes (R-MMU-3247509) | 183 | 49 | 17.34 | 2.83 | 7.15E-06 |
| Chromatin organization (R-MMU-4839726) | 183 | 49 | 17.34 | 2.83 | 3.58E-06 |
| Cilium Assembly (R-MMU-5617833) | 184 | 36 | 17.44 | 2.06 | 2.76E-02 |
| Axon guidance (R-MMU-422475) | 277 | 50 | 26.25 | 1.9 | 1.37E-02 |
| Neuronal System (R-MMU-112316) | 338 | 55 | 32.03 | 1.72 | 3.66E-02 |
| Developmental Biology (R-MMU-1266738) | 466 | 75 | 44.16 | 1.7 | 1.03E-02 |
| Generic Transcription Pathway (R-MMU-212436) | 733 | 113 | 69.47 | 1.63 | 1.17E-03 |
| Signaling by Receptor Tyrosine Kinases (R-MMU-9006934) | 403 | 62 | 38.19 | 1.62 | 4.92E-02 |
| Gene expression (Transcription) (R-MMU-74160) | 961 | 144 | 91.07 | 1.58 | 3.04E-04 |
| RNA Polymerase II Transcription (R-MMU-73857) | 851 | 127 | 80.65 | 1.57 | 1.09E-03 |
| Immune System (R-MMU-168256) | 1769 | 123 | 167.65 | 0.73 | 3.61E-02 |
| Innate Immune System (R-MMU-168249) | 1039 | 60 | 98.47 | 0.61 | 1.04E-02 |
| G alpha (s) signalling events (R-MMU-418555) | 720 | 36 | 68.23 | 0.53 | 9.83E-03 |
| Neutrophil degranulation (R-MMU-6798695) | 553 | 26 | 52.41 | 0.5 | 1.70E-02 |

| | | | | | |
|---|---|---|---|---|---|
| Olfactory Signaling Pathway (R-MMU-381753) | 585 | 22 | 55.44 | 0.4 | 4.57E-04 |
| Metabolism of amino acids and derivatives (R-MMU-71291) | 249 | 8 | 23.6 | 0.34 | 4.29E-02 |

| **Purple module** | **Mus musculus genes (22296)** | **Module genes (711)** | **Module genes (expected)** | **Module genes fold Enrichment** | **Module genes FDR** |
|---|---|---|---|---|---|
| RHO GTPases activate CIT (R-MMU-5625900) | 8 | 4 | 0.22 | 18.46 | 4.63E-02 |
| RHO GTPases activate PKNs (R-MMU-5625740) | 42 | 7 | 1.14 | 6.15 | 4.55E-02 |
| Oxidative Stress Induced Senescence (R-MMU-2559580) | 70 | 9 | 1.9 | 4.75 | 4.16E-02 |
| Death Receptor Signalling (R-MMU-73887) | 129 | 12 | 3.49 | 3.43 | 4.58E-02 |
| Asparagine N-linked glycosylation (R-MMU-446203) | 287 | 20 | 7.77 | 2.57 | 4.20E-02 |
| Membrane Trafficking (R-MMU-199991) | 570 | 34 | 15.44 | 2.2 | 1.78E-02 |
| Vesicle-mediated transport (R-MMU-5653656) | 646 | 36 | 17.5 | 2.06 | 2.67E-02 |
| Post-translational protein modification (R-MMU-597592) | 1325 | 64 | 35.89 | 1.78 | 7.13E-03 |
| Unclassified (UNCLASSIFIED) | 12915 | 305 | 349.87 | 0.87 | 4.36E-02 |
| GPCR downstream signalling (R-MMU-388396) | 1270 | 15 | 34.4 | 0.44 | 4.45E-02 |
| Signaling by GPCR (R-MMU-372790) | 1294 | 15 | 35.05 | 0.43 | 4.69E-02 |
| G alpha (s) signalling events (R-MMU-418555) | 720 | 2 | 19.5 | 0.1 | 1.09E-03 |
| Olfactory Signaling Pathway (R-MMU-381753) | 585 | 0 | 15.85 | < 0.01 | 4.27E-04 |

Supplementary Table S6. GO enrichment scores for the nine WGCNA modules tested. If no entries are shown for a module, no GO terms were returned at all.

| Black module | Mus musculus genes (22296) | Module genes (711) | Module genes (expected) | Module genes fold Enrichment | Module genes FDR |
|---|---|---|---|---|---|
| regulation of nucleobase-containing compound transport (GO:0032239) | 16 | 5 | 0.51 | 9.8 | 3.25E-02 |
| maturation of LSU-rRNA from tricistronic rRNA transcript (SSU-rRNA, 5.8S rRNA, LSU-rRNA) (GO:0000463) | 17 | 5 | 0.54 | 9.22 | 3.86E-02 |
| cytoplasmic translational initiation (GO:0002183) | 18 | 5 | 0.57 | 8.71 | 4.67E-02 |
| histone monoubiquitination (GO:0010390) | 30 | 7 | 0.96 | 7.32 | 1.30E-02 |
| positive regulation of mRNA catabolic process (GO:0061014) | 49 | 10 | 1.56 | 6.4 | 1.81E-03 |
| protein monoubiquitination (GO:0006513) | 63 | 12 | 2.01 | 5.97 | 5.15E-04 |
| positive regulation of cell cycle G1/S phase transition (GO:1902808) | 49 | 9 | 1.56 | 5.76 | 8.34E-03 |
| positive regulation of G1/S transition of mitotic cell cycle (GO:1900087) | 39 | 7 | 1.24 | 5.63 | 3.94E-02 |
| histone ubiquitination (GO:0016574) | 41 | 7 | 1.31 | 5.35 | 4.94E-02 |
| positive regulation of mRNA metabolic process (GO:1903313) | 90 | 15 | 2.87 | 5.23 | 1.67E-04 |
| translational initiation (GO:0006413) | 55 | 9 | 1.75 | 5.13 | 1.48E-02 |
| positive regulation of telomere maintenance (GO:0032206) | 50 | 8 | 1.59 | 5.02 | 3.36E-02 |
| positive regulation of cell cycle phase transition (GO:1901989) | 94 | 14 | 3 | 4.67 | 9.62E-04 |
| regulation of mRNA processing (GO:0050684) | 144 | 20 | 4.59 | 4.36 | 4.57E-05 |
| positive regulation of mitotic cell cycle phase transition (GO:1901992) | 80 | 11 | 2.55 | 4.31 | 1.26E-02 |
| positive regulation of actin filament polymerization (GO:0030838) | 81 | 11 | 2.58 | 4.26 | 1.31E-02 |

| | | | | | |
|---|---|---|---|---|---|
| RNA splicing (GO:0008380) | 315 | 42 | 10.05 | 4.18 | 8.82E-11 |
| regulation of mRNA catabolic process (GO:0061013) | 120 | 16 | 3.83 | 4.18 | 7.86E-04 |
| regulation of mRNA metabolic process (GO:1903311) | 251 | 33 | 8 | 4.12 | 3.47E-08 |
| mRNA processing (GO:0006397) | 401 | 50 | 12.79 | 3.91 | 4.34E-12 |
| regulation of RNA stability (GO:0043487) | 105 | 13 | 3.35 | 3.88 | 9.02E-03 |
| regulation of mRNA stability (GO:0043488) | 97 | 12 | 3.09 | 3.88 | 1.42E-02 |
| positive regulation of chromosome organization (GO:2001252) | 180 | 22 | 5.74 | 3.83 | 7.54E-05 |
| rRNA processing (GO:0006364) | 192 | 23 | 6.12 | 3.76 | 6.09E-05 |
| negative regulation of translation (GO:0017148) | 126 | 15 | 4.02 | 3.73 | 4.38E-03 |
| mRNA splicing, via spliceosome (GO:0000398) | 195 | 23 | 6.22 | 3.7 | 7.59E-05 |
| RNA splicing, via transesterification reactions with bulged adenosine as nucleophile (GO:0000377) | 195 | 23 | 6.22 | 3.7 | 7.47E-05 |
| RNA splicing, via transesterification reactions (GO:0000375) | 195 | 23 | 6.22 | 3.7 | 7.37E-05 |
| rRNA metabolic process (GO:0016072) | 201 | 23 | 6.41 | 3.59 | 1.13E-04 |
| positive regulation of protein polymerization (GO:0032273) | 114 | 13 | 3.64 | 3.58 | 1.57E-02 |
| ribosome biogenesis (GO:0042254) | 284 | 32 | 9.06 | 3.53 | 1.55E-06 |
| mRNA metabolic process (GO:0016071) | 522 | 58 | 16.65 | 3.48 | 2.96E-12 |
| regulation of cell cycle G1/S phase transition (GO:1902806) | 126 | 14 | 4.02 | 3.48 | 1.26E-02 |
| positive regulation of mitotic cell cycle (GO:0045931) | 163 | 18 | 5.2 | 3.46 | 2.03E-03 |
| regulation of G1/S transition of mitotic cell cycle (GO:2000045) | 111 | 12 | 3.54 | 3.39 | 3.67E-02 |
| ribonucleoprotein complex biogenesis (GO:0022613) | 418 | 44 | 13.33 | 3.3 | 2.18E-08 |
| RNA processing (GO:0006396) | 742 | 78 | 23.66 | 3.3 | 1.31E-15 |
| mRNA catabolic process (GO:0006402) | 116 | 12 | 3.7 | 3.24 | 4.88E-02 |
| negative regulation of cellular amide metabolic process (GO:0034249) | 145 | 15 | 4.62 | 3.24 | 1.40E-02 |
| protein folding (GO:0006457) | 149 | 15 | 4.75 | 3.16 | 1.77E-02 |

| | | | | | |
|---|---|---|---|---|---|
| regulation of ubiquitin-dependent protein catabolic process (GO:2000058) | 149 | 15 | 4.75 | 3.16 | 1.76E-02 |
| protein stabilization (GO:0050821) | 169 | 17 | 5.39 | 3.15 | 8.49E-03 |
| RNA localization (GO:0006403) | 160 | 16 | 5.1 | 3.14 | 1.30E-02 |
| regulation of cell cycle phase transition (GO:1901987) | 284 | 27 | 9.06 | 2.98 | 2.90E-04 |
| DNA conformation change (GO:0071103) | 179 | 17 | 5.71 | 2.98 | 1.36E-02 |
| regulation of mitotic cell cycle phase transition (GO:1901990) | 252 | 23 | 8.04 | 2.86 | 2.36E-03 |
| regulation of proteolysis involved in cellular protein catabolic process (GO:1903050) | 209 | 19 | 6.66 | 2.85 | 1.10E-02 |
| RNA metabolic process (GO:0016070) | 1222 | 111 | 38.97 | 2.85 | 5.02E-18 |
| regulation of translation (GO:0006417) | 320 | 29 | 10.2 | 2.84 | 2.85E-04 |
| ubiquitin-dependent protein catabolic process (GO:0006511) | 472 | 42 | 15.05 | 2.79 | 3.76E-06 |
| nucleobase-containing compound catabolic process (GO:0034655) | 227 | 20 | 7.24 | 2.76 | 1.09E-02 |
| regulation of cellular protein catabolic process (GO:1903362) | 240 | 21 | 7.65 | 2.74 | 8.32E-03 |
| ribonucleoprotein complex assembly (GO:0022618) | 206 | 18 | 6.57 | 2.74 | 2.12E-02 |
| modification-dependent macromolecule catabolic process (GO:0043632) | 494 | 43 | 15.75 | 2.73 | 4.46E-06 |
| proteasome-mediated ubiquitin-dependent protein catabolic process (GO:0043161) | 288 | 25 | 9.18 | 2.72 | 2.28E-03 |
| modification-dependent protein catabolic process (GO:0019941) | 484 | 42 | 15.43 | 2.72 | 6.50E-06 |
| ribonucleoprotein complex subunit organization (GO:0071826) | 219 | 19 | 6.98 | 2.72 | 1.62E-02 |
| positive regulation of cell cycle process (GO:0090068) | 247 | 21 | 7.88 | 2.67 | 1.12E-02 |
| protein ubiquitination (GO:0016567) | 414 | 35 | 13.2 | 2.65 | 1.27E-04 |
| posttranscriptional regulation of gene expression (GO:0010608) | 414 | 35 | 13.2 | 2.65 | 1.26E-04 |
| aromatic compound catabolic process (GO:0019439) | 285 | 24 | 9.09 | 2.64 | 4.91E-03 |

| | | | | | |
|---|---|---|---|---|---|
| cellular nitrogen compound catabolic process (GO:0044270) | 265 | 22 | 8.45 | 2.6 | 1.09E-02 |
| protein modification by small protein conjugation (GO:0032446) | 461 | 38 | 14.7 | 2.58 | 1.08E-04 |
| heterocycle catabolic process (GO:0046700) | 271 | 22 | 8.64 | 2.55 | 1.31E-02 |
| protein modification by small protein conjugation or removal (GO:0070647) | 582 | 47 | 18.56 | 2.53 | 7.66E-06 |
| ncRNA metabolic process (GO:0034660) | 412 | 33 | 13.14 | 2.51 | 7.52E-04 |
| regulation of mitotic cell cycle (GO:0007346) | 487 | 39 | 15.53 | 2.51 | 1.18E-04 |
| gene expression (GO:0010467) | 1591 | 127 | 50.74 | 2.5 | 3.77E-17 |
| cellular macromolecule catabolic process (GO:0044265) | 740 | 59 | 23.6 | 2.5 | 3.16E-07 |
| proteolysis involved in cellular protein catabolic process (GO:0051603) | 552 | 44 | 17.6 | 2.5 | 2.78E-05 |
| proteasomal protein catabolic process (GO:0010498) | 316 | 25 | 10.08 | 2.48 | 1.04E-02 |
| cellular protein catabolic process (GO:0044257) | 570 | 45 | 18.18 | 2.48 | 2.56E-05 |
| regulation of cellular amide metabolic process (GO:0034248) | 368 | 29 | 11.74 | 2.47 | 3.10E-03 |
| nucleic acid metabolic process (GO:0090304) | 1740 | 136 | 55.49 | 2.45 | 1.29E-17 |
| ncRNA processing (GO:0034470) | 336 | 26 | 10.71 | 2.43 | 9.33E-03 |
| regulation of protein stability (GO:0031647) | 275 | 21 | 8.77 | 2.39 | 3.86E-02 |
| organic cyclic compound catabolic process (GO:1901361) | 315 | 24 | 10.05 | 2.39 | 1.76E-02 |
| positive regulation of organelle organization (GO:0010638) | 578 | 44 | 18.43 | 2.39 | 8.77E-05 |
| transcription, DNA-templated (GO:0006351) | 348 | 26 | 11.1 | 2.34 | 1.30E-02 |
| regulation of chromosome organization (GO:0033044) | 349 | 26 | 11.13 | 2.34 | 1.33E-02 |
| nucleic acid-templated transcription (GO:0097659) | 349 | 26 | 11.13 | 2.34 | 1.32E-02 |
| translation (GO:0006412) | 310 | 23 | 9.89 | 2.33 | 2.94E-02 |
| amide biosynthetic process (GO:0043604) | 421 | 31 | 13.43 | 2.31 | 5.59E-03 |
| RNA biosynthetic process (GO:0032774) | 354 | 26 | 11.29 | 2.3 | 2.14E-02 |
| peptide biosynthetic process (GO:0043043) | 329 | 24 | 10.49 | 2.29 | 3.81E-02 |

| | | | | | |
|---|---|---|---|---|---|
| cell division (GO:0051301) | 480 | 35 | 15.31 | 2.29 | 2.09E-03 |
| regulation of cell cycle process (GO:0010564) | 579 | 42 | 18.46 | 2.27 | 4.62E-04 |
| regulation of cellular catabolic process (GO:0031329) | 676 | 49 | 21.56 | 2.27 | 7.50E-05 |
| positive regulation of cell cycle (GO:0045787) | 346 | 25 | 11.03 | 2.27 | 3.09E-02 |
| macromolecule catabolic process (GO:0009057) | 833 | 60 | 26.56 | 2.26 | 5.17E-06 |
| protein catabolic process (GO:0030163) | 626 | 45 | 19.96 | 2.25 | 2.33E-04 |
| nucleobase-containing compound metabolic process (GO:0006139) | 2162 | 154 | 68.94 | 2.23 | 3.89E-17 |
| positive regulation of cellular catabolic process (GO:0031331) | 354 | 25 | 11.29 | 2.21 | 3.59E-02 |
| heterocycle metabolic process (GO:0046483) | 2301 | 160 | 73.38 | 2.18 | 5.95E-17 |
| regulation of protein catabolic process (GO:0042176) | 379 | 26 | 12.09 | 2.15 | 3.67E-02 |
| cell cycle (GO:0007049) | 1182 | 81 | 37.69 | 2.15 | 1.91E-07 |
| cellular aromatic compound metabolic process (GO:0006725) | 2369 | 162 | 75.55 | 2.14 | 1.05E-16 |
| chromosome organization (GO:0051276) | 914 | 62 | 29.15 | 2.13 | 2.03E-05 |
| mitotic cell cycle (GO:0000278) | 509 | 34 | 16.23 | 2.09 | 1.31E-02 |
| cellular nitrogen compound biosynthetic process (GO:0044271) | 1108 | 74 | 35.33 | 2.09 | 2.39E-06 |
| nucleobase-containing compound biosynthetic process (GO:0034654) | 646 | 43 | 20.6 | 2.09 | 2.18E-03 |
| protein localization to organelle (GO:0033365) | 601 | 40 | 19.17 | 2.09 | 4.88E-03 |
| cellular nitrogen compound metabolic process (GO:0034641) | 2757 | 183 | 87.92 | 2.08 | 1.15E-17 |
| regulation of catabolic process (GO:0009894) | 819 | 54 | 26.12 | 2.07 | 3.32E-04 |
| organonitrogen compound catabolic process (GO:1901565) | 879 | 57 | 28.03 | 2.03 | 2.87E-04 |
| heterocycle biosynthetic process (GO:0018130) | 712 | 46 | 22.71 | 2.03 | 2.12E-03 |
| organic cyclic compound metabolic process (GO:1901360) | 2580 | 166 | 82.27 | 2.02 | 9.52E-15 |
| cell cycle process (GO:0022402) | 772 | 49 | 24.62 | 1.99 | 1.82E-03 |

| | | | | | |
|---|---|---|---|---|---|
| aromatic compound biosynthetic process (GO:0019438) | 727 | 46 | 23.18 | 1.98 | 3.85E-03 |
| regulation of cellular protein localization (GO:1903827) | 544 | 34 | 17.35 | 1.96 | 3.08E-02 |
| cellular protein localization (GO:0034613) | 1375 | 85 | 43.85 | 1.94 | 5.36E-06 |
| cellular macromolecule localization (GO:0070727) | 1382 | 85 | 44.07 | 1.93 | 5.88E-06 |
| chromatin organization (GO:0006325) | 620 | 38 | 19.77 | 1.92 | 2.11E-02 |
| microtubule-based process (GO:0007017) | 645 | 39 | 20.57 | 1.9 | 2.53E-02 |
| regulation of cell cycle (GO:0051726) | 946 | 57 | 30.17 | 1.89 | 1.48E-03 |
| intracellular transport (GO:0046907) | 1142 | 67 | 36.42 | 1.84 | 7.65E-04 |
| organic cyclic compound biosynthetic process (GO:1901362) | 838 | 49 | 26.72 | 1.83 | 1.10E-02 |
| cellular catabolic process (GO:0044248) | 1475 | 86 | 47.04 | 1.83 | 4.65E-05 |
| protein localization (GO:0008104) | 1907 | 109 | 60.81 | 1.79 | 2.43E-06 |
| macromolecule localization (GO:0033036) | 2208 | 126 | 70.41 | 1.79 | 1.70E-07 |
| embryo development ending in birth or egg hatching (GO:0009792) | 815 | 46 | 25.99 | 1.77 | 3.32E-02 |
| cellular component biogenesis (GO:0044085) | 2406 | 135 | 76.73 | 1.76 | 1.10E-07 |
| organic substance catabolic process (GO:1901575) | 1412 | 79 | 45.03 | 1.75 | 4.82E-04 |
| cellular localization (GO:0051641) | 2040 | 114 | 65.05 | 1.75 | 3.09E-06 |
| macromolecule metabolic process (GO:0043170) | 5051 | 278 | 161.07 | 1.73 | 7.38E-18 |
| cellular macromolecule biosynthetic process (GO:0034645) | 1203 | 66 | 38.36 | 1.72 | 4.89E-03 |
| cell projection organization (GO:0030030) | 1096 | 60 | 34.95 | 1.72 | 1.22E-02 |
| macromolecule biosynthetic process (GO:0009059) | 1235 | 67 | 39.38 | 1.7 | 5.97E-03 |
| positive regulation of cellular component organization (GO:0051130) | 1217 | 66 | 38.81 | 1.7 | 7.36E-03 |
| establishment of localization in cell (GO:0051649) | 1441 | 78 | 45.95 | 1.7 | 1.61E-03 |
| regulation of organelle organization (GO:0033043) | 1225 | 66 | 39.06 | 1.69 | 7.79E-03 |
| regulation of nucleobase-containing compound metabolic process (GO:0019219) | 3242 | 172 | 103.38 | 1.66 | 1.64E-08 |
| plasma membrane bounded cell projection organization (GO:0120036) | 1041 | 55 | 33.2 | 1.66 | 3.71E-02 |

| | | | | | |
|---|---|---|---|---|---|
| cellular macromolecule metabolic process (GO:0044260) | 3973 | 209 | 126.7 | 1.65 | 1.13E-10 |
| nitrogen compound metabolic process (GO:0006807) | 5705 | 300 | 181.93 | 1.65 | 1.45E-17 |
| regulation of RNA metabolic process (GO:0051252) | 3011 | 158 | 96.02 | 1.65 | 2.36E-07 |
| organelle organization (GO:0006996) | 3000 | 157 | 95.67 | 1.64 | 3.09E-07 |
| proteolysis (GO:0006508) | 1094 | 57 | 34.89 | 1.63 | 3.69E-02 |
| regulation of cellular macromolecule biosynthetic process (GO:2000112) | 3180 | 165 | 101.41 | 1.63 | 1.65E-07 |
| regulation of gene expression (GO:0010468) | 3652 | 189 | 116.46 | 1.62 | 8.78E-09 |
| catabolic process (GO:0009056) | 1701 | 88 | 54.24 | 1.62 | 1.92E-03 |
| negative regulation of gene expression (GO:0010629) | 1588 | 82 | 50.64 | 1.62 | 4.67E-03 |
| regulation of macromolecule biosynthetic process (GO:0010556) | 3266 | 168 | 104.15 | 1.61 | 2.28E-07 |
| establishment of protein localization (GO:0045184) | 1315 | 67 | 41.93 | 1.6 | 2.57E-02 |
| negative regulation of cellular macromolecule biosynthetic process (GO:2000113) | 1341 | 68 | 42.76 | 1.59 | 2.30E-02 |
| protein transport (GO:0015031) | 1230 | 62 | 39.22 | 1.58 | 4.93E-02 |
| cellular component assembly (GO:0022607) | 2184 | 110 | 69.65 | 1.58 | 5.20E-04 |
| cellular protein metabolic process (GO:0044267) | 2863 | 144 | 91.3 | 1.58 | 1.73E-05 |
| regulation of cellular biosynthetic process (GO:0031326) | 3407 | 170 | 108.65 | 1.56 | 1.48E-06 |
| negative regulation of macromolecule biosynthetic process (GO:0010558) | 1376 | 68 | 43.88 | 1.55 | 4.14E-02 |
| cellular metabolic process (GO:0044237) | 6339 | 312 | 202.15 | 1.54 | 9.58E-15 |
| cellular response to stress (GO:0033554) | 1425 | 70 | 45.44 | 1.54 | 3.94E-02 |
| negative regulation of cellular biosynthetic process (GO:0031327) | 1426 | 70 | 45.47 | 1.54 | 3.96E-02 |
| regulation of nucleic acid-templated transcription (GO:1903506) | 2763 | 135 | 88.11 | 1.53 | 1.86E-04 |
| regulation of biosynthetic process (GO:0009889) | 3480 | 170 | 110.97 | 1.53 | 4.58E-06 |
| regulation of RNA biosynthetic process (GO:2001141) | 2766 | 135 | 88.21 | 1.53 | 1.86E-04 |

| | | | | | |
|---|---|---|---|---|---|
| regulation of transcription, DNA-templated (GO:0006355) | 2755 | 134 | 87.85 | 1.53 | 2.32E-04 |
| primary metabolic process (GO:0044238) | 6255 | 303 | 199.47 | 1.52 | 4.24E-13 |
| organic substance metabolic process (GO:0071704) | 6683 | 318 | 213.12 | 1.49 | 4.18E-13 |
| macromolecule modification (GO:0043412) | 2438 | 116 | 77.75 | 1.49 | 2.81E-03 |
| positive regulation of nucleobase-containing compound metabolic process (GO:0045935) | 1787 | 85 | 56.99 | 1.49 | 3.11E-02 |
| cellular component organization or biogenesis (GO:0071840) | 5335 | 253 | 170.13 | 1.49 | 4.79E-09 |
| cellular protein modification process (GO:0006464) | 2264 | 107 | 72.2 | 1.48 | 7.36E-03 |
| protein modification process (GO:0036211) | 2264 | 107 | 72.2 | 1.48 | 7.30E-03 |
| regulation of nitrogen compound metabolic process (GO:0051171) | 4995 | 236 | 159.29 | 1.48 | 5.25E-08 |
| regulation of macromolecule metabolic process (GO:0060255) | 5277 | 248 | 168.28 | 1.47 | 2.21E-08 |
| metabolic process (GO:0008152) | 7193 | 338 | 229.38 | 1.47 | 1.19E-13 |
| regulation of cellular metabolic process (GO:0031323) | 5335 | 250 | 170.13 | 1.47 | 2.06E-08 |
| regulation of primary metabolic process (GO:0080090) | 5143 | 241 | 164.01 | 1.47 | 5.85E-08 |
| cellular biosynthetic process (GO:0044249) | 1989 | 93 | 63.43 | 1.47 | 2.86E-02 |
| organic substance biosynthetic process (GO:1901576) | 2080 | 97 | 66.33 | 1.46 | 2.34E-02 |
| regulation of metabolic process (GO:0019222) | 5740 | 264 | 183.04 | 1.44 | 2.94E-08 |
| regulation of cellular component organization (GO:0051128) | 2464 | 113 | 78.57 | 1.44 | 1.36E-02 |
| protein metabolic process (GO:0019538) | 3442 | 157 | 109.76 | 1.43 | 7.62E-04 |
| cellular component organization (GO:0016043) | 5145 | 230 | 164.07 | 1.4 | 7.56E-06 |
| regulation of cellular protein metabolic process (GO:0032268) | 2470 | 110 | 78.77 | 1.4 | 3.72E-02 |
| negative regulation of macromolecule metabolic process (GO:0010605) | 2428 | 108 | 77.43 | 1.39 | 4.12E-02 |
| negative regulation of cellular metabolic process (GO:0031324) | 2408 | 107 | 76.79 | 1.39 | 4.79E-02 |

| | | | | | |
|---|---|---|---|---|---|
| positive regulation of nitrogen compound metabolic process (GO:0051173) | 3022 | 132 | 96.37 | 1.37 | 2.30E-02 |
| organonitrogen compound metabolic process (GO:1901564) | 4283 | 187 | 136.58 | 1.37 | 9.01E-04 |
| positive regulation of macromolecule metabolic process (GO:0010604) | 3183 | 137 | 101.5 | 1.35 | 2.96E-02 |
| positive regulation of cellular process (GO:0048522) | 5297 | 225 | 168.92 | 1.33 | 4.62E-04 |
| positive regulation of cellular metabolic process (GO:0031325) | 3182 | 135 | 101.47 | 1.33 | 4.85E-02 |
| positive regulation of biological process (GO:0048518) | 5994 | 251 | 191.14 | 1.31 | 2.30E-04 |
| negative regulation of cellular process (GO:0048523) | 4522 | 185 | 144.2 | 1.28 | 2.50E-02 |
| negative regulation of biological process (GO:0048519) | 5045 | 203 | 160.88 | 1.26 | 2.93E-02 |
| cellular process (GO:0009987) | 13986 | 535 | 446 | 1.2 | 1.87E-09 |
| regulation of cellular process (GO:0050794) | 10522 | 388 | 335.54 | 1.16 | 1.26E-02 |
| regulation of biological process (GO:0050789) | 11232 | 409 | 358.18 | 1.14 | 1.79E-02 |
| biological_process (GO:0008150) | 20380 | 684 | 649.9 | 1.05 | 1.84E-04 |
| cell communication (GO:0007154) | 5145 | 121 | 164.07 | 0.74 | 1.26E-02 |
| signal transduction (GO:0007165) | 4739 | 104 | 151.12 | 0.69 | 1.63E-03 |
| signaling (GO:0023052) | 5018 | 110 | 160.02 | 0.69 | 7.44E-04 |
| Unclassified (UNCLASSIFIED) | 1916 | 27 | 61.1 | 0.44 | 1.86E-04 |
| system process (GO:0003008) | 2561 | 35 | 81.67 | 0.43 | 9.54E-07 |
| nervous system process (GO:0050877) | 2065 | 19 | 65.85 | 0.29 | 4.15E-09 |
| sensory perception (GO:0007600) | 1624 | 10 | 51.79 | 0.19 | 8.55E-10 |
| adaptive immune response (GO:0002250) | 490 | 3 | 15.63 | 0.19 | 2.70E-02 |
| G protein-coupled receptor signaling pathway (GO:0007186) | 1830 | 8 | 58.36 | 0.14 | 1.34E-13 |
| defense response to bacterium (GO:0042742) | 461 | 2 | 14.7 | 0.14 | 1.21E-02 |
| immune response-activating cell surface receptor signaling pathway (GO:0002429) | 330 | 1 | 10.52 | 0.1 | 4.80E-02 |
| humoral immune response (GO:0006959) | 374 | 1 | 11.93 | 0.08 | 2.03E-02 |

| | | | | | |
|---|---|---|---|---|---|
| immune response-activating signal transduction (GO:0002757) | 390 | 1 | 12.44 | 0.08 | 1.10E-02 |
| sensory perception of chemical stimulus (GO:0007606) | 1220 | 1 | 38.9 | 0.03 | 4.80E-13 |
| sensory perception of smell (GO:0007608) | 1122 | 0 | 35.78 | < 0.01 | 4.25E-13 |

| **Green module** | **Mus musculus genes (22296)** | **Module genes (711)** | **Module genes (expected)** | **Module genes fold Enrichment** | **Module genes FDR** |
|---|---|---|---|---|---|
| NADH dehydrogenase complex assembly (GO:0010257) | 44 | 22 | 2.56 | 8.61 | 9.14E-10 |
| mitochondrial respiratory chain complex I assembly (GO:0032981) | 44 | 22 | 2.56 | 8.61 | 8.95E-10 |
| ATP synthesis coupled proton transport (GO:0015986) | 20 | 10 | 1.16 | 8.61 | 3.98E-04 |
| energy coupled proton transport, down electrochemical gradient (GO:0015985) | 20 | 10 | 1.16 | 8.61 | 3.94E-04 |
| protein neddylation (GO:0045116) | 15 | 7 | 0.87 | 8.03 | 1.15E-02 |
| viral budding (GO:0046755) | 15 | 7 | 0.87 | 8.03 | 1.14E-02 |
| endoplasmic reticulum tubular network organization (GO:0071786) | 13 | 6 | 0.76 | 7.95 | 2.98E-02 |
| mitochondrial ATP synthesis coupled electron transport (GO:0042775) | 51 | 21 | 2.96 | 7.09 | 3.79E-08 |
| cellular oxidant detoxification (GO:0098869) | 15 | 6 | 0.87 | 6.89 | 4.93E-02 |
| mitochondrial respiratory chain complex assembly (GO:0033108) | 76 | 30 | 4.41 | 6.8 | 1.89E-11 |
| ATP biosynthetic process (GO:0006754) | 33 | 13 | 1.92 | 6.78 | 1.21E-04 |
| oxidative phosphorylation (GO:0006119) | 64 | 25 | 3.72 | 6.73 | 2.14E-09 |
| ribosomal small subunit assembly (GO:0000028) | 18 | 7 | 1.05 | 6.7 | 2.38E-02 |
| ATP synthesis coupled electron transport (GO:0042773) | 55 | 21 | 3.19 | 6.57 | 1.09E-07 |
| virion assembly (GO:0019068) | 22 | 8 | 1.28 | 6.26 | 1.42E-02 |
| ribosomal small subunit biogenesis (GO:0042274) | 63 | 22 | 3.66 | 6.01 | 1.64E-07 |
| proteasomal ubiquitin-independent protein catabolic process (GO:0010499) | 24 | 8 | 1.39 | 5.74 | 2.14E-02 |
| respiratory electron transport chain (GO:0022904) | 76 | 25 | 4.41 | 5.66 | 3.57E-08 |

| | | | | | |
|---|---|---|---|---|---|
| purine ribonucleoside triphosphate biosynthetic process (GO:0009206) | 43 | 14 | 2.5 | 5.61 | 2.82E-04 |
| electron transport chain (GO:0022900) | 80 | 26 | 4.65 | 5.6 | 2.01E-08 |
| purine nucleoside triphosphate biosynthetic process (GO:0009145) | 44 | 14 | 2.56 | 5.48 | 3.46E-04 |
| maturation of 5.8S rRNA (GO:0000460) | 26 | 8 | 1.51 | 5.3 | 3.04E-02 |
| ribonucleoside triphosphate biosynthetic process (GO:0009201) | 46 | 14 | 2.67 | 5.24 | 5.03E-04 |
| endosome transport via multivesicular body sorting pathway (GO:0032509) | 27 | 8 | 1.57 | 5.1 | 3.63E-02 |
| tricarboxylic acid cycle (GO:0006099) | 31 | 9 | 1.8 | 5 | 2.13E-02 |
| maturation of SSU-rRNA from tricistronic rRNA transcript (SSU-rRNA, 5.8S rRNA, LSU-rRNA) (GO:0000462) | 31 | 9 | 1.8 | 5 | 2.12E-02 |
| maturation of SSU-rRNA (GO:0030490) | 42 | 12 | 2.44 | 4.92 | 3.38E-03 |
| multivesicular body sorting pathway (GO:0071985) | 32 | 9 | 1.86 | 4.84 | 2.44E-02 |
| translational initiation (GO:0006413) | 55 | 15 | 3.19 | 4.7 | 6.94E-04 |
| protein targeting to ER (GO:0045047) | 33 | 9 | 1.92 | 4.7 | 2.87E-02 |
| establishment of protein localization to endoplasmic reticulum (GO:0072599) | 37 | 10 | 2.15 | 4.65 | 1.68E-02 |
| translation (GO:0006412) | 310 | 82 | 18.01 | 4.55 | 4.44E-23 |
| transcription initiation from RNA polymerase II promoter (GO:0006367) | 38 | 10 | 2.21 | 4.53 | 1.98E-02 |
| nucleoside triphosphate biosynthetic process (GO:0009142) | 57 | 15 | 3.31 | 4.53 | 9.49E-04 |
| aerobic respiration (GO:0009060) | 65 | 17 | 3.78 | 4.5 | 2.91E-04 |
| ATP metabolic process (GO:0046034) | 170 | 44 | 9.87 | 4.46 | 9.64E-12 |
| hydrogen peroxide metabolic process (GO:0042743) | 43 | 11 | 2.5 | 4.4 | 1.28E-02 |
| peptide biosynthetic process (GO:0043043) | 329 | 84 | 19.11 | 4.4 | 7.55E-23 |
| cellular respiration (GO:0045333) | 135 | 33 | 7.84 | 4.21 | 3.58E-08 |
| purine ribonucleoside triphosphate metabolic process (GO:0009205) | 58 | 14 | 3.37 | 4.16 | 3.60E-03 |

| | | | | | |
|---|---|---|---|---|---|
| proton transmembrane transport (GO:1902600) | 65 | 15 | 3.78 | 3.97 | 3.16E-03 |
| cytoplasmic translation (GO:0002181) | 65 | 15 | 3.78 | 3.97 | 3.14E-03 |
| peptide metabolic process (GO:0006518) | 451 | 103 | 26.2 | 3.93 | 5.67E-25 |
| ribosome assembly (GO:0042255) | 66 | 15 | 3.83 | 3.91 | 3.53E-03 |
| ribonucleoside triphosphate metabolic process (GO:0009199) | 62 | 14 | 3.6 | 3.89 | 6.20E-03 |
| protein targeting to mitochondrion (GO:0006626) | 58 | 13 | 3.37 | 3.86 | 1.09E-02 |
| amide biosynthetic process (GO:0043604) | 421 | 93 | 24.45 | 3.8 | 1.10E-21 |
| mitochondrial translation (GO:0032543) | 50 | 11 | 2.9 | 3.79 | 3.21E-02 |
| ribosomal large subunit biogenesis (GO:0042273) | 79 | 17 | 4.59 | 3.7 | 2.18E-03 |
| mitochondrion organization (GO:0007005) | 387 | 82 | 22.48 | 3.65 | 4.35E-18 |
| establishment of protein localization to mitochondrion (GO:0072655) | 67 | 14 | 3.89 | 3.6 | 1.15E-02 |
| cell redox homeostasis (GO:0045454) | 63 | 13 | 3.66 | 3.55 | 1.98E-02 |
| protein localization to mitochondrion (GO:0070585) | 73 | 15 | 4.24 | 3.54 | 8.36E-03 |
| energy derivation by oxidation of organic compounds (GO:0015980) | 196 | 40 | 11.38 | 3.51 | 4.56E-08 |
| mitochondrial transmembrane transport (GO:1990542) | 64 | 13 | 3.72 | 3.5 | 2.17E-02 |
| purine nucleoside triphosphate metabolic process (GO:0009144) | 69 | 14 | 4.01 | 3.49 | 1.42E-02 |
| protein folding (GO:0006457) | 149 | 30 | 8.65 | 3.47 | 8.69E-06 |
| RNA export from nucleus (GO:0006405) | 65 | 13 | 3.78 | 3.44 | 2.44E-02 |
| ribonucleoprotein complex export from nucleus (GO:0071426) | 61 | 12 | 3.54 | 3.39 | 4.15E-02 |
| ribonucleoprotein complex localization (GO:0071166) | 62 | 12 | 3.6 | 3.33 | 4.60E-02 |
| viral life cycle (GO:0019058) | 68 | 13 | 3.95 | 3.29 | 3.28E-02 |
| generation of precursor metabolites and energy (GO:0006091) | 285 | 54 | 16.55 | 3.26 | 5.50E-10 |
| mitochondrial transport (GO:0006839) | 166 | 31 | 9.64 | 3.22 | 2.17E-05 |
| ribosome biogenesis (GO:0042254) | 284 | 53 | 16.5 | 3.21 | 1.35E-09 |
| cellular amide metabolic process (GO:0043603) | 663 | 121 | 38.51 | 3.14 | 2.53E-22 |

| | | | | | |
|---|---|---|---|---|---|
| rRNA processing (GO:0006364) | 192 | 35 | 11.15 | 3.14 | 6.18E-06 |
| purine ribonucleotide biosynthetic process (GO:0009152) | 114 | 20 | 6.62 | 3.02 | 4.67E-03 |
| rRNA metabolic process (GO:0016072) | 201 | 35 | 11.67 | 3 | 1.61E-05 |
| purine nucleotide biosynthetic process (GO:0006164) | 124 | 21 | 7.2 | 2.92 | 4.76E-03 |
| nucleoside triphosphate metabolic process (GO:0009141) | 89 | 15 | 5.17 | 2.9 | 3.92E-02 |
| ribonucleoprotein complex biogenesis (GO:0022613) | 418 | 69 | 24.28 | 2.84 | 1.70E-10 |
| protein targeting (GO:0006605) | 208 | 34 | 12.08 | 2.81 | 8.04E-05 |
| purine-containing compound biosynthetic process (GO:0072522) | 129 | 21 | 7.49 | 2.8 | 7.58E-03 |
| ribonucleotide biosynthetic process (GO:0009260) | 123 | 20 | 7.14 | 2.8 | 1.05E-02 |
| ribose phosphate biosynthetic process (GO:0046390) | 130 | 21 | 7.55 | 2.78 | 8.25E-03 |
| protein maturation (GO:0051604) | 222 | 35 | 12.89 | 2.71 | 1.15E-04 |
| RNA localization (GO:0006403) | 160 | 25 | 9.29 | 2.69 | 3.41E-03 |
| vacuolar transport (GO:0007034) | 132 | 20 | 7.67 | 2.61 | 2.12E-02 |
| ribonucleoprotein complex assembly (GO:0022618) | 206 | 31 | 11.96 | 2.59 | 1.11E-03 |
| RNA transport (GO:0050658) | 140 | 21 | 8.13 | 2.58 | 2.47E-02 |
| nucleic acid transport (GO:0050657) | 140 | 21 | 8.13 | 2.58 | 2.45E-02 |
| cellular protein-containing complex assembly (GO:0034622) | 694 | 104 | 40.31 | 2.58 | 7.65E-14 |
| establishment of RNA localization (GO:0051236) | 142 | 21 | 8.25 | 2.55 | 2.60E-02 |
| organonitrogen compound biosynthetic process (GO:1901566) | 1079 | 159 | 62.67 | 2.54 | 4.00E-21 |
| ribonucleoprotein complex subunit organization (GO:0071826) | 219 | 32 | 12.72 | 2.52 | 1.19E-03 |
| RNA splicing (GO:0008380) | 315 | 46 | 18.3 | 2.51 | 2.24E-05 |
| gene expression (GO:0010467) | 1591 | 230 | 92.41 | 2.49 | 1.33E-30 |
| ncRNA processing (GO:0034470) | 336 | 48 | 19.52 | 2.46 | 1.88E-05 |
| cellular nitrogen compound biosynthetic process (GO:0044271) | 1108 | 156 | 64.36 | 2.42 | 3.95E-19 |

| | | | | | |
|---|---|---|---|---|---|
| mRNA splicing, via spliceosome (GO:0000398) | 195 | 27 | 11.33 | 2.38 | 1.29E-02 |
| RNA splicing, via transesterification reactions with bulged adenosine as nucleophile (GO:0000377) | 195 | 27 | 11.33 | 2.38 | 1.28E-02 |
| RNA splicing, via transesterification reactions (GO:0000375) | 195 | 27 | 11.33 | 2.38 | 1.27E-02 |
| protein processing (GO:0016485) | 161 | 22 | 9.35 | 2.35 | 3.61E-02 |
| RNA processing (GO:0006396) | 742 | 99 | 43.1 | 2.3 | 2.61E-10 |
| purine ribonucleotide metabolic process (GO:0009150) | 256 | 34 | 14.87 | 2.29 | 4.21E-03 |
| purine nucleotide metabolic process (GO:0006163) | 273 | 36 | 15.86 | 2.27 | 2.73E-03 |
| symbiotic process (GO:0044403) | 220 | 29 | 12.78 | 2.27 | 1.15E-02 |
| ribonucleotide metabolic process (GO:0009259) | 266 | 35 | 15.45 | 2.27 | 3.51E-03 |
| cellular macromolecule biosynthetic process (GO:0034645) | 1203 | 158 | 69.87 | 2.26 | 1.12E-16 |
| endosomal transport (GO:0016197) | 198 | 26 | 11.5 | 2.26 | 2.43E-02 |
| ncRNA metabolic process (GO:0034660) | 412 | 54 | 23.93 | 2.26 | 4.72E-05 |
| ribose phosphate metabolic process (GO:0019693) | 277 | 36 | 16.09 | 2.24 | 3.22E-03 |
| macromolecule biosynthetic process (GO:0009059) | 1235 | 159 | 71.73 | 2.22 | 3.09E-16 |
| cofactor metabolic process (GO:0051186) | 376 | 48 | 21.84 | 2.2 | 3.66E-04 |
| mRNA processing (GO:0006397) | 401 | 51 | 23.29 | 2.19 | 1.76E-04 |
| establishment of protein localization to organelle (GO:0072594) | 292 | 37 | 16.96 | 2.18 | 4.47E-03 |
| oxidation-reduction process (GO:0055114) | 791 | 100 | 45.94 | 2.18 | 3.01E-09 |
| Golgi vesicle transport (GO:0048193) | 248 | 31 | 14.4 | 2.15 | 1.56E-02 |
| purine-containing compound metabolic process (GO:0072521) | 313 | 39 | 18.18 | 2.15 | 3.43E-03 |
| nucleotide metabolic process (GO:0009117) | 349 | 43 | 20.27 | 2.12 | 2.06E-03 |
| regulation of translation (GO:0006417) | 320 | 39 | 18.59 | 2.1 | 5.69E-03 |
| nucleoside phosphate metabolic process (GO:0006753) | 359 | 43 | 20.85 | 2.06 | 3.70E-03 |
| cellular biosynthetic process (GO:0044249) | 1989 | 236 | 115.53 | 2.04 | 5.54E-21 |
| mRNA metabolic process (GO:0016071) | 522 | 61 | 30.32 | 2.01 | 2.09E-04 |

| | | | | | |
|---|---|---|---|---|---|
| nucleobase-containing small molecule metabolic process (GO:0055086) | 423 | 49 | 24.57 | 1.99 | 2.70E-03 |
| intracellular transport (GO:0046907) | 1142 | 131 | 66.33 | 1.97 | 9.10E-10 |
| cellular nitrogen compound metabolic process (GO:0034641) | 2757 | 316 | 160.13 | 1.97 | 8.20E-27 |
| intracellular protein transport (GO:0006886) | 752 | 86 | 43.68 | 1.97 | 5.54E-06 |
| regulation of cellular amide metabolic process (GO:0034248) | 368 | 42 | 21.37 | 1.96 | 1.02E-02 |
| drug metabolic process (GO:0017144) | 412 | 47 | 23.93 | 1.96 | 4.67E-03 |
| organic substance biosynthetic process (GO:1901576) | 2080 | 235 | 120.81 | 1.95 | 2.30E-18 |
| biosynthetic process (GO:0009058) | 2161 | 244 | 125.52 | 1.94 | 3.29E-19 |
| protein-containing complex assembly (GO:0065003) | 1334 | 150 | 77.48 | 1.94 | 8.74E-11 |
| protein localization to organelle (GO:0033365) | 601 | 67 | 34.91 | 1.92 | 3.81E-04 |
| protein transport (GO:0015031) | 1230 | 136 | 71.44 | 1.9 | 2.90E-09 |
| cellular protein catabolic process (GO:0044257) | 570 | 63 | 33.11 | 1.9 | 7.23E-04 |
| peptide transport (GO:0015833) | 1258 | 138 | 73.07 | 1.89 | 3.07E-09 |
| amide transport (GO:0042886) | 1279 | 140 | 74.29 | 1.88 | 2.93E-09 |
| carbohydrate derivative biosynthetic process (GO:1901137) | 458 | 50 | 26.6 | 1.88 | 7.07E-03 |
| posttranscriptional regulation of gene expression (GO:0010608) | 414 | 45 | 24.05 | 1.87 | 1.48E-02 |
| protein-containing complex subunit organization (GO:0043933) | 1522 | 164 | 88.4 | 1.86 | 1.70E-10 |
| modification-dependent macromolecule catabolic process (GO:0043632) | 494 | 53 | 28.69 | 1.85 | 6.12E-03 |
| protein catabolic process (GO:0030163) | 626 | 67 | 36.36 | 1.84 | 1.16E-03 |
| cellular macromolecule catabolic process (GO:0044265) | 740 | 79 | 42.98 | 1.84 | 1.73E-04 |
| RNA metabolic process (GO:0016070) | 1222 | 130 | 70.98 | 1.83 | 8.40E-08 |
| establishment of protein localization (GO:0045184) | 1315 | 139 | 76.38 | 1.82 | 2.48E-08 |
| modification-dependent protein catabolic process (GO:0019941) | 484 | 51 | 28.11 | 1.81 | 1.39E-02 |

| | | | | | |
|---|---|---|---|---|---|
| proteolysis involved in cellular protein catabolic process (GO:0051603) | 552 | 57 | 32.06 | 1.78 | 8.97E-03 |
| macromolecule catabolic process (GO:0009057) | 833 | 85 | 48.38 | 1.76 | 4.02E-04 |
| establishment of localization in cell (GO:0051649) | 1441 | 145 | 83.7 | 1.73 | 2.39E-07 |
| cellular protein metabolic process (GO:0044267) | 2863 | 284 | 166.29 | 1.71 | 1.26E-15 |
| cellular component biogenesis (GO:0044085) | 2406 | 237 | 139.75 | 1.7 | 3.62E-12 |
| nitrogen compound transport (GO:0071705) | 1526 | 149 | 88.63 | 1.68 | 8.46E-07 |
| nucleobase-containing compound metabolic process (GO:0006139) | 2162 | 211 | 125.57 | 1.68 | 3.43E-10 |
| heterocycle metabolic process (GO:0046483) | 2301 | 220 | 133.65 | 1.65 | 5.13E-10 |
| cellular macromolecule localization (GO:0070727) | 1382 | 132 | 80.27 | 1.64 | 1.80E-05 |
| cellular protein localization (GO:0034613) | 1375 | 131 | 79.86 | 1.64 | 2.24E-05 |
| nucleic acid metabolic process (GO:0090304) | 1740 | 165 | 101.06 | 1.63 | 7.27E-07 |
| cellular metabolic process (GO:0044237) | 6339 | 600 | 368.18 | 1.63 | 1.25E-35 |
| protein localization (GO:0008104) | 1907 | 178 | 110.76 | 1.61 | 5.05E-07 |
| organonitrogen compound catabolic process (GO:1901565) | 879 | 82 | 51.05 | 1.61 | 8.47E-03 |
| cellular aromatic compound metabolic process (GO:0006725) | 2369 | 220 | 137.6 | 1.6 | 6.93E-09 |
| cellular macromolecule metabolic process (GO:0044260) | 3973 | 363 | 230.76 | 1.57 | 1.25E-15 |
| heterocycle biosynthetic process (GO:0018130) | 712 | 65 | 41.35 | 1.57 | 4.97E-02 |
| metabolic process (GO:0008152) | 7193 | 653 | 417.79 | 1.56 | 2.67E-35 |
| organonitrogen compound metabolic process (GO:1901564) | 4283 | 388 | 248.77 | 1.56 | 1.50E-16 |
| carboxylic acid metabolic process (GO:0019752) | 773 | 70 | 44.9 | 1.56 | 4.58E-02 |
| organic cyclic compound metabolic process (GO:1901360) | 2580 | 233 | 149.85 | 1.55 | 1.47E-08 |
| nitrogen compound metabolic process (GO:0006807) | 5705 | 515 | 331.36 | 1.55 | 6.28E-24 |
| cellular component assembly (GO:0022607) | 2184 | 197 | 126.85 | 1.55 | 7.34E-07 |

| | | | | | |
|---|---|---|---|---|---|
| carbohydrate derivative metabolic process (GO:1901135) | 800 | 72 | 46.47 | 1.55 | 4.31E-02 |
| organic cyclic compound biosynthetic process (GO:1901362) | 838 | 75 | 48.67 | 1.54 | 3.45E-02 |
| macromolecule localization (GO:0033036) | 2208 | 197 | 128.25 | 1.54 | 1.48E-06 |
| protein metabolic process (GO:0019538) | 3442 | 307 | 199.92 | 1.54 | 2.46E-11 |
| macromolecule metabolic process (GO:0043170) | 5051 | 449 | 293.37 | 1.53 | 1.20E-18 |
| cellular localization (GO:0051641) | 2040 | 181 | 118.49 | 1.53 | 9.32E-06 |
| organic substance catabolic process (GO:1901575) | 1412 | 125 | 82.01 | 1.52 | 1.28E-03 |
| organic substance transport (GO:0071702) | 1842 | 162 | 106.99 | 1.51 | 7.84E-05 |
| organic substance metabolic process (GO:0071704) | 6683 | 585 | 388.16 | 1.51 | 2.02E-25 |
| small molecule metabolic process (GO:0044281) | 1438 | 125 | 83.52 | 1.5 | 2.48E-03 |
| primary metabolic process (GO:0044238) | 6255 | 543 | 363.3 | 1.49 | 4.13E-22 |
| cellular catabolic process (GO:0044248) | 1475 | 127 | 85.67 | 1.48 | 3.01E-03 |
| catabolic process (GO:0009056) | 1701 | 143 | 98.8 | 1.45 | 2.99E-03 |
| organelle organization (GO:0006996) | 3000 | 250 | 174.25 | 1.43 | 3.16E-06 |
| transport (GO:0006810) | 3523 | 268 | 204.62 | 1.31 | 8.54E-04 |
| establishment of localization (GO:0051234) | 3664 | 278 | 212.81 | 1.31 | 7.11E-04 |
| cellular component organization or biogenesis (GO:0071840) | 5335 | 396 | 309.87 | 1.28 | 2.36E-05 |
| cellular component organization (GO:0016043) | 5145 | 362 | 298.83 | 1.21 | 7.77E-03 |
| cellular process (GO:0009987) | 13986 | 931 | 812.34 | 1.15 | 4.26E-09 |
| biological_process (GO:0008150) | 20380 | 1238 | 1183.71 | 1.05 | 3.12E-06 |
| regulation of biological process (GO:0050789) | 11232 | 588 | 652.38 | 0.9 | 3.88E-02 |
| regulation of cellular process (GO:0050794) | 10522 | 548 | 611.14 | 0.9 | 4.56E-02 |
| system development (GO:0048731) | 4205 | 193 | 244.24 | 0.79 | 2.45E-02 |
| cellular response to stimulus (GO:0051716) | 6446 | 290 | 374.4 | 0.77 | 4.95E-05 |
| response to stimulus (GO:0050896) | 8284 | 371 | 481.15 | 0.77 | 8.64E-08 |
| animal organ development (GO:0048513) | 3040 | 133 | 176.57 | 0.75 | 3.17E-02 |
| cellular developmental process (GO:0048869) | 3769 | 163 | 218.91 | 0.74 | 3.77E-03 |

| | | | | | |
|---|---|---|---|---|---|
| cell differentiation (GO:0030154) | 3678 | 155 | 213.63 | 0.73 | 1.29E-03 |
| multicellular organismal process (GO:0032501) | 7231 | 296 | 419.99 | 0.7 | 7.78E-11 |
| regulation of multicellular organismal development (GO:2000026) | 2121 | 86 | 123.19 | 0.7 | 2.99E-02 |
| immune system process (GO:0002376) | 2274 | 92 | 132.08 | 0.7 | 1.73E-02 |
| neurogenesis (GO:0022008) | 1742 | 67 | 101.18 | 0.66 | 2.47E-02 |
| cell surface receptor signaling pathway (GO:0007166) | 1804 | 67 | 104.78 | 0.64 | 8.54E-03 |
| immune response (GO:0006955) | 1392 | 50 | 80.85 | 0.62 | 2.17E-02 |
| generation of neurons (GO:0048699) | 1636 | 56 | 95.02 | 0.59 | 2.35E-03 |
| cell development (GO:0048468) | 1746 | 59 | 101.41 | 0.58 | 8.11E-04 |
| cell communication (GO:0007154) | 5145 | 170 | 298.83 | 0.57 | 9.07E-16 |
| regulation of neurogenesis (GO:0050767) | 942 | 31 | 54.71 | 0.57 | 4.87E-02 |
| signal transduction (GO:0007165) | 4739 | 150 | 275.25 | 0.54 | 4.12E-16 |
| signaling (GO:0023052) | 5018 | 158 | 291.46 | 0.54 | 1.51E-17 |
| regulation of nervous system development (GO:0051960) | 1055 | 33 | 61.28 | 0.54 | 1.17E-02 |
| Unclassified (UNCLASSIFIED) | 1916 | 57 | 111.29 | 0.51 | 3.08E-06 |
| protein phosphorylation (GO:0006468) | 723 | 21 | 41.99 | 0.5 | 4.01E-02 |
| regulation of leukocyte activation (GO:0002694) | 698 | 19 | 40.54 | 0.47 | 2.44E-02 |
| animal organ morphogenesis (GO:0009887) | 1005 | 27 | 58.37 | 0.46 | 9.57E-04 |
| import into cell (GO:0098657) | 652 | 17 | 37.87 | 0.45 | 2.21E-02 |
| system process (GO:0003008) | 2561 | 61 | 148.75 | 0.41 | 4.78E-14 |
| regulation of lymphocyte activation (GO:0051249) | 602 | 14 | 34.97 | 0.4 | 1.02E-02 |
| adaptive immune response (GO:0002250) | 490 | 11 | 28.46 | 0.39 | 3.37E-02 |
| nervous system process (GO:0050877) | 2065 | 42 | 119.94 | 0.35 | 6.69E-14 |
| positive regulation of lymphocyte activation (GO:0051251) | 444 | 9 | 25.79 | 0.35 | 2.63E-02 |
| pattern specification process (GO:0007389) | 448 | 9 | 26.02 | 0.35 | 2.12E-02 |
| immune response-regulating signaling pathway (GO:0002764) | 403 | 8 | 23.41 | 0.34 | 4.33E-02 |

| | | | | | |
|---|---|---|---|---|---|
| defense response to bacterium (GO:0042742) | 461 | 9 | 26.78 | 0.34 | 1.69E-02 |
| cellular process involved in reproduction in multicellular organism (GO:0022412) | 424 | 8 | 24.63 | 0.32 | 2.09E-02 |
| immune response-activating signal transduction (GO:0002757) | 390 | 7 | 22.65 | 0.31 | 2.47E-02 |
| phagocytosis (GO:0006909) | 314 | 5 | 18.24 | 0.27 | 4.87E-02 |
| antigen receptor-mediated signaling pathway (GO:0050851) | 306 | 4 | 17.77 | 0.23 | 1.95E-02 |
| regulation of B cell activation (GO:0050864) | 313 | 4 | 18.18 | 0.22 | 1.44E-02 |
| immune response-activating cell surface receptor signaling pathway (GO:0002429) | 330 | 4 | 19.17 | 0.21 | 8.37E-03 |
| immune response-regulating cell surface receptor signaling pathway (GO:0002768) | 342 | 4 | 19.86 | 0.2 | 4.60E-03 |
| sensory perception (GO:0007600) | 1624 | 17 | 94.33 | 0.18 | 1.50E-19 |
| G protein-coupled receptor signaling pathway (GO:0007186) | 1830 | 17 | 106.29 | 0.16 | 6.66E-24 |
| positive regulation of B cell activation (GO:0050871) | 267 | 2 | 15.51 | 0.13 | 6.94E-03 |
| B cell receptor signaling pathway (GO:0050853) | 220 | 1 | 12.78 | 0.08 | 8.36E-03 |
| membrane invagination (GO:0010324) | 238 | 1 | 13.82 | 0.07 | 4.46E-03 |
| phagocytosis, engulfment (GO:0006911) | 223 | 0 | 12.95 | < 0.01 | 7.26E-04 |
| phagocytosis, recognition (GO:0006910) | 202 | 0 | 11.73 | < 0.01 | 2.16E-03 |
| plasma membrane invagination (GO:0099024) | 232 | 0 | 13.48 | < 0.01 | 5.07E-04 |
| sensory perception of smell (GO:0007608) | 1122 | 0 | 65.17 | < 0.01 | 2.60E-25 |
| sensory perception of chemical stimulus (GO:0007606) | 1220 | 0 | 70.86 | < 0.01 | 1.22E-27 |
| **Blue module** | **Mus musculus genes (22296)** | **Module genes (711)** | **Module genes (expected)** | **Module genes fold Enrichment** | **Module genes FDR** |
| inner ear receptor cell fate commitment (GO:0060120) | 5 | 5 | 0.47 | 10.55 | 3.09E-02 |
| auditory receptor cell fate commitment (GO:0009912) | 5 | 5 | 0.47 | 10.55 | 3.08E-02 |
| regulation of timing of cell differentiation (GO:0048505) | 14 | 7 | 1.33 | 5.28 | 4.75E-02 |
| paraxial mesoderm development (GO:0048339) | 22 | 10 | 2.08 | 4.8 | 1.12E-02 |

| | | | | | |
|---|---|---|---|---|---|
| central nervous system projection neuron axonogenesis (GO:0021952) | 28 | 12 | 2.65 | 4.52 | 4.98E-03 |
| auditory receptor cell morphogenesis (GO:0002093) | 22 | 9 | 2.08 | 4.32 | 3.37E-02 |
| lung epithelial cell differentiation (GO:0060487) | 32 | 13 | 3.03 | 4.29 | 4.09E-03 |
| telencephalon glial cell migration (GO:0022030) | 25 | 10 | 2.37 | 4.22 | 2.23E-02 |
| cerebral cortex radial glia guided migration (GO:0021801) | 25 | 10 | 2.37 | 4.22 | 2.22E-02 |
| hair cell differentiation (GO:0035315) | 48 | 19 | 4.55 | 4.18 | 1.86E-04 |
| lung cell differentiation (GO:0060479) | 33 | 13 | 3.13 | 4.16 | 5.05E-03 |
| negative regulation of stem cell differentiation (GO:2000737) | 23 | 9 | 2.18 | 4.13 | 4.13E-02 |
| synaptic membrane adhesion (GO:0099560) | 26 | 10 | 2.46 | 4.06 | 2.75E-02 |
| inner ear auditory receptor cell differentiation (GO:0042491) | 42 | 16 | 3.98 | 4.02 | 1.36E-03 |
| aorta morphogenesis (GO:0035909) | 29 | 11 | 2.75 | 4 | 1.79E-02 |
| auditory receptor cell development (GO:0060117) | 28 | 10 | 2.65 | 3.77 | 3.99E-02 |
| pharyngeal system development (GO:0060037) | 28 | 10 | 2.65 | 3.77 | 3.98E-02 |
| inner ear receptor cell stereocilium organization (GO:0060122) | 41 | 14 | 3.89 | 3.6 | 8.62E-03 |
| lung epithelium development (GO:0060428) | 44 | 15 | 4.17 | 3.6 | 5.64E-03 |
| regulation of heart morphogenesis (GO:2000826) | 42 | 14 | 3.98 | 3.52 | 1.03E-02 |
| inner ear receptor cell differentiation (GO:0060113) | 75 | 25 | 7.11 | 3.52 | 7.77E-05 |
| heart trabecula morphogenesis (GO:0061384) | 36 | 12 | 3.41 | 3.52 | 2.44E-02 |
| columnar/cuboidal epithelial cell development (GO:0002066) | 61 | 20 | 5.78 | 3.46 | 8.57E-04 |
| endocardial cushion development (GO:0003197) | 46 | 15 | 4.36 | 3.44 | 7.95E-03 |
| synapse assembly (GO:0007416) | 80 | 26 | 7.58 | 3.43 | 7.16E-05 |
| negative regulation of animal organ morphogenesis (GO:0110111) | 40 | 13 | 3.79 | 3.43 | 1.88E-02 |
| regulation of stem cell differentiation (GO:2000736) | 59 | 19 | 5.59 | 3.4 | 1.65E-03 |
| heart valve morphogenesis (GO:0003179) | 47 | 15 | 4.45 | 3.37 | 9.35E-03 |

| | | | | | |
|---|---|---|---|---|---|
| central nervous system neuron axonogenesis (GO:0021955) | 41 | 13 | 3.89 | 3.35 | 2.21E-02 |
| aorta development (GO:0035904) | 57 | 18 | 5.4 | 3.33 | 3.04E-03 |
| trabecula morphogenesis (GO:0061383) | 51 | 16 | 4.83 | 3.31 | 7.17E-03 |
| exocrine system development (GO:0035272) | 48 | 15 | 4.55 | 3.3 | 1.10E-02 |
| neuroepithelial cell differentiation (GO:0060563) | 64 | 20 | 6.07 | 3.3 | 1.48E-03 |
| inner ear receptor cell development (GO:0060119) | 58 | 18 | 5.5 | 3.27 | 3.56E-03 |
| mechanoreceptor differentiation (GO:0042490) | 81 | 25 | 7.68 | 3.26 | 2.21E-04 |
| digestive tract morphogenesis (GO:0048546) | 53 | 16 | 5.02 | 3.19 | 9.78E-03 |
| heart valve development (GO:0003170) | 57 | 17 | 5.4 | 3.15 | 7.44E-03 |
| somitogenesis (GO:0001756) | 71 | 21 | 6.73 | 3.12 | 1.82E-03 |
| ventricular septum morphogenesis (GO:0060412) | 44 | 13 | 4.17 | 3.12 | 3.46E-02 |
| response to epidermal growth factor (GO:0070849) | 44 | 13 | 4.17 | 3.12 | 3.45E-02 |
| histone H4 acetylation (GO:0043967) | 51 | 15 | 4.83 | 3.1 | 1.75E-02 |
| digestive system development (GO:0055123) | 130 | 38 | 12.32 | 3.08 | 3.41E-06 |
| mesenchyme morphogenesis (GO:0072132) | 48 | 14 | 4.55 | 3.08 | 2.68E-02 |
| ventricular septum development (GO:0003281) | 79 | 23 | 7.49 | 3.07 | 9.98E-04 |
| artery morphogenesis (GO:0048844) | 69 | 20 | 6.54 | 3.06 | 3.24E-03 |
| intraciliary transport (GO:0042073) | 45 | 13 | 4.26 | 3.05 | 3.99E-02 |
| positive regulation of synapse assembly (GO:0051965) | 77 | 22 | 7.3 | 3.01 | 1.84E-03 |
| establishment or maintenance of apical/basal cell polarity (GO:0035088) | 49 | 14 | 4.64 | 3.01 | 3.07E-02 |
| somatic stem cell population maintenance (GO:0035019) | 49 | 14 | 4.64 | 3.01 | 3.06E-02 |
| establishment or maintenance of bipolar cell polarity (GO:0061245) | 49 | 14 | 4.64 | 3.01 | 3.06E-02 |
| somite development (GO:0061053) | 88 | 25 | 8.34 | 3 | 6.51E-04 |
| digestive tract development (GO:0048565) | 117 | 33 | 11.09 | 2.98 | 4.21E-05 |
| positive regulation of glucose transmembrane transport (GO:0010828) | 50 | 14 | 4.74 | 2.95 | 3.47E-02 |
| inner ear morphogenesis (GO:0042472) | 111 | 31 | 10.52 | 2.95 | 9.87E-05 |

| | | | | | |
|---|---|---|---|---|---|
| epithelial to mesenchymal transition (GO:0001837) | 61 | 17 | 5.78 | 2.94 | 1.33E-02 |
| regulation of synapse assembly (GO:0051963) | 119 | 33 | 11.28 | 2.93 | 5.55E-05 |
| coronary vasculature development (GO:0060976) | 73 | 20 | 6.92 | 2.89 | 5.73E-03 |
| cardiac septum development (GO:0003279) | 121 | 33 | 11.47 | 2.88 | 7.39E-05 |
| cerebral cortex cell migration (GO:0021795) | 55 | 15 | 5.21 | 2.88 | 3.05E-02 |
| artery development (GO:0060840) | 99 | 27 | 9.38 | 2.88 | 5.61E-04 |
| histone lysine methylation (GO:0034968) | 59 | 16 | 5.59 | 2.86 | 2.30E-02 |
| ear morphogenesis (GO:0042471) | 133 | 36 | 12.6 | 2.86 | 4.78E-05 |
| oligodendrocyte differentiation (GO:0048709) | 63 | 17 | 5.97 | 2.85 | 1.75E-02 |
| positive regulation of dendritic spine development (GO:0060999) | 63 | 17 | 5.97 | 2.85 | 1.74E-02 |
| regulation of neuron migration (GO:2001222) | 52 | 14 | 4.93 | 2.84 | 4.52E-02 |
| neuron fate commitment (GO:0048663) | 78 | 21 | 7.39 | 2.84 | 4.96E-03 |
| positive regulation of neural precursor cell proliferation (GO:2000179) | 67 | 18 | 6.35 | 2.83 | 1.32E-02 |
| embryonic heart tube morphogenesis (GO:0003143) | 67 | 18 | 6.35 | 2.83 | 1.32E-02 |
| positive regulation of dendrite development (GO:1900006) | 108 | 29 | 10.24 | 2.83 | 3.65E-04 |
| heart looping (GO:0001947) | 60 | 16 | 5.69 | 2.81 | 2.63E-02 |
| roof of mouth development (GO:0060021) | 95 | 25 | 9 | 2.78 | 2.72E-03 |
| maintenance of cell number (GO:0098727) | 141 | 37 | 13.36 | 2.77 | 4.69E-05 |
| regulation of cardiocyte differentiation (GO:1905207) | 61 | 16 | 5.78 | 2.77 | 2.97E-02 |
| columnar/cuboidal epithelial cell differentiation (GO:0002065) | 119 | 31 | 11.28 | 2.75 | 3.77E-04 |
| stem cell population maintenance (GO:0019827) | 137 | 35 | 12.98 | 2.7 | 1.28E-04 |
| cardiac septum morphogenesis (GO:0060411) | 79 | 20 | 7.49 | 2.67 | 1.69E-02 |
| segmentation (GO:0035282) | 99 | 25 | 9.38 | 2.66 | 3.57E-03 |
| microtubule bundle formation (GO:0001578) | 100 | 25 | 9.48 | 2.64 | 3.88E-03 |
| ear development (GO:0043583) | 237 | 59 | 22.46 | 2.63 | 1.72E-07 |
| male gonad development (GO:0008584) | 114 | 28 | 10.8 | 2.59 | 1.91E-03 |

| | | | | | |
|---|---|---|---|---|---|
| mesenchyme development (GO:0060485) | 212 | 52 | 20.09 | 2.59 | 2.00E-06 |
| respiratory tube development (GO:0030323) | 204 | 50 | 19.33 | 2.59 | 3.56E-06 |
| histone methylation (GO:0016571) | 82 | 20 | 7.77 | 2.57 | 2.07E-02 |
| embryonic heart tube development (GO:0035050) | 82 | 20 | 7.77 | 2.57 | 2.06E-02 |
| lung development (GO:0030324) | 201 | 49 | 19.05 | 2.57 | 5.35E-06 |
| outflow tract morphogenesis (GO:0003151) | 78 | 19 | 7.39 | 2.57 | 2.80E-02 |
| development of primary male sexual characteristics (GO:0046546) | 115 | 28 | 10.9 | 2.57 | 2.10E-03 |
| respiratory system development (GO:0060541) | 231 | 56 | 21.89 | 2.56 | 1.02E-06 |
| Notch signaling pathway (GO:0007219) | 117 | 28 | 11.09 | 2.53 | 3.88E-03 |
| postsynapse organization (GO:0099173) | 92 | 22 | 8.72 | 2.52 | 1.36E-02 |
| regulation of dendritic spine development (GO:0060998) | 92 | 22 | 8.72 | 2.52 | 1.35E-02 |
| cerebral cortex development (GO:0021987) | 109 | 26 | 10.33 | 2.52 | 4.71E-03 |
| hair cycle process (GO:0022405) | 97 | 23 | 9.19 | 2.5 | 1.11E-02 |
| molting cycle process (GO:0022404) | 97 | 23 | 9.19 | 2.5 | 1.11E-02 |
| inner ear development (GO:0048839) | 207 | 49 | 19.62 | 2.5 | 1.21E-05 |
| synapse organization (GO:0050808) | 275 | 65 | 26.06 | 2.49 | 1.75E-07 |
| epithelial cell development (GO:0002064) | 216 | 51 | 20.47 | 2.49 | 7.07E-06 |
| neuron projection guidance (GO:0097485) | 217 | 51 | 20.57 | 2.48 | 7.65E-06 |
| histone acetylation (GO:0016573) | 94 | 22 | 8.91 | 2.47 | 1.64E-02 |
| neural tube development (GO:0021915) | 188 | 44 | 17.82 | 2.47 | 6.47E-05 |
| sensory perception of sound (GO:0007605) | 154 | 36 | 14.59 | 2.47 | 4.51E-04 |
| forebrain generation of neurons (GO:0021872) | 77 | 18 | 7.3 | 2.47 | 4.67E-02 |
| gland morphogenesis (GO:0022612) | 120 | 28 | 11.37 | 2.46 | 4.54E-03 |
| regulation of dendrite development (GO:0050773) | 189 | 44 | 17.91 | 2.46 | 6.92E-05 |
| axon guidance (GO:0007411) | 215 | 50 | 20.38 | 2.45 | 1.25E-05 |
| mesoderm development (GO:0007498) | 108 | 25 | 10.24 | 2.44 | 1.11E-02 |
| sensory organ morphogenesis (GO:0090596) | 286 | 66 | 27.1 | 2.44 | 3.23E-07 |
| peptidyl-threonine modification (GO:0018210) | 91 | 21 | 8.62 | 2.44 | 2.39E-02 |

| | | | | | |
|---|---|---|---|---|---|
| mesenchymal cell differentiation (GO:0048762) | 152 | 35 | 14.41 | 2.43 | 1.12E-03 |
| regulation of synapse organization (GO:0050807) | 265 | 61 | 25.11 | 2.43 | 9.96E-07 |
| regulation of neural precursor cell proliferation (GO:2000177) | 113 | 26 | 10.71 | 2.43 | 8.71E-03 |
| cell morphogenesis involved in neuron differentiation (GO:0048667) | 423 | 97 | 40.09 | 2.42 | 9.73E-11 |
| skin epidermis development (GO:0098773) | 96 | 22 | 9.1 | 2.42 | 2.80E-02 |
| camera-type eye morphogenesis (GO:0048593) | 131 | 30 | 12.41 | 2.42 | 3.08E-03 |
| peptidyl-threonine phosphorylation (GO:0018107) | 83 | 19 | 7.87 | 2.42 | 4.13E-02 |
| hair cycle (GO:0042633) | 105 | 24 | 9.95 | 2.41 | 1.62E-02 |
| molting cycle (GO:0042303) | 105 | 24 | 9.95 | 2.41 | 1.61E-02 |
| regulation of dendrite morphogenesis (GO:0048814) | 114 | 26 | 10.8 | 2.41 | 9.29E-03 |
| regulation of epithelial cell differentiation (GO:0030856) | 145 | 33 | 13.74 | 2.4 | 2.22E-03 |
| morphogenesis of embryonic epithelium (GO:0016331) | 176 | 40 | 16.68 | 2.4 | 2.98E-04 |
| cardiac ventricle development (GO:0003231) | 141 | 32 | 13.36 | 2.39 | 2.03E-03 |
| negative regulation of canonical Wnt signaling pathway (GO:0090090) | 119 | 27 | 11.28 | 2.39 | 7.41E-03 |
| axonogenesis (GO:0007409) | 335 | 76 | 31.75 | 2.39 | 3.95E-08 |
| epidermal cell differentiation (GO:0009913) | 168 | 38 | 15.92 | 2.39 | 5.40E-04 |
| regulation of synapse structure or activity (GO:0050803) | 275 | 62 | 26.06 | 2.38 | 1.57E-06 |
| limb morphogenesis (GO:0035108) | 169 | 38 | 16.02 | 2.37 | 5.79E-04 |
| appendage morphogenesis (GO:0035107) | 169 | 38 | 16.02 | 2.37 | 5.77E-04 |
| kidney epithelium development (GO:0072073) | 129 | 29 | 12.23 | 2.37 | 4.91E-03 |
| epidermis development (GO:0008544) | 272 | 61 | 25.78 | 2.37 | 2.31E-06 |
| primary neural tube formation (GO:0014020) | 116 | 26 | 10.99 | 2.37 | 1.07E-02 |
| hair follicle development (GO:0001942) | 94 | 21 | 8.91 | 2.36 | 4.04E-02 |
| cardiac chamber development (GO:0003205) | 188 | 42 | 17.82 | 2.36 | 3.23E-04 |
| developmental growth involved in morphogenesis (GO:0060560) | 130 | 29 | 12.32 | 2.35 | 5.31E-03 |
| embryonic organ morphogenesis (GO:0048562) | 314 | 70 | 29.76 | 2.35 | 3.12E-07 |

| | | | | | |
|---|---|---|---|---|---|
| post-embryonic development (GO:0009791) | 126 | 28 | 11.94 | 2.34 | 7.07E-03 |
| internal peptidyl-lysine acetylation (GO:0018393) | 99 | 22 | 9.38 | 2.34 | 3.20E-02 |
| negative regulation of cell morphogenesis involved in differentiation (GO:0010771) | 99 | 22 | 9.38 | 2.34 | 3.19E-02 |
| neural tube formation (GO:0001841) | 131 | 29 | 12.41 | 2.34 | 8.11E-03 |
| multicellular organism growth (GO:0035264) | 113 | 25 | 10.71 | 2.33 | 1.56E-02 |
| axon development (GO:0061564) | 363 | 80 | 34.4 | 2.33 | 3.94E-08 |
| embryonic placenta development (GO:0001892) | 109 | 24 | 10.33 | 2.32 | 2.06E-02 |
| neural tube closure (GO:0001843) | 109 | 24 | 10.33 | 2.32 | 2.06E-02 |
| renal system development (GO:0072001) | 279 | 61 | 26.44 | 2.31 | 5.19E-06 |
| tube closure (GO:0060606) | 110 | 24 | 10.42 | 2.3 | 2.21E-02 |
| limb development (GO:0060173) | 193 | 42 | 18.29 | 2.3 | 4.25E-04 |
| appendage development (GO:0048736) | 193 | 42 | 18.29 | 2.3 | 4.23E-04 |
| telencephalon development (GO:0021537) | 239 | 52 | 22.65 | 2.3 | 4.37E-05 |
| sensory perception of mechanical stimulus (GO:0050954) | 184 | 40 | 17.44 | 2.29 | 7.19E-04 |
| heart morphogenesis (GO:0003007) | 267 | 58 | 25.3 | 2.29 | 1.26E-05 |
| embryonic epithelial tube formation (GO:0001838) | 152 | 33 | 14.41 | 2.29 | 3.29E-03 |
| kidney development (GO:0001822) | 258 | 56 | 24.45 | 2.29 | 2.29E-05 |
| anterior/posterior pattern specification (GO:0009952) | 226 | 49 | 21.42 | 2.29 | 9.86E-05 |
| urogenital system development (GO:0001655) | 319 | 69 | 30.23 | 2.28 | 1.07E-06 |
| regulation of postsynapse organization (GO:0099175) | 125 | 27 | 11.85 | 2.28 | 1.61E-02 |
| internal protein amino acid acetylation (GO:0006475) | 102 | 22 | 9.67 | 2.28 | 3.87E-02 |
| peptidyl-lysine acetylation (GO:0018394) | 102 | 22 | 9.67 | 2.28 | 3.86E-02 |
| cell morphogenesis involved in differentiation (GO:0000904) | 563 | 121 | 53.36 | 2.27 | 1.10E-11 |
| eye morphogenesis (GO:0048592) | 163 | 35 | 15.45 | 2.27 | 3.19E-03 |
| tube formation (GO:0035148) | 173 | 37 | 16.4 | 2.26 | 1.93E-03 |
| male sex differentiation (GO:0046661) | 141 | 30 | 13.36 | 2.25 | 8.34E-03 |
| skin development (GO:0043588) | 254 | 54 | 24.07 | 2.24 | 5.16E-05 |

| | | | | | |
|---|---|---|---|---|---|
| pallium development (GO:0021543) | 157 | 33 | 14.88 | 2.22 | 6.24E-03 |
| dendrite development (GO:0016358) | 129 | 27 | 12.23 | 2.21 | 1.96E-02 |
| embryonic organ development (GO:0048568) | 478 | 100 | 45.3 | 2.21 | 3.93E-09 |
| epithelial tube formation (GO:0072175) | 158 | 33 | 14.97 | 2.2 | 6.53E-03 |
| cell-cell adhesion via plasma-membrane adhesion molecules (GO:0098742) | 183 | 38 | 17.34 | 2.19 | 2.92E-03 |
| anatomical structure maturation (GO:0071695) | 150 | 31 | 14.22 | 2.18 | 1.14E-02 |
| cardiac chamber morphogenesis (GO:0003206) | 141 | 29 | 13.36 | 2.17 | 1.96E-02 |
| neuron projection morphogenesis (GO:0048812) | 462 | 95 | 43.78 | 2.17 | 2.36E-08 |
| embryonic morphogenesis (GO:0048598) | 608 | 125 | 57.62 | 2.17 | 4.10E-11 |
| plasma membrane bounded cell projection morphogenesis (GO:0120039) | 467 | 96 | 44.26 | 2.17 | 1.93E-08 |
| cell projection morphogenesis (GO:0048858) | 473 | 97 | 44.83 | 2.16 | 2.15E-08 |
| developmental growth (GO:0048589) | 444 | 91 | 42.08 | 2.16 | 6.86E-08 |
| brain development (GO:0007420) | 610 | 125 | 57.81 | 2.16 | 4.63E-11 |
| determination of left/right symmetry (GO:0007368) | 122 | 25 | 11.56 | 2.16 | 3.58E-02 |
| regulation of axonogenesis (GO:0050770) | 205 | 42 | 19.43 | 2.16 | 1.25E-03 |
| regulation of extent of cell growth (GO:0061387) | 127 | 26 | 12.04 | 2.16 | 2.92E-02 |
| regulation of cell morphogenesis involved in differentiation (GO:0010769) | 343 | 70 | 32.51 | 2.15 | 5.11E-06 |
| cilium assembly (GO:0060271) | 256 | 52 | 24.26 | 2.14 | 2.26E-04 |
| determination of bilateral symmetry (GO:0009855) | 128 | 26 | 12.13 | 2.14 | 3.10E-02 |
| morphogenesis of an epithelium (GO:0002009) | 498 | 101 | 47.2 | 2.14 | 1.46E-08 |
| embryonic limb morphogenesis (GO:0030326) | 143 | 29 | 13.55 | 2.14 | 2.11E-02 |
| embryonic appendage morphogenesis (GO:0035113) | 143 | 29 | 13.55 | 2.14 | 2.11E-02 |
| placenta development (GO:0001890) | 168 | 34 | 15.92 | 2.14 | 9.40E-03 |
| plasma membrane bounded cell projection assembly (GO:0120031) | 346 | 70 | 32.79 | 2.13 | 7.89E-06 |
| head development (GO:0060322) | 664 | 134 | 62.93 | 2.13 | 2.20E-11 |
| nephron development (GO:0072006) | 124 | 25 | 11.75 | 2.13 | 4.03E-02 |

| | | | | | |
|---|---|---|---|---|---|
| forebrain development (GO:0030900) | 372 | 75 | 35.25 | 2.13 | 3.65E-06 |
| specification of symmetry (GO:0009799) | 129 | 26 | 12.23 | 2.13 | 4.47E-02 |
| regulation of embryonic development (GO:0045995) | 129 | 26 | 12.23 | 2.13 | 4.46E-02 |
| cilium organization (GO:0044782) | 289 | 58 | 27.39 | 2.12 | 8.85E-05 |
| cell projection assembly (GO:0030031) | 359 | 72 | 34.02 | 2.12 | 5.85E-06 |
| regulation of organelle assembly (GO:1902115) | 205 | 41 | 19.43 | 2.11 | 2.79E-03 |
| growth (GO:0040007) | 460 | 92 | 43.59 | 2.11 | 1.36E-07 |
| negative regulation of Wnt signaling pathway (GO:0030178) | 150 | 30 | 14.22 | 2.11 | 1.87E-02 |
| epithelial tube morphogenesis (GO:0060562) | 356 | 71 | 33.74 | 2.1 | 8.01E-06 |
| regulation of GTPase activity (GO:0043087) | 296 | 59 | 28.05 | 2.1 | 1.04E-04 |
| morphogenesis of a branching structure (GO:0001763) | 201 | 40 | 19.05 | 2.1 | 3.70E-03 |
| stem cell differentiation (GO:0048863) | 166 | 33 | 15.73 | 2.1 | 1.35E-02 |
| plasma membrane bounded cell projection organization (GO:0120036) | 1041 | 206 | 98.66 | 2.09 | 7.93E-17 |
| negative regulation of neuron differentiation (GO:0045665) | 258 | 51 | 24.45 | 2.09 | 5.68E-04 |
| bone development (GO:0060348) | 208 | 41 | 19.71 | 2.08 | 3.23E-03 |
| neuron projection development (GO:0031175) | 680 | 134 | 64.44 | 2.08 | 9.16E-11 |
| cell part morphogenesis (GO:0032990) | 498 | 98 | 47.2 | 2.08 | 8.73E-08 |
| regulation of canonical Wnt signaling pathway (GO:0060828) | 224 | 44 | 21.23 | 2.07 | 2.30E-03 |
| cell morphogenesis (GO:0000902) | 734 | 144 | 69.56 | 2.07 | 1.93E-11 |
| sensory organ development (GO:0007423) | 571 | 112 | 54.11 | 2.07 | 8.80E-09 |
| camera-type eye development (GO:0043010) | 320 | 62 | 30.33 | 2.04 | 1.05E-04 |
| morphogenesis of a branching epithelium (GO:0061138) | 191 | 37 | 18.1 | 2.04 | 9.15E-03 |
| cell projection organization (GO:0030030) | 1096 | 212 | 103.87 | 2.04 | 1.97E-16 |
| peptidyl-lysine modification (GO:0018205) | 238 | 46 | 22.56 | 2.04 | 2.41E-03 |
| tissue morphogenesis (GO:0048729) | 623 | 120 | 59.04 | 2.03 | 3.90E-09 |
| adult behavior (GO:0030534) | 182 | 35 | 17.25 | 2.03 | 1.53E-02 |

| | | | | | |
|---|---|---|---|---|---|
| animal organ morphogenesis (GO:0009887) | 1005 | 193 | 95.24 | 2.03 | 1.38E-14 |
| histone modification (GO:0016570) | 318 | 61 | 30.14 | 2.02 | 1.57E-04 |
| cell fate commitment (GO:0045165) | 266 | 51 | 25.21 | 2.02 | 1.12E-03 |
| epithelial cell differentiation (GO:0030855) | 517 | 99 | 49 | 2.02 | 2.40E-07 |
| regulation of animal organ morphogenesis (GO:2000027) | 209 | 40 | 19.81 | 2.02 | 7.19E-03 |
| heart development (GO:0007507) | 565 | 108 | 53.55 | 2.02 | 5.90E-08 |
| central nervous system development (GO:0007417) | 820 | 156 | 77.71 | 2.01 | 1.64E-11 |
| reproductive system development (GO:0061458) | 432 | 82 | 40.94 | 2 | 7.30E-06 |
| covalent chromatin modification (GO:0016569) | 327 | 62 | 30.99 | 2 | 1.98E-04 |
| gliogenesis (GO:0042063) | 227 | 43 | 21.51 | 2 | 5.64E-03 |
| reproductive structure development (GO:0048608) | 428 | 81 | 40.56 | 2 | 9.57E-06 |
| regulation of small GTPase mediated signal transduction (GO:0051056) | 259 | 49 | 24.55 | 2 | 2.05E-03 |
| regulation of Ras protein signal transduction (GO:0046578) | 228 | 43 | 21.61 | 1.99 | 5.78E-03 |
| cellular component morphogenesis (GO:0032989) | 841 | 158 | 79.7 | 1.98 | 1.94E-11 |
| skeletal system development (GO:0001501) | 487 | 91 | 46.15 | 1.97 | 2.45E-06 |
| positive regulation of neuron projection development (GO:0010976) | 359 | 67 | 34.02 | 1.97 | 1.49E-04 |
| epithelium development (GO:0060429) | 1029 | 192 | 97.52 | 1.97 | 1.11E-13 |
| regulation of protein localization to membrane (GO:1905475) | 188 | 35 | 17.82 | 1.96 | 2.63E-02 |
| negative regulation of neurogenesis (GO:0050768) | 335 | 62 | 31.75 | 1.95 | 3.73E-04 |
| connective tissue development (GO:0061448) | 227 | 42 | 21.51 | 1.95 | 8.35E-03 |
| central nervous system neuron differentiation (GO:0021953) | 200 | 37 | 18.95 | 1.95 | 1.77E-02 |
| tube development (GO:0035295) | 912 | 168 | 86.43 | 1.94 | 1.46E-11 |
| regulation of neuron differentiation (GO:0045664) | 766 | 141 | 72.59 | 1.94 | 1.25E-09 |
| positive regulation of cell morphogenesis involved in differentiation (GO:0010770) | 185 | 34 | 17.53 | 1.94 | 3.42E-02 |

| | | | | | |
|---|---|---|---|---|---|
| neuron differentiation (GO:0030182) | 1024 | 188 | 97.04 | 1.94 | 8.28E-13 |
| positive regulation of nervous system development (GO:0051962) | 643 | 118 | 60.94 | 1.94 | 7.58E-08 |
| neuron development (GO:0048666) | 830 | 152 | 78.66 | 1.93 | 2.97E-10 |
| regulation of nervous system development (GO:0051960) | 1055 | 193 | 99.98 | 1.93 | 4.52E-13 |
| regulation of Wnt signaling pathway (GO:0030111) | 290 | 53 | 27.48 | 1.93 | 2.26E-03 |
| establishment or maintenance of cell polarity (GO:0007163) | 197 | 36 | 18.67 | 1.93 | 2.39E-02 |
| eye development (GO:0001654) | 362 | 66 | 34.31 | 1.92 | 3.57E-04 |
| signal transduction by protein phosphorylation (GO:0023014) | 181 | 33 | 17.15 | 1.92 | 4.41E-02 |
| chromatin organization (GO:0006325) | 620 | 113 | 58.76 | 1.92 | 2.00E-07 |
| tube morphogenesis (GO:0035239) | 708 | 129 | 67.1 | 1.92 | 2.01E-08 |
| macromolecule methylation (GO:0043414) | 215 | 39 | 20.38 | 1.91 | 1.81E-02 |
| regulation of neuron projection development (GO:0010975) | 597 | 108 | 56.58 | 1.91 | 6.72E-07 |
| regulation of microtubule cytoskeleton organization (GO:0070507) | 199 | 36 | 18.86 | 1.91 | 3.41E-02 |
| visual system development (GO:0150063) | 365 | 66 | 34.59 | 1.91 | 3.82E-04 |
| regulation of cell morphogenesis (GO:0022604) | 522 | 94 | 49.47 | 1.9 | 6.07E-06 |
| cell-cell adhesion (GO:0098609) | 378 | 68 | 35.82 | 1.9 | 3.67E-04 |
| negative regulation of cell development (GO:0010721) | 384 | 69 | 36.39 | 1.9 | 2.97E-04 |
| sensory system development (GO:0048880) | 368 | 66 | 34.88 | 1.89 | 4.26E-04 |
| negative regulation of nervous system development (GO:0051961) | 358 | 64 | 33.93 | 1.89 | 6.63E-04 |
| regulation of neurogenesis (GO:0050767) | 942 | 168 | 89.27 | 1.88 | 1.20E-10 |
| positive regulation of neurogenesis (GO:0050769) | 573 | 102 | 54.3 | 1.88 | 3.38E-06 |
| positive regulation of cell projection organization (GO:0031346) | 463 | 82 | 43.88 | 1.87 | 7.69E-05 |

| | | | | | |
|---|---|---|---|---|---|
| positive regulation of neuron differentiation (GO:0045666) | 458 | 81 | 43.4 | 1.87 | 9.74E-05 |
| in utero embryonic development (GO:0001701) | 516 | 91 | 48.9 | 1.86 | 2.20E-05 |
| regulation of developmental growth (GO:0048638) | 392 | 69 | 37.15 | 1.86 | 5.07E-04 |
| positive regulation of developmental growth (GO:0048639) | 212 | 37 | 20.09 | 1.84 | 4.51E-02 |
| regulation of microtubule-based process (GO:0032886) | 235 | 41 | 22.27 | 1.84 | 2.80E-02 |
| positive regulation of GTPase activity (GO:0043547) | 218 | 38 | 20.66 | 1.84 | 3.76E-02 |
| chordate embryonic development (GO:0043009) | 798 | 139 | 75.63 | 1.84 | 4.04E-08 |
| pattern specification process (GO:0007389) | 448 | 78 | 42.46 | 1.84 | 2.20E-04 |
| embryo development (GO:0009790) | 1155 | 201 | 109.46 | 1.84 | 6.38E-12 |
| regulation of cytoskeleton organization (GO:0051493) | 535 | 93 | 50.7 | 1.83 | 2.48E-05 |
| positive regulation of cellular component biogenesis (GO:0044089) | 507 | 88 | 48.05 | 1.83 | 5.31E-05 |
| embryo development ending in birth or egg hatching (GO:0009792) | 815 | 141 | 77.24 | 1.83 | 5.70E-08 |
| positive regulation of cell development (GO:0010720) | 652 | 112 | 61.79 | 1.81 | 3.39E-06 |
| locomotory behavior (GO:0007626) | 239 | 41 | 22.65 | 1.81 | 3.10E-02 |
| gland development (GO:0048732) | 428 | 73 | 40.56 | 1.8 | 7.66E-04 |
| regulation of cell development (GO:0060284) | 1078 | 183 | 102.16 | 1.79 | 5.35E-10 |
| generation of neurons (GO:0048699) | 1636 | 277 | 155.04 | 1.79 | 3.07E-15 |
| positive regulation of transcription by RNA polymerase II (GO:0045944) | 1218 | 206 | 115.43 | 1.78 | 2.81E-11 |
| regionalization (GO:0003002) | 349 | 59 | 33.07 | 1.78 | 5.85E-03 |
| anatomical structure formation involved in morphogenesis (GO:0048646) | 914 | 154 | 86.62 | 1.78 | 3.99E-08 |
| positive regulation of RNA biosynthetic process (GO:1902680) | 1532 | 258 | 145.19 | 1.78 | 4.76E-14 |
| muscle cell differentiation (GO:0042692) | 268 | 45 | 25.4 | 1.77 | 2.86E-02 |
| positive regulation of nucleic acid-templated transcription (GO:1903508) | 1531 | 257 | 145.09 | 1.77 | 8.42E-14 |

| | | | | | |
|---|---|---|---|---|---|
| positive regulation of transcription, DNA-templated (GO:0045893) | 1527 | 256 | 144.71 | 1.77 | 1.07E-13 |
| striated muscle tissue development (GO:0014706) | 323 | 54 | 30.61 | 1.76 | 1.24E-02 |
| muscle tissue development (GO:0060537) | 341 | 57 | 32.32 | 1.76 | 7.18E-03 |
| anatomical structure morphogenesis (GO:0009653) | 2188 | 365 | 207.36 | 1.76 | 5.66E-20 |
| regulation of actin cytoskeleton organization (GO:0032956) | 342 | 57 | 32.41 | 1.76 | 9.38E-03 |
| regulation of cellular response to growth factor stimulus (GO:0090287) | 270 | 45 | 25.59 | 1.76 | 3.04E-02 |
| neurogenesis (GO:0022008) | 1742 | 290 | 165.09 | 1.76 | 3.41E-15 |
| tissue development (GO:0009888) | 1658 | 276 | 157.13 | 1.76 | 1.85E-14 |
| positive regulation of RNA metabolic process (GO:0051254) | 1629 | 271 | 154.38 | 1.76 | 3.75E-14 |
| microtubule-based movement (GO:0007018) | 259 | 43 | 24.55 | 1.75 | 4.39E-02 |
| circulatory system development (GO:0072359) | 903 | 149 | 85.58 | 1.74 | 2.26E-07 |
| regulation of supramolecular fiber organization (GO:1902903) | 346 | 57 | 32.79 | 1.74 | 1.02E-02 |
| muscle structure development (GO:0061061) | 486 | 80 | 46.06 | 1.74 | 8.62E-04 |
| regulation of actin filament-based process (GO:0032970) | 390 | 64 | 36.96 | 1.73 | 5.00E-03 |
| regulation of plasma membrane bounded cell projection organization (GO:0120035) | 765 | 125 | 72.5 | 1.72 | 5.73E-06 |
| positive regulation of nucleobase-containing compound metabolic process (GO:0045935) | 1787 | 291 | 169.35 | 1.72 | 2.36E-14 |
| blood vessel development (GO:0001568) | 522 | 85 | 49.47 | 1.72 | 6.51E-04 |
| regulation of cell projection organization (GO:0031344) | 774 | 126 | 73.35 | 1.72 | 6.31E-06 |
| organelle assembly (GO:0070925) | 639 | 104 | 60.56 | 1.72 | 8.87E-05 |
| behavior (GO:0007610) | 670 | 109 | 63.5 | 1.72 | 4.56E-05 |
| nervous system development (GO:0007399) | 2226 | 360 | 210.96 | 1.71 | 9.24E-18 |
| cardiovascular system development (GO:0072358) | 563 | 91 | 53.36 | 1.71 | 3.95E-04 |

| | | | | | |
|---|---|---|---|---|---|
| negative regulation of transcription by RNA polymerase II (GO:0000122) | 852 | 137 | 80.74 | 1.7 | 3.65E-06 |
| regulation of anatomical structure morphogenesis (GO:0022603) | 1066 | 171 | 101.03 | 1.69 | 1.10E-07 |
| vasculature development (GO:0001944) | 549 | 88 | 52.03 | 1.69 | 7.43E-04 |
| negative regulation of cell differentiation (GO:0045596) | 757 | 121 | 71.74 | 1.69 | 2.95E-05 |
| blood vessel morphogenesis (GO:0048514) | 421 | 67 | 39.9 | 1.68 | 7.93E-03 |
| regulation of epithelial cell proliferation (GO:0050678) | 359 | 57 | 34.02 | 1.68 | 2.21E-02 |
| cytoskeleton organization (GO:0007010) | 1028 | 163 | 97.42 | 1.67 | 4.87E-07 |
| regulation of transcription by RNA polymerase II (GO:0006357) | 1966 | 311 | 186.32 | 1.67 | 4.90E-14 |
| regulation of cellular component biogenesis (GO:0044087) | 923 | 146 | 87.47 | 1.67 | 3.39E-06 |
| protein phosphorylation (GO:0006468) | 723 | 114 | 68.52 | 1.66 | 9.89E-05 |
| regulation of chromosome organization (GO:0033044) | 349 | 55 | 33.07 | 1.66 | 3.22E-02 |
| microtubule cytoskeleton organization (GO:0000226) | 465 | 73 | 44.07 | 1.66 | 6.51E-03 |
| actin cytoskeleton organization (GO:0030036) | 474 | 74 | 44.92 | 1.65 | 7.26E-03 |
| peptidyl-amino acid modification (GO:0018193) | 705 | 110 | 66.81 | 1.65 | 2.37E-04 |
| actin filament-based process (GO:0030029) | 532 | 83 | 50.42 | 1.65 | 3.19E-03 |
| cellular response to growth factor stimulus (GO:0071363) | 391 | 61 | 37.06 | 1.65 | 2.04E-02 |
| cell development (GO:0048468) | 1746 | 272 | 165.47 | 1.64 | 1.39E-11 |
| negative regulation of transcription, DNA-templated (GO:0045892) | 1149 | 178 | 108.89 | 1.63 | 3.62E-07 |
| regulation of cellular component size (GO:0032535) | 394 | 61 | 37.34 | 1.63 | 2.70E-02 |
| negative regulation of nucleic acid-templated transcription (GO:1903507) | 1153 | 178 | 109.27 | 1.63 | 4.95E-07 |
| negative regulation of cellular macromolecule biosynthetic process (GO:2000113) | 1341 | 207 | 127.09 | 1.63 | 2.94E-08 |
| negative regulation of RNA biosynthetic process (GO:1902679) | 1154 | 178 | 109.36 | 1.63 | 5.00E-07 |

| | | | | | |
|---|---|---|---|---|---|
| positive regulation of cell differentiation (GO:0045597) | 1081 | 166 | 102.45 | 1.62 | 2.30E-06 |
| cell adhesion (GO:0007155) | 789 | 121 | 74.77 | 1.62 | 1.40E-04 |
| negative regulation of macromolecule biosynthetic process (GO:0010558) | 1376 | 211 | 130.4 | 1.62 | 2.74E-08 |
| response to growth factor (GO:0070848) | 405 | 62 | 38.38 | 1.62 | 3.05E-02 |
| positive regulation of macromolecule biosynthetic process (GO:0010557) | 1788 | 273 | 169.45 | 1.61 | 8.41E-11 |
| negative regulation of RNA metabolic process (GO:0051253) | 1240 | 189 | 117.52 | 1.61 | 3.76E-07 |
| chromosome organization (GO:0051276) | 914 | 139 | 86.62 | 1.6 | 4.67E-05 |
| positive regulation of cellular component organization (GO:0051130) | 1217 | 185 | 115.34 | 1.6 | 6.35E-07 |
| movement of cell or subcellular component (GO:0006928) | 1366 | 207 | 129.46 | 1.6 | 9.32E-08 |
| positive regulation of gene expression (GO:0010628) | 1954 | 296 | 185.18 | 1.6 | 1.89E-11 |
| biological adhesion (GO:0022610) | 799 | 121 | 75.72 | 1.6 | 2.69E-04 |
| negative regulation of nucleobase-containing compound metabolic process (GO:0045934) | 1343 | 203 | 127.28 | 1.59 | 1.62E-07 |
| regulation of multicellular organismal development (GO:2000026) | 2121 | 319 | 201.01 | 1.59 | 5.01E-12 |
| microtubule-based process (GO:0007017) | 645 | 97 | 61.13 | 1.59 | 2.70E-03 |
| animal organ development (GO:0048513) | 3040 | 457 | 288.1 | 1.59 | 5.03E-18 |
| regulation of cell differentiation (GO:0045595) | 1845 | 277 | 174.85 | 1.58 | 2.72E-10 |
| developmental process involved in reproduction (GO:0003006) | 740 | 111 | 70.13 | 1.58 | 7.99E-04 |
| positive regulation of cellular biosynthetic process (GO:0031328) | 1882 | 282 | 178.36 | 1.58 | 1.87E-10 |
| regulation of organelle organization (GO:0033043) | 1225 | 183 | 116.09 | 1.58 | 2.05E-06 |
| positive regulation of biosynthetic process (GO:0009891) | 1923 | 285 | 182.24 | 1.56 | 3.77E-10 |
| enzyme linked receptor protein signaling pathway (GO:0007167) | 528 | 78 | 50.04 | 1.56 | 1.75E-02 |

| | | | | | |
|---|---|---|---|---|---|
| negative regulation of cellular biosynthetic process (GO:0031327) | 1426 | 210 | 135.14 | 1.55 | 4.99E-07 |
| regulation of cellular protein localization (GO:1903827) | 544 | 80 | 51.56 | 1.55 | 1.64E-02 |
| positive regulation of developmental process (GO:0051094) | 1514 | 222 | 143.48 | 1.55 | 2.75E-07 |
| negative regulation of developmental process (GO:0051093) | 1023 | 150 | 96.95 | 1.55 | 9.20E-05 |
| locomotion (GO:0040011) | 1160 | 170 | 109.93 | 1.55 | 1.98E-05 |
| regulation of cell migration (GO:0030334) | 889 | 130 | 84.25 | 1.54 | 4.37E-04 |
| negative regulation of biosynthetic process (GO:0009890) | 1464 | 214 | 138.74 | 1.54 | 7.15E-07 |
| negative regulation of gene expression (GO:0010629) | 1588 | 231 | 150.5 | 1.53 | 2.18E-07 |
| regulation of transcription, DNA-templated (GO:0006355) | 2755 | 397 | 261.09 | 1.52 | 5.64E-13 |
| regulation of growth (GO:0040008) | 694 | 100 | 65.77 | 1.52 | 7.04E-03 |
| regulation of RNA biosynthetic process (GO:2001141) | 2766 | 398 | 262.13 | 1.52 | 6.08E-13 |
| regulation of developmental process (GO:0050793) | 2635 | 379 | 249.72 | 1.52 | 3.02E-12 |
| regulation of RNA metabolic process (GO:0051252) | 3011 | 433 | 285.35 | 1.52 | 3.64E-14 |
| regulation of nucleic acid-templated transcription (GO:1903506) | 2763 | 397 | 261.85 | 1.52 | 7.58E-13 |
| regulation of cellular component movement (GO:0051270) | 1024 | 147 | 97.04 | 1.51 | 3.04E-04 |
| regulation of cell motility (GO:2000145) | 936 | 134 | 88.71 | 1.51 | 8.36E-04 |
| regulation of cellular component organization (GO:0051128) | 2464 | 351 | 233.51 | 1.5 | 9.60E-11 |
| multicellular organism development (GO:0007275) | 4810 | 685 | 455.85 | 1.5 | 9.09E-24 |
| system development (GO:0048731) | 4205 | 598 | 398.51 | 1.5 | 5.32E-20 |
| regulation of nucleobase-containing compound metabolic process (GO:0019219) | 3242 | 461 | 307.25 | 1.5 | 1.83E-14 |
| cellular response to endogenous stimulus (GO:0071495) | 904 | 128 | 85.67 | 1.49 | 1.99E-03 |
| localization of cell (GO:0051674) | 958 | 135 | 90.79 | 1.49 | 1.62E-03 |

| | | | | | |
|---|---|---|---|---|---|
| cell motility (GO:0048870) | 958 | 135 | 90.79 | 1.49 | 1.62E-03 |
| regulation of cellular macromolecule biosynthetic process (GO:2000112) | 3180 | 446 | 301.37 | 1.48 | 3.20E-13 |
| cell differentiation (GO:0030154) | 3678 | 514 | 348.57 | 1.47 | 3.64E-15 |
| cell migration (GO:0016477) | 840 | 117 | 79.61 | 1.47 | 6.51E-03 |
| regulation of macromolecule biosynthetic process (GO:0010556) | 3266 | 452 | 309.52 | 1.46 | 1.35E-12 |
| cellular developmental process (GO:0048869) | 3769 | 521 | 357.19 | 1.46 | 1.07E-14 |
| regulation of locomotion (GO:0040012) | 1015 | 140 | 96.19 | 1.46 | 2.48E-03 |
| anatomical structure development (GO:0048856) | 5183 | 714 | 491.19 | 1.45 | 7.87E-22 |
| developmental process (GO:0032502) | 5470 | 751 | 518.39 | 1.45 | 5.47E-23 |
| regulation of cell adhesion (GO:0030155) | 685 | 94 | 64.92 | 1.45 | 3.39E-02 |
| regulation of gene expression (GO:0010468) | 3652 | 501 | 346.1 | 1.45 | 1.10E-13 |
| organelle organization (GO:0006996) | 3000 | 408 | 284.31 | 1.44 | 2.59E-10 |
| regulation of cellular biosynthetic process (GO:0031326) | 3407 | 460 | 322.88 | 1.42 | 1.92E-11 |
| regulation of biosynthetic process (GO:0009889) | 3480 | 469 | 329.8 | 1.42 | 1.42E-11 |
| positive regulation of nitrogen compound metabolic process (GO:0051173) | 3022 | 407 | 286.4 | 1.42 | 8.67E-10 |
| positive regulation of macromolecule metabolic process (GO:0010604) | 3183 | 428 | 301.65 | 1.42 | 2.69E-10 |
| cell-cell signaling (GO:0007267) | 784 | 105 | 74.3 | 1.41 | 3.61E-02 |
| cellular protein modification process (GO:0006464) | 2264 | 303 | 214.56 | 1.41 | 1.24E-06 |
| protein modification process (GO:0036211) | 2264 | 303 | 214.56 | 1.41 | 1.23E-06 |
| positive regulation of cellular metabolic process (GO:0031325) | 3182 | 425 | 301.56 | 1.41 | 7.81E-10 |
| phosphorylation (GO:0016310) | 985 | 131 | 93.35 | 1.4 | 1.35E-02 |
| regulation of cellular localization (GO:0060341) | 955 | 127 | 90.51 | 1.4 | 1.65E-02 |
| positive regulation of metabolic process (GO:0009893) | 3456 | 457 | 327.53 | 1.4 | 3.61E-10 |
| regulation of multicellular organismal process (GO:0051239) | 3161 | 417 | 299.57 | 1.39 | 5.87E-09 |

| | | | | | |
|---|---|---|---|---|---|
| negative regulation of cell communication (GO:0010648) | 1286 | 169 | 121.87 | 1.39 | 4.11E-03 |
| positive regulation of multicellular organismal process (GO:0051240) | 1893 | 248 | 179.4 | 1.38 | 1.01E-04 |
| negative regulation of signal transduction (GO:0009968) | 1168 | 153 | 110.69 | 1.38 | 8.83E-03 |
| negative regulation of signaling (GO:0023057) | 1291 | 169 | 122.35 | 1.38 | 4.25E-03 |
| macromolecule modification (GO:0043412) | 2438 | 319 | 231.05 | 1.38 | 3.28E-06 |
| negative regulation of macromolecule metabolic process (GO:0010605) | 2428 | 316 | 230.1 | 1.37 | 5.33E-06 |
| positive regulation of cell population proliferation (GO:0008284) | 963 | 125 | 91.26 | 1.37 | 3.89E-02 |
| reproductive process (GO:0022414) | 1390 | 180 | 131.73 | 1.37 | 4.14E-03 |
| reproduction (GO:0000003) | 1391 | 180 | 131.83 | 1.37 | 4.92E-03 |
| cellular macromolecule localization (GO:0070727) | 1382 | 178 | 130.97 | 1.36 | 5.60E-03 |
| negative regulation of nitrogen compound metabolic process (GO:0051172) | 2207 | 284 | 209.16 | 1.36 | 6.70E-05 |
| negative regulation of multicellular organismal process (GO:0051241) | 1298 | 167 | 123.01 | 1.36 | 9.88E-03 |
| cellular component organization (GO:0016043) | 5145 | 661 | 487.59 | 1.36 | 1.13E-13 |
| cellular protein localization (GO:0034613) | 1375 | 176 | 130.31 | 1.35 | 8.58E-03 |
| negative regulation of cellular metabolic process (GO:0031324) | 2408 | 308 | 228.21 | 1.35 | 3.45E-05 |
| intracellular transport (GO:0046907) | 1142 | 146 | 108.23 | 1.35 | 2.87E-02 |
| regulation of cell population proliferation (GO:0042127) | 1629 | 208 | 154.38 | 1.35 | 2.72E-03 |
| protein localization (GO:0008104) | 1907 | 243 | 180.73 | 1.34 | 7.35E-04 |
| regulation of macromolecule metabolic process (GO:0060255) | 5277 | 672 | 500.1 | 1.34 | 3.03E-13 |
| regulation of cellular metabolic process (GO:0031323) | 5335 | 674 | 505.6 | 1.33 | 1.07E-12 |
| regulation of signal transduction (GO:0009966) | 2763 | 349 | 261.85 | 1.33 | 1.33E-05 |
| negative regulation of metabolic process (GO:0009892) | 2686 | 339 | 254.55 | 1.33 | 2.32E-05 |
| intracellular signal transduction (GO:0035556) | 1388 | 175 | 131.54 | 1.33 | 1.46E-02 |

| | | | | | |
|---|---|---|---|---|---|
| cellular localization (GO:0051641) | 2040 | 257 | 193.33 | 1.33 | 7.68E-04 |
| regulation of nitrogen compound metabolic process (GO:0051171) | 4995 | 629 | 473.38 | 1.33 | 2.24E-11 |
| cell surface receptor signaling pathway (GO:0007166) | 1804 | 227 | 170.97 | 1.33 | 2.49E-03 |
| cellular component organization or biogenesis (GO:0071840) | 5335 | 670 | 505.6 | 1.33 | 4.27E-12 |
| regulation of primary metabolic process (GO:0080090) | 5143 | 645 | 487.4 | 1.32 | 1.94E-11 |
| regulation of metabolic process (GO:0019222) | 5740 | 719 | 543.98 | 1.32 | 4.51E-13 |
| positive regulation of cellular process (GO:0048522) | 5297 | 659 | 502 | 1.31 | 3.60E-11 |
| regulation of signaling (GO:0023051) | 3192 | 397 | 302.51 | 1.31 | 7.56E-06 |
| regulation of cell communication (GO:0010646) | 3177 | 395 | 301.09 | 1.31 | 8.32E-06 |
| negative regulation of cellular process (GO:0048523) | 4522 | 558 | 428.55 | 1.3 | 1.96E-08 |
| cellular component assembly (GO:0022607) | 2184 | 269 | 206.98 | 1.3 | 1.82E-03 |
| establishment of localization in cell (GO:0051649) | 1441 | 177 | 136.56 | 1.3 | 3.99E-02 |
| negative regulation of response to stimulus (GO:0048585) | 1565 | 190 | 148.32 | 1.28 | 3.87E-02 |
| regulation of localization (GO:0032879) | 2827 | 343 | 267.92 | 1.28 | 4.40E-04 |
| macromolecule localization (GO:0033036) | 2208 | 267 | 209.25 | 1.28 | 5.86E-03 |
| positive regulation of biological process (GO:0048518) | 5994 | 716 | 568.05 | 1.26 | 3.19E-09 |
| multicellular organismal process (GO:0032501) | 7231 | 863 | 685.28 | 1.26 | 4.42E-12 |
| negative regulation of biological process (GO:0048519) | 5045 | 600 | 478.12 | 1.25 | 5.89E-07 |
| regulation of biological quality (GO:0065008) | 3885 | 460 | 368.18 | 1.25 | 8.49E-05 |
| cellular macromolecule metabolic process (GO:0044260) | 3973 | 467 | 376.52 | 1.24 | 1.50E-04 |
| cellular component biogenesis (GO:0044085) | 2406 | 282 | 228.02 | 1.24 | 1.94E-02 |
| cellular protein metabolic process (GO:0044267) | 2863 | 330 | 271.33 | 1.22 | 1.64E-02 |
| regulation of response to stimulus (GO:0048583) | 3909 | 441 | 370.46 | 1.19 | 8.34E-03 |
| protein metabolic process (GO:0019538) | 3442 | 384 | 326.2 | 1.18 | 3.95E-02 |
| regulation of cellular process (GO:0050794) | 10522 | 1172 | 997.17 | 1.18 | 9.91E-11 |
| localization (GO:0051179) | 4795 | 531 | 454.42 | 1.17 | 6.99E-03 |

| | | | | | |
|---|---|---|---|---|---|
| macromolecule metabolic process (GO:0043170) | 5051 | 559 | 478.69 | 1.17 | 5.00E-03 |
| regulation of biological process (GO:0050789) | 11232 | 1241 | 1064.46 | 1.17 | 5.64E-11 |
| biological regulation (GO:0065007) | 11843 | 1294 | 1122.37 | 1.15 | 1.69E-10 |
| cellular process (GO:0009987) | 13986 | 1442 | 1325.46 | 1.09 | 3.87E-05 |
| biological_process (GO:0008150) | 20380 | 2000 | 1931.42 | 1.04 | 6.24E-06 |
| immune system process (GO:0002376) | 2274 | 162 | 215.51 | 0.75 | 7.05E-03 |
| G protein-coupled receptor signaling pathway (GO:0007186) | 1830 | 117 | 173.43 | 0.67 | 4.50E-04 |
| response to bacterium (GO:0009617) | 811 | 49 | 76.86 | 0.64 | 4.39E-02 |
| Unclassified (UNCLASSIFIED) | 1916 | 113 | 181.58 | 0.62 | 6.20E-06 |
| response to biotic stimulus (GO:0009607) | 1306 | 77 | 123.77 | 0.62 | 7.14E-04 |
| response to other organism (GO:0051707) | 1282 | 71 | 121.5 | 0.58 | 1.01E-04 |
| response to external biotic stimulus (GO:0043207) | 1284 | 71 | 121.69 | 0.58 | 1.01E-04 |
| immune effector process (GO:0002252) | 641 | 35 | 60.75 | 0.58 | 2.53E-02 |
| sensory perception of smell (GO:0007608) | 1122 | 57 | 106.33 | 0.54 | 3.61E-05 |
| defense response (GO:0006952) | 1317 | 66 | 124.81 | 0.53 | 2.03E-06 |
| inflammatory response (GO:0006954) | 446 | 22 | 42.27 | 0.52 | 4.04E-02 |
| sensory perception of chemical stimulus (GO:0007606) | 1220 | 60 | 115.62 | 0.52 | 3.41E-06 |
| immune response (GO:0006955) | 1392 | 68 | 131.92 | 0.52 | 3.10E-07 |
| peptide metabolic process (GO:0006518) | 451 | 22 | 42.74 | 0.51 | 3.36E-02 |
| defense response to bacterium (GO:0042742) | 461 | 21 | 43.69 | 0.48 | 1.32E-02 |
| humoral immune response (GO:0006959) | 374 | 16 | 35.44 | 0.45 | 2.73E-02 |
| peptide biosynthetic process (GO:0043043) | 329 | 14 | 31.18 | 0.45 | 4.49E-02 |
| defense response to other organism (GO:0098542) | 959 | 40 | 90.88 | 0.44 | 9.05E-07 |
| translation (GO:0006412) | 310 | 12 | 29.38 | 0.41 | 2.53E-02 |
| innate immune response (GO:0045087) | 733 | 28 | 69.47 | 0.4 | 6.25E-06 |
| production of molecular mediator of immune response (GO:0002440) | 281 | 10 | 26.63 | 0.38 | 2.69E-02 |
| immunoglobulin production (GO:0002377) | 257 | 9 | 24.36 | 0.37 | 3.34E-02 |
| antimicrobial humoral response (GO:0019730) | 104 | 1 | 9.86 | 0.1 | 4.76E-02 |

| Purple module | Mus musculus genes (22296) | Module genes (711) | Module genes (expected) | Module genes fold Enrichment | Module genes FDR |
|---|---|---|---|---|---|
| regulation of protein homooligomerization (GO:0032462) | 22 | 6 | 0.6 | 10.07 | 1.54E-02 |
| collagen catabolic process (GO:0030574) | 32 | 7 | 0.87 | 8.07 | 1.33E-02 |
| regulation of protein oligomerization (GO:0032459) | 41 | 7 | 1.11 | 6.3 | 3.82E-02 |
| collagen metabolic process (GO:0032963) | 57 | 8 | 1.54 | 5.18 | 4.43E-02 |
| neural tube closure (GO:0001843) | 109 | 11 | 2.95 | 3.73 | 4.96E-02 |
| regulation of histone modification (GO:0031056) | 150 | 14 | 4.06 | 3.45 | 2.26E-02 |
| regulation of chromatin organization (GO:1902275) | 194 | 18 | 5.26 | 3.43 | 4.28E-03 |
| epithelial tube morphogenesis (GO:0060562) | 356 | 26 | 9.64 | 2.7 | 3.72E-03 |
| regulation of neuron apoptotic process (GO:0043523) | 263 | 19 | 7.12 | 2.67 | 3.18E-02 |
| morphogenesis of an epithelium (GO:0002009) | 498 | 31 | 13.49 | 2.3 | 1.06E-02 |
| negative regulation of protein modification process (GO:0031400) | 562 | 33 | 15.22 | 2.17 | 1.46E-02 |
| negative regulation of transcription by RNA polymerase II (GO:0000122) | 852 | 47 | 23.08 | 2.04 | 3.50E-03 |
| tissue morphogenesis (GO:0048729) | 623 | 34 | 16.88 | 2.01 | 4.06E-02 |
| tube morphogenesis (GO:0035239) | 708 | 38 | 19.18 | 1.98 | 2.09E-02 |
| negative regulation of cellular component organization (GO:0051129) | 717 | 38 | 19.42 | 1.96 | 2.95E-02 |
| negative regulation of transcription, DNA-templated (GO:0045892) | 1149 | 60 | 31.13 | 1.93 | 1.65E-03 |
| negative regulation of nucleic acid-templated transcription (GO:1903507) | 1153 | 60 | 31.23 | 1.92 | 1.67E-03 |
| negative regulation of RNA biosynthetic process (GO:1902679) | 1154 | 60 | 31.26 | 1.92 | 1.58E-03 |
| tube development (GO:0035295) | 912 | 47 | 24.71 | 1.9 | 1.14E-02 |
| negative regulation of RNA metabolic process (GO:0051253) | 1240 | 63 | 33.59 | 1.88 | 1.92E-03 |
| negative regulation of nucleobase-containing compound metabolic process (GO:0045934) | 1343 | 68 | 36.38 | 1.87 | 1.40E-03 |

| | | | | | |
|---|---|---|---|---|---|
| regulation of organelle organization (GO:0033043) | 1225 | 60 | 33.19 | 1.81 | 5.78E-03 |
| negative regulation of macromolecule biosynthetic process (GO:0010558) | 1376 | 67 | 37.28 | 1.8 | 3.06E-03 |
| negative regulation of cellular macromolecule biosynthetic process (GO:2000113) | 1341 | 65 | 36.33 | 1.79 | 3.50E-03 |
| intracellular transport (GO:0046907) | 1142 | 55 | 30.94 | 1.78 | 1.32E-02 |
| negative regulation of biosynthetic process (GO:0009890) | 1464 | 70 | 39.66 | 1.77 | 2.70E-03 |
| negative regulation of cellular biosynthetic process (GO:0031327) | 1426 | 68 | 38.63 | 1.76 | 3.66E-03 |
| regulation of transcription by RNA polymerase II (GO:0006357) | 1966 | 92 | 53.26 | 1.73 | 6.23E-04 |
| protein transport (GO:0015031) | 1230 | 57 | 33.32 | 1.71 | 2.62E-02 |
| amide transport (GO:0042886) | 1279 | 59 | 34.65 | 1.7 | 2.48E-02 |
| peptide transport (GO:0015833) | 1258 | 58 | 34.08 | 1.7 | 2.35E-02 |
| regulation of cellular component organization (GO:0051128) | 2464 | 113 | 66.75 | 1.69 | 9.30E-05 |
| response to oxygen-containing compound (GO:1901700) | 1265 | 58 | 34.27 | 1.69 | 3.03E-02 |
| positive regulation of transcription, DNA-templated (GO:0045893) | 1527 | 70 | 41.37 | 1.69 | 8.63E-03 |
| positive regulation of nucleic acid-templated transcription (GO:1903508) | 1531 | 70 | 41.47 | 1.69 | 8.73E-03 |
| positive regulation of RNA biosynthetic process (GO:1902680) | 1532 | 70 | 41.5 | 1.69 | 8.65E-03 |
| negative regulation of cellular metabolic process (GO:0031324) | 2408 | 110 | 65.23 | 1.69 | 1.63E-04 |
| organic substance transport (GO:0071702) | 1842 | 84 | 49.9 | 1.68 | 2.14E-03 |
| establishment of protein localization (GO:0045184) | 1315 | 59 | 35.62 | 1.66 | 3.73E-02 |
| negative regulation of nitrogen compound metabolic process (GO:0051172) | 2207 | 99 | 59.79 | 1.66 | 1.23E-03 |

| | | | | | |
|---|---|---|---|---|---|
| positive regulation of RNA metabolic process (GO:0051254) | 1629 | 73 | 44.13 | 1.65 | 9.63E-03 |
| nitrogen compound transport (GO:0071705) | 1526 | 68 | 41.34 | 1.64 | 1.79E-02 |
| negative regulation of gene expression (GO:0010629) | 1588 | 70 | 43.02 | 1.63 | 1.80E-02 |
| regulation of protein modification process (GO:0031399) | 1756 | 77 | 47.57 | 1.62 | 1.14E-02 |
| negative regulation of macromolecule metabolic process (GO:0010605) | 2428 | 106 | 65.77 | 1.61 | 1.21E-03 |
| macromolecule localization (GO:0033036) | 2208 | 95 | 59.81 | 1.59 | 3.50E-03 |
| negative regulation of metabolic process (GO:0009892) | 2686 | 115 | 72.76 | 1.58 | 1.23E-03 |
| positive regulation of protein metabolic process (GO:0051247) | 1635 | 70 | 44.29 | 1.58 | 3.36E-02 |
| positive regulation of macromolecule biosynthetic process (GO:0010557) | 1788 | 76 | 48.44 | 1.57 | 2.61E-02 |
| regulation of molecular function (GO:0065009) | 2476 | 105 | 67.07 | 1.57 | 2.49E-03 |
| regulation of transcription, DNA-templated (GO:0006355) | 2755 | 116 | 74.63 | 1.55 | 1.64E-03 |
| regulation of nucleic acid-templated transcription (GO:1903506) | 2763 | 116 | 74.85 | 1.55 | 1.64E-03 |
| protein localization (GO:0008104) | 1907 | 80 | 51.66 | 1.55 | 2.41E-02 |
| regulation of RNA biosynthetic process (GO:2001141) | 2766 | 116 | 74.93 | 1.55 | 1.60E-03 |
| nervous system development (GO:0007399) | 2226 | 93 | 60.3 | 1.54 | 9.73E-03 |
| regulation of protein metabolic process (GO:0051246) | 2661 | 111 | 72.09 | 1.54 | 2.58E-03 |
| positive regulation of cellular biosynthetic process (GO:0031328) | 1882 | 78 | 50.98 | 1.53 | 4.19E-02 |
| negative regulation of cellular process (GO:0048523) | 4522 | 187 | 122.5 | 1.53 | 3.97E-06 |
| cellular localization (GO:0051641) | 2040 | 84 | 55.26 | 1.52 | 2.73E-02 |
| positive regulation of nitrogen compound metabolic process (GO:0051173) | 3022 | 124 | 81.87 | 1.51 | 1.68E-03 |
| positive regulation of gene expression (GO:0010628) | 1954 | 80 | 52.93 | 1.51 | 4.18E-02 |
| regulation of cellular protein metabolic process (GO:0032268) | 2470 | 101 | 66.91 | 1.51 | 1.12E-02 |

| | | | | | |
|---|---|---|---|---|---|
| positive regulation of cellular metabolic process (GO:0031325) | 3182 | 130 | 86.2 | 1.51 | 1.58E-03 |
| positive regulation of macromolecule metabolic process (GO:0010604) | 3183 | 129 | 86.23 | 1.5 | 1.64E-03 |
| regulation of macromolecule biosynthetic process (GO:0010556) | 3266 | 132 | 88.48 | 1.49 | 1.63E-03 |
| regulation of developmental process (GO:0050793) | 2635 | 106 | 71.38 | 1.48 | 1.10E-02 |
| positive regulation of metabolic process (GO:0009893) | 3456 | 139 | 93.62 | 1.48 | 1.46E-03 |
| cellular protein modification process (GO:0006464) | 2264 | 91 | 61.33 | 1.48 | 3.67E-02 |
| protein modification process (GO:0036211) | 2264 | 91 | 61.33 | 1.48 | 3.63E-02 |
| regulation of cellular macromolecule biosynthetic process (GO:2000112) | 3180 | 127 | 86.15 | 1.47 | 3.52E-03 |
| regulation of RNA metabolic process (GO:0051252) | 3011 | 120 | 81.57 | 1.47 | 5.40E-03 |
| transport (GO:0006810) | 3523 | 140 | 95.44 | 1.47 | 1.65E-03 |
| positive regulation of cellular process (GO:0048522) | 5297 | 210 | 143.5 | 1.46 | 5.76E-06 |
| protein metabolic process (GO:0019538) | 3442 | 136 | 93.24 | 1.46 | 2.61E-03 |
| animal organ development (GO:0048513) | 3040 | 120 | 82.35 | 1.46 | 8.82E-03 |
| regulation of cellular biosynthetic process (GO:0031326) | 3407 | 134 | 92.3 | 1.45 | 3.46E-03 |
| regulation of biosynthetic process (GO:0009889) | 3480 | 136 | 94.27 | 1.44 | 3.79E-03 |
| regulation of localization (GO:0032879) | 2827 | 110 | 76.58 | 1.44 | 2.32E-02 |
| localization (GO:0051179) | 4795 | 186 | 129.9 | 1.43 | 2.51E-04 |
| establishment of localization (GO:0051234) | 3664 | 142 | 99.26 | 1.43 | 3.65E-03 |
| negative regulation of biological process (GO:0048519) | 5045 | 195 | 136.67 | 1.43 | 1.36E-04 |
| positive regulation of biological process (GO:0048518) | 5994 | 231 | 162.38 | 1.42 | 5.49E-06 |
| regulation of nucleobase-containing compound metabolic process (GO:0019219) | 3242 | 124 | 87.83 | 1.41 | 1.77E-02 |
| regulation of biological quality (GO:0065008) | 3885 | 148 | 105.24 | 1.41 | 5.14E-03 |
| system development (GO:0048731) | 4205 | 160 | 113.91 | 1.4 | 2.51E-03 |
| regulation of multicellular organismal process (GO:0051239) | 3161 | 120 | 85.63 | 1.4 | 2.71E-02 |
| regulation of primary metabolic process (GO:0080090) | 5143 | 195 | 139.32 | 1.4 | 4.62E-04 |

| | | | | | |
|---|---|---|---|---|---|
| regulation of nitrogen compound metabolic process (GO:0051171) | 4995 | 188 | 135.31 | 1.39 | 1.30E-03 |
| regulation of gene expression (GO:0010468) | 3652 | 135 | 98.93 | 1.36 | 3.17E-02 |
| regulation of macromolecule metabolic process (GO:0060255) | 5277 | 195 | 142.95 | 1.36 | 1.44E-03 |
| regulation of cellular metabolic process (GO:0031323) | 5335 | 197 | 144.53 | 1.36 | 1.47E-03 |
| organonitrogen compound metabolic process (GO:1901564) | 4283 | 157 | 116.03 | 1.35 | 1.28E-02 |
| regulation of metabolic process (GO:0019222) | 5740 | 209 | 155.5 | 1.34 | 1.43E-03 |
| cellular component organization (GO:0016043) | 5145 | 187 | 139.38 | 1.34 | 4.01E-03 |
| multicellular organism development (GO:0007275) | 4810 | 173 | 130.3 | 1.33 | 1.30E-02 |
| cellular component organization or biogenesis (GO:0071840) | 5335 | 189 | 144.53 | 1.31 | 1.15E-02 |
| anatomical structure development (GO:0048856) | 5183 | 180 | 140.41 | 1.28 | 3.89E-02 |
| metabolic process (GO:0008152) | 7193 | 244 | 194.86 | 1.25 | 8.90E-03 |
| primary metabolic process (GO:0044238) | 6255 | 212 | 169.45 | 1.25 | 3.29E-02 |
| regulation of cellular process (GO:0050794) | 10522 | 335 | 285.04 | 1.18 | 1.31E-02 |
| regulation of biological process (GO:0050789) | 11232 | 353 | 304.28 | 1.16 | 2.00E-02 |
| biological regulation (GO:0065007) | 11843 | 368 | 320.83 | 1.15 | 2.53E-02 |
| cellular process (GO:0009987) | 13986 | 427 | 378.88 | 1.13 | 1.29E-02 |
| nervous system process (GO:0050877) | 2065 | 24 | 55.94 | 0.43 | 1.29E-03 |
| G protein-coupled receptor signaling pathway (GO:0007186) | 1830 | 14 | 49.57 | 0.28 | 5.54E-06 |
| sensory perception (GO:0007600) | 1624 | 6 | 43.99 | 0.14 | 6.94E-09 |
| sensory perception of smell (GO:0007608) | 1122 | 1 | 30.4 | 0.03 | 1.28E-08 |
| sensory perception of chemical stimulus (GO:0007606) | 1220 | 1 | 33.05 | 0.03 | 3.69E-09 |
| **Light green module** | **Mus musculus genes (22296)** | **Module genes (711)** | **Module genes (expected)** | **Module genes fold Enrichment** | **Module genes FDR** |
| negative regulation of cytoplasmic translation (GO:2000766) | 8 | 3 | 0.09 | 32.79 | 2.03E-02 |

| | | | | | |
|---|---|---|---|---|---|
| negative regulation of transcription by competitive promoter binding (GO:0010944) | 9 | 3 | 0.1 | 29.15 | 2.49E-02 |
| development of secondary sexual characteristics (GO:0045136) | 13 | 3 | 0.15 | 20.18 | 4.94E-02 |
| regulation of establishment of cell polarity (GO:2000114) | 19 | 4 | 0.22 | 18.41 | 1.32E-02 |
| adherens junction assembly (GO:0034333) | 34 | 7 | 0.39 | 18 | 1.32E-04 |
| regulation of establishment or maintenance of cell polarity (GO:0032878) | 24 | 4 | 0.27 | 14.57 | 2.37E-02 |
| cell-substrate adherens junction assembly (GO:0007045) | 25 | 4 | 0.29 | 13.99 | 2.57E-02 |
| regulation of protein exit from endoplasmic reticulum (GO:0070861) | 25 | 4 | 0.29 | 13.99 | 2.56E-02 |
| focal adhesion assembly (GO:0048041) | 25 | 4 | 0.29 | 13.99 | 2.55E-02 |
| negative regulation of endothelial cell apoptotic process (GO:2000352) | 31 | 4 | 0.35 | 11.28 | 4.51E-02 |
| epidermal growth factor receptor signaling pathway (GO:0007173) | 40 | 5 | 0.46 | 10.93 | 1.57E-02 |
| negative regulation of response to endoplasmic reticulum stress (GO:1903573) | 42 | 5 | 0.48 | 10.41 | 1.78E-02 |
| regulation of response to endoplasmic reticulum stress (GO:1905897) | 77 | 9 | 0.88 | 10.22 | 1.75E-04 |
| ERBB signaling pathway (GO:0038127) | 45 | 5 | 0.51 | 9.72 | 2.27E-02 |
| adherens junction organization (GO:0034332) | 70 | 7 | 0.8 | 8.74 | 4.05E-03 |
| negative regulation of fat cell differentiation (GO:0045599) | 53 | 5 | 0.61 | 8.25 | 3.81E-02 |
| negative regulation of transforming growth factor beta receptor signaling pathway (GO:0030512) | 64 | 6 | 0.73 | 8.2 | 1.48E-02 |
| negative regulation of cellular response to transforming growth factor beta stimulus (GO:1903845) | 66 | 6 | 0.75 | 7.95 | 1.68E-02 |
| positive regulation of blood vessel endothelial cell migration (GO:0043536) | 56 | 5 | 0.64 | 7.81 | 4.50E-02 |
| peptidyl-threonine phosphorylation (GO:0018107) | 83 | 6 | 0.95 | 6.32 | 3.89E-02 |

| | | | | | |
|---|---|---|---|---|---|
| positive regulation of endothelial cell migration (GO:0010595) | 97 | 7 | 1.11 | 6.31 | 1.78E-02 |
| negative regulation of transmembrane receptor protein serine/threonine kinase signaling pathway (GO:0090101) | 111 | 8 | 1.27 | 6.3 | 8.11E-03 |
| negative regulation of ossification (GO:0030279) | 84 | 6 | 0.96 | 6.25 | 4.04E-02 |
| epithelial cell proliferation (GO:0050673) | 99 | 7 | 1.13 | 6.18 | 1.92E-02 |
| negative regulation of cellular response to growth factor stimulus (GO:0090288) | 142 | 10 | 1.62 | 6.16 | 1.83E-03 |
| cell junction assembly (GO:0034329) | 131 | 9 | 1.5 | 6.01 | 4.72E-03 |
| regulation of transforming growth factor beta receptor signaling pathway (GO:0017015) | 103 | 7 | 1.18 | 5.94 | 2.32E-02 |
| gland morphogenesis (GO:0022612) | 120 | 8 | 1.37 | 5.83 | 1.21E-02 |
| hair cycle (GO:0042633) | 105 | 7 | 1.2 | 5.83 | 2.51E-02 |
| regulation of cellular response to transforming growth factor beta stimulus (GO:1903844) | 105 | 7 | 1.2 | 5.83 | 2.49E-02 |
| molting cycle (GO:0042303) | 105 | 7 | 1.2 | 5.83 | 2.48E-02 |
| positive regulation of protein catabolic process (GO:0045732) | 210 | 12 | 2.4 | 5 | 1.81E-03 |
| cell junction organization (GO:0034330) | 178 | 10 | 2.04 | 4.91 | 7.66E-03 |
| mammary gland development (GO:0030879) | 149 | 8 | 1.7 | 4.69 | 3.41E-02 |
| regulation of protein catabolic process (GO:0042176) | 379 | 20 | 4.33 | 4.61 | 2.34E-05 |
| regulation of endothelial cell migration (GO:0010594) | 155 | 8 | 1.77 | 4.51 | 4.03E-02 |
| regulation of cellular protein catabolic process (GO:1903362) | 240 | 12 | 2.74 | 4.37 | 4.76E-03 |
| regulation of canonical Wnt signaling pathway (GO:0060828) | 224 | 11 | 2.56 | 4.29 | 9.74E-03 |
| response to endoplasmic reticulum stress (GO:0034976) | 206 | 10 | 2.36 | 4.24 | 1.80E-02 |
| regulation of cellular response to growth factor stimulus (GO:0090287) | 270 | 13 | 3.09 | 4.21 | 3.57E-03 |
| regulation of proteolysis involved in cellular protein catabolic process (GO:1903050) | 209 | 10 | 2.39 | 4.18 | 1.93E-02 |

| | | | | | |
|---|---|---|---|---|---|
| positive regulation of catabolic process (GO:0009896) | 422 | 20 | 4.83 | 4.14 | 7.76E-05 |
| epithelial cell development (GO:0002064) | 216 | 10 | 2.47 | 4.05 | 2.37E-02 |
| positive regulation of epithelial cell proliferation (GO:0050679) | 198 | 9 | 2.26 | 3.97 | 4.38E-02 |
| positive regulation of cellular catabolic process (GO:0031331) | 354 | 16 | 4.05 | 3.95 | 1.12E-03 |
| regulation of protein stability (GO:0031647) | 275 | 12 | 3.15 | 3.82 | 1.26E-02 |
| regulation of transmembrane receptor protein serine/threonine kinase signaling pathway (GO:0090092) | 231 | 10 | 2.64 | 3.79 | 3.47E-02 |
| regulation of Wnt signaling pathway (GO:0030111) | 290 | 12 | 3.32 | 3.62 | 1.78E-02 |
| apoptotic signaling pathway (GO:0097190) | 275 | 11 | 3.15 | 3.5 | 3.47E-02 |
| negative regulation of cell differentiation (GO:0045596) | 757 | 30 | 8.66 | 3.47 | 7.43E-06 |
| positive regulation of protein kinase activity (GO:0045860) | 434 | 17 | 4.96 | 3.42 | 2.84E-03 |
| positive regulation of cell migration (GO:0030335) | 539 | 20 | 6.16 | 3.24 | 1.28E-03 |
| positive regulation of proteolysis (GO:0045862) | 325 | 12 | 3.72 | 3.23 | 3.73E-02 |
| regulation of catabolic process (GO:0009894) | 819 | 30 | 9.37 | 3.2 | 2.47E-05 |
| positive regulation of cellular component movement (GO:0051272) | 579 | 21 | 6.62 | 3.17 | 1.10E-03 |
| positive regulation of kinase activity (GO:0033674) | 472 | 17 | 5.4 | 3.15 | 6.12E-03 |
| cell migration (GO:0016477) | 840 | 30 | 9.61 | 3.12 | 3.60E-05 |
| positive regulation of cell motility (GO:2000147) | 561 | 20 | 6.42 | 3.12 | 2.11E-03 |
| regulation of cellular catabolic process (GO:0031329) | 676 | 24 | 7.73 | 3.1 | 3.70E-04 |
| regulation of cellular amide metabolic process (GO:0034248) | 368 | 13 | 4.21 | 3.09 | 3.42E-02 |
| positive regulation of transferase activity (GO:0051347) | 547 | 19 | 6.26 | 3.04 | 4.04E-03 |
| actin cytoskeleton organization (GO:0030036) | 474 | 16 | 5.42 | 2.95 | 1.64E-02 |
| positive regulation of locomotion (GO:0040017) | 593 | 20 | 6.78 | 2.95 | 3.74E-03 |
| localization of cell (GO:0051674) | 958 | 32 | 10.96 | 2.92 | 5.18E-05 |
| cell motility (GO:0048870) | 958 | 32 | 10.96 | 2.92 | 5.00E-05 |

| | | | | | |
|---|---|---|---|---|---|
| in utero embryonic development (GO:0001701) | 516 | 17 | 5.9 | 2.88 | 1.42E-02 |
| epithelial cell differentiation (GO:0030855) | 517 | 17 | 5.91 | 2.88 | 1.43E-02 |
| gland development (GO:0048732) | 428 | 14 | 4.9 | 2.86 | 4.01E-02 |
| regulation of cellular protein localization (GO:1903827) | 544 | 17 | 6.22 | 2.73 | 2.19E-02 |
| negative regulation of cellular component organization (GO:0051129) | 717 | 22 | 8.2 | 2.68 | 5.30E-03 |
| regulation of cell adhesion (GO:0030155) | 685 | 21 | 7.83 | 2.68 | 7.61E-03 |
| negative regulation of macromolecule biosynthetic process (GO:0010558) | 1376 | 42 | 15.74 | 2.67 | 9.13E-06 |
| regulation of cell migration (GO:0030334) | 889 | 27 | 10.17 | 2.66 | 1.18E-03 |
| enzyme linked receptor protein signaling pathway (GO:0007167) | 528 | 16 | 6.04 | 2.65 | 3.81E-02 |
| regulation of protein kinase activity (GO:0045859) | 660 | 20 | 7.55 | 2.65 | 1.16E-02 |
| regulation of cellular component movement (GO:0051270) | 1024 | 31 | 11.71 | 2.65 | 3.02E-04 |
| regulation of cellular response to stress (GO:0080135) | 664 | 20 | 7.59 | 2.63 | 1.24E-02 |
| actin filament-based process (GO:0030029) | 532 | 16 | 6.08 | 2.63 | 4.02E-02 |
| negative regulation of signal transduction (GO:0009968) | 1168 | 35 | 13.36 | 2.62 | 1.10E-04 |
| negative regulation of cellular macromolecule biosynthetic process (GO:2000113) | 1341 | 40 | 15.34 | 2.61 | 2.66E-05 |
| sensory organ development (GO:0007423) | 571 | 17 | 6.53 | 2.6 | 3.31E-02 |
| negative regulation of transcription, DNA-templated (GO:0045892) | 1149 | 34 | 13.14 | 2.59 | 1.73E-04 |
| negative regulation of cell communication (GO:0010648) | 1286 | 38 | 14.71 | 2.58 | 5.95E-05 |
| negative regulation of nucleic acid-templated transcription (GO:1903507) | 1153 | 34 | 13.19 | 2.58 | 1.79E-04 |
| negative regulation of RNA biosynthetic process (GO:1902679) | 1154 | 34 | 13.2 | 2.58 | 1.80E-04 |
| negative regulation of signaling (GO:0023057) | 1291 | 38 | 14.77 | 2.57 | 6.16E-05 |
| negative regulation of transcription by RNA polymerase II (GO:0000122) | 852 | 25 | 9.74 | 2.57 | 3.63E-03 |

| | | | | | |
|---|---|---|---|---|---|
| negative regulation of developmental process (GO:0051093) | 1023 | 30 | 11.7 | 2.56 | 7.40E-04 |
| locomotion (GO:0040011) | 1160 | 34 | 13.27 | 2.56 | 1.98E-04 |
| negative regulation of RNA metabolic process (GO:0051253) | 1240 | 36 | 14.18 | 2.54 | 1.39E-04 |
| regulation of kinase activity (GO:0043549) | 727 | 21 | 8.31 | 2.53 | 1.42E-02 |
| regulation of transferase activity (GO:0051338) | 831 | 24 | 9.5 | 2.53 | 5.86E-03 |
| regulation of cell motility (GO:2000145) | 936 | 27 | 10.71 | 2.52 | 2.51E-03 |
| chordate embryonic development (GO:0043009) | 798 | 23 | 9.13 | 2.52 | 8.06E-03 |
| protein catabolic process (GO:0030163) | 626 | 18 | 7.16 | 2.51 | 3.34E-02 |
| negative regulation of biosynthetic process (GO:0009890) | 1464 | 42 | 16.74 | 2.51 | 3.99E-05 |
| cellular macromolecule catabolic process (GO:0044265) | 740 | 21 | 8.46 | 2.48 | 1.69E-02 |
| embryo development ending in birth or egg hatching (GO:0009792) | 815 | 23 | 9.32 | 2.47 | 1.05E-02 |
| negative regulation of cellular biosynthetic process (GO:0031327) | 1426 | 40 | 16.31 | 2.45 | 9.08E-05 |
| animal organ morphogenesis (GO:0009887) | 1005 | 28 | 11.49 | 2.44 | 3.75E-03 |
| movement of cell or subcellular component (GO:0006928) | 1366 | 38 | 15.62 | 2.43 | 1.72E-04 |
| negative regulation of apoptotic process (GO:0043066) | 905 | 25 | 10.35 | 2.42 | 9.73E-03 |
| negative regulation of nucleobase-containing compound metabolic process (GO:0045934) | 1343 | 37 | 15.36 | 2.41 | 2.75E-04 |
| response to nitrogen compound (GO:1901698) | 837 | 23 | 9.57 | 2.4 | 1.75E-02 |
| epithelium development (GO:0060429) | 1029 | 28 | 11.77 | 2.38 | 4.76E-03 |
| negative regulation of programmed cell death (GO:0043069) | 922 | 25 | 10.54 | 2.37 | 1.11E-02 |
| response to organonitrogen compound (GO:0010243) | 738 | 20 | 8.44 | 2.37 | 4.15E-02 |
| positive regulation of intracellular signal transduction (GO:1902533) | 924 | 25 | 10.57 | 2.37 | 1.13E-02 |
| positive regulation of protein metabolic process (GO:0051247) | 1635 | 44 | 18.7 | 2.35 | 8.10E-05 |

| | | | | | |
|---|---|---|---|---|---|
| positive regulation of signal transduction (GO:0009967) | 1499 | 40 | 17.14 | 2.33 | 2.64E-04 |
| regulation of locomotion (GO:0040012) | 1015 | 27 | 11.61 | 2.33 | 7.86E-03 |
| regulation of protein localization (GO:0032880) | 1056 | 28 | 12.08 | 2.32 | 6.15E-03 |
| macromolecule catabolic process (GO:0009057) | 833 | 22 | 9.53 | 2.31 | 2.92E-02 |
| cytoskeleton organization (GO:0007010) | 1028 | 27 | 11.76 | 2.3 | 9.05E-03 |
| anatomical structure formation involved in morphogenesis (GO:0048646) | 914 | 24 | 10.45 | 2.3 | 1.84E-02 |
| negative regulation of cell death (GO:0060548) | 1032 | 27 | 11.8 | 2.29 | 9.51E-03 |
| regulation of intracellular signal transduction (GO:1902531) | 1577 | 41 | 18.04 | 2.27 | 2.75E-04 |
| negative regulation of gene expression (GO:0010629) | 1588 | 41 | 18.16 | 2.26 | 3.04E-04 |
| positive regulation of protein modification process (GO:0031401) | 1208 | 31 | 13.82 | 2.24 | 5.29E-03 |
| positive regulation of cellular protein metabolic process (GO:0032270) | 1528 | 39 | 17.48 | 2.23 | 6.80E-04 |
| positive regulation of protein phosphorylation (GO:0001934) | 985 | 25 | 11.27 | 2.22 | 2.63E-02 |
| positive regulation of signaling (GO:0023056) | 1697 | 43 | 19.41 | 2.22 | 3.06E-04 |
| regulation of signal transduction (GO:0009966) | 2763 | 70 | 31.6 | 2.22 | 6.49E-07 |
| tube development (GO:0035295) | 912 | 23 | 10.43 | 2.21 | 4.62E-02 |
| regulation of cellular localization (GO:0060341) | 955 | 24 | 10.92 | 2.2 | 3.69E-02 |
| embryo development (GO:0009790) | 1155 | 29 | 13.21 | 2.2 | 1.10E-02 |
| negative regulation of nitrogen compound metabolic process (GO:0051172) | 2207 | 55 | 25.24 | 2.18 | 2.65E-05 |
| negative regulation of response to stimulus (GO:0048585) | 1565 | 39 | 17.9 | 2.18 | 1.24E-03 |
| positive regulation of cell communication (GO:0010647) | 1690 | 42 | 19.33 | 2.17 | 5.63E-04 |
| negative regulation of protein metabolic process (GO:0051248) | 1052 | 26 | 12.03 | 2.16 | 2.47E-02 |
| regulation of apoptotic process (GO:0042981) | 1499 | 37 | 17.14 | 2.16 | 2.42E-03 |
| regulation of protein metabolic process (GO:0051246) | 2661 | 65 | 30.43 | 2.14 | 4.54E-06 |

| | | | | | |
|---|---|---|---|---|---|
| neuron differentiation (GO:0030182) | 1024 | 25 | 11.71 | 2.13 | 3.47E-02 |
| negative regulation of cellular protein metabolic process (GO:0032269) | 985 | 24 | 11.27 | 2.13 | 4.39E-02 |
| regulation of programmed cell death (GO:0043067) | 1522 | 37 | 17.41 | 2.13 | 2.87E-03 |
| tissue development (GO:0009888) | 1658 | 40 | 18.96 | 2.11 | 2.10E-03 |
| regulation of signaling (GO:0023051) | 3192 | 77 | 36.51 | 2.11 | 5.10E-07 |
| positive regulation of phosphorylation (GO:0042327) | 1039 | 25 | 11.88 | 2.1 | 3.90E-02 |
| positive regulation of catalytic activity (GO:0043085) | 1043 | 25 | 11.93 | 2.1 | 4.01E-02 |
| regulation of cell communication (GO:0010646) | 3177 | 76 | 36.34 | 2.09 | 7.40E-07 |
| positive regulation of molecular function (GO:0044093) | 1422 | 34 | 16.26 | 2.09 | 7.18E-03 |
| negative regulation of multicellular organismal process (GO:0051241) | 1298 | 31 | 14.85 | 2.09 | 1.44E-02 |
| regulation of localization (GO:0032879) | 2827 | 67 | 32.33 | 2.07 | 7.69E-06 |
| negative regulation of cellular metabolic process (GO:0031324) | 2408 | 57 | 27.54 | 2.07 | 6.59E-05 |
| response to endogenous stimulus (GO:0009719) | 1104 | 26 | 12.63 | 2.06 | 4.81E-02 |
| neurogenesis (GO:0022008) | 1742 | 41 | 19.92 | 2.06 | 2.17E-03 |
| regulation of protein phosphorylation (GO:0001932) | 1405 | 33 | 16.07 | 2.05 | 1.44E-02 |
| positive regulation of phosphorus metabolic process (GO:0010562) | 1107 | 26 | 12.66 | 2.05 | 4.89E-02 |
| positive regulation of phosphate metabolic process (GO:0045937) | 1107 | 26 | 12.66 | 2.05 | 4.86E-02 |
| regulation of cellular protein metabolic process (GO:0032268) | 2470 | 58 | 28.25 | 2.05 | 5.98E-05 |
| regulation of response to stress (GO:0080134) | 1321 | 31 | 15.11 | 2.05 | 1.68E-02 |
| regulation of cell death (GO:0010941) | 1670 | 39 | 19.1 | 2.04 | 3.76E-03 |
| generation of neurons (GO:0048699) | 1636 | 38 | 18.71 | 2.03 | 5.00E-03 |
| negative regulation of macromolecule metabolic process (GO:0010605) | 2428 | 56 | 27.77 | 2.02 | 1.65E-04 |
| intracellular signal transduction (GO:0035556) | 1388 | 32 | 15.87 | 2.02 | 1.99E-02 |
| cell development (GO:0048468) | 1746 | 40 | 19.97 | 2 | 4.73E-03 |

| | | | | | |
|---|---|---|---|---|---|
| response to organic substance (GO:0010033) | 2445 | 56 | 27.96 | 2 | 1.75E-04 |
| positive regulation of nitrogen compound metabolic process (GO:0051173) | 3022 | 69 | 34.56 | 2 | 1.70E-05 |
| regulation of protein modification process (GO:0031399) | 1756 | 40 | 20.08 | 1.99 | 5.01E-03 |
| negative regulation of cellular process (GO:0048523) | 4522 | 103 | 51.72 | 1.99 | 5.50E-09 |
| negative regulation of metabolic process (GO:0009892) | 2686 | 60 | 30.72 | 1.95 | 1.37E-04 |
| positive regulation of cellular metabolic process (GO:0031325) | 3182 | 71 | 36.39 | 1.95 | 1.97E-05 |
| positive regulation of macromolecule metabolic process (GO:0010604) | 3183 | 71 | 36.4 | 1.95 | 1.88E-05 |
| positive regulation of metabolic process (GO:0009893) | 3456 | 77 | 39.53 | 1.95 | 6.59E-06 |
| regulation of phosphorylation (GO:0042325) | 1529 | 34 | 17.49 | 1.94 | 2.32E-02 |
| cell surface receptor signaling pathway (GO:0007166) | 1804 | 40 | 20.63 | 1.94 | 8.86E-03 |
| positive regulation of RNA metabolic process (GO:0051254) | 1629 | 36 | 18.63 | 1.93 | 1.64E-02 |
| regulation of phosphate metabolic process (GO:0019220) | 1684 | 37 | 19.26 | 1.92 | 1.79E-02 |
| regulation of phosphorus metabolic process (GO:0051174) | 1685 | 37 | 19.27 | 1.92 | 1.77E-02 |
| anatomical structure morphogenesis (GO:0009653) | 2188 | 48 | 25.02 | 1.92 | 2.25E-03 |
| positive regulation of nucleobase-containing compound metabolic process (GO:0045935) | 1787 | 39 | 20.44 | 1.91 | 1.25E-02 |
| regulation of cell differentiation (GO:0045595) | 1845 | 40 | 21.1 | 1.9 | 1.45E-02 |
| negative regulation of biological process (GO:0048519) | 5045 | 109 | 57.7 | 1.89 | 1.36E-08 |
| positive regulation of nucleic acid-templated transcription (GO:1903508) | 1531 | 33 | 17.51 | 1.88 | 4.77E-02 |
| positive regulation of RNA biosynthetic process (GO:1902680) | 1532 | 33 | 17.52 | 1.88 | 4.77E-02 |
| positive regulation of response to stimulus (GO:0048584) | 2289 | 49 | 26.18 | 1.87 | 3.06E-03 |
| cellular response to organic substance (GO:0071310) | 1792 | 38 | 20.5 | 1.85 | 2.51E-02 |

| | | | | | |
|---|---|---|---|---|---|
| animal organ development (GO:0048513) | 3040 | 64 | 34.77 | 1.84 | 3.38E-04 |
| regulation of response to stimulus (GO:0048583) | 3909 | 82 | 44.71 | 1.83 | 1.69E-05 |
| positive regulation of cellular biosynthetic process (GO:0031328) | 1882 | 39 | 21.52 | 1.81 | 3.20E-02 |
| positive regulation of macromolecule biosynthetic process (GO:0010557) | 1788 | 37 | 20.45 | 1.81 | 3.67E-02 |
| nervous system development (GO:0007399) | 2226 | 46 | 25.46 | 1.81 | 1.07E-02 |
| regulation of transport (GO:0051049) | 1940 | 40 | 22.19 | 1.8 | 2.63E-02 |
| regulation of molecular function (GO:0065009) | 2476 | 51 | 28.32 | 1.8 | 5.03E-03 |
| positive regulation of gene expression (GO:0010628) | 1954 | 40 | 22.35 | 1.79 | 2.82E-02 |
| cellular developmental process (GO:0048869) | 3769 | 77 | 43.11 | 1.79 | 8.89E-05 |
| cell differentiation (GO:0030154) | 3678 | 75 | 42.07 | 1.78 | 1.44E-04 |
| regulation of transcription by RNA polymerase II (GO:0006357) | 1966 | 40 | 22.49 | 1.78 | 3.90E-02 |
| positive regulation of biosynthetic process (GO:0009891) | 1923 | 39 | 21.99 | 1.77 | 3.72E-02 |
| regulation of primary metabolic process (GO:0080090) | 5143 | 104 | 58.82 | 1.77 | 1.16E-06 |
| regulation of nitrogen compound metabolic process (GO:0051171) | 4995 | 101 | 57.13 | 1.77 | 1.86E-06 |
| regulation of macromolecule biosynthetic process (GO:0010556) | 3266 | 66 | 37.35 | 1.77 | 6.93E-04 |
| positive regulation of cellular process (GO:0048522) | 5297 | 106 | 60.58 | 1.75 | 1.28E-06 |
| regulation of RNA metabolic process (GO:0051252) | 3011 | 60 | 34.44 | 1.74 | 3.11E-03 |
| regulation of biosynthetic process (GO:0009889) | 3480 | 69 | 39.8 | 1.73 | 7.75E-04 |
| regulation of nucleobase-containing compound metabolic process (GO:0019219) | 3242 | 64 | 37.08 | 1.73 | 2.00E-03 |
| regulation of developmental process (GO:0050793) | 2635 | 52 | 30.14 | 1.73 | 1.06E-02 |
| regulation of macromolecule metabolic process (GO:0060255) | 5277 | 104 | 60.35 | 1.72 | 3.49E-06 |
| regulation of cellular component organization (GO:0051128) | 2464 | 48 | 28.18 | 1.7 | 2.41E-02 |

| | | | | | |
|---|---|---|---|---|---|
| cellular response to chemical stimulus (GO:0070887) | 2314 | 45 | 26.47 | 1.7 | 3.41E-02 |
| localization (GO:0051179) | 4795 | 93 | 54.84 | 1.7 | 3.67E-05 |
| regulation of cellular biosynthetic process (GO:0031326) | 3407 | 66 | 38.97 | 1.69 | 2.70E-03 |
| regulation of multicellular organismal process (GO:0051239) | 3161 | 61 | 36.15 | 1.69 | 5.90E-03 |
| response to chemical (GO:0042221) | 3433 | 66 | 39.26 | 1.68 | 2.88E-03 |
| regulation of cellular macromolecule biosynthetic process (GO:2000112) | 3180 | 61 | 36.37 | 1.68 | 6.18E-03 |
| regulation of nucleic acid-templated transcription (GO:1903506) | 2763 | 53 | 31.6 | 1.68 | 1.76E-02 |
| regulation of metabolic process (GO:0019222) | 5740 | 110 | 65.65 | 1.68 | 3.69E-06 |
| regulation of RNA biosynthetic process (GO:2001141) | 2766 | 53 | 31.63 | 1.68 | 1.77E-02 |
| regulation of cellular metabolic process (GO:0031323) | 5335 | 102 | 61.02 | 1.67 | 1.43E-05 |
| system development (GO:0048731) | 4205 | 80 | 48.09 | 1.66 | 4.24E-04 |
| regulation of gene expression (GO:0010468) | 3652 | 69 | 41.77 | 1.65 | 3.09E-03 |
| regulation of transcription, DNA-templated (GO:0006355) | 2755 | 52 | 31.51 | 1.65 | 3.00E-02 |
| positive regulation of biological process (GO:0048518) | 5994 | 113 | 68.55 | 1.65 | 5.05E-06 |
| multicellular organism development (GO:0007275) | 4810 | 90 | 55.01 | 1.64 | 1.75E-04 |
| anatomical structure development (GO:0048856) | 5183 | 94 | 59.28 | 1.59 | 2.88E-04 |
| developmental process (GO:0032502) | 5470 | 99 | 62.56 | 1.58 | 1.77E-04 |
| regulation of biological quality (GO:0065008) | 3885 | 70 | 44.43 | 1.58 | 1.04E-02 |
| cellular component organization (GO:0016043) | 5145 | 90 | 58.84 | 1.53 | 2.35E-03 |
| cellular macromolecule metabolic process (GO:0044260) | 3973 | 68 | 45.44 | 1.5 | 3.91E-02 |
| cellular component organization or biogenesis (GO:0071840) | 5335 | 90 | 61.02 | 1.48 | 7.95E-03 |
| multicellular organismal process (GO:0032501) | 7231 | 116 | 82.7 | 1.4 | 2.71E-03 |
| regulation of cellular process (GO:0050794) | 10522 | 161 | 120.34 | 1.34 | 1.38E-04 |
| regulation of biological process (GO:0050789) | 11232 | 170 | 128.46 | 1.32 | 8.17E-05 |
| biological regulation (GO:0065007) | 11843 | 177 | 135.45 | 1.31 | 7.88E-05 |

| | | | | | |
|---|---|---|---|---|---|
| cellular process (GO:0009987) | 13986 | 195 | 159.96 | 1.22 | 9.26E-04 |
| nervous system process (GO:0050877) | 2065 | 8 | 23.62 | 0.34 | 2.40E-02 |
| G protein-coupled receptor signaling pathway (GO:0007186) | 1830 | 6 | 20.93 | 0.29 | 1.77E-02 |
| sensory perception of smell (GO:0007608) | 1122 | 0 | 12.83 | < 0.01 | 7.69E-04 |
| sensory perception of chemical stimulus (GO:0007606) | 1220 | 0 | 13.95 | < 0.01 | 2.77E-04 |

Supplementary Table S7. Transcription factors predicted by oPOSSUM to be involved in the nine modules examined.

| Gene symbol | Ensembl | Mir96 seed regions | Mir183 seed regions | Mir182 seed regions | Deafness gene | Module |
|---|---|---|---|---|---|---|
| Nkx2-5 | ENSMUSG00000015579 | | | | | black, green |
| Foxi1 | ENSMUSG00000047861 | | | | Yes | black, blue, light green |
| Foxd3 | ENSMUSG00000067261 | | | | | black, blue, light green |
| Hoxa5 | ENSMUSG00000038253 | 2 | | 1 | | green |
| Sox5 | ENSMUSG00000041540 | 3 | | 2 | | black |
| Prrx2 | ENSMUSG00000039476 | | | | | black, green, light green |
| Sry | ENSMUSG00000069036 | | | | | black, green |
| Pdx1 | ENSMUSG00000029644 | | | 1 | | black, light green |
| Nobox | ENSMUSG00000029736 | | | | | black |
| Foxq1 | ENSMUSG00000038415 | 1 | | 1 | | black, blue |
| Nkx3-1 | ENSMUSG00000022061 | 1 | 1 | 1 | | green |
| Foxa2 | ENSMUSG00000037025 | | | | | blue |
| Nfatc2 | ENSMUSG00000027544 | | | | | light green |
| Cebpa | ENSMUSG00000034957 | 1 | | 1 | | light green |
| Plag1 | ENSMUSG00000003282 | | | 1 | | royal blue |
| Rela | ENSMUSG00000024927 | | | | | royal blue |
| Pparg;Rxra | ENSMUSG00000000440; ENSMUSG00000015846 | 1 (Rxra) | | | | dark green |

| | | | | | | |
|---|---|---|---|---|---|---|
| Pax5 | ENSMUSG00000014030 | 3 | 1 | 2 | | dark green, salmon |
| Tlx1;Nfic | ENSMUSG00000025215; ENSMUSG00000055053 | 1 (Nfic) | 1 (Nfic) | 1 (Nfic) | | dark green, royal blue |
| Arnt | ENSMUSG00000015522 | | | | | royal blue |
| MA0061.1 | Profile no longer in JASPAR database | | | | | royal blue |
| Ar | ENSMUSG00000046532 | 1 | 1 | 1 | | royal blue |
| Rxra;Vdr | ENSMUSG00000015846; ENSMUSG00000022479 | 1 (Rxra) | | | | royal blue |
| Nr2e3 | ENSMUSG00000032292 | 1 | | | | purple, dark green |
| Elk4 | ENSMUSG00000026436 | | 1 | | | purple |
| Elf5 | ENSMUSG00000027186 | | | | | purple |
| Hinfp | ENSMUSG00000032119 | 1 | 1 | 2 | | grey 60 |
| Esr1 | ENSMUSG00000019768 | 1 | 1 | 1 | | grey 60 |
| Myc;Max | ENSMUSG00000022346; ENSMUSG00000059436 | | | | Yes (Myc) | grey 60 |
