## Supplementary Data for "Hearing impairment due to *Mir183/96/182* mutations suggests both loss and gain of function effects"

**ABR thresholds and p values for Fig. 1**

*Mir183/96*$^{dko}$ ABR thresholds (mean and standard deviation)

| Stimulus/freq | Mean threshold (wildtype P14) | | Mean threshold (wildtype P21) | | Mean threshold (wildtype P28) | | Mean threshold (wildtype P56) | | Mean threshold (wildtype P90) | | Mean threshold (wildtype P180) | |
|---|---|---|---|---|---|---|---|---|---|---|---|---|
| | Mean | SD | Mean | SD | Mean | SD | Mean | SD | Mean | SD | Mean | SD |
| CLICK | 38.5 | 6.687467549 | 26.5 | 2.4152295 | 26.818182 | 2.5226249 | 23.333333 | 5 | 26 | 3.1622777 | 37.5 | 12.076147 |
| 3kHz | 90 | 9.428090416 | 70 | 7.0710678 | 62 | 14.40486 | 59.444444 | 8.4574096 | 59.5 | 9.846037 | 70.5 | 9.846037 |
| 6kHz | 71.5 | 16.33843458 | 36.5 | 7.0906825 | 33.181818 | 4.0451992 | 32.222222 | 6.6666667 | 30 | 4.0824829 | 41.5 | 10.287533 |
| 12kHz | 41 | 7.745966692 | 16.5 | 4.116363 | 16.363636 | 5.5185637 | 12.777778 | 5.6519417 | 13.5 | 6.6874675 | 33.5 | 22.242352 |
| 18kHz | 40 | 8.498365856 | 24 | 5.6764621 | 23.181818 | 4.6220814 | 22.222222 | 4.4095855 | 28.5 | 9.1439111 | 66 | 25.798363 |
| 24kHz | 40.5 | 13.21825337 | 25.5 | 3.6893239 | 32.272727 | 7.1984847 | 34.444444 | 12.360331 | 59.5 | 22.906816 | 84 | 22.705848 |
| 30kHz | 50 | 18.25741858 | 36 | 7.3786479 | 63.636364 | 21.80492 | 67.777778 | 22.515427 | 85.5 | 18.020051 | 95 | 0 |
| 36kHz | 60 | 17.32050808 | 45 | 11.303883 | 95 | 0 | 90.555556 | 9.1666667 | 94.5 | 1.5811388 | 95 | 0 |
| 42kHz | 76.5 | 17.48809119 | 58.5 | 21.21975 | 94 | 2.236068 | 94.444444 | 1.6666667 | 95 | 0 | 95 | 0 |

| | Mean threshold (heterozygote P14) | | Mean threshold (heterozygote P21) | | Mean threshold (heterozygote P28) | | Mean threshold (heterozygote P56) | | Mean threshold (heterozygote P90) | | Mean threshold (heterozygote P180) | |
|---|---|---|---|---|---|---|---|---|---|---|---|---|
| | Mean | SD | Mean | SD | Mean | SD | Mean | SD | Mean | SD | Mean | SD |
| CLICK | 42.85714286 | 6.362090103 | 27 | 4.2163702 | 27.5 | 3.7977726 | 24.583333 | 3.9648073 | 25.714286 | 4.3221891 | 32.1875 | 13.536217 |
| 3kHz | 90.71428571 | 11.33893419 | 68 | 5.3748385 | 62.142857 | 8.0917359 | 59.583333 | 10.326122 | 59.285714 | 9.1687477 | 67.8125 | 8.9384469 |
| 6kHz | 79.28571429 | 7.867957925 | 40.5 | 7.6194196 | 34.642857 | 6.3440723 | 33.75 | 8.0127739 | 33.928571 | 5.2545114 | 39.375 | 13.149778 |
| 12kHz | 48.57142857 | 10.69044968 | 20.5 | 8.9597867 | 17.5 | 7.7831178 | 15.416667 | 7.5252102 | 15.714286 | 8.0520286 | 24.0625 | 22.376978 |
| 18kHz | 40.71428571 | 15.11857892 | 26 | 10.749677 | 23.571429 | 7.1866788 | 24.166667 | 7.9296146 | 25.714286 | 10.716117 | 45.9375 | 22.892411 |
| 24kHz | 40 | 8.660254038 | 32 | 12.064641 | 36.785714 | 14.885091 | 37.916667 | 17.895953 | 38.214286 | 20.62552 | 60.3125 | 24.931155 |
| 30kHz | 56.42857143 | 12.81739889 | 41.5 | 15.101508 | 58.214286 | 20.995159 | 59.583333 | 23.880225 | 65 | 25.495098 | 90.3125 | 9.9110628 |
| 36kHz | 72.14285714 | 17.52549164 | 57 | 16.865481 | 92.857143 | 5.6694671 | 82.083333 | 15.588214 | 88.571429 | 11.338934 | 95 | 0 |
| 42kHz | 89.28571429 | 9.759000729 | 67 | 20.843331 | 94.285714 | 1.8898224 | 93.75 | 3.1079078 | 95 | 0 | 95 | 0 |

| | Mean threshold (homozygote P14) | | Mean threshold (homozygote P21) | | Mean threshold (homozygote P28) | | Mean threshold (homozygote P56) | | Mean threshold (homozygote P90) | | Mean threshold (homozygote P180) | |
|---|---|---|---|---|---|---|---|---|---|---|---|---|
| | Mean | SD | Mean | SD | Mean | SD | Mean | SD | Mean | SD | Mean | SD |
| CLICK | 95 | 0 | 95 | 0 | 95 | 0 | 95 | 0 | 95 | 0 | 95 | 0 |
| 3kHz | 95 | 0 | 95 | 0 | 95 | 0 | 95 | 0 | 95 | 0 | 95 | 0 |
| 6kHz | 95 | 0 | 95 | 0 | 95 | 0 | 95 | 0 | 95 | 0 | 95 | 0 |
| 12kHz | 95 | 0 | 95 | 0 | 94 | 3.1622777 | 95 | 0 | 95 | 0 | 95 | 0 |
| 18kHz | 95 | 0 | 95 | 0 | 95 | 0 | 95 | 0 | 95 | 0 | 95 | 0 |
| 24kHz | 95 | 0 | 95 | 0 | 95 | 0 | 95 | 0 | 95 | 0 | 95 | 0 |
| 30kHz | 95 | 0 | 95 | 0 | 95 | 0 | 95 | 0 | 95 | 0 | 95 | 0 |
| 36kHz | 95 | 0 | 95 | 0 | 95 | 0 | 95 | 0 | 95 | 0 | 95 | 0 |
| 42kHz | 95 | 0 | 95 | 0 | 95 | 0 | 95 | 0 | 95 | 0 | 95 | 0 |

Linear mixed model Bonferroni-corrected p-values

P14

| | Click | 3kHz | 6kHz | 12kHz | 18kHz | 24kHz | 30kHz | 36kHz | 42kHz |
|---|---|---|---|---|---|---|---|---|---|
| Wildtype vs heterozygote | 0.466 | 0.757 | 0.173 | 0.211 | 0.964 | 0.937 | 0.374 | 0.026 | 0.008 |
| Wildtype vs homozygote | <0.001 | 0.23 | <0.001 | <0.001 | <0.001 | <0.001 | <0.001 | <0.001 | <0.001 |
| Heterozygote vs homozygote | <0.001 | 0.401 | 0.001 | <0.001 | <0.001 | <0.001 | <0.001 | <0.001 | 0.219 |

P21

| | Click | 3kHz | 6kHz | 12kHz | 18kHz | 24kHz | 30kHz | 36kHz | 42kHz |
|---|---|---|---|---|---|---|---|---|---|
| Wildtype vs heterozygote | 0.929 | 0.679 | 0.457 | 0.384 | 0.707 | 0.21 | 0.296 | 0.018 | 0.061 |
| Wildtype vs homozygote | <0.001 | <0.001 | <0.001 | <0.001 | <0.001 | <0.001 | <0.001 | <0.001 | <0.001 |
| Heterozygote vs homozygote | <0.001 | <0.001 | <0.001 | <0.001 | <0.001 | <0.001 | <0.001 | <0.001 | <0.001 |

P28

| | Click | 3kHz | 6kHz | 12kHz | 18kHz | 24kHz | 30kHz | 36kHz | 42kHz |
|---|---|---|---|---|---|---|---|---|---|
| Wildtype vs heterozygote | 0.452 | 0.389 | 0.361 | 0.419 | 0.519 | 0.102 | 0.515 | 0.706 | 0.325 |
| Wildtype vs homozygote | <0.001 | <0.001 | <0.001 | <0.001 | <0.001 | <0.001 | <0.001 | 0.495 | 0.339 |
| Heterozygote vs homozygote | <0.001 | <0.001 | <0.001 | <0.001 | <0.001 | <0.001 | <0.001 | 0.719 | 0.961 |

P56

| | Click | 3kHz | 6kHz | 12kHz | 18kHz | 24kHz | 30kHz | 36kHz | 42kHz |
|---|---|---|---|---|---|---|---|---|---|
| Wildtype vs heterozygote | 0.748 | 0.947 | 0.737 | 0.499 | 0.69 | 0.442 | 0.106 | 0.028 | 0.84 |
| Wildtype vs homozygote | <0.001 | <0.001 | <0.001 | <0.001 | <0.001 | <0.001 | <0.001 | 0.207 | 0.75 |
| Heterozygote vs homozygote | <0.001 | <0.001 | <0.001 | <0.001 | <0.001 | <0.001 | <0.001 | 0.001 | 0.596 |

P90

| | Click | 3kHz | 6kHz | 12kHz | 18kHz | 24kHz | 30kHz | 36kHz | 42kHz |
|---|---|---|---|---|---|---|---|---|---|
| Wildtype vs heterozygote | 0.896 | 0.871 | 0.289 | 0.371 | 0.639 | <0.001 | <0.001 | 0.143 | 0.835 |
| Wildtype vs homozygote | <0.001 | <0.001 | <0.001 | <0.001 | <0.001 | <0.001 | 0.034 | 0.946 | 0.916 |
| Heterozygote vs homozygote | <0.001 | <0.001 | <0.001 | <0.001 | <0.001 | <0.001 | <0.001 | 0.144 | 0.756 |

P180

| | Click | 3kHz | 6kHz | 12kHz | 18kHz | 24kHz | 30kHz | 36kHz | 42kHz |
|---|---|---|---|---|---|---|---|---|---|
| Wildtype vs heterozygote | 0.223 | 0.502 | 0.625 | 0.016 | <0.001 | <0.001 | 0.14 | 0.96 | 0.96 |
| Wildtype vs homozygote | <0.001 | <0.001 | <0.001 | <0.001 | <0.001 | 0.019 | 0.916 | 0.916 | 0.916 |
| Heterozygote vs homozygote | <0.001 | <0.001 | <0.001 | <0.001 | <0.001 | <0.001 | 0.199 | 0.946 | 0.946 |

*Mir182*$^{ko}$ ABR thresholds (mean and standard deviation)

| Stimulus/freq | Mean threshold (wildtype P14) | | Mean threshold (wildtype P21) | | Mean threshold (wildtype P28) | | Mean threshold (wildtype P56) | | Mean threshold (wildtype P90) | | Mean threshold (wildtype P180) | |
|---|---|---|---|---|---|---|---|---|---|---|---|---|
| | Mean | SD | Mean | SD | Mean | SD | Mean | SD | Mean | SD | Mean | SD |
| CLICK | 42.5 | 5.244044241 | 27.5 | 2.7386128 | 27.9 | 4.9810246 | 21.666667 | 2.5819889 | 23.333333 | 2.5819889 | 26.666667 | 12.110601 |
| 3kHz | 95 | 0 | 69.16666667 | 5.845226 | 63.8 | 12.083986 | 60 | 10.954451 | 54.166667 | 10.206207 | 69.166667 | 12.006942 |
| 6kHz | 75 | 5.477225575 | 35.83333333 | 6.6458007 | 37.9 | 6.5568609 | 34.166667 | 9.1742393 | 28.333333 | 6.8313005 | 40.833333 | 13.93437 |
| 12kHz | 39.16666667 | 10.20620726 | 17.5 | 4.1833001 | 19.6 | 6.8947718 | 12.5 | 2.7386128 | 12.5 | 5.2440442 | 21.666667 | 16.931233 |
| 18kHz | 40.83333333 | 10.68488028 | 21.66666667 | 2.5819889 | 23.8 | 7.1111308 | 20.833333 | 7.3598007 | 18.333333 | 5.1639778 | 33.333333 | 21.369761 |
| 24kHz | 37.5 | 9.354143467 | 24.16666667 | 3.7638633 | 30.0 | 7.6870611 | 21.666667 | 6.0553007 | 29.166667 | 4.9159604 | 54.166667 | 26.910345 |
| 30kHz | 51.66666667 | 17.79513042 | 38.33333333 | 5.2 | 50.0 | 14.142136 | 38.333333 | 9.8319208 | 61.666667 | 21.602469 | 86.666667 | 16.02082 |
| 36kHz | 63.33333333 | 15.70562532 | 45 | 5.5 | 81.7 | 16.001894 | 75 | 17.029386 | 87.5 | 18.371173 | 90.833333 | 10.206207 |
| 42kHz | 83.33333333 | 14.37590577 | 60.83333333 | 10.68488 | 92.1 | 4.9810246 | 92.5 | 4.1833001 | 95 | 0 | 95 | 0 |

| | Mean threshold (heterozygote P14) | | Mean threshold (heterozygote P21) | | Mean threshold (heterozygote P28) | | Mean threshold (heterozygote P56) | | Mean threshold (heterozygote P90) | | Mean threshold (heterozygote P180) | |
|---|---|---|---|---|---|---|---|---|---|---|---|---|
| | Mean | SD | Mean | SD | Mean | SD | Mean | SD | Mean | SD | Mean | SD |
| CLICK | 46.5 | 14.72903406 | 25.90909091 | 2.0225996 | 28.6 | 4.9236596 | 22.5 | 5.2440442 | 25.714286 | 5.3452248 | 25 | 2.6726124 |
| 3kHz | 94.5 | 1.58113883 | 68.63636364 | 7.4467809 | 69.8 | 13.842419 | 55.833333 | 5.845226 | 62.142857 | 8.0917359 | 76.875 | 5.3033009 |
| 6kHz | 80 | 8.164965809 | 38.18181818 | 9.0201794 | 42.5 | 10.088891 | 32.5 | 8.2158384 | 30 | 2.8867513 | 38.125 | 9.2340596 |
| 12kHz | 55 | 12.24744871 | 13.63636364 | 5.5185637 | 18.9 | 9.9918798 | 10 | 3.1622777 | 13.571429 | 5.5634864 | 15 | 5.3452248 |
| 18kHz | 52 | 13.58103252 | 23.63636364 | 4.5226702 | 28.0 | 8.5438771 | 17.5 | 5.2440442 | 19.285714 | 3.4503278 | 28.125 | 14.376941 |
| 24kHz | 50.5 | 20.06240265 | 23.18181818 | 3.3709993 | 35.2 | 11.070905 | 25 | 7.7459667 | 37.142857 | 20.586635 | 68.75 | 29.36835 |
| 30kHz | 59.5 | 17.86523626 | 37.72727273 | 6.4666979 | 65.0 | 19.210612 | 49.166667 | 22.453656 | 70.714286 | 24.397502 | 80 | 27.902893 |
| 36kHz | 78.5 | 16.67499792 | 50 | 11.2 | 88.2 | 14.519728 | 76.666667 | 16.02082 | 73.571429 | 28.535692 | 81.875 | 24.339489 |
| 42kHz | 91 | 12.64911064 | 66.36363636 | 18.5 | 90.7 | 9.1671586 | 85.833333 | 10.68488 | 80 | 24.152295 | 84.375 | 19.719008 |

| | Mean threshold (homozygote P14) | | Mean threshold (homozygote P21) | | Mean threshold (homozygote P28) | | Mean threshold (homozygote P56) | | Mean threshold (homozygote P90) | | Mean threshold (homozygote P180) | |
|---|---|---|---|---|---|---|---|---|---|---|---|---|
| | Mean | SD | Mean | SD | Mean | SD | Mean | SD | Mean | SD | Mean | SD |
| CLICK | 56.92307692 | 21.46105596 | 30.71428571 | 4.5 | 28.7 | 5.4989176 | 28.888889 | 5.4645321 | 31.428571 | 4.7559487 | 40.833333 | 10.206207 |
| 3kHz | 94.23076923 | 1.87766904 | 60.71428571 | 6.7 | 65.7 | 10.668155 | 65.555556 | 9.1666667 | 74.285714 | 12.051477 | 79.166667 | 4.9159604 |
| 6kHz | 88.46153846 | 5.547001962 | 39.28571429 | 6.7 | 42.0 | 10.987005 | 36.666667 | 6.1237244 | 36.428571 | 13.758114 | 47.5 | 8.2158384 |
| 12kHz | 66.15384615 | 18.50155919 | 17.85714286 | 2.6726124 | 19.3 | 10.327956 | 16.666667 | 6.1237244 | 19.285714 | 12.72418 | 47.5 | 24.443813 |
| 18kHz | 63.84615385 | 20.22311447 | 24.28571429 | 4.4986771 | 35.0 | 18.41971 | 30 | 16.77051 | 51.428571 | 31.320159 | 95 | 0 |
| 24kHz | 62.69230769 | 28.40300634 | 28.57142857 | 4.7559487 | 61.7 | 23.042404 | 50.555556 | 22.699731 | 83.571429 | 14.920424 | 95 | 0 |
| 30kHz | 72.30769231 | 20.97770489 | 48.57142857 | 17.251639 | 84.3 | 15.453463 | 77.222222 | 18.219343 | 94.285714 | 1.8898224 | 95 | 0 |
| 36kHz | 84.23076923 | 15.92248853 | 77.14285714 | 20.987525 | 90.7 | 8.2085902 | 93.333333 | 3.5355339 | 95 | 0 | 95 | 0 |
| 42kHz | 95 | 0 | 91.42857143 | 7.4801324 | 90.7 | 8.6327174 | 91.666667 | 6.6143783 | 95 | 0 | 95 | 0 |

Linear mixed model Bonferroni-corrected p-values

P14

| | Click | 3kHz | 6kHz | 12kHz | 18kHz | 24kHz | 30kHz | 36kHz | 42kHz |
|---|---|---|---|---|---|---|---|---|---|
| Wildtype vs heterozygote | 1 | 1 | 1 | 0.122 | 0.428 | 0.21 | 0.781 | 0.056 | 0.449 |
| Wildtype vs homozygote | 0.097 | 1 | 0.48 | <0.001 | 0.003 | <0.001 | 0.003 | 0.001 | 0.089 |
| Heterozygote vs homozygote | 0.198 | 1 | 0.216 | 0.127 | 0.099 | 0.058 | 0.042 | 0.613 | 1 |

P21

| | Click | 3kHz | 6kHz | 12kHz | 18kHz | 24kHz | 30kHz | 36kHz | 42kHz |
|---|---|---|---|---|---|---|---|---|---|
| Wildtype vs heterozygote | 1 | 1 | 1 | 1 | 1 | 1 | 1 | 1 | 0.977 |
| Wildtype vs homozygote | 1 | 0.897 | 1 | 1 | 1 | 1 | 0.68 | <0.001 | <0.001 |
| Heterozygote vs homozygote | 1 | 0.773 | 1 | 1 | 1 | 1 | 0.419 | <0.001 | <0.001 |

P28

| | Click | 3kHz | 6kHz | 12kHz | 18kHz | 24kHz | 30kHz | 36kHz | 42kHz |
|---|---|---|---|---|---|---|---|---|---|
| Wildtype vs heterozygote | 1 | 1 | 1 | 1 | 1 | 0.991 | 0.009 | 0.295 | 1 |
| Wildtype vs homozygote | 1 | 0.609 | 1 | 1 | 0.074 | <0.001 | <0.001 | 0.093 | 1 |
| Heterozygote vs homozygote | 1 | 1 | 1 | 1 | 0.24 | <0.001 | <0.001 | 1 | 1 |

P56

| | Click | 3kHz | 6kHz | 12kHz | 18kHz | 24kHz | 30kHz | 36kHz | 42kHz |
|---|---|---|---|---|---|---|---|---|---|
| Wildtype vs heterozygote | 1 | 1 | 1 | 1 | 1 | 1 | 0.353 | 1 | 0.816 |
| Wildtype vs homozygote | 0.263 | 0.52 | 1 | 0.542 | 0.139 | <0.001 | <0.001 | <0.001 | 1 |
| Heterozygote vs homozygote | 0.382 | 0.166 | 0.788 | 0.199 | 0.039 | <0.001 | <0.001 | 0.001 | 0.278 |

P90

| | Click | 3kHz | 6kHz | 12kHz | 18kHz | 24kHz | 30kHz | 36kHz | 42kHz |
|---|---|---|---|---|---|---|---|---|---|
| Wildtype vs heterozygote | 1 | 0.448 | 1 | 1 | 1 | 0.463 | 0.14 | 0.224 | 0.125 |
| Wildtype vs homozygote | 1 | 0.029 | 1 | 1 | <0.001 | <0.001 | <0.001 | 1 | 1 |
| Heterozygote vs homozygote | 1 | 0.658 | 1 | 1 | <0.001 | <0.001 | 0.003 | 0.12 | 0.176 |

P180

| | Click | 3kHz | 6kHz | 12kHz | 18kHz | 24kHz | 30kHz | 36kHz | 42kHz |
|---|---|---|---|---|---|---|---|---|---|
| Wildtype vs heterozygote | 1 | 0.371 | 1 | 1 | 1 | 0.14 | 1 | 1 | 0.562 |
| Wildtype vs homozygote | 0.217 | 0.625 | 1 | 0.001 | <0.001 | <0.001 | 0.596 | 1 | 1 |
| Heterozygote vs homozygote | 0.292 | 1 | 1 | <0.001 | <0.001 | 0.001 | 0.219 | 0.374 | 0.689 |

**Noise exposure ABR thresholds (mean and standard deviation) for Fig. 2**

| Stimulus/frequency | Mean threshold; (wildtype unexposed pre noise exposure) | | Mean threshold (wildtype unexposed; noise +1 day) | | Mean threshold (wildtype unexposed; noise +3 days) | | Mean threshold (wildtype unexposed; noise +7 days) | | Mean threshold (wildtype unexposed; noise +14 days) | | Mean threshold (wildtype unexposed; noise +28 days) | |
|---|---|---|---|---|---|---|---|---|---|---|---|---|
| | Mean | SD | Mean | SD | Mean | SD | Mean | SD | Mean | SD | Mean | SD |
| CLICK | 24.16666667 | 3.763863264 | 23.33333333 | 2.581988897 | 23.33333333 | 2.581988897 | 23.33333333 | 4.082482905 | 23.33333333 | 2.581988897 | 26.66666667 | 2.581988897 |
| 3kHz | 59.16666667 | 4.915960401 | 56.66666667 | 4.082482905 | 59.16666667 | 4.915960401 | 56.66666667 | 5.163977795 | 65 | 6.32455532 | 68.33333333 | 6.055300708 |
| 6kHz | 30 | 6.32455532 | 34.16666667 | 4.915960401 | 29.16666667 | 5.845225972 | 30.83333333 | 3.763863264 | 30.83333333 | 3.763863264 | 30.83333333 | 3.763863264 |
| 12kHz | 11.66666667 | 6.831300511 | 12.5 | 2.738612788 | 12.5 | 2.738612788 | 10.83333333 | 2.041241452 | 10.83333333 | 2.041241452 | 15 | 4.472135955 |
| 18kHz | 18.33333333 | 2.581988897 | 18.33333333 | 4.082482905 | 17.5 | 2.738612788 | 19.16666667 | 6.645800679 | 18.33333333 | 5.163977795 | 20.83333333 | 2.041241452 |
| 24kHz | 26.66666667 | 5.163977795 | 24.16666667 | 5.845225972 | 25.83333333 | 4.915960401 | 22.5 | 6.892024376 | 26.66666667 | 5.163977795 | 28.33333333 | 6.055300708 |
| 30kHz | 43.33333333 | 10.8012345 | 42.5 | 9.354143467 | 43.33333333 | 10.32795559 | 40.83333333 | 14.97219645 | 43.33333333 | 16.32993162 | 50.83333333 | 22.67524348 |
| 36kHz | 53.33333333 | 21.13448998 | 57.5 | 20.18662924 | 53.33333333 | 22.28601953 | 56.66666667 | 19.14854216 | 57.5 | 19.93740204 | 62.5 | 18.64135188 |
| 42kHz | 62.5 | 20.67607313 | 65.83333333 | 18.28022611 | 66.66666667 | 21.13448998 | 67.5 | 20.43281674 | 72.5 | 26.02883017 | 78.33333333 | 19.91649233 |

| | Mean threshold; (heterozygote unexposed pre noise exposure) | | Mean threshold (heterozygote unexposed; noise +1 day) | | Mean threshold (heterozygote unexposed; noise +3 days) | | Mean threshold (heterozygote unexposed; noise +7 days) | | Mean threshold (heterozygote unexposed; noise +14 days) | | Mean threshold (heterozygote unexposed; noise +28 days) | |
|---|---|---|---|---|---|---|---|---|---|---|---|---|
| | Mean | SD | Mean | SD | Mean | SD | Mean | SD | Mean | SD | Mean | SD |
| CLICK | 26.66666667 | 4.082482905 | 30.83333333 | 7.359800722 | 29.16666667 | 2.041241452 | 28.33333333 | 4.082482905 | 27.5 | 2.738612788 | 29.16666667 | 3.763863264 |
| 3kHz | 66.66666667 | 8.755950358 | 68.33333333 | 7.527726527 | 65 | 4.472135955 | 65 | 5.477225575 | 65.83333333 | 3.763863264 | 62.5 | 2.738612788 |
| 6kHz | 33.33333333 | 6.831300511 | 38.33333333 | 6.831300511 | 31.66666667 | 6.055300708 | 36.66666667 | 4.082482905 | 33.33333333 | 4.082482905 | 35.83333333 | 3.763863264 |
| 12kHz | 21.66666667 | 8.164965809 | 19.16666667 | 3.763863264 | 18.33333333 | 2.581988897 | 18.33333333 | 2.581988897 | 18.33333333 | 5.163977795 | 17.5 | 6.892024376 |
| 18kHz | 28.33333333 | 6.831300511 | 27.5 | 7.582875444 | 25 | 3.16227766 | 28.33333333 | 5.163977795 | 26.66666667 | 8.164965809 | 25.83333333 | 5.845225972 |
| 24kHz | 33.33333333 | 8.755950358 | 36.66666667 | 7.527726527 | 29.16666667 | 3.763863264 | 33.33333333 | 5.163977795 | 30 | 6.32455532 | 37.5 | 6.892024376 |
| 30kHz | 57.5 | 18.64135188 | 66.66666667 | 12.90994449 | 54.16666667 | 17.72474729 | 55.83333333 | 17.72474729 | 59.16666667 | 23.11204592 | 75 | 19.23538406 |
| 36kHz | 88.33333333 | 14.02378931 | 95 | 0 | 87.5 | 18.37117307 | 90 | 10 | 90.83333333 | 8.010409894 | 95 | 0 |
| 42kHz | 95 | 0 | 95 | 0 | 95 | 0 | 94.16666667 | 2.041241452 | 95 | 0 | 95 | 0 |

| | Mean threshold; (wildtype noise pre noise exposure) | | Mean threshold (wildtype noise; noise +1 day) | | Mean threshold (wildtype noise; noise +3 days) | | Mean threshold (wildtype noise; noise +7 days) | | Mean threshold (wildtype noise; noise +14 days) | | Mean threshold (wildtype noise; noise +28 days) | |
|---|---|---|---|---|---|---|---|---|---|---|---|---|
| | Mean | SD | Mean | SD | Mean | SD | Mean | SD | Mean | SD | Mean | SD |
| CLICK | 22.5 | 2.738612788 | 37.5 | 4.183300133 | 31.66666667 | 4.082482905 | 28.33333333 | 4.082482905 | 24.16666667 | 2.041241452 | 25.83333333 | 2.041241452 |
| 3kHz | 54.16666667 | 9.703951085 | 59.16666667 | 13.93436998 | 55 | 5.477225575 | 51.66666667 | 6.055300708 | 57.5 | 15.08310313 | 57.5 | 8.215838363 |
| 6kHz | 28.33333333 | 2.581988897 | 30 | 3.16227766 | 30 | 0 | 28.33333333 | 4.082482905 | 30 | 6.32455532 | 30 | 3.16227766 |
| 12kHz | 10.83333333 | 3.763863264 | 36.66666667 | 10.32795559 | 18.33333333 | 6.055300708 | 14.16666667 | 3.763863264 | 11.66666667 | 2.581988897 | 12.5 | 4.183300133 |
| 18kHz | 17.5 | 4.183300133 | 72.5 | 8.215838363 | 52.5 | 18.9076704 | 34.16666667 | 19.34338819 | 17.5 | 2.738612788 | 22.5 | 2.738612788 |
| 24kHz | 25.83333333 | 3.763863264 | 83.33333333 | 4.082482905 | 65.83333333 | 17.72474729 | 48.33333333 | 21.83269719 | 25.83333333 | 2.041241452 | 29.16666667 | 3.763863264 |
| 30kHz | 34.16666667 | 5.845225972 | 95 | 0 | 83.33333333 | 12.51665557 | 64.16666667 | 19.34338819 | 44.16666667 | 5.845225972 | 55.83333333 | 11.14300977 |
| 36kHz | 49.16666667 | 8.612007122 | 95 | 0 | 92.5 | 6.123724357 | 74.16666667 | 17.15128761 | 70.83333333 | 10.68488028 | 89.16666667 | 9.174239296 |
| 42kHz | 73.33333333 | 19.66384161 | 95 | 0 | 91.66666667 | 5.163977795 | 84.16666667 | 13.57080199 | 87.5 | 12.5499004 | 95 | 0 |

| | Mean threshold; (heterozygote noise pre noise exposure) | | Mean threshold (heterozygote noise; noise +1 day) | | Mean threshold (heterozygote noise; noise +3 days) | | Mean threshold (heterozygote noise; noise +7 days) | | Mean threshold (heterozygote noise; noise +14 days) | | Mean threshold (heterozygote noise; noise +28 days) | |
|---|---|---|---|---|---|---|---|---|---|---|---|---|
| | Mean | SD | Mean | SD | Mean | SD | Mean | SD | Mean | SD | Mean | SD |
| CLICK | 23.33333333 | 2.581988897 | 40 | 3.16227766 | 34.16666667 | 7.359800722 | 28.33333333 | 2.581988897 | 25.83333333 | 5.845225972 | 28.33333333 | 2.581988897 |
| 3kHz | 60 | 7.745966692 | 63.33333333 | 6.831300511 | 66.66666667 | 5.163977795 | 64.16666667 | 7.359800722 | 63.33333333 | 4.082482905 | 55.83333333 | 3.763863264 |
| 6kHz | 30.83333333 | 4.915960401 | 33.33333333 | 6.055300708 | 38.33333333 | 5.163977795 | 38.33333333 | 9.309493363 | 35.83333333 | 5.845225972 | 33.33333333 | 6.055300708 |
| 12kHz | 12.5 | 2.738612788 | 44.16666667 | 7.359800722 | 23.33333333 | 8.755950358 | 18.33333333 | 4.082482905 | 15 | 4.472135955 | 15 | 0 |
| 18kHz | 21.66666667 | 2.581988897 | 77.5 | 5.244044241 | 55 | 15.49193338 | 27.5 | 8.215838363 | 21.66666667 | 4.082482905 | 25 | 3.16227766 |
| 24kHz | 27.5 | 2.738612788 | 88.33333333 | 4.082482905 | 62.5 | 13.32291259 | 46.66666667 | 20.16597795 | 32.5 | 6.123724357 | 38.33333333 | 18.07392228 |
| 30kHz | 44.16666667 | 16.55797894 | 94.16666667 | 2.041241452 | 80.83333333 | 14.63443428 | 61.66666667 | 29.26886856 | 50.83333333 | 18.55172948 | 65.83333333 | 27.46209509 |
| 36kHz | 72.5 | 24.84954728 | 95 | 0 | 95 | 0 | 81.66666667 | 17.22401424 | 78.33333333 | 19.91649233 | 85 | 15.8113883 |
| 42kHz | 88.33333333 | 10.8012345 | 95 | 0 | 92.5 | 4.183300133 | 90.83333333 | 10.20620726 | 93.33333333 | 4.082482905 | 95 | 0 |

| Mouse ID | Experiment date | Date of birth | Sex | *Mir183/96*$^{dko}$ geno | Age (days) | Age (weeks) | Noise status | Noise timing | Click threshold | 3kHz threshold | 6kHz threshold | 12kHz threshold | 18kHz threshold | 24kHz threshold | 30kHz threshold | 36kHz threshold | 42kHz threshold |
|---|---|---|---|---|---|---|---|---|---|---|---|---|---|---|---|---|---|
| CNTC20.1a | 13/07/2015 | 21/05/2015 | M | Wildtype | 53 | 7.6 | Control (no noise) | Pre noise exposure | 20 | 55 | 35 | 20 | 15 | 25 | 40 | 45 | 50 |
| CNTC20.1a | 15/07/2015 | 21/05/2015 | M | Wildtype | 55 | 7.9 | Control (no noise) | Noise +1 days | 20 | 55 | 35 | 15 | 15 | 20 | 35 | 40 | 55 |
| CNTC20.1a | 17/07/2015 | 21/05/2015 | M | Wildtype | 57 | 8.1 | Control (no noise) | Noise +3 days | 20 | 55 | 30 | 10 | 15 | 20 | 30 | 35 | 50 |
| CNTC20.1a | 21/07/2015 | 21/05/2015 | M | Wildtype | 61 | 8.7 | Control (no noise) | Noise + 7 days | 25 | 55 | 30 | 10 | 15 | 15 | 30 | 45 | 65 |
| CNTC20.1a | 28/07/2015 | 21/05/2015 | M | Wildtype | 68 | 9.7 | Control (no noise) | Noise + 14 days | 20 | 60 | 35 | 10 | 15 | 25 | 35 | 50 | 65 |
| CNTC20.1a | 11/08/2015 | 21/05/2015 | M | Wildtype | 82 | 11.7 | Control (no noise) | Noise + 28 days | 25 | 75 | 30 | 20 | 20 | 25 | 45 | 50 | 75 |
| CNTC20.1f | 13/07/2015 | 21/05/2015 | F | Wildtype | 53 | 7.6 | Control (no noise) | Pre noise exposure | 25 | 55 | 35 | 15 | 20 | 30 | 45 | 50 | 60 |
| CNTC20.1f | 15/07/2015 | 21/05/2015 | F | Wildtype | 55 | 7.9 | Control (no noise) | Noise +1 days | 25 | 50 | 35 | 10 | 15 | 25 | 35 | 60 | 65 |
| CNTC20.1f | 17/07/2015 | 21/05/2015 | F | Wildtype | 57 | 8.1 | Control (no noise) | Noise +3 days | 25 | 65 | 35 | 15 | 15 | 30 | 45 | 55 | 75 |
| CNTC20.1f | 21/07/2015 | 21/05/2015 | F | Wildtype | 61 | 8.7 | Control (no noise) | Noise + 7 days | 25 | 60 | 30 | 10 | 20 | 25 | 40 | 50 | 70 |
| CNTC20.1f | 28/07/2015 | 21/05/2015 | F | Wildtype | 68 | 9.7 | Control (no noise) | Noise + 14 days | 25 | 60 | 30 | 15 | 25 | 25 | 40 | 45 | 95 |
| CNTC20.1f | 11/08/2015 | 21/05/2015 | F | Wildtype | 82 | 11.7 | Control (no noise) | Noise + 28 days | 25 | 60 | 25 | 10 | 20 | 30 | 50 | 65 | 95 |
| CNTC20.1g | 13/07/2015 | 21/05/2015 | F | Wildtype | 53 | 7.6 | Control (no noise) | Pre noise exposure | 25 | 65 | 35 | 15 | 15 | 25 | 50 | 50 | 80 |
| CNTC20.1g | 15/07/2015 | 21/05/2015 | F | Wildtype | 55 | 7.9 | Control (no noise) | Noise +1 days | 25 | 55 | 40 | 10 | 15 | 20 | 45 | 55 | 80 |
| CNTC20.1g | 17/07/2015 | 21/05/2015 | F | Wildtype | 57 | 8.1 | Control (no noise) | Noise +3 days | 25 | 55 | 35 | 15 | 20 | 20 | 45 | 45 | 85 |
| CNTC20.1g | 21/07/2015 | 21/05/2015 | F | Wildtype | 61 | 8.7 | Control (no noise) | Noise + 7 days | 25 | 65 | 35 | 10 | 20 | 20 | 40 | 55 | 85 |
| CNTC20.1g | 28/07/2015 | 21/05/2015 | F | Wildtype | 68 | 9.7 | Control (no noise) | Noise + 14 days | 25 | 65 | 30 | 10 | 15 | 35 | 45 | 65 | 95 |
| CNTC20.1g | 11/08/2015 | 21/05/2015 | F | Wildtype | 82 | 11.7 | Control (no noise) | Noise + 28 days | 25 | 75 | 30 | 15 | 25 | 25 | 45 | 70 | 95 |
| DMKO44.1a | 07/09/2015 | 14/07/2015 | M | Wildtype | 55 | 7.9 | Control (no noise) | Pre noise exposure | 30 | 55 | 30 | 10 | 20 | 35 | 60 | 95 | 95 |
| DMKO44.1a | 09/09/2015 | 14/07/2015 | M | Wildtype | 57 | 8.1 | Control (no noise) | Noise +1 days | 25 | 60 | 35 | 15 | 20 | 35 | 60 | 95 | 95 |
| DMKO44.1a | 11/09/2015 | 14/07/2015 | M | Wildtype | 59 | 8.4 | Control (no noise) | Noise +3 days | 25 | 65 | 25 | 10 | 20 | 30 | 60 | 95 | 95 |
| DMKO44.1a | 15/09/2015 | 14/07/2015 | M | Wildtype | 63 | 9.0 | Control (no noise) | Noise + 7 days | 25 | 55 | 30 | 15 | 30 | 35 | 70 | 95 | 95 |
| DMKO44.1a | 22/09/2015 | 14/07/2015 | M | Wildtype | 70 | 10.0 | Control (no noise) | Noise + 14 days | 25 | 70 | 35 | 10 | 15 | 30 | 75 | 95 | 95 |
| DMKO44.1a | 06/10/2015 | 14/07/2015 | M | Wildtype | 84 | 12.0 | Control (no noise) | Noise + 28 days | 30 | 70 | 35 | 20 | 20 | 40 | 95 | 95 | 95 |
| DMKO44.1b | 07/09/2015 | 14/07/2015 | M | Heterozygote | 55 | 7.9 | Control (no noise) | Pre noise exposure | 30 | 70 | 30 | 20 | 35 | 30 | 60 | 95 | 95 |
| DMKO44.1b | 09/09/2015 | 14/07/2015 | M | Heterozygote | 57 | 8.1 | Control (no noise) | Noise +1 days | 25 | 65 | 35 | 15 | 25 | 30 | 55 | 95 | 95 |
| DMKO44.1b | 11/09/2015 | 14/07/2015 | M | Heterozygote | 59 | 8.4 | Control (no noise) | Noise +3 days | 30 | 70 | 35 | 20 | 30 | 25 | 55 | 95 | 95 |
| DMKO44.1b | 15/09/2015 | 14/07/2015 | M | Heterozygote | 63 | 9.0 | Control (no noise) | Noise + 7 days | 25 | 70 | 40 | 20 | 30 | 35 | 60 | 95 | 95 |
| DMKO44.1b | 22/09/2015 | 14/07/2015 | M | Heterozygote | 70 | 10.0 | Control (no noise) | Noise + 14 days | 30 | 70 | 30 | 15 | 30 | 25 | 50 | 95 | 95 |
| DMKO44.1b | 06/10/2015 | 14/07/2015 | M | Heterozygote | 84 | 12.0 | Control (no noise) | Noise + 28 days | 25 | 60 | 35 | 25 | 30 | 40 | 80 | 95 | 95 |
| DMKO44.1c | 07/09/2015 | 14/07/2015 | M | Wildtype | 55 | 7.9 | Noise | Pre noise exposure | 25 | 60 | 30 | 15 | 25 | 30 | 35 | 50 | 80 |
| DMKO44.1c | 09/09/2015 | 14/07/2015 | M | Wildtype | 57 | 8.1 | Noise | Noise +1 days | 35 | 60 | 30 | 35 | 75 | 85 | 95 | 95 | 95 |
| DMKO44.1c | 11/09/2015 | 14/07/2015 | M | Wildtype | 59 | 8.4 | Noise | Noise +3 days | 35 | 60 | 30 | 20 | 55 | 65 | 80 | 95 | 95 |
| DMKO44.1c | 15/09/2015 | 14/07/2015 | M | Wildtype | 63 | 9.0 | Noise | Noise + 7 days | 25 | 55 | 30 | 15 | 30 | 35 | 40 | 55 | 85 |
| DMKO44.1c | 22/09/2015 | 14/07/2015 | M | Wildtype | 70 | 10.0 | Noise | Noise + 14 days | 25 | 75 | 30 | 10 | 15 | 25 | 40 | 65 | 95 |
| DMKO44.1c | 06/10/2015 | 14/07/2015 | M | Wildtype | 84 | 12.0 | Noise | Noise + 28 days | 30 | 50 | 30 | 15 | 20 | 30 | 50 | 95 | 95 |
| DMKO44.1d | 07/09/2015 | 14/07/2015 | M | Heterozygote | 55 | 7.9 | Control (no noise) | Pre noise exposure | 30 | 70 | 45 | 25 | 30 | 35 | 70 | 95 | 95 |
| DMKO44.1d | 09/09/2015 | 14/07/2015 | M | Heterozygote | 57 | 8.1 | Control (no noise) | Noise +1 days | 30 | 75 | 35 | 20 | 20 | 30 | 75 | 95 | 95 |
| DMKO44.1d | 11/09/2015 | 14/07/2015 | M | Heterozygote | 59 | 8.4 | Control (no noise) | Noise +3 days | 30 | 65 | 25 | 15 | 25 | 30 | 80 | 95 | 95 |
| DMKO44.1d | 15/09/2015 | 14/07/2015 | M | Heterozygote | 63 | 9.0 | Control (no noise) | Noise + 7 days | 25 | 70 | 35 | 15 | 30 | 35 | 60 | 95 | 95 |
| DMKO44.1d | 22/09/2015 | 14/07/2015 | M | Heterozygote | 70 | 10.0 | Control (no noise) | Noise + 14 days | 30 | 70 | 35 | 25 | 35 | 40 | 95 | 95 | 95 |
| DMKO44.1d | 06/10/2015 | 14/07/2015 | M | Heterozygote | 84 | 12.0 | Control (no noise) | Noise + 28 days | 35 | 65 | 35 | 25 | 35 | 50 | 95 | 95 | 95 |
| DMKO44.1e | 07/09/2015 | 14/07/2015 | M | Heterozygote | 55 | 7.9 | Noise | Pre noise exposure | 25 | 55 | 35 | 15 | 25 | 30 | 40 | 55 | 95 |
| DMKO44.1e | 09/09/2015 | 14/07/2015 | M | Heterozygote | 57 | 8.1 | Noise | Noise +1 days | 40 | 70 | 45 | 50 | 80 | 85 | 95 | 95 | 95 |
| DMKO44.1e | 11/09/2015 | 14/07/2015 | M | Heterozygote | 59 | 8.4 | Noise | Noise +3 days | 45 | 70 | 45 | 30 | 50 | 55 | 75 | 95 | 95 |
| DMKO44.1e | 15/09/2015 | 14/07/2015 | M | Heterozygote | 63 | 9.0 | Noise | Noise + 7 days | 30 | 60 | 40 | 15 | 25 | 30 | 50 | 85 | 95 |
| DMKO44.1e | 22/09/2015 | 14/07/2015 | M | Heterozygote | 70 | 10.0 | Noise | Noise + 14 days | 25 | 65 | 30 | 20 | 20 | 30 | 45 | 60 | 95 |
| DMKO44.1e | 06/10/2015 | 14/07/2015 | M | Heterozygote | 84 | 12.0 | Noise | Noise + 28 days | 25 | 55 | 30 | 15 | 20 | 30 | 50 | 90 | 95 |
| DMKO44.1f | 07/09/2015 | 14/07/2015 | F | Heterozygote | 55 | 7.9 | Noise | Pre noise exposure | 25 | 65 | 35 | 15 | 20 | 30 | 75 | 95 | 95 |
| DMKO44.1f | 09/09/2015 | 14/07/2015 | F | Heterozygote | 57 | 8.1 | Noise | Noise +1 days | 45 | 65 | 35 | 55 | 80 | 90 | 95 | 95 | 95 |
| DMKO44.1f | 11/09/2015 | 14/07/2015 | F | Heterozygote | 59 | 8.4 | Noise | Noise +3 days | 35 | 70 | 35 | 25 | 50 | 65 | 95 | 95 | 95 |
| DMKO44.1f | 15/09/2015 | 14/07/2015 | F | Heterozygote | 63 | 9.0 | Noise | Noise + 7 days | 30 | 70 | 45 | 20 | 35 | 65 | 95 | 95 | 95 |
| DMKO44.1f | 22/09/2015 | 14/07/2015 | F | Heterozygote | 70 | 10.0 | Noise | Noise + 14 days | 35 | 60 | 40 | 20 | 30 | 40 | 85 | 95 | 95 |
| DMKO44.1f | 06/10/2015 | 14/07/2015 | F | Heterozygote | 84 | 12.0 | Noise | Noise + 28 days | 30 | 55 | 40 | 15 | 30 | 75 | 95 | 95 | 95 |
| DMKO44.1g | 07/09/2015 | 14/07/2015 | F | Wildtype | 55 | 7.9 | Noise | Pre noise exposure | 25 | 60 | 30 | 10 | 15 | 25 | 40 | 50 | 95 |
| DMKO44.1g | 09/09/2015 | 14/07/2015 | F | Wildtype | 57 | 8.1 | Noise | Noise +1 days | 30 | 45 | 30 | 25 | 60 | 80 | 95 | 95 | 95 |
| DMKO44.1g | 11/09/2015 | 14/07/2015 | F | Wildtype | 59 | 8.4 | Noise | Noise +3 days | 30 | 55 | 30 | 15 | 35 | 50 | 85 | 95 | 95 |
| DMKO44.1g | 15/09/2015 | 14/07/2015 | F | Wildtype | 63 | 9.0 | Noise | Noise + 7 days | 30 | 55 | 30 | 15 | 15 | 35 | 60 | 70 | 95 |
| DMKO44.1g | 22/09/2015 | 14/07/2015 | F | Wildtype | 70 | 10.0 | Noise | Noise + 14 days | 25 | 40 | 25 | 10 | 20 | 25 | 50 | 75 | 95 |
| DMKO44.1g | 06/10/2015 | 14/07/2015 | F | Wildtype | 84 | 12.0 | Noise | Noise + 28 days | 25 | 55 | 25 | 10 | 20 | 35 | 60 | 95 | 95 |
| DMKO54.1c | 08/08/2016 | 14/06/2016 | F | Heterozygote | 55 | 7.9 | Noise | Pre noise exposure | 20 | 65 | 30 | 10 | 20 | 25 | 35 | 50 | 80 |
| DMKO54.1c | 10/08/2016 | 14/06/2016 | F | Heterozygote | 57 | 8.1 | Noise | Noise +1 days | 40 | 70 | 30 | 35 | 70 | 90 | 95 | 95 | 95 |
| DMKO54.1c | 12/08/2016 | 14/06/2016 | F | Heterozygote | 59 | 8.4 | Noise | Noise +3 days | 40 | 70 | 40 | 30 | 65 | 60 | 70 | 95 | 95 |
| DMKO54.1c | 16/08/2016 | 14/06/2016 | F | Heterozygote | 63 | 9.0 | Noise | Noise + 7 days | 30 | 65 | 40 | 20 | 20 | 50 | 25 | 60 | 95 |
| DMKO54.1c | 23/08/2016 | 14/06/2016 | F | Heterozygote | 70 | 10.0 | Noise | Noise + 14 days | 30 | 60 | 35 | 15 | 20 | 40 | 45 | 75 | 85 |
| DMKO54.1c | 06/09/2016 | 14/06/2016 | F | Heterozygote | 84 | 12.0 | Noise | Noise + 28 days | 30 | 60 | 40 | 15 | 25 | 30 | 45 | 80 | 95 |
| DMKO54.1d | 08/08/2016 | 14/06/2016 | F | Wildtype | 55 | 7.9 | Noise | Pre noise exposure | 25 | 40 | 25 | 10 | 15 | 25 | 35 | 55 | 65 |
| DMKO54.1d | 10/08/2016 | 14/06/2016 | F | Wildtype | 57 | 8.1 | Noise | Noise +1 days | 40 | 55 | 25 | 25 | 75 | 80 | 95 | 95 | 95 |
| DMKO54.1d | 12/08/2016 | 14/06/2016 | F | Wildtype | 59 | 8.4 | Noise | Noise +3 days | 35 | 55 | 30 | 25 | 60 | 80 | 90 | 95 | 95 |
| DMKO54.1d | 16/08/2016 | 14/06/2016 | F | Wildtype | 63 | 9.0 | Noise | Noise + 7 days | 30 | 50 | 20 | 10 | 35 | 70 | 85 | 95 | 95 |
| DMKO54.1d | 23/08/2016 | 14/06/2016 | F | Wildtype | 70 | 10.0 | Noise | Noise + 14 days | 20 | 40 | 25 | 10 | 20 | 25 | 50 | 80 | 95 |
| DMKO54.1d | 06/09/2016 | 14/06/2016 | F | Wildtype | 84 | 12.0 | Noise | Noise + 28 days | 25 | 50 | 30 | 15 | 25 | 25 | 65 | 95 | 95 |
| DMKO54.1e | 08/08/2016 | 14/06/2016 | F | Heterozygote | 55 | 7.9 | Control (no noise) | Pre noise exposure | 20 | 55 | 35 | 20 | 30 | 30 | 40 | 90 | 95 |
| DMKO54.1e | 10/08/2016 | 14/06/2016 | F | Heterozygote | 57 | 8.1 | Control (no noise) | Noise +1 days | 30 | 60 | 40 | 20 | 30 | 35 | 55 | 95 | 95 |
| DMKO54.1e | 12/08/2016 | 14/06/2016 | F | Heterozygote | 59 | 8.4 | Control (no noise) | Noise +3 days | 25 | 65 | 40 | 20 | 25 | 35 | 50 | 95 | 95 |
| DMKO54.1e | 16/08/2016 | 14/06/2016 | F | Heterozygote | 63 | 9.0 | Control (no noise) | Noise + 7 days | 25 | 65 | 40 | 20 | 30 | 30 | 40 | 90 | 95 |
| DMKO54.1e | 23/08/2016 | 14/06/2016 | F | Heterozygote | 70 | 10.0 | Control (no noise) | Noise + 14 days | 30 | 60 | 40 | 20 | 35 | 30 | 45 | 95 | 95 |
| DMKO54.1e | 06/09/2016 | 14/06/2016 | F | Heterozygote | 84 | 12.0 | Control (no noise) | Noise + 28 days | 30 | 60 | 40 | 20 | 25 | 35 | 55 | 95 | 95 |
| DMKO54.1f | 08/08/2016 | 14/06/2016 | F | Wildtype | 55 | 7.9 | Control (no noise) | Pre noise exposure | 25 | 65 | 20 | 10 | 20 | 25 | 35 | 45 | 45 |
| DMKO54.1f | 10/08/2016 | 14/06/2016 | F | Wildtype | 57 | 8.1 | Control (no noise) | Noise +1 days | 25 | 60 | 35 | 15 | 25 | 25 | 40 | 55 | 50 |
| DMKO54.1f | 12/08/2016 | 14/06/2016 | F | Wildtype | 59 | 8.4 | Control (no noise) | Noise +3 days | 25 | 60 | 30 | 15 | 20 | 30 | 45 | 55 | 50 |
| DMKO54.1f | 16/08/2016 | 14/06/2016 | F | Wildtype | 63 | 9.0 | Control (no noise) | Noise + 7 days | 25 | 50 | 25 | 10 | 20 | 20 | 35 | 45 | 45 |
| DMKO54.1f | 23/08/2016 | 14/06/2016 | F | Wildtype | 70 | 10.0 | Control (no noise) | Noise + 14 days | 25 | 75 | 30 | 10 | 25 | 20 | 35 | 45 | 45 |
| DMKO54.1f | 06/09/2016 | 14/06/2016 | F | Wildtype | 84 | 12.0 | Control (no noise) | Noise + 28 days | 30 | 65 | 35 | 15 | 20 | 25 | 40 | 45 | 60 |
| DMKO55.1a | 08/08/2016 | 14/06/2016 | M | Wildtype | 55 | 7.9 | Control (no noise) | Pre noise exposure | 20 | 60 | 25 | 0 | 20 | 20 | 30 | 35 | 45 |
| DMKO55.1a | 10/08/2016 | 14/06/2016 | M | Wildtype | 57 | 8.1 | Control (no noise) | Noise +1 days | 20 | 60 | 25 | 10 | 20 | 20 | 40 | 40 | 50 |
| DMKO55.1a | 12/08/2016 | 14/06/2016 | M | Wildtype | 59 | 8.4 | Control (no noise) | Noise +3 days | 20 | 55 | 20 | 10 | 15 | 25 | 35 | 35 | 45 |
| DMKO55.1a | 16/08/2016 | 14/06/2016 | M | Wildtype | 63 | 9.0 | Control (no noise) | Noise + 7 days | 15 | 55 | 35 | 10 | 10 | 20 | 30 | 50 | 45 |
| DMKO55.1a | 23/08/2016 | 14/06/2016 | M | Wildtype | 70 | 10.0 | Control (no noise) | Noise + 14 days | 20 | 60 | 25 | 10 | 15 | 25 | 30 | 45 | 40 |
| DMKO55.1a | 06/09/2016 | 14/06/2016 | M | Wildtype | 84 | 12.0 | Control (no noise) | Noise + 28 days | 25 | 65 | 30 | 10 | 20 | 25 | 30 | 50 | 50 |
| DMKO55.1b | 08/08/2016 | 14/06/2016 | M | Wildtype | 55 | 7.9 | Noise | Pre noise exposure | 20 | 45 | 30 | 15 | 20 | 20 | 25 | 35 | 50 |
| DMKO55.1b | 10/08/2016 | 14/06/2016 | M | Wildtype | 57 | 8.1 | Noise | Noise +1 days | 40 | 85 | 35 | 50 | 80 | 85 | 95 | 95 | 95 |
| DMKO55.1b | 12/08/2016 | 14/06/2016 | M | Wildtype | 59 | 8.4 | Noise | Noise +3 days | 25 | 55 | 30 | 15 | 25 | 40 | 60 | 80 | 85 |
| DMKO55.1b | 16/08/2016 | 14/06/2016 | M | Wildtype | 63 | 9.0 | Noise | Noise + 7 days | 25 | 45 | 30 | 10 | 20 | 25 | 55 | 70 | 65 |
| DMKO55.1b | 23/08/2016 | 14/06/2016 | M | Wildtype | 70 | 10.0 | Noise | Noise + 14 days | 25 | 65 | 35 | 15 | 15 | 30 | 35 | 60 | 80 |
| DMKO55.1b | 06/09/2016 | 14/06/2016 | M | Wildtype | 84 | 12.0 | Noise | Noise + 28 days | 25 | 55 | 30 | 15 | 20 | 25 | 40 | 75 | 95 |
| DMKO55.1c | 08/08/2016 | 14/06/2016 | M | Heterozygote | 55 | 7.9 | Noise | Pre noise exposure | 25 | 50 | 25 | 10 | 20 | 25 | 30 | 45 | 70 |
| DMKO55.1c | 10/08/2016 | 14/06/2016 | M | Heterozygote | 57 | 8.1 | Noise | Noise +1 days | 40 | 55 | 30 | 40 | 75 | 85 | 90 | 95 | 95 |
| DMKO55.1c | 12/08/2016 | 14/06/2016 | M | Heterozygote | 59 | 8.4 | Noise | Noise +3 days | 25 | 60 | 30 | 10 | 30 | 45 | 60 | 95 | 90 |
| DMKO55.1c | 16/08/2016 | 14/06/2016 | M | Heterozygote | 63 | 9.0 | Noise | Noise + 7 days | 25 | 55 | 25 | 15 | 20 | 25 | 35 | 60 | 70 |
| DMKO55.1c | 23/08/2016 | 14/06/2016 | M | Heterozygote | 70 | 10.0 | Noise | Noise + 14 days | 20 | 70 | 30 | 10 | 20 | 30 | 30 | 50 | 95 |
| DMKO55.1c | 06/09/2016 | 14/06/2016 | M | Heterozygote | 84 | 12.0 | Noise | Noise + 28 days | 25 | 55 | 25 | 15 | 25 | 30 | 30 | 55 | 95 |
| DMKO55.1g | 08/08/2016 | 14/06/2016 | F | Heterozygote | 55 | 7.9 | Noise | Pre noise exposure | 20 | 70 | 35 | 15 | 25 | 25 | 35 | 95 | 95 |
| DMKO55.1g | 10/08/2016 | 14/06/2016 | F | Heterozygote | 57 | 8.1 | Noise | Noise +1 days | 35 | 65 | 30 | 40 | 75 | 85 | 95 | 95 | 95 |
| DMKO55.1g | 12/08/2016 | 14/06/2016 | F | Heterozygote | 59 | 8.4 | Noise | Noise +3 days | 30 | 60 | 40 | 30 | 75 | 85 | 95 | 95 | 85 |
| DMKO55.1g | 16/08/2016 | 14/06/2016 | F | Heterozygote | 63 | 9.0 | Noise | Noise + 7 days | 30 | 60 | 30 | 15 | 40 | 75 | 90 | 95 | 95 |
| DMKO55.1g | 23/08/2016 | 14/06/2016 | F | Heterozygote | 70 | 10.0 | Noise | Noise + 14 days | 20 | 60 | 35 | 10 | 20 | 30 | 45 | 95 | 95 |
| DMKO55.1g | 06/09/2016 | 14/06/2016 | F | Heterozygote | 84 | 12.0 | Noise | Noise + 28 days | 30 | 60 | 30 | 15 | 25 | 30 | 85 | 95 | 95 |
| DMKO55.1h | 08/08/2016 | 14/06/2016 | F | Wildtype | 55 | 7.9 | Noise | Pre noise exposure | 20 | 65 | 25 | 10 | 15 | 30 | 30 | 45 | 55 |
| DMKO55.1h | 10/08/2016 | 14/06/2016 | F | Wildtype | 57 | 8.1 | Noise | Noise +1 days | 40 | 50 | 30 | 45 | 65 | 80 | 95 | 95 | 95 |
| DMKO55.1h | 12/08/2016 | 14/06/2016 | F | Wildtype | 59 | 8.4 | Noise | Noise +3 days | 30 | 45 | 30 | 10 | 65 | 75 | 90 | 95 | 85 |
| DMKO55.1h | 16/08/2016 | 14/06/2016 | F | Wildtype | 63 | 9.0 | Noise | Noise + 7 days | 25 | 60 | 30 | 15 | 35 | 45 | 55 | 60 | 70 |
| DMKO55.1h | 23/08/2016 | 14/06/2016 | F | Wildtype | 70 | 10.0 | Noise | Noise + 14 days | 25 | 70 | 25 | 10 | 15 | 25 | 45 | 60 | 65 |
| DMKO55.1h | 06/09/2016 | 14/06/2016 | F | Wildtype | 84 | 12.0 | Noise | Noise + 28 days | 25 | 70 | 30 | 15 | 25 | 30 | 50 | 80 | 95 |
| DMKO55.1i | 08/08/2016 | 14/06/2016 | F | Heterozygote | 55 | 7.9 | Control (no noise) | Pre noise exposure | 25 | 60 | 25 | 10 | 20 | 25 | 35 | 60 | 95 |
| DMKO55.1i | 10/08/2016 | 14/06/2016 | F | Heterozygote | 57 | 8.1 | Control (no noise) | Noise +1 days | 45 | 75 | 50 | 25 | 40 | 50 | 80 | 95 | 95 |
| DMKO55.1i | 12/08/2016 | 14/06/2016 | F | Heterozygote | 59 | 8.4 | Control (no noise) | Noise +3 days | 30 | 70 | 25 | 20 | 25 | 25 | 25 | 50 | 95 |
| DMKO55.1i | 16/08/2016 | 14/06/2016 | F | Heterozygote | 63 | 9.0 | Control (no noise) | Noise + 7 days | 30 | 55 | 30 | 15 | 20 | 25 | 35 | 70 | 90 |
| DMKO55.1i | 23/08/2016 | 14/06/2016 | F | Heterozygote | 70 | 10.0 | Control (no noise) | Noise + 14 days | 25 | 65 | 30 | 20 | 25 | 25 | 35 | 75 | 95 |
| DMKO55.1i | 06/09/2016 | 14/06/2016 | F | Heterozygote | 84 | 12.0 | Control (no noise) | Noise + 28 days | 30 | 65 | 40 | 15 | 25 | 30 | 50 | 95 | 95 |
| DMKO57.1a | 03/10/2016 | 09/08/2016 | M | Heterozygote | 55 | 7.9 | Noise | Pre noise exposure | 25 | 55 | 25 | 10 | 20 | 30 | 50 | 95 | 95 |
| DMKO57.1a | 05/10/2016 | 09/08/2016 | M | Heterozygote | 57 | 8.1 | Noise | Noise +1 days | 40 | 55 | 30 | 45 | 85 | 95 | 95 | 95 | 95 |
| DMKO57.1a | 07/10/2016 | 09/08/2016 | M | Heterozygote | 59 | 8.4 | Noise | Noise +3 days | 30 | 70 | 40 | 15 | 60 | 65 | 90 | 95 | 95 |
| DMKO57.1a | 11/10/2016 | 09/08/2016 | M | Heterozygote | 63 | 9.0 | Noise | Noise + 7 days | 25 | 75 | 50 | 25 | 25 | 35 | 75 | 95 | 95 |
| DMKO57.1a | 18/10/2016 | 09/08/2016 | M | Heterozygote | 70 | 10.0 | Noise | Noise + 14 days | 25 | 65 | 45 | 15 | 20 | 25 | 55 | 95 | 95 |
| DMKO57.1a | 01/11/2016 | 09/08/2016 | M | Heterozygote | 84 | 12.0 | Noise | Noise + 28 days | 30 | 50 | 35 | 15 | 25 | 35 | 90 | 95 | 95 |
| DMKO57.1b | 03/10/2016 | 09/08/2016 | M | Wildtype | 55 | 7.9 | Noise | Pre noise exposure | 20 | 55 | 30 | 5 | 15 | 25 | 40 | 60 | 95 |
| DMKO57.1b | 05/10/2016 | 09/08/2016 | M | Wildtype | 57 | 8.1 | Noise | Noise +1 days | 40 | 60 | 30 | 40 | 80 | 90 | 95 | 95 | 95 |
| DMKO57.1b | 07/10/2016 | 09/08/2016 | M | Wildtype | 59 | 8.4 | Noise | Noise +3 days | 35 | 60 | 30 | 25 | 75 | 85 | 95 | 95 | 95 |
| DMKO57.1b | 11/10/2016 | 09/08/2016 | M | Wildtype | 63 | 9.0 | Noise | Noise + 7 days | 35 | 45 | 30 | 20 | 70 | 80 | 90 | 95 | 95 |
| DMKO57.1b | 18/10/2016 | 09/08/2016 | M | Wildtype | 70 | 10.0 | Noise | Noise + 14 days | 25 | 55 | 40 | 15 | 20 | 25 | 45 | 85 | 95 |
| DMKO57.1b | 01/11/2016 | 09/08/2016 | M | Wildtype | 84 | 12.0 | Noise | Noise + 28 days | 25 | 65 | 35 | 5 | 25 | 30 | 70 | 95 | 95 |
| DMKO57.1c | 03/10/2016 | 09/08/2016 | F | Heterozygote | 55 | 7.9 | Control (no noise) | Pre noise exposure | 25 | 65 | 30 | 20 | 20 | 30 | 55 | 95 | 95 |
| DMKO57.1c | 05/10/2016 | 09/08/2016 | F | Heterozygote | 57 | 8.1 | Control (no noise) | Noise +1 days | 30 | 60 | 30 | 15 | 20 | 35 | 55 | 95 | 95 |
| DMKO57.1c | 07/10/2016 | 09/08/2016 | F | Heterozygote | 59 | 8.4 | Control (no noise) | Noise +3 days | 30 | 60 | 30 | 15 | 25 | 30 | 55 | 95 | 95 |
| DMKO57.1c | 11/10/2016 | 09/08/2016 | F | Heterozygote | 63 | 9.0 | Control (no noise) | Noise + 7 days | 35 | 65 | 35 | 20 | 25 | 40 | 55 | 95 | 95 |
| DMKO57.1c | 18/10/2016 | 09/08/2016 | F | Heterozygote | 70 | 10.0 | Control (no noise) | Noise + 14 days | 25 | 65 | 30 | 10 | 15 | 25 | 50 | 90 | 95 |
| DMKO57.1c | 01/11/2016 | 09/08/2016 | F | Heterozygote | 84 | 12.0 | Control (no noise) | Noise + 28 days | 30 | 60 | 35 | 10 | 20 | 35 | 75 | 95 | 95 |
| DMKO57.1e | 03/10/2016 | 09/08/2016 | F | Heterozygote | 55 | 7.9 | Control (no noise) | Pre noise exposure | 30 | 80 | 35 | 35 | 35 | 50 | 85 | 95 | 95 |
| DMKO57.1e | 05/10/2016 | 09/08/2016 | F | Heterozygote | 57 | 8.1 | Control (no noise) | Noise +1 days | 25 | 75 | 40 | 20 | 30 | 40 | 80 | 95 | 95 |
| DMKO57.1e | 07/10/2016 | 09/08/2016 | F | Heterozygote | 59 | 8.4 | Control (no noise) | Noise +3 days | 30 | 60 | 35 | 20 | 20 | 30 | 60 | 95 | 95 |
| DMKO57.1e | 11/10/2016 | 09/08/2016 | F | Heterozygote | 63 | 9.0 | Control (no noise) | Noise + 7 days | 30 | 65 | 40 | 20 | 35 | 35 | 85 | 95 | 95 |
| DMKO57.1e | 18/10/2016 | 09/08/2016 | F | Heterozygote | 70 | 10.0 | Control (no noise) | Noise + 14 days | 25 | 65 | 35 | 20 | 20 | 35 | 80 | 95 | 95 |
| DMKO57.1e | 01/11/2016 | 09/08/2016 | F | Heterozygote | 84 | 12.0 | Control (no noise) | Noise + 28 days | 25 | 65 | 30 | 10 | 20 | 35 | 95 | 95 | 95 |

Linear mixed model Bonferroni-corrected p-values

Pre-noise

| | Click | 3kHz | 6kHz | 12kHz | 18kHz | 24kHz | 30kHz | 36kHz | 42kHz |
|---|---|---|---|---|---|---|---|---|---|
| Wildtype unexposed vs. wildtype noise-exposed | 1 | 1 | 1 | 1 | 1 | 1 | 1 | 1 | 0.4 |
| Wildtype unexposed vs. heterozygote unexposed | 1 | 1 | 1 | 0.121 | 0.543 | 1 | 0.183 | <0.001 | <0.001 |
| Wildtype unexposed vs. heterozygote noise-exposed | 1 | 1 | 1 | 1 | 1 | 1 | 1 | 0.006 | <0.001 |
| Heterozygote unexposed vs. wildtype noise-exposed | 1 | 0.356 | 1 | 0.202 | 0.373 | 1 | 0.003 | <0.001 | <0.001 |
| Wildtype noise-exposed vs. heterozygote noise-exposed | 1 | 1 | 1 | 1 | 1 | 1 | 0.795 | <0.001 | 0.042 |
| Heterozygote unexposed vs. heterozygote noise-exposed | 1 | 1 | 1 | 0.521 | 1 | 1 | 0.259 | 0.042 | 0.921 |

Noise+1day

| | Click | 3kHz | 6kHz | 12kHz | 18kHz | 24kHz | 30kHz | 36kHz | 42kHz |
|---|---|---|---|---|---|---|---|---|---|
| Wildtype unexposed vs. wildtype noise-exposed | 0.152 | 1 | 1 | <0.001 | <0.001 | <0.001 | <0.001 | <0.001 | <0.001 |
| Wildtype unexposed vs. heterozygote unexposed | 1 | 0.456 | 1 | 1 | 0.701 | 0.284 | 0.002 | <0.001 | <0.001 |
| Wildtype unexposed vs. heterozygote noise-exposed | 0.055 | 1 | 1 | <0.001 | <0.001 | <0.001 | <0.001 | <0.001 | <0.001 |
| Heterozygote unexposed vs. wildtype noise-exposed | 1 | 1 | 1 | 0.029 | <0.001 | <0.001 | <0.001 | 1 | 1 |
| Wildtype noise-exposed vs. heterozygote noise-exposed | 1 | 1 | 1 | 1 | 1 | 1 | 1 | 1 | 1 |
| Heterozygote unexposed vs. heterozygote noise-exposed | 0.948 | 1 | 1 | 0.001 | <0.001 | <0.001 | <0.001 | 1 | 1 |

Noise+3 days

| | Click | 3kHz | 6kHz | 12kHz | 18kHz | 24kHz | 30kHz | 36kHz | 42kHz |
|---|---|---|---|---|---|---|---|---|---|
| Wildtype unexposed vs. wildtype noise-exposed | 1 | 1 | 1 | 1 | <0.001 | <0.001 | <0.001 | <0.001 | <0.001 |
| Wildtype unexposed vs. heterozygote unexposed | 1 | 1 | 1 | 1 | 1 | 1 | 0.644 | <0.001 | <0.001 |
| Wildtype unexposed vs. heterozygote noise-exposed | 0.512 | 1 | 0.932 | 0.285 | <0.001 | <0.001 | <0.001 | <0.001 | <0.001 |
| Heterozygote unexposed vs. wildtype noise-exposed | 1 | 0.826 | 1 | 1 | <0.001 | <0.001 | <0.001 | 1 | 1 |
| Wildtype noise-exposed vs. heterozygote noise-exposed | 1 | 0.49 | 1 | 1 | 1 | 1 | 1 | 1 | 1 |
| Heterozygote unexposed vs. heterozygote noise-exposed | 1 | 1 | 1 | 1 | <0.001 | <0.001 | <0.001 | 1 | 1 |

Noise+7 days

| | Click | 3kHz | 6kHz | 12kHz | 18kHz | 24kHz | 30kHz | 36kHz | 42kHz |
|---|---|---|---|---|---|---|---|---|---|
| Wildtype unexposed vs. wildtype noise-exposed | 1 | 1 | 1 | 1 | 0.093 | <0.001 | 0.002 | 0.03 | 0.021 |
| Wildtype unexposed vs. heterozygote unexposed | 1 | 1 | 1 | 0.726 | 0.642 | 0.444 | 0.138 | <0.001 | <0.001 |
| Wildtype unexposed vs. heterozygote noise-exposed | 1 | 1 | 1 | 0.747 | 0.88 | 0.001 | 0.004 | <0.001 | <0.001 |
| Heterozygote unexposed vs. wildtype noise-exposed | 1 | 0.294 | 1 | 1 | 1 | 0.147 | 1 | 0.02 | 0.263 |
| Wildtype noise-exposed vs. heterozygote noise-exposed | 1 | 0.381 | 0.768 | 1 | 1 | 1 | 1 | 0.944 | 0.974 |
| Heterozygote unexposed vs. heterozygote noise-exposed | 1 | 1 | 1 | 1 | 1 | 0.292 | 1 | 0.737 | 1 |

Noise+14 days

| | Click | 3kHz | 6kHz | 12kHz | 18kHz | 24kHz | 30kHz | 36kHz | 42kHz |
|---|---|---|---|---|---|---|---|---|---|
| Wildtype unexposed vs. wildtype noise-exposed | 1 | 1 | 1 | 1 | 1 | 1 | 1 | 0.397 | 0.079 |
| Wildtype unexposed vs. heterozygote unexposed | 1 | 1 | 1 | 0.783 | 0.919 | 1 | 0.058 | <0.001 | <0.001 |
| Wildtype unexposed vs. heterozygote noise-exposed | 1 | 1 | 1 | 1 | 1 | 1 | 1 | 0.002 | 0.002 |
| Heterozygote unexposed vs. wildtype noise-exposed | 1 | 1 | 1 | 1 | 0.715 | 1 | 0.079 | 0.001 | 0.719 |
| Wildtype noise-exposed vs. heterozygote noise-exposed | 1 | 1 | 1 | 1 | 1 | 1 | 1 | 0.505 | 1 |
| Heterozygote unexposed vs. heterozygote noise-exposed | 1 | 1 | 1 | 1 | 1 | 1 | 0.899 | 0.159 | 1 |

Noise+28 days

| | Click | 3kHz | 6kHz | 12kHz | 18kHz | 24kHz | 30kHz | 36kHz | 42kHz |
|---|---|---|---|---|---|---|---|---|---|
| Wildtype unexposed vs. wildtype noise-exposed | 1 | 0.619 | 1 | 1 | 1 | 1 | 1 | <0.001 | 0.01 |
| Wildtype unexposed vs. heterozygote unexposed | 1 | 1 | 1 | 1 | 1 | 0.929 | 0.001 | <0.001 | 0.01 |
| Wildtype unexposed vs. heterozygote noise-exposed | 1 | 0.357 | 1 | 1 | 1 | 0.738 | 0.096 | 0.001 | 0.01 |
| Heterozygote unexposed vs. wildtype noise-exposed | 1 | 1 | 1 | 1 | 1 | 1 | 0.005 | 1 | 1 |
| Wildtype noise-exposed vs. heterozygote noise-exposed | 1 | 1 | 1 | 1 | 1 | 0.974 | 0.33 | 1 | 1 |
| Heterozygote unexposed vs. heterozygote noise-exposed | 1 | 1 | 1 | 1 | 1 | 1 | 0.897 | 0.254 | 1 |

**Noise latency data for Fig. 2**

Average

| | Noise status | Noise timing | kHz | 0dB | 5dB | 10dB | 15dB | 20dB | 25dB | 30dB | 35dB | 40dB | 45dB | 50dB | 55dB | 60dB | 65dB | 70dB | 75dB | 80dB | 85dB | 90dB | 95dB |
|---|---|---|---|---|---|---|---|---|---|---|---|---|---|---|---|---|---|---|---|---|---|---|---|
| Wildtype | Control | Pre noise exposure | 12 | 0.073333 | 0.183333 | 0.426667 | 0.468333 | 0.636667 | 0.725 | 0.936667 | 1.228333 | 1.44 | 1.61 | 1.983333 | 1.931667 | 2.361667 | 2.895 | 3.63 | 4.268333 | 4.401667 | 4.661667 | 5.074 | 5.172 |
| Heterozygote | Control | Pre noise exposure | 12 | 0.173333 | 0.291667 | 0.155 | 0.341667 | 0.396667 | 0.495 | 0.608333 | 0.795 | 0.963333 | 1.115 | 1.281667 | 1.411667 | 1.545 | 1.703333 | 2.046667 | 2.35 | 2.743333 | 3.18 | 3.485 | 3.573333 |
| Wildtype | Noise | Pre noise exposure | 12 | 0.238333 | 0.261667 | 0.458333 | 0.396667 | 0.63 | 0.823333 | 1.041667 | 1.251667 | 1.575 | 1.726667 | 2.018333 | 2.113333 | 2.38 | 3.051667 | 3.976667 | 4.21 | 4.191667 | 4.611667 | 4.9 | 4.995 |
| Heterozygote | Noise | Pre noise exposure | 12 | 0.251667 | 0.056667 | 0.238333 | 0.391667 | 0.481667 | 0.865 | 0.898333 | 1.106667 | 1.383333 | 1.536667 | 1.64 | 1.9 | 1.838333 | 2.385 | 2.845 | 3.34 | 3.586667 | 4.095 | 4.67 | 4.846667 |
| Wildtype | Control | Pre noise exposure | 24 | | | 0.078333 | 0.241667 | 0.253333 | 0.311667 | 0.498333 | 0.683333 | 0.683333 | 1.05 | 1.166667 | 1.378333 | 1.791667 | 1.916667 | 2.338333 | 2.783333 | 3.073333 | 3.3 | 3.546 | 3.798 |
| Heterozygote | Control | Pre noise exposure | 24 | | | 0.102 | 0.144 | 0.064 | 0.292 | 0.044 | 0.258 | 0.612 | 0.522 | 0.916 | 1.158 | 1.384 | 1.58 | 1.772 | 1.902 | 2.022 | 2.106 | 2.288 | 2.498 |
| Wildtype | Noise | Pre noise exposure | 24 | | | 0.14 | 0.068333 | 0.211667 | 0.335 | 0.348333 | 0.741667 | 0.816667 | 0.916667 | 1.335 | 1.42 | 1.776667 | 2.043333 | 2.506667 | 2.97 | 3.251667 | 3.518333 | 3.721667 | 4.173333 |
| Heterozygote | Noise | Pre noise exposure | 24 | | | 0.263333 | 0.121667 | 0.211667 | 0.311667 | 0.505 | 0.496667 | 0.705 | 0.878333 | 1.108333 | 1.316667 | 1.506667 | 1.901667 | 1.963333 | 2.19 | 2.441667 | 2.623333 | 2.926667 | 3.216667 |
| Wildtype | Control | Noise +1 day | 12 | 0.06 | 0.256667 | 0.245 | 0.538333 | 0.718333 | 0.813333 | 0.928333 | 1.251667 | 1.623333 | 1.771667 | 2.023333 | 2.213333 | 2.503333 | 3.083333 | 3.958333 | 4.514 | 4.724 | 5 | 5.6125 | 5.9975 |
| Heterozygote | Control | Noise +1 day | 12 | 0.13 | 0.115 | 0.101667 | 0.298333 | 0.36 | 0.5 | 0.636667 | 0.853333 | 1.058333 | 1.223333 | 1.426667 | 1.688333 | 1.706667 | 1.923333 | 2.363333 | 2.658333 | 3.236667 | 3.811667 | 4.268333 | 4.631667 |
| Wildtype | Noise | Noise +1 day | 12 | 0.0275 | 0.065 | 0.0675 | 0.046 | 0.196667 | 0.266667 | 0.293333 | 0.446667 | 0.518333 | 0.686667 | 0.888333 | 1.348333 | 1.87 | 2.168333 | 2.518333 | 2.836667 | 3.52 | 4.085 | 4.485 | 4.393333 |
| Heterozygote | Noise | Noise +1 day | 12 | 0.044 | 0.062 | 0.122 | 0.146 | 0.101667 | 0.166667 | 0.135 | 0.105 | 0.263333 | 0.258333 | 0.591667 | 0.793333 | 1.236667 | 1.43 | 1.786667 | 2.365 | 2.558333 | 3.15 | 3.678333 | 3.933333 |
| Wildtype | Control | Noise +1 day | 24 | | | 0.2 | 0.186667 | 0.233333 | 0.38 | 0.525 | 0.538333 | 0.781667 | 0.978333 | 1.355 | 1.703333 | 1.983333 | 2.381667 | 2.898333 | 2.808 | 3.398 | 3.628 | 4.0375 | 4.45 |
| Heterozygote | Control | Noise +1 day | 24 | | | 0.238333 | 0.101667 | 0.103333 | 0.211667 | 0.236667 | 0.236667 | 0.295 | 0.451667 | 0.735 | 0.93 | 1.121667 | 1.438333 | 1.518333 | 1.763333 | 1.915 | 2.13 | 2.188333 | 2.476667 |
| Wildtype | Noise | Noise +1 day | 24 | | | | | 0.04 | 0.22 | 0.21 | 0.66 | 0.01 | 0.13 | 0.05 | 0.08 | 0.13 | 0.05 | 0.14 | 0.39 | 0.27 | 0.77 | 0.77 | 1.01 |
| Heterozygote | Noise | Noise +1 day | 24 | | | | | | | | | | | | | | | | | | | | |
| Wildtype | Control | Noise +3 days | 12 | 0.201667 | 0.266667 | 0.285 | 0.483333 | 0.671667 | 0.796667 | 0.955 | 1.275 | 1.571667 | 1.85 | 2.04 | 2.286667 | 2.475 | 2.968333 | 4.096667 | 4.548333 | 4.685 | 4.948333 | 5.543333 | 5.668333 |
| Heterozygote | Control | Noise +3 days | 12 | 0.066667 | 0.116667 | 0.16 | 0.198333 | 0.341667 | 0.43 | 0.685 | 0.746667 | 0.893333 | 1.266667 | 1.275 | 1.516667 | 1.528333 | 1.63 | 2.043333 | 2.278333 | 2.6 | 2.951667 | 3.376667 | 3.58 |
| Wildtype | Noise | Noise +3 days | 12 | 0.136667 | 0.138333 | 0.271667 | 0.293333 | 0.413333 | 0.696667 | 0.823333 | 1.091667 | 1.358333 | 1.728333 | 2.055 | 2.233333 | 2.405 | 2.761667 | 2.918333 | 3.368333 | 3.546667 | 3.915 | 4.25 | 4.236667 |
| Heterozygote | Noise | Noise +3 days | 12 | 0.155 | 0.09 | 0.275 | 0.308333 | 0.273333 | 0.383333 | 0.653333 | 0.62 | 0.86 | 0.9 | 1.068333 | 1.35 | 1.48 | 1.851667 | 1.878333 | 2.168333 | 2.363333 | 2.866667 | 3.153333 | 3.145 |
| Wildtype | Control | Noise +3 days | 24 | | | 0.298333 | 0.151667 | 0.32 | 0.275 | 0.345 | 0.641667 | 0.871667 | 1.065 | 1.273333 | 1.475 | 1.876667 | 2.013333 | 2.493333 | 2.708333 | 3.038333 | 3.286667 | 3.628333 | 4.091667 |
| Heterozygote | Control | Noise +3 days | 24 | | | 0.108333 | 0.106667 | 0.113333 | 0.31 | 0.245 | 0.486667 | 0.51 | 0.483333 | 0.785 | 0.963333 | 1.236667 | 1.303333 | 1.623333 | 1.575 | 1.82 | 1.763333 | 1.953333 | 2.051667 |
| Wildtype | Noise | Noise +3 days | 24 | | | 0.25 | 0.226667 | 0.16 | 0.203333 | 0.133333 | 0.2 | 0.146667 | 0.083333 | 0.286667 | 0.38 | 0.49 | 0.473333 | 0.596667 | 0.833333 | 1.06 | 1.103333 | 1.413333 | 1.436667 |
| Heterozygote | Noise | Noise +3 days | 24 | | | -0.02 | 0.16 | 0 | 0.1 | 0.06 | 0.26 | 0.21 | 0.33 | -0.01 | 0.05 | 0.15 | 0.44 | 0.62 | 0.66 | 0.83 | 0.44 | 0.72 | 1.02 |
| Wildtype | Control | Noise +7 days | 12 | 0.073333 | 0.151667 | 0.331667 | 0.526667 | 0.576667 | 0.796667 | 1.098333 | 1.181667 | 1.288333 | 1.74 | 2.058333 | 2.15 | 2.525 | 3.021667 | 3.92 | 4.588333 | 4.8 | 4.888333 | 5.43 | 5.76 |
| Heterozygote | Control | Noise +7 days | 12 | 0.11 | 0.151667 | 0.261667 | 0.265 | 0.433333 | 0.451667 | 0.586667 | 0.771667 | 0.838333 | 1.071667 | 1.07 | 1.396667 | 1.386667 | 1.681667 | 1.928333 | 2.218333 | 2.508333 | 2.968333 | 3.203333 | 3.393333 |
| Wildtype | Noise | Noise +7 days | 12 | 0.118333 | 0.131667 | 0.151667 | 0.361667 | 0.396667 | 0.685 | 0.951667 | 1.106667 | 1.445 | 1.681667 | 1.971667 | 2.276667 | 2.448333 | 2.996667 | 3.328333 | 3.633333 | 3.94 | 4.203333 | 4.293333 | 4.298333 |
| Heterozygote | Noise | Noise +7 days | 12 | 0.076667 | 0.175 | 0.158333 | 0.193333 | 0.506667 | 0.528333 | 0.666667 | 0.766667 | 0.895 | 1.163333 | 1.275 | 1.425 | 1.683333 | 2.028333 | 2.275 | 2.315 | 2.688333 | 3.005 | 3.24 | 3.085 |
| Wildtype | Control | Noise +7 days | 24 | | | 0.096667 | 0.163333 | 0.171667 | 0.266667 | 0.47 | 0.665 | 0.77 | 1.048333 | 1.295 | 1.496667 | 1.658333 | 1.941667 | 2.266667 | 2.733333 | 3.093333 | 3.341667 | 3.608333 | 4.216667 |
| Heterozygote | Control | Noise +7 days | 24 | | | 0.114 | 0.288 | 0.212 | 0.202 | 0.384 | 0.454 | 0.444 | 0.654 | 0.854 | 1.076 | 1.18 | 1.418 | 1.75 | 1.778 | 1.844 | 1.938 | 2.134 | 2.198 |
| Wildtype | Noise | Noise +7 days | 24 | | | 0.114 | 0.128 | 0.16 | 0.042 | 0.336 | 0.292 | 0.348 | 0.476 | 0.644 | 0.72 | 0.808 | 1.14 | 1.038 | 1.476 | 1.51 | 1.522 | 1.844 | 2.322 |
| Heterozygote | Noise | Noise +7 days | 24 | | | 0.096667 | 0.16 | 0.113333 | 0.293333 | 0.38 | 0.48 | 0.533333 | 0.49 | 0.693333 | 0.713333 | 0.823333 | 0.856667 | 0.996667 | 1.223333 | 1.25 | 1.37 | 1.563333 | 1.766667 |
| Wildtype | Control | Noise +14 days | 12 | 0.203333 | 0.085 | 0.34 | 0.45 | 0.535 | 0.515 | 0.926667 | 0.948333 | 1.248333 | 1.478333 | 1.693333 | 1.818333 | 2.095 | 2.578333 | 3.291667 | 3.586667 | 3.67 | 3.901667 | 4.326667 | 4.39 |
| Heterozygote | Control | Noise +14 days | 12 | 0.163333 | 0.121667 | 0.148333 | 0.311667 | 0.28 | 0.386667 | 0.693333 | 0.76 | 0.946667 | 0.985 | 1.14 | 1.321667 | 1.536667 | 1.681667 | 1.988333 | 2.155 | 2.421667 | 2.783333 | 3.181667 | 3.186667 |
| Wildtype | Noise | Noise +14 days | 12 | 0.23 | 0.116667 | 0.406667 | 0.433333 | 0.6 | 0.75 | 0.851667 | 1.181667 | 1.323333 | 1.793333 | 1.851667 | 2.178333 | 2.573333 | 3.183333 | 3.933333 | 4.293333 | 4.205 | 4.216667 | 4.461667 | 4.116667 |
| Heterozygote | Noise | Noise +14 days | 12 | 0.196667 | 0.166667 | 0.281667 | 0.295 | 0.355 | 0.511667 | 0.626667 | 0.896667 | 0.98 | 1.121667 | 1.235 | 1.508333 | 1.716667 | 1.961667 | 2.301667 | 2.431667 | 2.515 | 2.9 | 3.168333 | 2.94 |
| Wildtype | Control | Noise +14 days | 24 | | | 0.208333 | 0.196667 | 0.331667 | 0.386667 | 0.408333 | 0.708333 | 0.746667 | 0.851667 | 0.941667 | 1.286667 | 1.418333 | 1.815 | 2.05 | 2.323333 | 2.618333 | 2.913333 | 3.308333 | 3.478333 |
| Heterozygote | Control | Noise +14 days | 24 | | | 0.126667 | 0.158333 | 0.205 | 0.255 | 0.338333 | 0.401667 | 0.538333 | 0.636667 | 0.773333 | 0.936667 | 1.12 | 1.323333 | 1.496667 | 1.678333 | 1.956667 | 1.931667 | 2.115 | 2.381667 |
| Wildtype | Noise | Noise +14 days | 24 | | | 0.118333 | 0.066667 | 0.195 | 0.345 | 0.411667 | 0.628333 | 0.878333 | 0.938333 | 1.333333 | 1.448333 | 1.573333 | 1.921667 | 2.12 | 2.365 | 2.625 | 2.755 | 2.945 | 3.306667 |
| Heterozygote | Noise | Noise +14 days | 24 | | | 0.176667 | 0.043333 | 0.133333 | 0.18 | 0.255 | 0.433333 | 0.385 | 0.603333 | 0.581667 | 0.876667 | 0.935 | 1.048333 | 1.13 | 1.203333 | 1.366667 | 1.436667 | 1.546667 | 1.718333 |
| Wildtype | Control | Noise +28 days | 12 | 0.138 | 0.228 | 0.216667 | 0.348333 | 0.498333 | 0.546667 | 0.673333 | 0.928333 | 1.041667 | 1.39 | 1.466667 | 1.795 | 1.856667 | 2.411667 | 2.936667 | 3.238333 | 3.483333 | 3.841667 | 4.18 | 4.234 |
| Heterozygote | Control | Noise +28 days | 12 | 0.163333 | 0.166667 | 0.188333 | 0.298333 | 0.315 | 0.478333 | 0.613333 | 0.698333 | 0.975 | 1.09 | 1.315 | 1.42 | 1.621667 | 1.905 | 1.981667 | 2.226667 | 2.493333 | 2.828333 | 3.23 | 3.266667 |
| Wildtype | Noise | Noise +28 days | 12 | 0.188333 | 0.228333 | 0.293333 | 0.391667 | 0.58 | 0.716667 | 0.888333 | 1.066667 | 1.43 | 1.666667 | 1.99 | 2.105 | 2.56 | 3.361667 | 3.885 | 4.043333 | 4.145 | 4.195 | 4.56 | 4.363333 |
| Heterozygote | Noise | Noise +28 days | 12 | 0.061667 | 0.016667 | 0.178333 | 0.291667 | 0.485 | 0.655 | 0.7 | 0.908333 | 1.166667 | 1.418333 | 1.635 | 1.89 | 2.08 | 2.51 | 2.733333 | 2.893333 | 3.343333 | 3.436667 | 3.678333 | 3.65 |
| Wildtype | Control | Noise +28 days | 24 | | | 0.183333 | 0.218333 | 0.155 | 0.331667 | 0.42 | 0.478333 | 0.681667 | 0.86 | 1.131667 | 1.115 | 1.183333 | 1.651667 | 1.835 | 2.061667 | 2.42 | 2.585 | 2.838333 | 3.268 |
| Heterozygote | Control | Noise +28 days | 24 | | | 0.161667 | 0.133333 | 0.12 | 0.165 | 0.128333 | 0.255 | 0.486667 | 0.408333 | 0.67 | 0.838333 | 0.961667 | 1.193333 | 1.348333 | 1.351667 | 1.533333 | 1.623333 | 1.663333 | 1.853333 |
| Wildtype | Noise | Noise +28 days | 24 | | | 0.16 | 0.203333 | -0.01333 | 0.163333 | 0.375 | 0.515 | 0.521667 | 0.838333 | 1.005 | 1.17 | 1.383333 | 1.55 | 1.898333 | 2.023333 | 2.276667 | 2.481667 | 2.633333 | 3.021667 |
| Heterozygote | Noise | Noise +28 days | 24 | | | 0.09 | 0.068 | 0.112 | 0.142 | 0.378 | 0.392 | 0.47 | 0.592 | 0.736 | 0.98 | 1.166 | 1.356 | 1.258 | 1.378 | 1.362 | 1.62 | 1.776 | 1.878 |

Standard deviation

| | Noise status | Noise timing | kHz | 0dB | 5dB | 10dB | 15dB | 20dB | 25dB | 30dB | 35dB | 40dB | 45dB | 50dB | 55dB | 60dB | 65dB | 70dB | 75dB | 80dB | 85dB | 90dB | 95dB |
|---|---|---|---|---|---|---|---|---|---|---|---|---|---|---|---|---|---|---|---|---|---|---|---|
| Wildtype | Control | Pre noise exposure | 12 | 0.075498 | 0.041633 | 0.14 | 0.175024 | 0.10504 | 0.096437 | 0.215484 | 0.411987 | 0.41509 | 0.470035 | 0.257164 | 0.164621 | 0.232881 | 0.202073 | 0.356417 | 0.632218 | 0.995507 | 1.205833 | 1.407142 | 1.704127 |
| Heterozygote | Control | Pre noise exposure | 12 | 0.112724 | 0.405828 | 0.279696 | 0.245065 | 0.15358 | 0.176607 | 0.15766 | 0.126293 | 0.245493 | 0.335425 | 0.279887 | 0.326767 | 0.338984 | 0.368004 | 0.623496 | 0.707842 | 0.818136 | 1.138051 | 1.029869 | 1.209027 |
| Wildtype | Noise | Pre noise exposure | 12 | 0.196409 | 0.09347 | 0.164732 | 0.157818 | 0.222711 | 0.244104 | 0.303079 | 0.256859 | 0.361925 | 0.39134 | 0.470209 | 0.425754 | 0.45396 | 0.467351 | 0.644505 | 0.80635 | 0.724498 | 0.983675 | 0.912338 | 1.056877 |
| Heterozygote | Noise | Pre noise exposure | 12 | 0.114091 | 0.194182 | 0.191981 | 0.245554 | 0.095167 | 0.120125 | 0.293967 | 0.241468 | 0.374789 | 0.504804 | 0.270333 | 0.46493 | 0.235917 | 0.222868 | 0.338866 | 0.413812 | 0.512627 | 0.693073 | 0.993821 | 1.110075 |
| Wildtype | Control | Pre noise exposure | 24 | | | 0.126623 | 0.072111 | 0.122882 | 0.15308 | 0.395769 | 0.217332 | 0.23516 | 0.377536 | 0.188768 | 0.459021 | 0.484286 | 0.618574 | 0.768635 | 0.869617 | 0.633745 | 0.951332 | 0.388909 | 0.572756 |
| Heterozygote | Control | Pre noise exposure | 24 | | | 0.105688 | 0.119708 | 0.116533 | 0.146356 | 0.112383 | 0.126965 | 0.208375 | 0.155788 | 0.207196 | 0.300366 | 0.260346 | 0.22902 | 0.12276 | 0.10663 | 0.340323 | 0.340632 | 0.246718 | 0.322056 |
| Wildtype | Noise | Pre noise exposure | 24 | | | 0.16 | 0.186485 | 0.133479 | 0.059917 | 0.19125 | 0.336536 | 0.274566 | 0.298038 | 0.311946 | 0.4155 | 0.35859 | 0.468259 | 0.39185 | 0.609196 | 0.659679 | 0.634206 | 0.936876 | 0.939929 |
| Heterozygote | Noise | Pre noise exposure | 24 | | | 0.173973 | 0.125605 | 0.281739 | 0.074944 | 0.165015 | 0.192319 | 0.212297 | 0.144695 | 0.174633 | 0.187688 | 0.343841 | 0.379706 | 0.445361 | 0.450156 | 0.605158 | 0.567121 | 0.63456 | 0.660747 |
| Wildtype | Control | Noise +1 day | 12 | 0.091652 | 0.073711 | 0.283784 | 0.355012 | 0.079373 | 0.271539 | 0.338575 | 0.157162 | 0.345302 | 0.236925 | 0.245832 | 0.26727 | 0.015275 | 0.305123 | 0.474377 | 0.661614 | 0.80829 | 1.32106 | 1.549097 | 1.7365 |
| Heterozygote | Control | Noise +1 day | 12 | 0.108074 | 0.098336 | 0.143724 | 0.159677 | 0.148728 | 0.159875 | 0.339156 | 0.314176 | 0.355382 | 0.384014 | 0.513602 | 0.62914 | 0.546175 | 0.736306 | 0.85301 | 1.056114 | 1.147356 | 0.884927 | 0.910328 | 0.998547 |
| Wildtype | Noise | Noise +1 day | 12 | 0.286167 | 0.224277 | 0.206942 | 0.221653 | 0.226156 | 0.133666 | 0.1541 | 0.196435 | 0.340201 | 0.320915 | 0.463354 | 0.555173 | 0.503428 | 0.718092 | 0.655665 | 0.786757 | 0.886138 | 0.833661 | 1.019642 | 1.1644 |
| Heterozygote | Noise | Noise +1 day | 12 | 0.179248 | 0.142548 | 0.105451 | 0.464467 | 0.140202 | 0.108012 | 0.243619 | 0.215569 | 0.373988 | 0.403902 | 0.442828 | 0.411712 | 0.516165 | 0.805034 | 0.822476 | 0.77291 | 0.93118 | 0.901443 | 0.75785 | 0.726021 |
| Wildtype | Control | Noise +1 day | 24 | | | 0.276225 | 0.247857 | 0.295466 | 0.086217 | 0.3188 | 0.219317 | 0.1823 | 0.160416 | 0.318957 | 0.357911 | 0.272274 | 0.521153 | 0.440341 | 0.406325 | 0.767876 | 0.769502 | 0.822456 | 0.721965 |
| Heterozygote | Control | Noise +1 day | 24 | | | 0.108151 | 0.206728 | 0.242872 | 0.128284 | 0.088015 | 0.093737 | 0.136931 | 0.296609 | 0.330015 | 0.451619 | 0.392195 | 0.431389 | 0.3867 | 0.561237 | 0.693361 | 0.819732 | 0.773884 | 0.739883 |
| Wildtype | Noise | Noise +1 day | 24 | | | | | | | | | | | | | | | | | | | | |
| Heterozygote | Noise | Noise +1 day | 24 | | | | | | | | | | | | | | | | | | | | |
| Wildtype | Control | Noise +3 days | 12 | 0.210087 | 0.15214 | 0.233302 | 0.180518 | 0.187768 | 0.169784 | 0.32953 | 0.385785 | 0.348392 | 0.283125 | 0.308804 | 0.256489 | 0.360597 | 0.726895 | 0.839111 | 0.971008 | 1.045079 | 1.000388 | 1.202375 | 0.949809 |
| Heterozygote | Control | Noise +3 days | 12 | 0.095847 | 0.081404 | 0.080994 | 0.214981 | 0.119234 | 0.16769 | 0.13896 | 0.298909 | 0.297568 | 0.129099 | 0.338925 | 0.370603 | 0.500297 | 0.453034 | 0.409422 | 0.474064 | 0.62514 | 0.601645 | 0.699333 | 0.768713 |
| Wildtype | Noise | Noise +3 days | 12 | 0.210016 | 0.133479 | 0.08085 | 0.328978 | 0.120941 | 0.330131 | 0.336908 | 0.454375 | 0.423765 | 0.542196 | 0.5362 | 0.413021 | 0.484221 | 0.443821 | 0.444136 | 0.58335 | 0.572806 | 0.733096 | 0.47879 | 0.771354 |
| Heterozygote | Noise | Noise +3 days | 12 | 0.299783 | 0.203568 | 0.180638 | 0.203413 | 0.146652 | 0.163422 | 0.336789 | 0.329302 | 0.2555 | 0.164924 | 0.365919 | 0.456508 | 0.396232 | 0.496283 | 0.43742 | 0.348449 | 0.531212 | 0.454474 | 0.698303 | 0.739453 |
| Wildtype | Control | Noise +3 days | 24 | | | 0.11089 | 0.116003 | 0.119666 | 0.188441 | 0.145705 | 0.194362 | 0.26057 | 0.227662 | 0.30605 | 0.383862 | 0.43953 | 0.515623 | 0.700619 | 0.874972 | 0.994393 | 1.07725 | 0.864024 | 1.013241 |
| Heterozygote | Control | Noise +3 days | 24 | | | 0.106286 | 0.156034 | 0.148279 | 0.181879 | 0.085499 | 0.094375 | 0.226097 | 0.290631 | 0.257119 | 0.318476 | 0.466719 | 0.370819 | 0.418649 | 0.586677 | 0.645446 | 0.553124 | 0.618633 | 0.432408 |
| Wildtype | Noise | Noise +3 days | 24 | | | 0.043589 | 0.172143 | 0.16 | 0.184481 | 0.196044 | 0.045826 | 0.205994 | 0.130128 | 0.251462 | 0.264575 | 0.327414 | 0.195533 | 0.536781 | 0.502925 | 0.491121 | 0.582866 | 0.754807 | 0.477214 |
| Heterozygote | Noise | Noise +3 days | 24 | | | | | | | | | | | | | | | | | | | | |
| Wildtype | Control | Noise +7 days | 12 | 0.135302 | 0.138046 | 0.074677 | 0.151877 | 0.128478 | 0.225359 | 0.416049 | 0.563646 | 0.328476 | 0.234521 | 0.367718 | 0.361054 | 0.28571 | 0.674638 | 0.676195 | 0.876457 | 0.946383 | 0.773936 | 0.924467 | 1.078796 |
| Heterozygote | Control | Noise +7 days | 12 | 0.151129 | 0.18957 | 0.167859 | 0.200674 | 0.046332 | 0.121724 | 0.219697 | 0.118561 | 0.273453 | 0.325786 | 0.321745 | 0.349781 | 0.420079 | 0.406764 | 0.539237 | 0.710195 | 0.601778 | 0.688256 | 0.441029 | 0.56578 |
| Wildtype | Noise | Noise +7 days | 12 | 0.188618 | 0.155231 | 0.195695 | 0.169165 | 0.150953 | 0.239144 | 0.14838 | 0.293235 | 0.347951 | 0.3657 | 0.442873 | 0.386713 | 0.387681 | 0.55601 | 0.596403 | 0.609579 | 0.823335 | 0.747119 | 0.554749 | 0.755206 |
| Heterozygote | Noise | Noise +7 days | 12 | 0.072572 | 0.158082 | 0.139344 | 0.351663 | 0.129409 | 0.176796 | 0.247279 | 0.1541 | 0.231409 | 0.356688 | 0.391804 | 0.216587 | 0.478316 | 0.268955 | 0.458378 | 0.736851 | 0.766875 | 0.779532 | 0.925592 | 0.918755 |
| Wildtype | Control | Noise +7 days | 24 | | | 0.260205 | 0.160083 | 0.246367 | 0.142782 | 0.180222 | 0.155274 | 0.237318 | 0.347069 | 0.412783 | 0.443832 | 0.32202 | 0.42282 | 0.520871 | 0.680225 | 0.709779 | 0.679718 | 0.663187 | 0.951854 |
| Heterozygote | Control | Noise +7 days | 24 | | | 0.041593 | 0.14025 | 0.224989 | 0.218105 | 0.188361 | 0.142934 | 0.153232 | 0.200325 | 0.12341 | 0.229412 | 0.36187 | 0.49937 | 0.439318 | 0.698119 | 0.725899 | 0.491854 | 0.76885 | 0.810969 |
| Wildtype | Noise | Noise +7 days | 24 | | | 0.136125 | 0.205961 | 0.083666 | 0.222081 | 0.134276 | 0.137186 | 0.157702 | 0.301877 | 0.268849 | 0.49553 | 0.479969 | 0.523116 | 0.659636 | 0.681087 | 0.660606 | 0.73326 | 0.578126 | 0.66047 |
| Heterozygote | Noise | Noise +7 days | 24 | | | 0.233524 | 0.07 | 0.305669 | 0.370045 | 0.20664 | 0.117898 | 0.28746 | 0.419404 | 0.460688 | 0.480139 | 0.678847 | 0.533042 | 0.481283 | 0.603517 | 0.482804 | 0.632218 | 0.479618 | 0.40501 |
| Wildtype | Control | Noise +14 days | 12 | 0.104626 | 0.229935 | 0.205037 | 0.22154 | 0.145155 | 0.22025 | 0.282111 | 0.249031 | 0.447903 | 0.29708 | 0.308847 | 0.161421 | 0.333392 | 0.358966 | 0.526362 | 0.72304 | 0.689957 | 0.898875 | 0.81532 | 0.903659 |
| Heterozygote | Control | Noise +14 days | 12 | 0.129718 | 0.245146 | 0.154844 | 0.095795 | 0.206495 | 0.201362 | 0.164154 | 0.231257 | 0.195721 | 0.344659 | 0.372075 | 0.342836 | 0.243283 | 0.384001 | 0.513202 | 0.670037 | 0.877324 | 0.869176 | 0.984793 | 0.80139 |
| Wildtype | Noise | Noise +14 days | 12 | 0.11296 | 0.117246 | 0.257734 | 0.134114 | 0.084143 | 0.241744 | 0.187874 | 0.117884 | 0.191694 | 0.267108 | 0.408285 | 0.241033 | 0.410544 | 0.369468 | 0.412779 | 0.667912 | 0.611972 | 0.598955 | 0.67131 | 0.685439 |
| Heterozygote | Noise | Noise +14 days | 12 | 0.202846 | 0.155778 | 0.169046 | 0.170265 | 0.110589 | 0.28379 | 0.191485 | 0.225004 | 0.262755 | 0.273965 | 0.356413 | 0.319213 | 0.214911 | 0.374295 | 0.38374 | 0.394331 | 0.649515 | 0.710971 | 0.891009 | 1.072343 |
| Wildtype | Control | Noise +14 days | 24 | | | 0.186806 | 0.219515 | 0.235152 | 0.305527 | 0.203609 | 0.148918 | 0.322532 | 0.368642 | 0.209897 | 0.185975 | 0.275856 | 0.376338 | 0.425253 | 0.530082 | 0.541606 | 0.568249 | 0.662825 | 0.58796 |
| Heterozygote | Control | Noise +14 days | 24 | | | 0.111116 | 0.0608 | 0.100747 | 0.159593 | 0.280387 | 0.157787 | 0.236001 | 0.181181 | 0.318476 | 0.337738 | 0.301463 | 0.502222 | 0.535637 | 0.623808 | 0.741314 | 0.727335 | 0.654546 | 0.798259 |
| Wildtype | Noise | Noise +14 days | 24 | | | 0.129987 | 0.057504 | 0.298446 | 0.142934 | 0.147298 | 0.136149 | 0.221035 | 0.122052 | 0.269493 | 0.160052 | 0.205102 | 0.180157 | 0.112783 | 0.25921 | 0.24615 | 0.368497 | 0.320734 | 0.462846 |
| Heterozygote | Noise | Noise +14 days | 24 | | | 0.18392 | 0.275947 | 0.101127 | 0.107703 | 0.148425 | 0.276671 | 0.281052 | 0.168008 | 0.286455 | 0.312132 | 0.328983 | 0.347874 | 0.530773 | 0.449563 | 0.498705 | 0.583701 | 0.596244 | 0.63041 |
| Wildtype | Control | Noise +28 days | 12 | 0.123572 | 0.10849 | 0.272078 | 0.135265 | 0.275639 | 0.393277 | 0.241219 | 0.206147 | 0.451328 | 0.568049 | 0.466247 | 0.454346 | 0.550297 | 0.678894 | 0.867379 | 0.922917 | 0.954226 | 1.115265 | 1.158775 | 1.294461 |
| Heterozygote | Control | Noise +28 days | 12 | 0.155906 | 0.15384 | 0.144971 | 0.191668 | 0.318292 | 0.364934 | 0.361755 | 0.360134 | 0.318418 | 0.585252 | 0.405648 | 0.645043 | 0.597475 | 0.689862 | 0.725959 | 0.731209 | 0.704746 | 0.803129 | 0.944542 | 0.813994 |
| Wildtype | Noise | Noise +28 days | 12 | 0.105151 | 0.112857 | 0.259281 | 0.146071 | 0.117303 | 0.218235 | 0.175205 | 0.193046 | 0.1772 | 0.227215 | 0.308934 | 0.232616 | 0.371537 | 0.414218 | 0.414861 | 0.468345 | 0.461638 | 0.439397 | 0.558963 | 0.485249 |
| Heterozygote | Noise | Noise +28 days | 12 | 0.187768 | 0.094163 | 0.216556 | 0.180601 | 0.200175 | 0.19685 | 0.088544 | 0.292672 | 0.312068 | 0.212548 | 0.270019 | 0.292233 | 0.319124 | 0.505925 | 0.297366 | 0.206559 | 0.629529 | 0.869659 | 0.895576 | 0.870885 |
| Wildtype | Control | Noise +28 days | 24 | | | 0.110212 | 0.137756 | 0.12661 | 0.272867 | 0.24257 | 0.372957 | 0.111967 | 0.241081 | 0.431575 | 0.365007 | 0.524277 | 0.56616 | 0.697273 | 0.65594 | 0.72208 | 0.695751 | 0.737981 | 0.919658 |
| Heterozygote | Control | Noise +28 days | 24 | | | 0.176569 | 0.13938 | 0.196469 | 0.093968 | 0.250073 | 0.113798 | 0.092232 | 0.147569 | 0.226186 | 0.32951 | 0.493373 | 0.523323 | 0.65052 | 0.585095 | 0.678754 | 0.833179 | 0.698303 | 0.753675 |
| Wildtype | Noise | Noise +28 days | 24 | | | 0.183412 | 0.141939 | 0.174318 | 0.224113 | 0.094816 | 0.151493 | 0.256392 | 0.209515 | 0.293172 | 0.457559 | 0.445092 | 0.539926 | 0.514875 | 0.483391 | 0.505081 | 0.275784 | 0.367677 | 0.497088 |
| Heterozygote | Noise | Noise +28 days | 24 | | | 0.178466 | 0.080747 | 0.159123 | 0.080436 | 0.119875 | 0.185122 | 0.225056 | 0.130652 | 0.295686 | 0.257391 | 0.329591 | 0.383901 | 0.373189 | 0.365677 | 0.438087 | 0.270832 | 0.372867 | 0.483808 |

| Mouse ID | *Mir183/96*$^{dko}$ genotype | Noise status | Noise timing | kHz | 0dB | 5dB | 10dB | 15dB | 20dB | 25dB | 30dB | 35dB | 40dB | 45dB | 50dB | 55dB | 60dB | 65dB | 70dB | 75dB | 80dB | 85dB | 90dB | 95dB |
|---|---|---|---|---|---|---|---|---|---|---|---|---|---|---|---|---|---|---|---|---|---|---|---|---|
| DMKO44.1a | Wildtype | Control | Pre noise exposure | 12 | 0.1 | 0.19 | 0.54 | 0.56 | 0.73 | 0.6 | 0.91 | 0.94 | 1.12 | 1.29 | 1.88 | 1.96 | 2.25 | 2.72 | 3.43 | 3.57 | 3.2 | 3.27 | 3.92 | 3.87 |
| DMKO44.1g | Wildtype | Noise | Pre noise exposure | 12 | 0.36 | 0.3 | 0.29 | 0.63 | 0.39 | 1.28 | 1.43 | 1.41 | 1.99 | 1.72 | 2.66 | 2.59 | 2.6 | 3.47 | 4.46 | 4.83 | 4.74 | 5.06 | 5.5 | 5.76 |
| DMKO44.1c | Wildtype | Noise | Pre noise exposure | 12 | 0.44 | 0.38 | 0.33 | 0.32 | 0.49 | 0.79 | 0.57 | 0.84 | 1.14 | 1.17 | 1.27 | 1.64 | 1.78 | 2.76 | 3.32 | 3.99 | 3.87 | 4.14 | 4.38 | 4.5 |
| DMKO54.1f | Wildtype | Control | Pre noise exposure | 12 | 0.04 | 0.27 | 0.4 | 0.39 | 0.83 | 0.78 | 1.32 | 1.76 | 1.55 | 2.23 | 2.36 | 2.25 | 2.68 | 2.97 | 3.73 | 4.67 | 4.85 | 4.88 | 5.91 | 6.28 |
| DMKO54.1d | Wildtype | Noise | Pre noise exposure | 12 | 0.18 | 0.33 | 0.71 | 0.42 | 0.64 | 0.71 | 1.01 | 1.44 | 1.51 | 1.83 | 2.04 | 2.12 | 2.46 | 3.16 | 4.09 | 3.96 | 3.92 | 4.65 | 5.03 | 4.51 |
| DMKO55.1b | Wildtype | Noise | Pre noise exposure | 12 | -0.09 | 0.22 | 0.55 | 0.21 | 0.64 | 0.63 | 1.06 | 1.24 | 1.51 | 1.72 | 1.93 | 2.06 | 1.96 | 2.38 | 3.25 | 3.4 | 3.68 | 4.36 | 4.93 | 4.78 |
| DMKO55.1a | Wildtype | Control | Pre noise exposure | 12 | -0.05 | 0.25 | 0.26 | 0.74 | 0.62 | 0.75 | 1 | 1.42 | 1.95 | 1.75 | 1.96 | 1.97 | 2.31 | 3.12 | 4.14 | 4.66 | 4.99 | 5.63 | | |
| CNTC20.1a | Wildtype | Control | Pre noise exposure | 12 | 0 | 0.3 | 0.6 | 0.44 | 0.65 | 0.9 | 0.77 | 1.44 | 1.45 | 1.41 | 1.98 | 1.84 | 2.6 | 3.01 | 3.39 | 4.28 | 4.14 | 4.84 | 5.42 | 5.45 |
| CNTC20.1f | Wildtype | Control | Pre noise exposure | 12 | 0.37 | 0 | 0.26 | 0.28 | 0.33 | 0.71 | 0.92 | 0.97 | 1.19 | 1.42 | 1.89 | 1.88 | 2.24 | 2.64 | 3.68 | 4.49 | 4.84 | 4.99 | 5.4 | 5.37 |
| CNTC20.1g | Wildtype | Control | Pre noise exposure | 12 | -0.02 | 0.09 | 0.5 | 0.4 | 0.66 | 0.61 | 0.7 | 0.84 | 1.38 | 1.56 | 1.83 | 1.69 | 2.09 | 2.91 | 3.41 | 3.94 | 4.39 | 4.36 | 4.72 | 4.89 |
| DMKO55.1h | Wildtype | Noise | Pre noise exposure | 12 | 0.16 | 0.22 | 0.53 | 0.52 | 1.04 | 0.89 | 1.29 | 1.51 | 2.02 | 2.37 | 2.34 | 2.61 | 3.04 | 3.64 | 4.9 | 5.51 | 5.4 | 6.2 | 6.1 | 6.69 |
| DMKO57.1b | Wildtype | Noise | Pre noise exposure | 12 | 0.38 | 0.12 | 0.34 | 0.28 | 0.58 | 0.64 | 0.89 | 1.07 | 1.28 | 1.55 | 1.87 | 1.66 | 2.44 | 2.9 | 3.84 | 3.57 | 3.54 | 3.26 | 3.46 | 3.73 |
| DMKO44.1b | Heterozygote | Control | Pre noise exposure | 12 | 0.2 | 0.5 | 0.28 | 0.29 | 0.64 | 0.51 | 0.53 | 0.68 | 0.97 | 1 | 1.29 | 1.4 | 1.66 | 2.02 | 2.27 | 2.74 | 3.47 | 4.64 | 4.69 | 5.16 |
| DMKO44.1d | Heterozygote | Control | Pre noise exposure | 12 | 0.02 | 0.1 | 0.09 | 0.24 | 0.28 | 0.58 | 0.68 | 0.75 | 0.6 | 0.94 | 1 | 1.22 | 1.45 | 1.42 | 1.96 | 2.74 | 3.49 | 4.34 | 4.61 | 4.69 |
| DMKO44.1f | Heterozygote | Noise | Pre noise exposure | 12 | 0.3 | 0.26 | 0.45 | 0.35 | 0.48 | 0.74 | 0.86 | 1.23 | 0.98 | 1.18 | 1.57 | 1.56 | 1.71 | 2.39 | 2.71 | 2.92 | 3 | 3.38 | 3.75 | 3.63 |
| DMKO44.1e | Heterozygote | Noise | Pre noise exposure | 12 | 0.08 | 0 | 0.43 | 0.4 | 0.52 | 0.73 | 0.6 | 0.68 | 1.05 | 0.86 | 1.26 | 1.65 | 1.8 | 2.49 | 2.69 | 3.52 | 3.88 | 4.7 | 5.84 | 6.26 |
| DMKO54.1e | Heterozygote | Control | Pre noise exposure | 12 | 0.21 | 1.01 | 0.16 | 0.81 | 0.33 | 0.14 | 0.64 | 0.89 | 0.96 | 1.17 | 1.39 | 1.83 | 1.89 | 2 | 2.14 | 2.2 | 2.29 | 2.72 | 2.59 | 2.98 |
| DMKO54.1c | Heterozygote | Noise | Pre noise exposure | 12 | 0.17 | 0.19 | -0.05 | 0.65 | 0.48 | 0.91 | 1.16 | 1.35 | 1.85 | 2.23 | 1.96 | 2.71 | 2 | 2.49 | 3.26 | 3.73 | 4.31 | 4.89 | 5.57 | 5.83 |
| DMKO55.1c | Heterozygote | Noise | Pre noise exposure | 12 | 0.29 | 0.01 | 0.16 | 0.67 | 0.63 | 1.01 | 1.17 | 1.26 | 1.77 | 1.9 | 1.86 | 2.13 | 1.7 | 2.47 | 3 | 3.59 | 3.62 | 4.33 | 5.26 | 5.27 |
| DMKO55.1i | Heterozygote | Control | Pre noise exposure | 12 | 0.05 | -0.13 | -0.34 | 0.38 | 0.23 | 0.56 | 0.85 | 0.94 | 0.87 | 1.3 | 1.19 | 1.38 | 1.39 | 1.78 | 2.37 | 2.49 | 2.73 | 2.36 | 2.86 | 2.72 |
| DMKO55.1g | Heterozygote | Noise | Pre noise exposure | 12 | 0.26 | -0.28 | 0.14 | 0.02 | 0.44 | 0.82 | 0.5 | 1.12 | 1.16 | 1.33 | 1.42 | 1.46 | 1.59 | 1.94 | 2.32 | 2.72 | 3 | 3.18 | 3.64 | 3.69 |
| DMKO57.1a | Heterozygote | Noise | Pre noise exposure | 12 | 0.41 | 0.16 | 0.3 | 0.26 | 0.34 | 0.98 | 1.1 | 1 | 1.49 | 1.72 | 1.77 | 1.89 | 2.23 | 2.53 | 3.09 | 3.56 | 3.71 | 4.09 | 3.96 | 4.4 |
| DMKO57.1e | Heterozygote | Control | Pre noise exposure | 12 | 0.28 | 0.15 | 0.24 | 0.12 | 0.39 | 0.58 | 0.38 | 0.63 | 1.02 | 0.65 | 1.05 | 0.93 | 1 | 1.1 | 0.87 | 1 | 1.36 | 1.73 | 2.35 | 2.03 |
| DMKO57.1c | Heterozygote | Control | Pre noise exposure | 12 | 0.28 | 0.12 | 0.5 | 0.21 | 0.51 | 0.6 | 0.57 | 0.88 | 1.36 | 1.63 | 1.77 | 1.71 | 1.88 | 1.9 | 2.67 | 2.93 | 3.12 | 3.29 | 3.81 | 3.86 |
| DMKO44.1a | Wildtype | Control | Pre noise exposure | 24 | | | 0 | 0.24 | 0.43 | 0.26 | 0.2 | 0.64 | 0.7 | 0.65 | 0.96 | 1.19 | 1.65 | 1.54 | 2.04 | 2.29 | 2.88 | 2.63 | 2.94 | 3.2 |
| DMKO44.1g | Wildtype | Noise | Pre noise exposure | 24 | | | 0.28 | -0.19 | 0.16 | 0.31 | 0.55 | 1.18 | 1.09 | 1.3 | 1.78 | 2.11 | 2.26 | 2.27 | 2.83 | 3.52 | 3.9 | 3.91 | 4.44 | 4.85 |
| DMKO44.1c | Wildtype | Noise | Pre noise exposure | 24 | | | 0.05 | -0.06 | 0.37 | 0.29 | 0.33 | 0.96 | 0.38 | 0.9 | 1.22 | 1.15 | 1.46 | 1.73 | 2.57 | 3.01 | 3.02 | 3.21 | 3.22 | 3.55 |
| DMKO54.1f | Wildtype | Control | Pre noise exposure | 24 | | | 0.25 | 0.2 | 0.2 | 0.27 | 0.3 | 0.56 | 0.65 | 0.89 | 1.21 | 1.2 | 1.51 | 1.72 | 1.84 | 2.22 | 2.53 | 3.17 | 3.49 | 4.01 |
| DMKO54.1d | Wildtype | Noise | Pre noise exposure | 24 | | | 0.09 | 0.19 | 0.17 | 0.33 | 0.21 | 0.51 | 0.93 | 0.79 | 1.38 | 1.22 | 1.6 | 2.13 | 2.71 | 3.06 | 2.99 | 3.6 | 3.77 | 4.37 |
| DMKO55.1b | Wildtype | Noise | Pre noise exposure | 24 | | | -0.11 | -0.03 | 0.33 | 0.38 | 0.07 | 0.91 | 1.04 | 1.02 | 1.09 | 1.41 | 1.58 | 2.18 | 2.51 | 2.97 | 3.26 | 3.57 | 3.87 | 4.1 |
| DMKO55.1a | Wildtype | Control | Pre noise exposure | 24 | | | 0.09 | 0.34 | 0.24 | 0.53 | 0.93 | 0.97 | 1.08 | 1.39 | 1.33 | 1.99 | 2.41 | 2.69 | 3.26 | 3.76 | 3.76 | 4.48 | | |
| CNTC20.1a | Wildtype | Control | Pre noise exposure | 24 | | | 0 | 0.27 | 0.14 | 0.29 | 0.54 | 0.62 | 0.68 | 1.28 | 0.98 | 1.29 | 1.67 | 2.02 | 2.31 | 2.65 | 3.38 | 3.29 | 4.34 | 4.11 |
| CNTC20.1f | Wildtype | Control | Pre noise exposure | 24 | | | 0.12 | 0.24 | 0.26 | 0.34 | 0.5 | 0.54 | 0.34 | 0.88 | 1.01 | 1.31 | 1.56 | 1.9 | 2.17 | 2.57 | 2.86 | 2.88 | 3.38 | 3.68 |
| CNTC20.1g | Wildtype | Control | Pre noise exposure | 24 | | | 0.01 | 0.16 | 0.25 | 0.18 | 0.52 | 0.77 | 0.65 | 1.21 | 1.51 | 1.29 | 1.95 | 1.63 | 2.41 | 3.21 | 3.03 | 3.35 | 3.58 | 3.99 |
| DMKO55.1h | Wildtype | Noise | Pre noise exposure | 24 | | | 0.31 | 0.26 | 0.24 | 0.43 | 0.56 | 0.63 | 0.86 | 1.07 | 1.59 | 1.67 | 2.21 | 2.65 | 2.68 | 3.44 | 4.07 | 4.34 | 4.84 | 5.41 |
| DMKO57.1b | Wildtype | Noise | Pre noise exposure | 24 | | | 0.22 | 0.24 | 0 | 0.27 | 0.37 | 0.26 | 0.6 | 0.42 | 0.95 | 0.96 | 1.55 | 1.3 | 1.74 | 1.82 | 2.27 | 2.48 | 2.19 | 2.76 |
| DMKO44.1b | Heterozygote | Control | Pre noise exposure | 24 | | | 0.17 | -0.02 | -0.05 | 0.29 | 0.15 | 0.38 | 0.75 | 0.55 | 0.91 | 1.44 | 1.76 | 1.87 | 1.88 | 1.95 | 2.12 | 1.76 | 2.26 | 2.4 |
| DMKO44.1d | Heterozygote | Control | Pre noise exposure | 24 | | | 0.13 | 0.12 | 0.13 | 0.41 | -0.12 | 0.38 | 0.38 | 0.37 | 0.73 | 0.7 | 1.11 | 1.29 | 1.62 | 1.75 | 1.47 | 1.85 | 1.96 | 2.2 |
| DMKO44.1f | Heterozygote | Noise | Pre noise exposure | 24 | | | 0.3 | 0.14 | -0.1 | 0.29 | 0.23 | 0.34 | 0.44 | 0.67 | 0.85 | 1.31 | 1.13 | 1.31 | 1.23 | 1.6 | 1.56 | 2.19 | 2.23 | 2.29 |
| DMKO44.1e | Heterozygote | Noise | Pre noise exposure | 24 | | | 0.46 | 0.26 | 0.36 | 0.36 | 0.62 | 0.37 | 0.55 | 0.87 | 1.24 | 1.08 | 1.26 | 1.81 | 1.86 | 2.42 | 2.67 | 2.94 | 3.45 | 3.54 |
| DMKO54.1e | Heterozygote | Control | Pre noise exposure | 24 | | | 0.18 | 0.3 | 0.19 | 0.46 | 0.06 | 0.16 | 0.41 | 0.72 | 1.18 | 1.22 | 1.35 | 1.54 | 1.83 | 2.04 | 2.4 | 2.63 | 2.65 | 2.94 |
| DMKO54.1c | Heterozygote | Noise | Pre noise exposure | 24 | | | -0.02 | 0.2 | 0.7 | 0.33 | 0.43 | 0.66 | 0.82 | 1.12 | 1.23 | 1.36 | 2.04 | 2.47 | 2.61 | 2.93 | 3.36 | 3.46 | 3.73 | 4.18 |
| DMKO55.1c | Heterozygote | Noise | Pre noise exposure | 24 | | | 0.37 | 0.16 | 0.1 | 0.41 | 0.46 | 0.42 | 0.85 | 0.91 | 1.07 | 1.64 | 1.7 | 1.99 | 1.94 | 2.1 | 2.58 | 2.78 | 3.17 | 3.4 |
| DMKO55.1i | Heterozygote | Control | Pre noise exposure | 24 | | | 0.11 | 0.21 | 0.12 | 0.11 | -0.01 | 0.1 | 0.67 | 0.36 | 0.7 | 1.04 | 1.19 | 1.46 | 1.87 | 1.9 | 2.1 | 2.12 | 2.33 | 2.72 |
| DMKO55.1g | Heterozygote | Noise | Pre noise exposure | 24 | | | 0.33 | 0.07 | 0.15 | 0.29 | 0.63 | 0.81 | 0.58 | 0.84 | 0.97 | 1.31 | 1.28 | 2.04 | 2.04 | 2.13 | 2.41 | 2.51 | 2.78 | 3.18 |
| DMKO57.1a | Heterozygote | Noise | Pre noise exposure | 24 | | | 0.14 | -0.1 | 0.06 | 0.19 | 0.66 | 0.38 | 0.99 | 0.86 | 1.29 | 1.2 | 1.63 | 1.79 | 2.1 | 1.96 | 2.07 | 1.86 | 2.2 | 2.71 |
| DMKO57.1e | Heterozygote | Control | Pre noise exposure | 24 | | | | | | | | | | | | | | | | | | | | |
| DMKO57.1c | Heterozygote | Control | Pre noise exposure | 24 | | | -0.08 | 0.11 | -0.07 | 0.19 | 0.14 | 0.27 | 0.85 | 0.61 | 1.06 | 1.39 | 1.51 | 1.74 | 1.66 | 1.87 | 2.02 | 2.17 | 2.24 | 2.23 |
| DMKO44.1a | Wildtype | Control | Noise +1 day | 12 | -0.11 | 0.17 | 0.24 | 0.59 | 0.68 | 0.8 | 0.54 | 1.07 | 1.07 | 1.63 | 1.48 | 1.82 | 2.36 | 2.8 | 2.94 | 3.25 | 3.26 | 3.17 | 3.71 | 3.82 |
| DMKO54.1f | Wildtype | Control | Noise +1 day | 12 | 0.07 | 0.06 | 0.04 | 0.95 | 0.8 | 0.32 | 0.96 | 1.11 | 1.74 | 1.61 | 1.84 | 2.25 | 2.39 | 2.31 | 2.75 | 3.47 | 3.86 | 3.81 | 4.82 | 5.29 |
| DMKO55.1a | Wildtype | Control | Noise +1 day | 12 | -0.05 | 0.2 | 0.6 | 0.24 | 0.65 | 0.78 | 1.21 | 1.36 | 1.55 | 2.03 | 1.95 | 2.31 | 2.37 | 2.87 | 3.65 | 4.49 | 4.86 | 5.71 | 6.77 | 7.28 |
| CNTC20.1a | Wildtype | Control | Noise +1 day | 12 | 0 | 0.3 | 0.18 | 0.62 | 1.15 | 1.09 | 1.03 | 1.55 | 1.88 | 2.02 | 2.59 | 2.66 | 2.68 | 4.06 | 5.1 | 5.82 | 5.84 | 6.43 | 7.15 | 7.6 |
| CNTC20.1f | Wildtype | Control | Noise +1 day | 12 | 0.26 | 0.11 | -0.05 | 0.33 | 0.23 | 0.88 | 0.68 | 1.23 | 1.45 | 1.32 | 1.83 | 2.04 | 2.46 | 2.64 | 4.84 | | | | | |
| CNTC20.1g | Wildtype | Control | Noise +1 day | 12 | 0.19 | 0.7 | 0.46 | 0.5 | 0.8 | 1.01 | 1.15 | 1.19 | 2.05 | 2.02 | 2.45 | 2.2 | 2.76 | 3.82 | 4.47 | 5.54 | 5.8 | 5.88 | | |
| DMKO44.1b | Heterozygote | Control | Noise +1 day | 12 | 0.13 | 0.21 | 0.16 | 0.1 | 0.61 | 0.47 | 0.87 | 0.96 | 1 | 1.22 | 1.45 | 1.68 | 1.55 | 1.99 | 2.33 | 2.73 | 3.54 | 3.91 | 4.34 | 4.82 |
| DMKO44.1d | Heterozygote | Control | Noise +1 day | 12 | 0.07 | 0.03 | 0.23 | 0.38 | 0.34 | 0.63 | 0.39 | 0.73 | 0.93 | 0.96 | 1.63 | 1.59 | 1.18 | 1.34 | 2.07 | 2.78 | 3.5 | 4.22 | 4.93 | 5.41 |
| DMKO54.1e | Heterozygote | Control | Noise +1 day | 12 | 0.13 | 0.09 | -0.01 | 0.34 | 0.22 | 0.48 | 0.74 | 1.08 | 1.16 | 1.03 | 1.12 | 1.65 | 1.7 | 1.91 | 2.3 | 2 | 2.61 | 3.1 | 3.51 | 3.83 |
| DMKO55.1i | Heterozygote | Control | Noise +1 day | 12 | -0.03 | 0 | 0.23 | 0.23 | 0.2 | 0.26 | 0.19 | 0.54 | 0.57 | 0.89 | 0.73 | 0.81 | 1.31 | 1.3 | 1.74 | 1.5 | 1.74 | 2.89 | 3.54 | 3.91 |
| DMKO57.1c | Heterozygote | Control | Noise +1 day | 12 | 0.19 | 0.25 | -0.13 | 0.55 | 0.4 | 0.72 | 1.12 | 1.3 | 1.66 | 1.94 | 2.26 | 2.78 | 2.72 | 3.31 | 4.02 | 4.58 | 5.15 | 5.31 | 5.72 | 6.14 |
| DMKO57.1e | Heterozygote | Control | Noise +1 day | 12 | 0.29 | 0.11 | 0.13 | 0.19 | 0.39 | 0.44 | 0.51 | 0.51 | 1.03 | 1.3 | 1.37 | 1.62 | 1.78 | 1.69 | 1.72 | 2.36 | 2.88 | 3.44 | 3.57 | 3.68 |
| DMKO44.1g | Wildtype | Noise | Noise +1 day | 12 | -0.38 | 0.15 | 0.19 | -0.2 | -0.02 | 0.41 | 0.53 | 0.77 | 0.79 | 1.28 | 1.74 | 2.03 | 2.53 | 3.06 | 2.99 | 3.3 | 3.86 | 4.3 | 4.36 | 4.1 |
| DMKO44.1c | Wildtype | Noise | Noise +1 day | 12 | . | . | . | . | -0.03 | 0.37 | 0.17 | 0.46 | 0.59 | 0.73 | 0.57 | 0.9 | 1.42 | 1.75 | 2.16 | 1.96 | 2.88 | 3.81 | 3.92 | 3.92 |
| DMKO54.1d | Wildtype | Noise | Noise +1 day | 12 | 0.05 | 0.06 | 0.19 | 0.19 | 0.24 | 0.05 | 0.24 | 0.42 | 0.57 | 0.49 | 0.82 | 1.62 | 2.04 | 2.41 | 2.82 | 3.3 | 4.38 | 4.51 | 5.4 | 5.17 |
| DMKO55.1b | Wildtype | Noise | Noise +1 day | 12 | 0.17 | -0.24 | 0.13 | 0.11 | 0.14 | 0.2 | 0.44 | 0.29 | 0.34 | 0.54 | 0.57 | 0.87 | 1.44 | 1.05 | 1.52 | 1.93 | 2.09 | 2.65 | 2.84 | 2.47 |
| DMKO55.1h | Wildtype | Noise | Noise +1 day | 12 | 0.27 | 0.29 | -0.24 | -0.17 | 0.27 | 0.23 | 0.22 | 0.53 | 0.88 | 0.71 | 1.07 | 1.87 | 2.35 | 2.7 | 3.33 | 3.86 | 4.3 | 5.14 | 5.59 | 5.76 |
| DMKO57.1b | Wildtype | Noise | Noise +1 day | 12 | . | . | . | 0.3 | 0.58 | 0.34 | 0.16 | 0.21 | -0.06 | 0.37 | 0.56 | 0.8 | 1.44 | 2.04 | 2.29 | 2.67 | 3.61 | 4.1 | 4.8 | 4.94 |
| DMKO44.1f | Heterozygote | Noise | Noise +1 day | 12 | 0 | 0 | 0.17 | 0.09 | 0 | 0.18 | -0.04 | 0.11 | 0.05 | -0.17 | 0.14 | 0.52 | 0.53 | 0.36 | 0.67 | 1.35 | 1.39 | 2.17 | 3.02 | 3.36 |
| DMKO44.1e | Heterozygote | Noise | Noise +1 day | 12 | 0.18 | -0.08 | 0.27 | 0.17 | 0.16 | 0.03 | 0.29 | -0.25 | -0.02 | 0.12 | 0.55 | 0.62 | 1.01 | 1.22 | 1.48 | 1.74 | 2 | 2.64 | 3.24 | 3.89 |
| DMKO54.1c | Heterozygote | Noise | Noise +1 day | 12 | 0.12 | 0.16 | -0.01 | 0.8 | 0.05 | 0.34 | 0.51 | 0.41 | 0.78 | 0.54 | 1.32 | 1.5 | 1.82 | 2.57 | 3.02 | 3.43 | 3.91 | 4.49 | 4.68 | 4.92 |
| DMKO55.1c | Heterozygote | Noise | Noise +1 day | 12 | 0.17 | 0.26 | 0.07 | 0.18 | 0.05 | 0.08 | 0.17 | 0.21 | 0.54 | 0.82 | 0.88 | 1.09 | 1.84 | 2.17 | 2.4 | 2.94 | 3.4 | 4.03 | 4.46 | 4.36 |
| DMKO55.1g | Heterozygote | Noise | Noise +1 day | 12 | -0.25 | -0.03 | 0.11 | -0.51 | 0.36 | 0.21 | 0.05 | 0.09 | 0.42 | 0.42 | 0.44 | 0.5 | 0.96 | 1.24 | 1.41 | 2.14 | 2.24 | 2.84 | 2.89 | 2.89 |
| DMKO57.1a | Heterozygote | Noise | Noise +1 day | 12 | . | . | . | . | -0.01 | 0.16 | -0.17 | 0.06 | -0.19 | -0.18 | 0.22 | 0.53 | 1.26 | 1.02 | 1.74 | 2.59 | 2.41 | 2.73 | 3.78 | 4.18 |
| DMKO44.1a | Wildtype | Control | Noise +1 day | 24 | | | -0.19 | -0.02 | 0.11 | 0.3 | 0.17 | 0.16 | 0.42 | 0.75 | 0.95 | 1.13 | 1.44 | 1.77 | 2.05 | 2.21 | 2.76 | 2.58 | 2.77 | 3.58 |
| DMKO54.1f | Wildtype | Control | Noise +1 day | 24 | | | 0.04 | 0.47 | -0.04 | 0.36 | 0.8 | 0.57 | 0.55 | 0.89 | 1.39 | 1.31 | 1.56 | 1.69 | 1.89 | 2.05 | 2.11 | 2.68 | 3.48 | 3.62 |
| DMKO55.1a | Wildtype | Control | Noise +1 day | 24 | | | 0.36 | 0.16 | 0.53 | 0.19 | 0.4 | 0.5 | 0.78 | 1.07 | 1.57 | 1.82 | 1.96 | 2.63 | 2.72 | 2.82 | 3.64 | 3.96 | 4.41 | 4.85 |
| CNTC20.1a | Wildtype | Control | Noise +1 day | 24 | | | 0.1 | 0.2 | 0.36 | 0.29 | 0.64 | 0.7 | 1.29 | 0.99 | 1.46 | 2.4 | 2.51 | 2.9 | 3.66 | 3.57 | 4.29 | 4.67 | 5.49 | 5.75 |
| CNTC20.1f | Wildtype | Control | Noise +1 day | 24 | | | 0.63 | 0.31 | 0.24 | 0.64 | 0.57 | 0.31 | 0.73 | 0.92 | 1.13 | 1.67 | 2.3 | 2.09 | 3.86 | | | | | |
| CNTC20.1g | Wildtype | Control | Noise +1 day | 24 | | | 0.26 | 0 | 0.2 | 0.5 | 0.57 | 0.99 | 0.92 | 1.25 | 1.63 | 1.89 | 2.13 | 3.21 | 3.21 | 3.39 | 4.19 | 4.25 | | |
| DMKO44.1b | Heterozygote | Control | Noise +1 day | 24 | | | 0.2 | -0.18 | -0.28 | 0.3 | 0.26 | 0.27 | 0.34 | 0.68 | 1.12 | 1.21 | 1.16 | 1.53 | 1.68 | 1.92 | 2.24 | 2.4 | 2.15 | 2.25 |
| DMKO44.1d | Heterozygote | Control | Noise +1 day | 24 | | | 0.31 | -0.13 | 0.41 | -0.01 | 0.33 | 0.29 | 0.29 | 0.47 | 0.65 | 0.9 | 1.13 | 1.49 | 1.53 | 1.72 | 1.61 | 1.73 | 2.17 | 2.84 |
| DMKO54.1e | Heterozygote | Control | Noise +1 day | 24 | | | 0.06 | 0.3 | 0 | 0.34 | 0.33 | 0.33 | 0.28 | 0.18 | 0.83 | 1.07 | 1.44 | 1.85 | 1.78 | 2.29 | 2.75 | 3.05 | 3.29 | 3.37 |
| DMKO55.1i | Heterozygote | Control | Noise +1 day | 24 | | | 0.2 | 0.28 | 0.1 | 0.27 | 0.11 | 0.23 | 0.13 | 0.05 | 0.24 | 0.31 | 0.57 | 0.65 | 0.8 | 0.91 | 1.38 | 1.03 | 1.19 | 1.49 |
| DMKO57.1c | Heterozygote | Control | Noise +1 day | 24 | | | 0.36 | 0.18 | 0.31 | 0.14 | 0.22 | 0.24 | 0.53 | 0.84 | 1.04 | 1.55 | 1.63 | 1.78 | 1.88 | 2.38 | 2.52 | 3 | 2.79 | 3.08 |
| DMKO57.1e | Heterozygote | Control | Noise +1 day | 24 | | | 0.3 | 0.16 | 0.08 | 0.23 | 0.17 | 0.06 | 0.2 | 0.49 | 0.53 | 0.54 | 0.8 | 1.33 | 1.44 | 1.36 | 0.99 | 1.57 | 1.54 | 1.83 |
| DMKO44.1g | Wildtype | Noise | Noise +1 day | 24 | | | | | | | | | | | | | | | | | | | | |
| DMKO44.1c | Wildtype | Noise | Noise +1 day | 24 | | | | | 0.04 | 0.22 | 0.21 | 0.66 | 0.01 | 0.13 | 0.05 | 0.08 | 0.13 | 0.05 | 0.14 | 0.39 | 0.27 | 0.77 | 0.77 | 1.01 |
| DMKO54.1d | Wildtype | Noise | Noise +1 day | 24 | | | | | | | | | | | | | | | | | | | | |
| DMKO55.1b | Wildtype | Noise | Noise +1 day | 24 | | | | | | | | | | | | | | | | | | | | |
| DMKO55.1h | Wildtype | Noise | Noise +1 day | 24 | | | | | | | | | | | | | | | | | | | | |
| DMKO57.1b | Wildtype | Noise | Noise +1 day | 24 | | | | | | | | | | | | | | | | | | | | |
| DMKO44.1f | Heterozygote | Noise | Noise +1 day | 24 | | | | | | | | | | | | | | | | | | | | |
| DMKO44.1e | Heterozygote | Noise | Noise +1 day | 24 | | | | | | | | | | | | | | | | | | | | |
| DMKO54.1c | Heterozygote | Noise | Noise +1 day | 24 | | | | | | | | | | | | | | | | | | | | |
| DMKO55.1c | Heterozygote | Noise | Noise +1 day | 24 | | | | | | | | | | | | | | | | | | | | |
| DMKO55.1g | Heterozygote | Noise | Noise +1 day | 24 | | | | | | | | | | | | | | | | | | | | |
| DMKO57.1a | Heterozygote | Noise | Noise +1 day | 24 | | | | | | | | | | | | | | | | | | | | |
| DMKO44.1a | Wildtype | Control | Noise +3 days | 12 | -0.01 | 0.27 | 0.5 | 0.54 | 0.47 | 0.66 | 0.73 | 1.25 | 1.23 | 1.66 | 2.05 | 2.39 | 2.52 | 3.03 | 3.98 | 4.14 | 3.83 | 4.17 | 4.54 | 4.62 |
| DMKO54.1f | Wildtype | Control | Noise +3 days | 12 | 0.12 | 0.5 | 0.2 | 0.56 | 0.58 | 0.65 | 1.13 | 0.97 | 1.44 | 1.94 | 2.03 | 2.24 | 2.03 | 2.08 | 3.07 | 3.6 | 3.87 | 3.92 | 5.02 | 5.41 |
| CNTC20.1a | Wildtype | Control | Noise +3 days | 12 | 0 | 0.09 | 0.53 | 0.64 | 1.01 | 0.94 | 1.54 | 1.96 | 2.19 | 2.39 | 2.51 | 2.75 | 3.08 | 4.11 | 5.45 | 5.84 | 5.88 | 6.37 | 7.82 | 7.4 |
| CNTC20.1f | Wildtype | Control | Noise +3 days | 12 | 0.46 | 0.37 | -0.11 | 0.61 | 0.72 | 0.82 | 0.69 | 0.98 | 1.67 | 1.69 | 1.68 | 2.11 | 2.62 | 3.01 | 4.06 | 4.23 | 4.56 | 5.14 | 5.47 | 5.74 |
| CNTC20.1g | Wildtype | Control | Noise +3 days | 12 | 0.45 | 0.13 | 0.26 | 0.39 | 0.68 | 1.05 | 0.74 | 1.46 | 1.61 | 1.72 | 2.23 | 2.19 | 2.27 | 3.28 | 4.56 | 5.69 | 6.09 | 5.84 | 5.72 | 5.75 |
| DMKO55.1a | Wildtype | Control | Noise +3 days | 12 | 0.19 | 0.24 | 0.33 | 0.16 | 0.57 | 0.66 | 0.9 | 1.03 | 1.29 | 1.7 | 1.74 | 2.04 | 2.33 | 2.3 | 3.46 | 3.79 | 3.88 | 4.25 | 4.69 | 5.09 |
| DMKO44.1b | Heterozygote | Control | Noise +3 days | 12 | 0.06 | -0.02 | 0.07 | 0.06 | 0.19 | 0.27 | 0.65 | 0.55 | 0.8 | 1.16 | 0.86 | 1.68 | 1.25 | 1.69 | 1.92 | 2.65 | 2.97 | 3.64 | 4.08 | 4.58 |
| DMKO44.1d | Heterozygote | Control | Noise +3 days | 12 | -0.08 | 0.17 | 0.09 | 0.29 | 0.4 | 0.51 | 0.72 | 0.61 | 0.88 | 1.19 | 1.22 | 1.19 | 1.64 | 1.77 | 2.19 | 2.41 | 2.87 | 3.35 | 4.2 | 4.34 |
| DMKO54.1e | Heterozygote | Control | Noise +3 days | 12 | 0 | 0.11 | 0.11 | -0.16 | 0.41 | 0.39 | 0.91 | 1.03 | 1.13 | 1.33 | 1.61 | 1.68 | 2.24 | 2.04 | 2.25 | 2.31 | 2.56 | 2.98 | 3.23 | 3.47 |
| DMKO55.1i | Heterozygote | Control | Noise +3 days | 12 | 0.19 | 0.13 | 0.19 | 0.28 | 0.2 | 0.22 | 0.52 | 0.31 | 0.4 | 1.12 | 0.9 | 0.95 | 0.75 | 0.81 | 1.28 | 1.45 | 1.43 | 1.87 | 2.28 | 2.66 |
| DMKO57.1c | Heterozygote | Control | Noise +3 days | 12 | 0.12 | 0.09 | 0.26 | 0.45 | 0.48 | 0.53 | 0.57 | 0.99 | 1.26 | 1.35 | 1.44 | 1.95 | 1.75 | 2 | 2.44 | 2.77 | 3.21 | 2.99 | 3.3 | 3.57 |
| DMKO57.1e | Heterozygote | Control | Noise +3 days | 12 | 0.11 | 0.22 | 0.24 | 0.27 | 0.37 | 0.66 | 0.74 | 0.99 | 0.89 | 1.45 | 1.62 | 1.65 | 1.54 | 1.47 | 2.18 | 2.08 | 2.56 | 2.88 | 3.17 | 2.86 |
| DMKO44.1g | Wildtype | Noise | Noise +3 days | 12 | 0.3 | -0.03 | 0.15 | -0.05 | 0.52 | 1.28 | 1.11 | 1.67 | 1.69 | 2.23 | 2.6 | 2.75 | 2.89 | 3.18 | 3.49 | 3.85 | 3.93 | 4.43 | 4.36 | 4.22 |
| DMKO44.1c | Wildtype | Noise | Noise +3 days | 12 | 0.26 | 0.18 | 0.32 | 0.28 | 0.49 | 0.56 | 0.77 | 0.92 | 1.12 | 1.38 | 1.95 | 1.88 | 1.97 | 2.38 | 2.55 | 2.82 | 2.85 | 3.57 | 4.36 | 4.13 |
| DMKO54.1d | Wildtype | Noise | Noise +3 days | 12 | 0.39 | 0.34 | 0.3 | 0.36 | 0.26 | 0.57 | 0.82 | 1.3 | 1.49 | 1.87 | 1.93 | 2.65 | 2.31 | 2.48 | 2.57 | 3.02 | 3.34 | 3.59 | 4.4 | 4.36 |

| | | | | | | | | | | | | | | | | | | | | | | | | |
|---|---|---|---|---|---|---|---|---|---|---|---|---|---|---|---|---|---|---|---|---|---|---|---|---|
| DMKO55.1b | Wildtype | Noise | Noise +3 days | 12 | 0.06 | 0.01 | 0.19 | 0.63 | 0.5 | 0.86 | 1.03 | 1.18 | 1.41 | 1.3 | 1.73 | 2.01 | 2.03 | 2.46 | 2.7 | 3 | 3.16 | 3.02 | 3.3 | 2.84 |
| DMKO55.1h | Wildtype | Noise | Noise +3 days | 12 | -0.07 | 0.15 | 0.33 | 0.66 | 0.45 | 0.57 | 1.02 | 1.17 | 1.8 | 2.46 | 2.77 | 2.35 | 3.12 | 3.45 | 3.48 | 4.31 | 4.45 | 5.09 | 4.66 | 5.04 |
| DMKO57.1b | Wildtype | Noise | Noise +3 days | 12 | -0.12 | 0.18 | 0.34 | -0.12 | 0.26 | 0.34 | 0.19 | 0.31 | 0.64 | 1.13 | 1.35 | 1.76 | 2.11 | 2.62 | 2.72 | 3.21 | 3.55 | 3.79 | 4.42 | 4.83 |
| DMKO44.1f | Heterozygote | Noise | Noise +3 days | 12 | 0 | 0 | 0.07 | 0 | 0.14 | 0.22 | 0.4 | 0.41 | 0.86 | 0.95 | 1.14 | 2.01 | 1.64 | 2.57 | 2.57 | 2.73 | 3.24 | 3.52 | 4.06 | 3.47 |
| DMKO44.1e | Heterozygote | Noise | Noise +3 days | 12 | 0.01 | -0.27 | 0.11 | 0.23 | 0.07 | 0.39 | 0.39 | 0.34 | 0.87 | 0.82 | 1.16 | 1.21 | 1.22 | 1.34 | 1.54 | 1.84 | 1.88 | 2.87 | 3.78 | 4.15 |
| DMKO54.1c | Heterozygote | Noise | Noise +3 days | 12 | 0.71 | 0.15 | 0.31 | 0.19 | 0.27 | 0.21 | 0.46 | 0.48 | 0.48 | 0.73 | 0.73 | 0.84 | 1.14 | 1.58 | 1.83 | 2.16 | 2.27 | 2.5 | 2.74 | 2.37 |
| DMKO55.1c | Heterozygote | Noise | Noise +3 days | 12 | 0.24 | 0.32 | 0.41 | 0.51 | 0.38 | 0.59 | 1.25 | 1.25 | 1.07 | 1.16 | 1.63 | 1.78 | 2.21 | 2.37 | 2.24 | 2.27 | 2.46 | 3.14 | 2.92 | 2.91 |
| DMKO55.1g | Heterozygote | Noise | Noise +3 days | 12 | -0.14 | 0.18 | 0.54 | 0.51 | 0.46 | 0.33 | 0.84 | 0.58 | 0.69 | 0.75 | 0.6 | 0.98 | 1.31 | 1.69 | 1.61 | 1.76 | 1.77 | 2.24 | 2.16 | 2.31 |
| DMKO57.1a | Heterozygote | Noise | Noise +3 days | 12 | 0.11 | 0.16 | 0.21 | 0.41 | 0.32 | 0.56 | 0.58 | 0.66 | 1.19 | 0.99 | 1.15 | 1.28 | 1.36 | 1.56 | 1.48 | 2.25 | 2.56 | 2.93 | 3.26 | 3.66 |
| DMKO44.1a | Wildtype | Control | Noise +3 days | 24 | | | 0.14 | 0 | 0.23 | 0.21 | 0.19 | 0.4 | 0.64 | 0.94 | 1.27 | 1.33 | 2.02 | 1.93 | 2.5 | 2.36 | 2.54 | 3.09 | 3.31 | 3.58 |
| DMKO54.1f | Wildtype | Control | Noise +3 days | 24 | | | 0.23 | 0.2 | 0.26 | 0.05 | 0.22 | 0.42 | 0.57 | 1.1 | 0.84 | 1.04 | 1.18 | 1.24 | 1.24 | 1.19 | 1.54 | 1.4 | 2.18 | 2.65 |
| CNTC20.1a | Wildtype | Control | Noise +3 days | 24 | | | 0.25 | 0.08 | 0.54 | 0.54 | 0.36 | 0.82 | 1.28 | 1.42 | 1.8 | 2.16 | 2.47 | 2.82 | 3.34 | 3.65 | 4.36 | 4.56 | 4.77 | 5.52 |
| CNTC20.1f | Wildtype | Control | Noise +3 days | 24 | | | 0.35 | 0.12 | 0.23 | 0.19 | 0.59 | 0.64 | 0.89 | 0.96 | 1.26 | 1.48 | 1.71 | 2.13 | 2.67 | 2.96 | 3.24 | 3.47 | 3.58 | 4.18 |
| CNTC20.1g | Wildtype | Control | Noise +3 days | 24 | | | 0.38 | 0.34 | 0.29 | 0.19 | 0.3 | 0.85 | 1.03 | 1.2 | 1.23 | 1.26 | 2.13 | 1.81 | 2.84 | 3.38 | 3.8 | 4.02 | 3.99 | 4.88 |
| DMKO55.1a | Wildtype | Control | Noise +3 days | 24 | | | 0.44 | 0.17 | 0.37 | 0.47 | 0.41 | 0.72 | 0.82 | 0.77 | 1.24 | 1.58 | 1.75 | 2.15 | 2.37 | 2.71 | 2.75 | 3.18 | 3.94 | 3.74 |
| DMKO44.1b | Heterozygote | Control | Noise +3 days | 24 | | | 0 | 0.06 | 0.09 | 0.42 | 0.23 | 0.62 | 0.23 | 0.45 | 0.46 | 0.91 | 1.12 | 1.15 | 1.48 | 1.04 | 1.32 | 1.64 | 1.58 | 2.07 |
| DMKO44.1d | Heterozygote | Control | Noise +3 days | 24 | | | 0.15 | 0.29 | 0.23 | 0.24 | 0.34 | 0.35 | 0.42 | 0.44 | 0.9 | 1.15 | 1.03 | 1.08 | 1.59 | 1.61 | 2 | 1.76 | 1.82 | 1.89 |
| DMKO54.1e | Heterozygote | Control | Noise +3 days | 24 | | | -0.04 | -0.07 | 0.15 | 0.16 | 0.15 | 0.51 | 0.83 | 0.9 | 1.22 | 1.49 | 2.01 | 1.95 | 2.35 | 2.47 | 2.98 | 2.64 | 3.1 | 2.84 |
| DMKO55.1i | Heterozygote | Control | Noise +3 days | 24 | | | 0.22 | 0.21 | 0.27 | 0.62 | 0.19 | 0.55 | 0.36 | 0.01 | 0.77 | 0.57 | 0.63 | 0.88 | 1.09 | 0.83 | 1.17 | 0.93 | 1.29 | 1.53 |
| DMKO57.1c | Heterozygote | Control | Noise +3 days | 24 | | | 0.2 | 0.22 | 0.09 | 0.14 | 0.2 | 0.44 | 0.5 | 0.47 | 0.67 | 0.86 | 1.48 | 1.33 | 1.76 | 1.84 | 1.81 | 1.96 | 1.99 | 2.05 |
| DMKO57.1e | Heterozygote | Control | Noise +3 days | 24 | | | 0.12 | -0.07 | -0.15 | 0.28 | 0.36 | 0.45 | 0.72 | 0.63 | 0.69 | 0.8 | 1.15 | 1.43 | 1.47 | 1.66 | 1.64 | 1.65 | 1.94 | 1.93 |
| DMKO44.1g | Wildtype | Noise | Noise +3 days | 24 | | | 0.27 | 0.35 | 0.32 | 0.36 | 0.34 | 0.25 | 0.07 | -0.05 | 0.2 | 0.18 | 0.41 | 0.49 | 0.79 | 0.66 | 0.94 | 0.85 | 1.06 | 1.29 |
| DMKO44.1c | Wildtype | Noise | Noise +3 days | 24 | | | 0.2 | 0.03 | 0 | 0 | -0.05 | 0.19 | -0.01 | 0.09 | 0.09 | 0.28 | 0.21 | 0.27 | -0.01 | 0.44 | 0.64 | 0.69 | 0.9 | 1.05 |
| DMKO54.1d | Wildtype | Noise | Noise +3 days | 24 | | | | | | | | | | | | | | | | | | | | |
| DMKO55.1b | Wildtype | Noise | Noise +3 days | 24 | | | 0.28 | 0.3 | 0.16 | 0.25 | 0.11 | 0.16 | 0.38 | 0.21 | 0.57 | 0.68 | 0.85 | 0.66 | 1.01 | 1.4 | 1.6 | 1.77 | 2.28 | 1.97 |
| DMKO55.1h | Wildtype | Noise | Noise +3 days | 24 | | | | | | | | | | | | | | | | | | | | |
| DMKO57.1b | Wildtype | Noise | Noise +3 days | 24 | | | | | | | | | | | | | | | | | | | | |
| DMKO44.1f | Heterozygote | Noise | Noise +3 days | 24 | | | | | | | | | | | | | | | | | | | | |
| DMKO44.1e | Heterozygote | Noise | Noise +3 days | 24 | | | -0.02 | 0.16 | 0 | 0.1 | 0.06 | 0.26 | 0.21 | 0.33 | -0.01 | 0.05 | 0.15 | 0.44 | 0.62 | 0.66 | 0.83 | 0.44 | 0.72 | 1.02 |
| DMKO54.1c | Heterozygote | Noise | Noise +3 days | 24 | | | | | | | | | | | | | | | | | | | | |
| DMKO55.1c | Heterozygote | Noise | Noise +3 days | 24 | | | | | | | | | | | | | | | | | | | | |
| DMKO55.1g | Heterozygote | Noise | Noise +3 days | 24 | | | | | | | | | | | | | | | | | | | | |
| DMKO57.1a | Heterozygote | Noise | Noise +3 days | 24 | | | | | | | | | | | | | | | | | | | | |
| DMKO44.1a | Wildtype | Control | Noise +7 days | 12 | -0.1 | 0.06 | 0.26 | 0.57 | 0.52 | 0.81 | 1.09 | 0.78 | 0.88 | 1.83 | 1.53 | 2.12 | 2.66 | 3.16 | 3.4 | 3.81 | 3.76 | 3.71 | 4.06 | 4.24 |
| DMKO54.1f | Wildtype | Control | Noise +7 days | 12 | 0.26 | 0.41 | 0.41 | 0.78 | 0.63 | 0.43 | 1.35 | 0.84 | 1.59 | 1.62 | 2.25 | 2.19 | 2.31 | 2.22 | 3.46 | 4.23 | 4.67 | 5.06 | 5.78 | 5.72 |
| CNTC20.1a | Wildtype | Control | Noise +7 days | 12 | 0.16 | 0.02 | 0.44 | 0.39 | 0.72 | 1.14 | 1.8 | 2.3 | 1.68 | 2.12 | 2.55 | 2.75 | 3 | 4.21 | 5.16 | 6.13 | 6.16 | 5.79 | 6.8 | 7.42 |
| CNTC20.1f | Wildtype | Control | Noise +7 days | 12 | 0.07 | 0.18 | 0.28 | 0.47 | 0.46 | 0.81 | 0.76 | 0.98 | 1.19 | 1.48 | 2.01 | 1.85 | 2.54 | 2.94 | 4.03 | 4.61 | 5.06 | 5.13 | 5.36 | 5.83 |
| CNTC20.1g | Wildtype | Control | Noise +7 days | 12 | -0.06 | 0.12 | 0.31 | 0.58 | 0.42 | 0.82 | 0.72 | 1.14 | 1.42 | 1.83 | 2.24 | 2.27 | 2.19 | 3.02 | 4.04 | 4.94 | 5.4 | 5.41 | 5.71 | 6.28 |
| DMKO55.1a | Wildtype | Control | Noise +7 days | 12 | 0.11 | 0.12 | 0.29 | 0.37 | 0.71 | 0.77 | 0.87 | 1.05 | 0.97 | 1.56 | 1.77 | 1.72 | 2.45 | 2.58 | 3.43 | 3.81 | 3.75 | 4.23 | 4.87 | 5.07 |
| DMKO44.1b | Heterozygote | Control | Noise +7 days | 12 | 0.11 | 0.04 | 0.39 | -0.08 | 0.4 | 0.49 | 0.38 | 0.73 | 0.91 | 0.99 | 1.01 | 1.31 | 1.48 | 1.78 | 1.8 | 2.35 | 2.92 | 3.15 | 3.54 | 3.42 |
| DMKO44.1d | Heterozygote | Control | Noise +7 days | 12 | 0.12 | 0.24 | 0.44 | 0.42 | 0.46 | 0.51 | 0.67 | 0.85 | 0.61 | 0.66 | 0.96 | 1.34 | 1.32 | 1.8 | 2.3 | 2.23 | 2.53 | 3.17 | 3.17 | 3.41 |
| DMKO54.1e | Heterozygote | Control | Noise +7 days | 12 | 0.12 | 0.44 | 0.3 | 0.27 | 0.45 | 0.25 | 0.45 | 0.7 | 1 | 1.43 | 1.35 | 1.76 | 1.51 | 1.82 | 1.81 | 2.4 | 2.67 | 3.1 | 3.14 | 3.11 |
| DMKO55.1i | Heterozygote | Control | Noise +7 days | 12 | -0.17 | 0.12 | 0.16 | 0.5 | 0.44 | 0.57 | 0.79 | 0.86 | 1.2 | 1.48 | 1.4 | 1.44 | 1.89 | 1.48 | 1.99 | 2.31 | 2.3 | 3.35 | 3.55 | 3.87 |
| DMKO57.1c | Heterozygote | Control | Noise +7 days | 12 | 0.21 | 0.19 | -0.02 | 0.27 | 0.36 | 0.53 | 0.87 | 0.9 | 0.87 | 1.04 | 1.18 | 1.73 | 1.5 | 2.21 | 2.63 | 3.1 | 3.18 | 3.45 | 3.44 | 4.07 |
| DMKO57.1e | Heterozygote | Control | Noise +7 days | 12 | 0.27 | -0.12 | 0.3 | 0.21 | 0.49 | 0.36 | 0.36 | 0.59 | 0.44 | 0.83 | 0.52 | 0.8 | 0.62 | 1 | 1.04 | 0.92 | 1.45 | 1.59 | 2.38 | 2.48 |
| DMKO44.1g | Wildtype | Noise | Noise +7 days | 12 | -0.06 | 0.39 | 0.29 | 0.51 | 0.19 | 1.16 | 1.06 | 1.35 | 1.81 | 2.14 | 2.3 | 2.66 | 2.98 | 3.88 | 4 | 4.26 | 4.66 | 4.84 | 4.49 | 5.04 |
| DMKO44.1c | Wildtype | Noise | Noise +7 days | 12 | 0.17 | -0.07 | 0.09 | 0.55 | 0.32 | 0.58 | 0.94 | 0.87 | 0.92 | 1.55 | 1.68 | 2.02 | 2.33 | 3.15 | 3.55 | 3.7 | 4.13 | 4.18 | 4.36 | 4.68 |
| DMKO54.1d | Wildtype | Noise | Noise +7 days | 12 | 0 | 0.09 | -0.18 | 0.22 | 0.33 | 0.54 | 0.72 | 0.93 | 1.38 | 1.19 | 1.44 | 1.8 | 2.26 | 2.32 | 2.37 | 2.6 | 2.66 | 2.97 | 3.46 | 3.47 |
| DMKO55.1b | Wildtype | Noise | Noise +7 days | 12 | 0.07 | 0.2 | 0.38 | 0.33 | 0.59 | 0.62 | 0.96 | 0.97 | 1.24 | 1.6 | 1.67 | 2.02 | 1.86 | 2.51 | 2.87 | 3.31 | 3.22 | 3.71 | 3.81 | 3.22 |
| DMKO55.1h | Wildtype | Noise | Noise +7 days | 12 | 0.47 | 0.05 | 0.22 | 0.44 | 0.4 | 0.68 | 1.15 | 1.59 | 1.49 | 2.09 | 2.59 | 2.75 | 2.57 | 3.2 | 3.67 | 4.15 | 4.69 | 4.78 | 4.9 | 4.68 |
| DMKO57.1b | Wildtype | Noise | Noise +7 days | 12 | 0.06 | 0.13 | 0.11 | 0.12 | 0.55 | 0.53 | 0.88 | 0.93 | 1.83 | 1.52 | 2.15 | 2.41 | 2.69 | 2.92 | 3.51 | 3.78 | 4.28 | 4.74 | 4.74 | 4.7 |
| DMKO44.1f | Heterozygote | Noise | Noise +7 days | 12 | 0.11 | 0.08 | 0.19 | 0.26 | 0.54 | 0.5 | 0.45 | 0.89 | 0.76 | 0.88 | 1.3 | 1.26 | 1.83 | 2.09 | 2.39 | 2.7 | 3.44 | 4 | 4.67 | 4.15 |
| DMKO44.1e | Heterozygote | Noise | Noise +7 days | 12 | 0.06 | 0.19 | -0.03 | -0.01 | 0.48 | 0.6 | 0.86 | 0.87 | 0.84 | 1.36 | 1.64 | 1.61 | 1.5 | 2.09 | 2.73 | 3.28 | 3.61 | 3.68 | 3.81 | 3.98 |
| DMKO54.1c | Heterozygote | Noise | Noise +7 days | 12 | 0 | 0.22 | 0 | -0.27 | 0.54 | 0.35 | 1.05 | 0.75 | 0.88 | 1.53 | 1.36 | 1.66 | 1.99 | 1.97 | 2.37 | 1.94 | 2.46 | 2.64 | 2.97 | 2.45 |
| DMKO55.1c | Heterozygote | Noise | Noise +7 days | 12 | 0.17 | -0.07 | 0.26 | 0.7 | 0.48 | 0.79 | 0.69 | 0.83 | 1.29 | 1.44 | 1.12 | 1.39 | 2.03 | 2.45 | 2.62 | 2.72 | 2.68 | 3.15 | 3.12 | 3.46 |
| DMKO55.1g | Heterozygote | Noise | Noise +7 days | 12 | 0.13 | 0.23 | 0.23 | 0.02 | 0.3 | 0.61 | 0.46 | 0.79 | 0.99 | 1.16 | 1.64 | 1.53 | 1.96 | 1.95 | 2.08 | 2.06 | 2.44 | 2.72 | 2.95 | 2.62 |
| DMKO57.1a | Heterozygote | Noise | Noise +7 days | 12 | -0.01 | 0.4 | 0.3 | 0.46 | 0.7 | 0.32 | 0.49 | 0.47 | 0.61 | 0.61 | 0.59 | 1.1 | 0.79 | 1.62 | 1.46 | 1.19 | 1.5 | 1.84 | 1.92 | 1.85 |
| DMKO44.1a | Wildtype | Control | Noise +7 days | 24 | | | 0 | 0.28 | 0.15 | 0.4 | 0.2 | 0.38 | 0.58 | 0.48 | 0.64 | 0.9 | 1.12 | 1.2 | 1.69 | 1.74 | 1.85 | 2.36 | 2.68 | 2.97 |
| DMKO54.1f | Wildtype | Control | Noise +7 days | 24 | | | -0.12 | 0.43 | -0.03 | 0.11 | 0.35 | 0.62 | 0.68 | 1.17 | 1.15 | 1.44 | 1.53 | 2.05 | 1.8 | 2.27 | 3.17 | 2.98 | 3.47 | 4.18 |
| CNTC20.1a | Wildtype | Control | Noise +7 days | 24 | 0.24 | 0.03 | 0 | 0.08 | 0.06 | 0.15 | 0.73 | 0.75 | 1.14 | 1.5 | 1.82 | 2.25 | 1.99 | 1.97 | 3.1 | 3.68 | 4.01 | 4.42 | 4.7 | 5.92 |
| CNTC20.1f | Wildtype | Control | Noise +7 days | 24 | | | 0.09 | 0.08 | -0.08 | 0.3 | 0.52 | 0.67 | 0.82 | 0.85 | 1.14 | 1.29 | 1.59 | 2.32 | 2.35 | 2.68 | 2.94 | 3.57 | 3.75 | 4.16 |
| CNTC20.1g | Wildtype | Control | Noise +7 days | 24 | | | 0 | 0 | 0.38 | 0.46 | 0.54 | 0.81 | 0.49 | 1.11 | 1.44 | 1.48 | 1.77 | 1.77 | 2.11 | 2.85 | 3.19 | 3.36 | 3.31 | 3.99 |
| DMKO55.1a | Wildtype | Control | Noise +7 days | 24 | | | 0.61 | 0.11 | 0.55 | 0.18 | 0.48 | 0.76 | 0.91 | 1.18 | 1.58 | 1.62 | 1.95 | 2.34 | 2.55 | 3.18 | 3.4 | 3.36 | 3.74 | 4.08 |
| DMKO44.1b | Heterozygote | Control | Noise +7 days | 24 | | | 0.17 | 0.38 | 0.58 | 0.1 | 0.58 | 0.43 | 0.38 | 0.42 | 0.75 | 0.72 | 0.73 | 1.17 | 1.23 | 1.28 | 1.4 | 1.68 | 1.64 | 1.87 |
| DMKO44.1d | Heterozygote | Control | Noise +7 days | 24 | | | 0.13 | 0.06 | 0.27 | 0.05 | 0.15 | 0.4 | 0.44 | 0.61 | 0.92 | 1 | 0.85 | 0.73 | 1.39 | 1.09 | 1.08 | 1.38 | 1.27 | 1.24 |
| DMKO54.1e | Heterozygote | Control | Noise +7 days | 24 | | | 0.06 | 0.27 | 0.07 | 0.03 | 0.35 | 0.28 | 0.46 | 0.71 | 0.79 | 1.19 | 1.48 | 1.69 | 1.85 | 2.13 | 2.5 | 2.31 | 2.81 | 3 |
| DMKO55.1i | Heterozygote | Control | Noise +7 days | 24 | | | 0.09 | 0.31 | 0.11 | 0.28 | 0.57 | 0.49 | 0.68 | 0.96 | 1.04 | 1.32 | 1.48 | 2.04 | 2.31 | 2.81 | 2.73 | 2.58 | 3.06 | 3.09 |
| DMKO57.1c | Heterozygote | Control | Noise +7 days | 24 | | | 0.12 | 0.42 | 0.03 | 0.55 | 0.27 | 0.67 | 0.26 | 0.57 | 0.77 | 1.15 | 1.36 | 1.46 | 1.97 | 1.58 | 1.51 | 1.74 | 1.89 | 1.79 |
| DMKO57.1e | Heterozygote | Control | Noise +7 days | 24 | | | | | | | | | | | | | | | | | | | | |
| DMKO44.1g | Wildtype | Noise | Noise +7 days | 24 | | | 0.23 | -0.19 | 0.25 | 0.28 | 0.22 | 0.21 | 0.5 | 0.65 | 0.88 | 1.08 | 1.2 | 1.45 | 1.48 | 2.11 | 1.58 | 2.14 | 2.26 | 3 |
| DMKO44.1c | Wildtype | Noise | Noise +7 days | 24 | | | 0.09 | 0.29 | 0.03 | 0.24 | 0.34 | 0.38 | 0.41 | 0.6 | 0.79 | 0.69 | 1.08 | 1.14 | 1.42 | 1.69 | 2.06 | 1.95 | 2.12 | 2.54 |
| DMKO54.1d | Wildtype | Noise | Noise +7 days | 24 | | | 0.27 | 0.33 | 0.17 | 0 | 0.21 | 0.09 | 0.09 | -0.05 | 0.27 | -0.08 | 0.01 | 0.24 | -0.12 | 0.35 | 0.45 | 0.5 | 0.91 | 1.53 |
| DMKO55.1b | Wildtype | Noise | Noise +7 days | 24 | | | 0.04 | 0.12 | 0.21 | -0.05 | 0.54 | 0.37 | 0.42 | 0.68 | 0.45 | 0.73 | 1.03 | 1.52 | 1.24 | 1.84 | 2.06 | 2.03 | 2.27 | 2.82 |
| DMKO55.1h | Wildtype | Noise | Noise +7 days | 24 | | | -0.06 | 0.09 | 0.14 | -0.26 | 0.37 | 0.41 | 0.32 | 0.5 | 0.83 | 1.18 | 0.72 | 1.35 | 1.17 | 1.39 | 1.4 | 0.99 | 1.66 | 1.72 |
| DMKO57.1b | Wildtype | Noise | Noise +7 days | 24 | | | | | | | | | | | | | | | | | | | | |
| DMKO44.1f | Heterozygote | Noise | Noise +7 days | 24 | | | 0.05 | 0.09 | 0.09 | 0.3 | 0.19 | 0.35 | 0.21 | 0.04 | 0.28 | 0.16 | 0.04 | 0.25 | 0.46 | 0.57 | 0.74 | 0.64 | 1.01 | 1.36 |
| DMKO44.1e | Heterozygote | Noise | Noise +7 days | 24 | | | -0.11 | 0.16 | -0.18 | -0.08 | 0.35 | 0.51 | 0.76 | 0.56 | 0.61 | 1.02 | 1.19 | 1.07 | 1.14 | 1.34 | 1.31 | 1.74 | 1.86 | 1.77 |
| DMKO54.1c | Heterozygote | Noise | Noise +7 days | 24 | | | | | | | | | | | | | | | | | | | | |
| DMKO55.1c | Heterozygote | Noise | Noise +7 days | 24 | | | 0.35 | 0.23 | 0.43 | 0.66 | 0.6 | 0.58 | 0.63 | 0.87 | 1.19 | 0.96 | 1.24 | 1.25 | 1.39 | 1.76 | 1.7 | 1.73 | 1.82 | 2.17 |
| DMKO55.1g | Heterozygote | Noise | Noise +7 days | 24 | | | | | | | | | | | | | | | | | | | | |
| DMKO57.1a | Heterozygote | Noise | Noise +7 days | 24 | | | | | | | | | | | | | | | | | | | | |
| DMKO44.1a | Wildtype | Control | Noise +14 days | 12 | 0.19 | 0.11 | -0.01 | 0.59 | 0.31 | 0.31 | 0.49 | 0.97 | 0.91 | 1.24 | 1.25 | 1.69 | 2.28 | 2.49 | 2.48 | 2.37 | 2.41 | 2.16 | 2.74 | 2.63 |
| DMKO54.1f | Wildtype | Control | Noise +14 days | 12 | 0.19 | -0.15 | 0.52 | 0.47 | 0.63 | 0.32 | 0.88 | 0.61 | 1.41 | 1.51 | 1.52 | 1.66 | 1.79 | 2.3 | 3.25 | 3.55 | 4.03 | 4.17 | 4.72 | 4.81 |
| CNTC20.1a | Wildtype | Control | Noise +14 days | 12 | 0.06 | 0.03 | 0.43 | 0.38 | 0.7 | 0.65 | 0.98 | 1.2 | 2 | 2.03 | 2.13 | 1.9 | 2.64 | 3.2 | 4.04 | 4.49 | 4.3 | 4.78 | 5.01 | 5.12 |
| CNTC20.1f | Wildtype | Control | Noise +14 days | 12 | 0.15 | -0.03 | 0.22 | 0.09 | 0.52 | 0.37 | 1.33 | 0.69 | 0.88 | 1.23 | 1.73 | 1.73 | 1.91 | 2.27 | 3.02 | 3.35 | 3.53 | 3.95 | 4.29 | 4.33 |
| CNTC20.1g | Wildtype | Control | Noise +14 days | 12 | 0.37 | 0.52 | 0.52 | 0.75 | 0.43 | 0.58 | 1.08 | 1.19 | 0.88 | 1.35 | 1.92 | 2.09 | 2.16 | 2.8 | 3.38 | 4.09 | 4.16 | 4.25 | 4.47 | 4.88 |
| DMKO55.1a | Wildtype | Control | Noise +14 days | 12 | 0.26 | 0.03 | 0.36 | 0.42 | 0.62 | 0.86 | 0.8 | 1.03 | 1.41 | 1.51 | 1.61 | 1.84 | 1.79 | 2.41 | 3.58 | 3.67 | 3.59 | 4.1 | 4.73 | 4.57 |
| DMKO44.1b | Heterozygote | Control | Noise +14 days | 12 | 0.31 | -0.23 | -0.06 | 0.17 | 0.15 | 0.14 | 0.56 | 0.62 | 0.86 | 0.42 | 1.08 | 1.01 | 1.27 | 1.52 | 1.72 | 1.97 | 2.96 | 2.86 | 3.35 | 3.3 |
| DMKO44.1d | Heterozygote | Control | Noise +14 days | 12 | -0.02 | 0.29 | 0.25 | 0.38 | 0.15 | 0.27 | 0.64 | 0.64 | 1.01 | 1.16 | 0.77 | 1.04 | 1.29 | 1.43 | 2.18 | 1.71 | 1.73 | 2 | 2.7 | 2.72 |
| DMKO54.1e | Heterozygote | Control | Noise +14 days | 12 | 0.1 | 0.03 | 0.23 | 0.45 | 0.15 | 0.22 | 0.72 | 0.56 | 1.06 | 1.16 | 1.54 | 1.51 | 1.74 | 2.02 | 2.26 | 2.33 | 2.62 | 3.24 | 3.25 | 3.17 |
| DMKO55.1i | Heterozygote | Control | Noise +14 days | 12 | 0.28 | 0.49 | 0.28 | 0.31 | 0.24 | 0.54 | 0.82 | 1.02 | 0.88 | 0.87 | 0.92 | 1.43 | 1.6 | 1.62 | 2.14 | 2.55 | 2.68 | 3.12 | 3.64 | 3.87 |
| DMKO57.1c | Heterozygote | Control | Noise +14 days | 12 | 0.08 | 0.08 | 0.23 | 0.28 | 0.31 | 0.63 | 0.93 | 1.09 | 1.22 | 1.42 | 1.66 | 1.87 | 1.87 | 2.26 | 2.54 | 3.14 | 3.48 | 3.93 | 4.55 | 4.14 |
| DMKO57.1e | Heterozygote | Control | Noise +14 days | 12 | 0.23 | 0.07 | -0.04 | 0.28 | 0.68 | 0.52 | 0.49 | 0.63 | 0.65 | 0.88 | 0.87 | 1.07 | 1.45 | 1.24 | 1.09 | 1.23 | 1.06 | 1.55 | 1.6 | 1.92 |
| DMKO44.1g | Wildtype | Noise | Noise +14 days | 12 | 0.35 | 0.17 | 0.62 | 0.47 | 0.62 | 0.85 | 0.91 | 1.25 | 1.62 | 1.9 | 2.05 | 2.48 | 2.88 | 3.72 | 4.4 | 4.92 | 4.83 | 4.55 | 5.02 | 4.63 |
| DMKO44.1c | Wildtype | Noise | Noise +14 days | 12 | 0.21 | 0.06 | 0.39 | 0.5 | 0.59 | 0.46 | 0.57 | 1.08 | 1.12 | 1.46 | 1.13 | 1.92 | 1.98 | 2.91 | 3.54 | 3.73 | 3.46 | 3.42 | 3.66 | 3.13 |
| DMKO54.1d | Wildtype | Noise | Noise +14 days | 12 | 0.07 | 0.15 | 0.54 | 0.44 | 0.72 | 1.12 | 1.11 | 1.28 | 1.41 | 2.25 | 2.18 | 2 | 2.87 | 3.51 | 4.3 | 5.01 | 4.74 | 4.8 | 5.17 | 4.62 |
| DMKO55.1b | Wildtype | Noise | Noise +14 days | 12 | 0.15 | -0.07 | 0.11 | 0.39 | 0.65 | 0.63 | 0.71 | 1.08 | 1.36 | 1.68 | 1.6 | 1.98 | 2.23 | 2.76 | 3.47 | 3.47 | 3.74 | 4.11 | 4.12 | 3.67 |
| DMKO55.1h | Wildtype | Noise | Noise +14 days | 12 | 0.36 | 0.28 | 0.69 | 0.6 | 0.54 | 0.87 | 0.88 | 1.33 | 1.11 | 1.79 | 2.11 | 2.29 | 2.48 | 3 | 4.2 | 4.72 | 4.69 | 4.8 | 4.98 | 4.86 |
| DMKO57.1b | Wildtype | Noise | Noise +14 days | 12 | 0.24 | 0.11 | 0.09 | 0.2 | 0.48 | 0.57 | 0.93 | 1.07 | 1.32 | 1.68 | 2.04 | 2.4 | 3 | 3.2 | 3.69 | 3.91 | 3.77 | 3.62 | 3.82 | 3.79 |
| DMKO44.1f | Heterozygote | Noise | Noise +14 days | 12 | -0.01 | -0.03 | 0.08 | 0.14 | 0.17 | 0.27 | 0.72 | 0.73 | 1.02 | 1.28 | 1.39 | 1.65 | 1.86 | 2.45 | 2.63 | 2.65 | 3.19 | 3.42 | 3.54 | 3.73 |
| DMKO44.1e | Heterozygote | Noise | Noise +14 days | 12 | 0.56 | 0.34 | 0.39 | 0.18 | 0.29 | 0.46 | 0.24 | 1.03 | 0.83 | 1.24 | 1.03 | 1.48 | 1.54 | 1.87 | 2.17 | 2.8 | 3.16 | 3.68 | 4.75 | 4.7 |
| DMKO54.1c | Heterozygote | Noise | Noise +14 days | 12 | 0.24 | 0.16 | 0.34 | 0.57 | 0.39 | 0.44 | 0.69 | 1.23 | 1.21 | 1.45 | 1.79 | 2.06 | 2.03 | 2.36 | 2.62 | 2.54 | 2.59 | 2.88 | 2.86 | 2.64 |
| DMKO55.1c | Heterozygote | Noise | Noise +14 days | 12 | 0.23 | 0.03 | 0.06 | 0.27 | 0.43 | 0.68 | 0.75 | 0.83 | 1.22 | 1.14 | 1.03 | 1.36 | 1.64 | 1.75 | 2.55 | 2.33 | 2.17 | 2.97 | 2.88 | 2.57 |
| DMKO55.1g | Heterozygote | Noise | Noise +14 days | 12 | 0.06 | 0.15 | 0.36 | 0.43 | 0.37 | 0.99 | 0.67 | 0.96 | 1.07 | 0.94 | 1.38 | 1.37 | 1.78 | 1.87 | 2.2 | 2.58 | 2.52 | 2.83 | 2.83 | 2.13 |
| DMKO57.1a | Heterozygote | Noise | Noise +14 days | 12 | 0.1 | 0.35 | 0.46 | 0.18 | 0.48 | 0.23 | 0.69 | 0.6 | 0.53 | 0.68 | 0.79 | 1.13 | 1.45 | 1.47 | 1.64 | 1.69 | 1.46 | 1.62 | 2.15 | 1.87 |
| DMKO44.1a | Wildtype | Control | Noise +14 days | 24 | | | 0.28 | 0.13 | -0.04 | -0.14 | 0.26 | 0.53 | 0.34 | 0.35 | 0.62 | 0.94 | 0.91 | 1.1 | 1.32 | 1.46 | 1.69 | 1.89 | 2.1 | 2.33 |
| DMKO54.1f | Wildtype | Control | Noise +14 days | 24 | | | 0.32 | -0.04 | 0.54 | 0.46 | 0.68 | 0.85 | 0.79 | 0.8 | 1.07 | 1.49 | 1.42 | 1.82 | 1.86 | 2.18 | 2.67 | 2.67 | 3.45 | 3.88 |
| CNTC20.1a | Wildtype | Control | Noise +14 days | 24 | | | 0.19 | 0.6 | 0.2 | 0.52 | 0.64 | 0.79 | 1.13 | 1.49 | 1.12 | 1.27 | 1.67 | 2.19 | 2.21 | 3.05 | 3.34 | 3.5 | 3.78 | 3.89 |
| CNTC20.1f | Wildtype | Control | Noise +14 days | 24 | | | 0.44 | 0.08 | 0.34 | 0.34 | 0.36 | 0.51 | 0.91 | 0.75 | 0.76 | 1.3 | 1.55 | 1.95 | 2.39 | 2.34 | 2.8 | 3.07 | 3.01 | 3.43 |
| CNTC20.1g | Wildtype | Control | Noise +14 days | 24 | | | 0.12 | 0.16 | 0.61 | 0.35 | 0.19 | 0.81 | 0.37 | 0.8 | 0.95 | 1.35 | 1.35 | 1.82 | 2.03 | 2.26 | 2.76 | 3.21 | 3.7 | 3.63 |
| DMKO55.1a | Wildtype | Control | Noise +14 days | 24 | | | -0.1 | 0.25 | 0.34 | 0.79 | 0.32 | 0.76 | 0.94 | 0.92 | 1.13 | 1.37 | 1.61 | 2.01 | 2.49 | 2.65 | 2.45 | 3.14 | 3.81 | 3.71 |
| DMKO44.1b | Heterozygote | Control | Noise +14 days | 24 | | | 0.29 | 0.16 | 0.17 | 0.06 | 0.52 | 0.58 | 0.9 | 0.47 | 0.8 | 1.05 | 1.19 | 1.49 | 1.58 | 1.48 | 2.1 | 1.99 | 2.09 | 2.46 |
| DMKO44.1d | Heterozygote | Control | Noise +14 days | 24 | | | 0.07 | 0.07 | 0.21 | 0.21 | 0.1 | 0.23 | 0.32 | 0.63 | 0.38 | 0.47 | 0.7 | 0.54 | 0.61 | 0.88 | 0.99 | 0.89 | 1.34 | 1.5 |
| DMKO54.1e | Heterozygote | Control | Noise +14 days | 24 | | | 0.23 | 0.22 | 0.13 | 0.31 | 0.09 | 0.46 | 0.46 | 0.64 | 1.13 | 1.39 | 1.27 | 1.54 | 1.94 | 2.28 | 2.5 | 2.71 | 2.71 | 3.36 |
| DMKO55.1i | Heterozygote | Control | Noise +14 days | 24 | | | 0.01 | 0.2 | 0.08 | 0.39 | 0.71 | 0.4 | 0.73 | 0.66 | 0.96 | 1.02 | 1.41 | 1.71 | 1.9 | 2.26 | 2.57 | 2.5 | 2.74 | 3.1 |
| DMKO57.1c | Heterozygote | Control | Noise +14 days | 24 | | | 0.04 | 0.1 | 0.35 | 0.46 | 0.53 | 0.54 | 0.52 | 0.96 | 0.98 | 1.08 | 1.36 | 1.79 | 1.85 | 2.11 | 2.52 | 2.28 | 2.5 | 2.45 |
| DMKO57.1e | Heterozygote | Control | Noise +14 days | 24 | | | 0.12 | 0.2 | 0.29 | 0.1 | 0.08 | 0.2 | 0.3 | 0.46 | 0.39 | 0.61 | 0.79 | 0.87 | 1.1 | 1.06 | 1.06 | 1.22 | 1.31 | 1.42 |
| DMKO44.1g | Wildtype | Noise | Noise +14 days | 24 | | | 0.02 | 0.01 | 0.17 | 0.57 | 0.33 | 0.75 | 1.21 | 0.91 | 1.64 | 1.34 | 1.46 | 2.26 | 2.01 | 2.59 | 2.93 | 2.96 | 3.07 | 3.5 |
| DMKO44.1c | Wildtype | Noise | Noise +14 days | 24 | | | 0.04 | 0.07 | -0.06 | 0.38 | 0.36 | 0.44 | 0.54 | 0.95 | 1.05 | 1.3 | 1.29 | 1.82 | 2.13 | 2.25 | 2.49 | 2.43 | 2.89 | 2.93 |
| DMKO54.1d | Wildtype | Noise | Noise +14 days | 24 | | | -0.02 | 0.13 | 0.35 | 0.15 | 0.61 | 0.69 | 0.95 | 0.88 | 1.62 | 1.71 | 1.7 | 1.87 | 2.24 | 2.54 | 2.68 | 3.27 | 3.39 | 3.96 |
| DMKO55.1b | Wildtype | Noise | Noise +14 days | 24 | | | 0.32 | 0.04 | 0.72 | 0.35 | 0.41 | 0.79 | 0.8 | 0.75 | 1.02 | 1.48 | 1.48 | 1.89 | 2.13 | 2.35 | 2.65 | 2.58 | 2.95 | 3.38 |
| DMKO55.1h | Wildtype | Noise | Noise +14 days | 24 | | | 0.21 | 0.01 | -0.01 | 0.38 | 0.55 | 0.54 | 0.95 | 1.07 | 1.4 | 1.54 | 1.87 | 1.95 | 2.24 | 2.55 | 2.78 | 2.97 | 2.97 | 3.43 |
| DMKO57.1b | Wildtype | Noise | Noise +14 days | 24 | | | 0.14 | 0.14 | 0 | 0.24 | 0.21 | 0.56 | 0.82 | 1.07 | 1.27 | 1.32 | 1.64 | 1.74 | 1.97 | 1.91 | 2.22 | 2.32 | 2.4 | 2.64 |
| DMKO44.1f | Heterozygote | Noise | Noise +14 days | 24 | | | 0.37 | -0.35 | 0.11 | 0.33 | 0.08 | 0.26 | 0.18 | 0.46 | 0.31 | 0.51 | 0.46 | 0.66 | 0.49 | 0.51 | 0.62 | 0.58 | 0.93 | 1.12 |
| DMKO44.1e | Heterozygote | Noise | Noise +14 days | 24 | | | 0.36 | -0.11 | 0 | 0.11 | 0.3 | 0.22 | 0.47 | 0.48 | 0.63 | 1.07 | 1.3 | 1.44 | 1.43 | 1.76 | 1.74 | 1.92 | 1.92 | 1.99 |
| DMKO54.1c | Heterozygote | Noise | Noise +14 days | 24 | | | -0.03 | 0.32 | 0.24 | 0.25 | 0.39 | 0.1 | -0.03 | 0.46 | 0.67 | 0.8 | 1.04 | 0.92 | 1.22 | 1.16 | 1.36 | 1.78 | 2.06 | 2.39 |
| DMKO55.1c | Heterozygote | Noise | Noise +14 days | 24 | | | 0.07 | 0.15 | 0.26 | 0.12 | 0.45 | 0.75 | 0.6 | 0.86 | 1.02 | 1.1 | 1.08 | 1.03 | 1.54 | 1.54 | 1.91 | 1.88 | 1.95 | 2.2 |
| DMKO55.1g | Heterozygote | Noise | Noise +14 days | 24 | | | 0 | 0.35 | 0.06 | 0.23 | 0.18 | 0.55 | 0.74 | 0.74 | 0.64 | 1.25 | 1.13 | 1.49 | 1.65 | 1.34 | 1.63 | 1.64 | 1.76 | 1.82 |
| DMKO57.1a | Heterozygote | Noise | Noise +14 days | 24 | | | 0.29 | -0.1 | 0.13 | 0.04 | 0.13 | 0.72 | 0.35 | 0.62 | 0.22 | 0.53 | 0.6 | 0.75 | 0.45 | 0.91 | 0.94 | 0.82 | 0.66 | 0.79 |
| DMKO44.1a | Wildtype | Control | Noise +28 days | 12 | | | 0.28 | 0.13 | -0.04 | -0.14 | 0.26 | 0.53 | 0.34 | 0.35 | 0.62 | 0.94 | 0.91 | 1.1 | 1.32 | 1.46 | 1.69 | 1.89 | 2.1 | 2.33 |
| DMKO54.1f | Wildtype | Control | Noise +28 days | 12 | 0.24 | 0.21 | 0.51 | 0.24 | 0.57 | 0.92 | 0.92 | 0.93 | 1.17 | 1.9 | 1.85 | 2.21 | 2.4 | 2.7 | 3.16 | 3.84 | 4.36 | 4.83 | 5.21 | 5.62 |
| CNTC20.1a | Wildtype | Control | Noise +28 days | 12 | 0.29 | 0.2 | 0.02 | 0.47 | 0.66 | 0.59 | 0.87 | 0.99 | 1.29 | 1.67 | 1.77 | 1.87 | 2.12 | 2.64 | 3.39 | 3.5 | 3.41 | 3.46 | 4.15 | 3.98 |
| CNTC20.1f | Wildtype | Control | Noise +28 days | 12 | 0 | 0.1 | 0.34 | 0.37 | 0.47 | 0.39 | 0.65 | 1.07 | 0.63 | 1.42 | 1.33 | 1.69 | 1.52 | 2.33 | 2.66 | 3.26 | 3.61 | 3.88 | 4.04 | 3.99 |
| CNTC20.1g | Wildtype | Control | Noise +28 days | 12 | 0.11 | 0.23 | -0.23 | 0.44 | 0.7 | 0.61 | 0.57 | 0.95 | 1.35 | 1.21 | 1.43 | 2.06 | 2.21 | 2.68 | 3.35 | 3.32 | 3.62 | 4.03 | 4.26 | |
| DMKO55.1a | Wildtype | Control | Noise +28 days | 12 | 0.05 | 0.4 | 0.38 | 0.44 | 0.63 | 0.91 | 0.77 | 1.1 | 1.47 | 1.79 | 1.8 | 2 | 1.98 | 3.02 | 3.74 | 4.05 | 4.21 | 4.96 | 5.32 | 5.25 |
| DMKO44.1b | Heterozygote | Control | Noise +28 days | 12 | 0.29 | -0.02 | 0.14 | 0.19 | 0.28 | 0.52 | 0.66 | 1.02 | 0.63 | 1.01 | 1.15 | 1.29 | 1.49 | 1.78 | 2.01 | 2.1 | 2.53 | 2.77 | 3.18 | 2.89 |
| DMKO44.1d | Heterozygote | Control | Noise +28 days | 12 | 0.01 | 0.06 | -0.06 | 0.12 | 0.06 | -0.16 | 0.28 | 0.21 | 0.58 | -0.02 | 0.68 | 0.28 | 0.49 | 0.64 | 0.72 | 1.01 | 1.4 | 1.64 | 1.55 | 2.03 |
| DMKO54.1e | Heterozygote | Control | Noise +28 days | 12 | 0.3 | 0.19 | 0.29 | 0.36 | -0.11 | 0.32 | 0.29 | 0.29 | 1.05 | 1.14 | 1.19 | 2.01 | 1.99 | 2.51 | 1.94 | 2.37 | 2.91 | 3.55 | 4.18 | 3.96 |
| DMKO55.1i | Heterozygote | Control | Noise +28 days | 12 | 0.25 | 0.08 | 0.35 | 0.16 | 0.33 | 0.85 | 0.43 | 0.75 | 1.04 | 1.39 | 1.49 | 1.23 | 1.91 | 2.04 | 2.33 | 2.76 | 2.79 | 3.17 | 3.5 | 3.75 |
| DMKO57.1c | Heterozygote | Control | Noise +28 days | 12 | -0.07 | 0.33 | 0.25 | 0.32 | 0.75 | 0.57 | 1.22 | 0.97 | 1.42 | 1.63 | 1.87 | 1.88 | 2.13 | 2.52 | 2.94 | 3.13 | 3.36 | 3.68 | 4 | 4.13 |
| DMKO57.1e | Heterozygote | Control | Noise +28 days | 12 | 0.2 | 0.36 | 0.16 | 0.64 | 0.58 | 0.77 | 0.8 | 0.95 | 1.13 | 1.39 | 1.51 | 1.83 | 1.72 | 1.94 | 1.95 | 1.99 | 1.97 | 2.16 | 2.97 | 2.84 |
| DMKO44.1g | Wildtype | Noise | Noise +28 days | 12 | 0.39 | 0.42 | 0.57 | 0.63 | 0.8 | 0.91 | 0.9 | 1.06 | 1.5 | 2.02 | 2.37 | 2.09 | 2.94 | 3.97 | 4.67 | 4.82 | 4.88 | 4.94 | 5.39 | 5.27 |
| DMKO44.1c | Wildtype | Noise | Noise +28 days | 12 | 0.19 | 0.2 | -0.19 | 0.32 | 0.45 | 0.41 | 0.69 | 0.79 | 1.39 | 1.35 | 1.74 | 1.68 | 2.39 | 2.96 | 3.46 | 3.35 | 3.64 | 3.64 | 3.85 | 3.86 |
| DMKO54.1d | Wildtype | Noise | Noise +28 days | 12 | 0.15 | 0.08 | 0.36 | 0.33 | 0.53 | 0.96 | 1.19 | 1.06 | 1.69 | 1.67 | 2.22 | 2.32 | 2.64 | 3.09 | 3.65 | 3.95 | 3.65 | 4.11 | 4.75 | 4.32 |
| DMKO55.1b | Wildtype | Noise | Noise +28 days | 12 | 0.13 | 0.27 | 0.39 | 0.38 | 0.58 | 0.6 | 0.81 | 0.98 | 1.35 | 1.57 | 1.58 | 2.06 | 2.16 | 3.14 | 3.88 | 4.09 | 4.26 | 4.4 | 4.71 | 4.32 |
| DMKO55.1h | Wildtype | Noise | Noise +28 days | 12 | 0.18 | 0.22 | 0.39 | 0.21 | 0.55 | 0.84 | 0.78 | 1.13 | 1.16 | 1.59 | 1.86 | 2.29 | 2.2 | 3.22 | 3.85 | 4.05 | 4.24 | 3.99 | 4.65 | 4.36 |
| DMKO57.1b | Wildtype | Noise | Noise +28 days | 12 | 0.09 | 0.18 | 0.24 | 0.48 | 0.57 | 0.58 | 0.96 | 1.38 | 1.49 | 1.8 | 2.17 | 2.19 | 3.03 | 3.79 | 3.8 | 4 | 4.2 | 4.09 | 4.01 | 4.05 |
| DMKO44.1f | Heterozygote | Noise | Noise +28 days | 12 | 0.01 | 0.06 | 0.28 | 0.02 | 0.75 | 0.36 | 0.58 | 1.29 | 1.44 | 1.42 | 1.93 | 2.27 | 2.33 | 2.96 | 2.84 | 2.87 | 2.99 | 2.75 | 2.74 | 3.33 |
| DMKO44.1e | Heterozygote | Noise | Noise +28 days | 12 | -0.02 | 0.14 | 0.37 | 0.31 | 0.47 | 0.77 | 0.81 | 1.25 | 0.9 | 1.41 | 1.78 | 1.99 | 2.19 | 2.31 | 2.81 | 3.29 | 4.57 | 5.11 | 5.34 | 5.32 |
| DMKO54.1c | Heterozygote | Noise | Noise +28 days | 12 | 0.2 | -0.08 | -0.07 | 0.54 | 0.51 | 0.91 | 0.67 | 0.75 | 1.51 | 1.67 | 1.92 | 1.96 | 2.34 | 3.09 | 3.02 | 2.8 | 3.28 | 3.6 | 3.77 | 3.11 |
| DMKO55.1c | Heterozygote | Noise | Noise +28 days | 12 | -0.01 | 0 | -0.01 | 0.19 | 0.52 | 0.64 | 0.79 | 0.7 | 1.09 | 1.3 | 1.4 | 1.53 | 1.85 | 1.76 | 2.45 | 2.82 | 2.85 | 3.1 | 3.7 | 3.65 |
| DMKO55.1g | Heterozygote | Noise | Noise +28 days | 12 | -0.17 | 0.08 | 0.05 | 0.42 | 0.53 | 0.74 | 0.71 | 0.6 | 0.73 | 1.09 | 1.39 | 1.55 | 1.54 | 2.21 | 2.29 | 2.69 | 3.02 | 3.17 | 3.33 | 2.87 |
| DMKO57.1a | Heterozygote | Noise | Noise +28 days | 12 | 0.36 | -0.1 | 0.45 | 0.27 | 0.13 | 0.51 | 0.64 | 0.86 | 1.33 | 1.62 | 1.39 | 2.04 | 2.23 | 2.73 | 2.99 | 2.89 | 3.35 | 2.89 | 3.19 | 3.62 |
| DMKO44.1a | Wildtype | Control | Noise +28 days | 24 | | | 0.05 | 0 | 0 | 0 | 0.46 | -0.18 | 0.62 | 0.46 | 0.43 | 0.44 | 0.2 | 0.71 | 0.62 | 1.06 | 1.15 | 1.5 | 1.81 | 1.96 |
| DMKO54.1f | Wildtype | Control | Noise +28 days | 24 | | | 0.17 | 0.3 | 0.28 | 0.3 | 0.28 | 0.63 | 0.68 | 0.96 | 1.07 | 1.2 | 1.15 | 1.43 | 1.61 | 1.83 | 2.11 | 2.54 | 2.76 | 3.83 |
| CNTC20.1a | Wildtype | Control | Noise +28 days | 24 | | | 0.34 | 0.27 | 0.29 | 0.7 | 0.28 | 0.52 | 0.74 | 0.87 | 1.3 | 1.11 | 1.34 | 1.74 | 1.93 | 2.16 | 2.66 | 2.94 | 2.89 | 3.09 |
| CNTC20.1f | Wildtype | Control | Noise +28 days | 24 | | | 0.23 | 0.36 | 0.01 | 0.07 | 0.64 | 0.31 | 0.55 | 0.72 | 1.43 | 1.22 | 1.31 | 1.94 | 2.33 | 1.9 | 2.6 | 2.12 | 2.64 | 3.06 |
| CNTC20.1g | Wildtype | Control | Noise +28 days | 24 | | | 0.07 | 0.28 | 0.19 | 0.57 | 0.11 | 0.73 | 0.63 | 1.01 | 0.91 | 1.17 | 1.33 | 1.68 | 1.88 | 2.38 | 2.74 | 2.95 | 2.82 | |
| DMKO55.1a | Wildtype | Control | Noise +28 days | 24 | | | 0.24 | 0.1 | 0.16 | 0.35 | 0.75 | 0.86 | 0.87 | 1.14 | 1.65 | 1.55 | 1.77 | 2.41 | 2.64 | 3.04 | 3.26 | 3.46 | 4.11 | 4.4 |
| DMKO44.1b | Heterozygote | Control | Noise +28 days | 24 | | | -0.19 | 0.13 | -0.16 | 0.03 | -0.14 | 0.15 | 0.53 | 0.41 | 0.6 | 0.71 | 0.73 | 0.96 | 1.31 | 1.08 | 1.34 | 1.28 | 1.26 | 1.26 |
| DMKO44.1d | Heterozygote | Control | Noise +28 days | 24 | | | 0.24 | 0.08 | -0.04 | 0.11 | 0.06 | 0.11 | 0.61 | 0.2 | 0.45 | 0.51 | 0.18 | 0.54 | 0.23 | 0.35 | 0.4 | 0.3 | 0.51 | 0.66 |
| DMKO54.1e | Heterozygote | Control | Noise +28 days | 24 | | | 0.23 | 0.12 | 0.24 | 0.12 | 0.25 | 0.39 | 0.5 | 0.43 | 0.99 | 1.09 | 1.44 | 1.77 | 1.61 | 1.68 | 2.19 | 2.46 | 2.28 | 2.41 |
| DMKO55.1i | Heterozygote | Control | Noise +28 days | 24 | | | 0.2 | 0.21 | 0.16 | 0.21 | 0.55 | 0.37 | 0.49 | 0.61 | 0.59 | 1.13 | 1.45 | 1.65 | 2.12 | 2.04 | 2.03 | 2.39 | 2.38 | 2.67 |
| DMKO57.1c | Heterozygote | Control | Noise +28 days | 24 | | | 0.19 | -0.08 | 0.39 | 0.28 | 0.13 | 0.28 | 0.46 | 0.51 | 0.91 | 1.16 | 1.19 | 1.53 | 1.72 | 1.58 | 2 | 2.05 | 1.89 | 2.03 |
| DMKO57.1e | Heterozygote | Control | Noise +28 days | 24 | | | 0.3 | 0.34 | 0.13 | 0.24 | -0.08 | 0.23 | 0.33 | 0.29 | 0.48 | 0.43 | 0.78 | 0.71 | 1.1 | 1.38 | 1.24 | 1.26 | 1.66 | 2.09 |
| DMKO44.1g | Wildtype | Noise | Noise +28 days | 24 | | | -0.08 | 0.09 | 0.07 | 0.22 | 0.25 | 0.78 | 0.78 | 0.87 | 1.16 | 1.43 | 1.81 | 1.98 | 2.09 | 2.42 | 2.47 | 2.77 | 2.7 | 3.08 |
| DMKO44.1c | Wildtype | Noise | Noise +28 days | 24 | | | 0.34 | -0.03 | 0.14 | -0.17 | 0.38 | 0.44 | 0.42 | 1.07 | 0.92 | 1.22 | 1.32 | 1.61 | 1.6 | 1.96 | 2.45 | 2.2 | 2.51 | 2.82 |
| DMKO54.1d | Wildtype | Noise | Noise +28 days | 24 | | | 0 | 0.27 | 0.14 | 0.45 | 0.47 | 0.54 | 0.41 | 0.76 | 0.92 | 1.08 | 1.61 | 1.34 | 1.88 | 1.82 | 2.22 | 2.45 | 2.69 | 3.12 |

| | | | | | | | | | | | | | | | | | | | | | | |
|---|---|---|---|---|---|---|---|---|---|---|---|---|---|---|---|---|---|---|---|---|---|---|
| DMKO55.1b | Wildtype | Noise | Noise +28 days | 24 | 0.23 | 0.27 | -0.32 | 0.24 | 0.39 | 0.42 | 0.67 | 0.62 | 1.17 | 1.6 | 1.44 | 1.7 | 2.33 | 2.38 | 2.58 | 2.68 | 2.88 | 3.51 |
| DMKO55.1h | Wildtype | Noise | Noise +28 days | 24 | 0.1 | 0.28 | -0.04 | 0.27 | 0.48 | 0.56 | 0.74 | 1.09 | 1.35 | 1.38 | 1.58 | 2.08 | 2.44 | 2.38 | 2.65 | 2.68 | 3.04 | 3.45 |
| DMKO57.1b | Wildtype | Noise | Noise +28 days | 24 | 0.37 | 0.34 | -0.07 | -0.03 | 0.28 | 0.35 | 0.11 | 0.62 | 0.51 | 0.31 | 0.54 | 0.59 | 1.05 | 1.18 | 1.29 | 2.11 | 1.98 | 2.15 |
| DMKO44.1f | Heterozygote | Noise | Noise +28 days | 24 | | | | | | | | | | | | | | | | | | |
| DMKO44.1e | Heterozygote | Noise | Noise +28 days | 24 | 0 | 0 | 0.25 | 0.09 | 0.44 | 0.69 | 0.8 | 0.68 | 1.17 | 1.28 | 1.62 | 1.87 | 1.57 | 1.71 | 1.79 | 1.49 | 2.02 | 2.11 |
| DMKO54.1c | Heterozygote | Noise | Noise +28 days | 24 | 0.14 | 0.18 | 0.05 | 0.27 | 0.33 | 0.44 | 0.22 | 0.55 | 0.68 | 1.03 | 1.17 | 1.29 | 1.46 | 1.55 | 1.58 | 1.68 | 1.94 | 2.01 |
| DMKO55.1c | Heterozygote | Noise | Noise +28 days | 24 | -0.15 | 0.08 | 0.29 | 0.14 | 0.53 | 0.34 | 0.57 | 0.67 | 0.77 | 1.08 | 1.31 | 1.5 | 1.41 | 1.33 | 1.48 | 1.89 | 2.1 | 2.38 |
| DMKO55.1g | Heterozygote | Noise | Noise +28 days | 24 | 0.13 | 0.1 | -0.1 | 0.15 | 0.38 | 0.25 | 0.33 | 0.68 | 0.72 | 0.93 | 0.98 | 1.31 | 1.22 | 1.53 | 1.32 | 1.82 | 1.63 | 1.79 |
| DMKO57.1a | Heterozygote | Noise | Noise +28 days | 24 | 0.33 | -0.02 | 0.07 | 0.06 | 0.21 | 0.24 | 0.43 | 0.38 | 0.34 | 0.58 | 0.75 | 0.81 | 0.63 | 0.77 | 0.64 | 1.22 | 1.19 | 1.1 |

**Network links and references for Fig. 7A**

***Mir183/96*** $^{dko}$ **network**

| Gene 1 | Interaction | Gene 2 | Reference |
|---|---|---|---|
| ENSMUSG00000025958 | upregulates | ENSMUSG00000048482 | TRRUST:11719924 |
| ENSMUSG00000005148 | downregulates | ENSMUSG00000061762 | ArrayExpress:E-GEOD-58719 |
| ENSMUSG00000024431 | downregulates | ENSMUSG00000028163 | manual:Lewis;TRRUST:17016446 |
| ENSMUSG00000020919 | upregulates | ENSMUSG00000006731 | ArrayExpress:E-GEOD-37646 |
| ENSMUSG00000004043 | downregulates | ENSMUSG00000061762 | ArrayExpress:E-GEOD-37646 |
| ENSMUSG00000065474 | downregulates | ENSMUSG00000024238 | miRTarBase:18411277;miRTarBase:19854497;miRTarBase:22012804;miRTarBase:25502084;miRTarBase:26160158;miRTarBase:21049046 |
| ENSMUSG00000065397 | downregulates | ENSMUSG00000022508 | miRTarBase:23041630;miRTarBase:26986233 |
| ENSMUSG00000029913 | downregulates | ENSMUSG00000005442 | ArrayExpress:E-GEOD-46821 |
| ENSMUSG00000065586 | downregulates | ENSMUSG00000031450 | manual:current |
| ENSMUSG00000065397 | downregulates | ENSMUSG00000041439 | ArrayExpress:E-GEOD-68511 |
| ENSMUSG00000025958 | downregulates | ENSMUSG00000021250 | TRRUST:1840296 |
| ENSMUSG00000065619 | downregulates | ENSMUSG00000031450 | manual:current |
| ENSMUSG00000000782 | downregulates | ENSMUSG00000030592 | ArrayExpress:E-GEOD-54976 |
| ENSMUSG00000017950 | downregulates | ENSMUSG00000075316 | ArrayExpress:E-MEXP-1729 |
| ENSMUSG00000026872 | downregulates | ENSMUSG00000065592 | TransmiR:22908280;TransmiR:25296715 |
| ENSMUSG00000005148 | downregulates | ENSMUSG00000051159 | ArrayExpress:E-GEOD-58719 |
| ENSMUSG00000032855 | downregulates | ENSMUSG00000026739 | ArrayExpress:E-TABM-411 |
| ENSMUSG00000032855 | downregulates | ENSMUSG00000056749 | ArrayExpress:E-TABM-411 |
| ENSMUSG00000022105 | downregulates | ENSMUSG00000018983 | TRRUST:9671466 |
| ENSMUSG00000065422 | downregulates | ENSMUSG00000004043 | miRTarBase:24736554 |
| ENSMUSG00000065422 | downregulates | ENSMUSG00000059552 | miRTarBase:21226887 |
| ENSMUSG00000024238 | downregulates | ENSMUSG00000065592 | TransmiR:22908280 |
| ENSMUSG00000028163 | upregulates | ENSMUSG00000027398 | TRRUST:12686724;14960579;8413223;8679226;9878621;TRRUST:10437651;11454276;14670622;16399630;16867259;24132147 |
| ENSMUSG00000044167 | upregulates | ENSMUSG00000022676 | ArrayExpress:E-MTAB-4023 |
| ENSMUSG00000047146 | downregulates | ENSMUSG00000041439 | ArrayExpress:E-MTAB-4578 |
| ENSMUSG00000005672 | downregulates | ENSMUSG00000048756 | ArrayExpress:E-MEXP-2282 |
| ENSMUSG00000065493 | downregulates | ENSMUSG00000024515 | miRTarBase:28348487;miRTarBase:26077733;miRTarBase:28340489 |
| ENSMUSG00000029687 | upregulates | ENSMUSG00000036202 | ArrayExpress:E-GEOD-34961 |
| ENSMUSG00000076361 | downregulates | ENSMUSG00000022676 | miRTarBase:23354685 |
| ENSMUSG00000044786 | upregulates | ENSMUSG00000004043 | ArrayExpress:E-GEOD-5324 |
| ENSMUSG00000034701 | downregulates | ENSMUSG00000024349 | ArrayExpress:E-GEOD-35396 |
| ENSMUSG00000009406 | upregulates | ENSMUSG00000032402 | ArrayExpress:E-GEOD-21546 |
| ENSMUSG00000020919 | downregulates | ENSMUSG00000061762 | ArrayExpress:E-GEOD-37646 |
| ENSMUSG00000029249 | upregulates | ENSMUSG00000055024 | ORegAnno:N/A |
| ENSMUSG00000065471 | downregulates | ENSMUSG00000004043 | miRTarBase:20489169;miRTarBase:24736554 |
| ENSMUSG00000027104 | upregulates | ENSMUSG00000022105 | ORegAnno:15618023 |
| ENSMUSG00000004043 | upregulates | ENSMUSG00000034220 | ArrayExpress:E-GEOD-37646 |
| ENSMUSG00000018983 | upregulates | ENSMUSG00000044167 | ArrayExpress:E-GEOD-52157 |
| ENSMUSG00000024515 | upregulates | ENSMUSG00000023067 | TRRUST:10878024;TRRUST:12429655 |
| ENSMUSG00000034701 | downregulates | ENSMUSG00000030307 | ArrayExpress:E-GEOD-35396 |
| ENSMUSG00000052684 | upregulates | ENSMUSG00000024431 | TRRUST:9657066 |
| ENSMUSG00000000782 | downregulates | ENSMUSG00000023067 | ArrayExpress:E-GEOD-54976 |
| ENSMUSG00000065397 | downregulates | ENSMUSG00000059552 | miRTarBase:27903673 |
| ENSMUSG00000065397 | downregulates | ENSMUSG00000024515 | miRTarBase:21036908 |
| ENSMUSG00000065619 | downregulates | ENSMUSG00000010066 | manual:current |
| ENSMUSG00000105904 | downregulates | ENSMUSG00000059970 | ArrayExpress:E-MTAB-7079 |
| ENSMUSG00000021250 | upregulates | ENSMUSG00000029618 | manual:Lewis |
| ENSMUSG00000065619 | downregulates | ENSMUSG00000026739 | miRTarBase:25295122 |
| ENSMUSG00000032855 | downregulates | ENSMUSG00000007872 | ArrayExpress:E-TABM-411 |
| ENSMUSG00000065592 | downregulates | ENSMUSG00000001280 | miRTarBase:24613927;miRTarBase:24510775 |
| ENSMUSG00000027104 | upregulates | ENSMUSG00000021250 | TRRUST:21069729 |
| ENSMUSG00000001280 | upregulates | ENSMUSG00000024515 | ORegAnno:21421563 |
| ENSMUSG00000028163 | downregulates | ENSMUSG00000023067 | TRRUST:18954936 |
| ENSMUSG00000044167 | upregulates | ENSMUSG00000002603 | manual:Lewis |
| ENSMUSG00000009406 | upregulates | ENSMUSG00000029249 | ArrayExpress:E-GEOD-21546 |
| ENSMUSG00000004043 | upregulates | ENSMUSG00000021194 | ArrayExpress:E-GEOD-37646 |
| ENSMUSG00000017146 | upregulates | ENSMUSG00000023067 | TRRUST:16735508;18025037;21245169;TRRUST:21245169 |
| ENSMUSG00000034701 | downregulates | ENSMUSG00000005148 | ArrayExpress:E-GEOD-35396 |
| ENSMUSG00000001280 | downregulates | ENSMUSG00000065400 | TransmiR:27550813 |
| ENSMUSG00000005442 | upregulates | ENSMUSG00000072825 | ArrayExpress:E-MTAB-6681 |
| ENSMUSG00000004043 | upregulates | ENSMUSG00000023067 | ArrayExpress:E-GEOD-37646 |
| ENSMUSG00000065619 | downregulates | ENSMUSG00000044167 | miRTarBase:20028871;miRTarBase:24368418 |
| ENSMUSG00000002603 | downregulates | ENSMUSG00000065400 | TransmiR:18698484 |
| ENSMUSG00000065592 | downregulates | ENSMUSG00000044167 | miRTarBase:24732799 |
| ENSMUSG00000048756 | upregulates | ENSMUSG00000044167 | manual:Lewis |
| ENSMUSG00000055024 | upregulates | ENSMUSG00000017146 | TRRUST:23353771 |
| ENSMUSG00000024401 | upregulates | ENSMUSG00000028163 | manual:Lewis |
| ENSMUSG00000065619 | downregulates | ENSMUSG00000024479 | manual:current |
| ENSMUSG00000028163 | downregulates | ENSMUSG00000059552 | TRRUST:15044535 |
| ENSMUSG00000065493 | downregulates | ENSMUSG00000027323 | miRTarBase:26670277 |
| ENSMUSG00000036202 | downregulates | ENSMUSG00000026407 | ArrayExpress:E-MTAB-3503 |
| ENSMUSG00000048482 | upregulates | ENSMUSG00000061762 | manual:Lewis |
| ENSMUSG00000024238 | upregulates | ENSMUSG00000035158 | TRRUST:19515996 |
| ENSMUSG00000044786 | downregulates | ENSMUSG00000027358 | ArrayExpress:E-GEOD-5324 |
| ENSMUSG00000004043 | downregulates | ENSMUSG00000027398 | ArrayExpress:E-GEOD-37646 |
| ENSMUSG00000025958 | upregulates | ENSMUSG00000035158 | TRRUST:19937254;TRRUST:17083486 |
| ENSMUSG00000001280 | upregulates | ENSMUSG00000029249 | TRRUST:21179468 |
| ENSMUSG00000001280 | upregulates | ENSMUSG00000038418 | TRRUST:8349585 |
| ENSMUSG00000065619 | downregulates | ENSMUSG00000026872 | miRTarBase:24245745 |
| ENSMUSG00000065400 | downregulates | ENSMUSG00000024238 | miRTarBase:18376396;miRTarBase:18411277;miRTarBase:19703993;miRTarBase:18381893;miRTarBase:19854497;miRTarBase:18829540;miRTarBase:21049046;miRTarBase:21389093;miRTarBase:21874049;miRTarBase:23381389;miRTarBase:24127332 |
| ENSMUSG00000023067 | downregulates | ENSMUSG00000024401 | manual:Lewis |
| ENSMUSG00000046733 | downregulates | ENSMUSG00000023067 | ArrayExpress:E-GEOD-21309 |
| ENSMUSG00000065408 | downregulates | ENSMUSG00000018983 | miRTarBase:27174918;miRTarBase:26997445 |
| ENSMUSG00000028163 | upregulates | ENSMUSG00000027358 | TRRUST:16835229 |
| ENSMUSG00000017950 | upregulates | ENSMUSG00000017950 | ORegAnno:N/A |
| ENSMUSG00000065493 | downregulates | ENSMUSG00000022508 | miRTarBase:23047694 |
| ENSMUSG00000029687 | downregulates | ENSMUSG00000022508 | ArrayExpress:E-GEOD-34961 |
| ENSMUSG00000044786 | upregulates | ENSMUSG00000024349 | ArrayExpress:E-GEOD-5324 |
| ENSMUSG00000029249 | downregulates | ENSMUSG00000061762 | TRRUST:12220737;19246391 |
| ENSMUSG00000052684 | downregulates | ENSMUSG00000065400 | TransmiR:27550813 |
| ENSMUSG00000065397 | downregulates | ENSMUSG00000030592 | ArrayExpress:E-GEOD-68511 |
| ENSMUSG00000024401 | downregulates | ENSMUSG00000001280 | manual:Lewis |
| ENSMUSG00000065474 | downregulates | ENSMUSG00000004043 | miRTarBase:25241899 |
| ENSMUSG00000005442 | upregulates | ENSMUSG00000023232 | ArrayExpress:E-MTAB-6681 |
| ENSMUSG00000020919 | upregulates | ENSMUSG00000021194 | ArrayExpress:E-GEOD-37646 |
| ENSMUSG00000029687 | downregulates | ENSMUSG00000022676 | TRRUST:23836662 |
| ENSMUSG00000065586 | downregulates | ENSMUSG00000024238 | miRTarBase:24277930 |
| ENSMUSG00000044167 | upregulates | ENSMUSG00000005148 | ArrayExpress:E-MTAB-4023 |
| ENSMUSG00000076361 | downregulates | ENSMUSG00000048482 | miRTarBase:23704927;miRTarBase:25955435 |
| ENSMUSG00000065493 | downregulates | ENSMUSG00000059552 | miRTarBase:26406332;miRTarBase:26177460 |
| ENSMUSG00000046733 | upregulates | ENSMUSG00000046532 | ArrayExpress:E-GEOD-21309 |
| ENSMUSG00000065586 | downregulates | ENSMUSG00000044167 | manual:Lewis;miRTarBase:19574223;miRTarBase:20028871;miRTarBase:26617698;miRTarBase:25369914 |
| ENSMUSG00000000782 | downregulates | ENSMUSG00000026872 | ArrayExpress:E-GEOD-54976 |
| ENSMUSG00000022508 | upregulates | ENSMUSG00000006731 | ArrayExpress:E-GEOD-58365 |
| ENSMUSG00000022676 | downregulates | ENSMUSG00000065474 | TransmiR:23447531 |
| ENSMUSG00000024406 | upregulates | ENSMUSG00000012396 | TRRUST:17068183;TRRUST:18687992;22745796 |
| ENSMUSG00000001280 | upregulates | ENSMUSG00000023067 | manual:Lewis;ORegAnno:N/A;TRRUST:10878024;12434298;12665570;14527674;15033443;15041706;15961274;17026958;TRRUST:8995437 |
| ENSMUSG00000055471 | upregulates | ENSMUSG00000052684 | manual:Lewis |
| ENSMUSG00000029249 | upregulates | ENSMUSG00000010066 | ORegAnno:N/A |
| ENSMUSG00000029249 | upregulates | ENSMUSG00000056812 | ORegAnno:N/A |
| ENSMUSG00000048756 | upregulates | ENSMUSG00000006818 | manual:Lewis |
| ENSMUSG00000018983 | upregulates | ENSMUSG00000006731 | ArrayExpress:E-GEOD-52157 |
| ENSMUSG00000059552 | downregulates | ENSMUSG00000065471 | TransmiR:26886852 |
| ENSMUSG00000051159 | downregulates | ENSMUSG00000004043 | ArrayExpress:E-MEXP-3202 |
| ENSMUSG00000076361 | downregulates | ENSMUSG00000024515 | miRTarBase:23226455 |
| ENSMUSG00000005672 | upregulates | ENSMUSG00000024401 | ArrayExpress:E-MEXP-2282 |
| ENSMUSG00000024401 | upregulates | ENSMUSG00000029687 | manual:Lewis |
| ENSMUSG00000048756 | upregulates | ENSMUSG00000003068 | ORegAnno:22412893 |
| ENSMUSG00000065586 | downregulates | ENSMUSG00000027323 | manual:Lewis;miRTarBase:22761336 |
| ENSMUSG00000024406 | downregulates | ENSMUSG00000065400 | TransmiR:23386720 |
| ENSMUSG00000065586 | upregulates | ENSMUSG00000034220 | manual:Lewis |
| ENSMUSG00000105904 | upregulates | ENSMUSG00000000782 | ArrayExpress:E-MTAB-7079 |
| ENSMUSG00000022105 | downregulates | ENSMUSG00000023067 | ArrayExpress:E-GEOD-52157;TRRUST:9671466 |
| ENSMUSG00000000782 | downregulates | ENSMUSG00000056749 | ArrayExpress:E-GEOD-54976 |
| ENSMUSG00000027398 | upregulates | ENSMUSG00000047146 | ArrayExpress:E-GEOD-70301 |
| ENSMUSG00000059552 | upregulates | ENSMUSG00000006818 | manual:Lewis;TRRUST:19226370 |
| ENSMUSG00000005442 | upregulates | ENSMUSG00000007872 | ArrayExpress:E-MTAB-6681 |
| ENSMUSG00000044167 | downregulates | ENSMUSG00000046532 | TRRUST:19074551 |
| ENSMUSG00000027358 | upregulates | ENSMUSG00000065408 | TransmiR:19285018 |
| ENSMUSG00000021250 | upregulates | ENSMUSG00000029015 | manual:Lewis |
| ENSMUSG00000022508 | upregulates | ENSMUSG00000027398 | ArrayExpress:E-GEOD-58365 |
| ENSMUSG00000028163 | upregulates | ENSMUSG00000024401 | TRRUST:10079106;10748079;11749970;12058956;12686724;15013838;16707469;16916598;17407192;18057724;20066113;7876168;8219230;8679226;TRRUST:10437651;10754326;11179036;11243852;11259437;11454276;12168659;15548379;15901350;15938622;18166486;18557935;20351193;8622636;9270030 |
| ENSMUSG00000065474 | downregulates | ENSMUSG00000005148 | miRTarBase:15131085 |
| ENSMUSG00000002603 | downregulates | ENSMUSG00000065474 | TransmiR:18698484 |
| ENSMUSG00000044786 | downregulates | ENSMUSG00000028163 | TRRUST:19654331 |
| ENSMUSG00000020919 | downregulates | ENSMUSG00000065592 | ArrayExpress:E-GEOD-37646 |
| ENSMUSG00000026739 | downregulates | ENSMUSG00000065408 | TransmiR:25737447 |
| ENSMUSG00000004043 | upregulates | ENSMUSG00000072825 | ArrayExpress:E-GEOD-37646 |
| ENSMUSG00000019982 | upregulates | ENSMUSG00000005672 | TRRUST:9416832 |
| ENSMUSG00000018983 | upregulates | ENSMUSG00000007872 | ArrayExpress:E-GEOD-52157 |
| ENSMUSG00000061762 | upregulates | ENSMUSG00000021250 | manual:Lewis |
| ENSMUSG00000046532 | upregulates | ENSMUSG00000024401 | TRRUST:19907077 |
| ENSMUSG00000044786 | downregulates | ENSMUSG00000029348 | ArrayExpress:E-GEOD-5324 |
| ENSMUSG00000000782 | downregulates | ENSMUSG00000024349 | ArrayExpress:E-GEOD-54976 |
| ENSMUSG00000018983 | upregulates | ENSMUSG00000024349 | ArrayExpress:E-GEOD-52157 |
| ENSMUSG00000034701 | downregulates | ENSMUSG00000023232 | ArrayExpress:E-GEOD-35396 |
| ENSMUSG00000065619 | downregulates | ENSMUSG00000041439 | manual:current |
| ENSMUSG00000034957 | upregulates | ENSMUSG00000017950 | manual:Lewis |
| ENSMUSG00000024401 | upregulates | ENSMUSG00000065408 | TransmiR:19949084 |
| ENSMUSG00000027323 | upregulates | ENSMUSG00000001280 | TRRUST:18769160 |
| ENSMUSG00000020919 | upregulates | ENSMUSG00000072825 | ArrayExpress:E-GEOD-37646 |
| ENSMUSG00000055024 | downregulates | ENSMUSG00000065474 | TransmiR:22384255 |
| ENSMUSG00000055980 | upregulates | ENSMUSG00000034957 | manual:Lewis |
| ENSMUSG00000065619 | downregulates | ENSMUSG00000022676 | miRTarBase:24277930 |
| ENSMUSG00000046733 | upregulates | ENSMUSG00000040270 | ArrayExpress:E-GEOD-21309 |
| ENSMUSG00000065619 | downregulates | ENSMUSG00000030307 | manual:current |
| ENSMUSG00000024431 | upregulates | ENSMUSG00000034957 | manual:Lewis |
| ENSMUSG00000075316 | downregulates | ENSMUSG00000024349 | ArrayExpress:E-GEOD-61373 |
| ENSMUSG00000021194 | downregulates | ENSMUSG00000021250 | ArrayExpress:E-GEOD-18304 |
| ENSMUSG00000020919 | upregulates | ENSMUSG00000034773 | ArrayExpress:E-GEOD-37646 |
| ENSMUSG00000038418 | upregulates | ENSMUSG00000023067 | TRRUST:12690110 |
| ENSMUSG00000065619 | downregulates | ENSMUSG00000038418 | miRTarBase:21118966 |
| ENSMUSG00000040270 | upregulates | ENSMUSG00000041642 | ArrayExpress:E-GEOD-30883 |
| ENSMUSG00000029687 | downregulates | ENSMUSG00000026739 | ArrayExpress:E-GEOD-34959;ArrayExpress:E-GEOD-34961;ArrayExpress:E-GEOD-74538 |
| ENSMUSG00000001280 | upregulates | ENSMUSG00000002603 | TRRUST:9580699 |
| ENSMUSG00000032855 | downregulates | ENSMUSG00000004043 | ArrayExpress:E-TABM-411 |
| ENSMUSG00000000782 | downregulates | ENSMUSG00000034957 | ArrayExpress:E-GEOD-54976 |
| ENSMUSG00000022676 | downregulates | ENSMUSG00000065592 | TransmiR:25249558 |
| ENSMUSG00000065397 | downregulates | ENSMUSG00000032402 | miRTarBase:21036908 |
| ENSMUSG00000003068 | downregulates | ENSMUSG00000029348 | ArrayExpress:E-MTAB-2791 |
| ENSMUSG00000056749 | downregulates | ENSMUSG00000027398 | TRRUST:8547328 |
| ENSMUSG00000022508 | downregulates | ENSMUSG00000000782 | ArrayExpress:E-GEOD-58365 |
| ENSMUSG00000076361 | downregulates | ENSMUSG00000025958 | miRTarBase:22325466 |
| ENSMUSG00000015605 | downregulates | ENSMUSG00000025958 | TRRUST:1531519 |
| ENSMUSG00000051159 | downregulates | ENSMUSG00000017950 | ArrayExpress:E-MEXP-3202 |
| ENSMUSG00000021194 | upregulates | ENSMUSG00000038418 | ArrayExpress:E-GEOD-18305 |
| ENSMUSG00000052684 | upregulates | ENSMUSG00000002603 | TRRUST:17270292 |
| ENSMUSG00000065397 | downregulates | ENSMUSG00000052684 | miRTarBase:22508041 |
| ENSMUSG00000027358 | upregulates | ENSMUSG00000105904 | TransmiR:19285018 |
| ENSMUSG00000005672 | downregulates | ENSMUSG00000024431 | ArrayExpress:E-MEXP-2282 |
| ENSMUSG00000034220 | downregulates | ENSMUSG00000022105 | manual:Lewis |
| ENSMUSG00000038418 | upregulates | ENSMUSG00000059552 | TRRUST:9242687;TRRUST:15958557 |
| ENSMUSG00000015605 | downregulates | ENSMUSG00000038418 | TRRUST:16456537 |
| ENSMUSG00000065587 | downregulates | ENSMUSG00000017950 | miRTarBase:22232426;miRTarBase:23298640 |
| ENSMUSG00000065619 | downregulates | ENSMUSG00000024515 | miRTarBase:23538390 |
| ENSMUSG00000059552 | upregulates | ENSMUSG00000059552 | TRRUST:22532570 |
| ENSMUSG00000052684 | upregulates | ENSMUSG00000052684 | TRRUST:9502786;TRRUST:9252389 |
| ENSMUSG00000028163 | upregulates | ENSMUSG00000019982 | TRRUST:10602492;9362453 |
| ENSMUSG00000001280 | upregulates | ENSMUSG00000059552 | ORegAnno:N/A |
| ENSMUSG00000028163 | upregulates | ENSMUSG00000021250 | TRRUST:19026727 |
| ENSMUSG00000005672 | upregulates | ENSMUSG00000021250 | ArrayExpress:E-MEXP-2282 |
| ENSMUSG00000007872 | downregulates | ENSMUSG00000029348 | ArrayExpress:E-GEOD-50933 |
| ENSMUSG00000038418 | downregulates | ENSMUSG00000019982 | TRRUST:7559553 |
| ENSMUSG00000047146 | downregulates | ENSMUSG00000034957 | ArrayExpress:E-MTAB-4578 |
| ENSMUSG00000001280 | upregulates | ENSMUSG00000003068 | ORegAnno:22412893 |
| ENSMUSG00000055980 | upregulates | ENSMUSG00000025958 | manual:Lewis |
| ENSMUSG00000044167 | upregulates | ENSMUSG00000052684 | manual:Lewis |
| ENSMUSG00000065400 | downregulates | ENSMUSG00000059552 | miRTarBase:23144891 |
| ENSMUSG00000046733 | downregulates | ENSMUSG00000029913 | ArrayExpress:E-GEOD-21309 |
| ENSMUSG00000044786 | upregulates | ENSMUSG00000023067 | ArrayExpress:E-GEOD-5324 |
| ENSMUSG00000032855 | downregulates | ENSMUSG00000027463 | ArrayExpress:E-TABM-411 |
| ENSMUSG00000032855 | downregulates | ENSMUSG00000006731 | ArrayExpress:E-TABM-411 |
| ENSMUSG00000065471 | downregulates | ENSMUSG00000059552 | miRTarBase:21226887 |
| ENSMUSG00000024515 | downregulates | ENSMUSG00000065586 | TransmiR:24375867 |
| ENSMUSG00000052684 | downregulates | ENSMUSG00000046532 | TRRUST:20148354 |
| ENSMUSG00000065400 | downregulates | ENSMUSG00000026872 | miRTarBase:18411277;miRTarBase:19703993;miRTarBase:18381893;miRTarBase:18376396;miRTarBase:19854497;miRTarBase:18829540;miRTarBase:19931509;miRTarBase:21049046;miRTarBase:21389093;miRTarBase:21529905;miRTarBase:23386720;miRTarBase:23381389;miRTarBase:24518294;miRTarBase:22012804;miRTarBase:24127332;miRTarBase:20826811;miRTarBase:24245745 |
| ENSMUSG00000076361 | downregulates | ENSMUSG00000044167 | miRTarBase:19574223 |
| ENSMUSG00000059552 | upregulates | ENSMUSG00000001280 | manual:Lewis |
| ENSMUSG00000012396 | upregulates | ENSMUSG00000024406 | TRRUST:22378194 |
| ENSMUSG00000044167 | upregulates | ENSMUSG00000039910 | ArrayExpress:E-MTAB-4023 |
| ENSMUSG00000000782 | upregulates | ENSMUSG00000047146 | ArrayExpress:E-GEOD-33292 |
| ENSMUSG00000038418 | upregulates | ENSMUSG00000021194 | TRRUST:12456801 |
| ENSMUSG00000029687 | downregulates | ENSMUSG00000030592 | ArrayExpress:E-GEOD-34961 |
| ENSMUSG00000065422 | downregulates | ENSMUSG00000026872 | miRTarBase:20516212 |
| ENSMUSG00000065619 | downregulates | ENSMUSG00000072825 | manual:current |
| ENSMUSG00000024431 | downregulates | ENSMUSG00000027104 | TRRUST:17016446 |
| ENSMUSG00000038418 | downregulates | ENSMUSG00000046532 | TRRUST:15517885 |
| ENSMUSG00000029249 | upregulates | ENSMUSG00000030592 | ORegAnno:N/A |
| ENSMUSG00000065408 | downregulates | ENSMUSG00000024515 | miRTarBase:26494556 |
| ENSMUSG00000002603 | upregulates | ENSMUSG00000032402 | manual:Lewis |
| ENSMUSG00000038418 | upregulates | ENSMUSG00000055024 | TRRUST:18798221;20414733 |
| ENSMUSG00000065586 | downregulates | ENSMUSG00000010066 | manual:current |
| ENSMUSG00000052684 | upregulates | ENSMUSG00000006731 | TRRUST:24102378 |
| ENSMUSG00000017146 | upregulates | ENSMUSG00000055024 | TRRUST:11782371 |
| ENSMUSG00000001280 | downregulates | ENSMUSG00000012396 | TRRUST:16541131 |
| ENSMUSG00000002603 | downregulates | ENSMUSG00000024401 | manual:Lewis |
| ENSMUSG00000032402 | upregulates | ENSMUSG00000023067 | TRRUST:10878024 |
| ENSMUSG00000025997 | upregulates | ENSMUSG00000029618 | manual:Lewis |
| ENSMUSG00000009406 | upregulates | ENSMUSG00000041642 | ArrayExpress:E-GEOD-21546 |
| ENSMUSG00000029249 | upregulates | ENSMUSG00000021194 | ORegAnno:N/A |
| ENSMUSG00000046532 | downregulates | ENSMUSG00000029249 | TRRUST:24163104 |
| ENSMUSG00000020919 | upregulates | ENSMUSG00000002603 | ArrayExpress:E-GEOD-37646 |
| ENSMUSG00000024431 | downregulates | ENSMUSG00000022105 | manual:Lewis |
| ENSMUSG00000028163 | upregulates | ENSMUSG00000065400 | TransmiR:23565224 |
| ENSMUSG00000005672 | downregulates | ENSMUSG00000041439 | ArrayExpress:E-MEXP-2282 |
| ENSMUSG00000029249 | upregulates | ENSMUSG00000055471 | ORegAnno:N/A |
| ENSMUSG00000005672 | downregulates | ENSMUSG00000032402 | ArrayExpress:E-MEXP-2282 |
| ENSMUSG00000022508 | downregulates | ENSMUSG00000028163 | TRRUST:15611242 |
| ENSMUSG00000004043 | upregulates | ENSMUSG00000056749 | ArrayExpress:E-GEOD-37646 |
| ENSMUSG00000065592 | downregulates | ENSMUSG00000055980 | miRTarBase:17827156;miRTarBase:19391107;miRTarBase:22431718;miRTarBase:23201159;miRTarBase:24690171;miRTarBase:19439999;miRTarBase:23714355;miRTarBase:25477374;miRTarBase:18064649 |
| ENSMUSG00000065397 | downregulates | ENSMUSG00000027323 | miRTarBase:24616504 |
| ENSMUSG00000035158 | downregulates | ENSMUSG00000065422 | TransmiR:26236014 |
| ENSMUSG00000059552 | upregulates | ENSMUSG00000023067 | manual:Lewis;TRRUST:10101800;10327058;10499591;10522058;10566557;10597242;11444825;11892838;12665570;12889596;12902982;14522900;15851483;15882981;15897882;17280505;18172861;18645021;19501055;19713748;20673369;21084835;21245169;21557991;22532570;22732402;7753560;9916993;TRRUST:10566557;10749849;11146550;11892838;12388558;12942774;14635196;14759370;18445702;18645021;20696891;21245169;22253232;22499945;22532570;8616838;9207798 |
| ENSMUSG00000030592 | upregulates | ENSMUSG00000026407 | ArrayExpress:E-MTAB-5755 |
| ENSMUSG00000003068 | downregulates | ENSMUSG00000010066 | ArrayExpress:E-MTAB-2791 |
| ENSMUSG00000032855 | downregulates | ENSMUSG00000024302 | ArrayExpress:E-TABM-411 |
| ENSMUSG00000065586 | downregulates | ENSMUSG00000035158 | manual:Lewis;miRTarBase:17597072 |
| ENSMUSG00000001280 | upregulates | ENSMUSG00000017950 | manual:Lewis |
| ENSMUSG00000065471 | downregulates | ENSMUSG00000006818 | miRTarBase:19487542 |
| ENSMUSG00000024401 | downregulates | ENSMUSG00000034957 | manual:Lewis |
| ENSMUSG00000065586 | downregulates | ENSMUSG00000059970 | manual:current;manual:Lewis |
| ENSMUSG00000018983 | downregulates | ENSMUSG00000036202 | ArrayExpress:E-GEOD-52157 |
| ENSMUSG00000017950 | upregulates | ENSMUSG00000005148 | ArrayExpress:E-MEXP-1729 |
| ENSMUSG00000024238 | downregulates | ENSMUSG00000065492 | TransmiR:22134354 |
| ENSMUSG00000028163 | downregulates | ENSMUSG00000038418 | TRRUST:21983014 |
| ENSMUSG00000034701 | downregulates | ENSMUSG00000038418 | ArrayExpress:E-GEOD-35396 |
| ENSMUSG00000030592 | downregulates | ENSMUSG00000029687 | ArrayExpress:E-MTAB-5755 |
| ENSMUSG00000065586 | downregulates | ENSMUSG00000055471 | manual:Lewis;miRTarBase:22414602 |
| ENSMUSG00000105904 | upregulates | ENSMUSG00000046532 | ArrayExpress:E-MTAB-7079 |
| ENSMUSG00000009406 | upregulates | ENSMUSG00000021250 | TRRUST:9813041 |
| ENSMUSG00000032855 | downregulates | ENSMUSG00000039910 | ArrayExpress:E-TABM-411 |
| ENSMUSG00000065586 | downregulates | ENSMUSG00000006731 | manual:current |
| ENSMUSG00000034701 | downregulates | ENSMUSG00000034957 | ArrayExpress:E-GEOD-35396 |
| ENSMUSG00000001280 | upregulates | ENSMUSG00000034957 | manual:Lewis |
| ENSMUSG00000028163 | upregulates | ENSMUSG00000028163 | TRRUST:1740105 |
| ENSMUSG00000065397 | downregulates | ENSMUSG00000048756 | ArrayExpress:E-GEOD-68511;miRTarBase:20371610;miRTarBase:21304824;miRTarBase:23970205;miRTarBase:27091996;miRTarBase:28006785;miRTarBase:28401267 |
| ENSMUSG00000026407 | downregulates | ENSMUSG00000063531 | ArrayExpress:E-MTAB-5755 |
| ENSMUSG00000030592 | upregulates | ENSMUSG00000006818 | ArrayExpress:E-MTAB-5755 |
| ENSMUSG00000065400 | downregulates | ENSMUSG00000032402 | miRTarBase:22020340 |
| ENSMUSG00000044167 | upregulates | ENSMUSG00000046733 | ArrayExpress:E-MTAB-4023 |
| ENSMUSG00000065592 | downregulates | ENSMUSG00000005148 | miRTarBase:23128394;miRTarBase:25273883;miRTarBase:24001939 |
| ENSMUSG00000024238 | upregulates | ENSMUSG00000024515 | ORegAnno:21421563 |
| ENSMUSG00000009406 | upregulates | ENSMUSG00000065493 | TransmiR:26667302 |
| ENSMUSG00000005672 | downregulates | ENSMUSG00000044167 | ArrayExpress:E-MEXP-2282 |
| ENSMUSG00000111314 | upregulates | ENSMUSG00000017950 | TRRUST:22068426 |
| ENSMUSG00000002603 | upregulates | ENSMUSG00000044786 | manual:Lewis |
| ENSMUSG00000055024 | upregulates | ENSMUSG00000023067 | TRRUST:21245169 |
| ENSMUSG00000035158 | upregulates | ENSMUSG00000023067 | TRRUST:15716956 |
| ENSMUSG00000065493 | downregulates | ENSMUSG00000048756 | miRTarBase:28321785 |
| ENSMUSG00000029249 | upregulates | ENSMUSG00000055980 | ORegAnno:N/A |
| ENSMUSG00000065592 | downregulates | ENSMUSG00000023067 | miRTarBase:20190813;miRTarBase:26540468 |
| ENSMUSG00000035158 | downregulates | ENSMUSG00000065471 | TransmiR:26236014 |
| ENSMUSG00000065592 | downregulates | ENSMUSG00000032402 | miRTarBase:24873879;miRTarBase:23632450;miRTarBase:25704091 |
| ENSMUSG00000022508 | upregulates | ENSMUSG00000072825 | ArrayExpress:E-GEOD-58365 |
| ENSMUSG00000028163 | downregulates | ENSMUSG00000061762 | manual:Lewis |
| ENSMUSG00000052684 | upregulates | ENSMUSG00000017146 | TRRUST:17434164 |
| ENSMUSG00000065492 | downregulates | ENSMUSG00000022508 | miRTarBase:23047694 |
| ENSMUSG00000000782 | downregulates | ENSMUSG00000052684 | ArrayExpress:E-GEOD-54976 |
| ENSMUSG00000065492 | downregulates | ENSMUSG00000025958 | miRTarBase:19258499;miRTarBase:19461653;miRTarBase:24816316 |
| ENSMUSG00000059552 | upregulates | ENSMUSG00000002603 | manual:Lewis |
| ENSMUSG00000030592 | downregulates | ENSMUSG00000036202 | ArrayExpress:E-MTAB-5755 |
| ENSMUSG00000015605 | upregulates | ENSMUSG00000065592 | TransmiR:19720868;TransmiR:19578358 |
| ENSMUSG00000051159 | downregulates | ENSMUSG00000024431 | ArrayExpress:E-MEXP-3202 |
| ENSMUSG00000059552 | downregulates | ENSMUSG00000029687 | TRRUST:15208672;21165554 |
| ENSMUSG00000019982 | upregulates | ENSMUSG00000065474 | TransmiR:24067373 |
| ENSMUSG00000048756 | upregulates | ENSMUSG00000022508 | TRRUST:15509806 |
| ENSMUSG00000022508 | downregulates | ENSMUSG00000070570 | ArrayExpress:E-GEOD-58365 |
| ENSMUSG00000076361 | downregulates | ENSMUSG00000035158 | miRTarBase:17597072;miRTarBase:22848417 |
| ENSMUSG00000029687 | downregulates | ENSMUSG00000059552 | TRRUST:20132185 |
| ENSMUSG00000024406 | upregulates | ENSMUSG00000065474 | TransmiR:23386720 |
| ENSMUSG00000059552 | upregulates | ENSMUSG00000048756 | TRRUST:10051583;20564353 |
| ENSMUSG00000065397 | downregulates | ENSMUSG00000035158 | miRTarBase:22771905;miRTarBase:25853464;miRTarBase:28359146 |
| ENSMUSG00000006818 | downregulates | ENSMUSG00000024749 | manual:Lewis |
| ENSMUSG00000009406 | upregulates | ENSMUSG00000000782 | ArrayExpress:E-GEOD-21546 |
| ENSMUSG00000020919 | upregulates | ENSMUSG00000056749 | ArrayExpress:E-GEOD-37646 |
| ENSMUSG00000026739 | downregulates | ENSMUSG00000012396 | TRRUST:25348805 |
| ENSMUSG00000009406 | upregulates | ENSMUSG00000025997 | ArrayExpress:E-GEOD-21546 |
| ENSMUSG00000005148 | upregulates | ENSMUSG00000038418 | TRRUST:15098654 |
| ENSMUSG00000040270 | upregulates | ENSMUSG00000021680 | ArrayExpress:E-GEOD-30883 |
| ENSMUSG00000059552 | downregulates | ENSMUSG00000032855 | TRRUST:16931520 |
| ENSMUSG00000020919 | upregulates | ENSMUSG00000052684 | ArrayExpress:E-GEOD-37646 |
| ENSMUSG00000025997 | upregulates | ENSMUSG00000029015 | manual:Lewis |
| ENSMUSG00000005672 | downregulates | ENSMUSG00000024349 | ArrayExpress:E-MEXP-2282 |
| ENSMUSG00000022676 | downregulates | ENSMUSG00000065400 | TransmiR:23447531 |
| ENSMUSG00000065586 | downregulates | ENSMUSG00000034723 | manual:current |
| ENSMUSG00000020919 | upregulates | ENSMUSG00000018983 | ArrayExpress:E-GEOD-37646 |
| ENSMUSG00000030592 | upregulates | ENSMUSG00000024302 | ArrayExpress:E-MTAB-5755 |
| ENSMUSG00000029249 | upregulates | ENSMUSG00000038457 | ORegAnno:N/A |
| ENSMUSG00000004043 | upregulates | ENSMUSG00000006731 | ArrayExpress:E-GEOD-37646 |
| ENSMUSG00000052684 | downregulates | ENSMUSG00000065474 | TransmiR:27550813 |
| ENSMUSG00000029249 | upregulates | ENSMUSG00000048482 | ORegAnno:N/A |
| ENSMUSG00000024406 | upregulates | ENSMUSG00000024406 | TRRUST:15988017 |
| ENSMUSG00000065397 | downregulates | ENSMUSG00000018983 | miRTarBase:20584899;miRTarBase:19759154;miRTarBase:26967247 |
| ENSMUSG00000020919 | downregulates | ENSMUSG00000027398 | ArrayExpress:E-GEOD-37646 |
| ENSMUSG00000024431 | downregulates | ENSMUSG00000061762 | manual:Lewis |
| ENSMUSG00000065408 | downregulates | ENSMUSG00000048756 | miRTarBase:27541266 |
| ENSMUSG00000052684 | downregulates | ENSMUSG00000021250 | TRRUST:10208434 |
| ENSMUSG00000046532 | upregulates | ENSMUSG00000076361 | TransmiR:27109471 |
| ENSMUSG00000065586 | downregulates | ENSMUSG00000029675 | manual:current |
| ENSMUSG00000046532 | downregulates | ENSMUSG00000023067 | TRRUST:16281084 |
| ENSMUSG00000044167 | upregulates | ENSMUSG00000022508 | ArrayExpress:E-MTAB-4023 |
| ENSMUSG00000044786 | upregulates | ENSMUSG00000002603 | ArrayExpress:E-GEOD-5324 |
| ENSMUSG00000029249 | upregulates | ENSMUSG00000029249 | ORegAnno:N/A |
| ENSMUSG00000024406 | downregulates | ENSMUSG00000023067 | TRRUST:19968627 |
| ENSMUSG00000005442 | upregulates | ENSMUSG00000021194 | ArrayExpress:E-MTAB-6681 |
| ENSMUSG00000039910 | upregulates | ENSMUSG00000032402 | TRRUST:16619037 |
| ENSMUSG00000039910 | upregulates | ENSMUSG00000034957 | TRRUST:18358466 |
| ENSMUSG00000004043 | upregulates | ENSMUSG00000034773 | ArrayExpress:E-GEOD-37646 |
| ENSMUSG00000044167 | upregulates | ENSMUSG00000024302 | ArrayExpress:E-MTAB-4023 |
| ENSMUSG00000022676 | downregulates | ENSMUSG00000065493 | TransmiR:26553365 |
| ENSMUSG00000065586 | downregulates | ENSMUSG00000038457 | manual:current |
| ENSMUSG00000055024 | upregulates | ENSMUSG00000004043 | TRRUST:21356387 |
| ENSMUSG00000024431 | upregulates | ENSMUSG00000017146 | TRRUST:22328717 |
| ENSMUSG00000051159 | downregulates | ENSMUSG00000059552 | ArrayExpress:E-MEXP-3202 |
| ENSMUSG00000017146 | downregulates | ENSMUSG00000065397 | TransmiR:25800897;TransmiR:27596294 |
| ENSMUSG00000065586 | downregulates | ENSMUSG00000024431 | manual:Lewis |
| ENSMUSG00000024238 | downregulates | ENSMUSG00000065587 | TransmiR:22134354 |
| ENSMUSG00000065619 | downregulates | ENSMUSG00000034723 | manual:current |
| ENSMUSG00000009406 | upregulates | ENSMUSG00000029687 | TRRUST:22222375 |
| ENSMUSG00000065586 | downregulates | ENSMUSG00000022676 | miRTarBase:24277930 |
| ENSMUSG00000001280 | upregulates | ENSMUSG00000048482 | manual:Lewis |
| ENSMUSG00000065493 | downregulates | ENSMUSG00000017950 | miRTarBase:20018894;miRTarBase:22232426;miRTarBase:23298640;miRTarBase:27135744 |
| ENSMUSG00000065619 | downregulates | ENSMUSG00000056749 | miRTarBase:24423866 |
| ENSMUSG00000065619 | downregulates | ENSMUSG00000024238 | miRTarBase:24277930 |
| ENSMUSG00000044786 | downregulates | ENSMUSG00000046733 | ArrayExpress:E-GEOD-5324 |
| ENSMUSG00000029249 | upregulates | ENSMUSG00000030307 | ORegAnno:N/A |
| ENSMUSG00000047146 | downregulates | ENSMUSG00000005148 | ArrayExpress:E-MTAB-4578 |
| ENSMUSG00000001280 | downregulates | ENSMUSG00000065474 | TransmiR:27550813 |
| ENSMUSG00000065587 | downregulates | ENSMUSG00000022508 | miRTarBase:23047694 |
| ENSMUSG00000105904 | downregulates | ENSMUSG00000025716 | ArrayExpress:E-MTAB-7079 |
| ENSMUSG00000032402 | downregulates | ENSMUSG00000036202 | TRRUST:25569105 |
| ENSMUSG00000044167 | upregulates | ENSMUSG00000059552 | manual:Lewis |
| ENSMUSG00000065586 | downregulates | ENSMUSG00000048482 | miRTarBase:25955435 |
| ENSMUSG00000044167 | upregulates | ENSMUSG00000041439 | ArrayExpress:E-MTAB-4023 |
| ENSMUSG00000027104 | upregulates | ENSMUSG00000024401 | TRRUST:10688670;10748079;10913190;20068037 |
| ENSMUSG00000047146 | downregulates | ENSMUSG00000096641 | ArrayExpress:E-MTAB-4578 |
| ENSMUSG00000020919 | upregulates | ENSMUSG00000023067 | ArrayExpress:E-GEOD-37646 |
| ENSMUSG00000017146 | upregulates | ENSMUSG00000027323 | TRRUST:10843985 |
| ENSMUSG00000046733 | downregulates | ENSMUSG00000044167 | ArrayExpress:E-GEOD-21309 |
| ENSMUSG00000065619 | downregulates | ENSMUSG00000024302 | manual:current |
| ENSMUSG00000044167 | upregulates | ENSMUSG00000056749 | ArrayExpress:E-MTAB-4023 |
| ENSMUSG00000046733 | upregulates | ENSMUSG00000027358 | ArrayExpress:E-GEOD-21309 |
| ENSMUSG00000024515 | upregulates | ENSMUSG00000007872 | TRRUST:18477564 |
| ENSMUSG00000029249 | upregulates | ENSMUSG00000017950 | ORegAnno:N/A |
| ENSMUSG00000024431 | downregulates | ENSMUSG00000015605 | TRRUST:17016446 |
| ENSMUSG00000015605 | upregulates | ENSMUSG00000021250 | manual:Lewis;TRRUST:10742173;9734357 |
| ENSMUSG00000024515 | upregulates | ENSMUSG00000111314 | TRRUST:12270938 |
| ENSMUSG00000046733 | downregulates | ENSMUSG00000034957 | ArrayExpress:E-GEOD-21309 |
| ENSMUSG00000051159 | downregulates | ENSMUSG00000002603 | ArrayExpress:E-MEXP-3202 |
| ENSMUSG00000076361 | downregulates | ENSMUSG00000023067 | miRTarBase:20190813 |
| ENSMUSG00000032855 | upregulates | ENSMUSG00000100980 | ArrayExpress:E-TABM-411 |
| ENSMUSG00000032855 | downregulates | ENSMUSG00000038418 | ArrayExpress:E-TABM-411 |
| ENSMUSG00000076361 | downregulates | ENSMUSG00000048756 | miRTarBase:19188590;miRTarBase:23262295;miRTarBase:22582938 |
| ENSMUSG00000024431 | upregulates | ENSMUSG00000023067 | manual:Lewis;TRRUST:17989362 |
| ENSMUSG00000018983 | upregulates | ENSMUSG00000052684 | ArrayExpress:E-GEOD-52157 |
| ENSMUSG00000017950 | upregulates | ENSMUSG00000024517 | ArrayExpress:E-GEOD-11759 |
| ENSMUSG00000024238 | downregulates | ENSMUSG00000076361 | TransmiR:24277930 |
| ENSMUSG00000024406 | upregulates | ENSMUSG00000036202 | TRRUST:25569105 |
| ENSMUSG00000065492 | downregulates | ENSMUSG00000017950 | miRTarBase:22232426 |
| ENSMUSG00000020919 | upregulates | ENSMUSG00000034220 | ArrayExpress:E-GEOD-37646 |
| ENSMUSG00000026739 | downregulates | ENSMUSG00000065474 | TransmiR:27105531 |
| ENSMUSG00000076361 | downregulates | ENSMUSG00000020919 | miRTarBase:25231457 |
| ENSMUSG00000024431 | downregulates | ENSMUSG00000009406 | TRRUST:17016446 |
| ENSMUSG00000065586 | downregulates | ENSMUSG00000056812 | manual:current;manual:Lewis |
| ENSMUSG00000065422 | downregulates | ENSMUSG00000027323 | miRTarBase:24036252 |
| ENSMUSG00000065422 | downregulates | ENSMUSG00000048756 | miRTarBase:20388878 |
| ENSMUSG00000059552 | downregulates | ENSMUSG00000012396 | TRRUST:15619621;24126347 |
| ENSMUSG00000065586 | downregulates | ENSMUSG00000048756 | manual:Lewis;miRTarBase:21203424;miRTarBase:25369914 |
| ENSMUSG00000005672 | upregulates | ENSMUSG00000036202 | ArrayExpress:E-MEXP-2282 |
| ENSMUSG00000065586 | downregulates | ENSMUSG00000023067 | miRTarBase:19422085;miRTarBase:26582573 |
| ENSMUSG00000065471 | downregulates | ENSMUSG00000044167 | miRTarBase:27746366 |
| ENSMUSG00000018983 | upregulates | ENSMUSG00000044786 | ArrayExpress:E-GEOD-52157 |
| ENSMUSG00000021250 | upregulates | ENSMUSG00000065408 | TransmiR:27926494 |
| ENSMUSG00000009406 | upregulates | ENSMUSG00000065408 | TransmiR:26747707 |
| ENSMUSG00000065474 | downregulates | ENSMUSG00000026872 | miRTarBase:19830559;miRTarBase:18411277;miRTarBase:19854497;miRTarBase:24245745;miRTarBase:25502084;miRTarBase:25975736;miRTarBase:21049046 |
| ENSMUSG00000004043 | upregulates | ENSMUSG00000017950 | ArrayExpress:E-GEOD-37646 |
| ENSMUSG00000051159 | upregulates | ENSMUSG00000034701 | ArrayExpress:E-MEXP-3202 |
| ENSMUSG00000038418 | upregulates | ENSMUSG00000007872 | TRRUST:17878368 |
| ENSMUSG00000059552 | downregulates | ENSMUSG00000021250 | TRRUST:8604347 |
| ENSMUSG00000065586 | downregulates | ENSMUSG00000072825 | manual:current |
| ENSMUSG00000005672 | downregulates | ENSMUSG00000039910 | ArrayExpress:E-MEXP-2282 |
| ENSMUSG00000017950 | upregulates | ENSMUSG00000034957 | manual:Lewis;TRRUST:15155787 |
| ENSMUSG00000065471 | downregulates | ENSMUSG00000048756 | miRTarBase:20388878 |
| ENSMUSG00000001280 | downregulates | ENSMUSG00000024401 | TRRUST:12759366 |
| ENSMUSG00000065586 | downregulates | ENSMUSG00000055980 | manual:Lewis |
| ENSMUSG00000019982 | downregulates | ENSMUSG00000023067 | TRRUST:8616838 |

***Mir183/96*** $^{dko}$ **known and predicted misregulation**

| Gene ID | fold change | status |
|---|---|---|
| ENSMUSG00000065400 | -1 | predicted |
| ENSMUSG00000056749 | 1 | predicted |
| ENSMUSG00000044786 | 1 | predicted |
| ENSMUSG00000021194 | 1 | predicted |
| ENSMUSG00000023067 | 1 | predicted |
| ENSMUSG00000025716 | 1.009585927 | known |
| ENSMUSG00000065422 | -1 | predicted |
| ENSMUSG00000027358 | -1 | predicted |
| ENSMUSG00000065587 | -1 | predicted |
| ENSMUSG00000015605 | -1 | predicted |
| ENSMUSG00000024515 | 1 | predicted |
| ENSMUSG00000047146 | -1 | predicted |
| ENSMUSG00000063531 | -1.141470271 | known |
| ENSMUSG00000024401 | -1 | predicted |
| ENSMUSG00000065592 | -1 | predicted |
| ENSMUSG00000065474 | -1 | predicted |
| ENSMUSG00000052684 | 1 | predicted |
| ENSMUSG00000032402 | 1 | predicted |
| ENSMUSG00000019982 | -1 | predicted |
| ENSMUSG00000024749 | -1.06239806 | known |
| ENSMUSG00000012396 | -1 | predicted |
| ENSMUSG00000070570 | -0.956503416 | known |
| ENSMUSG00000002603 | 1 | predicted |
| ENSMUSG00000026739 | 1 | predicted |
| ENSMUSG00000007872 | 1 | predicted |
| ENSMUSG00000065397 | -1 | predicted |
| ENSMUSG00000010066 | 0.547452704 | known |
| ENSMUSG00000032855 | -1 | predicted |
| ENSMUSG00000005672 | -1 | predicted |
| ENSMUSG00000017950 | 1 | predicted |
| ENSMUSG00000004043 | 1 | predicted |
| ENSMUSG00000034220 | 1 | predicted |
| ENSMUSG00000035158 | 1 | predicted |
| ENSMUSG00000034701 | -1 | predicted |
| ENSMUSG00000025997 | -1 | predicted |
| ENSMUSG00000006731 | 1.002457614 | known |
| ENSMUSG00000065586 | -100 | known |
| ENSMUSG00000020919 | 1 | predicted |
| ENSMUSG00000096641 | 2.32961413 | known |
| ENSMUSG00000065492 | -1 | predicted |
| ENSMUSG00000028163 | -1 | predicted |
| ENSMUSG00000105904 | -1 | predicted |
| ENSMUSG00000055024 | 1 | predicted |
| ENSMUSG00000005442 | 1 | predicted |
| ENSMUSG00000034773 | 1.502977144 | known |
| ENSMUSG00000029618 | -1.480031146 | known |
| ENSMUSG00000029348 | -1.401119206 | known |
| ENSMUSG00000065408 | -1 | predicted |
| ENSMUSG00000076361 | -1 | predicted |
| ENSMUSG00000029675 | 0.753649651 | known |
| ENSMUSG00000024349 | 1.67268314 | known |
| ENSMUSG00000024406 | -1 | predicted |
| ENSMUSG00000075316 | -1.716845583 | known |
| ENSMUSG00000048482 | 1 | predicted |
| ENSMUSG00000001280 | 1 | predicted |
| ENSMUSG00000072825 | 0.648798422 | known |
| ENSMUSG00000055471 | 1 | predicted |
| ENSMUSG00000029015 | -1.462827882 | known |
| ENSMUSG00000111314 | 1 | predicted |
| ENSMUSG00000065471 | -1 | predicted |
| ENSMUSG00000024238 | 1 | predicted |
| ENSMUSG00000027323 | 1 | predicted |
| ENSMUSG00000034723 | 0.581830334 | known |
| ENSMUSG00000034957 | 1 | predicted |
| ENSMUSG00000048756 | 1 | predicted |
| ENSMUSG00000021250 | -1 | predicted |
| ENSMUSG00000039910 | 1 | predicted |
| ENSMUSG00000024431 | 1 | predicted |
| ENSMUSG00000036202 | -1 | predicted |
| ENSMUSG00000026407 | 1 | predicted |
| ENSMUSG00000038418 | 1 | predicted |
| ENSMUSG00000059552 | 1 | predicted |
| ENSMUSG00000065493 | -1 | predicted |
| ENSMUSG00000038457 | 1.174878595 | known |
| ENSMUSG00000061762 | 0 | predicted |
| ENSMUSG00000005148 | 1 | predicted |
| ENSMUSG00000000782 | -1 | predicted |
| ENSMUSG00000100980 | -6.172404621 | known |
| ENSMUSG00000027398 | -1 | predicted |
| ENSMUSG00000040270 | -1 | predicted |
| ENSMUSG00000024517 | 1.894505585 | known |
| ENSMUSG00000023232 | 0.752432376 | known |
| ENSMUSG00000056812 | 0.713975665 | known |
| ENSMUSG00000025958 | 1 | predicted |
| ENSMUSG00000006818 | 1 | predicted |
| ENSMUSG00000009406 | 0 | predicted |
| ENSMUSG00000024479 | 0.650542374 | known |
| ENSMUSG00000026872 | 1 | predicted |
| ENSMUSG00000029913 | -1 | predicted |
| ENSMUSG00000030307 | 1.408440608 | known |
| ENSMUSG00000027104 | -1 | predicted |
| ENSMUSG00000046532 | -1 | predicted |
| ENSMUSG00000041439 | 0.604452755 | known |
| ENSMUSG00000044167 | 1 | predicted |
| ENSMUSG00000065619 | -100 | known |
| ENSMUSG00000017146 | 1 | predicted |
| ENSMUSG00000022676 | 1 | predicted |
| ENSMUSG00000024302 | 0.565905364 | known |
| ENSMUSG00000003068 | 1 | predicted |
| ENSMUSG00000022508 | 1 | predicted |
| ENSMUSG00000022105 | -1 | predicted |
| ENSMUSG00000018983 | 1 | predicted |
| ENSMUSG00000059970 | 1.219380605 | known |
| ENSMUSG00000041642 | -0.727605962 | known |
| ENSMUSG00000031450 | 1.662061997 | known |
| ENSMUSG00000030592 | 1 | predicted |
| ENSMUSG00000029687 | -1 | predicted |
| ENSMUSG00000051159 | -1 | predicted |
| ENSMUSG00000046733 | -1 | predicted |
| ENSMUSG00000029249 | 1 | predicted |
| ENSMUSG00000055980 | 1 | predicted |
| ENSMUSG00000021680 | -1.459858321 | known |
| ENSMUSG00000027463 | 1.661603767 | known |

**Network links and references for Fig. 7B**

***Mir182*** $^{ko}$ **network**

| Gene 1 | Interaction | Gene 2 | Reference |
|---|---|---|---|
| ENSMUSG00000017950 | upregulates | ENSMUSG00000024517 | ArrayExpress:E-GEOD-11759 |
| ENSMUSG00000111314 | upregulates | ENSMUSG00000017950 | TRRUST:22068426 |
| ENSMUSG00000059552 | downregulates | ENSMUSG00000017950 | TRRUST:16895524 |
| ENSMUSG00000065445 | downregulates | ENSMUSG00000017950 | miRTarBase:23298640 |
| ENSMUSG00000059552 | upregulates | ENSMUSG00000059552 | TRRUST:22532570 |
| ENSMUSG00000022676 | downregulates | ENSMUSG00000065493 | TransmiR:26553365 |
| ENSMUSG00000076361 | downregulates | ENSMUSG00000027784 | manual:current |
| ENSMUSG00000025958 | downregulates | ENSMUSG00000042821 | TRRUST:15955695 |
| ENSMUSG00000042821 | upregulates | ENSMUSG00000042821 | ORegAnno:16617148 |
| ENSMUSG00000002603 | upregulates | ENSMUSG00000065445 | TransmiR:21673106 |
| ENSMUSG00000048756 | downregulates | ENSMUSG00000059552 | manual:Lewis |
| ENSMUSG00000020919 | downregulates | ENSMUSG00000065445 | ArrayExpress:E-GEOD-37646 |
| ENSMUSG00000065493 | downregulates | ENSMUSG00000048756 | miRTarBase:28321785 |
| ENSMUSG00000076361 | downregulates | ENSMUSG00000025958 | miRTarBase:22325466 |
| ENSMUSG00000076361 | downregulates | ENSMUSG00000048756 | miRTarBase:19188590;miRTarBase:23262295;miRTarBase:22582938 |
| ENSMUSG00000059552 | downregulates | ENSMUSG00000024515 | ORegAnno:21421563 |
| ENSMUSG00000025958 | upregulates | ENSMUSG00000035158 | TRRUST:19937254;TRRUST:17083486 |
| ENSMUSG00000076361 | downregulates | ENSMUSG00000035158 | miRTarBase:17597072;miRTarBase:22848417 |
| ENSMUSG00000065493 | downregulates | ENSMUSG00000017950 | miRTarBase:20018894;miRTarBase:22232426;miRTarBase:23298640;miRTarBase:27135744 |
| ENSMUSG00000035158 | downregulates | ENSMUSG00000059552 | manual:Lewis |
| ENSMUSG00000042821 | downregulates | ENSMUSG00000017950 | TRRUST:16826572 |
| ENSMUSG00000024515 | upregulates | ENSMUSG00000111314 | TRRUST:12270938 |
| ENSMUSG00000002603 | downregulates | ENSMUSG00000017950 | manual:Lewis |
| ENSMUSG00000024515 | downregulates | ENSMUSG00000002603 | TRRUST:15312519 |
| ENSMUSG00000065493 | downregulates | ENSMUSG00000024515 | miRTarBase:28348487;miRTarBase:26077733;miRTarBase:28340489 |
| ENSMUSG00000059552 | upregulates | ENSMUSG00000002603 | manual:Lewis |
| ENSMUSG00000076361 | downregulates | ENSMUSG00000022676 | miRTarBase:23354685 |
| ENSMUSG00000076361 | downregulates | ENSMUSG00000020919 | miRTarBase:25231457 |
| ENSMUSG00000076361 | downregulates | ENSMUSG00000024515 | miRTarBase:23226455 |
| ENSMUSG00000017950 | upregulates | ENSMUSG00000017950 | ORegAnno:N/A |

***Mir182*** $^{ko}$ **known and predicted misregulation**

| Gene ID | fold change | status |
|---|---|---|
| ENSMUSG00000042821 | -1 | predicted |
| ENSMUSG00000076361 | -100 | known |
| ENSMUSG00000024517 | 0.891913549 | known |
| ENSMUSG00000024515 | 1 | predicted |
| ENSMUSG00000020919 | 1 | predicted |
| ENSMUSG00000022676 | 1 | predicted |
| ENSMUSG00000048756 | 1 | predicted |
| ENSMUSG00000111314 | 1 | predicted |
| ENSMUSG00000035158 | 1 | predicted |
| ENSMUSG00000025958 | 1 | predicted |
| ENSMUSG00000065445 | -1 | predicted |
| ENSMUSG00000027784 | 0.503151937 | known |
| ENSMUSG00000065493 | -1 | predicted |
| ENSMUSG00000017950 | 1 | predicted |
| ENSMUSG00000002603 | -1 | predicted |
| ENSMUSG00000059552 | -1 | predicted |

**Synapse counts for Fig. 5**

| | Synapses/nucleus | |
|---|---|---|
| *Mir183/96* *dko* | Average | Standard deviation |
| Wildtype | 14.10555556 | 2.278123169 |
| Heterozygote | 12.46785714 | 2.385022456 |
| Homozygote | 7.87797619 | 1.619121475 |

| | Synapses/nucleus | |
|---|---|---|
| *Mir182* *ko* | Average | Standard deviation |
| Wildtype | 12.81211844 | 2.403912048 |
| Heterozygote | 12.32414021 | 2.742892053 |
| Homozygote | 11.88503401 | 1.887978121 |

| | Allele | Genotype | Ribeye puncta (red) | GluR2 puncta (green) | Both (yellow) | Number of nuclei | Synapses per nucleus | Average of paired ears |
|---|---|---|---|---|---|---|---|---|
| DMKO69.3d ear 1 | *Mir183/96* *dko* | Homozygote | | | 49 | 6 | 8.166666667 | 8.166666667 |
| DMKO83.1d ear 1 | *Mir183/96* *dko* | Homozygote | 4 | 3 | 77 | 9 | 8.555555556 | 9.333333333 |
| DMKO83.1d ear 2 | *Mir183/96* *dko* | Homozygote | 4 | 3 | 91 | 9 | 10.11111111 | |
| DMKO83.1e ear 1 | *Mir183/96* *dko* | Homozygote | 10 | 4 | 41 | 8 | 5.125 | 6.133928571 |
| DMKO83.1e ear 2 | *Mir183/96* *dko* | Homozygote | 1 | 8 | 50 | 7 | 7.142857143 | |
| DMKO71.1b ear 1 | *Mir183/96* *dko* | Heterozygote | | | 84 | 6 | 14 | 14 |
| DMKO71.5a ear 1 | *Mir183/96* *dko* | Heterozygote | | | 74 | 5.5 | 13.45454545 | 14.02727273 |
| DMKO71.5a ear 2 | *Mir183/96* *dko* | Heterozygote | | | 73 | 5 | 14.6 | |
| DMKO71.5b ear 1 | *Mir183/96* *dko* | Heterozygote | 8 | 23 | 90 | 9 | 10 | 12.55555556 |
| DMKO71.5b ear 2 | *Mir183/96* *dko* | Heterozygote | | 12 | 136 | 9 | 15.11111111 | |
| DMKO71.5e ear 1 | *Mir183/96* *dko* | Heterozygote | | | 112 | 7 | 16 | 16 |
| DMKO78.3c ear 1 | *Mir183/96* *dko* | Heterozygote | | 18 | 89 | 9 | 9.888888889 | 10.04444444 |
| DMKO78.3c ear 2 | *Mir183/96* *dko* | Heterozygote | | | 102 | 10 | 10.2 | |
| DMKO78.3d ear 1 | *Mir183/96* *dko* | Heterozygote | | | 123 | 11 | 11.18181818 | 11.27272727 |
| DMKO78.3d ear 2 | *Mir183/96* *dko* | Heterozygote | | | 125 | 11 | 11.36363636 | |
| DMKO80.1h ear 1 | *Mir183/96* *dko* | Heterozygote | 15 | 4 | 75 | 8 | 9.375 | 9.375 |
| DMKO79.1f ear 1 | *Mir183/96* *dko* | Wildtype | | | 130 | 10 | 13 | 13 |
| DMKO78.3b ear 1 | *Mir183/96* *dko* | Wildtype | | | 163 | 9 | 18.11111111 | 17.22222222 |
| DMKO78.3b ear 2 | *Mir183/96* *dko* | Wildtype | | | 147 | 9 | 16.33333333 | |
| DMKO78.3a ear 1 | *Mir183/96* *dko* | Wildtype | | | 123 | 10 | 12.3 | 12.09444444 |
| DMKO78.3a ear 2 | *Mir183/96* *dko* | Wildtype | | | 107 | 9 | 11.88888889 | |
| DMKO74.2b ear 1 | *Mir183/96* *dko* | Wildtype | | | 82 | 7 | 11.71428571 | 11.71428571 |
| DMKO74.1b ear 1 | *Mir183/96* *dko* | Wildtype | | | 85 | 7 | 12.14285714 | 12.14285714 |
| DMOM61.4d ear 1 | *Mir182* *ko* | Homozygote | | | 81 | 8 | 10.125 | 10.125 |
| DMOM63.3c ear 1 | *Mir182* *ko* | Homozygote | | | 77 | 6 | 12.83333333 | 12.83333333 |
| DMOM61.5a ear 1 | *Mir182* *ko* | Homozygote | 13 | | 62 | 6 | 10.33333333 | 10.33333333 |
| DMOM68.2d ear 1 | *Mir182* *ko* | Homozygote | | | 102 | 10 | 10.2 | 11.1 |
| DMOM68.2d ear 2 | *Mir182* *ko* | Homozygote | | | 120 | 10 | 12 | |
| DMOM68.2a ear 1 | *Mir182* *ko* | Homozygote | | | 83 | 8 | 10.375 | 10.375 |
| DMOM63.3g ear 1 | *Mir182* *ko* | Homozygote | | | 94 | 7 | 13.42857143 | 13.42857143 |
| DMOM69.3c ear 1 | *Mir182* *ko* | Homozygote | | | 140 | 9 | 15.55555556 | 15 |
| DMOM69.3c ear 2 | *Mir182* *ko* | Homozygote | | | 130 | 9 | 14.44444444 | |
| DMOM63.3h ear 1 | *Mir182* *ko* | Heterozygote | | | 194 | 16 | 12.125 | 12.125 |
| DMOM63.3e ear 1 | *Mir182* *ko* | Heterozygote | | | 110 | 7 | 15.71428571 | 15.71428571 |
| DMOM68.2e ear 1 | *Mir182* *ko* | Heterozygote | | | 115 | 10 | 11.5 | 10.75 |
| DMOM68.2e ear 2 | *Mir182* *ko* | Heterozygote | | | 100 | 10 | 10 | |
| DMOM68.2c ear 1 | *Mir182* *ko* | Heterozygote | | | 73 | 9 | 8.111111111 | 10.10555556 |
| DMOM68.2c ear 2 | *Mir182* *ko* | Heterozygote | | | 121 | 10 | 12.1 | |
| DMOM68.2b ear 1 | *Mir182* *ko* | Heterozygote | | | 115 | 12 | 9.583333333 | 9.583333333 |
| DMOM69.3a ear 1 | *Mir182* *ko* | Heterozygote | | | 141 | 9 | 15.66666667 | 15.66666667 |
| DMOM69.3b ear 1 | *Mir182* *ko* | Wildtype | | | 95 | 7.5 | 12.66666667 | 14.61111111 |
| DMOM69.3b ear 2 | *Mir182* *ko* | Wildtype | | | 149 | 9 | 16.55555556 | |
| DMOM63.3b ear 1 | *Mir182* *ko* | Wildtype | | | 83 | 7 | 11.85714286 | 11.85714286 |
| DMOM61.4e ear 1 | *Mir182* *ko* | Wildtype | 16 | | 194 | 13 | 14.92307692 | 14.92307692 |
| DMOM61.5c ear 1 | *Mir182* *ko* | Wildtype | | | 69 | 7 | 9.857142857 | 9.857142857 |

**miRNA qRTPCR Ct data for Fig. S1**

| | *Mir183/96* $^{dko}$ average | | | *Mir183/96* $^{dko}$ standard deviation | | |
|---|---|---|---|---|---|---|
| | Wildtype | Heterozygote | Homozygote | Wildtype | Heterozygote | Homozygote |
| *Mir96* | 1.008721292 | 0.437368541 | 0 | 0.014966277 | 0.286609139 | 0 |
| *Mir182* | 1.022471681 | 1.186222892 | 1.440725879 | 0.017195795 | 0.601321939 | 0.731055593 |
| *Mir183* | 1.035990137 | 0.537156813 | 0 | 0.072634857 | 0.32929358 | 0 |

| | | | Ct | | | $2^{-ddCt}$ | | |
|---|---|---|---|---|---|---|---|---|
| miRNA | Mouse | *Mir183/96* $^{dko}$ genotype | Replicate 1 | Replicate 2 | Replicate 3 | Replicate 1 | Replicate 2 | Replicate 3 |
| Mir99a | DMKO84.3b | Wildtype | 29.74237518 | 29.291019 | 29.29022928 | | | |
| Mir99a | DMKO84.3h | Wildtype | 29.59775298 | 29.43968678 | 29.82496027 | | | |
| Mir99a | DMKO76.1d | Wildtype | 25.00924881 | 26.38940752 | 25.01435294 | | | |
| Mir99a | DMKO76.1e | Wildtype | 25.37267432 | 26.10423824 | 26.84473007 | | | |
| Mir99a | DMKO84.4g | Wildtype | 30.02486014 | 30.49457728 | 30.26551617 | | | |
| Mir99a | DMKO78.5a | Wildtype | 27.38497015 | 27.43188349 | 27.5625486 | | | |
| Mir99a | DMKO79.7b | Wildtype | 28.5130984 | 29.10563508 | 28.7366004 | | | |
| Mir99a | DMKO79.7b | Wildtype | 28.5130984 | 29.10563508 | 28.7366004 | | | |
| Mir99a | DMKO84.3e | Heterozygote | 27.7224639 | 27.72476811 | 27.71423828 | | | |
| Mir99a | DMKO84.3g | Heterozygote | 27.51702191 | 27.45347557 | 27.34834558 | | | |
| Mir99a | DMKO76.1a | Heterozygote | 25.70089698 | 25.85793511 | 25.00653265 | | | |
| Mir99a | DMKO76.1b | Heterozygote | 29.43447646 | 30.32040559 | 31.19176637 | | | |
| Mir99a | DMKO84.4d | Heterozygote | 29.84599034 | 29.82320062 | 29.9325089 | | | |
| Mir99a | DMKO84.3d | Homozygote | 28.57491971 | 28.58733021 | 28.74533606 | | | |
| Mir99a | DMKO84.3c | Homozygote | 27.67639082 | 27.53602972 | 27.61928208 | | | |
| Mir99a | DMKO78.5c | Homozygote | 32.98281471 | 33.05049156 | 32.50884834 | | | |
| Mir99a | DMKO84.4e | Homozygote | 30.34765799 | 29.77097699 | 30.53077364 | | | |
| Mir99a | DMKO79.7d | Homozygote | 31.88623183 | 29.70129936 | 31.63668707 | | | |
| Mir99a | DMKO79.7h | Homozygote | 29.49172319 | 29.47046834 | 30.01761859 | | | |
| Mir96 | DMKO84.3b | Wildtype | 33.21057164 | 33.14062557 | 33.0218502 | 0.941985982 | 0.988781439 | 1.073632 |
| Mir96 | DMKO84.3h | Wildtype | 33.62750852 | 33.55325697 | | 0.97459466 | 1.026067596 | |
| Mir96 | DMKO76.1d | Wildtype | 32.03729685 | 33.06309106 | 32.80541863 | 1.513587412 | 0.743383096 | 0.88875 |
| Mir96 | DMKO76.1e | Wildtype | 32.79291131 | 33.0349808 | 32.15789824 | 0.913210153 | 0.772148004 | 1.418171 |
| Mir96 | DMKO84.4g | Wildtype | 34.30311722 | 33.45968169 | 34.04273753 | 0.774889014 | 1.390397239 | 0.928157 |
| Mir96 | DMKO78.5a | Wildtype | 35.07344291 | 35.03759366 | 34.69321798 | 0.9083427 | 0.931196704 | 1.182249 |
| Mir96 | DMKO79.7b | Wildtype | 33.10619876 | 33.24497329 | 33.32266105 | 1.085539538 | 0.985985096 | 0.934295 |
| Mir96 | DMKO79.7b | Wildtype | 33.10619876 | 33.24497329 | 33.32266105 | 1.085539538 | 0.985985096 | 0.934295 |
| Mir96 | DMKO84.3e | Heterozygote | 33.40373685 | 33.63040619 | 33.67885779 | 0.249981728 | 0.213635783 | 0.20658 |
| Mir96 | DMKO84.3g | Heterozygote | 32.29016935 | 32.22556519 | 32.35927186 | 0.543001367 | 0.567869723 | 0.517606 |
| Mir96 | DMKO76.1a | Heterozygote | 32.49556412 | 32.92027674 | 33.60407731 | 1.141151142 | 0.8501429 | 0.529234 |
| Mir96 | DMKO76.1b | Heterozygote | 39.95949127 | - | 39.13768204 | 0.117503376 | 0 | 0.207701 |
| Mir96 | DMKO84.4d | Heterozygote | 34.51875172 | 34.46186377 | 34.93543826 | 0.507686282 | 0.528105157 | 0.38033 |
| Mir96 | DMKO84.3d | Homozygote | - | - | - | 0 | 0 | 0 |
| Mir96 | DMKO84.3c | Homozygote | - | - | - | 0 | 0 | 0 |
| Mir96 | DMKO78.5c | Homozygote | - | - | - | 0 | 0 | 0 |
| Mir96 | DMKO84.4e | Homozygote | - | - | - | 0 | 0 | 0 |
| Mir96 | DMKO79.7d | Homozygote | - | - | - | 0 | 0 | 0 |
| Mir96 | DMKO79.7h | Homozygote | - | - | - | 0 | 0 | 0 |
| Mir182 | DMKO84.3b | Wildtype | 28.0436346 | 27.23968444 | 28.14118482 | 0.849410208 | 1.482963936 | 0.793875 |
| Mir182 | DMKO84.3h | Wildtype | 28.13751322 | 28.09140637 | 28.22090161 | 1.008651069 | 1.041407013 | 0.952004 |
| Mir182 | DMKO76.1d | Wildtype | 23.9935081 | 24.13197816 | 24.45239988 | 1.147998387 | 1.042935946 | 0.83522 |
| Mir182 | DMKO76.1e | Wildtype | 24.09098127 | 24.38795621 | 24.47751758 | 1.17107793 | 0.953207459 | 0.895832 |
| Mir182 | DMKO84.4g | Wildtype | 28.01480277 | 27.41879578 | 28.01641659 | 0.871679051 | 1.317566639 | 0.870705 |
| Mir182 | DMKO78.5a | Wildtype | 24.27741088 | 24.77046575 | 24.88367771 | 1.289168257 | 0.915978533 | 0.846847 |
| Mir182 | DMKO79.7b | Wildtype | 27.11010521 | 27.43910462 | 26.63023425 | 0.965741841 | 0.768815896 | 1.346842 |
| Mir182 | DMKO79.7b | Wildtype | 27.11010521 | 27.43910462 | 26.63023425 | 0.965741841 | 0.768815896 | 1.346842 |
| Mir182 | DMKO84.3e | Heterozygote | 25.89418111 | 25.84564651 | 25.77608916 | 1.143351596 | 1.182470112 | 1.240878 |
| Mir182 | DMKO84.3g | Heterozygote | 25.7404782 | 25.61718934 | 25.49924412 | 1.171434705 | 1.275944345 | 1.38464 |
| Mir182 | DMKO76.1a | Heterozygote | 23.03046719 | 23.10857936 | 23.32896341 | 2.318106237 | 2.19593372 | 1.884851 |
| Mir182 | DMKO76.1b | Heterozygote | 28.54891813 | 29.357078 | 29.55328412 | 0.985027213 | 0.56255873 | 0.491025 |
| Mir182 | DMKO84.4d | Heterozygote | 28.31748307 | 28.08045512 | 27.77003211 | 0.537661747 | 0.633667832 | 0.785793 |
| Mir182 | DMKO84.3d | Homozygote | 26.5231856 | 26.54942651 | 26.42539987 | 1.394392322 | 1.369259243 | 1.492181 |
| Mir182 | DMKO84.3c | Homozygote | 25.86313862 | 26.01039364 | 26.0399779 | 1.211310992 | 1.093773713 | 1.071573 |
| Mir182 | DMKO78.5c | Homozygote | 29.04254588 | 28.90018513 | 29.15367065 | 1.984656989 | 2.19048516 | 1.837526 |
| Mir182 | DMKO84.4e | Homozygote | 29.19845104 | 28.35978586 | 25.04459421 | 0.371913498 | 0.665128131 | 6.620297 |
| Mir182 | DMKO79.7d | Homozygote | 30.22340855 | 30.26537121 | 30.37004271 | 0.545644904 | 0.530002701 | 0.492911 |
| Mir182 | DMKO79.7h | Homozygote | 27.8262283 | 27.67899201 | 28.27453748 | 1.078040828 | 1.193871949 | 0.790098 |
| Mir183 | DMKO84.3b | Wildtype | 25.19070015 | 26.48488099 | 27.07159807 | 2.082562203 | 0.849201453 | 0.565446 |
| Mir183 | DMKO84.3h | Wildtype | 28.08181953 | 28.14559937 | 28.12998737 | 1.02620282 | 0.981823804 | 0.992506 |
| Mir183 | DMKO76.1d | Wildtype | 25.06891695 | 25.29041759 | 25.27899634 | 1.104857272 | 0.947606086 | 0.955138 |
| Mir183 | DMKO76.1e | Wildtype | 25.48675153 | 25.65985317 | 25.56899812 | 1.060773084 | 0.940835524 | 1.001991 |
| Mir183 | DMKO84.4g | Wildtype | 28.63634923 | 28.92155253 | 28.41851366 | 1.015687019 | 0.833499137 | 1.181231 |
| Mir183 | DMKO78.5a | Wildtype | 26.54837902 | 26.56938991 | 26.26604043 | 0.941407182 | 0.92779622 | 1.144906 |
| Mir183 | DMKO79.7b | Wildtype | 26.87824813 | 26.84233068 | 26.70437102 | 0.952683218 | 0.976699003 | 1.074709 |
| Mir183 | DMKO79.7b | Wildtype | 26.87824813 | 26.84233068 | 26.70437102 | 0.952683218 | 0.976699003 | 1.074709 |
| Mir183 | DMKO84.3e | Heterozygote | 27.03212162 | 27.18281454 | 27.07739787 | 0.176314178 | 0.158826932 | 0.170867 |
| Mir183 | DMKO84.3g | Heterozygote | 28.28804664 | 27.15640218 | 27.02520842 | 0.196132854 | 0.429743461 | 0.470655 |
| Mir183 | DMKO76.1a | Heterozygote | 25.24578449 | 25.27698907 | 25.44293374 | 1.012398752 | 0.990736325 | 0.883088 |
| Mir183 | DMKO76.1b | Heterozygote | 31.38238507 | 30.72538755 | 31.37660324 | 0.329378387 | 0.519363065 | 0.330701 |
| Mir183 | DMKO84.4d | Heterozygote | 28.15404887 | 29.01560827 | 28.74723038 | 1.079481384 | 0.594100522 | 0.715566 |
| Mir183 | DMKO84.3d | Homozygote | - | - | - | 0 | 0 | 0 |
| Mir183 | DMKO84.3c | Homozygote | - | - | - | 0 | 0 | 0 |
| Mir183 | DMKO78.5c | Homozygote | - | - | - | 0 | 0 | 0 |
| Mir183 | DMKO84.4e | Homozygote | - | - | - | 0 | 0 | 0 |
| Mir183 | DMKO79.7d | Homozygote | - | - | - | 0 | 0 | 0 |
| Mir183 | DMKO79.7h | Homozygote | - | - | - | 0 | 0 | 0 |

| | *Mir182* $^{ko}$ average | | | *Mir182* $^{ko}$ standard deviation | | |
|---|---|---|---|---|---|---|
| | Wildtype | Heterozygote | Homozygote | Wildtype | Heterozygote | Homozygote |
| *Mir96* | 1.100839536 | 1.55904243 | 1.575991477 | 0.160658932 | 0.421350025 | 1.416257468 |
| *Mir182* | 1.093509066 | 0.72104922 | 1.44325E-05 | 0.102108236 | 0.541635729 | 2.8865E-05 |
| *Mir183* | 1.077714791 | 1.13265617 | 1.826481537 | 0.0925687 | 0.281911635 | 0.597587505 |

| | | | Ct | | | $2^{-ddCt}$ | | |
|---|---|---|---|---|---|---|---|---|
| miRNA | Mouse ID | *Mir182* $^{ko}$ genotype | Replicate 1 | Replicate 2 | Replicate 3 | Replicate 1 | Replicate 2 | Replicate 3 |
| Mir99a | DMOM61.3f | Wildtype | 31.39 | 31.48 | 31.61 | | | |
| Mir99a | DMOM64.3j | Wildtype | 26.60 | 26.54 | 26.00 | | | |
| Mir99a | DMOM64.3i | Wildtype | 28.58 | 28.85 | 28.60 | | | |
| Mir99a | DMOM60.3g | Wildtype | 29.59 | 27.97 | 28.29 | | | |
| Mir99a | DMOM61.3c | Heterozygote | 30.02 | 30.46 | 30.63 | | | |
| Mir99a | DMOM64.3d | Heterozygote | 27.87 | 28.04 | 26.30 | | | |
| Mir99a | DMOM64.3b | Heterozygote | 26.30 | 27.10 | 25.18 | | | |
| Mir99a | DMOM64.3e | Heterozygote | 28.93 | 28.73 | 29.27 | | | |
| Mir99a | DMOM61.3e | Homozygote | 31.02 | 34.13 | 35.14 | | | |
| Mir99a | DMOM61.3d | Homozygote | 32.50 | 33.05 | 32.19 | | | |
| Mir99a | DMOM60.3i | Homozygote | 26.84 | 27.20 | 26.35 | | | |
| Mir99a | DMOM64.3h | Homozygote | 30.53 | 30.13 | 30.35 | | | |
| Mir96 | DMOM61.3f | Wildtype | 29.14836608 | 28.60864682 | 28.99373437 | 0.851778177 | 1.238221 | 0.948146 |
| Mir96 | DMOM64.3j | Wildtype | 24.48468434 | 23.75717103 | 24.77593514 | 0.904115607 | 1.497019 | 0.738837 |
| Mir96 | DMOM64.3i | Wildtype | 25.90410795 | 26.14971106 | 26.18099453 | 1.128309787 | 0.951688 | 0.931273 |
| Mir96 | DMOM60.3g | Wildtype | 26.06470954 | 24.21935069 | 26.8026871 | 0.774250817 | 2.782211 | 0.464225 |
| Mir96 | DMOM61.3c | Heterozygote | 26.95472673 | 28.23315353 | 26.90022314 | 1.790739849 | 0.738223 | 1.859686 |
| Mir96 | DMOM64.3d | Heterozygote | 23.8130977 | 24.01624737 | 25.42584741 | 2.932611062 | 2.547419 | 0.958889 |
| Mir96 | DMOM64.3b | Heterozygote | 24.23492503 | 23.75168774 | 23.02848308 | 0.945875304 | 1.322217 | 2.182776 |
| Mir96 | DMOM64.3e | Heterozygote | 26.45129339 | 27.24144134 | 25.42645957 | 0.949367101 | 0.549005 | 1.931701 |
| Mir96 | DMOM61.3e | Homozygote | 34.37504185 | 32.59476991 | 33.20710187 | 0.087148387 | 0.299347 | 0.195814 |
| Mir96 | DMOM61.3d | Homozygote | 28.08159958 | 29.48185926 | 28.28856868 | 3.789940099 | 1.43586 | 3.283435 |
| Mir96 | DMOM60.3i | Homozygote | 23.42759967 | 28.13505286 | 26.7595073 | 1.360981081 | 0.052092 | 0.13516 |
| Mir96 | DMOM64.3h | Homozygote | 26.34910898 | 26.34439212 | 29.17707792 | 3.8580756 | 3.87071 | 0.543334 |
| Mir182 | DMOM61.3f | Wildtype | 32.06747386 | 30.4985933 | 29.6144754 | 0.394849195 | 1.171417 | 2.162008 |
| Mir182 | DMOM64.3j | Wildtype | 26.71239328 | 26.15856564 | 27.31840502 | 1.012130069 | 1.485782 | 0.66498 |
| Mir182 | DMOM64.3i | Wildtype | 28.93685814 | 28.12230756 | 27.66330271 | 0.617266184 | 1.085617 | 1.492282 |
| Mir182 | DMOM60.3g | Wildtype | 28.42501735 | 28.76876622 | 28.9579407 | 1.224526567 | 0.964916 | 0.846335 |
| Mir182 | DMOM61.3c | Heterozygote | 31.17408582 | 28.26510139 | 28.82529633 | 0.337071939 | 2.531711 | 1.717033 |
| Mir182 | DMOM64.3d | Heterozygote | 28.97274614 | 28.96427985 | 28.83203846 | 0.43019474 | 0.432727 | 0.474266 |
| Mir182 | DMOM64.3b | Heterozygote | 27.48944939 | 27.42165409 | 27.49412775 | 0.519689576 | 0.544694 | 0.518007 |
| Mir182 | DMOM64.3e | Heterozygote | 29.92674722 | 30.23128932 | 29.67286754 | 0.382130085 | 0.30941 | 0.455656 |
| Mir182 | DMOM61.3e | Homozygote | - | - | - | 0 | 0 | 0 |
| Mir182 | DMOM61.3d | Homozygote | - | - | - | 0 | 0 | 0 |
| Mir182 | DMOM60.3i | Homozygote | - | - | 39.38926052 | 0 | 0 | 0.000173 |
| Mir182 | DMOM64.3h | Homozygote | - | - | - | 0 | 0 | 0 |
| Mir183 | DMOM61.3f | Wildtype | 28.64520948 | 29.33010468 | 30.86674343 | 1.957225132 | 1.217494 | 0.419655 |
| Mir183 | DMOM64.3j | Wildtype | 27.61995568 | 27.62788625 | 26.31644594 | 0.741305579 | 0.737242 | 1.829754 |
| Mir183 | DMOM64.3i | Wildtype | 29.06585854 | 29.34332376 | 29.42164693 | 1.157558043 | 0.955031 | 0.904565 |
| Mir183 | DMOM60.3g | Wildtype | 28.21585864 | 28.45885259 | 28.52127933 | 1.135088574 | 0.959138 | 0.918521 |
| Mir183 | DMOM61.3c | Heterozygote | 31.04599957 | 27.73586347 | 30.34039046 | 0.170327864 | 1.689413 | 0.277776 |
| Mir183 | DMOM64.3d | Heterozygote | 28.234737 | 27.28606902 | 28.50032168 | 0.985808214 | 1.902698 | 0.820056 |
| Mir183 | DMOM64.3b | Heterozygote | 26.41107664 | 26.84535582 | 26.7641477 | 1.507755772 | 1.115835 | 1.180446 |
| Mir183 | DMOM64.3e | Heterozygote | 28.77475709 | 29.37476451 | 29.50259448 | 1.741398227 | 1.148888 | 1.051471 |
| Mir183 | DMOM61.3e | Homozygote | 32.06788944 | 30.26558985 | 30.33908972 | 0.699246793 | 2.438803 | 2.317667 |
| Mir183 | DMOM61.3d | Homozygote | 29.26657077 | 30.16400192 | 30.60029951 | 2.702523753 | 1.450828 | 1.072203 |
| Mir183 | DMOM60.3i | Homozygote | 26.68386327 | 26.48381219 | 26.04787531 | 0.927529127 | 1.065489 | 1.441382 |
| Mir183 | DMOM64.3h | Homozygote | 28.46238299 | 31.91900624 | 31.48020316 | 6.423857362 | 0.585125 | 0.793125 |

**ABR thresholds for Fig. S2**

| Mouse ID | Experiment date | Date of birth | Allele | Genotype | Weight (g) | Age (days) | Age (weeks) | Click threshold | 3kHz threshold | 6kHz threshold | 12kHz threshold | 18kHz threshold | 24kHz threshold | 30kHz threshold | 36kHz threshold | 42kHz threshold |
|---|---|---|---|---|---|---|---|---|---|---|---|---|---|---|---|---|
| DMKO63.1b | 05/10/2016 | 21/09/2016 | *Mir183/96*$^{dko}$ | Wildtype | 6 | 14 | 2.0 | 35 | 65 | 40 | 25 | 30 | 30 | 35 | 50 | 80 |
| DMKO57.4b | 27/10/2016 | 13/10/2016 | *Mir183/96*$^{dko}$ | Wildtype | 8.7 | 14 | 2.0 | 25 | 95 | 70 | 35 | 30 | 25 | 30 | 40 | 60 |
| DMKO57.4c | 27/10/2016 | 13/10/2016 | *Mir183/96*$^{dko}$ | Wildtype | 8 | 14 | 2.0 | 40 | 95 | 90 | 50 | 55 | 50 | 55 | 75 | 95 |
| DMKO57.4a | 27/10/2016 | 13/10/2016 | *Mir183/96*$^{dko}$ | Wildtype | 7.3 | 14 | 2.0 | 35 | 90 | 55 | 45 | 40 | 30 | 40 | 45 | 55 |
| DMKO68.2a | 14/11/2016 | 31/10/2016 | *Mir183/96*$^{dko}$ | Wildtype | 8 | 14 | 2.0 | 35 | 85 | 80 | 40 | 35 | 40 | 40 | 45 | 65 |
| DMKO68.2d | 14/11/2016 | 31/10/2016 | *Mir183/96*$^{dko}$ | Wildtype | 4.5 | 14 | 2.0 | 50 | 95 | 85 | 45 | 50 | 70 | 95 | 95 | 95 |
| DMKO68.2e | 14/11/2016 | 31/10/2016 | *Mir183/96*$^{dko}$ | Wildtype | 4.6 | 14 | 2.0 | 40 | 90 | 85 | 45 | 45 | 45 | 50 | 60 | 70 |
| DMKO66.2c | 20/12/2016 | 06/12/2016 | *Mir183/96*$^{dko}$ | Wildtype | 7.3 | 14 | 2.0 | 40 | 95 | 75 | 40 | 35 | 40 | 60 | 70 | 95 |
| DMKO66.5e | 21/04/2017 | 07/04/2017 | *Mir183/96*$^{dko}$ | Wildtype | 7.0 | 14 | 2.0 | 45 | 95 | 80 | 35 | 35 | 30 | 50 | 70 | 95 |
| DMKO76.3d | 20/09/2017 | 06/09/2017 | *Mir183/96*$^{dko}$ | Wildtype | 6.4 | 14 | 2.0 | 40 | 95 | 55 | 50 | 45 | 45 | 45 | 50 | 55 |
| DMKO63.1a | 05/10/2016 | 21/09/2016 | *Mir183/96*$^{dko}$ | Heterozygote | 5.6 | 14 | 2.0 | 30 | 65 | 65 | 30 | 15 | 25 | 30 | 45 | 75 |
| DMKO68.2c | 14/11/2016 | 31/10/2016 | *Mir183/96*$^{dko}$ | Heterozygote | 5.6 | 14 | 2.0 | 45 | 95 | 80 | 45 | 55 | 50 | 65 | 65 | 95 |
| DMKO66.2a | 20/12/2016 | 06/12/2016 | *Mir183/96*$^{dko}$ | Heterozygote | 7.7 | 14 | 2.0 | 45 | 95 | 75 | 45 | 40 | 40 | 60 | 80 | 95 |
| DMKO66.2b | 20/12/2016 | 06/12/2016 | *Mir183/96*$^{dko}$ | Heterozygote | 7.5 | 14 | 2.0 | 45 | 95 | 80 | 45 | 45 | 40 | 60 | 90 | 95 |
| DMKO66.2e | 20/12/2016 | 06/12/2016 | *Mir183/96*$^{dko}$ | Heterozygote | 7.6 | 14 | 2.0 | 40 | 95 | 90 | 55 | 40 | 50 | 65 | 95 | 95 |
| DMKO66.5a | 21/04/2017 | 07/04/2017 | *Mir183/96*$^{dko}$ | Heterozygote | 6.8 | 14 | 2.0 | 45 | 95 | 85 | 60 | 30 | 35 | 50 | 60 | 75 |
| DMKO66.5g | 21/04/2017 | 07/04/2017 | *Mir183/96*$^{dko}$ | Heterozygote | 6.6 | 14 | 2.0 | 50 | 95 | 80 | 60 | 60 | 40 | 65 | 70 | 95 |
| DMKO57.4d | 27/10/2016 | 13/10/2016 | *Mir183/96*$^{dko}$ | Homozygote | 6.9 | 14 | 2.0 | 95 | 95 | 95 | 95 | 95 | 95 | 95 | 95 | 95 |
| DMKO68.2b | 14/11/2016 | 31/10/2016 | *Mir183/96*$^{dko}$ | Homozygote | 7.3 | 14 | 2.0 | 95 | 95 | 95 | 95 | 95 | 95 | 95 | 95 | 95 |
| DMKO66.2d | 20/12/2016 | 06/12/2016 | *Mir183/96*$^{dko}$ | Homozygote | 6.4 | 14 | 2.0 | 95 | 95 | 95 | 95 | 95 | 95 | 95 | 95 | 95 |
| DMKO66.5c | 21/04/2017 | 07/04/2017 | *Mir183/96*$^{dko}$ | Homozygote | 4.6 | 14 | 2.0 | 95 | 95 | 95 | 95 | 95 | 95 | 95 | 95 | 95 |
| DMKO66.5f | 21/04/2017 | 07/04/2017 | *Mir183/96*$^{dko}$ | Homozygote | 6.2 | 14 | 2.0 | 95 | 95 | 95 | 95 | 95 | 95 | 95 | 95 | 95 |
| DMKO76.3i | 20/09/2017 | 06/09/2017 | *Mir183/96*$^{dko}$ | Homozygote | 5.9 | 14 | 2.0 | 95 | 95 | 95 | 95 | 95 | 95 | 95 | 95 | 95 |
| DMKO55.5d | 04/10/2016 | 13/09/2016 | *Mir183/96*$^{dko}$ | Wildtype | 7.6 | 21 | 3.0 | 25 | 65 | 30 | 15 | 15 | 20 | 25 | 35 | 35 |
| DMKO57.3a | 13/10/2016 | 22/09/2016 | *Mir183/96*$^{dko}$ | Wildtype | 11.5 | 21 | 3.0 | 30 | 75 | 35 | 15 | 30 | 25 | 40 | 50 | 70 |
| DMKO57.3b | 13/10/2016 | 22/09/2016 | *Mir183/96*$^{dko}$ | Wildtype | 9.5 | 21 | 3.0 | 25 | 65 | 45 | 10 | 30 | 30 | 35 | 45 | 70 |
| DMKO57.3c | 13/10/2016 | 22/09/2016 | *Mir183/96*$^{dko}$ | Wildtype | 9.3 | 21 | 3.0 | 25 | 65 | 30 | 15 | 25 | 25 | 30 | 50 | 50 |
| DMKO57.3d | 13/10/2016 | 22/09/2016 | *Mir183/96*$^{dko}$ | Wildtype | 10.8 | 21 | 3.0 | 30 | 70 | 40 | 15 | 25 | 30 | 45 | 70 | 85 |
| DMKO57.3f | 13/10/2016 | 22/09/2016 | *Mir183/96*$^{dko}$ | Wildtype | 10.4 | 21 | 3.0 | 25 | 60 | 30 | 15 | 15 | 20 | 45 | 30 | 30 |
| DMKO67.1c | 02/11/2016 | 12/10/2016 | *Mir183/96*$^{dko}$ | Wildtype | 8.9 | 21 | 3.0 | 30 | 75 | 50 | 20 | 30 | 30 | 45 | 50 | 90 |
| DMKO67.1h | 02/11/2016 | 12/10/2016 | *Mir183/96*$^{dko}$ | Wildtype | 8.6 | 21 | 3.0 | 25 | 70 | 30 | 20 | 20 | 25 | 35 | 35 | 70 |
| DMKO69.5b | 31/05/2017 | 10/05/2017 | *Mir183/96*$^{dko}$ | Wildtype | 7.5 | 21 | 3.0 | 25 | 85 | 40 | 25 | 25 | 25 | 30 | 40 | 45 |
| DMKO74.3e | 01/09/2017 | 11/08/2017 | *Mir183/96*$^{dko}$ | Wildtype | 9.1 | 21 | 3.0 | 25 | 70 | 35 | 15 | 25 | 25 | 30 | 45 | 40 |
| DMKO55.5h | 04/10/2016 | 13/09/2016 | *Mir183/96*$^{dko}$ | Heterozygote | 8 | 21 | 3.0 | 20 | 65 | 40 | 15 | 25 | 25 | 25 | 45 | 50 |
| DMKO67.1a | 02/11/2016 | 12/10/2016 | *Mir183/96*$^{dko}$ | Heterozygote | 7.7 | 21 | 3.0 | 35 | 65 | 35 | 15 | 25 | 30 | 45 | 70 | 95 |
| DMKO67.1b | 02/11/2016 | 12/10/2016 | *Mir183/96*$^{dko}$ | Heterozygote | 8.9 | 21 | 3.0 | 30 | 65 | 30 | 20 | 25 | 25 | 35 | 45 | 45 |
| DMKO67.1d | 02/11/2016 | 12/10/2016 | *Mir183/96*$^{dko}$ | Heterozygote | 8.7 | 21 | 3.0 | 25 | 65 | 35 | 15 | 20 | 35 | 50 | 55 | 60 |
| DMKO67.1e | 02/11/2016 | 12/10/2016 | *Mir183/96*$^{dko}$ | Heterozygote | 8.8 | 21 | 3.0 | 30 | 75 | 35 | 15 | 25 | 30 | 40 | 45 | 45 |
| DMKO67.1f | 02/11/2016 | 12/10/2016 | *Mir183/96*$^{dko}$ | Heterozygote | 8.7 | 21 | 3.0 | 25 | 65 | 50 | 20 | 25 | 30 | 35 | 55 | 90 |
| DMKO67.1g | 02/11/2016 | 12/10/2016 | *Mir183/96*$^{dko}$ | Heterozygote | 8.6 | 21 | 3.0 | 25 | 65 | 40 | 20 | 20 | 25 | 35 | 50 | 60 |
| DMKO69.3h | 27/03/2017 | 06/03/2017 | *Mir183/96*$^{dko}$ | Heterozygote | 9.0 | 21 | 3.0 | 25 | 65 | 45 | 20 | 25 | 30 | 35 | 70 | 80 |
| DMKO69.3f | 27/03/2017 | 06/03/2017 | *Mir183/96*$^{dko}$ | Heterozygote | 8.5 | 21 | 3.0 | 25 | 70 | 40 | 20 | 15 | 25 | 35 | 40 | 50 |
| DMKO69.4b | 19/04/2017 | 29/03/2017 | *Mir183/96*$^{dko}$ | Heterozygote | 6.7 | 21 | 3.0 | 30 | 80 | 55 | 45 | 55 | 65 | 80 | 95 | 95 |
| DMKO57.3e | 13/10/2016 | 22/09/2016 | *Mir183/96*$^{dko}$ | Homozygote | 11.5 | 21 | 3.0 | 95 | 95 | 95 | 95 | 95 | 95 | 95 | 95 | 95 |
| DMKO69.4g | 19/04/2017 | 29/03/2017 | *Mir183/96*$^{dko}$ | Homozygote | 8.5 | 21 | 3.0 | 95 | 95 | 95 | 95 | 95 | 95 | 95 | 95 | 95 |
| DMKO69.4a | 19/04/2017 | 29/03/2017 | *Mir183/96*$^{dko}$ | Homozygote | 8.1 | 21 | 3.0 | 95 | 95 | 95 | 95 | 95 | 95 | 95 | 95 | 95 |
| DMKO69.5f | 31/05/2017 | 10/05/2017 | *Mir183/96*$^{dko}$ | Homozygote | 7.7 | 21 | 3.0 | 95 | 95 | 95 | 95 | 95 | 95 | 95 | 95 | 95 |
| DMKO69.5g | 31/05/2017 | 10/05/2017 | *Mir183/96*$^{dko}$ | Homozygote | 7.9 | 21 | 3.0 | 95 | 95 | 95 | 95 | 95 | 95 | 95 | 95 | 95 |
| DMKO74.3j | 01/09/2017 | 11/08/2017 | *Mir183/96*$^{dko}$ | Homozygote | 8.5 | 21 | 3.0 | 95 | 95 | 95 | 95 | 95 | 95 | 95 | 95 | 95 |
| DMKO37.1e | 23/09/2014 | 26/08/2014 | *Mir183/96*$^{dko}$ | Wildtype | 16.9 | 28 | 4.0 | 25 | 50 | 30 | 10 | 20 | 40 | 90 | 95 | 90 |
| DMKO37.1h | 23/09/2014 | 26/08/2014 | *Mir183/96*$^{dko}$ | Wildtype | 16.1 | 28 | 4.0 | 25 | 50 | 35 | 15 | 15 | 35 | 95 | 95 | 95 |
| DMKO39.1c | 29/01/2015 | 01/01/2015 | *Mir183/96*$^{dko}$ | Wildtype | 14.3 | 28 | 4.0 | 25 | 80 | 30 | 10 | 20 | 30 | 45 | 95 | 95 |
| DMKO39.1g | 29/01/2015 | 01/01/2015 | *Mir183/96*$^{dko}$ | Wildtype | 12.6 | 28 | 4.0 | 25 | 75 | 40 | 10 | 25 | 45 | 95 | 95 | 95 |
| DMKO41.1f | 29/01/2015 | 01/01/2015 | *Mir183/96*$^{dko}$ | Wildtype | 15.7 | 28 | 4.0 | 30 | 55 | 30 | 15 | 25 | 25 | 70 | 95 | 95 |
| DMKO14.2d | 10/09/2012 | 13/08/2012 | *Mir183/96*$^{dko}$ | Wildtype | | 28 | 4.0 | 30 | | 35 | 25 | 30 | 30 | 65 | | |
| DMKO12.4b | 07/09/2012 | 09/08/2012 | *Mir183/96*$^{dko}$ | Wildtype | | 29 | 4.1 | 30 | | 40 | 25 | 30 | 35 | 45 | | |
| DMKO17.2g | 20/09/2012 | 22/08/2012 | *Mir183/96*$^{dko}$ | Wildtype | | 29 | 4.1 | 30 | | 35 | 20 | 25 | 40 | 65 | | |
| DMKO17.2h | 20/09/2012 | 22/08/2012 | *Mir183/96*$^{dko}$ | Wildtype | | 29 | 4.1 | 25 | | 30 | 15 | 20 | 25 | 50 | | |
| DMKO18.1d | 04/10/2012 | 05/09/2012 | *Mir183/96*$^{dko}$ | Wildtype | | 29 | 4.1 | 25 | | 30 | 20 | 20 | 25 | 35 | | |
| DMKO18.1b | 04/10/2012 | 05/09/2012 | *Mir183/96*$^{dko}$ | Wildtype | | 29 | 4.1 | 25 | | 30 | 15 | 25 | 25 | 45 | | |
| DMKO37.1c | 23/09/2014 | 26/08/2014 | *Mir183/96*$^{dko}$ | Heterozygote | 16.5 | 28 | 4.0 | 25 | 50 | 30 | 5 | 20 | 25 | 35 | 80 | 95 |
| DMKO37.1a | 23/09/2014 | 26/08/2014 | *Mir183/96*$^{dko}$ | Heterozygote | 17.3 | 28 | 4.0 | 20 | 75 | 30 | 20 | 40 | 85 | 95 | 95 | 95 |
| DMKO37.1g | 23/09/2014 | 26/08/2014 | *Mir183/96*$^{dko}$ | Heterozygote | 16.1 | 28 | 4.0 | 25 | 55 | 30 | 5 | 10 | 25 | 45 | 95 | 90 |
| DMKO39.1h | 29/01/2015 | 01/01/2015 | *Mir183/96*$^{dko}$ | Heterozygote | 13.8 | 28 | 4.0 | 25 | 60 | 30 | 10 | 20 | 30 | 45 | 95 | 95 |
| DMKO41.1b | 29/01/2015 | 01/01/2015 | *Mir183/96*$^{dko}$ | Heterozygote | 15.6 | 28 | 4.0 | 30 | 65 | 45 | 15 | 30 | 45 | 95 | 95 | 95 |
| DMKO41.1a | 29/01/2015 | 01/01/2015 | *Mir183/96*$^{dko}$ | Heterozygote | 15.5 | 28 | 4.0 | 25 | 65 | 40 | 15 | 15 | 30 | 45 | 95 | 95 |
| DMKO37.1b | 23/09/2014 | 26/08/2014 | *Mir183/96*$^{dko}$ | Heterozygote | 17 | 28 | 4.0 | 30 | 65 | 35 | 20 | 25 | 35 | 55 | 95 | 95 |
| DMKO12.4c | 07/09/2012 | 09/08/2012 | *Mir183/96*$^{dko}$ | Heterozygote | | 29 | 4.1 | 30 | | 30 | 20 | 20 | 40 | 90 | | |
| DMKO12.4d | 07/09/2012 | 09/08/2012 | *Mir183/96*$^{dko}$ | Heterozygote | | 29 | 4.1 | 35 | | 50 | 35 | 30 | 35 | 75 | | |
| DMKO17.2d | 20/09/2012 | 22/08/2012 | *Mir183/96*$^{dko}$ | Heterozygote | | 29 | 4.1 | 25 | | 35 | 20 | 25 | 30 | 50 | | |
| DMKO17.2a | 20/09/2012 | 22/08/2012 | *Mir183/96*$^{dko}$ | Heterozygote | | 29 | 4.1 | 30 | | 35 | 20 | 25 | 35 | 45 | | |
| DMKO17.2f | 20/09/2012 | 22/08/2012 | *Mir183/96*$^{dko}$ | Heterozygote | | 29 | 4.1 | 30 | | 35 | 15 | 25 | 35 | 45 | | |
| DMKO18.1a | 04/10/2012 | 05/09/2012 | *Mir183/96*$^{dko}$ | Heterozygote | | 29 | 4.1 | 25 | | 30 | 20 | 20 | 35 | 50 | | |
| DMKO15.2c | 04/09/2012 | 07/08/2012 | *Mir183/96*$^{dko}$ | Heterozygote | | 28 | 4.0 | 30 | | 30 | 25 | 25 | 30 | 45 | | |
| DMKO37.1d | 23/09/2014 | 26/08/2014 | *Mir183/96*$^{dko}$ | Homozygote | 15.5 | 28 | 4.0 | 95 | 95 | 95 | 95 | 95 | 95 | 95 | 95 | 95 |
| DMKO37.1f | 23/09/2014 | 26/08/2014 | *Mir183/96*$^{dko}$ | Homozygote | 13.9 | 28 | 4.0 | 95 | 95 | 95 | 95 | 95 | 95 | 95 | 95 | 95 |
| DMKO39.1f | 29/01/2015 | 01/01/2015 | *Mir183/96*$^{dko}$ | Homozygote | 11.1 | 28 | 4.0 | 95 | 95 | 95 | 95 | 95 | 95 | 95 | 95 | 95 |
| DMKO41.1e | 29/01/2015 | 01/01/2015 | *Mir183/96*$^{dko}$ | Homozygote | 9.8 | 28 | 4.0 | 95 | 95 | 95 | 95 | 95 | 95 | 95 | 95 | 95 |
| DMKO41.1d | 29/01/2015 | 01/01/2015 | *Mir183/96*$^{dko}$ | Homozygote | 13.4 | 28 | 4.0 | 95 | 95 | 95 | 95 | 95 | 95 | 95 | 95 | 95 |
| DMKO14.2b | 07/09/2012 | 09/08/2012 | *Mir183/96*$^{dko}$ | Homozygote | | 29 | 4.1 | 95 | | 95 | 85 | 95 | 95 | 95 | | |
| DMKO14.2c | 07/09/2012 | 09/08/2012 | *Mir183/96*$^{dko}$ | Homozygote | | 29 | 4.1 | 95 | | 95 | 95 | 95 | 95 | 95 | | |
| DMKO12.4a | 07/09/2012 | 09/08/2012 | *Mir183/96*$^{dko}$ | Homozygote | | 29 | 4.1 | 95 | | 95 | 95 | 95 | 95 | 95 | | |
| DMKO17.2e | 20/09/2012 | 22/08/2012 | *Mir183/96*$^{dko}$ | Homozygote | | 29 | 4.1 | 95 | | 95 | 95 | 95 | 95 | 95 | | |
| DMKO18.1c | 04/10/2012 | 05/09/2012 | *Mir183/96*$^{dko}$ | Homozygote | | 29 | 4.1 | 95 | | 95 | 95 | 95 | 95 | 95 | | |
| DMKO31.1d | 16/05/2014 | 22/03/2014 | *Mir183/96*$^{dko}$ | Wildtype | 23.2 | 55 | 7.9 | 20 | 55 | 30 | 10 | 20 | 15 | 40 | 80 | 90 |
| DMKO31.1e | 16/05/2014 | 22/03/2014 | *Mir183/96*$^{dko}$ | Wildtype | 20.5 | 55 | 7.9 | 20 | 45 | 25 | 10 | 20 | 20 | 40 | 70 | 95 |
| DMKO33.1a | 20/05/2014 | 23/03/2014 | *Mir183/96*$^{dko}$ | Wildtype | 28.3 | 58 | 8.3 | 25 | 55 | 30 | 10 | 20 | 45 | 75 | 95 | 95 |
| DMKO33.1c | 20/05/2014 | 23/03/2014 | *Mir183/96*$^{dko}$ | Wildtype | 21.6 | 58 | 8.3 | 15 | 55 | 25 | 10 | 15 | 25 | 40 | 95 | 95 |
| DMKO37.1e | 21/10/2014 | 26/08/2014 | *Mir183/96*$^{dko}$ | Wildtype | 23.1 | 56 | 8.0 | 20 | 65 | 40 | 25 | 30 | 40 | 70 | 95 | 95 |
| DMKO37.1h | 21/10/2014 | 26/08/2014 | *Mir183/96*$^{dko}$ | Wildtype | 22.1 | 56 | 8.0 | 30 | 65 | 30 | 5 | 20 | 50 | 95 | 95 | 95 |
| DMKO39.1c | 26/02/2015 | 01/01/2015 | *Mir183/96*$^{dko}$ | Wildtype | 23.3 | 56 | 8.0 | 25 | 55 | 30 | 15 | 25 | 40 | 75 | 95 | 95 |
| DMKO39.1g | 26/02/2015 | 01/01/2015 | *Mir183/96*$^{dko}$ | Wildtype | 19.1 | 56 | 8.0 | 30 | 70 | 45 | 15 | 25 | 45 | 95 | 95 | 95 |
| DMKO41.1f | 26/02/2015 | 01/01/2015 | *Mir183/96*$^{dko}$ | Wildtype | 27.4 | 56 | 8.0 | 25 | 70 | 35 | 15 | 25 | 30 | 80 | 95 | 95 |
| DMKO31.1b | 16/05/2014 | 22/03/2014 | *Mir183/96*$^{dko}$ | Heterozygote | 24.1 | 55 | 7.9 | 20 | 75 | 30 | 10 | 15 | 30 | 65 | 95 | 95 |
| DMKO33.1b | 20/05/2014 | 23/03/2014 | *Mir183/96*$^{dko}$ | Heterozygote | 28.8 | 58 | 8.3 | 25 | 60 | 30 | 20 | 30 | 70 | 95 | 95 | 95 |
| DMKO33.1e | 20/05/2014 | 23/03/2014 | *Mir183/96*$^{dko}$ | Heterozygote | 20.7 | 58 | 8.3 | 25 | 45 | 30 | 5 | 20 | 20 | 40 | 85 | 95 |
| DMKO33.1d | 20/05/2014 | 23/03/2014 | *Mir183/96*$^{dko}$ | Heterozygote | 23.7 | 58 | 8.3 | 15 | 50 | 25 | 10 | 10 | 20 | 50 | 90 | 95 |
| DMKO37.1c | 21/10/2014 | 26/08/2014 | *Mir183/96*$^{dko}$ | Heterozygote | 22.6 | 56 | 8.0 | 25 | 45 | 25 | 5 | 20 | 30 | 30 | 55 | 85 |
| DMKO37.1a | 21/10/2014 | 26/08/2014 | *Mir183/96*$^{dko}$ | Heterozygote | 24.3 | 56 | 8.0 | 25 | 70 | 30 | 20 | 20 | 75 | 95 | 95 | 95 |
| DMKO37.1g | 21/10/2014 | 26/08/2014 | *Mir183/96*$^{dko}$ | Heterozygote | 21 | 56 | 8.0 | 25 | 60 | 35 | 10 | 20 | 30 | 55 | 70 | 95 |
| DMKO34.3c | 03/12/2014 | 08/10/2014 | *Mir183/96*$^{dko}$ | Heterozygote | 26.2 | 56 | 8.0 | 30 | 70 | 40 | 25 | 35 | 50 | 95 | 95 | 95 |
| DMKO39.1h | 26/02/2015 | 01/01/2015 | *Mir183/96*$^{dko}$ | Heterozygote | 19.7 | 56 | 8.0 | 25 | 60 | 35 | 25 | 35 | 35 | 35 | 70 | 95 |
| DMKO41.1a | 26/02/2015 | 01/01/2015 | *Mir183/96*$^{dko}$ | Heterozygote | 23.5 | 56 | 8.0 | 25 | 70 | 55 | 15 | 30 | 30 | 50 | 85 | 95 |
| DMKO41.1b | 26/02/2015 | 01/01/2015 | *Mir183/96*$^{dko}$ | Heterozygote | 25.5 | 56 | 8.0 | 30 | 60 | 35 | 25 | 30 | 35 | 65 | 95 | 95 |
| DMKO37.1b | 21/10/2014 | 26/08/2014 | *Mir183/96*$^{dko}$ | Heterozygote | 22.7 | 56 | 8.0 | 25 | 50 | 35 | 15 | 25 | 30 | 40 | 55 | 90 |
| DMKO37.1d | 21/10/2014 | 26/08/2014 | *Mir183/96*$^{dko}$ | Homozygote | 22.9 | 56 | 8.0 | 95 | 95 | 95 | 95 | 95 | 95 | 95 | 95 | 95 |
| DMKO37.1f | 21/10/2014 | 26/08/2014 | *Mir183/96*$^{dko}$ | Homozygote | 18.3 | 56 | 8.0 | 95 | 95 | 95 | 95 | 95 | 95 | 95 | 95 | 95 |
| DMKO34.3a | 03/12/2014 | 08/10/2014 | *Mir183/96*$^{dko}$ | Homozygote | 26.8 | 56 | 8.0 | 95 | 95 | 95 | 95 | 95 | 95 | 95 | 95 | 95 |
| DMKO34.3b | 03/12/2014 | 08/10/2014 | *Mir183/96*$^{dko}$ | Homozygote | 24.8 | 56 | 8.0 | 95 | 95 | 95 | 95 | 95 | 95 | 95 | 95 | 95 |
| DMKO39.1f | 26/02/2015 | 01/01/2015 | *Mir183/96*$^{dko}$ | Homozygote | 19.1 | 56 | 8.0 | 95 | 95 | 95 | 95 | 95 | 95 | 95 | 95 | 95 |
| DMKO41.1e | 26/02/2015 | 01/01/2015 | *Mir183/96*$^{dko}$ | Homozygote | 22.5 | 56 | 8.0 | 95 | 95 | 95 | 95 | 95 | 95 | 95 | 95 | 95 |
| DMKO41.1d | 26/02/2015 | 01/01/2015 | *Mir183/96*$^{dko}$ | Homozygote | 23.7 | 56 | 8.0 | 95 | 95 | 95 | 95 | 95 | 95 | 95 | 95 | 95 |
| DMKO30.3e | 08/04/2014 | 07/01/2014 | *Mir183/96*$^{dko}$ | Wildtype | 27.8 | 91 | 13.0 | 20 | 55 | 25 | 5 | 20 | 25 | 45 | 90 | 95 |
| DMKO33.1a | 20/06/2014 | 23/03/2014 | *Mir183/96*$^{dko}$ | Wildtype | 32.1 | 89 | 12.7 | 30 | 50 | 25 | 15 | 35 | 80 | 95 | 95 | 95 |
| DMKO33.1c | 20/06/2014 | 23/03/2014 | *Mir183/96*$^{dko}$ | Wildtype | 24.7 | 89 | 12.7 | 25 | 55 | 30 | 10 | 20 | 55 | 95 | 95 | 95 |
| DMKO31.1d | 20/06/2014 | 22/03/2014 | *Mir183/96*$^{dko}$ | Wildtype | 30.4 | 90 | 12.9 | 30 | 50 | 30 | 20 | 40 | 80 | 95 | 95 | 95 |
| DMKO31.1e | 20/06/2014 | 22/03/2014 | *Mir183/96*$^{dko}$ | Wildtype | 21.8 | 90 | 12.9 | 25 | 45 | 30 | 5 | 20 | 30 | 60 | 95 | 95 |
| DMKO37.1e | 24/11/2014 | 26/08/2014 | *Mir183/96*$^{dko}$ | Wildtype | 26.8 | 90 | 12.9 | 25 | 65 | 25 | 15 | 30 | 70 | 95 | 95 | 95 |
| DMKO37.1h | 24/11/2014 | 26/08/2014 | *Mir183/96*$^{dko}$ | Wildtype | 24.7 | 90 | 12.9 | 25 | 75 | 30 | 25 | 30 | 85 | 95 | 95 | 95 |
| DMKO39.1c | 01/04/2015 | 01/01/2015 | *Mir183/96*$^{dko}$ | Wildtype | 25.9 | 90 | 12.9 | 25 | 65 | 35 | 10 | 25 | 55 | 95 | 95 | 95 |
| DMKO39.1g | 01/04/2015 | 01/01/2015 | *Mir183/96*$^{dko}$ | Wildtype | 22 | 90 | 12.9 | 30 | 65 | 35 | 20 | 45 | 80 | 95 | 95 | 95 |
| DMKO41.1f | 01/04/2015 | 01/01/2015 | *Mir183/96*$^{dko}$ | Wildtype | 33.8 | 90 | 12.9 | 25 | 70 | 35 | 10 | 20 | 35 | 85 | 95 | 95 |
| DMKO30.3b | 08/04/2014 | 07/01/2014 | *Mir183/96*$^{dko}$ | Heterozygote | 30.2 | 91 | 13.0 | 25 | 70 | 25 | 25 | 15 | 40 | 95 | 95 | 95 |
| DMKO30.3a | 08/04/2014 | 07/01/2014 | *Mir183/96*$^{dko}$ | Heterozygote | 27.2 | 91 | 13.0 | 25 | 60 | 30 | 15 | 15 | 30 | 55 | 95 | 95 |
| DMKO33.1d | 20/06/2014 | 23/03/2014 | *Mir183/96*$^{dko}$ | Heterozygote | 27.4 | 89 | 12.7 | 30 | 65 | 35 | 15 | 25 | 35 | 65 | 95 | 95 |
| DMKO33.1e | 20/06/2014 | 23/03/2014 | *Mir183/96*$^{dko}$ | Heterozygote | 24.6 | 89 | 12.7 | 25 | 70 | 35 | 15 | 25 | 25 | 70 | 95 | 95 |
| DMKO31.1b | 20/06/2014 | 22/03/2014 | *Mir183/96*$^{dko}$ | Heterozygote | 30.4 | 90 | 12.9 | 20 | 50 | 35 | 10 | 20 | 30 | 95 | 95 | 95 |
| DMKO33.1b | 20/06/2014 | 23/03/2014 | *Mir183/96*$^{dko}$ | Heterozygote | 33.7 | 89 | 12.7 | 30 | 75 | 45 | 15 | 35 | 80 | 95 | 95 | 95 |
| DMKO37.1c | 24/11/2014 | 26/08/2014 | *Mir183/96*$^{dko}$ | Heterozygote | 27.9 | 90 | 12.9 | 20 | 50 | 30 | 10 | 20 | 25 | 35 | 70 | 95 |
| DMKO37.1a | 24/11/2014 | 26/08/2014 | *Mir183/96*$^{dko}$ | Heterozygote | 27.7 | 90 | 12.9 | 35 | 55 | 30 | 40 | 55 | 85 | 95 | 95 | 95 |
| DMKO37.1g | 24/11/2014 | 26/08/2014 | *Mir183/96*$^{dko}$ | Heterozygote | 22.4 | 90 | 12.9 | 25 | 60 | 30 | 10 | 20 | 20 | 35 | 85 | 95 |
| DMKO37.3g | 16/01/2015 | 20/10/2014 | *Mir183/96*$^{dko}$ | Heterozygote | 23.4 | 88 | 12.6 | 25 | 55 | 35 | 10 | 25 | 30 | 35 | 65 | 95 |
| DMKO37.1b | 24/11/2014 | 26/08/2014 | *Mir183/96*$^{dko}$ | Heterozygote | 26.3 | 90 | 12.9 | 20 | 50 | 30 | 10 | 15 | 25 | 40 | 70 | 95 |
| DMKO39.1h | 01/04/2015 | 01/01/2015 | *Mir183/96*$^{dko}$ | Heterozygote | 23.1 | 90 | 12.9 | 25 | 50 | 40 | 15 | 25 | 25 | 45 | 95 | 95 |
| DMKO41.1b | 01/04/2015 | 01/01/2015 | *Mir183/96*$^{dko}$ | Heterozygote | 30.8 | 90 | 12.9 | 25 | 50 | 35 | 15 | 30 | 55 | 95 | 95 | 95 |
| DMKO41.1a | 01/04/2015 | 01/01/2015 | *Mir183/96*$^{dko}$ | Heterozygote | 27.9 | 90 | 12.9 | 30 | 70 | 40 | 15 | 35 | 30 | 55 | 95 | 95 |
| DMKO30.3c | 08/04/2014 | 07/01/2014 | *Mir183/96*$^{dko}$ | Homozygote | 28.6 | 91 | 13.0 | 95 | 95 | 95 | 95 | 95 | 95 | 95 | 95 | 95 |
| DMKO30.3d | 09/04/2014 | 07/01/2014 | *Mir183/96*$^{dko}$ | Homozygote | 22.5 | 92 | 13.1 | 95 | 95 | 95 | 95 | 95 | 95 | 95 | 95 | 95 |
| DMKO37.1d | 24/11/2014 | 26/08/2014 | *Mir183/96*$^{dko}$ | Homozygote | 25 | 90 | 12.9 | 95 | 95 | 95 | 95 | 95 | 95 | 95 | 95 | 95 |
| DMKO37.1f | 24/11/2014 | 26/08/2014 | *Mir183/96*$^{dko}$ | Homozygote | 20.7 | 90 | 12.9 | 95 | 95 | 95 | 95 | 95 | 95 | 95 | 95 | 95 |
| DMKO37.3h | 16/01/2015 | 20/10/2014 | *Mir183/96*$^{dko}$ | Homozygote | 26.6 | 88 | 12.6 | 95 | 95 | 95 | 95 | 95 | 95 | 95 | 95 | 95 |
| DMKO37.3i | 16/01/2015 | 20/10/2014 | *Mir183/96*$^{dko}$ | Homozygote | 18.3 | 88 | 12.6 | 95 | 95 | 95 | 95 | 95 | 95 | 95 | 95 | 95 |
| DMKO39.1f | 01/04/2015 | 01/01/2015 | *Mir183/96*$^{dko}$ | Homozygote | 21.7 | 90 | 12.9 | 95 | 95 | 95 | 95 | 95 | 95 | 95 | 95 | 95 |
| DMKO41.1e | 01/04/2015 | 01/01/2015 | *Mir183/96*$^{dko}$ | Homozygote | 25.2 | 90 | 12.9 | 95 | 95 | 95 | 95 | 95 | 95 | 95 | 95 | 95 |
| DMKO30.3e | 07/07/2014 | 07/01/2014 | *Mir183/96*$^{dko}$ | Wildtype | 35.4 | 181 | 25.9 | 25 | 60 | 35 | 20 | 35 | 55 | 95 | 95 | 95 |
| DMKO31.1e | 19/09/2014 | 22/03/2014 | *Mir183/96*$^{dko}$ | Wildtype | 28.6 | 181 | 25.9 | 25 | 55 | 30 | 10 | 35 | 30 | 95 | 95 | 95 |
| DMKO31.1d | 19/09/2014 | 22/03/2014 | *Mir183/96*$^{dko}$ | Wildtype | 39.9 | 181 | 25.9 | 35 | 60 | 25 | 20 | 80 | 95 | 95 | 95 | 95 |
| DMKO33.1a | 19/09/2014 | 23/03/2014 | *Mir183/96*$^{dko}$ | Wildtype | 40.8 | 180 | 25.7 | 40 | 65 | 35 | 25 | 85 | 95 | 95 | 95 | 95 |
| DMKO33.1c | 19/09/2014 | 23/03/2014 | *Mir183/96*$^{dko}$ | Wildtype | 35.2 | 180 | 25.7 | 30 | 80 | 40 | 20 | 50 | 95 | 95 | 95 | 95 |
| DMKO37.1e | 23/02/2015 | 26/08/2014 | *Mir183/96*$^{dko}$ | Wildtype | 34.1 | 181 | 25.9 | 50 | 80 | 50 | 40 | 95 | 95 | 95 | 95 | 95 |
| DMKO37.1h | 23/02/2015 | 26/08/2014 | *Mir183/96*$^{dko}$ | Wildtype | 34.4 | 181 | 25.9 | 50 | 75 | 55 | 75 | 95 | 95 | 95 | 95 | 95 |
| DMKO39.1g | 30/06/2015 | 01/01/2015 | *Mir183/96*$^{dko}$ | Wildtype | | 180 | 25.7 | 60 | 70 | 55 | 70 | 80 | 95 | 95 | 95 | 95 |
| DMKO39.1c | 30/06/2015 | 01/01/2015 | *Mir183/96*$^{dko}$ | Wildtype | | 180 | 25.7 | 30 | 80 | 45 | 20 | 75 | 95 | 95 | 95 | 95 |
| DMKO41.1f | 30/06/2015 | 01/01/2015 | *Mir183/96*$^{dko}$ | Wildtype | | 180 | 25.7 | 30 | 80 | 45 | 35 | 30 | 90 | 95 | 95 | 95 |
| DMKO30.3a | 07/07/2014 | 07/01/2014 | *Mir183/96*$^{dko}$ | Heterozygote | 34.9 | 181 | 25.9 | 25 | 65 | 35 | 15 | 20 | 40 | 85 | 95 | 95 |
| DMKO30.3f | 07/07/2014 | 07/01/2014 | *Mir183/96*$^{dko}$ | Heterozygote | 34.6 | 181 | 25.9 | 30 | 65 | 35 | 15 | 65 | 95 | 95 | 95 | 95 |
| DMKO30.3b | 07/07/2014 | 07/01/2014 | *Mir183/96*$^{dko}$ | Heterozygote | 38.7 | 181 | 25.9 | 20 | 65 | 35 | 10 | 35 | 80 | 95 | 95 | 95 |
| DMKO31.1b | 19/09/2014 | 22/03/2014 | *Mir183/96*$^{dko}$ | Heterozygote | 40.8 | 181 | 25.9 | 25 | 55 | 30 | 10 | 30 | 70 | 95 | 95 | 95 |
| DMKO33.1b | 19/09/2014 | 23/03/2014 | *Mir183/96*$^{dko}$ | Heterozygote | 41.6 | 180 | 25.7 | 50 | 70 | 35 | 80 | 95 | 95 | 95 | 95 | 95 |
| DMKO33.1d | 19/09/2014 | 23/03/2014 | *Mir183/96*$^{dko}$ | Heterozygote | 37.7 | 180 | 25.7 | 20 | 65 | 35 | 15 | 25 | 35 | 95 | 95 | 95 |
| DMKO33.1e | 19/09/2014 | 23/03/2014 | *Mir183/96*$^{dko}$ | Heterozygote | 32.2 | 180 | 25.7 | 30 | 70 | 30 | 5 | 35 | 45 | 95 | 95 | 95 |
| DMKO37.1c | 23/02/2015 | 26/08/2014 | *Mir183/96*$^{dko}$ | Heterozygote | 38.8 | 181 | 25.9 | 30 | 70 | 35 | 15 | 45 | 45 | 85 | 95 | 95 |
| DMKO37.1a | 23/02/2015 | 26/08/2014 | *Mir183/96*$^{dko}$ | Heterozygote | 37.2 | 181 | 25.9 | 75 | 75 | 75 | 75 | 95 | 95 | 95 | 95 | 95 |
| DMKO37.1g | 23/02/2015 | 26/08/2014 | *Mir183/96*$^{dko}$ | Heterozygote | 25.8 | 181 | 25.9 | 25 | 55 | 30 | 15 | 40 | 40 | 95 | 95 | 95 |
| DMKO37.1b | 23/02/2015 | 26/08/2014 | *Mir183/96*$^{dko}$ | Heterozygote | 34.8 | 181 | 25.9 | 30 | 65 | 35 | 15 | 35 | 30 | 75 | 95 | 95 |
| DMKO34.3c | 08/04/2015 | 08/10/2014 | *Mir183/96*$^{dko}$ | Heterozygote | 39.3 | 182 | 26.0 | 40 | 80 | 60 | 40 | 65 | 95 | 95 | 95 | 95 |
| DMKO37.3g | 17/04/2015 | 20/10/2014 | *Mir183/96*$^{dko}$ | Heterozygote | 36.8 | 179 | 25.6 | 25 | 55 | 35 | 10 | 30 | 40 | 60 | 95 | 95 |
| DMKO39.1h | 30/06/2015 | 01/01/2015 | *Mir183/96*$^{dko}$ | Heterozygote | | 180 | 25.7 | 30 | 65 | 25 | 25 | 50 | 55 | 95 | 95 | 95 |
| DMKO41.1b | 30/06/2015 | 01/01/2015 | *Mir183/96*$^{dko}$ | Heterozygote | | 180 | 25.7 | 30 | 80 | 45 | 25 | 40 | 70 | 95 | 95 | 95 |
| DMKO41.1a | 30/06/2015 | 01/01/2015 | *Mir183/96*$^{dko}$ | Heterozygote | | 180 | 25.7 | 30 | 85 | 55 | 15 | 30 | 35 | 95 | 95 | 95 |
| DMKO30.3d | 07/07/2014 | 07/01/2014 | *Mir183/96*$^{dko}$ | Homozygote | 25.5 | 181 | 25.9 | 95 | 95 | 95 | 95 | 95 | 95 | 95 | 95 | 95 |
| DMKO37.1d | 23/02/2015 | 26/08/2014 | *Mir183/96*$^{dko}$ | Homozygote | 33.1 | 181 | 25.9 | 95 | 95 | 95 | 95 | 95 | 95 | 95 | 95 | 95 |
| DMKO37.1f | 23/02/2015 | 26/08/2014 | *Mir183/96*$^{dko}$ | Homozygote | 25.6 | 181 | 25.9 | 95 | 95 | 95 | 95 | 95 | 95 | 95 | 95 | 95 |
| DMKO34.3a | 08/04/2015 | 08/10/2014 | *Mir183/96*$^{dko}$ | Homozygote | 36.2 | 182 | 26.0 | 95 | 95 | 95 | 95 | 95 | 95 | 95 | 95 | 95 |
| DMKO34.3b | 08/04/2015 | 08/10/2014 | *Mir183/96*$^{dko}$ | Homozygote | 36.1 | 182 | 26.0 | 95 | 95 | 95 | 95 | 95 | 95 | 95 | 95 | 95 |
| DMKO37.3h | 17/04/2015 | 20/10/2014 | *Mir183/96*$^{dko}$ | Homozygote | 34.4 | 179 | 25.6 | 95 | 95 | 95 | 95 | 95 | 95 | 95 | 95 | 95 |
| DMKO39.1f | 30/06/2015 | 01/01/2015 | *Mir183/96*$^{dko}$ | Homozygote | | 180 | 25.7 | 95 | 95 | 95 | 95 | 95 | 95 | 95 | 95 | 95 |
| DMKO41.1e | 30/06/2015 | 01/01/2015 | *Mir183/96*$^{dko}$ | Homozygote | | 180 | 25.7 | 95 | 95 | 95 | 95 | 95 | 95 | 95 | 95 | 95 |

**ABR thresholds for Fig. S3**

| Mouse ID | Experiment date | Date of birth | Allele | Genotype | Weight (g) | Age (days) | Age (weeks) | Click threshold | 3kHz threshold | 6kHz threshold | 12kHz threshold | 18kHz threshold | 24kHz threshold | 30kHz threshold | 36kHz threshold | 42kHz threshold |
|---|---|---|---|---|---|---|---|---|---|---|---|---|---|---|---|---|
| DMOM56.1b | 04/05/2016 | 20/04/2016 | *Mir182*$^{ko}$ | Wildtype | 6.9 | 14 | 2.0 | 40 | 95 | 75 | 45 | 60 | 40 | 75 | 85 | 95 |
| DMOM56.1e | 04/05/2016 | 20/04/2016 | *Mir182*$^{ko}$ | Wildtype | 6.4 | 14 | 2.0 | 50 | 95 | 75 | 55 | 45 | 45 | 70 | 80 | 95 |
| DMOM64.1b | 04/05/2017 | 20/04/2017 | *Mir182*$^{ko}$ | Wildtype | 7.5 | 14 | 2.0 | 45 | 95 | 70 | 35 | 30 | 30 | 40 | 55 | 80 |
| DMOM64.1e | 04/05/2017 | 20/04/2017 | *Mir182*$^{ko}$ | Wildtype | 6.8 | 14 | 2.0 | 35 | 95 | 75 | 25 | 35 | 25 | 35 | 45 | 60 |
| DMOM64.1d | 04/05/2017 | 20/04/2017 | *Mir182*$^{ko}$ | Wildtype | 7.0 | 14 | 2.0 | 40 | 95 | 70 | 40 | 35 | 35 | 35 | 55 | 75 |
| DMOM64.1a | 04/05/2017 | 20/04/2017 | *Mir182*$^{ko}$ | Wildtype | 7.1 | 14 | 2.0 | 45 | 95 | 85 | 35 | 40 | 50 | 55 | 60 | 95 |
| DMOM56.1d | 04/05/2016 | 20/04/2016 | *Mir182*$^{ko}$ | Heterozygote | 7.4 | 14 | 2.0 | 45 | 95 | 85 | 60 | 60 | 75 | 80 | 95 | 95 |
| DMOM56.1f | 04/05/2016 | 20/04/2016 | *Mir182*$^{ko}$ | Heterozygote | 7.2 | 14 | 2.0 | 40 | 95 | 80 | 55 | 40 | 35 | 55 | 70 | 95 |
| DMOM60.2f | 18/11/2016 | 04/11/2016 | *Mir182*$^{ko}$ | Heterozygote | 5.9 | 14 | 2.0 | 45 | 95 | 90 | 60 | 55 | 50 | 65 | 95 | 95 |
| DMOM60.2b | 18/11/2016 | 04/11/2016 | *Mir182*$^{ko}$ | Heterozygote | 6.2 | 14 | 2.0 | 55 | 95 | 70 | 55 | 50 | 55 | 65 | 80 | 95 |
| DMOM60.2i | 18/11/2016 | 04/11/2016 | *Mir182*$^{ko}$ | Heterozygote | 5.6 | 14 | 2.0 | 85 | 95 | 95 | 80 | 85 | 95 | 95 | 95 | 95 |
| DMOM58.4a | 23/11/2016 | 09/11/2016 | *Mir182*$^{ko}$ | Heterozygote | 7.7 | 14 | 2.0 | 45 | 95 | 80 | 60 | 50 | 45 | 50 | 85 | 95 |
| DMOM58.4b | 23/11/2016 | 09/11/2016 | *Mir182*$^{ko}$ | Heterozygote | 6.5 | 14 | 2.0 | 35 | 95 | 75 | 40 | 50 | 40 | 45 | 55 | 95 |
| DMOM58.4c | 23/11/2016 | 09/11/2016 | *Mir182*$^{ko}$ | Heterozygote | 7 | 14 | 2.0 | 40 | 95 | 80 | 50 | 50 | 40 | 45 | 90 | 95 |
| DMOM58.4e | 23/11/2016 | 09/11/2016 | *Mir182*$^{ko}$ | Heterozygote | 6.6 | 14 | 2.0 | 40 | 95 | 70 | 55 | 45 | 40 | 60 | 70 | 95 |
| DMOM58.4g | 23/11/2016 | 09/11/2016 | *Mir182*$^{ko}$ | Heterozygote | 6.1 | 14 | 2.0 | 35 | 90 | 75 | 35 | 35 | 30 | 35 | 50 | 55 |
| DMOM56.1a | 04/05/2016 | 20/04/2016 | *Mir182*$^{ko}$ | Homozygote | 6.8 | 14 | 2.0 | 80 | 95 | 95 | 85 | 85 | 95 | 95 | 95 | 95 |
| DMOM56.1c | 04/05/2016 | 20/04/2016 | *Mir182*$^{ko}$ | Homozygote | 6.4 | 14 | 2.0 | 65 | 95 | 95 | 80 | 75 | 95 | 95 | 95 | 95 |
| DMOM56.1g | 04/05/2016 | 20/04/2016 | *Mir182*$^{ko}$ | Homozygote | 6.7 | 14 | 2.0 | 40 | 95 | 85 | 55 | 45 | 35 | 50 | 75 | 95 |
| DMOM56.1i | 04/05/2016 | 20/04/2016 | *Mir182*$^{ko}$ | Homozygote | 7.5 | 14 | 2.0 | 35 | 95 | 85 | 55 | 45 | 45 | 55 | 60 | 95 |
| DMOM60.2a | 18/11/2016 | 04/11/2016 | *Mir182*$^{ko}$ | Homozygote | 5.5 | 14 | 2.0 | 95 | 95 | 95 | 95 | 95 | 95 | 95 | 95 | 95 |
| DMOM60.2c | 18/11/2016 | 04/11/2016 | *Mir182*$^{ko}$ | Homozygote | 6.3 | 14 | 2.0 | 45 | 95 | 85 | 60 | 45 | 45 | 60 | 95 | 95 |
| DMOM60.2d | 18/11/2016 | 04/11/2016 | *Mir182*$^{ko}$ | Homozygote | 6.2 | 14 | 2.0 | 50 | 95 | 85 | 45 | 45 | 45 | 65 | 85 | 95 |
| DMOM60.2e | 18/11/2016 | 04/11/2016 | *Mir182*$^{ko}$ | Homozygote | 5.8 | 14 | 2.0 | 60 | 95 | 90 | 70 | 60 | 70 | 85 | 95 | 95 |
| DMOM60.2k | 18/11/2016 | 04/11/2016 | *Mir182*$^{ko}$ | Homozygote | 5.5 | 14 | 2.0 | 95 | 95 | 95 | 95 | 95 | 95 | 95 | 95 | 95 |
| DMOM58.4d | 23/11/2016 | 09/11/2016 | *Mir182*$^{ko}$ | Homozygote | 7.2 | 14 | 2.0 | 40 | 95 | 90 | 45 | 55 | 35 | 50 | 60 | 95 |
| DMOM58.4f | 23/11/2016 | 09/11/2016 | *Mir182*$^{ko}$ | Homozygote | 6.4 | 14 | 2.0 | 40 | 90 | 80 | 65 | 55 | 40 | 55 | 95 | 95 |
| DMOM58.4h | 23/11/2016 | 09/11/2016 | *Mir182*$^{ko}$ | Homozygote | 6.4 | 14 | 2.0 | 60 | 95 | 90 | 70 | 85 | 95 | 95 | 95 | 95 |
| DMOM58.4i | 23/11/2016 | 09/11/2016 | *Mir182*$^{ko}$ | Homozygote | 7.1 | 14 | 2.0 | 35 | 90 | 80 | 40 | 45 | 25 | 45 | 55 | 95 |
| DMOM55.4g | 11/04/2016 | 21/03/2016 | *Mir182*$^{ko}$ | Wildtype | 9.9 | 21 | 3.0 | 25 | 60 | 25 | 20 | 20 | 25 | 45 | 40 | 50 |
| DMOM55.4c | 11/04/2016 | 21/03/2016 | *Mir182*$^{ko}$ | Wildtype | 9.5 | 21 | 3.0 | 25 | 70 | 45 | 15 | 20 | 20 | 35 | 45 | 65 |
| DMOM56.4e | 18/07/2016 | 27/06/2016 | *Mir182*$^{ko}$ | Wildtype | 11 | 21 | 3.0 | 30 | 65 | 40 | 15 | 20 | 30 | 35 | 45 | 75 |
| DMOM56.4d | 18/07/2016 | 27/06/2016 | *Mir182*$^{ko}$ | Wildtype | 11.5 | 21 | 3.0 | 30 | 75 | 35 | 15 | 25 | 25 | 35 | 55 | 70 |
| DMOM58.2c | 13/10/2016 | 22/09/2016 | *Mir182*$^{ko}$ | Wildtype | 10.7 | 21 | 3.0 | 30 | 70 | 35 | 15 | 20 | 25 | 45 | 40 | 55 |
| DMOM64.1i | 11/05/2017 | 20/04/2017 | *Mir182*$^{ko}$ | Wildtype | 10.3 | 21 | 3.0 | 25 | 75 | 35 | 25 | 25 | 20 | 35 | 45 | 50 |
| DMOM55.4a | 11/04/2016 | 21/03/2016 | *Mir182*$^{ko}$ | Heterozygote | 9.8 | 21 | 3.0 | 25 | 75 | 40 | 20 | 20 | 20 | 35 | 55 | 95 |
| DMOM55.4d | 11/04/2016 | 21/03/2016 | *Mir182*$^{ko}$ | Heterozygote | 9.9 | 21 | 3.0 | 30 | 75 | 35 | 15 | 30 | 25 | 55 | 40 | 50 |
| DMOM55.4e | 11/04/2016 | 21/03/2016 | *Mir182*$^{ko}$ | Heterozygote | 9.9 | 21 | 3.0 | 25 | 55 | 35 | 10 | 20 | 20 | 40 | 75 | 85 |
| DMOM55.4h | 11/04/2016 | 21/03/2016 | *Mir182*$^{ko}$ | Heterozygote | 9.1 | 21 | 3.0 | 25 | 65 | 30 | 15 | 20 | 25 | 40 | 65 | 95 |
| DMOM58.2a | 13/10/2016 | 22/09/2016 | *Mir182*$^{ko}$ | Heterozygote | 10.5 | 21 | 3.0 | 25 | 65 | 25 | 5 | 20 | 25 | 35 | 50 | 75 |
| DMOM58.2b | 13/10/2016 | 22/09/2016 | *Mir182*$^{ko}$ | Heterozygote | 10.5 | 21 | 3.0 | 25 | 60 | 35 | 10 | 20 | 20 | 35 | 40 | 50 |
| DMOM58.2d | 13/10/2016 | 22/09/2016 | *Mir182*$^{ko}$ | Heterozygote | 9.1 | 21 | 3.0 | 25 | 80 | 45 | 15 | 25 | 25 | 35 | 45 | 50 |
| DMOM58.2e | 13/10/2016 | 22/09/2016 | *Mir182*$^{ko}$ | Heterozygote | 9.2 | 21 | 3.0 | 25 | 70 | 30 | 5 | 20 | 20 | 30 | 40 | 50 |
| DMOM58.2f | 13/10/2016 | 22/09/2016 | *Mir182*$^{ko}$ | Heterozygote | 10.3 | 21 | 3.0 | 25 | 75 | 50 | 20 | 30 | 25 | 35 | 45 | 55 |
| DMOM63.2b | 08/05/2017 | 17/04/2017 | *Mir182*$^{ko}$ | Heterozygote | 10.6 | 21 | 3.0 | 30 | 65 | 55 | 20 | 25 | 20 | 35 | 45 | 55 |
| DMOM64.1c | 11/05/2017 | 20/04/2017 | *Mir182*$^{ko}$ | Heterozygote | 10.3 | 21 | 3.0 | 25 | 70 | 40 | 15 | 30 | 30 | 40 | 50 | 70 |
| DMOM55.4b | 11/04/2016 | 21/03/2016 | *Mir182*$^{ko}$ | Homozygote | 9.8 | 21 | 3.0 | 35 | 50 | 30 | 15 | 25 | 30 | 50 | 95 | 95 |
| DMOM55.4f | 11/04/2016 | 21/03/2016 | *Mir182*$^{ko}$ | Homozygote | 9.6 | 21 | 3.0 | 35 | 70 | 35 | 20 | 25 | 30 | 65 | 95 | 95 |
| DMOM55.4i | 11/04/2016 | 21/03/2016 | *Mir182*$^{ko}$ | Homozygote | 9.6 | 21 | 3.0 | 25 | 65 | 45 | 15 | 15 | 25 | 50 | 90 | 95 |
| DMOM56.4f | 18/07/2016 | 27/06/2016 | *Mir182*$^{ko}$ | Homozygote | 10.9 | 21 | 3.0 | 30 | 55 | 35 | 15 | 25 | 35 | 75 | 95 | 95 |
| DMOM63.2c | 08/05/2017 | 17/04/2017 | *Mir182*$^{ko}$ | Homozygote | 6.4 | 21 | 3.0 | 35 | 60 | 40 | 20 | 30 | 30 | 35 | 60 | 90 |
| DMOM63.2a | 08/05/2017 | 17/04/2017 | *Mir182*$^{ko}$ | Homozygote | 9.6 | 21 | 3.0 | 25 | 65 | 40 | 20 | 25 | 20 | 25 | 55 | 95 |
| DMOM63.2d | 08/05/2017 | 17/04/2017 | *Mir182*$^{ko}$ | Homozygote | 10.2 | 21 | 3.0 | 30 | 60 | 50 | 20 | 25 | 30 | 40 | 50 | 75 |
| DMOM35.2a | 08/10/2013 | 10/09/2013 | *Mir182*$^{ko}$ | Wildtype | 17.1 | 28 | 4.0 | 35 | 70 | 30 | 30 | 30 | 35 | 40 | 55 | 80 |
| DMOM35.2f | 08/10/2013 | 10/09/2013 | *Mir182*$^{ko}$ | Wildtype | 15 | 28 | 4.0 | 30 | 55 | 30 | 20 | 30 | 20 | 50 | 90 | 90 |
| DMOM36.2i | 09/10/2013 | 11/09/2013 | *Mir182*$^{ko}$ | Wildtype | 15.4 | 28 | 4.0 | 20 | 80 | 45 | 20 | 25 | 30 | 50 | 90 | 95 |
| DMOM36.2e | 09/10/2013 | 11/09/2013 | *Mir182*$^{ko}$ | Wildtype | 16.9 | 28 | 4.0 | 35 | 50 | 50 | 30 | 30 | 40 | 55 | 90 | 95 |
| DMOM36.2d | 09/10/2013 | 11/09/2013 | *Mir182*$^{ko}$ | Wildtype | 16 | 28 | 4.0 | 30 | 85 | 40 | 25 | 35 | 35 | 45 | 75 | 95 |
| DMOM31.5h | 10/10/2013 | 12/09/2013 | *Mir182*$^{ko}$ | Wildtype | 15.1 | 28 | 4.0 | 25 | 65 | 40 | 15 | 30 | 20 | 35 | 50 | 95 |
| DMOM34.2b | 16/10/2013 | 18/09/2013 | *Mir182*$^{ko}$ | Wildtype | 15.7 | 28 | 4.0 | 30 | 55 | 35 | 15 | 15 | 30 | 80 | 95 | 95 |
| DMOM34.2c | 16/10/2013 | 18/09/2013 | *Mir182*$^{ko}$ | Wildtype | 16 | 28 | 4.0 | 30 | 55 | 35 | 10 | 15 | 30 | 60 | 95 | 95 |
| DMOM39.2b | 29/04/2014 | 01/04/2014 | *Mir182*$^{ko}$ | Wildtype | 16.7 | 28 | 4.0 | 20 | 50 | 35 | 15 | 20 | 35 | 70 | 95 | 95 |
| DMOM39.2e | 29/04/2014 | 01/04/2014 | *Mir182*$^{ko}$ | Wildtype | 14.7 | 28 | 4.0 | 25 | 55 | 30 | 10 | 15 | 15 | 35 | 95 | 85 |
| DMOM47.3f | 04/02/2015 | 07/01/2015 | *Mir182*$^{ko}$ | Wildtype | 15.3 | 28 | 4.0 | 25 | 70 | 45 | 25 | 20 | 35 | 40 | 70 | 95 |
| DMOM47.3e | 04/02/2015 | 07/01/2015 | *Mir182*$^{ko}$ | Wildtype | 14 | 28 | 4.0 | 30 | 75 | 40 | 20 | 20 | 35 | 40 | 80 | 90 |
| DMOM37.1c | 18/09/2013 | 22/08/2013 | *Mir182*$^{ko}$ | Heterozygote | 13.8 | 27 | 3.9 | 30 | 50 | 30 | 10 | 15 | 30 | 35 | 80 | 85 |
| DMOM34.1a | 24/09/2013 | 27/08/2013 | *Mir182*$^{ko}$ | Heterozygote | 17.2 | 28 | 4.0 | 25 | 70 | 35 | 20 | 25 | 35 | 60 | 95 | 95 |
| DMOM35.2c | 08/10/2013 | 10/09/2013 | *Mir182*$^{ko}$ | Heterozygote | 16.2 | 28 | 4.0 | 20 | 80 | 55 | 15 | 25 | 30 | 85 | 95 | 95 |
| DMOM35.2b | 08/10/2013 | 10/09/2013 | *Mir182*$^{ko}$ | Heterozygote | 14.7 | 28 | 4.0 | 25 | 80 | 35 | 15 | 30 | 40 | 90 | 95 | 85 |
| DMOM35.2d | 08/10/2013 | 10/09/2013 | *Mir182*$^{ko}$ | Heterozygote | 16.6 | 28 | 4.0 | 35 | 60 | 45 | 30 | 40 | 35 | 45 | 90 | 90 |
| DMOM35.2h | 08/10/2013 | 10/09/2013 | *Mir182*$^{ko}$ | Heterozygote | 15.1 | 28 | 4.0 | 40 | 95 | 70 | 55 | 50 | 55 | 85 | 95 | 95 |
| DMOM35.2j | 08/10/2013 | 10/09/2013 | *Mir182*$^{ko}$ | Heterozygote | 13.7 | 28 | 4.0 | 25 | 50 | 45 | 10 | 20 | 25 | 60 | 90 | 90 |
| DMOM35.2e | 08/10/2013 | 10/09/2013 | *Mir182*$^{ko}$ | Heterozygote | 15 | 28 | 4.0 | 30 | 75 | 30 | 20 | 25 | 35 | 90 | 95 | 95 |
| DMOM35.2g | 08/10/2013 | 10/09/2013 | *Mir182*$^{ko}$ | Heterozygote | 15.9 | 28 | 4.0 | 25 | 95 | 40 | 20 | 25 | 25 | 30 | 70 | 95 |
| DMOM36.2f | 09/10/2013 | 11/09/2013 | *Mir182*$^{ko}$ | Heterozygote | 15.8 | 28 | 4.0 | 35 | 60 | 40 | 25 | 30 | 30 | 85 | 95 | 95 |
| DMOM36.2j | 09/10/2013 | 11/09/2013 | *Mir182*$^{ko}$ | Heterozygote | 15.9 | 28 | 4.0 | 35 | 55 | 40 | 20 | 35 | 40 | 75 | 95 | 95 |
| DMOM36.2g | 09/10/2013 | 11/09/2013 | *Mir182*$^{ko}$ | Heterozygote | 15.3 | 28 | 4.0 | 30 | 70 | 55 | 20 | 30 | 35 | 65 | 95 | 95 |
| DMOM36.2b | 09/10/2013 | 11/09/2013 | *Mir182*$^{ko}$ | Heterozygote | 17 | 28 | 4.0 | 30 | 70 | 45 | 20 | 30 | 30 | 55 | 90 | 95 |
| DMOM36.2c | 09/10/2013 | 11/09/2013 | *Mir182*$^{ko}$ | Heterozygote | 15.5 | 28 | 4.0 | 25 | 60 | 45 | 20 | 40 | 40 | 75 | 95 | 95 |
| DMOM36.2a | 09/10/2013 | 11/09/2013 | *Mir182*$^{ko}$ | Heterozygote | 16.6 | 28 | 4.0 | 25 | 75 | 40 | 15 | 30 | 25 | 50 | 45 | 55 |
| DMOM31.5e | 10/10/2013 | 12/09/2013 | *Mir182*$^{ko}$ | Heterozygote | 17.5 | 28 | 4.0 | 30 | 65 | 50 | 20 | 30 | 30 | 35 | 50 | 80 |
| DMOM31.5c | 10/10/2013 | 12/09/2013 | *Mir182*$^{ko}$ | Heterozygote | 16 | 28 | 4.0 | 30 | 95 | 45 | 25 | 25 | 35 | 75 | 95 | 95 |
| DMOM31.5d | 10/10/2013 | 12/09/2013 | *Mir182*$^{ko}$ | Heterozygote | 17.2 | 28 | 4.0 | 30 | 60 | 35 | 10 | 25 | 30 | 70 | 95 | 95 |
| DMOM34.2a | 16/10/2013 | 18/09/2013 | *Mir182*$^{ko}$ | Heterozygote | 16.7 | 28 | 4.0 | 30 | 75 | 55 | 15 | 30 | 75 | 95 | 95 | 95 |
| DMOM39.2a | 29/04/2014 | 01/04/2014 | *Mir182*$^{ko}$ | Heterozygote | 14.7 | 28 | 4.0 | 25 | 80 | 30 | 5 | 20 | 30 | 60 | 95 | 95 |
| DMOM39.2c | 29/04/2014 | 01/04/2014 | *Mir182*$^{ko}$ | Heterozygote | 16.4 | 28 | 4.0 | 20 | 55 | 30 | 10 | 10 | 30 | 50 | 95 | 85 |
| DMOM47.3b | 04/02/2015 | 07/01/2015 | *Mir182*$^{ko}$ | Heterozygote | 16.8 | 28 | 4.0 | 30 | 60 | 40 | 15 | 25 | 35 | 60 | 95 | 95 |
| DMOM37.1a | 18/09/2013 | 22/08/2013 | *Mir182*$^{ko}$ | Homozygote | 13.9 | 27 | 3.9 | 30 | 55 | 45 | 15 | 20 | 55 | 85 | 85 | 80 |
| DMOM37.1b | 18/09/2013 | 22/08/2013 | *Mir182*$^{ko}$ | Homozygote | 13.3 | 27 | 3.9 | 25 | 50 | 30 | 10 | 25 | 70 | 85 | 90 | 85 |
| DMOM34.1b | 24/09/2013 | 27/08/2013 | *Mir182*$^{ko}$ | Homozygote | 10.1 | 28 | 4.0 | 30 | 70 | 45 | 15 | 25 | 30 | 45 | 65 | 65 |
| DMOM35.2i | 08/10/2013 | 10/09/2013 | *Mir182*$^{ko}$ | Homozygote | 13.7 | 28 | 4.0 | 35 | 65 | 55 | 35 | 65 | 90 | 95 | 95 | 95 |
| DMOM36.2h | 09/10/2013 | 11/09/2013 | *Mir182*$^{ko}$ | Homozygote | 16.2 | 28 | 4.0 | 35 | 60 | 45 | 25 | 35 | 80 | 85 | 95 | 95 |
| DMOM31.5f | 10/10/2013 | 12/09/2013 | *Mir182*$^{ko}$ | Homozygote | 9.4 | 28 | 4.0 | 40 | 85 | 70 | 45 | 85 | 95 | 95 | 95 | 95 |
| DMOM31.5g | 10/10/2013 | 12/09/2013 | *Mir182*$^{ko}$ | Homozygote | 17.3 | 28 | 4.0 | 30 | 60 | 55 | 25 | 30 | 50 | 90 | 95 | 95 |
| DMOM31.5a | 10/10/2013 | 12/09/2013 | *Mir182*$^{ko}$ | Homozygote | 18.1 | 28 | 4.0 | 30 | 85 | 35 | 15 | 25 | 40 | 95 | 95 | 95 |
| DMOM31.5b | 10/10/2013 | 12/09/2013 | *Mir182*$^{ko}$ | Homozygote | 17.8 | 28 | 4.0 | 30 | 70 | 40 | 15 | 25 | 35 | 90 | 95 | 95 |
| DMOM34.2e | 16/10/2013 | 18/09/2013 | *Mir182*$^{ko}$ | Homozygote | 14.7 | 28 | 4.0 | 25 | 60 | 35 | 15 | 30 | 80 | 90 | 95 | 95 |
| DMOM34.2d | 16/10/2013 | 18/09/2013 | *Mir182*$^{ko}$ | Homozygote | 16.3 | 28 | 4.0 | 20 | 65 | 35 | 10 | 25 | 25 | 50 | 85 | 95 |
| DMOM39.2d | 29/04/2014 | 01/04/2014 | *Mir182*$^{ko}$ | Homozygote | 14.6 | 28 | 4.0 | 20 | 50 | 35 | 5 | 20 | 45 | 90 | 85 | 85 |
| DMOM47.3a | 04/02/2015 | 07/01/2015 | *Mir182*$^{ko}$ | Homozygote | 17.3 | 28 | 4.0 | 25 | 65 | 35 | 20 | 40 | 70 | 95 | 95 | 95 |
| DMOM47.3g | 04/02/2015 | 07/01/2015 | *Mir182*$^{ko}$ | Homozygote | 14.3 | 28 | 4.0 | 25 | 75 | 40 | 25 | 25 | 75 | 85 | 95 | 95 |
| DMOM47.3d | 04/02/2015 | 07/01/2015 | *Mir182*$^{ko}$ | Homozygote | 13.4 | 28 | 4.0 | 30 | 70 | 30 | 15 | 50 | 85 | 90 | 95 | 95 |
| DMOM34.2c | 14/11/2013 | 18/09/2013 | *Mir182*$^{ko}$ | Wildtype | 23.8 | 57 | 8.1 | 25 | 55 | 25 | 10 | 20 | 25 | 35 | 80 | 95 |
| DMOM34.2b | 14/11/2013 | 18/09/2013 | *Mir182*$^{ko}$ | Wildtype | 22.1 | 57 | 8.1 | 25 | 80 | 40 | 15 | 25 | 25 | 45 | 95 | 95 |
| DMOM39.2e | 28/05/2014 | 01/04/2014 | *Mir182*$^{ko}$ | Wildtype | 22.1 | 57 | 8.1 | 20 | 50 | 30 | 10 | 15 | 20 | 30 | 55 | 85 |
| DMOM39.2b | 28/05/2014 | 01/04/2014 | *Mir182*$^{ko}$ | Wildtype | 24.8 | 57 | 8.1 | 20 | 55 | 30 | 15 | 25 | 15 | 50 | 90 | 95 |
| DMOM47.3f | 04/03/2015 | 07/01/2015 | *Mir182*$^{ko}$ | Wildtype | 20.8 | 56 | 8.0 | 20 | 65 | 50 | 15 | 30 | 30 | 45 | 75 | 95 |
| DMOM47.3e | 04/03/2015 | 07/01/2015 | *Mir182*$^{ko}$ | Wildtype | 12.9 | 56 | 8.0 | 20 | 55 | 30 | 10 | 10 | 15 | 25 | 55 | 90 |
| DMOM34.2a | 14/11/2013 | 18/09/2013 | *Mir182*$^{ko}$ | Heterozygote | 27 | 57 | 8.1 | 25 | 55 | 30 | 10 | 20 | 35 | 85 | 95 | 95 |
| DMOM37.1c | 17/10/2013 | 22/08/2013 | *Mir182*$^{ko}$ | Heterozygote | 20.4 | 56 | 8.0 | 30 | 60 | 40 | 15 | 25 | 30 | 35 | 70 | 95 |
| DMOM34.1a | 22/10/2013 | 27/08/2013 | *Mir182*$^{ko}$ | Heterozygote | 28.2 | 56 | 8.0 | 20 | 55 | 25 | 10 | 15 | 15 | 25 | 60 | 70 |
| DMOM39.2a | 28/05/2014 | 01/04/2014 | *Mir182*$^{ko}$ | Heterozygote | 23.5 | 57 | 8.1 | 15 | 45 | 25 | 10 | 15 | 20 | 50 | 80 | 80 |
| DMOM39.2c | 28/05/2014 | 01/04/2014 | *Mir182*$^{ko}$ | Heterozygote | 26.7 | 57 | 8.1 | 20 | 60 | 30 | 5 | 10 | 20 | 35 | 60 | 80 |
| DMOM47.3b | 04/03/2015 | 07/01/2015 | *Mir182*$^{ko}$ | Heterozygote | 24.9 | 56 | 8.0 | 25 | 60 | 45 | 10 | 20 | 30 | 65 | 95 | 95 |
| DMOM34.2e | 14/11/2013 | 18/09/2013 | *Mir182*$^{ko}$ | Homozygote | 20.7 | 57 | 8.1 | 25 | 60 | 30 | 10 | 25 | 50 | 80 | 95 | 95 |
| DMOM37.1b | 17/10/2013 | 22/08/2013 | *Mir182*$^{ko}$ | Homozygote | 20.9 | 56 | 8.0 | 30 | 75 | 40 | 20 | 25 | 55 | 85 | 95 | 95 |
| DMOM37.1a | 17/10/2013 | 22/08/2013 | *Mir182*$^{ko}$ | Homozygote | 19.2 | 56 | 8.0 | 25 | 80 | 40 | 20 | 30 | 30 | 70 | 95 | 95 |
| DMOM34.1b | 23/10/2013 | 27/08/2013 | *Mir182*$^{ko}$ | Homozygote | 18.4 | 57 | 8.1 | 35 | 65 | 40 | 15 | 25 | 35 | 70 | 95 | 95 |
| DMOM34.2d | 14/11/2013 | 18/09/2013 | *Mir182*$^{ko}$ | Homozygote | 25.2 | 57 | 8.1 | 25 | 60 | 35 | 10 | 15 | 30 | 40 | 90 | 80 |
| DMOM39.2d | 28/05/2014 | 01/04/2014 | *Mir182*$^{ko}$ | Homozygote | 20.3 | 57 | 8.1 | 25 | 50 | 25 | 10 | 15 | 30 | 65 | 85 | 80 |
| DMOM47.3a | 04/03/2015 | 07/01/2015 | *Mir182*$^{ko}$ | Homozygote | 24.9 | 56 | 8.0 | 30 | 70 | 45 | 25 | 40 | 75 | 95 | 95 | 95 |
| DMOM47.3g | 04/03/2015 | 07/01/2015 | *Mir182*$^{ko}$ | Homozygote | 20.7 | 56 | 8.0 | 25 | 60 | 35 | 15 | 25 | 55 | 95 | 95 | 95 |
| DMOM47.3d | 04/03/2015 | 07/01/2015 | *Mir182*$^{ko}$ | Homozygote | 18.1 | 56 | 8.0 | 40 | 70 | 40 | 25 | 70 | 95 | 95 | 95 | 95 |
| DMOM34.6c | 08/04/2014 | 06/01/2014 | *Mir182*$^{ko}$ | Wildtype | 28.8 | 92 | 13.1 | 25 | 60 | 25 | 10 | 20 | 30 | 65 | 95 | 95 |
| DMOM34.6d | 08/04/2014 | 06/01/2014 | *Mir182*$^{ko}$ | Wildtype | 33 | 92 | 13.1 | 20 | 50 | 30 | 5 | 10 | 20 | 40 | 95 | 95 |
| DMOM39.1d | 14/05/2014 | 11/02/2014 | *Mir182*$^{ko}$ | Wildtype | 33.2 | 92 | 13.1 | 25 | 40 | 20 | 10 | 25 | 35 | 90 | 95 | 95 |
| DMOM39.2e | 01/07/2014 | 01/04/2014 | *Mir182*$^{ko}$ | Wildtype | 29.2 | 91 | 13.0 | 20 | 55 | 25 | 15 | 15 | 30 | 60 | 95 | 95 |
| DMOM47.3f | 08/04/2015 | 07/01/2015 | *Mir182*$^{ko}$ | Wildtype | 24.7 | 91 | 13.0 | 25 | 70 | 40 | 15 | 20 | 30 | 80 | 95 | 95 |
| DMOM63.1d | 23/06/2017 | 25/03/2017 | *Mir182*$^{ko}$ | Wildtype | 30.8 | 90 | 12.9 | 25 | 50 | 30 | 20 | 20 | 30 | 35 | 50 | 95 |
| DMOM39.1f | 14/05/2014 | 11/02/2014 | *Mir182*$^{ko}$ | Heterozygote | 30.8 | 92 | 13.1 | 25 | 50 | 30 | 15 | 20 | 75 | 95 | 95 | 95 |
| DMOM39.2c | 01/07/2014 | 01/04/2014 | *Mir182*$^{ko}$ | Heterozygote | 31.6 | 91 | 13.0 | 35 | 75 | 30 | 20 | 20 | 20 | 70 | 95 | 95 |
| DMOM39.2a | 01/07/2014 | 01/04/2014 | *Mir182*$^{ko}$ | Heterozygote | 29.1 | 91 | 13.0 | 20 | 55 | 25 | 10 | 15 | 35 | 95 | 95 | 95 |
| DMOM44.3a | 04/02/2015 | 06/11/2014 | *Mir182*$^{ko}$ | Heterozygote | 31.5 | 90 | 12.9 | 25 | 60 | 30 | 10 | 20 | 25 | 50 | 65 | 90 |
| DMOM44.3b | 04/02/2015 | 06/11/2014 | *Mir182*$^{ko}$ | Heterozygote | 33.3 | 90 | 12.9 | 20 | 65 | 30 | 5 | 15 | 20 | 50 | 35 | 50 |
| DMOM44.3c | 04/02/2015 | 06/11/2014 | *Mir182*$^{ko}$ | Heterozygote | 29.9 | 90 | 12.9 | 25 | 65 | 30 | 15 | 20 | 30 | 40 | 35 | 40 |
| DMOM47.3b | 08/04/2015 | 07/01/2015 | *Mir182*$^{ko}$ | Heterozygote | 28.7 | 91 | 13.0 | 30 | 65 | 35 | 20 | 25 | 55 | 95 | 95 | 95 |
| DMOM34.6a | 08/04/2014 | 06/01/2014 | *Mir182*$^{ko}$ | Homozygote | 36.1 | 92 | 13.1 | 25 | 80 | 35 | 10 | 25 | 95 | 95 | 95 | 95 |
| DMOM39.1b | 14/05/2014 | 11/02/2014 | *Mir182*$^{ko}$ | Homozygote | 37 | 92 | 13.1 | 30 | 55 | 25 | 15 | 50 | 70 | 95 | 95 | 95 |
| DMOM39.1c | 14/05/2014 | 11/02/2014 | *Mir182*$^{ko}$ | Homozygote | 34.3 | 92 | 13.1 | 30 | 70 | 30 | 10 | 35 | 95 | 95 | 95 | 95 |
| DMOM39.2d | 01/07/2014 | 01/04/2014 | *Mir182*$^{ko}$ | Homozygote | 23.9 | 91 | 13.0 | 30 | 65 | 25 | 10 | 20 | 75 | 90 | 95 | 95 |
| DMOM47.3a | 08/04/2015 | 07/01/2015 | *Mir182*$^{ko}$ | Homozygote | 28.6 | 91 | 13.0 | 30 | 90 | 65 | 20 | 40 | 60 | 95 | 95 | 95 |
| DMOM47.3d | 08/04/2015 | 07/01/2015 | *Mir182*$^{ko}$ | Homozygote | 20.7 | 91 | 13.0 | 40 | 85 | 40 | 45 | 95 | 95 | 95 | 95 | 95 |
| DMOM47.3g | 08/04/2015 | 07/01/2015 | *Mir182*$^{ko}$ | Homozygote | 23.8 | 91 | 13.0 | 35 | 75 | 35 | 25 | 95 | 95 | 95 | 95 | 95 |
| DMOM34.6d | 08/07/2014 | 06/01/2014 | *Mir182*$^{ko}$ | Wildtype | 43.1 | 183 | 26.1 | 25 | 80 | 30 | 10 | 15 | 30 | 85 | 95 | 95 |
| DMOM34.6c | 08/07/2014 | 06/01/2014 | *Mir182*$^{ko}$ | Wildtype | 38.8 | 183 | 26.1 | 15 | 50 | 30 | 20 | 30 | 55 | 95 | 95 | 95 |
| DMOM39.1d | 11/08/2014 | 11/02/2014 | *Mir182*$^{ko}$ | Wildtype | 42.4 | 181 | 25.9 | 50 | 70 | 50 | 55 | 75 | 95 | 95 | 95 | 95 |
| DMOM39.2e | 29/09/2014 | 01/04/2014 | *Mir182*$^{ko}$ | Wildtype | 42.8 | 181 | 25.9 | 20 | 75 | 35 | 15 | 30 | 45 | 95 | 95 | 95 |
| DMOM47.3f | 06/07/2015 | 07/01/2015 | *Mir182*$^{ko}$ | Wildtype | 41.4 | 180 | 25.7 | 25 | 80 | 65 | 20 | 30 | 75 | 95 | 95 | 95 |
| DMOM63.1d | 21/09/2017 | 25/03/2017 | *Mir182*$^{ko}$ | Wildtype | 37.0 | 180 | 25.7 | 25 | 60 | 35 | 10 | 20 | 25 | 55 | 70 | 95 |
| DMOM34.6b | 08/07/2014 | 06/01/2014 | *Mir182*$^{ko}$ | Heterozygote | 34.8 | 183 | 26.1 | 25 | 75 | 25 | 20 | 25 | 95 | 95 | 95 | 95 |
| DMOM34.6e | 08/07/2014 | 06/01/2014 | *Mir182*$^{ko}$ | Heterozygote | 31.8 | 183 | 26.1 | 25 | 80 | 45 | 15 | 10 | 55 | 95 | 95 | 95 |
| DMOM39.1f | 11/08/2014 | 11/02/2014 | *Mir182*$^{ko}$ | Heterozygote | 40.9 | 181 | 25.9 | 25 | 65 | 30 | 15 | 60 | 95 | 95 | 95 | 95 |
| DMOM39.2a | 29/09/2014 | 01/04/2014 | *Mir182*$^{ko}$ | Heterozygote | 38.3 | 181 | 25.9 | 25 | 75 | 30 | 5 | 25 | 95 | 95 | 95 | 95 |
| DMOM39.2c | 29/09/2014 | 01/04/2014 | *Mir182*$^{ko}$ | Heterozygote | 42.7 | 181 | 25.9 | 25 | 80 | 35 | 10 | 20 | 50 | 95 | 95 | 95 |
| DMOM44.3b | 06/05/2015 | 06/11/2014 | *Mir182*$^{ko}$ | Heterozygote | 40.8 | 181 | 25.9 | 25 | 80 | 45 | 15 | 25 | 25 | 30 | 40 | 50 |
| DMOM44.3a | 06/05/2015 | 06/11/2014 | *Mir182*$^{ko}$ | Heterozygote | 40.4 | 181 | 25.9 | 30 | 80 | 45 | 20 | 30 | 95 | 95 | 95 | 95 |
| DMOM44.3c | 06/05/2015 | 06/11/2014 | *Mir182*$^{ko}$ | Heterozygote | 41.1 | 181 | 25.9 | 20 | 80 | 50 | 20 | 30 | 40 | 40 | 45 | 55 |
| DMOM34.6a | 08/07/2014 | 06/01/2014 | *Mir182*$^{ko}$ | Homozygote | 46.8 | 183 | 26.1 | 35 | 80 | 45 | 30 | 95 | 95 | 95 | 95 | 95 |
| DMOM39.1c | 11/08/2014 | 11/02/2014 | *Mir182*$^{ko}$ | Homozygote | 46.4 | 181 | 25.9 | 45 | 85 | 60 | 50 | 95 | 95 | 95 | 95 | 95 |
| DMOM39.1b | 11/08/2014 | 11/02/2014 | *Mir182*$^{ko}$ | Homozygote | 47.1 | 181 | 25.9 | 25 | 70 | 45 | 30 | 95 | 95 | 95 | 95 | 95 |
| DMOM39.2d | 29/09/2014 | 01/04/2014 | *Mir182*$^{ko}$ | Homozygote | 33.7 | 181 | 25.9 | 45 | 80 | 50 | 40 | 95 | 95 | 95 | 95 | 95 |
| DMOM47.3d | 06/07/2015 | 07/01/2015 | *Mir182*$^{ko}$ | Homozygote | 29.2 | 180 | 25.7 | 55 | 80 | 50 | 95 | 95 | 95 | 95 | 95 | 95 |
| DMOM47.3g | 06/07/2015 | 07/01/2015 | *Mir182*$^{ko}$ | Homozygote | 35.3 | 180 | 25.7 | 40 | 80 | 35 | 40 | 95 | 95 | 95 | 95 | 95 |

**Waveform data for Fig. S4**

12kHz ABR , Threshold +20dB

| | *Mir183/96* $^{dko}$ Wildtype | | *Mir183/96* $^{dko}$ Heterozygote | |
|---|---|---|---|---|
| | Amplitude (µV) | | Amplitude (µV) | |
| Time (ms) | Mean | SD | Mean | SD |
| 0 | 0.096848172 | 0.172238297 | -0.048433818 | 0.22361604 |
| 0.01024 | 0.095891228 | 0.172800905 | -0.043074722 | 0.22223004 |
| 0.02048 | 0.095403605 | 0.174441305 | -0.038438347 | 0.22156425 |
| 0.03072 | 0.096137178 | 0.176747569 | -0.033956001 | 0.22196691 |
| 0.04096 | 0.097627216 | 0.178915352 | -0.029580846 | 0.2234001 |
| 0.0512 | 0.100022667 | 0.179860959 | -0.025493651 | 0.22518266 |
| 0.06144 | 0.102932988 | 0.180313765 | -0.021872902 | 0.22828993 |
| 0.07168 | 0.105116236 | 0.179643814 | -0.018883023 | 0.23072493 |
| 0.08192 | 0.10707329 | 0.177836917 | -0.016434281 | 0.23314875 |
| 0.09216 | 0.107224759 | 0.175721395 | -0.015431747 | 0.23461605 |
| 0.1024 | 0.105952465 | 0.172975176 | -0.015051493 | 0.23504922 |
| 0.11264 | 0.103250952 | 0.170984725 | -0.015955463 | 0.23449828 |
| 0.12288 | 0.099308667 | 0.16859782 | -0.017055569 | 0.23258972 |
| 0.13312 | 0.094429332 | 0.167973251 | -0.018844369 | 0.23007301 |
| 0.14336 | 0.089025766 | 0.167501846 | -0.020533871 | 0.22668646 |
| 0.1536 | 0.084439707 | 0.168230889 | -0.021711508 | 0.22399749 |
| 0.16384 | 0.080020587 | 0.169346148 | -0.022257814 | 0.2213498 |
| 0.174081 | 0.076675901 | 0.170030531 | -0.022238749 | 0.21837547 |
| 0.184321 | 0.074846759 | 0.170638434 | -0.021701032 | 0.21610614 |
| 0.194561 | 0.07348014 | 0.169207259 | -0.020515925 | 0.21389167 |
| 0.204801 | 0.073500569 | 0.167987955 | -0.019474206 | 0.21111543 |
| 0.215041 | 0.074035622 | 0.163980322 | -0.019134945 | 0.20868095 |
| 0.225281 | 0.075031998 | 0.15996719 | -0.019303011 | 0.20642401 |
| 0.235521 | 0.076631584 | 0.153730723 | -0.020849141 | 0.20471176 |
| 0.245761 | 0.078535503 | 0.147397618 | -0.023256493 | 0.20484048 |
| 0.256001 | 0.081101439 | 0.139750122 | -0.027106794 | 0.20577445 |
| 0.266241 | 0.08373323 | 0.132148504 | -0.031264089 | 0.2098273 |
| 0.276481 | 0.086846175 | 0.125043714 | -0.036004248 | 0.21519639 |
| 0.286721 | 0.089682881 | 0.119450067 | -0.040482976 | 0.22261844 |
| 0.296961 | 0.092527322 | 0.115889294 | -0.044712531 | 0.23116454 |
| 0.307201 | 0.094315081 | 0.114669757 | -0.048005165 | 0.23969896 |
| 0.317441 | 0.095458965 | 0.115760696 | -0.050464371 | 0.24780219 |
| 0.327681 | 0.095446492 | 0.117908663 | -0.052241594 | 0.25401777 |
| 0.337921 | 0.094195448 | 0.121242555 | -0.05305064 | 0.25835468 |
| 0.348161 | 0.09234321 | 0.124545302 | -0.053487518 | 0.25991206 |
| 0.358401 | 0.090085532 | 0.12857233 | -0.053416246 | 0.2599212 |
| 0.368641 | 0.087918071 | 0.131825314 | -0.053177735 | 0.25683111 |
| 0.378881 | 0.086317008 | 0.135551627 | -0.052767997 | 0.25384874 |
| 0.389121 | 0.084896555 | 0.139603319 | -0.052607856 | 0.24884873 |
| 0.399361 | 0.084662852 | 0.143812199 | -0.052850608 | 0.24501484 |
| 0.409601 | 0.084848256 | 0.148121068 | -0.053519745 | 0.24113899 |
| 0.419841 | 0.08505028 | 0.153243048 | -0.055090072 | 0.23816021 |
| 0.430081 | 0.085400944 | 0.158626104 | -0.057447314 | 0.23612865 |
| 0.440321 | 0.085192684 | 0.164035433 | -0.061101894 | 0.23513843 |
| 0.450561 | 0.084583529 | 0.16972857 | -0.065146218 | 0.23442334 |
| 0.460801 | 0.083327412 | 0.174925148 | -0.070210365 | 0.23453737 |
| 0.471041 | 0.081604779 | 0.179865004 | -0.075111642 | 0.23443119 |
| 0.481281 | 0.079824488 | 0.183466541 | -0.080113903 | 0.23477139 |
| 0.491521 | 0.078398348 | 0.186780521 | -0.083973352 | 0.23542891 |
| 0.501761 | 0.077399577 | 0.18845448 | -0.087043423 | 0.23536326 |
| 0.512001 | 0.077387546 | 0.18968183 | -0.089065033 | 0.23649063 |
| 0.522242 | 0.078366153 | 0.189285829 | -0.089091691 | 0.23697317 |
| 0.532482 | 0.080502568 | 0.188569332 | -0.088402206 | 0.23817475 |
| 0.542722 | 0.084143177 | 0.186044909 | -0.086254464 | 0.23943264 |
| 0.552962 | 0.088929916 | 0.183207101 | -0.083514822 | 0.2418731 |
| 0.563202 | 0.09426055 | 0.178739092 | -0.080202191 | 0.24435842 |
| 0.573442 | 0.100838654 | 0.174900517 | -0.07708163 | 0.24839648 |
| 0.583682 | 0.107838175 | 0.170428475 | -0.073485966 | 0.25333264 |
| 0.593922 | 0.114566173 | 0.167708754 | -0.071065845 | 0.25990722 |
| 0.604162 | 0.121414055 | 0.166310385 | -0.067809452 | 0.26732231 |
| 0.614402 | 0.127198298 | 0.167224899 | -0.065739101 | 0.27580117 |
| 0.624642 | 0.132410259 | 0.169624907 | -0.063022225 | 0.28480442 |
| 0.634882 | 0.135555008 | 0.173542524 | -0.060420949 | 0.29392937 |
| 0.645122 | 0.137488442 | 0.178297308 | -0.057244003 | 0.30235036 |
| 0.655362 | 0.137471559 | 0.182742756 | -0.053680361 | 0.31013816 |
| 0.665602 | 0.135907985 | 0.188050446 | -0.049096636 | 0.31680458 |
| 0.675842 | 0.131991339 | 0.192520688 | -0.043863448 | 0.32203389 |
| 0.686082 | 0.127465165 | 0.196524629 | -0.038209361 | 0.32550377 |
| 0.696322 | 0.120573705 | 0.20027892 | -0.031597375 | 0.32793303 |
| 0.706562 | 0.113190047 | 0.202918057 | -0.025125731 | 0.32886192 |
| 0.716802 | 0.104481997 | 0.204327013 | -0.018364159 | 0.32848193 |
| 0.727042 | 0.09494644 | 0.204542237 | -0.012521361 | 0.32772322 |
| 0.737282 | 0.085106422 | 0.202962646 | -0.007181175 | 0.32573017 |
| 0.747522 | 0.07478469 | 0.199941499 | -0.003299664 | 0.32395092 |
| 0.757762 | 0.06471544 | 0.195792322 | -0.000408884 | 0.32181733 |
| 0.768002 | 0.054258306 | 0.189791086 | 0.000214349 | 0.31967087 |
| 0.778242 | 0.044627728 | 0.183719943 | -5.2517E-05 | 0.31798417 |
| 0.788482 | 0.035143318 | 0.176230263 | -0.002731968 | 0.31658881 |
| 0.798722 | 0.027142309 | 0.169009252 | -0.006073007 | 0.31546297 |
| 0.808962 | 0.019100858 | 0.161195473 | -0.011387729 | 0.31489632 |
| 0.819202 | 0.012945229 | 0.153033116 | -0.016829322 | 0.31453333 |
| 0.829442 | 0.007784869 | 0.145285384 | -0.023106325 | 0.315016 |
| 0.839682 | 0.003829391 | 0.137074855 | -0.029101305 | 0.31527485 |
| 0.849922 | 0.001187204 | 0.129949677 | -0.034316683 | 0.31695283 |
| 0.860162 | 9.88079E-05 | 0.123426797 | -0.038458455 | 0.31769751 |
| 0.870403 | -0.000568313 | 0.119059576 | -0.04098608 | 0.3195203 |
| 0.880643 | 0.000148087 | 0.116365707 | -0.042269508 | 0.3210085 |
| 0.890883 | 0.001350828 | 0.11647908 | -0.041548112 | 0.32198324 |
| 0.901123 | 0.003172559 | 0.118637646 | -0.040220014 | 0.32233918 |
| 0.911363 | 0.005622126 | 0.123327745 | -0.037670818 | 0.32133286 |
| 0.921603 | 0.008651561 | 0.12900725 | -0.035284865 | 0.31961585 |
| 0.931843 | 0.012079557 | 0.136425103 | -0.032230396 | 0.31646922 |
| 0.942083 | 0.016050989 | 0.14418938 | -0.030069603 | 0.31270554 |
| 0.952323 | 0.020017134 | 0.152379901 | -0.027825977 | 0.30808839 |
| 0.962563 | 0.024258293 | 0.160544043 | -0.025695586 | 0.30418402 |
| 0.972803 | 0.0275666 | 0.16869423 | -0.024132959 | 0.30053128 |
| 0.983043 | 0.030331077 | 0.177366314 | -0.022200339 | 0.29781 |
| 0.993283 | 0.031981252 | 0.184943374 | -0.021288823 | 0.29658694 |
| 1.003523 | 0.032159815 | 0.192905801 | -0.019865519 | 0.29612736 |
| 1.013763 | 0.030851667 | 0.200288571 | -0.019843745 | 0.29710562 |
| 1.024003 | 0.028627708 | 0.206346092 | -0.01984463 | 0.29858621 |
| 1.034243 | 0.024941062 | 0.211103953 | -0.021306687 | 0.30103367 |
| 1.044483 | 0.020894681 | 0.214839451 | -0.02341663 | 0.30311987 |
| 1.054723 | 0.016572763 | 0.21587715 | -0.026256423 | 0.30549908 |
| 1.064963 | 0.012367063 | 0.215794539 | -0.03004263 | 0.30697148 |
| 1.075203 | 0.008272932 | 0.21288717 | -0.033724633 | 0.3079034 |
| 1.085443 | 0.005435293 | 0.209060046 | -0.037716169 | 0.30737406 |
| 1.095683 | 0.002883531 | 0.202984388 | -0.041229326 | 0.30607075 |
| 1.105923 | 0.001039347 | 0.196167801 | -0.044063036 | 0.3032477 |
| 1.116163 | -0.000263217 | 0.188714263 | -0.046040919 | 0.29968531 |
| 1.126403 | -0.001469271 | 0.181304629 | -0.047177598 | 0.29516306 |
| 1.136643 | -0.00249966 | 0.174420225 | -0.046840625 | 0.29016837 |
| 1.146883 | -0.003626594 | 0.168641925 | -0.046018628 | 0.28469357 |
| 1.157123 | -0.005704491 | 0.165353901 | -0.04401727 | 0.27919906 |
| 1.167363 | -0.007459948 | 0.163479302 | -0.041582464 | 0.27337197 |
| 1.177603 | -0.009867819 | 0.164259334 | -0.038996049 | 0.26833796 |
| 1.187843 | -0.012094107 | 0.167080358 | -0.036402687 | 0.26238462 |
| 1.198083 | -0.01423512 | 0.172002771 | -0.033900007 | 0.25728222 |
| 1.208323 | -0.015460957 | 0.178984896 | -0.032082941 | 0.25261299 |
| 1.218564 | -0.016036141 | 0.186412998 | -0.030106645 | 0.24793213 |
| 1.228804 | -0.015464544 | 0.195182516 | -0.028858036 | 0.24357932 |
| 1.239044 | -0.013851019 | 0.203442161 | -0.026778204 | 0.24020033 |
| 1.249284 | -0.01144743 | 0.210987224 | -0.024679467 | 0.23692157 |
| 1.259524 | -0.008184799 | 0.217352176 | -0.021429127 | 0.23355314 |
| 1.269764 | -0.004648712 | 0.221941097 | -0.017028988 | 0.23051701 |
| 1.280004 | -0.001107609 | 0.225858025 | -0.011306057 | 0.2272429 |
| 1.290244 | 0.002616997 | 0.227811015 | -0.003989832 | 0.2242081 |
| 1.300484 | 0.006323383 | 0.229374767 | 0.004284361 | 0.2207362 |
| 1.310724 | 0.010279074 | 0.23064943 | 0.01385027 | 0.21816509 |
| 1.320964 | 0.01444631 | 0.231352119 | 0.02431796 | 0.21611322 |
| 1.331204 | 0.019221585 | 0.232397362 | 0.035020615 | 0.21483038 |
| 1.341444 | 0.0253186 | 0.233516064 | 0.046221482 | 0.21478435 |
| 1.351684 | 0.031264617 | 0.234369798 | 0.056935532 | 0.21610868 |
| 1.361924 | 0.038335933 | 0.235138406 | 0.067658049 | 0.21810391 |
| 1.372164 | 0.045226583 | 0.235097235 | 0.076572918 | 0.22181229 |
| 1.382404 | 0.051545116 | 0.23453201 | 0.084784424 | 0.22584878 |
| 1.392644 | 0.057454263 | 0.232671556 | 0.09026274 | 0.23097398 |
| 1.402884 | 0.062063509 | 0.230481018 | 0.094900803 | 0.23614517 |
| 1.413124 | 0.065543248 | 0.226741781 | 0.096025788 | 0.24212641 |
| 1.423364 | 0.067985319 | 0.223313513 | 0.096842941 | 0.24825048 |
| 1.433604 | 0.06947074 | 0.219408762 | 0.095543556 | 0.25404076 |
| 1.443844 | 0.070220586 | 0.215739115 | 0.094229501 | 0.2597114 |
| 1.454084 | 0.07111467 | 0.213156345 | 0.093077372 | 0.26442896 |
| 1.464324 | 0.071255852 | 0.210574355 | 0.092247471 | 0.26837607 |
| 1.474564 | 0.072273616 | 0.209689886 | 0.0927537 | 0.27071632 |
| 1.484804 | 0.072954873 | 0.208051523 | 0.093809578 | 0.2720545 |
| 1.495044 | 0.073902328 | 0.207769589 | 0.095724887 | 0.27171116 |
| 1.505284 | 0.074646061 | 0.206818005 | 0.098133054 | 0.27049507 |
| 1.515524 | 0.074972766 | 0.205972544 | 0.100602825 | 0.2677736 |
| 1.525764 | 0.074504815 | 0.204823024 | 0.102867851 | 0.2640273 |
| 1.536004 | 0.072789812 | 0.203500743 | 0.10481328 | 0.25918338 |
| 1.546245 | 0.070147942 | 0.201871476 | 0.106411544 | 0.2530679 |
| 1.556484 | 0.066282251 | 0.200623295 | 0.107843888 | 0.24625904 |
| 1.566725 | 0.062252928 | 0.199222327 | 0.108665885 | 0.2382343 |
| 1.576964 | 0.057696351 | 0.198329705 | 0.109629805 | 0.23026684 |
| 1.587205 | 0.054398085 | 0.19663407 | 0.110766887 | 0.22162368 |
| 1.597445 | 0.05175539 | 0.19551602 | 0.111179065 | 0.21339403 |
| 1.607685 | 0.051467958 | 0.19258305 | 0.112514693 | 0.20611343 |
| 1.617925 | 0.052101675 | 0.189606296 | 0.113204163 | 0.19944868 |
| 1.628165 | 0.054581808 | 0.184225957 | 0.114176626 | 0.19430812 |
| 1.638405 | 0.057766741 | 0.178086613 | 0.114953616 | 0.19039591 |
| 1.648645 | 0.061384528 | 0.169888251 | 0.115899926 | 0.18778429 |
| 1.658885 | 0.064786388 | 0.161401647 | 0.116774467 | 0.18652977 |
| 1.669125 | 0.067364662 | 0.15232243 | 0.118073024 | 0.18663669 |
| 1.679365 | 0.069724768 | 0.144133648 | 0.119703113 | 0.18710941 |
| 1.689605 | 0.071643521 | 0.137231344 | 0.122036342 | 0.18924932 |
| 1.699845 | 0.072734723 | 0.13178445 | 0.124744596 | 0.19086261 |
| 1.710085 | 0.074736311 | 0.128536741 | 0.12811037 | 0.19338387 |
| 1.720325 | 0.076263546 | 0.126648394 | 0.131660543 | 0.19564682 |
| 1.730565 | 0.078637797 | 0.125636964 | 0.135339568 | 0.1974933 |
| 1.740805 | 0.080822717 | 0.126139511 | 0.138769745 | 0.19917895 |
| 1.751045 | 0.083374694 | 0.127030624 | 0.142108088 | 0.20050782 |
| 1.761285 | 0.085623006 | 0.129752495 | 0.144971671 | 0.20152758 |
| 1.771525 | 0.087463711 | 0.132367571 | 0.147646712 | 0.20272756 |
| 1.781765 | 0.088999583 | 0.136452494 | 0.149721286 | 0.20367113 |
| 1.792005 | 0.089947428 | 0.140584833 | 0.152261001 | 0.20495949 |
| 1.802245 | 0.090583376 | 0.144424119 | 0.15423384 | 0.20611599 |
| 1.812485 | 0.090324583 | 0.148015705 | 0.156047657 | 0.20722302 |
| 1.822725 | 0.089832494 | 0.150590462 | 0.158131228 | 0.20843564 |
| 1.832965 | 0.08901577 | 0.153010869 | 0.160130357 | 0.20866639 |
| 1.843205 | 0.087930922 | 0.154995758 | 0.162148057 | 0.20837823 |
| 1.853445 | 0.086390367 | 0.1572251 | 0.164063605 | 0.20726784 |
| 1.863685 | 0.085157467 | 0.160148088 | 0.166092317 | 0.20570316 |
| 1.873925 | 0.083755195 | 0.163714724 | 0.167873687 | 0.20327841 |
| 1.884165 | 0.082905943 | 0.16818294 | 0.169331326 | 0.2012227 |
| 1.894405 | 0.081855875 | 0.172990407 | 0.170559135 | 0.19969206 |
| 1.904645 | 0.082533102 | 0.177351944 | 0.171113993 | 0.19883739 |
| 1.914886 | 0.083297131 | 0.181541996 | 0.171439914 | 0.19934318 |
| 1.925126 | 0.085300516 | 0.184505627 | 0.171703008 | 0.20068784 |
| 1.935366 | 0.088881914 | 0.186094289 | 0.171776593 | 0.20305803 |
| 1.945606 | 0.093696368 | 0.186701057 | 0.17217793 | 0.20567907 |
| 1.955846 | 0.099037703 | 0.185706609 | 0.173014687 | 0.20826992 |
| 1.966086 | 0.105879317 | 0.183545414 | 0.173736423 | 0.21038658 |
| 1.976326 | 0.113120326 | 0.179727208 | 0.175124559 | 0.21137754 |
| 1.986566 | 0.120187653 | 0.17526963 | 0.176047633 | 0.21143939 |
| 1.996806 | 0.127500201 | 0.169298771 | 0.176668366 | 0.21040225 |
| 2.007046 | 0.134445432 | 0.163665062 | 0.176831127 | 0.20909781 |
| 2.017286 | 0.14082857 | 0.156798387 | 0.17629483 | 0.20715116 |
| 2.027526 | 0.146933164 | 0.151753167 | 0.175601117 | 0.20623993 |
| 2.037766 | 0.15304911 | 0.146831739 | 0.174153194 | 0.20572248 |
| 2.048006 | 0.158652445 | 0.143873191 | 0.173136754 | 0.20701886 |
| 2.058246 | 0.164469561 | 0.141549766 | 0.171964157 | 0.2090849 |
| 2.068486 | 0.170156491 | 0.140808276 | 0.171116535 | 0.21217402 |
| 2.078726 | 0.176240914 | 0.140316268 | 0.171098446 | 0.21534858 |
| 2.088966 | 0.182023606 | 0.141003552 | 0.171182911 | 0.21885066 |
| 2.099206 | 0.188069785 | 0.140974676 | 0.171873221 | 0.22084066 |
| 2.109446 | 0.194305 | 0.141948079 | 0.173297958 | 0.22248266 |
| 2.119686 | 0.199805665 | 0.142236318 | 0.175060138 | 0.22212099 |
| 2.129926 | 0.205149033 | 0.143195126 | 0.176900103 | 0.22104652 |
| 2.140166 | 0.209476015 | 0.143347113 | 0.179754895 | 0.21824187 |
| 2.150406 | 0.213051916 | 0.143583352 | 0.182390471 | 0.21504693 |
| 2.160646 | 0.21531948 | 0.143798737 | 0.185812686 | 0.21130609 |
| 2.170886 | 0.216285415 | 0.143924611 | 0.189519384 | 0.20793936 |
| 2.181126 | 0.216530597 | 0.144771952 | 0.193638544 | 0.20571297 |
| 2.191366 | 0.216163301 | 0.147614388 | 0.198381302 | 0.20531621 |
| 2.201606 | 0.2157036 | 0.152274274 | 0.204115247 | 0.20659222 |
| 2.211846 | 0.216258723 | 0.159881415 | 0.210067441 | 0.21050872 |
| 2.222086 | 0.217686377 | 0.170291158 | 0.217077465 | 0.2158041 |
| 2.232327 | 0.221041759 | 0.182821469 | 0.224485456 | 0.22282884 |
| 2.242567 | 0.22600018 | 0.197234666 | 0.232491384 | 0.23055985 |
| 2.252806 | 0.233251921 | 0.212312526 | 0.240775545 | 0.2389923 |
| 2.263047 | 0.241801444 | 0.227837324 | 0.249230372 | 0.24731732 |
| 2.273287 | 0.252131472 | 0.243165811 | 0.257814667 | 0.25599786 |
| 2.283527 | 0.263067675 | 0.257555899 | 0.266092084 | 0.26419139 |
| 2.293766 | 0.274602989 | 0.270680843 | 0.274469481 | 0.27188616 |
| 2.304007 | 0.285814262 | 0.282771941 | 0.282947867 | 0.2787272 |
| 2.314247 | 0.296949837 | 0.293073872 | 0.291831427 | 0.28465748 |
| 2.324487 | 0.306147695 | 0.301330681 | 0.30025364 | 0.28864761 |
| 2.334727 | 0.315413536 | 0.308134843 | 0.310524912 | 0.29129058 |
| 2.344967 | 0.322970451 | 0.313225447 | 0.320664497 | 0.29241156 |

12kHz ABR , Threshold +50dB

| | *Mir183/96* $^{dko}$ Wildtype | | *Mir183/96* $^{dko}$ Heterozygote | |
|---|---|---|---|---|
| | Amplitude (µV) | | Amplitude (µV) | |
| Time (ms) | Mean | SD | Mean | SD |
| 0 | 0.068306777 | 0.105836237 | 0.024726635 | 0.25384332 |
| 0.01024 | 0.06087216 | 0.112692789 | 0.019750927 | 0.256103087 |
| 0.02048 | 0.053403846 | 0.119975992 | 0.014106388 | 0.259589475 |
| 0.03072 | 0.045949456 | 0.127880789 | 0.007188478 | 0.263246207 |
| 0.04096 | 0.03898756 | 0.136105851 | -6.97183E-05 | 0.268970519 |
| 0.0512 | 0.032317049 | 0.144131365 | -0.008801095 | 0.276008025 |
| 0.06144 | 0.026075831 | 0.151851349 | -0.017318334 | 0.284284554 |
| 0.07168 | 0.020479463 | 0.159289613 | -0.026911219 | 0.294699677 |
| 0.08192 | 0.015561825 | 0.165664557 | -0.03542404 | 0.305416848 |
| 0.09216 | 0.011524199 | 0.170456785 | -0.044496385 | 0.316737506 |
| 0.1024 | 0.00759816 | 0.174103507 | -0.051959911 | 0.327103207 |
| 0.11264 | 0.004877058 | 0.175664793 | -0.058420396 | 0.335607119 |
| 0.12288 | 0.002008608 | 0.175844585 | -0.063435791 | 0.341748817 |
| 0.13312 | -0.001232307 | 0.174132614 | -0.0666409 | 0.344478731 |
| 0.14336 | -0.004285894 | 0.171218729 | -0.069179878 | 0.343781707 |
| 0.1536 | -0.008087371 | 0.166874242 | -0.070003924 | 0.339699873 |
| 0.16384 | -0.012810706 | 0.162097477 | -0.071110328 | 0.332182629 |
| 0.174081 | -0.017467837 | 0.155960434 | -0.071152438 | 0.322878349 |
| 0.184321 | -0.02267842 | 0.150661641 | -0.072110981 | 0.31107451 |
| 0.194561 | -0.027494415 | 0.145083653 | -0.072878571 | 0.299695436 |
| 0.204801 | -0.032548116 | 0.141236657 | -0.074367209 | 0.287300527 |
| 0.215041 | -0.036291004 | 0.139141398 | -0.075458313 | 0.276572015 |
| 0.225281 | -0.040160775 | 0.139721005 | -0.077074014 | 0.266308354 |
| 0.235521 | -0.043126803 | 0.142811386 | -0.077935811 | 0.258346039 |
| 0.245761 | -0.045724957 | 0.147865436 | -0.078474631 | 0.251008623 |
| 0.256001 | -0.048027676 | 0.154684277 | -0.078279997 | 0.245463008 |
| 0.266241 | -0.050298609 | 0.162675864 | -0.077279972 | 0.240723875 |
| 0.276481 | -0.051980689 | 0.169933602 | -0.075851687 | 0.237268236 |
| 0.286721 | -0.053752974 | 0.177606133 | -0.074021567 | 0.234772366 |
| 0.296961 | -0.054758225 | 0.183951 | -0.071868286 | 0.2332043 |
| 0.307201 | -0.05552153 | 0.189841668 | -0.069936697 | 0.232881492 |
| 0.317441 | -0.055402208 | 0.194429679 | -0.068404187 | 0.233697526 |
| 0.327681 | -0.054928808 | 0.198827609 | -0.067208987 | 0.235305326 |
| 0.337921 | -0.053657457 | 0.20155182 | -0.066899133 | 0.237404672 |
| 0.348161 | -0.051715905 | 0.204520979 | -0.067134554 | 0.239410162 |
| 0.358401 | -0.049774345 | 0.206208611 | -0.068762038 | 0.240913243 |
| 0.368641 | -0.04724488 | 0.208082304 | -0.070980455 | 0.241883163 |
| 0.378881 | -0.04501228 | 0.209539043 | -0.074743218 | 0.241712837 |
| 0.389121 | -0.04307843 | 0.210924365 | -0.079383785 | 0.240817943 |
| 0.399361 | -0.041507147 | 0.213233896 | -0.085467194 | 0.238690504 |
| 0.409601 | -0.040607539 | 0.21542499 | -0.092160494 | 0.235550793 |
| 0.419841 | -0.040726719 | 0.218456635 | -0.100085144 | 0.231168345 |
| 0.430081 | -0.041162565 | 0.222003238 | -0.107529543 | 0.226197682 |
| 0.440321 | -0.042771751 | 0.225424513 | -0.114820281 | 0.219664784 |
| 0.450561 | -0.044572966 | 0.229351902 | -0.121556477 | 0.213551876 |
| 0.460801 | -0.047100865 | 0.232851584 | -0.126431578 | 0.206182649 |
| 0.471041 | -0.04957405 | 0.235857859 | -0.129383087 | 0.19955775 |
| 0.481281 | -0.051975057 | 0.238029947 | -0.130745097 | 0.193197604 |
| 0.491521 | -0.053893082 | 0.23949904 | -0.129817705 | 0.187503175 |
| 0.501761 | -0.054942644 | 0.24011594 | -0.127247954 | 0.182001162 |
| 0.512001 | -0.054664304 | 0.239853538 | -0.123412033 | 0.178298622 |
| 0.522242 | -0.052794635 | 0.238662582 | -0.118983141 | 0.174605509 |
| 0.532482 | -0.049828798 | 0.237173483 | -0.115019913 | 0.172441221 |
| 0.542722 | -0.044697489 | 0.234715608 | -0.110903385 | 0.171235546 |
| 0.552962 | -0.03897402 | 0.232529664 | -0.108635698 | 0.171305934 |
| 0.563202 | -0.032312155 | 0.229634349 | -0.107796266 | 0.17307518 |
| 0.573442 | -0.025672117 | 0.227409879 | -0.108389902 | 0.176231804 |
| 0.583682 | -0.019922888 | 0.225254538 | -0.11110924 | 0.180528183 |
| 0.593922 | -0.014737951 | 0.223935891 | -0.114747133 | 0.186842795 |
| 0.604162 | -0.011703415 | 0.22319526 | -0.120264607 | 0.194167599 |
| 0.614402 | -0.009279197 | 0.224249642 | -0.126218903 | 0.202204765 |
| 0.624642 | -0.009220006 | 0.225752436 | -0.132605768 | 0.211654444 |
| 0.634882 | -0.009707458 | 0.229058898 | -0.138971604 | 0.220703041 |
| 0.645122 | -0.011952115 | 0.232971574 | -0.144805995 | 0.230771899 |
| 0.655362 | -0.014599828 | 0.237477265 | -0.149350099 | 0.239961084 |
| 0.665602 | -0.01818634 | 0.242185486 | -0.153003397 | 0.249337084 |
| 0.675842 | -0.022562428 | 0.246192942 | -0.155326032 | 0.257991808 |
| 0.686082 | -0.027341052 | 0.249181682 | -0.156115694 | 0.265706111 |
| 0.696322 | -0.033049349 | 0.250597843 | -0.156419866 | 0.272819175 |
| 0.706562 | -0.038751213 | 0.250400956 | -0.155877718 | 0.278173713 |
| 0.716802 | -0.045456686 | 0.24824379 | -0.154879949 | 0.28225915 |
| 0.727042 | -0.051688572 | 0.244774203 | -0.153429556 | 0.283830077 |
| 0.737282 | -0.05838966 | 0.239468419 | -0.151205216 | 0.28350372 |
| 0.747522 | -0.063802162 | 0.234030601 | -0.149015998 | 0.280458852 |
| 0.757762 | -0.069059663 | 0.228162848 | -0.145042278 | 0.274927395 |
| 0.768002 | -0.072834168 | 0.222107142 | -0.140792147 | 0.267124987 |
| 0.778242 | -0.075643548 | 0.216982239 | -0.1352454 | 0.258232586 |
| 0.788482 | -0.076762581 | 0.212738733 | -0.128845534 | 0.248199821 |
| 0.798722 | -0.077056438 | 0.209734209 | -0.121965847 | 0.237744432 |
| 0.808962 | -0.07661964 | 0.207887611 | -0.114334815 | 0.228006982 |
| 0.819202 | -0.075205758 | 0.208022098 | -0.106993514 | 0.219205385 |
| 0.829442 | -0.073531363 | 0.209597702 | -0.099373387 | 0.211319866 |
| 0.839682 | -0.072465821 | 0.212194167 | -0.091818407 | 0.204103408 |
| 0.849922 | -0.071155456 | 0.215766875 | -0.084662981 | 0.198366985 |
| 0.860162 | -0.070749438 | 0.219182458 | -0.077985005 | 0.193101801 |
| 0.870403 | -0.071033907 | 0.222199946 | -0.071191632 | 0.188662303 |
| 0.880643 | -0.071491523 | 0.223260116 | -0.065631178 | 0.184637228 |
| 0.890883 | -0.07294047 | 0.223303372 | -0.060631714 | 0.18129319 |
| 0.901123 | -0.074487803 | 0.220665508 | -0.056469919 | 0.178624234 |
| 0.911363 | -0.075947357 | 0.215663233 | -0.053861891 | 0.175924513 |
| 0.921603 | -0.077433586 | 0.20959224 | -0.052070331 | 0.173916629 |
| 0.931843 | -0.078954314 | 0.201944637 | -0.051090176 | 0.171905554 |
| 0.942083 | -0.080416684 | 0.19443379 | -0.052621153 | 0.169897105 |
| 0.952323 | -0.081658366 | 0.187485152 | -0.055205397 | 0.168266767 |
| 0.962563 | -0.083875737 | 0.181990296 | -0.058192691 | 0.165699931 |
| 0.972803 | -0.086424925 | 0.178535437 | -0.062523158 | 0.16399987 |
| 0.983043 | -0.089539239 | 0.1772643 | -0.067061934 | 0.161519972 |
| 0.993283 | -0.092747389 | 0.177492678 | -0.071985105 | 0.159528561 |
| 1.003523 | -0.096097841 | 0.179263304 | -0.076560941 | 0.15713334 |
| 1.013763 | -0.098885683 | 0.181783683 | -0.080203644 | 0.155158986 |
| 1.024003 | -0.100441054 | 0.184547319 | -0.083714361 | 0.152696096 |
| 1.034243 | -0.100520095 | 0.186477117 | -0.085441781 | 0.150648602 |
| 1.044483 | -0.099293534 | 0.188101651 | -0.086980685 | 0.147576233 |
| 1.054723 | -0.096206387 | 0.188445414 | -0.087126638 | 0.144065316 |
| 1.064963 | -0.092110862 | 0.187957903 | -0.086618545 | 0.140334324 |
| 1.075203 | -0.087316542 | 0.186675037 | -0.085421076 | 0.135765846 |
| 1.085443 | -0.081247837 | 0.185004832 | -0.082883063 | 0.131434476 |
| 1.095683 | -0.075606875 | 0.183982503 | -0.079728183 | 0.127294704 |
| 1.105923 | -0.069555651 | 0.182919481 | -0.074916068 | 0.124087373 |
| 1.116163 | -0.063662221 | 0.1834356 | -0.069083051 | 0.121621036 |
| 1.126403 | -0.058226986 | 0.18547653 | -0.061747563 | 0.119451913 |
| 1.136643 | -0.05327426 | 0.189195559 | -0.053334006 | 0.117664821 |
| 1.146883 | -0.048842704 | 0.194430948 | -0.044135906 | 0.115933683 |
| 1.157123 | -0.044872051 | 0.200600313 | -0.034332817 | 0.113452762 |
| 1.167363 | -0.041693177 | 0.207692593 | -0.024119499 | 0.111173372 |
| 1.177603 | -0.038220087 | 0.214419092 | -0.014458479 | 0.109704246 |
| 1.187843 | -0.034630639 | 0.221016826 | -0.00473197 | 0.108617395 |
| 1.198083 | -0.030521263 | 0.227247677 | 0.004295885 | 0.108301075 |
| 1.208323 | -0.025586577 | 0.233176549 | 0.012875744 | 0.110113132 |
| 1.218564 | -0.019458788 | 0.239277072 | 0.02022776 | 0.112813386 |
| 1.228804 | -0.012285485 | 0.246149101 | 0.027211048 | 0.116398732 |
| 1.239044 | -0.004559343 | 0.253384843 | 0.032871139 | 0.121054631 |
| 1.249284 | 0.003935749 | 0.26145934 | 0.037196977 | 0.125477919 |
| 1.259524 | 0.012522263 | 0.269989178 | 0.040789363 | 0.1300555 |
| 1.269764 | 0.020186899 | 0.278022212 | 0.042552428 | 0.133169321 |
| 1.280004 | 0.027357399 | 0.285934969 | 0.043025902 | 0.135219937 |
| 1.290244 | 0.032704333 | 0.292908791 | 0.042863702 | 0.135375171 |
| 1.300484 | 0.036721468 | 0.298837549 | 0.040756552 | 0.133668124 |
| 1.310724 | 0.038718718 | 0.303810432 | 0.038484375 | 0.130543701 |
| 1.320964 | 0.039087441 | 0.307783776 | 0.035048121 | 0.12603674 |
| 1.331204 | 0.038814578 | 0.310706038 | 0.032136051 | 0.121280593 |
| 1.341444 | 0.037659862 | 0.312059827 | 0.028779444 | 0.11698121 |
| 1.351684 | 0.036447727 | 0.312239668 | 0.026297339 | 0.112888685 |
| 1.361924 | 0.036849027 | 0.310027601 | 0.024815914 | 0.110212017 |
| 1.372164 | 0.037410665 | 0.305796275 | 0.024750377 | 0.107851694 |
| 1.382404 | 0.040136133 | 0.299153251 | 0.025593318 | 0.106267746 |
| 1.392644 | 0.043722135 | 0.290233791 | 0.028045314 | 0.10541192 |
| 1.402884 | 0.048563104 | 0.279762411 | 0.03198666 | 0.104694284 |
| 1.413124 | 0.054121793 | 0.267801616 | 0.036530716 | 0.104809594 |
| 1.423364 | 0.060119351 | 0.256165847 | 0.042208717 | 0.106407457 |
| 1.433604 | 0.06652908 | 0.244282732 | 0.048298566 | 0.107992065 |
| 1.443844 | 0.072980579 | 0.234614283 | 0.05491078 | 0.111052007 |
| 1.454084 | 0.0796793 | 0.226376897 | 0.06132937 | 0.114266376 |
| 1.464324 | 0.08744564 | 0.220951422 | 0.067980407 | 0.117507516 |
| 1.474564 | 0.09501365 | 0.217790111 | 0.073536552 | 0.120647738 |
| 1.484804 | 0.103750781 | 0.217034701 | 0.079049954 | 0.122748621 |
| 1.495044 | 0.112534747 | 0.218013913 | 0.082931222 | 0.12416057 |
| 1.505284 | 0.121818544 | 0.220410501 | 0.086257215 | 0.124426119 |
| 1.515524 | 0.130324318 | 0.223011663 | 0.088987013 | 0.124355338 |
| 1.525764 | 0.138041958 | 0.226316321 | 0.09048764 | 0.124036828 |
| 1.536004 | 0.144344712 | 0.228829302 | 0.092149284 | 0.123507869 |
| 1.546245 | 0.148541745 | 0.231995669 | 0.093508714 | 0.123956207 |
| 1.556484 | 0.150937205 | 0.234785315 | 0.095250444 | 0.124944492 |
| 1.566725 | 0.150971354 | 0.238043465 | 0.097028394 | 0.126786218 |
| 1.576964 | 0.149424711 | 0.242258158 | 0.099011004 | 0.129179543 |
| 1.587205 | 0.146126821 | 0.247014748 | 0.10093382 | 0.131865308 |
| 1.597445 | 0.142498742 | 0.252332971 | 0.10309112 | 0.135111032 |
| 1.607685 | 0.138638573 | 0.25792719 | 0.104424104 | 0.138519888 |
| 1.617925 | 0.134858422 | 0.263370306 | 0.106241242 | 0.142844101 |
| 1.628165 | 0.132404176 | 0.268978027 | 0.107834832 | 0.147651292 |
| 1.638405 | 0.13143461 | 0.273565555 | 0.109497575 | 0.153628992 |
| 1.648645 | 0.131895908 | 0.277883891 | 0.112015633 | 0.160593098 |
| 1.658885 | 0.133892298 | 0.281690635 | 0.114368347 | 0.16838475 |
| 1.669125 | 0.13789673 | 0.284783236 | 0.117978001 | 0.176001711 |
| 1.679365 | 0.14279627 | 0.28731286 | 0.121866345 | 0.183618107 |
| 1.689605 | 0.148987296 | 0.288873485 | 0.126251204 | 0.190725312 |
| 1.699845 | 0.1555388 | 0.290001664 | 0.131626943 | 0.196374697 |
| 1.710085 | 0.161913737 | 0.289817846 | 0.137524105 | 0.201280919 |
| 1.720325 | 0.168242453 | 0.28936977 | 0.143853093 | 0.204432054 |
| 1.730565 | 0.173700525 | 0.287645127 | 0.151494793 | 0.206036888 |
| 1.740805 | 0.177919163 | 0.286042616 | 0.159528454 | 0.206206348 |
| 1.751045 | 0.181732176 | 0.283886169 | 0.168417592 | 0.204472774 |
| 1.761285 | 0.184664796 | 0.281827718 | 0.177397187 | 0.200885046 |
| 1.771525 | 0.187015849 | 0.280237794 | 0.186617975 | 0.195576982 |
| 1.781765 | 0.1897309 | 0.279443449 | 0.195551701 | 0.188572746 |
| 1.792005 | 0.192710899 | 0.2787337 | 0.2039352 | 0.180189976 |
| 1.802245 | 0.19618753 | 0.279174827 | 0.211359428 | 0.17004409 |
| 1.812485 | 0.201094183 | 0.280185992 | 0.217890709 | 0.159803287 |
| 1.822725 | 0.206717246 | 0.281563447 | 0.223139873 | 0.148965972 |
| 1.832965 | 0.213843628 | 0.283172844 | 0.226927322 | 0.138358814 |
| 1.843205 | 0.221846456 | 0.284606939 | 0.23036514 | 0.12869258 |
| 1.853445 | 0.230766106 | 0.285324594 | 0.232252286 | 0.121608024 |
| 1.863685 | 0.240489716 | 0.285038955 | 0.234041732 | 0.116493408 |
| 1.873925 | 0.250895307 | 0.284486974 | 0.235456037 | 0.114885288 |
| 1.884165 | 0.26151523 | 0.282863152 | 0.237290748 | 0.116070422 |
| 1.894405 | 0.273038284 | 0.28032573 | 0.238847118 | 0.120015487 |
| 1.904645 | 0.28470305 | 0.278704158 | 0.241643415 | 0.124718227 |
| 1.914886 | 0.297368189 | 0.27581182 | 0.244119339 | 0.130034125 |
| 1.925126 | 0.310873158 | 0.274118216 | 0.247704067 | 0.134542027 |
| 1.935366 | 0.325647912 | 0.272206197 | 0.25098413 | 0.137766594 |
| 1.945606 | 0.341493182 | 0.271128962 | 0.255311193 | 0.140141085 |
| 1.955846 | 0.358584826 | 0.27041869 | 0.26005531 | 0.139791945 |
| 1.966086 | 0.377288708 | 0.270252273 | 0.265868645 | 0.139361077 |
| 1.976326 | 0.396723893 | 0.271163855 | 0.273190985 | 0.137423957 |
| 1.986566 | 0.417405992 | 0.272629051 | 0.282976527 | 0.135908521 |
| 1.996806 | 0.438775773 | 0.274746222 | 0.294781087 | 0.135302379 |
| 2.007046 | 0.461656202 | 0.278339376 | 0.308919461 | 0.137647397 |
| 2.017286 | 0.484295604 | 0.282439233 | 0.326297519 | 0.14224735 |
| 2.027526 | 0.509254461 | 0.287043289 | 0.345286529 | 0.150823705 |
| 2.037766 | 0.533941514 | 0.293087732 | 0.366608758 | 0.161616024 |
| 2.048006 | 0.560536377 | 0.298746811 | 0.38887495 | 0.173473476 |
| 2.058246 | 0.587767705 | 0.305215499 | 0.412495643 | 0.185813413 |
| 2.068486 | 0.616798513 | 0.311612888 | 0.437154136 | 0.196989861 |
| 2.078726 | 0.647012681 | 0.317018527 | 0.461635334 | 0.207060453 |
| 2.088966 | 0.678631643 | 0.322489177 | 0.487009074 | 0.214710506 |
| 2.099206 | 0.712427201 | 0.326316217 | 0.5134099 | 0.221840219 |
| 2.109446 | 0.747326622 | 0.329422134 | 0.540053231 | 0.22730786 |
| 2.119686 | 0.784861391 | 0.331564721 | 0.568459753 | 0.23327391 |
| 2.129926 | 0.823250165 | 0.332264392 | 0.597733698 | 0.238419517 |
| 2.140166 | 0.863820458 | 0.332661157 | 0.62872532 | 0.24444704 |
| 2.150406 | 0.905332188 | 0.332355717 | 0.661147952 | 0.250179818 |
| 2.160646 | 0.948102792 | 0.331555815 | 0.694619043 | 0.256493323 |
| 2.170886 | 0.990657111 | 0.330494316 | 0.729156145 | 0.261209316 |
| 2.181126 | 1.034063766 | 0.328962971 | 0.764206656 | 0.266264815 |
| 2.191366 | 1.075903157 | 0.326575973 | 0.798937257 | 0.269091781 |
| 2.201606 | 1.116792738 | 0.323501698 | 0.833919942 | 0.271217116 |
| 2.211846 | 1.155994028 | 0.319527517 | 0.867650662 | 0.271244855 |
| 2.222086 | 1.192682534 | 0.313562849 | 0.901002799 | 0.269701184 |
| 2.232327 | 1.227075438 | 0.306780246 | 0.93255806 | 0.266849168 |
| 2.242567 | 1.258729249 | 0.297667746 | 0.963500074 | 0.261500897 |
| 2.252806 | 1.287957062 | 0.286387801 | 0.992658206 | 0.255343141 |
| 2.263047 | 1.31449008 | 0.273275799 | 1.02045181 | 0.246350009 |
| 2.273287 | 1.338937054 | 0.257516864 | 1.046961589 | 0.236996395 |
| 2.283527 | 1.360383232 | 0.240246618 | 1.071014541 | 0.225717034 |
| 2.293766 | 1.379347434 | 0.222618893 | 1.09331555 | 0.214292538 |
| 2.304007 | 1.39510715 | 0.205003394 | 1.112507335 | 0.202710177 |
| 2.314247 | 1.407397827 | 0.19098969 | 1.128877767 | 0.192284666 |
| 2.324487 | 1.415456176 | 0.181649883 | 1.141428445 | 0.184235594 |
| 2.334727 | 1.419587155 | 0.180790119 | 1.149473071 | 0.178732541 |
| 2.344967 | 1.418562353 | 0.187671197 | 1.15383436 | 0.17723023 |

12kHz ABR , Threshold +20dB

| | *Mir182* $^{ko}$ Wildtype | | *Mir182* $^{ko}$ Homozygote | |
|---|---|---|---|---|
| | Amplitude (µV) | | Amplitude (µV) | |
| Time (ms) | Mean | SD | Mean | SD |
| 0 | -0.0575172 | 0.159209958 | 0.033157604 | 0.196578974 |
| 0.01024 | -0.06325792 | 0.158308691 | 0.034517024 | 0.200569361 |
| 0.02048 | -0.068223054 | 0.156113774 | 0.036334659 | 0.204919307 |
| 0.03072 | -0.072375632 | 0.153597495 | 0.039144052 | 0.210725085 |
| 0.04096 | -0.075183139 | 0.150369963 | 0.042593431 | 0.21686833 |
| 0.0512 | -0.077837246 | 0.146605394 | 0.046674672 | 0.224492476 |
| 0.06144 | -0.079714456 | 0.143263527 | 0.051153374 | 0.231449814 |
| 0.07168 | -0.081923896 | 0.140090833 | 0.055707958 | 0.239692569 |
| 0.08192 | -0.083806137 | 0.137745933 | 0.060145816 | 0.24704106 |
| 0.09216 | -0.08612564 | 0.136110921 | 0.064724225 | 0.254867611 |
| 0.1024 | -0.088290875 | 0.135689155 | 0.06915911 | 0.261988939 |
| 0.11264 | -0.090324241 | 0.135654495 | 0.073507019 | 0.268996331 |
| 0.12288 | -0.09161385 | 0.136511265 | 0.078804291 | 0.275129771 |
| 0.13312 | -0.092721261 | 0.137218225 | 0.083677561 | 0.280315107 |
| 0.14336 | -0.092487489 | 0.138150958 | 0.089505425 | 0.284083154 |
| 0.1536 | -0.091837281 | 0.139146152 | 0.094917235 | 0.286132156 |
| 0.16384 | -0.089759061 | 0.140258249 | 0.100701242 | 0.286501604 |
| 0.174081 | -0.086835352 | 0.141443017 | 0.10530407 | 0.284721329 |
| 0.184321 | -0.083300837 | 0.143800036 | 0.109561519 | 0.281615324 |
| 0.194561 | -0.07842801 | 0.146496239 | 0.112364522 | 0.276978997 |
| 0.204801 | -0.073010555 | 0.150318397 | 0.114302602 | 0.271555156 |
| 0.215041 | -0.066781055 | 0.154412891 | 0.115076724 | 0.266010228 |
| 0.225281 | -0.060144397 | 0.159216374 | 0.115362089 | 0.260453992 |
| 0.235521 | -0.053292224 | 0.164132398 | 0.115200723 | 0.255571375 |
| 0.245761 | -0.046689385 | 0.168398091 | 0.115266314 | 0.2518969 |
| 0.256001 | -0.040272372 | 0.172180008 | 0.115906701 | 0.249209541 |
| 0.266241 | -0.034691039 | 0.175384595 | 0.117565046 | 0.247618598 |
| 0.276481 | -0.030135484 | 0.177767025 | 0.120131329 | 0.247007334 |
| 0.286721 | -0.026284785 | 0.179880333 | 0.123727828 | 0.247145374 |
| 0.296961 | -0.02387385 | 0.181173622 | 0.128548697 | 0.247748057 |
| 0.307201 | -0.022448342 | 0.182552888 | 0.133937503 | 0.24848771 |
| 0.317441 | -0.022546203 | 0.183820886 | 0.139767143 | 0.24907154 |
| 0.327681 | -0.023356378 | 0.18443867 | 0.145892463 | 0.250036266 |
| 0.337921 | -0.025905669 | 0.184755174 | 0.151576907 | 0.250519318 |
| 0.348161 | -0.029047977 | 0.184922862 | 0.156904741 | 0.251795762 |
| 0.358401 | -0.033538915 | 0.184267267 | 0.161596879 | 0.25305057 |
| 0.368641 | -0.038765426 | 0.18327685 | 0.165053843 | 0.255191712 |
| 0.378881 | -0.044745727 | 0.181969002 | 0.167654416 | 0.25802711 |
| 0.389121 | -0.050759054 | 0.180320171 | 0.169044777 | 0.261545729 |
| 0.399361 | -0.056930593 | 0.178361497 | 0.168930576 | 0.265553451 |
| 0.409601 | -0.062298607 | 0.176843953 | 0.168366559 | 0.269922897 |
| 0.419841 | -0.066840571 | 0.175540359 | 0.166294147 | 0.273773774 |
| 0.430081 | -0.069798673 | 0.174787933 | 0.163804805 | 0.277448506 |
| 0.440321 | -0.071021173 | 0.175257619 | 0.160683584 | 0.279928514 |
| 0.450561 | -0.070801836 | 0.176025635 | 0.157506813 | 0.281611416 |
| 0.460801 | -0.06859485 | 0.177953466 | 0.154086553 | 0.281935796 |
| 0.471041 | -0.065390453 | 0.18062213 | 0.150738561 | 0.281067426 |
| 0.481281 | -0.061390261 | 0.183418267 | 0.147772568 | 0.279153682 |
| 0.491521 | -0.05730423 | 0.186253305 | 0.144905924 | 0.275800456 |
| 0.501761 | -0.053604767 | 0.188951667 | 0.142401712 | 0.27172217 |
| 0.512001 | -0.050681961 | 0.191158304 | 0.140583868 | 0.266197035 |
| 0.522242 | -0.048899455 | 0.192615908 | 0.139163722 | 0.259998243 |
| 0.532482 | -0.048203194 | 0.193352157 | 0.137991745 | 0.252561674 |
| 0.542722 | -0.048357762 | 0.193436977 | 0.13792617 | 0.244576734 |
| 0.552962 | -0.048941304 | 0.192510891 | 0.137577724 | 0.235867078 |
| 0.563202 | -0.049588563 | 0.190772462 | 0.137834687 | 0.226765853 |
| 0.573442 | -0.050208307 | 0.187831507 | 0.137454044 | 0.217833866 |
| 0.583682 | -0.050271825 | 0.184455916 | 0.136841304 | 0.209349207 |
| 0.593922 | -0.049541317 | 0.179832598 | 0.135289689 | 0.201949843 |
| 0.604162 | -0.048500287 | 0.174827508 | 0.132445845 | 0.195913179 |
| 0.614402 | -0.047019301 | 0.169303082 | 0.128176543 | 0.191106743 |
| 0.624642 | -0.04494244 | 0.164270186 | 0.122598377 | 0.188620695 |
| 0.634882 | -0.042554752 | 0.159732659 | 0.115444806 | 0.18736988 |
| 0.645122 | -0.039989729 | 0.155610105 | 0.106978932 | 0.187904449 |
| 0.655362 | -0.036772741 | 0.153228653 | 0.097676685 | 0.189560983 |
| 0.665602 | -0.03310321 | 0.151905511 | 0.087847858 | 0.19224789 |
| 0.675842 | -0.028964924 | 0.151462363 | 0.077490839 | 0.195174692 |
| 0.686082 | -0.02418448 | 0.151547391 | 0.067793568 | 0.198133927 |
| 0.696322 | -0.019261318 | 0.152734486 | 0.058345785 | 0.200432058 |
| 0.706562 | -0.014476463 | 0.153937532 | 0.049852525 | 0.202310102 |
| 0.716802 | -0.00997667 | 0.155836537 | 0.041962142 | 0.203318706 |
| 0.727042 | -0.006442911 | 0.158179916 | 0.035254875 | 0.203370787 |
| 0.737282 | -0.004040699 | 0.161485189 | 0.029541204 | 0.203038355 |
| 0.747522 | -0.003228937 | 0.165043326 | 0.024723371 | 0.20237874 |
| 0.757762 | -0.003516936 | 0.168862051 | 0.021054593 | 0.201469551 |
| 0.768002 | -0.005504877 | 0.172333117 | 0.018130985 | 0.20030649 |
| 0.778242 | -0.008116488 | 0.175089593 | 0.016299493 | 0.199446399 |
| 0.788482 | -0.011394942 | 0.176456355 | 0.015301746 | 0.198571353 |
| 0.798722 | -0.014938519 | 0.176404409 | 0.014824513 | 0.197909347 |
| 0.808962 | -0.017863906 | 0.174752356 | 0.015051751 | 0.197205191 |
| 0.819202 | -0.019989507 | 0.171930322 | 0.015219592 | 0.197046277 |
| 0.829442 | -0.021045928 | 0.167758157 | 0.015726199 | 0.196860953 |
| 0.839682 | -0.020883938 | 0.163316076 | 0.016093795 | 0.197236711 |
| 0.849922 | -0.018848487 | 0.158395413 | 0.016109436 | 0.197688522 |
| 0.860162 | -0.015503673 | 0.153740241 | 0.015851697 | 0.198875707 |
| 0.870403 | -0.011288939 | 0.149933288 | 0.015358861 | 0.199409888 |
| 0.880643 | -0.005483919 | 0.146617538 | 0.014569678 | 0.200537944 |
| 0.890883 | 0.000666706 | 0.144739918 | 0.013566286 | 0.2002105 |
| 0.901123 | 0.007200994 | 0.143528409 | 0.012069509 | 0.199167821 |
| 0.911363 | 0.013709054 | 0.143411909 | 0.01101387 | 0.196452521 |
| 0.921603 | 0.019713485 | 0.143874929 | 0.009931935 | 0.19258554 |
| 0.931843 | 0.02496188 | 0.145040743 | 0.009096419 | 0.187435882 |
| 0.942083 | 0.029351232 | 0.146491349 | 0.008998411 | 0.181082954 |
| 0.952323 | 0.032305663 | 0.148527998 | 0.009238272 | 0.175256863 |
| 0.962563 | 0.034316646 | 0.150882884 | 0.010456099 | 0.168603091 |
| 0.972803 | 0.034987995 | 0.153426503 | 0.011266988 | 0.163265113 |
| 0.983043 | 0.034498318 | 0.156269088 | 0.012546002 | 0.157787527 |
| 0.993283 | 0.033096328 | 0.158852305 | 0.013627871 | 0.153914466 |
| 1.003523 | 0.030946645 | 0.161514557 | 0.014157252 | 0.150007391 |
| 1.013763 | 0.028495782 | 0.163894139 | 0.013710565 | 0.14796584 |
| 1.024003 | 0.025787699 | 0.16553059 | 0.013160915 | 0.1468634 |
| 1.034243 | 0.023073871 | 0.166775371 | 0.011723651 | 0.147496062 |
| 1.044483 | 0.020858428 | 0.167468811 | 0.010168473 | 0.149754947 |
| 1.054723 | 0.018663988 | 0.167216607 | 0.008448816 | 0.153580538 |
| 1.064963 | 0.016819824 | 0.166734062 | 0.006565011 | 0.158973255 |
| 1.075203 | 0.015315872 | 0.165527269 | 0.004928076 | 0.165078624 |
| 1.085443 | 0.013849057 | 0.164328879 | 0.00340284 | 0.171807023 |
| 1.095683 | 0.012523164 | 0.162913327 | 0.00201106 | 0.178473494 |
| 1.105923 | 0.011299014 | 0.162123019 | 0.000683514 | 0.184737365 |
| 1.116163 | 0.010219111 | 0.161180542 | -0.000275957 | 0.190515247 |
| 1.126403 | 0.009273034 | 0.161527983 | -0.001230745 | 0.195244964 |
| 1.136643 | 0.008476638 | 0.161581779 | -0.00168685 | 0.199217606 |
| 1.146883 | 0.008312316 | 0.163091254 | -0.00233936 | 0.201854999 |
| 1.157123 | 0.008120617 | 0.164405429 | -0.002501655 | 0.204072204 |
| 1.167363 | 0.008263788 | 0.166403408 | -0.002840311 | 0.20468772 |
| 1.177603 | 0.008550064 | 0.168125324 | -0.00360378 | 0.205084346 |
| 1.187843 | 0.008908224 | 0.169902964 | -0.004589396 | 0.204429178 |
| 1.198083 | 0.009385412 | 0.171283608 | -0.006361121 | 0.203671449 |
| 1.208323 | 0.009988209 | 0.17197196 | -0.008780984 | 0.202217865 |
| 1.218564 | 0.010682574 | 0.172424533 | -0.012006305 | 0.200821819 |
| 1.228804 | 0.012007308 | 0.172249638 | -0.016086523 | 0.199176309 |
| 1.239044 | 0.013810129 | 0.172121723 | -0.020381401 | 0.198143903 |
| 1.249284 | 0.016478545 | 0.171865046 | -0.025344344 | 0.197170791 |
| 1.259524 | 0.019452044 | 0.171664221 | -0.030142252 | 0.197130967 |
| 1.269764 | 0.0234132 | 0.17201233 | -0.034594278 | 0.198063885 |
| 1.280004 | 0.027668714 | 0.17231385 | -0.038663008 | 0.199982485 |
| 1.290244 | 0.032266038 | 0.173567823 | -0.04185917 | 0.202489134 |
| 1.300484 | 0.036637392 | 0.174354176 | -0.044198189 | 0.205783248 |
| 1.310724 | 0.041571607 | 0.17648641 | -0.04558202 | 0.209047004 |
| 1.320964 | 0.046123721 | 0.17825188 | -0.046154614 | 0.211955088 |
| 1.331204 | 0.050585637 | 0.180660579 | -0.046051651 | 0.214068478 |
| 1.341444 | 0.055063138 | 0.182859552 | -0.045478073 | 0.214849302 |
| 1.351684 | 0.059236276 | 0.185194101 | -0.044281124 | 0.21431002 |
| 1.361924 | 0.063091051 | 0.186938237 | -0.04340079 | 0.211741829 |
| 1.372164 | 0.066827407 | 0.188514155 | -0.042270075 | 0.20790855 |
| 1.382404 | 0.069741 | 0.189399589 | -0.041155502 | 0.202604405 |
| 1.392644 | 0.072901758 | 0.190133965 | -0.040144126 | 0.196206756 |
| 1.402884 | 0.075257244 | 0.189838873 | -0.039162929 | 0.18958002 |
| 1.413124 | 0.077917682 | 0.18953888 | -0.038023281 | 0.183149985 |
| 1.423364 | 0.080810272 | 0.188440346 | -0.036579244 | 0.177719574 |
| 1.433604 | 0.083737563 | 0.187247365 | -0.034968982 | 0.173057207 |
| 1.443844 | 0.08724468 | 0.185510707 | -0.032905674 | 0.170263554 |
| 1.454084 | 0.090926 | 0.184002464 | -0.030433771 | 0.16810388 |
| 1.464324 | 0.095104714 | 0.182380783 | -0.027607644 | 0.167779886 |
| 1.474564 | 0.098925751 | 0.180692661 | -0.024711289 | 0.167714977 |
| 1.484804 | 0.102803791 | 0.179068215 | -0.021879706 | 0.168885489 |
| 1.495044 | 0.106275831 | 0.177363397 | -0.019126387 | 0.170700367 |
| 1.505284 | 0.109638456 | 0.175339375 | -0.017024944 | 0.173195264 |
| 1.515524 | 0.112214541 | 0.17297741 | -0.015433728 | 0.176155348 |
| 1.525764 | 0.114729311 | 0.169957057 | -0.01505425 | 0.180091676 |
| 1.536004 | 0.116875365 | 0.166991323 | -0.014905929 | 0.184100392 |
| 1.546245 | 0.118709483 | 0.16360509 | -0.016436379 | 0.188103474 |
| 1.556484 | 0.120655823 | 0.160673093 | -0.018021099 | 0.191875051 |
| 1.566725 | 0.12225691 | 0.157939367 | -0.020737726 | 0.194687951 |
| 1.576964 | 0.123707528 | 0.155947692 | -0.023289186 | 0.196515797 |
| 1.587205 | 0.125230237 | 0.155105263 | -0.02671233 | 0.196952483 |
| 1.597445 | 0.126259608 | 0.155407709 | -0.029724235 | 0.196018607 |
| 1.607685 | 0.127599394 | 0.15673934 | -0.032598692 | 0.193942124 |
| 1.617925 | 0.12888926 | 0.159142786 | -0.035111293 | 0.190620446 |
| 1.628165 | 0.129725916 | 0.16212399 | -0.037222804 | 0.186485206 |
| 1.638405 | 0.131249776 | 0.165323562 | -0.038307568 | 0.181881074 |
| 1.648645 | 0.131739331 | 0.168181794 | -0.038945931 | 0.176850115 |
| 1.658885 | 0.132732074 | 0.170018517 | -0.038402052 | 0.171890211 |
| 1.669125 | 0.132451467 | 0.170777318 | -0.037410818 | 0.167200839 |
| 1.679365 | 0.132053571 | 0.169506956 | -0.035736408 | 0.163041092 |
| 1.689605 | 0.130452693 | 0.166975033 | -0.033323018 | 0.159051736 |
| 1.699845 | 0.128623735 | 0.162795073 | -0.031134543 | 0.155799043 |
| 1.710085 | 0.125884004 | 0.15743005 | -0.02867757 | 0.152550088 |
| 1.720325 | 0.123721028 | 0.152279071 | -0.026307971 | 0.149593508 |
| 1.730565 | 0.121102728 | 0.146881217 | -0.023960946 | 0.146356671 |
| 1.740805 | 0.11976793 | 0.143049879 | -0.022028275 | 0.143468057 |
| 1.751045 | 0.118894089 | 0.139651334 | -0.019664781 | 0.140526109 |
| 1.761285 | 0.119085533 | 0.137898388 | -0.017496747 | 0.138102537 |
| 1.771525 | 0.119672354 | 0.13693605 | -0.015148584 | 0.136386266 |
| 1.781765 | 0.121069583 | 0.136230391 | -0.012690935 | 0.135696306 |
| 1.792005 | 0.122586984 | 0.135944047 | -0.010414625 | 0.135902469 |
| 1.802245 | 0.124249198 | 0.135499587 | -0.007942939 | 0.13703404 |
| 1.812485 | 0.125633036 | 0.134910187 | -0.005656721 | 0.138860833 |
| 1.822725 | 0.127389591 | 0.133868587 | -0.003389656 | 0.140575847 |
| 1.832965 | 0.128505782 | 0.132496666 | -0.001338317 | 0.142095326 |
| 1.843205 | 0.129959042 | 0.130666458 | 0.001194396 | 0.142497673 |
| 1.853445 | 0.131194068 | 0.128404302 | 0.003420626 | 0.142167082 |
| 1.863685 | 0.13236697 | 0.12547961 | 0.006008928 | 0.140356967 |
| 1.873925 | 0.13409819 | 0.122836813 | 0.008624927 | 0.137764181 |
| 1.884165 | 0.135543065 | 0.120043399 | 0.011213939 | 0.134311758 |
| 1.894405 | 0.137465833 | 0.116999129 | 0.013701959 | 0.131157412 |
| 1.904645 | 0.139729232 | 0.115102645 | 0.015887819 | 0.127885451 |
| 1.914886 | 0.142138204 | 0.113040084 | 0.017801297 | 0.125749614 |
| 1.925126 | 0.145246073 | 0.1120107 | 0.019364967 | 0.124482775 |
| 1.935366 | 0.148348909 | 0.111325699 | 0.020853013 | 0.124717237 |
| 1.945606 | 0.152177227 | 0.111858697 | 0.021884183 | 0.125295692 |
| 1.955846 | 0.155908209 | 0.112792618 | 0.022859725 | 0.126849607 |
| 1.966086 | 0.159755255 | 0.114824546 | 0.023606227 | 0.128659173 |
| 1.976326 | 0.163615367 | 0.117032328 | 0.024350637 | 0.130796111 |
| 1.986566 | 0.167315755 | 0.11975834 | 0.024617175 | 0.132186526 |
| 1.996806 | 0.170676066 | 0.122297813 | 0.024921653 | 0.133115837 |
| 2.007046 | 0.174094416 | 0.124822148 | 0.025018179 | 0.133042551 |
| 2.017286 | 0.177081709 | 0.126471518 | 0.024676391 | 0.131661336 |
| 2.027526 | 0.180021771 | 0.127998957 | 0.024572761 | 0.129376444 |
| 2.037766 | 0.183453731 | 0.128379586 | 0.024078797 | 0.126169199 |
| 2.048006 | 0.186600697 | 0.128791661 | 0.02347784 | 0.122690383 |
| 2.058246 | 0.190799948 | 0.128677675 | 0.022468913 | 0.119063266 |
| 2.068486 | 0.194886848 | 0.129338497 | 0.021353748 | 0.116538391 |
| 2.078726 | 0.199558281 | 0.130099511 | 0.019532714 | 0.115121532 |
| 2.088966 | 0.204616537 | 0.132343858 | 0.01696541 | 0.115121295 |
| 2.099206 | 0.209209309 | 0.135222655 | 0.013383617 | 0.117396394 |
| 2.109446 | 0.213748834 | 0.138984848 | 0.009129341 | 0.121301395 |
| 2.119686 | 0.217673773 | 0.1435683 | 0.00361054 | 0.127116783 |
| 2.129926 | 0.220948183 | 0.148127547 | -0.002532222 | 0.134310445 |
| 2.140166 | 0.223850807 | 0.153544795 | -0.009493358 | 0.141899595 |
| 2.150406 | 0.22648414 | 0.158560811 | -0.016936751 | 0.150220784 |
| 2.160646 | 0.228949377 | 0.164479656 | -0.024198928 | 0.157878582 |
| 2.170886 | 0.232155615 | 0.169507082 | -0.031490018 | 0.165227268 |
| 2.181126 | 0.235387982 | 0.174570055 | -0.038103872 | 0.171431421 |
| 2.191366 | 0.239534114 | 0.178873156 | -0.043977153 | 0.176937257 |
| 2.201606 | 0.243916654 | 0.182271523 | -0.048688013 | 0.181165745 |
| 2.211846 | 0.249248533 | 0.184267334 | -0.05271933 | 0.184246888 |
| 2.222086 | 0.254193931 | 0.185420181 | -0.054775412 | 0.186148591 |
| 2.232327 | 0.259882562 | 0.185967096 | -0.056352079 | 0.186835909 |
| 2.242567 | 0.265315982 | 0.186159614 | -0.056010306 | 0.186265557 |
| 2.252806 | 0.271225655 | 0.186388731 | -0.055034884 | 0.185064526 |
| 2.263047 | 0.277017501 | 0.187062643 | -0.052717722 | 0.182537266 |
| 2.273287 | 0.283356823 | 0.188110291 | -0.049062986 | 0.180243351 |
| 2.283527 | 0.289837144 | 0.189827856 | -0.044385796 | 0.177286677 |
| 2.293766 | 0.296775793 | 0.191207235 | -0.038545377 | 0.174545957 |
| 2.304007 | 0.303856242 | 0.192896742 | -0.031001876 | 0.172097381 |
| 2.314247 | 0.311356446 | 0.194559432 | -0.02263389 | 0.17026702 |
| 2.324487 | 0.319219972 | 0.195900435 | -0.012436305 | 0.168449173 |
| 2.334727 | 0.327077389 | 0.196985572 | -0.001230562 | 0.167857449 |
| 2.344967 | 0.335408482 | 0.197704513 | 0.011051025 | 0.167597812 |

12kHz ABR , Threshold +50dB

| | *Mir182* $^{ko}$ Wildtype | | *Mir182* $^{ko}$ Homozygote | |
|---|---|---|---|---|
| | Amplitude (µV) | | Amplitude (µV) | |
| Time (ms) | Mean | SD | Mean | SD |
| 0 | -0.23476716 | 0.217878906 | 0.098938526 | 0.178188784 |
| 0.01024 | -0.238101847 | 0.219822882 | 0.102997814 | 0.177702645 |
| 0.02048 | -0.241930349 | 0.223022537 | 0.107573256 | 0.177312663 |
| 0.03072 | -0.24592206 | 0.226972719 | 0.111375245 | 0.176396399 |
| 0.04096 | -0.249598324 | 0.232370637 | 0.114922388 | 0.175600655 |
| 0.0512 | -0.252504635 | 0.237615015 | 0.117703735 | 0.174041698 |
| 0.06144 | -0.254357559 | 0.243664122 | 0.11986815 | 0.172574347 |
| 0.07168 | -0.254386779 | 0.248510822 | 0.121603583 | 0.17080285 |
| 0.08192 | -0.253080501 | 0.252983317 | 0.122467356 | 0.169172432 |
| 0.09216 | -0.249221763 | 0.255215279 | 0.122874461 | 0.167040162 |
| 0.1024 | -0.243463727 | 0.256466355 | 0.122759077 | 0.165044938 |
| 0.11264 | -0.235311243 | 0.255522435 | 0.12202953 | 0.162458884 |
| 0.12288 | -0.225667146 | 0.253259867 | 0.121035683 | 0.159900562 |
| 0.13312 | -0.214563707 | 0.249808437 | 0.119695254 | 0.156956464 |
| 0.14336 | -0.202511992 | 0.245426488 | 0.117894255 | 0.154016968 |
| 0.1536 | -0.1906947 | 0.241241531 | 0.116010511 | 0.151177449 |
| 0.16384 | -0.17947841 | 0.237362896 | 0.114240554 | 0.148098443 |
| 0.174081 | -0.169420187 | 0.233928149 | 0.112589963 | 0.145462107 |
| 0.184321 | -0.160512994 | 0.231544877 | 0.110999654 | 0.142521641 |
| 0.194561 | -0.153499422 | 0.229945038 | 0.110511499 | 0.139766472 |
| 0.204801 | -0.147479153 | 0.228803227 | 0.110338318 | 0.137003315 |
| 0.215041 | -0.142254766 | 0.228152397 | 0.111221223 | 0.134469605 |
| 0.225281 | -0.137986001 | 0.227595641 | 0.112728274 | 0.131924062 |
| 0.235521 | -0.133783762 | 0.227275667 | 0.11543737 | 0.130102224 |
| 0.245761 | -0.12965007 | 0.226581872 | 0.118919971 | 0.1288803 |
| 0.256001 | -0.125567669 | 0.225992078 | 0.123467914 | 0.128332176 |
| 0.266241 | -0.121515241 | 0.225225647 | 0.1288186 | 0.128393039 |
| 0.276481 | -0.11743777 | 0.223726573 | 0.135066163 | 0.129340416 |
| 0.286721 | -0.113391209 | 0.222000253 | 0.141763293 | 0.130720613 |
| 0.296961 | -0.109667825 | 0.220019093 | 0.149309384 | 0.132442203 |
| 0.307201 | -0.106518444 | 0.217532234 | 0.156797196 | 0.134457557 |
| 0.317441 | -0.103501797 | 0.215581676 | 0.164186319 | 0.136395787 |
| 0.327681 | -0.101106374 | 0.214005048 | 0.170921474 | 0.1386187 |
| 0.337921 | -0.099597838 | 0.213967609 | 0.177090287 | 0.140311647 |
| 0.348161 | -0.098622606 | 0.214563732 | 0.181602221 | 0.142157453 |
| 0.358401 | -0.098345238 | 0.216805527 | 0.185182281 | 0.143509911 |
| 0.368641 | -0.098637297 | 0.219844709 | 0.186771376 | 0.144718638 |
| 0.378881 | -0.099420489 | 0.223415348 | 0.187369794 | 0.145396615 |
| 0.389121 | -0.100241165 | 0.227465714 | 0.186148945 | 0.145440004 |
| 0.399361 | -0.100697643 | 0.230930761 | 0.184182265 | 0.145567587 |
| 0.409601 | -0.10107767 | 0.234614358 | 0.181367672 | 0.144813007 |
| 0.419841 | -0.100425333 | 0.237389826 | 0.177854983 | 0.144387368 |
| 0.430081 | -0.099086171 | 0.239903457 | 0.174548812 | 0.144005387 |
| 0.440321 | -0.096859885 | 0.241864995 | 0.171217663 | 0.143857433 |
| 0.450561 | -0.093968551 | 0.243472069 | 0.167992516 | 0.143783365 |
| 0.460801 | -0.090362602 | 0.244982256 | 0.165191921 | 0.14478242 |
| 0.471041 | -0.086441565 | 0.246459801 | 0.162384787 | 0.145834726 |
| 0.481281 | -0.082900881 | 0.248199306 | 0.159452618 | 0.147446959 |
| 0.491521 | -0.079417719 | 0.250117249 | 0.156503201 | 0.149854438 |
| 0.501761 | -0.076366613 | 0.25257097 | 0.152596195 | 0.152402499 |
| 0.512001 | -0.074031762 | 0.25564993 | 0.148776458 | 0.155670012 |
| 0.522242 | -0.072247218 | 0.258966484 | 0.143625869 | 0.158869043 |
| 0.532482 | -0.070828949 | 0.263027829 | 0.138747619 | 0.162634439 |
| 0.542722 | -0.069178341 | 0.267094636 | 0.133176045 | 0.166319949 |
| 0.552962 | -0.06776865 | 0.271361604 | 0.128017177 | 0.170017927 |
| 0.563202 | -0.065579129 | 0.274744069 | 0.123156111 | 0.173613069 |
| 0.573442 | -0.062641951 | 0.277569984 | 0.119366384 | 0.176680821 |
| 0.583682 | -0.058783527 | 0.278702087 | 0.116576693 | 0.179302237 |
| 0.593922 | -0.05429228 | 0.277835547 | 0.115418978 | 0.181121037 |
| 0.604162 | -0.048322269 | 0.274709713 | 0.115628205 | 0.181864825 |
| 0.614402 | -0.042291509 | 0.268987211 | 0.117497988 | 0.181781546 |
| 0.624642 | -0.035405482 | 0.261390896 | 0.120177627 | 0.180966821 |
| 0.634882 | -0.028868742 | 0.25132542 | 0.123882303 | 0.179232819 |
| 0.645122 | -0.022336251 | 0.240144781 | 0.127741309 | 0.177263505 |
| 0.655362 | -0.016669531 | 0.228112626 | 0.130970917 | 0.175319577 |
| 0.665602 | -0.01213356 | 0.215520382 | 0.133869601 | 0.173711864 |
| 0.675842 | -0.008610324 | 0.203472451 | 0.135417795 | 0.17257447 |
| 0.686082 | -0.005983276 | 0.192299236 | 0.136095211 | 0.171618717 |
| 0.696322 | -0.00496293 | 0.182338085 | 0.135383769 | 0.172182141 |
| 0.706562 | -0.003832344 | 0.173901287 | 0.134409314 | 0.172166183 |
| 0.716802 | -0.003834574 | 0.167353859 | 0.132357296 | 0.173576298 |
| 0.727042 | -0.003534327 | 0.162148682 | 0.13035971 | 0.174371075 |
| 0.737282 | -0.003057944 | 0.158759501 | 0.128133501 | 0.175658075 |
| 0.747522 | -0.002765967 | 0.156134641 | 0.126203556 | 0.176189369 |
| 0.757762 | -0.001629143 | 0.154196062 | 0.124394737 | 0.176806739 |
| 0.768002 | 7.91727E-05 | 0.152544436 | 0.123237275 | 0.176110103 |
| 0.778242 | 0.001800202 | 0.150372434 | 0.121867773 | 0.175436341 |
| 0.788482 | 0.004621288 | 0.147931961 | 0.121326614 | 0.17389256 |
| 0.798722 | 0.007685934 | 0.144594741 | 0.120599503 | 0.172666703 |
| 0.808962 | 0.010985701 | 0.140778642 | 0.120200126 | 0.17120446 |
| 0.819202 | 0.014920477 | 0.13587627 | 0.119779304 | 0.170624645 |
| 0.829442 | 0.018846812 | 0.131250823 | 0.119295905 | 0.170089629 |
| 0.839682 | 0.022662028 | 0.126148834 | 0.11847358 | 0.170620692 |
| 0.849922 | 0.026214968 | 0.121453156 | 0.11777855 | 0.170913832 |
| 0.860162 | 0.029482127 | 0.117109546 | 0.116683298 | 0.171134442 |
| 0.870403 | 0.031854335 | 0.113461716 | 0.115412549 | 0.17087524 |
| 0.880643 | 0.033505841 | 0.110211105 | 0.114135295 | 0.170233083 |
| 0.890883 | 0.034376986 | 0.106972865 | 0.112393181 | 0.168888117 |
| 0.901123 | 0.034048391 | 0.104177343 | 0.11105606 | 0.167069902 |
| 0.911363 | 0.032966441 | 0.101347057 | 0.108912893 | 0.165620351 |
| 0.921603 | 0.031253922 | 0.099036244 | 0.10727396 | 0.164303432 |
| 0.931843 | 0.028952183 | 0.097915979 | 0.104897607 | 0.163761869 |
| 0.942083 | 0.026398576 | 0.098189989 | 0.102692735 | 0.163533635 |
| 0.952323 | 0.023993789 | 0.10058049 | 0.099742332 | 0.164415794 |
| 0.962563 | 0.02212552 | 0.105050306 | 0.09684202 | 0.164543723 |
| 0.972803 | 0.020829033 | 0.111425541 | 0.093528609 | 0.165505124 |
| 0.983043 | 0.020251236 | 0.119412392 | 0.090074118 | 0.164993221 |
| 0.993283 | 0.020669915 | 0.128276174 | 0.08672379 | 0.164144021 |
| 1.003523 | 0.021809496 | 0.137916369 | 0.083042388 | 0.162019694 |
| 1.013763 | 0.023957169 | 0.147140129 | 0.079951763 | 0.158918185 |
| 1.024003 | 0.026375201 | 0.156432816 | 0.076801866 | 0.154566086 |
| 1.034243 | 0.029705613 | 0.164258988 | 0.073543016 | 0.150005054 |
| 1.044483 | 0.033084692 | 0.171451823 | 0.070665243 | 0.14488424 |
| 1.054723 | 0.036785155 | 0.176807549 | 0.067514649 | 0.139908429 |
| 1.064963 | 0.04040079 | 0.180518529 | 0.064494229 | 0.135390666 |
| 1.075203 | 0.043228282 | 0.18246027 | 0.061301179 | 0.131619104 |
| 1.085443 | 0.045824142 | 0.182632389 | 0.058433476 | 0.128708188 |
| 1.095683 | 0.047211864 | 0.181379316 | 0.055474697 | 0.126944577 |
| 1.105923 | 0.047777979 | 0.178697229 | 0.052948891 | 0.126042094 |
| 1.116163 | 0.047172736 | 0.175460266 | 0.050319598 | 0.125720804 |
| 1.126403 | 0.046401382 | 0.171209065 | 0.048037823 | 0.126234558 |
| 1.136643 | 0.044699373 | 0.167036216 | 0.045388621 | 0.126404975 |
| 1.146883 | 0.043064471 | 0.163088388 | 0.043096211 | 0.126818542 |
| 1.157123 | 0.041596218 | 0.158895462 | 0.040216622 | 0.126202528 |
| 1.167363 | 0.040531214 | 0.155323071 | 0.037383357 | 0.125296171 |
| 1.177603 | 0.039729654 | 0.152342674 | 0.034764081 | 0.123210647 |
| 1.187843 | 0.039149727 | 0.14911296 | 0.031552969 | 0.121218095 |
| 1.198083 | 0.038704616 | 0.1462535 | 0.029234578 | 0.118318435 |
| 1.208323 | 0.03812956 | 0.143589495 | 0.026162999 | 0.11625498 |
| 1.218564 | 0.037603102 | 0.140626575 | 0.023848332 | 0.11424834 |
| 1.228804 | 0.036512809 | 0.137285697 | 0.020605808 | 0.11275313 |
| 1.239044 | 0.035617397 | 0.134539411 | 0.017791906 | 0.112210754 |
| 1.249284 | 0.034117941 | 0.131359506 | 0.014283803 | 0.111372182 |
| 1.259524 | 0.033026305 | 0.129185262 | 0.011226204 | 0.111422986 |
| 1.269764 | 0.032193048 | 0.127828089 | 0.008137554 | 0.111303348 |
| 1.280004 | 0.03117829 | 0.127905301 | 0.005707153 | 0.111645463 |
| 1.290244 | 0.031198484 | 0.129462806 | 0.003856706 | 0.1125437 |
| 1.300484 | 0.031152513 | 0.132197627 | 0.003361722 | 0.113735353 |
| 1.310724 | 0.032378277 | 0.136219158 | 0.003062534 | 0.115811249 |
| 1.320964 | 0.033539634 | 0.140368989 | 0.004003396 | 0.118204926 |
| 1.331204 | 0.035867869 | 0.144989013 | 0.005630048 | 0.121465488 |
| 1.341444 | 0.038217566 | 0.148448127 | 0.007506189 | 0.124805288 |
| 1.351684 | 0.041145423 | 0.151853515 | 0.0104547 | 0.12884204 |
| 1.361924 | 0.044239778 | 0.154007073 | 0.013827794 | 0.132643168 |
| 1.372164 | 0.047465547 | 0.155138055 | 0.018070267 | 0.136503256 |
| 1.382404 | 0.050531597 | 0.154976037 | 0.022790699 | 0.139512721 |
| 1.392644 | 0.053836922 | 0.15485176 | 0.028027971 | 0.142802562 |
| 1.402884 | 0.057104813 | 0.153480367 | 0.033189176 | 0.145099028 |
| 1.413124 | 0.060491072 | 0.15266023 | 0.038192219 | 0.147431464 |
| 1.423364 | 0.064116295 | 0.151818306 | 0.042059343 | 0.149792271 |
| 1.433604 | 0.067971764 | 0.151364905 | 0.04517071 | 0.151917604 |
| 1.443844 | 0.072431626 | 0.151601098 | 0.047259528 | 0.154408421 |
| 1.454084 | 0.07659894 | 0.152504305 | 0.048422466 | 0.1569876 |
| 1.464324 | 0.081454497 | 0.154052113 | 0.04934545 | 0.1593439 |
| 1.474564 | 0.085965032 | 0.155969591 | 0.050169696 | 0.162317037 |
| 1.484804 | 0.090531531 | 0.157921502 | 0.051548129 | 0.165204309 |
| 1.495044 | 0.094937875 | 0.160357653 | 0.053775693 | 0.168683324 |
| 1.505284 | 0.099060077 | 0.16173069 | 0.056968826 | 0.172274372 |
| 1.515524 | 0.102842967 | 0.163030034 | 0.061090454 | 0.176024677 |
| 1.525764 | 0.106400192 | 0.164004824 | 0.06551155 | 0.179269554 |
| 1.536004 | 0.109739831 | 0.164570141 | 0.070776581 | 0.182135292 |
| 1.546245 | 0.112962111 | 0.16584638 | 0.075671134 | 0.183728434 |
| 1.556484 | 0.11621684 | 0.167555731 | 0.080117438 | 0.183709883 |
| 1.566725 | 0.119472171 | 0.170659892 | 0.084621486 | 0.182989361 |
| 1.576964 | 0.123010964 | 0.175356764 | 0.088002323 | 0.180539664 |
| 1.587205 | 0.126855195 | 0.181053808 | 0.091288343 | 0.177816902 |
| 1.597445 | 0.131315422 | 0.187798512 | 0.094534588 | 0.174732825 |
| 1.607685 | 0.135932184 | 0.195403283 | 0.09783837 | 0.172215261 |
| 1.617925 | 0.141209096 | 0.202875064 | 0.1017969 | 0.170670669 |
| 1.628165 | 0.147016848 | 0.209982552 | 0.106046666 | 0.170288303 |
| 1.638405 | 0.152945447 | 0.216411934 | 0.11115693 | 0.171264233 |
| 1.648645 | 0.159437085 | 0.221823409 | 0.116938005 | 0.173012979 |
| 1.658885 | 0.166238211 | 0.225934072 | 0.122719712 | 0.176040057 |
| 1.669125 | 0.173081458 | 0.228329806 | 0.128192271 | 0.178829326 |
| 1.679365 | 0.180463367 | 0.229755728 | 0.133270769 | 0.181709613 |
| 1.689605 | 0.187830345 | 0.229800155 | 0.137458249 | 0.184415679 |
| 1.699845 | 0.195536285 | 0.228830728 | 0.140105866 | 0.186340343 |
| 1.710085 | 0.203161903 | 0.22732079 | 0.141546247 | 0.187860325 |
| 1.720325 | 0.210899195 | 0.225507538 | 0.14134008 | 0.18896106 |
| 1.730565 | 0.218109422 | 0.223338215 | 0.140219207 | 0.190112407 |
| 1.740805 | 0.225380031 | 0.221011012 | 0.137312513 | 0.191333677 |
| 1.751045 | 0.231620481 | 0.218691431 | 0.134086551 | 0.193024858 |
| 1.761285 | 0.237289875 | 0.21639553 | 0.129635523 | 0.195108358 |
| 1.771525 | 0.24243913 | 0.213942853 | 0.125214783 | 0.198121477 |
| 1.781765 | 0.246977073 | 0.211775246 | 0.120614923 | 0.201253748 |
| 1.792005 | 0.25122833 | 0.209670678 | 0.116300537 | 0.205248288 |
| 1.802245 | 0.255686282 | 0.207715342 | 0.112627331 | 0.208870472 |
| 1.812485 | 0.260187295 | 0.205805505 | 0.109544466 | 0.212634505 |
| 1.822725 | 0.265628436 | 0.203870209 | 0.10778325 | 0.216112357 |
| 1.832965 | 0.271567989 | 0.201784817 | 0.107080118 | 0.219595339 |
| 1.843205 | 0.278723543 | 0.199995505 | 0.10806269 | 0.222415041 |
| 1.853445 | 0.286310104 | 0.197848202 | 0.110485997 | 0.225292542 |
| 1.863685 | 0.295499256 | 0.195803088 | 0.114745191 | 0.228321944 |
| 1.873925 | 0.305452197 | 0.194217687 | 0.120950477 | 0.230945048 |
| 1.884165 | 0.316711841 | 0.192773005 | 0.12873078 | 0.233795496 |
| 1.894405 | 0.329267423 | 0.191773982 | 0.137634062 | 0.236333872 |
| 1.904645 | 0.343193218 | 0.190951132 | 0.148531764 | 0.238745409 |
| 1.914886 | 0.358866851 | 0.191044665 | 0.159306179 | 0.240717222 |
| 1.925126 | 0.375972645 | 0.1913574 | 0.17092336 | 0.242451511 |
| 1.935366 | 0.394709526 | 0.192416329 | 0.182821141 | 0.243887368 |
| 1.945606 | 0.415104779 | 0.194195472 | 0.194832925 | 0.244565706 |
| 1.955846 | 0.437386116 | 0.197118654 | 0.206777005 | 0.24478703 |
| 1.966086 | 0.460851093 | 0.200903266 | 0.219222624 | 0.245362397 |
| 1.976326 | 0.486426023 | 0.206649986 | 0.232324892 | 0.244887402 |
| 1.986566 | 0.51333417 | 0.213582788 | 0.245976532 | 0.24468532 |
| 1.996806 | 0.54262118 | 0.223334485 | 0.260909001 | 0.244600095 |
| 2.007046 | 0.572954956 | 0.234361842 | 0.276646097 | 0.245339908 |
| 2.017286 | 0.606016955 | 0.247981958 | 0.29421335 | 0.246534706 |
| 2.027526 | 0.640486801 | 0.262824815 | 0.312439807 | 0.249338519 |
| 2.037766 | 0.677536109 | 0.279096079 | 0.332591313 | 0.253397567 |
| 2.048006 | 0.715922755 | 0.295283712 | 0.353824753 | 0.25930877 |
| 2.058246 | 0.756840497 | 0.312520993 | 0.376832165 | 0.266114855 |
| 2.068486 | 0.799120393 | 0.327927738 | 0.400978367 | 0.27474822 |
| 2.078726 | 0.843054867 | 0.343397669 | 0.426987771 | 0.284146252 |
| 2.088966 | 0.888062233 | 0.356563156 | 0.454246392 | 0.294546768 |
| 2.099206 | 0.934497017 | 0.368389128 | 0.482919634 | 0.305373951 |
| 2.109446 | 0.981007129 | 0.377942102 | 0.512571994 | 0.317533718 |
| 2.119686 | 1.02820169 | 0.385379385 | 0.543225313 | 0.329870351 |
| 2.129926 | 1.075281671 | 0.390008481 | 0.574363 | 0.34329918 |
| 2.140166 | 1.122094232 | 0.39232215 | 0.605494745 | 0.356874649 |
| 2.150406 | 1.167960346 | 0.392363923 | 0.636875936 | 0.371217223 |
| 2.160646 | 1.2129152 | 0.389949526 | 0.667377328 | 0.385351177 |
| 2.170886 | 1.256020433 | 0.385962723 | 0.697997044 | 0.399338103 |
| 2.181126 | 1.297344375 | 0.380288246 | 0.727691784 | 0.412897583 |
| 2.191366 | 1.33608346 | 0.373975463 | 0.757673858 | 0.426012975 |
| 2.201606 | 1.372829115 | 0.367308029 | 0.78736522 | 0.437619241 |
| 2.211846 | 1.406777984 | 0.360840678 | 0.817721613 | 0.448810804 |
| 2.222086 | 1.438735729 | 0.355373161 | 0.848779581 | 0.458434809 |
| 2.232327 | 1.468214232 | 0.351126923 | 0.880516632 | 0.466675038 |
| 2.242567 | 1.496138394 | 0.348294092 | 0.913228645 | 0.473886748 |
| 2.252806 | 1.521986198 | 0.346896354 | 0.946165642 | 0.479335925 |
| 2.263047 | 1.545671594 | 0.346853926 | 0.979902775 | 0.483388787 |
| 2.273287 | 1.567644107 | 0.348022462 | 1.012764649 | 0.485889601 |
| 2.283527 | 1.586474288 | 0.349383546 | 1.045277216 | 0.486939062 |
| 2.293766 | 1.602585697 | 0.351306806 | 1.076394456 | 0.486821346 |
| 2.304007 | 1.614685118 | 0.353282383 | 1.10563906 | 0.485717064 |
| 2.314247 | 1.622741783 | 0.355174798 | 1.132749637 | 0.483868218 |
| 2.324487 | 1.626031494 | 0.357177113 | 1.157182969 | 0.481369668 |
| 2.334727 | 1.624452698 | 0.359476862 | 1.178409775 | 0.478430303 |
| 2.344967 | 1.617261815 | 0.361961724 | 1.19577679 | 0.474756023 |

| | | | | |
|---|---|---|---|---|
| 2.355207 | 0.329467472 | 0.317342584 | 0.332702815 | 0.29194631 |
| 2.365447 | 0.335713609 | 0.320249473 | 0.345418844 | 0.290255 |
| 2.375687 | 0.341195624 | 0.322802331 | 0.359512937 | 0.288091 |
| 2.385927 | 0.346135788 | 0.324878709 | 0.374305155 | 0.28504368 |
| 2.396167 | 0.351062155 | 0.327343218 | 0.38941215 | 0.28229408 |
| 2.406407 | 0.355239477 | 0.328938542 | 0.404261568 | 0.2789547 |
| 2.416647 | 0.358945788 | 0.330997312 | 0.418136586 | 0.27599683 |
| 2.426887 | 0.362318168 | 0.331312807 | 0.430970141 | 0.27220207 |
| 2.437127 | 0.364681075 | 0.33186185 | 0.441811926 | 0.26890446 |
| 2.447367 | 0.366991187 | 0.330665008 | 0.451654673 | 0.26490215 |
| 2.457607 | 0.369037834 | 0.328067184 | 0.459503538 | 0.26029252 |
| 2.467847 | 0.371851292 | 0.324273536 | 0.466831424 | 0.25544577 |
| 2.478087 | 0.374822346 | 0.319187348 | 0.473907884 | 0.24922408 |
| 2.488327 | 0.379071581 | 0.313235528 | 0.480568647 | 0.24317144 |
| 2.498567 | 0.383858698 | 0.306406624 | 0.488118585 | 0.23576476 |
| 2.508807 | 0.389970146 | 0.299076393 | 0.496198501 | 0.2283743 |
| 2.519047 | 0.396186486 | 0.29158372 | 0.504514158 | 0.22086079 |
| 2.529287 | 0.402804876 | 0.283985714 | 0.512661938 | 0.21376915 |
| 2.539527 | 0.408826758 | 0.276592041 | 0.521136791 | 0.20699201 |
| 2.549767 | 0.414684107 | 0.268984847 | 0.527949278 | 0.20152975 |
| 2.560007 | 0.419870898 | 0.261679555 | 0.533599202 | 0.19712957 |
| 2.570247 | 0.423489864 | 0.254106995 | 0.537079313 | 0.19374611 |
| 2.580487 | 0.427145878 | 0.246478216 | 0.53939437 | 0.19178917 |
| 2.590728 | 0.429383397 | 0.237819257 | 0.539334778 | 0.18953579 |
| 2.600967 | 0.430735633 | 0.229311111 | 0.537669995 | 0.18900053 |
| 2.611207 | 0.430837092 | 0.219373195 | 0.534933341 | 0.18689222 |
| 2.621448 | 0.429724368 | 0.209868917 | 0.531498343 | 0.18604887 |
| 2.631688 | 0.426911888 | 0.199447125 | 0.526663444 | 0.18470255 |
| 2.641928 | 0.422149276 | 0.189933947 | 0.522151611 | 0.18413666 |
| 2.652168 | 0.415716837 | 0.180023319 | 0.516717787 | 0.18460876 |
| 2.662408 | 0.407624205 | 0.171738482 | 0.51121484 | 0.18724093 |
| 2.672648 | 0.398096892 | 0.164150729 | 0.504910967 | 0.19146227 |
| 2.682888 | 0.387800989 | 0.158687669 | 0.498283635 | 0.19813105 |
| 2.693128 | 0.376986799 | 0.155933829 | 0.490651199 | 0.20643111 |
| 2.703368 | 0.36649742 | 0.156260186 | 0.482017189 | 0.21567957 |
| 2.713608 | 0.355781057 | 0.161165855 | 0.472643403 | 0.22517091 |
| 2.723848 | 0.345195481 | 0.169123147 | 0.461787389 | 0.23464999 |
| 2.734088 | 0.334777998 | 0.180690752 | 0.450109757 | 0.2423051 |
| 2.744328 | 0.323009414 | 0.193515927 | 0.436413438 | 0.24919727 |
| 2.754568 | 0.310018978 | 0.208070552 | 0.421898014 | 0.25405553 |
| 2.764808 | 0.295168793 | 0.221651438 | 0.405238901 | 0.25762794 |
| 2.775048 | 0.27737842 | 0.235856727 | 0.387267744 | 0.260226 |
| 2.785288 | 0.257125739 | 0.249048099 | 0.367225418 | 0.26144429 |
| 2.795528 | 0.233597356 | 0.262331577 | 0.345704543 | 0.2625322 |
| 2.805768 | 0.207917109 | 0.275820948 | 0.32237706 | 0.26291525 |
| 2.816008 | 0.179449677 | 0.288813849 | 0.298330788 | 0.26392687 |
| 2.826248 | 0.149590098 | 0.302915295 | 0.273370023 | 0.26524318 |
| 2.836488 | 0.118061161 | 0.316483681 | 0.247859473 | 0.26705741 |
| 2.846728 | 0.086526809 | 0.32934888 | 0.222711477 | 0.26962129 |
| 2.856968 | 0.053872014 | 0.341971508 | 0.197582628 | 0.27321764 |
| 2.867208 | 0.021443539 | 0.352796854 | 0.172731549 | 0.27679458 |
| 2.877448 | -0.010905822 | 0.36235929 | 0.147904385 | 0.28137312 |
| 2.887688 | -0.044136374 | 0.370475309 | 0.12279443 | 0.28542832 |
| 2.897928 | -0.076921888 | 0.376946002 | 0.097323422 | 0.28986059 |
| 2.908168 | -0.110625794 | 0.382611102 | 0.07112671 | 0.29390034 |
| 2.918408 | -0.144769599 | 0.386534303 | 0.04392977 | 0.29783189 |
| 2.928648 | -0.178972617 | 0.389351549 | 0.016579994 | 0.301931 |
| 2.938889 | -0.212976639 | 0.391117863 | -0.011233244 | 0.30573841 |
| 2.949129 | -0.246668667 | 0.391431292 | -0.038956572 | 0.3092908 |
| 2.959368 | -0.278838992 | 0.390510998 | -0.066081047 | 0.31256951 |
| 2.969609 | -0.309305554 | 0.388617299 | -0.092352798 | 0.31483017 |
| 2.979849 | -0.337730189 | 0.385033664 | -0.117316357 | 0.31564516 |
| 2.990089 | -0.36319537 | 0.381232036 | -0.141433608 | 0.31530071 |
| 3.000329 | -0.386284123 | 0.375777522 | -0.164117336 | 0.3131335 |
| 3.010569 | -0.406419093 | 0.369742974 | -0.185110335 | 0.30983571 |
| 3.020809 | -0.424652969 | 0.362565592 | -0.205266573 | 0.30523573 |
| 3.031049 | -0.440080086 | 0.354151815 | -0.223646095 | 0.30079339 |
| 3.041289 | -0.454584631 | 0.344095573 | -0.240246762 | 0.29600945 |
| 3.051529 | -0.467342968 | 0.332246259 | -0.255570973 | 0.29210453 |
| 3.061769 | -0.479774552 | 0.318955114 | -0.26928226 | 0.28908242 |
| 3.072009 | -0.491265758 | 0.303346863 | -0.281631623 | 0.28689875 |
| 3.082249 | -0.503297308 | 0.287505374 | -0.293152571 | 0.28581499 |
| 3.092489 | -0.514844211 | 0.270278827 | -0.303500052 | 0.28576667 |
| 3.102729 | -0.526444182 | 0.253441715 | -0.31362892 | 0.28587296 |
| 3.112969 | -0.538418638 | 0.237313454 | -0.322866342 | 0.28711026 |
| 3.123209 | -0.549764924 | 0.223020477 | -0.331745746 | 0.28809648 |
| 3.133449 | -0.561075514 | 0.210369726 | -0.339281429 | 0.28952107 |
| 3.143689 | -0.570801688 | 0.201430878 | -0.345940085 | 0.28991131 |
| 3.153929 | -0.580351437 | 0.194485184 | -0.350517812 | 0.29071442 |
| 3.164169 | -0.586821953 | 0.19206147 | -0.353182058 | 0.28967626 |
| 3.174409 | -0.592176874 | 0.19206474 | -0.35338743 | 0.28959747 |
| 3.184649 | -0.593642508 | 0.195679869 | -0.351358057 | 0.28823633 |
| 3.194889 | -0.592907091 | 0.202577126 | -0.346784869 | 0.28746767 |
| 3.205129 | -0.587349688 | 0.211335946 | -0.340125438 | 0.28673615 |
| 3.215369 | -0.578813573 | 0.222698571 | -0.331666199 | 0.28764163 |
| 3.225609 | -0.566347544 | 0.234907545 | -0.322089493 | 0.2892699 |
| 3.235849 | -0.550519556 | 0.24811295 | -0.311504201 | 0.29278961 |
| 3.246089 | -0.531236157 | 0.260796073 | -0.30038743 | 0.29756356 |
| 3.256329 | -0.510304707 | 0.273016317 | -0.289256051 | 0.30357211 |
| 3.266569 | -0.486525372 | 0.283764528 | -0.277605701 | 0.31050994 |
| 3.276809 | -0.461533638 | 0.293157521 | -0.265986487 | 0.31809481 |
| 3.28705 | -0.434933299 | 0.301548452 | -0.253577788 | 0.32575969 |
| 3.29729 | -0.407581442 | 0.308750615 | -0.240573438 | 0.33336795 |
| 3.307529 | -0.378936343 | 0.316142292 | -0.226287744 | 0.34102777 |
| 3.31777 | -0.350045411 | 0.322809627 | -0.21135377 | 0.34854886 |
| 3.32801 | -0.320718326 | 0.330431929 | -0.195124613 | 0.35589562 |
| 3.33825 | -0.292250517 | 0.337724868 | -0.178255071 | 0.36263758 |
| 3.34849 | -0.264159906 | 0.345378943 | -0.160701718 | 0.36905946 |
| 3.35873 | -0.237731118 | 0.352395566 | -0.143284379 | 0.37380387 |
| 3.36897 | -0.212954033 | 0.358850846 | -0.125718801 | 0.37711898 |
| 3.37921 | -0.190597169 | 0.364245593 | -0.108406642 | 0.37759956 |
| 3.38945 | -0.170308838 | 0.367984793 | -0.091688882 | 0.37594334 |
| 3.39969 | -0.15253782 | 0.370015286 | -0.074919374 | 0.3705796 |
| 3.40993 | -0.136624621 | 0.369747381 | -0.059115074 | 0.36269489 |
| 3.42017 | -0.122770485 | 0.367599471 | -0.04325485 | 0.3524812 |
| 3.43041 | -0.110736364 | 0.362555732 | -0.028150295 | 0.34057143 |
| 3.44065 | -0.099537255 | 0.355375775 | -0.013159962 | 0.32763084 |
| 3.45089 | -0.089532546 | 0.345866612 | 0.000521136 | 0.31561984 |
| 3.46113 | -0.079648768 | 0.334333939 | 0.014730541 | 0.30460034 |
| 3.47137 | -0.069838817 | 0.321398866 | 0.027450701 | 0.29575288 |
| 3.48161 | -0.059166545 | 0.307341702 | 0.040213536 | 0.28939854 |
| 3.49185 | -0.047928253 | 0.293084873 | 0.052115238 | 0.28559915 |
| 3.50209 | -0.035293785 | 0.27928078 | 0.063479958 | 0.28438796 |
| 3.51233 | -0.02151121 | 0.266539525 | 0.074338773 | 0.28459724 |
| 3.52257 | -0.006830638 | 0.254897067 | 0.0843722 | 0.28657663 |
| 3.53281 | 0.007985489 | 0.245671762 | 0.093441149 | 0.28975929 |
| 3.54305 | 0.022805118 | 0.238375504 | 0.101234986 | 0.2924919 |
| 3.55329 | 0.036317389 | 0.233266768 | 0.108138879 | 0.29568031 |
| 3.56353 | 0.047837753 | 0.230053491 | 0.113255472 | 0.29860589 |
| 3.57377 | 0.05680588 | 0.228567389 | 0.117530667 | 0.30073629 |
| 3.58401 | 0.063433494 | 0.229499446 | 0.120508425 | 0.30267087 |
| 3.59425 | 0.066556473 | 0.231717901 | 0.122731537 | 0.30449766 |
| 3.60449 | 0.066969747 | 0.235503599 | 0.123972572 | 0.30622132 |
| 3.61473 | 0.065124784 | 0.241094737 | 0.124582315 | 0.30846993 |
| 3.62497 | 0.060888734 | 0.24826622 | 0.124521914 | 0.31160047 |
| 3.635211 | 0.054850588 | 0.256641657 | 0.122957293 | 0.31509093 |
| 3.645451 | 0.047348678 | 0.267498517 | 0.120041399 | 0.31980478 |
| 3.65569 | 0.038773465 | 0.279101951 | 0.114555915 | 0.32418342 |
| 3.665931 | 0.029302413 | 0.293465919 | 0.107236654 | 0.32884944 |
| 3.676171 | 0.019186673 | 0.308637478 | 0.096207967 | 0.3326029 |
| 3.686411 | 0.009081652 | 0.325434938 | 0.083729396 | 0.33595056 |
| 3.696651 | -0.001340153 | 0.342129154 | 0.067699028 | 0.33811627 |
| 3.706891 | -0.01151077 | 0.359039599 | 0.051229649 | 0.33948199 |
| 3.717131 | -0.021181968 | 0.374229688 | 0.032816359 | 0.34068038 |
| 3.727371 | -0.030907732 | 0.387743144 | 0.01494822 | 0.34185685 |
| 3.737611 | -0.040928362 | 0.398576165 | -0.002847293 | 0.34309532 |
| 3.747851 | -0.051012276 | 0.406445674 | -0.020055349 | 0.34506839 |
| 3.758091 | -0.062166294 | 0.412180652 | -0.036502052 | 0.34766634 |
| 3.768331 | -0.074887708 | 0.415187234 | -0.052563071 | 0.35102193 |
| 3.778571 | -0.088899457 | 0.416779823 | -0.068169218 | 0.35500475 |
| 3.788811 | -0.104750104 | 0.417359162 | -0.084077132 | 0.35910819 |
| 3.799051 | -0.122521163 | 0.417248386 | -0.099981917 | 0.36384529 |
| 3.809291 | -0.141396012 | 0.41709656 | -0.116496616 | 0.3681197 |
| 3.819531 | -0.162197986 | 0.417069593 | -0.132477158 | 0.37279 |
| 3.829771 | -0.182724131 | 0.416712669 | -0.149024846 | 0.37651258 |
| 3.840011 | -0.204098217 | 0.417034871 | -0.164942703 | 0.38030152 |
| 3.850251 | -0.224801277 | 0.41628067 | -0.180418242 | 0.38415774 |
| 3.860491 | -0.245221411 | 0.416126646 | -0.195515758 | 0.38743292 |
| 3.870731 | -0.264186862 | 0.414639071 | -0.21006775 | 0.39114451 |
| 3.880971 | -0.281949256 | 0.413363029 | -0.224168652 | 0.39529178 |
| 3.891211 | -0.298455127 | 0.410971203 | -0.238282369 | 0.39970522 |
| 3.901451 | -0.312592403 | 0.408404894 | -0.252308657 | 0.40410723 |
| 3.911691 | -0.325831297 | 0.40551234 | -0.266745915 | 0.40839929 |
| 3.921931 | -0.336626196 | 0.401702111 | -0.280994504 | 0.41237802 |
| 3.932171 | -0.346722222 | 0.397698966 | -0.295925947 | 0.41441699 |
| 3.942411 | -0.355235764 | 0.39228899 | -0.310971527 | 0.41501588 |
| 3.952652 | -0.363650192 | 0.385580166 | -0.326027412 | 0.41326352 |
| 3.962891 | -0.371943248 | 0.376416293 | -0.341456054 | 0.4088572 |
| 3.973131 | -0.380099393 | 0.365702951 | -0.356369389 | 0.40141773 |
| 3.983371 | -0.38822501 | 0.352278562 | -0.370997139 | 0.39183563 |
| 3.993612 | -0.396101288 | 0.336516624 | -0.384139998 | 0.37984172 |
| 4.003851 | -0.403793267 | 0.319178911 | -0.396551487 | 0.36607934 |
| 4.014091 | -0.409548586 | 0.299858745 | -0.406617145 | 0.35124912 |
| 4.024332 | -0.414660816 | 0.28034712 | -0.415319278 | 0.33646248 |
| 4.034572 | -0.417881786 | 0.259876322 | -0.421745552 | 0.32186186 |
| 4.044812 | -0.420041079 | 0.240231706 | -0.426602641 | 0.30836225 |
| 4.055052 | -0.421066039 | 0.221482613 | -0.430203262 | 0.29640717 |
| 4.065292 | -0.420484234 | 0.204219117 | -0.432460614 | 0.28635969 |
| 4.075532 | -0.420496444 | 0.189724925 | -0.434382142 | 0.27845421 |
| 4.085772 | -0.41922351 | 0.177644651 | -0.435548306 | 0.27210898 |
| 4.096012 | -0.418008859 | 0.169676719 | -0.436289833 | 0.26826983 |
| 4.106252 | -0.415750707 | 0.165662637 | -0.43650791 | 0.26473629 |
| 4.116492 | -0.41368451 | 0.165376309 | -0.436286172 | 0.26330279 |
| 4.126732 | -0.409843631 | 0.169652164 | -0.43504398 | 0.26199859 |
| 4.136972 | -0.405698985 | 0.176836431 | -0.432947279 | 0.26143824 |
| 4.147212 | -0.39957806 | 0.186335601 | -0.429552009 | 0.26175651 |
| 4.157452 | -0.392852278 | 0.197555268 | -0.424748443 | 0.26252591 |
| 4.167692 | -0.384252388 | 0.209288539 | -0.418410059 | 0.26404082 |
| 4.177932 | -0.374905714 | 0.22142979 | -0.410490186 | 0.26593487 |
| 4.188172 | -0.36367051 | 0.232367388 | -0.400186558 | 0.26790449 |
| 4.198412 | -0.352377119 | 0.243340351 | -0.388235543 | 0.26974907 |
| 4.208652 | -0.339914513 | 0.252871144 | -0.374590546 | 0.27118158 |
| 4.218892 | -0.327620334 | 0.261993653 | -0.359499203 | 0.27153968 |
| 4.229132 | -0.315292033 | 0.269946055 | -0.343208858 | 0.2714408 |
| 4.239372 | -0.303241481 | 0.277887347 | -0.326768043 | 0.27111331 |
| 4.249612 | -0.292401209 | 0.28582058 | -0.310208487 | 0.27059861 |
| 4.259852 | -0.281713623 | 0.293624747 | -0.294218416 | 0.27036594 |
| 4.270092 | -0.271923698 | 0.302476691 | -0.279154839 | 0.27087744 |
| 4.280333 | -0.262539814 | 0.311852099 | -0.264502496 | 0.27205539 |
| 4.290572 | -0.252932109 | 0.322634805 | -0.251117126 | 0.27285481 |
| 4.300812 | -0.243165525 | 0.333939173 | -0.237917734 | 0.27400324 |
| 4.311052 | -0.232777332 | 0.346420067 | -0.225584987 | 0.27444052 |
| 4.321292 | -0.221582147 | 0.359012923 | -0.212878928 | 0.27424536 |
| 4.331532 | -0.210147767 | 0.371992918 | -0.200769259 | 0.2728593 |
| 4.341773 | -0.19850097 | 0.38428268 | -0.187946529 | 0.27077384 |
| 4.352013 | -0.187154309 | 0.39531984 | -0.175581565 | 0.26854118 |
| 4.362253 | -0.176656285 | 0.404533263 | -0.161675983 | 0.26516477 |
| 4.372493 | -0.16722572 | 0.412464044 | -0.148366602 | 0.26218435 |
| 4.382733 | -0.159249434 | 0.417257464 | -0.13394998 | 0.25926062 |
| 4.392972 | -0.152267939 | 0.419657811 | -0.119680224 | 0.25689354 |
| 4.403213 | -0.146393998 | 0.419896418 | -0.104637539 | 0.25468501 |
| 4.413453 | -0.140734411 | 0.41723524 | -0.090519734 | 0.25328821 |
| 4.423693 | -0.135091096 | 0.411958112 | -0.076041417 | 0.25216926 |
| 4.433933 | -0.128721283 | 0.404738254 | -0.062676542 | 0.25074744 |
| 4.444173 | -0.121773602 | 0.394710778 | -0.050199046 | 0.24897925 |
| 4.454413 | -0.113639825 | 0.382863676 | -0.039321704 | 0.24679606 |
| 4.464653 | -0.104564597 | 0.368601507 | -0.029219755 | 0.2440252 |
| 4.474893 | -0.094704124 | 0.353177684 | -0.021274354 | 0.24014849 |
| 4.485133 | -0.084598447 | 0.335868655 | -0.014771786 | 0.23594 |
| 4.495373 | -0.074617962 | 0.318273492 | -0.009721674 | 0.23220779 |
| 4.505613 | -0.064618585 | 0.299973491 | -0.00668694 | 0.22796792 |
| 4.515853 | -0.05569165 | 0.281517047 | -0.004814318 | 0.22446441 |
| 4.526093 | -0.047574619 | 0.264202946 | -0.005345187 | 0.22116214 |
| 4.536333 | -0.040303492 | 0.247895081 | -0.006760277 | 0.2194709 |
| 4.546573 | -0.034637482 | 0.233955406 | -0.010832624 | 0.21749526 |
| 4.556813 | -0.029096353 | 0.222797213 | -0.015623124 | 0.21667873 |
| 4.567053 | -0.025422359 | 0.215810375 | -0.022648438 | 0.21710577 |
| 4.577293 | -0.021900901 | 0.212301137 | -0.030257501 | 0.21787049 |
| 4.587533 | -0.01954064 | 0.212994138 | -0.039118031 | 0.2195347 |
| 4.597773 | -0.017705258 | 0.21667786 | -0.047957086 | 0.22203592 |
| 4.608013 | -0.016940592 | 0.222990705 | -0.057040737 | 0.22531924 |
| 4.618253 | -0.016513778 | 0.231504941 | -0.065639213 | 0.2291502 |
| 4.628493 | -0.017019798 | 0.240732239 | -0.074151998 | 0.23319072 |
| 4.638733 | -0.018092148 | 0.251763886 | -0.081917203 | 0.23825251 |
| 4.648973 | -0.01901415 | 0.262289043 | -0.089566715 | 0.24257854 |
| 4.659214 | -0.020792171 | 0.273785628 | -0.097320136 | 0.24772888 |
| 4.669454 | -0.021989703 | 0.284980241 | -0.104989881 | 0.25216664 |
| 4.679694 | -0.023244418 | 0.295484921 | -0.113305034 | 0.25635558 |
| 4.689933 | -0.024247507 | 0.305198416 | -0.12214452 | 0.26046798 |
| 4.700173 | -0.025036881 | 0.314066202 | -0.131522578 | 0.26342814 |
| 4.710413 | -0.025653666 | 0.321578228 | -0.141707619 | 0.26629081 |
| 4.720654 | -0.02605133 | 0.327631397 | -0.152499123 | 0.26807084 |
| 4.730894 | -0.026367249 | 0.33218934 | -0.163526799 | 0.26931696 |
| 4.741134 | -0.026928952 | 0.335884651 | -0.174847452 | 0.26950474 |
| 4.751374 | -0.02649112 | 0.337767804 | -0.185927963 | 0.26869569 |

| | | | | |
|---|---|---|---|---|
| 2.355207 | 1.413126667 | 0.202420994 | 1.152896856 | 0.179411024 |
| 2.365447 | 1.402891994 | 0.222823297 | 1.148019527 | 0.184971057 |
| 2.375687 | 1.3879505 | 0.24735268 | 1.138907773 | 0.193296452 |
| 2.385927 | 1.368769328 | 0.274295747 | 1.125086835 | 0.204019193 |
| 2.396167 | 1.34493951 | 0.302927744 | 1.108794025 | 0.216356596 |
| 2.406407 | 1.317702393 | 0.333482697 | 1.087889654 | 0.230348822 |
| 2.416647 | 1.286087881 | 0.365127881 | 1.064447692 | 0.245720985 |
| 2.426887 | 1.251311441 | 0.3983155 | 1.037390087 | 0.262537762 |
| 2.437127 | 1.212386698 | 0.433305558 | 1.007072823 | 0.280517332 |
| 2.447367 | 1.170863201 | 0.46894312 | 0.973604671 | 0.299213522 |
| 2.457607 | 1.125436743 | 0.506502115 | 0.936801485 | 0.319304137 |
| 2.467847 | 1.076407075 | 0.543924336 | 0.896499051 | 0.339537094 |
| 2.478087 | 1.024065966 | 0.582220405 | 0.853712133 | 0.360275315 |
| 2.488327 | 0.967387629 | 0.620752741 | 0.807275568 | 0.381624971 |
| 2.498567 | 0.906669433 | 0.659091828 | 0.75893883 | 0.403462769 |
| 2.508807 | 0.841366642 | 0.697172187 | 0.707843129 | 0.425112132 |
| 2.519047 | 0.771680772 | 0.734492573 | 0.655417775 | 0.447323245 |
| 2.529287 | 0.697476685 | 0.770305671 | 0.601155138 | 0.469225894 |
| 2.539527 | 0.61969621 | 0.803806249 | 0.54644336 | 0.490370005 |
| 2.549767 | 0.537191158 | 0.834327994 | 0.490430794 | 0.510553366 |
| 2.560007 | 0.452658847 | 0.860390534 | 0.434205902 | 0.528899033 |
| 2.570247 | 0.364823636 | 0.881980777 | 0.377554453 | 0.545603079 |
| 2.580487 | 0.275374404 | 0.898265244 | 0.319603703 | 0.559282159 |
| 2.590728 | 0.184544204 | 0.909438872 | 0.261174866 | 0.571308651 |
| 2.600967 | 0.093597477 | 0.914751079 | 0.201074046 | 0.57992442 |
| 2.611207 | 0.002944912 | 0.9159045 | 0.13940715 | 0.586197587 |
| 2.621448 | -0.086134362 | 0.911662577 | 0.076018915 | 0.58944319 |
| 2.631688 | -0.173689593 | 0.903791365 | 0.010841243 | 0.589864155 |
| 2.641928 | -0.258210584 | 0.891242525 | -0.055888415 | 0.587748627 |
| 2.652168 | -0.340452668 | 0.876075798 | -0.123764509 | 0.583003165 |
| 2.662408 | -0.419515145 | 0.856734119 | -0.191845438 | 0.576685997 |
| 2.672648 | -0.496022475 | 0.835559209 | -0.260286569 | 0.5687835 |
| 2.682888 | -0.569676687 | 0.811240412 | -0.327363114 | 0.560450706 |
| 2.693128 | -0.641479423 | 0.786131449 | -0.393626401 | 0.551492793 |
| 2.703368 | -0.710510572 | 0.758788436 | -0.457594098 | 0.542843935 |
| 2.713608 | -0.777451684 | 0.731623644 | -0.519586917 | 0.533544948 |
| 2.723848 | -0.841937626 | 0.703183964 | -0.579045595 | 0.524412602 |
| 2.734088 | -0.904007728 | 0.675328356 | -0.635972018 | 0.513866819 |
| 2.744328 | -0.962421574 | 0.645591102 | -0.689751281 | 0.502410392 |
| 2.754568 | -1.017979657 | 0.616401808 | -0.740897975 | 0.489236736 |
| 2.764808 | -1.069160917 | 0.585616148 | -0.788548448 | 0.47469793 |
| 2.775048 | -1.116074006 | 0.553622593 | -0.833168615 | 0.458388134 |
| 2.785288 | -1.158084497 | 0.520760997 | -0.873852896 | 0.441738198 |
| 2.795528 | -1.194313372 | 0.48775522 | -0.911322355 | 0.424286557 |
| 2.805768 | -1.224856347 | 0.455446936 | -0.945509821 | 0.407651837 |
| 2.816008 | -1.24872828 | 0.426771714 | -0.975353207 | 0.391716785 |
| 2.826248 | -1.266051958 | 0.402821811 | -1.002188619 | 0.378360619 |
| 2.836488 | -1.276317189 | 0.388454219 | -1.024122779 | 0.366264433 |
| 2.846728 | -1.279968401 | 0.383823915 | -1.042318293 | 0.358363193 |
| 2.856968 | -1.276306411 | 0.392993274 | -1.054679607 | 0.352602655 |
| 2.867208 | -1.266748577 | 0.413990629 | -1.062141495 | 0.35172819 |
| 2.877448 | -1.24999855 | 0.447240659 | -1.063288442 | 0.354045621 |
| 2.887688 | -1.228269468 | 0.489082758 | -1.058557212 | 0.361787889 |
| 2.897928 | -1.200348983 | 0.538786527 | -1.04702528 | 0.373614594 |
| 2.908168 | -1.168536017 | 0.592296511 | -1.030347228 | 0.390558237 |
| 2.918408 | -1.131514644 | 0.649749157 | -1.006296856 | 0.411634468 |
| 2.928648 | -1.092217465 | 0.706908731 | -0.977555522 | 0.435678662 |
| 2.938889 | -1.049461529 | 0.764174382 | -0.942366868 | 0.462570906 |
| 2.949129 | -1.005122721 | 0.818555117 | -0.903080076 | 0.490185801 |
| 2.959368 | -0.959171853 | 0.869272745 | -0.858347139 | 0.518050733 |
| 2.969609 | -0.91244594 | 0.91468758 | -0.810441532 | 0.544887004 |
| 2.979849 | -0.86535532 | 0.95404415 | -0.759889867 | 0.56980745 |
| 2.990089 | -0.818107867 | 0.986046912 | -0.707596737 | 0.59177244 |
| 3.000329 | -0.770447364 | 1.011540282 | -0.65468424 | 0.611010672 |
| 3.010569 | -0.723990458 | 1.029367957 | -0.602375109 | 0.625647179 |
| 3.020809 | -0.677435832 | 1.040974976 | -0.551408205 | 0.635954839 |
| 3.031049 | -0.63174591 | 1.046215535 | -0.501798715 | 0.641737563 |
| 3.041289 | -0.587168713 | 1.046745374 | -0.454603242 | 0.641064609 |
| 3.051529 | -0.543732574 | 1.042590568 | -0.409046091 | 0.635903625 |
| 3.061769 | -0.502201997 | 1.034246542 | -0.366221598 | 0.623966135 |
| 3.072009 | -0.462575669 | 1.022122347 | -0.325023032 | 0.606900788 |
| 3.082249 | -0.425709074 | 1.006410902 | -0.286731525 | 0.583966251 |
| 3.092489 | -0.391403233 | 0.986078469 | -0.250624664 | 0.556565507 |
| 3.102729 | -0.360534211 | 0.96181579 | -0.217534793 | 0.524944886 |
| 3.112969 | -0.332721248 | 0.933752782 | -0.186572653 | 0.489560539 |
| 3.123209 | -0.308303175 | 0.901513118 | -0.159323403 | 0.45232598 |
| 3.133449 | -0.287189636 | 0.867315058 | -0.134286428 | 0.413314266 |
| 3.143689 | -0.269633661 | 0.831649357 | -0.112997728 | 0.374169502 |
| 3.153929 | -0.254801391 | 0.79731337 | -0.093673769 | 0.33634087 |
| 3.164169 | -0.243906206 | 0.767191714 | -0.078285361 | 0.301069901 |
| 3.174409 | -0.235994143 | 0.742872567 | -0.064658163 | 0.271454258 |
| 3.184649 | -0.231489286 | 0.728530562 | -0.054794702 | 0.248913138 |
| 3.194889 | -0.229941919 | 0.723344725 | -0.046848791 | 0.236893757 |
| 3.205129 | -0.232245566 | 0.730908059 | -0.042221142 | 0.236806348 |
| 3.215369 | -0.236439755 | 0.746435419 | -0.039861026 | 0.24920734 |
| 3.225609 | -0.243961701 | 0.772760867 | -0.04144951 | 0.272390539 |
| 3.235849 | -0.253356928 | 0.803724456 | -0.045008373 | 0.303458573 |
| 3.246089 | -0.265134028 | 0.840329378 | -0.053245292 | 0.340225908 |
| 3.256329 | -0.278712746 | 0.878168491 | -0.063617972 | 0.380702477 |
| 3.266569 | -0.294342356 | 0.917679146 | -0.078291862 | 0.423001914 |
| 3.276809 | -0.311730258 | 0.955012769 | -0.094601806 | 0.466057494 |
| 3.28705 | -0.331980457 | 0.990857954 | -0.114325315 | 0.509690598 |
| 3.29729 | -0.35380767 | 1.021924525 | -0.134549093 | 0.552256079 |
| 3.307529 | -0.378583133 | 1.049769256 | -0.156587682 | 0.59472523 |
| 3.31777 | -0.405601457 | 1.071559329 | -0.178268078 | 0.635752963 |
| 3.32801 | -0.435633337 | 1.089320749 | -0.200734998 | 0.676766292 |
| 3.33825 | -0.468195831 | 1.101384716 | -0.222466098 | 0.71609084 |
| 3.34849 | -0.503588582 | 1.109683661 | -0.243423208 | 0.755712776 |
| 3.35873 | -0.542128007 | 1.11410142 | -0.265130832 | 0.7943685 |
| 3.36897 | -0.583128919 | 1.115055277 | -0.284976578 | 0.833419536 |
| 3.37921 | -0.62675408 | 1.114374939 | -0.306494383 | 0.871419628 |
| 3.38945 | -0.672717954 | 1.110068728 | -0.327006029 | 0.909160855 |
| 3.39969 | -0.720305872 | 1.10542277 | -0.349889253 | 0.945705898 |
| 3.40993 | -0.768546554 | 1.098689339 | -0.373194535 | 0.980881613 |
| 3.42017 | -0.817995516 | 1.091851403 | -0.400048277 | 1.013910959 |
| 3.43041 | -0.866475427 | 1.084051189 | -0.428591405 | 1.044715929 |
| 3.44065 | -0.915291165 | 1.077032717 | -0.461255489 | 1.072761027 |
| 3.45089 | -0.963058346 | 1.069998736 | -0.496326266 | 1.098490072 |
| 3.46113 | -1.010490062 | 1.064605227 | -0.535028429 | 1.121457434 |
| 3.47137 | -1.05738671 | 1.059195352 | -0.575779185 | 1.142270354 |
| 3.48161 | -1.103850547 | 1.05542192 | -0.619284845 | 1.160653874 |
| 3.49185 | -1.150513457 | 1.052645158 | -0.66366612 | 1.176653606 |
| 3.50209 | -1.196527728 | 1.049234534 | -0.709855548 | 1.189320867 |
| 3.51233 | -1.242470477 | 1.046469894 | -0.755844408 | 1.199246341 |
| 3.52257 | -1.287422496 | 1.042093008 | -0.802730931 | 1.204949948 |
| 3.53281 | -1.332162363 | 1.035812875 | -0.848884502 | 1.206385602 |
| 3.54305 | -1.374989213 | 1.026499553 | -0.895055788 | 1.202955971 |
| 3.55329 | -1.416847789 | 1.013383304 | -0.939947759 | 1.194369941 |
| 3.56353 | -1.456636565 | 0.995013769 | -0.983934388 | 1.18027462 |
| 3.57377 | -1.494353679 | 0.972482123 | -1.026325739 | 1.161249383 |
| 3.58401 | -1.529349096 | 0.943574482 | -1.066858517 | 1.136895032 |
| 3.59425 | -1.561496392 | 0.909745608 | -1.104766126 | 1.107596757 |
| 3.60449 | -1.5899711 | 0.87106444 | -1.140275595 | 1.073889063 |
| 3.61473 | -1.614827012 | 0.827572814 | -1.172900832 | 1.035978933 |
| 3.62497 | -1.635771692 | 0.780116383 | -1.202461862 | 0.994630754 |
| 3.635211 | -1.653035829 | 0.729529608 | -1.229303333 | 0.950054483 |
| 3.645451 | -1.666865438 | 0.675846938 | -1.253205309 | 0.902409914 |
| 3.65569 | -1.677612533 | 0.621692236 | -1.275719474 | 0.853088149 |
| 3.665931 | -1.686072568 | 0.566011941 | -1.295524806 | 0.801870106 |
| 3.676171 | -1.692954123 | 0.512445993 | -1.314708229 | 0.749978146 |
| 3.686411 | -1.697688818 | 0.462034378 | -1.331769313 | 0.698023133 |
| 3.696651 | -1.701679806 | 0.418924646 | -1.348204613 | 0.647390381 |
| 3.706891 | -1.703150491 | 0.385789071 | -1.361634518 | 0.598830021 |
| 3.717131 | -1.703746974 | 0.367903061 | -1.373440019 | 0.554334194 |
| 3.727371 | -1.700893541 | 0.367206818 | -1.380910311 | 0.514510987 |
| 3.737611 | -1.696454823 | 0.38544828 | -1.384948058 | 0.481590153 |
| 3.747851 | -1.688166181 | 0.419239591 | -1.383715468 | 0.456006317 |
| 3.758091 | -1.677509189 | 0.466118799 | -1.377654476 | 0.439037171 |
| 3.768331 | -1.663588663 | 0.522494679 | -1.366195406 | 0.430833178 |
| 3.778571 | -1.647327622 | 0.585349323 | -1.349779112 | 0.431315284 |
| 3.788811 | -1.629290402 | 0.653424589 | -1.328486017 | 0.440003865 |
| 3.799051 | -1.608914167 | 0.724630843 | -1.302568785 | 0.455033238 |
| 3.809291 | -1.587774356 | 0.797571856 | -1.272958125 | 0.475911467 |
| 3.819531 | -1.564611807 | 0.872067252 | -1.239192733 | 0.500879475 |
| 3.829771 | -1.540347651 | 0.945574877 | -1.202065442 | 0.528932425 |
| 3.840011 | -1.513493118 | 1.018509823 | -1.161242762 | 0.558576747 |
| 3.850251 | -1.485080485 | 1.088507576 | -1.117422696 | 0.590009056 |
| 3.860491 | -1.45253378 | 1.155482265 | -1.070192916 | 0.621462725 |
| 3.870731 | -1.418373086 | 1.217951275 | -1.019962496 | 0.653461208 |
| 3.880971 | -1.379149934 | 1.274897583 | -0.967163911 | 0.684400787 |
| 3.891211 | -1.338550155 | 1.326256785 | -0.911354021 | 0.715107308 |
| 3.901451 | -1.293985049 | 1.371779563 | -0.853664819 | 0.744010233 |
| 3.911691 | -1.24880452 | 1.410641613 | -0.793879732 | 0.772248976 |
| 3.921931 | -1.200683194 | 1.44426607 | -0.732719353 | 0.797331695 |
| 3.932171 | -1.153624178 | 1.472137235 | -0.670200187 | 0.820777236 |
| 3.942411 | -1.105035497 | 1.495109199 | -0.60724688 | 0.840180962 |
| 3.952652 | -1.05792168 | 1.513499878 | -0.54321833 | 0.8564039 |
| 3.962891 | -1.010487569 | 1.527244039 | -0.479166085 | 0.867617637 |
| 3.973131 | -0.965012296 | 1.537122676 | -0.415263316 | 0.874750362 |
| 3.983371 | -0.919635745 | 1.542956247 | -0.351079326 | 0.876998182 |
| 3.993612 | -0.875703711 | 1.544332375 | -0.288253278 | 0.875121227 |
| 4.003851 | -0.832876946 | 1.542419667 | -0.225960038 | 0.869575806 |
| 4.014091 | -0.7905289 | 1.536814748 | -0.165758737 | 0.86092778 |
| 4.024332 | -0.748797155 | 1.527382416 | -0.107822401 | 0.850671144 |
| 4.034572 | -0.707400794 | 1.515254055 | -0.052987184 | 0.838538279 |
| 4.044812 | -0.665780798 | 1.499632973 | -0.001482365 | 0.826112858 |
| 4.055052 | -0.623177998 | 1.480778937 | 0.045172236 | 0.812690066 |
| 4.065292 | -0.580347367 | 1.45887217 | 0.088431693 | 0.799310583 |
| 4.075532 | -0.535212199 | 1.433290472 | 0.126115258 | 0.784741298 |
| 4.085772 | -0.489356128 | 1.404397611 | 0.159901299 | 0.769480933 |
| 4.096012 | -0.442205302 | 1.371880362 | 0.188559911 | 0.751971121 |
| 4.106252 | -0.394356921 | 1.335920834 | 0.213797377 | 0.732634459 |
| 4.116492 | -0.34685759 | 1.297336516 | 0.23441925 | 0.710341559 |
| 4.126732 | -0.300590287 | 1.256772263 | 0.252031168 | 0.685150257 |
| 4.136972 | -0.256394713 | 1.215164253 | 0.266824576 | 0.65675149 |
| 4.147212 | -0.215777539 | 1.173301775 | 0.278676394 | 0.625499765 |
| 4.157452 | -0.179650378 | 1.132563535 | 0.288587721 | 0.592145609 |
| 4.167692 | -0.147710276 | 1.093292141 | 0.295558549 | 0.557596959 |
| 4.177932 | -0.12101403 | 1.056259408 | 0.300693244 | 0.523253455 |
| 4.188172 | -0.098952728 | 1.021344416 | 0.302998314 | 0.490258898 |
| 4.198412 | -0.081550998 | 0.988131714 | 0.302266435 | 0.460585273 |
| 4.208652 | -0.067502461 | 0.956650367 | 0.298622225 | 0.435266659 |
| 4.218892 | -0.057416979 | 0.926278128 | 0.291713514 | 0.416151637 |
| 4.229132 | -0.049175639 | 0.896050235 | 0.281912113 | 0.402996023 |
| 4.239372 | -0.043572536 | 0.866474054 | 0.269249727 | 0.396955105 |
| 4.249612 | -0.038974306 | 0.836083648 | 0.254063229 | 0.395930568 |
| 4.259852 | -0.034916829 | 0.805909952 | 0.237769933 | 0.400496406 |
| 4.270092 | -0.031473753 | 0.775326561 | 0.220102807 | 0.408155449 |
| 4.280333 | -0.0276454 | 0.74453894 | 0.202189206 | 0.418493371 |
| 4.290572 | -0.023899848 | 0.713533027 | 0.184587944 | 0.42978909 |
| 4.300812 | -0.019308098 | 0.683008465 | 0.167645586 | 0.44257292 |
| 4.311052 | -0.01525183 | 0.652823536 | 0.151679627 | 0.454385894 |
| 4.321292 | -0.010688478 | 0.62342695 | 0.136939304 | 0.466418212 |
| 4.331532 | -0.006382441 | 0.595543575 | 0.122765043 | 0.476970616 |
| 4.341773 | -0.002079914 | 0.569233556 | 0.109669541 | 0.48636636 |
| 4.352013 | 0.002265397 | 0.545486337 | 0.096829163 | 0.493779865 |
| 4.362253 | 0.007518868 | 0.523458102 | 0.083938782 | 0.499672065 |
| 4.372493 | 0.012717982 | 0.504549433 | 0.071017076 | 0.503077897 |
| 4.382733 | 0.019306513 | 0.486807877 | 0.057214839 | 0.504654522 |
| 4.392972 | 0.025784868 | 0.471792601 | 0.043113027 | 0.504040131 |
| 4.403213 | 0.033630267 | 0.457484024 | 0.028419871 | 0.501818957 |
| 4.413453 | 0.041192542 | 0.444527997 | 0.013100722 | 0.497900911 |
| 4.423693 | 0.049806034 | 0.431995378 | -0.002179933 | 0.492694552 |
| 4.433933 | 0.057955848 | 0.420331329 | -0.018049887 | 0.48667345 |
| 4.444173 | 0.067653883 | 0.409438886 | -0.033582526 | 0.479734826 |
| 4.454413 | 0.077336017 | 0.398426468 | -0.049492994 | 0.472641885 |
| 4.464653 | 0.088595832 | 0.387920085 | -0.064974661 | 0.465262909 |
| 4.474893 | 0.10068676 | 0.376956217 | -0.080558917 | 0.457979816 |
| 4.485133 | 0.113525812 | 0.365691251 | -0.095392719 | 0.451001252 |
| 4.495373 | 0.126824006 | 0.353088472 | -0.109970482 | 0.444568675 |
| 4.505613 | 0.140200676 | 0.339776048 | -0.123204491 | 0.438473305 |
| 4.515853 | 0.152050423 | 0.324541479 | -0.135447983 | 0.43296879 |
| 4.526093 | 0.163376331 | 0.309197247 | -0.145484707 | 0.427548168 |
| 4.536333 | 0.17176476 | 0.291630058 | -0.154003482 | 0.422437213 |
| 4.546573 | 0.178786485 | 0.27450397 | -0.159795061 | 0.416938212 |
| 4.556813 | 0.182820782 | 0.256071806 | -0.163703399 | 0.411278069 |
| 4.567053 | 0.185179579 | 0.238519477 | -0.165545536 | 0.404776226 |
| 4.577293 | 0.184923465 | 0.220719626 | -0.165569129 | 0.397973135 |
| 4.587533 | 0.183762953 | 0.204482395 | -0.164563985 | 0.390275924 |
| 4.597773 | 0.181072963 | 0.189899838 | -0.162287593 | 0.381995241 |
| 4.608013 | 0.178165369 | 0.177783626 | -0.159942697 | 0.37330422 |
| 4.618253 | 0.175478466 | 0.167965338 | -0.156435362 | 0.364459159 |
| 4.628493 | 0.172302148 | 0.161841749 | -0.153365107 | 0.355220757 |
| 4.638733 | 0.170709537 | 0.157667655 | -0.148720174 | 0.34664358 |
| 4.648973 | 0.169206466 | 0.155785016 | -0.144049957 | 0.338450758 |
| 4.659214 | 0.168741159 | 0.156974111 | -0.137180698 | 0.330974863 |
| 4.669454 | 0.168728169 | 0.161375366 | -0.129541369 | 0.324766872 |
| 4.679694 | 0.169657726 | 0.168587649 | -0.119187813 | 0.318963267 |
| 4.689933 | 0.170805343 | 0.179891831 | -0.106818359 | 0.314568164 |
| 4.700173 | 0.172905213 | 0.194946403 | -0.091952511 | 0.31043382 |
| 4.710413 | 0.174619589 | 0.212619832 | -0.074824691 | 0.30754667 |
| 4.720654 | 0.177974713 | 0.232254098 | -0.055151145 | 0.304643363 |
| 4.730894 | 0.180254932 | 0.253365419 | -0.03413627 | 0.303140282 |
| 4.741134 | 0.184551424 | 0.273636423 | -0.011623317 | 0.301491206 |
| 4.751374 | 0.188217945 | 0.293753069 | 0.011268997 | 0.300816937 |

| | | | | |
|---|---|---|---|---|
| 2.355207 | 0.343905523 | 0.198339239 | 0.024236043 | 0.16828366 |
| 2.365447 | 0.352613636 | 0.198427051 | 0.038187478 | 0.169078051 |
| 2.375687 | 0.361920602 | 0.197691391 | 0.052419559 | 0.170755075 |
| 2.385927 | 0.370766707 | 0.196911907 | 0.067316949 | 0.172896294 |
| 2.396167 | 0.380172423 | 0.195363986 | 0.082302791 | 0.174721518 |
| 2.406407 | 0.389598001 | 0.194114178 | 0.097670358 | 0.177218221 |
| 2.416647 | 0.399119017 | 0.192461168 | 0.113304306 | 0.179457269 |
| 2.426887 | 0.408510676 | 0.19163072 | 0.129034571 | 0.182293873 |
| 2.437127 | 0.418333008 | 0.191083873 | 0.14531612 | 0.185155248 |
| 2.447367 | 0.427770878 | 0.191492339 | 0.16148778 | 0.188558225 |
| 2.457607 | 0.437648848 | 0.191863586 | 0.178224449 | 0.192649437 |
| 2.467847 | 0.447325372 | 0.193325322 | 0.194567563 | 0.196525176 |
| 2.478087 | 0.45683136 | 0.194844177 | 0.211373788 | 0.200754241 |
| 2.488327 | 0.466466983 | 0.19629969 | 0.227835737 | 0.204905729 |
| 2.498567 | 0.475911213 | 0.197882068 | 0.244006884 | 0.20876886 |
| 2.508807 | 0.484723102 | 0.199256399 | 0.259982529 | 0.212190857 |
| 2.519047 | 0.493713066 | 0.200379055 | 0.27518844 | 0.215578687 |
| 2.529287 | 0.501732531 | 0.200967853 | 0.289660724 | 0.218799 |
| 2.539527 | 0.509361379 | 0.201691385 | 0.303142395 | 0.221664156 |
| 2.549767 | 0.515905395 | 0.201686566 | 0.315575707 | 0.224815896 |
| 2.560007 | 0.521393891 | 0.201792443 | 0.326770527 | 0.227750275 |
| 2.570247 | 0.525401257 | 0.201733542 | 0.336464361 | 0.230659664 |
| 2.580487 | 0.528216256 | 0.201650023 | 0.344716999 | 0.233185223 |
| 2.590728 | 0.529115967 | 0.201699597 | 0.351666502 | 0.235493628 |
| 2.600967 | 0.528771406 | 0.202032975 | 0.356994178 | 0.237708035 |
| 2.611207 | 0.526263285 | 0.202493602 | 0.360813002 | 0.239489675 |
| 2.621448 | 0.522534105 | 0.203936761 | 0.363564878 | 0.241730105 |
| 2.631688 | 0.516714293 | 0.205116175 | 0.364616413 | 0.243953536 |
| 2.641928 | 0.509375092 | 0.207109541 | 0.364824962 | 0.247125554 |
| 2.652168 | 0.500099994 | 0.208552761 | 0.363798542 | 0.25016376 |
| 2.662408 | 0.488908578 | 0.209963476 | 0.361952978 | 0.254340237 |
| 2.672648 | 0.476738023 | 0.210059312 | 0.359223117 | 0.258276463 |
| 2.682888 | 0.462408537 | 0.20936544 | 0.355848902 | 0.26275133 |
| 2.693128 | 0.44743864 | 0.206965927 | 0.351654224 | 0.266504186 |
| 2.703368 | 0.431398257 | 0.203698917 | 0.346712286 | 0.270523549 |
| 2.713608 | 0.414807451 | 0.199582912 | 0.340638239 | 0.273658447 |
| 2.723848 | 0.397233164 | 0.194866835 | 0.333410182 | 0.277064235 |
| 2.734088 | 0.379007776 | 0.19039832 | 0.325116742 | 0.279727651 |
| 2.744328 | 0.359822025 | 0.186807567 | 0.315443657 | 0.282507209 |
| 2.754568 | 0.339266298 | 0.184324386 | 0.304463999 | 0.284525498 |
| 2.764808 | 0.317226449 | 0.18304234 | 0.292249983 | 0.286522322 |
| 2.775048 | 0.293214608 | 0.183952576 | 0.279083146 | 0.287642574 |
| 2.785288 | 0.26749937 | 0.186126132 | 0.265069512 | 0.287779221 |
| 2.795528 | 0.240036417 | 0.190215471 | 0.249873458 | 0.28752515 |
| 2.805768 | 0.210972036 | 0.195529964 | 0.23376087 | 0.286107938 |
| 2.816008 | 0.181159186 | 0.202475585 | 0.216485037 | 0.284206062 |
| 2.826248 | 0.150867419 | 0.210253268 | 0.197938176 | 0.282028266 |
| 2.836488 | 0.120645227 | 0.218777593 | 0.177776246 | 0.279448777 |
| 2.846728 | 0.091116943 | 0.228151011 | 0.156770829 | 0.277641722 |
| 2.856968 | 0.062463453 | 0.237745599 | 0.133995637 | 0.276188041 |
| 2.867208 | 0.035125033 | 0.24792731 | 0.11056675 | 0.275648594 |
| 2.877448 | 0.008755438 | 0.258144631 | 0.086557196 | 0.276412277 |
| 2.887688 | -0.016896292 | 0.268342117 | 0.061858359 | 0.277912804 |
| 2.897928 | -0.041213291 | 0.278892122 | 0.036933492 | 0.281144247 |
| 2.908168 | -0.065759286 | 0.288760082 | 0.011781657 | 0.284489169 |
| 2.918408 | -0.089351846 | 0.298373013 | -0.01391858 | 0.288681662 |
| 2.928648 | -0.113140603 | 0.306633044 | -0.04003764 | 0.292642197 |
| 2.938889 | -0.136537449 | 0.313994258 | -0.067054718 | 0.296325911 |
| 2.949129 | -0.159737218 | 0.319402699 | -0.09452356 | 0.299464823 |
| 2.959368 | -0.182471075 | 0.322968789 | -0.122780714 | 0.301584669 |
| 2.969609 | -0.205089777 | 0.32472766 | -0.151431094 | 0.303394883 |
| 2.979849 | -0.226841941 | 0.324799336 | -0.179898661 | 0.304309019 |
| 2.990089 | -0.248021353 | 0.322953009 | -0.208258027 | 0.305203326 |
| 3.000329 | -0.268165577 | 0.320000038 | -0.23541451 | 0.305111183 |
| 3.010569 | -0.287143932 | 0.315294987 | -0.2614584 | 0.305923332 |
| 3.020809 | -0.305012991 | 0.310652135 | -0.286073344 | 0.305538407 |
| 3.031049 | -0.32095071 | 0.304433928 | -0.308812226 | 0.305886652 |
| 3.041289 | -0.335617457 | 0.298380559 | -0.329501824 | 0.3059749 |
| 3.051529 | -0.348278107 | 0.292524056 | -0.348703945 | 0.306488842 |
| 3.061769 | -0.359870034 | 0.286539591 | -0.365418196 | 0.306827882 |
| 3.072009 | -0.370003084 | 0.28105344 | -0.380739975 | 0.307981527 |
| 3.082249 | -0.379481244 | 0.27590934 | -0.393619563 | 0.308600609 |
| 3.092489 | -0.387922399 | 0.271022138 | -0.404764647 | 0.309958495 |
| 3.102729 | -0.396168713 | 0.26626878 | -0.41354445 | 0.310822391 |
| 3.112969 | -0.403606266 | 0.261313611 | -0.420416193 | 0.311417551 |
| 3.123209 | -0.410473119 | 0.256223489 | -0.424886741 | 0.311777012 |
| 3.133449 | -0.416197404 | 0.251589817 | -0.427254441 | 0.311482911 |
| 3.143689 | -0.420449091 | 0.24747188 | -0.427934088 | 0.31069457 |
| 3.153929 | -0.422820549 | 0.244254091 | -0.426605097 | 0.309461252 |
| 3.164169 | -0.423120787 | 0.242312972 | -0.424118398 | 0.307592923 |
| 3.174409 | -0.420768121 | 0.242119311 | -0.420348085 | 0.305771231 |
| 3.184649 | -0.415736505 | 0.243251075 | -0.415948291 | 0.303926665 |
| 3.194889 | -0.408071592 | 0.245575387 | -0.410494288 | 0.302691067 |
| 3.205129 | -0.397197792 | 0.248351845 | -0.40465836 | 0.30154203 |
| 3.215369 | -0.383595347 | 0.25176603 | -0.397843688 | 0.301140097 |
| 3.225609 | -0.367371226 | 0.255347237 | -0.39018527 | 0.301413084 |
| 3.235849 | -0.348465285 | 0.25866674 | -0.381289465 | 0.301521414 |
| 3.246089 | -0.327925148 | 0.261479671 | -0.371244939 | 0.30207936 |
| 3.256329 | -0.305385054 | 0.264740004 | -0.35947906 | 0.302576726 |
| 3.266569 | -0.282523478 | 0.266943735 | -0.346287954 | 0.303200596 |
| 3.276809 | -0.259228582 | 0.268838048 | -0.331692887 | 0.303344875 |
| 3.28705 | -0.235962536 | 0.270007321 | -0.315808625 | 0.303761559 |
| 3.29729 | -0.213634307 | 0.270363274 | -0.299029967 | 0.303775667 |
| 3.307529 | -0.191860281 | 0.26955158 | -0.281282087 | 0.303295805 |
| 3.31777 | -0.171043631 | 0.267439575 | -0.263306199 | 0.302366464 |
| 3.32801 | -0.150967081 | 0.264597915 | -0.245073279 | 0.300525121 |
| 3.33825 | -0.131560797 | 0.260098685 | -0.226599852 | 0.297735502 |
| 3.34849 | -0.112826386 | 0.254976546 | -0.208076059 | 0.293919542 |
| 3.35873 | -0.094812263 | 0.248727565 | -0.189381016 | 0.289054622 |
| 3.36897 | -0.077408534 | 0.24167793 | -0.17036184 | 0.283528455 |
| 3.37921 | -0.061221745 | 0.233551154 | -0.151144628 | 0.277322748 |
| 3.38945 | -0.045746937 | 0.224899016 | -0.131292511 | 0.271302354 |
| 3.39969 | -0.032137697 | 0.215321848 | -0.111003288 | 0.265329403 |
| 3.40993 | -0.019716587 | 0.204972875 | -0.090710454 | 0.25990918 |
| 3.42017 | -0.009069671 | 0.19488838 | -0.069782598 | 0.255154729 |
| 3.43041 | -8.33534E-05 | 0.184616747 | -0.049751751 | 0.251310423 |
| 3.44065 | 0.007309929 | 0.174645902 | -0.029965062 | 0.24817577 |
| 3.45089 | 0.013000596 | 0.16565483 | -0.011935365 | 0.245068004 |
| 3.46113 | 0.017456171 | 0.156936271 | 0.004818255 | 0.242733867 |
| 3.47137 | 0.020064908 | 0.14947765 | 0.019212236 | 0.239694851 |
| 3.48161 | 0.021642324 | 0.142387357 | 0.030750551 | 0.236269161 |
| 3.49185 | 0.021292561 | 0.137428764 | 0.039509265 | 0.231445675 |
| 3.50209 | 0.019810579 | 0.133608185 | 0.045950245 | 0.226093246 |
| 3.51233 | 0.016663134 | 0.133266471 | 0.049068294 | 0.218862176 |
| 3.52257 | 0.012388469 | 0.136301832 | 0.050129966 | 0.21163222 |
| 3.53281 | 0.006584252 | 0.143921135 | 0.04877218 | 0.203225771 |
| 3.54305 | 0.000357925 | 0.155684504 | 0.046188764 | 0.195385729 |
| 3.55329 | -0.006540986 | 0.171431992 | 0.041811126 | 0.187925637 |
| 3.56353 | -0.013956835 | 0.189657677 | 0.037026843 | 0.181720838 |
| 3.57377 | -0.0213733 | 0.209974104 | 0.031301012 | 0.176483725 |
| 3.58401 | -0.028707021 | 0.230766646 | 0.024960614 | 0.172708708 |
| 3.59425 | -0.036426859 | 0.251360476 | 0.018510202 | 0.170522627 |
| 3.60449 | -0.044582633 | 0.271042063 | 0.011690604 | 0.170307729 |
| 3.61473 | -0.053207579 | 0.289159717 | 0.004280947 | 0.171843404 |
| 3.62497 | -0.063158005 | 0.305970204 | -0.003124278 | 0.175607226 |
| 3.635211 | -0.073952051 | 0.320767618 | -0.011217062 | 0.181909069 |
| 3.645451 | -0.086214327 | 0.333856244 | -0.019650226 | 0.189774433 |
| 3.65569 | -0.100115881 | 0.34593929 | -0.028106667 | 0.199546086 |
| 3.665931 | -0.115061093 | 0.356205982 | -0.037529027 | 0.210217521 |
| 3.676171 | -0.131735586 | 0.36600183 | -0.047227583 | 0.221734858 |
| 3.686411 | -0.149279216 | 0.374305108 | -0.057529228 | 0.232555615 |
| 3.696651 | -0.167950486 | 0.382304745 | -0.06884429 | 0.243716582 |
| 3.706891 | -0.187726942 | 0.389289866 | -0.080785643 | 0.253510989 |
| 3.717131 | -0.208106232 | 0.395987546 | -0.094499535 | 0.262506361 |
| 3.727371 | -0.228868549 | 0.402169583 | -0.10931914 | 0.270166773 |
| 3.737611 | -0.250253442 | 0.40750424 | -0.12561801 | 0.276910144 |
| 3.747851 | -0.271660858 | 0.412592199 | -0.143857624 | 0.282880781 |
| 3.758091 | -0.293169692 | 0.416859641 | -0.163412421 | 0.288288426 |
| 3.768331 | -0.314459009 | 0.419519159 | -0.184387031 | 0.292877262 |
| 3.778571 | -0.335850354 | 0.421740722 | -0.206292822 | 0.297300089 |
| 3.788811 | -0.356897564 | 0.421795263 | -0.229139006 | 0.301940219 |
| 3.799051 | -0.377881001 | 0.421331771 | -0.251776042 | 0.305951079 |
| 3.809291 | -0.399059768 | 0.419097042 | -0.27450387 | 0.309925046 |
| 3.819531 | -0.420254801 | 0.416787749 | -0.296780675 | 0.313796161 |
| 3.829771 | -0.441326532 | 0.413564415 | -0.318495633 | 0.317247003 |
| 3.840011 | -0.462548489 | 0.411124308 | -0.339006374 | 0.32081613 |
| 3.850251 | -0.48355302 | 0.408368673 | -0.359279679 | 0.323298834 |
| 3.860491 | -0.503764764 | 0.406742786 | -0.378318947 | 0.325772303 |
| 3.870731 | -0.523149353 | 0.404963932 | -0.396806922 | 0.327667383 |
| 3.880971 | -0.541596403 | 0.403668563 | -0.414301655 | 0.328653905 |
| 3.891211 | -0.558034957 | 0.40204272 | -0.430856517 | 0.328936314 |
| 3.901451 | -0.573231261 | 0.399597571 | -0.446381444 | 0.328496053 |
| 3.911691 | -0.586870698 | 0.396290464 | -0.460447376 | 0.327523887 |
| 3.921931 | -0.598786741 | 0.392409447 | -0.473362872 | 0.325409656 |
| 3.932171 | -0.609203803 | 0.386912086 | -0.484505701 | 0.322591759 |
| 3.942411 | -0.618700472 | 0.380487158 | -0.493946854 | 0.318738161 |
| 3.952652 | -0.626757691 | 0.373238516 | -0.501806953 | 0.314217 |
| 3.962891 | -0.633760326 | 0.364806511 | -0.507990727 | 0.308405881 |
| 3.973131 | -0.639578432 | 0.355555918 | -0.512387641 | 0.301773591 |
| 3.983371 | -0.644566077 | 0.34471977 | -0.515576641 | 0.294429374 |
| 3.993612 | -0.648222068 | 0.33322917 | -0.516831797 | 0.286813453 |
| 4.003851 | -0.650729114 | 0.320561643 | -0.517346591 | 0.278572902 |
| 4.014091 | -0.6521345 | 0.306508874 | -0.516235404 | 0.270744165 |
| 4.024332 | -0.652360305 | 0.292516452 | -0.514088707 | 0.263203933 |
| 4.034572 | -0.651153518 | 0.277819087 | -0.511361774 | 0.256109738 |
| 4.044812 | -0.649201418 | 0.264284867 | -0.507600773 | 0.249728384 |
| 4.055052 | -0.64585292 | 0.251464304 | -0.503944025 | 0.243998009 |
| 4.065292 | -0.641930771 | 0.240575894 | -0.500132444 | 0.238961957 |
| 4.075532 | -0.636779308 | 0.231892819 | -0.49629 | 0.23491501 |
| 4.085772 | -0.631325746 | 0.225558707 | -0.493303329 | 0.231308304 |
| 4.096012 | -0.624903902 | 0.222100037 | -0.490172784 | 0.228330281 |
| 4.106252 | -0.617651147 | 0.220900626 | -0.487669943 | 0.226063244 |
| 4.116492 | -0.609278235 | 0.222035698 | -0.484839583 | 0.224110877 |
| 4.126732 | -0.599326345 | 0.225481589 | -0.481900617 | 0.222156376 |
| 4.136972 | -0.587664744 | 0.230251897 | -0.478225185 | 0.220713243 |
| 4.147212 | -0.573919551 | 0.23646487 | -0.473497658 | 0.218943823 |
| 4.157452 | -0.557933353 | 0.243803276 | -0.467606417 | 0.217906386 |
| 4.167692 | -0.539427957 | 0.251170872 | -0.46021646 | 0.216731582 |
| 4.177932 | -0.519570307 | 0.258554256 | -0.45136563 | 0.216471789 |
| 4.188172 | -0.497352552 | 0.265839429 | -0.440949582 | 0.216995698 |
| 4.198412 | -0.474569206 | 0.271481871 | -0.428910832 | 0.219303267 |
| 4.208652 | -0.450809877 | 0.276793547 | -0.415288635 | 0.223332169 |
| 4.218892 | -0.427352306 | 0.280033489 | -0.400137976 | 0.22954635 |
| 4.229132 | -0.404399752 | 0.283258428 | -0.383831276 | 0.238322925 |
| 4.239372 | -0.382332433 | 0.285056358 | -0.36617068 | 0.248963005 |
| 4.249612 | -0.361394303 | 0.286483596 | -0.347854253 | 0.261345267 |
| 4.259852 | -0.342086393 | 0.287800169 | -0.328793176 | 0.274422754 |
| 4.270092 | -0.323937979 | 0.28885371 | -0.30924446 | 0.287644983 |
| 4.280333 | -0.307095449 | 0.289688365 | -0.289438199 | 0.300368676 |
| 4.290572 | -0.291076133 | 0.29058869 | -0.26905707 | 0.31209352 |
| 4.300812 | -0.27537266 | 0.29065377 | -0.248688531 | 0.32254554 |
| 4.311052 | -0.260382665 | 0.291021895 | -0.227402344 | 0.331577541 |
| 4.321292 | -0.244294719 | 0.290318735 | -0.20582239 | 0.339192384 |
| 4.331532 | -0.228662326 | 0.289450008 | -0.183984461 | 0.345492883 |
| 4.341773 | -0.211722149 | 0.287971724 | -0.161688049 | 0.35010815 |
| 4.352013 | -0.194506435 | 0.286220472 | -0.139304173 | 0.353235373 |
| 4.362253 | -0.17689674 | 0.284217715 | -0.117072187 | 0.355219385 |
| 4.372493 | -0.158991158 | 0.281714549 | -0.095467382 | 0.355339667 |
| 4.382733 | -0.141099588 | 0.279381392 | -0.074303316 | 0.354229334 |
| 4.392972 | -0.123665285 | 0.276632105 | -0.054454052 | 0.351868223 |
| 4.403213 | -0.10644775 | 0.273640083 | -0.035504496 | 0.348428486 |
| 4.413453 | -0.090309586 | 0.270411166 | -0.018205194 | 0.343823493 |
| 4.423693 | -0.074645945 | 0.267114269 | -0.002052046 | 0.338611158 |
| 4.433933 | -0.060209022 | 0.263038636 | 0.012265868 | 0.332760843 |
| 4.444173 | -0.046584691 | 0.259316662 | 0.025196345 | 0.326750504 |
| 4.454413 | -0.034383142 | 0.254588107 | 0.036871691 | 0.320510278 |
| 4.464653 | -0.023485112 | 0.249851681 | 0.047311996 | 0.314836698 |
| 4.474893 | -0.013607891 | 0.244680704 | 0.0563289 | 0.309914099 |
| 4.485133 | -0.005863917 | 0.239019807 | 0.06508484 | 0.306225066 |
| 4.495373 | 0.00106757 | 0.232501262 | 0.072375795 | 0.30373981 |
| 4.505613 | 0.006157203 | 0.225702615 | 0.079240014 | 0.303386476 |
| 4.515853 | 0.010110394 | 0.21860158 | 0.085306241 | 0.304260445 |
| 4.526093 | 0.012948006 | 0.211415649 | 0.090675435 | 0.306778281 |
| 4.536333 | 0.014921066 | 0.204612788 | 0.095312002 | 0.310123172 |
| 4.546573 | 0.015628235 | 0.198407582 | 0.099100543 | 0.313779655 |
| 4.556813 | 0.015873067 | 0.193274854 | 0.102496989 | 0.31756309 |
| 4.567053 | 0.015556783 | 0.188518997 | 0.104908071 | 0.320971959 |
| 4.577293 | 0.013837532 | 0.185000074 | 0.106614092 | 0.323427803 |
| 4.587533 | 0.011861447 | 0.18202748 | 0.107450578 | 0.325729678 |
| 4.597773 | 0.008739104 | 0.179289014 | 0.107163543 | 0.326800328 |
| 4.608013 | 0.004577795 | 0.176861451 | 0.10595932 | 0.327060259 |
| 4.618253 | -0.000210684 | 0.174847327 | 0.103520596 | 0.326929083 |
| 4.628493 | -0.006286036 | 0.173131465 | 0.100138701 | 0.32561497 |
| 4.638733 | -0.012248598 | 0.17165419 | 0.096140678 | 0.323899693 |
| 4.648973 | -0.019503376 | 0.17096457 | 0.091379141 | 0.321255838 |
| 4.659214 | -0.026427997 | 0.170714879 | 0.086399704 | 0.318128187 |
| 4.669454 | -0.033401716 | 0.171141922 | 0.08178635 | 0.314259396 |
| 4.679694 | -0.040510823 | 0.172431668 | 0.077068804 | 0.309786267 |
| 4.689933 | -0.047238172 | 0.173607536 | 0.07280832 | 0.304816133 |
| 4.700173 | -0.054164282 | 0.175228544 | 0.06901665 | 0.299290881 |
| 4.710413 | -0.060910467 | 0.176846784 | 0.065831194 | 0.293761581 |
| 4.720654 | -0.068205291 | 0.178421817 | 0.062788443 | 0.287922341 |
| 4.730894 | -0.075827003 | 0.180115859 | 0.060327172 | 0.282501479 |
| 4.741134 | -0.084140967 | 0.182062094 | 0.058015709 | 0.277440997 |
| 4.751374 | -0.092560472 | 0.184319913 | 0.056189201 | 0.272874641 |

| | | | | |
|---|---|---|---|---|
| 2.355207 | 1.60491569 | 0.365302574 | 1.209662961 | 0.471040248 |
| 2.365447 | 1.586523592 | 0.368668636 | 1.218660037 | 0.466746845 |
| 2.375687 | 1.562642872 | 0.373273636 | 1.22337311 | 0.461792783 |
| 2.385927 | 1.532683504 | 0.377422962 | 1.222878049 | 0.456823604 |
| 2.396167 | 1.497081554 | 0.38223163 | 1.217866878 | 0.451240528 |
| 2.406407 | 1.455503356 | 0.386803109 | 1.207416087 | 0.445421421 |
| 2.416647 | 1.408157241 | 0.391875929 | 1.192388833 | 0.439484227 |
| 2.426887 | 1.35611586 | 0.396567048 | 1.172314823 | 0.433311455 |
| 2.437127 | 1.298604703 | 0.402144723 | 1.14794154 | 0.427357806 |
| 2.447367 | 1.236849999 | 0.407962517 | 1.118928487 | 0.421265121 |
| 2.457607 | 1.17115083 | 0.414621912 | 1.086223883 | 0.415410834 |
| 2.467847 | 1.101764181 | 0.421962317 | 1.04950541 | 0.409487228 |
| 2.478087 | 1.0292171 | 0.430301689 | 1.009011383 | 0.403677932 |
| 2.488327 | 0.953517361 | 0.438416151 | 0.964958695 | 0.397722781 |
| 2.498567 | 0.875308073 | 0.447199451 | 0.917383071 | 0.390991722 |
| 2.508807 | 0.794437526 | 0.454686496 | 0.866335874 | 0.384163118 |
| 2.519047 | 0.711738661 | 0.461874787 | 0.811639675 | 0.376715552 |
| 2.529287 | 0.626412209 | 0.46720063 | 0.753559918 | 0.369497264 |
| 2.539527 | 0.540452591 | 0.471302324 | 0.692283109 | 0.36239857 |
| 2.549767 | 0.452261491 | 0.473948464 | 0.628398746 | 0.356816969 |
| 2.560007 | 0.363876358 | 0.475258686 | 0.561664957 | 0.352606754 |
| 2.570247 | 0.274211 | 0.475428189 | 0.493204903 | 0.351155111 |
| 2.580487 | 0.18511833 | 0.475137578 | 0.422857787 | 0.352000182 |
| 2.590728 | 0.095067319 | 0.474209554 | 0.351586622 | 0.355646664 |
| 2.600967 | 0.006341478 | 0.473147092 | 0.279254207 | 0.361733624 |
| 2.611207 | -0.082609329 | 0.472060685 | 0.206314461 | 0.369645453 |
| 2.621448 | -0.170345476 | 0.471057137 | 0.132816332 | 0.379256929 |
| 2.631688 | -0.2572815 | 0.46972857 | 0.059069714 | 0.389590347 |
| 2.641928 | -0.343153393 | 0.468793359 | -0.015271951 | 0.400557755 |
| 2.652168 | -0.427520227 | 0.467471044 | -0.089620794 | 0.412127098 |
| 2.662408 | -0.510227309 | 0.466382498 | -0.164709132 | 0.422793356 |
| 2.672648 | -0.590454333 | 0.464750131 | -0.239808248 | 0.434834982 |
| 2.682888 | -0.667891631 | 0.463961343 | -0.315225009 | 0.445148986 |
| 2.693128 | -0.741683191 | 0.462898315 | -0.390509144 | 0.456686602 |
| 2.703368 | -0.811374328 | 0.462614347 | -0.465488116 | 0.466730595 |
| 2.713608 | -0.876126148 | 0.462774625 | -0.539225577 | 0.478057407 |
| 2.723848 | -0.935935119 | 0.464222349 | -0.611835749 | 0.487186445 |
| 2.734088 | -0.990731418 | 0.465917695 | -0.681849829 | 0.497388786 |
| 2.744328 | -1.040085423 | 0.469590325 | -0.74922194 | 0.505635521 |
| 2.754568 | -1.084366393 | 0.473305727 | -0.813324233 | 0.514376762 |
| 2.764808 | -1.123538786 | 0.479073386 | -0.874168667 | 0.520713142 |
| 2.775048 | -1.157397097 | 0.485131909 | -0.930295502 | 0.527376212 |
| 2.785288 | -1.185993707 | 0.493399262 | -0.982817533 | 0.531136648 |
| 2.795528 | -1.208354074 | 0.502016362 | -1.031139764 | 0.535079234 |
| 2.805768 | -1.22496205 | 0.51310384 | -1.07475168 | 0.535992589 |
| 2.816008 | -1.234211791 | 0.524765598 | -1.114098484 | 0.536084026 |
| 2.826248 | -1.236701471 | 0.538654617 | -1.148834156 | 0.533549906 |
| 2.836488 | -1.231707144 | 0.5529054 | -1.178915491 | 0.530732213 |
| 2.846728 | -1.218957704 | 0.569154319 | -1.204188337 | 0.525214429 |
| 2.856968 | -1.198890442 | 0.585107792 | -1.224874894 | 0.520365411 |
| 2.867208 | -1.17207787 | 0.601971066 | -1.240150186 | 0.51389959 |
| 2.877448 | -1.138164952 | 0.618314277 | -1.250657141 | 0.508982911 |
| 2.887688 | -1.099259803 | 0.634284565 | -1.255622536 | 0.503516889 |
| 2.897928 | -1.054013529 | 0.648941752 | -1.255547297 | 0.500339118 |
| 2.908168 | -1.005238146 | 0.663032804 | -1.249807961 | 0.497683406 |
| 2.918408 | -0.951369703 | 0.674805948 | -1.238413905 | 0.497762752 |
| 2.928648 | -0.895521326 | 0.686285535 | -1.221912238 | 0.499316712 |
| 2.938889 | -0.834821058 | 0.694659784 | -1.199578973 | 0.503630694 |
| 2.949129 | -0.773770203 | 0.703302355 | -1.171941129 | 0.509927129 |
| 2.959368 | -0.709163166 | 0.70889228 | -1.138622353 | 0.518936147 |
| 2.969609 | -0.644887394 | 0.714660023 | -1.099891106 | 0.529646682 |
| 2.979849 | -0.579104029 | 0.71726562 | -1.055124624 | 0.541944083 |
| 2.990089 | -0.515147794 | 0.719098019 | -1.005014922 | 0.555403704 |
| 3.000329 | -0.451355365 | 0.717490108 | -0.949220626 | 0.569263842 |
| 3.010569 | -0.391114553 | 0.714442637 | -0.888109258 | 0.582955965 |
| 3.020809 | -0.332854593 | 0.706567391 | -0.822988446 | 0.595921474 |
| 3.031049 | -0.279359701 | 0.697084495 | -0.753867444 | 0.608098697 |
| 3.041289 | -0.229373117 | 0.683256623 | -0.682653103 | 0.61809922 |
| 3.051529 | -0.184765366 | 0.667822796 | -0.610251507 | 0.626427957 |
| 3.061769 | -0.145534434 | 0.649628219 | -0.537766488 | 0.632059067 |
| 3.072009 | -0.112081778 | 0.631620394 | -0.466875422 | 0.635068698 |
| 3.082249 | -0.084513206 | 0.613234705 | -0.3978406 | 0.634701191 |
| 3.092489 | -0.063891893 | 0.596123923 | -0.332651064 | 0.631337024 |
| 3.102729 | -0.049897853 | 0.581145002 | -0.27076907 | 0.623969036 |
| 3.112969 | -0.042572423 | 0.568420107 | -0.213779004 | 0.613121533 |
| 3.123209 | -0.042429106 | 0.559019474 | -0.161253281 | 0.598820969 |
| 3.133449 | -0.048801596 | 0.552628407 | -0.114109271 | 0.581029395 |
| 3.143689 | -0.061890367 | 0.550406317 | -0.071442072 | 0.560905346 |
| 3.153929 | -0.080136954 | 0.551633154 | -0.034334119 | 0.538652971 |
| 3.164169 | -0.104789751 | 0.55741091 | -0.001343309 | 0.515844638 |
| 3.174409 | -0.13334757 | 0.566956959 | 0.02686909 | 0.493394686 |
| 3.184649 | -0.166280472 | 0.579887098 | 0.051247479 | 0.472504195 |
| 3.194889 | -0.20222585 | 0.596604512 | 0.071642528 | 0.454864925 |
| 3.205129 | -0.241280505 | 0.614780772 | 0.088568729 | 0.441033933 |
| 3.215369 | -0.281740674 | 0.635689177 | 0.102089102 | 0.432757651 |
| 3.225609 | -0.324301268 | 0.656828551 | 0.112159153 | 0.429183895 |
| 3.235849 | -0.367408738 | 0.679933687 | 0.118633319 | 0.431839938 |
| 3.246089 | -0.412248426 | 0.702379719 | 0.1213239 | 0.43882846 |
| 3.256329 | -0.456886838 | 0.726714201 | 0.120381926 | 0.45033156 |
| 3.266569 | -0.503030977 | 0.750135643 | 0.115036981 | 0.465187906 |
| 3.276809 | -0.548939535 | 0.774857922 | 0.105708807 | 0.482809905 |
| 3.28705 | -0.596119322 | 0.798284453 | 0.092098086 | 0.502702457 |
| 3.29729 | -0.643233681 | 0.822571605 | 0.07443559 | 0.524545874 |
| 3.307529 | -0.691475609 | 0.844516917 | 0.052226508 | 0.548071949 |
| 3.31777 | -0.7391347 | 0.866192631 | 0.026446737 | 0.573305297 |
| 3.32801 | -0.787763892 | 0.884982049 | -0.003631611 | 0.599617908 |
| 3.33825 | -0.835734871 | 0.902346303 | -0.036993963 | 0.627214282 |
| 3.34849 | -0.883629304 | 0.916046866 | -0.074108239 | 0.654735951 |
| 3.35873 | -0.930668333 | 0.927486356 | -0.11363547 | 0.682479321 |
| 3.36897 | -0.97729912 | 0.935483019 | -0.156281042 | 0.709093021 |
| 3.37921 | -1.022198717 | 0.940668389 | -0.200534847 | 0.734314786 |
| 3.38945 | -1.066859794 | 0.942416394 | -0.24671024 | 0.756662087 |
| 3.39969 | -1.109886877 | 0.941740428 | -0.29375567 | 0.776838785 |
| 3.40993 | -1.152503896 | 0.937386021 | -0.342463728 | 0.792926815 |
| 3.42017 | -1.194207489 | 0.930945414 | -0.391469897 | 0.805986272 |
| 3.43041 | -1.236106563 | 0.921143239 | -0.441989361 | 0.815256185 |
| 3.44065 | -1.278315687 | 0.909086632 | -0.49263493 | 0.821626759 |
| 3.45089 | -1.32084403 | 0.894444804 | -0.544880883 | 0.824785635 |
| 3.46113 | -1.365363514 | 0.877915047 | -0.597259981 | 0.825920599 |
| 3.47137 | -1.410496283 | 0.859905369 | -0.650524866 | 0.824640439 |
| 3.48161 | -1.458239275 | 0.841263257 | -0.70420012 | 0.821920111 |
| 3.49185 | -1.507286382 | 0.82236347 | -0.758854913 | 0.818198736 |
| 3.50209 | -1.558855489 | 0.804734812 | -0.813409152 | 0.813346444 |
| 3.51233 | -1.612144899 | 0.78808048 | -0.869745865 | 0.808381755 |
| 3.52257 | -1.667473137 | 0.773798798 | -0.926479394 | 0.803039203 |
| 3.53281 | -1.724828106 | 0.761546348 | -0.985023143 | 0.79910278 |
| 3.54305 | -1.783493958 | 0.752398187 | -1.044909785 | 0.79529471 |
| 3.55329 | -1.843846457 | 0.744925621 | -1.106775075 | 0.793357282 |
| 3.56353 | -1.904367029 | 0.740091786 | -1.169822142 | 0.792143597 |
| 3.57377 | -1.965587003 | 0.736792749 | -1.234015241 | 0.79246176 |
| 3.58401 | -2.025699601 | 0.734519955 | -1.298874619 | 0.792808663 |
| 3.59425 | -2.084189987 | 0.732597697 | -1.363236847 | 0.793590755 |
| 3.60449 | -2.139703402 | 0.730270524 | -1.427037097 | 0.79333892 |
| 3.61473 | -2.191625601 | 0.726682011 | -1.489110846 | 0.792365297 |
| 3.62497 | -2.238615656 | 0.720869783 | -1.549014638 | 0.78922437 |
| 3.635211 | -2.280529606 | 0.713173849 | -1.606938375 | 0.78443963 |
| 3.645451 | -2.316727173 | 0.701953295 | -1.662205651 | 0.777355945 |
| 3.65569 | -2.347453457 | 0.688795194 | -1.715148186 | 0.767735338 |
| 3.665931 | -2.372196019 | 0.671838482 | -1.76582594 | 0.756118404 |
| 3.676171 | -2.392235386 | 0.653292046 | -1.813996683 | 0.741279716 |
| 3.686411 | -2.406867504 | 0.631889062 | -1.859267563 | 0.724538494 |
| 3.696651 | -2.416253293 | 0.609956929 | -1.901905984 | 0.704749807 |
| 3.706891 | -2.421016204 | 0.587141335 | -1.940394272 | 0.682123442 |
| 3.717131 | -2.420304692 | 0.565516683 | -1.974370504 | 0.656899908 |
| 3.727371 | -2.414397514 | 0.545734371 | -2.003582587 | 0.62826199 |
| 3.737611 | -2.402830088 | 0.530401343 | -2.026390324 | 0.597553683 |
| 3.747851 | -2.385945308 | 0.518930893 | -2.044223597 | 0.563769407 |
| 3.758091 | -2.363672614 | 0.515062288 | -2.055117657 | 0.529347588 |
| 3.768331 | -2.33612721 | 0.51726651 | -2.060679972 | 0.493891286 |
| 3.778571 | -2.303908086 | 0.527419237 | -2.060084164 | 0.46047307 |
| 3.788811 | -2.266824639 | 0.544304268 | -2.054569234 | 0.430027161 |
| 3.799051 | -2.225359988 | 0.568106442 | -2.043134411 | 0.405795728 |
| 3.809291 | -2.179415607 | 0.597446153 | -2.027311007 | 0.388672952 |
| 3.819531 | -2.129650533 | 0.631796101 | -2.005639762 | 0.381874224 |
| 3.829771 | -2.074844515 | 0.669494357 | -1.97937573 | 0.383701245 |
| 3.840011 | -2.016026497 | 0.710543054 | -1.947755555 | 0.395967693 |
| 3.850251 | -1.952461737 | 0.753185116 | -1.91142711 | 0.414675337 |
| 3.860491 | -1.884205502 | 0.796570561 | -1.869840235 | 0.439925511 |
| 3.870731 | -1.811156392 | 0.840355699 | -1.824271351 | 0.46806146 |
| 3.880971 | -1.733369961 | 0.883099746 | -1.773729354 | 0.498752191 |
| 3.891211 | -1.651310748 | 0.924775908 | -1.719319344 | 0.529074874 |
| 3.901451 | -1.56454355 | 0.96403707 | -1.660487031 | 0.558871909 |
| 3.911691 | -1.474455179 | 1.001243133 | -1.598206858 | 0.586183716 |
| 3.921931 | -1.380297681 | 1.035074827 | -1.53143688 | 0.611227361 |
| 3.932171 | -1.284071586 | 1.066327604 | -1.461660802 | 0.632319271 |
| 3.942411 | -1.184454195 | 1.093638455 | -1.38784727 | 0.649876327 |
| 3.952652 | -1.083713284 | 1.11800798 | -1.311904998 | 0.66322372 |
| 3.962891 | -0.980856557 | 1.138505295 | -1.232902478 | 0.673434308 |
| 3.973131 | -0.87740501 | 1.156156945 | -1.152539647 | 0.679512246 |
| 3.983371 | -0.772469851 | 1.169841573 | -1.070685469 | 0.682914804 |
| 3.993612 | -0.66813637 | 1.18045684 | -0.988073681 | 0.683881251 |
| 4.003851 | -0.563193968 | 1.18741558 | -0.904850733 | 0.682777354 |
| 4.014091 | -0.459231494 | 1.191001582 | -0.82139415 | 0.680248456 |
| 4.024332 | -0.356020067 | 1.190843643 | -0.738089437 | 0.676240705 |
| 4.034572 | -0.254194197 | 1.187253927 | -0.654143178 | 0.671389561 |
| 4.044812 | -0.154222498 | 1.180705841 | -0.571090175 | 0.66525407 |
| 4.055052 | -0.056581595 | 1.170448266 | -0.487586503 | 0.658784954 |
| 4.065292 | 0.038529196 | 1.157655631 | -0.405037257 | 0.650673076 |
| 4.075532 | 0.130096765 | 1.1421021 | -0.322836058 | 0.642274819 |
| 4.085772 | 0.217801062 | 1.123998887 | -0.242030341 | 0.632751118 |
| 4.096012 | 0.301477279 | 1.104039461 | -0.162442335 | 0.622586532 |
| 4.106252 | 0.379790759 | 1.081760163 | -0.084961494 | 0.61229557 |
| 4.116492 | 0.453204502 | 1.05807703 | -0.009460191 | 0.601537591 |
| 4.126732 | 0.521051132 | 1.032672307 | 0.062941889 | 0.591267146 |
| 4.136972 | 0.583404572 | 1.005893742 | 0.132519204 | 0.581523669 |
| 4.147212 | 0.640148091 | 0.97794485 | 0.197808408 | 0.57300662 |
| 4.157452 | 0.691421437 | 0.948545893 | 0.259856535 | 0.565607875 |
| 4.167692 | 0.737130046 | 0.918541381 | 0.316389808 | 0.56056189 |
| 4.177932 | 0.77762056 | 0.887431074 | 0.368629254 | 0.556688173 |
| 4.188172 | 0.812774488 | 0.855948169 | 0.415112157 | 0.555709908 |
| 4.198412 | 0.843033445 | 0.824869856 | 0.456415314 | 0.556213629 |
| 4.208652 | 0.868446195 | 0.793634073 | 0.491389922 | 0.558665161 |
| 4.218892 | 0.889489833 | 0.764034855 | 0.521129712 | 0.562584718 |
| 4.229132 | 0.906552219 | 0.735601898 | 0.544809401 | 0.567916844 |
| 4.239372 | 0.919806355 | 0.709521604 | 0.563310145 | 0.57351969 |
| 4.249612 | 0.929913607 | 0.685764326 | 0.576700058 | 0.580210664 |
| 4.259852 | 0.936840925 | 0.664934743 | 0.585273235 | 0.586628829 |
| 4.270092 | 0.940647924 | 0.646867105 | 0.589770685 | 0.593415669 |
| 4.280333 | 0.941837154 | 0.631829364 | 0.590025763 | 0.599627791 |
| 4.290572 | 0.939978934 | 0.619592542 | 0.586920574 | 0.605239829 |
| 4.300812 | 0.935419952 | 0.610587509 | 0.580262872 | 0.609744372 |
| 4.311052 | 0.928088944 | 0.604151711 | 0.571120777 | 0.612827271 |
| 4.321292 | 0.91805919 | 0.600687157 | 0.558833563 | 0.613978361 |
| 4.331532 | 0.905510043 | 0.599983521 | 0.545300953 | 0.612503891 |
| 4.341773 | 0.890768944 | 0.602625389 | 0.529350693 | 0.609175071 |
| 4.352013 | 0.873491125 | 0.607206068 | 0.512375673 | 0.602899005 |
| 4.362253 | 0.853909365 | 0.614866483 | 0.494051562 | 0.594866793 |
| 4.372493 | 0.831885569 | 0.623630678 | 0.474586565 | 0.584650709 |
| 4.382733 | 0.80755816 | 0.63400126 | 0.453864399 | 0.573144745 |
| 4.392972 | 0.780234182 | 0.644463807 | 0.432180773 | 0.560056052 |
| 4.403213 | 0.750434155 | 0.654997238 | 0.409082108 | 0.546038743 |
| 4.413453 | 0.717355919 | 0.664649301 | 0.385463342 | 0.530758071 |
| 4.423693 | 0.681994803 | 0.673878234 | 0.360931024 | 0.514859245 |
| 4.433933 | 0.643572123 | 0.681170412 | 0.336534302 | 0.49755816 |
| 4.444173 | 0.603522193 | 0.687985648 | 0.312088991 | 0.48024674 |
| 4.454413 | 0.561424339 | 0.69284989 | 0.288575174 | 0.461593673 |
| 4.464653 | 0.51896832 | 0.696564741 | 0.265862056 | 0.443289094 |
| 4.474893 | 0.475924377 | 0.698473264 | 0.244544066 | 0.424002497 |
| 4.485133 | 0.434030945 | 0.698714158 | 0.22481815 | 0.405337429 |
| 4.495373 | 0.392739974 | 0.696746787 | 0.206035525 | 0.386195741 |
| 4.505613 | 0.353285025 | 0.693273867 | 0.189643186 | 0.367447398 |
| 4.515853 | 0.315249709 | 0.687238383 | 0.173949086 | 0.348478844 |
| 4.526093 | 0.278768755 | 0.679697169 | 0.160076317 | 0.329633793 |
| 4.536333 | 0.243676837 | 0.669820717 | 0.14724883 | 0.310684049 |
| 4.546573 | 0.209824845 | 0.658537367 | 0.135675963 | 0.292089048 |
| 4.556813 | 0.176982205 | 0.644940299 | 0.124856816 | 0.273228728 |
| 4.567053 | 0.145747516 | 0.629909559 | 0.11478341 | 0.255188589 |
| 4.577293 | 0.115094431 | 0.612578136 | 0.105066521 | 0.237480402 |
| 4.587533 | 0.086634664 | 0.593412705 | 0.096061091 | 0.220707993 |
| 4.597773 | 0.059334573 | 0.571966158 | 0.086817855 | 0.205268881 |
| 4.608013 | 0.034562886 | 0.54890855 | 0.07831036 | 0.190826821 |
| 4.618253 | 0.011543602 | 0.524327075 | 0.06960522 | 0.178902605 |
| 4.628493 | -0.008395677 | 0.498642832 | 0.061946214 | 0.168999137 |
| 4.638733 | -0.026292945 | 0.472330206 | 0.054291922 | 0.162154903 |
| 4.648973 | -0.040634112 | 0.446030644 | 0.047922892 | 0.158359649 |
| 4.659214 | -0.052199933 | 0.420519957 | 0.042541415 | 0.157830286 |
| 4.669454 | -0.060568611 | 0.395689953 | 0.03807077 | 0.159667006 |
| 4.679694 | -0.06545824 | 0.372253807 | 0.035097302 | 0.163572856 |
| 4.689933 | -0.067818177 | 0.350235821 | 0.032924236 | 0.167741257 |
| 4.700173 | -0.066996712 | 0.330203688 | 0.031671663 | 0.171531744 |
| 4.710413 | -0.064278287 | 0.3114785 | 0.031370108 | 0.173849697 |
| 4.720654 | -0.059070109 | 0.295084143 | 0.031278025 | 0.174887212 |
| 4.730894 | -0.052990331 | 0.280392953 | 0.031806983 | 0.17430351 |
| 4.741134 | -0.044858338 | 0.268650155 | 0.032690918 | 0.172885375 |
| 4.751374 | -0.0363896 | 0.259387934 | 0.034087876 | 0.171575936 |

| | | | | |
|---|---|---|---|---|
| 4.761614 | -0.025806397 | 0.338727166 | -0.196531384 | 0.26673682 |
| 4.771854 | -0.024665262 | 0.33800014 | -0.206148737 | 0.26349116 |
| 4.782094 | -0.021861942 | 0.336341993 | -0.214896943 | 0.25878399 |
| 4.792334 | -0.018854371 | 0.333233845 | -0.222628263 | 0.25320699 |
| 4.802574 | -0.014500021 | 0.328709312 | -0.22877559 | 0.24611279 |
| 4.812814 | -0.010513362 | 0.323100116 | -0.234577867 | 0.23868514 |
| 4.823054 | -0.005712321 | 0.316258217 | -0.238702014 | 0.23050543 |
| 4.833294 | -0.001539368 | 0.309229074 | -0.242235202 | 0.22204547 |
| 4.843534 | 0.001924941 | 0.30128435 | -0.244320616 | 0.21352519 |
| 4.853774 | 0.004986163 | 0.29337083 | -0.245071796 | 0.20455376 |
| 4.864014 | 0.007663139 | 0.286002565 | -0.244809853 | 0.19588772 |
| 4.874254 | 0.009871881 | 0.279031359 | -0.242665531 | 0.18733643 |
| 4.884494 | 0.011663897 | 0.272931774 | -0.240295546 | 0.17941392 |
| 4.894734 | 0.012956214 | 0.268105399 | -0.2370656 | 0.17256142 |
| 4.904974 | 0.013350634 | 0.264028185 | -0.234755701 | 0.16707195 |
| 4.915214 | 0.013654339 | 0.261708802 | -0.233178314 | 0.16354816 |
| 4.925454 | 0.011810832 | 0.26008263 | -0.233999891 | 0.16237971 |
| 4.935694 | 0.009614471 | 0.259623269 | -0.236400251 | 0.16253295 |
| 4.945934 | 0.005718725 | 0.25998473 | -0.241233615 | 0.1647909 |
| 4.956174 | 0.001779697 | 0.260856608 | -0.247907225 | 0.16744099 |
| 4.966414 | -0.003268549 | 0.261543508 | -0.256029068 | 0.17144202 |
| 4.976655 | -0.007537092 | 0.261780529 | -0.265025455 | 0.17358537 |
| 4.986895 | -0.01148717 | 0.261178126 | -0.274837488 | 0.17591008 |
| 4.997134 | -0.014240279 | 0.258595164 | -0.284810625 | 0.17668555 |
| 5.007374 | -0.016066296 | 0.255218658 | -0.29504173 | 0.17680838 |
| 5.017614 | -0.016637185 | 0.250568583 | -0.305179877 | 0.17627822 |
| 5.027854 | -0.015724443 | 0.245127209 | -0.314638155 | 0.17586628 |
| 5.038095 | -0.014297924 | 0.239500466 | -0.323210113 | 0.17643553 |
| 5.048335 | -0.011553115 | 0.234719304 | -0.32993103 | 0.17795895 |
| 5.058575 | -0.008504564 | 0.229899977 | -0.334301875 | 0.18021049 |
| 5.068815 | -0.005033432 | 0.226975606 | -0.335085735 | 0.18334086 |
| 5.079055 | -0.001354209 | 0.223824311 | -0.332635402 | 0.18657454 |
| 5.089295 | 0.002462352 | 0.222299712 | -0.325796497 | 0.18873383 |
| 5.099535 | 0.005741114 | 0.220983095 | -0.315987493 | 0.19037783 |
| 5.109775 | 0.009092898 | 0.221051848 | -0.302755159 | 0.1901708 |
| 5.120015 | 0.011642816 | 0.221730333 | -0.287697337 | 0.18915129 |
| 5.130255 | 0.013465856 | 0.224062642 | -0.270806024 | 0.18636212 |
| 5.140495 | 0.014307078 | 0.227652076 | -0.25366524 | 0.18337581 |
| 5.150735 | 0.014050159 | 0.232853604 | -0.236422966 | 0.1797068 |
| 5.160975 | 0.01262444 | 0.239782299 | -0.219071098 | 0.17631339 |
| 5.171215 | 0.010376277 | 0.246214816 | -0.202472041 | 0.17332896 |
| 5.181455 | 0.007518603 | 0.254539245 | -0.185868052 | 0.1711694 |
| 5.191695 | 0.005762293 | 0.260620171 | -0.16999813 | 0.16951482 |
| 5.201935 | 0.003772724 | 0.266883383 | -0.154145883 | 0.16877116 |
| 5.212175 | 0.00498934 | 0.270522822 | -0.138637868 | 0.16823017 |
| 5.222415 | 0.007640896 | 0.273447969 | -0.123850335 | 0.16850055 |
| 5.232655 | 0.013730235 | 0.273939794 | -0.109731263 | 0.16880652 |
| 5.242895 | 0.021445702 | 0.272957602 | -0.096317584 | 0.16999513 |
| 5.253135 | 0.031753692 | 0.270337585 | -0.084384789 | 0.17120676 |
| 5.263375 | 0.042933567 | 0.26647387 | -0.073362506 | 0.17279427 |
| 5.273615 | 0.055079775 | 0.261316106 | -0.063810193 | 0.17497879 |
| 5.283855 | 0.066986658 | 0.255106535 | -0.055250274 | 0.17747929 |
| 5.294095 | 0.079040004 | 0.248601438 | -0.047514444 | 0.18058767 |
| 5.304335 | 0.090822336 | 0.242216828 | -0.040535674 | 0.18378153 |
| 5.314575 | 0.102398081 | 0.236348639 | -0.03302669 | 0.18776851 |
| 5.324815 | 0.11444839 | 0.231284634 | -0.02588351 | 0.19197196 |
| 5.335055 | 0.126699071 | 0.22685804 | -0.017289078 | 0.19585146 |
| 5.345295 | 0.139705768 | 0.222596529 | -0.008251925 | 0.19947971 |
| 5.355536 | 0.153297304 | 0.218314211 | 0.001903656 | 0.20262839 |
| 5.365776 | 0.167471809 | 0.213923694 | 0.012962586 | 0.20459593 |
| 5.376016 | 0.181820926 | 0.208995168 | 0.024650574 | 0.20535354 |
| 5.386256 | 0.196524625 | 0.204073627 | 0.036528382 | 0.20520484 |
| 5.396495 | 0.210477088 | 0.199914091 | 0.048407813 | 0.20401878 |
| 5.406735 | 0.224068916 | 0.19618701 | 0.06016211 | 0.20167934 |
| 5.416975 | 0.236394779 | 0.195083526 | 0.070807159 | 0.19971734 |
| 5.427216 | 0.248009661 | 0.194677421 | 0.081254835 | 0.19706267 |
| 5.437456 | 0.257785313 | 0.197297638 | 0.090234736 | 0.19545678 |
| 5.447696 | 0.266461059 | 0.201039623 | 0.098450717 | 0.19397798 |
| 5.457936 | 0.274418972 | 0.206645325 | 0.105372795 | 0.19277742 |
| 5.468176 | 0.280703031 | 0.21289345 | 0.111340264 | 0.19278671 |
| 5.478416 | 0.287232752 | 0.219674335 | 0.115873659 | 0.19142279 |
| 5.488656 | 0.29310232 | 0.22653364 | 0.119564939 | 0.19023776 |
| 5.498896 | 0.299425528 | 0.233198836 | 0.122788802 | 0.18769086 |
| 5.509136 | 0.306084134 | 0.239546023 | 0.125326247 | 0.18396342 |
| 5.519376 | 0.313682061 | 0.245268526 | 0.128690534 | 0.17825265 |
| 5.529616 | 0.322466436 | 0.250762742 | 0.132545942 | 0.17110443 |
| 5.539856 | 0.331670096 | 0.25620749 | 0.137781941 | 0.16299341 |
| 5.550096 | 0.342079514 | 0.26171783 | 0.144793842 | 0.15375874 |
| 5.560336 | 0.352603421 | 0.267212795 | 0.153172765 | 0.14526605 |
| 5.570576 | 0.363551289 | 0.273040766 | 0.163215107 | 0.13707347 |
| 5.580816 | 0.373446556 | 0.279302434 | 0.173855521 | 0.13120243 |
| 5.591056 | 0.383471644 | 0.285704532 | 0.185067599 | 0.12686038 |
| 5.601296 | 0.391452336 | 0.292375247 | 0.195419763 | 0.12523524 |
| 5.611536 | 0.399380652 | 0.29933402 | 0.205117461 | 0.12519182 |
| 5.621776 | 0.40580658 | 0.305721231 | 0.213065982 | 0.127218 |
| 5.632016 | 0.411794952 | 0.312493874 | 0.219175171 | 0.13042774 |
| 5.642256 | 0.417706774 | 0.318722606 | 0.223305598 | 0.13485619 |
| 5.652496 | 0.423127469 | 0.324578992 | 0.225850999 | 0.13969933 |
| 5.662736 | 0.428953517 | 0.329516487 | 0.226849102 | 0.14494789 |
| 5.672976 | 0.434522899 | 0.333613478 | 0.226404124 | 0.15057713 |
| 5.683217 | 0.439804663 | 0.335960292 | 0.225521336 | 0.15575805 |
| 5.693457 | 0.444611615 | 0.336489993 | 0.224417744 | 0.15988372 |
| 5.703696 | 0.448535564 | 0.334301762 | 0.222971284 | 0.16298072 |
| 5.713936 | 0.451792133 | 0.330040452 | 0.222160022 | 0.16501005 |
| 5.724176 | 0.453986416 | 0.323227926 | 0.221516911 | 0.16542293 |
| 5.734416 | 0.455674831 | 0.31471528 | 0.221240415 | 0.16496395 |
| 5.744657 | 0.457125629 | 0.305290263 | 0.221105303 | 0.16324975 |
| 5.754897 | 0.458235737 | 0.295646768 | 0.220741706 | 0.16167826 |
| 5.765137 | 0.459806927 | 0.286017044 | 0.220624183 | 0.15953854 |
| 5.775377 | 0.461446357 | 0.277845389 | 0.219672239 | 0.15850133 |
| 5.785617 | 0.463677117 | 0.270896432 | 0.219011289 | 0.15704197 |
| 5.795857 | 0.46591115 | 0.265498525 | 0.217617535 | 0.15725019 |
| 5.806097 | 0.468420514 | 0.261242116 | 0.216628245 | 0.15783874 |
| 5.816337 | 0.470578585 | 0.258311609 | 0.214736661 | 0.1594826 |
| 5.826577 | 0.47317305 | 0.255679563 | 0.213682533 | 0.16197303 |
| 5.836817 | 0.474282991 | 0.25400316 | 0.211322693 | 0.16472046 |
| 5.847057 | 0.475114146 | 0.252422072 | 0.208527373 | 0.16814763 |
| 5.857297 | 0.475266835 | 0.251038736 | 0.204596445 | 0.17106847 |
| 5.867537 | 0.474027208 | 0.250381609 | 0.199210534 | 0.17304801 |
| 5.877777 | 0.471878267 | 0.25006939 | 0.19227008 | 0.17374938 |
| 5.888017 | 0.4689549 | 0.251184281 | 0.184092726 | 0.17377651 |
| 5.898257 | 0.465081968 | 0.252486953 | 0.175231675 | 0.17157484 |
| 5.908497 | 0.460359746 | 0.256089425 | 0.165397936 | 0.16989673 |
| 5.918737 | 0.455118501 | 0.258980767 | 0.156384044 | 0.16706607 |
| 5.928977 | 0.449496752 | 0.264798707 | 0.147334455 | 0.16637685 |
| 5.939217 | 0.44335326 | 0.269227368 | 0.139738842 | 0.16589206 |
| 5.949457 | 0.436863619 | 0.275431549 | 0.133272621 | 0.16851418 |
| 5.959697 | 0.430033055 | 0.281238277 | 0.127072692 | 0.17211036 |
| 5.969937 | 0.423084908 | 0.287743959 | 0.1221621 | 0.17794189 |
| 5.980177 | 0.415592689 | 0.294174039 | 0.116935831 | 0.18465344 |
| 5.990417 | 0.408169995 | 0.300911236 | 0.112251242 | 0.19235494 |
| 6.000657 | 0.400765821 | 0.308282818 | 0.106749629 | 0.20067575 |
| 6.010897 | 0.392875453 | 0.315480232 | 0.101015574 | 0.20914659 |
| 6.021137 | 0.38511386 | 0.32269929 | 0.094816791 | 0.21803013 |
| 6.031377 | 0.377169785 | 0.330724409 | 0.088244562 | 0.22726841 |
| 6.041617 | 0.368683522 | 0.338062597 | 0.08125868 | 0.23658195 |
| 6.051857 | 0.36016275 | 0.346014025 | 0.073926071 | 0.24628781 |
| 6.062098 | 0.350609347 | 0.354030751 | 0.066055137 | 0.25628884 |
| 6.072338 | 0.340468739 | 0.362534274 | 0.057832439 | 0.26569852 |
| 6.082578 | 0.32947805 | 0.371705284 | 0.048637109 | 0.27543511 |
| 6.092818 | 0.317579957 | 0.380794596 | 0.039250315 | 0.28352487 |
| 6.103057 | 0.304507824 | 0.390640752 | 0.028749741 | 0.29135872 |
| 6.113297 | 0.290773441 | 0.400807843 | 0.01834159 | 0.29764734 |
| 6.123538 | 0.275731554 | 0.410001748 | 0.007786116 | 0.30265584 |
| 6.133778 | 0.260783239 | 0.419728864 | -0.002510764 | 0.30689575 |
| 6.144018 | 0.244591847 | 0.427940816 | -0.011707173 | 0.30999439 |
| 6.154258 | 0.22836592 | 0.43554455 | -0.020079403 | 0.31242499 |
| 6.164498 | 0.21208575 | 0.442762827 | -0.027435937 | 0.31499614 |
| 6.174738 | 0.195361158 | 0.44928693 | -0.033792458 | 0.31764703 |
| 6.184978 | 0.178104064 | 0.455912662 | -0.039369871 | 0.32136188 |
| 6.195218 | 0.160565458 | 0.463154007 | -0.045015374 | 0.325856 |
| 6.205458 | 0.141904703 | 0.470999741 | -0.050385068 | 0.3318764 |
| 6.215698 | 0.122534984 | 0.479303172 | -0.056066105 | 0.33869947 |
| 6.225938 | 0.102195183 | 0.488600675 | -0.062175657 | 0.34610334 |
| 6.236178 | 0.081437739 | 0.497120737 | -0.068405436 | 0.35299752 |
| 6.246418 | 0.060403528 | 0.505676845 | -0.074913672 | 0.35890813 |
| 6.256658 | 0.039969737 | 0.512924499 | -0.08168195 | 0.36261688 |
| 6.266898 | 0.020455738 | 0.518513698 | -0.088418599 | 0.36342289 |
| 6.277138 | 0.002516622 | 0.522457714 | -0.095113371 | 0.36124625 |
| 6.287378 | -0.014425769 | 0.524854505 | -0.102466411 | 0.35594269 |
| 6.297618 | -0.029307144 | 0.525952324 | -0.109823935 | 0.34759903 |
| 6.307858 | -0.043630948 | 0.5262125 | -0.117442317 | 0.33729955 |
| 6.318098 | -0.056517233 | 0.526194142 | -0.126229203 | 0.32507156 |
| 6.328338 | -0.069149531 | 0.525832021 | -0.135070206 | 0.31215915 |
| 6.338578 | -0.081619686 | 0.526359654 | -0.145143033 | 0.29869987 |
| 6.348818 | -0.093773274 | 0.526611936 | -0.156205336 | 0.28571455 |
| 6.359058 | -0.106156218 | 0.527821553 | -0.168395038 | 0.27325615 |
| 6.369298 | -0.119008487 | 0.528682504 | -0.182231182 | 0.26191981 |
| 6.379539 | -0.131991687 | 0.529449931 | -0.197355498 | 0.25166973 |
| 6.389779 | -0.14520591 | 0.530020861 | -0.213807198 | 0.24257606 |
| 6.400019 | -0.159144044 | 0.529634845 | -0.231954174 | 0.23495946 |
| 6.410258 | -0.172811331 | 0.527607841 | -0.250964196 | 0.22772497 |
| 6.420498 | -0.186453086 | 0.524835788 | -0.27056463 | 0.22178999 |
| 6.430738 | -0.200005688 | 0.519782625 | -0.290905548 | 0.21554435 |
| 6.440979 | -0.212514578 | 0.513735481 | -0.311243344 | 0.20960229 |
| 6.451219 | -0.224714459 | 0.505948646 | -0.331600814 | 0.20352577 |
| 6.461459 | -0.235528097 | 0.497346809 | -0.35138935 | 0.19627368 |
| 6.471699 | -0.246591841 | 0.487718988 | -0.371498939 | 0.18893676 |
| 6.481939 | -0.255885963 | 0.477919336 | -0.390712148 | 0.18042843 |
| 6.492179 | -0.266210939 | 0.467075333 | -0.409643995 | 0.17127197 |
| 6.502419 | -0.274871497 | 0.456239992 | -0.428139389 | 0.1611 |
| 6.512659 | -0.28471426 | 0.444045768 | -0.445402418 | 0.15111571 |
| 6.522899 | -0.293679298 | 0.431268186 | -0.461349845 | 0.14099915 |
| 6.533139 | -0.302365791 | 0.417644876 | -0.476005737 | 0.13188075 |
| 6.543379 | -0.311265289 | 0.402736344 | -0.487853587 | 0.12432508 |
| 6.553619 | -0.319570744 | 0.387533319 | -0.497440875 | 0.11925019 |
| 6.563859 | -0.327115963 | 0.371832315 | -0.504250037 | 0.11703361 |
| 6.574099 | -0.334681481 | 0.356118533 | -0.508322009 | 0.11694649 |
| 6.584339 | -0.341434191 | 0.341657505 | -0.509985072 | 0.11921964 |
| 6.594579 | -0.347608849 | 0.327997072 | -0.509728653 | 0.12233417 |
| 6.604819 | -0.352581705 | 0.315765664 | -0.508368701 | 0.12567349 |
| 6.615059 | -0.357355352 | 0.304720729 | -0.506292539 | 0.12872524 |
| 6.625299 | -0.360533386 | 0.295501855 | -0.504660636 | 0.1305953 |
| 6.635539 | -0.36233371 | 0.286207723 | -0.503092455 | 0.1324421 |
| 6.645779 | -0.363114558 | 0.278221793 | -0.502649997 | 0.13318332 |
| 6.656019 | -0.362527311 | 0.269316936 | -0.502088053 | 0.1344892 |
| 6.666259 | -0.360117661 | 0.260766432 | -0.502539886 | 0.1359289 |
| 6.676499 | -0.356975733 | 0.251213215 | -0.502243893 | 0.13858638 |
| 6.686739 | -0.353120963 | 0.241685935 | -0.501809188 | 0.14233272 |
| 6.69698 | -0.348501886 | 0.23140778 | -0.50033451 | 0.14760442 |
| 6.707219 | -0.344107166 | 0.221436138 | -0.497619263 | 0.15339687 |
| 6.717459 | -0.33982515 | 0.211953469 | -0.493480206 | 0.16026964 |
| 6.727699 | -0.336389344 | 0.203231366 | -0.487563929 | 0.16720527 |
| 6.737939 | -0.333113052 | 0.196492201 | -0.479780674 | 0.17420946 |
| 6.748179 | -0.329537089 | 0.192324441 | -0.469877613 | 0.18137867 |
| 6.75842 | -0.32650838 | 0.191620218 | -0.458881649 | 0.18796035 |
| 6.76866 | -0.322025202 | 0.194507534 | -0.445557654 | 0.19527682 |
| 6.7789 | -0.316907831 | 0.201704271 | -0.431855072 | 0.20236922 |
| 6.78914 | -0.310467328 | 0.212009411 | -0.417044514 | 0.21036594 |
| 6.79938 | -0.302877103 | 0.224523794 | -0.402115721 | 0.2188617 |
| 6.809619 | -0.294463286 | 0.239323662 | -0.387120091 | 0.22806917 |
| 6.81986 | -0.285116828 | 0.254041134 | -0.372247499 | 0.23758521 |
| 6.8301 | -0.275709258 | 0.269349221 | -0.357921151 | 0.24706722 |
| 6.84034 | -0.265919573 | 0.283433971 | -0.343888005 | 0.25578603 |
| 6.85058 | -0.256291241 | 0.297240172 | -0.330294079 | 0.26411204 |
| 6.86082 | -0.247332028 | 0.309745584 | -0.317035926 | 0.27092824 |
| 6.87106 | -0.238383301 | 0.321169667 | -0.304406645 | 0.27643565 |
| 6.8813 | -0.230229574 | 0.331471306 | -0.29245144 | 0.28073971 |
| 6.89154 | -0.222897403 | 0.340561882 | -0.281004744 | 0.28344088 |
| 6.90178 | -0.216709634 | 0.348481632 | -0.270158388 | 0.28519607 |
| 6.91202 | -0.211004995 | 0.355414996 | -0.26032079 | 0.28603395 |
| 6.92226 | -0.206330751 | 0.360899936 | -0.250824555 | 0.28660383 |
| 6.9325 | -0.202503351 | 0.365696309 | -0.241915508 | 0.28707412 |
| 6.94274 | -0.199024916 | 0.369997579 | -0.233158836 | 0.28796525 |
| 6.95298 | -0.195508401 | 0.37403037 | -0.224739963 | 0.28925042 |
| 6.96322 | -0.192196339 | 0.378665668 | -0.21575472 | 0.29137461 |
| 6.97346 | -0.187604581 | 0.383565473 | -0.206271769 | 0.29398806 |
| 6.9837 | -0.182966416 | 0.389992443 | -0.195318749 | 0.29781927 |
| 6.99394 | -0.176998968 | 0.397161171 | -0.18285882 | 0.30153217 |
| 7.00418 | -0.169700334 | 0.405546395 | -0.168484034 | 0.30618975 |
| 7.01442 | -0.161218653 | 0.414711666 | -0.151091128 | 0.31085512 |
| 7.02466 | -0.151438956 | 0.424840142 | -0.131559782 | 0.31560711 |
| 7.0349 | -0.140873681 | 0.435193342 | -0.109281399 | 0.31999398 |
| 7.04514 | -0.129354293 | 0.445812661 | -0.084769337 | 0.32389267 |
| 7.05538 | -0.11765413 | 0.456177344 | -0.05879317 | 0.3272225 |
| 7.06562 | -0.106047454 | 0.466361161 | -0.031196879 | 0.32914836 |
| 7.075861 | -0.095103544 | 0.476277593 | -0.003730207 | 0.33087104 |
| 7.086101 | -0.084722353 | 0.485411272 | 0.02398114 | 0.33156689 |
| 7.096341 | -0.075232809 | 0.494504946 | 0.050907587 | 0.33165515 |
| 7.106581 | -0.066695268 | 0.502722468 | 0.076624433 | 0.33178957 |
| 7.11682 | -0.058166357 | 0.510994188 | 0.100659301 | 0.33226928 |
| 7.12706 | -0.050417982 | 0.518530644 | 0.123588819 | 0.33313574 |
| 7.1373 | -0.042359145 | 0.525373581 | 0.144111278 | 0.33422946 |
| 7.147541 | -0.034879855 | 0.531802496 | 0.16359578 | 0.33592303 |
| 7.157781 | -0.026753282 | 0.537263916 | 0.18126172 | 0.33784572 |

| | | | | |
|---|---|---|---|---|
| 4.761614 | 0.193440701 | 0.311608586 | 0.033796206 | 0.30026144 |
| 4.771854 | 0.199719741 | 0.328258273 | 0.055803085 | 0.300370711 |
| 4.782094 | 0.207363535 | 0.341889189 | 0.076107771 | 0.300650581 |
| 4.792334 | 0.217082767 | 0.354475836 | 0.095175746 | 0.301449626 |
| 4.802574 | 0.228311364 | 0.364488461 | 0.112594822 | 0.302178642 |
| 4.812814 | 0.241772253 | 0.373528424 | 0.128586062 | 0.303122193 |
| 4.823054 | 0.257256669 | 0.380206636 | 0.143895668 | 0.30338334 |
| 4.833294 | 0.274143357 | 0.386328507 | 0.158516593 | 0.302846018 |
| 4.843534 | 0.293012243 | 0.389602701 | 0.17298513 | 0.301579716 |
| 4.853774 | 0.312814637 | 0.392273446 | 0.187942072 | 0.299299157 |
| 4.864014 | 0.334212407 | 0.39237103 | 0.203268064 | 0.29628331 |
| 4.874254 | 0.357110682 | 0.391289875 | 0.219798834 | 0.29343479 |
| 4.884494 | 0.380935863 | 0.388976474 | 0.23688127 | 0.290631557 |
| 4.894734 | 0.406396583 | 0.386288777 | 0.254884343 | 0.288346895 |
| 4.904974 | 0.432730553 | 0.383235106 | 0.273695615 | 0.286675161 |
| 4.915214 | 0.460710364 | 0.380654265 | 0.293075985 | 0.285970392 |
| 4.925454 | 0.488577592 | 0.379305106 | 0.312517313 | 0.285409573 |
| 4.935694 | 0.51729174 | 0.378486522 | 0.332684483 | 0.285109586 |
| 4.945934 | 0.545272272 | 0.37935303 | 0.35255575 | 0.284778092 |
| 4.956174 | 0.573047516 | 0.380364559 | 0.373262492 | 0.284757329 |
| 4.966414 | 0.59891496 | 0.382568536 | 0.393804617 | 0.284523832 |
| 4.976655 | 0.624020398 | 0.383860876 | 0.414992192 | 0.284497403 |
| 4.986895 | 0.647652532 | 0.385611973 | 0.436490057 | 0.284182964 |
| 4.997134 | 0.669760196 | 0.384707539 | 0.458444084 | 0.284171263 |
| 5.007374 | 0.691208564 | 0.382913998 | 0.480158352 | 0.283709285 |
| 5.017614 | 0.712176888 | 0.378126819 | 0.501391202 | 0.282991901 |
| 5.027854 | 0.733171493 | 0.37175441 | 0.522417741 | 0.281803488 |
| 5.038095 | 0.754505977 | 0.363049504 | 0.541827159 | 0.280297429 |
| 5.048335 | 0.776571095 | 0.353505007 | 0.560473063 | 0.279283824 |
| 5.058575 | 0.799144134 | 0.343344881 | 0.577994623 | 0.27774535 |
| 5.068815 | 0.822167084 | 0.33373274 | 0.59440194 | 0.277238628 |
| 5.079055 | 0.845850403 | 0.32495652 | 0.610189489 | 0.27744343 |
| 5.089295 | 0.869730701 | 0.317883542 | 0.625970777 | 0.278553229 |
| 5.099535 | 0.893200487 | 0.312722352 | 0.641273771 | 0.27975372 |
| 5.109775 | 0.917095006 | 0.309702852 | 0.657478609 | 0.282348187 |
| 5.120015 | 0.940778166 | 0.30815688 | 0.673596838 | 0.284384487 |
| 5.130255 | 0.963794778 | 0.308268816 | 0.690276448 | 0.286526316 |
| 5.140495 | 0.987304817 | 0.308886674 | 0.707278622 | 0.287637277 |
| 5.150735 | 1.009530882 | 0.309796068 | 0.72447595 | 0.288161164 |
| 5.160975 | 1.031554292 | 0.310184645 | 0.741580946 | 0.287717741 |
| 5.171215 | 1.051963925 | 0.30970774 | 0.758872198 | 0.286025875 |
| 5.181455 | 1.070831885 | 0.30875494 | 0.775029591 | 0.284102009 |
| 5.191695 | 1.088253041 | 0.306663434 | 0.79175938 | 0.281100789 |
| 5.201935 | 1.102784256 | 0.304090392 | 0.807232959 | 0.278557897 |
| 5.212175 | 1.115916957 | 0.302071357 | 0.822369805 | 0.275176796 |
| 5.222415 | 1.126661827 | 0.299612323 | 0.837551858 | 0.272425837 |
| 5.232655 | 1.135508458 | 0.298969781 | 0.851277036 | 0.269591788 |
| 5.242895 | 1.143193344 | 0.298041508 | 0.865585932 | 0.266351822 |
| 5.253135 | 1.149570038 | 0.299031713 | 0.879120665 | 0.263728137 |
| 5.263375 | 1.155047466 | 0.300530954 | 0.892360032 | 0.260521586 |
| 5.273615 | 1.160246134 | 0.302629254 | 0.905437376 | 0.258031444 |
| 5.283855 | 1.165303429 | 0.305070037 | 0.917885329 | 0.255171539 |
| 5.294095 | 1.170419564 | 0.307740601 | 0.929528245 | 0.253587063 |
| 5.304335 | 1.175896972 | 0.310117108 | 0.940820694 | 0.251488486 |
| 5.314575 | 1.181842387 | 0.311853756 | 0.950431279 | 0.250331415 |
| 5.324815 | 1.18821307 | 0.313749914 | 0.958917805 | 0.249015369 |
| 5.335055 | 1.194909761 | 0.315450209 | 0.965707804 | 0.247756487 |
| 5.345295 | 1.201815486 | 0.317212547 | 0.971149172 | 0.246038403 |
| 5.355536 | 1.207899014 | 0.319328288 | 0.974248196 | 0.243991537 |
| 5.365776 | 1.213900884 | 0.322422125 | 0.976118284 | 0.241610102 |
| 5.376016 | 1.217943132 | 0.325832491 | 0.976665693 | 0.239156412 |
| 5.386256 | 1.220809042 | 0.33070652 | 0.975225236 | 0.236360402 |
| 5.396495 | 1.22121949 | 0.336352926 | 0.973301249 | 0.234903156 |
| 5.406735 | 1.219361464 | 0.3431316 | 0.970647165 | 0.233141327 |
| 5.416975 | 1.214886069 | 0.350876513 | 0.967558146 | 0.232597957 |
| 5.427216 | 1.208550622 | 0.359840741 | 0.964446902 | 0.232467681 |
| 5.437456 | 1.199669073 | 0.368887817 | 0.962481763 | 0.232656796 |
| 5.447696 | 1.189659933 | 0.378944523 | 0.960951754 | 0.232556202 |
| 5.457936 | 1.17847837 | 0.388521188 | 0.961283224 | 0.231944754 |
| 5.468176 | 1.167026222 | 0.39950072 | 0.962263763 | 0.230579933 |
| 5.478416 | 1.155158708 | 0.410291881 | 0.964830961 | 0.228447887 |
| 5.488656 | 1.143578241 | 0.422551316 | 0.968367176 | 0.226051951 |
| 5.498896 | 1.132134974 | 0.435852018 | 0.971767545 | 0.223084792 |
| 5.509136 | 1.120144854 | 0.450769851 | 0.975144982 | 0.221185751 |
| 5.519376 | 1.108140012 | 0.466600481 | 0.976690676 | 0.219955695 |
| 5.529616 | 1.094969531 | 0.484584043 | 0.977135147 | 0.221085827 |
| 5.539856 | 1.081028561 | 0.502708538 | 0.97414772 | 0.223563628 |
| 5.550096 | 1.065826302 | 0.522763888 | 0.968795581 | 0.229261521 |
| 5.560336 | 1.049320261 | 0.543614009 | 0.960084728 | 0.23732554 |
| 5.570576 | 1.031786511 | 0.5660764 | 0.948848546 | 0.247826104 |
| 5.580816 | 1.012385239 | 0.590293559 | 0.934530377 | 0.260878128 |
| 5.591056 | 0.992357636 | 0.6156164 | 0.918207135 | 0.275507743 |
| 5.601296 | 0.970046428 | 0.643304581 | 0.899750939 | 0.291780196 |
| 5.611536 | 0.94629699 | 0.670692597 | 0.880190347 | 0.308213963 |
| 5.621776 | 0.920423754 | 0.699616629 | 0.858837115 | 0.325844296 |
| 5.632016 | 0.893237602 | 0.727372942 | 0.836338942 | 0.341935686 |
| 5.642256 | 0.863266111 | 0.754991414 | 0.812633749 | 0.35830734 |
| 5.652496 | 0.833183477 | 0.78081444 | 0.787416965 | 0.37276145 |
| 5.662736 | 0.801309144 | 0.806287823 | 0.760612931 | 0.386644053 |
| 5.672976 | 0.769534461 | 0.830047941 | 0.732051937 | 0.398631618 |
| 5.683217 | 0.737275983 | 0.85315348 | 0.702517118 | 0.409881547 |
| 5.693457 | 0.705387647 | 0.876116426 | 0.671193808 | 0.419794672 |
| 5.703696 | 0.673580002 | 0.899134883 | 0.638931789 | 0.429660346 |
| 5.713936 | 0.641803789 | 0.922284526 | 0.606109243 | 0.438933696 |
| 5.724176 | 0.610079682 | 0.946888841 | 0.572012146 | 0.448999149 |
| 5.734416 | 0.577818774 | 0.97152076 | 0.537912455 | 0.459519778 |
| 5.744657 | 0.545120236 | 0.997852905 | 0.502848309 | 0.471045808 |
| 5.754897 | 0.512207069 | 1.023665146 | 0.467739514 | 0.48383296 |
| 5.765137 | 0.478788301 | 1.050295143 | 0.431806579 | 0.496733777 |
| 5.775377 | 0.445244302 | 1.075835849 | 0.396392277 | 0.510654968 |
| 5.785617 | 0.411920538 | 1.100345563 | 0.360164164 | 0.523803228 |
| 5.795857 | 0.379155949 | 1.122973566 | 0.324758814 | 0.5371672 |
| 5.806097 | 0.347309873 | 1.143343898 | 0.289049548 | 0.54866466 |
| 5.816337 | 0.316102624 | 1.161700407 | 0.254012545 | 0.559773908 |
| 5.826577 | 0.286187381 | 1.176934011 | 0.219048065 | 0.568503944 |
| 5.836817 | 0.257260561 | 1.190268513 | 0.184204887 | 0.575857781 |
| 5.847057 | 0.228823145 | 1.200749113 | 0.149584498 | 0.581069324 |
| 5.857297 | 0.201391319 | 1.209332045 | 0.114710767 | 0.583829464 |
| 5.867537 | 0.174027542 | 1.215165061 | 0.079487615 | 0.583658526 |
| 5.877777 | 0.147053142 | 1.219706997 | 0.044016023 | 0.580438227 |
| 5.888017 | 0.119541506 | 1.221527494 | 0.008487258 | 0.574312176 |
| 5.898257 | 0.091771126 | 1.222635301 | -0.028213938 | 0.565082572 |
| 5.908497 | 0.063529963 | 1.222459103 | -0.06446318 | 0.553078238 |
| 5.918737 | 0.034697846 | 1.221727628 | -0.101654551 | 0.53872924 |
| 5.928977 | 0.004623676 | 1.22074898 | -0.13873569 | 0.5229241 |
| 5.939217 | -0.025837243 | 1.220207411 | -0.176064093 | 0.506273827 |
| 5.949457 | -0.057218716 | 1.218955553 | -0.212903276 | 0.48842733 |
| 5.959697 | -0.089022882 | 1.218005732 | -0.249149882 | 0.470765039 |
| 5.969937 | -0.12104006 | 1.216271599 | -0.28415576 | 0.45251047 |
| 5.980177 | -0.153348825 | 1.213391853 | -0.317849487 | 0.434669278 |
| 5.990417 | -0.184290811 | 1.209146787 | -0.348799855 | 0.415983991 |
| 6.000657 | -0.215103855 | 1.202772222 | -0.37821354 | 0.397824646 |
| 6.010897 | -0.243977504 | 1.194968092 | -0.404498298 | 0.379640751 |
| 6.021137 | -0.272062972 | 1.184201167 | -0.428857446 | 0.362184952 |
| 6.031377 | -0.298091039 | 1.171401506 | -0.450715714 | 0.344906905 |
| 6.041617 | -0.323394775 | 1.156044295 | -0.470849797 | 0.329799583 |
| 6.051857 | -0.347173492 | 1.138239012 | -0.489433161 | 0.316209164 |
| 6.062098 | -0.370096554 | 1.11778039 | -0.506754277 | 0.304971806 |
| 6.072338 | -0.392166714 | 1.095481879 | -0.523102354 | 0.296874138 |
| 6.082578 | -0.413926139 | 1.071236575 | -0.538765018 | 0.291990505 |
| 6.092818 | -0.434570412 | 1.04657249 | -0.553601467 | 0.290801632 |
| 6.103057 | -0.45551074 | 1.020441019 | -0.567569706 | 0.292486961 |
| 6.113297 | -0.475553205 | 0.995299118 | -0.580785548 | 0.297263441 |
| 6.123538 | -0.494974504 | 0.969998695 | -0.593000811 | 0.303384772 |
| 6.133778 | -0.514329985 | 0.945682562 | -0.604126412 | 0.311167198 |
| 6.144018 | -0.532246945 | 0.920897627 | -0.61431356 | 0.317969377 |
| 6.154258 | -0.549557733 | 0.897105884 | -0.623539685 | 0.32503602 |
| 6.164498 | -0.565672858 | 0.872337232 | -0.632228093 | 0.33011863 |
| 6.174738 | -0.580564508 | 0.848007666 | -0.640450612 | 0.334371356 |
| 6.184978 | -0.594462759 | 0.822612752 | -0.648231315 | 0.336539799 |
| 6.195218 | -0.6077148 | 0.797429863 | -0.656652835 | 0.338590279 |
| 6.205458 | -0.62100252 | 0.772493339 | -0.66501422 | 0.33956209 |
| 6.215698 | -0.633704244 | 0.747866212 | -0.673978495 | 0.339755224 |
| 6.225938 | -0.64735087 | 0.723694302 | -0.683590323 | 0.340443133 |
| 6.236178 | -0.661158169 | 0.700859487 | -0.693502069 | 0.341125782 |
| 6.246418 | -0.675103863 | 0.678169588 | -0.703497546 | 0.341822553 |
| 6.256658 | -0.68869553 | 0.657241481 | -0.713286774 | 0.343551169 |
| 6.266898 | -0.701202817 | 0.636457051 | -0.722335632 | 0.34607258 |
| 6.277138 | -0.711901377 | 0.617828609 | -0.730462262 | 0.349770089 |
| 6.287378 | -0.720493053 | 0.600171576 | -0.736497692 | 0.355019952 |
| 6.297618 | -0.726328588 | 0.584255863 | -0.741641121 | 0.362180164 |
| 6.307858 | -0.729431643 | 0.569251836 | -0.744242328 | 0.371124431 |
| 6.318098 | -0.729929406 | 0.556642088 | -0.745882728 | 0.381492607 |
| 6.328338 | -0.728436703 | 0.544530094 | -0.745574666 | 0.39322909 |
| 6.338578 | -0.725017092 | 0.534446679 | -0.744488273 | 0.406456817 |
| 6.348818 | -0.720614987 | 0.524858094 | -0.742382314 | 0.419704705 |
| 6.359058 | -0.714881706 | 0.517188286 | -0.739600682 | 0.433820198 |
| 6.369298 | -0.70930286 | 0.510114704 | -0.736063763 | 0.448335408 |
| 6.379539 | -0.702877169 | 0.505211337 | -0.731592911 | 0.462736398 |
| 6.389779 | -0.696492657 | 0.501029556 | -0.725956091 | 0.478260238 |
| 6.400019 | -0.689881603 | 0.499209246 | -0.719493569 | 0.493690576 |
| 6.410258 | -0.683459397 | 0.498937188 | -0.71101807 | 0.509952619 |
| 6.420498 | -0.677429924 | 0.499533593 | -0.702135749 | 0.526516398 |
| 6.430738 | -0.671763885 | 0.502469323 | -0.69137057 | 0.543033669 |
| 6.440979 | -0.667146951 | 0.505617804 | -0.680328582 | 0.559004542 |
| 6.451219 | -0.663715457 | 0.509716111 | -0.668099592 | 0.573825442 |
| 6.461459 | -0.661274765 | 0.514413994 | -0.655986808 | 0.586835773 |
| 6.471699 | -0.660202452 | 0.519662943 | -0.642919193 | 0.597387415 |
| 6.481939 | -0.65928215 | 0.524736421 | -0.63004549 | 0.605494511 |
| 6.492179 | -0.658925374 | 0.530822083 | -0.616472358 | 0.610469039 |
| 6.502419 | -0.657343458 | 0.535868719 | -0.602550823 | 0.612963721 |
| 6.512659 | -0.654289991 | 0.541762456 | -0.587630161 | 0.612767504 |
| 6.522899 | -0.648977654 | 0.546213266 | -0.572296195 | 0.610590727 |
| 6.533139 | -0.640329389 | 0.550373132 | -0.555522091 | 0.607080988 |
| 6.543379 | -0.629022452 | 0.553529781 | -0.537515106 | 0.602027533 |
| 6.553619 | -0.614177557 | 0.55560294 | -0.518210132 | 0.595970163 |
| 6.563859 | -0.59710273 | 0.55734629 | -0.497321915 | 0.589166927 |
| 6.574099 | -0.577892991 | 0.558695165 | -0.475889483 | 0.581714076 |
| 6.584339 | -0.557720803 | 0.560437481 | -0.453181473 | 0.574077463 |
| 6.594579 | -0.537556791 | 0.562608281 | -0.431025113 | 0.565456185 |
| 6.604819 | -0.518673218 | 0.565801181 | -0.409141392 | 0.55676907 |
| 6.615059 | -0.501279128 | 0.56986171 | -0.388629894 | 0.54743416 |
| 6.625299 | -0.48623321 | 0.574947829 | -0.370274961 | 0.538235351 |
| 6.635539 | -0.474033646 | 0.579995424 | -0.353794111 | 0.527570455 |
| 6.645779 | -0.463867401 | 0.585450932 | -0.339493392 | 0.516670799 |
| 6.656019 | -0.456250207 | 0.590354857 | -0.327346572 | 0.504358049 |
| 6.666259 | -0.4496705 | 0.594349537 | -0.316911078 | 0.490783618 |
| 6.676499 | -0.444253923 | 0.59795286 | -0.307122784 | 0.475534972 |
| 6.686739 | -0.43888283 | 0.599826287 | -0.298487523 | 0.458113046 |
| 6.69698 | -0.433203648 | 0.601451405 | -0.289702156 | 0.439021275 |
| 6.707219 | -0.427027833 | 0.601609295 | -0.280919214 | 0.417461073 |
| 6.717459 | -0.419873341 | 0.601637582 | -0.271881755 | 0.394237101 |
| 6.727699 | -0.411507229 | 0.600952527 | -0.262141951 | 0.370138341 |
| 6.737939 | -0.402754123 | 0.60062171 | -0.252072026 | 0.344965137 |
| 6.748179 | -0.392434955 | 0.600485134 | -0.241253765 | 0.320235104 |
| 6.75842 | -0.381923792 | 0.601705182 | -0.230178785 | 0.296847786 |
| 6.76866 | -0.370512387 | 0.603712247 | -0.218470794 | 0.275522349 |
| 6.7789 | -0.358795558 | 0.607183328 | -0.206069207 | 0.256800954 |
| 6.78914 | -0.346656063 | 0.611986207 | -0.193464648 | 0.24203087 |
| 6.79938 | -0.335031115 | 0.617721317 | -0.179804787 | 0.230356963 |
| 6.809619 | -0.322965896 | 0.623773362 | -0.165544613 | 0.223161586 |
| 6.81986 | -0.311530789 | 0.630018232 | -0.150135429 | 0.219741118 |
| 6.8301 | -0.300304879 | 0.635311221 | -0.133733459 | 0.220136395 |
| 6.84034 | -0.289508334 | 0.639694327 | -0.11613615 | 0.223620403 |
| 6.85058 | -0.279281954 | 0.64246211 | -0.097193232 | 0.229633732 |
| 6.86082 | -0.268848307 | 0.643531377 | -0.077975548 | 0.237537257 |
| 6.87106 | -0.25887689 | 0.642583164 | -0.05815946 | 0.246499341 |
| 6.8813 | -0.248686197 | 0.640265375 | -0.038784335 | 0.256568946 |
| 6.89154 | -0.237652952 | 0.636438269 | -0.020084197 | 0.267065639 |
| 6.90178 | -0.225674224 | 0.630996237 | -0.003490794 | 0.278780178 |
| 6.91202 | -0.212309631 | 0.624980321 | 0.011397504 | 0.291272171 |
| 6.92226 | -0.197259081 | 0.618217904 | 0.023842614 | 0.305215491 |
| 6.9325 | -0.180725957 | 0.61087971 | 0.033714997 | 0.319616929 |
| 6.94274 | -0.162255103 | 0.603093918 | 0.041569459 | 0.335508977 |
| 6.95298 | -0.14291224 | 0.59523034 | 0.047045296 | 0.350856175 |
| 6.96322 | -0.122596893 | 0.587331915 | 0.051625541 | 0.36672638 |
| 6.97346 | -0.102066124 | 0.57913167 | 0.054437456 | 0.381513294 |
| 6.9837 | -0.08134647 | 0.571609361 | 0.057232669 | 0.395922444 |
| 6.99394 | -0.060681331 | 0.563759537 | 0.059565925 | 0.408531748 |
| 7.00418 | -0.04030171 | 0.5564877 | 0.062099239 | 0.420673866 |
| 7.01442 | -0.019685157 | 0.549134207 | 0.064631837 | 0.431396887 |
| 7.02466 | 0.001013438 | 0.541603457 | 0.067581975 | 0.440872614 |
| 7.0349 | 0.021535084 | 0.534294618 | 0.070506809 | 0.449183615 |
| 7.04514 | 0.042581047 | 0.52536592 | 0.073859136 | 0.456077526 |
| 7.05538 | 0.062959984 | 0.517215401 | 0.077122219 | 0.46158258 |
| 7.06562 | 0.082767785 | 0.507418362 | 0.080928458 | 0.465161207 |
| 7.075861 | 0.101853117 | 0.498133201 | 0.084711415 | 0.467699951 |
| 7.086101 | 0.119313518 | 0.488380763 | 0.089217344 | 0.468968656 |
| 7.096341 | 0.134604981 | 0.479330673 | 0.094419517 | 0.468750415 |
| 7.106581 | 0.148032834 | 0.470762844 | 0.099885469 | 0.468198245 |
| 7.11682 | 0.158970187 | 0.463420534 | 0.106783871 | 0.467258853 |
| 7.12706 | 0.167843044 | 0.45658192 | 0.113462215 | 0.465902092 |
| 7.1373 | 0.174411791 | 0.450805753 | 0.12147748 | 0.464646078 |
| 7.147541 | 0.179500687 | 0.445160503 | 0.128735924 | 0.463003022 |
| 7.157781 | 0.183416106 | 0.439784743 | 0.136311316 | 0.461562448 |

| | | | | |
|---|---|---|---|---|
| 4.761614 | -0.102013127 | 0.186716265 | 0.054178005 | 0.269220929 |
| 4.771854 | -0.111474722 | 0.189585472 | 0.052721737 | 0.265631468 |
| 4.782094 | -0.120950602 | 0.192533765 | 0.051167989 | 0.263250993 |
| 4.792334 | -0.130612046 | 0.195551804 | 0.049164096 | 0.260090021 |
| 4.802574 | -0.139735167 | 0.198660312 | 0.047257738 | 0.257363129 |
| 4.812814 | -0.148771613 | 0.201838434 | 0.044550256 | 0.253581253 |
| 4.823054 | -0.15711514 | 0.204887228 | 0.040998522 | 0.249601855 |
| 4.833294 | -0.164733095 | 0.207482331 | 0.036661547 | 0.244570632 |
| 4.843534 | -0.172027915 | 0.209917148 | 0.031044908 | 0.239260318 |
| 4.853774 | -0.178235229 | 0.211749501 | 0.024284893 | 0.23329182 |
| 4.864014 | -0.183423984 | 0.213087855 | 0.016496861 | 0.22743241 |
| 4.874254 | -0.187364399 | 0.214003142 | 0.007874398 | 0.221186322 |
| 4.884494 | -0.189876322 | 0.21391229 | -0.00123189 | 0.215246623 |
| 4.894734 | -0.190896066 | 0.213426929 | -0.010398473 | 0.209298611 |
| 4.904974 | -0.190377373 | 0.21194325 | -0.019353289 | 0.203918138 |
| 4.915214 | -0.188593996 | 0.209846601 | -0.027098874 | 0.19879918 |
| 4.925454 | -0.185776387 | 0.207000859 | -0.033783415 | 0.194709988 |
| 4.935694 | -0.182187495 | 0.203462626 | -0.038871682 | 0.191197599 |
| 4.945934 | -0.177902386 | 0.199404312 | -0.042202842 | 0.189276617 |
| 4.956174 | -0.174193304 | 0.194648779 | -0.043765183 | 0.187953065 |
| 4.966414 | -0.169682567 | 0.189590659 | -0.043685241 | 0.18785831 |
| 4.976655 | -0.165806877 | 0.184083614 | -0.04217015 | 0.188540558 |
| 4.986895 | -0.16170962 | 0.178088457 | -0.039296201 | 0.189479671 |
| 4.997134 | -0.157454941 | 0.172247638 | -0.035534084 | 0.190901989 |
| 5.007374 | -0.152837884 | 0.165882374 | -0.031304706 | 0.192413775 |
| 5.017614 | -0.147821858 | 0.159997177 | -0.026327883 | 0.193930932 |
| 5.027854 | -0.141791256 | 0.154068262 | -0.021204965 | 0.195508126 |
| 5.038095 | -0.135226393 | 0.148553741 | -0.016073732 | 0.197242509 |
| 5.048335 | -0.128132157 | 0.14410571 | -0.010840873 | 0.199265607 |
| 5.058575 | -0.120012086 | 0.139756296 | -0.005450177 | 0.200967891 |
| 5.068815 | -0.112094545 | 0.137575648 | -8.3313E-05 | 0.203030562 |
| 5.079055 | -0.103559358 | 0.136176524 | 0.005380286 | 0.204320141 |
| 5.089295 | -0.095223539 | 0.137183409 | 0.01134088 | 0.205774736 |
| 5.099535 | -0.086935093 | 0.13898633 | 0.017418997 | 0.206547103 |
| 5.109775 | -0.078776807 | 0.143367631 | 0.02399111 | 0.206948137 |
| 5.120015 | -0.070603437 | 0.147323347 | 0.030919887 | 0.207129403 |
| 5.130255 | -0.062584639 | 0.152656124 | 0.038160659 | 0.207467506 |
| 5.140495 | -0.054379642 | 0.157201559 | 0.045231518 | 0.20724652 |
| 5.150735 | -0.045843812 | 0.161133862 | 0.05254908 | 0.207480547 |
| 5.160975 | -0.037088907 | 0.164618229 | 0.05977068 | 0.207549794 |
| 5.171215 | -0.02792377 | 0.166692228 | 0.066262323 | 0.207555696 |
| 5.181455 | -0.017929967 | 0.168686222 | 0.073210587 | 0.207550832 |
| 5.191695 | -0.007547199 | 0.169399702 | 0.07934676 | 0.207108148 |
| 5.201935 | 0.00374897 | 0.170210029 | 0.085863744 | 0.206227689 |
| 5.212175 | 0.015591137 | 0.170430861 | 0.092546201 | 0.204399757 |
| 5.222415 | 0.02845735 | 0.170430808 | 0.099792771 | 0.201850748 |
| 5.232655 | 0.041725246 | 0.170904689 | 0.107732032 | 0.198087303 |
| 5.242895 | 0.055736179 | 0.171494994 | 0.116444702 | 0.19370089 |
| 5.253135 | 0.069667566 | 0.173477715 | 0.126079444 | 0.188227177 |
| 5.263375 | 0.084163566 | 0.176915067 | 0.136608151 | 0.183021469 |
| 5.273615 | 0.098169335 | 0.182238896 | 0.147771556 | 0.17809583 |
| 5.283855 | 0.112156154 | 0.189355031 | 0.159419346 | 0.173588854 |
| 5.294095 | 0.125631809 | 0.197992939 | 0.171055943 | 0.170002597 |
| 5.304335 | 0.139060589 | 0.207712223 | 0.182738757 | 0.16723153 |
| 5.314575 | 0.15210699 | 0.217349247 | 0.194060458 | 0.165183381 |
| 5.324815 | 0.165210069 | 0.226122217 | 0.20460791 | 0.163638122 |
| 5.335055 | 0.178239702 | 0.234007856 | 0.214814226 | 0.162793623 |
| 5.345295 | 0.191797267 | 0.239966417 | 0.224040023 | 0.162276288 |
| 5.355536 | 0.205150672 | 0.244596338 | 0.23252016 | 0.162879856 |
| 5.365776 | 0.218978757 | 0.247425613 | 0.240151472 | 0.163911658 |
| 5.376016 | 0.23251275 | 0.249434967 | 0.24688795 | 0.165853198 |
| 5.386256 | 0.246083468 | 0.250337951 | 0.252679018 | 0.168547725 |
| 5.396495 | 0.258798785 | 0.250852299 | 0.257264258 | 0.17172338 |
| 5.406735 | 0.27146668 | 0.250440798 | 0.261429826 | 0.175369339 |
| 5.416975 | 0.283506317 | 0.250436595 | 0.264303695 | 0.179451295 |
| 5.427216 | 0.295182757 | 0.249692532 | 0.266875645 | 0.183226396 |
| 5.437456 | 0.306631118 | 0.249687023 | 0.269242205 | 0.186991083 |
| 5.447696 | 0.318226156 | 0.249740738 | 0.271505482 | 0.190071435 |
| 5.457936 | 0.329857081 | 0.251270811 | 0.274519592 | 0.192185 |
| 5.468176 | 0.341911545 | 0.25351992 | 0.277685707 | 0.194004436 |
| 5.478416 | 0.353577429 | 0.258082073 | 0.28179494 | 0.194489926 |
| 5.488656 | 0.365084764 | 0.262862705 | 0.286612645 | 0.194840071 |
| 5.498896 | 0.375896996 | 0.269607496 | 0.291515357 | 0.194405382 |
| 5.509136 | 0.385796741 | 0.276501985 | 0.297212224 | 0.19407659 |
| 5.519376 | 0.394568308 | 0.283396706 | 0.302743729 | 0.192905305 |
| 5.529616 | 0.4021971 | 0.289685917 | 0.308132258 | 0.191706616 |
| 5.539856 | 0.40860911 | 0.295033595 | 0.312918325 | 0.190022175 |
| 5.550096 | 0.414186214 | 0.299100811 | 0.317851438 | 0.187810009 |
| 5.560336 | 0.41892058 | 0.301806738 | 0.321786546 | 0.185410855 |
| 5.570576 | 0.423016948 | 0.302844273 | 0.325208972 | 0.182939096 |
| 5.580816 | 0.42668255 | 0.302647956 | 0.328344156 | 0.180832023 |
| 5.591056 | 0.429638541 | 0.300905369 | 0.330909646 | 0.179226232 |
| 5.601296 | 0.432492676 | 0.297989263 | 0.332696991 | 0.179016036 |
| 5.611536 | 0.434929521 | 0.293842181 | 0.334131596 | 0.179728704 |
| 5.621776 | 0.436633654 | 0.289268132 | 0.334791199 | 0.182044435 |
| 5.632016 | 0.438591933 | 0.283595075 | 0.334789384 | 0.185744989 |
| 5.642256 | 0.440188299 | 0.277893439 | 0.334458196 | 0.190334849 |
| 5.652496 | 0.441465503 | 0.272361208 | 0.333348822 | 0.195941825 |
| 5.662736 | 0.443370847 | 0.267103993 | 0.332690396 | 0.202038295 |
| 5.672976 | 0.445406654 | 0.263008524 | 0.331731624 | 0.208298611 |
| 5.683217 | 0.447856948 | 0.259845628 | 0.331871752 | 0.214336986 |
| 5.693457 | 0.450877512 | 0.258379649 | 0.332776747 | 0.220264875 |
| 5.703696 | 0.454399021 | 0.258270708 | 0.334550806 | 0.225534147 |
| 5.713936 | 0.458085853 | 0.259970268 | 0.337456153 | 0.230148481 |
| 5.724176 | 0.461636597 | 0.263097215 | 0.34097318 | 0.234350782 |
| 5.734416 | 0.465026274 | 0.267121091 | 0.345075594 | 0.237893759 |
| 5.744657 | 0.467626469 | 0.272126324 | 0.348915802 | 0.241068098 |
| 5.754897 | 0.469283155 | 0.276981454 | 0.352617625 | 0.243908857 |
| 5.765137 | 0.469721487 | 0.281800438 | 0.355372828 | 0.246576913 |
| 5.775377 | 0.469407617 | 0.285490267 | 0.357471241 | 0.248992177 |
| 5.785617 | 0.46752933 | 0.288550481 | 0.358425104 | 0.251313891 |
| 5.795857 | 0.465166558 | 0.290135357 | 0.358954009 | 0.253141451 |
| 5.806097 | 0.461639018 | 0.290737083 | 0.358652116 | 0.254855861 |
| 5.816337 | 0.457966414 | 0.290217012 | 0.357863301 | 0.256528064 |
| 5.826577 | 0.453966959 | 0.28936897 | 0.356959599 | 0.257294558 |
| 5.836817 | 0.450172572 | 0.287751787 | 0.355720329 | 0.258814128 |
| 5.847057 | 0.446359293 | 0.286518856 | 0.354305708 | 0.259514464 |
| 5.857297 | 0.443165542 | 0.285377656 | 0.352567442 | 0.260806353 |
| 5.867537 | 0.439865789 | 0.284782126 | 0.350307368 | 0.261530163 |
| 5.877777 | 0.436615878 | 0.284710331 | 0.347722337 | 0.26240487 |
| 5.888017 | 0.432888359 | 0.284640496 | 0.343984156 | 0.263223165 |
| 5.898257 | 0.428508809 | 0.284634356 | 0.33949405 | 0.263800479 |
| 5.908497 | 0.422455935 | 0.283890733 | 0.334523562 | 0.264551246 |
| 5.918737 | 0.415976177 | 0.282538602 | 0.328451519 | 0.265376079 |
| 5.928977 | 0.407873893 | 0.280233245 | 0.322315862 | 0.266792681 |
| 5.939217 | 0.399333499 | 0.277415724 | 0.316110087 | 0.268722162 |
| 5.949457 | 0.390224457 | 0.273439235 | 0.309960759 | 0.271007011 |
| 5.959697 | 0.381371425 | 0.269723754 | 0.304851522 | 0.273933474 |
| 5.969937 | 0.372482157 | 0.265377585 | 0.300036615 | 0.276270179 |
| 5.980177 | 0.364402677 | 0.260995626 | 0.296530853 | 0.278760284 |
| 5.990417 | 0.356293419 | 0.256852696 | 0.293264361 | 0.280005227 |
| 6.000657 | 0.348781592 | 0.252198841 | 0.290726975 | 0.280649754 |
| 6.010897 | 0.340768988 | 0.247600416 | 0.288311071 | 0.279984608 |
| 6.021137 | 0.33277549 | 0.242856115 | 0.285603682 | 0.278186403 |
| 6.031377 | 0.323865281 | 0.23812373 | 0.282719157 | 0.275192726 |
| 6.041617 | 0.314827454 | 0.233517591 | 0.278689192 | 0.271222531 |
| 6.051857 | 0.304481594 | 0.229140107 | 0.273885101 | 0.266204414 |
| 6.062098 | 0.294062597 | 0.225838957 | 0.26811158 | 0.260630902 |
| 6.072338 | 0.282903022 | 0.223109499 | 0.260728816 | 0.254615611 |
| 6.082578 | 0.271700499 | 0.221221785 | 0.252456836 | 0.248412794 |
| 6.092818 | 0.259784868 | 0.22075823 | 0.242822724 | 0.242900731 |
| 6.103057 | 0.248288081 | 0.221289601 | 0.231852768 | 0.237343348 |
| 6.113297 | 0.236726509 | 0.223153193 | 0.219937305 | 0.232900635 |
| 6.123538 | 0.224769705 | 0.225788058 | 0.206964808 | 0.228864513 |
| 6.133778 | 0.213055117 | 0.229772565 | 0.193271179 | 0.225225227 |
| 6.144018 | 0.201202701 | 0.234497782 | 0.17896534 | 0.222160109 |
| 6.154258 | 0.188824088 | 0.239858008 | 0.164464931 | 0.219291329 |
| 6.164498 | 0.176571455 | 0.24629668 | 0.149620892 | 0.216230875 |
| 6.174738 | 0.163744226 | 0.252944668 | 0.134916461 | 0.213876469 |
| 6.184978 | 0.15069248 | 0.260173051 | 0.120119954 | 0.211684597 |
| 6.195218 | 0.137346875 | 0.267161033 | 0.105554029 | 0.210649195 |
| 6.205458 | 0.123675747 | 0.274149293 | 0.090988074 | 0.210454365 |
| 6.215698 | 0.110324961 | 0.2800236 | 0.076578942 | 0.211808085 |
| 6.225938 | 0.096883799 | 0.284896395 | 0.062595871 | 0.215072135 |
| 6.236178 | 0.084016871 | 0.288219829 | 0.048329176 | 0.219244652 |
| 6.246418 | 0.071641378 | 0.29037257 | 0.034653214 | 0.225453584 |
| 6.256658 | 0.059805252 | 0.291080146 | 0.020842634 | 0.232377694 |
| 6.266898 | 0.048810996 | 0.29116097 | 0.007549264 | 0.239962322 |
| 6.277138 | 0.037712011 | 0.290860065 | -0.005920346 | 0.247639464 |
| 6.287378 | 0.027100118 | 0.290423256 | -0.019034487 | 0.255128862 |
| 6.297618 | 0.015768616 | 0.290696122 | -0.032431158 | 0.26243751 |
| 6.307858 | 0.004036678 | 0.290974156 | -0.045516848 | 0.268307987 |
| 6.318098 | -0.008591896 | 0.292642761 | -0.058832994 | 0.273377617 |
| 6.328338 | -0.022097324 | 0.29412527 | -0.072376213 | 0.276953157 |
| 6.338578 | -0.03684635 | 0.296436352 | -0.085974305 | 0.278819749 |
| 6.348818 | -0.051542239 | 0.298477585 | -0.099669885 | 0.278498399 |
| 6.359058 | -0.067125003 | 0.301060358 | -0.113821488 | 0.276556612 |
| 6.369298 | -0.082292088 | 0.302696105 | -0.128128084 | 0.273134574 |
| 6.379539 | -0.097245081 | 0.304406567 | -0.143008062 | 0.268427043 |
| 6.389779 | -0.111019934 | 0.305505309 | -0.157994097 | 0.263760917 |
| 6.400019 | -0.124039237 | 0.306379638 | -0.173084198 | 0.259461594 |
| 6.410258 | -0.135466544 | 0.307061392 | -0.18835742 | 0.256077945 |
| 6.420498 | -0.145485358 | 0.307404337 | -0.202988251 | 0.254397345 |
| 6.430738 | -0.154190694 | 0.307952938 | -0.217551283 | 0.25396231 |
| 6.440979 | -0.161457392 | 0.30787978 | -0.23099087 | 0.255073962 |
| 6.451219 | -0.167688589 | 0.308114232 | -0.24384953 | 0.25711189 |
| 6.461459 | -0.172873139 | 0.30729505 | -0.256315988 | 0.260392917 |
| 6.471699 | -0.177519542 | 0.306617028 | -0.267649231 | 0.263826177 |
| 6.481939 | -0.181688452 | 0.304978297 | -0.27933132 | 0.268272133 |
| 6.492179 | -0.185508365 | 0.303418665 | -0.290359638 | 0.272430285 |
| 6.502419 | -0.189557889 | 0.301514188 | -0.301875999 | 0.277141493 |
| 6.512659 | -0.193369494 | 0.29968227 | -0.313128479 | 0.281793815 |
| 6.522899 | -0.197479429 | 0.297819265 | -0.324774989 | 0.28619569 |
| 6.533139 | -0.201564594 | 0.296532833 | -0.336235063 | 0.2900998 |
| 6.543379 | -0.206261458 | 0.295354063 | -0.347700647 | 0.293852724 |
| 6.553619 | -0.21055947 | 0.294268313 | -0.358745957 | 0.296530352 |
| 6.563859 | -0.215215227 | 0.293699268 | -0.370051635 | 0.297994271 |
| 6.574099 | -0.219609392 | 0.292542884 | -0.380580567 | 0.298815318 |
| 6.584339 | -0.223908839 | 0.292182555 | -0.391138505 | 0.298081277 |
| 6.594579 | -0.227551834 | 0.291418346 | -0.401155502 | 0.296539472 |
| 6.604819 | -0.230942702 | 0.29105503 | -0.411080058 | 0.294111587 |
| 6.615059 | -0.233432422 | 0.290702535 | -0.419731822 | 0.290961717 |
| 6.625299 | -0.23577973 | 0.290337116 | -0.428076347 | 0.287587531 |
| 6.635539 | -0.23737704 | 0.28998659 | -0.434605797 | 0.284168538 |
| 6.645779 | -0.239096805 | 0.289125561 | -0.44027351 | 0.280825657 |
| 6.656019 | -0.240661011 | 0.288006722 | -0.443835532 | 0.277509793 |
| 6.666259 | -0.24274517 | 0.286540139 | -0.445863707 | 0.274343044 |
| 6.676499 | -0.245078644 | 0.284334111 | -0.446029796 | 0.271022092 |
| 6.686739 | -0.247777742 | 0.28189153 | -0.444537533 | 0.267610059 |
| 6.69698 | -0.250855828 | 0.279459419 | -0.441257377 | 0.263781448 |
| 6.707219 | -0.2537413 | 0.276457605 | -0.436541429 | 0.259945476 |
| 6.717459 | -0.256265041 | 0.274442575 | -0.430256878 | 0.256378262 |
| 6.727699 | -0.258002311 | 0.272501372 | -0.422529574 | 0.253375 |
| 6.737939 | -0.258642178 | 0.27135272 | -0.413457751 | 0.251107431 |
| 6.748179 | -0.257709995 | 0.271296235 | -0.403358305 | 0.250351203 |
| 6.75842 | -0.25540465 | 0.272452927 | -0.392151872 | 0.250778834 |
| 6.76866 | -0.251565716 | 0.2744037 | -0.380406076 | 0.252624448 |
| 6.7789 | -0.246775576 | 0.278101386 | -0.368269934 | 0.25530563 |
| 6.78914 | -0.240741963 | 0.282689312 | -0.356286306 | 0.259199331 |
| 6.79938 | -0.234436927 | 0.288825912 | -0.344632076 | 0.262876573 |
| 6.809619 | -0.227993036 | 0.29599843 | -0.333499898 | 0.266863243 |
| 6.81986 | -0.221504162 | 0.303470049 | -0.323161017 | 0.270479502 |
| 6.8301 | -0.215299199 | 0.31174862 | -0.314015795 | 0.273251716 |
| 6.84034 | -0.209861491 | 0.319274807 | -0.305722574 | 0.27580848 |
| 6.85058 | -0.204333547 | 0.325656368 | -0.298536882 | 0.277338894 |
| 6.86082 | -0.199594847 | 0.330285147 | -0.292540247 | 0.278397975 |
| 6.87106 | -0.19492101 | 0.332583034 | -0.287608175 | 0.278538442 |
| 6.8813 | -0.189909446 | 0.332104749 | -0.283210903 | 0.278542998 |
| 6.89154 | -0.185196444 | 0.329389859 | -0.280022235 | 0.277681141 |
| 6.90178 | -0.179643913 | 0.324342607 | -0.276442953 | 0.276475955 |
| 6.91202 | -0.17398726 | 0.318283738 | -0.273563039 | 0.275295402 |
| 6.92226 | -0.167758375 | 0.311216981 | -0.270027014 | 0.274037967 |
| 6.9325 | -0.161370017 | 0.304787236 | -0.266148692 | 0.273010953 |
| 6.94274 | -0.154920778 | 0.298901142 | -0.26177576 | 0.272538143 |
| 6.95298 | -0.148456254 | 0.294538054 | -0.256580278 | 0.272074778 |
| 6.96322 | -0.142642094 | 0.2916959 | -0.250944201 | 0.272357688 |
| 6.97346 | -0.136851959 | 0.29017405 | -0.244224955 | 0.272667477 |
| 6.9837 | -0.131972121 | 0.290576164 | -0.237041168 | 0.272763269 |
| 6.99394 | -0.126792001 | 0.292051356 | -0.228551438 | 0.272521781 |
| 7.00418 | -0.122092601 | 0.294402841 | -0.219561063 | 0.272171975 |
| 7.01442 | -0.116967289 | 0.297670806 | -0.209384712 | 0.270920295 |
| 7.02466 | -0.111832008 | 0.301187437 | -0.198144562 | 0.269332965 |
| 7.0349 | -0.105897144 | 0.304630779 | -0.186305022 | 0.267040113 |
| 7.04514 | -0.100501708 | 0.308258135 | -0.173732363 | 0.264442569 |
| 7.05538 | -0.094743592 | 0.311560776 | -0.160880924 | 0.261249593 |
| 7.06562 | -0.089582411 | 0.314953434 | -0.147780623 | 0.258055835 |
| 7.075861 | -0.085580424 | 0.318039327 | -0.134775003 | 0.254631527 |
| 7.086101 | -0.082453986 | 0.321724244 | -0.122241161 | 0.251770915 |
| 7.096341 | -0.080751092 | 0.325003976 | -0.109654517 | 0.248851802 |
| 7.106581 | -0.080144931 | 0.328705494 | -0.097789685 | 0.246835177 |
| 7.11682 | -0.080619117 | 0.331745171 | -0.086052692 | 0.245678243 |
| 7.12706 | -0.08145681 | 0.334999051 | -0.074710056 | 0.245100186 |
| 7.1373 | -0.082713656 | 0.336908181 | -0.063428016 | 0.245223978 |
| 7.147541 | -0.083391081 | 0.338559325 | -0.0522104 | 0.245947056 |
| 7.157781 | -0.083363738 | 0.33883719 | -0.040830393 | 0.246761754 |

| | | | | |
|---|---|---|---|---|
| 4.761614 | -0.026518595 | 0.253503808 | 0.036669853 | 0.170830621 |
| 4.771854 | -0.016037294 | 0.250163544 | 0.039714036 | 0.171779577 |
| 4.782094 | -0.004377454 | 0.250086782 | 0.04496587 | 0.173940521 |
| 4.792334 | 0.007637114 | 0.252199678 | 0.051517454 | 0.177913952 |
| 4.802574 | 0.020689624 | 0.256441379 | 0.060227071 | 0.181633117 |
| 4.812814 | 0.033999611 | 0.261890054 | 0.070535251 | 0.186141907 |
| 4.823054 | 0.047908598 | 0.268475465 | 0.083026767 | 0.188963184 |
| 4.833294 | 0.061606256 | 0.274762491 | 0.096572317 | 0.191456916 |
| 4.843534 | 0.075716066 | 0.281183319 | 0.111387081 | 0.192020397 |
| 4.853774 | 0.089446701 | 0.286354783 | 0.126982844 | 0.192178387 |
| 4.864014 | 0.103406068 | 0.290893472 | 0.142626468 | 0.191228671 |
| 4.874254 | 0.117268558 | 0.29420425 | 0.15849783 | 0.190456625 |
| 4.884494 | 0.131577606 | 0.297036818 | 0.174514938 | 0.190074801 |
| 4.894734 | 0.146532443 | 0.299615816 | 0.190471037 | 0.190495839 |
| 4.904974 | 0.162240095 | 0.302923819 | 0.20686539 | 0.19177582 |
| 4.915214 | 0.179195841 | 0.30673253 | 0.22360588 | 0.194164087 |
| 4.925454 | 0.197332758 | 0.311606634 | 0.241076477 | 0.197196419 |
| 4.935694 | 0.217004356 | 0.31704014 | 0.25947869 | 0.200622906 |
| 4.945934 | 0.237657431 | 0.322839262 | 0.278530283 | 0.203630257 |
| 4.956174 | 0.259862506 | 0.328693222 | 0.298616266 | 0.20644271 |
| 4.966414 | 0.2824871 | 0.333893616 | 0.319026331 | 0.207865197 |
| 4.976655 | 0.305861804 | 0.338555082 | 0.339853998 | 0.20831187 |
| 4.986895 | 0.329315085 | 0.342674673 | 0.360746178 | 0.207532835 |
| 4.997134 | 0.352437564 | 0.346216074 | 0.38130146 | 0.205783746 |
| 5.007374 | 0.374937454 | 0.349536182 | 0.401202852 | 0.202895763 |
| 5.017614 | 0.396582646 | 0.353528525 | 0.420446409 | 0.199289301 |
| 5.027854 | 0.417042133 | 0.357382416 | 0.438712483 | 0.195103867 |
| 5.038095 | 0.436303313 | 0.362408116 | 0.455836238 | 0.19017282 |
| 5.048335 | 0.454703903 | 0.3679143 | 0.472071273 | 0.184450806 |
| 5.058575 | 0.472008041 | 0.374741103 | 0.487187728 | 0.178115251 |
| 5.068815 | 0.488790695 | 0.381841461 | 0.50171019 | 0.171398361 |
| 5.079055 | 0.504883783 | 0.390228971 | 0.515123451 | 0.164214793 |
| 5.089295 | 0.520996749 | 0.398606182 | 0.528489411 | 0.157016402 |
| 5.099535 | 0.536624838 | 0.408049902 | 0.541104297 | 0.150392677 |
| 5.109775 | 0.552268969 | 0.417458837 | 0.553555307 | 0.143800475 |
| 5.120015 | 0.567894875 | 0.426664334 | 0.565866716 | 0.138873681 |
| 5.130255 | 0.583075241 | 0.435454281 | 0.578311952 | 0.134643775 |
| 5.140495 | 0.598273934 | 0.443343921 | 0.590192907 | 0.131982121 |
| 5.150735 | 0.613264077 | 0.450113493 | 0.602382677 | 0.130583493 |
| 5.160975 | 0.628187437 | 0.455426179 | 0.613997288 | 0.130702057 |
| 5.171215 | 0.643121009 | 0.45923107 | 0.625715735 | 0.131643644 |
| 5.181455 | 0.658295451 | 0.461272708 | 0.637226097 | 0.133454332 |
| 5.191695 | 0.674092378 | 0.462096246 | 0.648548988 | 0.135807105 |
| 5.201935 | 0.690762227 | 0.461273425 | 0.660270353 | 0.138409911 |
| 5.212175 | 0.708037735 | 0.459414017 | 0.672440181 | 0.140873437 |
| 5.222415 | 0.726588036 | 0.456715678 | 0.684773463 | 0.143137349 |
| 5.232655 | 0.745936337 | 0.453558048 | 0.697873257 | 0.145147386 |
| 5.242895 | 0.766113108 | 0.450108012 | 0.71130681 | 0.146931014 |
| 5.253135 | 0.786331339 | 0.44682346 | 0.725340416 | 0.148547706 |
| 5.263375 | 0.806683531 | 0.443676748 | 0.738929195 | 0.150658599 |
| 5.273615 | 0.826616444 | 0.440834725 | 0.752544515 | 0.153353544 |
| 5.283855 | 0.84573614 | 0.4381862 | 0.765937751 | 0.157036324 |
| 5.294095 | 0.863879906 | 0.436224817 | 0.778416822 | 0.162348184 |
| 5.304335 | 0.881143534 | 0.434379732 | 0.790244882 | 0.168539571 |
| 5.314575 | 0.897356108 | 0.433092059 | 0.801214802 | 0.176817156 |
| 5.324815 | 0.912521385 | 0.43220989 | 0.811808402 | 0.1857586 |
| 5.335055 | 0.927016497 | 0.432858265 | 0.821596031 | 0.19609925 |
| 5.345295 | 0.940769859 | 0.433418953 | 0.830966455 | 0.206740724 |
| 5.355536 | 0.95370065 | 0.435701306 | 0.840973695 | 0.218574687 |
| 5.365776 | 0.965950423 | 0.438215464 | 0.850918936 | 0.230132686 |
| 5.376016 | 0.9780314 | 0.441795522 | 0.861674331 | 0.242406566 |
| 5.386256 | 0.989398107 | 0.44516956 | 0.873660803 | 0.254462284 |
| 5.396495 | 1.000229758 | 0.44839019 | 0.886728843 | 0.266317204 |
| 5.406735 | 1.010982493 | 0.450721671 | 0.900896966 | 0.278049615 |
| 5.416975 | 1.021050861 | 0.451827416 | 0.915747677 | 0.288813039 |
| 5.427216 | 1.030260301 | 0.451400272 | 0.931592281 | 0.298729158 |
| 5.437456 | 1.038848266 | 0.449346028 | 0.946935877 | 0.30759695 |
| 5.447696 | 1.045817146 | 0.445933169 | 0.962318252 | 0.315249987 |
| 5.457936 | 1.051303533 | 0.440587136 | 0.976366589 | 0.321396851 |
| 5.468176 | 1.054917929 | 0.434309586 | 0.989287918 | 0.326524736 |
| 5.478416 | 1.056769755 | 0.426936855 | 1.000373691 | 0.330029567 |
| 5.488656 | 1.056551334 | 0.418498334 | 1.009737641 | 0.33303033 |
| 5.498896 | 1.054622632 | 0.409084612 | 1.01689523 | 0.334731207 |
| 5.509136 | 1.051128098 | 0.399634589 | 1.022353033 | 0.336460052 |
| 5.519376 | 1.046345901 | 0.389080691 | 1.025453111 | 0.337807088 |
| 5.529616 | 1.040592977 | 0.379219418 | 1.027129998 | 0.339227941 |
| 5.539856 | 1.033985528 | 0.36907715 | 1.02731969 | 0.340667083 |
| 5.550096 | 1.026349652 | 0.359472338 | 1.025920694 | 0.342569598 |
| 5.560336 | 1.018881452 | 0.351249839 | 1.023608406 | 0.344264651 |
| 5.570576 | 1.010330218 | 0.343593483 | 1.02032649 | 0.346370246 |
| 5.580816 | 1.001735455 | 0.337741204 | 1.016032254 | 0.348477221 |
| 5.591056 | 0.992957914 | 0.333428008 | 1.01103197 | 0.351001956 |
| 5.601296 | 0.98376621 | 0.331233744 | 1.005485425 | 0.353048233 |
| 5.611536 | 0.974425411 | 0.331522203 | 0.999201064 | 0.355893852 |
| 5.621776 | 0.964552742 | 0.334974825 | 0.992492214 | 0.357989844 |
| 5.632016 | 0.954347739 | 0.341717296 | 0.985558103 | 0.360567201 |
| 5.642256 | 0.943141004 | 0.351595297 | 0.977872714 | 0.362624324 |
| 5.652496 | 0.931454389 | 0.364643403 | 0.97023804 | 0.364679455 |
| 5.662736 | 0.918290515 | 0.379371914 | 0.961502137 | 0.366656934 |
| 5.672976 | 0.904284093 | 0.395328569 | 0.951997166 | 0.368446727 |
| 5.683217 | 0.888938428 | 0.41104128 | 0.941141456 | 0.37029145 |
| 5.693457 | 0.872183389 | 0.425975117 | 0.928398602 | 0.372033348 |
| 5.703696 | 0.854155028 | 0.438967613 | 0.913506319 | 0.373970862 |
| 5.713936 | 0.83412597 | 0.450519413 | 0.896088059 | 0.375716082 |
| 5.724176 | 0.81255566 | 0.460494939 | 0.875671019 | 0.377413083 |
| 5.734416 | 0.788273302 | 0.469329337 | 0.852145389 | 0.378933907 |
| 5.744657 | 0.762168622 | 0.478050706 | 0.82537635 | 0.379908424 |
| 5.754897 | 0.732966962 | 0.486763363 | 0.795728296 | 0.380989028 |
| 5.765137 | 0.702146375 | 0.496618836 | 0.762763474 | 0.381816354 |
| 5.775377 | 0.66877721 | 0.507126377 | 0.727364659 | 0.382415516 |
| 5.785617 | 0.634486535 | 0.51841313 | 0.690154056 | 0.383685872 |
| 5.795857 | 0.59875609 | 0.530593324 | 0.651260879 | 0.385140553 |
| 5.806097 | 0.563052446 | 0.542483736 | 0.611746793 | 0.387501221 |
| 5.816337 | 0.527319187 | 0.55379929 | 0.571903002 | 0.390695447 |
| 5.826577 | 0.491534188 | 0.565019816 | 0.532201181 | 0.395146337 |
| 5.836817 | 0.456013948 | 0.574719553 | 0.493016872 | 0.400302183 |
| 5.847057 | 0.420238027 | 0.583851823 | 0.453965921 | 0.406789898 |
| 5.857297 | 0.384060116 | 0.59206949 | 0.415298222 | 0.413649875 |
| 5.867537 | 0.347114175 | 0.599238729 | 0.376203315 | 0.421144298 |
| 5.877777 | 0.30928444 | 0.605771455 | 0.336924383 | 0.428366559 |
| 5.888017 | 0.270621045 | 0.610874661 | 0.297080015 | 0.435663919 |
| 5.898257 | 0.23113435 | 0.615556213 | 0.256919107 | 0.442137601 |
| 5.908497 | 0.191444218 | 0.618525343 | 0.216012193 | 0.448357117 |
| 5.918737 | 0.151519415 | 0.620976157 | 0.175441926 | 0.453652584 |
| 5.928977 | 0.112072109 | 0.622101943 | 0.134751595 | 0.458075933 |
| 5.939217 | 0.072690639 | 0.622647496 | 0.094767599 | 0.46192867 |
| 5.949457 | 0.034282723 | 0.622307254 | 0.055198459 | 0.465199879 |
| 5.959697 | -0.004106065 | 0.621208146 | 0.016136652 | 0.467978132 |
| 5.969937 | -0.041552995 | 0.619623163 | -0.02246413 | 0.470648274 |
| 5.980177 | -0.078733258 | 0.617610168 | -0.060691418 | 0.473110432 |
| 5.990417 | -0.115542902 | 0.615023864 | -0.099223998 | 0.476562852 |
| 6.000657 | -0.15160151 | 0.612535434 | -0.137464195 | 0.479374093 |
| 6.010897 | -0.187320442 | 0.609480841 | -0.176047383 | 0.483347219 |
| 6.021137 | -0.222018528 | 0.606689883 | -0.214671359 | 0.486928169 |
| 6.031377 | -0.256037556 | 0.603272953 | -0.252656918 | 0.491622469 |
| 6.041617 | -0.289132709 | 0.599754877 | -0.290635366 | 0.495797335 |
| 6.051857 | -0.321160351 | 0.595067862 | -0.327376003 | 0.500575611 |
| 6.062098 | -0.35296045 | 0.589656358 | -0.363079447 | 0.505235842 |
| 6.072338 | -0.383928885 | 0.583423259 | -0.397581841 | 0.509990574 |
| 6.082578 | -0.414957277 | 0.575779564 | -0.430356026 | 0.514537789 |
| 6.092818 | -0.446007864 | 0.568058072 | -0.461855204 | 0.518533821 |
| 6.103057 | -0.47747781 | 0.559385697 | -0.492091199 | 0.522315814 |
| 6.113297 | -0.50921402 | 0.551278633 | -0.52064753 | 0.525545225 |
| 6.123538 | -0.540921982 | 0.542652063 | -0.548631745 | 0.52804983 |
| 6.133778 | -0.573155876 | 0.534987826 | -0.575414371 | 0.530317071 |
| 6.144018 | -0.603900403 | 0.526993513 | -0.601470816 | 0.532069724 |
| 6.154258 | -0.634463753 | 0.520073607 | -0.626511966 | 0.533388882 |
| 6.164498 | -0.662706502 | 0.513072511 | -0.65088837 | 0.534474727 |
| 6.174738 | -0.689240207 | 0.507628159 | -0.673589483 | 0.535479138 |
| 6.184978 | -0.713103324 | 0.502338923 | -0.695731757 | 0.536385732 |
| 6.195218 | -0.73397217 | 0.498734101 | -0.715936338 | 0.537313357 |
| 6.205458 | -0.752885534 | 0.495796719 | -0.735219103 | 0.538304906 |
| 6.215698 | -0.768328603 | 0.4940117 | -0.753513323 | 0.539142047 |
| 6.225938 | -0.782071032 | 0.492661611 | -0.770378016 | 0.53993388 |
| 6.236178 | -0.793384103 | 0.491971005 | -0.787069519 | 0.540575208 |
| 6.246418 | -0.8032129 | 0.491413177 | -0.802729365 | 0.540830426 |
| 6.256658 | -0.810958709 | 0.490569942 | -0.81785739 | 0.541003022 |
| 6.266898 | -0.818054919 | 0.489984577 | -0.831900381 | 0.541201596 |
| 6.277138 | -0.822977664 | 0.488330489 | -0.845014022 | 0.541401662 |
| 6.287378 | -0.826876307 | 0.487185541 | -0.856651836 | 0.541750955 |
| 6.297618 | -0.828975886 | 0.485174188 | -0.866467198 | 0.542794584 |
| 6.307858 | -0.829450396 | 0.483839711 | -0.874410467 | 0.543593381 |
| 6.318098 | -0.82808179 | 0.482402265 | -0.880905102 | 0.544909414 |
| 6.328338 | -0.825196504 | 0.481628373 | -0.884978908 | 0.545645075 |
| 6.338578 | -0.820766247 | 0.481601472 | -0.887765408 | 0.546147464 |
| 6.348818 | -0.81527389 | 0.482145782 | -0.888932191 | 0.545343615 |
| 6.359058 | -0.809472173 | 0.482687393 | -0.888754509 | 0.543524416 |
| 6.369298 | -0.803141128 | 0.483624755 | -0.887169488 | 0.540274105 |
| 6.379539 | -0.797308011 | 0.48371577 | -0.884542182 | 0.535408451 |
| 6.389779 | -0.792115706 | 0.483410119 | -0.880514892 | 0.529342064 |
| 6.400019 | -0.787131986 | 0.482035508 | -0.87531247 | 0.522419414 |
| 6.410258 | -0.782798632 | 0.480062155 | -0.868641439 | 0.514520269 |
| 6.420498 | -0.778912173 | 0.477474397 | -0.860721069 | 0.506876593 |
| 6.430738 | -0.774795544 | 0.475226954 | -0.851202081 | 0.499305223 |
| 6.440979 | -0.770279932 | 0.472813076 | -0.840406117 | 0.492496237 |
| 6.451219 | -0.765331161 | 0.47111507 | -0.828116362 | 0.486772627 |
| 6.461459 | -0.759236184 | 0.470559166 | -0.814265174 | 0.482503546 |
| 6.471699 | -0.752093562 | 0.47077497 | -0.799182855 | 0.479387219 |
| 6.481939 | -0.743011624 | 0.471835835 | -0.782332571 | 0.477561821 |
| 6.492179 | -0.732778946 | 0.473709654 | -0.764586184 | 0.476912559 |
| 6.502419 | -0.720404342 | 0.475571419 | -0.74508081 | 0.47686002 |
| 6.512659 | -0.706792662 | 0.477893772 | -0.724533565 | 0.477480666 |
| 6.522899 | -0.691516505 | 0.479446734 | -0.702721146 | 0.478162158 |
| 6.533139 | -0.675427638 | 0.481049012 | -0.680045455 | 0.479414174 |
| 6.543379 | -0.65849615 | 0.481932082 | -0.656101659 | 0.480730476 |
| 6.553619 | -0.641507562 | 0.483160057 | -0.631451216 | 0.48207367 |
| 6.563859 | -0.624016319 | 0.483878719 | -0.606374751 | 0.483588486 |
| 6.574099 | -0.607064436 | 0.485468292 | -0.580483121 | 0.484869622 |
| 6.584339 | -0.589651817 | 0.487173225 | -0.553875271 | 0.486313845 |
| 6.594579 | -0.572083551 | 0.489363129 | -0.527425866 | 0.486925647 |
| 6.604819 | -0.554504791 | 0.491953886 | -0.500369125 | 0.487524549 |
| 6.615059 | -0.536067834 | 0.494705488 | -0.473091246 | 0.487642886 |
| 6.625299 | -0.517512794 | 0.496384696 | -0.445447252 | 0.48811638 |
| 6.635539 | -0.498163229 | 0.498524452 | -0.417356854 | 0.488030244 |
| 6.645779 | -0.479001801 | 0.498795722 | -0.38914269 | 0.488651332 |
| 6.656019 | -0.459718189 | 0.498261772 | -0.359874296 | 0.489775288 |
| 6.666259 | -0.440775487 | 0.497051756 | -0.330212115 | 0.490949723 |
| 6.676499 | -0.422369486 | 0.494188358 | -0.300073684 | 0.492510158 |
| 6.686739 | -0.404795058 | 0.491047437 | -0.269340242 | 0.49414723 |
| 6.69698 | -0.388266433 | 0.486933646 | -0.238680709 | 0.495973502 |
| 6.707219 | -0.372077204 | 0.482426539 | -0.208348315 | 0.497651763 |
| 6.717459 | -0.357086139 | 0.477015477 | -0.179015782 | 0.499100795 |
| 6.727699 | -0.342034196 | 0.471382144 | -0.150281084 | 0.500590769 |
| 6.737939 | -0.327570105 | 0.464585637 | -0.123579271 | 0.502389017 |
| 6.748179 | -0.312392937 | 0.457096917 | -0.097785035 | 0.503543483 |
| 6.75842 | -0.297307831 | 0.448823656 | -0.073504491 | 0.504872014 |
| 6.76866 | -0.281281254 | 0.439655472 | -0.050501682 | 0.505637835 |
| 6.7789 | -0.264439974 | 0.429573541 | -0.028472243 | 0.506125698 |
| 6.78914 | -0.247283163 | 0.41978165 | -0.008127964 | 0.50552079 |
| 6.79938 | -0.229286952 | 0.409129436 | 0.011348659 | 0.504828387 |
| 6.809619 | -0.210746486 | 0.399309952 | 0.0292036 | 0.502873254 |
| 6.81986 | -0.191864286 | 0.389142926 | 0.045097311 | 0.500451536 |
| 6.8301 | -0.172594707 | 0.379747239 | 0.059499549 | 0.496812772 |
| 6.84034 | -0.153048477 | 0.370599963 | 0.072398673 | 0.492709313 |
| 6.85058 | -0.133028935 | 0.361653498 | 0.083708992 | 0.487118718 |
| 6.86082 | -0.112712872 | 0.353378275 | 0.095108558 | 0.481081411 |
| 6.87106 | -0.091898177 | 0.3454474 | 0.105881839 | 0.474028814 |
| 6.8813 | -0.070601982 | 0.33765241 | 0.117658013 | 0.467051739 |
| 6.89154 | -0.049229275 | 0.330781119 | 0.130441181 | 0.459904659 |
| 6.90178 | -0.027187005 | 0.323513962 | 0.144283529 | 0.453313837 |
| 6.91202 | -0.00539173 | 0.316994281 | 0.159050936 | 0.447823752 |
| 6.92226 | 0.016882078 | 0.310083731 | 0.17427148 | 0.443317347 |
| 6.9325 | 0.038446988 | 0.303401435 | 0.189625618 | 0.439916398 |
| 6.94274 | 0.05990326 | 0.296543541 | 0.204031409 | 0.437245803 |
| 6.95298 | 0.081767311 | 0.289763827 | 0.217231435 | 0.435505391 |
| 6.96322 | 0.102562968 | 0.283166651 | 0.228969308 | 0.433775265 |
| 6.97346 | 0.123968074 | 0.277379766 | 0.238886678 | 0.431852301 |
| 6.9837 | 0.145295103 | 0.271066209 | 0.247217546 | 0.429147076 |
| 6.99394 | 0.167046095 | 0.266249061 | 0.254163599 | 0.425872259 |
| 7.00418 | 0.188379079 | 0.261380466 | 0.260212315 | 0.421522866 |
| 7.01442 | 0.210044803 | 0.257452759 | 0.265380379 | 0.416037483 |
| 7.02466 | 0.231301844 | 0.253991208 | 0.270946097 | 0.409335433 |
| 7.0349 | 0.252234213 | 0.251121087 | 0.276366598 | 0.402342178 |
| 7.04514 | 0.272144721 | 0.248823143 | 0.282434537 | 0.393841376 |
| 7.05538 | 0.291302189 | 0.247187889 | 0.289467003 | 0.385657323 |
| 7.06562 | 0.309978338 | 0.245335861 | 0.297580777 | 0.376973907 |
| 7.075861 | 0.327498487 | 0.244013904 | 0.306192121 | 0.368672192 |
| 7.086101 | 0.344605246 | 0.242500786 | 0.316284191 | 0.360648136 |
| 7.096341 | 0.360825927 | 0.240866716 | 0.326747958 | 0.353599283 |
| 7.106581 | 0.376696501 | 0.238949676 | 0.337939085 | 0.347002692 |
| 7.11682 | 0.391393196 | 0.237233019 | 0.348794242 | 0.341460285 |
| 7.12706 | 0.405838168 | 0.235282317 | 0.36012452 | 0.336325446 |
| 7.1373 | 0.419005499 | 0.233988671 | 0.3705913 | 0.331965957 |
| 7.147541 | 0.432092613 | 0.232885374 | 0.380492072 | 0.328211176 |
| 7.157781 | 0.444575404 | 0.232387034 | 0.389463908 | 0.324069234 |

| | | | | |
|---|---|---|---|---|
| 7.168021 | -0.019482716 | 0.541434883 | 0.197280736 | 0.33920603 |
| 7.178261 | -0.012365162 | 0.54440984 | 0.212671944 | 0.3394331 |
| 7.188501 | -0.005287976 | 0.54513777 | 0.226018279 | 0.33935765 |
| 7.198741 | 0.000770579 | 0.544597654 | 0.238998191 | 0.33679736 |
| 7.208981 | 0.006663254 | 0.541175051 | 0.250162468 | 0.33286049 |
| 7.219221 | 0.012789929 | 0.536218549 | 0.260297862 | 0.32651476 |
| 7.229461 | 0.017881632 | 0.529301364 | 0.269433234 | 0.31871515 |
| 7.239701 | 0.023870391 | 0.520971421 | 0.277750494 | 0.30863561 |
| 7.249941 | 0.028976217 | 0.511974901 | 0.285673484 | 0.29704129 |
| 7.260181 | 0.03455682 | 0.502778722 | 0.293766078 | 0.28478475 |
| 7.270421 | 0.039308809 | 0.494114913 | 0.302382275 | 0.27138067 |
| 7.280661 | 0.043369755 | 0.486287749 | 0.311927952 | 0.25806721 |
| 7.290901 | 0.047411639 | 0.479701641 | 0.322628216 | 0.24509402 |
| 7.301141 | 0.050589719 | 0.474580924 | 0.334244108 | 0.23263847 |
| 7.311381 | 0.053661075 | 0.471173384 | 0.346397172 | 0.22156718 |
| 7.321621 | 0.057004407 | 0.468768817 | 0.359012589 | 0.21148473 |
| 7.331861 | 0.061314359 | 0.467997335 | 0.37100952 | 0.20407368 |
| 7.342101 | 0.066126786 | 0.468558009 | 0.381714663 | 0.19872466 |
| 7.352341 | 0.072421388 | 0.469308428 | 0.391288017 | 0.19550014 |
| 7.362581 | 0.07964961 | 0.4710851 | 0.399054672 | 0.19541744 |
| 7.372821 | 0.087043793 | 0.471921949 | 0.404809166 | 0.19676356 |
| 7.383061 | 0.094633453 | 0.472531019 | 0.408882039 | 0.19991357 |
| 7.393301 | 0.101421361 | 0.471777449 | 0.411765567 | 0.20351182 |
| 7.403542 | 0.107248692 | 0.47002563 | 0.413880695 | 0.20722851 |
| 7.413781 | 0.11186761 | 0.466848815 | 0.415608859 | 0.21026668 |
| 7.424021 | 0.115123616 | 0.462988297 | 0.417837828 | 0.21189125 |
| 7.434261 | 0.118122901 | 0.458707516 | 0.420312164 | 0.21174263 |
| 7.444501 | 0.120635729 | 0.454315239 | 0.423828185 | 0.21042083 |
| 7.454741 | 0.123298592 | 0.450901784 | 0.427448445 | 0.20712181 |
| 7.464982 | 0.127159093 | 0.447676455 | 0.431783908 | 0.20249474 |
| 7.475222 | 0.131333791 | 0.445869153 | 0.435294328 | 0.19757012 |
| 7.485462 | 0.137030395 | 0.444315722 | 0.438617751 | 0.19270394 |
| 7.495702 | 0.143049297 | 0.443193563 | 0.440197894 | 0.18936052 |
| 7.505942 | 0.15037851 | 0.442366931 | 0.440876335 | 0.18776759 |
| 7.516181 | 0.157620149 | 0.440781243 | 0.43950431 | 0.18801983 |
| 7.526422 | 0.165281326 | 0.438820061 | 0.437027561 | 0.18997393 |
| 7.536662 | 0.173015674 | 0.436236743 | 0.432961747 | 0.19264194 |
| 7.546902 | 0.180651667 | 0.432925309 | 0.428008935 | 0.19588115 |
| 7.557142 | 0.188401992 | 0.429501245 | 0.422686436 | 0.19900268 |
| 7.567382 | 0.195720283 | 0.424788682 | 0.416322722 | 0.20188578 |
| 7.577622 | 0.203261408 | 0.421414063 | 0.410008893 | 0.20554213 |
| 7.587862 | 0.210138167 | 0.417101056 | 0.403624783 | 0.20949384 |
| 7.598102 | 0.216517528 | 0.413080144 | 0.396701389 | 0.21431751 |
| 7.608342 | 0.222120672 | 0.408855814 | 0.389976743 | 0.22000454 |
| 7.618582 | 0.226084714 | 0.404460543 | 0.383089427 | 0.22669673 |
| 7.628822 | 0.229194428 | 0.399875552 | 0.376028258 | 0.23393491 |
| 7.639062 | 0.230840005 | 0.393715702 | 0.369328836 | 0.24149297 |
| 7.649302 | 0.2312361 | 0.387108316 | 0.36300536 | 0.24871869 |
| 7.659542 | 0.230659438 | 0.379512955 | 0.356946515 | 0.25555709 |
| 7.669782 | 0.229758427 | 0.370023463 | 0.351478987 | 0.2612983 |
| 7.680022 | 0.228743007 | 0.35976323 | 0.346566927 | 0.26604712 |
| 7.690262 | 0.227357802 | 0.348276901 | 0.342375602 | 0.26958484 |
| 7.700502 | 0.226416262 | 0.335572395 | 0.337713457 | 0.27088426 |
| 7.710742 | 0.22471079 | 0.321889434 | 0.333618671 | 0.27044765 |
| 7.720982 | 0.223267168 | 0.30760538 | 0.328873931 | 0.26835496 |
| 7.731222 | 0.220101563 | 0.292511321 | 0.323631326 | 0.26391631 |
| 7.741462 | 0.216968936 | 0.276658682 | 0.317815264 | 0.25801391 |
| 7.751702 | 0.211492582 | 0.2609396 | 0.31136376 | 0.25067544 |
| 7.761942 | 0.205756631 | 0.245146125 | 0.303976748 | 0.2433062 |
| 7.772182 | 0.198290383 | 0.229294394 | 0.295726011 | 0.23554682 |
| 7.782423 | 0.190747502 | 0.213972854 | 0.287257087 | 0.22930436 |
| 7.792663 | 0.182681275 | 0.198599363 | 0.277357891 | 0.223859 |
| 7.802903 | 0.174940703 | 0.184016851 | 0.267739428 | 0.22035832 |
| 7.813143 | 0.167716507 | 0.16906032 | 0.256695917 | 0.21783044 |
| 7.823382 | 0.161208095 | 0.155694804 | 0.246527447 | 0.21652728 |
| 7.833622 | 0.155872453 | 0.142880837 | 0.235571068 | 0.21638485 |
| 7.843863 | 0.151244743 | 0.132109278 | 0.225824161 | 0.21655157 |
| 7.854103 | 0.146955179 | 0.123630995 | 0.216070344 | 0.21791198 |
| 7.864343 | 0.142949918 | 0.117782858 | 0.208342744 | 0.22054546 |
| 7.874583 | 0.138950271 | 0.114634958 | 0.20089566 | 0.22374782 |
| 7.884823 | 0.134609653 | 0.114267265 | 0.195240251 | 0.22888434 |
| 7.895063 | 0.129566451 | 0.116501189 | 0.189230569 | 0.23432403 |
| 7.905303 | 0.124578223 | 0.120643904 | 0.184694175 | 0.24070773 |
| 7.915543 | 0.119375017 | 0.126902887 | 0.179163756 | 0.24643685 |
| 7.925783 | 0.11488606 | 0.134494214 | 0.173238258 | 0.25186834 |
| 7.936023 | 0.110883009 | 0.143018577 | 0.166552961 | 0.2563226 |
| 7.946263 | 0.108276575 | 0.151924218 | 0.158854158 | 0.25981743 |
| 7.956503 | 0.107338599 | 0.161043194 | 0.150251708 | 0.26221081 |
| 7.966743 | 0.107349815 | 0.169221733 | 0.140709943 | 0.26423697 |
| 7.976983 | 0.109059572 | 0.176538097 | 0.131401912 | 0.2658394 |
| 7.987223 | 0.111464525 | 0.182453668 | 0.121006171 | 0.26738975 |
| 7.997463 | 0.114296639 | 0.186742454 | 0.111859694 | 0.26900984 |
| 8.007703 | 0.117438762 | 0.190266276 | 0.10222918 | 0.2706816 |
| 8.017943 | 0.119907199 | 0.191992582 | 0.093726835 | 0.27242741 |
| 8.028183 | 0.121721163 | 0.192898154 | 0.085392096 | 0.27445276 |
| 8.038424 | 0.122575447 | 0.193698283 | 0.078092456 | 0.27605716 |
| 8.048663 | 0.121836416 | 0.193228066 | 0.0710146 | 0.2774214 |
| 8.058904 | 0.120035227 | 0.193760466 | 0.06494209 | 0.27791883 |
| 8.069143 | 0.116943869 | 0.193194728 | 0.059309416 | 0.27799529 |
| 8.079383 | 0.112431434 | 0.194329105 | 0.054115483 | 0.27650904 |
| 8.089623 | 0.10690144 | 0.195086729 | 0.049504926 | 0.27404205 |
| 8.099863 | 0.10074895 | 0.197949719 | 0.045250977 | 0.27007652 |
| 8.110104 | 0.093660595 | 0.200964707 | 0.041258453 | 0.26563267 |
| 8.120343 | 0.086135463 | 0.20611663 | 0.037438589 | 0.26008327 |
| 8.130584 | 0.077773245 | 0.212024615 | 0.03353573 | 0.25485708 |
| 8.140823 | 0.06899238 | 0.219137087 | 0.029815856 | 0.24980454 |
| 8.151064 | 0.059203131 | 0.226630904 | 0.025887883 | 0.24585317 |
| 8.161304 | 0.048995792 | 0.233773348 | 0.021399022 | 0.24267517 |
| 8.171543 | 0.037949358 | 0.240558446 | 0.016422654 | 0.24141626 |
| 8.181784 | 0.027011817 | 0.245349397 | 0.01095892 | 0.24091172 |
| 8.192023 | 0.015801663 | 0.24911543 | 0.004603775 | 0.24172278 |
| 8.202264 | 0.005845251 | 0.249838075 | -0.002383985 | 0.24359257 |
| 8.212503 | -0.003795619 | 0.249366752 | -0.010213758 | 0.2459023 |
| 8.222744 | -0.0117206 | 0.246101073 | -0.018148976 | 0.24818464 |
| 8.232984 | -0.019105278 | 0.242083175 | -0.026526858 | 0.25060408 |
| 8.243224 | -0.025650717 | 0.236036858 | -0.034778436 | 0.25212304 |
| 8.253464 | -0.031499835 | 0.23020014 | -0.042519517 | 0.25303862 |
| 8.263704 | -0.037820985 | 0.22333546 | -0.049282928 | 0.2529775 |
| 8.273944 | -0.044122772 | 0.218007195 | -0.055127901 | 0.25226778 |
| 8.284184 | -0.050940967 | 0.212687684 | -0.060116468 | 0.25094507 |
| 8.294424 | -0.058525997 | 0.209198439 | -0.063584869 | 0.24864449 |
| 8.304664 | -0.065751867 | 0.207010733 | -0.066540948 | 0.24631394 |
| 8.314904 | -0.073562309 | 0.20615519 | -0.068131876 | 0.24378224 |
| 8.325144 | -0.080703211 | 0.206817639 | -0.069608749 | 0.2403969 |
| 8.335384 | -0.087432953 | 0.208002222 | -0.069834726 | 0.23692889 |
| 8.345624 | -0.0936703 | 0.209805433 | -0.070428641 | 0.23366943 |
| 8.355865 | -0.099048682 | 0.211736966 | -0.070190757 | 0.22968495 |
| 8.366104 | -0.103625199 | 0.213598143 | -0.070240208 | 0.2262572 |
| 8.376344 | -0.108284351 | 0.215068135 | -0.069959959 | 0.22318407 |
| 8.386584 | -0.111879986 | 0.216551933 | -0.070802277 | 0.22070407 |
| 8.396824 | -0.116168747 | 0.217839777 | -0.071240642 | 0.21858013 |
| 8.407064 | -0.11985365 | 0.219287005 | -0.072629985 | 0.21702747 |
| 8.417304 | -0.1238163 | 0.220629533 | -0.074364438 | 0.21603719 |
| 8.427545 | -0.127791665 | 0.222010963 | -0.076000086 | 0.21506377 |
| 8.437784 | -0.132109455 | 0.22376746 | -0.077767297 | 0.21470346 |
| 8.448025 | -0.135671315 | 0.224972787 | -0.07873581 | 0.21436129 |
| 8.458264 | -0.139630553 | 0.226134933 | -0.07892383 | 0.2144958 |
| 8.468505 | -0.143724955 | 0.226643648 | -0.078204841 | 0.21441873 |
| 8.478745 | -0.146979797 | 0.227566478 | -0.076238165 | 0.21498497 |
| 8.488984 | -0.150774598 | 0.227296509 | -0.073278012 | 0.21559567 |
| 8.499225 | -0.154239724 | 0.227779127 | -0.069707084 | 0.2171183 |
| 8.509464 | -0.157476547 | 0.228308589 | -0.065265737 | 0.22013641 |
| 8.519705 | -0.160395142 | 0.230090121 | -0.060737995 | 0.22447465 |
| 8.529944 | -0.163087918 | 0.231781446 | -0.055551722 | 0.23105257 |
| 8.540185 | -0.164847121 | 0.234898174 | -0.050446463 | 0.2395545 |
| 8.550425 | -0.166517688 | 0.238513826 | -0.044921731 | 0.24904909 |
| 8.560665 | -0.167218237 | 0.241984415 | -0.039085204 | 0.25955177 |
| 8.570905 | -0.167940019 | 0.2451609 | -0.03284622 | 0.26959368 |
| 8.581144 | -0.168391916 | 0.247072236 | -0.026273687 | 0.279412 |
| 8.591385 | -0.16873211 | 0.247503796 | -0.019249666 | 0.28819348 |
| 8.601624 | -0.16986692 | 0.245618119 | -0.01225486 | 0.29567524 |
| 8.611865 | -0.171130088 | 0.241300312 | -0.004983075 | 0.30227503 |
| 8.622105 | -0.173151656 | 0.234706353 | 0.001662311 | 0.30735593 |
| 8.632345 | -0.176319805 | 0.226272266 | 0.007577718 | 0.31108283 |
| 8.642585 | -0.1800693 | 0.216306165 | 0.012952874 | 0.3134358 |
| 8.652825 | -0.184873829 | 0.206329581 | 0.016294899 | 0.31419566 |
| 8.663065 | -0.190495564 | 0.196083577 | 0.018801481 | 0.31449326 |
| 8.673305 | -0.196278547 | 0.187450865 | 0.019325175 | 0.31340129 |
| 8.683545 | -0.20272424 | 0.180247705 | 0.018662773 | 0.31226463 |
| 8.693785 | -0.208401529 | 0.17459614 | 0.016945288 | 0.3110595 |
| 8.704025 | -0.21375592 | 0.171080741 | 0.014237616 | 0.30953869 |
| 8.714265 | -0.217816687 | 0.168159619 | 0.011255993 | 0.30827946 |
| 8.724505 | -0.220411676 | 0.166699091 | 0.008218361 | 0.30655271 |
| 8.734745 | -0.222747792 | 0.16557337 | 0.00551562 | 0.30485889 |
| 8.744986 | -0.22276815 | 0.164617146 | 0.003101541 | 0.302573 |
| 8.755225 | -0.223166826 | 0.164572997 | 0.001612983 | 0.30038431 |
| 8.765466 | -0.221644038 | 0.164455463 | 0.001601191 | 0.29808878 |
| 8.775705 | -0.219930054 | 0.16480277 | 0.002128945 | 0.29550692 |
| 8.785945 | -0.217342606 | 0.165879092 | 0.004510657 | 0.29387965 |
| 8.796185 | -0.213550989 | 0.165746172 | 0.007558089 | 0.2921681 |
| 8.806425 | -0.209343178 | 0.166660466 | 0.012126265 | 0.29140988 |
| 8.816666 | -0.204038069 | 0.166174865 | 0.017122115 | 0.2910166 |
| 8.826905 | -0.198239433 | 0.16538757 | 0.022221104 | 0.29099748 |
| 8.837146 | -0.191850718 | 0.164161062 | 0.027328459 | 0.29151955 |
| 8.847385 | -0.185295509 | 0.162258958 | 0.031030956 | 0.29202045 |
| 8.857626 | -0.178828226 | 0.160331439 | 0.034356491 | 0.29242482 |
| 8.867866 | -0.17220557 | 0.157528307 | 0.035583516 | 0.29279349 |
| 8.878105 | -0.16608721 | 0.155703093 | 0.0361104 | 0.29279099 |
| 8.888346 | -0.160419705 | 0.153059275 | 0.035066662 | 0.29249346 |
| 8.898585 | -0.155070961 | 0.15174557 | 0.033265255 | 0.29212445 |
| 8.908826 | -0.150348438 | 0.150688425 | 0.031178159 | 0.29134354 |
| 8.919065 | -0.146085475 | 0.150169186 | 0.028852288 | 0.29028057 |
| 8.929306 | -0.142152876 | 0.151404005 | 0.027360515 | 0.28896789 |
| 8.939546 | -0.138681275 | 0.151159628 | 0.025869879 | 0.2869896 |
| 8.949786 | -0.134948922 | 0.152485878 | 0.025914717 | 0.28455649 |
| 8.960026 | -0.131449743 | 0.152146015 | 0.02597434 | 0.28136481 |
| 8.970266 | -0.127623712 | 0.151442202 | 0.027324903 | 0.277721 |
| 8.980506 | -0.124272295 | 0.149491321 | 0.028493504 | 0.27347215 |
| 8.990746 | -0.120269859 | 0.147214801 | 0.030413418 | 0.26860655 |
| 9.000986 | -0.117723504 | 0.143353105 | 0.032093951 | 0.26324646 |
| 9.011226 | -0.115248975 | 0.13951647 | 0.033498369 | 0.25742686 |
| 9.021466 | -0.114790529 | 0.135333393 | 0.034404777 | 0.2508319 |
| 9.031706 | -0.11467736 | 0.130831398 | 0.035244028 | 0.24400449 |
| 9.041946 | -0.116316569 | 0.125738132 | 0.035192662 | 0.2364135 |
| 9.052186 | -0.117876602 | 0.121615311 | 0.035089183 | 0.22880735 |
| 9.062427 | -0.120214549 | 0.117777804 | 0.034423397 | 0.22105219 |
| 9.072666 | -0.121492231 | 0.114965294 | 0.03395709 | 0.21409403 |
| 9.082906 | -0.121956256 | 0.114689662 | 0.033482256 | 0.20772129 |
| 9.093146 | -0.120827968 | 0.115975925 | 0.033376045 | 0.20308949 |
| 9.103386 | -0.11776783 | 0.119468085 | 0.033804778 | 0.20024674 |
| 9.113626 | -0.113234139 | 0.123174915 | 0.034511674 | 0.19958094 |
| 9.123866 | -0.106804948 | 0.127761149 | 0.036113486 | 0.20105062 |
| 9.134107 | -0.099374446 | 0.1314554 | 0.03782529 | 0.20391115 |
| 9.144346 | -0.090746858 | 0.134164687 | 0.039920596 | 0.20794227 |
| 9.154587 | -0.082127962 | 0.135807121 | 0.04216826 | 0.21259361 |
| 9.164826 | -0.072817791 | 0.137237 | 0.043737609 | 0.21685456 |
| 9.175066 | -0.064369344 | 0.137854045 | 0.044960555 | 0.22036088 |
| 9.185307 | -0.055788069 | 0.139044157 | 0.044883071 | 0.22291596 |
| 9.195546 | -0.048202509 | 0.1406477 | 0.043789555 | 0.22421818 |
| 9.205787 | -0.041483087 | 0.143149424 | 0.041388916 | 0.22460019 |
| 9.216026 | -0.035077745 | 0.146294276 | 0.037710633 | 0.22401255 |
| 9.226267 | -0.030216563 | 0.150544274 | 0.03319284 | 0.22281138 |
| 9.236506 | -0.025761768 | 0.155778248 | 0.028187652 | 0.22198481 |
| 9.246747 | -0.022703252 | 0.161636446 | 0.02340995 | 0.22123011 |
| 9.256987 | -0.020216903 | 0.168029276 | 0.019495685 | 0.2213121 |
| 9.267227 | -0.018944805 | 0.17455934 | 0.016316326 | 0.22232992 |
| 9.277467 | -0.01841006 | 0.180769114 | 0.014888373 | 0.2245963 |
| 9.287706 | -0.018900876 | 0.186815191 | 0.014572115 | 0.2275838 |
| 9.297947 | -0.020450219 | 0.192127993 | 0.015158445 | 0.23064708 |
| 9.308187 | -0.022625195 | 0.197113323 | 0.017090046 | 0.23441936 |
| 9.318427 | -0.025746713 | 0.201963163 | 0.018629359 | 0.23695894 |
| 9.328667 | -0.029350217 | 0.206598575 | 0.02075776 | 0.23891867 |
| 9.338907 | -0.033522636 | 0.211299582 | 0.022067862 | 0.23886179 |
| 9.349147 | -0.037337982 | 0.215949445 | 0.023739267 | 0.23825697 |
| 9.359387 | -0.041359179 | 0.219308899 | 0.024387445 | 0.23526062 |
| 9.369627 | -0.0445559 | 0.222343418 | 0.025495424 | 0.2319673 |
| 9.379867 | -0.047738496 | 0.22435011 | 0.026789144 | 0.22784099 |
| 9.390107 | -0.049257873 | 0.225088134 | 0.028076525 | 0.22388017 |
| 9.400347 | -0.05091295 | 0.22507949 | 0.030683584 | 0.2200493 |
| 9.410587 | -0.051300332 | 0.225490737 | 0.033980428 | 0.21715869 |
| 9.420827 | -0.050638935 | 0.225417602 | 0.038247275 | 0.21501844 |
| 9.431067 | -0.049092397 | 0.226514556 | 0.043330787 | 0.2131202 |
| 9.441307 | -0.046656846 | 0.229250563 | 0.049938155 | 0.21128197 |
| 9.451548 | -0.042540405 | 0.23296229 | 0.056520282 | 0.2088939 |
| 9.461787 | -0.037721954 | 0.238481348 | 0.064111171 | 0.20558044 |
| 9.472028 | -0.031391414 | 0.244984496 | 0.071231129 | 0.20120723 |
| 9.482267 | -0.024221449 | 0.252660614 | 0.078748511 | 0.19609542 |
| 9.492507 | -0.015861248 | 0.260429599 | 0.085340732 | 0.19001594 |
| 9.502748 | -0.007402067 | 0.268771375 | 0.090929767 | 0.18410249 |
| 9.512987 | 0.001307802 | 0.276449454 | 0.095059098 | 0.17872742 |
| 9.523228 | 0.01034794 | 0.284366974 | 0.097820407 | 0.17379686 |
| 9.533467 | 0.01878758 | 0.291221298 | 0.098186498 | 0.17016066 |
| 9.543708 | 0.027025923 | 0.297028199 | 0.096523248 | 0.16775525 |
| 9.553947 | 0.034319734 | 0.30189349 | 0.093128183 | 0.16634078 |
| 9.564188 | 0.041934656 | 0.304823462 | 0.088186874 | 0.16646572 |

| | | | | |
|---|---|---|---|---|
| 7.168021 | 0.186991652 | 0.43391296 | 0.143258172 | 0.45980375 |
| 7.178261 | 0.189673074 | 0.428018408 | 0.149330568 | 0.457400492 |
| 7.188501 | 0.193666716 | 0.421326061 | 0.15508614 | 0.455379786 |
| 7.198741 | 0.196333465 | 0.414675779 | 0.158888728 | 0.452880307 |
| 7.208981 | 0.200052917 | 0.407388574 | 0.162756825 | 0.450541924 |
| 7.219221 | 0.202936731 | 0.400365025 | 0.164899511 | 0.447775246 |
| 7.229461 | 0.205622548 | 0.393279995 | 0.166650663 | 0.445136605 |
| 7.239701 | 0.207642631 | 0.386431618 | 0.166976886 | 0.441415443 |
| 7.249941 | 0.209128341 | 0.379846503 | 0.166633111 | 0.43705836 |
| 7.260181 | 0.210306211 | 0.373361366 | 0.165128435 | 0.4305136 |
| 7.270421 | 0.211490963 | 0.366887375 | 0.162697291 | 0.423269722 |
| 7.280661 | 0.21294162 | 0.360337008 | 0.159390255 | 0.413671612 |
| 7.290901 | 0.214866645 | 0.353745017 | 0.154974995 | 0.40330055 |
| 7.301141 | 0.217622417 | 0.346716763 | 0.150470453 | 0.392031458 |
| 7.311381 | 0.220988769 | 0.338948695 | 0.145040022 | 0.381076725 |
| 7.321621 | 0.2253316 | 0.331172224 | 0.14027172 | 0.371257151 |
| 7.331861 | 0.23018944 | 0.323199161 | 0.135554859 | 0.362330525 |
| 7.342101 | 0.235819075 | 0.314106794 | 0.131191746 | 0.3551667 |
| 7.352341 | 0.241996219 | 0.305876661 | 0.127679703 | 0.349564771 |
| 7.362581 | 0.248290741 | 0.29709209 | 0.124706179 | 0.344764405 |
| 7.372821 | 0.255388136 | 0.288785697 | 0.122374603 | 0.340090466 |
| 7.383061 | 0.261624243 | 0.280693966 | 0.12012571 | 0.335344688 |
| 7.393301 | 0.268145142 | 0.272995995 | 0.118306112 | 0.329544209 |
| 7.403542 | 0.273545076 | 0.265959005 | 0.116424012 | 0.322561063 |
| 7.413781 | 0.278182056 | 0.259581996 | 0.114779713 | 0.313951411 |
| 7.424021 | 0.280903795 | 0.253519119 | 0.11286841 | 0.304345419 |
| 7.434261 | 0.282582595 | 0.248273806 | 0.111272496 | 0.293725372 |
| 7.444501 | 0.282352868 | 0.242559987 | 0.109091608 | 0.282304661 |
| 7.454741 | 0.280843057 | 0.237130298 | 0.107169782 | 0.270637186 |
| 7.464982 | 0.278062469 | 0.230845184 | 0.104866334 | 0.259285518 |
| 7.475222 | 0.274548798 | 0.223736938 | 0.101752912 | 0.247902406 |
| 7.485462 | 0.270573119 | 0.215605916 | 0.098224798 | 0.237472443 |
| 7.495702 | 0.267069042 | 0.206938141 | 0.094221993 | 0.227830906 |
| 7.505942 | 0.263389672 | 0.197972866 | 0.089540258 | 0.219167347 |
| 7.516181 | 0.26071364 | 0.188896377 | 0.084604533 | 0.212304753 |
| 7.526422 | 0.258406339 | 0.180647235 | 0.079651242 | 0.206313517 |
| 7.536662 | 0.256496571 | 0.173115519 | 0.074354475 | 0.201690776 |
| 7.546902 | 0.254577455 | 0.166454823 | 0.069611713 | 0.198112841 |
| 7.557142 | 0.252181459 | 0.160953555 | 0.06403642 | 0.194975352 |
| 7.567382 | 0.249309778 | 0.156007198 | 0.058497328 | 0.192879363 |
| 7.577622 | 0.245097094 | 0.151209473 | 0.051876233 | 0.190462782 |
| 7.587862 | 0.240106578 | 0.145992903 | 0.044533326 | 0.188574432 |
| 7.598102 | 0.233878544 | 0.140507562 | 0.035628298 | 0.187480319 |
| 7.608342 | 0.226719113 | 0.13434348 | 0.026261165 | 0.185592936 |
| 7.618582 | 0.218903987 | 0.128258189 | 0.015891497 | 0.183869143 |
| 7.628822 | 0.20988618 | 0.122345574 | 0.004729499 | 0.180954868 |
| 7.639062 | 0.200790685 | 0.11891593 | -0.006339673 | 0.178000974 |
| 7.649302 | 0.190431086 | 0.118307975 | -0.017478941 | 0.173847645 |
| 7.659542 | 0.17945526 | 0.121891234 | -0.028176554 | 0.169418928 |
| 7.669782 | 0.167520358 | 0.129171718 | -0.038900946 | 0.164436469 |
| 7.680022 | 0.15508421 | 0.139387161 | -0.048558431 | 0.160339164 |
| 7.690262 | 0.142146784 | 0.151220774 | -0.057880553 | 0.155973371 |
| 7.700502 | 0.128982301 | 0.163444042 | -0.066392132 | 0.152811622 |
| 7.710742 | 0.115896422 | 0.17465535 | -0.074386992 | 0.15014956 |
| 7.720982 | 0.103759501 | 0.185084248 | -0.081624977 | 0.148559567 |
| 7.731222 | 0.092269736 | 0.193140437 | -0.088250147 | 0.147692868 |
| 7.741462 | 0.08242862 | 0.200940117 | -0.094064671 | 0.148288899 |
| 7.751702 | 0.074019803 | 0.205461757 | -0.099992616 | 0.149770996 |
| 7.761942 | 0.067209166 | 0.209161343 | -0.105014047 | 0.152634387 |
| 7.772182 | 0.062419008 | 0.210954827 | -0.109807717 | 0.156894466 |
| 7.782423 | 0.058743134 | 0.21082857 | -0.114060314 | 0.162307526 |
| 7.792663 | 0.056835902 | 0.209686043 | -0.118054633 | 0.169353092 |
| 7.802903 | 0.055616491 | 0.206652372 | -0.121543337 | 0.176565864 |
| 7.813143 | 0.055391502 | 0.203146598 | -0.124838656 | 0.185122839 |
| 7.823382 | 0.055097871 | 0.198561276 | -0.128203206 | 0.19325978 |
| 7.833622 | 0.054759489 | 0.194780729 | -0.131764844 | 0.201574776 |
| 7.843863 | 0.053869781 | 0.191151865 | -0.135950578 | 0.208636816 |
| 7.854103 | 0.052016769 | 0.189197534 | -0.14075517 | 0.214186512 |
| 7.864343 | 0.04916834 | 0.189160753 | -0.146818223 | 0.218013826 |
| 7.874583 | 0.045091492 | 0.191578769 | -0.153124974 | 0.219637609 |
| 7.884823 | 0.040651811 | 0.196186779 | -0.161000797 | 0.219562297 |
| 7.895063 | 0.034719193 | 0.202932985 | -0.168737947 | 0.217639929 |
| 7.905303 | 0.029292858 | 0.210724492 | -0.177675951 | 0.214696793 |
| 7.915543 | 0.023307441 | 0.21884894 | -0.185740187 | 0.211282204 |
| 7.925783 | 0.017477056 | 0.22727322 | -0.194596979 | 0.208018102 |
| 7.936023 | 0.012093903 | 0.234466292 | -0.202768216 | 0.204931003 |
| 7.946263 | 0.006686519 | 0.240315302 | -0.21020194 | 0.202746492 |
| 7.956503 | 0.001422158 | 0.245377764 | -0.217561698 | 0.200519681 |
| 7.966743 | -0.00362697 | 0.249031707 | -0.223244897 | 0.199533427 |
| 7.976983 | -0.00906864 | 0.252013105 | -0.228896827 | 0.198628698 |
| 7.987223 | -0.014048947 | 0.254647481 | -0.232590071 | 0.198613897 |
| 7.997463 | -0.019580632 | 0.257006819 | -0.235933002 | 0.198966894 |
| 8.007703 | -0.024561795 | 0.259530234 | -0.237563704 | 0.199976141 |
| 8.017943 | -0.029594531 | 0.261563501 | -0.238843332 | 0.201315468 |
| 8.028183 | -0.034595286 | 0.263412282 | -0.238977918 | 0.203135934 |
| 8.038424 | -0.039805592 | 0.264626408 | -0.238663546 | 0.20515575 |
| 8.048663 | -0.045272658 | 0.264438022 | -0.238306698 | 0.207270219 |
| 8.058904 | -0.05065613 | 0.264546677 | -0.237974956 | 0.209708218 |
| 8.069143 | -0.056417429 | 0.263809259 | -0.238050341 | 0.212501143 |
| 8.079383 | -0.062529009 | 0.263413001 | -0.238743444 | 0.215622153 |
| 8.089623 | -0.068024056 | 0.263389977 | -0.239885876 | 0.220096447 |
| 8.099863 | -0.073618263 | 0.264853082 | -0.241660399 | 0.225287608 |
| 8.110104 | -0.079225979 | 0.267393336 | -0.24336798 | 0.231889562 |
| 8.120343 | -0.083837551 | 0.271644625 | -0.245725633 | 0.239849615 |
| 8.130584 | -0.089240784 | 0.276818274 | -0.247159994 | 0.248920628 |
| 8.140823 | -0.093554535 | 0.283002948 | -0.248630662 | 0.258945092 |
| 8.151064 | -0.098671782 | 0.289477121 | -0.249439577 | 0.268803301 |
| 8.161304 | -0.103351104 | 0.296508584 | -0.249869153 | 0.279369396 |
| 8.171543 | -0.108221603 | 0.302397114 | -0.249961766 | 0.28860671 |
| 8.181784 | -0.112674549 | 0.307678926 | -0.24996925 | 0.297259781 |
| 8.192023 | -0.117306801 | 0.312360644 | -0.249823814 | 0.304604844 |
| 8.202264 | -0.120605566 | 0.314953528 | -0.249884986 | 0.310065167 |
| 8.212503 | -0.124068221 | 0.317104145 | -0.250071235 | 0.314437658 |
| 8.222744 | -0.126213858 | 0.318002569 | -0.250333774 | 0.317425742 |
| 8.232984 | -0.127954653 | 0.317352177 | -0.250428472 | 0.318845846 |
| 8.243224 | -0.128601989 | 0.316074386 | -0.250273309 | 0.319565168 |
| 8.253464 | -0.128563703 | 0.313521207 | -0.249999489 | 0.319342483 |
| 8.263704 | -0.128346911 | 0.31015688 | -0.249385503 | 0.317978726 |
| 8.273944 | -0.127162522 | 0.306256209 | -0.248381334 | 0.316419214 |
| 8.284184 | -0.126602481 | 0.301563457 | -0.247392086 | 0.313444062 |
| 8.294424 | -0.125844899 | 0.297579458 | -0.24603946 | 0.310100226 |
| 8.304664 | -0.126071737 | 0.293098182 | -0.244568125 | 0.305983093 |
| 8.314904 | -0.126993757 | 0.289680194 | -0.243249346 | 0.301307085 |
| 8.325144 | -0.128795839 | 0.286649281 | -0.24145346 | 0.295913276 |
| 8.335384 | -0.132422802 | 0.284354944 | -0.239946349 | 0.290606827 |
| 8.345624 | -0.136236454 | 0.28194458 | -0.237847329 | 0.284872622 |
| 8.355865 | -0.141146112 | 0.279533558 | -0.235574722 | 0.278898165 |
| 8.366104 | -0.146423093 | 0.276978415 | -0.233395868 | 0.27273877 |
| 8.376344 | -0.151654433 | 0.272971911 | -0.230603196 | 0.266156171 |
| 8.386584 | -0.156672133 | 0.268516112 | -0.228358964 | 0.259671733 |
| 8.396824 | -0.160655234 | 0.262492918 | -0.225863118 | 0.252187013 |
| 8.407064 | -0.164183147 | 0.256199017 | -0.224147091 | 0.245191043 |
| 8.417304 | -0.166668303 | 0.2485234 | -0.222546646 | 0.238559688 |
| 8.427545 | -0.168139538 | 0.241442287 | -0.221754681 | 0.232152855 |
| 8.437784 | -0.168949933 | 0.234230758 | -0.221103578 | 0.22725839 |
| 8.448025 | -0.168898775 | 0.227872319 | -0.221324804 | 0.22312793 |
| 8.458264 | -0.168309249 | 0.222487283 | -0.221250614 | 0.220505098 |
| 8.468505 | -0.167293565 | 0.218451014 | -0.220993048 | 0.219221957 |
| 8.478745 | -0.165593951 | 0.215940368 | -0.220592471 | 0.218272235 |
| 8.488984 | -0.164551202 | 0.214899689 | -0.219123998 | 0.21844076 |
| 8.499225 | -0.162731214 | 0.215376461 | -0.217042787 | 0.218118047 |
| 8.509464 | -0.161698685 | 0.217743679 | -0.213072211 | 0.218121582 |
| 8.519705 | -0.160628095 | 0.22177439 | -0.20935072 | 0.217211408 |
| 8.529944 | -0.160047804 | 0.226801115 | -0.203672369 | 0.215637105 |
| 8.540185 | -0.159765236 | 0.23335491 | -0.19823713 | 0.213352637 |
| 8.550425 | -0.159439668 | 0.240513939 | -0.192198229 | 0.209551716 |
| 8.560665 | -0.159660057 | 0.247579129 | -0.186741606 | 0.20524751 |
| 8.570905 | -0.159065681 | 0.254278743 | -0.181461956 | 0.199042635 |
| 8.581144 | -0.159218738 | 0.260028495 | -0.176960035 | 0.19222496 |
| 8.591385 | -0.158645733 | 0.264599063 | -0.172946067 | 0.183880687 |
| 8.601624 | -0.158652539 | 0.26747654 | -0.169434142 | 0.175249645 |
| 8.611865 | -0.158313271 | 0.268446925 | -0.166398743 | 0.165646733 |
| 8.622105 | -0.15832428 | 0.268242495 | -0.162856621 | 0.156379007 |
| 8.632345 | -0.158424564 | 0.267068275 | -0.15949088 | 0.147418336 |
| 8.642585 | -0.158361105 | 0.265743037 | -0.15558668 | 0.139246287 |
| 8.652825 | -0.158090312 | 0.264900734 | -0.151276066 | 0.13188495 |
| 8.663065 | -0.15804148 | 0.265057319 | -0.146490272 | 0.125972116 |
| 8.673305 | -0.157659976 | 0.2671939 | -0.141284787 | 0.120753721 |
| 8.683545 | -0.15778325 | 0.27046664 | -0.135548054 | 0.116958667 |
| 8.693785 | -0.157631725 | 0.2749722 | -0.129327003 | 0.113722115 |
| 8.704025 | -0.158833252 | 0.280314031 | -0.122771702 | 0.111302455 |
| 8.714265 | -0.160538937 | 0.285887585 | -0.115745926 | 0.109279917 |
| 8.724505 | -0.162945242 | 0.290596177 | -0.108170055 | 0.107423947 |
| 8.734745 | -0.165885516 | 0.294411767 | -0.100637269 | 0.105793034 |
| 8.744986 | -0.16998784 | 0.296523195 | -0.092573428 | 0.104757306 |
| 8.755225 | -0.173870395 | 0.296902831 | -0.084791114 | 0.103586268 |
| 8.765466 | -0.177532457 | 0.295883635 | -0.076749326 | 0.103948682 |
| 8.775705 | -0.181241022 | 0.293554482 | -0.068671145 | 0.104296966 |
| 8.785945 | -0.183812623 | 0.290324508 | -0.060561178 | 0.10577629 |
| 8.796185 | -0.185802748 | 0.286688634 | -0.051633714 | 0.107711331 |
| 8.806425 | -0.1857616 | 0.283394088 | -0.042073857 | 0.109231653 |
| 8.816666 | -0.185448625 | 0.279890019 | -0.03155664 | 0.110683701 |
| 8.826905 | -0.182890188 | 0.277896434 | -0.020091589 | 0.110622047 |
| 8.837146 | -0.179880298 | 0.27552148 | -0.007441297 | 0.110217114 |
| 8.847385 | -0.175852648 | 0.274743026 | 0.005233987 | 0.108013607 |
| 8.857626 | -0.172186732 | 0.274096282 | 0.018456977 | 0.105440524 |
| 8.867866 | -0.168599419 | 0.273377975 | 0.031482274 | 0.102210132 |
| 8.878105 | -0.16660416 | 0.273163051 | 0.043252739 | 0.098859484 |
| 8.888346 | -0.165499882 | 0.272417347 | 0.054453214 | 0.095484602 |
| 8.898585 | -0.166297333 | 0.271065234 | 0.064462041 | 0.093342435 |
| 8.908826 | -0.168049493 | 0.268892481 | 0.073556099 | 0.092186181 |
| 8.919065 | -0.170510885 | 0.266124179 | 0.081419843 | 0.092206959 |
| 8.929306 | -0.172646286 | 0.262182765 | 0.088953088 | 0.09430135 |
| 8.939546 | -0.175002752 | 0.258857593 | 0.096203495 | 0.097821071 |
| 8.949786 | -0.175199082 | 0.253938726 | 0.103186902 | 0.102259477 |
| 8.960026 | -0.17505876 | 0.250537467 | 0.10953081 | 0.108134097 |
| 8.970266 | -0.172964454 | 0.246728895 | 0.116202467 | 0.113811698 |
| 8.980506 | -0.170007378 | 0.245008402 | 0.122055772 | 0.120099541 |
| 8.990746 | -0.165966659 | 0.243988389 | 0.127189658 | 0.125417345 |
| 9.000986 | -0.161609805 | 0.245245949 | 0.131551956 | 0.130052785 |
| 9.011226 | -0.156520455 | 0.247914338 | 0.135105447 | 0.133697176 |
| 9.021466 | -0.15151898 | 0.252882643 | 0.137280493 | 0.134964131 |
| 9.031706 | -0.146612003 | 0.25931875 | 0.138692906 | 0.135721132 |
| 9.041946 | -0.140974304 | 0.26741877 | 0.138993238 | 0.133237307 |
| 9.052186 | -0.136769476 | 0.276835155 | 0.138377364 | 0.129885275 |
| 9.062427 | -0.131279318 | 0.287454309 | 0.1369741 | 0.124010827 |
| 9.072666 | -0.127047124 | 0.297874593 | 0.134795391 | 0.117642157 |
| 9.082906 | -0.122016001 | 0.308956664 | 0.132422816 | 0.110713376 |
| 9.093146 | -0.117811599 | 0.317790785 | 0.12960289 | 0.104478689 |
| 9.103386 | -0.112675419 | 0.325636107 | 0.127166837 | 0.09903466 |
| 9.113626 | -0.107781843 | 0.330520922 | 0.125271437 | 0.096041767 |
| 9.123866 | -0.101953401 | 0.332707222 | 0.12420539 | 0.095584169 |
| 9.134107 | -0.096309726 | 0.332497782 | 0.12443542 | 0.096379296 |
| 9.144346 | -0.08897346 | 0.329768459 | 0.125781792 | 0.099768577 |
| 9.154587 | -0.082076738 | 0.326154987 | 0.128596855 | 0.104331994 |
| 9.164826 | -0.073670932 | 0.321699005 | 0.132006381 | 0.109963896 |
| 9.175066 | -0.06544098 | 0.317519568 | 0.136690933 | 0.116399663 |
| 9.185307 | -0.056105327 | 0.314456407 | 0.141644523 | 0.123694793 |
| 9.195546 | -0.046922653 | 0.312196418 | 0.146907416 | 0.131352122 |
| 9.205787 | -0.037254579 | 0.311685287 | 0.152272189 | 0.139777103 |
| 9.216026 | -0.027231098 | 0.31123863 | 0.157193954 | 0.148529077 |
| 9.226267 | -0.017711728 | 0.312252192 | 0.162405367 | 0.158062891 |
| 9.236506 | -0.007518848 | 0.313064485 | 0.166654415 | 0.168802055 |
| 9.246747 | 0.0025059 | 0.313956298 | 0.17065034 | 0.180184283 |
| 9.256987 | 0.012548242 | 0.314290008 | 0.173719369 | 0.192530233 |
| 9.267227 | 0.022018353 | 0.313777447 | 0.176553202 | 0.205541034 |
| 9.277467 | 0.031947619 | 0.312298968 | 0.177303347 | 0.219108657 |
| 9.287706 | 0.04086489 | 0.309947806 | 0.178239985 | 0.232148532 |
| 9.297947 | 0.049124133 | 0.305561335 | 0.177293455 | 0.244401729 |
| 9.308187 | 0.056170173 | 0.300464214 | 0.176104818 | 0.25583559 |
| 9.318427 | 0.062723028 | 0.294028092 | 0.173598397 | 0.265265152 |
| 9.328667 | 0.067331333 | 0.286087636 | 0.170967687 | 0.273532141 |
| 9.338907 | 0.071108745 | 0.277342862 | 0.167283461 | 0.279436135 |
| 9.349147 | 0.07399372 | 0.267783596 | 0.163441872 | 0.284059996 |
| 9.359387 | 0.076108798 | 0.257292471 | 0.158965096 | 0.287426219 |
| 9.369627 | 0.077613685 | 0.246342453 | 0.153901384 | 0.289069796 |
| 9.379867 | 0.079454583 | 0.235338516 | 0.148654494 | 0.289983367 |
| 9.390107 | 0.080793599 | 0.223910494 | 0.142562751 | 0.289697523 |
| 9.400347 | 0.083287084 | 0.213743099 | 0.136642514 | 0.289194589 |
| 9.410587 | 0.085486716 | 0.203634898 | 0.130672182 | 0.287872136 |
| 9.420827 | 0.088865579 | 0.195608208 | 0.124342523 | 0.286469569 |
| 9.431067 | 0.092822945 | 0.188675831 | 0.119300452 | 0.285294416 |
| 9.441307 | 0.096944449 | 0.184697702 | 0.114182213 | 0.284809855 |
| 9.451548 | 0.102347063 | 0.182130356 | 0.110118661 | 0.284497125 |
| 9.461787 | 0.107203191 | 0.182090724 | 0.107153429 | 0.285850776 |
| 9.472028 | 0.1124889 | 0.183955306 | 0.104050679 | 0.287368514 |
| 9.482267 | 0.117224321 | 0.187410153 | 0.102374818 | 0.290121179 |
| 9.492507 | 0.121302913 | 0.191995633 | 0.100206785 | 0.292500865 |
| 9.502748 | 0.124462259 | 0.197824841 | 0.098520534 | 0.295066264 |
| 9.512987 | 0.126427239 | 0.203785847 | 0.096038534 | 0.297011197 |
| 9.523228 | 0.12718149 | 0.210117473 | 0.093724967 | 0.297875668 |
| 9.533467 | 0.127550365 | 0.216288344 | 0.090409791 | 0.298241288 |
| 9.543708 | 0.12669281 | 0.222203503 | 0.086922842 | 0.2972869 |
| 9.553947 | 0.126622528 | 0.227855944 | 0.08256835 | 0.296355376 |
| 9.564188 | 0.125381606 | 0.232625918 | 0.078374011 | 0.295076084 |

| | | | | |
|---|---|---|---|---|
| 7.168021 | -0.082617256 | 0.3391536 | -0.029166084 | 0.247733986 |
| 7.178261 | -0.080564024 | 0.338339518 | -0.017409124 | 0.248234758 |
| 7.188501 | -0.077355026 | 0.337845426 | -0.00549498 | 0.248989048 |
| 7.198741 | -0.072885167 | 0.337626466 | 0.006684348 | 0.248780052 |
| 7.208981 | -0.067461845 | 0.337584702 | 0.018604176 | 0.248929084 |
| 7.219221 | -0.060917067 | 0.338799078 | 0.030202332 | 0.248867924 |
| 7.229461 | -0.053250133 | 0.339516162 | 0.041504038 | 0.24882236 |
| 7.239701 | -0.045264743 | 0.341570038 | 0.051979666 | 0.249275862 |
| 7.249941 | -0.035741424 | 0.342222662 | 0.062060177 | 0.249608045 |
| 7.260181 | -0.025935748 | 0.342850836 | 0.071368231 | 0.250849039 |
| 7.270421 | -0.014830326 | 0.34209993 | 0.079980721 | 0.251874977 |
| 7.280661 | -0.003521939 | 0.340076204 | 0.088026146 | 0.253128838 |
| 7.290901 | 0.008695801 | 0.336800069 | 0.095648661 | 0.254377525 |
| 7.301141 | 0.021141738 | 0.332768732 | 0.102689091 | 0.255126123 |
| 7.311381 | 0.033204207 | 0.327870233 | 0.109474419 | 0.255232372 |
| 7.321621 | 0.044908494 | 0.322997085 | 0.11555327 | 0.254979428 |
| 7.331861 | 0.055773045 | 0.318083895 | 0.121544266 | 0.253174661 |
| 7.342101 | 0.065239818 | 0.313730385 | 0.126782462 | 0.251080006 |
| 7.352341 | 0.073513294 | 0.309933556 | 0.131936051 | 0.247967517 |
| 7.362581 | 0.080344518 | 0.306549931 | 0.136571733 | 0.244235454 |
| 7.372821 | 0.086304851 | 0.304145128 | 0.141005531 | 0.240183592 |
| 7.383061 | 0.091204899 | 0.301595325 | 0.145532709 | 0.235911409 |
| 7.393301 | 0.095699749 | 0.299997771 | 0.150233981 | 0.232037038 |
| 7.403542 | 0.100124567 | 0.297241615 | 0.155357086 | 0.228061455 |
| 7.413781 | 0.104369264 | 0.294488994 | 0.161666206 | 0.224489376 |
| 7.424021 | 0.108965012 | 0.29076673 | 0.168609647 | 0.221200978 |
| 7.434261 | 0.113776206 | 0.28565237 | 0.176995696 | 0.218018872 |
| 7.444501 | 0.118702178 | 0.279694712 | 0.186606981 | 0.214898089 |
| 7.454741 | 0.124084213 | 0.272722888 | 0.196736487 | 0.212163658 |
| 7.464982 | 0.129606216 | 0.266057115 | 0.207598653 | 0.209509927 |
| 7.475222 | 0.135278165 | 0.258764839 | 0.218599452 | 0.20721669 |
| 7.485462 | 0.140923284 | 0.252868324 | 0.229174167 | 0.205827202 |
| 7.495702 | 0.147227377 | 0.247842877 | 0.238655159 | 0.204876876 |
| 7.505942 | 0.153234915 | 0.244130772 | 0.247514762 | 0.205253275 |
| 7.516181 | 0.159607328 | 0.241791825 | 0.254983622 | 0.206124136 |
| 7.526422 | 0.166108715 | 0.240676623 | 0.261389995 | 0.208173043 |
| 7.536662 | 0.172604771 | 0.240787858 | 0.266717331 | 0.210320075 |
| 7.546902 | 0.179085232 | 0.24134603 | 0.271377441 | 0.213290513 |
| 7.557142 | 0.18532415 | 0.242567778 | 0.274725207 | 0.215844895 |
| 7.567382 | 0.190935295 | 0.244250303 | 0.27772424 | 0.21844047 |
| 7.577622 | 0.196090038 | 0.246251254 | 0.279612369 | 0.220497138 |
| 7.587862 | 0.199966583 | 0.248254625 | 0.28054011 | 0.222038785 |
| 7.598102 | 0.203242752 | 0.250444578 | 0.280585678 | 0.223216776 |
| 7.608342 | 0.205234366 | 0.252472054 | 0.27946418 | 0.223259845 |
| 7.618582 | 0.206348084 | 0.254074771 | 0.277435471 | 0.223144946 |
| 7.628822 | 0.206564124 | 0.255210682 | 0.27404641 | 0.222619816 |
| 7.639062 | 0.206094942 | 0.255269666 | 0.270173137 | 0.222277928 |
| 7.649302 | 0.205282836 | 0.254430708 | 0.265073099 | 0.221330451 |
| 7.659542 | 0.204217982 | 0.252914935 | 0.260047816 | 0.22148795 |
| 7.669782 | 0.203020217 | 0.250441535 | 0.254454444 | 0.22157694 |
| 7.680022 | 0.201770631 | 0.248082315 | 0.249235671 | 0.222397402 |
| 7.690262 | 0.200966719 | 0.245217773 | 0.24471381 | 0.223420705 |
| 7.700502 | 0.200215421 | 0.24342051 | 0.240884717 | 0.224873176 |
| 7.710742 | 0.199896521 | 0.241956136 | 0.238286617 | 0.226086034 |
| 7.720982 | 0.199550714 | 0.241573118 | 0.237091523 | 0.227526664 |
| 7.731222 | 0.199520698 | 0.242439352 | 0.236948797 | 0.22882113 |
| 7.741462 | 0.199435663 | 0.244240372 | 0.237983696 | 0.229824921 |
| 7.751702 | 0.198696697 | 0.247464886 | 0.239678912 | 0.231280256 |
| 7.761942 | 0.197622155 | 0.250941175 | 0.241718358 | 0.233262965 |
| 7.772182 | 0.195696019 | 0.255695737 | 0.243511232 | 0.235701022 |
| 7.782423 | 0.193286192 | 0.259610449 | 0.244680475 | 0.239314312 |
| 7.792663 | 0.189810576 | 0.264016179 | 0.244809452 | 0.243485009 |
| 7.802903 | 0.186411104 | 0.267366994 | 0.243579173 | 0.24853864 |
| 7.813143 | 0.181908686 | 0.270379619 | 0.240769266 | 0.254052394 |
| 7.823382 | 0.178138818 | 0.272430019 | 0.236419143 | 0.259155729 |
| 7.833622 | 0.173429937 | 0.274572573 | 0.230717959 | 0.264193463 |
| 7.843863 | 0.169066814 | 0.276000546 | 0.223656266 | 0.268463218 |
| 7.854103 | 0.163812792 | 0.278256394 | 0.2158236 | 0.271497761 |
| 7.864343 | 0.158380457 | 0.280506715 | 0.207249237 | 0.27342996 |
| 7.874583 | 0.152036514 | 0.282965755 | 0.198727552 | 0.274144638 |
| 7.884823 | 0.144797409 | 0.285985761 | 0.189964853 | 0.273541263 |
| 7.895063 | 0.137016 | 0.288716897 | 0.181593049 | 0.2718798 |
| 7.905303 | 0.128839533 | 0.291467992 | 0.173492526 | 0.269381433 |
| 7.915543 | 0.120641614 | 0.293318982 | 0.165608928 | 0.266070629 |
| 7.925783 | 0.112240239 | 0.294265793 | 0.157956236 | 0.262468377 |
| 7.936023 | 0.104797191 | 0.294448176 | 0.150333005 | 0.258885111 |
| 7.946263 | 0.097165305 | 0.293119401 | 0.143029053 | 0.255006967 |
| 7.956503 | 0.090569959 | 0.291126274 | 0.135680286 | 0.251961854 |
| 7.966743 | 0.083801373 | 0.287827561 | 0.12883792 | 0.248997274 |
| 7.976983 | 0.077388158 | 0.28419153 | 0.122229421 | 0.246990615 |
| 7.987223 | 0.07085043 | 0.279897301 | 0.116236216 | 0.245641624 |
| 7.997463 | 0.064248202 | 0.275253538 | 0.11073315 | 0.245676699 |
| 8.007703 | 0.057923279 | 0.270976311 | 0.105510794 | 0.246525372 |
| 8.017943 | 0.050956846 | 0.266520413 | 0.100733222 | 0.248525937 |
| 8.028183 | 0.045016264 | 0.263438856 | 0.095745971 | 0.2516254 |
| 8.038424 | 0.038172604 | 0.261004374 | 0.091040318 | 0.25584089 |
| 8.048663 | 0.03280361 | 0.260307521 | 0.08545567 | 0.260345739 |
| 8.058904 | 0.026595737 | 0.260890377 | 0.079904018 | 0.265805785 |
| 8.069143 | 0.021302664 | 0.263329907 | 0.073914235 | 0.27123328 |
| 8.079383 | 0.015843435 | 0.266775825 | 0.067923576 | 0.277174059 |
| 8.089623 | 0.010796074 | 0.271317397 | 0.061464748 | 0.282417965 |
| 8.099863 | 0.005669812 | 0.275916008 | 0.055511069 | 0.287536015 |
| 8.110104 | 0.000670126 | 0.280360126 | 0.049876328 | 0.292116161 |
| 8.120343 | -0.004272183 | 0.283860691 | 0.044584004 | 0.296028124 |
| 8.130584 | -0.009301518 | 0.285737261 | 0.039755158 | 0.298822051 |
| 8.140823 | -0.014325082 | 0.286360658 | 0.035627381 | 0.301453806 |
| 8.151064 | -0.019615931 | 0.284798401 | 0.031636467 | 0.303210573 |
| 8.161304 | -0.024827198 | 0.281672988 | 0.02745273 | 0.304417532 |
| 8.171543 | -0.030167748 | 0.277155835 | 0.02362387 | 0.305655943 |
| 8.181784 | -0.035839669 | 0.271492757 | 0.019000277 | 0.306941263 |
| 8.192023 | -0.041417307 | 0.265285815 | 0.014036424 | 0.308970002 |
| 8.202264 | -0.047359978 | 0.258865278 | 0.008619179 | 0.311692475 |
| 8.212503 | -0.053154741 | 0.252710212 | 0.003139982 | 0.315379443 |
| 8.222744 | -0.058941146 | 0.246655058 | -0.002640251 | 0.319801577 |
| 8.232984 | -0.064841337 | 0.241568612 | -0.007876178 | 0.324372292 |
| 8.243224 | -0.069456674 | 0.236941635 | -0.012477498 | 0.329048292 |
| 8.253464 | -0.074202134 | 0.23335442 | -0.016713511 | 0.332576896 |
| 8.263704 | -0.077936111 | 0.229952074 | -0.020054617 | 0.335262953 |
| 8.273944 | -0.080765862 | 0.227874055 | -0.022939324 | 0.336390907 |
| 8.284184 | -0.083415309 | 0.225870881 | -0.025692014 | 0.336645981 |
| 8.294424 | -0.085511235 | 0.224741935 | -0.028462681 | 0.33605082 |
| 8.304664 | -0.088041055 | 0.223688741 | -0.030778238 | 0.334908927 |
| 8.314904 | -0.090589725 | 0.223306776 | -0.033702145 | 0.333371771 |
| 8.325144 | -0.093760803 | 0.222848289 | -0.035949198 | 0.331937955 |
| 8.335384 | -0.09716972 | 0.222492928 | -0.03822847 | 0.329806571 |
| 8.345624 | -0.101005861 | 0.222269806 | -0.039897011 | 0.328021452 |
| 8.355865 | -0.104662796 | 0.221934458 | -0.040967257 | 0.325338326 |
| 8.366104 | -0.108589713 | 0.222038616 | -0.041986026 | 0.323267521 |
| 8.376344 | -0.111794543 | 0.222428913 | -0.042453035 | 0.320735845 |
| 8.386584 | -0.115120611 | 0.223212781 | -0.0434328 | 0.319021235 |
| 8.396824 | -0.118212109 | 0.224495172 | -0.044816159 | 0.317879976 |
| 8.407064 | -0.121340876 | 0.22632732 | -0.047107835 | 0.317495526 |
| 8.417304 | -0.124401074 | 0.228265132 | -0.050233797 | 0.318329797 |
| 8.427545 | -0.127810479 | 0.230091454 | -0.054234652 | 0.320136323 |
| 8.437784 | -0.130960496 | 0.231135446 | -0.059146157 | 0.323075804 |
| 8.448025 | -0.134335314 | 0.231384369 | -0.064504634 | 0.326673703 |
| 8.458264 | -0.137257604 | 0.230584305 | -0.069784959 | 0.331119198 |
| 8.468505 | -0.139689539 | 0.228104182 | -0.075229527 | 0.335905448 |
| 8.478745 | -0.141727755 | 0.225222248 | -0.079923418 | 0.340868751 |
| 8.488984 | -0.142476634 | 0.221019337 | -0.0842184 | 0.345243883 |
| 8.499225 | -0.142957325 | 0.217186585 | -0.087823992 | 0.34912498 |
| 8.509464 | -0.142708683 | 0.21328757 | -0.090680307 | 0.352112035 |
| 8.519705 | -0.142185232 | 0.210663491 | -0.093026681 | 0.353685979 |
| 8.529944 | -0.141745196 | 0.208465774 | -0.095326249 | 0.354344639 |
| 8.540185 | -0.141800445 | 0.207891949 | -0.096944845 | 0.353923401 |
| 8.550425 | -0.142743616 | 0.207983606 | -0.099202383 | 0.353271486 |
| 8.560665 | -0.144479506 | 0.209041815 | -0.100781646 | 0.352409195 |
| 8.570905 | -0.147366729 | 0.210093507 | -0.102891162 | 0.352000598 |
| 8.581144 | -0.150981633 | 0.211508715 | -0.104273374 | 0.352093653 |
| 8.591385 | -0.15494413 | 0.212608544 | -0.105593871 | 0.353322065 |
| 8.601624 | -0.159022206 | 0.213844387 | -0.106453078 | 0.355152342 |
| 8.611865 | -0.163103349 | 0.214746821 | -0.10648316 | 0.357614191 |
| 8.622105 | -0.166612763 | 0.216403817 | -0.106364986 | 0.360204259 |
| 8.632345 | -0.169493388 | 0.218268357 | -0.105919951 | 0.362809431 |
| 8.642585 | -0.17181583 | 0.220986391 | -0.104990799 | 0.365353105 |
| 8.652825 | -0.173649871 | 0.224035383 | -0.104798922 | 0.366878382 |
| 8.663065 | -0.1753742 | 0.227851011 | -0.104467555 | 0.367759987 |
| 8.673305 | -0.176230321 | 0.231624896 | -0.104535073 | 0.367462272 |
| 8.683545 | -0.177506303 | 0.235041279 | -0.104845206 | 0.366140167 |
| 8.693785 | -0.178025352 | 0.237792239 | -0.105204477 | 0.362989786 |
| 8.704025 | -0.17885018 | 0.239550544 | -0.105635873 | 0.358727025 |
| 8.714265 | -0.179372328 | 0.240310101 | -0.1057599 | 0.352733313 |
| 8.724505 | -0.179681828 | 0.239135693 | -0.105931415 | 0.345193224 |
| 8.734745 | -0.18004379 | 0.236620659 | -0.105718048 | 0.336294005 |
| 8.744986 | -0.17976865 | 0.232793837 | -0.105646046 | 0.326058558 |
| 8.755225 | -0.179069467 | 0.227521387 | -0.105783972 | 0.3152064 |
| 8.765466 | -0.177640584 | 0.221968905 | -0.106015067 | 0.303127325 |
| 8.775705 | -0.17521605 | 0.21601181 | -0.106501992 | 0.291522753 |
| 8.785945 | -0.171662606 | 0.210664471 | -0.107234308 | 0.279342938 |
| 8.796185 | -0.16697467 | 0.205588204 | -0.108111567 | 0.268189886 |
| 8.806425 | -0.161733221 | 0.201018585 | -0.108810112 | 0.256894755 |
| 8.816666 | -0.155459074 | 0.196816825 | -0.10889764 | 0.247415308 |
| 8.826905 | -0.149618524 | 0.192780007 | -0.108710629 | 0.238302943 |
| 8.837146 | -0.143820599 | 0.18811857 | -0.107490617 | 0.231115797 |
| 8.847385 | -0.139175432 | 0.184081906 | -0.10555382 | 0.224529732 |
| 8.857626 | -0.135831981 | 0.179336983 | -0.102697041 | 0.220009984 |
| 8.867866 | -0.133915677 | 0.175594945 | -0.099165241 | 0.216616 |
| 8.878105 | -0.134019494 | 0.172422961 | -0.095123447 | 0.214533741 |
| 8.888346 | -0.134959356 | 0.17034646 | -0.090672904 | 0.213250727 |
| 8.898585 | -0.137427252 | 0.168795317 | -0.086040973 | 0.212589906 |
| 8.908826 | -0.140232636 | 0.168935796 | -0.081447186 | 0.211746633 |
| 8.919065 | -0.143188473 | 0.169617276 | -0.077131686 | 0.210150727 |
| 8.929306 | -0.145789103 | 0.17153871 | -0.072686293 | 0.207569685 |
| 8.939546 | -0.147264603 | 0.174146139 | -0.068873482 | 0.203113802 |
| 8.949786 | -0.147907794 | 0.17793116 | -0.065521556 | 0.197145411 |
| 8.960026 | -0.147243917 | 0.182969518 | -0.062414649 | 0.189698885 |
| 8.970266 | -0.145214026 | 0.188749096 | -0.060074236 | 0.181247758 |
| 8.980506 | -0.142327821 | 0.195152456 | -0.058140954 | 0.172247917 |
| 8.990746 | -0.138599306 | 0.202472078 | -0.056774217 | 0.163367925 |
| 9.000986 | -0.134261785 | 0.209290189 | -0.055811927 | 0.155655292 |
| 9.011226 | -0.129652976 | 0.216262194 | -0.054947052 | 0.149047237 |
| 9.021466 | -0.124947068 | 0.222056528 | -0.05467117 | 0.144019473 |
| 9.031706 | -0.120157167 | 0.226733999 | -0.054051609 | 0.140505441 |
| 9.041946 | -0.115588465 | 0.229687487 | -0.053790944 | 0.13865982 |
| 9.052186 | -0.110968182 | 0.231540705 | -0.053372406 | 0.137517601 |
| 9.062427 | -0.10644843 | 0.231684257 | -0.053080028 | 0.13755009 |
| 9.072666 | -0.101948461 | 0.230813247 | -0.053043132 | 0.138003323 |
| 9.082906 | -0.097490167 | 0.229812689 | -0.053105382 | 0.138615608 |
| 9.093146 | -0.092950619 | 0.228131125 | -0.05324693 | 0.13885662 |
| 9.103386 | -0.088569227 | 0.227359488 | -0.05389769 | 0.138889785 |
| 9.113626 | -0.084498733 | 0.226489064 | -0.054508645 | 0.13832883 |
| 9.123866 | -0.080576319 | 0.226152025 | -0.055174145 | 0.137158356 |
| 9.134107 | -0.077052141 | 0.225991173 | -0.055863444 | 0.13568837 |
| 9.144346 | -0.074354586 | 0.225610931 | -0.056368154 | 0.134531913 |
| 9.154587 | -0.072442948 | 0.224773965 | -0.056188023 | 0.133340994 |
| 9.164826 | -0.070941063 | 0.223730307 | -0.056067726 | 0.132553858 |
| 9.175066 | -0.070582699 | 0.22185143 | -0.054735544 | 0.132788242 |
| 9.185307 | -0.070275247 | 0.220202624 | -0.053581001 | 0.133736616 |
| 9.195546 | -0.070744121 | 0.218574847 | -0.051165477 | 0.136052404 |
| 9.205787 | -0.070894973 | 0.217226966 | -0.048848227 | 0.139431599 |
| 9.216026 | -0.071036939 | 0.216626907 | -0.045926571 | 0.144737041 |
| 9.226267 | -0.070817843 | 0.216273124 | -0.042814437 | 0.151075193 |
| 9.236506 | -0.07004243 | 0.216356852 | -0.039371864 | 0.159018945 |
| 9.246747 | -0.068944631 | 0.216365957 | -0.035878078 | 0.167764687 |
| 9.256987 | -0.067205308 | 0.21583979 | -0.032402325 | 0.176795157 |
| 9.267227 | -0.065716579 | 0.21475277 | -0.028838932 | 0.186008835 |
| 9.277467 | -0.063603955 | 0.212795616 | -0.025315401 | 0.194472537 |
| 9.287706 | -0.062006497 | 0.210366801 | -0.022083339 | 0.202334264 |
| 9.297947 | -0.060128852 | 0.207499964 | -0.018835872 | 0.20841486 |
| 9.308187 | -0.058694501 | 0.204526484 | -0.015921366 | 0.213375396 |
| 9.318427 | -0.057302479 | 0.201717312 | -0.012812097 | 0.216805674 |
| 9.328667 | -0.055831162 | 0.199322417 | -0.009675323 | 0.218558947 |
| 9.338907 | -0.054754687 | 0.197447407 | -0.006600575 | 0.218720494 |
| 9.349147 | -0.053428252 | 0.196010435 | -0.002918448 | 0.217920134 |
| 9.359387 | -0.05262739 | 0.19483018 | 0.000483767 | 0.216269872 |
| 9.369627 | -0.051755154 | 0.194549503 | 0.004607189 | 0.214216507 |
| 9.379867 | -0.051451851 | 0.19370127 | 0.008227338 | 0.212459356 |
| 9.390107 | -0.051699828 | 0.193522276 | 0.012078769 | 0.211120614 |
| 9.400347 | -0.052334079 | 0.192910159 | 0.015632371 | 0.210857595 |
| 9.410587 | -0.054103235 | 0.192621564 | 0.019270103 | 0.211408641 |
| 9.420827 | -0.056291895 | 0.191971448 | 0.021974945 | 0.213078442 |
| 9.431067 | -0.059485739 | 0.19133407 | 0.025189818 | 0.215790989 |
| 9.441307 | -0.063111711 | 0.190834461 | 0.027722983 | 0.218910428 |
| 9.451548 | -0.066865773 | 0.18996101 | 0.030175622 | 0.222536628 |
| 9.461787 | -0.070624624 | 0.189412234 | 0.032226541 | 0.226410579 |
| 9.472028 | -0.073857459 | 0.188398546 | 0.034238561 | 0.230315219 |
| 9.482267 | -0.076132758 | 0.187624243 | 0.036068413 | 0.233867165 |
| 9.492507 | -0.077385964 | 0.186601379 | 0.037339764 | 0.237471216 |
| 9.502748 | -0.077298897 | 0.18542359 | 0.038848114 | 0.240968139 |
| 9.512987 | -0.076508475 | 0.184262184 | 0.039940537 | 0.244196274 |
| 9.523228 | -0.074549446 | 0.183133707 | 0.041118406 | 0.247888584 |
| 9.533467 | -0.07262499 | 0.18161885 | 0.041955073 | 0.25149819 |
| 9.543708 | -0.070726263 | 0.181113305 | 0.042903563 | 0.255551179 |
| 9.553947 | -0.06937239 | 0.180623834 | 0.043888411 | 0.259282911 |
| 9.564188 | -0.06894957 | 0.181527726 | 0.044973735 | 0.263265551 |

| | | | | |
|---|---|---|---|---|
| 7.168021 | 0.457140135 | 0.232442217 | 0.39791033 | 0.32064483 |
| 7.178261 | 0.470072103 | 0.233392616 | 0.405190891 | 0.316745441 |
| 7.188501 | 0.483318249 | 0.234941139 | 0.411362428 | 0.31245925 |
| 7.198741 | 0.497571737 | 0.236682364 | 0.417037857 | 0.3081872 |
| 7.208981 | 0.511926475 | 0.239543254 | 0.421637896 | 0.303621415 |
| 7.219221 | 0.526620341 | 0.2424518 | 0.425688062 | 0.298916334 |
| 7.229461 | 0.541846241 | 0.246413753 | 0.429256163 | 0.294236221 |
| 7.239701 | 0.556627547 | 0.250208965 | 0.432514003 | 0.290290125 |
| 7.249941 | 0.57061223 | 0.254310722 | 0.435495248 | 0.287078013 |
| 7.260181 | 0.584506974 | 0.258663035 | 0.438742777 | 0.284667686 |
| 7.270421 | 0.596527005 | 0.262843329 | 0.441497476 | 0.283839407 |
| 7.280661 | 0.607710512 | 0.266452707 | 0.444448685 | 0.283782134 |
| 7.290901 | 0.616891369 | 0.269997168 | 0.447115219 | 0.284788539 |
| 7.301141 | 0.624251874 | 0.272841001 | 0.449139321 | 0.286375297 |
| 7.311381 | 0.629736175 | 0.274699408 | 0.450886485 | 0.288309882 |
| 7.321621 | 0.633208564 | 0.276048686 | 0.451900922 | 0.290335585 |
| 7.331861 | 0.635122013 | 0.2760606 | 0.452355802 | 0.291729094 |
| 7.342101 | 0.636214817 | 0.275331815 | 0.452480501 | 0.293525378 |
| 7.352341 | 0.63679149 | 0.27350247 | 0.452059183 | 0.294718366 |
| 7.362581 | 0.637349965 | 0.271256788 | 0.451163183 | 0.296062607 |
| 7.372821 | 0.638815434 | 0.268609466 | 0.450146703 | 0.297233116 |
| 7.383061 | 0.641096745 | 0.265307031 | 0.448312952 | 0.298993513 |
| 7.393301 | 0.64460246 | 0.262245511 | 0.445978086 | 0.300378959 |
| 7.403542 | 0.649012031 | 0.258581698 | 0.442704639 | 0.302498652 |
| 7.413781 | 0.653864169 | 0.254635016 | 0.438235516 | 0.30409237 |
| 7.424021 | 0.659343939 | 0.25009891 | 0.433244566 | 0.306497271 |
| 7.434261 | 0.664232922 | 0.244007474 | 0.426857239 | 0.308352024 |
| 7.444501 | 0.668321109 | 0.237628573 | 0.419905001 | 0.311172737 |
| 7.454741 | 0.671010633 | 0.22875654 | 0.412199919 | 0.313517016 |
| 7.464982 | 0.672310433 | 0.219887441 | 0.404175582 | 0.316619103 |
| 7.475222 | 0.671414945 | 0.209522682 | 0.39573279 | 0.319456872 |
| 7.485462 | 0.668514588 | 0.199825208 | 0.387569498 | 0.322622554 |
| 7.495702 | 0.663618094 | 0.190865094 | 0.379072539 | 0.325294409 |
| 7.505942 | 0.656972069 | 0.183516659 | 0.371610976 | 0.327662185 |
| 7.516181 | 0.648469025 | 0.179080759 | 0.363744082 | 0.329540634 |
| 7.526422 | 0.638796371 | 0.177002966 | 0.357229201 | 0.330622105 |
| 7.536662 | 0.628407535 | 0.177762852 | 0.351152211 | 0.331880066 |
| 7.546902 | 0.617499474 | 0.180741843 | 0.345635174 | 0.331715794 |
| 7.557142 | 0.606120224 | 0.185082172 | 0.340850268 | 0.332295028 |
| 7.567382 | 0.595305185 | 0.190250909 | 0.336500334 | 0.331378758 |
| 7.577622 | 0.584787129 | 0.195532204 | 0.332325989 | 0.331055162 |
| 7.587862 | 0.574673298 | 0.20118243 | 0.327927694 | 0.329218321 |
| 7.598102 | 0.565036602 | 0.206546736 | 0.323125659 | 0.327671844 |
| 7.608342 | 0.555557543 | 0.211805499 | 0.317689844 | 0.3247017 |
| 7.618582 | 0.546734849 | 0.216413463 | 0.311602612 | 0.321917449 |
| 7.628822 | 0.537382054 | 0.220500686 | 0.304359354 | 0.318072437 |
| 7.639062 | 0.528236769 | 0.223717708 | 0.296629044 | 0.314084129 |
| 7.649302 | 0.518582781 | 0.22593634 | 0.287861777 | 0.309555877 |
| 7.659542 | 0.509152418 | 0.226877954 | 0.278433178 | 0.304790174 |
| 7.669782 | 0.499288386 | 0.227587433 | 0.268069534 | 0.299398924 |
| 7.680022 | 0.489457008 | 0.227994622 | 0.257554774 | 0.293572848 |
| 7.690262 | 0.479927211 | 0.228519781 | 0.245962386 | 0.28756184 |
| 7.700502 | 0.469988275 | 0.230111233 | 0.234082304 | 0.281152365 |
| 7.710742 | 0.460164959 | 0.232661247 | 0.221738717 | 0.274339697 |
| 7.720982 | 0.449700061 | 0.236682456 | 0.209104494 | 0.267751403 |
| 7.731222 | 0.438852229 | 0.241968897 | 0.196463766 | 0.260753749 |
| 7.741462 | 0.426947174 | 0.248908896 | 0.183667934 | 0.25391342 |
| 7.751702 | 0.414508889 | 0.256573426 | 0.171162955 | 0.24674382 |
| 7.761942 | 0.401149113 | 0.265649447 | 0.158852449 | 0.239637554 |
| 7.772182 | 0.387623005 | 0.275018124 | 0.147161405 | 0.232146547 |
| 7.782423 | 0.373891866 | 0.285086586 | 0.135177251 | 0.224637891 |
| 7.792663 | 0.360388193 | 0.294786034 | 0.124366991 | 0.217150752 |
| 7.802903 | 0.347487223 | 0.304649658 | 0.113128821 | 0.210251975 |
| 7.813143 | 0.33504994 | 0.313426681 | 0.102426628 | 0.203856647 |
| 7.823382 | 0.323202077 | 0.321605708 | 0.091855559 | 0.198570339 |
| 7.833622 | 0.311456167 | 0.328184581 | 0.080984825 | 0.19469452 |
| 7.843863 | 0.299819829 | 0.333529962 | 0.070311766 | 0.191861331 |
| 7.854103 | 0.288112589 | 0.336712377 | 0.059359396 | 0.190462107 |
| 7.864343 | 0.276291109 | 0.338133348 | 0.048359199 | 0.18936485 |
| 7.874583 | 0.264044837 | 0.337579863 | 0.037236548 | 0.188590228 |
| 7.884823 | 0.252348763 | 0.334989618 | 0.026370824 | 0.187881993 |
| 7.895063 | 0.240983549 | 0.330940132 | 0.015070086 | 0.186321068 |
| 7.905303 | 0.230588129 | 0.325801913 | 0.004481022 | 0.185543188 |
| 7.915543 | 0.220732055 | 0.320034613 | -0.006304403 | 0.184521904 |
| 7.925783 | 0.212771024 | 0.314399417 | -0.016864125 | 0.184479247 |
| 7.936023 | 0.205386876 | 0.308638512 | -0.027325432 | 0.186740283 |
| 7.946263 | 0.198827125 | 0.303347479 | -0.037634669 | 0.190374268 |
| 7.956503 | 0.193238438 | 0.297946136 | -0.048317282 | 0.19715728 |
| 7.966743 | 0.187762654 | 0.292815331 | -0.058820242 | 0.205738555 |
| 7.976983 | 0.182811469 | 0.287631251 | -0.069152113 | 0.216489548 |
| 7.987223 | 0.177504467 | 0.282204406 | -0.079800705 | 0.22806137 |
| 7.997463 | 0.172893342 | 0.277229504 | -0.089675576 | 0.2399148 |
| 8.007703 | 0.168435528 | 0.272171613 | -0.099548348 | 0.2509569 |
| 8.017943 | 0.164349987 | 0.268124014 | -0.108171757 | 0.261698736 |
| 8.028183 | 0.161257322 | 0.264805668 | -0.11648202 | 0.270793203 |
| 8.038424 | 0.158967441 | 0.262232072 | -0.123621783 | 0.279797617 |
| 8.048663 | 0.157747294 | 0.260916285 | -0.130112263 | 0.2877962 |
| 8.058904 | 0.157265379 | 0.26033725 | -0.135516301 | 0.296005835 |
| 8.069143 | 0.157575646 | 0.260400719 | -0.140624107 | 0.303513273 |
| 8.079383 | 0.15816822 | 0.261244212 | -0.14483281 | 0.311421419 |
| 8.089623 | 0.158933086 | 0.262335747 | -0.148760281 | 0.318438514 |
| 8.099863 | 0.15883519 | 0.26398597 | -0.151806805 | 0.324987157 |
| 8.110104 | 0.158448638 | 0.265955651 | -0.15447324 | 0.330219705 |
| 8.120343 | 0.156361072 | 0.268012895 | -0.156313452 | 0.33398693 |
| 8.130584 | 0.153395254 | 0.269937511 | -0.157405543 | 0.336304323 |
| 8.140823 | 0.148493029 | 0.27195336 | -0.157669998 | 0.337003015 |
| 8.151064 | 0.142650227 | 0.273036705 | -0.157279683 | 0.336328944 |
| 8.161304 | 0.13511955 | 0.273938933 | -0.156248803 | 0.334772143 |
| 8.171543 | 0.126640623 | 0.274293197 | -0.155066422 | 0.332812029 |
| 8.181784 | 0.11717315 | 0.273738113 | -0.153473242 | 0.330871599 |
| 8.192023 | 0.10716561 | 0.273693643 | -0.152284923 | 0.329441451 |
| 8.202264 | 0.096483579 | 0.272703013 | -0.151300846 | 0.329084502 |
| 8.212503 | 0.08603584 | 0.27282353 | -0.150919291 | 0.329190632 |
| 8.222744 | 0.074987983 | 0.272687556 | -0.151366043 | 0.330527103 |
| 8.232984 | 0.064594916 | 0.273769645 | -0.152023857 | 0.332427714 |
| 8.243224 | 0.054662372 | 0.27474363 | -0.154012681 | 0.334995426 |
| 8.253464 | 0.044698921 | 0.277145381 | -0.156118178 | 0.338128505 |
| 8.263704 | 0.036062545 | 0.279587329 | -0.159041331 | 0.340946459 |
| 8.273944 | 0.027544649 | 0.283088186 | -0.161873627 | 0.344245542 |
| 8.284184 | 0.020631231 | 0.286722277 | -0.165534155 | 0.34729099 |
| 8.294424 | 0.013886823 | 0.290456526 | -0.168663255 | 0.350332918 |
| 8.304664 | 0.008886346 | 0.294188948 | -0.172244209 | 0.353203744 |
| 8.314904 | 0.004284477 | 0.296820644 | -0.17531476 | 0.356040774 |
| 8.325144 | 0.001022796 | 0.298491507 | -0.17823643 | 0.358934769 |
| 8.335384 | -0.00182345 | 0.298296508 | -0.180654118 | 0.361483156 |
| 8.345624 | -0.00384227 | 0.296371939 | -0.182405816 | 0.364183564 |
| 8.355865 | -0.005895743 | 0.292275316 | -0.183702119 | 0.366085179 |
| 8.366104 | -0.007747884 | 0.286752666 | -0.18382935 | 0.367997746 |
| 8.376344 | -0.01020655 | 0.279409582 | -0.183332422 | 0.369077096 |
| 8.386584 | -0.013504412 | 0.272115904 | -0.18220455 | 0.369015986 |
| 8.396824 | -0.017506148 | 0.263543055 | -0.180085453 | 0.368017318 |
| 8.407064 | -0.023048289 | 0.256798167 | -0.177646694 | 0.365501111 |
| 8.417304 | -0.030170396 | 0.250609813 | -0.174594561 | 0.361603013 |
| 8.427545 | -0.038215697 | 0.24664034 | -0.171271286 | 0.35633105 |
| 8.437784 | -0.048330559 | 0.244969097 | -0.167866797 | 0.350420081 |
| 8.448025 | -0.059281256 | 0.245149512 | -0.164011166 | 0.343707811 |
| 8.458264 | -0.071325735 | 0.247702831 | -0.160508876 | 0.33695789 |
| 8.468505 | -0.08401182 | 0.251242453 | -0.157187889 | 0.330573044 |
| 8.478745 | -0.09678517 | 0.255936746 | -0.154293216 | 0.324624718 |
| 8.488984 | -0.10982186 | 0.260401984 | -0.151729917 | 0.319278744 |
| 8.499225 | -0.122063805 | 0.264880136 | -0.15063961 | 0.31471535 |
| 8.509464 | -0.133981975 | 0.268768351 | -0.149776168 | 0.310130995 |
| 8.519705 | -0.145078997 | 0.271722915 | -0.150811598 | 0.306163446 |
| 8.529944 | -0.155469812 | 0.273900379 | -0.152154709 | 0.302086816 |
| 8.540185 | -0.164903865 | 0.275101026 | -0.154336018 | 0.297849465 |
| 8.550425 | -0.173830806 | 0.275248554 | -0.157209275 | 0.293497088 |
| 8.560665 | -0.182013571 | 0.274389004 | -0.159918385 | 0.289095811 |
| 8.570905 | -0.189595092 | 0.272950354 | -0.162189043 | 0.285293852 |
| 8.581144 | -0.196619662 | 0.270396103 | -0.164025974 | 0.281989683 |
| 8.591385 | -0.202931708 | 0.267752173 | -0.164856397 | 0.2800178 |
| 8.601624 | -0.208968892 | 0.264709949 | -0.164819452 | 0.279574539 |
| 8.611865 | -0.213829583 | 0.261638201 | -0.164141993 | 0.280601433 |
| 8.622105 | -0.218222049 | 0.259191152 | -0.162873834 | 0.282594221 |
| 8.632345 | -0.221820959 | 0.256782683 | -0.161736218 | 0.28623113 |
| 8.642585 | -0.224617443 | 0.25535725 | -0.160824846 | 0.289893154 |
| 8.652825 | -0.227139445 | 0.254818526 | -0.160607677 | 0.294007795 |
| 8.663065 | -0.228875916 | 0.254966692 | -0.161125726 | 0.29766498 |
| 8.673305 | -0.231074195 | 0.256041423 | -0.162527149 | 0.300770677 |
| 8.683545 | -0.232811873 | 0.25729136 | -0.164586395 | 0.303728416 |
| 8.693785 | -0.235273813 | 0.258834771 | -0.167129374 | 0.305722129 |
| 8.704025 | -0.23753837 | 0.259565609 | -0.169900012 | 0.307328505 |
| 8.714265 | -0.239846423 | 0.259582917 | -0.17237489 | 0.308700411 |
| 8.724505 | -0.24213015 | 0.258433348 | -0.174716513 | 0.309765948 |
| 8.734745 | -0.24349473 | 0.255982264 | -0.176522072 | 0.309982863 |
| 8.744986 | -0.243985414 | 0.252635579 | -0.177483509 | 0.309948809 |
| 8.755225 | -0.243412553 | 0.247926976 | -0.178363079 | 0.309281187 |
| 8.765466 | -0.241091541 | 0.242928246 | -0.178588355 | 0.307718902 |
| 8.775705 | -0.237633375 | 0.237226004 | -0.178351607 | 0.305821185 |
| 8.785945 | -0.232651002 | 0.23153166 | -0.178272744 | 0.303800261 |
| 8.796185 | -0.226856992 | 0.225463788 | -0.178136058 | 0.301000359 |
| 8.806425 | -0.220319028 | 0.219826813 | -0.178127579 | 0.298894893 |
| 8.816666 | -0.213873667 | 0.214182773 | -0.178246684 | 0.29609631 |
| 8.826905 | -0.207646142 | 0.2085458 | -0.17869981 | 0.293634658 |
| 8.837146 | -0.201896934 | 0.203677006 | -0.179221466 | 0.291310765 |
| 8.847385 | -0.197309854 | 0.198763999 | -0.17956305 | 0.288970186 |
| 8.857626 | -0.193832113 | 0.194530635 | -0.179813109 | 0.287018277 |
| 8.867866 | -0.190688872 | 0.190375825 | -0.179596895 | 0.28532986 |
| 8.878105 | -0.188411132 | 0.187217527 | -0.179031872 | 0.284146394 |
| 8.888346 | -0.186662053 | 0.184027872 | -0.177903531 | 0.28330031 |
| 8.898585 | -0.184838867 | 0.182216087 | -0.17670705 | 0.283471382 |
| 8.908826 | -0.183270447 | 0.180242876 | -0.175161781 | 0.283600749 |
| 8.919065 | -0.182067897 | 0.179215181 | -0.173491828 | 0.284514666 |
| 8.929306 | -0.180584075 | 0.178120617 | -0.171551005 | 0.28542389 |
| 8.939546 | -0.179988219 | 0.177421314 | -0.169830667 | 0.286390443 |
| 8.949786 | -0.17940591 | 0.175928493 | -0.167228051 | 0.287099415 |
| 8.960026 | -0.179211014 | 0.174794715 | -0.16464746 | 0.287409513 |
| 8.970266 | -0.179322029 | 0.172381834 | -0.161404339 | 0.288364168 |
| 8.980506 | -0.179394475 | 0.170331213 | -0.157758157 | 0.28843311 |
| 8.990746 | -0.179359996 | 0.167433202 | -0.153921618 | 0.289899328 |
| 9.000986 | -0.179103585 | 0.165070651 | -0.149944799 | 0.291131982 |
| 9.011226 | -0.17821004 | 0.162953882 | -0.145950058 | 0.293236754 |
| 9.021466 | -0.17695773 | 0.16208609 | -0.142332747 | 0.295322883 |
| 9.031706 | -0.17523651 | 0.161891029 | -0.139239117 | 0.298323813 |
| 9.041946 | -0.173029702 | 0.163465432 | -0.135961577 | 0.300689935 |
| 9.052186 | -0.170702586 | 0.165706759 | -0.133802136 | 0.303248428 |
| 9.062427 | -0.168293471 | 0.168992091 | -0.131987989 | 0.305814873 |
| 9.072666 | -0.166228178 | 0.1728251 | -0.130842882 | 0.308007277 |
| 9.082906 | -0.164350543 | 0.177262939 | -0.130966518 | 0.310568149 |
| 9.093146 | -0.16322666 | 0.181766444 | -0.131881 | 0.313361814 |
| 9.103386 | -0.162802162 | 0.186684957 | -0.134403708 | 0.316297065 |
| 9.113626 | -0.162921798 | 0.19182033 | -0.137774742 | 0.31954778 |
| 9.123866 | -0.164284347 | 0.197069451 | -0.142163225 | 0.322898963 |
| 9.134107 | -0.165986171 | 0.202181509 | -0.147509609 | 0.32529336 |
| 9.144346 | -0.168748235 | 0.206770938 | -0.153019274 | 0.327519955 |
| 9.154587 | -0.172534764 | 0.210473745 | -0.158600497 | 0.327580786 |
| 9.164826 | -0.176637561 | 0.213238214 | -0.163659963 | 0.326394708 |
| 9.175066 | -0.181413609 | 0.215046159 | -0.168654808 | 0.323400441 |
| 9.185307 | -0.186432459 | 0.21597224 | -0.172330004 | 0.318441735 |
| 9.195546 | -0.19141215 | 0.217132075 | -0.175626502 | 0.312209704 |
| 9.205787 | -0.19623707 | 0.218007435 | -0.177542343 | 0.304793002 |
| 9.216026 | -0.20040375 | 0.219421615 | -0.178412206 | 0.297035321 |
| 9.226267 | -0.204491725 | 0.221411176 | -0.177681921 | 0.289305696 |
| 9.236506 | -0.207842413 | 0.223728728 | -0.175802331 | 0.281821357 |
| 9.246747 | -0.211318322 | 0.226274282 | -0.172151429 | 0.275916953 |
| 9.256987 | -0.214429764 | 0.228660757 | -0.167257731 | 0.270630845 |
| 9.267227 | -0.217774566 | 0.230196594 | -0.161061064 | 0.267240427 |
| 9.277467 | -0.220775373 | 0.231427086 | -0.154042454 | 0.265463717 |
| 9.287706 | -0.223425904 | 0.231214762 | -0.146650824 | 0.264842366 |
| 9.297947 | -0.225288337 | 0.230505509 | -0.138861829 | 0.26548208 |
| 9.308187 | -0.225867119 | 0.229240881 | -0.131603235 | 0.267225299 |
| 9.318427 | -0.225620964 | 0.227596015 | -0.124718879 | 0.269152234 |
| 9.328667 | -0.223657881 | 0.226035177 | -0.11880841 | 0.271145375 |
| 9.338907 | -0.220622829 | 0.22406739 | -0.11367629 | 0.272956356 |
| 9.349147 | -0.216665368 | 0.222302263 | -0.109683599 | 0.274082925 |
| 9.359387 | -0.212246823 | 0.220446479 | -0.106543928 | 0.275057387 |
| 9.369627 | -0.207200094 | 0.218141362 | -0.104394145 | 0.275002415 |
| 9.379867 | -0.202489778 | 0.215851109 | -0.10278696 | 0.274576679 |
| 9.390107 | -0.19761207 | 0.212784661 | -0.101786098 | 0.273591135 |
| 9.400347 | -0.193451188 | 0.209741434 | -0.100760712 | 0.272359559 |
| 9.410587 | -0.18940973 | 0.206869784 | -0.099827397 | 0.270320398 |
| 9.420827 | -0.186186493 | 0.203828547 | -0.098287111 | 0.268926617 |
| 9.431067 | -0.183208471 | 0.201592424 | -0.096314793 | 0.267088126 |
| 9.441307 | -0.180984569 | 0.199996834 | -0.093490261 | 0.266223927 |
| 9.451548 | -0.179041543 | 0.199222798 | -0.090395442 | 0.265322518 |
| 9.461787 | -0.177281502 | 0.199228558 | -0.086229731 | 0.265593546 |
| 9.472028 | -0.1756965 | 0.199805671 | -0.081970828 | 0.265921745 |
| 9.482267 | -0.174082945 | 0.200953195 | -0.077101018 | 0.26689503 |
| 9.492507 | -0.172295712 | 0.202994521 | -0.072390424 | 0.267501341 |
| 9.502748 | -0.170494634 | 0.205246343 | -0.067609241 | 0.268143731 |
| 9.512987 | -0.168592411 | 0.208945247 | -0.062596442 | 0.268114778 |
| 9.523228 | -0.166182963 | 0.213053146 | -0.058035261 | 0.267312047 |
| 9.533467 | -0.163887459 | 0.217987227 | -0.05339983 | 0.2661313 |
| 9.543708 | -0.161373824 | 0.223259115 | -0.048851052 | 0.264013128 |
| 9.553947 | -0.15806866 | 0.228848314 | -0.044455455 | 0.261052802 |
| 9.564188 | -0.154521168 | 0.233806767 | -0.040158778 | 0.25746554 |

| | | | | | | | | | | | | | | | | | | | |
|---|---|---|---|---|---|---|---|---|---|---|---|---|---|---|---|---|---|---|---|
| 9.574428 | 0.048150481 | 0.305859137 | 0.082284717 | 0.16695095 | 9.574428 | 0.126100221 | 0.237470822 | 0.073654667 | 0.294350121 | 9.574428 | -0.069590886 | 0.183549711 | 0.0461996 | 0.266707413 | 9.574428 | -0.150119299 | 0.238680613 | -0.036139049 | 0.253258478 |
| 9.584667 | 0.054655463 | 0.305121936 | 0.076038457 | 0.16918029 | 9.584667 | 0.126166359 | 0.241139603 | 0.069418728 | 0.294251861 | 9.584667 | -0.071125462 | 0.186760457 | 0.04808609 | 0.270212016 | 9.584667 | -0.144464116 | 0.242526743 | -0.032090551 | 0.248565556 |
| 9.594908 | 0.060385132 | 0.302585284 | 0.070800574 | 0.17150791 | 9.594908 | 0.127511052 | 0.245354557 | 0.06537433 | 0.295130865 | 9.594908 | -0.073527056 | 0.191183203 | 0.050437279 | 0.273184376 | 9.594908 | -0.138133986 | 0.245678645 | -0.028492601 | 0.242939424 |
| 9.605147 | 0.065603328 | 0.298318456 | 0.065364912 | 0.17455708 | 9.605147 | 0.128185588 | 0.248013722 | 0.062398688 | 0.29661861 | 9.605147 | -0.076199973 | 0.195812656 | 0.053444051 | 0.276474528 | 9.605147 | -0.130622757 | 0.248006759 | -0.024751554 | 0.237573063 |
| 9.615388 | 0.070482935 | 0.292916772 | 0.061553647 | 0.1772633 | 9.615388 | 0.129599019 | 0.25083682 | 0.060215673 | 0.298741309 | 9.615388 | -0.079300947 | 0.200637825 | 0.057305706 | 0.279108683 | 9.615388 | -0.122259528 | 0.250160417 | -0.021738711 | 0.232413581 |
| 9.625628 | 0.07449045 | 0.287026959 | 0.0580914 | 0.18011954 | 9.625628 | 0.129937132 | 0.253017044 | 0.059654632 | 0.300813491 | 9.625628 | -0.082080283 | 0.205029657 | 0.061871308 | 0.282065368 | 9.625628 | -0.113605582 | 0.251900913 | -0.018108468 | 0.227578388 |
| 9.635868 | 0.078055882 | 0.280014947 | 0.055560679 | 0.18226146 | 9.635868 | 0.130010901 | 0.255097537 | 0.060304222 | 0.30305737 | 9.635868 | -0.084708763 | 0.208811761 | 0.066801331 | 0.284202634 | 9.635868 | -0.104636817 | 0.253630843 | -0.015395816 | 0.224036076 |
| 9.646108 | 0.080616383 | 0.273360301 | 0.0526988 | 0.18438728 | 9.646108 | 0.129199487 | 0.256893391 | 0.06250073 | 0.304461625 | 9.646108 | -0.086446859 | 0.211217409 | 0.07256646 | 0.285932238 | 9.646108 | -0.096372097 | 0.25552604 | -0.01222231 | 0.221855235 |
| 9.656348 | 0.083158522 | 0.26582747 | 0.050119818 | 0.18668443 | 9.656348 | 0.127796985 | 0.258924477 | 0.065320919 | 0.304893835 | 9.656348 | -0.087886723 | 0.213499412 | 0.078304051 | 0.286837326 | 9.656348 | -0.088550009 | 0.257538707 | -0.009395631 | 0.221621792 |
| 9.666588 | 0.084461444 | 0.258624979 | 0.047376608 | 0.1881742 | 9.666588 | 0.125944643 | 0.261547143 | 0.069326548 | 0.305063656 | 9.666588 | -0.088412157 | 0.215170234 | 0.084072528 | 0.286459888 | 9.666588 | -0.081872361 | 0.258534983 | -0.006441286 | 0.222955317 |
| 9.676828 | 0.085776115 | 0.250541234 | 0.04449578 | 0.18991827 | 9.676828 | 0.123789817 | 0.264120436 | 0.073603531 | 0.303814011 | 9.676828 | -0.088103788 | 0.216642072 | 0.089765201 | 0.285522701 | 9.676828 | -0.076417413 | 0.259405426 | -0.003311139 | 0.226726027 |
| 9.687068 | 0.086850711 | 0.24240368 | 0.041975492 | 0.19132932 | 9.687068 | 0.122184837 | 0.267074539 | 0.07788203 | 0.302011432 | 9.687068 | -0.087232867 | 0.218732977 | 0.094922913 | 0.283393495 | 9.687068 | -0.07172313 | 0.25884809 | -0.000269752 | 0.232036313 |
| 9.697308 | 0.087055919 | 0.233941276 | 0.039963671 | 0.191873 | 9.697308 | 0.12040844 | 0.269962395 | 0.082398453 | 0.2991414 | 9.697308 | -0.085081166 | 0.22130761 | 0.099312334 | 0.280956128 | 9.697308 | -0.067883723 | 0.257087092 | 0.003142586 | 0.23828403 |
| 9.707548 | 0.087499507 | 0.225191395 | 0.038621764 | 0.19233513 | 9.707548 | 0.119679217 | 0.272546319 | 0.086409581 | 0.295616645 | 9.707548 | -0.082898069 | 0.224500186 | 0.103342741 | 0.278172565 | 9.707548 | -0.064168814 | 0.253947066 | 0.007035295 | 0.245756484 |
| 9.717788 | 0.086913131 | 0.215802413 | 0.038257358 | 0.19139571 | 9.717788 | 0.119666999 | 0.274301832 | 0.090782904 | 0.291489908 | 9.717788 | -0.079565975 | 0.228647975 | 0.105924163 | 0.275995536 | 9.717788 | -0.060957149 | 0.250344957 | 0.010906191 | 0.25260047 |
| 9.728028 | 0.086172682 | 0.206379144 | 0.039092357 | 0.19052111 | 9.728028 | 0.119962569 | 0.275367098 | 0.094746786 | 0.286517253 | 9.728028 | -0.076487659 | 0.233149188 | 0.107869384 | 0.273841932 | 9.728028 | -0.056896847 | 0.245960937 | 0.01520369 | 0.258909773 |
| 9.738268 | 0.084444508 | 0.196623396 | 0.040545319 | 0.18946843 | 9.738268 | 0.121295202 | 0.275359169 | 0.099261874 | 0.280695828 | 9.738268 | -0.073052982 | 0.237923507 | 0.108130336 | 0.272703878 | 9.738268 | -0.053248734 | 0.242116134 | 0.019466901 | 0.263895274 |
| 9.748508 | 0.081530826 | 0.186480442 | 0.042632803 | 0.18847126 | 9.748508 | 0.12239791 | 0.274037963 | 0.103612204 | 0.274381046 | 9.748508 | -0.070481284 | 0.242937588 | 0.107710738 | 0.271851328 | 9.748508 | -0.048888005 | 0.239254966 | 0.023564187 | 0.266888431 |
| 9.758748 | 0.078178717 | 0.176790086 | 0.044921626 | 0.18833069 | 9.758748 | 0.123874983 | 0.272048863 | 0.107942746 | 0.268254964 | 9.758748 | -0.068281583 | 0.247497026 | 0.105574817 | 0.271999255 | 9.758748 | -0.044436902 | 0.237826676 | 0.027491339 | 0.268066862 |
| 9.768989 | 0.073632529 | 0.167514997 | 0.047177435 | 0.18873712 | 9.768989 | 0.124340372 | 0.268533897 | 0.11200557 | 0.261260829 | 9.768989 | -0.067202597 | 0.251936784 | 0.102349484 | 0.272587939 | 9.768989 | -0.039965668 | 0.238250548 | 0.030577199 | 0.266911246 |
| 9.779228 | 0.0680612 | 0.159718416 | 0.048768436 | 0.19036094 | 9.779228 | 0.124609971 | 0.264587596 | 0.11536103 | 0.255402482 | 9.779228 | -0.067169272 | 0.255592114 | 0.097559878 | 0.273819343 | 9.779228 | -0.03549647 | 0.240962161 | 0.033467941 | 0.264609531 |
| 9.789468 | 0.062123977 | 0.153710515 | 0.049712835 | 0.1928196 | 9.789468 | 0.123249958 | 0.259782184 | 0.11781151 | 0.249879267 | 9.789468 | -0.068007912 | 0.259075359 | 0.091959921 | 0.275893793 | 9.789468 | -0.031292181 | 0.2455338 | 0.035821695 | 0.260926963 |
| 9.799708 | 0.055659837 | 0.149372413 | 0.049863905 | 0.19569081 | 9.799708 | 0.121254775 | 0.254637957 | 0.118859805 | 0.245795529 | 9.799708 | -0.069625657 | 0.261273174 | 0.084578366 | 0.278650305 | 9.799708 | -0.027185912 | 0.252007365 | 0.038050301 | 0.256694488 |
| 9.809948 | 0.048984477 | 0.146629582 | 0.048537941 | 0.19952531 | 9.809948 | 0.117675389 | 0.249438703 | 0.118533622 | 0.24224586 | 9.809948 | -0.071725275 | 0.263323406 | 0.076285442 | 0.28203166 | 9.809948 | -0.023411963 | 0.259136528 | 0.040722826 | 0.253144228 |
| 9.820189 | 0.042507714 | 0.145583602 | 0.046801945 | 0.20338776 | 9.820189 | 0.113983862 | 0.244699471 | 0.117229288 | 0.240506892 | 9.820189 | -0.073892096 | 0.264893552 | 0.067133745 | 0.285859367 | 9.820189 | -0.020333033 | 0.267369947 | 0.043530384 | 0.249472277 |
| 9.830428 | 0.036021112 | 0.144810254 | 0.043201681 | 0.20737406 | 9.830428 | 0.109295378 | 0.240265168 | 0.114062887 | 0.239689825 | 9.830428 | -0.075766128 | 0.266163967 | 0.057108835 | 0.290411133 | 9.830428 | -0.017815195 | 0.275062868 | 0.04729559 | 0.247549979 |
| 9.840669 | 0.030661321 | 0.144319117 | 0.039401781 | 0.21150147 | 9.840669 | 0.104873384 | 0.237320738 | 0.110711076 | 0.239461705 | 9.840669 | -0.077238885 | 0.267431049 | 0.046821097 | 0.294712067 | 9.840669 | -0.016011847 | 0.282545597 | 0.05219679 | 0.246144573 |
| 9.850908 | 0.025113568 | 0.143200103 | 0.034157681 | 0.21509306 | 9.850908 | 0.100624962 | 0.235398346 | 0.106481976 | 0.240112699 | 9.850908 | -0.077797915 | 0.26910466 | 0.036508474 | 0.29972027 | 9.850908 | -0.015388333 | 0.288928659 | 0.057574685 | 0.245896554 |
| 9.861149 | 0.020544889 | 0.141721829 | 0.028451758 | 0.2188508 | 9.861149 | 0.097352949 | 0.234940254 | 0.102847955 | 0.240259385 | 9.861149 | -0.077851556 | 0.271203687 | 0.027030964 | 0.303712796 | 9.861149 | -0.015707041 | 0.294469923 | 0.063826193 | 0.246690404 |
| 9.871388 | 0.016213008 | 0.139595072 | 0.022256065 | 0.22135048 | 9.871388 | 0.094533884 | 0.236174998 | 0.098773115 | 0.240691792 | 9.871388 | -0.077027843 | 0.273574583 | 0.018097531 | 0.307987171 | 9.871388 | -0.01699436 | 0.299010693 | 0.070514799 | 0.247756671 |
| 9.881628 | 0.012329225 | 0.13668913 | 0.015831207 | 0.22410418 | 9.881628 | 0.093513596 | 0.239140927 | 0.096350202 | 0.240249992 | 9.881628 | -0.076520901 | 0.276550226 | 0.011255775 | 0.311043948 | 9.881628 | -0.019343624 | 0.302909423 | 0.077275174 | 0.249498565 |
| 9.891869 | 0.00828726 | 0.13374474 | 0.009277802 | 0.22537447 | 9.891869 | 0.092882617 | 0.243668458 | 0.094357571 | 0.239683502 | 9.891869 | -0.075164046 | 0.279138683 | 0.005130241 | 0.313555763 | 9.891869 | -0.022066516 | 0.306798652 | 0.08396971 | 0.251537685 |
| 9.902108 | 0.004358847 | 0.130014951 | 0.003338015 | 0.22615908 | 9.902108 | 0.093158613 | 0.249645928 | 0.093473285 | 0.238163709 | 9.902108 | -0.074617085 | 0.28201261 | 0.001420975 | 0.314930527 | 9.902108 | -0.025368004 | 0.309877366 | 0.09004819 | 0.253296503 |
| 9.912349 | -2.72652E-05 | 0.126670555 | -0.002172201 | 0.22574823 | 9.912349 | 0.094497959 | 0.257436323 | 0.093421261 | 0.236715057 | 9.912349 | -0.07367196 | 0.283790458 | -0.001069401 | 0.315261001 | 9.912349 | -0.02817594 | 0.313500863 | 0.0957201 | 0.255219034 |
| 9.922588 | -0.004615972 | 0.121772104 | -0.006576979 | 0.22402024 | 9.922588 | 0.095683571 | 0.266287714 | 0.093985185 | 0.234816461 | 9.922588 | -0.073727252 | 0.284855373 | -0.002220662 | 0.31445327 | 9.922588 | -0.031060351 | 0.316669534 | 0.100690781 | 0.2567384 |
| 9.932829 | -0.009977536 | 0.117535924 | -0.01035907 | 0.22116068 | 9.932829 | 0.097909514 | 0.276010449 | 0.094923328 | 0.233008688 | 9.932829 | -0.073140354 | 0.284746516 | -0.002350525 | 0.312976818 | 9.932829 | -0.032661361 | 0.319859307 | 0.104788685 | 0.257914016 |
| 9.943069 | -0.015356892 | 0.11235718 | -0.013479586 | 0.21689455 | 9.943069 | 0.099691502 | 0.285844483 | 0.095508324 | 0.230692611 | 9.943069 | -0.07239719 | 0.283081025 | -0.001660665 | 0.310427926 | 9.943069 | -0.033812167 | 0.322356358 | 0.108588891 | 0.259106023 |
| 9.953309 | -0.021381366 | 0.107311158 | -0.015541239 | 0.21149989 | 9.953309 | 0.102367811 | 0.29538744 | 0.095567354 | 0.228482761 | 9.953309 | -0.071048038 | 0.280394112 | -0.000725388 | 0.307463268 | 9.953309 | -0.033584908 | 0.323947485 | 0.111567949 | 0.259859566 |
| 9.963549 | -0.026927634 | 0.102164787 | -0.017608331 | 0.20535773 | 9.963549 | 0.105215166 | 0.303512443 | 0.094960815 | 0.226465001 | 9.963549 | -0.069163929 | 0.276198527 | 9.4444E-05 | 0.303892875 | 9.963549 | -0.032829077 | 0.324728144 | 0.114103782 | 0.260978329 |
| 9.973789 | -0.032849577 | 0.097714748 | -0.018839708 | 0.19831013 | 9.973789 | 0.10899897 | 0.310333804 | 0.092894501 | 0.223858684 | 9.973789 | -0.066187523 | 0.270906387 | 0.000638103 | 0.300368786 | 9.973789 | -0.030906948 | 0.323846002 | 0.116340663 | 0.262000846 |
| 9.984029 | -0.037994474 | 0.093383839 | -0.020266186 | 0.19134294 | 9.984029 | 0.112779864 | 0.31482604 | 0.090015112 | 0.221447425 | 9.984029 | -0.062401649 | 0.265002665 | 0.000693023 | 0.296392891 | 9.984029 | -0.028644558 | 0.321913347 | 0.118195027 | 0.263549338 |
| 9.994268 | -0.042858718 | 0.090265681 | -0.021117285 | 0.18410962 | 9.994268 | 0.117253665 | 0.317323194 | 0.086159357 | 0.219188896 | 9.994268 | -0.058514246 | 0.258144043 | -0.000426468 | 0.292678807 | 9.994268 | -0.02580854 | 0.31884138 | 0.119688155 | 0.264739385 |
| 10.00451 | -0.046934616 | 0.087225094 | -0.022141907 | 0.1773562 | 10.00451 | 0.12219186 | 0.317517331 | 0.081753783 | 0.216689016 | 10.00451 | -0.054322707 | 0.251260103 | -0.002039538 | 0.288478773 | 10.00451 | -0.022643275 | 0.314253809 | 0.120954522 | 0.266316627 |
| 10.01475 | -0.050382982 | 0.085406522 | -0.023133303 | 0.17182841 | 10.01475 | 0.127042226 | 0.315581738 | 0.076895696 | 0.213791605 | 10.01475 | -0.050439803 | 0.244402765 | -0.004778747 | 0.284662658 | 10.01475 | -0.019411325 | 0.309312352 | 0.121725134 | 0.267551767 |
| 10.02499 | -0.053654448 | 0.083716136 | -0.024039977 | 0.16693588 | 10.02499 | 0.13138479 | 0.312549015 | 0.072774413 | 0.2106839 | 10.02499 | -0.047235468 | 0.238125782 | -0.007930366 | 0.280853293 | 10.02499 | -0.015623277 | 0.303601589 | 0.122537814 | 0.267766506 |
| 10.03523 | -0.056037924 | 0.08331223 | -0.025393146 | 0.16364616 | 10.03523 | 0.135819774 | 0.307778131 | 0.069391077 | 0.205735208 | 10.03523 | -0.044763423 | 0.232665458 | -0.011609749 | 0.277044238 | 10.03523 | -0.011924204 | 0.29728097 | 0.122806465 | 0.267278228 |
| 10.04547 | -0.058528535 | 0.084568389 | -0.027666849 | 0.16202943 | 10.04547 | 0.139005437 | 0.303164005 | 0.067680499 | 0.200342514 | 10.04547 | -0.043150308 | 0.228240182 | -0.01531162 | 0.274016019 | 10.04547 | -0.007279244 | 0.291079813 | 0.122781806 | 0.265903296 |
| 10.05571 | -0.060462397 | 0.087425176 | -0.030703281 | 0.16082185 | 10.05571 | 0.141825262 | 0.298018195 | 0.06736822 | 0.192875868 | 10.05571 | -0.042471087 | 0.22547889 | -0.018882892 | 0.271230933 | 10.05571 | -0.002035972 | 0.284697844 | 0.123143634 | 0.263383977 |
| 10.06595 | -0.062355867 | 0.092211037 | -0.03484905 | 0.16124193 | 10.06595 | 0.143708033 | 0.294102626 | 0.069278504 | 0.184200093 | 10.06595 | -0.042482274 | 0.223479976 | -0.02200111 | 0.268948908 | 10.06595 | 0.003902919 | 0.278789674 | 0.123102006 | 0.260007138 |
| 10.07619 | -0.063922625 | 0.098626244 | -0.040279053 | 0.16189336 | 10.07619 | 0.144214626 | 0.290579935 | 0.072725855 | 0.174681208 | 10.07619 | -0.042884026 | 0.222851739 | -0.024736684 | 0.267625809 | 10.07619 | 0.01091172 | 0.272780747 | 0.123757458 | 0.256749849 |
| 10.08643 | -0.064948308 | 0.106564183 | -0.04695081 | 0.1631434 | 10.08643 | 0.143949383 | 0.288376433 | 0.077818688 | 0.164495008 | 10.08643 | -0.04403315 | 0.222677686 | -0.027445124 | 0.266179683 | 10.08643 | 0.0193356 | 0.268337442 | 0.124281247 | 0.253294511 |
| 10.09667 | -0.066177105 | 0.114695601 | -0.054471182 | 0.16477238 | 10.09667 | 0.142121582 | 0.286992228 | 0.083975043 | 0.15465846 | 10.09667 | -0.044695618 | 0.223845046 | -0.029478161 | 0.265657575 | 10.09667 | 0.027875058 | 0.264211725 | 0.125659275 | 0.250824681 |
| 10.10691 | -0.066884805 | 0.123300279 | -0.062622048 | 0.16646446 | 10.10691 | 0.138780287 | 0.285998529 | 0.090677431 | 0.145963707 | 10.10691 | -0.046046267 | 0.225033904 | -0.031549918 | 0.264007617 | 10.10691 | 0.037843161 | 0.262055892 | 0.126983878 | 0.249337376 |
| 10.11715 | -0.067384978 | 0.131221065 | -0.070789189 | 0.16874036 | 10.11715 | 0.134001252 | 0.285784305 | 0.097796012 | 0.138361283 | 10.11715 | -0.046411249 | 0.226991459 | -0.033481326 | 0.263127069 | 10.11715 | 0.047373166 | 0.260910766 | 0.128536114 | 0.249421266 |
| 10.12739 | -0.068112727 | 0.139256469 | -0.079135548 | 0.17125066 | 10.12739 | 0.128263969 | 0.285585672 | 0.104374798 | 0.132849263 | 10.12739 | -0.046905765 | 0.228937548 | -0.035561296 | 0.260505115 | 10.12739 | 0.05710822 | 0.261434343 | 0.130230597 | 0.250865002 |
| 10.13763 | -0.06917304 | 0.14603548 | -0.086898804 | 0.17382707 | 10.13763 | 0.121076239 | 0.285265618 | 0.11121305 | 0.129419127 | 10.13763 | -0.046849601 | 0.231300123 | -0.037524617 | 0.25802139 | 10.13763 | 0.065389753 | 0.263155282 | 0.131283358 | 0.253891409 |
| 10.14787 | -0.070777992 | 0.15312089 | -0.093700427 | 0.17730234 | 10.14787 | 0.113854821 | 0.284899985 | 0.116810792 | 0.12794857 | 10.14787 | -0.046957791 | 0.233347791 | -0.039528805 | 0.253959838 | 10.14787 | 0.07285087 | 0.265337678 | 0.13215161 | 0.258071044 |
| 10.15811 | -0.072714732 | 0.159088235 | -0.09995794 | 0.17982293 | 10.15811 | 0.10639558 | 0.28494755 | 0.122121508 | 0.128419621 | 10.15811 | -0.046418855 | 0.234826379 | -0.041876474 | 0.249822149 | 10.15811 | 0.07807111 | 0.26856755 | 0.132188629 | 0.26361128 |
| 10.16835 | -0.075647619 | 0.165001549 | -0.105124698 | 0.18311784 | 10.16835 | 0.099288725 | 0.284957477 | 0.126772815 | 0.130605375 | 10.16835 | -0.046628053 | 0.235730558 | -0.044460215 | 0.244466982 | 10.16835 | 0.081901783 | 0.27138986 | 0.131768424 | 0.269608514 |
| 10.17859 | -0.079004937 | 0.169873707 | -0.108743436 | 0.18564153 | 10.17859 | 0.093417237 | 0.285709356 | 0.130157117 | 0.134710136 | 10.17859 | -0.046922057 | 0.23573479 | -0.047526837 | 0.239246802 | 10.17859 | 0.0835276 | 0.274336746 | 0.131034261 | 0.275939007 |
| 10.18883 | -0.082492748 | 0.17365361 | -0.111264695 | 0.18750012 | 10.18883 | 0.088245146 | 0.286509987 | 0.133074891 | 0.140249118 | 10.18883 | -0.047292858 | 0.234770086 | -0.051076107 | 0.233713644 | 10.18883 | 0.083836426 | 0.276984017 | 0.129960809 | 0.282550326 |
| 10.19907 | -0.086227951 | 0.176843327 | -0.112031772 | 0.18907965 | 10.19907 | 0.084502659 | 0.287805696 | 0.134227382 | 0.147274302 | 10.19907 | -0.048681387 | 0.233281364 | -0.054868975 | 0.228532636 | 10.19907 | 0.083301619 | 0.279114169 | 0.129149291 | 0.288770947 |
| 10.20931 | -0.089492303 | 0.177289483 | -0.111613302 | 0.18927485 | 10.20931 | 0.081486281 | 0.288971712 | 0.13497889 | 0.155286173 | 10.20931 | -0.049718264 | 0.231101397 | -0.059221262 | 0.224032325 | 10.20931 | 0.081926073 | 0.280754209 | 0.128418789 | 0.294174704 |
| 10.21955 | -0.091933912 | 0.176926421 | -0.109020479 | 0.18910264 | 10.21955 | 0.079955593 | 0.289884989 | 0.134020899 | 0.164470418 | 10.21955 | -0.051235931 | 0.228687964 | -0.063183459 | 0.220098066 | 10.21955 | 0.080610329 | 0.282354189 | 0.128774959 | 0.299263957 |
| 10.22979 | -0.093357605 | 0.174275198 | -0.106094737 | 0.187446 | 10.22979 | 0.078866462 | 0.290117807 | 0.132615318 | 0.173334188 | 10.22979 | -0.052257453 | 0.226322102 | -0.067124236 | 0.217650237 | 10.22979 | 0.079476012 | 0.283410385 | 0.129117244 | 0.303414443 |
| 10.24003 | -0.093236104 | 0.170341414 | -0.101369873 | 0.1854873 | 10.24003 | 0.07827362 | 0.28967503 | 0.130439263 | 0.18231006 | 10.24003 | -0.053044856 | 0.223590486 | -0.070326375 | 0.216075971 | 10.24003 | 0.078736284 | 0.284193222 | 0.130891233 | 0.307200582 |
| 10.25027 | -0.09238725 | 0.164310543 | -0.096605246 | 0.18241875 | 10.25027 | 0.078661339 | 0.28823878 | 0.128275795 | 0.190300201 | 10.25027 | -0.053071282 | 0.220922085 | -0.072608349 | 0.215755937 | 10.25027 | 0.078378422 | 0.284787629 | 0.132795462 | 0.310755402 |
| 10.26051 | -0.090113417 | 0.15822444 | -0.091283368 | 0.17980754 | 10.26051 | 0.079116321 | 0.28571818 | 0.126300473 | 0.196211648 | 10.26051 | -0.052568491 | 0.218462272 | -0.073851802 | 0.216584577 | 10.26051 | 0.0783143 | 0.285506199 | 0.135014319 | 0.314340138 |
| 10.27075 | -0.087295859 | 0.150740106 | -0.086337034 | 0.1762719 | 10.27075 | 0.080595007 | 0.282943449 | 0.124784615 | 0.200690174 | 10.27075 | -0.051520997 | 0.216068924 | -0.07426194 | 0.218317992 | 10.27075 | 0.078473037 | 0.285929153 | 0.137309369 | 0.317867696 |
| 10.28099 | -0.084423993 | 0.143536364 | -0.082003613 | 0.1733592 | 10.28099 | 0.082484017 | 0.27912612 | 0.124109939 | 0.203316321 | 10.28099 | -0.050301428 | 0.214289655 | -0.073303884 | 0.221243078 | 10.28099 | 0.078690744 | 0.287022013 | 0.139096401 | 0.321570621 |
| 10.29123 | -0.082225226 | 0.136417811 | -0.078326288 | 0.17024897 | 10.29123 | 0.085324613 | 0.275385123 | 0.124512588 | 0.203835956 | 10.29123 | -0.049036537 | 0.213162055 | -0.071697061 | 0.224490511 | 10.29123 | 0.078480304 | 0.28844626 | 0.140533247 | 0.324932726 |
| 10.30147 | -0.080575859 | 0.130309725 | -0.075842299 | 0.16872336 | 10.30147 | 0.089287746 | 0.271612639 | 0.125659729 | 0.20332581 | 10.30147 | -0.047927303 | 0.212695894 | -0.069245478 | 0.228798089 | 10.30147 | 0.078506881 | 0.290851899 | 0.140827215 | 0.327908297 |
| 10.31171 | -0.08027785 | 0.124545379 | -0.073755197 | 0.16690788 | 10.31171 | 0.09338527 | 0.267975457 | 0.127884354 | 0.201405749 | 10.31171 | -0.047954693 | 0.213180466 | -0.06632814 | 0.232983683 | 10.31171 | 0.07836307 | 0.294028174 | 0.140357549 | 0.329913085 |
| 10.32195 | -0.080989231 | 0.120491637 | -0.072534851 | 0.16746327 | 10.32195 | 0.098591183 | 0.264701989 | 0.131206816 | 0.198660447 | 10.32195 | -0.0480178 | 0.214461378 | -0.062687891 | 0.237705477 | 10.32195 | 0.078078231 | 0.298056303 | 0.139312101 | 0.331104079 |
| 10.33219 | -0.083511193 | 0.116946664 | -0.07183512 | 0.16889216 | 10.33219 | 0.10364255 | 0.262034489 | 0.135050742 | 0.195621398 | 10.33219 | -0.049134323 | 0.216029424 | -0.059033696 | 0.241481543 | 10.33219 | 0.078179076 | 0.302719975 | 0.137097829 | 0.331562633 |
| 10.34243 | -0.086139845 | 0.114817792 | -0.071094363 | 0.17241454 | 10.34243 | 0.108474341 | 0.259683624 | 0.140233882 | 0.19272675 | 10.34243 | -0.05040623 | 0.218272657 | -0.054918715 | 0.245025592 | 10.34243 | 0.078437598 | 0.307395139 | 0.134662528 | 0.331075984 |
| 10.35267 | -0.089658868 | 0.113131952 | -0.07055333 | 0.17699574 | 10.35267 | 0.11281254 | 0.257471119 | 0.1458923 | 0.18961461 | 10.35267 | -0.05205668 | 0.220159961 | -0.050852643 | 0.247432704 | 10.35267 | 0.079337706 | 0.312108908 | 0.131662803 | 0.330644958 |
| 10.36291 | -0.092778004 | 0.112483106 | -0.070000411 | 0.18255858 | 10.36291 | 0.116004208 | 0.255815059 | 0.152276845 | 0.187577476 | 10.36291 | -0.053298895 | 0.221918614 | -0.046057197 | 0.249095791 | 10.36291 | 0.080496775 | 0.316230183 | 0.128471643 | 0.330105646 |
| 10.37315 | -0.095351819 | 0.111760122 | -0.069073285 | 0.18879388 | 10.37315 | 0.117979394 | 0.253467918 | 0.159071873 | 0.185352028 | 10.37315 | -0.054366591 | 0.222664223 | -0.041617885 | 0.249578726 | 10.37315 | 0.082182887 | 0.319575488 | 0.125205493 | 0.329976818 |
| 10.38339 | -0.09675008 | 0.111195234 | -0.06891798 | 0.19402226 | 10.38339 | 0.118593248 | 0.251134779 | 0.166097821 | 0.184785276 | 10.38339 | -0.05449759 | 0.222497431 | -0.0370259 | 0.249420831 | 10.38339 | 0.084030779 | 0.322695058 | 0.121887421 | 0.33012047 |
| 10.39363 | -0.096659857 | 0.110503643 | -0.067795523 | 0.19874809 | 10.39363 | 0.11784112 | 0.247355677 | 0.173115507 | 0.184905543 | 10.39363 | -0.054565437 | 0.221282173 | -0.0324566 | 0.248214267 | 10.39363 | 0.085521612 | 0.324942653 | 0.119409864 | 0.331045764 |
| 10.40387 | -0.09505407 | 0.109435882 | -0.067829258 | 0.2007414 | 10.40387 | 0.115973163 | 0.24293225 | 0.179284016 | 0.186024507 | 10.40387 | -0.053494265 | 0.218482726 | -0.028502825 | 0.24704809 | 10.40387 | 0.08663263 | 0.327221988 | 0.11686093 | 0.33171065 |
| 10.41411 | -0.091870191 | 0.108271414 | -0.067256094 | 0.2014355 | 10.41411 | 0.11288565 | 0.236895599 | 0.185295766 | 0.188684642 | 10.41411 | -0.052348046 | 0.214885262 | -0.024995344 | 0.245123411 | 10.41411 | 0.087143271 | 0.32915438 | 0.115432284 | 0.332692743 |
| 10.42435 | -0.087830584 | 0.107392529 | -0.066972442 | 0.19925058 | 10.42435 | 0.109085342 | 0.229685525 | 0.189629351 | 0.192011967 | 10.42435 | -0.050624535 | 0.210823186 | -0.02301794 | 0.242803717 | 10.42435 | 0.08596729 | 0.331008818 | 0.115133765 | 0.333339061 |
| 10.43459 | -0.082875165 | 0.106066811 | -0.066946177 | 0.19517549 | 10.43459 | 0.104581773 | 0.221675294 | 0.192811073 | 0.197099294 | 10.43459 | -0.049042921 | 0.206243898 | -0.021644489 | 0.240918746 | 10.43459 | 0.083870679 | 0.332793759 | 0.115702302 | 0.33341396 |
| 10.44483 | -0.078539378 | 0.105753193 | -0.066383405 | 0.19022391 | 10.44483 | 0.100369353 | 0.213581988 | 0.194769015 | 0.201675521 | 10.44483 | -0.046763211 | 0.202472217 | -0.021966639 | 0.238466299 | 10.44483 | 0.08019671 | 0.334125291 | 0.117520234 | 0.333169068 |
| 10.45507 | -0.07443412 | 0.105478061 | -0.066471438 | 0.18413853 | 10.45507 | 0.095792328 | 0.20650971 | 0.195218343 | 0.208072037 | 10.45507 | -0.044606076 | 0.199479421 | -0.022960015 | 0.236482115 | 10.45507 | 0.075655921 | 0.334999342 | 0.12012244 | 0.332528223 |
| 10.46531 | -0.071640475 | 0.106104527 | -0.065886424 | 0.17866229 | 10.46531 | 0.092942069 | 0.201013896 | 0.194581884 | 0.214296753 | 10.46531 | -0.042205282 | 0.197622914 | -0.025008341 | 0.234801563 | 10.46531 | 0.070099659 | 0.335161871 | 0.123168678 | 0.33181716 |
| 10.47555 | -0.070353806 | 0.107441281 | -0.066252319 | 0.17309349 | 10.47555 | 0.089658656 | 0.198252927 | 0.193030457 | 0.221195607 | 10.47555 | -0.039255853 | 0.197428707 | -0.027341564 | 0.233728742 | 10.47555 | 0.064225651 | 0.334646232 | 0.126514784 | 0.331107099 |
| 10.48579 | -0.070522807 | 0.109526183 | -0.066185988 | 0.16916298 | 10.48579 | 0.088717199 | 0.197879417 | 0.191064986 | 0.228182821 | 10.48579 | -0.036307093 | 0.198361723 | -0.029498032 | 0.233152692 | 10.48579 | 0.058261468 | 0.333405869 | 0.129320147 | 0.330945172 |
| 10.49603 | -0.07242945 | 0.112221295 | -0.066651428 | 0.16592006 | 10.49603 | 0.087650221 | 0.199979688 | 0.188986874 | 0.235255814 | 10.49603 | -0.032850057 | 0.200666793 | -0.031643372 | 0.233264985 | 10.49603 | 0.052622636 | 0.331998624 | 0.1317007 | 0.331266858 |
| 10.50627 | -0.074814562 | 0.114678043 | -0.06702646 | 0.16333282 | 10.50627 | 0.087901883 | 0.204007971 | 0.186651534 | 0.242724149 | 10.50627 | -0.029385995 | 0.203130627 | -0.032834331 | 0.2336861 | 10.50627 | 0.047125278 | 0.330356381 | 0.133283955 | 0.332109663 |
| 10.51651 | -0.07858479 | 0.117547698 | -0.06799591 | 0.16178658 | 10.51651 | 0.087954008 | 0.208370213 | 0.184890507 | 0.249525538 | 10.51651 | -0.025908221 | 0.206020018 | -0.033374804 | 0.234514766 | 10.51651 | 0.042246836 | 0.329283618 | 0.133954754 | 0.333763828 |
| 10.52675 | -0.083024252 | 0.1193248 | -0.068500881 | 0.16101636 | 10.52675 | 0.088393108 | 0.213280285 | 0.182801877 | 0.25653477 | 10.52675 | -0.022665328 | 0.208677192 | -0.033270285 | 0.234905837 | 10.52675 | 0.037283149 | 0.328988883 | 0.134276568 | 0.335227016 |
| 10.53699 | -0.087679117 | 0.121225735 | -0.069610608 | 0.15979113 | 10.53699 | 0.088247892 | 0.217466051 | 0.180972938 | 0.263386537 | 10.53699 | -0.019350025 | 0.210376449 | -0.032460544 | 0.23537485 | 10.53699 | 0.032178956 | 0.32940413 | 0.13407147 | 0.336672419 |
| 10.54723 | -0.092926625 | 0.12242343 | -0.071028731 | 0.15953799 | 10.54723 | 0.087730352 | 0.220569325 | 0.178485346 | 0.269185093 | 10.54723 | -0.016606639 | 0.211300997 | -0.031135875 | 0.235399953 | 10.54723 | 0.027344566 | 0.330883811 | 0.133940208 | 0.338268027 |
| 10.55747 | -0.098162733 | 0.12302937 | -0.072677267 | 0.15893552 | 10.55747 | 0.086365867 | 0.222601156 | 0.175676025 | 0.2745862 | 10.55747 | -0.013851852 | 0.21110727 | -0.029756746 | 0.235268696 | 10.55747 | 0.021476187 | 0.33237754 | 0.134053696 | 0.338997614 |
| 10.56771 | -0.103046065 | 0.123830483 | -0.075384977 | 0.15904065 | 10.56771 | 0.085068432 | 0.223414278 | 0.172166258 | 0.278443164 | 10.56771 | -0.011313119 | 0.210020831 | -0.028196695 | 0.234797089 | 10.56771 | 0.016058839 | 0.334386466 | 0.134735106 | 0.339623745 |
| 10.57795 | -0.108056691 | 0.123577283 | -0.078460313 | 0.15909622 | 10.57795 | 0.082956217 | 0.223728585 | 0.168119529 | 0.28175106 | 10.57795 | -0.009165248 | 0.20749209 | -0.026760554 | 0.234446995 | 10.57795 | 0.010172869 | 0.335412284 | 0.135807557 | 0.339976191 |
| 10.58819 | -0.111461568 | 0.123549214 | -0.081662886 | 0.15971416 | 10.58819 | 0.080586178 | 0.222528902 | 0.163934524 | 0.283044683 | 10.58819 | -0.007032469 | 0.204186538 | -0.025699124 | 0.234314365 | 10.58819 | 0.004194745 | 0.336067638 | 0.137680174 | 0.339721028 |
| 10.59843 | -0.114106049 | 0.123252075 | -0.08565399 | 0.16110505 | 10.59843 | 0.078291799 | 0.221461755 | 0.159273336 | 0.283746328 | 10.59843 | -0.005369236 | 0.200040064 | -0.024811054 | 0.233542725 | 10.59843 | -0.001128389 | 0.335499105 | 0.140114068 | 0.339537224 |
| 10.60867 | -0.115200013 | 0.123291351 | -0.088921944 | 0.16310956 | 10.60867 | 0.075456784 | 0.219667361 | 0.155779182 | 0.283206809 | 10.60867 | -0.003930561 | 0.195283171 | -0.024382506 | 0.233243445 | 10.60867 | -0.006474447 | 0.33421839 | 0.142606412 | 0.338637468 |
| 10.61891 | -0.11516111 | 0.124138582 | -0.091425336 | 0.16680193 | 10.61891 | 0.072410951 | 0.217252893 | 0.152019543 | 0.282504099 | 10.61891 | -0.003258828 | 0.189997306 | -0.023933908 | 0.232140558 | 10.61891 | -0.011182102 | 0.332240544 | 0.145595037 | 0.336700943 |
| 10.62915 | -0.113491094 | 0.12634241 | -0.093420086 | 0.17077102 | 10.62915 | 0.069223767 | 0.214075095 | 0.14946513 | 0.280311729 | 10.62915 | -0.002850897 | 0.184575234 | -0.023811345 | 0.231023015 | 10.62915 | -0.015010179 | 0.329551367 | 0.14806363 | 0.334253669 |
| 10.63939 | -0.111273849 | 0.129892228 | -0.094488557 | 0.17652177 | 10.63939 | 0.065796421 | 0.209479906 | 0.147654836 | 0.279022215 | 10.63939 | -0.003251092 | 0.179149645 | -0.023844217 | 0.228932819 | 10.63939 | -0.018367025 | 0.326553122 | 0.150493483 | 0.329622771 |
| 10.64963 | -0.108820098 | 0.134748179 | -0.095043827 | 0.18351108 | 10.64963 | 0.061785386 | 0.203982129 | 0.146544165 | 0.276431963 | 10.64963 | -0.003703316 | 0.173881196 | -0.023474996 | 0.226934567 | 10.64963 | -0.020193215 | 0.323562526 | 0.152234406 | 0.324407728 |
| 10.65987 | -0.105827427 | 0.140938792 | -0.095343863 | 0.19118007 | 10.65987 | 0.058114739 | 0.19653456 | 0.146100394 | 0.274034989 | 10.65987 | -0.004855168 | 0.169026423 | -0.023062032 | 0.224490832 | 10.65987 | -0.021357981 | 0.320438338 | 0.153404771 | 0.317371024 |
| 10.67011 | -0.103464897 | 0.147450819 | -0.095612158 | 0.1998271 | 10.67011 | 0.054294953 | 0.188404751 | 0.146404166 | 0.271328313 | 10.67011 | -0.005851703 | 0.164462062 | -0.022526375 | 0.221876924 | 10.67011 | -0.020771518 | 0.316585175 | 0.153475143 | 0.309804198 |
| 10.68035 | -0.100879329 | 0.155447308 | -0.096936528 | 0.20862705 | 10.68035 | 0.050619264 | 0.17988904 | 0.147371325 | 0.268508514 | 10.68035 | -0.007210452 | 0.160413302 | -0.0214851 | 0.219978911 | 10.68035 | -0.01914716 | 0.313465414 | 0.153088601 | 0.30130216 |
| 10.69059 | -0.098964314 | 0.162156936 | -0.098155984 | 0.21714322 | 10.69059 | 0.047405348 | 0.171736901 | 0.148406293 | 0.265605964 | 10.69059 | -0.008347118 | 0.156681622 | -0.020324354 | 0.218407041 | 10.69059 | -0.01605818 | 0.309428228 | 0.151507335 | 0.293238887 |
| 10.70083 | -0.096034342 | 0.169430553 | -0.100230473 | 0.2251065 | 10.70083 | 0.044987383 | 0.165174001 | 0.150736249 | 0.262285994 | 10.70083 | -0.00965376 | 0.153542874 | -0.018922596 | 0.217546764 | 10.70083 | -0.012604636 | 0.305494689 | 0.149228291 | 0.285193783 |
| 10.71107 | -0.093334301 | 0.175123657 | -0.102226245 | 0.23185369 | 10.71107 | 0.042887233 | 0.160569566 | 0.152861229 | 0.259509543 | 10.71107 | -0.01082203 | 0.150549818 | -0.01732483 | 0.217708922 | 10.71107 | -0.00804541 | 0.300908865 | 0.146026558 | 0.27848347 |
| 10.72131 | -0.089627255 | 0.179853499 | -0.10404569 | 0.23684511 | 10.72131 | 0.041865135 | 0.158579896 | 0.156062846 | 0.256017156 | 10.72131 | -0.012055045 | 0.148339283 | -0.015968684 | 0.218677408 | 10.72131 | -0.003909917 | 0.296202074 | 0.141738606 | 0.272280238 |
| 10.73155 | -0.085459213 | 0.181752501 | -0.104690344 | 0.24033619 | 10.73155 | 0.041142367 | 0.159081217 | 0.15823601 | 0.253175965 | 10.73155 | -0.013051598 | 0.146221999 | -0.01441511 | 0.220509194 | 10.73155 | 2.90535E-05 | 0.290916005 | 0.136944975 | 0.267917716 |
| 10.74179 | -0.080522478 | 0.182053384 | -0.104297388 | 0.24162434 | 10.74179 | 0.041333313 | 0.162071941 | 0.1611207 | 0.249459579 | 10.74179 | -0.013801495 | 0.145342822 | -0.012955688 | 0.223321386 | 10.74179 | 0.003361716 | 0.285014356 | 0.130684853 | 0.264397251 |
| 10.75203 | -0.075210551 | 0.179073789 | -0.101726722 | 0.24145748 | 10.75203 | 0.041527121 | 0.1669146 | 0.163203049 | 0.246307272 | 10.75203 | -0.014495738 | 0.144669752 | -0.011518314 | 0.226641609 | 10.75203 | 0.00582329 | 0.278584437 | 0.123949335 | 0.262488449 |
| 10.76227 | -0.069751438 | 0.173674109 | -0.09739569 | 0.23944415 | 10.76227 | 0.041540614 | 0.1734341 | 0.164122897 | 0.242480972 | 10.76227 | -0.015131258 | 0.145547593 | -0.010050792 | 0.230779448 | 10.76227 | 0.007409757 | 0.271696604 | 0.11616619 | 0.261416985 |
| 10.77251 | -0.064155437 | 0.165512821 | -0.090893794 | 0.23596794 | 10.77251 | 0.041354254 | 0.180332499 | 0.164377064 | 0.239152575 | 10.77251 | -0.014862592 | 0.146536623 | -0.00843027 | 0.235345266 | 10.77251 | 0.008460261 | 0.264522099 | 0.107588859 | 0.261302561 |
| 10.78275 | -0.058454855 | 0.155397857 | -0.082735989 | 0.23131581 | 10.78275 | 0.040483918 | 0.187536772 | 0.162991224 | 0.235281316 | 10.78275 | -0.01514357 | 0.148134429 | -0.006806374 | 0.240155963 | 10.78275 | 0.008838891 | 0.25743173 | 0.098938751 | 0.261610778 |
| 10.79299 | -0.053334718 | 0.144358069 | -0.072882908 | 0.22563949 | 10.79299 | 0.038781478 | 0.194328816 | 0.161090153 | 0.231676424 | 10.79299 | -0.014649981 | 0.150182322 | -0.004706313 | 0.245268641 | 10.79299 | 0.008752448 | 0.250319082 | 0.089842701 | 0.26251811 |
| 10.80323 | -0.048294998 | 0.132555217 | -0.062569513 | 0.2189747 | 10.80323 | 0.036379075 | 0.199524571 | 0.157578142 | 0.228146089 | 10.80323 | -0.013940027 | 0.151489319 | -0.003283129 | 0.249913516 | 10.80323 | 0.008340231 | 0.244124079 | 0.081197865 | 0.26338592 |
| 10.81347 | -0.043503514 | 0.122004964 | -0.051857984 | 0.21212259 | 10.81347 | 0.033101353 | 0.20358943 | 0.153155644 | 0.224697512 | 10.81347 | -0.012835224 | 0.15294017 | -0.001758444 | 0.254283973 | 10.81347 | 0.007435744 | 0.23852032 | 0.072794727 | 0.264656985 |
| 10.82371 | -0.039424283 | 0.112347217 | -0.041848541 | 0.2043073 | 10.82371 | 0.030018788 | 0.205830225 | 0.148101538 | 0.221370268 | 10.82371 | -0.011465582 | 0.153986797 | -0.001150002 | 0.257679233 | 10.82371 | 0.006458456 | 0.234089494 | 0.065123109 | 0.266084816 |
| 10.83395 | -0.035985485 | 0.105939501 | -0.033408014 | 0.19663432 | 10.83395 | 0.026038783 | 0.206369719 | 0.142566611 | 0.218621472 | 10.83395 | -0.009150334 | 0.154818892 | -0.00099635 | 0.259930295 | 10.83395 | 0.004483977 | 0.230829364 | 0.058382916 | 0.267681038 |
| 10.84419 | -0.03342914 | 0.10179211 | -0.026307543 | 0.18886597 | 10.84419 | 0.022794784 | 0.205196121 | 0.136736114 | 0.215657359 | 10.84419 | -0.006735545 | 0.155656079 | -0.001803099 | 0.260519481 | 10.84419 | 0.00250923 | 0.228537569 | 0.052267724 | 0.269392721 |
| 10.85443 | -0.031891314 | 0.100452091 | -0.021558404 | 0.18164736 | 10.85443 | 0.019403145 | 0.203230111 | 0.130534221 | 0.213308352 | 10.85443 | -0.003417447 | 0.156703118 | -0.003634243 | 0.25950111 | 10.85443 | -0.000204327 | 0.227245798 | 0.046655961 | 0.270639241 |
| 10.86467 | -0.031012974 | 0.101772648 | -0.018866793 | 0.17521046 | 10.86467 | 0.015970468 | 0.200018582 | 0.12487385 | 0.21065836 | 10.86467 | -0.000105402 | 0.157744039 | -0.005818525 | 0.256829051 | 10.86467 | -0.00341185 | 0.226960648 | 0.042315353 | 0.272032817 |
| 10.87491 | -0.031551274 | 0.104548335 | -0.018111659 | 0.17034893 | 10.87491 | 0.012614186 | 0.196452233 | 0.118994737 | 0.208392382 | 10.87491 | 0.003460334 | 0.159169915 | -0.008649967 | 0.252665811 | 10.87491 | -0.006899274 | 0.227012771 | 0.037816967 | 0.271839143 |
| 10.88515 | -0.032437291 | 0.107318409 | -0.018770433 | 0.1675789 | 10.88515 | 0.008572155 | 0.192848313 | 0.113165434 | 0.205577447 | 10.88515 | 0.007029684 | 0.160590109 | -0.011188676 | 0.24750869 | 10.88515 | -0.010331036 | 0.228267477 | 0.034210544 | 0.271404831 |
| 10.89539 | -0.033743375 | 0.110569416 | -0.021192695 | 0.16739985 | 10.89539 | 0.004598298 | 0.189877522 | 0.107627983 | 0.202970568 | 10.89539 | 0.009850677 | 0.162157409 | -0.013819152 | 0.241742834 | 10.89539 | -0.013994513 | 0.229475252 | 0.030853543 | 0.269386493 |
| 10.90563 | -0.034978212 | 0.112668176 | -0.02419508 | 0.1692904 | 10.90563 | -0.000680686 | 0.187592724 | 0.102392371 | 0.200628248 | 10.90563 | 0.012706133 | 0.163422889 | -0.015399347 | 0.236117519 | 10.90563 | -0.017245577 | 0.231501951 | 0.027699406 | 0.266599469 |
| 10.91587 | -0.036363565 | 0.114203683 | -0.027704721 | 0.17387284 | 10.91587 | -0.006156919 | 0.186688924 | 0.096792065 | 0.198124202 | 10.91587 | 0.014759333 | 0.164758441 | -0.016939625 | 0.230378471 | 10.91587 | -0.019942427 | 0.233707189 | 0.024746791 | 0.262493435 |
| 10.92611 | -0.037105414 | 0.11512213 | -0.031931211 | 0.18008514 | 10.92611 | -0.011849198 | 0.187401874 | 0.091872194 | 0.195569445 | 10.92611 | 0.016716496 | 0.165610834 | -0.016960805 | 0.225816091 | 10.92611 | -0.022366812 | 0.235675371 | 0.021730513 | 0.258030775 |
| 10.93635 | -0.037785656 | 0.115521906 | -0.036192388 | 0.18753713 | 10.93635 | -0.017461768 | 0.189264733 | 0.087366139 | 0.193691357 | 10.93635 | 0.018296134 | 0.166297199 | -0.016734344 | 0.221347727 | 10.93635 | -0.023975632 | 0.2378621 | 0.018641353 | 0.252728613 |
| 10.94659 | -0.0375769 | 0.116116011 | -0.040268824 | 0.19503282 | 10.94659 | -0.023249128 | 0.192553031 | 0.082799219 | 0.191521517 | 10.94659 | 0.020053 | 0.166663099 | -0.015642618 | 0.217737202 | 10.94659 | -0.025176591 | 0.239574218 | 0.015505284 | 0.247153856 |
| 10.95683 | -0.037539764 | 0.116871859 | -0.044480412 | 0.20270977 | 10.95683 | -0.027916487 | 0.1963567 | 0.079927205 | 0.189132533 | 10.95683 | 0.021669754 | 0.166557289 | -0.014470722 | 0.21444622 | 10.95683 | -0.025862802 | 0.240831809 | 0.012203126 | 0.241683986 |
| 10.96707 | -0.03647378 | 0.11805644 | -0.047498464 | 0.20965322 | 10.96707 | -0.032306199 | 0.200720456 | 0.077234135 | 0.186541289 | 10.96707 | 0.023803906 | 0.16613122 | -0.012726125 | 0.211329461 | 10.96707 | -0.02643034 | 0.241324966 | 0.008523256 | 0.236042343 |
| 10.97731 | -0.035534588 | 0.119437364 | -0.050066808 | 0.21532532 | 10.97731 | -0.035396456 | 0.205022077 | 0.076128748 | 0.183708174 | 10.97731 | 0.02583286 | 0.165295023 | -0.011226562 | 0.208280501 | 10.97731 | -0.02668063 | 0.241021094 | 0.005231379 | 0.231134982 |
| 10.98755 | -0.033612512 | 0.121115939 | -0.051474704 | 0.22023897 | 10.98755 | -0.038473742 | 0.208306067 | 0.075521232 | 0.180243239 | 10.98755 | 0.027883621 | 0.164207458 | -0.00969513 | 0.204981715 | 10.98755 | -0.027235879 | 0.240080167 | 0.001366029 | 0.226531611 |
| 10.99779 | -0.031564127 | 0.123607729 | -0.051703685 | 0.22381671 | 10.99779 | -0.040684597 | 0.211739795 | 0.076372408 | 0.176584365 | 10.99779 | 0.030316371 | 0.162495022 | -0.00817445 | 0.201919218 | 10.99779 | -0.027818074 | 0.237574088 | -0.00185723 | 0.222650958 |
| 11.00803 | -0.0291724 | 0.126397714 | -0.050497073 | 0.22676112 | 11.00803 | -0.04242345 | 0.214096055 | 0.077947971 | 0.172581523 | 11.00803 | 0.032014808 | 0.161275183 | -0.006930003 | 0.198689471 | 11.00803 | -0.028967284 | 0.23463711 | -0.005599106 | 0.219682116 |
| 11.01827 | -0.026062917 | 0.129995874 | -0.049138151 | 0.22847807 | 11.01827 | -0.044544999 | 0.21542845 | 0.08032351 | 0.168541739 | 11.01827 | 0.033965141 | 0.159367629 | -0.005349985 | 0.195530726 | 11.01827 | -0.030799702 | 0.230708224 | -0.008830138 | 0.217106271 |
| 11.02851 | -0.022834904 | 0.134720585 | -0.047010063 | 0.22886908 | 11.02851 | -0.046770031 | 0.2170566 | 0.083376398 | 0.164499674 | 11.02851 | 0.035437593 | 0.158104256 | -0.003886825 | 0.192860054 | 11.02851 | -0.032821757 | 0.226014784 | -0.012359354 | 0.216017347 |
| 11.03875 | -0.018965666 | 0.140225662 | -0.045280337 | 0.22834702 | 11.03875 | -0.049520272 | 0.218213665 | 0.087087988 | 0.160719815 | 11.03875 | 0.036875788 | 0.156440622 | -0.002522013 | 0.190670176 | 11.03875 | -0.035668662 | 0.221272507 | -0.015570151 | 0.215247881 |
| 11.04899 | -0.015509751 | 0.146061821 | -0.044675033 | 0.22611329 | 11.04899 | -0.053170125 | 0.219889168 | 0.090746819 | 0.1574009 | 11.04899 | 0.038217876 | 0.155281698 | -0.000863826 | 0.188587579 | 11.04899 | -0.038642386 | 0.216716345 | -0.018612471 | 0.215471039 |
| 11.05923 | -0.011645446 | 0.151827202 | -0.045051278 | 0.22312739 | 11.05923 | -0.05825027 | 0.222060322 | 0.095278372 | 0.154410056 | 11.05923 | 0.039442862 | 0.153782522 | 0.000100845 | 0.187345027 | 11.05923 | -0.042384675 | 0.212846416 | -0.021660174 | 0.216053154 |
| 11.06947 | -0.008517005 | 0.15808039 | -0.047464861 | 0.21870933 | 11.06947 | -0.064009233 | 0.225788446 | 0.099967173 | 0.152222317 | 11.06947 | 0.040803403 | 0.152681873 | 0.000947194 | 0.186005215 | 11.06947 | -0.04558749 | 0.209956371 | -0.023494205 | 0.21692721 |
| 11.07971 | -0.005700316 | 0.163531081 | -0.051525217 | 0.21388156 | 11.07971 | -0.071208656 | 0.231025547 | 0.104757749 | 0.150463064 | 11.07971 | 0.042282842 | 0.151295956 | 0.000926916 | 0.184983437 | 11.07971 | -0.049553834 | 0.207948696 | -0.025381979 | 0.217191627 |
| 11.08995 | -0.004201821 | 0.168308488 | -0.05697697 | 0.20952234 | 11.08995 | -0.078431089 | 0.237806042 | 0.109520755 | 0.149101486 | 11.08995 | 0.043700627 | 0.150000939 | 0.000298082 | 0.183724024 | 11.08995 | -0.053273454 | 0.2072741 | -0.025686796 | 0.2169711 |
| 11.10019 | -0.003932979 | 0.173480626 | -0.064215688 | 0.20481803 | 11.10019 | -0.086454725 | 0.246001327 | 0.114554535 | 0.14812762 | 11.10019 | 0.045034475 | 0.148632373 | -0.000977115 | 0.18261239 | 11.10019 | -0.057110843 | 0.207905263 | -0.025850214 | 0.215737964 |
| 11.11043 | -0.004593301 | 0.178087441 | -0.072009729 | 0.20179014 | 11.11043 | -0.093729299 | 0.255594437 | 0.118584517 | 0.146102419 | 11.11043 | 0.046412 | 0.14726701 | -0.002912275 | 0.181615848 | 11.11043 | -0.061352839 | 0.208941313 | -0.024929362 | 0.213362155 |
| 11.12067 | -0.007060052 | 0.18262258 | -0.080179741 | 0.19913045 | 11.12067 | -0.100254776 | 0.265848358 | 0.122039474 | 0.143330285 | 11.12067 | 0.047604979 | 0.145931995 | -0.005040085 | 0.180820458 | 11.12067 | -0.065725874 | 0.211111622 | -0.024125241 | 0.210143705 |
| 11.13091 | -0.009966041 | 0.186603025 | -0.088335635 | 0.19822785 | 11.13091 | -0.105534046 | 0.276092798 | 0.123946743 | 0.139217939 | 11.13091 | 0.048455133 | 0.144681804 | -0.00725532 | 0.18030482 | 11.13091 | -0.070634022 | 0.214180927 | -0.023144076 | 0.20646031 |
| 11.14115 | -0.013767191 | 0.189477788 | -0.095492803 | 0.19825595 | 11.14115 | -0.10960938 | 0.286094859 | 0.12476913 | 0.133945147 | 11.14115 | 0.048967731 | 0.143725946 | -0.009373524 | 0.180795656 | 11.14115 | -0.076105621 | 0.21739989 | -0.023103669 | 0.202158733 |
| 11.15139 | -0.017585096 | 0.191788833 | -0.101760949 | 0.20011129 | 11.15139 | -0.112454701 | 0.29567402 | 0.123469247 | 0.127331702 | 11.15139 | 0.049380249 | 0.143591029 | -0.011618712 | 0.183119159 | 11.15139 | -0.081986712 | 0.221543861 | -0.023857741 | 0.197851647 |
| 11.16163 | -0.021290277 | 0.191982607 | -0.106470096 | 0.20336753 | 11.16163 | -0.113864853 | 0.303227297 | 0.120466506 | 0.1213228 | 11.16163 | 0.049700631 | 0.144034938 | -0.013213128 | 0.182687673 | 11.16163 | -0.087730241 | 0.225417235 | -0.026070005 | 0.193851482 |
| 11.17187 | -0.024099411 | 0.191948148 | -0.109430872 | 0.20850187 | 11.17187 | -0.114577639 | 0.309798922 | 0.116349483 | 0.115111989 | 11.17187 | 0.04955883 | 0.146321405 | -0.015043619 | 0.184585222 | 11.17187 | -0.093868825 | 0.229625367 | -0.029543621 | 0.189780058 |
| 11.18211 | -0.026444297 | 0.190995414 | -0.110872169 | 0.21400262 | 11.18211 | -0.114333407 | 0.313505443 | 0.110908356 | 0.116670449 | 11.18211 | 0.050183127 | 0.148907081 | -0.016862978 | 0.187115211 | 11.18211 | -0.099541679 | 0.233319797 | -0.034545007 | 0.18648976 |
| 11.19235 | -0.028384889 | 0.189729026 | -0.110833427 | 0.22185038 | 11.19235 | -0.113425103 | 0.315228558 | 0.104914222 | 0.106915886 | 11.19235 | 0.050469379 | 0.152953873 | -0.018681491 | 0.190212082 | 11.19235 | -0.104694895 | 0.235864078 | -0.040632431 | 0.183098832 |
| 11.20259 | -0.029634521 | 0.188924043 | -0.109611107 | 0.22923579 | 11.20259 | -0.111914353 | 0.313996136 | 0.098955197 | 0.104623639 | 11.20259 | 0.051805799 | 0.15731125 | -0.020525493 | 0.194085848 | 11.20259 | -0.108941973 | 0.237367728 | -0.047244216 | 0.180335221 |
| 11.21283 | -0.031144149 | 0.188559365 | -0.107284486 | 0.23760342 | 11.21283 | -0.109637256 | 0.310894732 | 0.093044071 | 0.102997612 | 11.21283 | 0.053172636 | 0.162093254 | -0.02264616 | 0.198353831 | 11.21283 | -0.112600023 | 0.237354926 | -0.054376355 | 0.177647547 |
| 11.22307 | -0.032761021 | 0.188534763 | -0.104933393 | 0.24459316 | 11.22307 | -0.106338811 | 0.305689206 | 0.088253609 | 0.101765782 | 11.22307 | 0.05550256 | 0.165454545 | -0.024550846 | 0.202407289 | 11.22307 | -0.115353819 | 0.235588259 | -0.06107667 | 0.17541989 |
| 11.23331 | -0.034733395 | 0.188884641 | -0.102335372 | 0.25122311 | 11.23331 | -0.102190246 | 0.298844762 | 0.08389749 | 0.101006921 | 11.23331 | 0.058076925 | 0.16882309 | -0.0263743 | 0.205802324 | 11.23331 | -0.117485311 | 0.232398789 | -0.067318132 | 0.173069881 |
| 11.24355 | -0.037262244 | 0.189327659 | -0.10017917 | 0.25662564 | 11.24355 | -0.097027722 | 0.291784709 | 0.081042022 | 0.099797116 | 11.24355 | 0.061428696 | 0.170588255 | -0.027794586 | 0.208377113 | 11.24355 | -0.119197279 | 0.22788459 | -0.072408081 | 0.170578255 |
| 11.25379 | -0.039903381 | 0.19024681 | -0.09869891 | 0.25986242 | 11.25379 | -0.090782332 | 0.284033411 | 0.078986571 | 0.099449257 | 11.25379 | 0.064844224 | 0.171452839 | -0.029308544 | 0.20909602 | 11.25379 | -0.120540826 | 0.222557838 | -0.076856378 | 0.168246663 |
| 11.26403 | -0.043175335 | 0.191439857 | -0.097812146 | 0.26222001 | 11.26403 | -0.083458942 | 0.277121907 | 0.07853426 | 0.098834595 | 11.26403 | 0.068715928 | 0.171017254 | -0.029873958 | 0.208296784 | 11.26403 | -0.122245689 | 0.215928073 | -0.08026494 | 0.165160237 |
| 11.27427 | -0.045996522 | 0.19354175 | -0.097666027 | 0.26322296 | 11.27427 | -0.075620556 | 0.271230273 | 0.078830318 | 0.099235782 | 11.27427 | 0.072683982 | 0.169932743 | -0.030659161 | 0.205902391 | 11.27427 | -0.123825493 | 0.209550196 | -0.082767737 | 0.161564989 |
| 11.28451 | -0.048648103 | 0.196483188 | -0.09734756 | 0.26328235 | 11.28451 | -0.067319074 | 0.266153472 | 0.080429438 | 0.100190722 | 11.28451 | 0.077284823 | 0.168736184 | -0.030586744 | 0.20237784 | 11.28451 | -0.125911127 | 0.202764977 | -0.085273184 | 0.157094174 |
| 11.29475 | -0.050326553 | 0.199787467 | -0.097415745 | 0.26207241 | 11.29475 | -0.059266075 | 0.262673595 | 0.08266187 | 0.101790374 | 11.29475 | 0.08148956 | 0.167343308 | -0.030820474 | 0.197910829 | 11.29475 | -0.128634943 | 0.195852363 | -0.087041441 | 0.151988815 |
| 11.30499 | -0.051532052 | 0.203590406 | -0.09654873 | 0.26065055 | 11.30499 | -0.051217485 | 0.259958016 | 0.085459769 | 0.104868759 | 11.30499 | 0.086405348 | 0.166693749 | -0.030368984 | 0.193276826 | 11.30499 | -0.131165545 | 0.189307301 | -0.089487682 | 0.146732558 |
| 11.31523 | -0.050958807 | 0.207207634 | -0.094432663 | 0.25841768 | 11.31523 | -0.044623101 | 0.259347362 | 0.087858963 | 0.108811865 | 11.31523 | 0.091182282 | 0.166341852 | -0.029935259 | 0.188913795 | 11.31523 | -0.134450663 | 0.182891661 | -0.091633385 | 0.140973876 |
| 11.32547 | -0.049074066 | 0.210757488 | -0.091319915 | 0.25529653 | 11.32547 | -0.038780475 | 0.258716246 | 0.089764491 | 0.113056747 | 11.32547 | 0.09633626 | 0.166737818 | -0.029217093 | 0.184905495 | 11.32547 | -0.137427122 | 0.176828519 | -0.094221651 | 0.135930226 |
| 11.33571 | -0.046006819 | 0.214046618 | -0.086612848 | 0.25200527 | 11.33571 | -0.035078999 | 0.259217756 | 0.090466629 | 0.118441869 | 11.33571 | 0.101481454 | 0.167150182 | -0.028421658 | 0.181378818 | 11.33571 | -0.140688085 | 0.170564629 | -0.096917718 | 0.131274346 |
| 11.34595 | -0.041149604 | 0.217206348 | -0.08097848 | 0.24814864 | 11.34595 | -0.032019752 | 0.260474606 | 0.089886679 | 0.123808541 | 11.34595 | 0.106230254 | 0.166926362 | -0.027179135 | 0.178129413 | 11.34595 | -0.143577349 | 0.165010984 | -0.100008115 | 0.127348256 |
| 11.35619 | -0.035798902 | 0.220940542 | -0.07465586 | 0.24435907 | 11.35619 | -0.030996632 | 0.261810522 | 0.087729596 | 0.129390045 | 11.35619 | 0.110944082 | 0.166596046 | -0.025807444 | 0.175341136 | 11.35619 | -0.146757853 | 0.159066 | -0.102912487 | 0.12455397 |
| 11.36643 | -0.029362785 | 0.224932029 | -0.067933883 | 0.24082673 | 11.36643 | -0.030569687 | 0.263640958 | 0.084525274 | 0.134953057 | 11.36643 | 0.115208854 | 0.164891737 | -0.024036668 | 0.17190461 | 11.36643 | -0.149234774 | 0.153873052 | -0.106112508 | 0.122488886 |
| 11.37667 | -0.023089721 | 0.229364473 | -0.06208289 | 0.2377229 | 11.37667 | -0.031584341 | 0.26485946 | 0.080485733 | 0.140036695 | 11.37667 | 0.119083719 | 0.162243939 | -0.021793362 | 0.168465966 | 11.37667 | -0.152260699 | 0.148360831 | -0.108877728 | 0.121431449 |
| 11.38691 | -0.017396792 | 0.23406031 | -0.057084032 | 0.235389 | 11.38691 | -0.032447185 | 0.266569778 | 0.076292985 | 0.145155564 | 11.38691 | 0.122268305 | 0.15856788 | -0.019242855 | 0.164384018 | 11.38691 | -0.15504831 | 0.144517456 | -0.111730489 | 0.1216155 |
| 11.39715 | -0.012819282 | 0.239099086 | -0.053019421 | 0.23379657 | 11.39715 | -0.034049811 | 0.26754012 | 0.072196346 | 0.14993508 | 11.39715 | 0.125140245 | 0.154083537 | -0.016043589 | 0.160341613 | 11.39715 | -0.158112561 | 0.140395348 | -0.113898434 | 0.122871434 |
| 11.40739 | -0.009410881 | 0.243801613 | -0.050626984 | 0.23241926 | 11.40739 | -0.035653521 | 0.268094043 | 0.068764434 | 0.154386039 | 11.40739 | 0.127679424 | 0.148934263 | -0.012381967 | 0.156406698 | 11.40739 | -0.161595003 | 0.137580862 | -0.115853281 | 0.125341348 |
| 11.41763 | -0.007997632 | 0.248458719 | -0.049407467 | 0.23206649 | 11.41763 | -0.036722266 | 0.268464427 | 0.066362556 | 0.159217756 | 11.41763 | 0.130156011 | 0.143733431 | -0.008284316 | 0.152678355 | 11.41763 | -0.165051471 | 0.135459993 | -0.116542572 | 0.128645671 |
| 11.42787 | -0.007611639 | 0.252974462 | -0.04939625 | 0.23170123 | 11.42787 | -0.038351892 | 0.26791297 | 0.064680341 | 0.16359621 | 11.42787 | 0.132436616 | 0.13872607 | -0.004128507 | 0.150567859 | 11.42787 | -0.169445424 | 0.1337355 | -0.116864868 | 0.132655449 |
| 11.43811 | -0.008595501 | 0.256817828 | -0.050160296 | 0.23156118 | 11.43811 | -0.038726915 | 0.266780944 | 0.063766375 | 0.167761965 | 11.43811 | 0.13485267 | 0.134095186 | 0.000480107 | 0.148971308 | 11.43811 | -0.173897809 | 0.132736448 | -0.116311024 | 0.136901022 |
| 11.44835 | -0.010081293 | 0.260731121 | -0.051425438 | 0.23123445 | 11.44835 | -0.040221017 | 0.265255777 | 0.063283249 | 0.171725911 | 11.44835 | 0.137076289 | 0.130100152 | 0.004583392 | 0.149102017 | 11.44835 | -0.178999222 | 0.1314373 | -0.114844187 | 0.1410948 |
| 11.45859 | -0.011014885 | 0.263695029 | -0.052984916 | 0.23156231 | 11.45859 | -0.04072088 | 0.263388202 | 0.06333809 | 0.175225301 | 11.45859 | 0.139144503 | 0.12703312 | 0.008933331 | 0.150481663 | 11.45859 | -0.184497424 | 0.130508796 | -0.113528483 | 0.144797521 |
| 11.46883 | -0.011951089 | 0.266645379 | -0.054416384 | 0.23137085 | 11.46883 | -0.042342934 | 0.260699924 | 0.062956405 | 0.177859183 | 11.46883 | 0.14077228 | 0.124106711 | 0.011258058 | 0.153473244 | 11.46883 | -0.190224601 | 0.129419601 | -0.111674683 | 0.148224528 |
| 11.47907 | -0.010785426 | 0.267702896 | -0.055458625 | 0.23183362 | 11.47907 | -0.043640999 | 0.258105485 | 0.062155398 | 0.179551359 | 11.47907 | 0.141831327 | 0.122389536 | 0.013567595 | 0.157865146 | 11.47907 | -0.196470034 | 0.128339054 | -0.110464899 | 0.150754915 |
| 11.48931 | -0.008961696 | 0.269093383 | -0.05615304 | 0.23290722 | 11.48931 | -0.046157512 | 0.254937911 | 0.060744451 | 0.180511118 | 11.48931 | 0.141753837 | 0.121255727 | 0.014234205 | 0.163730142 | 11.48931 | -0.202777121 | 0.127359224 | -0.10967545 | 0.153194646 |
| 11.49955 | -0.004847055 | 0.268049165 | -0.056337136 | 0.23448323 | 11.49955 | -0.048506813 | 0.251623158 | 0.057759173 | 0.180326647 | 11.49955 | 0.141244179 | 0.12063894 | 0.013970742 | 0.170888916 | 11.49955 | -0.208909889 | 0.126959303 | -0.109908693 | 0.155192459 |
| 11.50979 | 2.46534E-05 | 0.26673105 | -0.056522012 | 0.23699797 | 11.50979 | -0.0513696 | 0.248173749 | 0.053881509 | 0.179695676 | 11.50979 | 0.139626924 | 0.120733228 | 0.012639556 | 0.178843298 | 11.50979 | -0.215437062 | 0.127413159 | -0.110178192 | 0.157126126 |
| 11.52003 | 0.006293256 | 0.263800471 | -0.056209335 | 0.2402419 | 11.52003 | -0.053830215 | 0.244366998 | 0.048343644 | 0.178494899 | 11.52003 | 0.137353248 | 0.120702965 | 0.010723063 | 0.187388584 | 11.52003 | -0.221094142 | 0.128682178 | -0.111942581 | 0.159151109 |
| 11.53027 | 0.012439761 | 0.259945231 | -0.056223181 | 0.24442451 | 11.53027 | -0.055695499 | 0.241093732 | 0.041271533 | 0.178169575 | 11.53027 | 0.134560967 | 0.120330036 | 0.008068445 | 0.19598723 | 11.53027 | -0.226426185 | 0.131353284 | -0.113321229 | 0.161678756 |
| 11.54051 | 0.019009562 | 0.256130592 | -0.056572117 | 0.24875336 | 11.54051 | -0.056852 | 0.237448009 | 0.0327048 | 0.1775278 | 11.54051 | 0.131451277 | 0.119150712 | 0.005814701 | 0.204261832 | 11.54051 | -0.231184517 | 0.135012228 | -0.115394522 | 0.164947443 |
| 11.55075 | 0.024553531 | 0.251528938 | -0.057560877 | 0.25307165 | 11.55075 | -0.056485539 | 0.234374332 | 0.023306508 | 0.179275964 | 11.55075 | 0.12846004 | 0.116181357 | 0.003496892 | 0.212186522 | 11.55075 | -0.235035408 | 0.139433138 | -0.116888273 | 0.168939965 |
| 11.56099 | 0.029110222 | 0.247539148 | -0.059388208 | 0.25730866 | 11.56099 | -0.054766038 | 0.231368616 | 0.012840443 | 0.181491023 | 11.56099 | 0.125077043 | 0.11267776 | 0.001943151 | 0.219473921 | 11.56099 | -0.238100365 | 0.144070054 | -0.11862419 | 0.173842051 |
| 11.57123 | 0.03229141 | 0.243454762 | -0.061686863 | 0.26052363 | 11.57123 | -0.052005614 | 0.228841118 | 0.002357222 | 0.185831306 | 11.57123 | 0.121843383 | 0.107177273 | 0.000417631 | 0.226059599 | 11.57123 | -0.240226465 | 0.149276455 | -0.120096378 | 0.178992789 |
| 11.58147 | 0.033794843 | 0.240915003 | -0.065748096 | 0.26269005 | 11.58147 | -0.048557527 | 0.225313931 | -0.008704821 | 0.190638457 | 11.58147 | 0.119311931 | 0.102071098 | 2.78847E-05 | 0.23178551 | 11.58147 | -0.241394281 | 0.153630551 | -0.120879034 | 0.183981796 |
| 11.59171 | 0.03394374 | 0.238928208 | -0.070192745 | 0.26319792 | 11.59171 | -0.044389403 | 0.222966653 | -0.018854859 | 0.196927842 | 11.59171 | 0.11649883 | 0.096148588 | -0.000683778 | 0.236980184 | 11.59171 | -0.241763391 | 0.15824147 | -0.122377424 | 0.187895695 |
| 11.60195 | 0.032450621 | 0.237590238 | -0.075231791 | 0.26255854 | 11.60195 | -0.040263332 | 0.219934418 | -0.028604757 | 0.202927867 | 11.60195 | 0.114361428 | 0.091165534 | -0.001214761 | 0.24083077 | 11.60195 | -0.241000333 | 0.16157402 | -0.123796094 | 0.190184493 |
| 11.61219 | 0.029546715 | 0.23758953 | -0.080500383 | 0.26053959 | 11.61219 | -0.03607745 | 0.216582921 | -0.03735984 | 0.208961665 | 11.61219 | 0.112950763 | 0.086702285 | -0.001788454 | 0.244064759 | 11.61219 | -0.239217176 | 0.165547303 | -0.12610379 | 0.190687703 |
| 11.62243 | 0.026074225 | 0.237948622 | -0.08539703 | 0.25712093 | 11.62243 | -0.032997105 | 0.212299321 | -0.045046762 | 0.214041998 | 11.62243 | 0.111629145 | 0.082938808 | -0.002778168 | 0.245753952 | 11.62243 | -0.236033451 | 0.168911638 | -0.128824726 | 0.189559044 |
| 11.63267 | 0.020975096 | 0.238721089 | -0.089929927 | 0.25351037 | 11.63267 | -0.02998899 | 0.208635409 | -0.051634418 | 0.21872461 | 11.63267 | 0.111417276 | 0.079528767 | -0.004459939 | 0.246683378 | 11.63267 | -0.231819452 | 0.172478393 | -0.132667782 | 0.186751974 |
| 11.64291 | 0.016011706 | 0.238763909 | -0.092854887 | 0.24919775 | 11.64291 | -0.02773544 | 0.203247995 | -0.057141896 | 0.222028954 | 11.64291 | 0.111249317 | 0.076740681 | -0.006339558 | 0.246317902 | 11.64291 | -0.225636584 | 0.176406477 | -0.136792224 | 0.183023209 |
| 11.65315 | 0.009954764 | 0.238718826 | -0.094729385 | 0.24526377 | 11.65315 | -0.026512287 | 0.197877775 | -0.061880912 | 0.225313071 | 11.65315 | 0.111963713 | 0.074107027 | -0.008443895 | 0.245463933 | 11.65315 | -0.219119554 | 0.180399155 | -0.141463886 | 0.179935298 |
| 11.66339 | 0.004040085 | 0.236570703 | -0.095279641 | 0.24133341 | 11.66339 | -0.025218626 | 0.192450578 | -0.065591947 | 0.228015365 | 11.66339 | 0.112861487 | 0.071849728 | -0.010928801 | 0.24350786 | 11.66339 | -0.210568476 | 0.183776096 | -0.146122147 | 0.17683919 |
| 11.67363 | -0.002117834 | 0.233401056 | -0.094578312 | 0.23783022 | 11.67363 | -0.025010784 | 0.187646041 | -0.069193635 | 0.231466298 | 11.67363 | 0.11403767 | 0.070854187 | -0.013108439 | 0.241056617 | 11.67363 | -0.202004462 | 0.18682079 | -0.150516149 | 0.174954956 |
| 11.68387 | -0.00834896 | 0.227548831 | -0.093103934 | 0.23509594 | 11.68387 | -0.024870758 | 0.183577374 | -0.072470581 | 0.2348666 | 11.68387 | 0.115671531 | 0.070677347 | -0.015252703 | 0.239026547 | 11.68387 | -0.192874174 | 0.188433411 | -0.154178603 | 0.173868773 |
| 11.69411 | -0.013813741 | 0.220653844 | -0.091463016 | 0.23218174 | 11.69411 | -0.025249374 | 0.181748406 | -0.075447783 | 0.239578034 | 11.69411 | 0.117425412 | 0.071975409 | -0.016919712 | 0.236179943 | 11.69411 | -0.18439398 | 0.188819226 | -0.157051051 | 0.173922382 |
| 11.70435 | -0.019290249 | 0.211945712 | -0.08973708 | 0.23049397 | 11.70435 | -0.026295836 | 0.181513229 | -0.078955532 | 0.244382684 | 11.70435 | 0.12005914 | 0.074926186 | -0.018233006 | 0.234090962 | 11.70435 | -0.176254845 | 0.187879832 | -0.159030487 | 0.173671387 |
| 11.71459 | -0.023944496 | 0.203458127 | -0.088846432 | 0.22882844 | 11.71459 | -0.027334145 | 0.184193676 | -0.081844792 | 0.249580244 | 11.71459 | 0.122741232 | 0.079060479 | -0.019239414 | 0.232242143 | 11.71459 | -0.16962994 | 0.185754252 | -0.160235209 | 0.173644628 |
| 11.72483 | -0.028290887 | 0.195081345 | -0.088863443 | 0.22775221 | 11.72483 | -0.028882125 | 0.189201252 | -0.08470386 | 0.254198456 | 11.72483 | 0.126818764 | 0.084470258 | -0.019665976 | 0.230701917 | 11.72483 | -0.163689852 | 0.182988352 | -0.160389507 | 0.173300555 |
| 11.73507 | -0.031605504 | 0.188334142 | -0.089693828 | 0.22711161 | 11.73507 | -0.030796528 | 0.196074674 | -0.087470662 | 0.258554467 | 11.73507 | 0.131649964 | 0.090632584 | -0.019575794 | 0.2295865 | 11.73507 | -0.159590764 | 0.179883919 | -0.159875253 | 0.172391605 |
| 11.74531 | -0.034880307 | 0.182559719 | -0.091252526 | 0.22740218 | 11.74531 | -0.032564235 | 0.205153611 | -0.089531803 | 0.262511183 | 11.74531 | 0.137638908 | 0.09771039 | -0.019623882 | 0.228724808 | 11.74531 | -0.155980274 | 0.17717744 | -0.158422887 | 0.171031429 |
| 11.75555 | -0.037757743 | 0.17855832 | -0.093980132 | 0.22739959 | 11.75555 | -0.034351746 | 0.214989778 | -0.09082836 | 0.265462703 | 11.75555 | 0.144836239 | 0.105713474 | -0.019076821 | 0.22779889 | 11.75555 | -0.153734071 | 0.175056125 | -0.156550147 | 0.168818021 |
| 11.76579 | -0.039862814 | 0.175356918 | -0.096291621 | 0.22866228 | 11.76579 | -0.03641105 | 0.225406612 | -0.091711646 | 0.268441709 | 11.76579 | 0.152893011 | 0.11382367 | -0.01883238 | 0.226608837 | 11.76579 | -0.151812316 | 0.174248839 | -0.153605845 | 0.166798376 |
| 11.77603 | -0.042184347 | 0.172543706 | -0.09888014 | 0.22998058 | 11.77603 | -0.038012365 | 0.235304446 | -0.092134411 | 0.271069759 | 11.77603 | 0.161446706 | 0.122405768 | -0.018674504 | 0.225598884 | 11.77603 | -0.150486859 | 0.174700249 | -0.150723013 | 0.164032853 |
| 11.78627 | -0.044092073 | 0.169714747 | -0.100688214 | 0.23283835 | 11.78627 | -0.039823102 | 0.244320813 | -0.092102984 | 0.273523874 | 11.78627 | 0.170263571 | 0.13040677 | -0.018843242 | 0.224254621 | 11.78627 | -0.149296527 | 0.176020656 | -0.147504883 | 0.161328764 |
| 11.79651 | -0.045542189 | 0.166412757 | -0.101940658 | 0.23650723 | 11.79651 | -0.041339897 | 0.251787653 | -0.091928693 | 0.275437034 | 11.79651 | 0.178675565 | 0.137730501 | -0.019778069 | 0.223274755 | 11.79651 | -0.148203578 | 0.17915588 | -0.144538698 | 0.158258008 |
| 11.80675 | -0.046413791 | 0.162794253 | -0.101677024 | 0.24145372 | 11.80675 | -0.043155042 | 0.257846868 | -0.091814452 | 0.277386875 | 11.80675 | 0.185903516 | 0.143270685 | -0.021098013 | 0.222669259 | 11.80675 | -0.147011133 | 0.182177044 | -0.142048593 | 0.155369294 |
| 11.81699 | -0.047268206 | 0.15914629 | -0.100982425 | 0.24762089 | 11.81699 | -0.044709122 | 0.262335831 | -0.092041539 | 0.278112925 | 11.81699 | 0.191657125 | 0.147013858 | -0.022904447 | 0.222919529 | 11.81699 | -0.145321197 | 0.186321436 | -0.140479075 | 0.152148695 |
| 11.82723 | -0.04731387 | 0.155741557 | -0.098810316 | 0.2551894 | 11.82723 | -0.04678981 | 0.265479018 | -0.092313335 | 0.277809379 | 11.82723 | 0.196057722 | 0.148811418 | -0.025472153 | 0.224403333 | 11.82723 | -0.143407514 | 0.190074103 | -0.139481095 | 0.148840693 |
| 11.83747 | -0.046376123 | 0.153336607 | -0.096121926 | 0.26369822 | 11.83747 | -0.04867899 | 0.267052148 | -0.092978535 | 0.27638824 | 11.83747 | 0.198410521 | 0.148582207 | -0.027655999 | 0.227155889 | 11.83747 | -0.141232347 | 0.193603526 | -0.139439482 | 0.145990916 |
| 11.84771 | -0.045242545 | 0.152864187 | -0.09282899 | 0.27271016 | 11.84771 | -0.050627794 | 0.26866631 | -0.093742044 | 0.274480006 | 11.84771 | 0.1988822 | 0.146618012 | -0.030574904 | 0.231149117 | 11.84771 | -0.138787322 | 0.196650074 | -0.13955515 | 0.142945951 |
| 11.85795 | -0.043405022 | 0.154190404 | -0.088935114 | 0.28153327 | 11.85795 | -0.052978694 | 0.268297818 | -0.094323907 | 0.270809387 | 11.85795 | 0.198171983 | 0.143191904 | -0.032876762 | 0.236559046 | 11.85795 | -0.136171236 | 0.198403599 | -0.140400906 | 0.140703186 |
| 11.86819 | -0.040599999 | 0.157982867 | -0.085578436 | 0.29038291 | 11.86819 | -0.054631459 | 0.268081019 | -0.094896859 | 0.267394439 | 11.86819 | 0.195749082 | 0.138994748 | -0.034824769 | 0.242602058 | 11.86819 | -0.134208475 | 0.199587511 | -0.140681491 | 0.139161643 |
| 11.87843 | -0.037643747 | 0.163309712 | -0.081436132 | 0.29762449 | 11.87843 | -0.056570772 | 0.266679664 | -0.094599842 | 0.263057934 | 11.87843 | 0.192191716 | 0.133773864 | -0.036386404 | 0.249207754 | 11.87843 | -0.132500078 | 0.199053032 | -0.140898936 | 0.138675669 |
| 11.88867 | -0.033830983 | 0.170901382 | -0.078417316 | 0.30402672 | 11.88867 | -0.057377209 | 0.264291584 | -0.093631623 | 0.259106961 | 11.88867 | 0.188118953 | 0.128349448 | -0.037394064 | 0.255375682 | 11.88867 | -0.132104288 | 0.19732222 | -0.140739038 | 0.139042299 |
| 11.89891 | -0.030470852 | 0.178494143 | -0.076097782 | 0.30880434 | 11.89891 | -0.057886102 | 0.260782395 | -0.092328156 | 0.254658685 | 11.89891 | 0.183371073 | 0.122172648 | -0.037497333 | 0.261775114 | 11.89891 | -0.132163127 | 0.19417894 | -0.140182322 | 0.14056603 |
| 11.90915 | -0.026676396 | 0.18639992 | -0.074272569 | 0.31247642 | 11.90915 | -0.056992748 | 0.256320602 | -0.089133799 | 0.250393372 | 11.90915 | 0.178426833 | 0.115688238 | -0.037200132 | 0.26721861 | 11.90915 | -0.13375548 | 0.190033764 | -0.139150127 | 0.142378984 |
| 11.91939 | -0.023454669 | 0.194526649 | -0.074056648 | 0.31557545 | 11.91939 | -0.055203078 | 0.250038386 | -0.085815231 | 0.245535698 | 11.91939 | 0.173976494 | 0.10902592 | -0.036244352 | 0.271676824 | 11.91939 | -0.135473868 | 0.184562471 | -0.138131188 | 0.144720729 |
| 11.92963 | -0.021525179 | 0.201710206 | -0.074870778 | 0.31782542 | 11.92963 | -0.052089794 | 0.242941651 | -0.08120007 | 0.240555532 | 11.92963 | 0.169596507 | 0.101629613 | -0.034837801 | 0.275518959 | 11.92963 | -0.138100311 | 0.17890327 | -0.137141959 | 0.146426354 |
| 11.93987 | -0.020126895 | 0.208268907 | -0.077066152 | 0.31975294 | 11.93987 | -0.04784642 | 0.23522224 | -0.075853961 | 0.235186276 | 11.93987 | 0.166387051 | 0.094848855 | -0.0329482 | 0.278371433 | 11.93987 | -0.140737397 | 0.173093281 | -0.135984844 | 0.148252084 |
| 11.95011 | -0.020283063 | 0.213601191 | -0.079737839 | 0.32163426 | 11.95011 | -0.042171912 | 0.226689154 | -0.069830907 | 0.229881071 | 11.95011 | 0.163935613 | 0.087575261 | -0.031061307 | 0.280715214 | 11.95011 | -0.142600076 | 0.1673458 | -0.135221668 | 0.148871502 |
| 11.96035 | -0.021903145 | 0.218393608 | -0.082768044 | 0.32299987 | 11.96035 | -0.036036571 | 0.21882192 | -0.06322555 | 0.224715344 | 11.96035 | 0.1622068 | 0.081642227 | -0.029194237 | 0.281912485 | 11.96035 | -0.144344403 | 0.162556007 | -0.134471773 | 0.149268085 |
| 11.97059 | -0.024573382 | 0.221804992 | -0.085360242 | 0.32454837 | 11.97059 | -0.029309408 | 0.211014753 | -0.05620326 | 0.220177573 | 11.97059 | 0.160981937 | 0.0762656 | -0.027160862 | 0.283153904 | 11.97059 | -0.144514984 | 0.158683422 | -0.133690623 | 0.149214404 |

| | | | | |
|---|---|---|---|---|
| 11.98083 | -0.028140382 | 0.223579562 | -0.086987947 | 0.32525099 |
| 11.99107 | -0.03284863 | 0.223878646 | -0.087080758 | 0.32592407 |
| 12.00131 | -0.037519143 | 0.2227086 | -0.08585099 | 0.32626785 |
| 12.01155 | -0.04301513 | 0.219041296 | -0.083291738 | 0.32610972 |
| 12.02179 | -0.047764188 | 0.21397171 | -0.079896973 | 0.32577538 |
| 12.03203 | -0.053115333 | 0.207641216 | -0.075950803 | 0.32529851 |
| 12.04227 | -0.057438138 | 0.199617805 | -0.072661819 | 0.32514097 |
| 12.05252 | -0.061762071 | 0.191169835 | -0.069516865 | 0.3255674 |
| 12.06275 | -0.065258754 | 0.181949366 | -0.068063785 | 0.32681299 |
| 12.073 | -0.068180189 | 0.173044467 | -0.06729671 | 0.32944821 |
| 12.08323 | -0.070779006 | 0.163703711 | -0.067718581 | 0.33340128 |
| 12.09348 | -0.07268457 | 0.155781358 | -0.069065819 | 0.33931651 |
| 12.10371 | -0.074899554 | 0.148506843 | -0.070632239 | 0.34649404 |
| 12.11395 | -0.076396377 | 0.143556263 | -0.073139139 | 0.35443166 |
| 12.1242 | -0.078819118 | 0.140508237 | -0.075081168 | 0.36343506 |
| 12.13443 | -0.081464991 | 0.140190311 | -0.077845707 | 0.37184927 |
| 12.14468 | -0.084823194 | 0.142096337 | -0.080225545 | 0.38016761 |
| 12.15491 | -0.08871957 | 0.145890621 | -0.083248424 | 0.38749092 |
| 12.16516 | -0.093090268 | 0.15091141 | -0.08645805 | 0.39421094 |
| 12.1754 | -0.098427041 | 0.15738627 | -0.090379092 | 0.40017264 |
| 12.18564 | -0.103435817 | 0.163447244 | -0.094682236 | 0.40547178 |
| 12.19588 | -0.108751116 | 0.170098667 | -0.099410719 | 0.41047047 |
| 12.20611 | -0.112966373 | 0.175708061 | -0.104509506 | 0.41524657 |
| 12.21636 | -0.116735624 | 0.180805413 | -0.109124506 | 0.41967996 |
| 12.22659 | -0.118778723 | 0.184386158 | -0.113924137 | 0.42443098 |
| 12.23684 | -0.118480268 | 0.186631226 | -0.117127188 | 0.4281908 |
| 12.24708 | -0.116494779 | 0.187180733 | -0.119863671 | 0.43224623 |
| 12.25732 | -0.111710839 | 0.186146029 | -0.12088 | 0.43555301 |
| 12.26756 | -0.105041045 | 0.183396717 | -0.120559724 | 0.43906209 |
| 12.2778 | -0.09668039 | 0.179146087 | -0.119013794 | 0.44208508 |
| 12.28804 | -0.087393361 | 0.174272432 | -0.11627753 | 0.44452632 |
| 12.29828 | -0.078121526 | 0.168880076 | -0.112616469 | 0.44795168 |
| 12.30852 | -0.069795373 | 0.163555515 | -0.108867653 | 0.44992325 |
| 12.31876 | -0.062880038 | 0.159792628 | -0.104580062 | 0.45236541 |
| 12.329 | -0.05889683 | 0.157476137 | -0.101161329 | 0.45348854 |
| 12.33924 | -0.056903339 | 0.157462748 | -0.097797988 | 0.45456326 |
| 12.34948 | -0.058123988 | 0.159795496 | -0.095336008 | 0.45367265 |
| 12.35972 | -0.061138035 | 0.164764931 | -0.093468401 | 0.45151629 |
| 12.36996 | -0.066662931 | 0.172111955 | -0.091865823 | 0.44785012 |
| 12.3802 | -0.073228223 | 0.181172054 | -0.091111159 | 0.44251024 |
| 12.39044 | -0.080189244 | 0.191351389 | -0.089716422 | 0.43519865 |
| 12.40068 | -0.087515717 | 0.202418318 | -0.089047925 | 0.42735694 |
| 12.41092 | -0.093794584 | 0.212939334 | -0.087645111 | 0.41812308 |
| 12.42116 | -0.099629668 | 0.223167864 | -0.086111144 | 0.4091383 |
| 12.4314 | -0.103429181 | 0.23183447 | -0.084236856 | 0.39988831 |
| 12.44164 | -0.106708733 | 0.239529342 | -0.082056172 | 0.39176527 |
| 12.45188 | -0.10790955 | 0.245694118 | -0.079868203 | 0.38375259 |
| 12.46212 | -0.108183985 | 0.251012262 | -0.0776582 | 0.37745467 |
| 12.47236 | -0.106608084 | 0.255451687 | -0.075737617 | 0.37193634 |
| 12.4826 | -0.104470267 | 0.258925845 | -0.074781885 | 0.36782152 |
| 12.49284 | -0.100774862 | 0.26214137 | -0.074392721 | 0.36454415 |
| 12.50308 | -0.096288705 | 0.264251423 | -0.075242604 | 0.36175847 |
| 12.51332 | -0.091206845 | 0.265815941 | -0.077225547 | 0.35978922 |
| 12.52356 | -0.085972651 | 0.266029096 | -0.080347555 | 0.35724887 |
| 12.5338 | -0.080977772 | 0.265756166 | -0.083820671 | 0.35428484 |
| 12.54404 | -0.076725021 | 0.264473315 | -0.088452981 | 0.3501022 |
| 12.55428 | -0.073642006 | 0.262412948 | -0.092679328 | 0.3454121 |
| 12.56452 | -0.07194994 | 0.26024434 | -0.09655976 | 0.33950571 |
| 12.57476 | -0.071731611 | 0.257373408 | -0.099878037 | 0.33359386 |
| 12.585 | -0.072965205 | 0.254255874 | -0.102028673 | 0.32751262 |
| 12.59524 | -0.074951571 | 0.250184495 | -0.103390923 | 0.32252115 |
| 12.60548 | -0.077628599 | 0.24553795 | -0.103104869 | 0.31848528 |
| 12.61572 | -0.07962089 | 0.239234817 | -0.102554138 | 0.3162127 |
| 12.62596 | -0.081560473 | 0.232401434 | -0.10059642 | 0.31431067 |
| 12.6362 | -0.081248613 | 0.223722817 | -0.099036656 | 0.31409925 |
| 12.64644 | -0.080689567 | 0.214416821 | -0.09700215 | 0.31300394 |
| 12.65668 | -0.077103763 | 0.204024283 | -0.094748676 | 0.31190406 |
| 12.66692 | -0.07304692 | 0.193829077 | -0.093089978 | 0.30924677 |
| 12.67716 | -0.067460114 | 0.183803896 | -0.091193713 | 0.30526677 |
| 12.6874 | -0.061512393 | 0.174627109 | -0.089546199 | 0.2996731 |
| 12.69764 | -0.055296088 | 0.166557229 | -0.087832799 | 0.2929131 |
| 12.70788 | -0.049972399 | 0.160687959 | -0.086193671 | 0.28494205 |
| 12.71812 | -0.045693093 | 0.15584209 | -0.084601549 | 0.27688888 |
| 12.72836 | -0.042544881 | 0.153564884 | -0.083836919 | 0.26870723 |
| 12.7386 | -0.040672526 | 0.151821028 | -0.083179701 | 0.26110221 |
| 12.74884 | -0.039792469 | 0.152072239 | -0.083547877 | 0.25442478 |
| 12.75908 | -0.039706274 | 0.152784688 | -0.084463977 | 0.24897881 |
| 12.76932 | -0.039000692 | 0.15483294 | -0.086386777 | 0.24434212 |
| 12.77956 | -0.038798999 | 0.157306902 | -0.088646425 | 0.24126564 |
| 12.7898 | -0.036770599 | 0.160366671 | -0.091246005 | 0.23866418 |
| 12.80004 | -0.034298329 | 0.163717744 | -0.094306544 | 0.23719402 |
| 12.81028 | -0.030305287 | 0.167082211 | -0.09646098 | 0.23618007 |
| 12.82052 | -0.025675445 | 0.170258923 | -0.09903245 | 0.23527124 |
| 12.83076 | -0.01981989 | 0.172651112 | -0.100816426 | 0.23475347 |
| 12.841 | -0.013427449 | 0.175014574 | -0.102329456 | 0.23369225 |
| 12.85124 | -0.006473726 | 0.175362972 | -0.102857011 | 0.23265803 |
| 12.86148 | 0.001032977 | 0.176636395 | -0.103336678 | 0.23149144 |
| 12.87172 | 0.009189137 | 0.175736256 | -0.10305675 | 0.22966702 |
| 12.88196 | 0.017609753 | 0.175895908 | -0.10253956 | 0.22799618 |
| 12.8922 | 0.026603491 | 0.174904156 | -0.101703493 | 0.22646068 |
| 12.90244 | 0.03511672 | 0.173981949 | -0.101247536 | 0.22474239 |
| 12.91268 | 0.044108353 | 0.172086758 | -0.100597988 | 0.22322194 |
| 12.92292 | 0.051914149 | 0.169660066 | -0.100719605 | 0.22173007 |
| 12.93316 | 0.059722943 | 0.166091083 | -0.100739686 | 0.2205734 |
| 12.9434 | 0.066354542 | 0.162416407 | -0.101133782 | 0.21886689 |
| 12.95364 | 0.072704895 | 0.158223349 | -0.101280171 | 0.21719187 |
| 12.96388 | 0.078695352 | 0.154658402 | -0.101215435 | 0.21507149 |
| 12.97412 | 0.084451807 | 0.151666765 | -0.100543958 | 0.21202021 |
| 12.98436 | 0.089994111 | 0.149012042 | -0.099028172 | 0.20806007 |
| 12.9946 | 0.095742182 | 0.147304686 | -0.096548028 | 0.20395898 |
| 13.00484 | 0.101308286 | 0.145810015 | -0.093482411 | 0.19774572 |
| 13.01508 | 0.107078636 | 0.144971143 | -0.08906943 | 0.19158228 |
| 13.02532 | 0.112490506 | 0.144575393 | -0.084017084 | 0.18422717 |
| 13.03556 | 0.117302116 | 0.145116606 | -0.078084107 | 0.17773541 |
| 13.0458 | 0.121765241 | 0.146353649 | -0.070771396 | 0.17031954 |
| 13.05604 | 0.124635707 | 0.148722427 | -0.06378926 | 0.1649338 |
| 13.06628 | 0.126479135 | 0.151557898 | -0.054964187 | 0.15951395 |
| 13.07652 | 0.12653038 | 0.154490486 | -0.046738233 | 0.15584865 |
| 13.08676 | 0.124691044 | 0.157870445 | -0.037508556 | 0.15293139 |
| 13.097 | 0.120788114 | 0.160719424 | -0.028861767 | 0.15123779 |
| 13.10724 | 0.115993893 | 0.163369711 | -0.020432432 | 0.15058211 |
| 13.11748 | 0.109391456 | 0.165264837 | -0.012391813 | 0.1504802 |
| 13.12772 | 0.102733356 | 0.167282187 | -0.00518676 | 0.1515481 |
| 13.13796 | 0.095475266 | 0.168185352 | 0.000842876 | 0.15246539 |
| 13.1482 | 0.088197295 | 0.168689205 | 0.006104434 | 0.15439765 |
| 13.15844 | 0.081170391 | 0.168481092 | 0.010170355 | 0.15514956 |
| 13.16868 | 0.07413446 | 0.167609218 | 0.013301354 | 0.15674893 |
| 13.17892 | 0.066784085 | 0.166476195 | 0.015461915 | 0.1575501 |
| 13.18916 | 0.059257826 | 0.165058279 | 0.017541602 | 0.15895015 |
| 13.1994 | 0.05123877 | 0.16418242 | 0.018955604 | 0.16055045 |
| 13.20964 | 0.043948899 | 0.163102565 | 0.021471511 | 0.16272292 |
| 13.21988 | 0.035800618 | 0.161768632 | 0.02402234 | 0.16611201 |
| 13.23012 | 0.028889347 | 0.159377806 | 0.028388782 | 0.16996711 |
| 13.24036 | 0.023012736 | 0.155991373 | 0.033069917 | 0.17416413 |
| 13.2506 | 0.018176202 | 0.150831452 | 0.039395179 | 0.17821376 |
| 13.26084 | 0.015194022 | 0.144831723 | 0.045555418 | 0.1822313 |
| 13.27108 | 0.013814777 | 0.1390517 | 0.052558417 | 0.18545978 |
| 13.28132 | 0.01462879 | 0.134486762 | 0.058809574 | 0.18813575 |
| 13.29156 | 0.016479171 | 0.131536767 | 0.064261257 | 0.19024347 |
| 13.3018 | 0.020231762 | 0.131190933 | 0.068749326 | 0.19240167 |
| 13.31204 | 0.024759074 | 0.132600983 | 0.072425591 | 0.19413264 |
| 13.32228 | 0.030039694 | 0.135078791 | 0.074401872 | 0.19557899 |
| 13.33252 | 0.035458528 | 0.138219361 | 0.076094655 | 0.19721647 |
| 13.34276 | 0.040514923 | 0.139819294 | 0.076842604 | 0.19801638 |
| 13.353 | 0.045057703 | 0.141399855 | 0.076973181 | 0.19798096 |
| 13.36324 | 0.049350329 | 0.141909626 | 0.077315982 | 0.19766308 |
| 13.37348 | 0.05326445 | 0.14154411 | 0.076181915 | 0.19616153 |
| 13.38372 | 0.056717894 | 0.141605449 | 0.075569076 | 0.19433454 |
| 13.39396 | 0.060349992 | 0.141313639 | 0.073460107 | 0.19195662 |
| 13.4042 | 0.064580652 | 0.141869133 | 0.070239412 | 0.19001505 |
| 13.41444 | 0.069015966 | 0.142532681 | 0.065758965 | 0.18744679 |
| 13.42468 | 0.073484881 | 0.144188458 | 0.060118327 | 0.18635822 |
| 13.43492 | 0.078779667 | 0.145777536 | 0.052424705 | 0.18489025 |
| 13.44516 | 0.083549343 | 0.147761002 | 0.044281702 | 0.18444331 |
| 13.4554 | 0.08828673 | 0.150294804 | 0.034746514 | 0.1839124 |
| 13.46564 | 0.092543626 | 0.153319465 | 0.025930108 | 0.18447996 |
| 13.47588 | 0.096192049 | 0.157000527 | 0.016912417 | 0.18535826 |
| 13.48612 | 0.099369203 | 0.160966319 | 0.009604849 | 0.18640452 |
| 13.49636 | 0.101649784 | 0.166198776 | 0.003634197 | 0.18788259 |
| 13.5066 | 0.103646878 | 0.171742895 | -0.000213203 | 0.18922087 |
| 13.51684 | 0.104635665 | 0.17765356 | -0.002120939 | 0.18982206 |
| 13.52708 | 0.105225385 | 0.184779736 | -0.00195373 | 0.19003399 |
| 13.53732 | 0.104829586 | 0.191309425 | -0.000613447 | 0.18899394 |
| 13.54756 | 0.103845815 | 0.197903749 | 0.002249425 | 0.18762398 |
| 13.5578 | 0.102244419 | 0.204608561 | 0.005669604 | 0.18474009 |
| 13.56804 | 0.099876494 | 0.209674074 | 0.009519941 | 0.18214346 |
| 13.57828 | 0.097340937 | 0.214160479 | 0.013727876 | 0.17895451 |
| 13.58852 | 0.094503957 | 0.216974389 | 0.017220582 | 0.17539549 |
| 13.59876 | 0.091658315 | 0.218795885 | 0.020830424 | 0.17172036 |
| 13.609 | 0.088582053 | 0.219612665 | 0.0240287 | 0.16769482 |
| 13.61924 | 0.085781531 | 0.218740189 | 0.026217128 | 0.16312467 |
| 13.62948 | 0.082493115 | 0.217586414 | 0.028253353 | 0.15836297 |
| 13.63972 | 0.078885905 | 0.215442801 | 0.029325518 | 0.15358857 |
| 13.64996 | 0.074827634 | 0.212549455 | 0.029669145 | 0.14974179 |
| 13.6602 | 0.069548622 | 0.209602573 | 0.029431597 | 0.14717706 |
| 13.67044 | 0.06398016 | 0.20585716 | 0.028190219 | 0.1470526 |
| 13.68068 | 0.057344404 | 0.202693669 | 0.026576698 | 0.14842644 |
| 13.69092 | 0.050345444 | 0.199289062 | 0.023930429 | 0.15192863 |
| 13.70116 | 0.04320928 | 0.197008411 | 0.020825096 | 0.15620995 |
| 13.7114 | 0.036394277 | 0.195154724 | 0.017303757 | 0.16126486 |
| 13.72164 | 0.030351478 | 0.194125513 | 0.012565725 | 0.16535217 |
| 13.73188 | 0.025079789 | 0.193569093 | 0.007985406 | 0.16996229 |
| 13.74212 | 0.021050221 | 0.193429494 | 0.002525176 | 0.17304723 |
| 13.75236 | 0.018694575 | 0.192429199 | -0.00260162 | 0.17615056 |
| 13.7626 | 0.017158591 | 0.19183779 | -0.007643751 | 0.17890411 |
| 13.77284 | 0.017629319 | 0.189604294 | -0.012399527 | 0.1814079 |
| 13.78308 | 0.018426659 | 0.187597693 | -0.016463667 | 0.18429008 |
| 13.79332 | 0.021793228 | 0.183423283 | -0.019379362 | 0.18796438 |
| 13.80356 | 0.025467244 | 0.180407561 | -0.021332077 | 0.1922373 |
| 13.8138 | 0.031117777 | 0.175802024 | -0.022061165 | 0.19745721 |
| 13.82404 | 0.03737724 | 0.172680798 | -0.022099262 | 0.20322532 |
| 13.83428 | 0.044929894 | 0.169959082 | -0.020743475 | 0.20942305 |
| 13.84452 | 0.052869084 | 0.169093696 | -0.018949108 | 0.21514491 |
| 13.85476 | 0.061077036 | 0.169938971 | -0.01691827 | 0.22090734 |
| 13.865 | 0.069070581 | 0.172829601 | -0.015106119 | 0.22498197 |
| 13.87524 | 0.076553404 | 0.177833482 | -0.013204884 | 0.22845688 |
| 13.88548 | 0.083589766 | 0.184666589 | -0.01200754 | 0.23091013 |
| 13.89572 | 0.089690404 | 0.193079233 | -0.011709527 | 0.23221513 |
| 13.90596 | 0.095890909 | 0.202675124 | -0.011634644 | 0.23332021 |
| 13.9162 | 0.101291717 | 0.212896278 | -0.012005735 | 0.23341813 |
| 13.92644 | 0.107004099 | 0.223685397 | -0.012614811 | 0.23427685 |
| 13.93668 | 0.112135946 | 0.233741211 | -0.013113671 | 0.23433169 |
| 13.94692 | 0.117178172 | 0.242773783 | -0.013501693 | 0.23477376 |
| 13.95716 | 0.121894963 | 0.250919422 | -0.013511986 | 0.2350585 |
| 13.9674 | 0.12569808 | 0.256961732 | -0.013750441 | 0.23583252 |
| 13.97764 | 0.128284295 | 0.261144241 | -0.013817845 | 0.23634897 |
| 13.98788 | 0.13037492 | 0.263416613 | -0.01371245 | 0.23726968 |
| 13.99812 | 0.130625337 | 0.263074012 | -0.014029805 | 0.23783203 |
| 14.00836 | 0.12990497 | 0.261104991 | -0.014242604 | 0.23902341 |
| 14.0186 | 0.127644584 | 0.2569925 | -0.014179369 | 0.23964353 |
| 14.02884 | 0.124638943 | 0.251865397 | -0.013759723 | 0.24027806 |
| 14.03908 | 0.120627588 | 0.245438873 | -0.012594851 | 0.24090224 |
| 14.04932 | 0.116201154 | 0.238333113 | -0.010770044 | 0.24092431 |
| 14.05956 | 0.111373181 | 0.231513267 | -0.008319193 | 0.24103957 |
| 14.0698 | 0.106927977 | 0.22504357 | -0.004827866 | 0.2413281 |
| 14.08004 | 0.102623727 | 0.219592743 | -0.001619484 | 0.24187057 |
| 14.09028 | 0.099016608 | 0.214584318 | 0.001937149 | 0.24301083 |
| 14.10052 | 0.095894299 | 0.211796502 | 0.005193056 | 0.24489631 |
| 14.11076 | 0.093185318 | 0.209420252 | 0.007489761 | 0.24707439 |
| 14.121 | 0.090848486 | 0.209058241 | 0.009275097 | 0.24971417 |
| 14.13124 | 0.0886993 | 0.208938012 | 0.00960513 | 0.25213044 |
| 14.14148 | 0.08649383 | 0.209437673 | 0.009612053 | 0.25420169 |
| 14.15172 | 0.084679299 | 0.209289399 | 0.007929097 | 0.25578348 |
| 14.16196 | 0.0828265 | 0.209083924 | 0.006244674 | 0.25719603 |
| 14.1722 | 0.081326681 | 0.207519578 | 0.003444757 | 0.2585872 |
| 14.18244 | 0.080538101 | 0.205490906 | 0.000989186 | 0.26024037 |
| 14.19268 | 0.080258947 | 0.203004317 | -0.001447914 | 0.26264704 |
| 14.20292 | 0.080301876 | 0.200448293 | -0.003829959 | 0.2658856 |
| 14.21316 | 0.081007274 | 0.1990062 | -0.005305844 | 0.27000045 |
| 14.2234 | 0.081896668 | 0.197757214 | -0.006311601 | 0.27512913 |
| 14.23364 | 0.082441153 | 0.197885609 | -0.006344038 | 0.28065452 |
| 14.24388 | 0.082732793 | 0.198325025 | -0.006073133 | 0.28723244 |
| 14.25412 | 0.082127633 | 0.198982675 | -0.005289855 | 0.29342137 |
| 14.26436 | 0.08062633 | 0.198936019 | -0.004055643 | 0.29933525 |
| 14.2746 | 0.077409973 | 0.198377422 | -0.002950525 | 0.30439485 |
| 14.28484 | 0.07292616 | 0.196361096 | -0.002247552 | 0.30765991 |
| 14.29508 | 0.067174447 | 0.193713511 | -0.001481502 | 0.30867098 |
| 14.30532 | 0.060316107 | 0.190495018 | -0.00142068 | 0.3077183 |
| 14.31556 | 0.05247819 | 0.187888075 | -0.001487217 | 0.30413391 |
| 14.3258 | 0.04499891 | 0.185358613 | -0.00163356 | 0.29891462 |
| 14.33604 | 0.037915484 | 0.184658826 | -0.001202298 | 0.29197404 |
| 14.34628 | 0.032167012 | 0.185115906 | -0.000860431 | 0.28451801 |
| 14.35652 | 0.028292029 | 0.187071337 | 0.001039365 | 0.27759738 |
| 14.36676 | 0.026511865 | 0.189881187 | 0.003886719 | 0.27061706 |
| 14.377 | 0.027685642 | 0.192973461 | 0.007502144 | 0.26572896 |

| | | | | |
|---|---|---|---|---|
| 11.98083 | -0.022352899 | 0.204884563 | -0.049080335 | 0.216849766 |
| 11.99107 | -0.014918684 | 0.199675644 | -0.041554755 | 0.214398248 |
| 12.00131 | -0.008194848 | 0.195752389 | -0.03429579 | 0.212819807 |
| 12.01155 | -0.001304066 | 0.193222657 | -0.026433838 | 0.211951058 |
| 12.02179 | 0.006021585 | 0.191556318 | -0.019077637 | 0.211752969 |
| 12.03203 | 0.012553697 | 0.190261095 | -0.011249904 | 0.210600877 |
| 12.04227 | 0.019926179 | 0.189256835 | -0.002981016 | 0.209532342 |
| 12.05252 | 0.026667607 | 0.187779206 | 0.005666597 | 0.207247346 |
| 12.06275 | 0.033819992 | 0.185896933 | 0.015172914 | 0.204171974 |
| 12.073 | 0.040332942 | 0.183254719 | 0.025788348 | 0.199942871 |
| 12.08323 | 0.046066117 | 0.179742999 | 0.036642946 | 0.195130232 |
| 12.09348 | 0.051027992 | 0.175031786 | 0.048387126 | 0.189297264 |
| 12.10371 | 0.05456743 | 0.169794049 | 0.059727381 | 0.183877707 |
| 12.11395 | 0.05652598 | 0.163529018 | 0.070324866 | 0.178055989 |
| 12.1242 | 0.056815133 | 0.156965659 | 0.079481855 | 0.173497143 |
| 12.13443 | 0.055119132 | 0.150349171 | 0.086541374 | 0.170024605 |
| 12.14468 | 0.052124072 | 0.143725214 | 0.091174967 | 0.167730389 |
| 12.15491 | 0.047497576 | 0.138136804 | 0.093375385 | 0.167466443 |
| 12.16516 | 0.041585147 | 0.133764251 | 0.093154129 | 0.16837807 |
| 12.1754 | 0.035039841 | 0.129837567 | 0.090867541 | 0.169492752 |
| 12.18564 | 0.027490112 | 0.127582256 | 0.087229707 | 0.172003504 |
| 12.19588 | 0.020110679 | 0.125622514 | 0.082550876 | 0.173094922 |
| 12.20611 | 0.012860961 | 0.123760196 | 0.077659243 | 0.174428703 |
| 12.21636 | 0.005425047 | 0.121829378 | 0.072401388 | 0.1741652 |
| 12.22659 | -0.000620455 | 0.119035726 | 0.067477199 | 0.173817646 |
| 12.23684 | -0.006709767 | 0.116022202 | 0.062551714 | 0.172756818 |
| 12.24708 | -0.010630132 | 0.112442643 | 0.058421445 | 0.171770079 |
| 12.25732 | -0.013907321 | 0.10975466 | 0.053706865 | 0.170994203 |
| 12.26756 | -0.015103084 | 0.107341584 | 0.049594572 | 0.171092694 |
| 12.2778 | -0.014744228 | 0.107782428 | 0.045008088 | 0.171491439 |
| 12.28804 | -0.012325456 | 0.108839185 | 0.040332346 | 0.172069098 |
| 12.29828 | -0.008679103 | 0.112402084 | 0.034926475 | 0.17264232 |
| 12.30852 | -0.00332515 | 0.116344119 | 0.029535415 | 0.172867228 |
| 12.31876 | 0.00222526 | 0.120647844 | 0.022944053 | 0.172444763 |
| 12.329 | 0.008013388 | 0.124770565 | 0.016006233 | 0.172142614 |
| 12.33924 | 0.013351679 | 0.12850156 | 0.008366204 | 0.171938155 |
| 12.34948 | 0.017400047 | 0.132355337 | 0.00012398 | 0.172665669 |
| 12.35972 | 0.020807558 | 0.13579796 | -0.008784832 | 0.174288073 |
| 12.36996 | 0.0226615 | 0.139518499 | -0.017793252 | 0.17683909 |
| 12.3802 | 0.024294697 | 0.142705979 | -0.026997109 | 0.180893188 |
| 12.39044 | 0.025025063 | 0.145813389 | -0.036567086 | 0.184850143 |
| 12.40068 | 0.025654884 | 0.147842492 | -0.045461687 | 0.189826809 |
| 12.41092 | 0.026587664 | 0.149484958 | -0.054534051 | 0.194808185 |
| 12.42116 | 0.027221435 | 0.150119446 | -0.062867223 | 0.199289095 |
| 12.4314 | 0.028104252 | 0.150386159 | -0.070860348 | 0.203897475 |
| 12.44164 | 0.028692662 | 0.150513618 | -0.078301772 | 0.207807777 |
| 12.45188 | 0.029224571 | 0.150491343 | -0.084560429 | 0.210832191 |
| 12.46212 | 0.029514273 | 0.150675506 | -0.090162911 | 0.213406591 |
| 12.47236 | 0.030026582 | 0.150546862 | -0.094485214 | 0.214724673 |
| 12.4826 | 0.030855322 | 0.150938429 | -0.09728334 | 0.216449116 |
| 12.49284 | 0.032610743 | 0.151122659 | -0.098829358 | 0.21705367 |
| 12.50308 | 0.035055922 | 0.151239023 | -0.099135209 | 0.219112976 |
| 12.51332 | 0.038517637 | 0.151785568 | -0.098299992 | 0.220713257 |
| 12.52356 | 0.043115119 | 0.152275422 | -0.096717334 | 0.223632765 |
| 12.5338 | 0.048573398 | 0.153093881 | -0.094676958 | 0.226811645 |
| 12.54404 | 0.054454315 | 0.154345248 | -0.092348035 | 0.229874039 |
| 12.55428 | 0.061049918 | 0.155312537 | -0.08969568 | 0.232139762 |
| 12.56452 | 0.067847587 | 0.157304252 | -0.086668884 | 0.232453973 |
| 12.57476 | 0.075129189 | 0.158823145 | -0.083496525 | 0.230832659 |
| 12.585 | 0.082455738 | 0.160769914 | -0.079459033 | 0.226040544 |
| 12.59524 | 0.090211373 | 0.163029321 | -0.07490186 | 0.218868728 |
| 12.60548 | 0.098150883 | 0.165057296 | -0.069758807 | 0.207817664 |
| 12.61572 | 0.10687088 | 0.167681619 | -0.064307814 | 0.195843733 |
| 12.62596 | 0.114923677 | 0.169710659 | -0.058879665 | 0.18127944 |
| 12.6362 | 0.123789191 | 0.172460888 | -0.054332819 | 0.167082156 |
| 12.64644 | 0.131587181 | 0.175213486 | -0.050301468 | 0.153599099 |
| 12.65668 | 0.138883687 | 0.177608825 | -0.048208409 | 0.142339144 |
| 12.66692 | 0.144818384 | 0.179891932 | -0.046849315 | 0.134087218 |
| 12.67716 | 0.149144817 | 0.182865296 | -0.04770435 | 0.129298897 |
| 12.6874 | 0.151200819 | 0.185104873 | -0.049514765 | 0.127935101 |
| 12.69764 | 0.151046503 | 0.187786775 | -0.052368897 | 0.12926121 |
| 12.70788 | 0.148905161 | 0.190788809 | -0.056279535 | 0.132255306 |
| 12.71812 | 0.144600507 | 0.193120566 | -0.060540069 | 0.136892513 |
| 12.72836 | 0.13866314 | 0.196266311 | -0.065316573 | 0.141474582 |
| 12.7386 | 0.131710337 | 0.1982638 | -0.070506761 | 0.146728397 |
| 12.74884 | 0.123475113 | 0.20087192 | -0.075783129 | 0.151682572 |
| 12.75908 | 0.115336403 | 0.202072839 | -0.081420665 | 0.155756786 |
| 12.76932 | 0.106797701 | 0.203548715 | -0.087271493 | 0.159706788 |
| 12.77956 | 0.097910284 | 0.203722222 | -0.092878125 | 0.16189538 |
| 12.7898 | 0.089740617 | 0.204218895 | -0.09886353 | 0.163768812 |
| 12.80004 | 0.081209786 | 0.203870603 | -0.104223813 | 0.16433496 |
| 12.81028 | 0.073834365 | 0.203294299 | -0.109547637 | 0.164522153 |
| 12.82052 | 0.066648643 | 0.202420183 | -0.114682034 | 0.164243652 |
| 12.83076 | 0.060883765 | 0.200836564 | -0.118981217 | 0.164482105 |
| 12.841 | 0.056607445 | 0.199506248 | -0.123433788 | 0.164558776 |
| 12.85124 | 0.053984506 | 0.197498037 | -0.127301928 | 0.166466758 |
| 12.86148 | 0.053102924 | 0.196391517 | -0.131056093 | 0.169032306 |
| 12.87172 | 0.054880705 | 0.195295215 | -0.134118967 | 0.173280522 |
| 12.88196 | 0.057557865 | 0.195740823 | -0.137083741 | 0.178328561 |
| 12.8922 | 0.062723827 | 0.196942973 | -0.139417625 | 0.184716434 |
| 12.90244 | 0.068237308 | 0.199785369 | -0.14157447 | 0.190267232 |
| 12.91268 | 0.074996149 | 0.20293075 | -0.142470472 | 0.196055681 |
| 12.92292 | 0.08159823 | 0.207412586 | -0.143897086 | 0.200216834 |
| 12.93316 | 0.088010461 | 0.211867309 | -0.144656157 | 0.203522637 |
| 12.9434 | 0.094168607 | 0.216770644 | -0.145070247 | 0.205360821 |
| 12.95364 | 0.099297656 | 0.22100153 | -0.145716315 | 0.206638702 |
| 12.96388 | 0.103976196 | 0.224953723 | -0.145921546 | 0.207408895 |
| 12.97412 | 0.106679556 | 0.227525053 | -0.145981654 | 0.208292911 |
| 12.98436 | 0.108953036 | 0.228685486 | -0.145470872 | 0.209701531 |
| 12.9946 | 0.108576351 | 0.228248564 | -0.144404514 | 0.211077833 |
| 13.00484 | 0.106860747 | 0.225857452 | -0.141858995 | 0.213412663 |
| 13.01508 | 0.102728828 | 0.222605585 | -0.138665415 | 0.215212488 |
| 13.02532 | 0.096622407 | 0.218010329 | -0.133608212 | 0.217270965 |
| 13.03556 | 0.088936145 | 0.213161785 | -0.127319566 | 0.217767829 |
| 13.0458 | 0.079925163 | 0.20864022 | -0.119520853 | 0.217682103 |
| 13.05604 | 0.07012672 | 0.204983511 | -0.110439207 | 0.216493688 |
| 13.06628 | 0.061274555 | 0.202434276 | -0.100371626 | 0.214283608 |
| 13.07652 | 0.052552354 | 0.201938607 | -0.089355611 | 0.212071186 |
| 13.08676 | 0.045404333 | 0.20308524 | -0.07805352 | 0.209892622 |
| 13.097 | 0.039613771 | 0.206528105 | -0.066618537 | 0.208696825 |
| 13.10724 | 0.035080822 | 0.211489908 | -0.055612735 | 0.208648654 |
| 13.11748 | 0.03183371 | 0.217834914 | -0.045259046 | 0.210068467 |
| 13.12772 | 0.028791736 | 0.224968518 | -0.035981207 | 0.212642847 |
| 13.13796 | 0.026482252 | 0.231485851 | -0.028423402 | 0.215906889 |
| 13.1482 | 0.024608376 | 0.237688009 | -0.021571717 | 0.219700777 |
| 13.15844 | 0.022056344 | 0.242623301 | -0.017041723 | 0.223707633 |
| 13.16868 | 0.019926769 | 0.246403071 | -0.013630198 | 0.227083078 |
| 13.17892 | 0.017574146 | 0.249162453 | -0.011745651 | 0.230526761 |
| 13.18916 | 0.01493992 | 0.251017061 | -0.010862926 | 0.23291616 |
| 13.1994 | 0.011236199 | 0.253277434 | -0.010979925 | 0.235853296 |
| 13.20964 | 0.007563593 | 0.254624299 | -0.011903537 | 0.237277368 |
| 13.21988 | 0.001747446 | 0.256809242 | -0.012990489 | 0.238868218 |
| 13.23012 | -0.004676266 | 0.258282025 | -0.014846057 | 0.238838128 |
| 13.24036 | -0.013280269 | 0.25940118 | -0.015894646 | 0.238877618 |
| 13.2506 | -0.022722596 | 0.258917146 | -0.01809308 | 0.237539047 |
| 13.26084 | -0.032933652 | 0.257856617 | -0.019767332 | 0.235032219 |
| 13.27108 | -0.043556377 | 0.254132372 | -0.021790096 | 0.23178111 |
| 13.28132 | -0.052954173 | 0.250035181 | -0.023708396 | 0.227420456 |
| 13.29156 | -0.061023218 | 0.244075191 | -0.02604634 | 0.221776064 |
| 13.3018 | -0.066424293 | 0.237821811 | -0.028057385 | 0.21531045 |
| 13.31204 | -0.069526702 | 0.230694326 | -0.029559954 | 0.209031192 |
| 13.32228 | -0.068945491 | 0.22424387 | -0.031222456 | 0.202268925 |
| 13.33252 | -0.06608657 | 0.217301897 | -0.031980275 | 0.196524096 |
| 13.34276 | -0.060723932 | 0.211071585 | -0.032812419 | 0.192146931 |
| 13.353 | -0.054009206 | 0.205724449 | -0.033331503 | 0.189277926 |
| 13.36324 | -0.046991185 | 0.200914422 | -0.034760561 | 0.187658412 |
| 13.37348 | -0.039759301 | 0.19793658 | -0.036415121 | 0.186949724 |
| 13.38372 | -0.034099939 | 0.195722547 | -0.039569345 | 0.187029321 |
| 13.39396 | -0.029078182 | 0.195228489 | -0.043470264 | 0.187185479 |
| 13.4042 | -0.026124866 | 0.196132687 | -0.048870597 | 0.187475059 |
| 13.41444 | -0.023761963 | 0.199074194 | -0.054163385 | 0.187224474 |
| 13.42468 | -0.022690186 | 0.202081485 | -0.060199238 | 0.187528648 |
| 13.43492 | -0.021609165 | 0.207193944 | -0.065639245 | 0.187456264 |
| 13.44516 | -0.020754258 | 0.212420099 | -0.070769657 | 0.187899061 |
| 13.4554 | -0.019236937 | 0.217945228 | -0.074593026 | 0.188720443 |
| 13.46564 | -0.017661883 | 0.223281589 | -0.078141559 | 0.189779786 |
| 13.47588 | -0.015589513 | 0.227556913 | -0.080367002 | 0.191244022 |
| 13.48612 | -0.013131457 | 0.231306712 | -0.082313882 | 0.192536009 |
| 13.49636 | -0.010568378 | 0.233063069 | -0.083267272 | 0.193673614 |
| 13.5066 | -0.00814192 | 0.234256397 | -0.084067145 | 0.194031559 |
| 13.51684 | -0.006410614 | 0.233874763 | -0.083720156 | 0.193726763 |
| 13.52708 | -0.004390021 | 0.233375867 | -0.083386401 | 0.192053647 |
| 13.53732 | -0.003484533 | 0.231993187 | -0.08226218 | 0.189534137 |
| 13.54756 | -0.003407182 | 0.231860227 | -0.080932016 | 0.187592505 |
| 13.5578 | -0.003376948 | 0.232255159 | -0.079461256 | 0.184676536 |
| 13.56804 | -0.004359453 | 0.233723406 | -0.077707303 | 0.183020655 |
| 13.57828 | -0.005392781 | 0.23620133 | -0.076529926 | 0.182208543 |
| 13.58852 | -0.006479428 | 0.239623391 | -0.075285672 | 0.182458516 |
| 13.59876 | -0.007368394 | 0.24277243 | -0.074114952 | 0.182436487 |
| 13.609 | -0.007633047 | 0.24590026 | -0.073358533 | 0.182829016 |
| 13.61924 | -0.007448363 | 0.247559736 | -0.072042716 | 0.182456677 |
| 13.62948 | -0.006392161 | 0.248812097 | -0.071076227 | 0.180966364 |
| 13.63972 | -0.004800475 | 0.247944698 | -0.069946685 | 0.17837805 |
| 13.64996 | -0.003139226 | 0.246464862 | -0.068347116 | 0.174998535 |
| 13.6602 | -0.00049182 | 0.244063013 | -0.067020058 | 0.171421528 |
| 13.67044 | 0.001446059 | 0.240712428 | -0.065814751 | 0.166993454 |
| 13.68068 | 0.004323194 | 0.237541497 | -0.064686921 | 0.163078544 |
| 13.69092 | 0.006269528 | 0.234328138 | -0.064317659 | 0.159185384 |
| 13.70116 | 0.00899053 | 0.232100089 | -0.064172593 | 0.155167509 |
| 13.7114 | 0.011384229 | 0.230538253 | -0.064871473 | 0.151131992 |
| 13.72164 | 0.013915207 | 0.231177068 | -0.066755936 | 0.147376256 |
| 13.73188 | 0.016303502 | 0.232956979 | -0.068734316 | 0.143860863 |
| 13.74212 | 0.018705569 | 0.236503218 | -0.072479034 | 0.140631171 |
| 13.75236 | 0.020887212 | 0.241476945 | -0.075923497 | 0.138518606 |
| 13.7626 | 0.022371802 | 0.246360161 | -0.080218734 | 0.136728042 |
| 13.77284 | 0.024210945 | 0.252005581 | -0.084492399 | 0.136278275 |
| 13.78308 | 0.02503366 | 0.255640846 | -0.088120831 | 0.136754278 |
| 13.79332 | 0.025689685 | 0.258712831 | -0.090888424 | 0.138999616 |
| 13.80356 | 0.026286746 | 0.260144656 | -0.092564333 | 0.14264584 |
| 13.8138 | 0.026907419 | 0.26032091 | -0.09263228 | 0.147852777 |
| 13.82404 | 0.027518995 | 0.259687306 | -0.09163048 | 0.154007736 |
| 13.83428 | 0.029274014 | 0.258254922 | -0.088958623 | 0.1599939 |
| 13.84452 | 0.031645147 | 0.256710496 | -0.085468759 | 0.166309511 |
| 13.85476 | 0.035537048 | 0.254697364 | -0.081343036 | 0.170950168 |
| 13.865 | 0.040641166 | 0.253164299 | -0.076381352 | 0.175766667 |
| 13.87524 | 0.047189525 | 0.251821606 | -0.072130744 | 0.179513795 |
| 13.88548 | 0.054660673 | 0.25068652 | -0.067509025 | 0.182960139 |
| 13.89572 | 0.062880863 | 0.250276594 | -0.063993659 | 0.18637671 |
| 13.90596 | 0.071060572 | 0.250438231 | -0.060837384 | 0.189651264 |
| 13.9162 | 0.078988439 | 0.251410424 | -0.058776784 | 0.192364781 |
| 13.92644 | 0.085915434 | 0.252778886 | -0.05771741 | 0.194171556 |
| 13.93668 | 0.091522816 | 0.255059712 | -0.057691133 | 0.195631456 |
| 13.94692 | 0.096052157 | 0.257854566 | -0.05823641 | 0.19451133 |
| 13.95716 | 0.099071738 | 0.261094398 | -0.059665268 | 0.193247054 |
| 13.9674 | 0.100976119 | 0.264991841 | -0.061439956 | 0.189957277 |
| 13.97764 | 0.101670142 | 0.268605838 | -0.063028688 | 0.186980286 |
| 13.98788 | 0.102221808 | 0.272668263 | -0.06440452 | 0.183433343 |
| 13.99812 | 0.101822296 | 0.275756316 | -0.06502358 | 0.180878621 |
| 14.00836 | 0.101077236 | 0.278509903 | -0.065610658 | 0.179229267 |
| 14.0186 | 0.100383693 | 0.280154852 | -0.065071514 | 0.178905158 |
| 14.02884 | 0.098882833 | 0.280374029 | -0.064854246 | 0.179297726 |
| 14.03908 | 0.097307354 | 0.279508216 | -0.064282451 | 0.180723192 |
| 14.04932 | 0.095133533 | 0.276850978 | -0.063931256 | 0.181649896 |
| 14.05956 | 0.092283449 | 0.273016101 | -0.063964789 | 0.183343733 |
| 14.0698 | 0.089582888 | 0.267641424 | -0.064360879 | 0.183575962 |
| 14.08004 | 0.085880072 | 0.26145472 | -0.065060672 | 0.184378805 |
| 14.09028 | 0.081936768 | 0.253561389 | -0.06575629 | 0.184100815 |
| 14.10052 | 0.077693199 | 0.245867658 | -0.066371497 | 0.18471501 |
| 14.11076 | 0.072464368 | 0.237341347 | -0.066809821 | 0.185000048 |
| 14.121 | 0.067363452 | 0.229113824 | -0.066406189 | 0.186254176 |
| 14.13124 | 0.060854351 | 0.221023016 | -0.065468072 | 0.187900398 |
| 14.14148 | 0.055179603 | 0.214037397 | -0.063585579 | 0.190199155 |
| 14.15172 | 0.048821967 | 0.207371343 | -0.06060537 | 0.193407852 |
| 14.16196 | 0.043826633 | 0.20129172 | -0.05686304 | 0.196085544 |
| 14.1722 | 0.039774302 | 0.196166698 | -0.052389124 | 0.199147132 |
| 14.18244 | 0.037601314 | 0.190663196 | -0.046293969 | 0.200881513 |
| 14.19268 | 0.037509215 | 0.186566858 | -0.040202471 | 0.2023817 |
| 14.20292 | 0.038911333 | 0.182794942 | -0.033457471 | 0.201826492 |
| 14.21316 | 0.042074364 | 0.180711268 | -0.025891586 | 0.200842095 |
| 14.2234 | 0.04549105 | 0.180370604 | -0.01799472 | 0.198503516 |
| 14.23364 | 0.049277851 | 0.181588695 | -0.01000825 | 0.195204507 |
| 14.24388 | 0.051999633 | 0.185090534 | -0.001921336 | 0.191419128 |
| 14.25412 | 0.053804557 | 0.190072348 | 0.006298458 | 0.187374648 |
| 14.26436 | 0.054108032 | 0.195526341 | 0.014457335 | 0.183283459 |
| 14.2746 | 0.052826214 | 0.202026413 | 0.0216702 | 0.178936366 |
| 14.28484 | 0.050771486 | 0.207628753 | 0.02916517 | 0.175038194 |
| 14.29508 | 0.047544561 | 0.212284825 | 0.035480905 | 0.171064418 |
| 14.30532 | 0.044277679 | 0.21565559 | 0.041367092 | 0.167092091 |
| 14.31556 | 0.040743118 | 0.21703515 | 0.046405371 | 0.16361574 |
| 14.3258 | 0.037668705 | 0.216779122 | 0.051318605 | 0.159984196 |
| 14.33604 | 0.035207245 | 0.21499472 | 0.055831574 | 0.157557376 |
| 14.34628 | 0.03333995 | 0.212227109 | 0.060087771 | 0.155598854 |
| 14.35652 | 0.032003557 | 0.208653343 | 0.064528793 | 0.155278305 |
| 14.36676 | 0.031432837 | 0.204726195 | 0.06916031 | 0.156801288 |
| 14.377 | 0.031202698 | 0.200448096 | 0.073947671 | 0.159993441 |

| | | | | |
|---|---|---|---|---|
| 11.98083 | 0.160268028 | 0.072587557 | -0.025221781 | 0.283612978 |
| 11.99107 | 0.159645049 | 0.069933857 | -0.023470741 | 0.284450339 |
| 12.00131 | 0.158433338 | 0.068549738 | -0.021405549 | 0.28502175 |
| 12.01155 | 0.157475843 | 0.067824843 | -0.018970532 | 0.285539615 |
| 12.02179 | 0.156190552 | 0.067783238 | -0.016640339 | 0.286232162 |
| 12.03203 | 0.154420429 | 0.068259942 | -0.013084653 | 0.286320266 |
| 12.04227 | 0.153079288 | 0.068895298 | -0.009812193 | 0.286051836 |
| 12.05252 | 0.151519645 | 0.070419199 | -0.00583361 | 0.284177572 |
| 12.06275 | 0.150311217 | 0.072205745 | -0.002072588 | 0.281352284 |
| 12.073 | 0.149111221 | 0.074608354 | 0.001359297 | 0.277116169 |
| 12.08323 | 0.148032425 | 0.076956845 | 0.004464818 | 0.271626271 |
| 12.09348 | 0.147059324 | 0.080057248 | 0.006563434 | 0.264844259 |
| 12.10371 | 0.146078656 | 0.083131019 | 0.008234553 | 0.257932465 |
| 12.11395 | 0.144860434 | 0.08618154 | 0.009144214 | 0.250666748 |
| 12.1242 | 0.143799496 | 0.089004228 | 0.010138995 | 0.24380262 |
| 12.13443 | 0.142264073 | 0.091297805 | 0.010726394 | 0.237595393 |
| 12.14468 | 0.141166826 | 0.093729796 | 0.012354523 | 0.232628239 |
| 12.15491 | 0.139149347 | 0.095198383 | 0.014326705 | 0.228350576 |
| 12.16516 | 0.137912632 | 0.097418765 | 0.017679535 | 0.225447837 |
| 12.1754 | 0.135749886 | 0.099741758 | 0.021438577 | 0.223287607 |
| 12.18564 | 0.134267865 | 0.103799187 | 0.026585337 | 0.222157704 |
| 12.19588 | 0.131926439 | 0.108700734 | 0.031613642 | 0.221663082 |
| 12.20611 | 0.129814064 | 0.115681513 | 0.036889461 | 0.221310974 |
| 12.21636 | 0.126905148 | 0.123588559 | 0.041643291 | 0.221664533 |
| 12.22659 | 0.123909931 | 0.132465432 | 0.04533266 | 0.221730398 |
| 12.23684 | 0.120080891 | 0.141123272 | 0.048108364 | 0.221744863 |
| 12.24708 | 0.115802927 | 0.148794765 | 0.049484372 | 0.221497602 |
| 12.25732 | 0.110990953 | 0.15530401 | 0.049592417 | 0.220870646 |
| 12.26756 | 0.106090211 | 0.160116721 | 0.04885189 | 0.219675711 |
| 12.2778 | 0.100965213 | 0.163620032 | 0.047180942 | 0.217966638 |
| 12.28804 | 0.0956672 | 0.165873243 | 0.045033911 | 0.215229154 |
| 12.29828 | 0.090711958 | 0.167820577 | 0.042876829 | 0.211929126 |
| 12.30852 | 0.085392596 | 0.16965577 | 0.040369894 | 0.207308224 |
| 12.31876 | 0.080225256 | 0.172124981 | 0.038269469 | 0.201804809 |
| 12.329 | 0.075023492 | 0.175422549 | 0.036230831 | 0.195473847 |
| 12.33924 | 0.069890297 | 0.179416133 | 0.034231653 | 0.188675548 |
| 12.34948 | 0.064849499 | 0.183917114 | 0.032604398 | 0.181067434 |
| 12.35972 | 0.059729271 | 0.188580322 | 0.030784352 | 0.173702524 |
| 12.36996 | 0.055129332 | 0.19266486 | 0.029471224 | 0.166377406 |
| 12.3802 | 0.050600513 | 0.196341645 | 0.027619792 | 0.159247773 |
| 12.39044 | 0.046055804 | 0.198682153 | 0.026661209 | 0.152378158 |
| 12.40068 | 0.041266153 | 0.20001141 | 0.025496144 | 0.146567286 |
| 12.41092 | 0.036942539 | 0.200266656 | 0.024357783 | 0.141254937 |
| 12.42116 | 0.032015651 | 0.199388777 | 0.024098764 | 0.137639413 |
| 12.4314 | 0.027052441 | 0.197883535 | 0.023574358 | 0.134970408 |
| 12.44164 | 0.022620333 | 0.195774894 | 0.023449436 | 0.134003338 |
| 12.45188 | 0.018211893 | 0.194517496 | 0.023551058 | 0.134083453 |
| 12.46212 | 0.014737124 | 0.193624151 | 0.023514404 | 0.134968231 |
| 12.47236 | 0.011711561 | 0.194608679 | 0.023336049 | 0.136258651 |
| 12.4826 | 0.010031533 | 0.19734457 | 0.02307998 | 0.137873468 |
| 12.49284 | 0.008872015 | 0.201549067 | 0.022091823 | 0.139136409 |
| 12.50308 | 0.009288347 | 0.207301784 | 0.020997744 | 0.140352752 |
| 12.51332 | 0.010056384 | 0.213715198 | 0.018983898 | 0.141589631 |
| 12.52356 | 0.011847255 | 0.220546216 | 0.016565706 | 0.142666109 |
| 12.5338 | 0.014107897 | 0.227136 | 0.013528415 | 0.144476062 |
| 12.54404 | 0.016678967 | 0.233494325 | 0.009989701 | 0.146609514 |
| 12.55428 | 0.018951987 | 0.23880692 | 0.006388309 | 0.149430618 |
| 12.56452 | 0.021082896 | 0.243667232 | 0.002498265 | 0.152723075 |
| 12.57476 | 0.022902164 | 0.247068861 | -0.000642607 | 0.15610906 |
| 12.585 | 0.023845223 | 0.249575358 | -0.003804881 | 0.159566116 |
| 12.59524 | 0.024713522 | 0.250441121 | -0.00607951 | 0.162391886 |
| 12.60548 | 0.025008145 | 0.250492873 | -0.008079693 | 0.164889121 |
| 12.61572 | 0.025014434 | 0.249123712 | -0.009689328 | 0.165978453 |
| 12.62596 | 0.024705057 | 0.247548239 | -0.011389171 | 0.166661101 |
| 12.6362 | 0.023843547 | 0.244901525 | -0.013724717 | 0.166017221 |
| 12.64644 | 0.022456127 | 0.242497355 | -0.016901424 | 0.164488988 |
| 12.65668 | 0.020233405 | 0.240379106 | -0.020554449 | 0.161898668 |
| 12.66692 | 0.017288498 | 0.238300072 | -0.025342301 | 0.1583776 |
| 12.67716 | 0.013653234 | 0.237089289 | -0.030348633 | 0.153365278 |
| 12.6874 | 0.009664037 | 0.236240045 | -0.035579572 | 0.147560616 |
| 12.69764 | 0.005792779 | 0.236244725 | -0.03996218 | 0.140968337 |
| 12.70788 | 0.002423643 | 0.236887156 | -0.043711887 | 0.134022715 |
| 12.71812 | -0.000161787 | 0.237837615 | -0.046215178 | 0.127358773 |
| 12.72836 | -0.001697178 | 0.239408347 | -0.047942601 | 0.122355995 |
| 12.7386 | -0.002496396 | 0.240950253 | -0.048194441 | 0.118921847 |
| 12.74884 | -0.002190697 | 0.242677769 | -0.047970722 | 0.11818987 |
| 12.75908 | -0.001889339 | 0.24384814 | -0.046779479 | 0.120074214 |
| 12.76932 | -0.001543449 | 0.245090525 | -0.045015302 | 0.124293305 |
| 12.77956 | -0.001653902 | 0.245278258 | -0.042404919 | 0.130040408 |
| 12.7898 | -0.002344776 | 0.245080962 | -0.039126251 | 0.13655546 |
| 12.80004 | -0.003994942 | 0.243872912 | -0.034932455 | 0.14340966 |
| 12.81028 | -0.006518822 | 0.242044533 | -0.029733667 | 0.149363692 |
| 12.82052 | -0.010138896 | 0.239519234 | -0.024441158 | 0.154128018 |
| 12.83076 | -0.014356733 | 0.236391611 | -0.018266232 | 0.1577022 |
| 12.841 | -0.019789255 | 0.233200517 | -0.012578617 | 0.159369758 |
| 12.85124 | -0.025660972 | 0.229450245 | -0.00694621 | 0.159890019 |
| 12.86148 | -0.032302223 | 0.226059736 | -0.002042038 | 0.158941097 |
| 12.87172 | -0.039206675 | 0.222106426 | 0.002294374 | 0.157125955 |
| 12.88196 | -0.046142014 | 0.218913281 | 0.005566228 | 0.154777148 |
| 12.8922 | -0.052673977 | 0.215214206 | 0.00824571 | 0.152072489 |
| 12.90244 | -0.058692367 | 0.212481466 | 0.010192036 | 0.149413356 |
| 12.91268 | -0.063280896 | 0.209703645 | 0.011338951 | 0.147130571 |
| 12.92292 | -0.066763554 | 0.208047926 | 0.011551803 | 0.145937112 |
| 12.93316 | -0.068860557 | 0.206881723 | 0.011144577 | 0.145154971 |
| 12.9434 | -0.069320444 | 0.206693506 | 0.010038237 | 0.145777763 |
| 12.95364 | -0.069042918 | 0.207501858 | 0.007957904 | 0.147282471 |
| 12.96388 | -0.067512731 | 0.208625071 | 0.005626576 | 0.149292753 |
| 12.97412 | -0.065665597 | 0.210385508 | 0.002725629 | 0.152061691 |
| 12.98436 | -0.06336071 | 0.212058664 | -0.000433733 | 0.154268469 |
| 12.9946 | -0.060744954 | 0.213583939 | -0.003509001 | 0.156534389 |
| 13.00484 | -0.057607145 | 0.214642899 | -0.006378698 | 0.158100891 |
| 13.01508 | -0.054479111 | 0.215227162 | -0.008945238 | 0.158977587 |
| 13.02532 | -0.049994001 | 0.215596986 | -0.011216115 | 0.159096584 |
| 13.03556 | -0.045594742 | 0.215630734 | -0.012844868 | 0.15902906 |
| 13.0458 | -0.039965924 | 0.215584124 | -0.013922834 | 0.15793938 |
| 13.05604 | -0.034383384 | 0.215642543 | -0.01488159 | 0.157002482 |
| 13.06628 | -0.028683369 | 0.216068121 | -0.015169472 | 0.155441908 |
| 13.07652 | -0.023236537 | 0.216697377 | -0.015634947 | 0.154168585 |
| 13.08676 | -0.018500521 | 0.217178964 | -0.015960222 | 0.152480191 |
| 13.097 | -0.01538059 | 0.218270254 | -0.016670166 | 0.151182463 |
| 13.10724 | -0.012901464 | 0.219087365 | -0.017646945 | 0.150127065 |
| 13.11748 | -0.012290804 | 0.219493205 | -0.019210095 | 0.149299445 |
| 13.12772 | -0.012427972 | 0.219421857 | -0.021133439 | 0.149060528 |
| 13.13796 | -0.014134702 | 0.218926419 | -0.023487524 | 0.149278242 |
| 13.1482 | -0.015580293 | 0.217411067 | -0.025949351 | 0.15012436 |
| 13.15844 | -0.017972428 | 0.21532133 | -0.028440064 | 0.151526231 |
| 13.16868 | -0.019821481 | 0.212924319 | -0.030440716 | 0.153579926 |
| 13.17892 | -0.0211268 | 0.210606245 | -0.031834582 | 0.156567931 |
| 13.18916 | -0.021437969 | 0.208077275 | -0.032776542 | 0.159681513 |
| 13.1994 | -0.020962366 | 0.20655901 | -0.032259082 | 0.163208514 |
| 13.20964 | -0.018813071 | 0.204909669 | -0.031360327 | 0.166885465 |
| 13.21988 | -0.016046518 | 0.204315107 | -0.029143039 | 0.169783184 |
| 13.23012 | -0.011783825 | 0.203800092 | -0.026748361 | 0.172283902 |
| 13.24036 | -0.007275901 | 0.203372153 | -0.023363824 | 0.173203206 |
| 13.2506 | -0.001859703 | 0.20321353 | -0.019902892 | 0.173674645 |
| 13.26084 | 0.003255757 | 0.202986934 | -0.016286165 | 0.17236724 |
| 13.27108 | 0.008491824 | 0.20233719 | -0.012747713 | 0.171351446 |
| 13.28132 | 0.012957916 | 0.201616856 | -0.009024939 | 0.169361662 |
| 13.29156 | 0.016570581 | 0.201108855 | -0.005784719 | 0.168236126 |
| 13.3018 | 0.019036894 | 0.200244362 | -0.002812725 | 0.167184872 |
| 13.31204 | 0.020496579 | 0.200486927 | -0.000144186 | 0.166568593 |
| 13.32228 | 0.019852537 | 0.200509941 | 0.001883138 | 0.166630288 |
| 13.33252 | 0.017882003 | 0.201573426 | 0.003169356 | 0.166512908 |
| 13.34276 | 0.014464138 | 0.202187154 | 0.003664458 | 0.166578635 |
| 13.353 | 0.009318591 | 0.2034647 | 0.003364501 | 0.166265122 |
| 13.36324 | 0.003199703 | 0.204034895 | 0.002073976 | 0.166363266 |
| 13.37348 | -0.003768732 | 0.203584835 | -9.24957E-05 | 0.166029769 |
| 13.38372 | -0.011130567 | 0.202771493 | -0.002717109 | 0.166470795 |
| 13.39396 | -0.018265551 | 0.200863839 | -0.00610335 | 0.16617359 |
| 13.4042 | -0.025253334 | 0.198339264 | -0.009491371 | 0.166818848 |
| 13.41444 | -0.031603672 | 0.19498243 | -0.013354767 | 0.16664443 |
| 13.42468 | -0.037288023 | 0.191621349 | -0.017015578 | 0.166723683 |
| 13.43492 | -0.042039794 | 0.187332753 | -0.020857602 | 0.166567183 |
| 13.44516 | -0.046086016 | 0.183476464 | -0.02459761 | 0.166294379 |
| 13.4554 | -0.049298391 | 0.179062779 | -0.028216204 | 0.166134595 |
| 13.46564 | -0.051719127 | 0.174985245 | -0.032210509 | 0.166406495 |
| 13.47588 | -0.053423223 | 0.171342824 | -0.035811184 | 0.167088388 |
| 13.48612 | -0.054512387 | 0.168149493 | -0.040082921 | 0.168464976 |
| 13.49636 | -0.055029297 | 0.165835393 | -0.043924297 | 0.170992814 |
| 13.5066 | -0.055465425 | 0.164461366 | -0.048015452 | 0.173714664 |
| 13.51684 | -0.055446499 | 0.163934496 | -0.051835124 | 0.177941188 |
| 13.52708 | -0.056190721 | 0.164378051 | -0.055678325 | 0.182189352 |
| 13.53732 | -0.056393021 | 0.165385797 | -0.058953645 | 0.186497521 |
| 13.54756 | -0.057341818 | 0.167413042 | -0.062272279 | 0.190390758 |
| 13.5578 | -0.057924143 | 0.17009554 | -0.064596757 | 0.192872089 |
| 13.56804 | -0.058992085 | 0.173333658 | -0.067394399 | 0.194101825 |
| 13.57828 | -0.059312453 | 0.177735101 | -0.069316506 | 0.193175496 |
| 13.58852 | -0.059539087 | 0.181910846 | -0.071207239 | 0.190263215 |
| 13.59876 | -0.059452683 | 0.187471597 | -0.0727651 | 0.185981191 |
| 13.609 | -0.05896533 | 0.192695286 | -0.074305783 | 0.180659312 |
| 13.61924 | -0.058099542 | 0.198214455 | -0.075711799 | 0.174684681 |
| 13.62948 | -0.056866978 | 0.203722227 | -0.076621619 | 0.169320282 |
| 13.63972 | -0.055466718 | 0.209389002 | -0.078092857 | 0.164560958 |
| 13.64996 | -0.053643309 | 0.215391029 | -0.079028077 | 0.160998877 |
| 13.6602 | -0.051105292 | 0.221789745 | -0.079997306 | 0.157683054 |
| 13.67044 | -0.048356043 | 0.2287248 | -0.080800878 | 0.155696962 |
| 13.68068 | -0.045118197 | 0.236375176 | -0.081195575 | 0.153524778 |
| 13.69092 | -0.04128704 | 0.244296822 | -0.081211566 | 0.151168969 |
| 13.70116 | -0.037520922 | 0.252257302 | -0.080533859 | 0.149367685 |
| 13.7114 | -0.033943361 | 0.260028367 | -0.079468563 | 0.147105857 |
| 13.72164 | -0.030913953 | 0.267054917 | -0.077766262 | 0.146137025 |
| 13.73188 | -0.02828846 | 0.273248461 | -0.075307157 | 0.145531274 |
| 13.74212 | -0.02615872 | 0.278205432 | -0.073218743 | 0.14662518 |
| 13.75236 | -0.024995097 | 0.282290971 | -0.070216736 | 0.14923135 |
| 13.7626 | -0.023602091 | 0.28480019 | -0.067641779 | 0.153526319 |
| 13.77284 | -0.022896491 | 0.28664261 | -0.064810126 | 0.159068025 |
| 13.78308 | -0.021473639 | 0.287477142 | -0.06254048 | 0.165437107 |
| 13.79332 | -0.020385098 | 0.286789541 | -0.060063513 | 0.172069957 |
| 13.80356 | -0.019118318 | 0.285982267 | -0.058095133 | 0.178376551 |
| 13.8138 | -0.017838496 | 0.283625867 | -0.056476784 | 0.183736484 |
| 13.82404 | -0.016725181 | 0.280782288 | -0.054949863 | 0.188038792 |
| 13.83428 | -0.01636811 | 0.276894207 | -0.053721148 | 0.190748757 |
| 13.84452 | -0.016374514 | 0.272328082 | -0.052537798 | 0.191834226 |
| 13.85476 | -0.016914644 | 0.267147292 | -0.05115784 | 0.192138102 |
| 13.865 | -0.018603916 | 0.261341702 | -0.049566052 | 0.190518223 |
| 13.87524 | -0.020338347 | 0.255437637 | -0.048215207 | 0.189001194 |
| 13.88548 | -0.022492034 | 0.249164435 | -0.046130525 | 0.186908012 |
| 13.89572 | -0.024610423 | 0.243501405 | -0.044842237 | 0.18439138 |
| 13.90596 | -0.027089787 | 0.238129411 | -0.043319582 | 0.182283692 |
| 13.9162 | -0.028946421 | 0.233746091 | -0.043302701 | 0.180312481 |
| 13.92644 | -0.030901357 | 0.230559794 | -0.043646156 | 0.17857441 |
| 13.93668 | -0.03286678 | 0.228740899 | -0.04541671 | 0.177325292 |
| 13.94692 | -0.034786496 | 0.227736443 | -0.048155514 | 0.176680362 |
| 13.95716 | -0.037267035 | 0.228598163 | -0.052203982 | 0.176880484 |
| 13.9674 | -0.039910827 | 0.229479058 | -0.056363032 | 0.177795614 |
| 13.97764 | -0.043222421 | 0.231410277 | -0.061198385 | 0.179695755 |
| 13.98788 | -0.04670857 | 0.233717134 | -0.066160937 | 0.182222844 |
| 13.99812 | -0.050765686 | 0.235993682 | -0.070954391 | 0.185582133 |
| 14.00836 | -0.054516517 | 0.238592797 | -0.075341242 | 0.189211524 |
| 14.0186 | -0.058266519 | 0.241451589 | -0.079628449 | 0.19324789 |
| 14.02884 | -0.061643992 | 0.2440141 | -0.083426357 | 0.197354 |
| 14.03908 | -0.064829283 | 0.246558738 | -0.086680053 | 0.201334918 |
| 14.04932 | -0.067377625 | 0.248590312 | -0.089279684 | 0.205201578 |
| 14.05956 | -0.069756984 | 0.250136297 | -0.091187427 | 0.20850744 |
| 14.0698 | -0.07147831 | 0.250744301 | -0.091788328 | 0.21146583 |
| 14.08004 | -0.072970901 | 0.24993043 | -0.091250039 | 0.214113519 |
| 14.09028 | -0.073617298 | 0.248365574 | -0.089502365 | 0.216056568 |
| 14.10052 | -0.073437161 | 0.244858918 | -0.085747357 | 0.218189123 |
| 14.11076 | -0.072591414 | 0.240458797 | -0.081275755 | 0.219534959 |
| 14.121 | -0.070673325 | 0.234340465 | -0.07535699 | 0.221227955 |
| 14.13124 | -0.067761669 | 0.227878869 | -0.068811168 | 0.222499468 |
| 14.14148 | -0.064207559 | 0.220144219 | -0.06195003 | 0.223727246 |
| 14.15172 | -0.060293665 | 0.212408814 | -0.055309609 | 0.224724379 |
| 14.16196 | -0.05575952 | 0.20490563 | -0.049158365 | 0.225289597 |
| 14.1722 | -0.051746334 | 0.197512536 | -0.043907603 | 0.226085907 |
| 14.18244 | -0.047596307 | 0.191201055 | -0.039592903 | 0.226066396 |
| 14.19268 | -0.043925787 | 0.185605445 | -0.036342768 | 0.226596398 |
| 14.20292 | -0.041018192 | 0.181541575 | -0.033879808 | 0.226292721 |
| 14.21316 | -0.038116114 | 0.177988807 | -0.031791238 | 0.226086426 |
| 14.2234 | -0.036400971 | 0.175974693 | -0.030357465 | 0.225517477 |
| 14.23364 | -0.034555611 | 0.174504407 | -0.028525725 | 0.224588965 |
| 14.24388 | -0.033363377 | 0.173916958 | -0.026617845 | 0.223006834 |
| 14.25412 | -0.03226533 | 0.173924635 | -0.024434962 | 0.221202617 |
| 14.26436 | -0.031340775 | 0.174632157 | -0.021590471 | 0.219105053 |
| 14.2746 | -0.030656301 | 0.175796473 | -0.018359616 | 0.216862547 |
| 14.28484 | -0.029644893 | 0.178036125 | -0.01469264 | 0.215043241 |
| 14.29508 | -0.029293611 | 0.179967761 | -0.011004835 | 0.213811411 |
| 14.30532 | -0.029021358 | 0.182293226 | -0.007137081 | 0.213506404 |
| 14.31556 | -0.029501891 | 0.184348417 | -0.003552135 | 0.213650334 |
| 14.3258 | -0.03095146 | 0.185582823 | -0.00053218 | 0.214857347 |
| 14.33604 | -0.033550783 | 0.186352076 | 0.001796862 | 0.216198102 |
| 14.34628 | -0.037434993 | 0.185779402 | 0.002997915 | 0.217905657 |
| 14.35652 | -0.042849404 | 0.184496387 | 0.003688388 | 0.219563337 |
| 14.36676 | -0.04930933 | 0.183059199 | 0.002760527 | 0.221270735 |
| 14.377 | -0.057024161 | 0.180834891 | 0.001336936 | 0.222427309 |

| | | | | |
|---|---|---|---|---|
| 11.98083 | -0.143839121 | 0.155711665 | -0.132851668 | 0.148332429 |
| 11.99107 | -0.141851894 | 0.153749172 | -0.131835021 | 0.147807017 |
| 12.00131 | -0.138896499 | 0.15299172 | -0.130426435 | 0.1467641 |
| 12.01155 | -0.134660774 | 0.152918792 | -0.128713013 | 0.145736253 |
| 12.02179 | -0.130502985 | 0.15399 | -0.126400586 | 0.143951607 |
| 12.03203 | -0.125876171 | 0.155633365 | -0.124047622 | 0.14177966 |
| 12.04227 | -0.121818471 | 0.158361338 | -0.121060356 | 0.138985271 |
| 12.05252 | -0.11828594 | 0.161495003 | -0.118032605 | 0.13573484 |
| 12.06275 | -0.115843261 | 0.165795334 | -0.114709323 | 0.132031019 |
| 12.073 | -0.114472798 | 0.170254589 | -0.111485666 | 0.128877644 |
| 12.08323 | -0.114105784 | 0.175972228 | -0.108126529 | 0.125877034 |
| 12.09348 | -0.114614906 | 0.18163318 | -0.104737512 | 0.123893461 |
| 12.10371 | -0.115799897 | 0.188493681 | -0.101611675 | 0.122992198 |
| 12.11395 | -0.117619211 | 0.194899085 | -0.098565543 | 0.12291133 |
| 12.1242 | -0.119537662 | 0.202297842 | -0.095961525 | 0.123640892 |
| 12.13443 | -0.121535829 | 0.209135382 | -0.094001361 | 0.124855724 |
| 12.14468 | -0.123723364 | 0.216752681 | -0.092837803 | 0.126566598 |
| 12.15491 | -0.125534522 | 0.224274787 | -0.092563116 | 0.127771913 |
| 12.16516 | -0.127151254 | 0.231396511 | -0.093913863 | 0.129011299 |
| 12.1754 | -0.128728248 | 0.238501505 | -0.096343532 | 0.129274286 |
| 12.18564 | -0.129993722 | 0.244483203 | -0.100097707 | 0.129645868 |
| 12.19588 | -0.131341866 | 0.249237133 | -0.104979373 | 0.128723338 |
| 12.20611 | -0.132773772 | 0.252044442 | -0.110604412 | 0.128214949 |
| 12.21636 | -0.133966088 | 0.252804901 | -0.116789819 | 0.127219386 |
| 12.22659 | -0.135868006 | 0.251554771 | -0.122456416 | 0.12684865 |
| 12.23684 | -0.137110929 | 0.24797441 | -0.127771975 | 0.127114879 |
| 12.24708 | -0.138859096 | 0.242803328 | -0.132045219 | 0.128353903 |
| 12.25732 | -0.139790779 | 0.23680463 | -0.134640682 | 0.131158954 |
| 12.26756 | -0.141008697 | 0.229696991 | -0.135747297 | 0.134969621 |
| 12.2778 | -0.14115722 | 0.223027562 | -0.135008769 | 0.140413131 |
| 12.28804 | -0.141177018 | 0.21650422 | -0.133059927 | 0.146593985 |
| 12.29828 | -0.140718047 | 0.211591284 | -0.129579355 | 0.15385287 |
| 12.30852 | -0.140057736 | 0.207329985 | -0.125527545 | 0.161572518 |
| 12.31876 | -0.139203962 | 0.204424174 | -0.121098541 | 0.169920002 |
| 12.329 | -0.138077299 | 0.202867946 | -0.116758294 | 0.179002551 |
| 12.33924 | -0.136914051 | 0.202543194 | -0.113042401 | 0.18796562 |
| 12.34948 | -0.135451636 | 0.202833117 | -0.110029085 | 0.197570962 |
| 12.35972 | -0.133263946 | 0.203862388 | -0.107780808 | 0.206786874 |
| 12.36996 | -0.130345175 | 0.205716026 | -0.106241189 | 0.215238069 |
| 12.3802 | -0.126223397 | 0.207533409 | -0.105499679 | 0.223147482 |
| 12.39044 | -0.120728139 | 0.209554022 | -0.10503357 | 0.229095079 |
| 12.40068 | -0.113958439 | 0.211818273 | -0.105498599 | 0.233931687 |
| 12.41092 | -0.105512615 | 0.213727813 | -0.105656265 | 0.237161747 |
| 12.42116 | -0.096168153 | 0.215770661 | -0.106988179 | 0.238867711 |
| 12.4314 | -0.085751587 | 0.217868473 | -0.108429201 | 0.239945534 |
| 12.44164 | -0.075536592 | 0.220263916 | -0.110234239 | 0.24005131 |
| 12.45188 | -0.065269179 | 0.222727355 | -0.112523606 | 0.240087784 |
| 12.46212 | -0.056481731 | 0.225420653 | -0.114611967 | 0.239695436 |
| 12.47236 | -0.048250467 | 0.227992722 | -0.116834645 | 0.239685611 |
| 12.4826 | -0.04231799 | 0.23013228 | -0.118336415 | 0.23926675 |
| 12.49284 | -0.037288046 | 0.231891957 | -0.11964743 | 0.238928486 |
| 12.50308 | -0.033915004 | 0.232420348 | -0.120217482 | 0.238116671 |
| 12.51332 | -0.031225986 | 0.231979368 | -0.120557221 | 0.236620453 |
| 12.52356 | -0.029424636 | 0.230571598 | -0.120624608 | 0.234493673 |
| 12.5338 | -0.027521492 | 0.227997793 | -0.120679692 | 0.231579733 |
| 12.54404 | -0.026032359 | 0.224605898 | -0.120181127 | 0.228341625 |
| 12.55428 | -0.024214546 | 0.220834315 | -0.120061093 | 0.224803817 |
| 12.56452 | -0.022368749 | 0.2164627 | -0.119394554 | 0.221504691 |
| 12.57476 | -0.020440111 | 0.212267645 | -0.118425193 | 0.218772738 |
| 12.585 | -0.019067826 | 0.207946464 | -0.117296635 | 0.217170609 |
| 12.59524 | -0.017251706 | 0.204460497 | -0.11614068 | 0.216568481 |
| 12.60548 | -0.016393287 | 0.201177196 | -0.114973427 | 0.217259321 |
| 12.61572 | -0.015230016 | 0.199180704 | -0.113946833 | 0.218526273 |
| 12.62596 | -0.014853872 | 0.197387235 | -0.113560179 | 0.220411361 |
| 12.6362 | -0.013987382 | 0.19713294 | -0.113670277 | 0.222139525 |
| 12.64644 | -0.013798948 | 0.197517254 | -0.114111017 | 0.223118062 |
| 12.65668 | -0.013124435 | 0.198905523 | -0.114394097 | 0.223347215 |
| 12.66692 | -0.012640904 | 0.201140156 | -0.114909332 | 0.221918057 |
| 12.67716 | -0.011849192 | 0.204173787 | -0.114466302 | 0.21979683 |
| 12.6874 | -0.01100896 | 0.207670519 | -0.113820854 | 0.216301431 |
| 12.69764 | -0.010401542 | 0.211465326 | -0.112196286 | 0.212421663 |
| 12.70788 | -0.009039179 | 0.215391652 | -0.110534702 | 0.207909181 |
| 12.71812 | -0.00828388 | 0.218628714 | -0.108332176 | 0.20347949 |
| 12.72836 | -0.00633013 | 0.221684924 | -0.10657079 | 0.199319931 |
| 12.7386 | -0.004762428 | 0.223072295 | -0.104779745 | 0.19559413 |
| 12.74884 | -0.002051209 | 0.224380402 | -0.103340533 | 0.192888665 |
| 12.75908 | 0.001336435 | 0.223617393 | -0.102496502 | 0.190905794 |
| 12.76932 | 0.005534472 | 0.222504786 | -0.101621693 | 0.19057326 |
| 12.77956 | 0.010719038 | 0.220367658 | -0.100985736 | 0.190988759 |
| 12.7898 | 0.016812641 | 0.217777911 | -0.100222165 | 0.192725419 |
| 12.80004 | 0.023728121 | 0.215208142 | -0.099702247 | 0.194528323 |
| 12.81028 | 0.031087013 | 0.212704339 | -0.098854293 | 0.196628177 |
| 12.82052 | 0.038954544 | 0.21076685 | -0.098158208 | 0.198240034 |
| 12.83076 | 0.047336972 | 0.209133013 | -0.097153693 | 0.198952933 |
| 12.841 | 0.055380913 | 0.207691623 | -0.095986461 | 0.198989105 |
| 12.85124 | 0.063369078 | 0.20612091 | -0.094633162 | 0.198185076 |
| 12.86148 | 0.071463336 | 0.204647992 | -0.092745434 | 0.197014046 |
| 12.87172 | 0.078447399 | 0.201984509 | -0.090222482 | 0.195479278 |
| 12.88196 | 0.084960417 | 0.19915499 | -0.087016506 | 0.194591112 |
| 12.8922 | 0.090759086 | 0.195342613 | -0.083041976 | 0.193696419 |
| 12.90244 | 0.095012999 | 0.191085231 | -0.078256583 | 0.193284602 |
| 12.91268 | 0.098682399 | 0.186514678 | -0.073020868 | 0.193088758 |
| 12.92292 | 0.100889106 | 0.181621984 | -0.067460094 | 0.192861013 |
| 12.93316 | 0.102534484 | 0.177767009 | -0.061886547 | 0.191925887 |
| 12.9434 | 0.10351182 | 0.173481504 | -0.056690209 | 0.190546185 |
| 12.95364 | 0.104361285 | 0.170361693 | -0.052431301 | 0.187802585 |
| 12.96388 | 0.105223798 | 0.167825035 | -0.04889842 | 0.1846531 |
| 12.97412 | 0.106766914 | 0.165558139 | -0.046716321 | 0.180271568 |
| 12.98436 | 0.109043614 | 0.163630969 | -0.045510943 | 0.175405912 |
| 12.9946 | 0.111608049 | 0.16170515 | -0.045147693 | 0.170447485 |
| 13.00484 | 0.115746509 | 0.159314171 | -0.045523721 | 0.165070743 |
| 13.01508 | 0.119823765 | 0.156385146 | -0.046007211 | 0.160323284 |
| 13.02532 | 0.124419185 | 0.153003346 | -0.046427151 | 0.155474346 |
| 13.03556 | 0.12898479 | 0.149226901 | -0.046597508 | 0.151552197 |
| 13.0458 | 0.133251521 | 0.145085805 | -0.04590932 | 0.147889052 |
| 13.05604 | 0.136762305 | 0.141203181 | -0.044896973 | 0.145020478 |
| 13.06628 | 0.139471756 | 0.138247008 | -0.04339906 | 0.142997606 |
| 13.07652 | 0.141110136 | 0.13588548 | -0.041022134 | 0.141974565 |
| 13.08676 | 0.141784076 | 0.134875193 | -0.038808082 | 0.141743957 |
| 13.097 | 0.14132968 | 0.13529003 | -0.035820677 | 0.142225114 |
| 13.10724 | 0.140229088 | 0.13695377 | -0.032684616 | 0.142996734 |
| 13.11748 | 0.138620721 | 0.140206052 | -0.029348214 | 0.144182512 |
| 13.12772 | 0.136737628 | 0.144220965 | -0.025282417 | 0.144871187 |
| 13.13796 | 0.13479516 | 0.149646592 | -0.021155068 | 0.145609571 |
| 13.1482 | 0.133319382 | 0.155383635 | -0.016714437 | 0.146232068 |
| 13.15844 | 0.132186102 | 0.160991893 | -0.0119099 | 0.146759487 |
| 13.16868 | 0.130970168 | 0.166864341 | -0.00724495 | 0.147429906 |
| 13.17892 | 0.130425834 | 0.171253984 | -0.002779448 | 0.14851607 |
| 13.18916 | 0.129848357 | 0.174991837 | 0.001241459 | 0.150246588 |
| 13.1994 | 0.128994196 | 0.177806685 | 0.004793134 | 0.152675036 |
| 13.20964 | 0.12772271 | 0.179914668 | 0.007479673 | 0.155834624 |
| 13.21988 | 0.12624857 | 0.1819195 | 0.009450243 | 0.159633033 |
| 13.23012 | 0.123939996 | 0.18431141 | 0.010928185 | 0.163858582 |
| 13.24036 | 0.121698594 | 0.18799219 | 0.011411097 | 0.168557128 |
| 13.2506 | 0.119030069 | 0.193073369 | 0.012066848 | 0.17324259 |
| 13.26084 | 0.116836086 | 0.200132265 | 0.01235739 | 0.178281268 |
| 13.27108 | 0.115203227 | 0.20870417 | 0.012930926 | 0.183418396 |
| 13.28132 | 0.114295482 | 0.218417382 | 0.013937467 | 0.188020667 |
| 13.29156 | 0.114317997 | 0.228866049 | 0.015819563 | 0.193445927 |
| 13.3018 | 0.115222651 | 0.2393329 | 0.017918749 | 0.198210821 |
| 13.31204 | 0.116926251 | 0.249053068 | 0.020676947 | 0.20291599 |
| 13.32228 | 0.11896219 | 0.257075489 | 0.024007282 | 0.207337388 |
| 13.33252 | 0.121365265 | 0.263188094 | 0.026702614 | 0.211567601 |
| 13.34276 | 0.123572607 | 0.266778607 | 0.029762429 | 0.215255502 |
| 13.353 | 0.125497655 | 0.267702447 | 0.03192753 | 0.218886139 |
| 13.36324 | 0.12736595 | 0.266442227 | 0.033805536 | 0.221959541 |
| 13.37348 | 0.12854807 | 0.263801373 | 0.035039023 | 0.22543351 |
| 13.38372 | 0.129820646 | 0.259782634 | 0.036166597 | 0.228360178 |
| 13.39396 | 0.130606886 | 0.255753683 | 0.037135559 | 0.231781534 |
| 13.4042 | 0.131076548 | 0.252206989 | 0.038361418 | 0.234660726 |
| 13.41444 | 0.131724946 | 0.24876912 | 0.039946592 | 0.237661452 |
| 13.42468 | 0.13184933 | 0.246764158 | 0.042420405 | 0.23983663 |
| 13.43492 | 0.131602527 | 0.244642404 | 0.045088414 | 0.241367494 |
| 13.44516 | 0.131113459 | 0.243241871 | 0.048975998 | 0.241759957 |
| 13.4554 | 0.129867173 | 0.242012196 | 0.053176377 | 0.240692224 |
| 13.46564 | 0.127989307 | 0.24074202 | 0.057740005 | 0.23889345 |
| 13.47588 | 0.125289572 | 0.239939999 | 0.062784322 | 0.235716473 |
| 13.48612 | 0.121460922 | 0.238604178 | 0.067885818 | 0.232054739 |
| 13.49636 | 0.116975187 | 0.238348251 | 0.073121055 | 0.227926106 |
| 13.5066 | 0.111339882 | 0.237451158 | 0.078116422 | 0.224249595 |
| 13.51684 | 0.104919433 | 0.237605673 | 0.082888275 | 0.220797805 |
| 13.52708 | 0.098270976 | 0.237565501 | 0.087432939 | 0.21820394 |
| 13.53732 | 0.091487791 | 0.237810367 | 0.091204485 | 0.216609685 |
| 13.54756 | 0.08490863 | 0.238635118 | 0.094616071 | 0.216206421 |
| 13.5578 | 0.079064197 | 0.239216334 | 0.097018125 | 0.216722835 |
| 13.56804 | 0.073982622 | 0.240090456 | 0.098586264 | 0.218248271 |
| 13.57828 | 0.070309821 | 0.240800823 | 0.099276712 | 0.220698518 |
| 13.58852 | 0.067127225 | 0.241237006 | 0.099262239 | 0.223497826 |
| 13.59876 | 0.065256266 | 0.240958498 | 0.098484828 | 0.22665089 |
| 13.609 | 0.063504854 | 0.240172425 | 0.097335378 | 0.229461162 |
| 13.61924 | 0.062314947 | 0.238588441 | 0.095863104 | 0.232207032 |
| 13.62948 | 0.060898617 | 0.236052229 | 0.094773898 | 0.234272715 |
| 13.63972 | 0.059227534 | 0.232418444 | 0.093255041 | 0.236088352 |
| 13.64996 | 0.057098283 | 0.22830007 | 0.092436739 | 0.236818023 |
| 13.6602 | 0.054387913 | 0.223639824 | 0.091572211 | 0.237457312 |
| 13.67044 | 0.051749382 | 0.218873924 | 0.091056066 | 0.237458323 |
| 13.68068 | 0.048536767 | 0.214634121 | 0.090548978 | 0.236938443 |
| 13.69092 | 0.046301304 | 0.210824777 | 0.090083544 | 0.236528991 |
| 13.70116 | 0.044357523 | 0.208178766 | 0.089727161 | 0.236173272 |
| 13.7114 | 0.043774329 | 0.206427655 | 0.089379201 | 0.235899921 |
| 13.72164 | 0.044042271 | 0.205472836 | 0.089078286 | 0.236204925 |
| 13.73188 | 0.046333304 | 0.204970357 | 0.088361835 | 0.236830645 |
| 13.74212 | 0.049019081 | 0.204016108 | 0.087983799 | 0.237833508 |
| 13.75236 | 0.052876387 | 0.202662547 | 0.086909931 | 0.239015929 |
| 13.7626 | 0.057497407 | 0.200061636 | 0.086111553 | 0.240785663 |
| 13.77284 | 0.061901444 | 0.196670187 | 0.084314215 | 0.242163137 |
| 13.78308 | 0.066504444 | 0.192431123 | 0.083081377 | 0.243764074 |
| 13.79332 | 0.070317722 | 0.187976838 | 0.08079013 | 0.245186654 |
| 13.80356 | 0.074045242 | 0.18337168 | 0.079378464 | 0.246028433 |
| 13.8138 | 0.076824292 | 0.179719019 | 0.07742489 | 0.24668135 |
| 13.82404 | 0.07894255 | 0.176784206 | 0.076425846 | 0.246577661 |
| 13.83428 | 0.080243957 | 0.174916916 | 0.07588467 | 0.24644293 |
| 13.84452 | 0.080839919 | 0.173982886 | 0.075784781 | 0.245958562 |
| 13.85476 | 0.08011856 | 0.174313169 | 0.076633048 | 0.24610409 |
| 13.865 | 0.078615541 | 0.17515845 | 0.077573143 | 0.246270399 |
| 13.87524 | 0.076012057 | 0.176345537 | 0.079004755 | 0.24728906 |
| 13.88548 | 0.072342636 | 0.177767761 | 0.080350689 | 0.24873043 |
| 13.89572 | 0.06775805 | 0.179086671 | 0.08168971 | 0.250669985 |
| 13.90596 | 0.06263061 | 0.17996412 | 0.082457836 | 0.252449216 |
| 13.9162 | 0.057014642 | 0.180617771 | 0.083266221 | 0.254506643 |
| 13.92644 | 0.05104347 | 0.1803501 | 0.083475782 | 0.25592299 |
| 13.93668 | 0.045441596 | 0.179762597 | 0.083780725 | 0.25752541 |
| 13.94692 | 0.040209287 | 0.178179484 | 0.084099495 | 0.258608314 |
| 13.95716 | 0.035662979 | 0.176411014 | 0.084065491 | 0.260072647 |
| 13.9674 | 0.031870333 | 0.173987057 | 0.084587093 | 0.262239049 |
| 13.97764 | 0.029799563 | 0.171455464 | 0.084936383 | 0.265030233 |
| 13.98788 | 0.028735046 | 0.169012789 | 0.085882183 | 0.268630085 |
| 13.99812 | 0.029482115 | 0.167179503 | 0.086251451 | 0.273474165 |
| 14.00836 | 0.031455434 | 0.16487351 | 0.086857133 | 0.278444479 |
| 14.0186 | 0.034953171 | 0.163052274 | 0.087681289 | 0.283600802 |
| 14.02884 | 0.039470614 | 0.160998078 | 0.087986501 | 0.288194454 |
| 14.03908 | 0.044518326 | 0.158233133 | 0.088717978 | 0.291797508 |
| 14.04932 | 0.049976124 | 0.154624895 | 0.089458449 | 0.293945446 |
| 14.05956 | 0.055502769 | 0.150127509 | 0.090206759 | 0.294893763 |
| 14.0698 | 0.060682629 | 0.144877566 | 0.09102565 | 0.29424952 |
| 14.08004 | 0.065310463 | 0.139419237 | 0.091775491 | 0.292666551 |
| 14.09028 | 0.06979314 | 0.133615886 | 0.091903596 | 0.290223557 |
| 14.10052 | 0.073704916 | 0.128746491 | 0.091554391 | 0.286772716 |
| 14.11076 | 0.077466599 | 0.124731857 | 0.089931137 | 0.283326662 |
| 14.121 | 0.08123474 | 0.121708869 | 0.087783508 | 0.279395543 |
| 14.13124 | 0.084885505 | 0.120265827 | 0.084022565 | 0.275411758 |
| 14.14148 | 0.088663336 | 0.119181017 | 0.080279242 | 0.27136257 |
| 14.15172 | 0.092100705 | 0.119249971 | 0.075616413 | 0.26717875 |
| 14.16196 | 0.095855133 | 0.119097337 | 0.071235216 | 0.26315544 |
| 14.1722 | 0.098696809 | 0.118826309 | 0.066884385 | 0.259231572 |
| 14.18244 | 0.101462362 | 0.117310502 | 0.063377883 | 0.254925906 |
| 14.19268 | 0.103227629 | 0.115230363 | 0.060537052 | 0.250871864 |
| 14.20292 | 0.10487764 | 0.111856536 | 0.057724809 | 0.246160027 |
| 14.21316 | 0.105265122 | 0.10861717 | 0.055974294 | 0.241317093 |
| 14.2234 | 0.105759923 | 0.105122525 | 0.053979821 | 0.235824089 |
| 14.23364 | 0.105291669 | 0.103260936 | 0.052255805 | 0.230184386 |
| 14.24388 | 0.10535356 | 0.102513565 | 0.050020821 | 0.223527028 |
| 14.25412 | 0.104999438 | 0.102881116 | 0.047865321 | 0.217349686 |
| 14.26436 | 0.105778602 | 0.104186331 | 0.045692328 | 0.210177653 |
| 14.2746 | 0.107179617 | 0.106070091 | 0.043395652 | 0.203933162 |
| 14.28484 | 0.110285756 | 0.107875656 | 0.041816897 | 0.196977838 |
| 14.29508 | 0.114808919 | 0.109443461 | 0.040675088 | 0.191125209 |
| 14.30532 | 0.12114367 | 0.110999417 | 0.040407684 | 0.185679958 |
| 14.31556 | 0.129209657 | 0.112291612 | 0.040932156 | 0.181057902 |
| 14.3258 | 0.1380582 | 0.113355804 | 0.04237871 | 0.177566786 |
| 14.33604 | 0.148251018 | 0.114445457 | 0.044656307 | 0.175079769 |
| 14.34628 | 0.158010092 | 0.115221597 | 0.047051743 | 0.173525791 |
| 14.35652 | 0.16733112 | 0.116500419 | 0.050388162 | 0.172538278 |
| 14.36676 | 0.175574517 | 0.117358719 | 0.053588242 | 0.172578088 |
| 14.377 | 0.181815091 | 0.11900914 | 0.057049164 | 0.172568975 |

| | | | | | | | | | | | | | | | | | | | |
|---|---|---|---|---|---|---|---|---|---|---|---|---|---|---|---|---|---|---|---|
| 14.38724 | 0.030767161 | 0.196435785 | 0.012455909 | 0.26197535 | 14.38724 | 0.031360301 | 0.195791369 | 0.078206318 | 0.165084882 | 14.38724 | -0.065145095 | 0.179699078 | -0.00105762 | 0.223895876 | 14.38724 | 0.186445657 | 0.120741106 | 0.060559053 | 0.17332805 |
| 14.39748 | 0.036545391 | 0.199525431 | 0.01830693 | 0.26012334 | 14.39748 | 0.031291569 | 0.19100996 | 0.08303527 | 0.17122596 | 14.39748 | -0.073421792 | 0.178615897 | -0.003865685 | 0.224950467 | 14.39748 | 0.189095293 | 0.12257603 | 0.064174092 | 0.174580725 |
| 14.40772 | 0.043691628 | 0.202138203 | 0.024716958 | 0.25956715 | 14.40772 | 0.031318917 | 0.185563906 | 0.086667347 | 0.178560631 | 14.40772 | -0.081958679 | 0.17821523 | -0.007100533 | 0.225729182 | 14.40772 | 0.189678531 | 0.124385286 | 0.06721202 | 0.176006357 |
| 14.41796 | 0.052528287 | 0.204342517 | 0.031791848 | 0.26016395 | 14.41796 | 0.03077819 | 0.180115816 | 0.08977278 | 0.185293159 | 14.41796 | -0.089104932 | 0.179036528 | -0.010605623 | 0.22666721 | 14.41796 | 0.18916005 | 0.126076684 | 0.070083217 | 0.178428949 |
| 14.4282 | 0.061630799 | 0.206265808 | 0.038445823 | 0.26178604 | 14.4282 | 0.029839599 | 0.174820982 | 0.092035145 | 0.190866043 | 14.4282 | -0.096315301 | 0.179571151 | -0.014718579 | 0.22705578 | 14.4282 | 0.187286184 | 0.126955067 | 0.0723672 | 0.181032479 |
| 14.43844 | 0.070632281 | 0.208531624 | 0.045705628 | 0.26416179 | 14.43844 | 0.02788186 | 0.170381365 | 0.093046056 | 0.195148118 | 14.43844 | -0.101867723 | 0.180234997 | -0.01892869 | 0.227352985 | 14.43844 | 0.184876588 | 0.127784305 | 0.073714577 | 0.183971187 |
| 14.44868 | 0.079047552 | 0.210398227 | 0.052576566 | 0.26720976 | 14.44868 | 0.025072794 | 0.166275194 | 0.093406853 | 0.197218988 | 14.44868 | -0.106940994 | 0.180570444 | -0.023967174 | 0.226991282 | 14.44868 | 0.181979402 | 0.127788578 | 0.073994082 | 0.187114007 |
| 14.45892 | 0.08648225 | 0.212521048 | 0.059445745 | 0.27055486 | 14.45892 | 0.021486965 | 0.164112256 | 0.093192343 | 0.197645149 | 14.45892 | -0.110611448 | 0.179735762 | -0.029721218 | 0.226386951 | 14.45892 | 0.179574096 | 0.127967775 | 0.073545929 | 0.189855298 |
| 14.46916 | 0.092593435 | 0.214724078 | 0.066177162 | 0.274426 | 14.46916 | 0.016688468 | 0.162948528 | 0.092778716 | 0.19680939 | 14.46916 | -0.113880714 | 0.178305728 | -0.036077973 | 0.224926642 | 14.46916 | 0.177166211 | 0.127773133 | 0.071913676 | 0.192750745 |
| 14.4794 | 0.096952667 | 0.216713696 | 0.072739299 | 0.2779085 | 14.4794 | 0.011357127 | 0.163310302 | 0.092152954 | 0.195107286 | 14.4794 | -0.116065651 | 0.176378359 | -0.043924043 | 0.222650342 | 14.4794 | 0.175577425 | 0.12741268 | 0.069683695 | 0.194601634 |
| 14.48964 | 0.100160965 | 0.218369803 | 0.078452757 | 0.28122637 | 14.48964 | 0.005538475 | 0.164493845 | 0.092681688 | 0.193102022 | 14.48964 | -0.11807671 | 0.17352598 | -0.052095441 | 0.219860675 | 14.48964 | 0.174443885 | 0.127445447 | 0.067038074 | 0.196759987 |
| 14.49988 | 0.102230848 | 0.219788307 | 0.083792799 | 0.28327786 | 14.49988 | -2.01811E-05 | 0.166714967 | 0.093783341 | 0.191089849 | 14.49988 | -0.119403891 | 0.171515029 | -0.061053399 | 0.216372776 | 14.49988 | 0.173895862 | 0.127235381 | 0.064436496 | 0.197633939 |
| 14.51012 | 0.102929552 | 0.220742643 | 0.087464456 | 0.28435814 | 14.51012 | -0.005309663 | 0.169172199 | 0.09624602 | 0.189122054 | 14.51012 | -0.120391936 | 0.169039544 | -0.070273695 | 0.212977663 | 14.51012 | 0.173574671 | 0.126425063 | 0.062405408 | 0.198098423 |
| 14.52036 | 0.102842839 | 0.221275928 | 0.089757225 | 0.28375579 | 14.52036 | -0.009120283 | 0.172204339 | 0.100070769 | 0.186780265 | 14.52036 | -0.121261244 | 0.167606086 | -0.078897372 | 0.209365903 | 14.52036 | 0.173388393 | 0.125860852 | 0.061120602 | 0.197309814 |
| 14.5306 | 0.102105228 | 0.221450514 | 0.090034602 | 0.28151015 | 14.5306 | -0.012270665 | 0.174399191 | 0.104850357 | 0.183737363 | 14.5306 | -0.121931204 | 0.166316685 | -0.086809129 | 0.206103139 | 14.5306 | 0.173265582 | 0.124449418 | 0.061201447 | 0.195555657 |
| 14.54084 | 0.100777825 | 0.221324105 | 0.088452439 | 0.27762957 | 14.54084 | -0.013288582 | 0.177377191 | 0.110968255 | 0.179539816 | 14.54084 | -0.122192731 | 0.16567505 | -0.09304111 | 0.203611992 | 14.54084 | 0.172674544 | 0.122744844 | 0.062667647 | 0.192398154 |
| 14.55108 | 0.098723984 | 0.220832749 | 0.084785003 | 0.27239748 | 14.55108 | -0.013696923 | 0.179629873 | 0.117073303 | 0.174861514 | 14.55108 | -0.122108664 | 0.16452225 | -0.097621102 | 0.201334377 | 14.55108 | 0.171610851 | 0.120550274 | 0.065727639 | 0.188036968 |
| 14.56132 | 0.096339439 | 0.220245593 | 0.079717532 | 0.26615875 | 14.56132 | -0.012953301 | 0.182092499 | 0.123886398 | 0.168222119 | 14.56132 | -0.121510233 | 0.163989552 | -0.099709844 | 0.199610436 | 14.56132 | 0.170052129 | 0.118669571 | 0.069787981 | 0.182771935 |
| 14.57156 | 0.09356074 | 0.219307998 | 0.073500711 | 0.25967828 | 14.57156 | -0.012215847 | 0.184348451 | 0.12954435 | 0.16212929 | 14.57156 | -0.120710209 | 0.162378278 | -0.099989777 | 0.197716946 | 14.57156 | 0.168021052 | 0.116722364 | 0.07510108 | 0.175906113 |
| 14.5818 | 0.090550413 | 0.218267347 | 0.06645068 | 0.25312562 | 14.5818 | -0.011972523 | 0.186652696 | 0.134352081 | 0.15464116 | 14.5818 | -0.119026499 | 0.160398407 | -0.097867438 | 0.196220383 | 14.5818 | 0.165381126 | 0.1152797 | 0.080719717 | 0.169372142 |
| 14.59204 | 0.087321817 | 0.216659831 | 0.059274853 | 0.24711755 | 14.59204 | -0.01263821 | 0.188295189 | 0.137551438 | 0.148838555 | 14.59204 | -0.11668755 | 0.157973374 | -0.094341438 | 0.194014445 | 14.59204 | 0.162570043 | 0.114751152 | 0.086423416 | 0.16225055 |
| 14.60228 | 0.084218933 | 0.21472362 | 0.052223807 | 0.24240632 | 14.60228 | -0.01410353 | 0.190161665 | 0.139013745 | 0.144005162 | 14.60228 | -0.113871562 | 0.154892867 | -0.089096587 | 0.191766957 | 14.60228 | 0.159585286 | 0.114746924 | 0.091972554 | 0.155787416 |
| 14.61252 | 0.081287065 | 0.212509966 | 0.045741468 | 0.23844988 | 14.61252 | -0.017253538 | 0.190826848 | 0.138532597 | 0.14124531 | 14.61252 | -0.110536903 | 0.151243598 | -0.083347583 | 0.18911714 | 14.61252 | 0.156583463 | 0.116027662 | 0.09669455 | 0.150326487 |
| 14.62276 | 0.07890371 | 0.20948064 | 0.039445608 | 0.23606636 | 14.62276 | -0.020452325 | 0.191004404 | 0.136726478 | 0.141363876 | 14.62276 | -0.10643699 | 0.147967274 | -0.076760722 | 0.186729798 | 14.62276 | 0.154059712 | 0.118111668 | 0.101031881 | 0.146392079 |
| 14.633 | 0.076368191 | 0.206343291 | 0.03427925 | 0.23421089 | 14.633 | -0.0242866 | 0.190071393 | 0.133417164 | 0.143579522 | 14.633 | -0.102249497 | 0.14430753 | -0.070206573 | 0.184378202 | 14.633 | 0.151698811 | 0.121320959 | 0.104329497 | 0.144056535 |
| 14.64324 | 0.07463938 | 0.202388132 | 0.029576088 | 0.23392068 | 14.64324 | -0.027433201 | 0.188454106 | 0.129901661 | 0.147856389 | 14.64324 | -0.097592164 | 0.141336757 | -0.063936914 | 0.182813008 | 14.64324 | 0.150438941 | 0.12575234 | 0.107190059 | 0.143326404 |
| 14.65348 | 0.07264037 | 0.198324781 | 0.025814394 | 0.23390071 | 14.65348 | -0.029611376 | 0.185922396 | 0.125626173 | 0.153109607 | 14.65348 | -0.092982755 | 0.138583339 | -0.058019686 | 0.182214073 | 14.65348 | 0.149539467 | 0.13066767 | 0.109221081 | 0.143488982 |
| 14.66372 | 0.070587474 | 0.193560283 | 0.022697165 | 0.2344984 | 14.66372 | -0.030732349 | 0.18232071 | 0.12179768 | 0.159164218 | 14.66372 | -0.08852216 | 0.136593958 | -0.052815 | 0.182195897 | 14.66372 | 0.149743343 | 0.136063121 | 0.110931796 | 0.144215811 |
| 14.67396 | 0.068292824 | 0.188863576 | 0.020458984 | 0.23630089 | 14.67396 | -0.030365255 | 0.178966592 | 0.118391276 | 0.165853687 | 14.67396 | -0.08439508 | 0.135390649 | -0.048441284 | 0.183397991 | 14.67396 | 0.150418899 | 0.141288512 | 0.111487661 | 0.145013825 |
| 14.6842 | 0.065554312 | 0.18349761 | 0.01895587 | 0.2381915 | 14.6842 | -0.028844503 | 0.17522547 | 0.115284384 | 0.172357028 | 14.6842 | -0.080854492 | 0.135004805 | -0.045034556 | 0.184642777 | 14.6842 | 0.152361786 | 0.146289799 | 0.111368963 | 0.14547017 |
| 14.69444 | 0.062228859 | 0.178623131 | 0.018387086 | 0.24114898 | 14.69444 | -0.026944753 | 0.171620014 | 0.112736226 | 0.179744582 | 14.69444 | -0.078479591 | 0.13582549 | -0.041851284 | 0.186736969 | 14.69444 | 0.15440494 | 0.150499548 | 0.110518196 | 0.145631577 |
| 14.70468 | 0.058934357 | 0.173440409 | 0.018320334 | 0.24372451 | 14.70468 | -0.024582689 | 0.169027994 | 0.110589771 | 0.187424825 | 14.70468 | -0.076674415 | 0.137142212 | -0.039408464 | 0.187977509 | 14.70468 | 0.157263891 | 0.154066115 | 0.107865223 | 0.145864033 |
| 14.71492 | 0.055397241 | 0.169024069 | 0.018985651 | 0.24737461 | 14.71492 | -0.022571528 | 0.166503094 | 0.109419781 | 0.194799324 | 14.71492 | -0.076080491 | 0.139432576 | -0.037516957 | 0.189607289 | 14.71492 | 0.160144743 | 0.157152558 | 0.104651822 | 0.145934369 |
| 14.72516 | 0.052148632 | 0.164628005 | 0.020335443 | 0.24995374 | 14.72516 | -0.021075996 | 0.164974165 | 0.108380681 | 0.202788552 | 14.72516 | -0.07624052 | 0.141732356 | -0.035239903 | 0.190521949 | 14.72516 | 0.163086974 | 0.159520017 | 0.09984422 | 0.146678278 |
| 14.7354 | 0.049490696 | 0.161242369 | 0.02165691 | 0.25264892 | 14.7354 | -0.020262468 | 0.163318945 | 0.108230656 | 0.209676967 | 14.7354 | -0.076943264 | 0.144706028 | -0.0333994 | 0.191624641 | 14.7354 | 0.165378744 | 0.161103558 | 0.094262322 | 0.147683787 |
| 14.74564 | 0.047341984 | 0.158065374 | 0.023776885 | 0.25470081 | 14.74564 | -0.020676808 | 0.161868574 | 0.108746154 | 0.216390676 | 14.74564 | -0.077605744 | 0.146930812 | -0.031408287 | 0.192597502 | 14.74564 | 0.167327103 | 0.162060664 | 0.087777146 | 0.149519072 |
| 14.75588 | 0.045641826 | 0.155992584 | 0.024929035 | 0.25690933 | 14.75588 | -0.021301051 | 0.159991504 | 0.109659924 | 0.221240625 | 14.75588 | -0.078172174 | 0.149528934 | -0.029476011 | 0.193917501 | 14.75588 | 0.168290736 | 0.162006164 | 0.080879376 | 0.15191869 |
| 14.76612 | 0.044749088 | 0.154983849 | 0.026945244 | 0.25864239 | 14.76612 | -0.022447211 | 0.157900176 | 0.111331433 | 0.225411187 | 14.76612 | -0.077613016 | 0.151283899 | -0.02789569 | 0.196057918 | 14.76612 | 0.16838989 | 0.160985254 | 0.074496694 | 0.154918128 |
| 14.77636 | 0.043709848 | 0.154721197 | 0.027693416 | 0.26112619 | 14.77636 | -0.023466514 | 0.155185965 | 0.11290289 | 0.227487327 | 14.77636 | -0.076337903 | 0.152598649 | -0.026681495 | 0.198409368 | 14.77636 | 0.167905436 | 0.158962478 | 0.068307558 | 0.158416696 |
| 14.7866 | 0.043223232 | 0.156331912 | 0.028362035 | 0.26372714 | 14.7866 | -0.024288577 | 0.152974453 | 0.114557818 | 0.227729614 | 14.7866 | -0.073673795 | 0.153196518 | -0.026123471 | 0.201378755 | 14.7866 | 0.166862729 | 0.156288491 | 0.063525185 | 0.162375202 |
| 14.79684 | 0.042288642 | 0.158862257 | 0.028545336 | 0.26719336 | 14.79684 | -0.023646424 | 0.150403833 | 0.115717459 | 0.22632162 | 14.79684 | -0.06986045 | 0.15386307 | -0.026050412 | 0.204030988 | 14.79684 | 0.16508957 | 0.152548548 | 0.060244266 | 0.166427901 |
| 14.80708 | 0.04176711 | 0.16295566 | 0.028037978 | 0.2709913 | 14.80708 | -0.022024377 | 0.149474209 | 0.117222067 | 0.223781098 | 14.80708 | -0.065121836 | 0.153643091 | -0.027096319 | 0.206374894 | 14.80708 | 0.163872131 | 0.148471501 | 0.058623421 | 0.170309854 |
| 14.81732 | 0.041530626 | 0.168510178 | 0.02743006 | 0.27496361 | 14.81732 | -0.018311826 | 0.148446018 | 0.117920344 | 0.219511151 | 14.81732 | -0.05969073 | 0.153731794 | -0.028839181 | 0.207616911 | 14.81732 | 0.162243126 | 0.143922518 | 0.058814997 | 0.17413248 |
| 14.82756 | 0.041043384 | 0.174464163 | 0.026444503 | 0.27950874 | 14.82756 | -0.01359801 | 0.149295845 | 0.119054047 | 0.214860544 | 14.82756 | -0.053996465 | 0.153950386 | -0.031211205 | 0.207747293 | 14.82756 | 0.161083146 | 0.139107792 | 0.060316675 | 0.177809518 |
| 14.8378 | 0.041922176 | 0.181159693 | 0.026108054 | 0.28335327 | 14.8378 | -0.00713405 | 0.150414802 | 0.120858805 | 0.209957641 | 14.8378 | -0.048690955 | 0.154838448 | -0.034168769 | 0.20726235 | 14.8378 | 0.160043274 | 0.134378623 | 0.063060242 | 0.181239724 |
| 14.84804 | 0.042409908 | 0.187371345 | 0.025562204 | 0.28608039 | 14.84804 | -0.00023044 | 0.15253949 | 0.123182791 | 0.205377269 | 14.84804 | -0.043447276 | 0.156046643 | -0.037558159 | 0.205587228 | 14.84804 | 0.158589413 | 0.129474527 | 0.066050989 | 0.184844512 |
| 14.85828 | 0.044957531 | 0.192026602 | 0.026404016 | 0.28790134 | 14.85828 | 0.007242944 | 0.154873783 | 0.127239468 | 0.201193311 | 14.85828 | -0.039200469 | 0.158140431 | -0.041107537 | 0.203677354 | 14.85828 | 0.157297518 | 0.125118332 | 0.068956567 | 0.188339537 |
| 14.86852 | 0.046869744 | 0.195664229 | 0.028228751 | 0.28780384 | 14.86852 | 0.013570211 | 0.157336548 | 0.132383355 | 0.197474187 | 14.86852 | -0.035645629 | 0.161005774 | -0.044225328 | 0.201806676 | 14.86852 | 0.15517672 | 0.121760041 | 0.071641784 | 0.191986223 |
| 14.87876 | 0.050238017 | 0.196354112 | 0.031793296 | 0.28539094 | 14.87876 | 0.01953197 | 0.159711567 | 0.139288329 | 0.194629872 | 14.87876 | -0.032552956 | 0.16455727 | -0.047962146 | 0.200077113 | 14.87876 | 0.152301519 | 0.11917082 | 0.073172345 | 0.195394982 |
| 14.889 | 0.05319803 | 0.195934907 | 0.036266335 | 0.28096425 | 14.889 | 0.023921867 | 0.161307184 | 0.146919411 | 0.19185202 | 14.889 | -0.030386455 | 0.168082671 | -0.050631066 | 0.199105266 | 14.889 | 0.149095719 | 0.117799354 | 0.074430004 | 0.199180269 |
| 14.89924 | 0.05607444 | 0.192461682 | 0.042623422 | 0.27512994 | 14.89924 | 0.027106128 | 0.162729825 | 0.155814315 | 0.189711237 | 14.89924 | -0.028499384 | 0.17241645 | -0.053926055 | 0.198129903 | 14.89924 | 0.145038007 | 0.117898108 | 0.074525231 | 0.202767558 |
| 14.90948 | 0.058644524 | 0.188417046 | 0.049673135 | 0.26746749 | 14.90948 | 0.029238733 | 0.163187953 | 0.164683454 | 0.187096176 | 14.90948 | -0.026943613 | 0.176209995 | -0.056347353 | 0.198510656 | 14.90948 | 0.140704169 | 0.119481068 | 0.074641696 | 0.206908545 |
| 14.91972 | 0.059778331 | 0.183202097 | 0.057269007 | 0.25972669 | 14.91972 | 0.030660681 | 0.163365986 | 0.173908256 | 0.185147491 | 14.91972 | -0.025958094 | 0.179931114 | -0.058851923 | 0.199145381 | 14.91972 | 0.135784704 | 0.122040602 | 0.074373003 | 0.211287652 |
| 14.92996 | 0.060156397 | 0.177574401 | 0.064935265 | 0.25177629 | 14.92996 | 0.031085001 | 0.162569052 | 0.182293577 | 0.18246797 | 14.92996 | -0.025418483 | 0.183145988 | -0.061320367 | 0.200524665 | 14.92996 | 0.131403845 | 0.126449161 | 0.074002575 | 0.216445079 |
| 14.9402 | 0.058993777 | 0.172146835 | 0.071902176 | 0.24499041 | 14.9402 | 0.030976031 | 0.162187746 | 0.190570998 | 0.179727353 | 14.9402 | -0.025404993 | 0.185844251 | -0.063211432 | 0.202170079 | 14.9402 | 0.127252343 | 0.131233053 | 0.074033076 | 0.221839997 |
| 14.95044 | 0.056726836 | 0.167214083 | 0.077932583 | 0.23855868 | 14.95044 | 0.03026017 | 0.160992605 | 0.197682094 | 0.176922172 | 14.95044 | -0.026406421 | 0.188765045 | -0.064690164 | 0.204191531 | 14.95044 | 0.123302963 | 0.137030037 | 0.074132173 | 0.228181773 |
| 14.96068 | 0.053744975 | 0.163208646 | 0.082960992 | 0.23422308 | 14.96068 | 0.028766612 | 0.161007441 | 0.204351278 | 0.173307508 | 14.96068 | -0.028125122 | 0.190935089 | -0.065705392 | 0.206101371 | 14.96068 | 0.1202518 | 0.142659242 | 0.074143685 | 0.234974655 |
| 14.97092 | 0.049667256 | 0.159940094 | 0.086762112 | 0.23029397 | 14.97092 | 0.026614459 | 0.161994768 | 0.210563023 | 0.169572377 | 14.97092 | -0.030833672 | 0.193207134 | -0.065895651 | 0.207868084 | 14.97092 | 0.118088843 | 0.148127321 | 0.074555515 | 0.24212226 |
| 14.98116 | 0.045378096 | 0.158211891 | 0.089422848 | 0.22752409 | 14.98116 | 0.023710006 | 0.163503586 | 0.215709879 | 0.165428205 | 14.98116 | -0.034152514 | 0.195370208 | -0.065419317 | 0.209225369 | 14.98116 | 0.11637956 | 0.152517343 | 0.074996588 | 0.249386149 |
| 14.9914 | 0.041356385 | 0.157787343 | 0.091736791 | 0.22542814 | 14.9914 | 0.020376341 | 0.167137708 | 0.221100976 | 0.160358464 | 14.9914 | -0.037873765 | 0.196738375 | -0.06405302 | 0.210138986 | 14.9914 | 0.115455865 | 0.155999333 | 0.075622877 | 0.257037528 |
| 15.00164 | 0.036677099 | 0.157988676 | 0.092418339 | 0.22340957 | 15.00164 | 0.016789077 | 0.170904415 | 0.22514929 | 0.155316054 | 15.00164 | -0.041496074 | 0.197906071 | -0.062133691 | 0.210705824 | 15.00164 | 0.115134577 | 0.157840109 | 0.076937549 | 0.263930383 |
| 15.01188 | 0.032798637 | 0.159498491 | 0.093453225 | 0.22232789 | 15.01188 | 0.013300659 | 0.175416456 | 0.22948983 | 0.149834875 | 15.01188 | -0.044791293 | 0.197820629 | -0.059257725 | 0.210737453 | 15.01188 | 0.115069641 | 0.1584883 | 0.078953592 | 0.271233602 |
| 15.02212 | 0.028684116 | 0.161292717 | 0.0930065 | 0.22068964 | 15.02212 | 0.010216247 | 0.178615924 | 0.232429219 | 0.14508082 | 15.02212 | -0.047311695 | 0.196942007 | -0.056165625 | 0.210948843 | 15.02212 | 0.115427139 | 0.157376959 | 0.081722665 | 0.277072677 |
| 15.03236 | 0.024696841 | 0.163173055 | 0.092535545 | 0.22001748 | 15.03236 | 0.007127887 | 0.181379507 | 0.235071458 | 0.14054907 | 15.03236 | -0.048873022 | 0.194848677 | -0.051819414 | 0.211009473 | 15.03236 | 0.115948484 | 0.155595576 | 0.085890037 | 0.282766785 |
| 15.0426 | 0.021112305 | 0.165112141 | 0.091040195 | 0.21966437 | 15.0426 | 0.004654492 | 0.181544799 | 0.23721088 | 0.138027831 | 15.0426 | -0.049746593 | 0.192035154 | -0.047256882 | 0.211410144 | 15.0426 | 0.116877858 | 0.15260254 | 0.090980772 | 0.286772275 |
| 15.05284 | 0.017290746 | 0.166866706 | 0.08941225 | 0.22057958 | 15.05284 | 0.00245387 | 0.180946618 | 0.23801255 | 0.136860616 | 15.05284 | -0.049398257 | 0.188155565 | -0.041745001 | 0.212369226 | 15.05284 | 0.117606246 | 0.149853229 | 0.097445754 | 0.289608473 |
| 15.06308 | 0.014037931 | 0.168708399 | 0.087184531 | 0.22158216 | 15.06308 | 0.000481686 | 0.177766268 | 0.238740652 | 0.138479376 | 15.06308 | -0.048999918 | 0.184395016 | -0.03548851 | 0.212988142 | 15.06308 | 0.119063229 | 0.14693426 | 0.10456075 | 0.290582682 |
| 15.07332 | 0.01100345 | 0.171108227 | 0.084748371 | 0.22478008 | 15.07332 | -0.001301409 | 0.173772458 | 0.238603318 | 0.141412702 | 15.07332 | -0.047768969 | 0.180477832 | -0.028892383 | 0.214168107 | 15.07332 | 0.119906931 | 0.144599755 | 0.112455245 | 0.290431163 |
| 15.08356 | 0.008710137 | 0.173958522 | 0.082535406 | 0.22789093 | 15.08356 | -0.002937083 | 0.168506555 | 0.237929561 | 0.147077423 | 15.08356 | -0.046595228 | 0.177617879 | -0.021893324 | 0.214513612 | 15.08356 | 0.121557652 | 0.14251632 | 0.12070732 | 0.288409155 |
| 15.0938 | 0.006830117 | 0.178389543 | 0.079864319 | 0.23241355 | 15.0938 | -0.004440935 | 0.162222484 | 0.236637541 | 0.15331336 | 15.0938 | -0.044978399 | 0.175059839 | -0.014850256 | 0.214339849 | 15.0938 | 0.122370671 | 0.140941632 | 0.129118991 | 0.285678571 |
| 15.10404 | 0.006108425 | 0.183304035 | 0.077787608 | 0.23690595 | 15.10404 | -0.005742328 | 0.156239597 | 0.23492797 | 0.160683565 | 15.10404 | -0.043177132 | 0.174416286 | -0.008228302 | 0.212431654 | 15.10404 | 0.123157604 | 0.139377434 | 0.13708135 | 0.282436272 |
| 15.11428 | 0.00588152 | 0.189565805 | 0.075233174 | 0.24185477 | 15.11428 | -0.006980556 | 0.149498368 | 0.23250237 | 0.166934439 | 15.11428 | -0.040698591 | 0.174504699 | -0.00210899 | 0.209585757 | 15.11428 | 0.123431969 | 0.137397328 | 0.144642181 | 0.279459271 |
| 15.12452 | 0.007010622 | 0.196101939 | 0.073118532 | 0.24612537 | 15.12452 | -0.00863299 | 0.143593276 | 0.228868006 | 0.173515183 | 15.12452 | -0.037597023 | 0.176191435 | 0.002644239 | 0.205176785 | 15.12452 | 0.123267043 | 0.135821821 | 0.151284957 | 0.276848329 |
| 15.13476 | 0.008812392 | 0.201769465 | 0.070209154 | 0.24907815 | 15.13476 | -0.010100499 | 0.137190065 | 0.224679449 | 0.177376149 | 15.13476 | -0.033572484 | 0.178161727 | 0.007312216 | 0.199576231 | 15.13476 | 0.122491458 | 0.133714729 | 0.156525351 | 0.275156699 |
| 15.145 | 0.011168972 | 0.206834401 | 0.067204807 | 0.25166736 | 15.145 | -0.011745323 | 0.13217961 | 0.219271995 | 0.180098811 | 15.145 | -0.028871509 | 0.180687707 | 0.010271725 | 0.193241758 | 15.145 | 0.120699279 | 0.132029784 | 0.160427634 | 0.274629834 |
| 15.15524 | 0.014698944 | 0.209699892 | 0.063720993 | 0.25179101 | 15.15524 | -0.013707091 | 0.126729442 | 0.213336747 | 0.18117712 | 15.15524 | -0.022748188 | 0.1829509 | 0.01322861 | 0.187250489 | 15.15524 | 0.118840832 | 0.131013531 | 0.16235388 | 0.275122726 |
| 15.16548 | 0.017510227 | 0.210516167 | 0.059900077 | 0.25137961 | 15.16548 | -0.015157354 | 0.121825074 | 0.20592151 | 0.18015955 | 15.16548 | -0.016741921 | 0.184375878 | 0.015466884 | 0.181215273 | 15.16548 | 0.116101751 | 0.130751105 | 0.162732203 | 0.276228522 |
| 15.17572 | 0.021155431 | 0.20804304 | 0.055855249 | 0.24943328 | 15.17572 | -0.016590744 | 0.117889779 | 0.198611039 | 0.178160227 | 15.17572 | -0.009571032 | 0.184544333 | 0.017856018 | 0.176329599 | 15.17572 | 0.112987382 | 0.131902626 | 0.161298148 | 0.278342114 |
| 15.18596 | 0.023460191 | 0.204049493 | 0.051439919 | 0.24663395 | 15.18596 | -0.017873676 | 0.114474759 | 0.190595296 | 0.174973863 | 15.18596 | -0.002832477 | 0.183795721 | 0.020405804 | 0.172216688 | 15.18596 | 0.109911078 | 0.134448751 | 0.158547864 | 0.280384579 |
| 15.1962 | 0.025443751 | 0.197609548 | 0.047539112 | 0.24367305 | 15.1962 | -0.018645041 | 0.112283511 | 0.182941723 | 0.171615038 | 15.1962 | 0.004367763 | 0.181617034 | 0.023289028 | 0.169040942 | 15.1962 | 0.106399653 | 0.137580072 | 0.155200748 | 0.28281157 |
| 15.20644 | 0.026242395 | 0.189910611 | 0.043646512 | 0.24075105 | 15.20644 | -0.019738927 | 0.111622119 | 0.174960492 | 0.168109248 | 15.20644 | 0.010477473 | 0.178730752 | 0.026566225 | 0.166520611 | 15.20644 | 0.103395981 | 0.142114809 | 0.15135214 | 0.285007985 |
| 15.21668 | 0.02571632 | 0.181303036 | 0.040098677 | 0.23845615 | 15.21668 | -0.021558289 | 0.111438296 | 0.167559063 | 0.165492177 | 15.21668 | 0.016724414 | 0.174971131 | 0.029929049 | 0.16413653 | 15.21668 | 0.100725702 | 0.146987174 | 0.147905855 | 0.287060662 |
| 15.22692 | 0.024076903 | 0.172930968 | 0.037279865 | 0.23645013 | 15.22692 | -0.023984438 | 0.111925439 | 0.160160514 | 0.163549871 | 15.22692 | 0.021534891 | 0.171171153 | 0.033575265 | 0.16211518 | 15.22692 | 0.098603766 | 0.152313648 | 0.144684637 | 0.289257728 |
| 15.23716 | 0.021199692 | 0.164812483 | 0.034391936 | 0.23553366 | 15.23716 | -0.027556958 | 0.113080308 | 0.153065683 | 0.163385779 | 15.23716 | 0.025680297 | 0.167041971 | 0.036710498 | 0.159775762 | 15.23716 | 0.096881064 | 0.157922918 | 0.142202862 | 0.290632516 |
| 15.2474 | 0.017455776 | 0.157219461 | 0.032461789 | 0.23480596 | 15.2474 | -0.033076286 | 0.113561678 | 0.145761472 | 0.164152019 | 15.2474 | 0.028457001 | 0.163192886 | 0.039701152 | 0.15752213 | 15.2474 | 0.095482693 | 0.164231083 | 0.140130967 | 0.29219516 |
| 15.25764 | 0.012248584 | 0.150668021 | 0.030813563 | 0.23479635 | 15.25764 | -0.039423715 | 0.114409448 | 0.138522758 | 0.166570939 | 15.25764 | 0.030346779 | 0.159812849 | 0.042161051 | 0.155031273 | 15.25764 | 0.094470169 | 0.170083764 | 0.138779182 | 0.293553389 |
| 15.26788 | 0.006748253 | 0.14505453 | 0.029867967 | 0.23426755 | 15.26788 | -0.047332334 | 0.115306602 | 0.131617742 | 0.170509615 | 15.26788 | 0.030877787 | 0.157289087 | 0.044270526 | 0.152996468 | 15.26788 | 0.09321209 | 0.176658484 | 0.136993169 | 0.293995806 |
| 15.27812 | 0.000143262 | 0.140064436 | 0.029409495 | 0.23409659 | 15.27812 | -0.055669559 | 0.116340587 | 0.124498039 | 0.175284623 | 15.27812 | 0.030466396 | 0.155790987 | 0.046307838 | 0.150551041 | 15.27812 | 0.091850146 | 0.182329199 | 0.13561516 | 0.294549859 |
| 15.28836 | -0.006678353 | 0.136107836 | 0.029928592 | 0.23343111 | 15.28836 | -0.064713683 | 0.118350209 | 0.117511532 | 0.18044905 | 15.28836 | 0.029220531 | 0.156133543 | 0.048210275 | 0.148669434 | 15.28836 | 0.090340231 | 0.188127841 | 0.134096548 | 0.2939776 |
| 15.2986 | -0.014685581 | 0.133492956 | 0.03109242 | 0.23280094 | 15.2986 | -0.072755155 | 0.120974686 | 0.11110602 | 0.186575122 | 15.2986 | 0.027257726 | 0.157629849 | 0.050654538 | 0.146754455 | 15.2986 | 0.088431736 | 0.192946844 | 0.132543831 | 0.292964051 |
| 15.30884 | -0.02235464 | 0.131954281 | 0.032937842 | 0.23162133 | 15.30884 | -0.080065307 | 0.12373864 | 0.104800858 | 0.192849849 | 15.30884 | 0.024802312 | 0.160938101 | 0.053061311 | 0.144939786 | 15.30884 | 0.08724448 | 0.196922686 | 0.131149481 | 0.290847243 |
| 15.31908 | -0.030141571 | 0.132553426 | 0.035696412 | 0.23091566 | 15.31908 | -0.085298956 | 0.126373527 | 0.099246809 | 0.199510903 | 15.31908 | 0.022042179 | 0.165139947 | 0.05638973 | 0.142968503 | 15.31908 | 0.086072337 | 0.199772129 | 0.130439455 | 0.288363046 |
| 15.32932 | -0.038304523 | 0.134905679 | 0.038473661 | 0.22999992 | 15.32932 | -0.088198205 | 0.128193141 | 0.093742185 | 0.206027241 | 15.32932 | 0.018882796 | 0.170192501 | 0.059612496 | 0.140956801 | 15.32932 | 0.085526231 | 0.201956371 | 0.130070989 | 0.284745368 |
| 15.33956 | -0.045796899 | 0.139562887 | 0.042276422 | 0.2297597 | 15.33956 | -0.088208321 | 0.12882459 | 0.089193424 | 0.213304279 | 15.33956 | 0.015678686 | 0.175258055 | 0.063522 | 0.138946635 | 15.33956 | 0.085668144 | 0.202608086 | 0.130775414 | 0.281079383 |
| 15.3498 | -0.053583687 | 0.145231409 | 0.045776619 | 0.22952836 | 15.3498 | -0.085348828 | 0.128492636 | 0.084945459 | 0.220189086 | 15.3498 | 0.01247772 | 0.179935473 | 0.067004146 | 0.136423427 | 15.3498 | 0.086177618 | 0.202775737 | 0.132508728 | 0.277163523 |
| 15.36004 | -0.060493418 | 0.152698152 | 0.050013291 | 0.22977166 | 15.36004 | -0.080030409 | 0.126725923 | 0.081309195 | 0.227305483 | 15.36004 | 0.00930802 | 0.183760892 | 0.070675429 | 0.134180402 | 15.36004 | 0.086883173 | 0.201927009 | 0.135156501 | 0.273003519 |
| 15.37028 | -0.067885209 | 0.159403645 | 0.054471061 | 0.22961281 | 15.37028 | -0.072421573 | 0.124408588 | 0.078814193 | 0.233847422 | 15.37028 | 0.006654196 | 0.186379297 | 0.073361411 | 0.131461265 | 15.37028 | 0.08762928 | 0.200748035 | 0.138993543 | 0.269779554 |
| 15.38052 | -0.075166537 | 0.165729211 | 0.058768223 | 0.22937667 | 15.38052 | -0.06352401 | 0.122140885 | 0.07672471 | 0.240025837 | 15.38052 | 0.004646507 | 0.187591131 | 0.076155871 | 0.129078388 | 15.38052 | 0.087758591 | 0.199583613 | 0.143246357 | 0.266552602 |
| 15.39076 | -0.082305003 | 0.170365871 | 0.063511078 | 0.22836322 | 15.39076 | -0.054392674 | 0.11946892 | 0.075937366 | 0.245339326 | 15.39076 | 0.003032556 | 0.18729572 | 0.07763556 | 0.126459293 | 15.39076 | 0.087605265 | 0.198724919 | 0.148597557 | 0.263672488 |
| 15.401 | -0.089033207 | 0.172965785 | 0.067956213 | 0.22681862 | 15.401 | -0.045261917 | 0.11797616 | 0.076419698 | 0.249491951 | 15.401 | 0.003102081 | 0.185118823 | 0.078672971 | 0.124460672 | 15.401 | 0.086896381 | 0.198019307 | 0.153784539 | 0.261724563 |
| 15.41124 | -0.095532574 | 0.173646223 | 0.071879771 | 0.22520955 | 15.41124 | -0.036755274 | 0.117217553 | 0.077757787 | 0.252224714 | 15.41124 | 0.003589903 | 0.181926896 | 0.078324062 | 0.121944512 | 15.41124 | 0.085826315 | 0.198302281 | 0.159257113 | 0.259843165 |
| 15.42148 | -0.10095762 | 0.172026514 | 0.075003449 | 0.22316099 | 15.42148 | -0.029529728 | 0.118251407 | 0.080507964 | 0.253841255 | 15.42148 | 0.005539019 | 0.177942329 | 0.07751559 | 0.119985931 | 15.42148 | 0.084337001 | 0.199609971 | 0.164194377 | 0.258622462 |
| 15.43172 | -0.105272347 | 0.16953943 | 0.077297742 | 0.22166374 | 15.43172 | -0.023044551 | 0.1196919 | 0.084631603 | 0.253090779 | 15.43172 | 0.007997585 | 0.173479485 | 0.074910603 | 0.117552273 | 15.43172 | 0.082376231 | 0.200883864 | 0.16878549 | 0.257834772 |
| 15.44196 | -0.108490905 | 0.165548916 | 0.07791001 | 0.22039478 | 15.44196 | -0.017664457 | 0.12263755 | 0.088978708 | 0.251139531 | 15.44196 | 0.011214693 | 0.169511587 | 0.071724842 | 0.115144261 | 15.44196 | 0.080164953 | 0.203393675 | 0.172655318 | 0.257716699 |
| 15.4522 | -0.110476573 | 0.161086048 | 0.077307277 | 0.21963167 | 15.4522 | -0.012655402 | 0.125988046 | 0.094293829 | 0.247446462 | 15.4522 | 0.014763459 | 0.16605205 | 0.067435635 | 0.11262923 | 15.4522 | 0.077048168 | 0.205459161 | 0.175875361 | 0.258219701 |
| 15.46244 | -0.111093109 | 0.157012216 | 0.076151378 | 0.21869198 | 15.46244 | -0.008346481 | 0.129593497 | 0.099712887 | 0.242281673 | 15.46244 | 0.018380366 | 0.163936984 | 0.06240241 | 0.109844337 | 15.46244 | 0.073599475 | 0.20789081 | 0.17857053 | 0.259486112 |
| 15.47268 | -0.111599214 | 0.152732928 | 0.07302522 | 0.21789907 | 15.47268 | -0.003636396 | 0.132210558 | 0.104715881 | 0.236041458 | 15.47268 | 0.021704621 | 0.162569254 | 0.057061813 | 0.107216944 | 15.47268 | 0.068619555 | 0.209847787 | 0.180606249 | 0.261542054 |
| 15.48292 | -0.111209831 | 0.149021514 | 0.070644357 | 0.21742818 | 15.48292 | 0.000978707 | 0.135002501 | 0.109015993 | 0.228399957 | 15.48292 | 0.024894289 | 0.162786271 | 0.051945042 | 0.104446307 | 15.48292 | 0.063919608 | 0.211323327 | 0.182465278 | 0.264455536 |
| 15.49317 | -0.110780664 | 0.145621402 | 0.067123089 | 0.21707338 | 15.49317 | 0.006458784 | 0.136572699 | 0.112183931 | 0.220165157 | 15.49317 | 0.027323211 | 0.163577286 | 0.047655787 | 0.102227696 | 15.49317 | 0.05766909 | 0.212797626 | 0.184305306 | 0.268031101 |
| 15.5034 | -0.110102911 | 0.143256765 | 0.065108781 | 0.21659764 | 15.5034 | 0.012602961 | 0.138037943 | 0.114795163 | 0.21128238 | 15.5034 | 0.029430144 | 0.165260819 | 0.043910016 | 0.100097581 | 15.5034 | 0.052077529 | 0.214209233 | 0.185892791 | 0.27238122 |
| 15.51365 | -0.109138723 | 0.140224707 | 0.063048347 | 0.21624955 | 15.51365 | 0.019613643 | 0.138943316 | 0.116125093 | 0.201791745 | 15.51365 | 0.030973269 | 0.167582341 | 0.042019503 | 0.098442819 | 15.51365 | 0.046299473 | 0.215343073 | 0.187660338 | 0.276535152 |
| 15.52388 | -0.107225705 | 0.138294058 | 0.063167687 | 0.21603765 | 15.52388 | 0.027461634 | 0.140743147 | 0.117410635 | 0.192834728 | 15.52388 | 0.031796361 | 0.170588642 | 0.041303377 | 0.097642243 | 15.52388 | 0.041074627 | 0.217103757 | 0.189802065 | 0.280901108 |
| 15.53412 | -0.104431527 | 0.135977295 | 0.063581845 | 0.21489893 | 15.53412 | 0.03613335 | 0.14254858 | 0.118777446 | 0.184638987 | 15.53412 | 0.032505027 | 0.173707778 | 0.042047264 | 0.096618437 | 15.53412 | 0.036321411 | 0.218770253 | 0.191586555 | 0.28449167 |
| 15.54436 | -0.100023453 | 0.133066063 | 0.066449961 | 0.21340625 | 15.54436 | 0.045044947 | 0.144944882 | 0.120310023 | 0.177600965 | 15.54436 | 0.032745209 | 0.177657726 | 0.044011233 | 0.096259718 | 15.54436 | 0.032101053 | 0.221109021 | 0.193534252 | 0.287651296 |
| 15.5546 | -0.094105357 | 0.129595845 | 0.069816664 | 0.21102098 | 15.5546 | 0.053570468 | 0.14813749 | 0.123122287 | 0.172746645 | 15.5546 | 0.032547278 | 0.182066646 | 0.046610976 | 0.09587052 | 15.5546 | 0.0286039 | 0.223789139 | 0.195111973 | 0.289122814 |
| 15.56485 | -0.086766675 | 0.125391502 | 0.074688219 | 0.20837043 | 15.56485 | 0.062097984 | 0.151224429 | 0.126532732 | 0.170152409 | 15.56485 | 0.032479032 | 0.18681575 | 0.049854029 | 0.095750709 | 15.56485 | 0.025637054 | 0.226671848 | 0.196174361 | 0.289943019 |
| 15.57508 | -0.077944123 | 0.12040431 | 0.080177388 | 0.20523606 | 15.57508 | 0.069364664 | 0.154381119 | 0.13067704 | 0.169559904 | 15.57508 | 0.031871239 | 0.191492407 | 0.053053592 | 0.095732341 | 15.57508 | 0.023580756 | 0.230430417 | 0.196456955 | 0.289184946 |
| 15.58533 | -0.068456113 | 0.114552782 | 0.085942875 | 0.20196082 | 15.58533 | 0.075421456 | 0.157177639 | 0.134953133 | 0.171203978 | 15.58533 | 0.031171258 | 0.195682351 | 0.056358289 | 0.095967433 | 15.58533 | 0.022199698 | 0.234209968 | 0.195614826 | 0.287758296 |
| 15.59556 | -0.058612367 | 0.109593618 | 0.09230824 | 0.19882176 | 15.59556 | 0.079755566 | 0.159647609 | 0.138609545 | 0.173733494 | 15.59556 | 0.030051912 | 0.199669926 | 0.059194711 | 0.096556603 | 15.59556 | 0.022295908 | 0.238345879 | 0.193589394 | 0.284995215 |
| 15.60581 | -0.048700421 | 0.104491324 | 0.098133755 | 0.1955065 | 15.60581 | 0.082660212 | 0.161219498 | 0.141455733 | 0.177253151 | 15.60581 | 0.028813891 | 0.202184981 | 0.062238723 | 0.097144112 | 15.60581 | 0.023563343 | 0.242413303 | 0.190492291 | 0.282109725 |
| 15.61604 | -0.039787243 | 0.100336226 | 0.104652214 | 0.19332101 | 15.61604 | 0.083918496 | 0.161695441 | 0.142064555 | 0.181001567 | 15.61604 | 0.027004038 | 0.203533498 | 0.064824915 | 0.098122873 | 15.61604 | 0.026333516 | 0.246166529 | 0.18621396 | 0.27833473 |
| 15.62629 | -0.031251704 | 0.098107596 | 0.111038683 | 0.19108423 | 15.62629 | 0.083524588 | 0.161671831 | 0.141080309 | 0.184237496 | 15.62629 | 0.025174471 | 0.204104012 | 0.067393145 | 0.09901718 | 15.62629 | 0.029969226 | 0.249612174 | 0.1811926 | 0.27513503 |
| 15.63653 | -0.023555235 | 0.09672017 | 0.118189628 | 0.18928773 | 15.63653 | 0.081753728 | 0.160382209 | 0.137849668 | 0.186996572 | 15.63653 | 0.023255368 | 0.203613498 | 0.06972375 | 0.100182526 | 15.63653 | 0.03427783 | 0.252306513 | 0.176013375 | 0.271982693 |
| 15.64676 | -0.017328167 | 0.096577514 | 0.125618309 | 0.18745318 | 15.64676 | 0.079457173 | 0.159533628 | 0.133029815 | 0.189189434 | 15.64676 | 0.021486603 | 0.202459682 | 0.071663725 | 0.101525281 | 15.64676 | 0.038658382 | 0.254508826 | 0.170925246 | 0.270103451 |
| 15.65701 | -0.011670143 | 0.097924152 | 0.134191805 | 0.18660099 | 15.65701 | 0.075912492 | 0.158405146 | 0.126464386 | 0.190964854 | 15.65701 | 0.020112057 | 0.200974885 | 0.073034282 | 0.104090691 | 15.65701 | 0.04212543 | 0.256241931 | 0.166259012 | 0.269513534 |
| 15.66724 | -0.007326595 | 0.099517969 | 0.142956534 | 0.18502107 | 15.66724 | 0.072499411 | 0.159439921 | 0.119868282 | 0.192510843 | 15.66724 | 0.019616927 | 0.199818161 | 0.07346954 | 0.107145593 | 15.66724 | 0.044963753 | 0.257381788 | 0.162669051 | 0.270375935 |
| 15.67749 | -0.004085797 | 0.101490767 | 0.152734957 | 0.18345942 | 15.67749 | 0.068457191 | 0.160754477 | 0.112141696 | 0.193973235 | 15.67749 | 0.019608208 | 0.199318277 | 0.073143653 | 0.11179942 | 15.67749 | 0.046062044 | 0.258658361 | 0.160462511 | 0.273266949 |
| 15.68773 | -0.00176872 | 0.103956318 | 0.162046228 | 0.18176849 | 15.68773 | 0.064818509 | 0.164434549 | 0.104964408 | 0.195481439 | 15.68773 | 0.020700351 | 0.199754299 | 0.071260152 | 0.116885922 | 15.68773 | 0.04581945 | 0.259869642 | 0.158901242 | 0.277079746 |
| 15.69797 | -0.001656584 | 0.105524228 | 0.171407438 | 0.17994301 | 15.69797 | 0.06073795 | 0.168254125 | 0.09742657 | 0.197098914 | 15.69797 | 0.022249487 | 0.201438676 | 0.068713395 | 0.123147705 | 15.69797 | 0.044415414 | 0.260855922 | 0.158694624 | 0.282709246 |
| 15.70821 | -0.00165977 | 0.107315844 | 0.17963631 | 0.17749277 | 15.70821 | 0.056759911 | 0.172620877 | 0.090846884 | 0.198059417 | 15.70821 | 0.024450266 | 0.204278045 | 0.064835117 | 0.129109644 | 15.70821 | 0.04207541 | 0.261957941 | 0.159213326 | 0.28852758 |
| 15.71845 | -0.004226639 | 0.108101044 | 0.187087706 | 0.1747906 | 15.71845 | 0.052189568 | 0.176341133 | 0.083418242 | 0.199274163 | 15.71845 | 0.026501032 | 0.208443057 | 0.060608171 | 0.135137912 | 15.71845 | 0.039700373 | 0.263190168 | 0.160214499 | 0.295225392 |
| 15.72869 | -0.007029049 | 0.108037646 | 0.192758671 | 0.17244398 | 15.72869 | 0.048005098 | 0.179974303 | 0.07650586 | 0.19965228 | 15.72869 | 0.028479114 | 0.213199973 | 0.056085296 | 0.140063292 | 15.72869 | 0.036774697 | 0.264089745 | 0.161136218 | 0.301254172 |
| 15.73892 | -0.011409632 | 0.107685562 | 0.197203342 | 0.16912535 | 15.73892 | 0.043074964 | 0.182893355 | 0.069607313 | 0.200582871 | 15.73892 | 0.029804493 | 0.218481722 | 0.051236591 | 0.144664946 | 15.73892 | 0.03476116 | 0.264818437 | 0.16152849 | 0.307326126 |
| 15.74917 | -0.016383215 | 0.105432154 | 0.200166332 | 0.1663036 | 15.74917 | 0.037851994 | 0.185862127 | 0.061775725 | 0.200696891 | 15.74917 | 0.029964448 | 0.223734895 | 0.046895124 | 0.14778302 | 15.74917 | 0.033191221 | 0.265347095 | 0.161692159 | 0.312276143 |
| 15.75941 | -0.022409199 | 0.103153188 | 0.201649405 | 0.16319142 | 15.75941 | 0.033418136 | 0.18938841 | 0.054667755 | 0.201208838 | 15.75941 | 0.029320172 | 0.228883417 | 0.042932424 | 0.150610171 | 15.75941 | 0.032287815 | 0.265822298 | 0.160376122 | 0.316753494 |
| 15.76965 | -0.028577246 | 0.100156237 | 0.202817676 | 0.16009457 | 15.76965 | 0.028312935 | 0.193513684 | 0.047101066 | 0.200817351 | 15.76965 | 0.027681924 | 0.233491581 | 0.039171206 | 0.152763701 | 15.76965 | 0.03186106 | 0.26632143 | 0.158045881 | 0.31997071 |
| 15.77989 | -0.035457323 | 0.097866434 | 0.202538937 | 0.15735179 | 15.77989 | 0.024271775 | 0.198867963 | 0.040099796 | 0.199801718 | 15.77989 | 0.02518207 | 0.237517828 | 0.036005298 | 0.15494322 | 15.77989 | 0.032018889 | 0.266802206 | 0.154904124 | 0.322059432 |
| 15.79013 | -0.042301473 | 0.096857148 | 0.202937035 | 0.15496596 | 15.79013 | 0.019479393 | 0.205121008 | 0.033380804 | 0.197444164 | 15.79013 | 0.022008245 | 0.240785643 | 0.032882004 | 0.157102753 | 15.79013 | 0.031899928 | 0.267636749 | 0.150458241 | 0.323362269 |
| 15.80037 | -0.049015142 | 0.097514851 | 0.202101914 | 0.15399152 | 15.80037 | 0.016632983 | 0.212291874 | 0.027806953 | 0.193221719 | 15.80037 | 0.018478009 | 0.242847738 | 0.030110735 | 0.159860022 | 15.80037 | 0.032127003 | 0.268817879 | 0.145688521 | 0.322836228 |
| 15.81061 | -0.055669848 | 0.10120678 | 0.202543176 | 0.15387138 | 15.81061 | 0.01315707 | 0.219900108 | 0.02289639 | 0.18764952 | 15.81061 | 0.014829718 | 0.244384406 | 0.027548285 | 0.162996112 | 15.81061 | 0.031393643 | 0.270062531 | 0.140292336 | 0.321658653 |
| 15.82085 | -0.061211541 | 0.107690284 | 0.202465987 | 0.15434782 | 15.82085 | 0.011221842 | 0.227575302 | 0.019379024 | 0.179716979 | 15.82085 | 0.010806857 | 0.245285809 | 0.024766146 | 0.165878967 | 15.82085 | 0.030379142 | 0.271632643 | 0.134861235 | 0.318501122 |
| 15.83109 | -0.066067131 | 0.116491503 | 0.203161098 | 0.15618894 | 15.83109 | 0.009501789 | 0.235060388 | 0.017349543 | 0.170870582 | 15.83109 | 0.007001434 | 0.245191339 | 0.02276633 | 0.169382998 | 15.83109 | 0.028784824 | 0.27335858 | 0.129784607 | 0.314621745 |
| 15.84133 | -0.069866159 | 0.127359445 | 0.203825036 | 0.15826956 | 15.84133 | 0.008914185 | 0.242201542 | 0.016185394 | 0.161217715 | 15.84133 | 0.002986767 | 0.245345505 | 0.020782554 | 0.171511267 | 15.84133 | 0.026048698 | 0.275372432 | 0.125067594 | 0.309594364 |
| 15.85157 | -0.072333626 | 0.138371211 | 0.205094402 | 0.1601026 | 15.85157 | 0.009585243 | 0.248462576 | 0.016054432 | 0.152051368 | 15.85157 | -0.001085039 | 0.24526084 | 0.019458695 | 0.173851856 | 15.85157 | 0.023080663 | 0.277317795 | 0.120621192 | 0.303757764 |
| 15.86181 | -0.073533045 | 0.149746558 | 0.206739418 | 0.16206256 | 15.86181 | 0.010533215 | 0.253988139 | 0.016397064 | 0.144017173 | 15.86181 | -0.005651209 | 0.244542886 | 0.018827687 | 0.174645063 | 15.86181 | 0.018845391 | 0.279658505 | 0.117154191 | 0.2978548 |
| 15.87205 | -0.073639048 | 0.159654853 | 0.207976265 | 0.16301332 | 15.87205 | 0.012795338 | 0.258577934 | 0.01689388 | 0.138307964 | 15.87205 | -0.010078362 | 0.243855114 | 0.019023815 | 0.175182049 | 15.87205 | 0.014720901 | 0.282285558 | 0.114021794 | 0.292170144 |
| 15.88229 | -0.072694703 | 0.167914439 | 0.209785043 | 0.16386138 | 15.88229 | 0.0158313 | 0.261967906 | 0.016568697 | 0.134792064 | 15.88229 | -0.014631708 | 0.24173143 | 0.019878878 | 0.174240649 | 15.88229 | 0.009425205 | 0.285200379 | 0.11160426 | 0.286888849 |
| 15.89253 | -0.070882812 | 0.174268869 | 0.211153998 | 0.16345584 | 15.89253 | 0.019147448 | 0.264057584 | 0.015151951 | 0.133734976 | 15.89253 | -0.019268959 | 0.239029014 | 0.021211264 | 0.173040949 | 15.89253 | 0.004326753 | 0.288132582 | 0.110220974 | 0.282378198 |
| 15.90277 | -0.068210664 | 0.178534023 | 0.212594067 | 0.16322164 | 15.90277 | 0.023429813 | 0.264760897 | 0.012670113 | 0.134178226 | 15.90277 | -0.023670701 | 0.234946643 | 0.02288319 | 0.171177811 | 15.90277 | -0.001409016 | 0.291203127 | 0.109572124 | 0.278592451 |
| 15.91301 | -0.065168073 | 0.180982189 | 0.21354621 | 0.16243088 | 15.91301 | 0.028082068 | 0.263967731 | 0.007910667 | 0.135767267 | 15.91301 | -0.027176566 | 0.22980166 | 0.024792319 | 0.168939604 | 15.91301 | -0.0068268 | 0.294155405 | 0.110114549 | 0.275043575 |
| 15.92325 | -0.062109633 | 0.181934157 | 0.214059918 | 0.16222766 | 15.92325 | 0.033405719 | 0.261592738 | 0.002099186 | 0.137913088 | 15.92325 | -0.030098545 | 0.224295934 | 0.026234895 | 0.166788475 | 15.92325 | -0.012377424 | 0.297018266 | 0.111601765 | 0.271860187 |
| 15.93349 | -0.059072035 | 0.181755076 | 0.213773998 | 0.16170965 | 15.93349 | 0.038859129 | 0.258736604 | -0.005484017 | 0.139775136 | 15.93349 | -0.031599525 | 0.218428221 | 0.027556608 | 0.164288444 | 15.93349 | -0.017062519 | 0.299259996 | 0.114188859 | 0.268888805 |
| 15.94373 | -0.056851429 | 0.180895885 | 0.212977566 | 0.16229199 | 15.94373 | 0.044719301 | 0.254162379 | -0.013342952 | 0.141725747 | 15.94373 | -0.031813349 | 0.212861515 | 0.027778552 | 0.162241998 | 15.94373 | -0.021208308 | 0.300818648 | 0.117692954 | 0.26546155 |
| 15.95397 | -0.055201877 | 0.179792364 | 0.211227023 | 0.16229763 | 15.95397 | 0.051214986 | 0.248944161 | -0.021991384 | 0.143006078 | 15.95397 | -0.030156851 | 0.208364704 | 0.0280825 | 0.159849083 | 15.95397 | -0.024241109 | 0.30126218 | 0.12162062 | 0.262705108 |
| 15.96421 | -0.054771543 | 0.178432056 | 0.208824499 | 0.16173456 | 15.96421 | 0.057025567 | 0.24383046 | -0.029514491 | 0.143995298 | 15.96421 | -0.027426746 | 0.205174388 | 0.027052202 | 0.158117385 | 15.96421 | -0.026008129 | 0.300970488 | 0.125761977 | 0.259925866 |
| 15.97445 | -0.055730206 | 0.177635125 | 0.205706587 | 0.16114588 | 15.97445 | 0.06278715 | 0.238369361 | -0.036316926 | 0.144382235 | 15.97445 | -0.022884716 | 0.203312398 | 0.025688885 | 0.156339035 | 15.97445 | -0.026399366 | 0.298794294 | 0.129906431 | 0.25745595 |
| 15.98469 | -0.057160155 | 0.176706326 | 0.202551298 | 0.15879937 | 15.98469 | 0.068048929 | 0.233605721 | -0.041257665 | 0.14491737 | 15.98469 | -0.017383611 | 0.202807914 | 0.023553963 | 0.154990328 | 15.98469 | -0.025520069 | 0.295955309 | 0.133043665 | 0.255225943 |
| 15.99493 | -0.059803019 | 0.177322947 | 0.199177845 | 0.1560146 | 15.99493 | 0.072660203 | 0.229816102 | -0.044486946 | 0.14493852 | 15.99493 | -0.010868854 | 0.203703093 | 0.020982125 | 0.154003485 | 15.99493 | -0.023240963 | 0.291421328 | 0.135729017 | 0.253416225 |
| 16.00517 | -0.062307373 | 0.178074803 | 0.196672834 | 0.15172895 | 16.00517 | 0.076240634 | 0.227135251 | -0.045623382 | 0.145429994 | 16.00517 | -0.003895886 | 0.205052288 | 0.017963213 | 0.153684878 | 16.00517 | -0.019960683 | 0.286071728 | 0.137175256 | 0.250970964 |
| 16.01541 | -0.064893596 | 0.180311214 | 0.195006931 | 0.14732262 | 16.01541 | 0.079522384 | 0.226658105 | -0.045057858 | 0.145272563 | 16.01541 | 0.003547029 | 0.206535662 | 0.014791345 | 0.153616995 | 16.01541 | -0.016732643 | 0.279888387 | 0.137457136 | 0.248616944 |
| 16.02565 | -0.066228366 | 0.182753864 | 0.194574983 | 0.1426582 | 16.02565 | 0.08221285 | 0.227197987 | -0.043208948 | 0.145174348 | 16.02565 | 0.01064671 | 0.207562162 | 0.011575329 | 0.154248336 | 16.02565 | -0.012957938 | 0.273478707 | 0.136428653 | 0.245794911 |
| 16.03589 | -0.067355068 | 0.184935235 | 0.195534302 | 0.13782229 | 16.03589 | 0.08459008 | 0.229468778 | -0.040537757 | 0.14489403 | 16.03589 | 0.018256836 | 0.207521698 | 0.008663795 | 0.155306652 | 16.03589 | -0.009789977 | 0.26730889 | 0.134467484 | 0.242549689 |
| 16.04613 | -0.066476217 | 0.186674113 | 0.197357391 | 0.1343987 | 16.04613 | 0.086758117 | 0.232854338 | -0.036788861 | 0.144313781 | 16.04613 | 0.024870715 | 0.205938379 | 0.006615409 | 0.156926017 | 16.04613 | -0.007647898 | 0.261368173 | 0.131183249 | 0.239393422 |
| 16.05637 | -0.064485996 | 0.186530454 | 0.201022717 | 0.13178729 | 16.05637 | 0.088965535 | 0.236910974 | -0.033263735 | 0.143613664 | 16.05637 | 0.031386742 | 0.202886927 | 0.005257639 | 0.159538238 | 16.05637 | -0.00638894 | 0.256900277 | 0.127245244 | 0.23612665 |
| 16.06661 | -0.061571968 | 0.18415485 | 0.204401627 | 0.13039162 | 16.06661 | 0.091666818 | 0.240793778 | -0.02957685 | 0.143282703 | 16.06661 | 0.037248746 | 0.198140677 | 0.004878685 | 0.161791056 | 16.06661 | -0.005921019 | 0.253351126 | 0.122435158 | 0.233170496 |
| 16.07685 | -0.057266152 | 0.180331289 | 0.207867047 | 0.13050234 | 16.07685 | 0.094127936 | 0.244123567 | -0.026025751 | 0.143894484 | 16.07685 | 0.042565696 | 0.191900324 | 0.005560425 | 0.165120662 | 16.07685 | -0.006679969 | 0.251640882 | 0.117354348 | 0.23069546 |
| 16.08709 | -0.052973351 | 0.174140108 | 0.210974809 | 0.13118429 | 16.08709 | 0.096933631 | 0.246106108 | -0.0224638 | 0.144774022 | 16.08709 | 0.047196667 | 0.184886745 | 0.007141626 | 0.168236805 | 16.08709 | -0.007158655 | 0.250960715 | 0.111858975 | 0.228116502 |
| 16.09733 | -0.048673671 | 0.167578075 | 0.21296126 | 0.13331176 | 16.09733 | 0.099391118 | 0.247449977 | -0.019213878 | 0.146933381 | 16.09733 | 0.051620427 | 0.177254392 | 0.009180902 | 0.171470159 | 16.09733 | -0.008584789 | 0.251746826 | 0.106716582 | 0.226228914 |
| 16.10757 | -0.045339732 | 0.160860257 | 0.212925339 | 0.13571275 | 16.10757 | 0.101720286 | 0.247119204 | -0.015850452 | 0.149553119 | 16.10757 | 0.055617601 | 0.169706151 | 0.011267062 | 0.174165413 | 16.10757 | -0.008728269 | 0.25353189 | 0.101497174 | 0.224702805 |
| 16.11781 | -0.043451909 | 0.155529833 | 0.211452304 | 0.1380106 | 16.11781 | 0.103729285 | 0.247024841 | -0.012573591 | 0.153012833 | 16.11781 | 0.059568546 | 0.162352011 | 0.01326279 | 0.176870824 | 16.11781 | -0.009025352 | 0.255995381 | 0.097007221 | 0.222902531 |
| 16.12805 | -0.043500368 | 0.152197703 | 0.20836755 | 0.14023018 | 16.12805 | 0.104445219 | 0.246504072 | -0.009730758 | 0.155772267 | 16.12805 | 0.063570011 | 0.155508708 | 0.014625408 | 0.178699579 | 16.12805 | -0.0081949 | 0.258765908 | 0.093200478 | 0.222441383 |
| 16.13829 | -0.045146301 | 0.150995955 | 0.203634251 | 0.14250832 | 16.13829 | 0.105077202 | 0.247400069 | -0.00637184 | 0.159509326 | 16.13829 | 0.067372143 | 0.149209652 | 0.014768658 | 0.180485041 | 16.13829 | -0.007220505 | 0.261525067 | 0.089874451 | 0.221525343 |
| 16.14853 | -0.049209259 | 0.151628187 | 0.198330822 | 0.14431704 | 16.14853 | 0.104361027 | 0.249554466 | -0.003573806 | 0.162484638 | 16.14853 | 0.071060949 | 0.143605765 | 0.014597914 | 0.181356337 | 16.14853 | -0.005088973 | 0.263661181 | 0.08772927 | 0.221679224 |
| 16.15877 | -0.054376226 | 0.15432995 | 0.19236852 | 0.14597452 | 16.15877 | 0.103665081 | 0.253890785 | -0.000724988 | 0.165745207 | 16.15877 | 0.074501267 | 0.138777276 | 0.012902194 | 0.182255053 | 16.15877 | -0.003349745 | 0.265290912 | 0.085909252 | 0.221605281 |
| 16.16901 | -0.06033475 | 0.157235344 | 0.186527259 | 0.14791232 | 16.16901 | 0.102296529 | 0.260136117 | 0.001467278 | 0.169237628 | 16.16901 | 0.077142418 | 0.135722549 | 0.010799814 | 0.182249043 | 16.16901 | -0.001024937 | 0.265638627 | 0.085165687 | 0.221512918 |
| 16.17925 | -0.066383938 | 0.160548716 | 0.181056742 | 0.1491755 | 16.17925 | 0.101062146 | 0.268072078 | 0.003083129 | 0.173000706 | 16.17925 | 0.079440628 | 0.133791756 | 0.00811503 | 0.182397735 | 16.17925 | 0.000886038 | 0.264788318 | 0.084675539 | 0.220543254 |
| 16.18949 | -0.071203774 | 0.163857321 | 0.176656582 | 0.15133728 | 16.18949 | 0.100340334 | 0.277564177 | 0.003967066 | 0.176809325 | 16.18949 | 0.080125473 | 0.133873234 | 0.005478502 | 0.181913982 | 16.18949 | 0.003298967 | 0.263300304 | 0.08476868 | 0.218405827 |
| 16.19973 | -0.074854103 | 0.165315189 | 0.172317219 | 0.15312061 | 16.19973 | 0.100343415 | 0.287307473 | 0.00375049 | 0.180296041 | 16.19973 | 0.080496479 | 0.13568082 | 0.003305103 | 0.181704138 | 16.19973 | 0.005047438 | 0.260933188 | 0.085265576 | 0.214792844 |
| 16.20997 | -0.076472441 | 0.166702794 | 0.16914299 | 0.15601296 | 16.20997 | 0.101902085 | 0.297031269 | 0.003075802 | 0.183921505 | 16.20997 | 0.079232883 | 0.138560835 | 0.001785582 | 0.181160224 | 16.20997 | 0.007452897 | 0.258243614 | 0.085527319 | 0.20943331 |
| 16.22021 | -0.076010471 | 0.165629991 | 0.166093995 | 0.15868801 | 16.22021 | 0.104260592 | 0.305708844 | 0.001184134 | 0.18629682 | 16.22021 | 0.077936003 | 0.141817435 | 0.000874355 | 0.180953279 | 16.22021 | 0.009931556 | 0.255416481 | 0.086160221 | 0.202960082 |
| 16.23045 | -0.07351777 | 0.163866209 | 0.163734741 | 0.1622257 | 16.23045 | 0.108727736 | 0.312713499 | -0.000636756 | 0.18795913 | 16.23045 | 0.075250413 | 0.14524564 | 0.001271 | 0.1813258 | 16.23045 | 0.012986169 | 0.252995002 | 0.086659399 | 0.1957724 |
| 16.24069 | -0.070078798 | 0.16045949 | 0.161610644 | 0.16547508 | 16.24069 | 0.114207941 | 0.317509232 | -0.00298388 | 0.188564594 | 16.24069 | 0.072511113 | 0.147948147 | 0.002465389 | 0.181649696 | 16.24069 | 0.016485453 | 0.250569079 | 0.086643392 | 0.188459566 |
| 16.25093 | -0.06534812 | 0.156887123 | 0.159890312 | 0.1686663 | 16.25093 | 0.120616587 | 0.320697894 | -0.005152253 | 0.188602107 | 16.25093 | 0.069515495 | 0.150219519 | 0.004433194 | 0.183115018 | 16.25093 | 0.020367634 | 0.248498554 | 0.086403108 | 0.181848994 |
| 16.26117 | -0.060943597 | 0.15277075 | 0.158775802 | 0.17062822 | 16.26117 | 0.127616783 | 0.320784698 | -0.00713615 | 0.187686956 | 16.26117 | 0.066546256 | 0.151550937 | 0.007012986 | 0.184747679 | 16.26117 | 0.024489273 | 0.246332759 | 0.086180334 | 0.176074936 |
| 16.27141 | -0.056771193 | 0.149608558 | 0.157600888 | 0.17170386 | 16.27141 | 0.134568977 | 0.319323232 | -0.008641113 | 0.186875611 | 16.27141 | 0.063754206 | 0.152721291 | 0.010016349 | 0.187052656 | 16.27141 | 0.028279951 | 0.244872949 | 0.084815431 | 0.171878296 |
| 16.28165 | -0.053873853 | 0.147358322 | 0.157557403 | 0.17140358 | 16.28165 | 0.14082195 | 0.314981707 | -0.009563483 | 0.18581655 | 16.28165 | 0.061293112 | 0.153018553 | 0.013419865 | 0.189577612 | 16.28165 | 0.031370078 | 0.242510068 | 0.083351238 | 0.169007602 |
| 16.29189 | -0.052065191 | 0.146267865 | 0.157183037 | 0.1694742 | 16.29189 | 0.145732097 | 0.309556595 | -0.010296859 | 0.184990156 | 16.29189 | 0.059432423 | 0.153612415 | 0.01666943 | 0.192548989 | 16.29189 | 0.03369812 | 0.240868967 | 0.081105281 | 0.167537164 |
| 16.30213 | -0.050981927 | 0.146680624 | 0.157058264 | 0.1667705 | 16.30213 | 0.14965891 | 0.301426049 | -0.009726053 | 0.184051899 | 16.30213 | 0.057754056 | 0.154246274 | 0.020113268 | 0.195213457 | 16.30213 | 0.034710049 | 0.238617029 | 0.078412386 | 0.167883039 |
| 16.31237 | -0.051295582 | 0.147551969 | 0.156659754 | 0.1628929 | 16.31237 | 0.151927864 | 0.292573597 | -0.009071378 | 0.183971035 | 16.31237 | 0.056662333 | 0.154902092 | 0.023514362 | 0.197116483 | 16.31237 | 0.034064847 | 0.236594473 | 0.075159895 | 0.169165509 |
| 16.32261 | -0.051277755 | 0.150180509 | 0.156544807 | 0.1585722 | 16.32261 | 0.152602187 | 0.282088238 | -0.007428336 | 0.183231258 | 16.32261 | 0.055721609 | 0.155155848 | 0.026639351 | 0.19868029 | 16.32261 | 0.032464216 | 0.233871315 | 0.071261077 | 0.172025752 |
| 16.33285 | -0.051727719 | 0.15309889 | 0.155755991 | 0.15419818 | 16.33285 | 0.152308287 | 0.27239057 | -0.005291871 | 0.183471005 | 16.33285 | 0.055676153 | 0.155931255 | 0.029806561 | 0.19896774 | 16.33285 | 0.029252512 | 0.230682357 | 0.067337454 | 0.175718817 |
| 16.34309 | -0.050700375 | 0.155908109 | 0.155215326 | 0.1504368 | 16.34309 | 0.15086782 | 0.26240087 | -0.003079101 | 0.183277264 | 16.34309 | 0.055605938 | 0.155475827 | 0.032836666 | 0.198401561 | 16.34309 | 0.025139339 | 0.22715016 | 0.063089507 | 0.179777036 |
| 16.35333 | -0.050011147 | 0.159539129 | 0.15402592 | 0.14690341 | 16.35333 | 0.14863673 | 0.254754895 | -0.000628405 | 0.183763331 | 16.35333 | 0.056163049 | 0.1547023 | 0.035528483 | 0.196865992 | 16.35333 | 0.019938807 | 0.222250944 | 0.058578243 | 0.183760933 |
| 16.36357 | -0.047727524 | 0.162989232 | 0.153268095 | 0.14368229 | 16.36357 | 0.145587898 | 0.248830948 | 0.001372864 | 0.18388166 | 16.36357 | 0.057133038 | 0.153214845 | 0.038304782 | 0.194750261 | 16.36357 | 0.013837519 | 0.216966109 | 0.054163252 | 0.187147284 |
| 16.37381 | -0.045020017 | 0.166417407 | 0.152514269 | 0.14121067 | 16.37381 | 0.14247059 | 0.245263548 | 0.00355821 | 0.18415277 | 16.37381 | 0.058510557 | 0.151502511 | 0.04053706 | 0.19220691 | 16.37381 | 0.00748159 | 0.210377715 | 0.049924901 | 0.188817465 |
| 16.38405 | -0.041549361 | 0.169371763 | 0.151643924 | 0.13897951 | 16.38405 | 0.138530736 | 0.243657841 | 0.004999482 | 0.184660158 | 16.38405 | 0.060055206 | 0.149511599 | 0.042723911 | 0.189664123 | 16.38405 | 0.000284817 | 0.203722698 | 0.045642505 | 0.189899576 |
| 16.39429 | -0.038621222 | 0.171673923 | 0.151024408 | 0.1375635 | 16.39429 | 0.134749932 | 0.243454036 | 0.006655891 | 0.184772959 | 16.39429 | 0.061915565 | 0.148229556 | 0.044534072 | 0.187684324 | 16.39429 | -0.006742593 | 0.196737352 | 0.041910529 | 0.188793984 |
| 16.40453 | -0.034959128 | 0.173332235 | 0.150045625 | 0.13593938 | 16.40453 | 0.130493057 | 0.244695355 | 0.008654336 | 0.18476849 | 16.40453 | 0.063748358 | 0.147224794 | 0.045981412 | 0.18618884 | 16.40453 | -0.013787307 | 0.189938715 | 0.038589078 | 0.187352889 |
| 16.41477 | -0.032017436 | 0.173854585 | 0.149622397 | 0.13591982 | 16.41477 | 0.126465287 | 0.244988698 | 0.010793874 | 0.184645162 | 16.41477 | 0.065212134 | 0.147474198 | 0.046695986 | 0.185607074 | 16.41477 | -0.020613479 | 0.183932944 | 0.036046192 | 0.184572653 |
| 16.42501 | -0.02890036 | 0.174037466 | 0.148174586 | 0.1359109 | 16.42501 | 0.122441523 | 0.245973979 | 0.013945436 | 0.184537396 | 16.42501 | 0.066738879 | 0.147793873 | 0.047523729 | 0.185696744 | 16.42501 | -0.026396361 | 0.178279684 | 0.034357442 | 0.182164254 |
| 16.43525 | -0.026750147 | 0.173179329 | 0.147538483 | 0.13707461 | 16.43525 | 0.118618247 | 0.245574798 | 0.017335014 | 0.184334398 | 16.43525 | 0.067173507 | 0.149125458 | 0.047607873 | 0.186104368 | 16.43525 | -0.031714721 | 0.173690146 | 0.033736023 | 0.179700499 |
| 16.44549 | -0.024627917 | 0.172745042 | 0.146266473 | 0.13820393 | 16.44549 | 0.114848285 | 0.245021629 | 0.021805222 | 0.184168546 | 16.44549 | 0.067296723 | 0.150588197 | 0.047855963 | 0.186403818 | 16.44549 | -0.035801084 | 0.169779761 | 0.034320506 | 0.177677286 |
| 16.45573 | -0.022513879 | 0.171066707 | 0.145546604 | 0.1401743 | 16.45573 | 0.111680453 | 0.24363916 | 0.026454951 | 0.184188809 | 16.45573 | 0.066224707 | 0.152203471 | 0.047878115 | 0.186529643 | 16.45573 | -0.039120033 | 0.166486533 | 0.035926939 | 0.176797489 |
| 16.46597 | -0.020569524 | 0.170837347 | 0.144441342 | 0.14260367 | 16.46597 | 0.108607703 | 0.242273884 | 0.031445708 | 0.184372757 | 16.46597 | 0.064378019 | 0.153497912 | 0.048738165 | 0.185653589 | 16.46597 | -0.040958491 | 0.16322281 | 0.038345803 | 0.176462094 |
| 16.47621 | -0.017990837 | 0.170116905 | 0.14454238 | 0.145097 | 16.47621 | 0.105802344 | 0.241149041 | 0.036912238 | 0.184944951 | 16.47621 | 0.061164673 | 0.154776364 | 0.049724604 | 0.184775216 | 16.47621 | -0.041854825 | 0.160907811 | 0.041686992 | 0.176731093 |
| 16.48645 | -0.014804361 | 0.170515729 | 0.144368221 | 0.14813629 | 16.48645 | 0.1032296 | 0.240357116 | 0.042365277 | 0.185234794 | 16.48645 | 0.057236544 | 0.155494913 | 0.051918406 | 0.18334885 | 16.48645 | -0.041474099 | 0.157959436 | 0.045124521 | 0.17803858 |
| 16.49669 | -0.010268228 | 0.172428805 | 0.144682523 | 0.15149739 | 16.49669 | 0.100986343 | 0.240252581 | 0.04785554 | 0.186757106 | 16.49669 | 0.052445713 | 0.155481695 | 0.054902001 | 0.182257508 | 16.49669 | -0.039803265 | 0.155085848 | 0.048482387 | 0.179445168 |
| 16.50693 | -0.004972 | 0.175116255 | 0.145677969 | 0.15515002 | 16.50693 | 0.098785945 | 0.240301333 | 0.052824724 | 0.188190499 | 16.50693 | 0.047807128 | 0.154736761 | 0.05908203 | 0.180976087 | 16.50693 | -0.03709903 | 0.151873859 | 0.051195672 | 0.181432551 |
| 16.51717 | 0.002590475 | 0.179499182 | 0.146550045 | 0.15793198 | 16.51717 | 0.096572019 | 0.240674312 | 0.057065758 | 0.191159401 | 16.51717 | 0.042957008 | 0.153622226 | 0.063988425 | 0.180419399 | 16.51717 | -0.03347435 | 0.148735717 | 0.053332671 | 0.183301813 |
| 16.52741 | 0.010322572 | 0.184711122 | 0.14765415 | 0.16153055 | 16.52741 | 0.094222715 | 0.240653568 | 0.060461 | 0.194577292 | 16.52741 | 0.039249508 | 0.15183684 | 0.069646605 | 0.179654637 | 16.52741 | -0.029077842 | 0.145860343 | 0.054098318 | 0.185095956 |
| 16.53765 | 0.019977796 | 0.190679756 | 0.148681085 | 0.1630156 | 16.53765 | 0.092413078 | 0.2399735 | 0.061897506 | 0.199129437 | 16.53765 | 0.036565031 | 0.150384085 | 0.075651522 | 0.178568621 | 16.53765 | -0.024574364 | 0.142887805 | 0.053568804 | 0.186700631 |
| 16.54789 | 0.029553708 | 0.196648082 | 0.149412658 | 0.16485922 | 16.54789 | 0.089411769 | 0.238407508 | 0.060771456 | 0.204396433 | 16.54789 | 0.035208015 | 0.148802923 | 0.081472599 | 0.176964188 | 16.54789 | -0.020085838 | 0.141044897 | 0.05205958 | 0.187560362 |
| 16.55813 | 0.039682851 | 0.202237678 | 0.149597455 | 0.16417243 | 16.55813 | 0.086973519 | 0.236006244 | 0.057899135 | 0.210053943 | 16.55813 | 0.035020369 | 0.148322903 | 0.087199427 | 0.174212403 | 16.55813 | -0.016337225 | 0.140313554 | 0.049467126 | 0.187372601 |
| 16.56837 | 0.049289613 | 0.207109919 | 0.149513779 | 0.1631741 | 16.56837 | 0.083813743 | 0.232982321 | 0.051968324 | 0.215628021 | 16.56837 | 0.035992247 | 0.148375285 | 0.091504981 | 0.170838481 | 16.56837 | -0.013190979 | 0.140505423 | 0.046268121 | 0.186797772 |
| 16.57861 | 0.058138134 | 0.210592933 | 0.148351857 | 0.15974553 | 16.57861 | 0.080880307 | 0.230042895 | 0.044034734 | 0.220290921 | 16.57861 | 0.037368278 | 0.149031943 | 0.095217475 | 0.166275909 | 16.57861 | -0.01150516 | 0.142143506 | 0.042735461 | 0.185517311 |
| 16.58885 | 0.066488675 | 0.213586247 | 0.146573931 | 0.15526473 | 16.58885 | 0.077176126 | 0.227759304 | 0.033934572 | 0.2244768 | 16.58885 | 0.039250115 | 0.150348681 | 0.097473621 | 0.161791515 | 16.58885 | -0.010932935 | 0.144811838 | 0.039062284 | 0.183380545 |
| 16.59909 | 0.073494053 | 0.214997749 | 0.143868302 | 0.1489355 | 16.59909 | 0.073527062 | 0.227242309 | 0.022922637 | 0.227508915 | 16.59909 | 0.040722407 | 0.152370369 | 0.098378292 | 0.157412715 | 16.59909 | -0.011577918 | 0.149229248 | 0.036004527 | 0.1816675 |
| 16.60933 | 0.079251597 | 0.215386935 | 0.140147425 | 0.14098727 | 16.60933 | 0.069974032 | 0.228004838 | 0.010851799 | 0.229474302 | 16.60933 | 0.04224424 | 0.155182534 | 0.098051237 | 0.154519396 | 16.60933 | -0.013251196 | 0.153459474 | 0.03297765 | 0.179376575 |
| 16.61957 | 0.084179356 | 0.215749059 | 0.135813001 | 0.13172614 | 16.61957 | 0.066198509 | 0.231173118 | -0.000625031 | 0.229861121 | 16.61957 | 0.042949272 | 0.158368395 | 0.096988825 | 0.152816406 | 16.61957 | -0.015868434 | 0.157941748 | 0.031222175 | 0.178053243 |
| 16.62981 | 0.088101181 | 0.215551688 | 0.130208472 | 0.1216206 | 16.62981 | 0.06263404 | 0.235545245 | -0.011927971 | 0.229887928 | 16.62981 | 0.043809654 | 0.162235256 | 0.095026753 | 0.153643851 | 16.62981 | -0.018929453 | 0.162347125 | 0.029255941 | 0.177013884 |
| 16.64005 | 0.090308905 | 0.215797126 | 0.124678639 | 0.110973 | 16.64005 | 0.059466828 | 0.241168167 | -0.021646285 | 0.228798461 | 16.64005 | 0.043767675 | 0.16588913 | 0.092518206 | 0.155898362 | 16.64005 | -0.022414051 | 0.165995796 | 0.028676338 | 0.177002727 |
| 16.65029 | 0.092293571 | 0.216605618 | 0.118218964 | 0.10070181 | 16.65029 | 0.056751035 | 0.246234852 | -0.030302453 | 0.227321821 | 16.65029 | 0.044407508 | 0.169055714 | 0.089950774 | 0.160313734 | 16.65029 | -0.025739178 | 0.168895544 | 0.027731285 | 0.177684084 |
| 16.66053 | 0.092639673 | 0.218860954 | 0.111676991 | 0.09183229 | 16.66053 | 0.054331978 | 0.250681069 | -0.036980336 | 0.225624105 | 16.66053 | 0.044177983 | 0.171446393 | 0.087013662 | 0.16555962 | 16.66053 | -0.028708495 | 0.171313095 | 0.02800517 | 0.179267226 |
| 16.67077 | 0.092393596 | 0.221871219 | 0.105787023 | 0.0842426 | 16.67077 | 0.053228991 | 0.25296253 | -0.042092133 | 0.22479781 | 16.67077 | 0.044152219 | 0.172591313 | 0.084238232 | 0.17141081 | 16.67077 | -0.031301092 | 0.173597026 | 0.028138926 | 0.18146585 |
| 16.68101 | 0.090080697 | 0.226365994 | 0.099928386 | 0.07936793 | 16.68101 | 0.052883255 | 0.253200752 | -0.045249418 | 0.224519773 | 16.68101 | 0.043664138 | 0.173059459 | 0.080740475 | 0.17767784 | 16.68101 | -0.033217652 | 0.175344238 | 0.028500495 | 0.184460092 |
| 16.69125 | 0.087512459 | 0.2312274 | 0.095484447 | 0.07607539 | 16.69125 | 0.053723167 | 0.250524685 | -0.046919738 | 0.225763054 | 16.69125 | 0.042855925 | 0.17276508 | 0.077561093 | 0.183278655 | 16.69125 | -0.034236556 | 0.176414147 | 0.029572807 | 0.187616771 |
| 16.70149 | 0.082560393 | 0.236596636 | 0.092031838 | 0.07476767 | 16.70149 | 0.056080127 | 0.245666849 | -0.047181715 | 0.228107727 | 16.70149 | 0.041387087 | 0.172019331 | 0.073649322 | 0.188464717 | 16.70149 | -0.03472692 | 0.177192331 | 0.030439423 | 0.191065931 |
| 16.71173 | 0.076533564 | 0.241323015 | 0.090699433 | 0.07449893 | 16.71173 | 0.060231831 | 0.237976591 | -0.04573289 | 0.230983296 | 16.71173 | 0.039557005 | 0.171575024 | 0.06972824 | 0.191798483 | 16.71173 | -0.034563943 | 0.176781474 | 0.031678062 | 0.194422201 |
| 16.72197 | 0.069054766 | 0.245683668 | 0.090706507 | 0.0747602 | 16.72197 | 0.065857352 | 0.228733754 | -0.043131455 | 0.234210711 | 16.72197 | 0.036963996 | 0.171634158 | 0.065550369 | 0.194035854 | 16.72197 | -0.033615904 | 0.175399649 | 0.032570106 | 0.197364385 |
| 16.73221 | 0.060384739 | 0.247894619 | 0.093052257 | 0.07570944 | 16.73221 | 0.072889889 | 0.21778608 | -0.039080235 | 0.236893978 | 16.73221 | 0.034431062 | 0.172056171 | 0.061547745 | 0.193772992 | 16.73221 | -0.032217547 | 0.172761476 | 0.033879589 | 0.19970892 |
| 16.74245 | 0.050901654 | 0.248878514 | 0.096737474 | 0.0769182 | 16.74245 | 0.081586888 | 0.205459724 | -0.033945601 | 0.23901893 | 16.74245 | 0.031600117 | 0.172948203 | 0.057561584 | 0.192055903 | 16.74245 | -0.030429867 | 0.169441163 | 0.034551784 | 0.20094428 |
| 16.75269 | 0.041142339 | 0.247748109 | 0.102302922 | 0.07848862 | 16.75269 | 0.090822148 | 0.192689459 | -0.028281287 | 0.240372933 | 16.75269 | 0.029318987 | 0.174581284 | 0.054059062 | 0.187756633 | 16.75269 | -0.028548861 | 0.165441766 | 0.035062246 | 0.201107803 |
| 16.76293 | 0.03212061 | 0.245464448 | 0.108463871 | 0.08013056 | 16.76293 | 0.100745668 | 0.179569466 | -0.021358859 | 0.241285797 | 16.76293 | 0.027320867 | 0.175725164 | 0.050860206 | 0.182683355 | 16.76293 | -0.026607496 | 0.161140359 | 0.034783804 | 0.200213571 |
| 16.77317 | 0.023563626 | 0.241242545 | 0.115143189 | 0.08209102 | 16.77317 | 0.110066352 | 0.166917691 | -0.015086889 | 0.241451965 | 16.77317 | 0.02631095 | 0.177486371 | 0.047879733 | 0.175843307 | 16.77317 | -0.024732482 | 0.15735515 | 0.034011072 | 0.198090208 |
| 16.78341 | 0.016521851 | 0.236384656 | 0.121901822 | 0.0838088 | 16.78341 | 0.119180848 | 0.154716977 | -0.008688258 | 0.242373426 | 16.78341 | 0.0257479 | 0.178185463 | 0.045086659 | 0.168437608 | 16.78341 | -0.023398543 | 0.153825123 | 0.032031738 | 0.195347472 |

| | | | | |
|---|---|---|---|---|
| 16.79365 | 0.011531598 | 0.231397173 | 0.127848571 | 0.08573898 |
| 16.80389 | 0.007914761 | 0.225940123 | 0.133056508 | 0.08755178 |
| 16.81413 | 0.006461562 | 0.22135158 | 0.136882101 | 0.09006116 |
| 16.82437 | 0.006490102 | 0.217585608 | 0.139998673 | 0.09287352 |
| 16.83461 | 0.007992352 | 0.214821275 | 0.14146544 | 0.09714833 |
| 16.84485 | 0.01023239 | 0.212555641 | 0.142163115 | 0.10239936 |
| 16.85509 | 0.013071928 | 0.212020804 | 0.141692018 | 0.10934615 |
| 16.86533 | 0.015651944 | 0.21140898 | 0.140675736 | 0.11745536 |
| 16.87557 | 0.017432101 | 0.211653319 | 0.138050115 | 0.12595775 |
| 16.88581 | 0.018378077 | 0.212026956 | 0.135424318 | 0.13440683 |
| 16.89605 | 0.017557954 | 0.212344488 | 0.130763311 | 0.14248075 |
| 16.90629 | 0.015014407 | 0.212889996 | 0.126267623 | 0.14893178 |
| 16.91653 | 0.010658666 | 0.212878234 | 0.120111004 | 0.154002 |
| 16.92677 | 0.004876494 | 0.213317994 | 0.113827997 | 0.15706269 |
| 16.93701 | -0.0023087 | 0.213864495 | 0.106630636 | 0.15839712 |
| 16.94725 | -0.010099112 | 0.214473351 | 0.0997657 | 0.15865426 |
| 16.95749 | -0.017841197 | 0.215052735 | 0.092857001 | 0.15751825 |
| 16.96773 | -0.024854591 | 0.216005347 | 0.086893311 | 0.15624778 |
| 16.97797 | -0.030737877 | 0.216475325 | 0.081180626 | 0.15471764 |
| 16.98821 | -0.034551101 | 0.217183097 | 0.077294147 | 0.15305979 |
| 16.99845 | -0.03650338 | 0.21668722 | 0.074084524 | 0.15121239 |
| 17.00869 | -0.035558926 | 0.217086627 | 0.07203735 | 0.14923263 |
| 17.01893 | -0.033164189 | 0.215212249 | 0.070921954 | 0.14651438 |
| 17.02917 | -0.027936863 | 0.213777823 | 0.07065905 | 0.14259452 |
| 17.03941 | -0.021585932 | 0.21129655 | 0.07061612 | 0.13758987 |
| 17.04965 | -0.013899003 | 0.207908805 | 0.070355263 | 0.1312596 |
| 17.05989 | -0.005720042 | 0.204650162 | 0.070488843 | 0.12375447 |
| 17.07013 | 0.002474319 | 0.200683377 | 0.069865148 | 0.11506825 |
| 17.08037 | 0.010071412 | 0.197487889 | 0.068759846 | 0.10606828 |
| 17.09061 | 0.016658646 | 0.193832189 | 0.067089864 | 0.09710152 |
| 17.10085 | 0.022229333 | 0.192053327 | 0.065109903 | 0.08847012 |
| 17.11109 | 0.025881031 | 0.190041416 | 0.062537108 | 0.08108563 |
| 17.12133 | 0.02860243 | 0.189973392 | 0.0602018 | 0.07609952 |
| 17.13157 | 0.029533088 | 0.190327118 | 0.057255513 | 0.07333341 |
| 17.14181 | 0.029145988 | 0.191965518 | 0.055284749 | 0.07398991 |
| 17.15205 | 0.027649076 | 0.194532466 | 0.053005099 | 0.0775008 |
| 17.16229 | 0.025170309 | 0.197473727 | 0.051565014 | 0.08347568 |
| 17.17253 | 0.022266838 | 0.201478367 | 0.050546515 | 0.09109877 |
| 17.18277 | 0.018351764 | 0.20613682 | 0.04988477 | 0.10044314 |
| 17.19301 | 0.014985392 | 0.211057861 | 0.049900016 | 0.11031573 |
| 17.20325 | 0.011100701 | 0.216658197 | 0.049642571 | 0.12061868 |
| 17.21349 | 0.007602711 | 0.22328661 | 0.04981376 | 0.13105578 |
| 17.22373 | 0.004045165 | 0.229433158 | 0.050023249 | 0.1404975 |
| 17.23397 | 0.000796226 | 0.236329843 | 0.049275447 | 0.14914683 |
| 17.24421 | -0.001958973 | 0.242914439 | 0.049384983 | 0.15679805 |
| 17.25445 | -0.005012862 | 0.24891216 | 0.047893957 | 0.1632341 |
| 17.26469 | -0.007399867 | 0.253721424 | 0.046862475 | 0.16895212 |
| 17.27493 | -0.009548749 | 0.257295051 | 0.044523066 | 0.17414579 |
| 17.28517 | -0.011169158 | 0.259026944 | 0.041837631 | 0.17951024 |
| 17.29541 | -0.011976488 | 0.258615588 | 0.038436055 | 0.18543371 |
| 17.30565 | -0.012170651 | 0.256698272 | 0.033768651 | 0.19127878 |
| 17.31589 | -0.010875517 | 0.252828473 | 0.028847881 | 0.19798544 |
| 17.32613 | -0.008800854 | 0.248404366 | 0.023132942 | 0.20460642 |
| 17.33637 | -0.00540413 | 0.243655379 | 0.016989372 | 0.21055139 |
| 17.34661 | -0.001032608 | 0.239235555 | 0.011497906 | 0.21566169 |
| 17.35685 | 0.004477977 | 0.235892903 | 0.006250735 | 0.21898349 |
| 17.36709 | 0.010093939 | 0.234069906 | 0.002250948 | 0.22091502 |
| 17.37733 | 0.016627812 | 0.233499027 | -0.000236175 | 0.22070869 |
| 17.38757 | 0.02313465 | 0.233603627 | -0.000789016 | 0.21895901 |
| 17.39781 | 0.029387027 | 0.235259302 | 0.000115082 | 0.21570778 |
| 17.40805 | 0.035607697 | 0.236291517 | 0.002559045 | 0.211182 |
| 17.41829 | 0.041346701 | 0.237646706 | 0.006927217 | 0.20615196 |
| 17.42853 | 0.046799782 | 0.238654745 | 0.011721315 | 0.20063093 |
| 17.43877 | 0.05131069 | 0.239146229 | 0.016748328 | 0.19474582 |
| 17.44901 | 0.055369033 | 0.239297283 | 0.022109864 | 0.18801567 |
| 17.45925 | 0.058248548 | 0.239622924 | 0.026532756 | 0.18136435 |
| 17.46949 | 0.060302705 | 0.238912772 | 0.029719802 | 0.17351448 |
| 17.47973 | 0.061548804 | 0.238598118 | 0.031987244 | 0.16490602 |
| 17.48997 | 0.061383274 | 0.237405474 | 0.032278372 | 0.15577573 |
| 17.50021 | 0.060817031 | 0.235336743 | 0.03137473 | 0.14579993 |
| 17.51045 | 0.059837841 | 0.232362377 | 0.028592845 | 0.13572277 |
| 17.52069 | 0.058296534 | 0.228640312 | 0.025139903 | 0.12529388 |
| 17.53093 | 0.057259544 | 0.224246281 | 0.02065972 | 0.11496461 |
| 17.54117 | 0.056196821 | 0.219998752 | 0.01565665 | 0.10553657 |
| 17.55141 | 0.05609828 | 0.216595396 | 0.010747833 | 0.09601217 |
| 17.56165 | 0.056031892 | 0.21477446 | 0.006107812 | 0.08786765 |
| 17.57189 | 0.056979829 | 0.21493021 | 0.002108161 | 0.08132062 |
| 17.58213 | 0.058540089 | 0.217420611 | -0.001746655 | 0.07548548 |
| 17.59237 | 0.060523393 | 0.222302165 | -0.004898311 | 0.07127072 |
| 17.60261 | 0.063147603 | 0.228983274 | -0.007836224 | 0.0678198 |
| 17.61285 | 0.066039339 | 0.236946956 | -0.01078936 | 0.06602559 |
| 17.62309 | 0.069029323 | 0.245326597 | -0.014172232 | 0.06442573 |
| 17.63333 | 0.072124377 | 0.253590883 | -0.016621168 | 0.06545991 |
| 17.64357 | 0.074678846 | 0.260551843 | -0.020385177 | 0.06778294 |
| 17.65381 | 0.076485387 | 0.266450441 | -0.022704994 | 0.07282973 |
| 17.66405 | 0.077777021 | 0.270320043 | -0.025317822 | 0.07966893 |
| 17.67429 | 0.077939231 | 0.272446264 | -0.026387845 | 0.0889281 |
| 17.68453 | 0.07746121 | 0.272950032 | -0.027541179 | 0.09924902 |
| 17.69477 | 0.076544405 | 0.271606476 | -0.027257733 | 0.11052785 |
| 17.70501 | 0.075285774 | 0.26864341 | -0.027134225 | 0.12222724 |
| 17.71525 | 0.074517161 | 0.263930906 | -0.026181751 | 0.13373494 |
| 17.72549 | 0.073913725 | 0.258735386 | -0.025874426 | 0.1437707 |
| 17.73573 | 0.074677291 | 0.252340041 | -0.025753657 | 0.15375345 |
| 17.74597 | 0.07560854 | 0.246274771 | -0.02666604 | 0.16189926 |
| 17.75621 | 0.077376581 | 0.240607389 | -0.027662833 | 0.16941136 |
| 17.76645 | 0.079854597 | 0.236276864 | -0.029754181 | 0.17640008 |
| 17.77669 | 0.082478056 | 0.233245684 | -0.032203513 | 0.18188883 |
| 17.78693 | 0.085114752 | 0.232120994 | -0.035017741 | 0.18761671 |
| 17.79717 | 0.087210223 | 0.232370695 | -0.037822743 | 0.19185078 |
| 17.80741 | 0.089479946 | 0.234240899 | -0.04081316 | 0.1953859 |
| 17.81765 | 0.090684183 | 0.237568168 | -0.043230486 | 0.19844615 |
| 17.82789 | 0.091274456 | 0.241126793 | -0.04546959 | 0.20008879 |
| 17.83813 | 0.091472991 | 0.245778736 | -0.046952652 | 0.20205876 |
| 17.84837 | 0.090657232 | 0.249423144 | -0.048019526 | 0.20313464 |
| 17.85861 | 0.089107557 | 0.25293548 | -0.048725228 | 0.20522931 |
| 17.86885 | 0.087534782 | 0.25503622 | -0.048401109 | 0.20769267 |
| 17.87909 | 0.085248116 | 0.25511313 | -0.04827414 | 0.21209498 |
| 17.88933 | 0.083587711 | 0.254058136 | -0.047319122 | 0.21752272 |
| 17.89957 | 0.082119268 | 0.250805319 | -0.046449476 | 0.22507105 |
| 17.90981 | 0.081856554 | 0.246243471 | -0.045191843 | 0.23347407 |
| 17.92005 | 0.082440471 | 0.239677145 | -0.044361062 | 0.24297531 |
| 17.93029 | 0.085194373 | 0.232895597 | -0.04319587 | 0.25261455 |
| 17.94053 | 0.088473718 | 0.224226372 | -0.043109713 | 0.26168917 |
| 17.95077 | 0.093528944 | 0.21527872 | -0.043552499 | 0.27008664 |
| 17.96101 | 0.099279473 | 0.205027737 | -0.045181335 | 0.27724805 |
| 17.97125 | 0.105257356 | 0.194749064 | -0.048108354 | 0.28271089 |
| 17.98149 | 0.111115379 | 0.183533194 | -0.05216164 | 0.28713379 |
| 17.99173 | 0.116145906 | 0.172256177 | -0.057675785 | 0.28934788 |
| 18.00197 | 0.120148927 | 0.160760955 | -0.063609767 | 0.29092722 |
| 18.01221 | 0.123176524 | 0.150051837 | -0.070181414 | 0.2908987 |
| 18.02245 | 0.12482942 | 0.139861931 | -0.076607204 | 0.2899476 |
| 18.03269 | 0.126243918 | 0.13144992 | -0.081998612 | 0.28878328 |
| 18.04293 | 0.127592282 | 0.123919684 | -0.087360582 | 0.28668604 |
| 18.05317 | 0.129230467 | 0.118852462 | -0.090680982 | 0.28510931 |
| 18.06341 | 0.132025427 | 0.114585939 | -0.093428935 | 0.28290365 |
| 18.07365 | 0.136148874 | 0.112131389 | -0.094595681 | 0.2811354 |
| 18.08389 | 0.14178547 | 0.109499851 | -0.095059734 | 0.27879335 |
| 18.09413 | 0.148316533 | 0.106920719 | -0.094435583 | 0.2757221 |
| 18.10437 | 0.155469847 | 0.104289368 | -0.093679675 | 0.27244372 |
| 18.11461 | 0.162528161 | 0.10054843 | -0.092320255 | 0.26768377 |
| 18.12485 | 0.169309966 | 0.09727449 | -0.091020071 | 0.26241789 |
| 18.13509 | 0.174566764 | 0.09370025 | -0.089854647 | 0.25687514 |
| 18.14533 | 0.178203311 | 0.090690834 | -0.088747765 | 0.25121478 |
| 18.15557 | 0.180150062 | 0.087659144 | -0.088368129 | 0.24594814 |
| 18.16581 | 0.180104817 | 0.084883831 | -0.088085873 | 0.24161067 |
| 18.17605 | 0.178315544 | 0.082108581 | -0.088933041 | 0.238635 |
| 18.18629 | 0.174818405 | 0.079070883 | -0.090246301 | 0.23622847 |
| 18.19653 | 0.170245045 | 0.07591214 | -0.092682627 | 0.23529759 |
| 18.20677 | 0.164769647 | 0.072699531 | -0.095663002 | 0.23473702 |
| 18.21701 | 0.158767767 | 0.069486752 | -0.100036002 | 0.23460042 |
| 18.22725 | 0.152713333 | 0.066586621 | -0.104303502 | 0.23450277 |
| 18.23749 | 0.146955928 | 0.063590267 | -0.109625305 | 0.23441018 |
| 18.24773 | 0.141351935 | 0.061633845 | -0.114737459 | 0.23399779 |
| 18.25797 | 0.136586109 | 0.059074554 | -0.120134422 | 0.23337584 |
| 18.26821 | 0.132244192 | 0.057982318 | -0.124870892 | 0.23274486 |
| 18.27845 | 0.1286105 | 0.056492906 | -0.129597106 | 0.23204637 |
| 18.28869 | 0.125781648 | 0.056329228 | -0.133528746 | 0.23161034 |
| 18.29893 | 0.1234663 | 0.055694667 | -0.136760281 | 0.23122677 |
| 18.30917 | 0.122679915 | 0.056776823 | -0.139188939 | 0.23212125 |
| 18.31941 | 0.122597161 | 0.057482062 | -0.140617299 | 0.23236125 |
| 18.32965 | 0.123829875 | 0.059016057 | -0.140595072 | 0.23362015 |
| 18.33989 | 0.126130146 | 0.060566263 | -0.139146738 | 0.23434653 |
| 18.35013 | 0.129341479 | 0.062142918 | -0.13669864 | 0.23451043 |
| 18.36037 | 0.132659073 | 0.0637184 | -0.133071186 | 0.2342592 |
| 18.37061 | 0.135888391 | 0.065642703 | -0.128184875 | 0.23322805 |
| 18.38085 | 0.138353146 | 0.068042012 | -0.122929962 | 0.23209907 |
| 18.39109 | 0.140386812 | 0.070651532 | -0.117854699 | 0.23127128 |
| 18.40133 | 0.140603219 | 0.074019445 | -0.112734702 | 0.23116379 |
| 18.41157 | 0.140372112 | 0.077464644 | -0.108615894 | 0.23238086 |
| 18.42181 | 0.139316211 | 0.080876776 | -0.105265794 | 0.23451457 |
| 18.43205 | 0.137344459 | 0.084267495 | -0.103341575 | 0.23769445 |
| 18.44229 | 0.135690667 | 0.086961064 | -0.101742539 | 0.24068384 |
| 18.45253 | 0.133712543 | 0.089542868 | -0.101390694 | 0.24368403 |
| 18.46277 | 0.131654165 | 0.091729467 | -0.101091918 | 0.2450962 |
| 18.47301 | 0.129789852 | 0.094475307 | -0.100873596 | 0.24623089 |
| 18.48325 | 0.127650799 | 0.097545706 | -0.100146668 | 0.24610386 |
| 18.49349 | 0.125561432 | 0.101218705 | -0.099261255 | 0.24585951 |
| 18.50373 | 0.123593751 | 0.104807149 | -0.097912217 | 0.24494305 |
| 18.51397 | 0.121621517 | 0.109095047 | -0.096600214 | 0.24484251 |
| 18.52421 | 0.119823087 | 0.111444493 | -0.095386105 | 0.24459899 |
| 18.53445 | 0.118724366 | 0.113886554 | -0.094030172 | 0.24478151 |
| 18.54469 | 0.117499049 | 0.113768922 | -0.093875208 | 0.24518409 |
| 18.55493 | 0.117292479 | 0.113274594 | -0.093170202 | 0.24468177 |
| 18.56517 | 0.116695092 | 0.111587697 | -0.092517286 | 0.24317949 |
| 18.57541 | 0.116994722 | 0.108798959 | -0.091156191 | 0.23983562 |
| 18.58565 | 0.116814861 | 0.106180332 | -0.088704186 | 0.23496813 |
| 18.59589 | 0.116976369 | 0.102852304 | -0.084746659 | 0.22693465 |
| 18.60613 | 0.11656203 | 0.099065951 | -0.079442991 | 0.21670385 |
| 18.61637 | 0.115933011 | 0.094740671 | -0.072937136 | 0.20429129 |
| 18.62661 | 0.115010908 | 0.089936738 | -0.065759238 | 0.19028368 |
| 18.63685 | 0.113264346 | 0.084753545 | -0.058273359 | 0.17437295 |
| 18.64709 | 0.111333662 | 0.079357876 | -0.051443739 | 0.15859322 |
| 18.65733 | 0.109086135 | 0.074886256 | -0.044957425 | 0.14204198 |
| 18.66757 | 0.106636753 | 0.070959554 | -0.04002931 | 0.1264165 |
| 18.67781 | 0.104957879 | 0.069049442 | -0.035536373 | 0.11210418 |
| 18.68805 | 0.103433051 | 0.069056918 | -0.032465259 | 0.09915711 |
| 18.69829 | 0.10308693 | 0.070465079 | -0.030031811 | 0.08874473 |
| 18.70853 | 0.104271895 | 0.07306178 | -0.028132244 | 0.08100112 |
| 18.71877 | 0.106160893 | 0.076709056 | -0.02700512 | 0.07626588 |
| 18.72901 | 0.109791767 | 0.080420482 | -0.026179055 | 0.07412776 |
| 18.73925 | 0.11337194 | 0.085374992 | -0.026282041 | 0.07459641 |
| 18.74949 | 0.117919922 | 0.090539559 | -0.02641537 | 0.07667288 |
| 18.75973 | 0.12181813 | 0.098254861 | -0.028062611 | 0.08027991 |
| 18.76997 | 0.124717565 | 0.107233644 | -0.03001518 | 0.08421181 |
| 18.78021 | 0.126515764 | 0.117739451 | -0.032614584 | 0.08907828 |
| 18.79045 | 0.126388854 | 0.128706246 | -0.036368065 | 0.09433013 |
| 18.80069 | 0.124922351 | 0.139936075 | -0.039743974 | 0.10005787 |
| 18.81093 | 0.121198035 | 0.148772823 | -0.043668562 | 0.10656573 |
| 18.82117 | 0.116854208 | 0.155050084 | -0.047236555 | 0.11398939 |
| 18.83142 | 0.110910053 | 0.157595074 | -0.050482134 | 0.12194434 |
| 18.84165 | 0.104900744 | 0.156628616 | -0.053059076 | 0.1303696 |
| 18.85189 | 0.098077105 | 0.151238874 | -0.055509932 | 0.13829647 |
| 18.86213 | 0.091755988 | 0.143819797 | -0.057557147 | 0.14547046 |
| 18.87237 | 0.085215285 | 0.133617836 | -0.059482326 | 0.15161971 |
| 18.88261 | 0.079076107 | 0.123186428 | -0.061797924 | 0.15585265 |
| 18.89285 | 0.072362019 | 0.113359342 | -0.063918776 | 0.1583038 |
| 18.9031 | 0.066560314 | 0.104980908 | -0.066598904 | 0.15925589 |
| 18.91333 | 0.060249165 | 0.098960924 | -0.068729382 | 0.15803795 |
| 18.92357 | 0.054154461 | 0.096164365 | -0.071017849 | 0.15572201 |
| 18.93382 | 0.049239067 | 0.095845037 | -0.073135984 | 0.15290549 |
| 18.94406 | 0.043900981 | 0.098519913 | -0.074603232 | 0.14890068 |
| 18.95429 | 0.040748192 | 0.103630735 | -0.076032072 | 0.14558188 |
| 18.96453 | 0.037482298 | 0.11064749 | -0.07727755 | 0.1430316 |
| 18.97478 | 0.03663669 | 0.118580125 | -0.078600024 | 0.14183593 |
| 18.98501 | 0.035705208 | 0.127687986 | -0.080001306 | 0.14119653 |
| 18.99525 | 0.036795851 | 0.136026766 | -0.081103926 | 0.1417651 |
| 19.0055 | 0.038087446 | 0.143774682 | -0.082593115 | 0.14187774 |
| 19.01574 | 0.040702873 | 0.150634174 | -0.083095309 | 0.14227823 |
| 19.02597 | 0.043380791 | 0.157030166 | -0.083583701 | 0.14261416 |
| 19.03621 | 0.047387011 | 0.162714576 | -0.083224608 | 0.14204677 |
| 19.04646 | 0.051455035 | 0.169481143 | -0.082332428 | 0.14269367 |
| 19.05669 | 0.05678926 | 0.176243316 | -0.081414698 | 0.14385549 |
| 19.06693 | 0.062286763 | 0.184251333 | -0.079701307 | 0.14696988 |
| 19.07718 | 0.068210461 | 0.192529975 | -0.078992608 | 0.15159415 |
| 19.08742 | 0.074799968 | 0.200890534 | -0.077903508 | 0.15804013 |
| 19.09765 | 0.080734212 | 0.208525027 | -0.07736307 | 0.16532687 |
| 19.10789 | 0.087762856 | 0.215293518 | -0.076650853 | 0.17363628 |
| 19.11814 | 0.093327684 | 0.22064453 | -0.075843236 | 0.18115892 |
| 19.12838 | 0.099978343 | 0.223948038 | -0.074565843 | 0.18846203 |
| 19.13861 | 0.105082036 | 0.225230014 | -0.072399963 | 0.19446196 |
| 19.14886 | 0.110812564 | 0.224171619 | -0.069452723 | 0.19909935 |
| 19.1591 | 0.115160607 | 0.221161312 | -0.065834895 | 0.20214957 |
| 19.16933 | 0.119177709 | 0.21550032 | -0.061117373 | 0.20431546 |
| 19.17957 | 0.121830041 | 0.208552447 | -0.056774259 | 0.20514968 |
| 19.18982 | 0.122926739 | 0.199758843 | -0.052178805 | 0.20587895 |

| | | | | |
|---|---|---|---|---|
| 16.79365 | 0.127409285 | 0.144987735 | -0.003751971 | 0.242934208 |
| 16.80389 | 0.134185696 | 0.137691725 | -0.000364312 | 0.244394486 |
| 16.81413 | 0.139994628 | 0.133314126 | 0.001183643 | 0.24596816 |
| 16.82437 | 0.144508771 | 0.133080642 | -0.000310719 | 0.248395551 |
| 16.83461 | 0.147649556 | 0.135270486 | -0.003480066 | 0.249948081 |
| 16.84485 | 0.149186262 | 0.140264941 | -0.009857813 | 0.251917645 |
| 16.85509 | 0.148849165 | 0.146083078 | -0.017474606 | 0.252394022 |
| 16.86533 | 0.146357242 | 0.152801827 | -0.027208297 | 0.252331899 |
| 16.87557 | 0.141902698 | 0.159089603 | -0.037208915 | 0.251338428 |
| 16.88581 | 0.134825426 | 0.164055272 | -0.047920301 | 0.24911017 |
| 16.89605 | 0.125755862 | 0.168253377 | -0.057631261 | 0.246600235 |
| 16.90629 | 0.114897498 | 0.170579336 | -0.066983334 | 0.242979386 |
| 16.91653 | 0.102638631 | 0.17155704 | -0.074374974 | 0.239842695 |
| 16.92677 | 0.089948223 | 0.17116827 | -0.080941824 | 0.235717262 |
| 16.93701 | 0.077530353 | 0.169971741 | -0.085509081 | 0.232550598 |
| 16.94725 | 0.064707786 | 0.168053615 | -0.089509373 | 0.228874708 |
| 16.95749 | 0.053946159 | 0.165688809 | -0.092634787 | 0.226406785 |
| 16.96773 | 0.043430528 | 0.163269471 | -0.095599843 | 0.224058594 |
| 16.97797 | 0.034981071 | 0.160623831 | -0.098849641 | 0.223158145 |
| 16.98821 | 0.027408048 | 0.157493923 | -0.102993152 | 0.223039169 |
| 16.99845 | 0.021653325 | 0.153509968 | -0.107389584 | 0.224386725 |
| 17.00869 | 0.01763705 | 0.148629139 | -0.112692968 | 0.226596142 |
| 17.01893 | 0.014960199 | 0.142975685 | -0.11821897 | 0.230141943 |
| 17.02917 | 0.013902479 | 0.136735967 | -0.123261687 | 0.234593661 |
| 17.03941 | 0.014087061 | 0.130995879 | -0.128109184 | 0.238728986 |
| 17.04965 | 0.015680811 | 0.125776543 | -0.131579014 | 0.243583442 |
| 17.05989 | 0.017847201 | 0.123020219 | -0.134298541 | 0.247496374 |
| 17.07013 | 0.020186099 | 0.121892913 | -0.135493779 | 0.250899547 |
| 17.08037 | 0.02258313 | 0.123002503 | -0.136012148 | 0.252882262 |
| 17.09061 | 0.024512174 | 0.125245708 | -0.135595647 | 0.25412493 |
| 17.10085 | 0.025304401 | 0.128258607 | -0.135006607 | 0.253814639 |
| 17.11109 | 0.025456746 | 0.131048959 | -0.134125744 | 0.252434926 |
| 17.12133 | 0.02371996 | 0.133414579 | -0.133925609 | 0.249607119 |
| 17.13157 | 0.021756567 | 0.134690213 | -0.133836831 | 0.245303872 |
| 17.14181 | 0.018048306 | 0.135011214 | -0.134576394 | 0.239935086 |
| 17.15205 | 0.013826044 | 0.134579156 | -0.136494614 | 0.232466897 |
| 17.16229 | 0.009497806 | 0.132771222 | -0.138805383 | 0.224503279 |
| 17.17253 | 0.00480222 | 0.130615588 | -0.142318589 | 0.214511334 |
| 17.18277 | 0.000610855 | 0.127599389 | -0.146131717 | 0.204413721 |
| 17.19301 | -0.002706831 | 0.125222619 | -0.150595057 | 0.193347934 |
| 17.20325 | -0.005003231 | 0.12313463 | -0.154394406 | 0.182520193 |
| 17.21349 | -0.00610978 | 0.122443282 | -0.157706014 | 0.172183922 |
| 17.22373 | -0.006208883 | 0.124227676 | -0.159896617 | 0.162287814 |
| 17.23397 | -0.005050521 | 0.12745896 | -0.160864252 | 0.153396841 |
| 17.24421 | -0.003360766 | 0.132492547 | -0.159920842 | 0.145652018 |
| 17.25445 | -0.001637793 | 0.13867128 | -0.157707053 | 0.13909812 |
| 17.26469 | 0.000348621 | 0.145496465 | -0.154575052 | 0.133337067 |
| 17.27493 | 0.001764003 | 0.151695277 | -0.15022358 | 0.129152383 |
| 17.28517 | 0.00281316 | 0.157157922 | -0.1456186 | 0.125144381 |
| 17.29541 | 0.002975284 | 0.160684847 | -0.141019836 | 0.121415531 |
| 17.30565 | 0.002739515 | 0.162703188 | -0.136584233 | 0.117451069 |
| 17.31589 | 0.002141019 | 0.162955196 | -0.132933216 | 0.112664049 |
| 17.32613 | 0.000813039 | 0.161538403 | -0.130604188 | 0.10718857 |
| 17.33637 | -0.000571938 | 0.158978158 | -0.128830753 | 0.100621009 |
| 17.34661 | -0.002047142 | 0.155584243 | -0.128802598 | 0.093602907 |
| 17.35685 | -0.00391462 | 0.152785792 | -0.129601547 | 0.086875052 |
| 17.36709 | -0.005753131 | 0.149208008 | -0.131749628 | 0.080588839 |
| 17.37733 | -0.007579374 | 0.146161783 | -0.133930293 | 0.075516793 |
| 17.38757 | -0.009687382 | 0.143656056 | -0.136659992 | 0.072658924 |
| 17.39781 | -0.011794528 | 0.140908205 | -0.139418482 | 0.071498948 |
| 17.40805 | -0.013819781 | 0.138827312 | -0.141302684 | 0.071798894 |
| 17.41829 | -0.015890318 | 0.136822858 | -0.142975115 | 0.073930251 |
| 17.42853 | -0.017794611 | 0.134893687 | -0.143252091 | 0.07637225 |
| 17.43877 | -0.019480729 | 0.133620386 | -0.142701559 | 0.079642939 |
| 17.44901 | -0.021255361 | 0.131922898 | -0.140659633 | 0.082391687 |
| 17.45925 | -0.022305799 | 0.129848948 | -0.137455663 | 0.085332823 |
| 17.46949 | -0.023748817 | 0.127833001 | -0.132894884 | 0.086659676 |
| 17.47973 | -0.024972025 | 0.125059077 | -0.127044288 | 0.087537757 |
| 17.48997 | -0.026670027 | 0.122076049 | -0.120161034 | 0.086883011 |
| 17.50021 | -0.028698078 | 0.12013885 | -0.112602608 | 0.085004989 |
| 17.51045 | -0.031245011 | 0.118765771 | -0.105090381 | 0.081984473 |
| 17.52069 | -0.034948169 | 0.119235343 | -0.09703682 | 0.078676313 |
| 17.53093 | -0.03916294 | 0.121656445 | -0.090254725 | 0.07529823 |
| 17.54117 | -0.043944242 | 0.125817689 | -0.083880132 | 0.072432194 |
| 17.55141 | -0.049374003 | 0.131335642 | -0.078348555 | 0.071855245 |
| 17.56165 | -0.05517671 | 0.137316537 | -0.07403783 | 0.073174047 |
| 17.57189 | -0.060324851 | 0.143676493 | -0.070726111 | 0.0773056 |
| 17.58213 | -0.065594606 | 0.149022265 | -0.068177407 | 0.083240354 |
| 17.59237 | -0.069918831 | 0.153862724 | -0.06672389 | 0.091465869 |
| 17.60261 | -0.073553136 | 0.156970215 | -0.066311652 | 0.09994621 |
| 17.61285 | -0.07673709 | 0.158990563 | -0.066397157 | 0.108515324 |
| 17.62309 | -0.078811069 | 0.159133389 | -0.067076354 | 0.116265246 |
| 17.63333 | -0.081283558 | 0.158331403 | -0.068244529 | 0.121894359 |
| 17.64357 | -0.082582266 | 0.156076779 | -0.069194781 | 0.125905431 |
| 17.65381 | -0.084714155 | 0.153486373 | -0.070041369 | 0.127219331 |
| 17.66405 | -0.086982776 | 0.150188312 | -0.070120746 | 0.126482899 |
| 17.67429 | -0.089118715 | 0.146970342 | -0.069580047 | 0.123434454 |
| 17.68453 | -0.09228663 | 0.143431711 | -0.068357805 | 0.119494841 |
| 17.69477 | -0.094960235 | 0.139913586 | -0.066236436 | 0.114176447 |
| 17.70501 | -0.098056424 | 0.136040321 | -0.063953373 | 0.109204793 |
| 17.71525 | -0.100321329 | 0.132121563 | -0.061362533 | 0.104713282 |
| 17.72549 | -0.102546327 | 0.127307802 | -0.059098887 | 0.100898934 |
| 17.73573 | -0.103811962 | 0.122149188 | -0.057245619 | 0.098867007 |
| 17.74597 | -0.104194544 | 0.116696604 | -0.056273666 | 0.097509676 |
| 17.75621 | -0.104194259 | 0.111296039 | -0.056133334 | 0.097999851 |
| 17.76645 | -0.103609875 | 0.10587779 | -0.056712326 | 0.099499453 |
| 17.77669 | -0.102655566 | 0.101768377 | -0.05830991 | 0.10258824 |
| 17.78693 | -0.101024194 | 0.098363109 | -0.060935627 | 0.107296097 |
| 17.79717 | -0.100032734 | 0.095911888 | -0.063628148 | 0.113768392 |
| 17.80741 | -0.097939389 | 0.094313184 | -0.067513161 | 0.120695434 |
| 17.81765 | -0.096315825 | 0.092191999 | -0.071478774 | 0.128808061 |
| 17.82789 | -0.094449875 | 0.089832325 | -0.076095809 | 0.13566871 |
| 17.83813 | -0.092448682 | 0.086692378 | -0.080246615 | 0.142249566 |
| 17.84837 | -0.090604285 | 0.08232016 | -0.085002444 | 0.146971893 |
| 17.85861 | -0.089131113 | 0.078268684 | -0.089223742 | 0.150163015 |
| 17.86885 | -0.08767182 | 0.073968108 | -0.093048216 | 0.151645062 |
| 17.87909 | -0.086476088 | 0.071506246 | -0.096467157 | 0.151844661 |
| 17.88933 | -0.085389758 | 0.071030823 | -0.099305612 | 0.151286476 |
| 17.89957 | -0.084016168 | 0.073487545 | -0.101241403 | 0.150328695 |
| 17.90981 | -0.082128276 | 0.077431953 | -0.102326344 | 0.149139813 |
| 17.92005 | -0.079939209 | 0.083660743 | -0.102261937 | 0.148455757 |
| 17.93029 | -0.07718766 | 0.090013175 | -0.10109124 | 0.148445894 |
| 17.94053 | -0.073760115 | 0.096674076 | -0.098898513 | 0.148616014 |
| 17.95077 | -0.070543871 | 0.103113215 | -0.095719607 | 0.149648382 |
| 17.96101 | -0.066889635 | 0.10798893 | -0.092055071 | 0.150763447 |
| 17.97125 | -0.064149959 | 0.113131622 | -0.088205084 | 0.152968772 |
| 17.98149 | -0.062026106 | 0.116473025 | -0.084647814 | 0.155158105 |
| 17.99173 | -0.060188215 | 0.11972738 | -0.08185323 | 0.158632331 |
| 18.00197 | -0.059720293 | 0.122627506 | -0.07974946 | 0.163084706 |
| 18.01221 | -0.059184724 | 0.125625207 | -0.078204115 | 0.169248597 |
| 18.02245 | -0.0591352 | 0.128974705 | -0.077734071 | 0.176359944 |
| 18.03269 | -0.058472894 | 0.132237036 | -0.076717945 | 0.18475207 |
| 18.04293 | -0.056767466 | 0.136306685 | -0.075915622 | 0.193611586 |
| 18.05317 | -0.054104979 | 0.139667533 | -0.07403055 | 0.202561638 |
| 18.06341 | -0.049763942 | 0.142871193 | -0.071323636 | 0.210539468 |
| 18.07365 | -0.043220185 | 0.145156911 | -0.067205979 | 0.217426548 |
| 18.08389 | -0.035834409 | 0.146656681 | -0.061901202 | 0.222305549 |
| 18.09413 | -0.026622025 | 0.146718231 | -0.055690415 | 0.225565061 |
| 18.10437 | -0.016849575 | 0.146498763 | -0.04856839 | 0.226614496 |
| 18.11461 | -0.006584085 | 0.145124872 | -0.041509112 | 0.226172744 |
| 18.12485 | 0.003674944 | 0.143317384 | -0.034660227 | 0.223768518 |
| 18.13509 | 0.013792415 | 0.14190981 | -0.028713013 | 0.220504674 |
| 18.14533 | 0.023180484 | 0.139426016 | -0.023807733 | 0.215957242 |
| 18.15557 | 0.032129549 | 0.13750521 | -0.020197136 | 0.211181527 |
| 18.16581 | 0.040118798 | 0.135046466 | -0.018046465 | 0.206272166 |
| 18.17605 | 0.04717428 | 0.13240207 | -0.016242452 | 0.202154032 |
| 18.18629 | 0.053551368 | 0.13001676 | -0.015561852 | 0.198494817 |
| 18.19653 | 0.058021839 | 0.128037712 | -0.014640803 | 0.197167524 |
| 18.20677 | 0.061483888 | 0.127486232 | -0.013749217 | 0.196437953 |
| 18.21701 | 0.063188343 | 0.128875724 | -0.013004464 | 0.197828484 |
| 18.22725 | 0.063389487 | 0.13259164 | -0.011597097 | 0.200295631 |
| 18.23749 | 0.061913317 | 0.138138328 | -0.010884868 | 0.203575007 |
| 18.24773 | 0.05929668 | 0.145693103 | -0.010449523 | 0.208108819 |
| 18.25797 | 0.056064545 | 0.153350808 | -0.01126578 | 0.212560674 |
| 18.26821 | 0.052597352 | 0.161239345 | -0.012861385 | 0.218383305 |
| 18.27845 | 0.049199572 | 0.168295572 | -0.015865523 | 0.223929688 |
| 18.28869 | 0.047081137 | 0.173895512 | -0.01993185 | 0.230208376 |
| 18.29893 | 0.045722412 | 0.178153033 | -0.024729816 | 0.236190431 |
| 18.30917 | 0.045574418 | 0.181238576 | -0.030443789 | 0.241867237 |
| 18.31941 | 0.04660733 | 0.182958042 | -0.035416588 | 0.247051276 |
| 18.32965 | 0.048361712 | 0.184102837 | -0.040957644 | 0.25113251 |
| 18.33989 | 0.049876074 | 0.184576416 | -0.045410022 | 0.254633714 |
| 18.35013 | 0.051381168 | 0.184720918 | -0.049307601 | 0.256836196 |
| 18.36037 | 0.051842915 | 0.185311069 | -0.052572378 | 0.2591238 |
| 18.37061 | 0.051557916 | 0.185874845 | -0.055554791 | 0.260627747 |
| 18.38085 | 0.04939235 | 0.18745201 | -0.058395727 | 0.262658022 |
| 18.39109 | 0.046527193 | 0.189358854 | -0.061243354 | 0.265306623 |
| 18.40133 | 0.043315627 | 0.192144624 | -0.064472629 | 0.268919912 |
| 18.41157 | 0.038482447 | 0.196096302 | -0.068263071 | 0.273033773 |
| 18.42181 | 0.034446751 | 0.200056074 | -0.072521069 | 0.277695038 |
| 18.43205 | 0.029965357 | 0.204552872 | -0.076643965 | 0.282629301 |
| 18.44229 | 0.025725865 | 0.209334226 | -0.08138506 | 0.287046607 |
| 18.45253 | 0.021525954 | 0.214162741 | -0.085422733 | 0.290492751 |
| 18.46277 | 0.01714002 | 0.218401306 | -0.089548584 | 0.292194546 |
| 18.47301 | 0.012538823 | 0.223344964 | -0.092444628 | 0.29246875 |
| 18.48325 | 0.007485049 | 0.227329686 | -0.095322827 | 0.290358267 |
| 18.49349 | 0.001773103 | 0.231617302 | -0.09721972 | 0.287604759 |
| 18.50373 | -0.004408191 | 0.23491418 | -0.098928296 | 0.282741739 |
| 18.51397 | -0.010589598 | 0.237800717 | -0.099794215 | 0.278186269 |
| 18.52421 | -0.017411993 | 0.239486525 | -0.100873687 | 0.272552239 |
| 18.53445 | -0.023308676 | 0.240373004 | -0.101647756 | 0.267383264 |
| 18.54469 | -0.029285969 | 0.239731378 | -0.102348383 | 0.262154513 |
| 18.55493 | -0.034620921 | 0.238011495 | -0.10297497 | 0.25669939 |
| 18.56517 | -0.039538074 | 0.235595979 | -0.103537 | 0.251879664 |
| 18.57541 | -0.043940241 | 0.232203708 | -0.104082964 | 0.246672514 |
| 18.58565 | -0.048561071 | 0.228658083 | -0.104017797 | 0.241732044 |
| 18.59589 | -0.052781577 | 0.224585772 | -0.104050063 | 0.236386204 |
| 18.60613 | -0.057274036 | 0.220188951 | -0.103086935 | 0.231504973 |
| 18.61637 | -0.061443932 | 0.21588282 | -0.102132921 | 0.225528387 |
| 18.62661 | -0.066144653 | 0.212018961 | -0.100076875 | 0.219805393 |
| 18.63685 | -0.070204695 | 0.208089474 | -0.097529245 | 0.213057529 |
| 18.64709 | -0.074152514 | 0.205330403 | -0.094608765 | 0.206123472 |
| 18.65733 | -0.078339948 | 0.203605829 | -0.090849764 | 0.198280732 |
| 18.66757 | -0.081778002 | 0.202961047 | -0.087032561 | 0.190653052 |
| 18.67781 | -0.08562997 | 0.203237896 | -0.083010898 | 0.182644973 |
| 18.68805 | -0.089319781 | 0.204088023 | -0.079156312 | 0.1757111 |
| 18.69829 | -0.092936157 | 0.205185656 | -0.07540777 | 0.169926879 |
| 18.70853 | -0.096527211 | 0.205862802 | -0.072850447 | 0.165757811 |
| 18.71877 | -0.09993305 | 0.205821597 | -0.070007043 | 0.164480236 |
| 18.72901 | -0.102884096 | 0.205016127 | -0.069223409 | 0.165453978 |
| 18.73925 | -0.105315126 | 0.202873683 | -0.068669555 | 0.168804049 |
| 18.74949 | -0.10680207 | 0.200147155 | -0.069864788 | 0.173970912 |
| 18.75973 | -0.107543886 | 0.19636209 | -0.071926704 | 0.180615232 |
| 18.76997 | -0.107812632 | 0.192257118 | -0.075661398 | 0.18727096 |
| 18.78021 | -0.107463948 | 0.188332359 | -0.080114704 | 0.194034733 |
| 18.79045 | -0.106484832 | 0.18496662 | -0.085965085 | 0.199997965 |
| 18.80069 | -0.105890552 | 0.183233637 | -0.091561062 | 0.204765766 |
| 18.81093 | -0.105852682 | 0.182405305 | -0.097943584 | 0.209179159 |
| 18.82117 | -0.105986765 | 0.183914076 | -0.103340559 | 0.213596281 |
| 18.83142 | -0.107409586 | 0.186343265 | -0.108182705 | 0.218481741 |
| 18.84165 | -0.109703269 | 0.190607087 | -0.111874969 | 0.223827306 |
| 18.85189 | -0.1130838 | 0.195218546 | -0.114173768 | 0.230767727 |
| 18.86213 | -0.117307994 | 0.200320252 | -0.115021158 | 0.237118988 |
| 18.87237 | -0.121791926 | 0.204681562 | -0.113935068 | 0.24444163 |
| 18.88261 | -0.126921338 | 0.208361852 | -0.111994391 | 0.249613212 |
| 18.89285 | -0.130821512 | 0.209543033 | -0.107494692 | 0.253713071 |
| 18.9031 | -0.134186556 | 0.208935148 | -0.10208206 | 0.25533511 |
| 18.91333 | -0.135829184 | 0.205331626 | -0.094396571 | 0.254774699 |
| 18.92357 | -0.136287293 | 0.199772819 | -0.086276863 | 0.251966052 |
| 18.93382 | -0.134884331 | 0.191752571 | -0.07646996 | 0.24795351 |
| 18.94406 | -0.132230817 | 0.182363968 | -0.066547873 | 0.242916169 |
| 18.95429 | -0.129054278 | 0.172278332 | -0.056059948 | 0.237670958 |
| 18.96453 | -0.125362234 | 0.161880508 | -0.046389733 | 0.233071714 |
| 18.97478 | -0.121803288 | 0.152041437 | -0.036866803 | 0.229488939 |
| 18.98501 | -0.119171836 | 0.142735473 | -0.028653705 | 0.22681538 |
| 18.99525 | -0.116979244 | 0.134420893 | -0.020932562 | 0.225046675 |
| 19.0055 | -0.115248123 | 0.127030444 | -0.014694891 | 0.224491637 |
| 19.01574 | -0.11420255 | 0.120977928 | -0.009120353 | 0.224448752 |
| 19.02597 | -0.112787701 | 0.116273943 | -0.004387258 | 0.224953135 |
| 19.03621 | -0.112044013 | 0.113220673 | -0.000591358 | 0.226155623 |
| 19.04646 | -0.110018905 | 0.113247312 | 0.002547762 | 0.226896021 |
| 19.05669 | -0.108297513 | 0.115024382 | 0.004718567 | 0.228245038 |
| 19.06693 | -0.10564559 | 0.119951394 | 0.006769489 | 0.228338669 |
| 19.07718 | -0.103290133 | 0.125924202 | 0.007896836 | 0.228656089 |
| 19.08742 | -0.100165952 | 0.133124566 | 0.00855484 | 0.226980439 |
| 19.09765 | -0.097436989 | 0.139799329 | 0.009518009 | 0.22481483 |
| 19.10789 | -0.094988471 | 0.14647276 | 0.009376358 | 0.221016191 |
| 19.11814 | -0.092807369 | 0.15183336 | 0.00968539 | 0.216111724 |
| 19.12838 | -0.09146178 | 0.156674524 | 0.00920553 | 0.209692992 |
| 19.13861 | -0.091204717 | 0.160726329 | 0.008374418 | 0.202244334 |
| 19.14886 | -0.091967908 | 0.164041456 | 0.006756106 | 0.193212294 |
| 19.1591 | -0.094068046 | 0.167613145 | 0.004734818 | 0.1833112 |
| 19.16933 | -0.097491054 | 0.170681567 | 0.002106223 | 0.172489146 |
| 19.17957 | -0.102476249 | 0.173968725 | -0.001081052 | 0.161386382 |
| 19.18982 | -0.108623482 | 0.177720464 | -0.004110878 | 0.150984826 |

| | | | | |
|---|---|---|---|---|
| 16.79365 | 0.026036794 | 0.179697817 | 0.041914234 | 0.161146822 |
| 16.80389 | 0.026948695 | 0.179969364 | 0.038880584 | 0.153423621 |
| 16.81413 | 0.027970013 | 0.180680834 | 0.034471388 | 0.146967792 |
| 16.82437 | 0.029230633 | 0.181430009 | 0.030726438 | 0.14068965 |
| 16.83461 | 0.030874343 | 0.18190158 | 0.025857554 | 0.135801087 |
| 16.84485 | 0.032215811 | 0.183359395 | 0.021221609 | 0.132064703 |
| 16.85509 | 0.033936981 | 0.184234259 | 0.017168584 | 0.129676894 |
| 16.86533 | 0.035476383 | 0.18578266 | 0.01344868 | 0.128811827 |
| 16.87557 | 0.037380603 | 0.187275988 | 0.010855161 | 0.129557318 |
| 16.88581 | 0.039241873 | 0.188584963 | 0.008871968 | 0.131785079 |
| 16.89605 | 0.041146018 | 0.189602752 | 0.007946099 | 0.135032282 |
| 16.90629 | 0.042728076 | 0.190291371 | 0.007475984 | 0.139199441 |
| 16.91653 | 0.044315654 | 0.190396737 | 0.007167382 | 0.143658836 |
| 16.92677 | 0.045322745 | 0.190112549 | 0.007220996 | 0.147928463 |
| 16.93701 | 0.046217325 | 0.189260787 | 0.006907586 | 0.151862389 |
| 16.94725 | 0.046270849 | 0.187812004 | 0.006132323 | 0.154999489 |
| 16.95749 | 0.046171035 | 0.186400548 | 0.005170074 | 0.157218617 |
| 16.96773 | 0.045557805 | 0.18378122 | 0.004053124 | 0.158921537 |
| 16.97797 | 0.044484035 | 0.181416289 | 0.002562135 | 0.15961694 |
| 16.98821 | 0.04285709 | 0.178506211 | 0.001184535 | 0.159439598 |
| 16.99845 | 0.041079804 | 0.175631432 | 0.000155535 | 0.158938278 |
| 17.00869 | 0.038895357 | 0.172244013 | -0.000847399 | 0.157577448 |
| 17.01893 | 0.036949401 | 0.169390352 | -0.001173223 | 0.155801565 |
| 17.02917 | 0.03492516 | 0.165767016 | -0.001235392 | 0.153734791 |
| 17.03941 | 0.033417862 | 0.162530619 | -0.000564245 | 0.151381047 |
| 17.04965 | 0.032457599 | 0.158772051 | 0.000322712 | 0.149291302 |
| 17.05989 | 0.031970818 | 0.155376593 | 0.001937679 | 0.1471536 |
| 17.07013 | 0.031627699 | 0.152306431 | 0.003572023 | 0.145491348 |
| 17.08037 | 0.031830781 | 0.149664045 | 0.005300055 | 0.144184874 |
| 17.09061 | 0.031420616 | 0.147852919 | 0.007194263 | 0.143387721 |
| 17.10085 | 0.031164851 | 0.147218789 | 0.008596324 | 0.14312516 |
| 17.11109 | 0.029783806 | 0.14728273 | 0.010009638 | 0.143310253 |
| 17.12133 | 0.028208015 | 0.148269286 | 0.010749748 | 0.144399974 |
| 17.13157 | 0.026401652 | 0.149646536 | 0.011163007 | 0.145454084 |
| 17.14181 | 0.024192635 | 0.151217863 | 0.011226264 | 0.147562368 |
| 17.15205 | 0.022431641 | 0.152447336 | 0.010478911 | 0.149624957 |
| 17.16229 | 0.021519056 | 0.153359881 | 0.009475361 | 0.151924489 |
| 17.17253 | 0.02117487 | 0.153619995 | 0.007625788 | 0.154121895 |
| 17.18277 | 0.022651512 | 0.153103037 | 0.005317416 | 0.156088645 |
| 17.19301 | 0.024991094 | 0.152817634 | 0.002250403 | 0.157958912 |
| 17.20325 | 0.028884155 | 0.152088245 | -0.000975415 | 0.158894662 |
| 17.21349 | 0.033579964 | 0.152344285 | -0.004342004 | 0.160051566 |
| 17.22373 | 0.039558629 | 0.153404829 | -0.007771744 | 0.160362008 |
| 17.23397 | 0.04537784 | 0.155984241 | -0.010706112 | 0.16039895 |
| 17.24421 | 0.051298696 | 0.159380741 | -0.012701429 | 0.160120373 |
| 17.25445 | 0.056860983 | 0.163426897 | -0.014607809 | 0.159624234 |
| 17.26469 | 0.061591959 | 0.167200242 | -0.015424745 | 0.158834519 |
| 17.27493 | 0.065265339 | 0.169616671 | -0.015904104 | 0.157546408 |
| 17.28517 | 0.06790806 | 0.170909597 | -0.01609122 | 0.156736691 |
| 17.29541 | 0.070011299 | 0.170756467 | -0.016520992 | 0.155507429 |
| 17.30565 | 0.071270156 | 0.169796648 | -0.016998378 | 0.154477025 |
| 17.31589 | 0.072203818 | 0.168289559 | -0.017836604 | 0.153837173 |
| 17.32613 | 0.07342482 | 0.167128326 | -0.019413827 | 0.153548854 |
| 17.33637 | 0.07487166 | 0.166997722 | -0.02128258 | 0.153850725 |
| 17.34661 | 0.076279687 | 0.167251241 | -0.023536385 | 0.154219222 |
| 17.35685 | 0.078167761 | 0.168365297 | -0.02656044 | 0.155284261 |
| 17.36709 | 0.079530367 | 0.169759817 | -0.029783933 | 0.155794662 |
| 17.37733 | 0.080381881 | 0.171099091 | -0.033216326 | 0.156402837 |
| 17.38757 | 0.080244304 | 0.171890637 | -0.03701737 | 0.155796673 |
| 17.39781 | 0.078706191 | 0.172326999 | -0.040668499 | 0.154686993 |
| 17.40805 | 0.076290736 | 0.172109906 | -0.044393178 | 0.151867185 |
| 17.41829 | 0.072502507 | 0.172146803 | -0.047762248 | 0.148543953 |
| 17.42853 | 0.068557982 | 0.17183714 | -0.050830712 | 0.143684238 |
| 17.43877 | 0.064298383 | 0.172701986 | -0.053943882 | 0.138749589 |
| 17.44901 | 0.060939617 | 0.174092731 | -0.056348204 | 0.132932868 |
| 17.45925 | 0.058212184 | 0.176744267 | -0.059184731 | 0.1273092 |
| 17.46949 | 0.05686518 | 0.179991774 | -0.061350253 | 0.122154684 |
| 17.47973 | 0.056271579 | 0.18421665 | -0.063604759 | 0.118083147 |
| 17.48997 | 0.056802569 | 0.188311687 | -0.065532845 | 0.115180877 |
| 17.50021 | 0.057785461 | 0.19170692 | -0.06717817 | 0.114040817 |
| 17.51045 | 0.059194397 | 0.194289029 | -0.068385412 | 0.114532801 |
| 17.52069 | 0.060238019 | 0.194764195 | -0.069010134 | 0.116725308 |
| 17.53093 | 0.061661502 | 0.193860701 | -0.069189303 | 0.120336476 |
| 17.54117 | 0.062499882 | 0.190593336 | -0.068888123 | 0.125416318 |
| 17.55141 | 0.062904519 | 0.185614006 | -0.068085583 | 0.130923921 |
| 17.56165 | 0.063090021 | 0.179409917 | -0.067249747 | 0.138068247 |
| 17.57189 | 0.062921654 | 0.172331225 | -0.065913429 | 0.144216204 |
| 17.58213 | 0.062197464 | 0.165014353 | -0.064803655 | 0.151654795 |
| 17.59237 | 0.061234074 | 0.157845647 | -0.063131181 | 0.157930195 |
| 17.60261 | 0.060647571 | 0.151063435 | -0.061705869 | 0.164230988 |
| 17.61285 | 0.060048059 | 0.145405551 | -0.059642427 | 0.169499058 |
| 17.62309 | 0.060328988 | 0.14054761 | -0.057383257 | 0.174457931 |
| 17.63333 | 0.06120068 | 0.136761151 | -0.054063499 | 0.178003209 |
| 17.64357 | 0.062725343 | 0.134322511 | -0.050225415 | 0.180753642 |
| 17.65381 | 0.065068282 | 0.13278606 | -0.04533546 | 0.182529056 |
| 17.66405 | 0.067147941 | 0.132324074 | -0.039761483 | 0.183331269 |
| 17.67429 | 0.069433922 | 0.132541508 | -0.033309539 | 0.183111424 |
| 17.68453 | 0.070757983 | 0.133016802 | -0.026665225 | 0.181973962 |
| 17.69477 | 0.07156836 | 0.133684292 | -0.020085551 | 0.179756357 |
| 17.70501 | 0.071278355 | 0.134485899 | -0.013509635 | 0.176633145 |
| 17.71525 | 0.07069489 | 0.135794648 | -0.007727731 | 0.172632968 |
| 17.72549 | 0.068841118 | 0.137549522 | -0.002261098 | 0.167810181 |
| 17.73573 | 0.067373956 | 0.140780342 | 0.002427895 | 0.162808679 |
| 17.74597 | 0.06551901 | 0.144510482 | 0.006831585 | 0.157465118 |
| 17.75621 | 0.063372054 | 0.149866647 | 0.011220403 | 0.152422879 |
| 17.76645 | 0.061536763 | 0.155278208 | 0.015468948 | 0.147799869 |
| 17.77669 | 0.059830067 | 0.16100433 | 0.019743479 | 0.144209828 |
| 17.78693 | 0.057740936 | 0.166630613 | 0.02426966 | 0.140448484 |
| 17.79717 | 0.056050093 | 0.171814718 | 0.02895093 | 0.137264446 |
| 17.80741 | 0.054365842 | 0.176120113 | 0.033486998 | 0.133954826 |
| 17.81765 | 0.052594257 | 0.179625931 | 0.038257918 | 0.130435303 |
| 17.82789 | 0.051094382 | 0.182197327 | 0.04284342 | 0.126739928 |
| 17.83813 | 0.049533762 | 0.183565462 | 0.047307133 | 0.122958148 |
| 17.84837 | 0.047908393 | 0.184096594 | 0.051353602 | 0.119388864 |
| 17.85861 | 0.046017814 | 0.183123737 | 0.055022589 | 0.116284412 |
| 17.86885 | 0.043990313 | 0.18128595 | 0.057515504 | 0.113603469 |
| 17.87909 | 0.041522931 | 0.178098107 | 0.059403909 | 0.111605084 |
| 17.88933 | 0.038476262 | 0.173592633 | 0.059611276 | 0.11011755 |
| 17.89957 | 0.035275061 | 0.168770995 | 0.058962115 | 0.108976777 |
| 17.90981 | 0.031299384 | 0.163294302 | 0.056990847 | 0.108084612 |
| 17.92005 | 0.027343718 | 0.15762396 | 0.054064859 | 0.107483628 |
| 17.93029 | 0.023037013 | 0.153105718 | 0.050675116 | 0.10690502 |
| 17.94053 | 0.018761827 | 0.148941979 | 0.046839369 | 0.106761422 |
| 17.95077 | 0.014667941 | 0.1463119 | 0.043345396 | 0.106696216 |
| 17.96101 | 0.011379481 | 0.144631282 | 0.039823734 | 0.107128454 |
| 17.97125 | 0.008549174 | 0.14400483 | 0.037174254 | 0.107878632 |
| 17.98149 | 0.007016898 | 0.144189574 | 0.035105349 | 0.109134012 |
| 17.99173 | 0.006286663 | 0.144550915 | 0.033803765 | 0.110811033 |
| 18.00197 | 0.006863631 | 0.145830881 | 0.033283549 | 0.11272019 |
| 18.01221 | 0.008161419 | 0.14671533 | 0.033276344 | 0.114775388 |
| 18.02245 | 0.009961534 | 0.148247721 | 0.034304825 | 0.116864028 |
| 18.03269 | 0.01239803 | 0.149881946 | 0.035254206 | 0.118683512 |
| 18.04293 | 0.014470152 | 0.152184679 | 0.036295114 | 0.120676726 |
| 18.05317 | 0.016203225 | 0.154638174 | 0.037570713 | 0.122422714 |
| 18.06341 | 0.01782182 | 0.158044863 | 0.038092764 | 0.124418894 |
| 18.07365 | 0.018719489 | 0.161327123 | 0.038664447 | 0.126598974 |
| 18.08389 | 0.01933842 | 0.164751337 | 0.038480955 | 0.128760537 |
| 18.09413 | 0.019493281 | 0.167915314 | 0.037881325 | 0.130823886 |
| 18.10437 | 0.019686904 | 0.170207509 | 0.03751156 | 0.132513886 |
| 18.11461 | 0.019659946 | 0.171457402 | 0.036551918 | 0.13302374 |
| 18.12485 | 0.020365532 | 0.171340876 | 0.035729659 | 0.133101033 |
| 18.13509 | 0.020946147 | 0.170187751 | 0.034931976 | 0.131933265 |
| 18.14533 | 0.022439513 | 0.1674343 | 0.034007492 | 0.130072482 |
| 18.15557 | 0.023996832 | 0.164099307 | 0.032946988 | 0.127549299 |
| 18.16581 | 0.026446907 | 0.159454151 | 0.031843583 | 0.125170162 |
| 18.17605 | 0.028963728 | 0.154912436 | 0.030566765 | 0.123037786 |
| 18.18629 | 0.031987995 | 0.149992791 | 0.029438075 | 0.121096408 |
| 18.19653 | 0.035483691 | 0.145398384 | 0.028491037 | 0.119972698 |
| 18.20677 | 0.039305946 | 0.140500037 | 0.027916756 | 0.119076781 |
| 18.21701 | 0.043619198 | 0.136415333 | 0.028436804 | 0.118479119 |
| 18.22725 | 0.048680281 | 0.131713046 | 0.029586406 | 0.118055023 |
| 18.23749 | 0.054509073 | 0.127073392 | 0.032031688 | 0.118364121 |
| 18.24773 | 0.060418279 | 0.12229449 | 0.035712989 | 0.119095171 |
| 18.25797 | 0.067127647 | 0.117405395 | 0.040305497 | 0.120783408 |
| 18.26821 | 0.073547609 | 0.11232494 | 0.045887126 | 0.123522578 |
| 18.27845 | 0.079370832 | 0.108206463 | 0.051687697 | 0.126796858 |
| 18.28869 | 0.084401755 | 0.104827 | 0.057779697 | 0.130966195 |
| 18.29893 | 0.087871817 | 0.10290337 | 0.063704643 | 0.13559549 |
| 18.30917 | 0.089624685 | 0.103007904 | 0.068705696 | 0.140319168 |
| 18.31941 | 0.089674999 | 0.104007523 | 0.073128941 | 0.144891145 |
| 18.32965 | 0.087710566 | 0.107002097 | 0.076961309 | 0.14901372 |
| 18.33989 | 0.084198201 | 0.110157817 | 0.079593805 | 0.151808437 |
| 18.35013 | 0.079638879 | 0.114255803 | 0.081857718 | 0.153920414 |
| 18.36037 | 0.073940516 | 0.117935361 | 0.083527452 | 0.154367904 |
| 18.37061 | 0.068391234 | 0.121882041 | 0.085065226 | 0.154036403 |
| 18.38085 | 0.062748806 | 0.125222283 | 0.086240287 | 0.152768771 |
| 18.39109 | 0.058167082 | 0.128849133 | 0.087319796 | 0.151622377 |
| 18.40133 | 0.054247751 | 0.131460964 | 0.088236652 | 0.15062384 |
| 18.41157 | 0.052025589 | 0.134150337 | 0.089021127 | 0.150210841 |
| 18.42181 | 0.050766463 | 0.136240775 | 0.089235035 | 0.15030349 |
| 18.43205 | 0.050856626 | 0.138106111 | 0.08924798 | 0.150887323 |
| 18.44229 | 0.051758347 | 0.139343078 | 0.088353905 | 0.152133839 |
| 18.45253 | 0.053366788 | 0.140363584 | 0.086906115 | 0.153233063 |
| 18.46277 | 0.055755918 | 0.140935978 | 0.084414229 | 0.154780072 |
| 18.47301 | 0.057816723 | 0.141492407 | 0.081019082 | 0.156441866 |
| 18.48325 | 0.060099185 | 0.141479462 | 0.076350602 | 0.15888789 |
| 18.49349 | 0.061949108 | 0.14144241 | 0.070935153 | 0.161597634 |
| 18.50373 | 0.06310042 | 0.141500142 | 0.064308109 | 0.165209517 |
| 18.51397 | 0.063529893 | 0.141051622 | 0.057669231 | 0.169393999 |
| 18.52421 | 0.062714393 | 0.141018612 | 0.050573204 | 0.173807987 |
| 18.53445 | 0.060420188 | 0.1404923 | 0.043924595 | 0.178364765 |
| 18.54469 | 0.056910031 | 0.14042918 | 0.038097269 | 0.182841053 |
| 18.55493 | 0.05188543 | 0.139657488 | 0.033083313 | 0.187032685 |
| 18.56517 | 0.045612648 | 0.139430431 | 0.029379484 | 0.190823512 |
| 18.57541 | 0.038686377 | 0.138858102 | 0.026335587 | 0.194167785 |
| 18.58565 | 0.031130337 | 0.138674226 | 0.024397033 | 0.196969352 |
| 18.59589 | 0.024100647 | 0.138070815 | 0.022692855 | 0.199195243 |
| 18.60613 | 0.017313442 | 0.137570067 | 0.021067104 | 0.20059793 |
| 18.61637 | 0.011534319 | 0.137084025 | 0.019445076 | 0.201437054 |
| 18.62661 | 0.006907816 | 0.136486069 | 0.017593495 | 0.201735802 |
| 18.63685 | 0.003103092 | 0.135655289 | 0.015469659 | 0.201287624 |
| 18.64709 | 0.000481024 | 0.135131592 | 0.013525828 | 0.20112013 |
| 18.65733 | -0.001483499 | 0.134529667 | 0.01187179 | 0.200695978 |
| 18.66757 | -0.002745628 | 0.134272974 | 0.011264052 | 0.200768283 |
| 18.67781 | -0.003655131 | 0.134000922 | 0.011306096 | 0.201493568 |
| 18.68805 | -0.004289931 | 0.134421488 | 0.012976061 | 0.202810219 |
| 18.69829 | -0.004671939 | 0.134887299 | 0.01554116 | 0.20465522 |
| 18.70853 | -0.005083946 | 0.136154343 | 0.019140954 | 0.207138558 |
| 18.71877 | -0.005538011 | 0.137334606 | 0.023620233 | 0.209231805 |
| 18.72901 | -0.006534031 | 0.139371475 | 0.027866586 | 0.211249054 |
| 18.73925 | -0.007107925 | 0.140669057 | 0.032282659 | 0.212308275 |
| 18.74949 | -0.008674537 | 0.142944361 | 0.036004956 | 0.212344292 |
| 18.75973 | -0.010047783 | 0.14482746 | 0.039390154 | 0.211253403 |
| 18.76997 | -0.011976739 | 0.1470535 | 0.041724493 | 0.20912255 |
| 18.78021 | -0.013834439 | 0.14988972 | 0.043975305 | 0.206050087 |
| 18.79045 | -0.015977598 | 0.153279594 | 0.045692065 | 0.202320681 |
| 18.80069 | -0.017974666 | 0.157402411 | 0.046817197 | 0.198029655 |
| 18.81093 | -0.020283492 | 0.162203563 | 0.048405425 | 0.193204162 |
| 18.82117 | -0.021794509 | 0.167279057 | 0.049125461 | 0.188304757 |
| 18.83142 | -0.02397759 | 0.172588737 | 0.049912336 | 0.182928953 |
| 18.84165 | -0.02533449 | 0.177419965 | 0.050172018 | 0.177256932 |
| 18.85189 | -0.026612893 | 0.181729451 | 0.04980526 | 0.171928148 |
| 18.86213 | -0.027229402 | 0.185045132 | 0.049137353 | 0.166593258 |
| 18.87237 | -0.027448619 | 0.187747964 | 0.047703849 | 0.161488568 |
| 18.88261 | -0.027206862 | 0.18873758 | 0.045925278 | 0.156575383 |
| 18.89285 | -0.026263455 | 0.189285961 | 0.043970403 | 0.151940338 |
| 18.9031 | -0.02513989 | 0.188956762 | 0.041538585 | 0.147677323 |
| 18.91333 | -0.023551636 | 0.187778369 | 0.039520109 | 0.143495749 |
| 18.92357 | -0.022255998 | 0.186705857 | 0.03745005 | 0.139902292 |
| 18.93382 | -0.020594484 | 0.185282022 | 0.035881995 | 0.136868649 |
| 18.94406 | -0.019539366 | 0.184021138 | 0.034882861 | 0.134644296 |
| 18.95429 | -0.018599116 | 0.182613006 | 0.03434718 | 0.133492643 |
| 18.96453 | -0.017831981 | 0.181172072 | 0.034628616 | 0.133476435 |
| 18.97478 | -0.017096076 | 0.179474786 | 0.035380284 | 0.134789285 |
| 18.98501 | -0.015982011 | 0.177650149 | 0.036542182 | 0.137137488 |
| 18.99525 | -0.014658473 | 0.175777059 | 0.037895012 | 0.140427963 |
| 19.0055 | -0.012324449 | 0.174024069 | 0.039690662 | 0.144187736 |
| 19.01574 | -0.009315504 | 0.171679648 | 0.041006525 | 0.148176337 |
| 19.02597 | -0.005078452 | 0.170307402 | 0.042345066 | 0.152055537 |
| 19.03621 | -0.000418471 | 0.167777631 | 0.043531875 | 0.155277916 |
| 19.04646 | 0.005146105 | 0.16582882 | 0.044051776 | 0.158120013 |
| 19.05669 | 0.010386381 | 0.163078934 | 0.044174569 | 0.160356975 |
| 19.06693 | 0.015919847 | 0.160546158 | 0.043870932 | 0.162329863 |
| 19.07718 | 0.020146986 | 0.157430601 | 0.042650169 | 0.16413196 |
| 19.08742 | 0.02420454 | 0.154361352 | 0.041143151 | 0.166205853 |
| 19.09765 | 0.026707821 | 0.151617154 | 0.039074017 | 0.168402486 |
| 19.10789 | 0.028689203 | 0.148698753 | 0.036736043 | 0.170639569 |
| 19.11814 | 0.030035936 | 0.146614634 | 0.034920951 | 0.173309065 |
| 19.12838 | 0.031141126 | 0.144387855 | 0.033432161 | 0.175935626 |
| 19.13861 | 0.031990382 | 0.143069682 | 0.0330211 | 0.178409262 |
| 19.14886 | 0.033259993 | 0.141949271 | 0.034203514 | 0.180390904 |
| 19.1591 | 0.035063339 | 0.141322216 | 0.036370308 | 0.1827762 |
| 19.16933 | 0.037161304 | 0.140763597 | 0.040209004 | 0.184427573 |
| 19.17957 | 0.040215432 | 0.140517968 | 0.045182307 | 0.186096767 |
| 19.18982 | 0.043342488 | 0.139758229 | 0.051188127 | 0.18751457 |

| | | | | |
|---|---|---|---|---|
| 16.79365 | -0.022592911 | 0.151058536 | 0.029330811 | 0.192538927 |
| 16.80389 | -0.022408385 | 0.149042998 | 0.025817715 | 0.189405459 |
| 16.81413 | -0.022648658 | 0.147705185 | 0.021339824 | 0.187150395 |
| 16.82437 | -0.023528832 | 0.14673869 | 0.016733065 | 0.185325968 |
| 16.83461 | -0.024751391 | 0.14647921 | 0.011779026 | 0.184032767 |
| 16.84485 | -0.026104132 | 0.146380947 | 0.007131879 | 0.183389077 |
| 16.85509 | -0.027672977 | 0.147052344 | 0.003006647 | 0.182450402 |
| 16.86533 | -0.029567806 | 0.148391613 | -0.000241542 | 0.182149828 |
| 16.87557 | -0.031644818 | 0.150689363 | -0.00305894 | 0.181238151 |
| 16.88581 | -0.033702535 | 0.153827054 | -0.004524455 | 0.180490056 |
| 16.89605 | -0.036579302 | 0.157569969 | -0.005990896 | 0.179154566 |
| 16.90629 | -0.039508682 | 0.162179976 | -0.006491114 | 0.177438087 |
| 16.91653 | -0.043084193 | 0.166547597 | -0.007374462 | 0.175656295 |
| 16.92677 | -0.04645606 | 0.170733861 | -0.008401998 | 0.173035963 |
| 16.93701 | -0.050848608 | 0.174440613 | -0.009872801 | 0.170466885 |
| 16.94725 | -0.054570335 | 0.177640274 | -0.012396027 | 0.167533172 |
| 16.95749 | -0.059075491 | 0.179744771 | -0.01567025 | 0.164278257 |
| 16.96773 | -0.063122279 | 0.181145895 | -0.019643264 | 0.161326942 |
| 16.97797 | -0.067306717 | 0.18202212 | -0.024442832 | 0.158337702 |
| 16.98821 | -0.07131409 | 0.18159696 | -0.029509218 | 0.156161677 |
| 16.99845 | -0.074829846 | 0.180625973 | -0.034534423 | 0.154542663 |
| 17.00869 | -0.077661233 | 0.178976291 | -0.039226402 | 0.153826446 |
| 17.01893 | -0.080140945 | 0.176938098 | -0.043361032 | 0.154454343 |
| 17.02917 | -0.081809672 | 0.175100899 | -0.046471447 | 0.155532415 |
| 17.03941 | -0.082575271 | 0.17304738 | -0.048855924 | 0.158110768 |
| 17.04965 | -0.083605949 | 0.172328997 | -0.050123248 | 0.160989464 |
| 17.05989 | -0.083884622 | 0.172156649 | -0.050439792 | 0.164721328 |
| 17.07013 | -0.084864659 | 0.173345776 | -0.05011554 | 0.169109835 |
| 17.08037 | -0.086200847 | 0.175655176 | -0.049037981 | 0.17385773 |
| 17.09061 | -0.088331194 | 0.178948714 | -0.047328021 | 0.179132139 |
| 17.10085 | -0.091267264 | 0.182635966 | -0.045614738 | 0.184480928 |
| 17.11109 | -0.095094506 | 0.186530212 | -0.043294985 | 0.190228433 |
| 17.12133 | -0.099268756 | 0.190254429 | -0.041436571 | 0.195687544 |
| 17.13157 | -0.103934181 | 0.193266682 | -0.039310388 | 0.201269455 |
| 17.14181 | -0.108304923 | 0.195615964 | -0.037931854 | 0.206710926 |
| 17.15205 | -0.111949718 | 0.197327448 | -0.036890594 | 0.211889745 |
| 17.16229 | -0.114711074 | 0.198168577 | -0.036357581 | 0.216798896 |
| 17.17253 | -0.115994478 | 0.198047503 | -0.037072275 | 0.221283045 |
| 17.18277 | -0.115865404 | 0.197444282 | -0.038229519 | 0.225378712 |
| 17.19301 | -0.114028384 | 0.195833177 | -0.040795626 | 0.228685783 |
| 17.20325 | -0.111421346 | 0.193567706 | -0.043736454 | 0.231186405 |
| 17.21349 | -0.107686726 | 0.190940309 | -0.047978984 | 0.233480799 |
| 17.22373 | -0.104302718 | 0.18786091 | -0.05221825 | 0.234229321 |
| 17.23397 | -0.101031389 | 0.185208009 | -0.057201242 | 0.235257578 |
| 17.24421 | -0.099058718 | 0.182739095 | -0.061587116 | 0.235078398 |
| 17.25445 | -0.097968099 | 0.18132916 | -0.065496055 | 0.235003459 |
| 17.26469 | -0.098412291 | 0.181133842 | -0.069112816 | 0.233776576 |
| 17.27493 | -0.099581179 | 0.181953559 | -0.071323596 | 0.23308201 |
| 17.28517 | -0.101561572 | 0.184100489 | -0.072876579 | 0.230981793 |
| 17.29541 | -0.103455918 | 0.187319403 | -0.073310128 | 0.228848396 |
| 17.30565 | -0.105160425 | 0.191387928 | -0.07322188 | 0.225514254 |
| 17.31589 | -0.106224034 | 0.196178288 | -0.071754477 | 0.221920495 |
| 17.32613 | -0.106416627 | 0.201427399 | -0.070050209 | 0.216959473 |
| 17.33637 | -0.106255433 | 0.206447129 | -0.066980411 | 0.211923578 |
| 17.34661 | -0.105546189 | 0.211912728 | -0.063447635 | 0.206363782 |
| 17.35685 | -0.104299211 | 0.216880989 | -0.058449564 | 0.200972998 |
| 17.36709 | -0.103811154 | 0.220912056 | -0.052866173 | 0.195685191 |
| 17.37733 | -0.103110772 | 0.224934302 | -0.046363616 | 0.191458086 |
| 17.38757 | -0.103062488 | 0.228323489 | -0.039513189 | 0.187439316 |
| 17.39781 | -0.103215248 | 0.231273798 | -0.032344705 | 0.184566478 |
| 17.40805 | -0.103935529 | 0.234641985 | -0.025797134 | 0.182078308 |
| 17.41829 | -0.104286106 | 0.238318642 | -0.0202133 | 0.180474715 |
| 17.42853 | -0.104486652 | 0.243075853 | -0.015697639 | 0.179100776 |
| 17.43877 | -0.104237169 | 0.24847003 | -0.012508932 | 0.178347787 |
| 17.44901 | -0.103242492 | 0.255100361 | -0.010699801 | 0.178163583 |
| 17.45925 | -0.101401887 | 0.262067074 | -0.010091164 | 0.178085049 |
| 17.46949 | -0.098591348 | 0.269510703 | -0.010134493 | 0.179126078 |
| 17.47973 | -0.095395981 | 0.276372112 | -0.010539419 | 0.180358297 |
| 17.48997 | -0.091521605 | 0.283044651 | -0.010987395 | 0.182417336 |
| 17.50021 | -0.087605321 | 0.288444961 | -0.011336812 | 0.184441223 |
| 17.51045 | -0.08329402 | 0.292854823 | -0.01043212 | 0.186762647 |
| 17.52069 | -0.07959874 | 0.296102182 | -0.009200473 | 0.188976603 |
| 17.53093 | -0.075791708 | 0.298154559 | -0.006830158 | 0.190384085 |
| 17.54117 | -0.071952894 | 0.299288245 | -0.003642688 | 0.191767266 |
| 17.55141 | -0.068318225 | 0.30011435 | 0.000261518 | 0.192364529 |
| 17.56165 | -0.064224485 | 0.300731253 | 0.004755592 | 0.192958559 |
| 17.57189 | -0.059753217 | 0.30138422 | 0.009217958 | 0.192747457 |
| 17.58213 | -0.054780119 | 0.302680051 | 0.013684055 | 0.192890165 |
| 17.59237 | -0.049579092 | 0.304702951 | 0.017534535 | 0.192644067 |
| 17.60261 | -0.044166511 | 0.307060138 | 0.020563905 | 0.192827401 |
| 17.61285 | -0.038589302 | 0.310169039 | 0.022557109 | 0.192479896 |
| 17.62309 | -0.03389262 | 0.313152314 | 0.023673875 | 0.192182035 |
| 17.63333 | -0.029645634 | 0.31692231 | 0.023332267 | 0.191761129 |
| 17.64357 | -0.02670854 | 0.32005709 | 0.022181304 | 0.191203862 |
| 17.65381 | -0.025075613 | 0.323296381 | 0.019544975 | 0.190879292 |
| 17.66405 | -0.024491851 | 0.325401485 | 0.016126968 | 0.190778698 |
| 17.67429 | -0.025950374 | 0.327016424 | 0.011671382 | 0.191818493 |
| 17.68453 | -0.027881783 | 0.327581377 | 0.00653685 | 0.193305038 |
| 17.69477 | -0.031357072 | 0.326996772 | 0.000912473 | 0.196223415 |
| 17.70501 | -0.035190786 | 0.325293612 | -0.004697898 | 0.199548749 |
| 17.71525 | -0.03938822 | 0.322872441 | -0.010090972 | 0.20366024 |
| 17.72549 | -0.043242349 | 0.319639023 | -0.015029446 | 0.207758737 |
| 17.73573 | -0.046997564 | 0.315805257 | -0.019157526 | 0.211966007 |
| 17.74597 | -0.050044384 | 0.311386991 | -0.022475183 | 0.215475402 |
| 17.75621 | -0.052548915 | 0.306796537 | -0.0247279 | 0.218681148 |
| 17.76645 | -0.054486818 | 0.301835172 | -0.026223565 | 0.221185337 |
| 17.77669 | -0.056195914 | 0.296913455 | -0.026780499 | 0.223237473 |
| 17.78693 | -0.057964433 | 0.291533733 | -0.026519934 | 0.22503792 |
| 17.79717 | -0.059614022 | 0.286907972 | -0.025363614 | 0.22665114 |
| 17.80741 | -0.061867836 | 0.281838496 | -0.023422722 | 0.228654925 |
| 17.81765 | -0.063801631 | 0.277452638 | -0.021034053 | 0.230692156 |
| 17.82789 | -0.066539858 | 0.272982196 | -0.017457921 | 0.233548056 |
| 17.83813 | -0.068908216 | 0.269174726 | -0.01381396 | 0.236696192 |
| 17.84837 | -0.071553604 | 0.265184272 | -0.009673995 | 0.239978033 |
| 17.85861 | -0.073761924 | 0.261662528 | -0.005429567 | 0.243404993 |
| 17.86885 | -0.076124595 | 0.258342389 | -0.000965228 | 0.246632354 |
| 17.87909 | -0.077743577 | 0.254650845 | 0.003268983 | 0.249509373 |
| 17.88933 | -0.078971168 | 0.251941222 | 0.007246276 | 0.251061296 |
| 17.89957 | -0.079557519 | 0.248597284 | 0.011053309 | 0.252208704 |
| 17.90981 | -0.078889961 | 0.245979975 | 0.014356644 | 0.252481004 |
| 17.92005 | -0.07744357 | 0.243743805 | 0.017620458 | 0.251102503 |
| 17.93029 | -0.074361196 | 0.241399705 | 0.020188091 | 0.249757989 |
| 17.94053 | -0.069984196 | 0.239712173 | 0.023061082 | 0.246814558 |
| 17.95077 | -0.064418947 | 0.23800923 | 0.025838231 | 0.244299205 |
| 17.96101 | -0.057274237 | 0.23639462 | 0.029036837 | 0.240137455 |
| 17.97125 | -0.049483597 | 0.234640349 | 0.032533637 | 0.236596709 |
| 17.98149 | -0.040844826 | 0.232622988 | 0.036991584 | 0.231754635 |
| 17.99173 | -0.032596965 | 0.230355736 | 0.041272485 | 0.227327485 |
| 18.00197 | -0.024401122 | 0.227640704 | 0.046196039 | 0.22165987 |
| 18.01221 | -0.017548216 | 0.224915681 | 0.050719838 | 0.216522617 |
| 18.02245 | -0.012049083 | 0.222439034 | 0.05481719 | 0.211055277 |
| 18.03269 | -0.008690461 | 0.220251306 | 0.057941631 | 0.206338372 |
| 18.04293 | -0.006931287 | 0.2190886 | 0.060173884 | 0.201935029 |
| 18.05317 | -0.007354825 | 0.218689471 | 0.061090208 | 0.198136423 |
| 18.06341 | -0.009085378 | 0.219668004 | 0.061342065 | 0.195035393 |
| 18.07365 | -0.012791125 | 0.221535699 | 0.060409172 | 0.192729026 |
| 18.08389 | -0.016381852 | 0.224474402 | 0.058496599 | 0.190393673 |
| 18.09413 | -0.021317873 | 0.227359387 | 0.056067847 | 0.189184371 |
| 18.10437 | -0.026222067 | 0.231059127 | 0.052699042 | 0.187605096 |
| 18.11461 | -0.03093845 | 0.233915235 | 0.048541916 | 0.187049793 |
| 18.12485 | -0.036000652 | 0.236908625 | 0.043601353 | 0.186416196 |
| 18.13509 | -0.040608706 | 0.239160216 | 0.038000173 | 0.186485021 |
| 18.14533 | -0.045770356 | 0.241763342 | 0.031666354 | 0.186460376 |
| 18.15557 | -0.050388048 | 0.243682795 | 0.024804858 | 0.186626631 |
| 18.16581 | -0.055723055 | 0.246119317 | 0.018251784 | 0.186688256 |
| 18.17605 | -0.060738953 | 0.248053994 | 0.011214918 | 0.18645907 |
| 18.18629 | -0.066213627 | 0.250233809 | 0.00493377 | 0.185803719 |
| 18.19653 | -0.071460635 | 0.251729728 | -0.00108485 | 0.18471705 |
| 18.20677 | -0.076296389 | 0.25264371 | -0.006397877 | 0.183310787 |
| 18.21701 | -0.080967056 | 0.25255037 | -0.011292404 | 0.181539626 |
| 18.22725 | -0.084324124 | 0.251655445 | -0.015558545 | 0.180115254 |
| 18.23749 | -0.087123386 | 0.249680784 | -0.01926225 | 0.179005814 |
| 18.24773 | -0.088472771 | 0.247141925 | -0.022666247 | 0.178972023 |
| 18.25797 | -0.088675413 | 0.24403504 | -0.025445279 | 0.180006856 |
| 18.26821 | -0.087860583 | 0.240840401 | -0.02788792 | 0.182690941 |
| 18.27845 | -0.085916346 | 0.2374676 | -0.029870295 | 0.186719344 |
| 18.28869 | -0.083352521 | 0.234524941 | -0.031489023 | 0.192564163 |
| 18.29893 | -0.080351784 | 0.231386643 | -0.0329769 | 0.199254661 |
| 18.30917 | -0.076989926 | 0.229020151 | -0.033953359 | 0.20739493 |
| 18.31941 | -0.07386989 | 0.227091594 | -0.035332326 | 0.216047241 |
| 18.32965 | -0.070636612 | 0.225942813 | -0.036081732 | 0.224993297 |
| 18.33989 | -0.068245619 | 0.22582769 | -0.03722112 | 0.234103904 |
| 18.35013 | -0.066193747 | 0.226276991 | -0.038152009 | 0.242598043 |
| 18.36037 | -0.065585719 | 0.227658272 | -0.038620564 | 0.250455846 |
| 18.37061 | -0.065749184 | 0.228856839 | -0.039118316 | 0.256635629 |
| 18.38085 | -0.06732132 | 0.23034311 | -0.038702005 | 0.261603379 |
| 18.39109 | -0.07024633 | 0.231003126 | -0.038085041 | 0.264271625 |
| 18.40133 | -0.073984436 | 0.231254419 | -0.037006228 | 0.264839817 |
| 18.41157 | -0.078900635 | 0.230902416 | -0.035240558 | 0.26352111 |
| 18.42181 | -0.083983523 | 0.229661232 | -0.033322756 | 0.260135173 |
| 18.43205 | -0.089345606 | 0.227844073 | -0.031161927 | 0.255485402 |
| 18.44229 | -0.094304829 | 0.225666358 | -0.028832201 | 0.249383831 |
| 18.45253 | -0.0990433 | 0.222830978 | -0.026398371 | 0.242875974 |
| 18.46277 | -0.103044398 | 0.220366032 | -0.023832735 | 0.236314322 |
| 18.47301 | -0.106370142 | 0.217993504 | -0.021793698 | 0.229032298 |
| 18.48325 | -0.109086849 | 0.216917178 | -0.019815736 | 0.222959351 |
| 18.49349 | -0.111168845 | 0.21638683 | -0.018226436 | 0.216421447 |
| 18.50373 | -0.112887128 | 0.217452929 | -0.017375608 | 0.211294495 |
| 18.51397 | -0.114134139 | 0.219356153 | -0.017078396 | 0.206494532 |
| 18.52421 | -0.115099926 | 0.222372738 | -0.017457129 | 0.202862686 |
| 18.53445 | -0.11633129 | 0.226395134 | -0.018300723 | 0.200888066 |
| 18.54469 | -0.117454776 | 0.230457939 | -0.019984575 | 0.199710955 |
| 18.55493 | -0.118945647 | 0.235592777 | -0.021385395 | 0.199786084 |
| 18.56517 | -0.120835927 | 0.240658561 | -0.02394094 | 0.200756482 |
| 18.57541 | -0.123177796 | 0.246075775 | -0.025686557 | 0.201879948 |
| 18.58565 | -0.125555493 | 0.251307804 | -0.028336869 | 0.203116528 |
| 18.59589 | -0.127909768 | 0.256804369 | -0.030758749 | 0.203198077 |
| 18.60613 | -0.130030899 | 0.26180438 | -0.033462686 | 0.203325799 |
| 18.61637 | -0.131622137 | 0.266880048 | -0.036302846 | 0.201611343 |
| 18.62661 | -0.132065506 | 0.271358974 | -0.039213773 | 0.199529959 |
| 18.63685 | -0.132139579 | 0.275119956 | -0.042113876 | 0.19619582 |
| 18.64709 | -0.130783535 | 0.278447035 | -0.045038122 | 0.192545439 |
| 18.65733 | -0.129093363 | 0.280765377 | -0.04733005 | 0.187469775 |
| 18.66757 | -0.126953033 | 0.282346793 | -0.049112922 | 0.18216599 |
| 18.67781 | -0.124324651 | 0.283582931 | -0.050290283 | 0.175539835 |
| 18.68805 | -0.122114978 | 0.284033214 | -0.050181966 | 0.168464917 |
| 18.69829 | -0.119742755 | 0.284007399 | -0.049361632 | 0.160142067 |
| 18.70853 | -0.11783305 | 0.283853636 | -0.047358372 | 0.151265976 |
| 18.71877 | -0.115919394 | 0.283182177 | -0.044775373 | 0.141634543 |
| 18.72901 | -0.114160411 | 0.281923454 | -0.04121575 | 0.132198937 |
| 18.73925 | -0.112381319 | 0.280466768 | -0.037706129 | 0.122652657 |
| 18.74949 | -0.110328085 | 0.278056153 | -0.033960412 | 0.114118863 |
| 18.75973 | -0.108431841 | 0.275646359 | -0.030780991 | 0.107086922 |
| 18.76997 | -0.105944928 | 0.272392144 | -0.027607117 | 0.101641717 |
| 18.78021 | -0.103850988 | 0.269120986 | -0.025666776 | 0.098432341 |
| 18.79045 | -0.100731245 | 0.266076586 | -0.02404394 | 0.09643862 |
| 18.80069 | -0.097912051 | 0.2632843 | -0.023445173 | 0.095974011 |
| 18.81093 | -0.09460902 | 0.260445957 | -0.023261172 | 0.095896033 |
| 18.82117 | -0.090803563 | 0.258193456 | -0.023511744 | 0.095549734 |
| 18.83142 | -0.086668582 | 0.256157785 | -0.024385312 | 0.095371332 |
| 18.84165 | -0.082397365 | 0.254074804 | -0.024863702 | 0.09457195 |
| 18.85189 | -0.078002678 | 0.251688294 | -0.025272944 | 0.094250294 |
| 18.86213 | -0.073153253 | 0.249150849 | -0.025384366 | 0.093892678 |
| 18.87237 | -0.068791998 | 0.246127461 | -0.024504331 | 0.094321041 |
| 18.88261 | -0.064536021 | 0.242510609 | -0.023491026 | 0.095512363 |
| 18.89285 | -0.059933966 | 0.238373369 | -0.021421536 | 0.097029017 |
| 18.9031 | -0.055517539 | 0.233374035 | -0.019644849 | 0.098656617 |
| 18.91333 | -0.050878807 | 0.228240239 | -0.017608757 | 0.1002765 |
| 18.92357 | -0.045848406 | 0.222373015 | -0.015843086 | 0.101264699 |
| 18.93382 | -0.040248561 | 0.216588081 | -0.01528142 | 0.101812983 |
| 18.94406 | -0.034857597 | 0.211208386 | -0.015327169 | 0.102222501 |
| 18.95429 | -0.028813251 | 0.206521153 | -0.016846371 | 0.102913197 |
| 18.96453 | -0.023379493 | 0.202688094 | -0.019254105 | 0.10415801 |
| 18.97478 | -0.018132603 | 0.200398929 | -0.022643074 | 0.106632854 |
| 18.98501 | -0.01350428 | 0.199904618 | -0.026833059 | 0.110700776 |
| 18.99525 | -0.010221373 | 0.200331486 | -0.031426468 | 0.11550859 |
| 19.0055 | -0.007312581 | 0.202534469 | -0.035793575 | 0.121919497 |
| 19.01574 | -0.005914774 | 0.205062379 | -0.040064349 | 0.128506543 |
| 19.02597 | -0.004763351 | 0.207953318 | -0.043690113 | 0.134895776 |
| 19.03621 | -0.004707551 | 0.209880045 | -0.04659384 | 0.141440125 |
| 19.04646 | -0.005157537 | 0.211516084 | -0.048290879 | 0.14693001 |
| 19.05669 | -0.005966211 | 0.211516702 | -0.049115397 | 0.151924245 |
| 19.06693 | -0.00692734 | 0.210554856 | -0.049150958 | 0.15601761 |
| 19.07718 | -0.008186639 | 0.208286639 | -0.048242646 | 0.15957027 |
| 19.08742 | -0.009619328 | 0.205804574 | -0.046864536 | 0.162001434 |
| 19.09765 | -0.010932267 | 0.20254685 | -0.045405227 | 0.163605567 |
| 19.10789 | -0.012115225 | 0.199739047 | -0.043966223 | 0.163985113 |
| 19.11814 | -0.013422641 | 0.196555456 | -0.042838817 | 0.162811762 |
| 19.12838 | -0.014555673 | 0.193728726 | -0.041997333 | 0.160485057 |
| 19.13861 | -0.015476053 | 0.19104906 | -0.041824116 | 0.156359336 |
| 19.14886 | -0.01610771 | 0.187603245 | -0.041434614 | 0.151071323 |
| 19.1591 | -0.016480634 | 0.183960942 | -0.041481788 | 0.14526118 |
| 19.16933 | -0.016364253 | 0.179445551 | -0.04109735 | 0.138603233 |
| 19.17957 | -0.01540233 | 0.17464814 | -0.040672116 | 0.132540358 |
| 19.18982 | -0.014019601 | 0.169418408 | -0.039914103 | 0.126790084 |

| | | | | |
|---|---|---|---|---|
| 19.20006 | 0.122159428 | 0.190619322 | -0.048594574 | 0.20624002 |
| 19.21029 | 0.119742678 | 0.181155801 | -0.045651738 | 0.20735623 |
| 19.22054 | 0.115173871 | 0.171927548 | -0.044183773 | 0.20928591 |
| 19.23078 | 0.108257403 | 0.164033763 | -0.044375374 | 0.21185606 |
| 19.24102 | 0.099935359 | 0.15674612 | -0.04569214 | 0.2151001 |
| 19.25126 | 0.089209481 | 0.150641689 | -0.048589692 | 0.21836778 |
| 19.2615 | 0.078155865 | 0.144832566 | -0.052757877 | 0.22146825 |
| 19.27174 | 0.0653086 | 0.140181696 | -0.057967294 | 0.22342895 |
| 19.28197 | 0.052782462 | 0.134892988 | -0.063844375 | 0.2243858 |
| 19.29222 | 0.040233116 | 0.130282607 | -0.070440783 | 0.22419097 |
| 19.30246 | 0.028916182 | 0.125942233 | -0.076732796 | 0.22253167 |
| 19.3127 | 0.018706105 | 0.122785847 | -0.083122188 | 0.22043637 |
| 19.32294 | 0.010271096 | 0.121313213 | -0.088269405 | 0.21741912 |
| 19.33318 | 0.004004729 | 0.121465709 | -0.092941057 | 0.21414527 |
| 19.34342 | -0.000384816 | 0.124397998 | -0.095804627 | 0.21035648 |
| 19.35366 | -0.003069229 | 0.129137515 | -0.0970537 | 0.20662902 |
| 19.3639 | -0.003834291 | 0.135525884 | -0.096975056 | 0.20233898 |
| 19.37414 | -0.003260966 | 0.143368088 | -0.095110555 | 0.19855769 |
| 19.38438 | -0.001569831 | 0.151097512 | -0.092066175 | 0.19488955 |
| 19.39462 | 0.001386443 | 0.159514134 | -0.087654751 | 0.19186279 |
| 19.40486 | 0.005603325 | 0.167198131 | -0.082666162 | 0.18974914 |
| 19.4151 | 0.010479828 | 0.174814144 | -0.077296298 | 0.18885113 |
| 19.42534 | 0.016668702 | 0.181690323 | -0.071990946 | 0.18841787 |
| 19.43558 | 0.02349836 | 0.188426472 | -0.066861314 | 0.18862646 |
| 19.44582 | 0.031480785 | 0.194337282 | -0.062712398 | 0.18934598 |
| 19.45606 | 0.039411716 | 0.199433859 | -0.059258264 | 0.18954604 |
| 19.4663 | 0.047369485 | 0.203385815 | -0.056902574 | 0.19050541 |
| 19.47654 | 0.055650985 | 0.205920454 | -0.055512282 | 0.19053244 |
| 19.48678 | 0.062636034 | 0.207023998 | -0.055069475 | 0.19153842 |
| 19.49702 | 0.068957984 | 0.206892748 | -0.054962696 | 0.19213697 |
| 19.50726 | 0.07435012 | 0.205607979 | -0.054544204 | 0.19348251 |
| 19.5175 | 0.078848198 | 0.204099857 | -0.054376348 | 0.19467418 |
| 19.52774 | 0.081955379 | 0.201860474 | -0.052455179 | 0.19695021 |
| 19.53798 | 0.084050835 | 0.200792783 | -0.050060874 | 0.19890103 |
| 19.54822 | 0.085478066 | 0.198942228 | -0.04574707 | 0.20180545 |
| 19.55846 | 0.085401396 | 0.197856793 | -0.040237754 | 0.20430839 |
| 19.5687 | 0.084775083 | 0.19637549 | -0.034385019 | 0.20698083 |
| 19.57894 | 0.082467247 | 0.194745788 | -0.027278901 | 0.2100476 |
| 19.58918 | 0.079683717 | 0.192360216 | -0.020539794 | 0.21284879 |
| 19.59942 | 0.07562719 | 0.188993904 | -0.014082047 | 0.21612374 |
| 19.60966 | 0.070878723 | 0.185083541 | -0.008762978 | 0.21952789 |
| 19.6199 | 0.065837639 | 0.180909864 | -0.004298471 | 0.22337301 |
| 19.63014 | 0.060393182 | 0.177107183 | -0.000710631 | 0.22698967 |
| 19.64038 | 0.055048905 | 0.173975384 | 0.001257957 | 0.23061534 |
| 19.65062 | 0.050266246 | 0.172746905 | 0.002525633 | 0.23347306 |
| 19.66086 | 0.045921103 | 0.172756359 | 0.00320905 | 0.23621187 |
| 19.6711 | 0.042333342 | 0.173665516 | 0.002876722 | 0.23736931 |
| 19.68134 | 0.03973429 | 0.17519274 | 0.002077878 | 0.23748791 |
| 19.69158 | 0.037508788 | 0.176804326 | 0.000773598 | 0.23725214 |
| 19.70182 | 0.036280239 | 0.17760666 | -0.001108927 | 0.23518846 |
| 19.71206 | 0.035523308 | 0.178080837 | -0.003349009 | 0.23279679 |
| 19.7223 | 0.034685053 | 0.178226286 | -0.00521865 | 0.2299454 |
| 19.73254 | 0.033821605 | 0.178105011 | -0.007431416 | 0.22684664 |
| 19.74278 | 0.032207581 | 0.179131532 | -0.00900796 | 0.22373326 |
| 19.75302 | 0.029588894 | 0.180761381 | -0.009846101 | 0.22074832 |
| 19.76326 | 0.025875336 | 0.1839633 | -0.009517638 | 0.21815707 |
| 19.7735 | 0.020953464 | 0.188056358 | -0.008467813 | 0.21490489 |
| 19.78374 | 0.015628717 | 0.193062942 | -0.006246435 | 0.21218915 |
| 19.79398 | 0.010274753 | 0.198007924 | -0.003237884 | 0.20868968 |
| 19.80422 | 0.004955778 | 0.202192649 | -0.000111142 | 0.20517811 |
| 19.81446 | 0.001177721 | 0.205011699 | 0.003741328 | 0.20100322 |
| 19.8247 | -0.000785053 | 0.206202489 | 0.007121519 | 0.19716795 |
| 19.83494 | -0.001963047 | 0.204983743 | 0.010004039 | 0.19306321 |
| 19.84518 | -0.000935487 | 0.202519029 | 0.012923611 | 0.18895557 |
| 19.85542 | 0.000603556 | 0.197650536 | 0.015043635 | 0.18514378 |
| 19.86566 | 0.002067654 | 0.192361984 | 0.016580561 | 0.18132859 |
| 19.8759 | 0.004078271 | 0.18480356 | 0.018249419 | 0.17799171 |
| 19.88614 | 0.003895116 | 0.177320772 | 0.019277828 | 0.17503486 |
| 19.89638 | 0.003083054 | 0.169049114 | 0.020619547 | 0.17284589 |
| 19.90662 | 0.000331235 | 0.159887738 | 0.021930207 | 0.17148926 |
| 19.91686 | -0.003566894 | 0.151023045 | 0.023333766 | 0.17266305 |
| 19.9271 | -0.008475401 | 0.142011119 | 0.024773818 | 0.1747804 |
| 19.93734 | -0.01398673 | 0.132415197 | 0.02623472 | 0.17940487 |
| 19.94758 | -0.019761 | 0.123867766 | 0.027463626 | 0.18509152 |
| 19.95782 | -0.024657092 | 0.115655401 | 0.027897682 | 0.19244462 |
| 19.96806 | -0.029570332 | 0.108523545 | 0.027784901 | 0.19963746 |
| 19.9783 | -0.033579869 | 0.103461386 | 0.026107227 | 0.2069528 |
| 19.98854 | -0.03657325 | 0.099594129 | 0.023274607 | 0.21234248 |

| | | | | |
|---|---|---|---|---|
| 19.20006 | -0.115612525 | 0.180891013 | -0.006877533 | 0.141693581 |
| 19.21029 | -0.123513499 | 0.18386246 | -0.009371934 | 0.135039117 |
| 19.22054 | -0.1316179 | 0.185735209 | -0.011172718 | 0.131006861 |
| 19.23078 | -0.140237795 | 0.187143179 | -0.011902826 | 0.129125507 |
| 19.24102 | -0.14772815 | 0.186239831 | -0.012016059 | 0.129908514 |
| 19.25126 | -0.15513286 | 0.184697354 | -0.011455978 | 0.130532694 |
| 19.2615 | -0.161545342 | 0.181736473 | -0.009920277 | 0.132817958 |
| 19.27174 | -0.167078414 | 0.178148457 | -0.007922512 | 0.134655494 |
| 19.28197 | -0.171193017 | 0.174092946 | -0.005439475 | 0.137585324 |
| 19.29222 | -0.174626869 | 0.170224567 | -0.002425114 | 0.141811071 |
| 19.30246 | -0.176441692 | 0.167569412 | 0.001021636 | 0.147889077 |
| 19.3127 | -0.177503393 | 0.165508911 | 0.004548071 | 0.155646276 |
| 19.32294 | -0.177463919 | 0.164827857 | 0.008409448 | 0.165834093 |
| 19.33318 | -0.175675175 | 0.165862048 | 0.012644745 | 0.176834323 |
| 19.34342 | -0.173247935 | 0.167892177 | 0.016692424 | 0.188899729 |
| 19.35366 | -0.169413378 | 0.171216052 | 0.020775967 | 0.200680079 |
| 19.3639 | -0.164442588 | 0.175198359 | 0.024785028 | 0.211687531 |
| 19.37414 | -0.15860094 | 0.178882452 | 0.028336144 | 0.220409815 |
| 19.38438 | -0.152320044 | 0.182746155 | 0.031365041 | 0.22768457 |
| 19.39462 | -0.145795564 | 0.185214346 | 0.033288239 | 0.231909061 |
| 19.40486 | -0.138846275 | 0.186878131 | 0.034327332 | 0.233641111 |
| 19.4151 | -0.133509931 | 0.186538218 | 0.033585713 | 0.232767031 |
| 19.42534 | -0.12823766 | 0.185714421 | 0.03160296 | 0.230844064 |
| 19.43558 | -0.124850074 | 0.182993769 | 0.028150697 | 0.227122807 |
| 19.44582 | -0.122308054 | 0.179859241 | 0.023592116 | 0.223455577 |
| 19.45606 | -0.121974869 | 0.175891882 | 0.018005428 | 0.220353855 |
| 19.4663 | -0.122426938 | 0.171999564 | 0.012163805 | 0.217599648 |
| 19.47654 | -0.123661038 | 0.167831721 | 0.006581329 | 0.21666306 |
| 19.48678 | -0.125621366 | 0.16396496 | 0.001439707 | 0.216119482 |
| 19.49702 | -0.127714583 | 0.160903211 | -0.002413446 | 0.216692605 |
| 19.50726 | -0.129295099 | 0.157879122 | -0.005457247 | 0.217014275 |
| 19.5175 | -0.1302101 | 0.15614599 | -0.006436943 | 0.217383734 |
| 19.52774 | -0.130277026 | 0.154556692 | -0.006973524 | 0.216693585 |
| 19.53798 | -0.12926936 | 0.15340939 | -0.006167085 | 0.214884108 |
| 19.54822 | -0.12693399 | 0.153017116 | -0.004996505 | 0.212647688 |
| 19.55846 | -0.124248323 | 0.152136021 | -0.003793409 | 0.209462186 |
| 19.5687 | -0.119902647 | 0.152159167 | -0.00259091 | 0.206667069 |
| 19.57894 | -0.115256218 | 0.151842978 | -0.002367115 | 0.204036317 |
| 19.58918 | -0.109717589 | 0.152047088 | -0.002486238 | 0.202506793 |
| 19.59942 | -0.104138066 | 0.151774617 | -0.003003371 | 0.202004227 |
| 19.60966 | -0.09815285 | 0.151844405 | -0.004428728 | 0.20270006 |
| 19.6199 | -0.092111176 | 0.151647965 | -0.005332413 | 0.204046266 |
| 19.63014 | -0.086794019 | 0.151451634 | -0.00644422 | 0.206348326 |
| 19.64038 | -0.081217894 | 0.150721429 | -0.006815014 | 0.207585749 |
| 19.65062 | -0.076444549 | 0.150851583 | -0.00664418 | 0.209258762 |
| 19.66086 | -0.071593083 | 0.15034302 | -0.005241649 | 0.208925846 |
| 19.6711 | -0.067601395 | 0.150971935 | -0.003489067 | 0.207950459 |
| 19.68134 | -0.063499981 | 0.151525919 | -0.000323644 | 0.205841604 |
| 19.69158 | -0.059688757 | 0.152901038 | 0.002915236 | 0.202054363 |
| 19.70182 | -0.056308253 | 0.155815177 | 0.0069917 | 0.198823113 |
| 19.71206 | -0.052849295 | 0.158662138 | 0.010780884 | 0.19484649 |
| 19.7223 | -0.049689772 | 0.162377718 | 0.014352071 | 0.191412545 |
| 19.73254 | -0.046387072 | 0.165858984 | 0.01762708 | 0.189388412 |
| 19.74278 | -0.043181757 | 0.169236065 | 0.019910398 | 0.187468708 |
| 19.75302 | -0.039886869 | 0.170537638 | 0.021414404 | 0.187271384 |
| 19.76326 | -0.036395129 | 0.171115798 | 0.022295145 | 0.187590251 |
| 19.7735 | -0.032708977 | 0.169713416 | 0.022240151 | 0.189262815 |
| 19.78374 | -0.028992487 | 0.167502391 | 0.022264728 | 0.191669387 |
| 19.79398 | -0.024857584 | 0.164201256 | 0.021793032 | 0.195548555 |
| 19.80422 | -0.020386406 | 0.161870677 | 0.021293344 | 0.200881632 |
| 19.81446 | -0.016110087 | 0.159328522 | 0.021516742 | 0.206434696 |
| 19.8247 | -0.011570591 | 0.157885571 | 0.021650093 | 0.21333467 |
| 19.83494 | -0.007480694 | 0.15671204 | 0.022054233 | 0.219424221 |
| 19.84518 | -0.003982891 | 0.155108064 | 0.022870039 | 0.225086991 |
| 19.85542 | -0.001484459 | 0.152152398 | 0.023806794 | 0.228961295 |
| 19.86566 | 0.000242719 | 0.147541276 | 0.024620792 | 0.231661073 |
| 19.8759 | 0.000298355 | 0.141246447 | 0.025624172 | 0.233007359 |
| 19.88614 | -0.00046786 | 0.133633158 | 0.026621635 | 0.23405214 |
| 19.89638 | -0.001682365 | 0.125039436 | 0.027638188 | 0.234944129 |
| 19.90662 | -0.003093016 | 0.117368554 | 0.029087909 | 0.23615223 |
| 19.91686 | -0.004170484 | 0.111424548 | 0.030449924 | 0.238988836 |
| 19.9271 | -0.004162114 | 0.107335075 | 0.031995926 | 0.241533922 |
| 19.93734 | -0.002726789 | 0.106028795 | 0.033840107 | 0.245294186 |
| 19.94758 | -0.000449119 | 0.107028738 | 0.034864922 | 0.24825209 |
| 19.95782 | 0.003489087 | 0.110069097 | 0.035780929 | 0.251191736 |
| 19.96806 | 0.008073161 | 0.114532924 | 0.035740986 | 0.252296491 |
| 19.9783 | 0.01352462 | 0.120298411 | 0.034958103 | 0.253372872 |
| 19.98854 | 0.01884101 | 0.126641342 | 0.033144988 | 0.252366183 |

| | | | | |
|---|---|---|---|---|
| 19.20006 | 0.04719408 | 0.139150941 | 0.057427629 | 0.189162893 |
| 19.21029 | 0.050924757 | 0.137814396 | 0.064135122 | 0.190183704 |
| 19.22054 | 0.054725657 | 0.136503283 | 0.07036386 | 0.19101342 |
| 19.23078 | 0.057978486 | 0.134365564 | 0.076643769 | 0.190827046 |
| 19.24102 | 0.060507326 | 0.132754364 | 0.081929428 | 0.190058523 |
| 19.25126 | 0.062132034 | 0.130838255 | 0.087181533 | 0.187825005 |
| 19.2615 | 0.062990739 | 0.129568216 | 0.091561841 | 0.184894443 |
| 19.27174 | 0.062478928 | 0.129118579 | 0.095681908 | 0.180967408 |
| 19.28197 | 0.061577737 | 0.12980881 | 0.098952402 | 0.177528446 |
| 19.29222 | 0.060137531 | 0.131971571 | 0.101910108 | 0.174198156 |
| 19.30246 | 0.058550118 | 0.135042838 | 0.104111552 | 0.172254803 |
| 19.3127 | 0.057363748 | 0.139827755 | 0.105342385 | 0.171808528 |
| 19.32294 | 0.056831418 | 0.144577987 | 0.106066895 | 0.17336786 |
| 19.33318 | 0.057128303 | 0.150252285 | 0.105957655 | 0.176887473 |
| 19.34342 | 0.058472693 | 0.155682637 | 0.105326387 | 0.18181518 |
| 19.35366 | 0.060904186 | 0.160966365 | 0.104403597 | 0.188502299 |
| 19.3639 | 0.063990803 | 0.16544784 | 0.103303525 | 0.195080613 |
| 19.37414 | 0.067626927 | 0.169183839 | 0.102652245 | 0.202186977 |
| 19.38438 | 0.071758928 | 0.172405078 | 0.102085411 | 0.208385451 |
| 19.39462 | 0.075533365 | 0.174057604 | 0.102205589 | 0.213947937 |
| 19.40486 | 0.079064528 | 0.175912276 | 0.102411542 | 0.217841953 |
| 19.4151 | 0.081569806 | 0.175844695 | 0.103066303 | 0.221022305 |
| 19.42534 | 0.083494185 | 0.176510039 | 0.103528302 | 0.222964041 |
| 19.43558 | 0.08403427 | 0.175992136 | 0.103559534 | 0.224247048 |
| 19.44582 | 0.083617488 | 0.175581993 | 0.10292133 | 0.224850136 |
| 19.45606 | 0.081968121 | 0.174783901 | 0.101482743 | 0.224919463 |
| 19.4663 | 0.079354531 | 0.174070162 | 0.098766103 | 0.224955112 |
| 19.47654 | 0.075623902 | 0.173779344 | 0.094965824 | 0.224318468 |
| 19.48678 | 0.071345574 | 0.17412227 | 0.089740521 | 0.223391855 |
| 19.49702 | 0.066493615 | 0.174661989 | 0.083867173 | 0.221982539 |
| 19.50726 | 0.060813997 | 0.176041267 | 0.076892733 | 0.220235784 |
| 19.5175 | 0.055527903 | 0.178336087 | 0.069480792 | 0.217800047 |
| 19.52774 | 0.049448023 | 0.18011178 | 0.06198197 | 0.215321163 |
| 19.53798 | 0.043762491 | 0.182126921 | 0.054725175 | 0.212782959 |
| 19.54822 | 0.037620242 | 0.183549006 | 0.048269405 | 0.210699563 |
| 19.55846 | 0.031846311 | 0.183853734 | 0.042417962 | 0.209366252 |
| 19.5687 | 0.025592934 | 0.1838802 | 0.038235284 | 0.208909624 |
| 19.57894 | 0.019985289 | 0.182807269 | 0.034996036 | 0.209591339 |
| 19.58918 | 0.014070887 | 0.18192777 | 0.03328543 | 0.21053957 |
| 19.59942 | 0.009035845 | 0.181011571 | 0.033124927 | 0.212033217 |
| 19.60966 | 0.004520796 | 0.180502522 | 0.034391478 | 0.212668624 |
| 19.6199 | 0.000964312 | 0.179924551 | 0.036739241 | 0.212995865 |
| 19.63014 | -0.001616851 | 0.179848914 | 0.040518329 | 0.211330926 |
| 19.64038 | -0.002829557 | 0.179480046 | 0.04517607 | 0.209022206 |
| 19.65062 | -0.00308224 | 0.178874885 | 0.050961488 | 0.204551213 |
| 19.66086 | -0.002244486 | 0.178520813 | 0.056843972 | 0.199983544 |
| 19.6711 | -0.000512391 | 0.177532331 | 0.063190058 | 0.194282287 |
| 19.68134 | 0.002039938 | 0.176990721 | 0.069245586 | 0.188734141 |
| 19.69158 | 0.00496317 | 0.176592119 | 0.075023904 | 0.18395529 |
| 19.70182 | 0.008444645 | 0.176364958 | 0.07969386 | 0.179574204 |
| 19.71206 | 0.012161474 | 0.176888515 | 0.083304008 | 0.176460622 |
| 19.7223 | 0.016414364 | 0.17767211 | 0.086421723 | 0.174540604 |
| 19.73254 | 0.020756507 | 0.179395155 | 0.088242627 | 0.173453831 |
| 19.74278 | 0.025557018 | 0.181800909 | 0.089479765 | 0.173215551 |
| 19.75302 | 0.030759696 | 0.185407916 | 0.089744036 | 0.173237637 |
| 19.76326 | 0.036251778 | 0.189846602 | 0.090019616 | 0.173513272 |
| 19.7735 | 0.042029288 | 0.195560462 | 0.089255122 | 0.173118478 |
| 19.78374 | 0.047884665 | 0.201786112 | 0.087898159 | 0.172385998 |
| 19.79398 | 0.053162012 | 0.208601735 | 0.086184851 | 0.170912263 |
| 19.80422 | 0.057846093 | 0.215417468 | 0.083489993 | 0.169209309 |
| 19.81446 | 0.061456863 | 0.221334191 | 0.080450921 | 0.166725868 |
| 19.8247 | 0.063397774 | 0.226775088 | 0.077077114 | 0.164140248 |
| 19.83494 | 0.063750393 | 0.230607062 | 0.07385233 | 0.1611854 |
| 19.84518 | 0.061971372 | 0.233376256 | 0.070778234 | 0.158379873 |
| 19.85542 | 0.058488534 | 0.234192124 | 0.068449623 | 0.155406792 |
| 19.86566 | 0.053469628 | 0.234233842 | 0.067043071 | 0.153322836 |
| 19.8759 | 0.046712153 | 0.232877127 | 0.066285978 | 0.15160014 |
| 19.88614 | 0.039176114 | 0.230851318 | 0.06651471 | 0.151020646 |
| 19.89638 | 0.030654645 | 0.228534521 | 0.067224348 | 0.151363751 |
| 19.90662 | 0.021711766 | 0.225944947 | 0.068554499 | 0.152809216 |
| 19.91686 | 0.012386961 | 0.223261824 | 0.069896444 | 0.155090641 |
| 19.9271 | 0.003198772 | 0.220371239 | 0.071287965 | 0.158326882 |
| 19.93734 | -0.005890556 | 0.217756996 | 0.072295645 | 0.161970263 |
| 19.94758 | -0.014713772 | 0.214793331 | 0.072787712 | 0.1660154 |
| 19.95782 | -0.022970942 | 0.211815863 | 0.072550303 | 0.170192749 |
| 19.96806 | -0.030238051 | 0.209176641 | 0.071752241 | 0.173972433 |
| 19.9783 | -0.037141284 | 0.20707115 | 0.069967421 | 0.177412181 |
| 19.98854 | -0.042521073 | 0.205239981 | 0.067757531 | 0.179880298 |

| | | | | |
|---|---|---|---|---|
| 19.20006 | -0.011712223 | 0.164088217 | -0.03881125 | 0.122157962 |
| 19.21029 | -0.009114685 | 0.158938497 | -0.037729507 | 0.118347649 |
| 19.22054 | -0.005883268 | 0.154635176 | -0.036788818 | 0.115877029 |
| 19.23078 | -0.002902096 | 0.151174268 | -0.035687508 | 0.114614756 |
| 19.24102 | -9.10886E-05 | 0.148609838 | -0.034725295 | 0.11386423 |
| 19.25126 | 0.002065613 | 0.147529343 | -0.034147316 | 0.114477315 |
| 19.2615 | 0.003674568 | 0.147141184 | -0.033346661 | 0.115850624 |
| 19.27174 | 0.004310806 | 0.148432239 | -0.032925194 | 0.117896272 |
| 19.28197 | 0.004389819 | 0.149753498 | -0.032290273 | 0.120460631 |
| 19.29222 | 0.004496041 | 0.151960803 | -0.031965905 | 0.123195961 |
| 19.30246 | 0.004191391 | 0.154305709 | -0.031646245 | 0.125937312 |
| 19.3127 | 0.004463531 | 0.15614807 | -0.031452481 | 0.128509058 |
| 19.32294 | 0.005107348 | 0.157180171 | -0.031442987 | 0.130958758 |
| 19.33318 | 0.006622657 | 0.157193155 | -0.031347469 | 0.132989162 |
| 19.34342 | 0.008322787 | 0.155723438 | -0.031867111 | 0.134815874 |
| 19.35366 | 0.01103368 | 0.153078904 | -0.031981702 | 0.136540721 |
| 19.3639 | 0.013665695 | 0.148654194 | -0.032834196 | 0.138286053 |
| 19.37414 | 0.016779088 | 0.144185776 | -0.033897939 | 0.139955057 |
| 19.38438 | 0.019476241 | 0.138349354 | -0.035461842 | 0.142058666 |
| 19.39462 | 0.022541418 | 0.133117657 | -0.037505321 | 0.144356061 |
| 19.40486 | 0.024872773 | 0.128254511 | -0.040487545 | 0.147489858 |
| 19.4151 | 0.027208615 | 0.124363502 | -0.043824496 | 0.15111166 |
| 19.42534 | 0.028706337 | 0.12167876 | -0.048111984 | 0.155336964 |
| 19.43558 | 0.029929032 | 0.119785688 | -0.052238158 | 0.160455436 |
| 19.44582 | 0.030350106 | 0.11939642 | -0.056927638 | 0.165624385 |
| 19.45606 | 0.03012436 | 0.11939603 | -0.061150506 | 0.171081178 |
| 19.4663 | 0.029590732 | 0.119959647 | -0.064607618 | 0.175938292 |
| 19.47654 | 0.028479374 | 0.120899848 | -0.067411569 | 0.180383575 |
| 19.48678 | 0.028168739 | 0.121956603 | -0.068695802 | 0.183411132 |
| 19.49702 | 0.02754517 | 0.12309987 | -0.069064071 | 0.185725316 |
| 19.50726 | 0.028463969 | 0.12449168 | -0.067827023 | 0.186784569 |
| 19.5175 | 0.030075654 | 0.12587868 | -0.065924393 | 0.18691144 |
| 19.52774 | 0.033264012 | 0.127194091 | -0.062270316 | 0.186946974 |
| 19.53798 | 0.037605576 | 0.128337148 | -0.058834248 | 0.1864437 |
| 19.54822 | 0.043253817 | 0.129159437 | -0.05404678 | 0.186535868 |
| 19.55846 | 0.0496422 | 0.130280754 | -0.049483723 | 0.186966573 |
| 19.5687 | 0.05706159 | 0.130508859 | -0.044412684 | 0.187604071 |
| 19.57894 | 0.064866671 | 0.130925239 | -0.039506655 | 0.18923012 |
| 19.58918 | 0.073140165 | 0.131245482 | -0.034737513 | 0.190816681 |
| 19.59942 | 0.081485925 | 0.131642543 | -0.030267137 | 0.192634273 |
| 19.60966 | 0.09024816 | 0.131974084 | -0.026287963 | 0.1942131 |
| 19.6199 | 0.099088778 | 0.132770547 | -0.022726851 | 0.195228858 |
| 19.63014 | 0.108121777 | 0.133745787 | -0.020070483 | 0.195286249 |
| 19.64038 | 0.117034484 | 0.134952797 | -0.017820376 | 0.194532603 |
| 19.65062 | 0.126181451 | 0.135743623 | -0.016506725 | 0.192598899 |
| 19.66086 | 0.134690261 | 0.136582568 | -0.015567087 | 0.189932304 |
| 19.6711 | 0.142772944 | 0.136792545 | -0.015132474 | 0.18645593 |
| 19.68134 | 0.149820978 | 0.135905472 | -0.014957049 | 0.182475419 |
| 19.69158 | 0.155932513 | 0.135025595 | -0.014900669 | 0.178701734 |
| 19.70182 | 0.160811991 | 0.133274356 | -0.014777921 | 0.174596027 |
| 19.71206 | 0.164153125 | 0.131512004 | -0.014714153 | 0.171450696 |
| 19.7223 | 0.16724166 | 0.129863056 | -0.01416269 | 0.168451503 |
| 19.73254 | 0.168877235 | 0.128833933 | -0.013755757 | 0.165765515 |
| 19.74278 | 0.170799251 | 0.128436193 | -0.013364018 | 0.163645205 |
| 19.75302 | 0.171663389 | 0.128928139 | -0.012466011 | 0.161348577 |
| 19.76326 | 0.173339896 | 0.130485124 | -0.012112502 | 0.159136295 |
| 19.7735 | 0.174417597 | 0.132711637 | -0.01186505 | 0.156543737 |
| 19.78374 | 0.175321358 | 0.135600203 | -0.011575639 | 0.153813355 |
| 19.79398 | 0.17600005 | 0.138551743 | -0.011724784 | 0.150722595 |
| 19.80422 | 0.176334696 | 0.140940225 | -0.01202908 | 0.148147049 |
| 19.81446 | 0.175955445 | 0.142957478 | -0.012559159 | 0.145621302 |
| 19.8247 | 0.175296149 | 0.143552088 | -0.013329401 | 0.144325286 |
| 19.83494 | 0.174545755 | 0.143402137 | -0.014136408 | 0.144687739 |
| 19.84518 | 0.173803866 | 0.142065193 | -0.015434875 | 0.146129471 |
| 19.85542 | 0.173437636 | 0.140140863 | -0.016744469 | 0.149899556 |
| 19.86566 | 0.173743375 | 0.137975395 | -0.018087659 | 0.154264624 |
| 19.8759 | 0.175205669 | 0.134967568 | -0.019994189 | 0.159711035 |
| 19.88614 | 0.177052235 | 0.132952582 | -0.021892709 | 0.165351735 |
| 19.89638 | 0.18018424 | 0.130653475 | -0.024122531 | 0.170642587 |
| 19.90662 | 0.184147124 | 0.128857052 | -0.026203901 | 0.175443411 |
| 19.91686 | 0.188685271 | 0.127438345 | -0.028581171 | 0.179091262 |
| 19.9271 | 0.193873553 | 0.126044602 | -0.030576567 | 0.18259642 |
| 19.93734 | 0.199510998 | 0.12510418 | -0.032253892 | 0.184499034 |
| 19.94758 | 0.205557049 | 0.124383347 | -0.033178834 | 0.186484969 |
| 19.95782 | 0.211370965 | 0.123362865 | -0.033436542 | 0.187758293 |
| 19.96806 | 0.217298654 | 0.123415394 | -0.032977948 | 0.189312837 |
| 19.9783 | 0.222706165 | 0.123138787 | -0.031550652 | 0.190860067 |
| 19.98854 | 0.227549195 | 0.123541147 | -0.029800685 | 0.193343826 |

**Vestibular counts for Fig. S5**

| Age | *Mir183/96* $^{dko}$ genotype | Number of animals without hyperactivity | Number of animals displaying hyperactivity |
|---|---|---|---|
| P14 | Wildtype | 9 | 0 |
| P14 | Heterozygote | 7 | 0 |
| P14 | Homozygote | 4 | 1 |
| P21 | Wildtype | 9 | 0 |
| P21 | Heterozygote | 10 | 0 |
| P21 | Homozygote | 5 | 0 |
| P28 | Wildtype | 5 | 0 |
| P28 | Heterozygote | 7 | 0 |
| P28 | Homozygote | 5 | 0 |
| P56 | Wildtype | 12 | 0 |
| P56 | Heterozygote | 16 | 0 |
| P56 | Homozygote | 4 | 3 |
| P90 | Wildtype | 10 | 0 |
| P90 | Heterozygote | 14 | 0 |
| P90 | Homozygote | 4 | 4 |
| P180 | Wildtype | 10 | 0 |
| P180 | Heterozygote | 16 | 0 |
| P180 | Homozygote | 1 | 8 |

**DPOAE measurements for Fig. S6**

*Mir183/96 [dko]*

| F2 (kHz) | Wildtype Mean | Wildtype Standard deviation | Heterozygote Mean | Heterozygote Standard deviation | Homozygote Mean | Homozygote Standard deviation |
|---|---|---|---|---|---|---|
| 6 | 35.625 | 5.629958132 | 40 | 5.477225575 | 68.75 | 7.5 |
| 12 | 16.875 | 3.720119046 | 17.5 | 4.183300133 | 67.5 | 5 |
| 18 | 18.75 | 3.535533906 | 17.5 | 6.123724357 | 75 | 0 |
| 24 | 35 | 7.071067812 | 34.16666667 | 6.645800679 | 71.25 | 4.787135539 |
| 30 | 50 | 10.35098339 | 45 | 8.366600265 | 75 | 0 |
| 36 | 61.25 | 13.29607891 | 56.66666667 | 8.164965809 | 75 | 0 |

| F1 level (dB SPL) | 6k Wildtype mean | 12k Wildtype mean | 18k Wildtype mean | 24k Wildtype mean | 30k Wildtype mean | 36k Wildtype mean |
|---|---|---|---|---|---|---|
| -10 | -28.60504243 | -15.59303219 | -16.22379187 | -15.81019366 | -17.73758131 | -17.61273114 |
| -5 | -29.91969997 | -25.02354253 | -18.18287859 | -9.892037552 | -19.09570022 | -14.45834288 |
| 0 | -25.91785523 | -17.98873402 | -16.70365197 | -10.9342201 | -23.49198003 | -15.89474161 |
| 5 | -24.58478976 | -17.73438323 | -12.27310362 | -11.89813459 | -21.23430093 | -15.84286272 |
| 10 | -23.99436992 | -15.50737258 | -15.33309005 | -21.83058131 | -21.18706061 | -14.59999782 |
| 15 | -28.24848486 | -6.511125216 | -8.242061264 | -10.18451752 | -19.78857741 | -16.27325453 |
| 20 | -23.65389261 | 0.775626056 | 2.169595635 | -15.59622688 | -20.01142106 | -17.13162835 |
| 25 | -24.99917005 | 8.00819872 | 9.614540416 | -10.58033997 | -14.1886151 | -17.98111386 |
| 30 | -18.88343667 | 14.31248422 | 17.17009677 | -5.34025492 | -15.13157158 | -18.28752819 |
| 35 | -13.51655968 | 19.34759296 | 22.98884557 | 1.376710896 | -12.21729203 | -14.59254195 |
| 40 | -9.062787743 | 23.26121728 | 27.78720282 | 6.774633758 | -7.871867856 | -12.62222186 |
| 45 | -2.632197104 | 26.22046464 | 30.32547251 | 9.863048624 | -5.942917184 | -11.58820744 |
| 50 | 1.110681722 | 26.57835904 | 30.82839296 | 15.4245273 | -3.227742976 | -5.676161285 |
| 55 | 5.795209936 | 23.51807667 | 29.36526157 | 17.5437833 | 2.011541424 | -0.540573152 |
| 60 | 7.623096608 | 26.73435981 | 31.4697152 | 20.49590861 | 6.07438198 | -1.917242208 |
| 65 | 9.228725436 | 33.8242816 | 35.20741862 | 22.15885222 | 5.716994432 | 5.496362144 |
| 70 | 10.7589517 | 39.55936768 | 43.75753933 | 23.83612566 | 12.32985088 | 11.29347168 |
| 75 | 17.2321913 | 42.3905792 | 46.98271642 | 35.18217549 | 18.79629894 | 12.7343247 |

| F1 level (dB SPL) | 6k Standard deviation | 12k Standard deviation | 18k Standard deviation | 24k Standard deviation | 30k Standard deviation | 36k Standard deviation |
|---|---|---|---|---|---|---|
| -10 | 9.939418778 | 5.148797691 | 4.70077768 | 7.207204204 | 2.357838239 | 5.247587605 |
| -5 | 8.641413568 | 6.110121312 | 3.857311786 | 7.236148726 | 8.241190335 | 7.176945578 |
| 0 | 3.784875772 | 6.612754897 | 11.12635825 | 3.897157798 | 3.489014212 | 12.40596658 |
| 5 | 4.88589446 | 7.149300889 | 5.33892892 | 7.212496673 | 5.228264713 | 8.195340152 |
| 10 | 3.743116223 | 7.08360937 | 3.912088726 | 12.09799055 | 8.568947117 | 8.395666634 |
| 15 | 7.701786117 | 3.105363663 | 5.67813224 | 5.500522222 | 4.716456711 | 4.961002684 |
| 20 | 6.388171329 | 1.551760897 | 3.428911072 | 6.126908812 | 5.338103172 | 8.466238323 |
| 25 | 6.407732209 | 0.890129228 | 3.457287686 | 7.03830162 | 6.389450832 | 5.376984381 |
| 30 | 8.352100219 | 0.586718082 | 2.530364095 | 5.10019795 | 6.544700375 | 7.718647185 |
| 35 | 10.88390206 | 0.670837386 | 2.235260284 | 2.940356933 | 4.358940702 | 4.332864182 |
| 40 | 11.95987787 | 0.904796519 | 1.813818718 | 3.885692112 | 6.089002343 | 7.986051818 |
| 45 | 7.624682154 | 0.982446628 | 1.551051907 | 5.688074767 | 8.507523366 | 7.464888487 |
| 50 | 7.137861691 | 1.532281309 | 1.508876522 | 4.257599122 | 11.01884361 | 12.03925758 |
| 55 | 4.881019457 | 4.030431426 | 2.023270216 | 4.849434499 | 6.539343634 | 10.64160816 |
| 60 | 6.674487318 | 1.588895655 | 2.633451157 | 4.132365645 | 5.641455934 | 11.85721789 |
| 65 | 6.847105404 | 1.614295557 | 4.442339952 | 4.601266984 | 6.882403517 | 12.4669861 |
| 70 | 8.511443718 | 1.35369334 | 2.81482245 | 9.628295428 | 7.100463219 | 10.2266848 |
| 75 | 6.341094285 | 1.591639899 | 3.037745746 | 4.498761688 | 6.575341056 | 12.78414224 |

| F1 level (dB SPL) | 6k Heterozygote mean | 12k Heterozygote mean | 18k Heterozygote mean | 24k Heterozygote mean | 30k Heterozygote mean | 36k Heterozygote mean |
|---|---|---|---|---|---|---|
| -10 | -22.59748932 | -18.59428881 | -24.07568077 | -21.65452433 | -21.28703198 | -15.42728525 |
| -5 | -26.24983654 | -22.19625131 | -16.16412177 | -9.708395701 | -19.43931921 | -17.53704107 |
| 0 | -28.57224602 | -19.32768666 | -22.7372695 | -14.60982434 | -19.8381597 | -15.69259431 |
| 5 | -23.28568832 | -20.09328179 | -18.02074061 | -18.0100096 | -20.9855203 | -17.13576875 |
| 10 | -26.49007036 | -15.04839441 | -11.88398677 | -16.00686609 | -17.50688495 | -15.00448165 |
| 15 | -26.89067383 | -9.191314005 | -4.937489899 | -9.095874475 | -17.07013956 | -16.04901077 |
| 20 | -28.00528486 | -4.868920299 | 3.807684693 | -16.84906914 | -18.14447445 | -13.7014173 |
| 25 | -23.42038153 | 5.4492416 | 11.01720789 | -12.3812867 | -15.31691742 | -19.24909107 |
| 30 | -24.37696802 | 12.48946253 | 16.52135851 | -4.203731435 | -15.59852352 | -20.09548766 |
| 35 | -18.7816361 | 17.66631509 | 21.7581469 | -3.291677669 | -12.61822716 | -18.00373734 |
| 40 | -13.97706573 | 21.75939413 | 25.94899763 | 5.020035733 | -5.411789835 | -15.99350016 |
| 45 | -7.425318443 | 24.80150221 | 28.976 | 10.57828523 | -3.855835936 | -17.81165252 |
| 50 | -5.292836437 | 26.08625391 | 30.22412868 | 14.67608218 | 2.588823253 | -5.684484043 |
| 55 | 0.939323647 | 23.01549568 | 29.85113463 | 17.0113833 | 5.185461259 | -2.409339285 |
| 60 | 1.305556245 | 22.00843366 | 27.6624425 | 18.57465873 | 3.393848256 | 2.310411422 |
| 65 | 3.20162088 | 29.9498083 | 32.02040422 | 17.74529365 | 6.955926229 | 9.156767573 |
| 70 | 1.297553024 | 35.75497694 | 41.3197353 | 27.33520725 | 15.32584764 | 13.76796797 |
| 75 | 7.63896896 | 38.76544205 | 46.98233924 | 36.28801502 | 21.94782805 | 16.56626688 |

| F1 level (dB SPL) | 6k Standard deviation | 12k Standard deviation | 18k Standard deviation | 24k Standard deviation | 30k Standard deviation | 36k Standard deviation |
|---|---|---|---|---|---|---|
| -10 | 3.745830443 | 6.523476411 | 13.12408753 | 13.02776887 | 8.131034328 | 6.484678859 |
| -5 | 4.562469915 | 10.20083193 | 3.019089987 | 6.096884253 | 3.326240122 | 10.43823764 |
| 0 | 4.602010864 | 3.703536294 | 14.55760935 | 7.999382653 | 5.956560034 | 14.0696886 |
| 5 | 4.069175272 | 4.880103023 | 6.927954814 | 9.73237983 | 4.637676232 | 7.901711976 |
| 10 | 4.351720654 | 4.842236893 | 2.746869478 | 6.481100179 | 6.940147253 | 9.910194962 |
| 15 | 1.469405118 | 3.550910733 | 5.397697715 | 4.103426785 | 4.357627293 | 6.742476129 |
| 20 | 5.869171558 | 11.41187758 | 5.321339126 | 8.777659828 | 5.884481436 | 4.35734649 |
| 25 | 7.151337557 | 4.451198171 | 4.35330277 | 6.962412654 | 4.470884947 | 2.260601376 |
| 30 | 6.846638814 | 2.576919086 | 4.163570183 | 2.877052894 | 12.96534813 | 6.856523799 |
| 35 | 5.119373535 | 2.321795467 | 3.377311595 | 7.792584039 | 9.415442858 | 7.253742872 |
| 40 | 7.016004298 | 2.631313933 | 3.34346312 | 2.769220035 | 8.267873748 | 8.538262525 |
| 45 | 5.986382336 | 2.727045846 | 3.007505514 | 2.277932812 | 8.851613226 | 15.14705486 |
| 50 | 8.116688525 | 1.749958115 | 2.730163111 | 2.0011126 | 7.302477606 | 10.82232236 |
| 55 | 5.649818571 | 1.501768836 | 1.946015215 | 3.114739879 | 8.712145317 | 10.32393025 |
| 60 | 7.48385561 | 3.985791468 | 2.407081912 | 4.377305389 | 6.204527463 | 7.499088249 |
| 65 | 7.161038465 | 3.494653135 | 6.547592327 | 10.7743953 | 4.497631452 | 8.645491469 |
| 70 | 14.18550069 | 3.130601993 | 4.208917769 | 3.959719221 | 4.1734699 | 7.790309014 |
| 75 | 7.289264207 | 3.607835932 | 3.402196649 | 2.956769127 | 3.514682907 | 7.659687374 |

| F1 level (dB SPL) | 6k Homozygote mean | 12k Homozygote mean | 18k Homozygote mean | 24k Homozygote mean | 30k Homozygote mean | 36k Homozygote mean |
|---|---|---|---|---|---|---|
| -10 | -24.73370982 | -13.45420659 | -16.78011443 | -12.98019277 | -18.07485248 | -18.2525894 |
| -5 | -26.46031565 | -17.68467059 | -13.40537216 | -6.588580885 | -17.6470377 | -15.09050402 |
| 0 | -20.80160179 | -18.19788742 | -15.72453517 | -6.216698368 | -18.6537449 | -14.50087339 |
| 5 | -23.2760041 | -13.90969958 | -12.77372928 | -10.35087616 | -18.43997402 | -16.55710498 |
| 10 | -20.34305536 | -14.29422208 | -11.18592464 | -12.89664858 | -13.86693862 | -6.894117888 |
| 15 | -29.38425267 | -13.27928499 | -20.35016013 | -7.728410752 | -17.86225152 | -14.33038729 |
| 20 | -19.37546854 | -12.53912858 | -10.6959415 | -11.87149747 | -13.98498867 | -8.247599915 |
| 25 | -22.51792026 | -20.40751024 | -10.11802662 | -6.401827136 | -11.71660032 | -11.69005022 |
| 30 | -28.68081152 | -13.34366106 | -14.80303552 | -5.836304064 | -14.66108442 | -19.91144004 |
| 35 | -21.72810112 | -10.9926185 | -11.1754592 | -5.824848336 | -12.40490195 | -13.01642752 |
| 40 | -18.41911987 | -10.36244602 | -10.09516083 | -4.072267864 | -10.42546867 | -9.390651136 |
| 45 | -10.20835635 | -1.468863744 | -5.867087328 | -7.278336384 | -13.15488518 | -10.37898048 |
| 50 | -13.83362893 | 0.242695808 | -1.605260928 | -0.085188352 | -6.782936512 | -6.35591456 |
| 55 | -8.25367104 | 0.484946176 | -8.049076928 | 2.23832864 | 4.405894488 | -2.493708992 |
| 60 | -16.02685363 | -13.08023629 | -9.355657328 | 4.328088864 | 3.478205344 | -4.894721312 |
| 65 | -12.09594803 | -1.724287904 | -7.083865584 | -5.358553088 | -3.399582872 | -8.516053056 |
| 70 | -13.85290598 | 2.934963456 | -0.898195344 | -4.58440736 | -4.150060224 | -12.83901403 |
| 75 | -6.98767616 | -4.276202368 | -1.210145312 | -5.75122192 | -0.994297088 | -5.823424736 |

| F1 level (dB SPL) | 6k Standard deviation | 12k Standard deviation | 18k Standard deviation | 24k Standard deviation | 30k Standard deviation | 36k Standard deviation |
|---|---|---|---|---|---|---|
| -10 | 0.745167829 | 7.849559886 | 8.140136543 | 3.597386767 | 10.68099851 | 8.766002922 |
| -5 | 7.911167207 | 7.488326847 | 3.900931082 | 9.48555632 | 7.670566402 | 6.634681355 |
| 0 | 6.618301078 | 13.00354556 | 12.03275212 | 7.848766847 | 6.490433951 | 9.052557364 |
| 5 | 6.533851416 | 4.874811537 | 5.291967503 | 5.745570073 | 9.34617236 | 13.54686828 |
| 10 | 9.615627902 | 8.713585766 | 8.349121655 | 5.68271995 | 2.570913531 | 5.859575945 |
| 15 | 11.12900562 | 5.769720772 | 8.168954446 | 3.170621819 | 10.63711261 | 9.991703181 |
| 20 | 8.132807772 | 8.021382811 | 6.237329818 | 4.704577803 | 4.547204794 | 7.53730375 |
| 25 | 10.36745704 | 20.43365895 | 6.411378622 | 3.131051742 | 9.838966186 | 4.937828596 |
| 30 | 5.580598775 | 7.380421952 | 6.872989913 | 3.69467094 | 6.523775901 | 6.875899979 |
| 35 | 8.890284169 | 8.790224371 | 5.690531649 | 5.076600625 | 7.744361137 | 6.780266898 |
| 40 | 8.348954157 | 10.46628549 | 4.866455199 | 8.703226594 | 4.492165198 | 3.501110194 |
| 45 | 4.751577628 | 6.057479422 | 8.913740114 | 6.236771573 | 9.203705366 | 3.363230789 |
| 50 | 5.876521462 | 6.375622964 | 11.83560956 | 7.051324094 | 13.42633925 | 12.26247774 |
| 55 | 3.136174232 | 9.595411891 | 3.146047192 | 9.110459188 | 7.316052598 | 8.500027993 |
| 60 | 3.676577808 | 3.293570716 | 15.52203277 | 8.305291609 | 7.209826414 | 5.426512518 |
| 65 | 3.94288767 | 3.496662906 | 12.75174432 | 5.489650227 | 11.20662314 | 8.249530314 |
| 70 | 3.549622004 | 6.305805652 | 8.315295148 | 10.74497296 | 10.88675448 | 11.28335072 |
| 75 | 0.582924415 | 11.75832576 | 7.613084438 | 19.55870974 | 8.591828883 | 9.393650352 |

| F2 (kHz) | Homozygote #1 | Homozygote #2 | Homozygote #3 | Homozygote #4 |
|---|---|---|---|---|
| 6 | 65 | 75 | 75 | 60 |
| 12 | 65 | 65 | 75 | 65 |
| 18 | 75 | 75 | 75 | 75 |
| 24 | 65 | 70 | 75 | 75 |
| 30 | 75 | 75 | 75 | 75 |
| 36 | 75 | 75 | 75 | 75 |

| F1 Level (dB SPL) | 6k | 12k | 18k | 24k | 30k | 36k |
|---|---|---|---|---|---|---|
| Homozygote #1 | | | | | | |
| -10 | -23.8341673 | -10.50221773 | -13.96156621 | -9.363416064 | -28.13754778 | |
| -5 | -29.34220595 | -16.03280998 | -10.31487283 | -1.43654592 | -17.11845786 | |
| 0 | -29.9172311 | -35.22516582 | -6.654916096 | -6.660761088 | -15.56555162 | |
| 5 | -32.39358464 | -15.04105574 | -7.872191488 | -7.35159296 | -20.41197773 | |
| 10 | -18.88266854 | -15.19432909 | -10.31960576 | -10.02951373 | -11.0271959 | |
| 15 | -25.21674342 | -10.45917696 | -15.58265754 | -4.833416704 | -15.11532851 | |
| 20 | -18.65044378 | -8.302582784 | -6.817736704 | -8.99570176 | -11.76814694 | |
| 25 | -28.9507799 | -21.02487654 | -8.906492928 | -3.489758464 | -8.48039424 | |
| 30 | -36.86479462 | -14.74206822 | -13.59854387 | -1.704630528 | -11.58008115 | |
| 35 | -27.83743181 | -3.295328256 | -11.76418099 | -0.5910864 | -14.7472599 | |
| 40 | -26.56629965 | -3.982614272 | -7.305137152 | -4.349349376 | -10.74169958 | -10.20895334 |
| 45 | -11.00954829 | 3.355889664 | 1.971518464 | -6.571396096 | -7.030930944 | -9.182174208 |
| 50 | -22.39503155 | 6.1897088 | 8.800388096 | 3.941559552 | 4.075100928 | 2.393741056 |
| 55 | -5.920345088 | 8.718603264 | -3.481161984 | 11.2499671 | 8.8917504 | 4.2109632 |
| 60 | -16.66645094 | -9.509387264 | 0.165412672 | 10.32671232 | 12.13665587 | -0.153108608 |
| 65 | -9.046459392 | -0.749558656 | -1.253436032 | 0.93906176 | 2.490024192 | -9.23267584 |
| 70 | -12.49635123 | 8.583293952 | 5.52855552 | 2.420285696 | 8.0074624 | -0.73744032 |
| 75 | -6.857849856 | 3.543699712 | 6.125352448 | 6.976641024 | 5.19693568 | 2.138883584 |
| Homozygote #2 | | | | | | |
| -10 | -25.20832 | -15.55206349 | -20.69615206 | -10.46167142 | -20.64402432 | -27.49666918 |
| -5 | -23.20389734 | -20.93628416 | -13.79813581 | -4.381480448 | -20.46549606 | -8.674577408 |
| 0 | -18.19091149 | -17.59514624 | -15.69968538 | -9.332760576 | -15.56688486 | -4.929741312 |
| 5 | -17.00538061 | -12.73906176 | -9.891004416 | -9.477989376 | -19.04556851 | -9.492943872 |
| 10 | -20.09575014 | -15.25391258 | -11.4780631 | -20.35218637 | -17.18129459 | -5.675869184 |
| 15 | -38.62352691 | -10.21250867 | -12.12580864 | -5.7835136 | -20.02130739 | -7.313411584 |
| 20 | -22.29067981 | -10.46406963 | -7.041419776 | -10.07849882 | -12.33277645 | -7.288014848 |
| 25 | -14.92702925 | -47.51211725 | -5.582951424 | -5.479550464 | -6.55374592 | -9.054030848 |
| 30 | -27.59226778 | -8.738176 | -21.34485197 | -5.776718336 | -16.63227494 | -13.5032105 |
| 35 | -24.29555098 | -5.219492864 | -5.111574016 | -3.57660672 | -11.23594547 | -12.64518963 |
| 40 | -20.74082714 | -5.555109376 | -5.043423232 | 0.36216592 | -9.268513792 | -8.170758144 |
| 45 | -8.96499712 | 2.202440192 | 1.602354304 | -4.429963776 | -3.728589568 | -11.63055718 |
| 50 | -12.03284787 | 5.259535872 | 8.370907648 | 4.552355328 | 5.324174848 | 2.478185984 |
| 55 | -7.576989184 | 7.787968512 | -9.60290304 | 8.764832768 | 12.21046067 | 5.481603584 |
| 60 | -20.90949427 | -17.05912422 | 1.545234688 | 11.73086003 | 6.60477184 | -1.93084672 |
| 65 | -9.306525696 | 2.587361024 | 0.83850144 | -8.206142464 | 4.311279104 | 2.961531648 |
| 70 | -15.1218647 | 8.202481152 | 4.381147648 | 5.838775808 | 1.608050432 | -12.75175526 |
| 75 | -6.654415872 | 3.06733696 | 1.58616768 | 5.954778112 | 5.243588608 | 0.82045248 |
| Homozygote #3 | | | | | | |
| -10 | -25.46632499 | -4.624740864 | -6.879801344 | -15.50997094 | -2.966573568 | -17.20169062 |
| -5 | -17.46117632 | -8.119981568 | -10.73507021 | 0.013732141 | -7.400782848 | -14.6727383 |
| 0 | -14.36022886 | -3.578498816 | -7.806857216 | 4.74391808 | -15.09876019 | -15.64722688 |
| 5 | -20.96650445 | -8.071003136 | -19.9355863 | -5.892422656 | -5.87120896 | -8.002380288 |
| 10 | -9.497993216 | -2.773782528 | -1.271965824 | -7.24479232 | -13.0521088 | -1.739429888 |
| 15 | -15.53531187 | -10.5138217 | -30.41205658 | -8.37597952 | -5.368210432 | -9.907565568 |
| 20 | -8.61070336 | -7.010019328 | -8.98755072 | -9.514663936 | -11.07688038 | -1.236041088 |
| 25 | -12.54282138 | 1.617681536 | -6.480922112 | -5.79624192 | -5.475451392 | -8.629623808 |
| 30 | -24.9161687 | -6.72822272 | -5.683625472 | -5.18473728 | -7.676596224 | -27.17498163 |
| 35 | -8.570627072 | -12.80335565 | -9.16749312 | -6.660804608 | -2.534240512 | -6.429406208 |
| 40 | -6.76013568 | -5.900178944 | -12.14571213 | 3.820407296 | -5.426112512 | -5.436033024 |
| 45 | -4.709000192 | -1.402326528 | -12.11401626 | -1.923011584 | -22.78224896 | -6.418805248 |
| 50 | -9.002958848 | -6.047630336 | -13.33471539 | 1.661606144 | -20.77764198 | -6.724223488 |
| 55 | -12.84690944 | -11.43034061 | -10.5231401 | -4.034150144 | -0.32289808 | -10.30392013 |
| 60 | -12.53949338 | -11.48390502 | -7.221760512 | -6.002375168 | -3.351664896 | -12.4478976 |
| 65 | -17.5166423 | -3.109175808 | -1.784603008 | -2.800534016 | -0.449587552 | -11.24921446 |
| 70 | -18.0406825 | -2.347883776 | -0.846096448 | -9.009439744 | -10.45098803 | -27.91124992 |
| 75 | -7.845422592 | -21.46862694 | -0.756925952 | -34.55268864 | -1.422039552 | -8.100972032 |
| Homozygote #4 | | | | | | |
| -10 | -24.42602701 | -23.13780429 | -25.58293811 | -16.58571264 | -20.55126426 | -10.05940838 |
| -5 | -35.83398298 | -25.64960666 | -18.77340979 | -20.55002931 | -25.60341402 | -21.92419635 |
| 0 | -20.73803571 | -16.39273882 | -32.73668198 | -13.61718989 | -28.38378291 | -22.92565197 |
| 5 | -22.73854669 | -19.7876777 | -13.39613491 | -18.68149965 | -28.43114086 | -32.17599078 |
| 10 | -32.89580954 | -23.95486413 | -21.67406387 | -13.96010189 | -14.2071552 | -13.26705459 |
| 15 | -38.16142848 | -21.93163264 | -23.28011776 | -11.92073318 | -30.94415974 | -25.7701847 |
| 20 | -27.95004723 | -24.37984256 | -19.93705882 | -18.89712538 | -20.76215091 | -16.21874381 |
| 25 | -33.6510505 | -14.7107287 | -19.50174003 | -10.8417577 | -26.35680973 | -17.386496 |
| 30 | -25.35001498 | -23.16617728 | -18.58512077 | -10.67913011 | -22.75538534 | -19.056128 |
| 35 | -26.20879462 | -22.65229722 | -18.65858867 | -12.47089562 | -21.10216192 | -19.97468672 |
| 40 | -19.60921702 | -26.01188147 | -15.88637082 | -16.1222953 | -16.2655488 | -13.74686003 |
| 45 | -16.14987981 | -10.0314583 | -14.92820582 | -16.18897408 | -19.07777126 | -14.28438528 |
| 50 | -11.90367744 | -4.430831104 | -10.25762406 | -10.49627443 | -15.75337984 | -23.57136179 |
| 55 | -6.670440448 | -3.136446464 | -8.589102592 | -7.027335168 | -3.15573504 | -9.363482624 |
| 60 | -13.99197594 | -14.26852864 | -31.91151616 | 1.257158272 | -1.47694144 | -5.04703232 |
| 65 | -12.51416474 | -5.625778176 | -26.13592474 | -11.36659763 | -19.95004723 | -16.54385357 |
| 70 | -9.752725504 | -2.698037504 | -12.6563881 | -17.5872512 | -15.7647657 | -9.955610624 |
| 75 | -6.59301632 | -2.2472192 | -11.79517542 | -1.383618176 | -12.99567309 | -18.15206298 |

*Mir182* [ko]

| F2 (kHz) | Wildtype Mean | Wildtype Standard deviation | Homozygote Mean | Homozygote Standard deviation |
|---|---|---|---|---|
| 6 | 35 | 7.071067812 | 35.71428571 | 13.04753215 |
| 12 | 15 | 0 | 15.71428571 | 3.450327797 |
| 18 | 17.5 | 3.535533906 | 21.42857143 | 11.44344271 |
| 24 | 30 | 0 | 48.57142857 | 17.96159925 |
| 30 | 45 | 7.071067812 | 65.71428571 | 9.759000729 |
| 36 | 57.5 | 3.535533906 | 74.28571429 | 1.889822365 |

| F1 level (dB SPL) | 6k Wildtype mean | 12k Wildtype mean | 18k Wildtype mean | 24k Wildtype mean | 30k Wildtype mean | 36k Wildtype mean |
|---|---|---|---|---|---|---|
| -10 | -16.61378867 | -18.6973143 | -29.70295603 | -16.5484201 | -17.07278234 | -14.69168486 |
| -5 | -19.12209971 | -16.09989018 | -17.27779482 | -9.518831616 | -30.60211763 | -11.31794842 |
| 0 | -22.08681882 | -20.09578957 | -12.0385111 | -17.1195561 | -19.42643098 | -20.70280243 |
| 5 | -18.61590374 | -15.791296 | -14.85435955 | -15.08882637 | -18.74843853 | -11.89100544 |
| 10 | -23.26356582 | -11.20642483 | -12.69898086 | -11.55952282 | -19.19724544 | -17.9691776 |
| 15 | -15.65026586 | -5.955756928 | -5.045882624 | -7.387621632 | -15.83997338 | -17.02843392 |
| 20 | -16.20068608 | 1.176392768 | 4.837203456 | -9.2146688 | -24.8018647 | -22.88244326 |
| 25 | -23.39330253 | 7.664882688 | 12.94976256 | -12.00318771 | -10.14735514 | -15.42831206 |
| 30 | -9.587675136 | 13.80743322 | 20.10187264 | -4.516891472 | -13.60274381 | -18.93371392 |
| 35 | -7.96535168 | 18.62227149 | 25.53630925 | 2.633602304 | -9.353751296 | -18.38449869 |
| 40 | -3.882922592 | 22.43405517 | 30.20599603 | 10.03048192 | -5.647394048 | -28.44390707 |
| 45 | 1.934013312 | 25.81721395 | 33.49317325 | 13.29099315 | -5.470170496 | -18.31625421 |
| 50 | 2.573619264 | 25.96092006 | 35.03631974 | 17.7729321 | 3.202734912 | -8.787328256 |
| 55 | 5.0978248 | 21.13095578 | 34.33674342 | 19.78335539 | 10.54276966 | -3.66294016 |
| 60 | 4.700080672 | 26.06391501 | 31.92613274 | 22.08820736 | 8.275179392 | 5.65623168 |
| 65 | 11.77416909 | 33.50180454 | 37.02258688 | 22.44813824 | 11.33379328 | 10.95288038 |
| 70 | 19.07127808 | 39.67147213 | 44.93717094 | 25.71877478 | 22.2437417 | 12.08129331 |
| 75 | 20.11331686 | 41.63500851 | 48.44839117 | 35.86578637 | 29.51753114 | 19.33110886 |

| F1 level (dB SPL) | 6k Homozygote mean | 12k Homozygote mean | 18k Homozygote mean | 24k Homozygote mean | 30k Homozygote mean | 36k Homozygote mean |
|---|---|---|---|---|---|---|
| -10 | -24.64104389 | -12.13527523 | -19.12844932 | -12.98715209 | -18.92710224 | -17.66517029 |
| -5 | -20.31748233 | -21.09528635 | -23.78229263 | -7.088762441 | -19.14404718 | -12.22393147 |
| 0 | -19.05633317 | -20.1486984 | -13.44034904 | -9.826304805 | -16.30070243 | -8.239865417 |
| 5 | -25.54375431 | -14.59747094 | -14.82250152 | -11.14441918 | -17.06694334 | -14.07142473 |
| 10 | -24.1726502 | -13.45779266 | -12.96929236 | -13.0066364 | -16.65766912 | -9.698553417 |
| 15 | -23.37994679 | -7.223519973 | -8.804589778 | -6.703108133 | -20.36164228 | -13.51971387 |
| 20 | -18.98451747 | 0.20873387 | 1.623842304 | -12.98244176 | -18.34721587 | -18.24704453 |
| 25 | -18.45713646 | 6.26727755 | 7.313984 | -7.532049408 | -13.42873322 | -14.10396877 |
| 30 | -8.986426761 | 12.11699463 | 13.03906494 | -5.987147282 | -10.92349213 | -22.48329787 |
| 35 | -6.045972361 | 16.85055269 | 17.58167976 | -3.527675904 | -20.99075365 | -15.08156687 |
| 40 | 0.801836279 | 20.65155306 | 21.93142257 | -6.25278603 | -12.2815838 | -13.75733292 |
| 45 | 4.59588245 | 23.98110983 | 24.03612423 | 1.299675877 | -11.78424565 | -10.53551419 |
| 50 | 6.741502903 | 24.9174645 | 25.84024854 | 3.264272585 | -10.67656671 | -7.215606089 |
| 55 | 7.419629349 | 25.40974343 | 25.01547813 | 8.004244718 | 3.27294219 | -10.15198947 |
| 60 | 7.577243282 | 25.51650977 | 23.60877861 | 2.184221897 | 8.641933605 | -10.23809192 |
| 65 | 11.98125005 | 30.35788756 | 27.23129929 | 6.679700306 | -0.170492489 | -2.986088594 |
| 70 | 15.96716924 | 36.83977743 | 38.27887397 | 5.961791378 | 3.107838519 | 0.867368997 |
| 75 | 20.23892641 | 40.7377054 | 38.93651368 | 16.51488582 | 11.03596134 | 7.536692005 |

| F1 level (dB SPL) | 6k Standard deviation | 12k Standard deviation | 18k Standard deviation | 24k Standard deviation | 30k Standard deviation | 36k Standard deviation |
|---|---|---|---|---|---|---|
| -10 | 7.363225055 | 0.399085365 | 4.202001846 | 5.947952892 | 6.669237469 | 2.768083243 |
| -5 | 4.086050273 | 8.841634591 | 4.832870886 | 1.010377526 | 19.41698477 | 2.038392127 |
| 0 | 5.589189222 | 5.186588461 | 0.813101929 | 2.962993279 | 6.675242967 | 9.586253285 |
| 5 | 10.11499628 | 1.604015957 | 5.550593727 | 1.271440716 | 9.289373609 | 3.625681173 |
| 10 | 5.890708061 | 4.81066814 | 0.04403766 | 3.236625003 | 14.61186922 | 0.935764252 |
| 15 | 10.32345055 | 5.18898751 | 0.76834345 | 0.639963813 | 4.835941698 | 0.869912314 |
| 20 | 5.748541991 | 0.859916516 | 1.79742498 | 0.309906551 | 10.17983958 | 2.015518524 |
| 25 | 3.239968068 | 0.363000247 | 1.85975573 | 3.246910522 | 0.189900145 | 1.433386406 |
| 30 | 1.234320167 | 0.144289065 | 2.118550386 | 5.725550612 | 1.810422892 | 9.937010108 |
| 35 | 4.632059258 | 0.035963474 | 2.223592286 | 0.77588725 | 2.98569419 | 0.503902802 |
| 40 | 6.703581046 | 0.76481828 | 1.836771343 | 0.141898161 | 2.669242703 | 5.684382222 |
| 45 | 4.204431307 | 1.056352093 | 0.788877922 | 0.539121199 | 2.48818119 | 7.410127889 |
| 50 | 6.207225348 | 1.371557442 | 0.252824638 | 0.847765684 | 4.350288089 | 8.081646993 |
| 55 | 5.729614564 | 0.724148304 | 0.24086288 | 0.620912252 | 1.739767425 | 13.52293799 |
| 60 | 8.130656485 | 3.209380145 | 1.910173959 | 0.062011432 | 5.967841305 | 5.826206165 |
| 65 | 0.945956364 | 0.341524113 | 0.495109606 | 1.590771858 | 4.108291032 | 3.703420651 |
| 70 | 0.292615584 | 2.676039255 | 1.688415136 | 1.372669625 | 1.023686067 | 3.874691372 |
| 75 | 2.566696335 | 0.466021971 | 1.030011607 | 0.79052737 | 1.479133613 | 2.661940021 |

| F1 level (dB SPL) | 6k Standard deviation | 12k Standard deviation | 18k Standard deviation | 24k Standard deviation | 30k Standard deviation | 36k Standard deviation |
|---|---|---|---|---|---|---|
| -10 | 7.407544065 | 3.807399573 | 5.688287094 | 3.462613459 | 4.159953311 | 5.97621344 |
| -5 | 9.407545617 | 4.462422683 | 9.797956845 | 5.952012603 | 1.253371377 | 2.957360882 |
| 0 | 8.356714093 | 4.83448585 | 4.863538457 | 2.095921491 | 4.479634416 | 4.957996073 |
| 5 | 6.298076723 | 3.720377462 | 3.895333045 | 3.578664575 | 5.69139776 | 4.05157113 |
| 10 | 6.976833762 | 3.817813577 | 3.209444311 | 4.939816415 | 4.101681179 | 5.096307106 |
| 15 | 5.160330386 | 4.38841455 | 7.333232203 | 2.912823852 | 2.858001191 | 3.394487431 |
| 20 | 8.215772979 | 4.202617061 | 6.986503667 | 4.677572868 | 8.859481244 | 5.124167187 |
| 25 | 10.65035084 | 3.3124644 | 9.373696265 | 2.34279177 | 4.352285782 | 5.067885247 |
| 30 | 6.513966653 | 2.287495821 | 13.24663434 | 5.832121472 | 3.949236102 | 6.454927601 |
| 35 | 5.179112454 | 1.58715276 | 14.88650346 | 2.394985104 | 5.666874859 | 7.879911061 |
| 40 | 3.424029631 | 1.541008808 | 13.34892955 | 11.62752809 | 7.664585209 | 3.789116005 |
| 45 | 3.007050299 | 1.51057899 | 12.32523761 | 6.469933924 | 8.461276422 | 4.328438881 |
| 50 | 3.003214728 | 1.582667974 | 9.66047634 | 8.339159994 | 8.953483123 | 2.653737033 |
| 55 | 3.979539267 | 1.173386507 | 5.909592302 | 6.252299654 | 7.058029822 | 5.883327987 |
| 60 | 6.699262697 | 3.007966984 | 4.878868609 | 12.15199843 | 6.877005218 | 9.080782789 |
| 65 | 2.720034987 | 4.010004643 | 11.91609209 | 8.11421065 | 5.561748876 | 9.90459868 |
| 70 | 2.207735293 | 2.037723448 | 7.360570836 | 12.67811549 | 9.325802205 | 7.899699643 |
| 75 | 3.341882941 | 1.406170191 | 7.564520097 | 15.84524308 | 7.304172258 | 6.322959939 |

| F2 (kHz) | Homozygote #1 | Homozygote #2 | Homozygote #3 | Homozygote #4 | Homozygote #5 | Homozygote #6 | Homozygote #7 |
|---|---|---|---|---|---|---|---|
| 6 | 30 | 30 | 65 | 35 | 30 | 30 | 30 |
| 12 | 15 | 15 | 20 | 20 | 10 | 15 | 15 |
| 18 | 25 | 10 | 20 | 20 | 15 | 45 | 15 |
| 24 | 45 | 30 | 55 | 45 | 25 | 75 | 65 |
| 30 | 50 | 65 | 75 | 70 | 55 | 75 | 70 |
| 36 | 75 | 75 | 75 | 70 | 75 | 75 | 75 |

| F1 level (dB SPL) | | 6k | 12k | 18k | 24k | 30k | 36k |
|---|---|---|---|---|---|---|---|
| Homozygote #1 | | | | | | | |
| | -10 | -27.64908749 | -17.60192512 | -30.25189478 | -19.67733965 | -26.23993856 | -21.12421683 |
| | -5 | -24.2576384 | -26.56876339 | -21.6035328 | -4.676018688 | -21.01763891 | -12.78771302 |
| | 0 | -21.59984435 | -21.16568883 | -20.7159767 | -10.34393907 | -10.7593513 | -9.403067392 |
| | 5 | -21.72766413 | -19.3460183 | -12.56960614 | -6.339152384 | -19.23704832 | -14.41727693 |
| | 10 | -22.50814874 | -14.37907866 | -11.54521805 | -8.306459648 | -15.93631334 | -8.546596352 |
| | 15 | -31.41717402 | -6.34570496 | -15.81900083 | -4.53679872 | -21.33934285 | -15.63427021 |
| | 20 | -25.92454451 | 0.799326784 | -4.697943552 | -5.810490368 | -14.85745766 | -25.46874982 |
| | 25 | -21.96344832 | 6.3648256 | 1.807489024 | -6.05570816 | -7.999814656 | -10.93112627 |
| | 30 | -10.84315546 | 12.50144768 | 12.86503936 | -18.5377792 | -10.27365376 | -29.76895795 |
| | 35 | -8.979267584 | 17.60586752 | 18.49234022 | -6.224278016 | -12.05562982 | -13.96796416 |
| | 40 | -1.153664768 | 22.02967245 | 23.50098227 | -7.334552064 | -8.066144768 | -11.45211904 |
| | 45 | 2.257275904 | 25.83692083 | 26.2098985 | -0.160440896 | -3.445157632 | -11.4497577 |
| | 50 | 3.830863104 | 25.81134131 | 29.57936435 | -2.122972544 | 2.844809472 | -8.991446016 |
| | 55 | 4.160482304 | 23.93554944 | 30.61346509 | 2.030913152 | 5.876500992 | -9.30765312 |
| | 60 | 4.330344448 | 24.3117527 | 27.6706304 | -8.268619776 | 10.19474125 | -12.37435597 |
| | 65 | 8.028916224 | 29.346304 | 16.89232998 | -0.907566528 | 4.9021056 | -4.220446208 |
| | 70 | 13.80660531 | 36.72254054 | 32.85520589 | -8.192881664 | -9.80929536 | 2.700506624 |
| | 75 | 18.33623757 | 41.95088794 | 40.89670451 | -5.353689088 | 5.203359744 | 9.209357312 |
| Homozygote #2 | | | | | | | |
| | -10 | -21.13138483 | -17.36061952 | -17.44407142 | -14.8686121 | -18.63124378 | -25.5429079 |
| | -5 | -21.02230426 | -24.57779405 | -27.57013709 | -20.10247578 | -19.59822336 | -16.46575206 |
| | 0 | -18.84531302 | -28.2188841 | -19.27703757 | -13.51184282 | -21.82382797 | -18.74471322 |
| | 5 | -27.97835469 | -15.69625088 | -19.73288755 | -10.44796826 | -28.86130074 | -22.63949722 |
| | 10 | -23.95919974 | -20.18596454 | -9.06528768 | -18.4944681 | -21.32151091 | -21.16297728 |
| | 15 | -22.54838374 | -7.761071616 | -1.697079168 | -12.3465257 | -18.45446451 | -16.01114419 |
| | 20 | -20.3343401 | 1.341296 | 8.543230976 | -17.83210394 | -15.69594061 | -23.65945446 |
| | 25 | -17.72602573 | 7.98037504 | 16.73245798 | -12.29572813 | -21.03390618 | -16.90468557 |
| | 30 | -6.174033408 | 12.5552384 | 23.3820631 | -6.997887488 | -18.39908864 | -28.41910272 |
| | 35 | -1.172211584 | 17.2298752 | 28.25880166 | 1.061471232 | -25.69802957 | -24.43596595 |
| | 40 | 4.359714816 | 21.19156122 | 30.94161818 | 6.583508992 | -24.73926451 | -11.5096832 |
| | 45 | 7.431392256 | 24.60707635 | 31.29461965 | 11.65487718 | -25.30100224 | -18.10485862 |
| | 50 | 9.370464256 | 27.20894566 | 29.26887526 | 15.60574771 | -16.78750208 | -9.73279232 |
| | 55 | 12.20730778 | 26.51193139 | 24.88085299 | 16.93569229 | -12.06807757 | -21.34088294 |
| | 60 | 14.02765722 | 19.57995315 | 27.97040435 | 15.28705843 | -6.691228672 | -23.91256883 |
| | 65 | 14.57093632 | 33.24716851 | 37.75107482 | 7.036681728 | -3.711981056 | -20.83728998 |
| | 70 | 17.298432 | 36.80934298 | 42.29288755 | 19.22810266 | 4.646132224 | -9.57631488 |
| | 75 | 21.64332134 | 40.43626496 | 39.76402534 | 31.94741555 | 12.79063347 | 6.079606784 |
| Homozygote #3 | | | | | | | |
| | -10 | -9.45421312 | -10.66257408 | -15.32304896 | -11.24837786 | -20.45213286 | -12.89746125 |
| | -5 | 0.452399776 | -23.39858432 | -11.89157888 | -6.541189632 | -20.2107945 | -8.287191552 |
| | 0 | -1.13256704 | -13.69655603 | -12.30180045 | -11.60362291 | -13.68286822 | -5.394006016 |
| | 5 | -15.84023552 | -9.032977408 | -13.73886157 | -15.60431206 | -15.96732928 | -12.51165696 |
| | 10 | -9.689995264 | -15.86094899 | -17.69066906 | -15.42034534 | -19.10662758 | -6.901038592 |
| | 15 | -14.88424858 | -10.74743603 | -10.88148378 | -7.6086784 | -17.17764506 | -10.67431117 |
| | 20 | -1.71563712 | 1.09052032 | 1.481405312 | -14.15571968 | -15.14250752 | -14.14672691 |
| | 25 | 3.416367872 | 6.967519744 | 6.018333184 | -6.716038144 | -13.68104346 | -15.25399245 |
| | 30 | -0.345875136 | 13.83885005 | 12.27000218 | -4.025946624 | -7.713544192 | -17.90732493 |
| | 35 | -11.42686618 | 18.0013609 | 18.30545613 | -3.687610112 | -26.49030042 | -11.18371226 |
| | 40 | -0.871789376 | 21.54847437 | 21.41894042 | -28.32821043 | -20.06679347 | -9.580819456 |
| | 45 | 4.895071744 | 24.17416192 | 21.58178714 | -4.359098368 | -21.84446771 | -13.09870285 |
| | 50 | 6.74439168 | 24.46757069 | 22.80034714 | -2.412376576 | -10.84694118 | -7.15451136 |
| | 55 | 3.031922176 | 26.30043034 | 21.2465664 | -0.567693824 | 3.02924672 | -7.124320256 |
| | 60 | -5.313637888 | 27.09086413 | 20.91173069 | -1.568806528 | 12.05151027 | -20.41111142 |
| | 65 | 12.23771546 | 27.68274022 | 15.89685965 | 0.713706752 | -9.955467264 | -2.964335616 |
| | 70 | 13.13553101 | 38.22842675 | 35.85143603 | -1.41259392 | 1.549217664 | -8.55704576 |
| | 75 | 13.75916442 | 39.88617626 | 33.48041523 | -0.252051648 | 3.363800576 | -5.005576704 |
| Homozygote #4 | | | | | | | |
| | -10 | -26.88052224 | -9.803859968 | -15.1481129 | -9.724545024 | -13.11725158 | -24.47127552 |
| | -5 | -20.73847603 | -14.39117005 | -20.18009088 | -2.863773952 | -19.05321984 | -10.63870157 |
| | 0 | -21.57037978 | -18.43607962 | -10.91384218 | -8.580215296 | -16.33270272 | -5.801374208 |
| | 5 | -24.37126144 | -16.93259571 | -9.792111616 | -7.815130624 | -12.77517107 | -11.82170112 |
| | 10 | -25.78901402 | -12.2184192 | -14.57758208 | -7.90568448 | -11.95261133 | -8.196502528 |
| | 15 | -22.63049421 | -14.12128768 | -11.72129485 | -7.728851968 | -20.72565555 | -16.74693837 |
| | 20 | -24.02490163 | -5.9858048 | -1.363753856 | -16.73222349 | -15.00101734 | -18.56268083 |
| | 25 | -29.72880077 | 1.499912576 | 7.611206656 | -7.793049088 | -11.80236186 | -9.244148736 |
| | 30 | -19.81551821 | 8.952556544 | 16.73662771 | -4.088012288 | -14.13169664 | -25.00691763 |
| | 35 | -12.32490189 | 15.06317619 | 21.93968128 | -2.799624704 | -21.45115136 | -27.05128038 |
| | 40 | -2.675719424 | 19.23188941 | 25.16010189 | -5.9644928 | -13.82978048 | -15.15744051 |
| | 45 | 1.981928576 | 22.45009203 | 25.73002138 | -1.59590656 | -11.31228979 | -11.26127411 |
| | 50 | 4.7136 | 23.62030285 | 25.98388941 | 5.195354112 | -9.5920128 | -2.563309312 |
| | 55 | 5.837298688 | 25.98975283 | 24.77848576 | 9.577406464 | 8.378008064 | -11.26217011 |
| | 60 | 7.166247936 | 28.83933594 | 24.75501158 | 10.6456023 | 9.175279616 | -2.867259136 |
| | 65 | 10.16225382 | 33.44559104 | 37.53304474 | 14.61556326 | -1.84105536 | 1.51998848 |
| | 70 | 16.36874957 | 37.22371482 | 43.62689741 | 18.38885274 | 12.74720666 | 11.27231078 |
| | 75 | 23.15367014 | 40.72194458 | 43.37328538 | 30.26284749 | 19.98682112 | 9.347672064 |
| Homozygote #5 | | | | | | | |
| | -10 | -27.67599002 | -10.79551795 | -22.39428813 | -13.56793446 | -16.80982221 | -14.52777779 |
| | -5 | -24.76326707 | -16.20651827 | -43.37665229 | -5.450250752 | -17.76549069 | -13.24653875 |
| | 0 | -21.26524416 | -23.60612659 | -10.10718925 | -8.492069376 | -11.88024013 | -4.913915392 |
| | 5 | -35.09405286 | -12.65528218 | -17.46223923 | -14.72694784 | -15.30450534 | -9.996862464 |
| | 10 | -27.3753129 | -10.51781734 | -9.762169856 | -14.68615578 | -11.6555776 | -7.20967424 |
| | 15 | -21.42897562 | -0.853128 | -3.520350208 | -5.536847872 | -22.97844736 | -8.03703808 |
| | 20 | -17.00086989 | 6.44968448 | 7.223341056 | -13.86924032 | -19.09573632 | -10.63040205 |
| | 25 | -16.23629414 | 11.18689178 | 13.77647718 | -8.111675392 | -9.516336128 | -15.64981658 |
| | 30 | -5.380273152 | 15.42171955 | 21.02391808 | -0.73151552 | -8.090686976 | -17.85627443 |
| | 35 | 0.3046832 | 19.12162509 | 26.17322496 | -3.2543872 | -25.48194304 | -6.972995072 |
| | 40 | 4.88415232 | 22.56891904 | 30.35726234 | 6.14163968 | -5.849782784 | -14.34796237 |
| | 45 | 8.268426752 | 25.75758336 | 33.35591526 | 9.352737792 | -8.304759808 | -5.800700416 |
| | 50 | 10.38869811 | 26.49870541 | 34.9690839 | 13.63994931 | -3.405088768 | -4.825409024 |
| | 55 | 12.19629978 | 23.79459379 | 31.0821417 | 14.35821978 | 6.72662528 | -2.987562496 |
| | 60 | 13.76071782 | 25.17064499 | 27.42581658 | 17.68611226 | 11.59822234 | -6.136561664 |
| | 65 | 16.17950822 | 32.58906624 | 37.7860055 | 16.75129139 | 6.3268992 | -8.341807616 |
| | 70 | 19.73828403 | 38.78601933 | 44.42139853 | 7.403080192 | -6.806765056 | 3.512897024 |
| | 75 | 23.1503319 | 41.73531546 | 46.99796275 | 27.24248166 | 5.122346496 | 9.945848832 |
| Homozygote #6 | | | | | | | |
| | -10 | -27.75374029 | -10.94221926 | -19.35982387 | -11.62535014 | -16.43858944 | -10.4292608 |
| | -5 | -26.32532378 | -19.99986074 | -21.9984896 | -3.023033088 | -17.54534707 | -14.81503437 |
| | 0 | -21.25222298 | -15.96342886 | -7.430185984 | -8.468157952 | -17.7974313 | -4.978439168 |
| | 5 | -30.57832755 | -17.33158912 | -11.48434432 | -9.400352768 | -15.17964698 | -13.96476928 |
| | 10 | -29.8431488 | -12.53794611 | -15.93460326 | -18.57157939 | -21.47923149 | -7.394358784 |
| | 15 | -23.1414784 | -7.242122752 | -18.53903872 | -5.615433728 | -24.57874637 | -11.55771085 |
| | 20 | -24.17234534 | -4.487868928 | -8.854540288 | -7.114545664 | -10.93265203 | -17.17789798 |
| | 25 | -21.6312279 | 2.49404544 | -10.03662029 | -5.008505856 | -13.66280192 | -22.63987814 |
| | 30 | -5.960902144 | 9.441886208 | -15.46843443 | -4.305919488 | -8.342243328 | -12.44597965 |
| | 35 | -1.69726464 | 14.79926579 | -15.07916595 | -3.968308992 | -14.66247168 | -7.377903616 |
| | 40 | 3.863694848 | 18.8027433 | -7.203997696 | -8.281459712 | -4.155806976 | -13.04489984 |
| | 45 | 6.719371776 | 22.68535808 | -2.228980736 | -2.087172224 | -5.933057536 | -6.98848768 |
| | 50 | 9.266531328 | 23.53988403 | 5.876779008 | -3.669132032 | -11.9508439 | -7.768297984 |
| | 55 | 10.15446016 | 26.37548339 | 14.22445466 | 6.630353408 | 3.369737216 | -6.41811968 |
| | 60 | 11.09805978 | 27.35905587 | 14.58934067 | -12.22886195 | 13.06172621 | -6.393871872 |
| | 65 | 11.70098586 | 22.84267725 | 11.39513037 | -3.655792128 | 0.95141312 | 3.71826816 |
| | 70 | 15.34211789 | 32.53834547 | 25.01743821 | -10.28451738 | 3.72790784 | -1.4099136 |
| | 75 | 21.76191898 | 38.21196902 | 24.63941018 | 5.778966016 | 9.430007808 | 7.364583424 |
| Homozygote #7 | | | | | | | |
| | -10 | -31.94236928 | -7.780210688 | -13.97790515 | -10.19790541 | -20.80073728 | -14.6632919 |
| | -5 | -25.56776653 | -22.5243136 | -19.85556685 | -6.9645952 | -18.81761587 | -9.326588928 |
| | 0 | -27.72876083 | -19.9541248 | -13.33641114 | -7.784286208 | -21.82849536 | -8.443542528 |
| | 5 | -23.216384 | -11.18758298 | -18.97746022 | -13.67707034 | -12.14360166 | -13.14820915 |
| | 10 | -30.04373197 | -8.50437376 | -12.20951654 | -7.661762048 | -15.15181158 | -8.478726144 |
| | 15 | -27.60887296 | -3.493888768 | 0.546119104 | -3.548620544 | -17.27719424 | -15.97658419 |
| | 20 | -19.71898368 | 2.253983232 | 9.03515648 | -15.3627689 | -37.70519962 | -18.08339968 |
| | 25 | -25.33052621 | 7.377372672 | 15.28854426 | -6.743641088 | -16.30486835 | -8.104133632 |
| | 30 | -14.38522982 | 12.107264 | 20.46423859 | -3.222970368 | -9.513531392 | -25.97852774 |
| | 35 | -7.025977856 | 16.13269811 | 24.98142003 | -5.820993536 | -21.09574963 | -14.58114662 |
| | 40 | -2.793534464 | 19.18761165 | 29.34505062 | -6.585935872 | -9.2635136 | -21.20840602 |
| | 45 | 0.617710144 | 22.35657626 | 32.30960845 | -3.707265792 | -6.348984832 | -7.04481792 |
| | 50 | 2.87597184 | 23.27550157 | 32.4034007 | -3.386661888 | -24.99838771 | -9.473476608 |
| | 55 | 4.34963456 | 24.96046285 | 28.28238029 | 7.06482176 | 7.598554624 | -12.62321766 |
| | 60 | 7.971313664 | 26.2639616 | 21.93851597 | -6.262931456 | 11.10328422 | 0.429085472 |
| | 65 | 10.98843443 | 33.35166566 | 33.36464998 | 12.20401766 | 2.134638336 | 10.22300262 |
| | 70 | 16.0804649 | 37.5700521 | 43.88685414 | 16.60249702 | 15.70046566 | 8.129142784 |
| | 75 | 19.86784051 | 42.22137958 | 43.40379238 | 25.97823078 | 21.35476019 | 15.81535232 |

**Latency data for Fig. S7**

Average

| | | 0dB | 5dB | 10dB | 15dB | 20dB | 25dB | 30dB | 35dB | 40dB | 45dB | 50dB | 55dB | 60dB | 65dB | 70dB | 75dB | 80dB | 85dB | 90dB | 95dB |
|---|---|---|---|---|---|---|---|---|---|---|---|---|---|---|---|---|---|---|---|---|---|
| Wildtype | 12 | 0.117143 | 0.044286 | 0.062857 | 0.351429 | 0.434286 | 0.547143 | 0.774286 | 0.817143 | 1.018571 | 1.36 | 1.632857 | 1.867143 | 2.42 | 2.671429 | 2.961429 | 3.155714 | 3.182857 | 3.565714 | 3.984286 | 3.944286 |
| Wildtype | 24 | | | 0.146 | 0.124 | 0.226 | 0.068 | 0.21 | 0.288 | 0.228 | 0.332 | 0.478 | 0.388 | 0.54 | 0.548 | 0.824 | 1.004 | 1.084 | 1.04 | 1.492 | 1.882 |
| Heterozygote | 12 | 0.198182 | 0.130909 | 0.183636 | 0.274545 | 0.408182 | 0.481818 | 0.564545 | 0.755455 | 0.850909 | 1.134545 | 1.289091 | 1.558182 | 1.832727 | 2.275455 | 2.635455 | 2.899091 | 3.09 | 3.561818 | 4.027273 | 4.037273 |
| Heterozygote | 24 | | | 0.115556 | 0.194444 | 0.146667 | 0.152222 | 0.237778 | 0.372222 | 0.552222 | 0.573333 | 0.781111 | 0.782222 | 1.066667 | 1.345556 | 1.561111 | 1.667778 | 1.743333 | 1.777778 | 2.02 | 2.295556 |

Standard deviation

| 0dB | 5dB | 10dB | 15dB | 20dB | 25dB | 30dB | 35dB | 40dB | 45dB | 50dB | 55dB | 60dB | 65dB | 70dB | 75dB | 80dB | 85dB | 90dB | 95dB |
|---|---|---|---|---|---|---|---|---|---|---|---|---|---|---|---|---|---|---|---|
| 0.180066 | 0.263746 | 0.144189 | 0.106055 | 0.178499 | 0.200143 | 0.378984 | 0.382087 | 0.41083 | 0.37372 | 0.524332 | 0.764847 | 0.654879 | 0.92485 | 1.188226 | 1.361382 | 1.126687 | 1.258939 | 1.467297 | 1.476323 |
| | | 0.120748 | 0.153069 | 0.209237 | 0.097826 | 0.187617 | 0.205597 | 0.255969 | 0.258592 | 0.375792 | 0.277435 | 0.345905 | 0.563977 | 0.805376 | 0.935136 | 1.036234 | 0.948683 | 1.106219 | 1.259829 |
| 0.083525 | 0.117427 | 0.186025 | 0.148685 | 0.156769 | 0.171103 | 0.205395 | 0.351009 | 0.229497 | 0.231187 | 0.29122 | 0.352415 | 0.429327 | 0.544323 | 0.457173 | 0.624731 | 0.649415 | 0.653342 | 0.845057 | 0.823348 |
| | | 0.068394 | 0.124007 | 0.141421 | 0.257137 | 0.158728 | 0.159121 | 0.245804 | 0.291204 | 0.15268 | 0.245091 | 0.372022 | 0.38112 | 0.450651 | 0.448352 | 0.431393 | 0.400805 | 0.420357 | 0.49576 |

| Mouse ID | *Mir183/96* [dko] genotype | Age | kHz | 0dB | 5dB | 10dB | 15dB | 20dB | 25dB | 30dB | 35dB | 40dB | 45dB | 50dB | 55dB | 60dB | 65dB | 70dB | 75dB | 80dB | 85dB | 90dB | 95dB |
|---|---|---|---|---|---|---|---|---|---|---|---|---|---|---|---|---|---|---|---|---|---|---|---|
| DMKO31.1d | Wildtype | P55 | 12 | 0.24 | 0.4 | -0.2 | -0.11 | 0.29 | 1.2 | 1.02 | 1.14 | 1.47 | 1.38 | 1.19 | 1.34 | 2.19 | 2.65 | 3.37 | 2.99 | 3.4 | 3.91 | 4.54 | 4.74 |
| DMKO31.1e | Wildtype | P55 | 12 | 0.11 | 0.18 | 0.46 | 0.52 | 0.59 | 0.99 | 0.84 | 1.38 | 1.32 | 1.52 | 1.74 | 2.33 | 2.97 | 3.57 | 4.55 | 4.59 | 5.16 | 5.03 | 5.66 | 5.57 |
| DMKO33.1a | Wildtype | P58 | 12 | 0 | 0.42 | 0.3 | 0.4 | 0.26 | 1.2 | 1.13 | 1.09 | 1.35 | 1.8 | 2.05 | 1.99 | 2.58 | 3.65 | 4.16 | 4.51 | 4.63 | 4.71 | 5.25 | 4.91 |
| DMKO33.1c | Wildtype | P58 | 12 | 0.11 | 0.54 | 0.3 | 0.33 | 0.4 | 0.92 | 1.57 | 1.46 | 1.45 | 1.91 | 2.12 | 2.3 | 2.34 | 2.99 | 3.68 | 3.95 | 4.35 | 4.38 | 4.91 | 4.72 |
| DMKO37.1h | Wildtype | P56 | 12 | 0.05 | 0.31 | 0.21 | 0.33 | 0.75 | 0.76 | 0.89 | 1.08 | 1.19 | 1.58 | 1.68 | 1.99 | 2.38 | 2.61 | 2.94 | 3.35 | 3.46 | 3.82 | 3.95 | 4.16 |
| DMKO37.1e | Wildtype | P56 | 12 | 0.01 | -0.07 | -0.07 | -0.12 | 0.56 | 0.31 | 0.73 | 0.98 | 0.94 | 1.13 | 1.72 | 1.81 | 2.33 | 2.47 | 2.51 | 3.21 | 3.58 | 4.05 | 4.57 | 4.69 |
| DMKO31.1b | Heterozygote | P55 | 12 | 0.14 | 0.24 | 0.18 | 0.31 | 0.74 | 0.26 | 0.5 | 1.11 | 1.29 | 1.44 | 1.41 | 1.84 | 1.72 | 2.12 | 2.04 | 2.7 | 2.76 | 2.76 | 2.81 | 3.16 |
| DMKO33.1b | Heterozygote | P58 | 12 | 0.33 | 0.14 | 0.07 | 0.18 | 0.34 | 0.33 | 0.61 | 0.46 | 0.96 | 1.09 | 1.08 | 1.58 | 1.78 | 1.97 | 2.12 | 2.23 | 2.14 | 2.49 | 2.27 | 2.26 |
| DMKO33.1d | Heterozygote | P58 | 12 | 0.16 | 0.2 | 0 | 0.9 | 0.58 | 0.45 | 0.98 | 1.74 | 1.18 | 1.57 | 1.53 | 1.67 | 1.84 | 2.36 | 2.97 | 4.11 | 4.18 | 5.06 | 5.9 | 5.73 |
| DMKO33.1e | Heterozygote | P58 | 12 | 0.26 | 0.74 | 0.18 | 0.68 | 0.19 | 1.05 | 1.2 | 1.14 | 1.19 | 0.77 | 1.36 | 2.17 | 1.99 | 2.12 | 3.22 | 4.03 | 4.02 | 4.62 | 5.09 | 5.22 |
| DMKO34.3c | Heterozygote | P56 | 12 | 0.18 | 0.24 | 0.1 | 0.19 | -0.06 | 0.02 | 0.32 | 0.21 | 0.7 | 0.91 | 0.95 | 1.27 | 1.16 | 1.23 | 1.28 | 1.77 | 2.32 | 2.4 | 2.84 | 3.15 |
| DMKO37.1a | Heterozygote | P56 | 12 | -0.01 | 0.29 | 0.05 | -0.09 | 0.36 | 0.28 | 0.68 | 0.41 | 0.69 | 0.91 | 0.93 | 1.13 | 1.52 | 1.95 | 2.2 | 2.48 | 2.7 | 3.34 | 3.72 | 3.45 |
| DMKO37.1b | Heterozygote | P56 | 12 | 0.29 | 0.07 | 0.36 | 0.36 | 0.36 | 0.66 | 0.79 | 1.06 | 1.3 | 1.39 | 1.28 | 1.86 | 1.68 | 1.84 | 2.38 | 3.09 | 3.47 | 3.8 | 4.38 | 5.03 |
| DMKO37.1c | Heterozygote | P56 | 12 | 0.22 | 0.42 | 0.35 | 0.14 | 0.49 | 0.54 | 0.9 | 0.94 | 1.12 | 1.17 | 1.29 | 1.41 | 1.98 | 2.22 | 3.5 | 3.36 | 3.96 | 4.56 | 5.41 | 5.53 |
| DMKO37.1g | Heterozygote | P56 | 12 | 0.11 | 0.34 | 0.47 | 0.47 | 0.43 | 0.72 | 0.69 | 0.48 | 0.88 | 0.96 | 1.11 | 1.11 | 1.7 | 1.89 | 2.03 | 2.72 | 3.04 | 3.54 | 3.77 | 3.9 |
| DMKO31.1d | Wildtype | P55 | 24 | | | 0.59 | 0.51 | -0.22 | 0.84 | 0.69 | 0.48 | 0.69 | 0.73 | 1.1 | 1.51 | 1.91 | 1.96 | 2.46 | 2.22 | 2.05 | 2.32 | 2.67 | 3.32 |
| DMKO31.1e | Wildtype | P55 | 24 | | | 0.27 | 0.11 | 0.01 | 0.13 | 0.38 | 0.68 | 0.97 | 1.11 | 1.27 | 1.76 | 2.69 | 2.77 | 2.8 | 2.9 | 3.38 | 3.42 | 3.47 | 3.95 |
| DMKO33.1a | Wildtype | P58 | 24 | | | -0.02 | 0.13 | 0.12 | 0.24 | -0.01 | 0.05 | -0.13 | 0.15 | 0.41 | 0.33 | 0.83 | 0.68 | 0.74 | 1.05 | 0.99 | 0.96 | 1.96 | 2.43 |
| DMKO33.1c | Wildtype | P58 | 24 | | | 0.15 | 0.23 | 0.08 | 0.36 | -0.19 | 0.51 | 0.51 | 0.89 | 0.97 | 1.15 | 1.26 | 1.04 | 1.06 | 1.63 | 1.77 | 2.16 | 3.06 | 3.56 |
| DMKO37.1h | Wildtype | P56 | 24 | | | 0 | 0.42 | 0.08 | 0.19 | 0.44 | 0.26 | 0.1 | 0.2 | 0.27 | 0.34 | 0.55 | 0.51 | 1.04 | 0.85 | 1.23 | 1.7 | 2.27 | 2.52 |
| DMKO37.1e | Wildtype | P56 | 24 | | | 0.14 | 0.07 | -0.1 | 0.21 | 0.32 | 0.39 | 0.37 | 0.24 | 0.42 | 0.48 | 0.8 | 0.85 | 1.01 | 1.07 | 1.17 | 1.38 | 1.55 | 2.02 |
| DMKO31.1b | Heterozygote | P55 | 24 | | | -0.19 | -0.01 | 0.39 | 0.02 | 0.22 | 0.33 | 0.75 | 0.42 | 0.61 | 0.45 | 0.72 | 0.76 | 1.15 | 1.39 | 1.5 | 1.99 | 2.39 | 2.69 |
| DMKO33.1b | Heterozygote | P58 | 24 | | | | | | | | | | | | | | | | | | | | |
| DMKO33.1d | Heterozygote | P58 | 24 | | | 0.34 | 0.19 | -0.05 | 0.39 | 0.5 | 0.69 | 1.1 | 0.99 | 1.47 | 1.28 | 1.96 | 2.18 | 2.3 | 2.41 | 2.57 | 2.81 | 2.95 | 3.14 |
| DMKO33.1e | Heterozygote | P58 | 24 | | | -0.03 | 0.14 | 0.29 | 0.29 | 0.8 | 0.67 | 0.69 | 0.97 | 0.71 | 1.05 | 2.1 | 1.97 | 1.93 | 2.21 | 2.29 | 2.23 | 2.77 | 3.45 |
| DMKO34.3c | Heterozygote | P56 | 24 | | | 0.08 | 0.09 | 0.2 | 0.42 | 0.28 | 0.01 | 0.26 | 0.07 | -0.11 | 0.13 | 0.52 | 0.59 | 0.84 | 1.1 | 1.17 | 1.43 | 1.5 | 1.88 |
| DMKO37.1a | Heterozygote | P56 | 24 | | | | | | | | | | | | | | | | | | | | |
| DMKO37.1b | Heterozygote | P56 | 24 | | | 0 | 0.09 | 0.15 | 0.19 | 0.32 | 0.5 | 0.61 | 0.87 | 0.99 | 1.15 | 1.39 | 1.59 | 1.43 | 1.58 | 1.83 | 1.83 | 2.37 | 2.43 |
| DMKO37.1c | Heterozygote | P56 | 24 | | | 0.27 | 0.41 | 0.07 | -0.14 | 0.5 | 0.39 | 0.75 | 1.08 | 1.24 | 1.3 | 1.37 | 1.63 | 2.12 | 2.11 | 2.92 | 2.88 | 3.13 | 3.29 |
| DMKO37.1g | Heterozygote | P56 | 24 | | | 0.13 | 0.24 | 0.26 | -0.1 | 0.32 | 0.5 | 0.61 | 0.49 | 0.69 | 0.79 | 0.72 | 1.11 | 0.99 | 1.5 | 1.84 | 1.57 | 2 | 2.31 |
| DMKO30.3e | Wildtype | P91 | 12 | 0.23 | 0.29 | -0.16 | 0.3 | 0.55 | 0.48 | 0.78 | 0.4 | 0.6 | 1.21 | 1.31 | 1.32 | 2.07 | 2.03 | 2.05 | 2.64 | 2.31 | 2.9 | 2.78 | 2.57 |
| DMKO31.1d | Wildtype | P90 | 12 | 0.06 | -0.38 | 0.11 | 0.25 | 0.12 | 0.28 | 0.47 | 0.46 | 0.66 | 1.09 | 1.32 | 1.72 | 1.91 | 2.37 | 2.52 | 2.55 | 3.01 | 3.55 | 4.22 | 3.84 |
| DMKO31.1e | Wildtype | P90 | 12 | 0.31 | 0.26 | 0.15 | 0.43 | 0.62 | 0.81 | 1.14 | 1.33 | 1.72 | 1.98 | 2.14 | 3.05 | 3.5 | 4.43 | 5.35 | 6.06 | 5.53 | 6.3 | 7.13 | 7.13 |
| DMKO33.1a | Wildtype | P89 | 12 | -0.24 | 0.19 | 0.14 | 0.45 | 0.46 | 0.76 | 1.3 | 1.14 | 1.29 | 1.61 | 1.86 | 2.46 | 2.59 | 2.89 | 3.27 | 3.24 | 3.31 | 3.29 | 3.41 | 3.43 |
| DMKO33.1c | Wildtype | P89 | 12 | 0.22 | -0.27 | 0.26 | 0.46 | 0.53 | 0.66 | 0.97 | 1.13 | 1.24 | 1.6 | 2.04 | 2.07 | 2.62 | 3.01 | 3.17 | 3.07 | 3.33 | 3.46 | 3.79 | 3.96 |
| DMKO37.1e | Wildtype | P90 | 12 | 0.16 | 0.08 | -0.07 | 0.38 | 0.5 | 0.37 | 0.45 | 0.77 | 0.87 | 1.02 | 2.03 | 1.76 | 2.75 | 2.45 | 2.63 | 2.74 | 2.53 | 2.96 | 3.56 | 3.47 |
| DMKO37.1h | Wildtype | P90 | 12 | 0.08 | 0.14 | 0.01 | 0.19 | 0.26 | 0.47 | 0.31 | 0.49 | 0.75 | 1.01 | 0.73 | 0.69 | 1.5 | 1.52 | 1.74 | 1.79 | 2.26 | 2.5 | 3 | 3.21 |
| DMKO30.3a | Heterozygote | P91 | 12 | 0.13 | 0.15 | 0.36 | 0.57 | 0.52 | 0.54 | 0.62 | 1.02 | 0.89 | 0.98 | 1.11 | 1.46 | 1.7 | 2.15 | 2.64 | 3.06 | 3.3 | 3.59 | 3.85 | 3.82 |
| DMKO30.3b | Heterozygote | P91 | 12 | 0.3 | -0.07 | -0.05 | 0.37 | 0.47 | 0.26 | 0.72 | 0.95 | 0.81 | 1.28 | 1.95 | 2.53 | 2.67 | 3.18 | 3.21 | 3.85 | 3.45 | 4.16 | 3.86 | 3.93 |
| DMKO31.1b | Heterozygote | P90 | 12 | 0.08 | 0.18 | 0.1 | 0.15 | 0.46 | 0.57 | 0.91 | 0.87 | 1.07 | 1.21 | 1.58 | 1.64 | 2.37 | 2.43 | 2.4 | 2.86 | 2.78 | 3.16 | 3.18 | 2.9 |
| DMKO33.1b | Heterozygote | P89 | 12 | 0.18 | 0.08 | 0.01 | 0.24 | 0.33 | 0.54 | 0.69 | 0.63 | 0.91 | 1.1 | 1.16 | 1.54 | 1.66 | 1.88 | 2.16 | 1.69 | 1.87 | 1.94 | 2.02 | 2.26 |
| DMKO33.1d | Heterozygote | P89 | 12 | 0.23 | 0.22 | 0.41 | 0.2 | 0.58 | 0.73 | 0.41 | 0.94 | 0.78 | 1.05 | 1.29 | 1.55 | 1.74 | 2.03 | 2.39 | 2.98 | 3.36 | 3.85 | 4.64 | 4.61 |
| DMKO33.1e | Heterozygote | P89 | 12 | 0.11 | 0.24 | 0.24 | 0.24 | 0.27 | 0.54 | 0.72 | 1.02 | 0.99 | 1.53 | 1.34 | 1.7 | 2.27 | 3.13 | 3.54 | 3.8 | 3.94 | 3.95 | 4.72 | 4.38 |
| DMKO37.1a | Heterozygote | P90 | 12 | 0.19 | 0.24 | 0.28 | 0.15 | 0.18 | 0.23 | 0.18 | -0.15 | 0.43 | 0.69 | 1.17 | 1.42 | 1.91 | 2.7 | 2.9 | 3.07 | 3.63 | 3.77 | 4.26 | 4.11 |
| DMKO37.1b | Heterozygote | P90 | 12 | 0.12 | 0.16 | -0.03 | 0.11 | 0.51 | 0.62 | 0.55 | 1.03 | 0.94 | 1.41 | 1.3 | 1.45 | 1.4 | 1.96 | 2.7 | 3.01 | 3.18 | 4.08 | 4.92 | 5.17 |
| DMKO37.1c | Heterozygote | P90 | 12 | 0.25 | 0.1 | 0.4 | 0.4 | 0.58 | 0.63 | 0.61 | 0.85 | 1.16 | 1.11 | 1.4 | 1.24 | 1.63 | 2.33 | 2.73 | 2.91 | 3.53 | 4.09 | 4.6 | 4.37 |
| DMKO37.1g | Heterozygote | P90 | 12 | 0.34 | -0.09 | -0.03 | 0.15 | 0.13 | 0.38 | 0.46 | 0.47 | 0.45 | 0.93 | 0.89 | 1.32 | 1.46 | 1.71 | 1.98 | 2.36 | 2.02 | 3.02 | 3.75 | 4.19 |
| DMKO37.3g | Heterozygote | P88 | 12 | 0.25 | 0.23 | 0.33 | 0.44 | 0.46 | 0.26 | 0.34 | 0.68 | 0.93 | 1.19 | 0.99 | 1.29 | 1.35 | 1.53 | 2.34 | 2.3 | 2.93 | 3.57 | 4.5 | 4.67 |
| DMKO30.3e | Wildtype | P91 | 24 | | | 0.29 | 0.27 | 0.37 | 0.02 | 0.5 | 0.54 | 0.58 | 0.71 | 0.86 | 0.71 | 0.76 | 1.03 | 1.58 | 1.24 | 1.57 | 1.15 | 1.72 | 1.55 |
| DMKO31.1d | Wildtype | P90 | 24 | | | 0.06 | 0 | 0.06 | 0 | 0.23 | 0.2 | 0 | 0.12 | 0.14 | 0.22 | 0.14 | -0.08 | 0 | 0.24 | 0.19 | 0.41 | 0.58 | 1.45 |
| DMKO31.1e | Wildtype | P90 | 24 | | | 0.24 | 0.31 | 0.5 | 0.23 | 0.22 | 0.41 | 0.4 | 0.38 | 0.9 | 0.66 | 0.99 | 1.2 | 1.71 | 2.52 | 2.57 | 2.54 | 3.22 | 4.03 |
| DMKO33.1a | Wildtype | P89 | 24 | | | 0.14 | 0 | 0.2 | 0 | 0.1 | 0.29 | 0 | 0.06 | 0.15 | 0.1 | 0.28 | 0.08 | 0.08 | 0.32 | 0.06 | 0.07 | 0.48 | 0.7 |
| DMKO33.1c | Wildtype | P89 | 24 | | | 0 | 0.04 | 0 | 0.09 | 0 | 0 | 0.16 | 0.39 | 0.34 | 0.25 | 0.53 | 0.51 | 0.75 | 0.7 | 1.03 | 1.03 | 1.46 | 1.68 |
| DMKO37.1e | Wildtype | P90 | 24 | | | | | | | | | | | | | | | | | | | | |
| DMKO37.1h | Wildtype | P90 | 24 | | | | | | | | | | | | | | | | | | | | |
| DMKO30.3a | Heterozygote | P91 | 24 | | | 0.12 | 0.03 | 0.12 | 0.18 | 0.16 | 0.24 | 0.48 | 0.97 | 0.78 | 1.21 | 1.27 | 1.94 | 2.2 | 2.36 | 2.57 | 2.57 | 2.18 | 2.64 |
| DMKO30.3b | Heterozygote | P91 | 24 | | | 0.15 | 0.1 | 0.06 | 0.17 | 0.36 | 0.51 | 0.34 | 0.76 | 0.59 | 0.93 | 1.04 | 1.35 | 1.49 | 1.73 | 1.71 | 1.76 | 1.84 | 2.18 |
| DMKO31.1b | Heterozygote | P90 | 24 | | | 0.09 | 0.25 | 0.49 | -0.01 | 0.09 | 0.32 | 0.18 | 0.11 | 0.81 | 0.56 | 0.51 | 0.84 | 0.81 | 0.94 | 1 | 1.04 | 1.18 | 1.16 |
| DMKO33.1b | Heterozygote | P89 | 24 | | | | | | | | | | | | | | | | | | | | |
| DMKO33.1d | Heterozygote | P89 | 24 | | | 0.15 | 0.13 | 0.08 | 0.16 | 0.08 | 0.34 | 0.42 | 0.26 | 0.76 | 0.91 | 1.04 | 1.56 | 1.62 | 1.5 | 1.5 | 1.73 | 1.56 | 2.03 |
| DMKO33.1e | Heterozygote | P89 | 24 | | | 0.04 | 0.24 | 0.09 | -0.38 | 0.08 | 0.54 | 0.62 | 0.76 | 0.88 | 0.97 | 1.9 | 1.72 | 2.1 | 2.29 | 1.94 | 1.6 | 2.09 | 2.53 |
| DMKO37.1a | Heterozygote | P90 | 24 | | | | | | | | | | | | | | | | | | | | |
| DMKO37.1b | Heterozygote | P90 | 24 | | | 0.1 | 0.43 | 0.2 | 0.5 | 0.33 | 0.51 | 1.03 | 0.34 | 0.79 | 0.6 | 1.02 | 1.44 | 1.52 | 1.61 | 1.74 | 1.87 | 2.2 | 2.22 |
| DMKO37.1c | Heterozygote | P90 | 24 | | | 0.23 | 0.3 | 0.11 | 0.46 | 0.43 | 0.55 | 0.75 | 0.83 | 0.84 | 0.81 | 0.99 | 1.41 | 1.86 | 1.8 | 2.05 | 2.02 | 2.49 | 2.84 |
| DMKO37.1g | Heterozygote | P90 | 24 | | | 0 | 0.09 | 0 | 0.11 | 0.14 | 0.2 | 0.62 | 0.47 | 0.53 | 0.46 | 0.86 | 0.93 | 1.09 | 1.44 | 1.67 | 1.73 | 2.33 | 2.59 |
| DMKO37.3g | Heterozygote | P88 | 24 | | | 0.16 | 0.18 | 0.17 | 0.18 | 0.47 | 0.14 | 0.53 | 0.66 | 1.05 | 0.59 | 0.97 | 0.92 | 1.36 | 1.34 | 1.51 | 1.68 | 2.31 | 2.47 |

**ABR thresholds at P56 for Fig. S8**

*Mir183/96*$^{dko}$ ABR thresholds (mean and standard deviation)

| Stimulus/frequency | Mean threshold (wildtype P56) | | Mean threshold (heterozygote P56) | | Bonferroni-corrected p-value for wildtype vs heterozygote, mixed linear model pairwise comparison |
|---|---|---|---|---|---|
| | Mean | SD | Mean | SD | |
| CLICK | 23.33333333 | 3.97911213 | 24.79166667 | 3.753018 | 0.703 |
| 3kHz | 57.85714286 | 7.99553447 | 61.45833333 | 9.495899 | 0.401 |
| 6kHz | 30.47619048 | 5.6799564 | 32.91666667 | 6.902531 | 0.564 |
| 12kHz | 11.9047619 | 5.35634915 | 16.25 | 7.408867 | 0.142 |
| 18kHz | 19.76190476 | 4.32324834 | 24.58333333 | 6.902531 | 0.215 |
| 24kHz | 29.76190476 | 9.41756212 | 34.16666667 | 13.80506 | 0.288 |
| 30kHz | 51.19047619 | 21.6739914 | 55.20833333 | 21.23774 | 0.33 |
| 36kHz | 68.0952381 | 23.7421666 | 81.25 | 18.07292 | 0.001 |
| 42kHz | 79.28571429 | 20.0801964 | 92.70833333 | 6.075461 | <0.001 |

| Mouse ID | Experiment date | Date of birth | Sex | *Mir183/96*$^{dko}$ genotype | Weight (g) | Age (days) | Age (weeks) | Click threshold | 3kHz threshold | 6kHz threshold | 12kHz threshold | 18kHz threshold | 24kHz threshold | 30kHz threshold | 36kHz threshold | 42kHz threshold |
|---|---|---|---|---|---|---|---|---|---|---|---|---|---|---|---|---|
| DMKO31.1d | 16/05/2014 | 22/03/2014 | F | Wildtype | 23.2 | 55 | 7.9 | 20 | 55 | 30 | 10 | 20 | 15 | 40 | 80 | 90 |
| DMKO31.1e | 16/05/2014 | 22/03/2014 | F | Wildtype | 20.5 | 55 | 7.9 | 20 | 45 | 25 | 10 | 20 | 20 | 40 | 70 | 95 |
| DMKO33.1a | 20/05/2014 | 23/03/2014 | M | Wildtype | 28.3 | 58 | 8.3 | 25 | 55 | 30 | 10 | 20 | 45 | 75 | 95 | 95 |
| DMKO33.1c | 20/05/2014 | 23/03/2014 | F | Wildtype | 21.6 | 58 | 8.3 | 15 | 55 | 25 | 10 | 15 | 25 | 40 | 95 | 95 |
| DMKO37.1e | 21/10/2014 | 26/08/2014 | M | Wildtype | 23.1 | 56 | 8.0 | 20 | 65 | 40 | 25 | 30 | 40 | 70 | 95 | 95 |
| DMKO37.1h | 21/10/2014 | 26/08/2014 | F | Wildtype | 22.1 | 56 | 8.0 | 30 | 65 | 30 | 5 | 20 | 50 | 95 | 95 | 95 |
| DMKO39.1c | 26/02/2015 | 01/01/2015 | M | Wildtype | 23.3 | 56 | 8.0 | 25 | 55 | 30 | 15 | 25 | 40 | 75 | 95 | 95 |
| DMKO39.1g | 26/02/2015 | 01/01/2015 | F | Wildtype | 19.1 | 56 | 8.0 | 30 | 70 | 45 | 15 | 25 | 45 | 95 | 95 | 95 |
| DMKO41.1f | 26/02/2015 | 01/01/2015 | M | Wildtype | 27.4 | 56 | 8.0 | 25 | 70 | 35 | 15 | 25 | 30 | 80 | 95 | 95 |
| DMKO44.1a | 07/09/2015 | 14/07/2015 | M | Wildtype | 25.7 | 55 | 7.9 | 30 | 55 | 30 | 10 | 20 | 35 | 60 | 95 | 95 |
| DMKO54.1f | 08/08/2016 | 14/06/2016 | F | Wildtype | 17.0 | 55 | 7.9 | 25 | 65 | 20 | 10 | 20 | 25 | 35 | 45 | 45 |
| DMKO55.1a | 08/08/2016 | 14/06/2016 | M | Wildtype | 24.0 | 55 | 7.9 | 20 | 60 | 25 | 0 | 20 | 20 | 30 | 35 | 45 |
| CNTC20.1a | 13/07/2015 | 21/05/2015 | M | Wildtype | 26.7 | 53 | 7.6 | 20 | 55 | 35 | 20 | 15 | 25 | 40 | 45 | 50 |
| CNTC20.1f | 13/07/2015 | 21/05/2015 | F | Wildtype | 20 | 53 | 7.6 | 25 | 55 | 35 | 15 | 20 | 30 | 45 | 50 | 60 |
| CNTC20.1g | 13/07/2015 | 21/05/2015 | F | Wildtype | 17.7 | 53 | 7.6 | 25 | 65 | 35 | 15 | 15 | 25 | 50 | 50 | 80 |
| DMKO44.1g | 07/09/2015 | 14/07/2015 | F | Wildtype | 21.4 | 55 | 7.9 | 25 | 60 | 30 | 10 | 15 | 25 | 40 | 50 | 95 |
| DMKO44.1c | 07/09/2015 | 14/07/2015 | M | Wildtype | 21.9 | 55 | 7.9 | 25 | 60 | 30 | 15 | 25 | 30 | 35 | 50 | 80 |
| DMKO54.1d | 08/08/2016 | 14/06/2016 | F | Wildtype | 19.8 | 55 | 7.9 | 25 | 40 | 25 | 10 | 15 | 25 | 35 | 55 | 65 |
| DMKO55.1b | 08/08/2016 | 14/06/2016 | M | Wildtype | 22.7 | 55 | 7.9 | 20 | 45 | 30 | 15 | 20 | 20 | 25 | 35 | 50 |
| DMKO55.1h | 08/08/2016 | 14/06/2016 | F | Wildtype | 17.4 | 55 | 7.9 | 20 | 65 | 25 | 10 | 15 | 30 | 30 | 45 | 55 |
| DMKO57.1b | 03/10/2016 | 09/08/2016 | M | Wildtype | 24.0 | 55 | 7.9 | 20 | 55 | 30 | 5 | 15 | 25 | 40 | 60 | 95 |
| DMKO31.1b | 16/05/2014 | 22/03/2014 | M | Heterozygote | 24.1 | 55 | 7.9 | 20 | 75 | 30 | 10 | 15 | 30 | 65 | 95 | 95 |
| DMKO33.1b | 20/05/2014 | 23/03/2014 | M | Heterozygote | 28.8 | 58 | 8.3 | 25 | 60 | 30 | 20 | 30 | 70 | 95 | 95 | 95 |
| DMKO33.1e | 20/05/2014 | 23/03/2014 | F | Heterozygote | 20.7 | 58 | 8.3 | 25 | 45 | 30 | 5 | 20 | 20 | 40 | 85 | 95 |
| DMKO33.1d | 20/05/2014 | 23/03/2014 | F | Heterozygote | 23.7 | 58 | 8.3 | 15 | 50 | 25 | 10 | 10 | 20 | 50 | 90 | 95 |
| DMKO37.1c | 21/10/2014 | 26/08/2014 | M | Heterozygote | 22.6 | 56 | 8.0 | 25 | 45 | 25 | 5 | 20 | 30 | 30 | 55 | 85 |
| DMKO37.1a | 21/10/2014 | 26/08/2014 | M | Heterozygote | 24.3 | 56 | 8.0 | 25 | 70 | 30 | 20 | 20 | 75 | 95 | 95 | 95 |
| DMKO37.1g | 21/10/2014 | 26/08/2014 | F | Heterozygote | 21 | 56 | 8.0 | 25 | 60 | 35 | 10 | 20 | 30 | 55 | 70 | 95 |
| DMKO34.3c | 03/12/2014 | 08/10/2014 | M | Heterozygote | 26.2 | 56 | 8.0 | 30 | 70 | 40 | 25 | 35 | 50 | 95 | 95 | 95 |
| DMKO39.1h | 26/02/2015 | 01/01/2015 | F | Heterozygote | 19.7 | 56 | 8.0 | 25 | 60 | 35 | 25 | 35 | 35 | 35 | 70 | 95 |
| DMKO41.1a | 26/02/2015 | 01/01/2015 | M | Heterozygote | 23.5 | 56 | 8.0 | 25 | 70 | 55 | 15 | 30 | 30 | 50 | 85 | 95 |
| DMKO41.1b | 26/02/2015 | 01/01/2015 | M | Heterozygote | 25.5 | 56 | 8.0 | 30 | 60 | 35 | 25 | 30 | 35 | 65 | 95 | 95 |
| DMKO37.1b | 21/10/2014 | 26/08/2014 | M | Heterozygote | 22.7 | 56 | 8.0 | 25 | 50 | 35 | 15 | 25 | 30 | 40 | 55 | 90 |
| DMKO44.1b | 07/09/2015 | 14/07/2015 | M | Heterozygote | 22.5 | 55 | 7.9 | 30 | 70 | 30 | 20 | 35 | 30 | 60 | 95 | 95 |
| DMKO44.1d | 07/09/2015 | 14/07/2015 | M | Heterozygote | 22.7 | 55 | 7.9 | 30 | 70 | 45 | 25 | 30 | 35 | 70 | 95 | 95 |
| DMKO54.1e | 08/08/2016 | 14/06/2016 | F | Heterozygote | 17.5 | 55 | 7.9 | 20 | 55 | 35 | 20 | 30 | 30 | 40 | 90 | 95 |
| DMKO55.1i | 08/08/2016 | 14/06/2016 | F | Heterozygote | 18.4 | 55 | 7.9 | 25 | 60 | 25 | 10 | 20 | 25 | 35 | 60 | 95 |
| DMKO57.1c | 03/10/2016 | 09/08/2016 | F | Heterozygote | 20.2 | 55 | 7.9 | 25 | 65 | 30 | 20 | 20 | 30 | 55 | 95 | 95 |
| DMKO57.1e | 03/10/2016 | 09/08/2016 | F | Heterozygote | 20.8 | 55 | 7.9 | 30 | 80 | 35 | 35 | 35 | 50 | 85 | 95 | 95 |
| DMKO44.1f | 07/09/2015 | 14/07/2015 | F | Heterozygote | 20.4 | 55 | 7.9 | 25 | 65 | 35 | 15 | 20 | 30 | 75 | 95 | 95 |
| DMKO44.1e | 07/09/2015 | 14/07/2015 | M | Heterozygote | 25.1 | 55 | 7.9 | 25 | 55 | 35 | 15 | 25 | 30 | 40 | 55 | 95 |
| DMKO54.1c | 08/08/2016 | 14/06/2016 | F | Heterozygote | 17.7 | 55 | 7.9 | 20 | 65 | 30 | 10 | 20 | 25 | 35 | 50 | 80 |
| DMKO55.1c | 08/08/2016 | 14/06/2016 | M | Heterozygote | 21.8 | 55 | 7.9 | 25 | 50 | 25 | 10 | 20 | 25 | 30 | 45 | 70 |
| DMKO55.1g | 08/08/2016 | 14/06/2016 | F | Heterozygote | 18.6 | 55 | 7.9 | 20 | 70 | 35 | 15 | 25 | 25 | 35 | 95 | 95 |
| DMKO57.1a | 03/10/2016 | 09/08/2016 | M | Heterozygote | 24.3 | 55 | 7.9 | 25 | 55 | 25 | 10 | 20 | 30 | 50 | 95 | 95 |

**qRTPCR Ct data for Fig. S11**

A) *Mir183/96* [dko]

| | Jag1 | Ccer2 | Slc52a3 | Grk1 | Grp | Myo3a | Ttc21a | BC030867 | Slc6a11 | Eln | Hspa2 | Tmem173 | Mfsd6 | Dtna | St8sia3 | Kif21b | P2rx3 | Tmc1 | Slc26a5 | Ocm |
|---|---|---|---|---|---|---|---|---|---|---|---|---|---|---|---|---|---|---|---|---|
| Wildtype average | 1.005481 | 1.007472425 | 1.03311408 | 1.006444 | 1.008277 | 1.019665 | 1.03155 | 1.014502 | 1.012597 | 1.002947 | 1.008174 | 1.008329 | 1.00436751 | 1.006095633 | 1.023311391 | 1.015956 | 1.00732 | 1.003119 | 1.005431 | 1.001528 |
| Homozygote average | 1.045032 | 5.515322009 | 4.924421877 | 4.740007 | 3.610864 | 3.376527 | 2.875462 | 2.467803 | 2.113168 | 2.059305 | 2.193834 | 1.807445 | 1.60935045 | 1.573913858 | 1.522939381 | 1.012014 | 0.658516 | 0.363632 | 0.320597 | 0.065459 |
| Wildtype standard deviation | 0.004368 | 0.005796808 | 0.040439808 | 0.007684 | 0.014791 | 0.036845 | 0.057549 | 0.022922 | 0.019741 | 0.002031 | 0.004929 | 0.01341 | 0.00394601 | 0.005363478 | 0.034415519 | 0.01917 | 0.013452 | 0.002425 | 0.006713 | 0.001687 |
| Homozygote standard deviation | 0.085378 | 3.498814743 | 2.2124677 | 4.094574 | 1.151317 | 1.667027 | 2.167747 | 0.926756 | 0.566731 | 1.304808 | 0.387214 | 0.328471 | 0.28495659 | 0.512060856 | 0.311023997 | 0.421186 | 0.434764 | 0.181457 | 0.178453 | 0.033972 |

B) *Mir182* [ko]

| | Jag1 | Ccer2 | Grp | Ppm1l | Slc26a5 | Ocm |
|---|---|---|---|---|---|---|
| Wildtype average | 1.008485 | 1.013499172 | 1.004850246 | 1.001611 | 1.005361 | 1.000644 |
| Homozygote average | 0.975165 | 1.640896639 | 1.581649817 | 1.11729 | 0.678917 | 0.657407 |
| Wildtype standard deviation | 0.016179 | 0.008289648 | 0.004434253 | 0.001748 | 0.006539 | 0.000441 |
| Homozygote standard deviation | 0.101639 | 0.611220972 | 0.208604576 | 0.280979 | 0.263848 | 0.214133 |

C) *Mir183/96* [dko]

| | Jag1 | Foxo1 | Ikzf2 | Cdkn1a | Sp1 | Nr3c1 | Tgfb1 | Fos | Foxo3 | Trp53 | Rest |
|---|---|---|---|---|---|---|---|---|---|---|---|
| Wildtype average | 1.002316 | 1.01928484 | 1.01414305 | 1.024584 | 1.015719 | 1.009408 | 1.015919 | 1.011069 | 1.020668 | 1.00513 | 1.01443 |
| Homozygote average | 1.013318 | 1.043187227 | 1.118276433 | 1.140098 | 1.203657 | 1.207774 | 1.224245 | 1.328771 | 1.424536 | 1.4806 | 1.520181 |
| Wildtype standard deviation | 0.003006 | 0.013062659 | 0.017484294 | 0.026406 | 0.014873 | 0.020859 | 0.022709 | 0.011332 | 0.025728 | 0.00406 | 0.02485 |
| Homozygote standard deviation | 0.119187 | 0.454319468 | 0.395424969 | 0.119516 | 0.258332 | 0.394734 | 0.233249 | 0.68684 | 0.366405 | 0.980021 | 0.824362 |

D) *Mir182* [ko]

| | Jag1 | Foxo3 | Trp53 |
|---|---|---|---|
| Wildtype average | 1.009846 | 1.018277649 | 1.010721442 |
| Homozygote average | 0.945848 | 1.161318931 | 1.491698685 |
| Wildtype standard deviation | 0.007327 | 0.018180215 | 0.013544322 |
| Homozygote standard deviation | 0.086052 | 0.207587438 | 1.050147422 |

| Gene | Mouse | *Mir183/96* [dko] genotype | Ct Replicate 1 | Ct Replicate 2 | Ct Replicate 3 | $2^{-ddCt}$ Replicate 1 | $2^{-ddCt}$ Replicate 2 | $2^{-ddCt}$ Replicate 3 | Average $2^{-ddC}$ | Standard deviation $2^{-ddCt}$ |
|---|---|---|---|---|---|---|---|---|---|---|
| Hprt | DMKO38.2b | Wildtype | 28.65449 | 28.56373 | 28.55193 | | | | | |
| Hprt | DMKO39.2f | Wildtype | 27.61983 | 27.68711 | 27.66412 | | | | | |
| Hprt | DMKO41.2d | Wildtype | 28.22992 | 28.25393 | 28.25211 | | | | | |
| Hprt | DMKO58.2f | Wildtype | 27.07676 | 26.55697 | 26.56961 | | | | | |
| Hprt | DMKO58.2h | Wildtype | 27.62372 | 27.22909 | 27.32687 | | | | | |
| Hprt | DMKO58.2g | Wildtype | 27.19318 | 27.26333 | 27.62018 | | | | | |
| Hprt | DMKO67.2b | Wildtype | 27.14409 | 27.19825 | 26.94221 | | | | | |
| Hprt | DMKO67.2b | Wildtype | 27.58209 | 27.3884 | 27.78502 | | | | | |
| Hprt | DMKO67.2b | Wildtype | 27.45715 | 26.96537 | 27.45595 | | | | | |
| Hprt | DMKO68.3a | Wildtype | 28.40713 | 27.71088 | 27.80468 | | | | | |
| Hprt | DMKO68.3c | Wildtype | 27.24363 | 27.27145 | 27.36257 | | | | | |
| Hprt | DMKO57.5c | Wildtype | 28.05228 | 27.70047 | 26.7566 | | | | | |
| Hprt | DMKO64.3d | Wildtype | 27.45231 | 27.4612 | 27.40158 | | | | | |
| Hprt | DMKO64.3g | Wildtype | 27.14047 | 27.17262 | 26.97001 | | | | | |
| Hprt | DMKO57.5c | Wildtype | 27.70923 | 27.74686 | 27.75536 | | | | | |
| Hprt | DMKO64.3d | Wildtype | 27.96971 | 28.162 | 27.91608 | | | | | |
| Hprt | DMKO57.5c | Wildtype | 27.70608 | 27.72333 | 27.46559 | | | | | |
| Hprt | DMKO64.3d | Wildtype | 28.04883 | 27.9313 | 27.47581 | | | | | |
| Hprt | DMKO67.2b | Wildtype | 26.93533 | 26.96548 | 26.81702 | | | | | |
| Hprt | DMKO64.3g | Wildtype | 28.22616 | 28.25739 | 28.29503 | | | | | |
| Hprt | DMKO64.3g | Wildtype | 28.11863 | 28.13544 | 28.07336 | | | | | |
| Hprt | DMKO64.3g | Wildtype | 27.68222 | 27.41399 | 27.38067 | | | | | |
| Hprt | DMKO57.5c | Wildtype | 27.43734 | 27.45647 | 27.60829 | | | | | |
| Hprt | DMKO72.1b | Wildtype | 26.93583 | 26.76998 | 26.69966 | | | | | |
| Hprt | DMKO72.1d | Wildtype | 26.92922 | 26.90076 | 26.64881 | | | | | |
| Hprt | DMKO64.5f | Wildtype | 26.68794 | 26.39857 | 26.51666 | | | | | |
| Hprt | DMKO72.1b | Wildtype | 27.33877 | 27.57915 | 27.22809 | | | | | |
| Hprt | DMKO72.1d | Wildtype | 27.33291 | 27.39105 | 27.48566 | | | | | |
| Hprt | DMKO84.3b | Wildtype | 28.30715 | 28.09015 | 27.84374 | | | | | |
| Hprt | DMKO84.3h | Wildtype | 28.30868 | 28.15989 | 28.3593 | | | | | |
| Hprt | DMKO84.3b | Wildtype | 29.26495 | 28.69223 | 29.40181 | | | | | |
| Hprt | DMKO84.3h | Wildtype | 29.75953 | 28.98486 | 29.51179 | | | | | |
| Hprt | DMKO79.5h | Wildtype | 28.49064 | 28.41735 | 28.34778 | | | | | |
| Hprt | DMKO79.7b | Wildtype | 28.83333 | 28.66846 | 28.6845 | | | | | |
| Hprt | DMKO79.5h | Wildtype | 30.04369 | 30.75275 | 30.63287 | | | | | |
| Hprt | DMKO79.7b | Wildtype | 30.32916 | 30.35471 | 30.14703 | | | | | |
| Hprt | DMKO84.4g | Wildtype | 28.48775 | 28.37004 | 28.65772 | | | | | |
| Hprt | DMKO84.4g | Wildtype | 32.08428 | 32.10975 | 32.2431 | | | | | |
| Hprt | DMKO84.3b | Wildtype | 29.34196 | 29.37716 | 30.24417 | | | | | |
| Hprt | DMKO84.3h | Wildtype | 28.18839 | 28.98182 | 28.39637 | | | | | |
| Hprt | DMKO79.5h | Wildtype | 28.50408 | 28.16057 | 28.63904 | | | | | |
| Hprt | DMKO79.7b | Wildtype | 28.33969 | 29.25135 | 29.32662 | | | | | |
| Hprt | DMKO84.4g | Wildtype | 28.29078 | 29.40119 | 29.15772 | | | | | |
| Hprt | DMKO57.5c | Wildtype | 27.40464 | 27.57983 | 27.51966 | | | | | |
| Hprt | DMKO84.3b | Wildtype | 29.76308 | 29.66631 | 29.23487 | | | | | |
| Hprt | DMKO84.3h | Wildtype | 28.819 | 28.99916 | 29.03204 | | | | | |
| Jag1 | DMKO38.2b | Wildtype | 30.18453 | 30.37494 | 30.17586 | 1.042885 | 0.913939 | 1.049171 | 1.001998 | 0.076326 |
| Jag1 | DMKO39.2f | Wildtype | 27.436 | 27.45442 | 27.37805 | 0.990906 | 0.978338 | 1.031522 | 1.000255 | 0.027797 |
| Jag1 | DMKO41.2d | Wildtype | 28.09524 | 27.95975 | 27.5748 | 0.859375 | 0.943993 | 1.232674 | 1.012014 | 0.195725 |
| Jag1 | DMKO58.2f | Wildtype | 26.03896 | 26.07059 | 25.98234 | 0.994243 | 0.972681 | 1.03404 | 1.000321 | 0.031128 |
| Jag1 | DMKO58.2h | Wildtype | 27.8503 | 27.64945 | 27.65777 | 0.913117 | 1.049515 | 1.043482 | 1.002038 | 0.077067 |
| Jag1 | DMKO58.2g | Wildtype | 27.58897 | 27.6053 | 27.61166 | 1.009053 | 0.9977 | 0.993313 | 1.000022 | 0.008123 |
| Jag1 | DMKO67.2b | Wildtype | 27.11436 | 26.71231 | 26.74764 | 0.837256 | 1.106341 | 1.079575 | 1.007724 | 0.148235 |
| Jag1 | DMKO67.2b | Wildtype | 27.11436 | 26.71231 | 26.74764 | 0.837256 | 1.106341 | 1.079575 | 1.007724 | 0.148235 |
| Jag1 | DMKO68.3a | Wildtype | 27.82571 | 26.93157 | 27.10509 | 0.688602 | 1.279772 | 1.134747 | 1.034374 | 0.308102 |
| Jag1 | DMKO68.3c | Wildtype | 26.7013 | 26.8463 | 26.7617 | 1.048601 | 0.948332 | 1.00561 | 1.000847 | 0.050304 |
| Jag1 | DMKO57.5c | Wildtype | 26.95803 | 26.95612 | 27.36847 | 1.098985 | 1.100447 | 0.826873 | 1.008769 | 0.157528 |
| Jag1 | DMKO57.5c | Wildtype | 26.95803 | 26.95612 | 27.36847 | 1.098985 | 1.100447 | 0.826873 | 1.008769 | 0.157528 |
| Jag1 | DMKO64.3d | Wildtype | 27.12913 | 27.36761 | 27.16579 | 1.065637 | 0.903274 | 1.038894 | 1.002602 | 0.087053 |
| Jag1 | DMKO64.3g | Wildtype | 26.42263 | 26.4505 | 26.32757 | 0.984597 | 0.96576 | 1.051653 | 1.00067 | 0.045146 |
| Jag1 | DMKO72.1b | Wildtype | 26.16011 | 26.25815 | 26.15098 | 1.020755 | 0.953693 | 1.027236 | 1.000561 | 0.040718 |
| Jag1 | DMKO72.1d | Wildtype | 26.21499 | 26.35766 | 25.97612 | 0.978019 | 0.885931 | 1.154125 | 1.006025 | 0.136273 |
| Jag1 | DMKO64.5f | Wildtype | 25.47948 | 25.58664 | 25.70669 | 1.080317 | 1.002983 | 0.922901 | 1.002067 | 0.078712 |
| Jag1 | DMKO84.3b | Wildtype | 26.29189 | 26.3753 | 26.48079 | 1.064935 | 1.005118 | 0.934243 | 1.001432 | 0.065424 |
| Jag1 | DMKO84.3h | Wildtype | 27.36836 | 27.47976 | 27.41642 | 1.037531 | 0.960433 | 1.003534 | 1.000499 | 0.038639 |
| Jag1 | DMKO79.5h | Wildtype | 27.19626 | 27.14839 | 27.13007 | 0.97399 | 1.006853 | 1.019717 | 1.000186 | 0.023581 |
| Jag1 | DMKO79.7b | Wildtype | 26.55694 | 26.3586 | 26.37437 | 0.915754 | 1.050716 | 1.039288 | 1.001919 | 0.07484 |
| Jag1 | DMKO84.4g | Wildtype | 26.20126 | 26.11426 | 26.25211 | 0.991683 | 1.053326 | 0.957337 | 1.000782 | 0.048637 |
| Ccer2 | DMKO41.2d | Wildtype | 30.28443 | 30.18028 | 30.42218 | 1.007791 | 1.08324 | 0.91602 | 1.00235 | 0.083743 |
| Ccer2 | DMKO67.2b | Wildtype | 31.03841 | 31.04556 | 30.75466 | 0.938091 | 0.933455 | 1.141989 | 1.004511 | 0.119081 |
| Ccer2 | DMKO57.5c | Wildtype | 32.55011 | 32.88577 | 33.15139 | 1.241698 | 0.983949 | 0.818487 | 1.014711 | 0.213276 |
| Ccer2 | DMKO64.3d | Wildtype | 31.28249 | 31.08549 | 31.40853 | 0.983739 | 1.12767 | 0.901442 | 1.004284 | 0.114504 |
| Ccer2 | DMKO64.3g | Wildtype | 31.33987 | 30.83389 | 30.76935 | 0.779797 | 1.107376 | 1.15804 | 1.015071 | 0.205322 |
| Ccer2 | DMKO72.1b | Wildtype | 29.93814 | 29.69004 | 29.64563 | 0.882577 | 1.048191 | 1.080954 | 1.003907 | 0.106345 |
| Grp | DMKO67.2b | Wildtype | 31.63535 | 31.99523 | 31.04287 | 0.947678 | 0.738457 | 1.42894 | 1.038358 | 0.35406 |
| Grp | DMKO57.5c | Wildtype | 32.13673 | 31.96987 | 32.06437 | 0.946226 | 1.062251 | 0.994897 | 1.001125 | 0.058262 |
| Grp | DMKO64.3d | Wildtype | 32.05445 | 31.77691 | 31.75582 | 0.875357 | 1.061043 | 1.076668 | 1.004356 | 0.11199 |
| Grp | DMKO64.3g | Wildtype | 32.56173 | 32.67293 | 32.52448 | 1.017233 | 0.941772 | 1.04384 | 1.000948 | 0.052946 |
| Grp | DMKO72.1b | Wildtype | 31.78894 | 31.61151 | 31.80666 | 0.963775 | 1.089897 | 0.952004 | 1.001892 | 0.076442 |
| Grp | DMKO72.1d | Wildtype | 31.13125 | 31.17437 | 31.38813 | 1.071772 | 1.040215 | 0.896963 | 1.002983 | 0.093162 |
| Grk1 | DMKO41.2d | Wildtype | 33.15539 | 33.54646 | 33.15656 | 1.094863 | 0.834898 | 1.093974 | 1.007911 | 0.149835 |
| Grk1 | DMKO67.2b | Wildtype | 32.54285 | 32.37772 | 32.25089 | 0.899775 | 1.008888 | 1.101597 | 1.00342 | 0.101022 |
| Grk1 | DMKO57.5c | Wildtype | 34.25124 | 34.97764 | 34.51653 | 1.2575 | 0.76005 | 1.046285 | 1.021278 | 0.249666 |
| Grk1 | DMKO64.3d | Wildtype | 34.07312 | 34.23113 | 34.29321 | 1.091291 | 0.97808 | 0.936882 | 1.002084 | 0.079954 |
| Grk1 | DMKO64.3g | Wildtype | 32.90565 | 33.19352 | 32.97277 | 1.085477 | 0.889127 | 1.036133 | 1.003579 | 0.102142 |
| Grk1 | DMKO72.1d | Wildtype | 31.62999 | 31.67181 | 31.72814 | 1.032865 | 1.003359 | 0.964939 | 1.000388 | 0.034061 |
| Tmc1 | DMKO41.2d | Wildtype | 32.20263 | 32.33488 | 32.63598 | 1.139603 | 1.039785 | 0.843923 | 1.00777 | 0.150417 |
| Tmc1 | DMKO67.2b | Wildtype | 31.26389 | 30.98793 | 31.05763 | 0.894565 | 1.083141 | 1.032056 | 1.003254 | 0.097532 |
| Tmc1 | DMKO57.5c | Wildtype | 32.41901 | 32.39014 | 32.59687 | 1.035023 | 1.055943 | 0.914976 | 1.001981 | 0.076071 |
| Tmc1 | DMKO64.3d | Wildtype | 32.88405 | 32.75922 | 32.94391 | 0.985102 | 1.074134 | 0.945063 | 1.001433 | 0.066067 |
| Tmc1 | DMKO64.3g | Wildtype | 31.97822 | 32.08951 | 32.14962 | 1.067495 | 0.988242 | 0.947918 | 1.001218 | 0.060836 |
| Tmc1 | DMKO72.1b | Wildtype | 31.38084 | 31.48466 | 31.65548 | 1.09138 | 1.0156 | 0.902197 | 1.003059 | 0.095213 |
| Hspa2 | DMKO41.2d | Wildtype | 30.61335 | 30.30042 | 30.20495 | 0.846485 | 1.051529 | 1.123465 | 1.00716 | 0.143722 |
| Hspa2 | DMKO67.2b | Wildtype | 30.49815 | 30.3161 | 29.94283 | 0.843352 | 0.956781 | 1.239306 | 1.013146 | 0.203906 |
| Hspa2 | DMKO57.5c | Wildtype | 29.65721 | 29.63886 | 29.72432 | 1.011331 | 1.024276 | 0.965361 | 1.000323 | 0.030962 |
| Hspa2 | DMKO64.3d | Wildtype | 30.98466 | 30.57911 | 30.757 | 0.863894 | 1.144313 | 1.011567 | 1.006591 | 0.140276 |
| Hspa2 | DMKO64.3g | Wildtype | 30.75126 | 30.32494 | 30.40431 | 0.836388 | 1.123941 | 1.063772 | 1.008034 | 0.151663 |
| Hspa2 | DMKO72.1d | Wildtype | 30.19966 | 29.68342 | 29.68029 | 0.787194 | 1.12587 | 1.128315 | 1.013793 | 0.196244 |
| Myo3a | DMKO41.2d | Wildtype | 35.09957 | 34.88467 | 35.11373 | 0.954677 | 1.108023 | 0.945355 | 1.002685 | 0.091344 |
| Myo3a | DMKO67.2b | Wildtype | 32.50246 | 32.76159 | 32.87892 | 1.158185 | 0.967765 | 0.892179 | 1.006043 | 0.137072 |
| Myo3a | DMKO57.5c | Wildtype | 35.63433 | 35.36374 | 35.3673 | 0.883188 | 1.06539 | 1.062767 | 1.003782 | 0.104445 |
| Myo3a | DMKO64.3d | Wildtype | 36.64275 | 36.87088 | 35.50748 | 0.810915 | 0.692307 | 1.781253 | 1.094825 | 0.597414 |
| Myo3a | DMKO64.3g | Wildtype | 33.24956 | 33.57753 | 33.47065 | 1.13526 | 0.904412 | 0.973954 | 1.004542 | 0.118425 |
| Myo3a | DMKO72.1b | Wildtype | 32.34849 | 32.15551 | 32.546 | 1.001049 | 1.144318 | 0.872968 | 1.006111 | 0.135746 |
| St8sia3 | DMKO41.2d | Wildtype | 31.44184 | 31.39661 | 31.54865 | 1.014329 | 1.046635 | 0.941946 | 1.00097 | 0.053608 |
| St8sia3 | DMKO67.2b | Wildtype | 30.62685 | 30.71422 | 30.46375 | 0.982655 | 0.924911 | 1.100269 | 1.002612 | 0.089366 |
| St8sia3 | DMKO57.5c | Wildtype | 31.30768 | 31.19479 | 30.36607 | 0.78377 | 0.847563 | 1.505356 | 1.045563 | 0.399468 |
| St8sia3 | DMKO64.3d | Wildtype | 31.30114 | 31.25967 | 30.09148 | 0.748959 | 0.7708 | 1.732209 | 1.083989 | 0.561481 |
| St8sia3 | DMKO64.3g | Wildtype | 31.09323 | 31.12572 | 31.09885 | 1.008844 | 0.986376 | 1.004925 | 1.000048 | 0.012001 |
| St8sia3 | DMKO72.1d | Wildtype | 29.87166 | 29.58478 | 29.97511 | 0.958503 | 1.169374 | 0.892181 | 1.006686 | 0.144742 |
| Tmem173 | DMKO67.2b | Wildtype | 30.77789 | 30.77857 | 30.47887 | 0.933392 | 0.932952 | 1.148356 | 1.0049 | 0.124237 |
| Tmem173 | DMKO57.5c | Wildtype | 31.01962 | 30.9096 | 30.82297 | 0.931593 | 1.005421 | 1.067642 | 1.001552 | 0.068107 |
| Tmem173 | DMKO64.3d | Wildtype | 31.38153 | 31.15348 | 31.04587 | 0.877879 | 1.028219 | 1.107847 | 1.004648 | 0.116782 |
| Tmem173 | DMKO64.3g | Wildtype | 31.42186 | 31.43967 | 31.31641 | 0.979956 | 0.967931 | 1.054264 | 1.000717 | 0.046761 |
| Tmem173 | DMKO72.1b | Wildtype | 29.41228 | 29.70179 | 30.34591 | 1.326584 | 1.085381 | 0.694517 | 1.035494 | 0.318973 |
| Tmem173 | DMKO72.1d | Wildtype | 29.89157 | 29.68161 | 29.65369 | 0.901703 | 1.042956 | 1.063336 | 1.002665 | 0.088027 |
| Slc52a3 | DMKO67.2b | Wildtype | 33.73807 | 32.81538 | 32.76663 | 0.645561 | 1.22375 | 1.265815 | 1.045042 | 0.3466 |
| Slc52a3 | DMKO57.5c | Wildtype | 34.08005 | 33.7583 | 33.7045 | 0.851197 | 1.063867 | 1.104288 | 1.006451 | 0.135964 |
| Slc52a3 | DMKO64.3d | Wildtype | 33.89269 | 33.11461 | 33.0331 | 0.684967 | 1.174619 | 1.242893 | 1.034159 | 0.30433 |
| Slc52a3 | DMKO64.3g | Wildtype | 34.19437 | 33.8845 | 34.08055 | 0.906745 | 1.124002 | 0.981177 | 1.003975 | 0.110408 |
| Slc52a3 | DMKO72.1b | Wildtype | 33.0251 | 31.6364 | 31.46857 | 0.506367 | 1.325889 | 1.489455 | 1.107237 | 0.526756 |
| Slc52a3 | DMKO72.1d | Wildtype | 32.21798 | 32.01265 | 32.16153 | 0.94131 | 1.085281 | 0.97887 | 1.00182 | 0.074679 |
| Kif21b | DMKO67.2b | Wildtype | 32.35884 | 31.84219 | 31.22254 | 0.682555 | 0.976485 | 1.500365 | 1.053135 | 0.414258 |
| Kif21b | DMKO57.5c | Wildtype | 33.02939 | 33.20622 | 33.06653 | 1.050681 | 0.929477 | 1.023978 | 1.001379 | 0.063684 |
| Kif21b | DMKO64.3d | Wildtype | 33.83798 | 33.50055 | 33.35698 | 0.827707 | 1.045808 | 1.155238 | 1.009584 | 0.166743 |
| Kif21b | DMKO64.3g | Wildtype | 32.78496 | 32.47887 | 32.45042 | 0.862416 | 1.066253 | 1.087485 | 1.005384 | 0.124269 |
| Kif21b | DMKO72.1b | Wildtype | 30.28746 | 30.4109 | 30.00589 | 0.964122 | 0.885064 | 1.171908 | 1.007031 | 0.148158 |
| Kif21b | DMKO72.1d | Wildtype | 31.09026 | 31.52177 | 30.8309 | 1.040578 | 0.771571 | 1.245516 | 1.019222 | 0.237694 |
| Mfsd6 | DMKO67.2b | Wildtype | 29.24338 | 29.67825 | 29.53086 | 1.181633 | 0.874128 | 0.968149 | 1.00797 | 0.157572 |
| Mfsd6 | DMKO57.5c | Wildtype | 29.86934 | 29.51641 | 29.98285 | 0.946184 | 1.208423 | 0.874592 | 1.009733 | 0.175755 |
| Mfsd6 | DMKO64.3d | Wildtype | 30.15003 | 30.14664 | 30.4664 | 1.074994 | 1.07752 | 0.863314 | 1.005276 | 0.12295 |
| Mfsd6 | DMKO64.3g | Wildtype | 30.39676 | 30.15384 | 30.24411 | 0.912654 | 1.080023 | 1.01452 | 1.002399 | 0.08434 |
| Mfsd6 | DMKO72.1b | Wildtype | 29.45518 | 29.40792 | 29.53494 | 1.007536 | 1.041089 | 0.953347 | 1.000658 | 0.044274 |
| Mfsd6 | DMKO72.1d | Wildtype | 29.51206 | 29.55445 | 29.57583 | 1.024834 | 0.995157 | 0.980516 | 1.000169 | 0.02258 |
| BC030867 | DMKO67.2b | Wildtype | 30.89148 | 30.78696 | 30.89205 | 0.976269 | 1.04962 | 0.975885 | 1.000591 | 0.04246 |
| BC030867 | DMKO57.5c | Wildtype | 30.74742 | 30.97256 | 30.72588 | 1.048167 | 0.896718 | 1.063932 | 1.002939 | 0.092327 |
| BC030867 | DMKO64.3d | Wildtype | 30.46859 | 30.60202 | 30.78107 | 1.108521 | 1.010598 | 0.892643 | 1.003921 | 0.108094 |
| BC030867 | DMKO64.3g | Wildtype | 31.74179 | 31.53838 | 31.58478 | 0.9201 | 1.059415 | 1.025885 | 1.0018 | 0.072713 |
| BC030867 | DMKO72.1b | Wildtype | 31.06692 | 29.85083 | 30.28717 | 0.630567 | 1.464907 | 1.082577 | 1.05935 | 0.417655 |
| BC030867 | DMKO72.1d | Wildtype | 29.69575 | 30.21396 | 30.3138 | 1.300217 | 0.907862 | 0.847158 | 1.018412 | 0.24593 |
| Eln | DMKO67.2b | Wildtype | 31.53917 | 31.25773 | 31.50791 | 0.9303 | 1.130694 | 0.950674 | 1.00389 | 0.110287 |
| Eln | DMKO57.5c | Wildtype | 32.20869 | 32.30182 | 32.11461 | 0.99978 | 0.937279 | 1.067153 | 1.001404 | 0.064952 |
| Eln | DMKO64.3d | Wildtype | 32.14819 | 32.20694 | 31.84614 | 0.945337 | 0.907614 | 1.165501 | 1.00615 | 0.139284 |
| Eln | DMKO64.3g | Wildtype | 31.67535 | 31.67156 | 31.56444 | 0.973846 | 0.976412 | 1.051664 | 1.00064 | 0.044206 |
| Eln | DMKO72.1b | Wildtype | 30.9357 | 31.12566 | 31.11715 | 1.089603 | 0.955179 | 0.96083 | 1.001871 | 0.076031 |
| Eln | DMKO72.1d | Wildtype | 31.47223 | 31.16816 | 31.33471 | 0.903002 | 1.114866 | 0.993318 | 1.003729 | 0.106315 |
| P2rx3 | DMKO67.2b | Wildtype | 29.27173 | 29.32901 | 29.22694 | 1.002888 | 0.963852 | 1.034516 | 1.000419 | 0.035396 |
| P2rx3 | DMKO57.5c | Wildtype | 32.02866 | 32.10827 | 32.24814 | 1.071547 | 1.014021 | 0.920326 | 1.001965 | 0.076328 |
| P2rx3 | DMKO64.3d | Wildtype | 31.58471 | 31.36956 | 31.19523 | 0.869621 | 1.009476 | 1.139132 | 1.006076 | 0.134788 |
| P2rx3 | DMKO64.3g | Wildtype | 30.17877 | 30.09123 | 30.1173 | 0.966157 | 1.026596 | 1.008214 | 1.000322 | 0.030982 |
| P2rx3 | DMKO72.1b | Wildtype | 29.21069 | 28.5728 | 29.44706 | 0.911403 | 1.418188 | 0.773669 | 1.03442 | 0.339413 |
| P2rx3 | DMKO72.1d | Wildtype | 28.87928 | 28.98207 | 29.0043 | 1.054046 | 0.981559 | 0.96655 | 1.000718 | 0.046789 |
| Ttc21a | DMKO67.2b | Wildtype | 33.99881 | 33.55774 | 33.64191 | 0.831628 | 1.129027 | 1.065042 | 1.008565 | 0.156536 |
| Ttc21a | DMKO57.5c | Wildtype | 34.17649 | 33.76692 | 32.44354 | 0.609555 | 0.809666 | 2.026196 | 1.148472 | 0.766688 |
| Ttc21a | DMKO64.3d | Wildtype | 33.58269 | 33.4668 | 33.75503 | 1.013129 | 1.097872 | 0.89905 | 1.00335 | 0.099771 |
| Ttc21a | DMKO64.3g | Wildtype | 33.44197 | 34.00116 | 33.93965 | 1.276584 | 0.866397 | 0.904136 | 1.015706 | 0.226714 |
| Ttc21a | DMKO72.1b | Wildtype | 32.1625 | 32.64662 | 32.14659 | 1.114247 | 0.796613 | 1.126603 | 1.012488 | 0.187055 |
| Ttc21a | DMKO72.1d | Wildtype | 28.87928 | 28.98207 | 29.0043 | 1.054046 | 0.981559 | 0.96655 | 1.000718 | 0.046789 |
| Slc6a11 | DMKO57.5c | Wildtype | 32.60035 | 33.07581 | 33.17561 | 1.274776 | 0.916861 | 0.855584 | 1.01574 | 0.226414 |
| Slc6a11 | DMKO64.3d | Wildtype | 34.4994 | 33.41 | 33.5986 | 0.631388 | 1.343502 | 1.178869 | 1.051253 | 0.372815 |
| Slc6a11 | DMKO67.2b | Wildtype | 32.16826 | 31.95087 | 32.11182 | 0.938691 | 1.091349 | 0.976143 | 1.002061 | 0.079561 |
| Slc6a11 | DMKO64.3g | Wildtype | 33.6421 | 33.63966 | 33.6662 | 1.005017 | 1.006721 | 0.988365 | 1.000034 | 0.010142 |
| Slc6a11 | DMKO72.1b | Wildtype | 32.4471 | 32.56643 | 32.30893 | 0.995659 | 0.916617 | 1.095726 | 1.002667 | 0.08976 |
| Slc6a11 | DMKO72.1d | Wildtype | 32.38721 | 32.58811 | 32.68892 | 1.123139 | 0.977141 | 0.911191 | 1.003824 | 0.108464 |
| Dtna | DMKO67.2b | Wildtype | 30.51637 | 30.20213 | 30.55489 | 0.938285 | 1.166616 | 0.91356 | 1.006154 | 0.139513 |
| Dtna | DMKO57.5c | Wildtype | 30.26108 | 30.73006 | 30.70351 | 1.234392 | 0.891818 | 0.908387 | 1.011532 | 0.19318 |
| Dtna | DMKO64.3d | Wildtype | 30.60675 | 30.53252 | 30.46136 | 0.95052 | 1.000712 | 1.051308 | 1.000846 | 0.050394 |
| Dtna | DMKO64.3g | Wildtype | 31.30451 | 30.73056 | 30.91581 | 0.800582 | 1.191736 | 1.048127 | 1.013482 | 0.197865 |
| Dtna | DMKO72.1b | Wildtype | 29.92555 | 29.76528 | 30.0606 | 0.99419 | 1.111005 | 0.905346 | 1.003514 | 0.103146 |
| Dtna | DMKO72.1d | Wildtype | 29.77657 | 29.83851 | 29.67857 | 0.991704 | 0.950028 | 1.061405 | 1.001046 | 0.056273 |
| Ocm | DMKO38.2b | Wildtype | 27.37797 | 27.52657 | 27.37096 | 1.033255 | 0.932127 | 1.038286 | 1.001223 | 0.059891 |
| Ocm | DMKO39.2f | Wildtype | 27.65162 | 27.70675 | 27.60203 | 1.001282 | 0.963741 | 1.036295 | 1.000439 | 0.036284 |
| Ocm | DMKO41.2d | Wildtype | 29.26785 | 29.19595 | 29.11311 | 0.948981 | 0.997476 | 1.056428 | 1.000962 | 0.053808 |
| Ocm | DMKO67.2b | Wildtype | 27.51367 | 27.24413 | 27.56597 | 0.951047 | 1.146413 | 0.917184 | 1.004882 | 0.123734 |
| Ocm | DMKO67.2b | Wildtype | 27.51367 | 27.24413 | 27.56597 | 0.951047 | 1.146413 | 0.917184 | 1.004882 | 0.123734 |
| Ocm | DMKO64.3d | Wildtype | 28.00847 | 28.02742 | 27.86437 | 0.971497 | 0.958823 | 1.073545 | 1.001288 | 0.062897 |
| Ocm | DMKO64.3g | Wildtype | 28.11934 | 28.20237 | 28.11763 | 1.018966 | 0.961979 | 1.020175 | 1.000373 | 0.033256 |
| Slc26a5 | DMKO38.2b | Wildtype | 35.02212 | 34.77209 | 35.43646 | 1.038693 | 1.235246 | 0.779398 | 1.017779 | 0.228642 |
| Slc26a5 | DMKO39.2f | Wildtype | 32.81283 | 32.97109 | 32.57956 | 0.98282 | 0.880708 | 1.155298 | 1.006275 | 0.13879 |
| Slc26a5 | DMKO41.2d | Wildtype | 33.73436 | 33.65879 | 33.351 | 0.899393 | 0.947759 | 1.173147 | 1.006766 | 0.146105 |
| Slc26a5 | DMKO67.2b | Wildtype | 31.49571 | 31.4993 | 31.39936 | 0.978796 | 0.976362 | 1.046398 | 1.000519 | 0.039751 |
| Slc26a5 | DMKO67.2b | Wildtype | 31.49571 | 31.4993 | 31.39936 | 0.978796 | 0.976362 | 1.046398 | 1.000519 | 0.039751 |
| Slc26a5 | DMKO64.3d | Wildtype | 32.6913 | 32.58419 | 32.56087 | 0.946597 | 1.019549 | 1.03616 | 1.000769 | 0.047644 |
| Slc26a5 | DMKO64.3g | Wildtype | 31.16838 | 31.27458 | 31.19754 | 1.03177 | 0.958546 | 1.011124 | 1.00048 | 0.037755 |
| Nr3c1 | DMKO64.3g | Wildtype | 30.73126 | 30.78571 | 31.00442 | 1.078632 | 1.038682 | 0.892573 | 1.003296 | 0.097947 |
| Nr3c1 | DMKO57.5c | Wildtype | 31.1274 | 31.17375 | 31.25406 | 1.040782 | 1.007879 | 0.953305 | 1.000655 | 0.044184 |
| Nr3c1 | DMKO72.1b | Wildtype | 28.42977 | 28.43651 | 28.42017 | 0.99934 | 0.994682 | 1.00601 | 1.000011 | 0.005694 |
| Nr3c1 | DMKO72.1d | Wildtype | 28.37523 | 28.34712 | 28.40686 | 1.000813 | 1.020507 | 0.97911 | 1.000143 | 0.020707 |
| Nr3c1 | DMKO84.3b | Wildtype | 32.07067 | 31.97033 | 32.04129 | 0.970474 | 1.040374 | 0.990436 | 1.000428 | 0.036006 |
| Nr3c1 | DMKO84.3h | Wildtype | 31.22272 | 30.16978 | 30.97451 | 0.740353 | 1.536048 | 0.879339 | 1.051913 | 0.424993 |
| Fos | DMKO84.3b | Wildtype | 32.1891 | 32.7083 | 32.72568 | 1.27626 | 0.890524 | 0.879863 | 1.015549 | 0.225845 |
| Fos | DMKO84.3h | Wildtype | 34.16832 | 34.38467 | 34.21327 | 1.062233 | 0.914308 | 1.029645 | 1.002062 | 0.077724 |
| Fos | DMKO79.5h | Wildtype | 35.32495 | 36.09562 | 35.96112 | 1.384097 | 0.811282 | 0.890557 | 1.028645 | 0.310372 |
| Fos | DMKO79.7b | Wildtype | 33.81935 | 34.01155 | 33.888 | 1.062118 | 0.929646 | 1.012767 | 1.00151 | 0.06695 |
| Fos | DMKO84.4g | Wildtype | 36.76333 | 36.33849 | 36.61633 | 0.876233 | 1.176276 | 0.970222 | 1.007577 | 0.15347 |
| Foxo1 | DMKO84.3b | Wildtype | 28.67429 | 28.80707 | 28.45831 | 0.980961 | 0.894704 | 1.13938 | 1.005015 | 0.124099 |
| Foxo1 | DMKO84.3h | Wildtype | 29.01787 | 28.34093 | 28.48066 | 0.755384 | 1.207667 | 1.096188 | 1.019746 | 0.235632 |
| Foxo1 | DMKO79.5h | Wildtype | 29.60728 | 30.07573 | 29.95513 | 1.207562 | 0.872757 | 0.948849 | 1.009723 | 0.175507 |
| Foxo1 | DMKO79.7b | Wildtype | 29.84325 | 29.0883 | 29.31308 | 0.743099 | 1.254035 | 1.073109 | 1.023414 | 0.259068 |
| Foxo1 | DMKO84.4g | Wildtype | 31.19572 | 30.46104 | 30.23761 | 0.676299 | 1.125386 | 1.313892 | 1.038526 | 0.327551 |
| Foxo3 | DMKO79.5h | Wildtype | 30.99277 | 31.13555 | 31.0228 | 1.040736 | 0.94267 | 1.019295 | 1.0009 | 0.051556 |
| Foxo3 | DMKO79.7b | Wildtype | 30.09465 | 29.93741 | 29.5586 | 0.851988 | 0.950094 | 1.235377 | 1.012487 | 0.199164 |
| Foxo3 | DMKO84.4g | Wildtype | 32.1113 | 32.28711 | 31.62108 | 0.929933 | 0.82324 | 1.306237 | 1.019803 | 0.25373 |
| Foxo3 | DMKO84.3b | Wildtype | 30.16914 | 30.09033 | 30.44003 | 1.04538 | 1.104073 | 0.866418 | 1.005291 | 0.123796 |
| Foxo3 | DMKO84.3h | Wildtype | 29.36301 | 30.31861 | 30.49661 | 1.620458 | 0.83555 | 0.738567 | 1.064858 | 0.483601 |
| Ikzf2 | DMKO84.3b | Wildtype | 30.52567 | 30.53295 | 30.53668 | 1.004234 | 0.99918 | 0.9966 | 1.000005 | 0.003883 |
| Ikzf2 | DMKO84.3h | Wildtype | 32.43371 | 32.36364 | 32.1452 | 0.920488 | 0.966299 | 1.124269 | 1.003685 | 0.106911 |
| Ikzf2 | DMKO79.5h | Wildtype | 33.03057 | 33.00634 | 32.91408 | 0.968012 | 0.984401 | 1.049415 | 1.000609 | 0.043054 |
| Ikzf2 | DMKO79.7b | Wildtype | 31.31521 | 31.18008 | 30.50593 | 0.803962 | 0.882903 | 1.408808 | 1.031891 | 0.328797 |
| Ikzf2 | DMKO84.4g | Wildtype | 33.8496 | 33.09851 | 32.96679 | 0.685566 | 1.153851 | 1.264157 | 1.034525 | 0.307199 |

| Gene | Mouse ID | *Mir182* [ko] genotype | Ct Replicate 1 | Ct Replicate 2 | Ct Replicate 3 | $2^{-ddCt}$ Replicate 1 | $2^{-ddCt}$ Replicate 2 | $2^{-ddCt}$ Replicate 3 | Average $2^{-ddC}$ | Standard deviation $2^{-ddCt}$ |
|---|---|---|---|---|---|---|---|---|---|---|
| Hprt | DMOM54.1d | Wildtype | 26.14841 | 26.19626 | 26.09924 | | | | | |
| Hprt | DMOM52.3a | Wildtype | 27.15403 | 27.03504 | 27.18665 | | | | | |
| Hprt | DMOM54.1d | Wildtype | 26.71876 | 26.90545 | 27.08586 | | | | | |
| Hprt | DMOM50.4c | Wildtype | 26.23787 | 26.22742 | 26.51878 | | | | | |
| Hprt | DMOM50.4g | Wildtype | 27.35753 | 27.2917 | 27.8006 | | | | | |
| Hprt | DMOM46.3a | Wildtype | 28.03483 | 27.94598 | 28.0563 | | | | | |
| Hprt | DMOM46.3b | Wildtype | 27.87051 | 28.02845 | 27.75346 | | | | | |
| Hprt | DMOM50.4c | Wildtype | 25.75084 | 25.69805 | 26.18007 | | | | | |
| Hprt | DMOM50.4g | Wildtype | 26.60113 | 26.39601 | 26.60486 | | | | | |
| Hprt | DMOM52.3a | Wildtype | 29.62052 | 27.69259 | 27.96158 | | | | | |
| Hprt | DMOM54.1d | Wildtype | 27.26237 | 26.9743 | 27.04224 | | | | | |
| Hprt | DMOM50.4c | Wildtype | 27.58713 | 26.51554 | 26.36686 | | | | | |
| Hprt | DMOM50.4g | Wildtype | 27.79914 | 27.68033 | 26.74189 | | | | | |
| Hprt | DMOM46.3a | Wildtype | 31.0735 | 30.63691 | 30.76636 | | | | | |
| Hprt | DMOM46.3b | Wildtype | 31.12016 | 31.03339 | 31.11023 | | | | | |
| Hprt | DMOM55.3c | Wildtype | 26.66119 | 26.29266 | 26.61846 | | | | | |
| Hprt | DMOM55.1e | Wildtype | 26.73371 | 26.70854 | 26.70537 | | | | | |
| Hprt | DMOM55.1e | Wildtype | 26.98497 | 27.1803 | 27.0051 | | | | | |
| Hprt | DMOM60.3g | Wildtype | 28.36831 | 27.52606 | 28.98774 | | | | | |
| Hprt | DMOM64.4g | Wildtype | 29.08284 | 28.55286 | 28.63271 | | | | | |
| Hprt | DMOM64.3j | Wildtype | 27.0571 | 27.50674 | 27.1076 | | | | | |
| Hprt | DMOM64.3k | Wildtype | 27.89768 | 28.09013 | 27.95354 | | | | | |
| Hprt | DMOM60.3g | Wildtype | 29.68915 | 30.11913 | 30.24427 | | | | | |
| Hprt | DMOM64.4g | Wildtype | 29.20975 | 28.92574 | 29.79115 | | | | | |
| Hprt | DMOM64.3j | Wildtype | 26.72104 | 27.32246 | 27.25836 | | | | | |
| Hprt | DMOM64.3k | Wildtype | 28.13574 | 28.52435 | 28.48747 | | | | | |
| Jag1 | DMOM54.1d | Wildtype | 25.40928 | 25.70821 | 25.6002 | 1.11983 | 0.910264 | 0.981026 | 1.003707 | 0.106608 |
| Jag1 | DMOM52.3a | Wildtype | 27.87731 | 27.839 | 27.84433 | 0.983665 | 1.010131 | 1.00641 | 1.000069 | 0.014327 |
| Jag1 | DMOM50.4c | Wildtype | 25.5248 | 25.66475 | 25.5181 | 1.031268 | 0.935923 | 1.036068 | 1.001086 | 0.056484 |
| Jag1 | DMOM50.4g | Wildtype | 26.48132 | 26.75843 | 27.48496 | 1.344357 | 1.109421 | 0.670485 | 1.041421 | 0.342044 |
| Jag1 | DMOM46.3a | Wildtype | 27.49265 | 27.34216 | 27.31863 | 0.927763 | 1.029769 | 1.046702 | 1.001411 | 0.064341 |
| Jag1 | DMOM46.3b | Wildtype | 27.37436 | 27.33083 | 27.43742 | 1.004525 | 1.035291 | 0.961561 | 1.000459 | 0.037033 |
| Jag1 | DMOM55.3c | Wildtype | 27.92942 | 27.98965 | 28.02244 | 1.036044 | 0.993679 | 0.97135 | 1.000358 | 0.03286 |
| Jag1 | DMOM55.1e | Wildtype | 26.67297 | 26.56424 | 26.40947 | 0.917589 | 0.989417 | 1.101469 | 1.002825 | 0.09267 |
| Jag1 | DMOM55.1e | Wildtype | 26.67297 | 26.56424 | 26.40947 | 0.917589 | 0.989417 | 1.101469 | 1.002825 | 0.09267 |
| Jag1 | DMOM55.1e | Wildtype | 26.67297 | 26.56424 | 26.40947 | 0.917589 | 0.989417 | 1.101469 | 1.002825 | 0.09267 |
| Jag1 | DMOM60.3g | Wildtype | 25.99046 | 26.46083 | 25.96663 | 1.108685 | 0.800222 | 1.127149 | 1.012019 | 0.183654 |
| Jag1 | DMOM60.3g | Wildtype | 25.99046 | 26.46083 | 25.96663 | 1.108685 | 0.800222 | 1.127149 | 1.012019 | 0.183654 |
| Jag1 | DMOM64.4g | Wildtype | 27.609 | 27.50865 | 27.14518 | 0.87779 | 0.941017 | 1.210632 | 1.009813 | 0.176764 |
| Jag1 | DMOM64.4g | Wildtype | 27.609 | 27.50865 | 27.14518 | 0.87779 | 0.941017 | 1.210632 | 1.009813 | 0.176764 |
| Jag1 | DMOM64.3j | Wildtype | | 25.69074 | 25.69312 | | 1.000826 | 0.999175 | 1 | 0.001168 |
| Jag1 | DMOM64.3k | Wildtype | 26.61623 | 27.26715 | 27.00985 | 1.272954 | 0.810711 | 0.968995 | 1.017553 | 0.234916 |
| Ccer2 | DMOM52.3a | Wildtype | 33.20763 | 32.67981 | 31.95737 | 0.663104 | 0.956028 | 1.577422 | 1.065518 | 0.466889 |
| Ccer2 | DMOM54.1d | Wildtype | 30.25723 | 29.60096 | 30.2054 | 0.849075 | 1.338145 | 0.880138 | 1.022453 | 0.273839 |
| Ccer2 | DMOM50.4c | Wildtype | 31.08822 | 30.77205 | 31.02866 | 0.916851 | 1.141497 | 0.955491 | 1.004613 | 0.120109 |
| Ccer2 | DMOM50.4g | Wildtype | 30.72798 | 31.11439 | 31.13694 | 1.20174 | 0.919366 | 0.90511 | 1.008738 | 0.167296 |
| Ccer2 | DMOM46.3a | Wildtype | 35.00272 | 34.5838 | 34.24784 | 0.762461 | 1.019353 | 1.286642 | 1.022819 | 0.262108 |
| Ccer2 | DMOM46.3b | Wildtype | 34.44548 | 33.89781 | 34.44863 | 0.881784 | 1.288921 | 0.879855 | 1.016854 | 0.235619 |
| Ccer2 | DMOM55.1e | Wildtype | 31.53582 | 31.23944 | 31.18819 | 0.861744 | 1.058271 | 1.096541 | 1.005519 | 0.125974 |
| Grp | DMOM54.1d | Wildtype | 31.10299 | 31.37894 | 31.20072 | 1.090179 | 0.900382 | 1.018768 | 1.00311 | 0.095862 |
| Grp | DMOM52.3a | Wildtype | 32.25992 | 32.12395 | 32.15455 | 0.945763 | 1.039237 | 1.017427 | 1.000809 | 0.048903 |
| Grp | DMOM50.4c | Wildtype | 31.34363 | 31.30589 | 31.33451 | 0.989233 | 1.015449 | 0.995504 | 1.000062 | 0.013689 |
| Grp | DMOM50.4g | Wildtype | 31.80773 | 32.00733 | 32.30878 | 1.175724 | 1.023811 | 0.830758 | 1.010098 | 0.172891 |
| Grp | DMOM46.3a | Wildtype | 32.0731 | 31.90238 | 31.59899 | 0.861581 | 0.969813 | 1.196785 | 1.009393 | 0.171071 |
| Grp | DMOM46.3b | Wildtype | 31.46348 | 31.48017 | 31.52254 | 1.017654 | 1.005954 | 0.976836 | 1.000148 | 0.021019 |
| Grp | DMOM55.1e | Wildtype | 32.09122 | 32.36815 | 32.47055 | 1.163729 | 0.960477 | 0.894667 | 1.006291 | 0.140259 |
| Ocm | DMOM54.1d | Wildtype | 27.35832 | 27.45432 | 27.34722 | 1.019811 | 0.954156 | 1.027688 | 1.000551 | 0.040372 |
| Ocm | DMOM52.3a | Wildtype | 24.0548 | 24.1316 | 24.16616 | 1.044434 | 0.990288 | 0.966847 | 1.000523 | 0.039793 |
| Ocm | DMOM50.4c | Wildtype | 25.32775 | 25.17541 | 25.24205 | 0.946487 | 1.051899 | 1.00441 | 1.000932 | 0.052792 |
| Ocm | DMOM50.4g | Wildtype | 26.48482 | 26.42514 | 26.39272 | 0.965541 | 1.006316 | 1.029188 | 1.000348 | 0.032241 |
| Ocm | DMOM46.3a | Wildtype | 26.67401 | 26.489 | 26.57702 | 0.936921 | 1.065119 | 1.002073 | 1.001371 | 0.064102 |
| Ocm | DMOM46.3b | Wildtype | 26.54209 | 26.48659 | 26.53234 | 0.985037 | 1.023669 | 0.991717 | 1.000141 | 0.020648 |
| Ppm1l | DMOM54.1d | Wildtype | 27.76723 | 27.74191 | 27.72612 | 0.984768 | 1.002205 | 1.013233 | 1.000069 | 0.014352 |
| Ppm1l | DMOM52.3a | Wildtype | 27.99434 | 27.66026 | 27.94864 | 0.915994 | 1.154674 | 0.94547 | 1.00538 | 0.13013 |
| Ppm1l | DMOM50.4c | Wildtype | 27.99407 | 28.05472 | 27.87873 | 0.987443 | 0.946792 | 1.069629 | 1.001288 | 0.062578 |
| Ppm1l | DMOM50.4g | Wildtype | 28.73045 | 28.40436 | 28.43953 | 0.867137 | 1.08705 | 1.060871 | 1.005019 | 0.120125 |
| Ppm1l | DMOM46.3a | Wildtype | 30.01462 | 29.87882 | 29.84668 | 0.932226 | 1.024242 | 1.047312 | 1.00126 | 0.060888 |
| Ppm1l | DMOM46.3b | Wildtype | 30.46282 | 30.34207 | 30.27602 | 0.931407 | 1.01272 | 1.060159 | 1.001429 | 0.065115 |
| Ppm1l | DMOM55.1e | Wildtype | 29.27794 | 29.22767 | 29.15622 | 0.961039 | 0.99512 | 1.045642 | 1.000601 | 0.042567 |
| Slc26a5 | DMOM54.1d | Wildtype | 30.93993 | 30.5862 | 30.26954 | 0.789289 | 1.008604 | 1.256155 | 1.018016 | 0.233575 |
| Slc26a5 | DMOM52.3a | Wildtype | 29.19758 | 29.04002 | 28.97496 | 0.915906 | 1.021603 | 1.068727 | 1.002079 | 0.078259 |
| Slc26a5 | DMOM50.4c | Wildtype | 29.18809 | 29.1711 | 29.52917 | 1.077754 | 1.090525 | 0.850834 | 1.006371 | 0.134851 |
| Slc26a5 | DMOM50.4g | Wildtype | 30.1836 | 30.28731 | 30.19069 | 1.02593 | 0.954767 | 1.020905 | 1.000534 | 0.039715 |
| Slc26a5 | DMOM46.3a | Wildtype | 30.91171 | 30.73566 | 30.86354 | 0.949514 | 1.072748 | 0.98175 | 1.001337 | 0.063909 |
| Slc26a5 | DMOM46.3b | Wildtype | 31.97466 | 31.70474 | 31.96438 | 0.937308 | 1.130157 | 0.944015 | 1.003827 | 0.109456 |
| Foxo3 | DMOM60.3g | Wildtype | 29.86983 | 29.45343 | 28.83538 | 0.715185 | 0.954475 | 1.464932 | 1.044864 | 0.382959 |
| Foxo3 | DMOM60.3g | Wildtype | 29.86983 | 29.45343 | 28.83538 | 0.715185 | 0.954475 | 1.464932 | 1.044864 | 0.382959 |
| Foxo3 | DMOM64.4g | Wildtype | 28.74861 | 29.34929 | 28.98846 | 1.214343 | 0.800792 | 1.028346 | 1.014493 | 0.207123 |
| Foxo3 | DMOM64.4g | Wildtype | 28.74861 | 29.34929 | 28.98846 | 1.214343 | 0.800792 | 1.028346 | 1.014493 | 0.207123 |
| Foxo3 | DMOM64.3j | Wildtype | 28.49721 | 28.03779 | 28.13471 | 0.827037 | 1.137168 | 1.063287 | 1.009164 | 0.161994 |
| Foxo3 | DMOM64.3k | Wildtype | 29.13781 | 29.47569 | 29.31559 | 1.126532 | 0.891315 | 0.995921 | 1.00459 | 0.117848 |
| Trp53 | DMOM60.3g | Wildtype | 29.49399 | 29.74066 | 29.71395 | 1.113842 | 0.938788 | 0.956333 | 1.002987 | 0.096403 |
| Trp53 | DMOM60.3g | Wildtype | 29.49399 | 29.74066 | 29.71395 | 1.113842 | 0.938788 | 0.956333 | 1.002987 | 0.096403 |
| Trp53 | DMOM64.4g | Wildtype | 30.31176 | 31.04743 | 31.03183 | 1.399825 | 0.840649 | 0.849789 | 1.030088 | 0.320235 |
| Trp53 | DMOM64.4g | Wildtype | 30.31176 | 31.04743 | 31.03183 | 1.399825 | 0.840649 | 0.849789 | 1.030088 | 0.320235 |
| Trp53 | DMOM64.3j | Wildtype | 29.08395 | 29.10348 | 29.09026 | 1.005991 | 0.992459 | 1.001598 | 1.000016 | 0.006903 |
| Trp53 | DMOM64.3k | Wildtype | 29.86842 | 30.09714 | 30.36301 | 1.181896 | 1.00862 | 0.838867 | 1.009794 | 0.171518 |
| Hprt | DMOM54.1c | Homozygote | 25.04048 | 25.25753 | 25.31346 | | | | | |
| Hprt | DMOM52.3g | Homozygote | 26.70874 | 26.60121 | 26.8809 | | | | | |
| Hprt | DMOM54.1c | Homozygote | 25.87349 | 25.98086 | 26.05728 | | | | | |
| Hprt | DMOM50.4e | Homozygote | 26.56974 | 26.31716 | 26.74572 | | | | | |
| Hprt | DMOM50.4a | Homozygote | 26.72024 | 26.44968 | 26.59585 | | | | | |
| Hprt | DMOM46.3d | Homozygote | 27.80695 | 27.67345 | 27.83338 | | | | | |
| Hprt | DMOM46.3g | Homozygote | 28.18409 | 28.30339 | 28.27247 | | | | | |
| Hprt | DMOM50.4e | Homozygote | 26.15469 | 26.06134 | 26.09448 | | | | | |
| Hprt | DMOM50.4a | Homozygote | 26.25367 | 26.14156 | 26.3241 | | | | | |
| Hprt | DMOM52.3g | Homozygote | 26.64358 | 26.05621 | 26.83537 | | | | | |
| Hprt | DMOM54.1c | Homozygote | 26.82115 | 27.24319 | 26.32646 | | | | | |
| Hprt | DMOM50.4e | Homozygote | 26.79548 | 25.83286 | 26.86509 | | | | | |
| Hprt | DMOM50.4a | Homozygote | 28.01923 | 27.32953 | 27.55432 | | | | | |
| Hprt | DMOM46.3d | Homozygote | 31.99244 | 31.4353 | 31.51541 | | | | | |
| Hprt | DMOM46.3g | Homozygote | 32.07618 | 31.00969 | 31.34781 | | | | | |
| Hprt | DMOM55.3f | Homozygote | 27.01052 | 27.12545 | 27.00247 | | | | | |
| Hprt | DMOM55.1d | Homozygote | 27.01117 | 26.91164 | 26.86996 | | | | | |
| Hprt | DMOM55.1h | Homozygote | 26.79156 | 26.39383 | 26.41894 | | | | | |
| Hprt | DMOM55.1g | Homozygote | 27.34133 | 26.90015 | 27.06306 | | | | | |
| Hprt | DMOM55.1g | Homozygote | 27.25964 | 27.35621 | 27.11935 | | | | | |
| Hprt | DMOM60.3i | Homozygote | 28.46703 | 28.32319 | 28.47138 | | | | | |
| Hprt | DMOM60.3d | Homozygote | 27.14293 | 27.29452 | 27.18028 | | | | | |
| Hprt | DMOM64.4f | Homozygote | 30.63079 | 30.52602 | 31.4458 | | | | | |
| Hprt | DMOM64.4d | Homozygote | 28.67832 | 28.51526 | 28.49936 | | | | | |
| Hprt | DMOM64.3h | Homozygote | 27.58028 | 27.53399 | 28.38148 | | | | | |
| Hprt | DMOM64.3a | Homozygote | 25.75523 | 26.17001 | 25.707 | | | | | |
| Hprt | DMOM60.3i | Homozygote | 30.59656 | 29.42235 | 30.63222 | | | | | |
| Hprt | DMOM60.3d | Homozygote | 29.12622 | 28.75751 | 28.34243 | | | | | |
| Hprt | DMOM64.4f | Homozygote | 30.65695 | 32.26803 | 31.16251 | | | | | |
| Hprt | DMOM64.4d | Homozygote | 28.94352 | 30.10225 | 29.7222 | | | | | |
| Hprt | DMOM64.3h | Homozygote | 27.83923 | 27.87006 | 28.23049 | | | | | |
| Hprt | DMOM64.3a | Homozygote | 25.90026 | 26.2099 | 25.94469 | | | | | |
| Jag1 | DMOM54.1c | Homozygote | 24.53808 | 24.6899 | 24.55941 | 1.064623 | 0.958278 | 1.048994 | 1.023965 | 0.057421 |
| Jag1 | DMOM52.3g | Homozygote | 27.6443 | 27.99749 | 27.86283 | 0.879217 | 0.688296 | 0.755632 | 0.774382 | 0.096832 |
| Jag1 | DMOM50.4e | Homozygote | 25.72016 | 25.7161 | 25.90248 | 1.04626 | 1.049208 | 0.922051 | 1.00584 | 0.072578 |
| Jag1 | DMOM50.4a | Homozygote | 26.36133 | 26.22039 | 26.1098 | 0.785792 | 0.866432 | 0.935457 | 0.86256 | 0.074907 |
| Jag1 | DMOM46.3d | Homozygote | 27.07851 | 27.09757 | 27.18948 | 1.045981 | 1.032253 | 0.968542 | 1.015592 | 0.041321 |
| Jag1 | DMOM46.3g | Homozygote | 27.96471 | 27.83206 | 28.23003 | 0.861745 | 0.944736 | 0.716982 | 0.841154 | 0.115265 |
| Jag1 | DMOM55.3f | Homozygote | 27.38831 | 27.5236 | 27.20537 | 2.1648 | 1.971026 | 2.457472 | 2.197766 | 0.244893 |
| Jag1 | DMOM55.1d | Homozygote | 26.71377 | 26.67907 | 26.56217 | 1.035393 | 1.060599 | 1.150112 | 1.082035 | 0.060289 |
| Jag1 | DMOM55.1h | Homozygote | 25.77387 | 26.34818 | 26.42204 | 1.509347 | 1.01369 | 0.9631 | 1.162045 | 0.301834 |
| Jag1 | DMOM55.1g | Homozygote | 26.97194 | 26.73707 | 26.69019 | 0.974406 | 1.146683 | 1.184552 | 1.10188 | 0.112008 |
| Jag1 | DMOM60.3i | Homozygote | 26.17444 | 26.49128 | 26.78478 | 1.065374 | 0.855311 | 0.697866 | 0.872851 | 0.184381 |
| Jag1 | DMOM60.3d | Homozygote | 25.29635 | 24.86771 | 25.51995 | 0.84372 | 1.135614 | 0.722579 | 0.900637 | 0.212319 |
| Jag1 | DMOM64.4f | Homozygote | 29.36256 | 29.3103 | 29.50662 | 1.124893 | 1.166383 | 1.017993 | 1.10309 | 0.07656 |
| Jag1 | DMOM64.4d | Homozygote | 27.34694 | 27.24348 | 27.19622 | 0.921582 | 0.990095 | 1.023068 | 0.978248 | 0.05177 |
| Jag1 | DMOM64.3h | Homozygote | 26.23715 | 26.18081 | 26.86293 | 1.044548 | 1.086151 | 0.67694 | 0.93588 | 0.225211 |
| Jag1 | DMOM64.3a | Homozygote | 24.6603 | 25.31317 | 25.23386 | 1.149557 | 0.731135 | 0.772453 | 0.884382 | 0.230576 |
| Ccer2 | DMOM52.3g | Homozygote | 32.14469 | 32.13035 | 31.63658 | 0.36782 | 0.371495 | 0.523108 | 0.420808 | 0.088613 |
| Ccer2 | DMOM54.1c | Homozygote | 29.2108 | 29.44863 | 29.22003 | 1.428358 | 1.21128 | 1.419251 | 1.352963 | 0.122786 |
| Ccer2 | DMOM50.4e | Homozygote | 29.85634 | 29.82508 | 29.57093 | 1.718644 | 1.756291 | 2.094614 | 1.856516 | 0.207056 |
| Ccer2 | DMOM50.4a | Homozygote | 30.48186 | 30.22483 | 30.39018 | 1.668411 | 1.993783 | 1.777882 | 1.813359 | 0.165561 |
| Ccer2 | DMOM46.3d | Homozygote | 33.87792 | 34.16165 | 34.06898 | 2.939688 | 2.414839 | 2.575057 | 2.643194 | 0.268977 |
| Ccer2 | DMOM46.3g | Homozygote | 34.96225 | 34.51618 | 35.16401 | 0.80759 | 1.100194 | 0.70219 | 0.869991 | 0.206209 |
| Ccer2 | DMOM55.1g | Homozygote | 31.08387 | 31.01881 | 31.27073 | 1.343096 | 1.405045 | 1.179928 | 1.309356 | 0.11629 |
| Grp | DMOM54.1c | Homozygote | 29.7266 | 29.58597 | 29.48123 | 1.471011 | 1.621614 | 1.743727 | 1.612117 | 0.136606 |
| Grp | DMOM52.3g | Homozygote | 31.57609 | 31.22756 | 31.34059 | 1.155426 | 1.471163 | 1.3603 | 1.328963 | 0.160185 |
| Grp | DMOM50.4e | Homozygote | 30.59819 | 30.61676 | 30.56465 | 1.926522 | 1.90189 | 1.971836 | 1.933416 | 0.035479 |
| Grp | DMOM50.4a | Homozygote | 30.8056 | 30.65013 | 30.81586 | 1.266625 | 1.410749 | 1.257652 | 1.311675 | 0.085918 |
| Grp | DMOM46.3d | Homozygote | 30.99672 | 31.20863 | 31.06208 | 1.53722 | 1.327226 | 1.469136 | 1.444527 | 0.107138 |
| Grp | DMOM46.3g | Homozygote | 31.29658 | 31.13978 | 31.18214 | 1.475626 | 1.645046 | 1.597442 | 1.572705 | 0.087377 |
| Grp | DMOM55.1g | Homozygote | 31.60149 | 31.92884 | 31.91272 | 1.861875 | 1.48391 | 1.50059 | 1.615459 | 0.213566 |
| Ocm | DMOM54.1c | Homozygote | 27.68865 | 27.95739 | 27.98163 | 0.41214 | 0.342093 | 0.336395 | 0.363542 | 0.042183 |
| Ocm | DMOM52.3g | Homozygote | 23.93027 | 24.03106 | 24.2087 | 1.123955 | 1.048107 | 0.926682 | 1.032915 | 0.099511 |
| Ocm | DMOM50.4e | Homozygote | 26.02999 | 25.93146 | 26.41374 | 0.673045 | 0.720615 | 0.515849 | 0.636503 | 0.107162 |
| Ocm | DMOM50.4a | Homozygote | 26.46748 | 26.33694 | 26.44102 | 0.610907 | 0.668759 | 0.622214 | 0.63396 | 0.030663 |
| Ocm | DMOM46.3d | Homozygote | 26.76923 | 27.07378 | 26.99605 | 0.731108 | 0.591973 | 0.624742 | 0.649274 | 0.07274 |
| Ocm | DMOM46.3g | Homozygote | 27.85654 | 27.77484 | 27.82678 | 0.612111 | 0.647774 | 0.624866 | 0.62825 | 0.018071 |
| Ppm1l | DMOM54.1c | Homozygote | 26.20853 | 26.41556 | 26.46008 | 1.507765 | 1.306205 | 1.266515 | 1.360161 | 0.129359 |
| Ppm1l | DMOM52.3g | Homozygote | 27.46737 | 27.27969 | 27.44372 | 1.003767 | 1.143222 | 1.020355 | 1.055781 | 0.076179 |
| Ppm1l | DMOM50.4e | Homozygote | 28.0248 | 27.94772 | 27.93333 | 1.122896 | 1.184522 | 1.196398 | 1.167939 | 0.039458 |
| Ppm1l | DMOM50.4a | Homozygote | 28.13623 | 28.16249 | 28.20358 | 0.704105 | 0.691404 | 0.671989 | 0.689166 | 0.016174 |
| Ppm1l | DMOM46.3d | Homozygote | 29.36323 | 29.5927 | 29.70984 | 1.238879 | 1.0567 | 0.974289 | 1.089956 | 0.135394 |
| Ppm1l | DMOM46.3g | Homozygote | 30.10044 | 30.27438 | 30.18371 | 1.546535 | 1.370875 | 1.459795 | 1.459068 | 0.087832 |
| Ppm1l | DMOM55.1g | Homozygote | 29.53969 | 29.41166 | 29.55939 | 0.91332 | 0.998083 | 0.900937 | 0.937447 | 0.052876 |
| Slc26a5 | DMOM54.1c | Homozygote | 30.64809 | 30.3653 | 30.87056 | 0.490967 | 0.597285 | 0.420806 | 0.503019 | 0.088855 |
| Slc26a5 | DMOM52.3g | Homozygote | 28.99544 | 28.78355 | 29.22617 | 1.040113 | 1.20467 | 0.886391 | 1.043725 | 0.15917 |
| Slc26a5 | DMOM50.4e | Homozygote | 30.44543 | 30.54912 | 30.0969 | 0.521613 | 0.485437 | 0.66415 | 0.557067 | 0.094485 |
| Slc26a5 | DMOM50.4a | Homozygote | 30.92719 | 30.48647 | 30.78268 | 0.383053 | 0.519912 | 0.423409 | 0.442125 | 0.070323 |
| Slc26a5 | DMOM46.3d | Homozygote | 31.56701 | 31.48828 | 31.33186 | 0.502548 | 0.530737 | 0.591516 | 0.5416 | 0.045468 |
| Slc26a5 | DMOM46.3g | Homozygote | 32.82401 | 32.43401 | 32.37144 | 0.804022 | 1.05359 | 1.100286 | 0.985966 | 0.159288 |
| Foxo3 | DMOM60.3i | Homozygote | 29.66299 | 28.85815 | 28.78771 | 0.947861 | 1.655875 | 1.738719 | 1.447485 | 0.434665 |
| Foxo3 | DMOM60.3d | Homozygote | 28.75573 | 27.43433 | 28.53168 | 0.639503 | 1.598168 | 0.746942 | 0.994871 | 0.525225 |
| Foxo3 | DMOM64.4f | Homozygote | 32.01801 | 31.38028 | 30.05761 | 0.522823 | 0.813446 | 2.034672 | 1.123647 | 0.802241 |
| Foxo3 | DMOM64.4d | Homozygote | 28.74231 | 28.33838 | 29.81275 | 1.481354 | 1.959989 | 0.705385 | 1.382243 | 0.633147 |
| Foxo3 | DMOM64.3h | Homozygote | 29.01626 | 29.01092 | 28.94977 | 1.061637 | 1.065568 | 1.11171 | 1.079638 | 0.027845 |
| Foxo3 | DMOM64.3a | Homozygote | 27.0416 | 27.0109 | 27.05185 | 0.93553 | 0.955655 | 0.928906 | 0.94003 | 0.013931 |
| Trp53 | DMOM60.3i | Homozygote | 30.2736 | 30.59733 | 31.23566 | 0.745079 | 0.595316 | 0.382465 | 0.574287 | 0.182219 |
| Trp53 | DMOM60.3d | Homozygote | 29.17548 | 29.43105 | 30.14596 | 0.573789 | 0.480637 | 0.292825 | 0.449084 | 0.143115 |
| Trp53 | DMOM64.4f | Homozygote | 31.52747 | 31.6122 | 31.14311 | 2.502113 | 2.359393 | 3.26596 | 2.709155 | 0.487459 |
| Trp53 | DMOM64.4d | Homozygote | 29.40641 | 29.3638 | 29.98206 | 3.184435 | 3.279886 | 2.136704 | 2.867008 | 0.634261 |
| Trp53 | DMOM64.3h | Homozygote | 29.5294 | 29.92374 | 29.5276 | 1.358929 | 1.033923 | 1.36063 | 1.251161 | 0.188135 |
| Trp53 | DMOM64.3a | Homozygote | 27.88841 | 27.52515 | 27.44885 | 0.905553 | 1.164838 | 1.228101 | 1.099497 | 0.170913 |

| | | | | | | | | | | |
|---|---|---|---|---|---|---|---|---|---|---|
| Rest | DMKO84.3b | Wildtype | 28.07881 | 28.05892 | 28.04812 | 0.988379 | 1.002103 | 1.009634 | 1.000039 | 0.010776 |
| Rest | DMKO84.3h | Wildtype | 28.60687 | 28.7197 | 28.36367 | 0.970327 | 0.897331 | 1.148496 | 1.005384 | 0.1292 |
| Rest | DMKO79.5h | Wildtype | 31.3661 | 31.46673 | 31.3309 | 1.015232 | 0.946834 | 1.040305 | 1.00079 | 0.04838 |
| Rest | DMKO79.7b | Wildtype | 30.13778 | 29.32094 | 28.93379 | 0.626935 | 1.104374 | 1.444311 | 1.05854 | 0.410611 |
| Rest | DMKO84.4g | Wildtype | 32.00204 | 32.15786 | 31.73586 | 0.974825 | 0.875024 | 1.172339 | 1.007396 | 0.15131 |
| Sp1 | DMKO84.3b | Wildtype | 28.18131 | 28.08126 | 28.16305 | 0.973037 | 1.042909 | 0.985427 | 1.000457 | 0.037282 |
| Sp1 | DMKO84.3h | Wildtype | 27.96627 | 28.20633 | 28.27288 | 1.134634 | 0.960704 | 0.917391 | 1.004243 | 0.11498 |
| Sp1 | DMKO79.5h | Wildtype | 29.3871 | 29.96055 | 29.65264 | 1.213912 | 0.81576 | 1.009835 | 1.013169 | 0.199097 |
| Sp1 | DMKO79.7b | Wildtype | 29.43227 | 28.85835 | 28.47281 | 0.701675 | 1.044486 | 1.364461 | 1.036874 | 0.331458 |
| Sp1 | DMKO84.4g | Wildtype | 31.14898 | 30.42768 | 30.51043 | 0.730377 | 1.204154 | 1.137028 | 1.023853 | 0.256364 |
| Trp53 | DMKO84.3b | Wildtype | 30.02116 | 29.8334 | 30.21291 | 1.00092 | 1.140044 | 0.876352 | 1.005772 | 0.131913 |
| Trp53 | DMKO84.3h | Wildtype | 30.21728 | 30.70245 | 30.26414 | 1.130798 | 0.807858 | 1.094662 | 1.011106 | 0.176943 |
| Trp53 | DMKO79.5h | Wildtype | 33.4585 | 33.51764 | 33.33879 | 0.986105 | 0.946494 | 1.071417 | 1.001339 | 0.06384 |
| Trp53 | DMKO79.7b | Wildtype | 31.7069 | 31.64723 | 31.5289 | 0.946569 | 0.986538 | 1.070863 | 1.001323 | 0.063452 |
| Trp53 | DMKO84.4g | Wildtype | 34.34626 | 34.45222 | 34.07698 | 0.962965 | 0.894777 | 1.160578 | 1.006107 | 0.138052 |
| Cdkn1a | DMKO84.3b | Wildtype | 28.83523 | 29.97632 | 29.5782 | 1.545441 | 0.700731 | 0.923414 | 1.056528 | 0.437805 |
| Cdkn1a | DMKO84.3h | Wildtype | 28.75955 | 28.21844 | 28.3014 | 0.793837 | 1.155104 | 1.090555 | 1.013165 | 0.192666 |
| Cdkn1a | DMKO79.5h | Wildtype | 27.18834 | 27.17468 | 27.3802 | 1.042035 | 1.051944 | 0.912273 | 1.002084 | 0.077936 |
| Cdkn1a | DMKO79.7b | Wildtype | 26.67596 | 27.35597 | 27.74963 | 1.499581 | 0.93598 | 0.712465 | 1.049342 | 0.405618 |
| Cdkn1a | DMKO84.4g | Wildtype | 27.28678 | 27.26742 | 27.46128 | 1.036499 | 1.050496 | 0.91841 | 1.001802 | 0.072557 |
| Tgfb1 | DMKO84.3b | Wildtype | 31.67937 | 31.74914 | 31.84888 | 1.056842 | 1.00695 | 0.939684 | 1.001159 | 0.058793 |
| Tgfb1 | DMKO84.3h | Wildtype | 30.44337 | 31.29773 | 31.52515 | 1.564152 | 0.865151 | 0.738974 | 1.056092 | 0.444493 |
| Tgfb1 | DMKO79.5h | Wildtype | 30.03353 | 30.2237 | 30.5464 | 1.176372 | 1.031094 | 0.824436 | 1.010634 | 0.176858 |
| Tgfb1 | DMKO79.7b | Wildtype | 30.1859 | 30.28513 | 30.57359 | 1.119075 | 1.044691 | 0.855368 | 1.006378 | 0.135964 |
| Tgfb1 | DMKO84.4g | Wildtype | 30.35371 | 30.52641 | 30.71869 | 1.132279 | 1.004534 | 0.879189 | 1.005334 | 0.126547 |
| Hprt | DMKO38.2a | Homozygote | 28.5967 | 28.62321 | 28.50305 | | | | | |
| Hprt | DMKO39.2c | Homozygote | 27.67002 | 27.91795 | 27.55215 | | | | | |
| Hprt | DMKO41.2a | Homozygote | 28.6818 | 28.67433 | 28.77185 | | | | | |
| Hprt | DMKO58.2a | Homozygote | 26.76881 | 26.84965 | 26.95884 | | | | | |
| Hprt | DMKO58.2b | Homozygote | 27.33711 | 27.36186 | 27.363 | | | | | |
| Hprt | DMKO58.2d | Homozygote | 26.86488 | 26.37959 | 26.4056 | | | | | |
| Hprt | DMKO67.2d | Homozygote | 28.49966 | 28.66849 | 28.64527 | | | | | |
| Hprt | DMKO67.2g | Homozygote | 28.29271 | 28.28047 | 28.01288 | | | | | |
| Hprt | DMKO67.2d | Homozygote | 28.89261 | 28.63633 | 28.51769 | | | | | |
| Hprt | DMKO67.2d | Homozygote | 29.035 | 28.88365 | 28.7126 | | | | | |
| Hprt | DMKO68.3b | Homozygote | 27.5315 | 27.61124 | 27.2083 | | | | | |
| Hprt | DMKO68.3b | Homozygote | 27.5315 | 27.61124 | 27.2083 | | | | | |
| Hprt | DMKO57.5a | Homozygote | 26.77817 | 26.4602 | 26.82506 | | | | | |
| Hprt | DMKO57.5b | Homozygote | 26.63454 | 27.06166 | 27.01111 | | | | | |
| Hprt | DMKO64.3j | Homozygote | 27.05621 | 26.97465 | 27.25704 | | | | | |
| Hprt | DMKO64.3h | Homozygote | 27.56824 | 27.53732 | 27.15408 | | | | | |
| Hprt | DMKO57.5a | Homozygote | 27.87911 | 27.41002 | 27.47129 | | | | | |
| Hprt | DMKO64.3j | Homozygote | 27.68712 | 27.60658 | 27.71867 | | | | | |
| Hprt | DMKO57.5a | Homozygote | 27.29452 | 27.16443 | 26.89761 | | | | | |
| Hprt | DMKO64.3j | Homozygote | 27.80187 | 27.98158 | 27.66775 | | | | | |
| Hprt | DMKO67.2d | Homozygote | 28.19913 | 28.26423 | 28.01288 | | | | | |
| Hprt | DMKO64.3h | Homozygote | 29.45914 | 29.77705 | 28.61122 | | | | | |
| Hprt | DMKO64.3h | Homozygote | 27.68561 | 27.89285 | 27.58192 | | | | | |
| Hprt | DMKO64.3h | Homozygote | 27.85409 | 27.97827 | 27.93016 | | | | | |
| Hprt | DMKO57.5a | Homozygote | 27.22128 | 27.13648 | 27.26281 | | | | | |
| Hprt | DMKO72.1g | Homozygote | 26.99057 | 26.73003 | 26.67386 | | | | | |
| Hprt | DMKO72.1h | Homozygote | 27.04569 | 26.87855 | 26.9864 | | | | | |
| Hprt | DMKO64.5e | Homozygote | 26.62678 | 26.27061 | 26.2502 | | | | | |
| Hprt | DMKO72.1g | Homozygote | 27.48529 | 27.5659 | 27.66234 | | | | | |
| Hprt | DMKO72.1h | Homozygote | 27.66336 | 27.59297 | 27.49896 | | | | | |
| Hprt | DMKO84.3d | Homozygote | 27.60034 | 27.83304 | 27.70722 | | | | | |
| Hprt | DMKO84.3c | Homozygote | 27.64088 | 27.84097 | 27.31917 | | | | | |
| Hprt | DMKO84.3d | Homozygote | 28.50393 | 29.00712 | 28.8278 | | | | | |
| Hprt | DMKO84.3c | Homozygote | 28.1628 | 28.39522 | 28.48962 | | | | | |
| Hprt | DMKO79.5f | Homozygote | 28.08275 | 28.00044 | 28.00611 | | | | | |
| Hprt | DMKO79.7h | Homozygote | 28.69641 | 28.98546 | 28.85411 | | | | | |
| Hprt | DMKO79.5f | Homozygote | 28.44677 | 29.65682 | 28.80385 | | | | | |
| Hprt | DMKO79.7h | Homozygote | 33.10671 | 32.57412 | 31.67217 | | | | | |
| Hprt | DMKO84.4e | Homozygote | 28.42901 | 28.36796 | 28.6414 | | | | | |
| Hprt | DMKO84.4e | Homozygote | 31.71713 | 32.72098 | 30.53402 | | | | | |
| Hprt | DMKO84.3d | Homozygote | 29.76553 | 29.70862 | 28.51871 | | | | | |
| Hprt | DMKO84.3c | Homozygote | 29.16386 | 28.73922 | 29.10246 | | | | | |
| Hprt | DMKO79.5f | Homozygote | 27.57496 | 28.23117 | 28.33534 | | | | | |
| Hprt | DMKO79.7h | Homozygote | 28.86989 | 30.01496 | 29.86333 | | | | | |
| Hprt | DMKO84.4e | Homozygote | 28.60417 | 29.30976 | 29.48699 | | | | | |
| Hprt | DMKO57.5a | Homozygote | 27.34122 | 27.85121 | 27.79706 | | | | | |
| Hprt | DMKO84.3d | Homozygote | 28.97995 | 29.29376 | 29.41326 | | | | | |
| Hprt | DMKO84.3c | Homozygote | 29.05723 | 29.18172 | 29.00042 | | | | | |
| Jag1 | DMKO38.2a | Homozygote | 29.1621 | 29.06372 | 29.14983 | 2.09549 | 2.243362 | 2.113382 | 2.150744 | 0.080707 |
| Jag1 | DMKO39.2c | Homozygote | 29.0156 | 28.50421 | 28.72615 | 0.344741 | 0.491401 | 0.421334 | 0.419158 | 0.073354 |
| Jag1 | DMKO41.2a | Homozygote | 28.64993 | 28.18159 | 28.08297 | 0.807017 | 1.116515 | 1.19551 | 1.039681 | 0.205327 |
| Jag1 | DMKO58.2a | Homozygote | 26.74241 | 26.70579 | 26.28033 | 0.665668 | 0.682781 | 0.916978 | 0.755142 | 0.140415 |
| Jag1 | DMKO58.2b | Homozygote | 28.17285 | 28.02185 | 28.14635 | 0.710592 | 0.788995 | 0.723764 | 0.741117 | 0.041983 |
| Jag1 | DMKO58.2d | Homozygote | 26.412 | 26.18329 | 26.08293 | 1.302337 | 1.526064 | 1.635997 | 1.488133 | 0.170034 |
| Jag1 | DMKO67.2d | Homozygote | 28.25752 | 28.33603 | 28.28794 | 1.079383 | 1.022216 | 1.05686 | 1.05282 | 0.028797 |
| Jag1 | DMKO67.2g | Homozygote | 27.16249 | 27.2864 | 27.03867 | 1.736421 | 1.593512 | 1.892037 | 1.740657 | 0.149308 |
| Jag1 | DMKO68.3b | Homozygote | 27.44201 | 27.4485 | 27.29087 | 0.62484 | 0.622035 | 0.693852 | 0.646909 | 0.040678 |
| Jag1 | DMKO68.3b | Homozygote | 27.44201 | 27.4485 | 27.29087 | 0.70006 | 0.696918 | 0.77738 | 0.724786 | 0.045575 |
| Jag1 | DMKO57.5a | Homozygote | 26.72517 | 26.18501 | 26.22408 | 0.733942 | 1.067243 | 1.038732 | 0.946639 | 0.184752 |
| Jag1 | DMKO57.5b | Homozygote | 27.05272 | 26.86112 | 27.24078 | 0.67868 | 0.775075 | 0.595738 | 0.683164 | 0.089752 |
| Jag1 | DMKO64.3j | Homozygote | 26.55331 | 26.96431 | 27.15465 | 1.252784 | 0.942218 | 0.825761 | 1.006921 | 0.220742 |
| Jag1 | DMKO64.3h | Homozygote | 26.52588 | 26.44902 | 26.56709 | 1.148589 | 1.211443 | 1.116249 | 1.15876 | 0.048405 |
| Jag1 | DMKO72.1g | Homozygote | 26.12885 | 26.27205 | 26.35791 | 1.040469 | 0.942157 | 0.887722 | 0.956783 | 0.077417 |
| Jag1 | DMKO72.1h | Homozygote | 26.16481 | 26.08627 | 26.11215 | 1.118888 | 1.18149 | 1.160484 | 1.153621 | 0.031861 |
| Jag1 | DMKO64.5e | Homozygote | 26.04489 | 26.1445 | 26.09021 | 0.6571 | 0.613262 | 0.636777 | 0.635713 | 0.021938 |
| Jag1 | DMKO84.3d | Homozygote | 26.16515 | 26.20662 | 26.15425 | 0.901686 | 0.876142 | 0.908528 | 0.895452 | 0.01707 |
| Jag1 | DMKO84.3c | Homozygote | 26.60693 | 26.58307 | 26.60088 | 1.101112 | 1.119476 | 1.105743 | 1.108777 | 0.009551 |
| Jag1 | DMKO79.5f | Homozygote | 26.89748 | 26.78445 | 26.91372 | 0.915058 | 0.989636 | 0.904818 | 0.936504 | 0.046298 |
| Jag1 | DMKO79.7h | Homozygote | 26.801 | 26.45949 | 26.02336 | 0.838295 | 1.062188 | 1.437112 | 1.112532 | 0.302566 |
| Jag1 | DMKO84.4e | Homozygote | 26.49059 | 26.50108 | 26.21659 | 0.797139 | 0.791366 | 0.963868 | 0.850791 | 0.09797 |
| Ccer2 | DMKO41.2a | Homozygote | 28.13833 | 28.02358 | 27.77591 | 6.153062 | 6.662446 | 7.910237 | 6.908582 | 0.904076 |
| Ccer2 | DMKO67.2d | Homozygote | 29.70284 | 29.35747 | 29.86484 | 5.064494 | 6.43429 | 4.526548 | 5.341777 | 0.983633 |
| Ccer2 | DMKO57.5a | Homozygote | 29.22516 | 29.06203 | 29.33709 | 11.2116 | 12.55381 | 10.37465 | 11.38002 | 1.099297 |
| Ccer2 | DMKO64.3j | Homozygote | 28.37843 | 28.49098 | 28.98714 | 5.796863 | 5.361791 | 3.801482 | 4.986712 | 1.049238 |
| Ccer2 | DMKO64.3h | Homozygote | 31.6241 | 32.34176 | 32.15506 | 1.301233 | 0.791257 | 0.900577 | 0.997689 | 0.268499 |
| Ccer2 | DMKO72.1g | Homozygote | 28.23603 | 28.44379 | 27.83525 | 3.274056 | 2.834941 | 4.322463 | 3.477153 | 0.764275 |
| Grp | DMKO67.2d | Homozygote | 30.53704 | 30.58989 | 30.51146 | 4.340372 | 4.184235 | 4.418009 | 4.314205 | 0.119064 |
| Grp | DMKO57.5a | Homozygote | 30.3239 | 30.28176 | 30.28644 | 2.995398 | 3.084193 | 3.074201 | 3.051264 | 0.048639 |
| Grp | DMKO64.3j | Homozygote | 29.32699 | 29.45423 | 29.21853 | 4.563879 | 4.178624 | 4.920223 | 4.554242 | 0.370894 |
| Grp | DMKO64.3h | Homozygote | 32.23045 | 31.52714 | 30.62555 | 2.600646 | 4.23447 | 7.91053 | 4.915215 | 2.71961 |
| Grp | DMKO72.1g | Homozygote | 30.04637 | 30.61047 | 30.68001 | 3.676927 | 2.487002 | 2.369959 | 2.844629 | 0.723163 |
| Grp | DMKO72.1h | Homozygote | 30.52101 | 30.26869 | 30.49449 | 1.855919 | 2.210617 | 1.890342 | 1.985626 | 0.195607 |
| Grk1 | DMKO41.2a | Homozygote | 32.0977 | 31.73798 | 31.77911 | 3.143644 | 4.033842 | 3.920478 | 3.699321 | 0.484557 |
| Grk1 | DMKO67.2d | Homozygote | 32.406 | 32.50169 | 32.61583 | 2.11628 | 1.980469 | 1.829814 | 1.975521 | 0.143297 |
| Grk1 | DMKO57.5a | Homozygote | 30.73253 | 30.56526 | 31.07734 | 12.98645 | 14.58286 | 10.22566 | 12.59832 | 2.204379 |
| Grk1 | DMKO64.3j | Homozygote | 31.49953 | 31.43065 | 31.35438 | 5.114103 | 5.364224 | 5.655407 | 5.377911 | 0.270911 |
| Grk1 | DMKO64.3h | Homozygote | 35.18208 | 33.89863 | 32.63234 | 0.455285 | 1.108255 | 2.665823 | 1.409788 | 1.135699 |
| Grk1 | DMKO72.1h | Homozygote | 30.00071 | 30.00274 | 30.32575 | 3.624595 | 3.619505 | 2.893435 | 3.379178 | 0.420674 |
| Tmc1 | DMKO41.2a | Homozygote | 34.82138 | 35.00135 | 34.55904 | 0.255923 | 0.225909 | 0.306959 | 0.262931 | 0.040977 |
| Tmc1 | DMKO67.2d | Homozygote | 34.27915 | 34.07614 | 33.8796 | 0.236685 | 0.272448 | 0.312211 | 0.273781 | 0.03778 |
| Tmc1 | DMKO57.5a | Homozygote | 33.054 | 32.79128 | 32.55704 | 0.600539 | 0.720488 | 0.847501 | 0.722843 | 0.123498 |
| Tmc1 | DMKO64.3j | Homozygote | 34.21688 | 34.36972 | 34.32769 | 0.307867 | 0.27692 | 0.285106 | 0.289964 | 0.016035 |
| Tmc1 | DMKO64.3h | Homozygote | 34.43198 | 35.0947 | 36.16327 | 0.395956 | 0.25012 | 0.119255 | 0.25511 | 0.138418 |
| Tmc1 | DMKO72.1g | Homozygote | 33.23046 | 33.02726 | 33.05914 | 0.345247 | 0.397464 | 0.388778 | 0.377163 | 0.027979 |
| Hspa2 | DMKO41.2a | Homozygote | 29.52384 | 29.48448 | 29.26512 | 2.484691 | 2.55342 | 2.972723 | 2.670278 | 0.264169 |
| Hspa2 | DMKO67.2d | Homozygote | 29.92075 | 30.12141 | 30.08658 | 2.691947 | 2.342401 | 2.399639 | 2.477996 | 0.187485 |
| Hspa2 | DMKO57.5a | Homozygote | 29.11786 | 28.72654 | 28.44834 | 1.324325 | 1.736977 | 2.1064 | 1.722567 | 0.391237 |
| Hspa2 | DMKO64.3j | Homozygote | 29.25062 | 29.23718 | 29.17791 | 2.262368 | 2.283543 | 2.379305 | 2.308405 | 0.062307 |
| Hspa2 | DMKO64.3h | Homozygote | 32.48864 | 31.45115 | 29.64098 | 0.509728 | 1.046297 | 3.669204 | 1.741743 | 1.690653 |
| Hspa2 | DMKO72.1h | Homozygote | 29.08069 | 29.09583 | 28.51661 | 1.939461 | 1.919203 | 2.867377 | 2.242013 | 0.541675 |
| Myo3a | DMKO41.2a | Homozygote | 33.15449 | 32.65945 | 33.08007 | 5.07064 | 7.146366 | 5.339078 | 5.852028 | 1.128937 |
| Myo3a | DMKO67.2d | Homozygote | 32.81292 | 32.93235 | 33.18071 | 1.997845 | 1.839125 | 1.548265 | 1.795078 | 0.228003 |
| Myo3a | DMKO57.5a | Homozygote | 33.10336 | 33.0687 | 33.53467 | 4.599333 | 4.711174 | 3.410812 | 4.24044 | 0.720652 |
| Myo3a | DMKO64.3j | Homozygote | 33.4625 | 33.94534 | 34.41369 | 5.786677 | 4.140762 | 2.992895 | 4.306778 | 1.40427 |
| Myo3a | DMKO64.3h | Homozygote | 34.65246 | 34.18968 | 32.30133 | 0.872398 | 1.202333 | 4.45119 | 2.175307 | 1.977864 |
| Myo3a | DMKO72.1g | Homozygote | 31.54321 | 31.77394 | 31.55766 | 1.994408 | 1.699641 | 1.974538 | 1.889529 | 0.164748 |
| St8sia3 | DMKO41.2a | Homozygote | 31.12698 | 31.1788 | 31.44031 | 1.740362 | 1.678962 | 1.400614 | 1.606646 | 0.181051 |
| St8sia3 | DMKO67.2d | Homozygote | 31.26216 | 31.26108 | 31.33558 | 1.89697 | 1.898397 | 1.802853 | 1.866073 | 0.054755 |
| St8sia3 | DMKO57.5a | Homozygote | 29.9802 | 29.58629 | 29.31711 | 1.378566 | 1.811368 | 2.182923 | 1.790952 | 0.402567 |
| St8sia3 | DMKO64.3j | Homozygote | 30.32467 | 30.61379 | 30.06092 | 1.472072 | 1.204745 | 1.767369 | 1.481395 | 0.281428 |
| St8sia3 | DMKO64.3h | Homozygote | 30.53949 | 30.1167 | 30.1213 | 1.130866 | 1.515949 | 1.511124 | 1.38598 | 0.220948 |
| St8sia3 | DMKO72.1h | Homozygote | 29.98619 | 29.96186 | 30.00101 | 1.004319 | 1.0214 | 0.994051 | 1.00659 | 0.013815 |
| Tmem173 | DMKO67.2d | Homozygote | 30.52634 | 30.99981 | 30.7593 | 2.377001 | 1.711987 | 2.022564 | 2.037184 | 0.332748 |
| Tmem173 | DMKO57.5a | Homozygote | 30.17009 | 29.80637 | 30.19069 | 1.512536 | 1.946231 | 1.491086 | 1.649951 | 0.25681 |
| Tmem173 | DMKO64.3j | Homozygote | 29.75645 | 29.71761 | 29.47389 | 2.131744 | 2.189914 | 2.592947 | 2.304868 | 0.251173 |
| Tmem173 | DMKO64.3h | Homozygote | 32.91603 | 32.34565 | 30.45513 | 0.706892 | 1.04967 | 3.891856 | 1.882806 | 1.748309 |
| Tmem173 | DMKO72.1g | Homozygote | 29.32175 | 29.51404 | 29.43132 | 1.610399 | 1.409441 | 1.492618 | 1.504153 | 0.100974 |
| Tmem173 | DMKO72.1h | Homozygote | 29.36897 | 29.38576 | 29.36311 | 1.469381 | 1.452381 | 1.475359 | 1.465707 | 0.011922 |
| Slc52a3 | DMKO67.2d | Homozygote | 31.80484 | 32.05053 | 31.38165 | 5.273994 | 4.448157 | 7.071846 | 5.597999 | 1.341518 |
| Slc52a3 | DMKO57.5a | Homozygote | 31.78732 | 31.91683 | 31.26423 | 3.757986 | 3.435315 | 5.400323 | 4.197875 | 1.053775 |
| Slc52a3 | DMKO64.3j | Homozygote | 30.75134 | 30.59022 | 31.04325 | 4.757884 | 5.320038 | 3.886338 | 4.654753 | 0.722392 |
| Slc52a3 | DMKO64.3h | Homozygote | 34.38315 | 32.02865 | 30.98495 | 1.616561 | 8.267386 | 17.04333 | 8.97576 | 7.737743 |
| Slc52a3 | DMKO72.1g | Homozygote | 30.56454 | 30.77249 | 30.77304 | 3.177717 | 2.75116 | 2.750103 | 2.892993 | 0.246579 |
| Slc52a3 | DMKO72.1h | Homozygote | 30.20126 | 30.73473 | 31.06456 | 4.32095 | 2.985307 | 2.375197 | 3.227151 | 0.995166 |
| Kif21b | DMKO67.2d | Homozygote | 32.68186 | 33.79518 | 33.0743 | 1.167189 | 0.539508 | 0.889212 | 0.865303 | 0.314523 |
| Kif21b | DMKO57.5a | Homozygote | 32.71023 | 32.41737 | 31.71778 | 1.181105 | 1.446938 | 2.34988 | 1.659308 | 0.612645 |
| Kif21b | DMKO64.3j | Homozygote | 33.19985 | 32.69629 | 32.43222 | 1.01409 | 1.437684 | 1.726453 | 1.392743 | 0.358301 |
| Kif21b | DMKO64.3h | Homozygote | 36.04849 | 33.91187 | 33.73009 | 0.182487 | 0.802455 | 0.910212 | 0.631718 | 0.392758 |
| Kif21b | DMKO72.1g | Homozygote | 31.02969 | 31.19739 | 31.00913 | 0.657122 | 0.585009 | 0.666551 | 0.636228 | 0.044606 |
| Kif21b | DMKO72.1h | Homozygote | 31.44027 | 31.54774 | 31.52281 | 0.926115 | 0.85963 | 0.874614 | 0.886786 | 0.034874 |
| Mfsd6 | DMKO67.2d | Homozygote | 30.34956 | 30.41337 | 30.34961 | 1.645887 | 1.57467 | 1.645824 | 1.622127 | 0.041099 |
| Mfsd6 | DMKO57.5a | Homozygote | 28.48494 | 28.57768 | 28.5677 | 2.225697 | 2.087116 | 2.101612 | 2.138141 | 0.076171 |
| Mfsd6 | DMKO64.3j | Homozygote | 29.14039 | 29.33639 | 29.43249 | 1.703886 | 1.48744 | 1.391588 | 1.527638 | 0.159983 |
| Mfsd6 | DMKO64.3h | Homozygote | 29.27673 | 29.21306 | 29.01905 | 1.514836 | 1.58318 | 1.811069 | 1.636362 | 0.155112 |
| Mfsd6 | DMKO72.1g | Homozygote | 29.17806 | 29.23772 | 29.16313 | 1.391972 | 1.335581 | 1.406452 | 1.378002 | 0.037444 |
| Mfsd6 | DMKO72.1h | Homozygote | 29.29434 | 29.40208 | 29.18834 | 1.3519 | 1.254621 | 1.454977 | 1.353833 | 0.100192 |
| BC030867 | DMKO67.2d | Homozygote | 30.21085 | 30.85805 | 30.45319 | 4.692104 | 2.996003 | 3.966572 | 3.884893 | 0.850995 |
| BC030867 | DMKO57.5a | Homozygote | 29.05903 | 29.07934 | 29.14501 | 2.367622 | 2.334536 | 2.230648 | 2.310935 | 0.071472 |
| BC030867 | DMKO64.3j | Homozygote | 29.01396 | 29.0021 | 28.77272 | 3.034985 | 3.060032 | 3.587385 | 3.227468 | 0.31195 |
| BC030867 | DMKO64.3h | Homozygote | 30.08801 | 29.94236 | 30.20763 | 2.210897 | 2.445757 | 2.034976 | 2.230543 | 0.206094 |
| BC030867 | DMKO72.1g | Homozygote | 29.51679 | 29.76242 | 30.24673 | 2.105293 | 1.775701 | 1.269336 | 1.716777 | 0.421082 |
| BC030867 | DMKO72.1h | Homozygote | 29.61662 | 29.63597 | 29.97758 | 1.558078 | 1.537323 | 1.213196 | 1.436199 | 0.193405 |
| Eln | DMKO67.2d | Homozygote | 30.99369 | 30.9622 | 30.56613 | 4.071342 | 4.161161 | 5.475752 | 4.569419 | 0.786192 |
| Eln | DMKO57.5a | Homozygote | 30.66067 | 30.76599 | 31.06293 | 2.634185 | 2.448749 | 1.993222 | 2.358719 | 0.329829 |
| Eln | DMKO64.3j | Homozygote | 31.14351 | 30.99033 | 31.08678 | 1.493233 | 1.660504 | 1.553124 | 1.568954 | 0.084752 |
| Eln | DMKO64.3h | Homozygote | 31.07326 | 31.07025 | 30.58218 | 1.128841 | 1.1312 | 1.586581 | 1.282207 | 0.263598 |
| Eln | DMKO72.1g | Homozygote | 31.14528 | 31.19421 | 31.08611 | 1.074302 | 1.038475 | 1.119277 | 1.077351 | 0.040488 |
| Eln | DMKO72.1h | Homozygote | 30.85001 | 30.96274 | 30.95833 | 1.576697 | 1.458187 | 1.462654 | 1.499179 | 0.067169 |
| P2rx3 | DMKO67.2d | Homozygote | 31.64209 | 31.65483 | 31.38208 | 0.581583 | 0.576471 | 0.696441 | 0.618165 | 0.067837 |
| P2rx3 | DMKO57.5a | Homozygote | 31.9031 | 31.14395 | 31.24404 | 1.0533 | 1.782711 | 1.663225 | 1.499745 | 0.391221 |
| P2rx3 | DMKO64.3j | Homozygote | 31.45081 | 32.04012 | 32.89597 | 0.751177 | 0.499278 | 0.275871 | 0.508775 | 0.237795 |
| P2rx3 | DMKO64.3h | Homozygote | 31.67822 | 32.01229 | 31.69277 | 0.260953 | 0.207012 | 0.258334 | 0.2421 | 0.030415 |
| P2rx3 | DMKO72.1g | Homozygote | 29.90222 | 29.80261 | 30.18959 | 0.643404 | 0.689398 | 0.527204 | 0.620002 | 0.083591 |
| P2rx3 | DMKO72.1h | Homozygote | 29.98265 | 30.14192 | 30.7274 | 0.556499 | 0.498332 | 0.332103 | 0.462311 | 0.116454 |
| Ttc21a | DMKO67.2d | Homozygote | 32.4189 | 32.70428 | 32.54162 | 7.454817 | 6.116867 | 6.846901 | 6.806195 | 0.669904 |
| Ttc21a | DMKO57.5a | Homozygote | 32.06703 | 31.79651 | 31.56197 | 1.843537 | 2.223746 | 2.61632 | 2.227868 | 0.386408 |
| Ttc21a | DMKO64.3j | Homozygote | 32.20987 | 32.372 | 31.72159 | 2.620901 | 2.342301 | 3.676513 | 2.879905 | 0.703806 |
| Ttc21a | DMKO64.3h | Homozygote | 31.87538 | 31.89975 | 31.36409 | 2.887586 | 2.839217 | 4.115736 | 3.280846 | 0.72344 |
| Ttc21a | DMKO72.1g | Homozygote | 31.35449 | 31.82595 | 32.56865 | 2.224144 | 1.604132 | 0.958657 | 1.595644 | 0.632786 |
| Ttc21a | DMKO72.1h | Homozygote | 29.98265 | 30.14192 | 30.7274 | 0.556499 | 0.498332 | 0.332103 | 0.462311 | 0.116454 |
| Slc6a11 | DMKO57.5a | Homozygote | 31.17439 | 31.38398 | 30.92032 | 2.400593 | 2.075985 | 2.862865 | 2.446481 | 0.395442 |
| Slc6a11 | DMKO64.3j | Homozygote | 32.36569 | 32.32121 | 32.00967 | 2.767787 | 2.854455 | 3.542461 | 3.054901 | 0.424457 |
| Slc6a11 | DMKO67.2d | Homozygote | 32.83209 | 32.92491 | 32.18275 | 1.411959 | 1.323978 | 2.214581 | 1.650173 | 0.490767 |
| Slc6a11 | DMKO64.3h | Homozygote | 33.02567 | 32.39283 | 32.33378 | 1.176612 | 1.824471 | 1.900689 | 1.633924 | 0.397873 |
| Slc6a11 | DMKO72.1g | Homozygote | 31.98386 | 31.6175 | 31.98931 | 1.564958 | 2.017385 | 1.559062 | 1.713802 | 0.262927 |
| Slc6a11 | DMKO72.1h | Homozygote | 31.3919 | 31.51604 | 31.99771 | 2.539837 | 2.330419 | 1.668931 | 2.179729 | 0.454588 |
| Dtna | DMKO67.2d | Homozygote | 30.88736 | 30.67055 | 30.90368 | 2.175524 | 2.528309 | 2.151048 | 2.284961 | 0.211101 |
| Dtna | DMKO57.5a | Homozygote | 29.33154 | 29.37113 | 29.27595 | 2.118445 | 2.061095 | 2.201664 | 2.127068 | 0.07068 |
| Dtna | DMKO64.3j | Homozygote | 29.50255 | 29.89089 | 29.9237 | 1.608657 | 1.229023 | 1.201388 | 1.346356 | 0.227579 |
| Dtna | DMKO64.3h | Homozygote | 30.29731 | 30.19014 | 29.70204 | 1.228846 | 1.323605 | 1.856489 | 1.469647 | 0.338349 |
| Dtna | DMKO72.1g | Homozygote | 29.82428 | 29.78699 | 30.49996 | 1.215911 | 1.247751 | 0.761203 | 1.074955 | 0.272183 |
| Dtna | DMKO72.1h | Homozygote | 29.49608 | 29.69263 | 30.1594 | 1.366379 | 1.192348 | 0.862765 | 1.140497 | 0.25578 |
| Ocm | DMKO38.2a | Homozygote | 31.27682 | 31.04092 | 31.2004 | 0.031872 | 0.037534 | 0.033606 | 0.034338 | 0.002901 |
| Ocm | DMKO39.2c | Homozygote | 31.85348 | 32.08194 | 31.92475 | 0.13639 | 0.116416 | 0.129816 | 0.127541 | 0.01018 |
| Ocm | DMKO41.2a | Homozygote | 34.60426 | 35.32113 | 34.22675 | 0.031604 | 0.019229 | 0.041057 | 0.03063 | 0.010947 |
| Ocm | DMKO67.2d | Homozygote | 32.71619 | 32.92509 | 32.70498 | 0.069869 | 0.06045 | 0.070414 | 0.066911 | 0.005602 |
| Ocm | DMKO67.2g | Homozygote | 32.22837 | 32.1963 | 32.50831 | 0.044727 | 0.045733 | 0.036838 | 0.042433 | 0.004871 |
| Ocm | DMKO64.3j | Homozygote | 31.27431 | 31.61207 | 31.30807 | 0.080345 | 0.063575 | 0.078487 | 0.074136 | 0.009193 |
| Ocm | DMKO64.3h | Homozygote | 31.88849 | 31.99399 | 31.72382 | 0.080869 | 0.075166 | 0.090646 | 0.082227 | 0.007829 |
| Slc26a5 | DMKO38.2a | Homozygote | 35.00383 | 34.92635 | 34.28729 | 0.484028 | 0.510733 | 0.795373 | 0.596711 | 0.172563 |
| Slc26a5 | DMKO39.2c | Homozygote | 35.36695 | 35.12891 | 34.78858 | 0.419489 | 0.49474 | 0.626364 | 0.513531 | 0.10471 |
| Slc26a5 | DMKO41.2a | Homozygote | 36.0535 | 35.55165 | 35.80568 | 0.242511 | 0.343402 | 0.287961 | 0.291291 | 0.050528 |
| Slc26a5 | DMKO67.2d | Homozygote | 35.05893 | 35.37808 | 35.28166 | 0.224002 | 0.179548 | 0.191958 | 0.198503 | 0.022939 |
| Slc26a5 | DMKO67.2g | Homozygote | 34.44985 | 35.28061 | 35.16247 | 0.155972 | 0.087692 | 0.095176 | 0.112947 | 0.037448 |
| Slc26a5 | DMKO64.3j | Homozygote | 33.78928 | 33.91761 | 33.74797 | 0.351782 | 0.321842 | 0.361999 | 0.345208 | 0.02087 |
| Slc26a5 | DMKO64.3h | Homozygote | 33.53623 | 33.34553 | 34.72601 | 0.21629 | 0.246855 | 0.094815 | 0.185987 | 0.080423 |
| Nr3c1 | DMKO64.3h | Homozygote | 30.89635 | 30.77327 | 30.8532 | 1.294749 | 1.410047 | 1.334057 | 1.346284 | 0.058614 |
| Nr3c1 | DMKO57.5a | Homozygote | 29.86713 | 29.97875 | 30.13069 | 2.033695 | 1.882281 | 1.694128 | 1.870035 | 0.170114 |
| Nr3c1 | DMKO72.1g | Homozygote | 28.96077 | 28.43474 | 28.38756 | 0.689863 | 0.993379 | 1.026403 | 0.903215 | 0.185505 |
| Nr3c1 | DMKO72.1h | Homozygote | 28.43165 | 28.53987 | 28.60527 | 1.063411 | 0.986565 | 0.942837 | 0.997604 | 0.06104 |
| Nr3c1 | DMKO84.3d | Homozygote | 31.8779 | 32.08718 | 30.47461 | 0.886418 | 0.766721 | 2.344598 | 1.332579 | 0.878475 |
| Nr3c1 | DMKO84.3c | Homozygote | 31.75404 | 31.92698 | 31.20811 | 0.714304 | 0.633613 | 1.042856 | 0.796925 | 0.216771 |
| Fos | DMKO84.3d | Homozygote | 33.00533 | 33.54767 | 33.15594 | 0.57262 | 0.393194 | 0.515858 | 0.493891 | 0.091707 |
| Fos | DMKO84.3c | Homozygote | 32.55436 | 32.53872 | 32.90079 | 1.549226 | 1.566109 | 1.218506 | 1.444614 | 0.195997 |
| Fos | DMKO79.5f | Homozygote | 33.59575 | 34.03182 | 32.85482 | 1.614237 | 1.19316 | 2.697811 | 1.835069 | 0.776253 |
| Fos | DMKO79.7h | Homozygote | 36.46377 | 36.43049 | 36.50588 | 0.766609 | 0.784497 | 0.744556 | 0.765221 | 0.020007 |
| Fos | DMKO84.4e | Homozygote | 35.14068 | 36.57686 | 34.20432 | 1.923466 | 0.710806 | 3.680912 | 2.105061 | 1.493357 |
| Foxo1 | DMKO84.3d | Homozygote | 28.26147 | 28.64892 | 28.12583 | 1.031715 | 0.788726 | 1.133423 | 0.984621 | 0.177108 |
| Foxo1 | DMKO84.3c | Homozygote | 28.14467 | 28.11429 | 28.06557 | 0.659285 | 0.673315 | 0.696439 | 0.676346 | 0.018762 |
| Foxo1 | DMKO79.5f | Homozygote | 28.57379 | 28.59264 | 28.18886 | 0.869524 | 0.858232 | 1.135422 | 0.954392 | 0.156878 |
| Foxo1 | DMKO79.7h | Homozygote | 31.11174 | 30.85459 | 30.31693 | 1.392011 | 1.663614 | 2.414925 | 1.823517 | 0.529872 |
| Foxo1 | DMKO84.4e | Homozygote | 31.76889 | 31.23836 | 29.52106 | 0.324038 | 0.468062 | 1.539079 | 0.77706 | 0.663845 |
| Foxo3 | DMKO79.5f | Homozygote | 30.05858 | 29.53374 | 29.17561 | 0.699551 | 1.006492 | 1.290087 | 0.99871 | 0.295345 |
| Foxo3 | DMKO79.7h | Homozygote | 32.02045 | 31.32918 | 31.18443 | 1.011948 | 1.633993 | 1.806446 | 1.484129 | 0.417913 |
| Foxo3 | DMKO84.4e | Homozygote | 31.2117 | 30.9001 | 30.37001 | 1.236681 | 1.534821 | 2.216303 | 1.662602 | 0.502156 |
| Foxo3 | DMKO84.3d | Homozygote | 30.33347 | 30.13601 | 29.12005 | 0.745471 | 0.854816 | 1.72865 | 1.109646 | 0.538854 |
| Foxo3 | DMKO84.3c | Homozygote | 29.57129 | 29.66804 | 29.67671 | 1.955819 | 1.82896 | 1.818 | 1.867593 | 0.076602 |
| Ikzf2 | DMKO84.3d | Homozygote | 30.42928 | 30.33022 | 30.22438 | 0.848182 | 0.908466 | 0.97762 | 0.911423 | 0.06477 |
| Ikzf2 | DMKO84.3c | Homozygote | 30.38146 | 30.43117 | 30.43837 | 1.819079 | 1.757475 | 1.74873 | 1.775094 | 0.038342 |
| Ikzf2 | DMKO79.5f | Homozygote | 31.16257 | 31.63124 | 31.19708 | 1.242974 | 0.898213 | 1.213601 | 1.118263 | 0.191133 |

| | | | | | | | | | | |
|---|---|---|---|---|---|---|---|---|---|---|
| Ikzf2 | DMKO79.7h | Homozygote | 33.38468 | 33.20995 | 32.78021 | 0.864397 | 0.975688 | 1.314243 | 1.051443 | 0.234295 |
| Ikzf2 | DMKO84.4e | Homozygote | 34.18273 | 32.68481 | 33.28685 | 0.387939 | 1.095682 | 0.721858 | 0.73516 | 0.354059 |
| Rest | DMKO84.3d | Homozygote | 27.53031 | 27.53009 | 27.54638 | 1.142024 | 1.142201 | 1.129373 | 1.137866 | 0.007356 |
| Rest | DMKO84.3c | Homozygote | 27.81706 | 27.55046 | 27.48909 | 0.799318 | 0.961555 | 1.003339 | 0.921404 | 0.107774 |
| Rest | DMKO79.5f | Homozygote | 28.92216 | 28.53737 | 28.49206 | 1.943233 | 2.53722 | 2.618169 | 2.366207 | 0.368536 |
| Rest | DMKO79.7h | Homozygote | 31.96522 | 32.12332 | 32.23061 | 0.797185 | 0.714439 | 0.663237 | 0.724954 | 0.06759 |
| Rest | DMKO84.4e | Homozygote | 30.75828 | 30.32936 | 29.67353 | 1.645628 | 2.215391 | 3.490408 | 2.450476 | 0.944591 |
| Sp1 | DMKO84.3d | Homozygote | 27.37247 | 27.50617 | 27.397 | 1.346634 | 1.227444 | 1.323935 | 1.299337 | 0.063288 |
| Sp1 | DMKO84.3c | Homozygote | 27.44165 | 27.55816 | 27.45081 | 0.777726 | 0.717383 | 0.772804 | 0.755971 | 0.033508 |
| Sp1 | DMKO79.5f | Homozygote | 27.75841 | 28.06222 | 27.7771 | 1.320478 | 1.069734 | 1.30348 | 1.231231 | 0.140118 |
| Sp1 | DMKO79.7h | Homozygote | 30.74044 | 30.68666 | 30.65393 | 1.278757 | 1.32732 | 1.357782 | 1.321286 | 0.039856 |
| Sp1 | DMKO84.4e | Homozygote | 30.39415 | 29.64221 | 29.30181 | 0.878559 | 1.47955 | 1.873269 | 1.410459 | 0.500941 |
| Trp53 | DMKO84.3d | Homozygote | 29.40244 | 29.4141 | 29.58065 | 1.214198 | 1.204422 | 1.073108 | 1.163909 | 0.078788 |
| Trp53 | DMKO84.3c | Homozygote | 29.57012 | 29.56318 | 29.50608 | 0.843812 | 0.84788 | 0.882111 | 0.857934 | 0.021036 |
| Trp53 | DMKO79.5f | Homozygote | 31.00746 | 30.03831 | 30.28612 | 1.896786 | 3.713314 | 3.127255 | 2.912452 | 0.927119 |
| Trp53 | DMKO79.7h | Homozygote | 34.84985 | 34.72501 | 35.17656 | 0.48359 | 0.527301 | 0.385594 | 0.465495 | 0.072566 |
| Trp53 | DMKO84.4e | Homozygote | 32.59067 | 33.06865 | 32.78378 | 2.317901 | 1.664212 | 2.027514 | 2.003209 | 0.327521 |
| Cdkn1a | DMKO84.3d | Homozygote | 29.05178 | 28.5891 | 29.26026 | 1.062892 | 1.464768 | 0.919877 | 1.149179 | 0.282508 |
| Cdkn1a | DMKO84.3c | Homozygote | 28.71496 | 28.93978 | 29.11823 | 1.141678 | 0.976934 | 0.863268 | 0.99396 | 0.139984 |
| Cdkn1a | DMKO79.5f | Homozygote | 26.12722 | 26.60378 | 26.76244 | 1.662226 | 1.194626 | 1.070212 | 1.309021 | 0.312146 |
| Cdkn1a | DMKO79.7h | Homozygote | 27.49232 | 27.74837 | 28.17278 | 1.299879 | 1.088487 | 0.81108 | 1.066482 | 0.245141 |
| Cdkn1a | DMKO84.4e | Homozygote | 26.8972 | 27.43327 | 27.612 | 1.542249 | 1.063609 | 0.93968 | 1.181846 | 0.31821 |
| Tgfb1 | DMKO84.3d | Homozygote | 31.73383 | 31.48772 | 31.42078 | 0.813281 | 0.964552 | 1.010368 | 0.929401 | 0.103138 |
| Tgfb1 | DMKO84.3c | Homozygote | 31.70515 | 31.76253 | 30.58247 | 0.909568 | 0.8741 | 1.980587 | 1.254751 | 0.628842 |
| Tgfb1 | DMKO79.5f | Homozygote | 29.25918 | 30.08347 | 30.30703 | 1.538252 | 0.868744 | 0.744034 | 1.050344 | 0.427117 |
| Tgfb1 | DMKO79.7h | Homozygote | 30.20444 | 30.13288 | 30.96295 | 1.686386 | 1.772152 | 0.996833 | 1.485123 | 0.425041 |
| Tgfb1 | DMKO84.4e | Homozygote | 30.11331 | 30.21438 | 30.37287 | 1.519257 | 1.416465 | 1.269102 | 1.401608 | 0.125737 |
